## Supplementary Table 1 for "Mapping Cryptic Phosphorylation Sites in the Human Proteome"

### SUPP. TABLE 1

| SUB_ACCID | SUB_GN | SITE | RESPOS | SASA | pLDDT | LENGTH | RSA | PP_SASA | PP_RSA | PROP_IC |
|---|---|---|---|---|---|---|---|---|---|---|
| O43684 | BUB3 | TSQFLLVsSWDTSVR | S33 | 0 | 98.84 | 328 | 0 | 0 | 0 | 1 |
| O75347 | TBCA | QAEILQEsrMMIPDC | S60 | 0.077 | 89.9 | 108 | 0.061208 | 0.077 | 0.0612 | 1 |
| O75347 | TBCA | QRRLEAAyLDLQRIL | Y75 | 0.253 | 97.8 | 108 | 0.110987 | 0.253 | 0.111 | 1 |
| O75347 | TBCA | DLEEAEEykEARLVL | Y94 | 0.326 | 97.84 | 108 | 0.143012 | 0.326 | 0.143 | 1 |
| O60716 | CTNND1 | KARDMDLtEVITGTL | T469 | 0.061 | 98.49 | 968 | 0.041664 | 0.061 | 0.0417 | 1 |
| P18206 | VCL | RDyLIDGsRGILSGT | S112 | 0.008 | 93.38 | 1134 | 0.006359 | 0.008 | 0.0064 | 0.8571 |
| Q9H6R0 | DHX33 | DTRIkFLtDGMLLRE | T171 | 0.189 | 96.14 | 707 | 0.12909 | 0.189 | 0.1291 | 1 |
| Q14117 | DPYS | FPFMGSRsIDDFHQG | S79 | 0.064 | 98.21 | 519 | 0.050875 | 0.064 | 0.0509 | 0.8 |
| Q00005 | PPP2R2B | RyIMTRDyLTVKVWD | Y301 | 0.092 | 97.77 | 443 | 0.040359 | 0.092 | 0.0404 | 1 |
| Q9BRQ4 | CFAP300 | LPQKtFQsLsSKEIT | S24 | 0.084 | 93.83 | 267 | 0.066773 | 0.084 | 0.0668 | 1 |
| Q9BRQ4 | CFAP300 | SAGMCYPsAKNHEQT | S234 | 0.033 | 95.84 | 267 | 0.026232 | 0.033 | 0.0262 | 0.8571 |
| Q9BZH6 | WDR11 | SNGSVLVyHLtsGLL | Y500 | 0.082 | 93.09 | 1224 | 0.035972 | 0.268 | 0.1176 | 0.9 |
| Q9NRX2 | MRPL17 | HLLRNLLtGLVRHER | T34 | 0.053 | 98.55 | 175 | 0.0362 | 0.062 | 0.0531 | 1 |
| Q9NRX2 | MRPL17 | FQVLAPRyKDQtGGy | Y97 | 0.025 | 97.22 | 175 | 0.010967 | 0.025 | 0.011 | 1 |
| Q9NRX2 | MRPL17 | yKDQtGGytRMLQIP | Y104 | 0.044 | 97.15 | 175 | 0.019302 | 0.044 | 0.0193 | 1 |
| Q9NRX2 | MRPL17 | KDQtGGytRMLQIPN | T105 | 0.065 | 97.63 | 175 | 0.044396 | 0.105 | 0.0778 | 1 |
| Q14520 | HABP2 | GKRPGVYtQVtKFLN | T541 | 0.036 | 96.48 | 560 | 0.024588 | 0.036 | 0.0246 | 0.8333 |
| Q8TER0 | SNED1 | ALLPGKRyTIQLTTL | Y1073 | 0.176 | 91.62 | 1413 | 0.077209 | 0.176 | 0.0772 | 1 |
| Q5MY95 | ENTPD8 | TPTFLGAtAGMRLLS | T125 | 0.085 | 96.4 | 495 | 0.058056 | 0.085 | 0.0581 | 1 |
| Q8TDG4 | HELQ | LYLSFVLyTLLKETN | Y942 | 0.231 | 96.18 | 1101 | 0.101336 | 0.231 | 0.1013 | 1 |
| P35579 | MYH9 | MGLLRVIsGVLQLGN | S344 | 0 | 94.78 | 1960 | 0 | 0 | 0 | 1 |
| Q9NZW5 | PALS2 | ADIKAGkyLEHGEyE | Y404 | 0.201 | 95.23 | 540 | 0.088176 | 0.201 | 0.0882 | 0.9286 |
| Q9NZW5 | PALS2 | LKKTVDEsARIQRAy | S493 | 0.05 | 97.79 | 540 | 0.039746 | 0.05 | 0.0397 | 1 |
| Q8WWZ7 | ABCA5 | GGQkRKLsLGIAVLG | S624 | 0.064 | 92.25 | 1642 | 0.050875 | 0.116 | 0.0986 | 1 |
| Q8WWZ7 | ABCA5 | ILLLDEPtAGMDPCs | T642 | 0.012 | 86.04 | 1642 | 0.008196 | 0.012 | 0.0082 | 1 |
| P01833 | PIGR | GSFSVVItGLRKEDA | T313 | 0.002 | 91.89 | 764 | 0.001366 | 0.002 | 0.0014 | 0.9 |
| P01833 | PIGR | RKEDAGRyLCGAHSD | Y323 | 0.004 | 95.19 | 764 | 0.001755 | 0.004 | 0.0018 | 0.8667 |
| Q8TDM6 | DLG5 | NLsGQKDsGIsLENG | S730 | 0.005 | 82 | 1919 | 0.003975 | 0.005 | 0.004 | 1 |

| | | | | | | | | | | |
|---|---|---|---|---|---|---|---|---|---|---|
| Q6WBX8 | RAD9B | EANFILAtLADEQsR | T272 | 0.006 | 95.56 | 426 | 0.004098 | 0.154 | 0.1052 | 1 |
| Q9UGN5 | PARP2 | GLLQGkHsTkGLGKM | S514 | 0 | 98.61 | 583 | 0 | 0 | 0 | 1 |
| Q9Y6Q5 | AP1M2 | GkSRVPPtVtNAVSW | T152 | 0.058 | 90.25 | 423 | 0.039615 | 0.145 | 0.113 | 1 |
| Q9Y6Q5 | AP1M2 | YITQSGDyQLRTS__ | Y418 | 0.057 | 96.2 | 423 | 0.025005 | 0.057 | 0.025 | 1 |
| Q6DKJ4 | NXN | DPQGEVItRQGRVEV | T289 | 0.084 | 93.96 | 435 | 0.057373 | 0.084 | 0.0574 | 1 |
| Q8N4V2 | SVOP | RKTGLKIsVLWtLYY | S156 | 0.003 | 94.38 | 548 | 0.002385 | 0.003 | 0.0024 | 1 |
| Q8N4V2 | SVOP | LKIsVLWtLYYGILS | T160 | 0.045 | 96.29 | 548 | 0.030736 | 0.045 | 0.0307 | 1 |
| Q7Z4H7 | HAUS6 | QVLDQsLtKEVFKFC | T70 | 0.013 | 90.44 | 955 | 0.008879 | 0.013 | 0.0089 | 0.8889 |
| Q15075 | EEA1 | CSAKNALtPSSKKPV | T1392 | 0.094 | 82.66 | 1411 | 0.064203 | 0.094 | 0.0642 | 1 |
| Q9Y3R5 | DOP1B | TVKSLRAsLLDSNVL | S234 | 0.004 | 95.14 | 2298 | 0.00318 | 0.004 | 0.0032 | 1 |
| Q9Y3R5 | DOP1B | ILSAATQtLLRRDMS | T284 | 0.003 | 92.82 | 2298 | 0.002049 | 0.003 | 0.002 | 1 |
| Q6ZQW0 | IDO2 | GIRHSKEsGDFLYRM | S289 | 0.055 | 94.78 | 420 | 0.04372 | 0.055 | 0.0437 | 1 |
| P01889 | HLA-B | IEQEGPEyWDrNTQI | Y83 | 0.126 | 97.96 | 362 | 0.055274 | 0.126 | 0.0553 | 1 |
| P01889 | HLA-B | YAYDGkDyIALNEDL | Y147 | 0.118 | 98.73 | 362 | 0.051765 | 0.118 | 0.0518 | 1 |
| Q6P4A7 | SFXN4 | VIFLVQAsGMNVYMS | S209 | 0.008 | 97.32 | 337 | 0.006359 | 0.008 | 0.0064 | 0.8 |
| Q9UBX1 | CTSF | KNLGGLEtEDDYSYQ | T353 | 0.205 | 98.66 | 484 | 0.140018 | 0.205 | 0.14 | 0.875 |
| P25705 | ATP5F1A | TGIkAVDsLVPIGrG | S198 | 0.068 | 95.66 | 553 | 0.054054 | 0.068 | 0.0541 | 1 |
| P25705 | ATP5F1A | kTsIAIDtIINQkRF | T225 | 0.009 | 98.34 | 553 | 0.006147 | 0.009 | 0.0061 | 0.8462 |
| P25705 | ATP5F1A | SDEkkkLyCIyVAIG | Y243 | 0.249 | 98.45 | 553 | 0.109233 | 0.249 | 0.1092 | 1 |
| P25705 | ATP5F1A | kkkLyCIyVAIGQKR | Y246 | 0.055 | 98.72 | 553 | 0.024128 | 0.055 | 0.0241 | 1 |
| P25705 | ATP5F1A | GkHALIIyDDLSkQA | Y311 | 0.004 | 98.73 | 553 | 0.001755 | 0.004 | 0.0018 | 1 |
| P25705 | ATP5F1A | RPPGREAyPGDVFyL | Y337 | 0.191 | 96.86 | 553 | 0.083789 | 0.191 | 0.0838 | 1 |
| P12236 | SLC25A6 | GGIAAAIsktAVAPI | S22 | 0.131 | 92.43 | 298 | 0.104134 | 0.131 | 0.1041 | 1 |
| P12236 | SLC25A6 | QIAADkQykGIVDCI | Y51 | 0.147 | 96.34 | 298 | 0.064487 | 0.246 | 0.1079 | 1 |
| P12236 | SLC25A6 | PkEQGVLsFWRGNLA | S69 | 0.072 | 95.45 | 298 | 0.057234 | 0.072 | 0.0572 | 1 |
| P51784 | USP11 | NHkRRNDsVIVDTFH | S452 | 0 | 97.51 | 963 | 0 | 0 | 0 | 1 |
| Q8WVR3 | TRAPPC14 | tVAGKHLtVLKVLNS | T238 | 0.06 | 93.14 | 580 | 0.040981 | 0.182 | 0.1243 | 1 |
| Q8WVR3 | TRAPPC14 | YERFTVTyTLLNNLQ | Y390 | 0.001 | 89.43 | 580 | 0.000439 | 0.001 | 0.0004 | 1 |
| O14647 | CHD2 | GATGASTtVYAIEAN | T282 | 0.03 | 84.11 | 1828 | 0.02049 | 0.03 | 0.0205 | 0.875 |
| O95455 | TGDS | LIQLIKEtNsESEME | T279 | 0.129 | 95.59 | 350 | 0.088109 | 0.129 | 0.0881 | 1 |
| O00533 | CHL1 | QNGPGLEyRVTWKPQ | Y747 | 0.037 | 95.4 | 1208 | 0.016231 | 0.037 | 0.0162 | 1 |

| | | | | | | | | | | |
|---|---|---|---|---|---|---|---|---|---|---|
| Q6IAN0 | DHRS7B | YRGtIMDtTVDVDkR | T154 | 0.197 | 96.89 | 325 | 0.134554 | 0.197 | 0.1346 | 1 |
| Q04760 | GLO1 | rVKDPKksLDFYTRV | S45 | 0.007 | 98.89 | 184 | 0.005564 | 0.007 | 0.0056 | 0.9 |
| Q14999 | CUL7 | FNILQNSsSELFGPR | S1183 | 0.026 | 83.02 | 1698 | 0.020668 | 0.036 | 0.0343 | 1 |
| Q9NPH3 | IL1RAP | DVLWFRPtLLNDTGN | T104 | 0.004 | 95.77 | 570 | 0.002732 | 0.004 | 0.0027 | 1 |
| Q9NPH3 | IL1RAP | LLNDTGNytCMLRNT | Y112 | 0.042 | 94.79 | 570 | 0.018425 | 0.042 | 0.0184 | 1 |
| Q9NPH3 | IL1RAP | LNDTGNytCMLRNTT | T113 | 0.103 | 94.27 | 570 | 0.07035 | 0.103 | 0.0703 | 1 |
| P32121 | ARRB2 | TRVFKKssPNCKLtV | S14 | 0.039 | 96.03 | 409 | 0.031002 | 0.039 | 0.031 | 1 |
| P32121 | ARRB2 | ssPNCKLtVYLGKRD | T20 | 0.119 | 97.77 | 409 | 0.081279 | 0.119 | 0.0813 | 1 |
| Q9Y6L7 | TLL2 | SPNWPDKyPsRRECT | Y796 | 0.034 | 89.95 | 1015 | 0.014915 | 0.034 | 0.0149 | 1 |
| Q9Y6L7 | TLL2 | SLMIRFRtDDTINKK | T989 | 0.159 | 94.33 | 1015 | 0.108599 | 0.159 | 0.1086 | 1 |
| P0CW24 | PNMA6A | MLGLVREsEAWEASL | S325 | 0.138 | 95.08 | 399 | 0.109698 | 0.138 | 0.1097 | 1 |
| Q12906 | ILF3 | SLTIHLTsPVVREEM | S173 | 0 | 96.7 | 894 | 0 | 0.014 | 0.0111 | 0.8571 |
| Q9BVP2 | GNL3 | LkkVIEAsDVVLEVL | S141 | 0.067 | 94.83 | 549 | 0.053259 | 0.067 | 0.0533 | 1 |
| Q9BVP2 | GNL3 | NLESWLNyLkkELPT | Y194 | 0.207 | 94.73 | 549 | 0.090808 | 0.207 | 0.0908 | 1 |
| Q9BVP2 | GNL3 | VGVSMGLtRsMQVVP | T289 | 0.175 | 86.32 | 549 | 0.119527 | 0.175 | 0.1195 | 1 |
| Q9H0R1 | AP5M1 | ISLSGTVtFGAKSHE | T408 | 0.204 | 94.09 | 490 | 0.139335 | 0.204 | 0.1393 | 1 |
| P31327 | CPS1 | kEPLFGIstGNLITG | S294 | 0.012 | 98.46 | 1500 | 0.009539 | 0.012 | 0.0095 | 0.8889 |
| P31327 | CPS1 | EPLFGIstGNLITGL | T295 | 0.056 | 98.4 | 1500 | 0.038249 | 0.056 | 0.0382 | 1 |
| P31327 | CPS1 | sKVLILGsGGLSIGQ | S430 | 0.075 | 90.72 | 1500 | 0.059619 | 0.075 | 0.0596 | 1 |
| P31327 | CPS1 | sVESIMAtEDrQLFS | T544 | 0.023 | 97.96 | 1500 | 0.015709 | 0.046 | 0.0314 | 0.875 |
| P31327 | CPS1 | GYPVMIRsAyALGGL | S588 | 0.059 | 96.61 | 1500 | 0.0469 | 0.117 | 0.0931 | 1 |
| P31327 | CPS1 | LASkATGyPLAFIAA | Y732 | 0.04 | 98.17 | 1500 | 0.017547 | 0.04 | 0.0175 | 0.8571 |
| P31327 | CPS1 | RIGSSMksVGEVMAI | S794 | 0.026 | 96.98 | 1500 | 0.020668 | 0.026 | 0.0207 | 0.8889 |
| P31327 | CPS1 | GFsDkQIskCLGLtE | S918 | 0.084 | 98.48 | 1500 | 0.066773 | 0.114 | 0.1099 | 1 |
| P31327 | CPS1 | sRSFPFVsktLGVDF | S1268 | 0.055 | 97.48 | 1500 | 0.04372 | 0.087 | 0.0717 | 0.875 |
| P31327 | CPS1 | SFPFVsktLGVDFID | T1270 | 0 | 96.71 | 1500 | 0 | 0 | 0 | 0.8182 |
| P31327 | CPS1 | ILRCEMAstGEVACF | S1331 | 0.021 | 92.29 | 1500 | 0.016693 | 0.021 | 0.0167 | 0.875 |
| P31327 | CPS1 | DSkSLFHyRQYSAGK | Y1491 | 0.054 | 90.18 | 1500 | 0.023689 | 0.268 | 0.0876 | 1 |
| Q969K3 | RNF34 | kDFCSVCsVLQENLR | S90 | 0.102 | 95.58 | 372 | 0.081081 | 0.102 | 0.0811 | 1 |
| O15540 | FABP7 | DVVAVRHyEKA____ | Y129 | 0.055 | 98.46 | 132 | 0.024128 | 0.055 | 0.0241 | 1 |
| O95786 | RIGI | kEkMMNDsILRLQtW | S764 | 0.006 | 96.62 | 925 | 0.004769 | 0.006 | 0.0048 | 1 |

| | | | | | | | | | | |
|---|---|---|---|---|---|---|---|---|---|---|
| Q9BYF1 | ACE2 | HKCDIsNstEAGQKL | S547 | 0.106 | 97.57 | 805 | 0.084261 | 0.106 | 0.0843 | 1 |
| Q9BYF1 | ACE2 | ANLkPRIsFNFFVTA | S680 | 0.184 | 93.33 | 805 | 0.146264 | 0.184 | 0.1463 | 1 |
| P09327 | VIL1 | LKTQSCCyLWCGKGC | Y555 | 0.147 | 94.93 | 827 | 0.064487 | 0.236 | 0.1035 | 1 |
| Q9UQB8 | BAIAP2 | HRLTENVyKTIMEQF | Y17 | 0.134 | 98.39 | 552 | 0.058784 | 0.134 | 0.0588 | 1 |
| Q9NRS6 | SNX15 | HIPALNNsPQLKEFF | S116 | 0.045 | 92.62 | 342 | 0.035771 | 0.178 | 0.1415 | 1 |
| Q9NZM1 | MYOF | GEEMSDIyVkGWIPG | Y1816 | 0.067 | 93.08 | 2061 | 0.029392 | 0.067 | 0.0294 | 1 |
| O14497 | ARID1A | PLDLYRLyVsVKEIG | Y1055 | 0.135 | 94.11 | 2285 | 0.059223 | 0.312 | 0.1369 | 1 |
| Q96Q05 | TRAPPC9 | RMGNPALsVRHLSFL | S480 | 0.001 | 95.4 | 1148 | 0.000795 | 0.001 | 0.0008 | 1 |
| Q8NE71 | ABCF1 | PNGVGKstLLLLLtG | T666 | 0.111 | 92.88 | 845 | 0.075815 | 0.111 | 0.0758 | 1 |
| Q8NE71 | ABCF1 | MEETPTEyLQRGFNL | Y710 | 0.275 | 94.83 | 845 | 0.120639 | 0.275 | 0.1206 | 1 |
| Q9NR22 | PRMT8 | GSGTGILsMFAAKAG | S126 | 0 | 98.89 | 394 | 0 | 0 | 0 | 1 |
| Q9NR22 | PRMT8 | WMGyCLFyESMLNTV | Y193 | 0.229 | 95.68 | 394 | 0.100459 | 0.326 | 0.143 | 1 |
| P16383 | GCFC2 | SAVENKTsPHSKFQE | S637 | 0.164 | 86.2 | 781 | 0.130366 | 0.164 | 0.1304 | 1 |
| Q86XP3 | DDX42 | DIPVLVAtDVAARGL | T555 | 0.117 | 91.69 | 938 | 0.079913 | 0.117 | 0.0799 | 1 |
| Q86XP3 | DDX42 | GVAYtLLtPkDSNFA | T603 | 0.149 | 92.09 | 938 | 0.101769 | 0.149 | 0.1018 | 0.8333 |
| O75683 | SURF6 | LEAKMKWtNLLyKAE | T271 | 0.105 | 92.25 | 361 | 0.071716 | 0.105 | 0.0717 | 1 |
| O75683 | SURF6 | MKWtNLLyKAEGVKI | Y275 | 0.171 | 92.16 | 361 | 0.075015 | 0.171 | 0.075 | 1 |
| Q6NUN0 | ACSM5 | VSVACMRtGTVMIPG | T138 | 0.157 | 98.52 | 579 | 0.107233 | 0.157 | 0.1072 | 1 |
| Q6NUN0 | ACSM5 | QSLRHCLtGGEALNP | T345 | 0.001 | 92.87 | 579 | 0.000683 | 0.001 | 0.0007 | 1 |
| Q6NUN0 | ACSM5 | LESAVVSsPDPIRGE | S505 | 0.011 | 95.64 | 579 | 0.008744 | 0.153 | 0.1216 | 1 |
| O75306 | NDUFS2 | kVDDAKVsPPKRAEM | S364 | 0.106 | 91.36 | 463 | 0.084261 | 0.106 | 0.0843 | 0.8571 |
| P23258 | TUBG1 | LCAEHGIsPEGIVEE | S32 | 0.11 | 96.71 | 451 | 0.087441 | 0.11 | 0.0874 | 1 |
| P23258 | TUBG1 | MANHTSIsSLFERTC | S385 | 0 | 95.44 | 451 | 0 | 0 | 0 | 1 |
| P23258 | TUBG1 | HAATRPDyISWGTQE | Y443 | 0.176 | 93.13 | 451 | 0.077209 | 0.176 | 0.0772 | 1 |
| Q8TE60 | ADAMTS18 | CKFYKGLyLNQHKAN | Y755 | 0.237 | 91.97 | 1221 | 0.103968 | 0.237 | 0.104 | 1 |
| Q8TE60 | ADAMTS18 | QHKANEyyPVVLIPA | Y765 | 0.288 | 92.32 | 1221 | 0.126341 | 0.288 | 0.1263 | 1 |
| Q8TE60 | ADAMTS18 | SSyLAVRsLsQKYYL | S793 | 0.009 | 92.17 | 1221 | 0.007154 | 0.009 | 0.0072 | 1 |
| Q8TBY0 | RBM46 | LDEMKKVtEGVVDVI | T165 | 0.089 | 93.5 | 533 | 0.060788 | 0.089 | 0.0608 | 1 |
| Q9HC84 | MUC5B | ETHGMAVsWDRKTSV | S1003 | 0.009 | 92.865 | 5762 | 0.007154 | 0.009 | 0.0072 | 1 |
| A1L020 | MEX3A | VVGPKGAtIKRIQQQ | T245 | 0.106 | 93.74 | 520 | 0.072399 | 0.106 | 0.0724 | 1 |
| Q7Z449 | CYP2U1 | WRQQRKFsHSTLRHF | S181 | 0 | 96.7 | 544 | 0 | 0 | 0 | 1 |

| | | | | | | | | | | |
|---|---|---|---|---|---|---|---|---|---|---|
| P09914 | IFIT1 | DVNAIIHyLKAIkIE | Y402 | 0.006 | 98.5 | 478 | 0.002632 | 0.006 | 0.0026 | 0.9474 |
| P09914 | IFIT1 | ASLTRDksINSLkKL | S418 | 0 | 97.88 | 478 | 0 | 0 | 0 | 1 |
| Q16772 | GSTA3 | GKLRNDGsLMFQQVP | S49 | 0.069 | 97.99 | 222 | 0.054849 | 0.069 | 0.0548 | 0.8 |
| P35499 | SCN4A | GFCVDDFtFLRDPWN | T187 | 0.052 | 82.75 | 1836 | 0.035517 | 0.052 | 0.0355 | 1 |
| A0AV02 | SLC12A8 | SIGSGGVySMISSVL | Y107 | 0.038 | 92.84 | 714 | 0.01667 | 0.038 | 0.0167 | 1 |
| Q6P597 | KLC3 | KNNLAsAyLKQNKYQ | Y385 | 0.292 | 93.93 | 504 | 0.128096 | 0.292 | 0.1281 | 0.8889 |
| Q5JY77 | GPRASP1 | IRMIRHLtTTTDYHT | T1265 | 0.004 | 93.15 | 1395 | 0.002732 | 0.004 | 0.0027 | 1 |
| Q9NP60 | IL1RAPL2 | IKWKGSKsskLNSkF | S541 | 0.141 | 73.24 | 686 | 0.112083 | 0.141 | 0.1121 | 1 |
| Q8NBJ5 | COLGALT1 | YYKRTPAyIPIRKRD | Y217 | 0.097 | 95.95 | 622 | 0.042553 | 0.097 | 0.0426 | 1 |
| Q8NBJ5 | COLGALT1 | YVCNKEEyGFLPVPL | Y288 | 0.029 | 98.31 | 622 | 0.012722 | 0.029 | 0.0127 | 1 |
| Q8NBJ5 | COLGALT1 | KHPPAEPsRFISAPt | S325 | 0.063 | 97.33 | 622 | 0.05008 | 0.063 | 0.0501 | 1 |
| O00571 | DDX3X | HQFRsGksPILVATA | S492 | 0.11 | 91.92 | 662 | 0.087441 | 0.11 | 0.0874 | 1 |
| Q16540 | MRPL23 | TRVDLrNyLEGIyNV | Y56 | 0.316 | 97.32 | 153 | 0.138625 | 0.316 | 0.1386 | 1 |
| P61160 | ACTR2 | DMkHLWDytFGPEkL | Y91 | 0.23 | 97.37 | 394 | 0.100898 | 0.23 | 0.1009 | 1 |
| Q13304 | GPR17 | CYLLIIRsLRQGLRV | S246 | 0.123 | 90.76 | 367 | 0.097775 | 0.123 | 0.0978 | 1 |
| Q13304 | GPR17 | GALDPIMyFFVAEKF | Y325 | 0 | 90.09 | 367 | 0 | 0.274 | 0.1202 | 0.9 |
| Q53HC9 | EIPR1 | AIRFLVGtQSLKYDN | T34 | 0 | 97.58 | 387 | 0 | 0 | 0 | 1 |
| Q8WV44 | TRIM41 | DLTLDPDtAHPALML | T439 | 0.045 | 90.83 | 630 | 0.030736 | 0.045 | 0.0307 | 0.9091 |
| Q8WV44 | TRIM41 | RFGVyLDyEAGRLGF | Y585 | 0.274 | 92.97 | 630 | 0.1202 | 0.274 | 0.1202 | 1 |
| P05121 | SERPINE1 | LVLVNALyFNGQWKt | Y193 | 0.039 | 94.93 | 402 | 0.017109 | 0.039 | 0.0171 | 0.8889 |
| Q8IVV2 | LOXHD1 | ATLKNFRyHISLKtG | Y686 | 0 | 87.46 | 2067 | 0 | 0 | 0 | 1 |
| Q8IVV2 | LOXHD1 | RyHISLKtGDVsGAs | T692 | 0.01 | 91.44 | 2067 | 0.00683 | 0.01 | 0.0068 | 1 |
| Q8IVV2 | LOXHD1 | GDVsGAstDSRVYIK | T700 | 0.07 | 87.08 | 2067 | 0.047811 | 0.07 | 0.0478 | 1 |
| Q8IVV2 | LOXHD1 | EIQKLVHyEVEIWTG | Y816 | 0 | 80.67 | 2067 | 0 | 0 | 0 | 1 |
| P35251 | RFC1 | LLsRAADsICDGDLV | S914 | 0.032 | 96.9 | 1148 | 0.025437 | 0.032 | 0.0254 | 1 |
| O43731 | KDELR3 | FRILGDLsHLLAMIL | S11 | 0.083 | 96.29 | 214 | 0.065978 | 0.083 | 0.066 | 1 |
| Q8NB78 | KDM1B | KKHSVLMsVIAGEAV | S711 | 0 | 98.83 | 822 | 0 | 0 | 0 | 1 |
| Q8NB78 | KDM1B | DPtKYFVtRWSTDPW | T755 | 0.073 | 98.7 | 822 | 0.04986 | 0.073 | 0.0499 | 0.8571 |
| P14780 | MMP9 | FIFEGRSySACTTDG | Y241 | 0.306 | 84.66 | 707 | 0.134238 | 0.306 | 0.1342 | 1 |
| P51606 | RENBP | LFYQVAEytFRQFRD | Y355 | 0.248 | 98.78 | 427 | 0.108794 | 0.248 | 0.1088 | 1 |
| P51606 | RENBP | FYQVAEytFRQFRDP | T356 | 0 | 98.83 | 427 | 0 | 0 | 0 | 1 |

| | | | | | | | | | | |
|---|---|---|---|---|---|---|---|---|---|---|
| Q8WXX0 | DNAH7 | VEADIGNyWRGLYKL | Y803 | 0.16 | 90.07 | 4024 | 0.07019 | 0.309 | 0.1316 | 1 |
| Q8WXX0 | DNAH7 | LERVMINsIHKVTGD | S1154 | 0.018 | 96.725 | 4024 | 0.014308 | 0.018 | 0.0143 | 1 |
| Q6ZU15 | SEPTIN14 | NILCVGEtGIGKSTL | T61 | 0.168 | 90.44 | 432 | 0.114746 | 0.168 | 0.1147 | 0.9091 |
| P0DJD0 | RGPD1 | LIKIIDDsDsNLsVV | S716 | 0.067 | 86.24 | 1748 | 0.053259 | 0.083 | 0.066 | 1 |
| Q8IUR5 | TMTC1 | TKEAEKMtNHIVSEE | T740 | 0 | 96.63 | 882 | 0 | 0 | 0 | 1 |
| Q8IV04 | TBC1D10C | MtsHWEktMsRRYKK | T77 | 0.002 | 97.53 | 446 | 0.001366 | 0.002 | 0.0014 | 1 |
| Q13868 | EXOSC2 | RLRNCIIsLVTQRMM | S243 | 0.006 | 91.36 | 293 | 0.004769 | 0.006 | 0.0048 | 1 |
| Q9Y2X7 | GIT1 | GVTAKDLskQLHSSV | S134 | 0.002 | 96.72 | 761 | 0.00159 | 0.002 | 0.0016 | 1 |
| Q9Y2X7 | GIT1 | FHPEKGttPLHVAAK | T169 | 0.052 | 98.43 | 761 | 0.035517 | 0.052 | 0.0355 | 1 |
| Q9BR39 | JPH2 | KRSGFGVsERssGLR | S306 | 0.107 | 95.55 | 696 | 0.085056 | 0.107 | 0.0851 | 1 |
| Q9BZ72 | PITPNM2 | VFPIMCAyKLCKVEF | Y191 | 0 | 98.11 | 1349 | 0 | 0 | 0 | 0.9333 |
| Q6ZVZ8 | ASB18 | ARNGRGEtALsAACG | T254 | 0.025 | 96.47 | 466 | 0.017075 | 0.025 | 0.0171 | 1 |
| Q6ZVZ8 | ASB18 | GRGEtALsAACGAAR | S257 | 0.025 | 94.88 | 466 | 0.019873 | 0.025 | 0.0199 | 1 |
| Q8TD57 | DNAH3 | RKRLFIEsIPRLFPQ | S313 | 0.128 | 83.58 | 4116 | 0.101749 | 0.1395 | 0.1109 | 0.8333 |
| Q96S86 | HAPLN3 | QDKEHDVsLEIQDLR | S130 | 0.057 | 97.68 | 360 | 0.04531 | 0.057 | 0.0453 | 1 |
| Q96S86 | HAPLN3 | RHRRLHRyDVFCFAt | Y255 | 0.331 | 98.69 | 360 | 0.145205 | 0.331 | 0.1452 | 1 |
| Q9BZQ8 | NIBAN1 | LMNLPLHsVkMEPCY | S403 | 0.119 | 95.18 | 928 | 0.094595 | 0.119 | 0.0946 | 1 |
| P07199 | CENPB | NKRAILAsERkyGVA | S55 | 0.101 | 89.36 | 599 | 0.080286 | 0.101 | 0.0803 | 1 |
| P68032 | ACTC1 | APrAVFPsIVGRPRH | S35 | 0.003 | 97.78 | 377 | 0.002385 | 0.003 | 0.0024 | 1 |
| P68032 | ACTC1 | GMGQkDsyVGDEAQs | Y55 | 0.041 | 96.07 | 377 | 0.017986 | 0.12 | 0.0526 | 1 |
| P68032 | ACTC1 | MEkIWHHtFyNELRV | T91 | 0.003 | 98.66 | 377 | 0.002049 | 0.003 | 0.002 | 0.9 |
| P68032 | ACTC1 | DLAGRDLtDyLMkIL | T188 | 0.017 | 98.3 | 377 | 0.011611 | 0.017 | 0.0116 | 0.9 |
| P68032 | ACTC1 | ERFRCPEtLFQPsFI | T262 | 0.003 | 98.41 | 377 | 0.002049 | 0.003 | 0.002 | 0.8889 |
| P68032 | ACTC1 | IDIRkDLyANNVLSG | Y296 | 0.004 | 98.37 | 377 | 0.001755 | 0.004 | 0.0018 | 1 |
| P68032 | ACTC1 | MWISKQEyDEAGPsI | Y364 | 0.127 | 98.35 | 377 | 0.055713 | 0.251 | 0.1094 | 1 |
| P56199 | ITGA1 | KEEQGKVyVYALNQT | Y527 | 0.065 | 98.43 | 1179 | 0.028515 | 0.065 | 0.0285 | 1 |
| P56199 | ITGA1 | DDHGGAVyIYHGSGK | Y604 | 0 | 97.38 | 1179 | 0 | 0 | 0 | 1 |
| P24723 | PRKCH | FYAAEIIsALMFLHD | S465 | 0.119 | 98.4 | 683 | 0.094595 | 0.119 | 0.0946 | 0.8 |
| P24723 | PRKCH | NPTMRLGsLtQGGEH | S599 | 0.168 | 92.22 | 683 | 0.133546 | 0.168 | 0.1335 | 1 |
| P51617 | IRAK1 | CERSGQRtASVLWPW | T66 | 0 | 94.18 | 712 | 0 | 0 | 0 | 0.875 |
| Q06203 | PPAT | QESAGIVtSDGSSVP | T52 | 0.004 | 97.28 | 517 | 0.002732 | 0.021 | 0.0143 | 1 |

| | | | | | | | | | | |
|---|---|---|---|---|---|---|---|---|---|---|
| Q9H871 | RMND5A | LVCGHIIsRDALNkM | S359 | 0.001 | 95.13 | 391 | 0.000795 | 0.001 | 0.0008 | 1 |
| P19367 | HK1 | LPVGFTFsFPCQQSK | S155 | 0.071 | 97.88 | 917 | 0.056439 | 0.071 | 0.0564 | 1 |
| P19367 | HK1 | QLFEKMVsGMyLGEL | S298 | 0 | 97.83 | 917 | 0 | 0 | 0 | 1 |
| P19367 | HK1 | EKMVsGMyLGELVRL | Y301 | 0.108 | 98.25 | 917 | 0.047378 | 0.108 | 0.0474 | 1 |
| P19367 | HK1 | kFNTsDVsAIEkNkE | S340 | 0.124 | 98.03 | 917 | 0.098569 | 0.124 | 0.0986 | 1 |
| P19367 | HK1 | VGVDGSLyktHPQYS | Y417 | 0.23 | 97.89 | 917 | 0.100898 | 0.23 | 0.1009 | 1 |
| P19367 | HK1 | EKMISGMyLGEIVRN | Y749 | 0.109 | 97.27 | 917 | 0.047817 | 0.109 | 0.0478 | 1 |
| P19367 | HK1 | KtVCGVVsRRAAQLC | S827 | 0.036 | 98.72 | 917 | 0.028617 | 0.036 | 0.0286 | 1 |
| P19367 | HK1 | CNVSFLLsEDGSGKG | S893 | 0.053 | 80.63 | 917 | 0.04213 | 0.053 | 0.0421 | 1 |
| P49821 | NDUFV1 | NVETVAVsPTICRrG | S251 | 0 | 98.65 | 464 | 0 | 0 | 0 | 0.9167 |
| P49821 | NDUFV1 | VIPGGSstPLIPKSV | T324 | 0.063 | 97.73 | 464 | 0.04303 | 0.063 | 0.043 | 0.9167 |
| P21333 | FLNA | FEDRkDGsCGVAYVV | S2292 | 0.018 | 72.56 | 2647 | 0.014308 | 0.018 | 0.0143 | 0.8333 |
| O95749 | GGPS1 | GIPsVINsANyVYFL | S91 | 0.007 | 98.64 | 300 | 0.005564 | 0.007 | 0.0056 | 1 |
| Q9UNA0 | ADAMTS5 | TEDKRLMssILtSID | S440 | 0.062 | 93.85 | 930 | 0.049285 | 0.108 | 0.0916 | 1 |
| P09917 | ALOX5 | ERGAVDSyDVTVDEE | Y54 | 0.175 | 97.28 | 674 | 0.07677 | 0.175 | 0.0768 | 1 |
| P09917 | ALOX5 | DWYLKYItLKTPHGD | T87 | 0.084 | 98.59 | 674 | 0.057373 | 0.084 | 0.0574 | 1 |
| P09917 | ALOX5 | DyIEFPCyRWItGDV | Y101 | 0.231 | 97.82 | 674 | 0.101336 | 0.29 | 0.1272 | 0.8889 |
| P09917 | ALOX5 | GVDFVLNysKAMENL | Y182 | 0.113 | 95.81 | 674 | 0.049571 | 0.226 | 0.0991 | 0.9231 |
| P09917 | ALOX5 | RAMKDLtyAsLCFPE | Y446 | 0.172 | 98.76 | 674 | 0.075454 | 0.331 | 0.1406 | 1 |
| P09917 | ALOX5 | WEAIRTFtAEVVDIY | T486 | 0 | 98.88 | 674 | 0 | 0 | 0 | 1 |
| P09917 | ALOX5 | GMRGRKSsGFPkSVK | S524 | 0.013 | 97.62 | 674 | 0.010334 | 0.013 | 0.0103 | 1 |
| P09917 | ALOX5 | QLPYYYLsPDRIPNs | S664 | 0.096 | 98.74 | 674 | 0.076312 | 0.096 | 0.0763 | 1 |
| P09917 | ALOX5 | sPDRIPNsVAI____ | S671 | 0.092 | 98.73 | 674 | 0.073132 | 0.1 | 0.0889 | 1 |
| Q9H8V3 | ECT2 | IGDIFLkySKDLVkT | Y539 | 0.169 | 95.5 | 914 | 0.074138 | 0.169 | 0.0741 | 1 |
| Q15648 | MED1 | KMYLALQsLEQDLSK | S179 | 0.043 | 92.22 | 1581 | 0.034181 | 0.043 | 0.0342 | 1 |
| Q15648 | MED1 | GHLMNLKyyVSPSDL | Y224 | 0.003 | 92.56 | 1581 | 0.001316 | 0.003 | 0.0013 | 0.9167 |
| Q15648 | MED1 | kVVQRCMsIPVtMRA | S503 | 0.108 | 94.76 | 1581 | 0.085851 | 0.108 | 0.0859 | 0.8 |
| Q15648 | MED1 | RCMsIPVtMRAIRRK | T507 | 0.014 | 95.31 | 1581 | 0.009562 | 0.014 | 0.0096 | 1 |
| P59045 | NLRP11 | AANILLPsNTHKDRy | S439 | 0.172 | 79.42 | 1033 | 0.136725 | 0.172 | 0.1367 | 1 |
| P29353 | SHC1 | GDFLVREsTTTPGQy | S513 | 0.17 | 84.89 | 583 | 0.135135 | 0.17 | 0.1351 | 1 |
| Q13882 | PTK6 | PsADyVLsVRDTQAV | S117 | 0 | 95.81 | 451 | 0 | 0 | 0 | 1 |

| | | | | | | | | | | |
|---|---|---|---|---|---|---|---|---|---|---|
| Q13882 | PTK6 | DTQAVRHyKIWRRAG | Y127 | 0 | 93.27 | 451 | 0 | 0.102 | 0.0447 | 1 |
| Q13882 | PTK6 | IKEDVyLsHDHNIPy | S344 | 0.13 | 86.48 | 451 | 0.103339 | 0.13 | 0.1033 | 1 |
| Q9Y5S9 | RBM8A | KGyTLVEyEtYkEAQ | Y121 | 0 | 98.76 | 174 | 0 | 0 | 0 | 0.9375 |
| Q8WVL7 | ANKRD49 | EKAtHVNtRDEDEYT | T103 | 0.168 | 96.57 | 239 | 0.114746 | 0.168 | 0.1147 | 1 |
| Q02763 | TEK | GEAIRIRtMKMRQQA | T115 | 0.004 | 97.16 | 1124 | 0.002732 | 0.004 | 0.0027 | 1 |
| P25092 | GUCY2C | VGIKMPRyCLFGDTV | Y940 | 0.131 | 89.57 | 1073 | 0.057468 | 0.131 | 0.0575 | 1 |
| P25092 | GUCY2C | LPLRIHVsGSTIAIL | S965 | 0.015 | 95.67 | 1073 | 0.011924 | 0.015 | 0.0119 | 1 |
| O76031 | CLPX | tNILFVAsGAFNGLD | S420 | 0 | 94.32 | 633 | 0 | 0 | 0 | 0.875 |
| O76031 | CLPX | RNAVIPQyQALFSMD | Y529 | 0.278 | 92.86 | 633 | 0.121955 | 0.278 | 0.122 | 1 |
| Q8IWI9 | MGA | SPVDNHRyKWNGRWW | Y135 | 0.281 | 97.67 | 3065 | 0.123271 | 0.281 | 0.1233 | 1 |
| Q8IWI9 | MGA | SFYKLKLtNNTLDQE | T182 | 0 | 96.26 | 3065 | 0 | 0 | 0 | 1 |
| Q9HCL0 | PCDH18 | QKTyENNyLILTNAT | Y415 | 0.117 | 91.14 | 1135 | 0.051326 | 0.117 | 0.0513 | 1 |
| Q8NB49 | ATP11C | IPVsMYVtVEMQKFL | T365 | 0.127 | 89.71 | 1132 | 0.086743 | 0.127 | 0.0867 | 0.8 |
| A0PJK1 | SLC5A10 | SVLsLLLsVFtKISL | S145 | 0.08 | 90.46 | 596 | 0.063593 | 0.08 | 0.0636 | 1 |
| A0PJK1 | SLC5A10 | yGQLEAAyAQAIPsR | Y228 | 0.039 | 96.96 | 596 | 0.017109 | 0.039 | 0.0171 | 1 |
| A0PJK1 | SLC5A10 | LNHAKAGsILASYLK | S301 | 0.001 | 96.53 | 596 | 0.000795 | 0.001 | 0.0008 | 1 |
| A0PJK1 | SLC5A10 | IIMPGMIsRALFPDD | S322 | 0 | 97.55 | 596 | 0 | 0 | 0 | 1 |
| A0PJK1 | SLC5A10 | NSSSTLFtMDIWRRL | T393 | 0 | 94.8 | 596 | 0 | 0 | 0 | 1 |
| Q08881 | ITK | LAtGCAQyDPTkNAS | Y146 | 0.205 | 91.7 | 620 | 0.089931 | 0.205 | 0.0899 | 0.9091 |
| Q9Y6C9 | MTCH2 | KYCGLCDsIITIYRE | S165 | 0.004 | 95.42 | 303 | 0.00318 | 0.004 | 0.0032 | 1 |
| Q9NX01 | TXNL4B | RGkLIVQsPIDPkNI | S132 | 0.177 | 97.49 | 149 | 0.1407 | 0.177 | 0.1407 | 1 |
| P37023 | ACVRL1 | LWEIARRtIVNGIVE | T412 | 0 | 98.12 | 503 | 0 | 0 | 0 | 1 |
| O43451 | MGAM | TRSTFAGsGKFAAHW | S618 | 0.012 | 98.8 | 1857 | 0.009539 | 0.012 | 0.0095 | 1 |
| O43451 | MGAM | LLNSSRHyLNIRYTL | Y712 | 0 | 98.78 | 1857 | 0 | 0 | 0 | 1 |
| O43451 | MGAM | NSYGVHPyyMGLEED | Y1172 | 0.001 | 98.57 | 1857 | 0.000439 | 0.001 | 0.0004 | 1 |
| O43451 | MGAM | SYGVHPyyMGLEEDG | Y1173 | 0.009 | 98.15 | 1857 | 0.003948 | 0.009 | 0.0039 | 1 |
| O43451 | MGAM | YFPRARWyDyytGVD | Y1683 | 0.019 | 98.32 | 1857 | 0.008335 | 0.019 | 0.0083 | 1 |
| P52907 | CAPZA1 | TVSNEAQtAkEFIkI | T224 | 0.008 | 96.99 | 286 | 0.005464 | 0.008 | 0.0055 | 1 |
| Q96QF7 | GCNA | DELVQRIyDLFNRSV | Y526 | 0.108 | 97.98 | 691 | 0.047378 | 0.108 | 0.0474 | 1 |
| P30040 | ERP29 | FKRLAENsASsDDLL | S80 | 0.099 | 88.55 | 261 | 0.078697 | 0.099 | 0.0787 | 1 |
| P30040 | ERP29 | LAENsASsDDLLVAE | S83 | 0.104 | 85.66 | 261 | 0.082671 | 0.104 | 0.0827 | 0.8571 |

| | | | | | | | | | | |
|---|---|---|---|---|---|---|---|---|---|---|
| P30040 | ERP29 | KKEELQksLNILtAF | S244 | 0.123 | 92.31 | 261 | 0.097775 | 0.142 | 0.1129 | 1 |
| Q9UKA2 | FBXL4 | IKQINFPtNLIRLEV | T202 | 0 | 96.43 | 621 | 0 | 0 | 0 | 1 |
| Q9Y2G5 | POFUT2 | YRGWFWGyEEtRGLN | Y182 | 0.107 | 96.3 | 429 | 0.046939 | 0.107 | 0.0469 | 1 |
| Q9Y2G5 | POFUT2 | WFWGyEEtRGLNVsC | T185 | 0.013 | 97.3 | 429 | 0.008879 | 0.013 | 0.0089 | 1 |
| Q9Y2G5 | POFUT2 | EtRGLNVsCLsVQGS | S191 | 0.163 | 97.08 | 429 | 0.129571 | 0.163 | 0.1296 | 1 |
| Q9Y2G5 | POFUT2 | GLNVsCLsVQGSASI | S194 | 0.009 | 97.88 | 429 | 0.007154 | 0.009 | 0.0072 | 1 |
| Q8WVM8 | SCFD1 | RNQLYEsyyLNFISA | Y114 | 0.018 | 98.01 | 642 | 0.007896 | 0.018 | 0.0079 | 1 |
| P31645 | SLC6A4 | FKERIIKsItPEtPT | S611 | 0 | 93.84 | 630 | 0 | 0 | 0 | 1 |
| Q8TAG9 | EXOC6 | IYASLKFsEsLHRSS | S503 | 0.072 | 83.52 | 804 | 0.057234 | 0.072 | 0.0572 | 0.8571 |
| Q15399 | TLR1 | IItCIEKsyKSIFVL | S690 | 0.087 | 85.95 | 786 | 0.069158 | 0.087 | 0.0692 | 0.9091 |
| Q12907 | LMAN2 | ERVFPYIsVMVNNGS | S178 | 0.094 | 98.16 | 356 | 0.074722 | 0.094 | 0.0747 | 0.8333 |
| Q12907 | LMAN2 | SAGTGDLsDNHDIIS | S263 | 0.176 | 96.95 | 356 | 0.139905 | 0.176 | 0.1399 | 1 |
| Q99570 | PIK3R4 | yGAVGFItVVARQIS | T680 | 0.009 | 92.2 | 1358 | 0.006147 | 0.009 | 0.0061 | 1 |
| Q8NEY4 | ATP6V1C2 | LIRRMAQsVVEVMED | S74 | 0.125 | 92.86 | 427 | 0.099364 | 0.125 | 0.0994 | 1 |
| O14964 | HGS | KykVVQDtyQIMkVE | T131 | 0.064 | 95.15 | 777 | 0.043713 | 0.064 | 0.0437 | 1 |
| O14964 | HGS | ykVVQDtyQIMkVEG | Y132 | 0.226 | 95.41 | 777 | 0.099143 | 0.226 | 0.0991 | 1 |
| Q09472 | EP300 | SANNRAEyyHLLAEk | Y629 | 0 | 94.65 | 2414 | 0 | 0.332 | 0.1217 | 1 |
| O95870 | ABHD16A | NAEQLCRyQGPVLLI | Y418 | 0.025 | 97.62 | 558 | 0.010967 | 0.025 | 0.011 | 0.9333 |
| O95870 | ABHD16A | TVPEDIMsNRGNDLL | S443 | 0.118 | 96.87 | 558 | 0.0938 | 0.118 | 0.0938 | 1 |
| Q14896 | MYBPC3 | RRVHSGTyQVTVRIE | Y1043 | 0.06 | 94.5 | 1274 | 0.026321 | 0.06 | 0.0263 | 1 |
| Q14896 | MYBPC3 | ELIIGNGyyFRVFSQ | Y1135 | 0.144 | 91.45 | 1274 | 0.063171 | 0.245 | 0.1075 | 1 |
| Q5VVC0 | SPO16 | KEKIKWttTIIISSS | T13 | 0.061 | 86.55 | 180 | 0.041664 | 0.061 | 0.0417 | 1 |
| Q6PL18 | ATAD2 | EAKRtAPsIVyVPHI | S855 | 0.002 | 89.03 | 1390 | 0.00159 | 0.034 | 0.027 | 1 |
| Q6PL18 | ATAD2 | RtAPsIVyVPHIHVW | Y858 | 0.16 | 94.21 | 1390 | 0.07019 | 0.16 | 0.0702 | 1 |
| Q9H2X6 | HIPK2 | tyLQsRYyRAPEIIL | Y367 | 0.146 | 98.75 | 1198 | 0.064048 | 0.146 | 0.064 | 1 |
| F5H284 | PPIAL4D | sVVFFEItRDGKPLG | T11 | 0.172 | 98.54 | 164 | 0.117478 | 0.172 | 0.1175 | 1 |
| Q9UNL2 | SSR3 | YILSISAsSGLIALL | S173 | 0 | 95 | 185 | 0 | 0 | 0 | 1 |
| Q6GTS8 | PM20D1 | KGAIQIPtVtFSSEK | T63 | 0.001 | 98.06 | 502 | 0.000683 | 0.001 | 0.0007 | 1 |
| Q6GTS8 | PM20D1 | AIQIPtVtFSSEKSN | T65 | 0.049 | 95.82 | 502 | 0.033468 | 0.049 | 0.0335 | 1 |
| P49247 | RPIA | NQVLGIGsGsTIVHA | S104 | 0.15 | 97.63 | 311 | 0.119237 | 0.15 | 0.1192 | 0.8889 |
| Q9Y4A5 | TRRAP | VRKQAMHsLQVLAQV | S1263 | 0.011 | 96.67 | 3859 | 0.008744 | 0.011 | 0.0087 | 1 |

| | | | | | | | | | | |
|---|---|---|---|---|---|---|---|---|---|---|
| Q17RR3 | PNLIPRP3 | KLSGSEVtQGTVFLR | T369 | 0.031 | 92.58 | 467 | 0.021173 | 0.031 | 0.0212 | 0.8571 |
| P23434 | GCSH | SELYSPLsGEVTEIN | S117 | 0.181 | 98.38 | 173 | 0.14388 | 0.181 | 0.1439 | 1 |
| Q9NUB1 | ACSS1 | CAPESMGsEDMLFML | S285 | 0.039 | 98.79 | 689 | 0.031002 | 0.039 | 0.031 | 1 |
| Q9NUB1 | ACSS1 | HQRFVDAyFkAYPGY | Y521 | 0.144 | 95.03 | 689 | 0.063171 | 0.144 | 0.0632 | 0.9167 |
| Q96C24 | SYTL4 | AAKAGGtsDSFVKGy | S547 | 0 | 85.49 | 671 | 0 | 0 | 0 | 1 |
| Q96C24 | SYTL4 | LPMRNKAsKRKtPVM | S563 | 0.091 | 78.37 | 671 | 0.072337 | 0.091 | 0.0723 | 0.8571 |
| Q9BZG8 | DPH1 | GEILGCTsPRLSkEV | S216 | 0.102 | 96.78 | 443 | 0.081081 | 0.102 | 0.0811 | 1 |
| Q9BZG8 | DPH1 | PNVPAYRyDPYSkVL | Y251 | 0.016 | 98.65 | 443 | 0.007019 | 0.26 | 0.1141 | 1 |
| Q2VWP7 | PRTG | RLQHAGVyVCRATtP | Y302 | 0.03 | 95.84 | 1150 | 0.013161 | 0.03 | 0.0132 | 1 |
| Q2VWP7 | PRTG | VyVCRATtPGTRNFT | T308 | 0.008 | 91.53 | 1150 | 0.005464 | 0.008 | 0.0055 | 1 |
| A8MVW5 | HEPACAM2 | SFSPQNNtLHIAPVt | T203 | 0.168 | 98.53 | 462 | 0.114746 | 0.168 | 0.1147 | 1 |
| A8MVW5 | HEPACAM2 | tLHIAPVtKEDIGNY | T210 | 0.18 | 96.71 | 462 | 0.122942 | 0.18 | 0.1229 | 1 |
| Q7Z3T8 | ZFYVE16 | LGAEykAyPAPLtsI | Y1204 | 0.164 | 93.55 | 1539 | 0.071944 | 0.164 | 0.0719 | 1 |
| Q7Z3T8 | ZFYVE16 | AyPAPLtsIRGRKPL | S1210 | 0.001 | 96.66 | 1539 | 0.000795 | 0.001 | 0.0008 | 1 |
| Q99715 | COL12A1 | NVGVEVFsLGIKAAD | S273 | 0.01 | 93.785 | 3063 | 0.007949 | 0.01 | 0.0079 | 1 |
| P63104 | YWHAZ | kkGIVDQsQQAyQEA | S145 | 0 | 98.6 | 245 | 0 | 0 | 0 | 1 |
| P63104 | YWHAZ | VDQsQQAyQEAFEIS | Y149 | 0 | 98.85 | 245 | 0 | 0 | 0 | 1 |
| P61962 | DCAF7 | HGKRKEIyKyEAPWT | Y11 | 0.109 | 96.83 | 342 | 0.047817 | 0.305 | 0.1338 | 0.8 |
| P61962 | DCAF7 | KRKEIyKyEAPWTVY | Y13 | 0.254 | 97.93 | 342 | 0.111426 | 0.254 | 0.1114 | 1 |
| P61962 | DCAF7 | RFRLALGsFVEEYNN | S38 | 0.007 | 97.68 | 342 | 0.005564 | 0.007 | 0.0056 | 1 |
| P61962 | DCAF7 | TFDHPyPtTkLMWIP | T71 | 0.023 | 97.68 | 342 | 0.015709 | 0.023 | 0.0157 | 1 |
| P61962 | DCAF7 | ASVGADGsVRMFDLR | S200 | 0.029 | 98.31 | 342 | 0.023053 | 0.051 | 0.0405 | 1 |
| P61962 | DCAF7 | LEHSTIIyEDPQHHP | Y216 | 0.05 | 98.11 | 342 | 0.021934 | 0.05 | 0.0219 | 1 |
| Q9GZU3 | TMEM39B | MKEVLVSsMLsAYYV | S293 | 0.003 | 93.6 | 492 | 0.002385 | 0.003 | 0.0024 | 1 |
| Q9GZU3 | TMEM39B | CELFLLVsIsTsVIL | S327 | 0.148 | 94.96 | 492 | 0.117647 | 0.148 | 0.1176 | 1 |
| Q9HAT8 | PELI2 | ISCkGQHsIsyTLsR | S78 | 0.024 | 95.22 | 420 | 0.019078 | 0.097 | 0.0938 | 0.8889 |
| O00308 | WWP2 | TGFTLPFykRMLNKR | Y631 | 0.005 | 92.67 | 870 | 0.002193 | 0.005 | 0.0022 | 1 |
| Q96A23 | CPNE4 | GDDGILRsPKGEPVL | S492 | 0.106 | 94.51 | 557 | 0.084261 | 0.071 | 0.1067 | 1 |
| O43586 | PSTPIP1 | LYkKAMEsKKTYEQK | S140 | 0.123 | 98.31 | 416 | 0.097775 | 0.123 | 0.0978 | 1 |
| O43586 | PSTPIP1 | HQKQVEKsQNKARQC | S173 | 0.121 | 92.31 | 416 | 0.096185 | 0.121 | 0.0962 | 1 |
| O43586 | PSTPIP1 | WEQEHRttCEAFQLQ | T212 | 0.026 | 98.43 | 416 | 0.017758 | 0.026 | 0.0178 | 1 |

| | | | | | | | | | | |
|---|---|---|---|---|---|---|---|---|---|---|
| O00422 | SAP18 | PEARKKGtHFNFAIV | T85 | 0.022 | 96.57 | 153 | 0.015026 | 0.022 | 0.015 | 1 |
| O14709 | ZNF197 | RQLRyHEtsGPQEAL | T53 | 0.19 | 91.79 | 1029 | 0.129773 | 0.19 | 0.1298 | 1 |
| Q9UBC5 | MYO1A | yALANVAyQSLRDRD | Y85 | 0.091 | 90.17 | 1043 | 0.03992 | 0.091 | 0.0399 | 1 |
| Q99558 | MAP3K14 | VSSGQAHsLtSLAKT | S352 | 0.114 | 92.27 | 947 | 0.09062 | 0.114 | 0.0906 | 1 |
| Q96ME1 | FBXL18 | ATKNLLDsMARNVVL | S285 | 0.178 | 90.09 | 805 | 0.141495 | 0.178 | 0.1415 | 1 |
| Q15238 | PSG5 | tRNETGPyECEIRDR | Y215 | 0.019 | 94.53 | 335 | 0.008335 | 0.019 | 0.0083 | 1 |
| Q9Y646 | CPQ | AKVGALAsLIRSVAS | S202 | 0 | 98.91 | 472 | 0 | 0 | 0 | 1 |
| Q9Y646 | CPQ | PEQVVLVsGHLDsWD | S288 | 0 | 98.69 | 472 | 0 | 0 | 0 | 1 |
| Q9Y646 | CPQ | LVsGHLDsWDVGQGA | S293 | 0.014 | 98.5 | 472 | 0.011129 | 0.014 | 0.0111 | 0.9286 |
| Q11130 | FUT7 | LVPTVAQyRFYLSFE | Y222 | 0.016 | 98.67 | 342 | 0.007019 | 0.016 | 0.007 | 1 |
| P05387 | RPLP2 | ______MRyVAsyLLA | Y3 | 0.337 | 91.08 | 115 | 0.147837 | 0.337 | 0.1478 | 1 |
| P05387 | RPLP2 | __MRyVAsyLLAALG | S6 | 0.037 | 91.23 | 115 | 0.029412 | 0.037 | 0.0294 | 1 |
| P05387 | RPLP2 | _MRyVAsyLLAALGG | Y7 | 0.044 | 91.25 | 115 | 0.019302 | 0.044 | 0.0193 | 0.8667 |
| Q68E01 | INTS3 | IHNPQALsPQFTGIL | S283 | 0.061 | 96.08 | 1043 | 0.04849 | 0.061 | 0.0485 | 1 |
| P49674 | CSNK1E | RFDDkPDySYLRQLF | Y266 | 0.114 | 98.78 | 416 | 0.05001 | 0.114 | 0.05 | 1 |
| O43776 | NARS1 | LKARsMVtRCFRDHF | T253 | 0.212 | 98.86 | 548 | 0.144799 | 0.212 | 0.1448 | 1 |
| O75355 | ENTPD3 | YTHSFQCyGRNEAEK | Y262 | 0.121 | 98.37 | 529 | 0.053081 | 0.121 | 0.0531 | 1 |
| P23468 | PTPRD | RyANVIAyDHsRVLL | Y1392 | 0.247 | 95.82 | 1912 | 0.108355 | 0.392 | 0.1356 | 1 |
| P23468 | PTPRD | RLVNIMPyEStRVCL | Y1681 | 0.088 | 89.99 | 1912 | 0.038604 | 0.231 | 0.1013 | 0.8571 |
| P23468 | PTPRD | GREKCHQyWPAERSA | Y1757 | 0.025 | 96.2 | 1912 | 0.010967 | 0.025 | 0.011 | 1 |
| P23468 | PTPRD | AEYNMPQyILREFkV | Y1782 | 0.196 | 94.55 | 1912 | 0.085982 | 0.196 | 0.086 | 1 |
| O95486 | SEC24A | NPELFRCtLtsIPQt | T389 | 0 | 98.64 | 1093 | 0 | 0 | 0 | 1 |
| O95486 | SEC24A | ELFRCtLtsIPQtQA | T391 | 0.107 | 97.88 | 1093 | 0.073082 | 0.107 | 0.0731 | 1 |
| O95486 | SEC24A | LFRCtLtsIPQtQAL | S392 | 0.096 | 97.62 | 1093 | 0.076312 | 0.096 | 0.0763 | 1 |
| Q8NBQ5 | HSD17B11 | FLLAYCSsKFAAVGF | S188 | 0 | 98.43 | 300 | 0 | 0 | 0 | 1 |
| Q9NUQ7 | UFSP2 | IyVVQGIyGyHHYMQ | Y285 | 0 | 98.6 | 469 | 0 | 0 | 0 | 1 |
| Q9NUQ7 | UFSP2 | VVQGIyGyHHYMQDR | Y287 | 0.005 | 98.68 | 469 | 0.002193 | 0.005 | 0.0022 | 1 |
| P08758 | ANXA5 | GtNEkVLtEIIAsRt | T111 | 0.096 | 98.46 | 320 | 0.065569 | 0.192 | 0.1311 | 0.8182 |
| P08758 | ANXA5 | LRAIkQVyEEEyGss | Y129 | 0 | 98.57 | 320 | 0 | 0 | 0 | 1 |
| O43660 | PLRG1 | HYTFAsGsPDNIkQW | S391 | 0.001 | 95.97 | 514 | 0.000795 | 0.001 | 0.0008 | 1 |
| P51692 | STAT5B | LkIkLGHyAtQLQNT | Y90 | 0.284 | 95.21 | 787 | 0.124587 | 0.284 | 0.1246 | 0.875 |

| | | | | | | | | | | |
|---|---|---|---|---|---|---|---|---|---|---|
| Q14112 | NID2 | PFLADIDtsHGRGRV | T115 | 0.046 | 89.8 | 1375 | 0.031419 | 0.046 | 0.0314 | 0.9167 |
| Q9H6Y2 | WDR55 | RFALRAEsIDCMVPV | S255 | 0.069 | 97.68 | 383 | 0.054849 | 0.069 | 0.0548 | 1 |
| O94779 | CNTN5 | SQETGNLyISKVQTS | Y255 | 0.14 | 97.38 | 1100 | 0.061416 | 0.14 | 0.0614 | 1 |
| O94779 | CNTN5 | QTSDVGSyICLVKNT | Y267 | 0.068 | 96.06 | 1100 | 0.029831 | 0.068 | 0.0298 | 1 |
| Q8TDI0 | CHD5 | DKKGDVHyLIKWKDL | Y610 | 0.066 | 92.08 | 1954 | 0.028953 | 0.066 | 0.029 | 1 |
| P12235 | SLC25A4 | GGVAAAVsktAVAPI | S22 | 0.099 | 92.3 | 298 | 0.078697 | 0.099 | 0.0787 | 1 |
| P12235 | SLC25A4 | AFFkGAWsNVLRGMG | S276 | 0.018 | 91.67 | 298 | 0.014308 | 0.018 | 0.0143 | 0.8571 |
| Q9H0B6 | KLC2 | KAEEVEyyyRRALEI | Y346 | 0.29 | 97.23 | 622 | 0.127219 | 0.29 | 0.1272 | 1 |
| Q9H0B6 | KLC2 | AEEVEyyyRRALEIY | Y347 | 0.069 | 97.01 | 622 | 0.030269 | 0.069 | 0.0303 | 1 |
| Q9Y6W3 | CAPN7 | RWGkLPLsPkQKTTF | S251 | 0.102 | 93 | 813 | 0.081081 | 0.102 | 0.0811 | 1 |
| Q9Y6W3 | CAPN7 | HLRWKGRysENDVkN | Y489 | 0.127 | 96.01 | 813 | 0.055713 | 0.127 | 0.0557 | 1 |
| Q9Y6W3 | CAPN7 | LRWKGRysENDVkNW | S490 | 0.028 | 95.54 | 813 | 0.022258 | 0.028 | 0.0223 | 1 |
| Q9Y6W3 | CAPN7 | KTDGKkVyyPADPPP | Y619 | 0.084 | 94.99 | 813 | 0.03685 | 0.084 | 0.0369 | 1 |
| Q9Y6W3 | CAPN7 | TDGKkVyyPADPPPy | Y620 | 0.173 | 95.01 | 813 | 0.075893 | 0.173 | 0.0759 | 1 |
| Q9Y6W3 | CAPN7 | IDGIRINsPHYLTkI | S635 | 0.158 | 88.35 | 813 | 0.125597 | 0.158 | 0.1256 | 1 |
| Q9UL51 | HCN2 | GtIGKKMyFIQHGVV | Y582 | 0.085 | 97.13 | 889 | 0.037288 | 0.085 | 0.0373 | 1 |
| P27797 | CALR | FGkFVLssGkFyGDE | S53 | 0.143 | 97.42 | 417 | 0.113673 | 0.143 | 0.1137 | 1 |
| P27797 | CALR | TSQDArFyALsAsFE | Y75 | 0.004 | 98.28 | 417 | 0.001755 | 0.004 | 0.0018 | 1 |
| P27797 | CALR | DArFyALsAsFEPFs | S78 | 0.001 | 98.51 | 417 | 0.000795 | 0.001 | 0.0008 | 1 |
| P27797 | CALR | DDEFtHLyTLIVRPD | Y172 | 0.028 | 98.81 | 417 | 0.012283 | 0.028 | 0.0123 | 1 |
| P27797 | CALR | IVRPDNtyEVKIDNS | Y182 | 0 | 98.64 | 417 | 0 | 0 | 0 | 1 |
| P27797 | CALR | IFDNFLItNDEAYAE | T333 | 0.001 | 98.62 | 417 | 0.000683 | 0.001 | 0.0007 | 1 |
| Q9Y5X9 | LIPG | KGtVGRItGLDPAGP | T190 | 0.003 | 98.88 | 500 | 0.002049 | 0.003 | 0.002 | 1 |
| Q9Y5X9 | LIPG | kMRNKRNsKMYLKtR | S332 | 0.014 | 91.35 | 500 | 0.011129 | 0.014 | 0.0111 | 0.8333 |
| P25445 | FAS | PCQEGKEyTDKAHFS | Y91 | 0.167 | 98.21 | 335 | 0.07326 | 0.208 | 0.0912 | 1 |
| Q9BQL6 | FERMT1 | MNPECFVsPRCAKRH | S520 | 0.065 | 88.55 | 677 | 0.051669 | 0.065 | 0.0517 | 1 |
| Q9HAY6 | BCO1 | IVVsEFGtMAyPDPC | T95 | 0.079 | 92.74 | 547 | 0.053958 | 0.079 | 0.054 | 1 |
| Q9HAY6 | BCO1 | IPTKIIKyDILtKsS | Y438 | 0 | 98.78 | 547 | 0 | 0 | 0 | 1 |
| Q9NYY1 | IL20 | LLQWMEEtE_______ | T175 | 0.008 | 87.01 | 176 | 0.005464 | 0.008 | 0.0055 | 1 |
| O75165 | DNAJC13 | IRVIHALsENELCVR | S2077 | 0.002 | 88.42 | 2243 | 0.00159 | 0.002 | 0.0016 | 1 |
| Q68CK6 | ACSM2B | ILGSLLEsWtLGACT | S276 | 0 | 97.94 | 577 | 0 | 0 | 0 | 1 |

| | | | | | | | | | | |
|---|---|---|---|---|---|---|---|---|---|---|
| P68133 | ACTA1 | APrAVFPsIVGRPRH | S35 | 0.001 | 97.85 | 377 | 0.000795 | 0.001 | 0.0008 | 1 |
| P68133 | ACTA1 | GMGQkDsyVGDEAQs | Y55 | 0.034 | 96.04 | 377 | 0.014915 | 0.15 | 0.0658 | 1 |
| P68133 | ACTA1 | MEkIWHHtFyNELRV | T91 | 0.002 | 98.65 | 377 | 0.001366 | 0.002 | 0.0014 | 0.8889 |
| P68133 | ACTA1 | DLAGRDLtDyLMkIL | T188 | 0.015 | 98.1 | 377 | 0.010245 | 0.015 | 0.0102 | 0.9 |
| P68133 | ACTA1 | ErFRCPEtLFQPsFI | T262 | 0.006 | 97.95 | 377 | 0.004098 | 0.006 | 0.0041 | 0.8889 |
| P68133 | ACTA1 | IDIRKDLyANNVMSG | Y296 | 0.005 | 98.26 | 377 | 0.002193 | 0.005 | 0.0022 | 1 |
| P68133 | ACTA1 | NVMSGGttMyPGIAD | T306 | 0.014 | 98.45 | 377 | 0.009562 | 0.014 | 0.0096 | 0.8889 |
| P68133 | ACTA1 | MWITKQEyDEAGPsI | Y364 | 0.167 | 98.3 | 377 | 0.07326 | 0.332 | 0.1452 | 1 |
| Q92824 | PCSK5 | MGRRGLGsVFVWASG | S305 | 0.006 | 97 | 1860 | 0.004769 | 0.006 | 0.0048 | 1 |
| Q07817 | BCL2L1 | GRIVAFFsFGGALCV | S145 | 0 | 93.01 | 233 | 0 | 0.052 | 0.0413 | 0.8889 |
| Q9H4D0 | CLSTN2 | FPDIKIVstFAKtEA | S666 | 0.023 | 93.34 | 955 | 0.018283 | 0.023 | 0.0183 | 1 |
| Q9H4D0 | CLSTN2 | PDIKIVstFAKtEAP | T667 | 0.181 | 91.18 | 955 | 0.123625 | 0.181 | 0.1236 | 1 |
| Q8WWQ0 | PHIP | RHDNtVItAVNNMtL | T441 | 0 | 98.13 | 1821 | 0 | 0 | 0 | 1 |
| Q8WWQ0 | PHIP | ItAVNNMtLKVWNSY | T447 | 0.166 | 92.58 | 1821 | 0.11338 | 0.202 | 0.1234 | 1 |
| Q05655 | PRKCD | EkARLsysDKNLIDs | S647 | 0.063 | 66.44 | 676 | 0.05008 | 0.063 | 0.0501 | 1 |
| P78563 | ADARB1 | VKDAKVIsVSTGtKC | S370 | 0 | 98.74 | 741 | 0 | 0 | 0 | 1 |
| P78563 | ADARB1 | KRsIFQksERGGFRL | S429 | 0.158 | 96.3 | 741 | 0.125597 | 0.158 | 0.1256 | 0.8333 |
| P78563 | ADARB1 | SKLAAKEyQAAKARL | Y708 | 0.18 | 98.5 | 741 | 0.078963 | 0.18 | 0.079 | 0.9231 |
| P04049 | RAF1 | IVLYELMtGELPYSH | T543 | 0.127 | 95.18 | 648 | 0.086743 | 0.127 | 0.0867 | 1 |
| Q16740 | CLPP | MRHsLPNsRIMIHQP | S173 | 0.077 | 98.39 | 277 | 0.061208 | 0.077 | 0.0612 | 1 |
| P20839 | IMPDH1 | ITDPVVLsPSHTVGD | S122 | 0.071 | 97.49 | 514 | 0.056439 | 0.071 | 0.0564 | 1 |
| P20839 | IMPDH1 | LCGAAVGtREDDKYR | T252 | 0.145 | 96.63 | 514 | 0.099037 | 0.145 | 0.099 | 1 |
| P20839 | IMPDH1 | KkYRGMGsLDAMEkS | S416 | 0.023 | 87.47 | 514 | 0.018283 | 0.023 | 0.0183 | 1 |
| P20839 | IMPDH1 | SSSSQkRyFsEGDkV | Y430 | 0.285 | 84.55 | 514 | 0.125025 | 0.285 | 0.125 | 1 |
| P20839 | IMPDH1 | SSQkRyFsEGDkVkI | S432 | 0.063 | 76.72 | 514 | 0.05008 | 0.063 | 0.0501 | 1 |
| Q9HC10 | OTOF | FDKIIKIsVIHsKNL | S320 | 0.042 | 95.03 | 1997 | 0.033386 | 0.042 | 0.0334 | 0.8571 |
| Q9NZN4 | EHD2 | VTKDKskyDEIFyNL | Y453 | 0.036 | 95.84 | 543 | 0.015793 | 0.036 | 0.0158 | 0.8182 |
| Q6ZW31 | SYDE1 | ITGILKDyLRELPtP | Y478 | 0.005 | 97.08 | 735 | 0.002193 | 0.005 | 0.0022 | 1 |
| Q4VNC0 | ATP13A5 | LSEARIRtVMITGDN | T715 | 0.155 | 94.26 | 1218 | 0.105867 | 0.155 | 0.1059 | 1 |
| Q96GD4 | AURKB | HPWVRANsRRVLPPS | S331 | 0.165 | 98.01 | 344 | 0.131161 | 0.165 | 0.1312 | 1 |
| P21333 | FLNA | GPyHIGGsPFkAKVT | S2510 | 0.032 | 70.81 | 2647 | 0.025437 | 0.102 | 0.0811 | 0.8333 |

| | | | | | | | | | | |
|---|---|---|---|---|---|---|---|---|---|---|
| Q14318 | FKBP8 | CLNNLAAsQLkLDHy | S281 | 0 | 98.4 | 412 | 0 | 0 | 0 | 1 |
| Q13813 | SPTAN1 | WsQLLANsAARKKkL | S2072 | 0.021 | 78.56 | 2472 | 0.016693 | 0.021 | 0.0167 | 1 |
| O15305 | PMM2 | RNGMLNVsPIGRSCs | S130 | 0 | 98.64 | 246 | 0 | 0 | 0 | 1 |
| P13639 | EEF2 | PVVSyREtVsEEsNV | T582 | 0.098 | 97.58 | 858 | 0.066935 | 0.098 | 0.0669 | 1 |
| P13639 | EEF2 | REtVsEEsNVLCLsk | S587 | 0.035 | 98.25 | 858 | 0.027822 | 0.035 | 0.0278 | 1 |
| P13639 | EEF2 | EsNVLCLsksPNkHN | S593 | 0 | 97.52 | 858 | 0 | 0 | 0 | 1 |
| P13639 | EEF2 | NVLCLsksPNkHNRL | S595 | 0.048 | 95.83 | 858 | 0.038156 | 0.048 | 0.0382 | 1 |
| P13639 | EEF2 | PtArRCLyAsVLTAQ | Y730 | 0.14 | 98.4 | 858 | 0.061416 | 0.183 | 0.0803 | 1 |
| P13639 | EEF2 | ArRCLyAsVLTAQPr | S732 | 0 | 98.5 | 858 | 0 | 0 | 0 | 1 |
| Q8N5L8 | RPP25L | PAVWVLLsRDPLDPN | S122 | 0 | 97.69 | 163 | 0 | 0 | 0 | 1 |
| Q96JA3 | PLEKHA8 | TNKEEFttLQkIVLH | T381 | 0.094 | 98.28 | 519 | 0.064203 | 0.094 | 0.0642 | 1 |
| Q96JA3 | PLEKHA8 | QtALNNAyGKTLRQH | Y437 | 0.091 | 95.31 | 519 | 0.03992 | 0.091 | 0.0399 | 1 |
| P45974 | USP5 | EALLsADsASRkQEV | S159 | 0.067 | 77.06 | 858 | 0.053259 | 0.067 | 0.0533 | 0.8333 |
| Q96PE3 | INPP4A | ILEQMAEsVLSLHVP | S252 | 0 | 94.41 | 977 | 0 | 0 | 0 | 1 |
| Q96PE3 | INPP4A | DGGSDQNyDIVTIGA | Y355 | 0.312 | 94.49 | 977 | 0.13687 | 0.312 | 0.1369 | 1 |
| Q96PE3 | INPP4A | EGCRRENtMKNVGSR | T942 | 0 | 93.03 | 977 | 0 | 0 | 0 | 1 |
| Q96PE3 | INPP4A | KNVGSRKyAFNsLQL | Y951 | 0.001 | 92.14 | 977 | 0.000439 | 0.001 | 0.0004 | 1 |
| Q96PE3 | INPP4A | LkAFPkHyRPPEGTy | Y965 | 0.033 | 94.99 | 977 | 0.014477 | 0.033 | 0.0145 | 1 |
| P12931 | SRC | TFVALyDyEsRTETD | Y95 | 0.062 | 96.86 | 536 | 0.027199 | 0.062 | 0.0272 | 0.9286 |
| P12931 | SRC | tGyIPSNyVAPSDSI | Y139 | 0.101 | 95.82 | 536 | 0.044307 | 0.101 | 0.0443 | 1 |
| P12931 | SRC | NAENPRGtFLVRESE | T174 | 0.005 | 98.43 | 536 | 0.003415 | 0.005 | 0.0034 | 1 |
| P12931 | SRC | LQQLVAyySKHADGL | Y233 | 0 | 97.77 | 536 | 0 | 0 | 0 | 1 |
| P12931 | SRC | FGEVWMGtWNGTTRV | T288 | 0.164 | 98.07 | 536 | 0.112014 | 0.195 | 0.1332 | 1 |
| P12931 | SRC | IkTLKPGtMsPEAFL | T304 | 0.133 | 85.82 | 536 | 0.090841 | 0.133 | 0.0908 | 1 |
| P12931 | SRC | QIASGMAyVERMNyV | Y379 | 0.189 | 97.48 | 536 | 0.082912 | 0.253 | 0.111 | 1 |
| Q96NJ5 | KLHL32 | GAMEEyLyAVGGRNE | Y401 | 0.023 | 98.28 | 620 | 0.01009 | 0.023 | 0.0101 | 1 |
| Q16775 | HAGH | VYCGHEytINNLkFA | T224 | 0.011 | 98.76 | 308 | 0.007513 | 0.011 | 0.0075 | 0.9091 |
| Q15293 | RCN1 | ArHLVyEsDkNKDEK | S292 | 0.027 | 92.41 | 331 | 0.021463 | 0.027 | 0.0215 | 1 |
| P62081 | RPS7 | LEDLVFPsEIVGkRI | S137 | 0.069 | 91.29 | 194 | 0.054849 | 0.123 | 0.1031 | 1 |
| Q14831 | GRM7 | LGWNYVStLAsEGsY | T223 | 0.006 | 97.41 | 915 | 0.004098 | 0.006 | 0.0041 | 1 |
| Q14831 | GRM7 | NYVStLAsEGsYGEK | S226 | 0.034 | 91.49 | 915 | 0.027027 | 0.034 | 0.027 | 1 |

| | | | | | | | | | | |
|---|---|---|---|---|---|---|---|---|---|---|
| Q14831 | GRM7 | ISYAALLtKtNRIYR | T675 | 0.069 | 93.66 | 915 | 0.047128 | 0.069 | 0.0471 | 1 |
| Q14831 | GRM7 | YAALLtKtNRIYRIF | T677 | 0.018 | 90.43 | 915 | 0.012294 | 0.204 | 0.1393 | 1 |
| Q86VI1 | EXOC3L1 | sDALIRFsRDHFRGK | S471 | 0.066 | 94.2 | 746 | 0.052464 | 0.066 | 0.0525 | 1 |
| Q86VI1 | EXOC3L1 | ERAVVLQyLSALMQG | Y597 | 0.005 | 94.98 | 746 | 0.002193 | 0.005 | 0.0022 | 1 |
| Q6IA69 | NADSYN1 | NLQRILksIEIAKNR | S29 | 0.002 | 98.93 | 706 | 0.00159 | 0.002 | 0.0016 | 1 |
| Q6IA69 | NADSYN1 | VDLVtMVtSKNGGIy | T220 | 0.033 | 97.86 | 706 | 0.022539 | 0.043 | 0.035 | 1 |
| Q6IA69 | NADSYN1 | tSKNGGIyLLANQKG | Y227 | 0.023 | 98.88 | 706 | 0.01009 | 0.023 | 0.0101 | 1 |
| Q6IA69 | NADSYN1 | SkNssQEtCtRAREL | T427 | 0.143 | 93.43 | 706 | 0.097671 | 0.143 | 0.0977 | 1 |
| Q6IA69 | NADSYN1 | VKRFFSkysMNRHkM | Y643 | 0.046 | 98.43 | 706 | 0.02018 | 0.046 | 0.0202 | 1 |
| Q6IA69 | NADSYN1 | KRFFSkysMNRHkMT | S644 | 0.032 | 98.46 | 706 | 0.025437 | 0.032 | 0.0254 | 1 |
| Q6IA69 | NADSYN1 | MTTLTPAyHAENYSP | Y657 | 0.028 | 98.81 | 706 | 0.012283 | 0.028 | 0.0123 | 1 |
| P35858 | IGFALS | LGRLRyLsLRNNsLR | S512 | 0 | 97.27 | 605 | 0 | 0 | 0 | 0.8 |
| Q86XJ1 | GAS2L3 | EYLSEGRyRLGDKIL | Y242 | 0.052 | 96.42 | 694 | 0.022812 | 0.052 | 0.0228 | 1 |
| Q05707 | COL14A1 | EYKISVytKLQEIEG | T989 | 0.001 | 90.53 | 1796 | 0.000683 | 0.001 | 0.0007 | 1 |
| Q16572 | SLC18A3 | GLGVMFAstVLFAFA | S165 | 0.007 | 96.83 | 532 | 0.005564 | 0.007 | 0.0056 | 1 |
| Q16572 | SLC18A3 | LGVMFAstVLFAFAE | T166 | 0.007 | 95.16 | 532 | 0.004781 | 0.176 | 0.1202 | 0.8333 |
| Q9NRR6 | INPP5E | VAERLLDyTRTVQAL | Y439 | 0.125 | 98.65 | 644 | 0.054836 | 0.125 | 0.0548 | 1 |
| Q6DN12 | MCTP2 | RKQItQRyCLQNSLK | Y496 | 0.141 | 76.34 | 878 | 0.061855 | 0.183 | 0.0803 | 1 |
| Q9Y285 | FARSA | ALQLPMDyVQRVkRt | Y292 | 0.004 | 97.01 | 508 | 0.001755 | 0.004 | 0.0018 | 1 |
| Q9Y285 | FARSA | yVQRVkRtHsQGGyG | T299 | 0.02 | 97.09 | 508 | 0.01366 | 0.02 | 0.0137 | 1 |
| Q9Y285 | FARSA | NLQMVyDsPLCRLDA | S490 | 0.113 | 78.17 | 508 | 0.089825 | 0.129 | 0.1025 | 1 |
| P18440 | NAT1 | SkYRkIYsFTLKPRT | S191 | 0.044 | 98.91 | 290 | 0.034976 | 0.044 | 0.035 | 1 |
| P18440 | NAT1 | sSVFTSKsFCSLQtP | S221 | 0.029 | 98.87 | 290 | 0.023053 | 0.029 | 0.0231 | 1 |
| P18440 | NAT1 | KsFCSLQtPDGVHCL | T227 | 0.172 | 98.85 | 290 | 0.117478 | 0.172 | 0.1175 | 0.8333 |
| Q9Y617 | PSAT1 | IQkELLDykGVGIsV | Y32 | 0.193 | 97.8 | 370 | 0.084666 | 0.193 | 0.0847 | 1 |
| Q9Y617 | PSAT1 | DykGVGIsVLEMSHr | S38 | 0.065 | 98.04 | 370 | 0.051669 | 0.065 | 0.0517 | 1 |
| Q9Y617 | PSAT1 | SVGGIRAsLyNAVTI | S344 | 0.006 | 98.73 | 370 | 0.004769 | 0.006 | 0.0048 | 0.8889 |
| O14976 | GAK | HYAVYNLsPRTYRPS | S456 | 0.031 | 90.96 | 1311 | 0.024642 | 0.031 | 0.0246 | 0.8333 |
| P34925 | RYK | QAsEIQVtMMLtEsC | T376 | 0.078 | 95.68 | 607 | 0.053275 | 0.102 | 0.0697 | 1 |
| P34925 | RYK | VtMMLtEsCKLRGLH | S382 | 0.02 | 96.39 | 607 | 0.015898 | 0.02 | 0.0159 | 1 |
| Q9UKQ9 | KLK9 | IsGWGAVssPKALFP | S152 | 0.044 | 91.66 | 250 | 0.034976 | 0.044 | 0.035 | 0.875 |

| | | | | | | | | | | |
|---|---|---|---|---|---|---|---|---|---|---|
| Q9UKQ9 | KLK9 | sGWGAVssPKALFPV | S153 | 0.021 | 90.72 | 250 | 0.016693 | 0.021 | 0.0167 | 1 |
| Q15269 | PWP2 | SDTLPLAtRYNVkCV | T54 | 0.048 | 94.68 | 919 | 0.032785 | 0.096 | 0.0656 | 1 |
| Q15269 | PWP2 | DGRKFVVtkGNIAQM | T114 | 0 | 94.44 | 919 | 0 | 0 | 0 | 1 |
| Q15269 | PWP2 | PMKSVLAsASWDkTV | S518 | 0 | 97.29 | 919 | 0 | 0 | 0 | 1 |
| Q15269 | PWP2 | MSKFVCIyHVREQIL | Y632 | 0.023 | 95.48 | 919 | 0.01009 | 0.023 | 0.0101 | 1 |
| Q15269 | PWP2 | kPEIRVTsLRFsPTG | S707 | 0.122 | 96.36 | 919 | 0.09698 | 0.122 | 0.097 | 1 |
| P00533 | EGFR | IIRGNMyyENSyALA | Y113 | 0.171 | 91.46 | 1210 | 0.075015 | 0.171 | 0.075 | 0.8182 |
| P00533 | EGFR | NMyyENSyALAVLSN | Y117 | 0.341 | 96.36 | 1210 | 0.149592 | 0.341 | 0.1496 | 1 |
| P49792 | RANBP2 | IAsVQGstPsPRQKs | T19 | 0.094 | 71.86 | 3224 | 0.064203 | 0.153 | 0.1006 | 1 |
| P49792 | RANBP2 | GFYFAkLyYEAkEyD | Y36 | 0.098 | 92.77 | 3224 | 0.042991 | 0.098 | 0.043 | 1 |
| P00451 | F8 | KKVRFMAytDETFKT | Y450 | 0.162 | 94.22 | 2351 | 0.071067 | 0.314 | 0.1377 | 1 |
| Q9UK58 | CCNL1 | PEERLsPtPSMQDGL | T67 | 0.02 | 92.88 | 526 | 0.01366 | 0.02 | 0.0137 | 1 |
| O75452 | RDH16 | LSHLRDKyVFITGCD | Y31 | 0.258 | 98.77 | 317 | 0.113181 | 0.258 | 0.1132 | 1 |
| P19801 | AOC1 | tttMAKNtVFLIEML | T68 | 0.11 | 98.73 | 751 | 0.075131 | 0.215 | 0.1463 | 1 |
| Q12789 | GTF3C1 | QLLYKRRyIFsVVEN | Y961 | 0.122 | 83.2 | 2109 | 0.05352 | 0.122 | 0.0535 | 1 |
| Q12789 | GTF3C1 | LQCVCLNtPLGVVRC | T1048 | 0.039 | 86.28 | 2109 | 0.026638 | 0.039 | 0.0266 | 1 |
| Q8N1I0 | DOCK4 | LLDILLRtILEItSR | T889 | 0.015 | 92.16 | 1966 | 0.010245 | 0.112 | 0.078 | 0.875 |
| A2VCK2 | DCDC2B | AAKRVVVyRNGDPFF | Y14 | 0.044 | 91.67 | 349 | 0.019302 | 0.321 | 0.1408 | 1 |
| Q15773 | MLF2 | RAHILQRsRNHRTGD | S167 | 0.145 | 91.81 | 248 | 0.115263 | 0.145 | 0.1153 | 1 |
| P17252 | PRKCA | RkGtEELyAIkILKK | Y365 | 0.034 | 94.68 | 672 | 0.014915 | 0.252 | 0.1015 | 1 |
| P17252 | PRKCA | HMMDGVttRtFCGtP | T495 | 0.181 | 96.74 | 672 | 0.123625 | 0.181 | 0.1236 | 1 |
| P17252 | PRKCA | tFCGtPDyIAPEIIA | Y504 | 0.176 | 98.04 | 672 | 0.077209 | 0.176 | 0.0772 | 1 |
| Q9Y4C0 | NRXN3 | DGRSGTIsVNsRRTP | S557 | 0.13 | 91.33 | 1643 | 0.103339 | 0.069 | 0.1229 | 1 |
| Q9Y4C0 | NRXN3 | AGLAQGMysNLPKLV | Y1002 | 0.191 | 88.26 | 1643 | 0.083789 | 0.191 | 0.0838 | 1 |
| Q9Y4C0 | NRXN3 | NLPKLVAsRDGFQGC | S1011 | 0.077 | 87.25 | 1643 | 0.061208 | 0.077 | 0.0612 | 1 |
| Q9H344 | OR51I2 | GLGAAARsFItLFPL | S152 | 0 | 97.68 | 312 | 0 | 0 | 0 | 1 |
| Q9NZQ7 | CD274 | KLQDAGVyRCMISYG | Y112 | 0.001 | 98.43 | 290 | 0.000439 | 0.001 | 0.0004 | 1 |
| Q9NZQ7 | CD274 | GkTtTTNsKREEKLF | S184 | 0.098 | 93.9 | 290 | 0.077902 | 0.098 | 0.0779 | 1 |
| Q9NZQ7 | CD274 | EKLFNVtsTLRINTT | S195 | 0.001 | 98.62 | 290 | 0.000795 | 0.001 | 0.0008 | 1 |
| Q96EP1 | CHFR | WKNMLtEsLVALQRG | S578 | 0.023 | 98.39 | 664 | 0.018283 | 0.023 | 0.0183 | 0.8571 |
| P53602 | MVD | LPINSSLsVTLHQDQ | S40 | 0 | 98.68 | 400 | 0 | 0 | 0 | 1 |

| | | | | | | | | | | |
|---|---|---|---|---|---|---|---|---|---|---|
| P53602 | MVD | GsGsACRsLyGGFVE | S162 | 0 | 98.23 | 400 | 0 | 0 | 0 | 1 |
| P53602 | MVD | MRASVEtsPLLRFRA | S222 | 0.066 | 98.44 | 400 | 0.052464 | 0.11 | 0.0953 | 1 |
| Q8IZU9 | KIRREL3 | RTTVDYTyFSEPVSC | Y307 | 0.175 | 95.39 | 778 | 0.07677 | 0.175 | 0.0768 | 1 |
| Q7Z7A4 | PXK | SWQIVRRysDFDLLN | Y56 | 0.3 | 96.75 | 578 | 0.131606 | 0.3 | 0.1316 | 1 |
| O14654 | IRS4 | DAPARLEyYENARKF | Y111 | 0.183 | 94.04 | 1257 | 0.080279 | 0.183 | 0.0803 | 0.9 |
| Q5VYX0 | RNLS | VGAGMtGsLCAALLR | S14 | 0 | 98.87 | 342 | 0 | 0 | 0 | 1 |
| Q5VYX0 | RNLS | QQLEAVsyssRYALG | Y190 | 0.329 | 98.58 | 342 | 0.144328 | 0.329 | 0.1443 | 0.8125 |
| Q5VYX0 | RNLS | LEAVsyssRYALGLF | S192 | 0.087 | 98.54 | 342 | 0.069158 | 0.087 | 0.0692 | 0.875 |
| Q8TCP9 | FAM200A | KDDFPLLsRKSILLL | S510 | 0 | 96.4 | 573 | 0 | 0 | 0 | 1 |
| P31153 | MAT2A | kAVVPAkyLDEDTIy | Y235 | 0.238 | 98.46 | 395 | 0.104407 | 0.238 | 0.1044 | 0.8889 |
| P31153 | MAT2A | RVLVQVsyAIGVSHP | Y320 | 0.071 | 98.55 | 395 | 0.031147 | 0.071 | 0.0311 | 1 |
| P23921 | RRM1 | LTTkHPDyAILAArI | Y74 | 0.009 | 89.35 | 792 | 0.003948 | 0.009 | 0.0039 | 1 |
| P23921 | RRM1 | GGIGVAVsCIRATGS | S253 | 0.071 | 98.44 | 792 | 0.056439 | 0.138 | 0.1097 | 1 |
| P23921 | RRM1 | sNQQNLGtIkCSNLC | T422 | 0.138 | 98.59 | 792 | 0.094256 | 0.138 | 0.0943 | 0.8889 |
| P23921 | RRM1 | LAkEQGPyEtyEGsP | Y553 | 0.085 | 98.58 | 792 | 0.037288 | 0.085 | 0.0373 | 1 |
| P23921 | RRM1 | EQGPyEtyEGsPVSk | Y556 | 0.147 | 98.66 | 792 | 0.064487 | 0.294 | 0.129 | 1 |
| P23921 | RRM1 | PyEtyEGsPVSkGIL | S559 | 0 | 98.55 | 792 | 0 | 0 | 0 | 1 |
| P23921 | RRM1 | LLIAPMPtASTAQIL | T604 | 0.025 | 94.13 | 792 | 0.017075 | 0.041 | 0.0302 | 1 |
| Q9H0H3 | KLHL25 | TFMCDKIyQVDHkAk | Y311 | 0.31 | 92.87 | 589 | 0.135993 | 0.31 | 0.136 | 1 |
| Q9H0H3 | KLHL25 | PSPRkEFsASAIGCk | S334 | 0.137 | 95.37 | 589 | 0.108903 | 0.137 | 0.1089 | 1 |
| Q9H0H3 | KLHL25 | SAIGCkVyVTGGRGS | Y343 | 0.079 | 96.31 | 589 | 0.034656 | 0.079 | 0.0347 | 0.9167 |
| Q96P47 | AGAP3 | DSRARKLsTDLKRCT | S219 | 0.113 | 94.03 | 875 | 0.089825 | 0.113 | 0.0898 | 1 |
| Q8TCS8 | PNPT1 | DPyEIIEsFNVVAKE | S339 | 0 | 97.89 | 783 | 0 | 0 | 0 | 1 |
| Q8TCS8 | PNPT1 | FPFTIRVtsEVLESN | T473 | 0.104 | 98.64 | 783 | 0.071033 | 0.104 | 0.071 | 1 |
| Q8TCS8 | PNPT1 | PFTIRVtsEVLESNG | S474 | 0 | 98.74 | 783 | 0 | 0 | 0 | 1 |
| O96006 | ZBED1 | SENQNRAyVtLAAHF | Y210 | 0.082 | 94.89 | 694 | 0.035972 | 0.082 | 0.036 | 0.8462 |
| O96006 | ZBED1 | SAKVFGAttNyGkDI | T268 | 0.001 | 97.82 | 694 | 0.000683 | 0.001 | 0.0007 | 0.8462 |
| O96006 | ZBED1 | AKVFGAttNyGkDIV | T269 | 0.008 | 95.99 | 694 | 0.005464 | 0.008 | 0.0055 | 1 |
| P46109 | CRKL | DNLEyVRtLyDFPGN | T130 | 0.029 | 86.86 | 303 | 0.019807 | 0.029 | 0.0198 | 0.9 |
| P16435 | POR | EESSIRQyELVVHtD | Y245 | 0.112 | 94.82 | 677 | 0.049133 | 0.112 | 0.0491 | 1 |
| P16435 | POR | YRTALTyyLDITNPP | Y374 | 0.003 | 98.48 | 677 | 0.001316 | 0.003 | 0.0013 | 1 |

| | | | | | | | | | | |
|---|---|---|---|---|---|---|---|---|---|---|
| P16435 | POR | VGETLLYyGCRRSDE | Y564 | 0.004 | 98.79 | 677 | 0.001755 | 0.004 | 0.0018 | 1 |
| P16435 | POR | CRRSDEDyLyREELA | Y573 | 0.22 | 98.21 | 677 | 0.096511 | 0.22 | 0.0965 | 1 |
| P16435 | POR | RSDEDyLyREELAQF | Y575 | 0.08 | 98.57 | 677 | 0.035095 | 0.08 | 0.0351 | 1 |
| O75570 | MTRF1 | NDVILEVtAGRTTGG | T190 | 0.216 | 94.64 | 445 | 0.147531 | 0.216 | 0.1475 | 1 |
| Q9NYU2 | UGGT1 | MLsVLKNtKTPVkFW | T1282 | 0.119 | 94.12 | 1555 | 0.081279 | 0.119 | 0.0813 | 0.9091 |
| P48047 | ATP5PO | GRyAtALysAASkQN | Y46 | 0.1 | 93.74 | 213 | 0.043869 | 0.172 | 0.0655 | 1 |
| P48047 | ATP5PO | RyAtALysAASkQNk | S47 | 0.034 | 92.44 | 213 | 0.027027 | 0.034 | 0.027 | 1 |
| P48047 | ATP5PO | VksLNDItAkERFSP | T96 | 0.083 | 84.38 | 213 | 0.05669 | 0.132 | 0.0864 | 1 |
| P48047 | ATP5PO | GEkyVDMsVktKIQK | S197 | 0.028 | 91.91 | 213 | 0.022258 | 0.028 | 0.0223 | 1 |
| P52209 | PGD | IIIDGGNsEyRDTTR | S104 | 0.07 | 97.39 | 483 | 0.055644 | 0.07 | 0.0556 | 1 |
| P52209 | PGD | IDGGNsEyRDTTRRC | Y106 | 0.068 | 97.99 | 483 | 0.029831 | 0.068 | 0.0298 | 1 |
| P52209 | PGD | kGILFVGsGVsGGEE | S126 | 0.01 | 98.82 | 483 | 0.007949 | 0.01 | 0.0079 | 1 |
| P52209 | PGD | LFVGsGVsGGEEGAR | S129 | 0.065 | 97.04 | 483 | 0.051669 | 0.065 | 0.0517 | 1 |
| P12004 | PCNA | GIPEQEysCVVkMPS | S134 | 0.186 | 96.7 | 261 | 0.147854 | 0.186 | 0.1479 | 1 |
| P12004 | PCNA | QLTFALryLNFFtkA | Y211 | 0.321 | 98.2 | 261 | 0.140818 | 0.321 | 0.1408 | 0.9 |
| P12004 | PCNA | LryLNFFtkATPLSS | T216 | 0.191 | 98.63 | 261 | 0.130456 | 0.191 | 0.1305 | 1 |
| P12004 | PCNA | DMGHLkYyLAPkIED | Y250 | 0.277 | 97.6 | 261 | 0.121516 | 0.277 | 0.1215 | 0.8889 |
| Q86TX2 | ACOT1 | GVGLLGIsKGGELCL | S232 | 0.16 | 98.48 | 421 | 0.127186 | 0.16 | 0.1272 | 1 |
| Q86TX2 | ACOT1 | AAVVINGsVANVGGT | S257 | 0.001 | 98.84 | 421 | 0.000795 | 0.001 | 0.0008 | 1 |
| Q9HAU5 | UPF2 | AMHLDARyVtMVENA | Y744 | 0.184 | 92.6 | 1272 | 0.080718 | 0.184 | 0.0807 | 1 |
| Q96G28 | CFAP36 | QFQEACTsPLAkTHT | S85 | 0.182 | 91.41 | 342 | 0.144674 | 0.182 | 0.1447 | 1 |
| Q7KZI7 | MARK2 | AKFRQIVsAVQyCHQ | S161 | 0.138 | 98.59 | 788 | 0.109698 | 0.138 | 0.1097 | 1 |
| Q7KZI7 | MARK2 | KIADFGFsNEFtFGN | S197 | 0.016 | 77.82 | 788 | 0.012719 | 0.016 | 0.0127 | 1 |
| Q7KZI7 | MARK2 | DyKDPRRtELMVSMG | T329 | 0.175 | 94.36 | 788 | 0.119527 | 0.175 | 0.1195 | 1 |
| Q7KZI7 | MARK2 | ELMVSMGytREEIQD | Y337 | 0.145 | 96 | 788 | 0.063609 | 0.293 | 0.1285 | 1 |
| Q7KZI7 | MARK2 | DSLVGQRyNEVMATy | Y351 | 0.192 | 96.79 | 788 | 0.084228 | 0.192 | 0.0842 | 0.8333 |
| Q7KZI7 | MARK2 | yNEVMATyLLLGyKs | Y358 | 0.013 | 96.25 | 788 | 0.005703 | 0.013 | 0.0057 | 1 |
| Q13620 | CUL4B | QEREVPEyLHHVNkR | Y413 | 0.049 | 97.79 | 913 | 0.021496 | 0.049 | 0.0215 | 1 |
| P13051 | UNG | ERKHytVyPPPHQVF | Y128 | 0.31 | 98.46 | 313 | 0.135993 | 0.31 | 0.136 | 1 |
| Q9BR11 | ZSWIM1 | GHLARAVyFAIPAKE | Y113 | 0 | 97.74 | 485 | 0 | 0 | 0 | 0.8571 |
| P07357 | C8A | DQDCLDGsDEDDCED | S125 | 0.098 | 93.08 | 584 | 0.077902 | 0.098 | 0.0779 | 1 |

| | | | | | | | | | | |
|---|---|---|---|---|---|---|---|---|---|---|
| P07357 | C8A | INDYGTHyITSGsMG | Y341 | 0.056 | 94.77 | 584 | 0.024566 | 0.056 | 0.0246 | 1 |
| P07357 | C8A | THyITSGsMGGIYEY | S346 | 0.096 | 91.27 | 584 | 0.076312 | 0.096 | 0.0763 | 1 |
| O15357 | INPPL1 | LSTVsHDyLKGsyGL | Y185 | 0.32 | 80.06 | 1258 | 0.140379 | 0.32 | 0.1404 | 1 |
| O75191 | XYLB | IHAEGLGyRVMSkTk | Y428 | 0.016 | 97.85 | 536 | 0.007019 | 0.016 | 0.007 | 1 |
| O75191 | XYLB | SPGASQVyEALLPQy | Y513 | 0 | 98.1 | 536 | 0 | 0 | 0 | 1 |
| O75191 | XYLB | yEALLPQyAKLEQRI | Y520 | 0.023 | 98.34 | 536 | 0.01009 | 0.023 | 0.0101 | 1 |
| Q02413 | DSG1 | RNtGEIRtMNNFLDR | T220 | 0.01 | 96.65 | 1049 | 0.00683 | 0.01 | 0.0068 | 1 |
| Q92752 | TNR | VDFILLKyGLVGGEG | Y538 | 0.227 | 90.53 | 1358 | 0.099582 | 0.227 | 0.0996 | 1 |
| O75563 | SKAP2 | SKtVFYyyGsDKDKQ | Y152 | 0 | 97.3 | 359 | 0 | 0 | 0 | 1 |
| P0C7U0 | ELFN1 | VKQLTQNsATITVQL | S328 | 0.118 | 88.99 | 828 | 0.0938 | 0.118 | 0.0938 | 1 |
| O00300 | TNFRSF11B | YHFPKtVtQsLKKTI | T362 | 0.194 | 90.75 | 401 | 0.132505 | 0.194 | 0.1325 | 1 |
| Q15858 | SCN9A | LYMLSPFsPLRRISI | S113 | 0.179 | 82.14 | 1988 | 0.14229 | 0.179 | 0.1423 | 1 |
| O75808 | CAPN15 | APLRRREsMHVEQRR | S456 | 0.059 | 94.76 | 1086 | 0.0469 | 0.059 | 0.0469 | 0.9 |
| O75808 | CAPN15 | MkVDDSAyESLGLRP | Y708 | 0.118 | 92.96 | 1086 | 0.051765 | 0.118 | 0.0518 | 1 |
| O75808 | CAPN15 | LRPRHAYsILDVRDV | S720 | 0.019 | 93.77 | 1086 | 0.015103 | 0.019 | 0.0151 | 1 |
| P47712 | PLA2G4A | kLLGPkNsEGLHsAR | S178 | 0.114 | 87.54 | 749 | 0.09062 | 0.114 | 0.0906 | 1 |
| P47712 | PLA2G4A | ATYVAGLsGSTWYMS | S228 | 0.054 | 97.15 | 749 | 0.042925 | 0.054 | 0.0429 | 1 |
| P47712 | PLA2G4A | MAPDLFGskFFMGtV | S370 | 0.021 | 95.93 | 749 | 0.016693 | 0.179 | 0.1423 | 0.875 |
| P47712 | PLA2G4A | GskFFMGtVVKKYEE | T376 | 0.158 | 91.62 | 749 | 0.107916 | 0.158 | 0.1079 | 1 |
| P47712 | PLA2G4A | IKEAMVEsIEyRRQN | S715 | 0.062 | 92.36 | 749 | 0.049285 | 0.062 | 0.0493 | 1 |
| P49815 | TSC2 | NEHGSYRyTEFLTGL | Y1571 | 0.027 | 94.08 | 1807 | 0.011845 | 0.027 | 0.0118 | 1 |
| P49815 | TSC2 | HLGNDFVsIVyNDsG | S1647 | 0.004 | 94.51 | 1807 | 0.00318 | 0.004 | 0.0032 | 1 |
| P49815 | TSC2 | NDFVsIVyNDsGEDF | Y1650 | 0.027 | 95.66 | 1807 | 0.011845 | 0.027 | 0.0118 | 1 |
| P49815 | TSC2 | VsIVyNDsGEDFkLG | S1653 | 0.162 | 84.13 | 1807 | 0.128776 | 0.162 | 0.1288 | 1 |
| P0DMM9 | SULT1A3 | PLALLPQtLLDQkVk | T117 | 0.206 | 98.25 | 295 | 0.140701 | 0.206 | 0.1407 | 0.8333 |
| Q14671 | PUM1 | RGNVLVLsQHKFASN | S1073 | 0 | 98.55 | 1186 | 0 | 0 | 0 | 1 |
| Q14671 | PUM1 | DGPHSALyTMMKDQy | Y1113 | 0.24 | 94.94 | 1186 | 0.105285 | 0.24 | 0.1053 | 1 |
| Q5JR12 | PPM1J | yVANAGDsRAIIVRN | S280 | 0 | 98.28 | 505 | 0 | 0 | 0 | 1 |
| Q5JR12 | PPM1J | EPNDHSRyTALAQAL | Y458 | 0.07 | 95.86 | 505 | 0.030708 | 0.172 | 0.0755 | 1 |
| Q8N6M6 | AOPEP | AIRIWYKtKPEGRSV | T250 | 0.043 | 97.15 | 819 | 0.02937 | 0.091 | 0.0622 | 1 |
| Q8N6M6 | AOPEP | KRLGDETyFSFLRKF | Y605 | 0.001 | 95.11 | 819 | 0.000439 | 0.001 | 0.0004 | 1 |

| | | | | | | | | | | |
|---|---|---|---|---|---|---|---|---|---|---|
| Q8IZU8 | DSEL | FPNWGVVtYGAGLPN | T419 | 0.008 | 98.41 | 1212 | 0.005464 | 0.008 | 0.0055 | 1 |
| Q8IZU8 | DSEL | SGEAVsAySSAMRLK | Y560 | 0.105 | 98.4 | 1212 | 0.046062 | 0.199 | 0.0846 | 1 |
| Q13885 | TUBB2A | MPGFAPLtsrGsQQy | T274 | 0.188 | 89.23 | 445 | 0.128407 | 0.237 | 0.1463 | 0.8571 |
| Q13885 | TUBB2A | QELFkRIsEQFtAMF | S382 | 0.139 | 95.41 | 445 | 0.110493 | 0.139 | 0.1105 | 1 |
| Q9UM07 | PADI4 | GGIsGLDsFGNLEVS | S406 | 0.106 | 98.24 | 663 | 0.084261 | 0.106 | 0.0843 | 1 |
| Q75T13 | PGAP1 | SANDAEDsLRMHSTV | S812 | 0.021 | 92.36 | 922 | 0.016693 | 0.056 | 0.0445 | 0.8889 |
| Q75T13 | PGAP1 | SSKLLKttsQFPLPL | T883 | 0.006 | 92.62 | 922 | 0.004098 | 0.148 | 0.1011 | 1 |
| Q07837 | SLC3A1 | RTGKYTDyYIWHDCt | Y236 | 0.004 | 98.44 | 685 | 0.001755 | 0.004 | 0.0018 | 1 |
| Q07837 | SLC3A1 | TSRLGNQyVNVMNML | Y461 | 0.156 | 96.59 | 685 | 0.068435 | 0.156 | 0.0684 | 1 |
| O75691 | UTP20 | SKELGILskISKFMK | S1337 | 0.108 | 82.35 | 2785 | 0.085851 | 0.108 | 0.0859 | 1 |
| Q76I76 | SSH2 | MGVSRsAstVIAyAM | S401 | 0 | 98.42 | 1423 | 0 | 0 | 0 | 1 |
| Q76I76 | SSH2 | GVSRsAstVIAyAMK | T402 | 0.023 | 98.44 | 1423 | 0.015709 | 0.023 | 0.0157 | 1 |
| Q9BXU1 | STK31 | MQQKLyMsVEDFILE | S451 | 0.124 | 90.9 | 1019 | 0.098569 | 0.124 | 0.0986 | 1 |
| P25398 | RPS12 | SNCDEPMyVKLVEAL | Y61 | 0.116 | 86.3 | 132 | 0.050888 | 0.116 | 0.0509 | 0.9167 |
| Q96JK4 | HHIPL1 | ISRVVLtsPWEGDER | S242 | 0.116 | 95.71 | 782 | 0.09221 | 0.116 | 0.0922 | 1 |
| Q8NB59 | SYT14 | PNTYVKLtLLNSMGQ | T458 | 0.013 | 94.69 | 555 | 0.008879 | 0.065 | 0.0444 | 0.8333 |
| Q8NA58 | PNLDC1 | FLKNGIPyMNEEQEK | Y121 | 0.103 | 96.04 | 520 | 0.045185 | 0.103 | 0.0452 | 1 |
| Q96GJ1 | TRMT2B | TAGAEMLyRTVGELT | Y333 | 0.158 | 97.12 | 504 | 0.069312 | 0.301 | 0.1284 | 1 |
| Q9Y3D2 | MSRB2 | yHCVCCDsPLFSsEK | S95 | 0.134 | 96.57 | 182 | 0.106519 | 0.134 | 0.1065 | 0.8 |
| P48147 | PREP | DETAVQDyHGHkICD | Y19 | 0.306 | 98.69 | 710 | 0.134238 | 0.306 | 0.1342 | 1 |
| P48147 | PREP | GHkICDPyAWLEDPD | Y28 | 0.036 | 98.87 | 710 | 0.015793 | 0.036 | 0.0158 | 1 |
| P48147 | PREP | WMGGAELsDDGRyVL | S241 | 0.003 | 98.88 | 710 | 0.002385 | 0.003 | 0.0024 | 1 |
| P48147 | PREP | ELsDDGRyVLLsIRE | Y246 | 0.092 | 98.91 | 710 | 0.040359 | 0.092 | 0.0404 | 1 |
| P48147 | PREP | kGIDASDyQTVQIFy | Y435 | 0.151 | 97.17 | 710 | 0.066242 | 0.299 | 0.1312 | 1 |
| P48147 | PREP | yQTVQIFyPSKDGTK | Y442 | 0.003 | 98.9 | 710 | 0.001316 | 0.003 | 0.0013 | 1 |
| P48147 | PREP | FEWLVKYsPLHNVkL | S615 | 0 | 98.95 | 710 | 0 | 0 | 0 | 1 |
| Q96PD2 | DCBLD2 | HFNYLRIyNGIGVSR | Y134 | 0.276 | 97.5 | 775 | 0.121077 | 0.276 | 0.1211 | 1 |
| Q96PD2 | DCBLD2 | KkITGIItTGSTMVE | T366 | 0.015 | 98.11 | 775 | 0.010245 | 0.015 | 0.0102 | 1 |
| Q8WYA1 | BMAL2 | RGkILFVsksVskIL | S208 | 0.118 | 88.31 | 636 | 0.0938 | 0.137 | 0.1089 | 0.8333 |
| Q6UB28 | METAP1D | SRGAQILtKLPHEA_ | T329 | 0 | 98.67 | 335 | 0 | 0 | 0 | 1 |
| Q8NBN3 | TMEM87A | KAEEVELyLEkLKEK | Y104 | 0.235 | 81.85 | 555 | 0.103091 | 0.235 | 0.1031 | 1 |

| | | | | | | | | | | |
|---|---|---|---|---|---|---|---|---|---|---|
| P54578 | USP14 | GEMAsAQyITAALRD | Y151 | 0.235 | 91.05 | 494 | 0.103091 | 0.235 | 0.1031 | 1 |
| Q8IVB5 | LIX1L | VPSNSPPyVCYVTLP | Y136 | 0.324 | 98.21 | 337 | 0.142134 | 0.324 | 0.1421 | 1 |
| Q13394 | MAB21L1 | LVyHLNKyyNEKCQA | Y15 | 0.044 | 98.46 | 359 | 0.019302 | 0.044 | 0.0193 | 1 |
| Q9NTI2 | ATP8A2 | RGTQLRNtQWVFGIV | T279 | 0.006 | 84.23 | 1188 | 0.004098 | 0.006 | 0.0041 | 1 |
| Q9NTI2 | ATP8A2 | WVFGIVVytGHDtKL | Y288 | 0.014 | 93.47 | 1188 | 0.006142 | 0.014 | 0.0061 | 1 |
| Q9NTI2 | ATP8A2 | VFGIVVytGHDtKLM | T289 | 0 | 93.71 | 1188 | 0 | 0 | 0 | 1 |
| Q9NTI2 | ATP8A2 | VVytGHDtKLMQNst | T293 | 0 | 89.68 | 1188 | 0 | 0 | 0 | 1 |
| P35228 | NOS2 | QLKSQKRyHEDIFGA | Y1123 | 0.143 | 92.47 | 1153 | 0.062732 | 0.143 | 0.0627 | 1 |
| Q13618 | CUL3 | TPEDRKVyTYVA___ | Y764 | 0.086 | 97.1 | 768 | 0.037727 | 0.086 | 0.0377 | 0.8571 |
| O15118 | NPC1 | RFFkNSYsPLLLKDW | S826 | 0 | 88.32 | 1278 | 0 | 0 | 0 | 1 |
| Q8TD84 | DSCAML1 | NAAGKIRsPNIRVKA | S114 | 0.004 | 86.08 | 2053 | 0.00318 | 0.004 | 0.0032 | 1 |
| Q06418 | TYRO3 | CANALGPsPYADWVP | S307 | 0.064 | 96.55 | 890 | 0.050875 | 0.064 | 0.0509 | 1 |
| Q96G04 | EEF2KMT | kMCRPRAyIFSDCHs | Y184 | 0.003 | 98.25 | 330 | 0.001316 | 0.003 | 0.0013 | 1 |
| Q9BPX5 | ARPC5L | LAVGGLGsIIRVLTA | S142 | 0 | 97.25 | 153 | 0 | 0 | 0 | 1 |
| Q7Z2Q7 | LRRC70 | ALRILDLsNNNILRI | S188 | 0.137 | 98.46 | 622 | 0.108903 | 0.137 | 0.1089 | 0.875 |
| P53355 | DAPK1 | EKSTGLQyAAKFIKK | Y39 | 0.255 | 94.7 | 1430 | 0.111865 | 0.255 | 0.1119 | 1 |
| Q99956 | DUSP9 | YLGSARDsANLESLA | S219 | 0.04 | 95.96 | 384 | 0.031797 | 0.04 | 0.0318 | 0.9 |
| Q99956 | DUSP9 | LAKLGIRyILNVtPN | Y232 | 0.121 | 98.72 | 384 | 0.053081 | 0.121 | 0.0531 | 1 |
| O43930 | PRKY | AKKLVDRtWtLCGtP | T201 | 0.058 | 92.99 | 277 | 0.039615 | 0.058 | 0.0396 | 1 |
| Q9UHD8 | SEPTIN9 | IPFAVVGsDHEyQVN | S502 | 0.038 | 89.61 | 586 | 0.030207 | 0.038 | 0.0302 | 0.8 |
| P31415 | CASQ1 | GyFKsKDsEHYKAFE | S191 | 0.164 | 97.86 | 396 | 0.130366 | 0.164 | 0.1304 | 1 |
| P31415 | CASQ1 | PYIPFFAtFDsKVAK | T213 | 0.005 | 98.72 | 396 | 0.003415 | 0.005 | 0.0034 | 1 |
| O75185 | ATP2C2 | VIGtGEssQFGEVFK | S268 | 0.015 | 84.88 | 946 | 0.011924 | 0.015 | 0.0119 | 1 |
| O75185 | ATP2C2 | NAVMGQPtEGALMAL | T457 | 0.138 | 89.31 | 946 | 0.094256 | 0.138 | 0.0943 | 0.8333 |
| O75185 | ATP2C2 | GVSVKMItGDALEtA | T600 | 0.051 | 90.02 | 946 | 0.034834 | 0.051 | 0.0348 | 1 |
| P49221 | TGM4 | GVFCCGPsPLtAIRK | S366 | 0 | 98.46 | 684 | 0 | 0 | 0 | 0.9 |
| P98172 | EFNB1 | PYEYYKLyLVRPEQA | Y79 | 0.008 | 96.22 | 346 | 0.003509 | 0.008 | 0.0035 | 1 |
| Q7Z7F0 | KHDC4 | QDEISRLsGAAVsTR | S133 | 0.035 | 96.54 | 614 | 0.027822 | 0.035 | 0.0278 | 1 |
| Q8TF05 | PPP4R1 | LLPIVVRyLADQNNQ | Y139 | 0.045 | 96.23 | 950 | 0.019741 | 0.045 | 0.0197 | 1 |
| Q9NZN1 | IL1RAPL1 | GGFVVRRtTELTVTA | T228 | 0.057 | 97.55 | 696 | 0.038932 | 0.057 | 0.0389 | 1 |
| P56192 | MARS1 | __MRLFVsDGVPGCL | S6 | 0.061 | 89.54 | 900 | 0.04849 | 0.061 | 0.0485 | 1 |

| | | | | | | | | | | |
|---|---|---|---|---|---|---|---|---|---|---|
| P56192 | MARS1 | PLLQDPAyLPEELSA | Y161 | 0.119 | 93.31 | 900 | 0.052204 | 0.119 | 0.0522 | 1 |
| Q9ULH1 | ASAP1 | VHISRIQsLELDKLG | S493 | 0.01 | 96.4 | 1129 | 0.007949 | 0.01 | 0.0079 | 1 |
| Q6ZVX7 | NCCRP1 | QHHVAPRtSGRGPPG | T218 | 0.216 | 88.4 | 275 | 0.147531 | 0.216 | 0.1475 | 1 |
| P17480 | UBTF | QQsVIGDyLARFKND | Y500 | 0.33 | 95.92 | 764 | 0.144766 | 0.33 | 0.1448 | 0.8182 |
| Q13618 | CUL3 | LNSRSPEyLSLFIDD | Y387 | 0.217 | 93.92 | 768 | 0.095195 | 0.217 | 0.0952 | 1 |
| Q27J81 | INF2 | QGYVRQLsQALDtsN | S129 | 0.001 | 92.69 | 1249 | 0.000795 | 0.001 | 0.0008 | 1 |
| P42768 | WAS | QLtDAEtsKLIyDFI | S287 | 0.011 | 87.75 | 502 | 0.008744 | 0.011 | 0.0087 | 1 |
| Q8TAQ9 | SUN3 | EHISEKVsPSGNISS | S272 | 0.179 | 91.76 | 357 | 0.14229 | 0.179 | 0.1423 | 1 |
| Q6PF15 | KLHL35 | ARQGGVNtDkVQCFD | T456 | 0.003 | 97.95 | 583 | 0.002049 | 0.003 | 0.002 | 0.9 |
| Q9Y6T7 | DGKB | PYSIINNyFsIGVDA | Y586 | 0.183 | 96.25 | 804 | 0.080279 | 0.183 | 0.0803 | 0.8125 |
| Q9Y6T7 | DGKB | SIINNyFsIGVDAsI | S588 | 0.015 | 96.82 | 804 | 0.011924 | 0.015 | 0.0119 | 1 |
| Q9P2L0 | WDR35 | QYQKLTTsDENGLII | S89 | 0 | 96.09 | 1181 | 0 | 0 | 0 | 1 |
| Q5T6X5 | GPRC6A | SFLRTVPsDFHQIKA | S191 | 0.061 | 96.1 | 926 | 0.04849 | 0.061 | 0.0485 | 1 |
| Q5T6X5 | GPRC6A | IEVRINRtLKKIILE | T261 | 0.011 | 98.22 | 926 | 0.007513 | 0.011 | 0.0075 | 1 |
| Q5T6X5 | GPRC6A | WSTATKIttIPNVKK | T312 | 0.155 | 93.71 | 926 | 0.105867 | 0.155 | 0.1059 | 1 |
| Q9UKD2 | MRTO4 | GEVGLLFtNRTkEEV | T105 | 0 | 98.26 | 239 | 0 | 0 | 0 | 1 |
| Q9UKD2 | MRTO4 | LRQLGLPtALkRGVV | T160 | 0.106 | 97.32 | 239 | 0.072399 | 0.106 | 0.0724 | 1 |
| Q8IWW6 | ARHGAP12 | GLDIDGIyRVsGNLA | Y691 | 0.008 | 98.61 | 846 | 0.003509 | 0.008 | 0.0035 | 1 |
| P43378 | PTPN9 | MRTQRAFsIQTPEQY | S557 | 0.01 | 98.32 | 593 | 0.007949 | 0.01 | 0.0079 | 0.9 |
| Q8N1F7 | NUP93 | DRRLsPAtENkLRLH | T367 | 0.203 | 86.7 | 819 | 0.138652 | 0.203 | 0.1387 | 1 |
| Q8N1F7 | NUP93 | DPYkRAVyCIIGRCD | Y391 | 0.103 | 89.36 | 819 | 0.045185 | 0.103 | 0.0452 | 1 |
| Q17RG1 | KCTD19 | LESVKPLyTMALGLL | Y287 | 0.16 | 94.76 | 926 | 0.07019 | 0.16 | 0.0702 | 1 |
| P46777 | RPL5 | MEEDEDAykkQFSQY | Y219 | 0.281 | 95.87 | 297 | 0.123271 | 0.281 | 0.1233 | 1 |
| P46777 | RPL5 | PDMMEEMykkAHAAI | Y240 | 0.003 | 97.26 | 297 | 0.001316 | 0.003 | 0.0013 | 1 |
| Q02241 | KIF23 | AsDGEIEtKLIkGDI | T873 | 0.087 | 87.52 | 960 | 0.059422 | 0.087 | 0.0594 | 1 |
| Q9HC84 | MUC5B | CLKAVTLsLDGGDtA | S484 | 0.067 | 89.66 | 5762 | 0.053259 | 0.067 | 0.0533 | 1 |
| Q96BS2 | TESC | HRRFkQLsGDQPTIR | S36 | 0.05 | 95.19 | 214 | 0.039746 | 0.05 | 0.0397 | 1 |
| P22392 | NME2 | EEHLkQHyIDLkDrP | Y52 | 0.207 | 94.71 | 152 | 0.090808 | 0.207 | 0.0908 | 0.8 |
| P22392 | NME2 | GrNIIhGsDsVkSAE | S120 | 0.006 | 98.41 | 152 | 0.004769 | 0.006 | 0.0048 | 1 |
| Q9UPQ0 | LIMCH1 | AEGILQQyIERFTIS | Y446 | 0.232 | 91.14 | 1083 | 0.101775 | 0.232 | 0.1018 | 0.8 |
| P05156 | CFI | PTYCQQKsLECLHPG | S90 | 0 | 96.31 | 583 | 0 | 0 | 0 | 1 |

| | | | | | | | | | | |
|---|---|---|---|---|---|---|---|---|---|---|
| H3BPM6 | MKRN2OS | ASKFItLyRAIREHG | Y196 | 0.198 | 95.6 | 223 | 0.08686 | 0.263 | 0.1154 | 1 |
| Q6P2Q9 | PRPF8 | VDHNIADyMtAkNNV | Y1008 | 0.015 | 96.66 | 2335 | 0.00658 | 0.015 | 0.0066 | 1 |
| Q6P2Q9 | PRPF8 | HNIADyMtAkNNVVI | T1010 | 0.136 | 95.95 | 2335 | 0.09289 | 0.1 | 0.1103 | 1 |
| Q6P2Q9 | PRPF8 | ARDLIQRyLTEHPDP | Y1113 | 0.172 | 93.75 | 2335 | 0.075454 | 0.172 | 0.0755 | 1 |
| Q6P2Q9 | PRPF8 | NNENIVGyNNkKCWP | Y1128 | 0.183 | 87.52 | 2335 | 0.080279 | 0.183 | 0.0803 | 1 |
| Q86VZ5 | SGMS1 | YPMEWGKtFLAFLYA | T135 | 0.067 | 91.93 | 413 | 0.045762 | 0.067 | 0.0458 | 0.9 |
| Q86VZ5 | SGMS1 | WLLLKYKsIIsRRFF | S209 | 0.06 | 88.2 | 413 | 0.047695 | 0.06 | 0.0477 | 1 |
| P23458 | JAK1 | TVTRLVNtLkEGkRL | T1107 | 0.104 | 92.75 | 1154 | 0.071033 | 0.104 | 0.071 | 1 |
| E9PAV3 | NACA | VITkPDVyksPAsDt | Y1975 | 0.116 | 96.41 | 2078 | 0.050888 | 0.116 | 0.0509 | 1 |
| P21817 | RYR1 | GLFFPVVsFSAGVKV | S781 | 0 | 96.835 | 5038 | 0 | 0 | 0 | 1 |
| Q4G0J3 | LARP7 | GVIVKIIstEPLPGR | S461 | 0.13 | 82.14 | 582 | 0.103339 | 0.144 | 0.1274 | 1 |
| P59045 | NLRP11 | GVAKLLEsLIsPDCV | S940 | 0.087 | 95.23 | 1033 | 0.069158 | 0.087 | 0.0692 | 1 |
| P38935 | IGHMBP2 | IIHGPPGtGKTTTVV | T218 | 0.009 | 96.34 | 993 | 0.006147 | 0.009 | 0.0061 | 0.8 |
| P38935 | IGHMBP2 | ARDIAVVsPYNLQVD | S539 | 0.002 | 93.92 | 993 | 0.00159 | 0.002 | 0.0016 | 1 |
| P30153 | PPP2R1A | DANQHVksALASVIM | S343 | 0.027 | 97.12 | 589 | 0.021463 | 0.027 | 0.0215 | 1 |
| P30153 | PPP2R1A | ASVIMGLsPILGKDN | S353 | 0 | 94.34 | 589 | 0 | 0 | 0 | 1 |
| P10412 | H1-4 | kGtGAsGsFkLNkkA | S104 | 0.154 | 96.25 | 219 | 0.122417 | 0.154 | 0.1224 | 0.875 |
| Q9H0E2 | TOLLIP | GMtRMDPyCRLRLGy | Y76 | 0.169 | 98.6 | 274 | 0.074138 | 0.169 | 0.0741 | 0.9 |
| Q5TA45 | INTS11 | MNLkVPIyFSTGLTE | Y266 | 0.282 | 93.62 | 600 | 0.123709 | 0.282 | 0.1237 | 1 |
| O14965 | AURKA | KIADFGWsVHAPssR | S278 | 0.006 | 66.63 | 403 | 0.004769 | 0.006 | 0.0048 | 1 |
| Q9UGC7 | MTRF1L | QGRVHTStMTVAILP | T218 | 0.006 | 97.28 | 380 | 0.004098 | 0.006 | 0.0041 | 1 |
| Q9NP74 | PALMD | QIQVLEQsILRLEKE | S81 | 0.146 | 98.35 | 551 | 0.116058 | 0.084 | 0.1409 | 1 |
| O14815 | CAPN9 | EKALKRGsLLGCFID | S228 | 0.079 | 92.11 | 690 | 0.062798 | 0.129 | 0.1025 | 1 |
| Q9BXW9 | FANCD2 | ISFGSLLyKYAFKFF | Y458 | 0.028 | 97.65 | 1451 | 0.012283 | 0.028 | 0.0123 | 0.8125 |
| O60423 | ATP8B3 | NSTRKRMsVLVRKPE | S645 | 0 | 87.78 | 1300 | 0 | 0.035 | 0.0278 | 1 |
| P15309 | ACP3 | FQELESEtLKSEEFQ | T171 | 0.026 | 98.31 | 386 | 0.017758 | 0.026 | 0.0178 | 1 |
| P15309 | ACP3 | PSWATEDtMtKLREL | T231 | 0.138 | 98.2 | 386 | 0.094256 | 0.138 | 0.0943 | 1 |
| P15309 | ACP3 | MtKLRELsELSLLSL | S239 | 0.002 | 98.51 | 386 | 0.00159 | 0.002 | 0.0016 | 0.9 |
| Q9H3H3 | C11orf68 | VyTYLGIyRANRWHL | Y255 | 0.312 | 93.03 | 292 | 0.13687 | 0.312 | 0.1369 | 0.8571 |
| Q9HBU9 | POPDC2 | PEEFDLLyKTLCLPL | Y112 | 0.338 | 90.63 | 364 | 0.148276 | 0.338 | 0.1483 | 1 |
| P02774 | GC | FRKDPKEyANQFMWE | Y155 | 0.28 | 98.54 | 474 | 0.122832 | 0.28 | 0.1228 | 0.8462 |

| | | | | | | | | | | |
|---|---|---|---|---|---|---|---|---|---|---|
| P02774 | GC | LKHLsLLttLSNRVC | T213 | 0 | 94.44 | 474 | 0 | 0 | 0 | 1 |
| Q8TCT0 | CERK | GCFVCRQsKQQLEEE | S357 | 0.087 | 94.82 | 537 | 0.069158 | 0.087 | 0.0692 | 0.9 |
| Q8TCT0 | CERK | RRSPRGLsPAAHLGD | S408 | 0 | 94.55 | 537 | 0 | 0 | 0 | 1 |
| Q6ZW49 | PAXIP1 | PEPKGEKyECALKRA | Y149 | 0.012 | 92.96 | 1069 | 0.005264 | 0.012 | 0.0053 | 1 |
| Q05586 | GRIN1 | SDKFIYAtVKQSSVD | T683 | 0.021 | 95.68 | 938 | 0.014343 | 0.021 | 0.0143 | 1 |
| P01024 | C3 | QRIsLPEsLkRIPIE | S287 | 0.053 | 79.99 | 1663 | 0.04213 | 0.053 | 0.0421 | 0.875 |
| Q7Z2K6 | ERMP1 | DALQARDyLEHITSI | Y115 | 0.119 | 98.69 | 904 | 0.052204 | 0.119 | 0.0522 | 1 |
| P23526 | AHCY | AKAGIPVyAWkGETD | Y100 | 0.017 | 98.89 | 432 | 0.007458 | 0.017 | 0.0075 | 1 |
| P23526 | AHCY | LDDGGDLtNLIHTky | T136 | 0.013 | 98.87 | 432 | 0.008879 | 0.013 | 0.0089 | 1 |
| P23526 | AHCY | LPGIrGIsEETtTGV | S154 | 0 | 98.93 | 432 | 0 | 0 | 0 | 1 |
| P23526 | AHCY | PAINVNDsVTkSkFD | S183 | 0.054 | 98.64 | 432 | 0.042925 | 0.054 | 0.0429 | 1 |
| P23526 | AHCY | NLyGCREsLIDGIkR | S198 | 0.025 | 98.82 | 432 | 0.019873 | 0.025 | 0.0199 | 0.9 |
| P23526 | AHCY | KVAVVAGyGDVGkGC | Y221 | 0.097 | 98.88 | 432 | 0.042553 | 0.097 | 0.0426 | 1 |
| P23526 | AHCY | LWTHPDKyPVGVHFL | Y379 | 0.167 | 98.56 | 432 | 0.07326 | 0.167 | 0.0733 | 0.8333 |
| Q9BQ04 | RBM4B | KLHVGNIsPtCTNQE | S86 | 0.122 | 87.78 | 359 | 0.09698 | 0.122 | 0.097 | 1 |
| P56277 | CMC4 | AQYPKGRsVVCsGFE | S47 | 0.025 | 89.67 | 68 | 0.019873 | 0.025 | 0.0199 | 1 |
| P07510 | CHRNG | LTFLLVVtILIVVNA | T316 | 0.041 | 90.21 | 517 | 0.028004 | 0.041 | 0.028 | 1 |
| O95837 | GNA14 | CYDRRREyQLSDSAK | Y147 | 0.33 | 96.13 | 355 | 0.144766 | 0.33 | 0.1448 | 1 |
| Q14392 | LRRC32 | LASPLGFyTALRHLD | Y72 | 0.031 | 94.74 | 662 | 0.013599 | 0.031 | 0.0136 | 1 |
| Q14392 | LRRC32 | LRDLPPytFANLASL | T405 | 0.068 | 92.39 | 662 | 0.046445 | 0.105 | 0.0793 | 1 |
| Q01518 | CAP1 | NDAAMFytNRVLKEY | T165 | 0.006 | 97.75 | 475 | 0.004098 | 0.006 | 0.0041 | 1 |
| Q8NB49 | ATP11C | LkNTEKIyGVAVYTG | Y261 | 0.13 | 94.82 | 1132 | 0.057029 | 0.13 | 0.057 | 1 |
| Q9Y6L7 | TLL2 | NMRVEFKsDNtVSKR | S716 | 0.147 | 91.23 | 1015 | 0.116852 | 0.147 | 0.1169 | 1 |
| Q86U06 | RBM23 | VEFCEIQsVPLAIGL | S219 | 0.046 | 90.71 | 439 | 0.036566 | 0.046 | 0.0366 | 1 |
| Q9NX02 | NLRP2 | GAKLLYTtLRHPKCF | T834 | 0.048 | 88.96 | 1062 | 0.032785 | 0.048 | 0.0328 | 1 |
| O00566 | MPHOSPH10 | AQKRPENsLLEETLH | S399 | 0.103 | 92 | 681 | 0.081876 | 0.103 | 0.0819 | 1 |
| Q6RFH5 | WDR74 | DRTVkHFsTEDGIFQ | S71 | 0.048 | 91.14 | 385 | 0.038156 | 0.048 | 0.0382 | 1 |
| Q8N3C7 | CLIP4 | SVKLHEGsQVLLTSS | S627 | 0.058 | 92.14 | 705 | 0.046105 | 0.058 | 0.0461 | 0.8 |
| P46781 | RPS9 | sQkHIDFsLrsPyGG | S160 | 0.064 | 92.04 | 194 | 0.050875 | 0.102 | 0.0811 | 1 |
| P46781 | RPS9 | HIDFsLrsPyGGGRP | S163 | 0.026 | 89.06 | 194 | 0.020668 | 0.026 | 0.0207 | 0.875 |
| O60706 | ABCC9 | SNLRFCItGMMVILN | T170 | 0.004 | 91.9 | 1549 | 0.002732 | 0.004 | 0.0027 | 1 |

| | | | | | | | | | | |
|---|---|---|---|---|---|---|---|---|---|---|
| Q9UPU5 | USP24 | RDVySSDyFSFVLSL | Y2138 | 0.001 | 93.85 | 2620 | 0.000439 | 0.001 | 0.0004 | 0.9167 |
| Q9UPU5 | USP24 | TKLKHPyyPCMAKVS | Y2158 | 0.088 | 92.74 | 2620 | 0.038604 | 0.167 | 0.0733 | 1 |
| O75072 | FKTN | EAVAFRKsAKELLQL | S276 | 0.173 | 97.14 | 461 | 0.13752 | 0.173 | 0.1375 | 1 |
| O75072 | FKTN | KFGKVEDsLELSFQG | S352 | 0.006 | 97.13 | 461 | 0.004769 | 0.006 | 0.0048 | 1 |
| P24723 | PRKCH | EAVGLQPtRWsLRHs | T25 | 0.053 | 79.33 | 683 | 0.0362 | 0.217 | 0.1482 | 1 |
| Q9Y316 | MEMO1 | QEFGkLFsKYLADPS | S174 | 0.107 | 98.6 | 297 | 0.085056 | 0.107 | 0.0851 | 1 |
| Q9Y316 | MEMO1 | DESQGEIyRSIEHLD | Y210 | 0.22 | 98.57 | 297 | 0.096511 | 0.22 | 0.0965 | 1 |
| P04406 | GAPDH | WGDAGAEyVVEstGV | Y94 | 0.038 | 98.8 | 335 | 0.01667 | 0.038 | 0.0167 | 1 |
| P04406 | GAPDH | RVIIsAPsADAPMFV | S125 | 0.081 | 97.26 | 335 | 0.064388 | 0.081 | 0.0644 | 1 |
| P04406 | GAPDH | IIsNAsCttNCLAPL | T153 | 0.072 | 98.69 | 335 | 0.049177 | 0.139 | 0.0949 | 0.8 |
| P04406 | GAPDH | IsNAsCttNCLAPLA | T154 | 0.005 | 98.75 | 335 | 0.003415 | 0.005 | 0.0034 | 0.9167 |
| P04406 | GAPDH | DHFVKLIsWyDNEFG | S312 | 0.003 | 98.95 | 335 | 0.002385 | 0.003 | 0.0024 | 1 |
| P51805 | PLXNA3 | CsWRGVEyRLVQSAH | Y289 | 0.048 | 93.93 | 1871 | 0.021057 | 0.048 | 0.0211 | 1 |
| Q9UGP5 | POLL | QAIGLKHysDFLERM | Y380 | 0.28 | 98.43 | 575 | 0.122832 | 0.28 | 0.1228 | 1 |
| Q9UGP5 | POLL | GPGRVLPtPTEKDVF | T553 | 0.11 | 97.16 | 575 | 0.075131 | 0.11 | 0.0751 | 1 |
| Q9UG63 | ABCF2 | LDLSPLEyMMKCyPE | Y482 | 0.254 | 93.18 | 623 | 0.111426 | 0.254 | 0.1114 | 1 |
| P17480 | UBTF | sEKKKAKyKAREAAL | Y456 | 0.268 | 91.99 | 764 | 0.117568 | 0.268 | 0.1176 | 0.8333 |
| Q8TCA0 | LRRC20 | IKFDMLMsPEGARAP | S175 | 0.113 | 93.27 | 184 | 0.089825 | 0.113 | 0.0898 | 0.8571 |
| O75521 | ECI2 | PFSHLGQsPEGCssy | S278 | 0.177 | 98.42 | 394 | 0.1407 | 0.177 | 0.1407 | 1 |
| Q9NPB8 | GPCPD1 | GVSVQYRyFKGyFLE | Y68 | 0 | 98.53 | 672 | 0 | 0 | 0 | 1 |
| Q96I99 | SUCLG2 | LDIsRETyLAILMDR | Y153 | 0.105 | 98.31 | 432 | 0.046062 | 0.105 | 0.0461 | 1 |
| P04899 | GNAI2 | LLLGAGEsGkSTIVk | S44 | 0.029 | 97.47 | 355 | 0.023053 | 0.029 | 0.0231 | 1 |
| P04899 | GNAI2 | ERIAQSDyIPTQQDV | Y168 | 0.106 | 97.55 | 355 | 0.046501 | 0.106 | 0.0465 | 1 |
| Q13349 | ITGAD | GQVLLLRsLPVLKVG | S614 | 0.038 | 90.73 | 1161 | 0.030207 | 0.038 | 0.0302 | 1 |
| O60840 | CACNA1F | LVLHPSAyIRNGWNL | Y158 | 0.012 | 84.21 | 1977 | 0.005264 | 0.012 | 0.0053 | 0.9167 |
| Q96C24 | SYTL4 | AKKRSNPyVKTYLLP | Y396 | 0.169 | 96.22 | 671 | 0.074138 | 0.169 | 0.0741 | 0.8182 |
| Q96HY6 | DDRGK1 | VGLRtQDtINRIQDL | T245 | 0 | 92.95 | 314 | 0 | 0 | 0 | 0.9091 |
| Q96PV4 | PNMA5 | sPLSVRStDMIRLKH | T284 | 0.1 | 86.27 | 448 | 0.068301 | 0.1 | 0.0683 | 1 |
| O75600 | GCAT | LGGASGGyTTGPGPL | Y274 | 0.014 | 98.78 | 419 | 0.006142 | 0.014 | 0.0061 | 0.9333 |
| O75600 | GCAT | LRQRARPyLFSNSLP | Y292 | 0.175 | 98.67 | 419 | 0.07677 | 0.175 | 0.0768 | 1 |
| Q96PF2 | TSSK2 | TVLRKKGyIVGINLG | Y12 | 0.033 | 97.82 | 358 | 0.014477 | 0.033 | 0.0145 | 1 |

| | | | | | | | | | | |
|---|---|---|---|---|---|---|---|---|---|---|
| Q96PF2 | TSSK2 | ETSDGRIyIIMELGV | Y87 | 0.004 | 96.33 | 358 | 0.001755 | 0.004 | 0.0018 | 1 |
| P05120 | SERPINB2 | NsWVKtQtKGKIPNL | T177 | 0.156 | 86.74 | 415 | 0.10655 | 0.156 | 0.1066 | 1 |
| Q9UNK9 | ANGEL1 | TDGCAVCykPTRFRL | Y336 | 0.131 | 98.78 | 670 | 0.057468 | 0.131 | 0.0575 | 1 |
| Q9UNK9 | ANGEL1 | RFRLLCAsPVEyFRP | S347 | 0.182 | 98.34 | 670 | 0.144674 | 0.182 | 0.1447 | 1 |
| Q9BPY3 | FAM118B | KGIkVISyGDDyADL | Y303 | 0.121 | 96.06 | 351 | 0.053081 | 0.121 | 0.0531 | 1 |
| P07225 | PROS1 | VFENDPEtDYFYPKY | T78 | 0.03 | 83.56 | 676 | 0.02049 | 0.03 | 0.0205 | 0.8182 |
| Q8TAT6 | NPLOC4 | sIIIRVQsPDGVkRI | S11 | 0.133 | 86.69 | 608 | 0.105724 | 0.133 | 0.1057 | 1 |
| Q9UKZ4 | TENM1 | STSPAHKyyLAMDPV | Y1239 | 0.01 | 94.22 | 2725 | 0.004387 | 0.01 | 0.0044 | 1 |
| Q13042 | CDC16 | FKMCYKLtSVVMEkD | T254 | 0.009 | 96.52 | 620 | 0.006147 | 0.009 | 0.0061 | 1 |
| Q13042 | CDC16 | RRYLskAttLEKTYG | T327 | 0.098 | 95.43 | 620 | 0.066935 | 0.098 | 0.0669 | 1 |
| Q92828 | CORO2A | PASkENCyDSVPITR | Y26 | 0.07 | 98.62 | 525 | 0.030708 | 0.07 | 0.0307 | 1 |
| Q03468 | ERCC6 | CHGILITsySYIRLM | S624 | 0.011 | 92.32 | 1493 | 0.008744 | 0.011 | 0.0087 | 1 |
| Q03468 | ERCC6 | HGILITsySYIRLMQ | Y625 | 0.242 | 91.37 | 1493 | 0.106162 | 0.242 | 0.1062 | 1 |
| P23919 | DTYMK | EKLsQGVtLVVDRYA | T92 | 0.041 | 98.68 | 212 | 0.028004 | 0.082 | 0.056 | 1 |
| Q8WU03 | GLYATL2 | NVISWEQtLQIQGCQ | T100 | 0.075 | 98.33 | 294 | 0.051226 | 0.075 | 0.0512 | 1 |
| Q99447 | PCYT2 | DSGsNLttDLIVQRI | T342 | 0.172 | 94.71 | 389 | 0.117478 | 0.172 | 0.1175 | 0.8182 |
| Q6PIF6 | MYO7B | CQILKQLtHNSNRHS | T1719 | 0.014 | 90.12 | 2116 | 0.009562 | 0.096 | 0.0546 | 1 |
| O43861 | ATP9B | EELGRLVyLLTDKTG | Y464 | 0.138 | 95.36 | 1147 | 0.060539 | 0.138 | 0.0605 | 1 |
| O43861 | ATP9B | WTEsVGLtLVsRDLT | T604 | 0.161 | 90.41 | 1147 | 0.109965 | 0.18 | 0.1229 | 1 |
| O43861 | ATP9B | GKEGKQAsLAADFsI | S900 | 0 | 81.4 | 1147 | 0 | 0 | 0 | 1 |
| O43861 | ATP9B | AsLAADFsItQFRHI | S906 | 0 | 91.48 | 1147 | 0 | 0 | 0 | 1 |
| Q8TF76 | HASPIN | QMRTkLSsLATAKSI | S621 | 0.166 | 95.61 | 798 | 0.131956 | 0.166 | 0.132 | 1 |
| Q8TBG4 | ETNPPL | HADSAsAyADEVKKI | Y189 | 0.006 | 98.72 | 499 | 0.002632 | 0.006 | 0.0026 | 1 |
| P0C869 | PLA2G4B | VLVARELsCLHVQLE | S150 | 0.115 | 90.35 | 781 | 0.091415 | 0.115 | 0.0914 | 1 |
| P0C869 | PLA2G4B | APsQLQRyRQELAER | Y382 | 0 | 92.37 | 781 | 0 | 0.023 | 0.0101 | 1 |
| P0C869 | PLA2G4B | KYGAFIPsELFGSEF | S473 | 0.048 | 98.2 | 781 | 0.038156 | 0.048 | 0.0382 | 1 |
| P0C869 | PLA2G4B | VNLSSSDsPYHYTKV | S732 | 0.059 | 89.9 | 781 | 0.0469 | 0.059 | 0.0469 | 1 |
| Q96MV8 | ZDHHC15 | SYTDKERyENEERPE | Y97 | 0.327 | 94.25 | 337 | 0.14345 | 0.327 | 0.1434 | 0.8182 |
| Q12986 | NFX1 | RCCPCGQtPLsQLLE | T599 | 0.018 | 80.04 | 1120 | 0.012294 | 0.018 | 0.0123 | 1 |
| Q5JXB2 | UBE2NL | VRFMTkIyHPNVDKL | Y77 | 0.321 | 93.18 | 153 | 0.140818 | 0.321 | 0.1408 | 0.8333 |
| Q7Z5J8 | ANKAR | GEAVIALsKDSRMHQ | S1019 | 0 | 91.37 | 1434 | 0 | 0 | 0 | 1 |

| | | | | | | | | | | |
|---|---|---|---|---|---|---|---|---|---|---|
| P40145 | ADCY8 | EIyVKGIsEQEGKIK | S1159 | 0.166 | 78.76 | 1251 | 0.131956 | 0.166 | 0.132 | 1 |
| Q96A32 | MYL11 | ADPEDVItGAFkVLD | T101 | 0.209 | 92.72 | 169 | 0.14275 | 0.209 | 0.1427 | 1 |
| Q9Y6N7 | ROBO1 | NMVGEREsEVAELTV | S251 | 0.148 | 95.28 | 1651 | 0.117647 | 0.148 | 0.1176 | 1 |
| Q9UEW8 | STK39 | DKEMMKKyGKsFRkL | Y306 | 0.069 | 96.96 | 545 | 0.030269 | 0.333 | 0.0948 | 1 |
| Q9Y5C1 | ANGPTL3 | RRGLSWKsQNGRLYS | S436 | 0.04 | 83.03 | 460 | 0.031797 | 0.04 | 0.0318 | 1 |
| Q9Y5C1 | ANGPTL3 | RLYSIKstKMLIHPT | T447 | 0.014 | 95.84 | 460 | 0.009562 | 0.141 | 0.0963 | 0.8889 |
| Q9Y6Q1 | CAPN6 | YAKLLGCyEALDGLt | Y174 | 0.059 | 92.62 | 641 | 0.025882 | 0.059 | 0.0259 | 1 |
| Q6GMV2 | SMYD5 | VAGRARVsVEVRFVS | S22 | 0.066 | 92.08 | 418 | 0.052464 | 0.066 | 0.0525 | 1 |
| Q14197 | MRPL58 | QNVNKVNskAEVRFH | S97 | 0.077 | 93.81 | 206 | 0.061208 | 0.077 | 0.0612 | 1 |
| P13591 | NCAM1 | SAASEFKtQPVQGEP | T605 | 0.066 | 88.7 | 858 | 0.045079 | 0.066 | 0.0451 | 1 |
| O15350 | TP73 | stPPPPGtAIRAMPV | T173 | 0.063 | 96.88 | 636 | 0.04303 | 0.063 | 0.043 | 1 |
| Q15173 | PPP2R5B | GVLIEPVyPDIIRMI | Y126 | 0.02 | 95.16 | 497 | 0.008774 | 0.02 | 0.0088 | 1 |
| Q15173 | PPP2R5B | QVARCVSsPHFQVAE | S368 | 0.168 | 95.53 | 497 | 0.133546 | 0.168 | 0.1335 | 1 |
| Q96NU7 | AMDHD1 | GYGLDLEtELKMLRV | T165 | 0.1 | 98.69 | 426 | 0.068301 | 0.1 | 0.0683 | 1 |
| Q8N5V2 | NGEF | LVTSEASyyKsLNLL | Y288 | 0 | 94.57 | 710 | 0 | 0.139 | 0.061 | 1 |
| Q8N5V2 | NGEF | RQIPGDKyQVFDSAP | Y537 | 0.194 | 95.97 | 710 | 0.085105 | 0.194 | 0.0851 | 1 |
| Q9BRJ6 | C7orf50 | GAELALDyLCRWAQK | Y105 | 0.059 | 95.16 | 194 | 0.025882 | 0.059 | 0.0259 | 1 |
| Q9UNS2 | COPS3 | MLESYKKyILVSLIL | Y227 | 0.018 | 97.59 | 423 | 0.007896 | 0.018 | 0.0079 | 1 |
| Q8TEA1 | NSUN6 | AGDVISVySDIkGKC | Y151 | 0.153 | 97.3 | 469 | 0.067119 | 0.153 | 0.0671 | 1 |
| P05388 | RPLP0 | YkRVLALsVETDyTF | S252 | 0.058 | 83.78 | 317 | 0.046105 | 0.058 | 0.0461 | 1 |
| Q9Y2A9 | B3GNT3 | GGGGFLLsRFTAAAL | S276 | 0 | 98.87 | 372 | 0 | 0 | 0 | 0.8182 |
| Q15034 | HERC3 | FVDAYVNyVFQISVH | Y896 | 0.181 | 94.45 | 1050 | 0.079402 | 0.181 | 0.0794 | 0.8 |
| Q15034 | HERC3 | QISVHEWyTAFSSGF | Y906 | 0.223 | 92.41 | 1050 | 0.097827 | 0.223 | 0.0978 | 0.8667 |
| P23378 | GLDC | SETNIVRyMKKLENk | Y537 | 0.037 | 98.8 | 1020 | 0.016231 | 0.037 | 0.0162 | 1 |
| P23378 | GLDC | GKGLKQAtEtAILNA | T830 | 0.018 | 98.81 | 1020 | 0.012294 | 0.018 | 0.0123 | 1 |
| P23378 | GLDC | GLKQAtEtAILNANy | T832 | 0.04 | 98.83 | 1020 | 0.027321 | 0.04 | 0.0273 | 1 |
| P23378 | GLDC | tAILNANyMAKRLET | Y839 | 0.077 | 98.92 | 1020 | 0.033779 | 0.077 | 0.0338 | 1 |
| P23378 | GLDC | RFCDAMIsIRQEIAD | S925 | 0.178 | 98.84 | 1020 | 0.141495 | 0.178 | 0.1415 | 1 |
| Q7Z5N4 | SDK1 | TYSSEYKyIIPsLQK | Y154 | 0.227 | 81.66 | 2213 | 0.099582 | 0.227 | 0.0996 | 0.8333 |
| Q7Z5N4 | SDK1 | QKLDAGFyRCVVRNR | Y167 | 0 | 85.01 | 2213 | 0 | 0.036 | 0.0158 | 1 |
| Q7Z5N4 | SDK1 | RRyEVIMtAYNIIGE | T1678 | 0 | 84.96 | 2213 | 0 | 0 | 0 | 1 |

| | | | | | | | | | | |
|---|---|---|---|---|---|---|---|---|---|---|
| Q8NFY4 | SEMA6D | LLEEIEAyNHAKCSA | Y472 | 0.046 | 95.8 | 1073 | 0.02018 | 0.166 | 0.0728 | 1 |
| Q8NCS7 | SLC44A5 | CPEKFLtyVEMQLLy | Y128 | 0.015 | 81.85 | 719 | 0.00658 | 0.015 | 0.0066 | 1 |
| Q8NCS7 | SLC44A5 | GRAIRyHtGSLAFGS | T513 | 0.21 | 96.37 | 719 | 0.143433 | 0.21 | 0.1434 | 1 |
| Q8NCS7 | SLC44A5 | YGRNFCRsAKDAFNL | S587 | 0.02 | 95.72 | 719 | 0.015898 | 0.02 | 0.0159 | 1 |
| P11908 | PRPS2 | sAGAtKVyAILTHGI | Y245 | 0.173 | 98.79 | 318 | 0.075893 | 0.173 | 0.0759 | 1 |
| P35716 | SOX11 | MNAFMVWsKIERRKI | S60 | 0.014 | 97.67 | 441 | 0.011129 | 0.014 | 0.0111 | 1 |
| O75128 | COBL | EELELSksLNELGIK | S212 | 0.021 | 93.66 | 1261 | 0.016693 | 0.021 | 0.0167 | 1 |
| O14910 | LIN7A | MGGKEQNsPIyISRI | S130 | 0.181 | 93.76 | 233 | 0.14388 | 0.181 | 0.1439 | 0.8571 |
| O14910 | LIN7A | KEQNsPIyISRIIPG | Y133 | 0.054 | 97.59 | 233 | 0.023689 | 0.054 | 0.0237 | 1 |
| O14672 | ADAM10 | VKAIDTIyQTTDFSG | Y257 | 0 | 94.26 | 748 | 0 | 0 | 0 | 1 |
| O14672 | ADAM10 | YSDGKKKsLNTGIIT | S357 | 0 | 91.35 | 748 | 0 | 0 | 0 | 1 |
| O14672 | ADAM10 | NNKFSLCsIRNISQV | S436 | 0.002 | 97.01 | 748 | 0.00159 | 0.002 | 0.0016 | 1 |
| Q0VDF9 | HSPA14 | LtMKRDGsLHVTCTD | S487 | 0.167 | 87.85 | 509 | 0.132751 | 0.167 | 0.1328 | 1 |
| P28838 | LAP3 | ETPANEMtPTRFAEI | T211 | 0.209 | 98.89 | 519 | 0.14275 | 0.209 | 0.1427 | 1 |
| P28838 | LAP3 | GKGITFDsGGISIKA | S288 | 0.077 | 98.63 | 519 | 0.061208 | 0.077 | 0.0612 | 1 |
| P28838 | LAP3 | ADMGGAAtICSAIVS | T311 | 0.001 | 98.94 | 519 | 0.000683 | 0.001 | 0.0007 | 1 |
| P28838 | LAP3 | kTIQVDNtDAEGRLI | T363 | 0.052 | 98.58 | 519 | 0.035517 | 0.052 | 0.0355 | 1 |
| Q2TAC6 | KIF19 | NYRDSKLtRLLKDsL | T304 | 0 | 92.3 | 998 | 0 | 0 | 0 | 0.9167 |
| P27824 | CANX | tLsGWILskAKKDDt | S86 | 0.13 | 91.39 | 592 | 0.103339 | 0.13 | 0.1033 | 1 |
| P27824 | CANX | QFHDkTPyTIMFGPD | Y185 | 0.153 | 95.55 | 592 | 0.067119 | 0.153 | 0.0671 | 1 |
| P27824 | CANX | LELWSMTsDIFFDNF | S431 | 0.067 | 90.73 | 592 | 0.053259 | 0.067 | 0.0533 | 1 |
| P54753 | EPHB3 | TERRVHIsHLLAHTR | S413 | 0.134 | 92.71 | 998 | 0.106519 | 0.134 | 0.1065 | 1 |
| P54753 | EPHB3 | YAAVNITtNQAAPSE | T448 | 0.07 | 85.68 | 998 | 0.047811 | 0.07 | 0.0478 | 1 |
| Q9UBP9 | GULP1 | NAKFLGStEVEQPkG | T35 | 0.13 | 97.38 | 304 | 0.088792 | 0.13 | 0.0888 | 1 |
| Q14651 | PLS1 | LCKMINLsEPDTIDE | S172 | 0.115 | 96.67 | 629 | 0.091415 | 0.115 | 0.0914 | 0.9091 |
| Q14651 | PLS1 | DIKDSRAyFHLLNQI | Y301 | 0.037 | 98.17 | 629 | 0.016231 | 0.037 | 0.0162 | 1 |
| Q14651 | PLS1 | ALVIFQLyEMIRVPV | Y431 | 0 | 98.49 | 629 | 0 | 0 | 0 | 1 |
| P12429 | ANXA3 | RQLIVkEyQAAyGkE | Y60 | 0.002 | 98.3 | 323 | 0.000877 | 0.002 | 0.0009 | 1 |
| P12429 | ANXA3 | MKDISQAyYTVYkks | Y132 | 0 | 98.59 | 323 | 0 | 0 | 0 | 1 |
| P12429 | ANXA3 | LkLTFDEyRNISQkD | Y216 | 0.004 | 98.25 | 323 | 0.001755 | 0.004 | 0.0018 | 1 |
| P12429 | ANXA3 | KSDTsGDyEITLLKI | Y311 | 0.196 | 98.57 | 323 | 0.085982 | 0.196 | 0.086 | 0.8462 |

| | | | | | | | | | | |
|---|---|---|---|---|---|---|---|---|---|---|
| Q9Y694 | SLC22A7 | RYAGRRLtQAGtLLG | T405 | 0.084 | 95.51 | 548 | 0.057373 | 0.084 | 0.0574 | 1 |
| Q9Y694 | SLC22A7 | RRLtQAGtLLGTALA | T409 | 0.132 | 97.14 | 548 | 0.090158 | 0.132 | 0.0902 | 1 |
| Q9HCL2 | GPAM | LLsVVVDtLsTNVIP | T338 | 0.016 | 91.49 | 828 | 0.010928 | 0.016 | 0.0109 | 0.9167 |
| Q7LFX5 | CHST15 | GQNTTDPyLTNSyVL | Y213 | 0.062 | 95.09 | 561 | 0.027199 | 0.062 | 0.0272 | 1 |
| Q7LFX5 | CHST15 | DPyLTNSyVLySKRF | Y218 | 0.15 | 91.57 | 561 | 0.065803 | 0.15 | 0.0658 | 1 |
| P09960 | LTA4H | VDTCSLAsPAsVCRT | S12 | 0.171 | 96.1 | 611 | 0.13593 | 0.171 | 0.1359 | 1 |
| P09960 | LTA4H | sVDFTRRtLTGTAAL | T34 | 0.037 | 98.49 | 611 | 0.025272 | 0.074 | 0.0505 | 1 |
| P09960 | LTA4H | KELVALMsAIRDGET | S172 | 0 | 98.68 | 611 | 0 | 0 | 0 | 1 |
| P09960 | LTA4H | kASMHPVtAMLVGkD | T600 | 0.001 | 88 | 611 | 0.000683 | 0.001 | 0.0007 | 1 |
| Q96NZ1 | FOXN4 | PKPIYSYsCLIAMAL | S199 | 0.141 | 97.65 | 517 | 0.112083 | 0.141 | 0.1121 | 1 |
| Q6ZN66 | GBP6 | VQRAADyySQQMAQR | Y334 | 0.002 | 98 | 633 | 0.000877 | 0.002 | 0.0009 | 1 |
| Q6ZN66 | GBP6 | KERIEQDyWQVPRKG | Y446 | 0.003 | 96.9 | 633 | 0.001316 | 0.003 | 0.0013 | 1 |
| P09525 | ANXA4 | IRRIsQTyQQQyGRS | Y128 | 0.001 | 98.72 | 319 | 0.000439 | 0.001 | 0.0004 | 1 |
| P09525 | ANXA4 | LLHVFDEykRISQkD | Y212 | 0.006 | 98.6 | 319 | 0.002632 | 0.006 | 0.0026 | 1 |
| P29144 | TPP2 | kSTVLRNyKEAQEYG | Y227 | 0.111 | 97.89 | 1249 | 0.048694 | 0.111 | 0.0487 | 1 |
| P29144 | TPP2 | QNKRQMGsGDAYPHQ | S869 | 0.104 | 96.17 | 1249 | 0.082671 | 0.097 | 0.1166 | 1 |
| Q2WGJ9 | FER1L6 | KIGDKPIsFEVSIGN | S501 | 0.089 | 94.7 | 1857 | 0.070747 | 0.055 | 0.0875 | 1 |
| Q9H0C2 | SLC25A31 | GGVAAAVsKtAVAPI | S34 | 0.123 | 95.12 | 315 | 0.097775 | 0.081 | 0.1232 | 1 |
| Q9H0C2 | SLC25A31 | QIsPEARyKGMVDCL | Y63 | 0.074 | 97.57 | 315 | 0.032463 | 0.074 | 0.0325 | 1 |
| Q9H0C2 | SLC25A31 | SGEAKRQyKGtLDCF | Y261 | 0.165 | 96.36 | 315 | 0.072383 | 0.256 | 0.1123 | 0.9167 |
| Q8TE04 | PANK1 | LHFIRFPsCAMHRFI | S314 | 0.135 | 94.85 | 598 | 0.107313 | 0.135 | 0.1073 | 1 |
| Q8TE04 | PANK1 | LGLCCLLtGCEtFEE | T451 | 0.06 | 98.02 | 598 | 0.040981 | 0.06 | 0.041 | 1 |
| P45381 | ASPA | KMsEDLPyEVRRAQE | Y88 | 0.127 | 98.19 | 313 | 0.055713 | 0.127 | 0.0557 | 1 |
| Q49MI3 | CERKL | RLHPRLIsLYGGSME | S546 | 0.067 | 95.42 | 558 | 0.053259 | 0.067 | 0.0533 | 1 |
| Q9NVD3 | SETD4 | DTCALAPyLDLLNHS | Y231 | 0.021 | 96.78 | 440 | 0.009212 | 0.021 | 0.0092 | 1 |
| Q9NVD3 | SETD4 | ELKILRAsAEtLHsL | S428 | 0 | 97.58 | 440 | 0 | 0 | 0 | 1 |
| Q9Y3E7 | CHMP3 | KELVNEWsLKIRKEM | S20 | 0.095 | 97.58 | 222 | 0.075517 | 0.095 | 0.0755 | 1 |
| Q9ULV4 | CORO1C | KGDSSIryFEITDEs | Y292 | 0.013 | 98.05 | 474 | 0.005703 | 0.013 | 0.0057 | 0.8462 |
| O75874 | IDH1 | EMQGDEMtRIIWELI | T19 | 0.023 | 96.21 | 414 | 0.015709 | 0.023 | 0.0157 | 1 |
| O75874 | IDH1 | LSKGWPLyLSTkNtI | Y208 | 0.22 | 98.79 | 414 | 0.096511 | 0.22 | 0.0965 | 1 |
| O75874 | IDH1 | kDIFQEIyDkQYksQ | Y231 | 0.223 | 98.72 | 414 | 0.097827 | 0.223 | 0.0978 | 1 |

| | | | | | | | | | | |
|---|---|---|---|---|---|---|---|---|---|---|
| O75874 | IDH1 | MyQkGQEtSTNPIAS | T325 | 0.038 | 95.91 | 414 | 0.025955 | 0.038 | 0.026 | 1 |
| O75874 | IDH1 | PNVQRsDyLNTFEFM | Y391 | 0.28 | 97.37 | 414 | 0.122832 | 0.28 | 0.1228 | 1 |
| O96005 | CLPTM1 | FDAVSGDyYPIIyFN | Y275 | 0.007 | 91.1 | 669 | 0.003071 | 0.007 | 0.0031 | 0.8 |
| O96005 | CLPTM1 | GDyYPIIyFNDyWNL | Y280 | 0.048 | 91.39 | 669 | 0.021057 | 0.048 | 0.0211 | 0.9091 |
| P25098 | GRK2 | FyEEIKKyEkLETEE | Y92 | 0.003 | 95.89 | 689 | 0.001316 | 0.003 | 0.0013 | 1 |
| P25098 | GRK2 | AsVGtHGyMAPEVLQ | Y356 | 0.22 | 96.75 | 689 | 0.096511 | 0.22 | 0.0965 | 1 |
| Q9UQE7 | SMC3 | ICRsMEVsTQLArAF | S639 | 0 | 87.05 | 1217 | 0 | 0 | 0 | 1 |
| Q9UQE7 | SMC3 | GyyDTRKsRLELQKD | S674 | 0.103 | 88.42 | 1217 | 0.081876 | 0.103 | 0.0819 | 1 |
| Q9BY07 | SLC4A5 | IPDLALMsFILFFGT | S747 | 0.005 | 90.04 | 1137 | 0.003975 | 0.005 | 0.004 | 1 |
| P08887 | IL6R | RSVQLHDsGNYSCYR | S91 | 0.132 | 93.73 | 468 | 0.104929 | 0.132 | 0.1049 | 1 |
| P08887 | IL6R | RStPsLTtKAVLLVR | T144 | 0.157 | 95.34 | 468 | 0.107233 | 0.157 | 0.1072 | 0.8889 |
| Q5SZK8 | FREM2 | INHHKIAyRPPGQEL | Y786 | 0.001 | 94.475 | 3169 | 0.000439 | 0.001 | 0.0004 | 1 |
| Q9UHX1 | PUF60 | KGFAFVEyEVPEAAQ | Y177 | 0.018 | 93.02 | 559 | 0.007896 | 0.018 | 0.0079 | 1 |
| O15027 | SEC16A | QVFKFIYsCRLAEMG | S1810 | 0.009 | 95.07 | 2357 | 0.007154 | 0.009 | 0.0072 | 0.8182 |
| Q15029 | EFTUD2 | RYDQDLCytDILFTE | Y167 | 0.019 | 92.07 | 972 | 0.008335 | 0.019 | 0.0083 | 1 |
| Q15029 | EFTUD2 | YDQDLCytDILFTEQ | T168 | 0.018 | 93.62 | 972 | 0.012294 | 0.018 | 0.0123 | 1 |
| Q15029 | EFTUD2 | RGVGIkStPVTVVLP | T184 | 0.056 | 96.49 | 972 | 0.038249 | 0.056 | 0.0382 | 1 |
| Q15029 | EFTUD2 | kRLWGDIyFNPktRK | Y348 | 0.253 | 94.46 | 972 | 0.110987 | 0.253 | 0.111 | 1 |
| Q15029 | EFTUD2 | HAFGRVLsGTIHAGQ | S498 | 0.001 | 95.75 | 972 | 0.000795 | 0.001 | 0.0008 | 1 |
| Q15029 | EFTUD2 | TVGRLWIsVARYHIE | S533 | 0.072 | 95.13 | 972 | 0.057234 | 0.072 | 0.0572 | 1 |
| Q15029 | EFTUD2 | VDQPIVktAtItEPr | T562 | 0.069 | 94.05 | 972 | 0.047128 | 0.069 | 0.0471 | 1 |
| Q15029 | EFTUD2 | QPIVktAtItEPrGN | T564 | 0.017 | 96.99 | 972 | 0.011611 | 0.017 | 0.0116 | 1 |
| Q15029 | EFTUD2 | IVktAtItEPrGNEE | T566 | 0.009 | 95.72 | 972 | 0.006147 | 0.009 | 0.0061 | 1 |
| Q15029 | EFTUD2 | HDLRkMYsEIDIKVA | S649 | 0.15 | 89.61 | 972 | 0.119237 | 0.15 | 0.1192 | 1 |
| Q15029 | EFTUD2 | PNILVDDtLPsEVDk | T749 | 0.094 | 90.74 | 972 | 0.064203 | 0.094 | 0.0642 | 1 |
| P47870 | GABRB2 | WRDKRLSyNVIPLNL | Y98 | 0.024 | 92.84 | 512 | 0.010528 | 0.024 | 0.0105 | 1 |
| P55010 | EIF5 | kkQTIGNsCKACGYR | S121 | 0.138 | 86.16 | 431 | 0.109698 | 0.138 | 0.1097 | 0.8571 |
| Q7Z4W1 | DCXR | NHsVyCstkGALDML | T152 | 0 | 98.73 | 244 | 0 | 0 | 0 | 1 |
| Q86UT6 | NLRX1 | LEDsRGAtLNPWRKA | T947 | 0.17 | 75.98 | 975 | 0.116112 | 0.17 | 0.1161 | 1 |
| Q9NSV4 | DIAPH3 | IKVNENkyENVDLLC | Y673 | 0.164 | 86.29 | 1193 | 0.071944 | 0.164 | 0.0719 | 1 |
| Q9Y3F4 | STRAP | PDGELYAsGSEDGTL | S282 | 0 | 98.68 | 350 | 0 | 0 | 0 | 0.8 |

| | | | | | | | | | | |
|---|---|---|---|---|---|---|---|---|---|---|
| Q8ND04 | SMG8 | HQLCEERsLTDQHCV | S570 | 0 | 94.47 | 991 | 0 | 0.083 | 0.066 | 1 |
| Q14CM0 | FRMPD4 | VGPRyGIsHVINtKT | S448 | 0.04 | 96.33 | 1322 | 0.031797 | 0.04 | 0.0318 | 1 |
| Q15061 | WDR43 | IPLLQELtkRLQGHP | T510 | 0.175 | 85.37 | 677 | 0.119527 | 0.175 | 0.1195 | 1 |
| Q8WZ82 | OVCA2 | CLAGFRQsERGFREK | S18 | 0.088 | 94.06 | 227 | 0.069952 | 0.175 | 0.1391 | 1 |
| Q8WZ82 | OVCA2 | PRFILLVsGFCPRGI | S149 | 0.001 | 98.54 | 227 | 0.000795 | 0.001 | 0.0008 | 1 |
| Q8N3J2 | METTL4 | ItLMGQKyLLPPKSS | Y255 | 0.034 | 94.03 | 472 | 0.014915 | 0.341 | 0.1496 | 1 |
| Q8N3J2 | METTL4 | PKSSFLLsDISCMQP | S266 | 0.05 | 93.16 | 472 | 0.039746 | 0.115 | 0.0914 | 1 |
| A7E2Y1 | MYH7B | NTKRVIQyFAIVAAL | Y235 | 0.004 | 88.13 | 1983 | 0.001755 | 0.004 | 0.0018 | 1 |
| A6NHZ5 | LRRC14B | KFLGNPLsARALRRL | S401 | 0.092 | 96.92 | 514 | 0.073132 | 0.092 | 0.0731 | 1 |
| Q9NQW1 | SEC31B | PEDIKALsWNRQAQH | S175 | 0.065 | 96.29 | 1179 | 0.051669 | 0.065 | 0.0517 | 1 |
| P18031 | PTPN1 | DNDyINAsLIkMEEA | S70 | 0 | 98.94 | 435 | 0 | 0 | 0 | 0.8 |
| P18031 | PTPN1 | AGIGRsGtFCLADTC | T224 | 0.006 | 98.81 | 435 | 0.004098 | 0.006 | 0.0041 | 0.8333 |
| A6NGN9 | IGLON5 | VQTERTRsMLLFANV | S274 | 0 | 96.53 | 336 | 0 | 0 | 0 | 1 |
| Q9NYI0 | PSD3 | LEFEKTRyEMYVsIL | Y981 | 0.063 | 95.68 | 1048 | 0.027637 | 0.297 | 0.1303 | 0.9091 |
| Q6PIF6 | MYO7B | VHNLLIRyQQHKIYT | Y89 | 0.07 | 91.42 | 2116 | 0.030708 | 0.07 | 0.0307 | 1 |
| P48553 | TRAPPC10 | IPQRVKFtVttGHYT | T807 | 0.167 | 87.15 | 1259 | 0.114063 | 0.167 | 0.1141 | 1 |
| P48553 | TRAPPC10 | RVKFtVttGHYTIKN | T810 | 0.013 | 79.48 | 1259 | 0.008879 | 0.013 | 0.0089 | 1 |
| Q9NSI6 | BRWD1 | RPVPLIEsELyFLIA | S16 | 0.072 | 87.06 | 2320 | 0.057234 | 0.072 | 0.0572 | 1 |
| O96017 | CHEK2 | IAYIEDHsGNGTFVN | S164 | 0.022 | 96.39 | 543 | 0.017488 | 0.022 | 0.0175 | 0.8571 |
| O00139 | KIF2A | NQTFRFDyAFDDSAP | Y278 | 0.023 | 97.87 | 706 | 0.01009 | 0.023 | 0.0101 | 1 |
| O00139 | KIF2A | EGAEINKsLLALkEC | S484 | 0.087 | 91.5 | 706 | 0.069158 | 0.087 | 0.0692 | 1 |
| P51124 | GZMM | RVLARVLsFsSRVCT | S224 | 0.045 | 94.14 | 257 | 0.035771 | 0.045 | 0.0358 | 0.8571 |
| A5D6W6 | FITM1 | ATRRVAVtARHLSRL | T121 | 0.098 | 91.9 | 292 | 0.066935 | 0.098 | 0.0669 | 1 |
| Q9BZC7 | ABCA2 | TSTFKMLtGDESTTG | T2101 | 0.007 | 86.43 | 2435 | 0.004781 | 0.111 | 0.0699 | 1 |
| Q9BZC7 | ABCA2 | EAFVNGHsVLKELLQ | S2117 | 0.053 | 88.48 | 2435 | 0.04213 | 0.053 | 0.0421 | 1 |
| Q9BZC7 | ABCA2 | ADKPAGTySGGNKRK | Y2186 | 0.18 | 74.93 | 2435 | 0.078963 | 0.282 | 0.0959 | 1 |
| Q86VZ1 | P2RY8 | VLLAHIVsRLFYGKS | S263 | 0.087 | 89.46 | 359 | 0.069158 | 0.087 | 0.0692 | 1 |
| O60488 | ACSL4 | RRVNNFGsGLtALGL | S140 | 0.041 | 98.69 | 711 | 0.032592 | 0.064 | 0.0529 | 1 |
| O60488 | ACSL4 | SCVkHIIyVDNkAIN | Y227 | 0.226 | 97.83 | 711 | 0.099143 | 0.226 | 0.0991 | 1 |
| O60488 | ACSL4 | LAHVLELtAEISCFT | T328 | 0.018 | 98.04 | 711 | 0.012294 | 0.018 | 0.0123 | 0.8 |
| O60488 | ACSL4 | EEkTAEDySVDENGQ | Y541 | 0.171 | 98.32 | 711 | 0.075015 | 0.171 | 0.075 | 1 |

| | | | | | | | | | | |
|---|---|---|---|---|---|---|---|---|---|---|
| O60488 | ACSL4 | IPIkVRLsPEPWtPE | S674 | 0.047 | 95.86 | 711 | 0.037361 | 0.047 | 0.0374 | 1 |
| O00755 | WNT7A | EEDPVtGsVGTQGRA | S286 | 0.07 | 88.59 | 349 | 0.055644 | 0.083 | 0.066 | 1 |
| Q8N271 | PROM2 | ARILRNVsECFLARE | S727 | 0.093 | 89.13 | 834 | 0.073927 | 0.138 | 0.1097 | 1 |
| Q8N271 | PROM2 | IIFAVKtsKyFRPIR | S803 | 0.01 | 87.78 | 834 | 0.007949 | 0.058 | 0.0461 | 1 |
| Q8N5D0 | WDTC1 | SGPFVRLyDIRMIHN | Y212 | 0.014 | 97.41 | 677 | 0.006142 | 0.014 | 0.0061 | 1 |
| Q8TDX7 | NEK7 | VGtPyyMsPERIHEN | S204 | 0.004 | 91.28 | 302 | 0.00318 | 0.004 | 0.0032 | 1 |
| Q6N021 | TET2 | kkLAPDAyNNQIEYE | Y1345 | 0.127 | 98.74 | 2002 | 0.055713 | 0.127 | 0.0557 | 1 |
| Q9Y2X3 | NOP58 | LFETSVGyAIFKVLN | Y11 | 0.115 | 90.16 | 529 | 0.050449 | 0.115 | 0.0504 | 1 |
| Q86X95 | CIR1 | KkDFHPAskSNIKkV | S18 | 0.094 | 91.28 | 450 | 0.074722 | 0.094 | 0.0747 | 1 |
| P11182 | DBT | kAQIMNVsWSADHRV | S447 | 0.011 | 97.11 | 482 | 0.008744 | 0.011 | 0.0087 | 1 |
| Q15046 | KARS1 | CDHPQIMsPLAKWHR | S470 | 0.049 | 97.2 | 597 | 0.038951 | 0.049 | 0.039 | 1 |
| Q15046 | KARS1 | VAMFLTDsNNIKEVL | S561 | 0.026 | 98.27 | 597 | 0.020668 | 0.026 | 0.0207 | 1 |
| Q9H6R4 | NOL6 | SLRDtAAsLRLERDT | S824 | 0 | 91.95 | 1146 | 0 | 0 | 0 | 1 |
| O60885 | BRD4 | VkLNLPDyykIIKtP | Y97 | 0.175 | 96.77 | 1362 | 0.07677 | 0.311 | 0.1269 | 0.8 |
| O00160 | MYO1F | ENQCVIIsGESGAGk | S109 | 0.15 | 88.49 | 1098 | 0.119237 | 0.15 | 0.1192 | 1 |
| O00160 | MYO1F | VNMEPDQyQMGsTkV | Y667 | 0.291 | 90.28 | 1098 | 0.127658 | 0.291 | 0.1277 | 1 |
| O00160 | MYO1F | FADSVTKyDRRFKPI | Y771 | 0.142 | 94.81 | 1098 | 0.062293 | 0.142 | 0.0623 | 1 |
| O00160 | MYO1F | ILTPKCVyVIGREKV | Y790 | 0.021 | 95.87 | 1098 | 0.009212 | 0.021 | 0.0092 | 1 |
| Q13206 | DDX10 | NINILVCtPGRLLQH | T197 | 0.131 | 93.8 | 875 | 0.089475 | 0.131 | 0.0895 | 1 |
| Q13206 | DDX10 | LLQHMDEtVSFHATD | T208 | 0.126 | 88.79 | 875 | 0.08606 | 0.126 | 0.0861 | 1 |
| Q13206 | DDX10 | NLPKKRQtLLFSAtQ | T249 | 0.012 | 97.06 | 875 | 0.008196 | 0.012 | 0.0082 | 1 |
| Q9P273 | TENM3 | GTDLKLSyLSSRAAG | Y1006 | 0.05 | 95.38 | 2699 | 0.021934 | 0.05 | 0.0219 | 1 |
| Q9P273 | TENM3 | RTAILQGyELDASNM | Y1104 | 0.246 | 92.72 | 2699 | 0.107917 | 0.246 | 0.1079 | 1 |
| Q9P273 | TENM3 | DVQNGILyKGNGENQ | Y1130 | 0.017 | 95.25 | 2699 | 0.007458 | 0.017 | 0.0075 | 1 |
| Q9P273 | TENM3 | LyVGDFNyVRRIFPS | Y1193 | 0.194 | 91.89 | 2699 | 0.085105 | 0.309 | 0.1356 | 1 |
| Q9P273 | TENM3 | RMPVRVVsPDNQVIW | S1579 | 0.009 | 87.97 | 2699 | 0.007154 | 0.009 | 0.0072 | 1 |
| Q9P273 | TENM3 | ETAGVLKtVNLQsDG | T1979 | 0.217 | 95.38 | 2699 | 0.148214 | 0.217 | 0.1482 | 1 |
| Q9P273 | TENM3 | PFANttKyAYEYDVD | Y2128 | 0.219 | 95.27 | 2699 | 0.096072 | 0.219 | 0.0961 | 1 |
| Q9P273 | TENM3 | RGTEIFEySSKGLLT | Y2207 | 0.228 | 95.05 | 2699 | 0.10002 | 0.228 | 0.1 | 1 |
| Q9P273 | TENM3 | GSGWTVIyRyDGLGR | Y2227 | 0.096 | 95.61 | 2699 | 0.042114 | 0.096 | 0.0421 | 1 |
| Q9P273 | TENM3 | GWTVIyRyDGLGRRV | Y2229 | 0.137 | 95.53 | 2699 | 0.0601 | 0.137 | 0.0601 | 1 |

| | | | | | | | | | | |
|---|---|---|---|---|---|---|---|---|---|---|
| P0DJI8 | SAA1 | NyIGSDKyFHARGNY | Y53 | 0.097 | 98.63 | 122 | 0.042553 | 0.097 | 0.0426 | 1 |
| P0DJI8 | SAA1 | FGHGAEDsLADQAAN | S94 | 0.014 | 92.63 | 122 | 0.011129 | 0.014 | 0.0111 | 1 |
| P34949 | MPI | QIAEDkPyAELWMGT | Y46 | 0.079 | 98.75 | 423 | 0.034656 | 0.158 | 0.0693 | 1 |
| P34949 | MPI | PFLFkVLsVETPLsI | S102 | 0 | 98.88 | 423 | 0 | 0 | 0 | 1 |
| P34949 | MPI | YKVLALDsASILLMV | S366 | 0.021 | 97.45 | 423 | 0.016693 | 0.021 | 0.0167 | 1 |
| O43464 | HTRA2 | IHkVILGsPAHRAGL | S400 | 0.039 | 95.13 | 458 | 0.031002 | 0.039 | 0.031 | 1 |
| Q5T3U5 | ABCC10 | TIAHRLNtILNSDRV | T1442 | 0.191 | 71.25 | 1492 | 0.130456 | 0.191 | 0.1305 | 1 |
| P55809 | OXCT1 | QIKRMVSsyVGENAE | S114 | 0 | 97.87 | 520 | 0 | 0 | 0 | 0.8 |
| P55809 | OXCT1 | NAEFERQyLSGELEV | Y126 | 0.026 | 95.12 | 520 | 0.011406 | 0.026 | 0.0114 | 0.9286 |
| P55809 | OXCT1 | EGGsPIkyNkDGSVA | Y174 | 0.301 | 95.7 | 520 | 0.132044 | 0.301 | 0.132 | 0.8333 |
| P55809 | OXCT1 | GAMDLVssAKtKVVV | S432 | 0.15 | 97.42 | 520 | 0.119237 | 0.15 | 0.1192 | 1 |
| P55809 | OXCT1 | DLVssAKtKVVVTME | T435 | 0.063 | 98.65 | 520 | 0.04303 | 0.063 | 0.043 | 1 |
| Q8WYL5 | SSH1 | MGVSRsAstVIAyAM | S402 | 0 | 98.17 | 1049 | 0 | 0 | 0 | 1 |
| Q8WYL5 | SSH1 | GVSRsAstVIAyAMK | T403 | 0.021 | 98.19 | 1049 | 0.014343 | 0.021 | 0.0143 | 1 |
| Q8WYL5 | SSH1 | VKQKRsItRPNAGFM | T430 | 0.022 | 93.33 | 1049 | 0.015026 | 0.022 | 0.015 | 1 |
| Q8WYL5 | SSH1 | FMRQLsEyEGILDAS | Y443 | 0.081 | 96.66 | 1049 | 0.035534 | 0.081 | 0.0355 | 1 |
| Q5JU85 | IQSEC2 | KYGGsFLsRRAARTI | S347 | 0.175 | 82.96 | 1488 | 0.13911 | 0.175 | 0.1391 | 1 |
| Q9UDY4 | DNAJB4 | PKFKRDGsNIIyTAK | S248 | 0.174 | 96.43 | 337 | 0.138315 | 0.174 | 0.1383 | 1 |
| O60806 | TBX19 | SFSKVKLtNKLNGGG | T148 | 0.016 | 95.08 | 448 | 0.010928 | 0.016 | 0.0109 | 1 |
| Q14995 | NR1D2 | LFDAKERtVtFLSGk | T460 | 0.198 | 91.43 | 579 | 0.135237 | 0.198 | 0.1352 | 1 |
| O00425 | IGF2BP3 | EksITILstPEGTSA | S248 | 0.154 | 89.13 | 579 | 0.122417 | 0.154 | 0.1224 | 1 |
| Q9UJX6 | ANAPC2 | IVRDFPDsRPAIEDL | S354 | 0.008 | 94.92 | 822 | 0.006359 | 0.158 | 0.1256 | 1 |
| Q9NTX5 | ECHDC1 | ACDFRLMtPESkIRF | T175 | 0.029 | 98.51 | 307 | 0.019807 | 0.029 | 0.0198 | 1 |
| Q14919 | DRAP1 | AAVPVIIsRALELFL | S39 | 0.018 | 97.3 | 205 | 0.014308 | 0.018 | 0.0143 | 0.8889 |
| Q92824 | PCSK5 | IGALKDyyHFyHSRT | Y72 | 0.032 | 90.16 | 1860 | 0.014038 | 0.032 | 0.014 | 1 |
| Q9UHL9 | GTF2IRD1 | PFKRPCtyGVPKLKR | Y404 | 0.148 | 93.04 | 959 | 0.064925 | 0.148 | 0.0649 | 0.9231 |
| Q9H1E1 | RNASE7 | EKRQNKSyVVACKPP | Y130 | 0.032 | 98.74 | 156 | 0.014038 | 0.032 | 0.014 | 0.875 |
| Q96M27 | PRRC1 | QDYNLRWsGLLVTVG | S390 | 0.013 | 97.78 | 445 | 0.010334 | 0.013 | 0.0103 | 1 |
| Q96M27 | PRRC1 | ARAIAGMyKQRLPPR | Y436 | 0.02 | 96.56 | 445 | 0.008774 | 0.02 | 0.0088 | 1 |
| Q9NP99 | TREM1 | MPKtLACtERPsKNS | T70 | 0.074 | 94.93 | 234 | 0.050543 | 0.074 | 0.0505 | 1 |
| Q9NP99 | TREM1 | LACtERPsKNSHPVQ | S74 | 0.085 | 93.35 | 234 | 0.067568 | 0.085 | 0.0676 | 1 |

| | | | | | | | | | | |
|---|---|---|---|---|---|---|---|---|---|---|
| Q38SD2 | LRRK1 | TQQTEEQyFQFLAKF | Y971 | 0 | 86.76 | 2015 | 0 | 0.468 | 0.1398 | 1 |
| Q86W50 | METTL16 | VkRIIHDsLQLkKRL | S235 | 0 | 96.46 | 562 | 0 | 0 | 0 | 1 |
| Q13838 | DDX39B | SkEYERFskYMPNVK | S137 | 0.135 | 97.94 | 428 | 0.107313 | 0.135 | 0.1073 | 1 |
| Q13838 | DDX39B | CPHIVVGtPGRILAL | T172 | 0.159 | 96.67 | 428 | 0.108599 | 0.159 | 0.1086 | 1 |
| Q13838 | DDX39B | EKQVMMFsATLSKEI | S228 | 0.025 | 91.76 | 428 | 0.019873 | 0.043 | 0.0342 | 1 |
| Q96PQ0 | SORCS2 | DMGKVLEssLWRsSD | S179 | 0.173 | 90.75 | 1159 | 0.13752 | 0.173 | 0.1375 | 1 |
| Q96PQ0 | SORCS2 | MGKVLEssLWRsSDF | S180 | 0.102 | 91.23 | 1159 | 0.081081 | 0.102 | 0.0811 | 1 |
| Q96PQ0 | SORCS2 | LEssLWRsSDFGTSY | S184 | 0.003 | 93.7 | 1159 | 0.002385 | 0.003 | 0.0024 | 1 |
| Q96PQ0 | SORCS2 | RDQSLFLsADEGAtF | S234 | 0.018 | 94.84 | 1159 | 0.014308 | 0.018 | 0.0143 | 1 |
| Q96PQ0 | SORCS2 | EDKVLAYtKESKLYV | T267 | 0.004 | 93.87 | 1159 | 0.002732 | 0.004 | 0.0027 | 1 |
| O15235 | MRPS12 | PPRKLGPtEGRPQLK | T52 | 0.121 | 95.06 | 138 | 0.082645 | 0.121 | 0.0826 | 1 |
| P42261 | GRIA1 | LEKNGIGyHyILANL | Y226 | 0.203 | 88.8 | 906 | 0.089053 | 0.203 | 0.0891 | 1 |
| P42261 | GRIA1 | KNGIGyHyILANLGF | Y228 | 0.004 | 96.28 | 906 | 0.001755 | 0.004 | 0.0018 | 1 |
| P42261 | GRIA1 | KYTSALTyDGVKVMA | Y294 | 0.267 | 94.35 | 906 | 0.117129 | 0.267 | 0.1171 | 1 |
| Q9Y6D5 | ARFGEF2 | NKMTkEQyIKMNRGI | Y797 | 0 | 92.25 | 1785 | 0 | 0 | 0 | 0.9231 |
| O60658 | PDE8A | SEEYFSQtDEEKQQG | T752 | 0.027 | 95.2 | 829 | 0.018441 | 0.027 | 0.0184 | 1 |
| Q96NB2 | SFXN2 | SVRQMALsyFTATTT | S149 | 0.004 | 90.71 | 322 | 0.00318 | 0.004 | 0.0032 | 1 |
| Q96NB2 | SFXN2 | GItQVVIsRITMSAP | S231 | 0.086 | 91.91 | 322 | 0.068363 | 0.086 | 0.0684 | 0.8 |
| Q6PCE3 | PGM2L1 | SQRLAKLtAAVLLAK | T134 | 0 | 98.8 | 622 | 0 | 0 | 0 | 1 |
| Q6PCE3 | PGM2L1 | LAKDVPVyLFSRYVP | Y146 | 0.308 | 98.8 | 622 | 0.135115 | 0.308 | 0.1351 | 1 |
| Q6PCE3 | PGM2L1 | GEsVLELsLRLAEKE | S316 | 0.006 | 98.28 | 622 | 0.004769 | 0.006 | 0.0048 | 1 |
| Q6PCE3 | PGM2L1 | ILHVRDVttGYDSSQ | T532 | 0.129 | 94.92 | 622 | 0.088109 | 0.129 | 0.0881 | 1 |
| P00915 | CA1 | sVsYNPAtAKEIINV | T56 | 0 | 98.58 | 261 | 0 | 0 | 0 | 1 |
| P00915 | CA1 | FEDNDNRsVLKGGPF | S78 | 0.017 | 98.68 | 261 | 0.013514 | 0.017 | 0.0135 | 0.8889 |
| P00915 | CA1 | GGPFsDsyRLFQFHF | Y89 | 0.081 | 98.75 | 261 | 0.035534 | 0.081 | 0.0355 | 0.8462 |
| P15559 | NQO1 | AykEGHLsPDIVAEQ | S82 | 0.176 | 98.69 | 274 | 0.139905 | 0.176 | 0.1399 | 1 |
| P15559 | NQO1 | kNKkFGLsVGHHLGk | S255 | 0 | 98.78 | 274 | 0 | 0 | 0 | 0.8889 |
| O14929 | HAT1 | DDETAFGykGLKILL | Y69 | 0.159 | 95.84 | 419 | 0.069751 | 0.256 | 0.1123 | 0.9286 |
| Q86VD7 | SLC25A42 | QVLssLLsGALAGAL | S39 | 0 | 90.48 | 318 | 0 | 0 | 0 | 1 |
| Q86VD7 | SLC25A42 | VVRRRMQtAGVTGyP | T256 | 0 | 97 | 318 | 0 | 0 | 0 | 1 |
| Q8N1G2 | CMTR1 | HAKGFGMtLkGPNDF | T301 | 0.083 | 97 | 835 | 0.05669 | 0.083 | 0.0567 | 1 |

| | | | | | | | | | | |
|---|---|---|---|---|---|---|---|---|---|---|
| Q8N1G2 | CMTR1 | GGHFICKtFDLFTPF | T405 | 0.065 | 97.18 | 835 | 0.044396 | 0.065 | 0.0444 | 1 |
| Q8N1G2 | CMTR1 | PIRVkEVyRLEEMEk | Y707 | 0.278 | 96.4 | 835 | 0.121955 | 0.278 | 0.122 | 1 |
| Q9UHQ1 | NARF | EWLEGINsPKAREVL | S431 | 0.152 | 95.83 | 456 | 0.120827 | 0.152 | 0.1208 | 1 |
| Q96M93 | ADAD1 | DGLSGKItESSPFkS | T494 | 0.208 | 94.15 | 576 | 0.142067 | 0.208 | 0.1421 | 1 |
| Q96M93 | ADAD1 | kSGMsMAsRLCkAAM | S507 | 0.008 | 97.94 | 576 | 0.006359 | 0.008 | 0.0064 | 1 |
| P51512 | MMP16 | TGKVDRNtIDWMKKP | T92 | 0.006 | 89.67 | 607 | 0.004098 | 0.006 | 0.0041 | 0.8182 |
| Q5TEU4 | NDUFAF5 | MLAAAAVyREMyRNE | Y286 | 0.013 | 98.24 | 345 | 0.005703 | 0.013 | 0.0057 | 1 |
| Q8TAE8 | GADD45GIP[illegible] | PDPEDLLtPRWQLGP | T47 | 0.207 | 94.49 | 222 | 0.141384 | 0.207 | 0.1414 | 1 |
| Q03001 | DST | ANRVQRDsVICEDkL | S397 | 0.004 | 94.28 | 7570 | 0.00318 | 0.004 | 0.0032 | 1 |
| Q13162 | PRDX4 | THQISKDyGVyLEDS | Y188 | 0.059 | 98.75 | 271 | 0.025882 | 0.059 | 0.0259 | 1 |
| Q93033 | CD101 | LEKMDWAtFQLEItF | T493 | 0.145 | 88.54 | 1021 | 0.099037 | 0.145 | 0.099 | 1 |
| Q93033 | CD101 | FQLEItFtAItDSGT | T501 | 0.045 | 80.46 | 1021 | 0.030736 | 0.045 | 0.0307 | 1 |
| Q93033 | CD101 | AItDSGTyECRVsEK | Y509 | 0.002 | 91.05 | 1021 | 0.000877 | 0.042 | 0.0184 | 1 |
| Q13724 | MOGS | FPPVPLFtAVPSRSF | T436 | 0.01 | 98.86 | 837 | 0.00683 | 0.01 | 0.0068 | 1 |
| Q13724 | MOGS | ADsRHLWsPFGLRSL | S724 | 0.007 | 98.79 | 837 | 0.005564 | 0.007 | 0.0056 | 1 |
| Q8IWV2 | CNTN4 | ENARKIRtVGNQTST | T855 | 0.045 | 89.86 | 1026 | 0.030736 | 0.045 | 0.0307 | 1 |
| Q6PKC3 | TXNDC11 | EVVLLFFyAPWCGQS | Y133 | 0.011 | 97.24 | 985 | 0.004826 | 0.011 | 0.0048 | 1 |
| Q4ZHG4 | FNDC1 | PVYRAALtKRKIsEE | T250 | 0.041 | 85.3 | 1894 | 0.028004 | 0.041 | 0.028 | 0.9231 |
| Q9NY15 | STAB1 | PFTVFAPsNEAVDsL | S547 | 0.049 | 84.71 | 2570 | 0.038951 | 0.049 | 0.039 | 1 |
| O60825 | PFKFB2 | ILEVKVSsPDYPERN | S175 | 0.15 | 83.35 | 505 | 0.119237 | 0.15 | 0.1192 | 1 |
| O60825 | PFKFB2 | ALRDQEKyLyRYPGG | Y356 | 0.198 | 94.76 | 505 | 0.08686 | 0.198 | 0.0869 | 0.9091 |
| O60825 | PFKFB2 | RDQEKyLyRYPGGES | Y358 | 0.234 | 93.89 | 505 | 0.102652 | 0.234 | 0.1027 | 1 |
| O60825 | PFKFB2 | RYPGGESyQDLVQRL | Y366 | 0.168 | 96.93 | 505 | 0.073699 | 0.168 | 0.0737 | 1 |
| Q9H6R0 | DHX33 | ItITGIKyVVDtGMV | Y368 | 0.18 | 97.15 | 707 | 0.078963 | 0.18 | 0.079 | 1 |
| Q9H6R0 | DHX33 | GIKyVVDtGMVkAKk | T372 | 0.057 | 97.86 | 707 | 0.038932 | 0.057 | 0.0389 | 1 |
| Q9H6R0 | DHX33 | GMVkAKkyNPDSGLE | Y380 | 0.243 | 88.1 | 707 | 0.106601 | 0.243 | 0.1066 | 1 |
| Q9H6R0 | DHX33 | SGICYRLytEDEFEk | Y418 | 0.077 | 97.99 | 707 | 0.033779 | 0.077 | 0.0338 | 1 |
| Q9H6R0 | DHX33 | GICYRLytEDEFEkF | T419 | 0.151 | 97.32 | 707 | 0.103135 | 0.151 | 0.1031 | 1 |
| Q9H6R0 | DHX33 | MsMPIAssRGDVESV | S613 | 0.177 | 93.35 | 707 | 0.1407 | 0.177 | 0.1407 | 1 |
| Q9Y3B7 | MRPL11 | KEGIPLPtKILVkPD | T68 | 0.024 | 90.03 | 192 | 0.016392 | 0.024 | 0.0164 | 1 |
| Q8WY54 | PPM1E | VNGSLSVsRAIGDAE | S400 | 0 | 98.12 | 755 | 0 | 0 | 0 | 1 |

| | | | | | | | | | | |
|---|---|---|---|---|---|---|---|---|---|---|
| P39748 | FEN1 | MDCLTFGsPVLMrHL | S187 | 0 | 98.75 | 380 | 0 | 0 | 0 | 1 |
| P39748 | FEN1 | RRLDPNkyPVPENWL | Y268 | 0.272 | 97.09 | 380 | 0.119322 | 0.272 | 0.1193 | 1 |
| P43897 | TSFM | VKVPSGFyVGSyVHG | Y212 | 0.313 | 97.13 | 325 | 0.137309 | 0.313 | 0.1373 | 0.9286 |
| P48163 | ME1 | VGLACQQySLVFRKP | Y113 | 0.274 | 94.37 | 572 | 0.1202 | 0.274 | 0.1202 | 1 |
| P48163 | ME1 | VIKAIVVtDGERILG | T151 | 0 | 98.69 | 572 | 0 | 0 | 0 | 1 |
| P48163 | ME1 | ELLKDPLyIGLRQRR | Y208 | 0.104 | 97.91 | 572 | 0.045623 | 0.104 | 0.0456 | 1 |
| P48163 | ME1 | LNTIRDVsLkIAEkI | S510 | 0.001 | 98.64 | 572 | 0.000795 | 0.001 | 0.0008 | 1 |
| P59910 | DNAJB13 | VLGITRNsEDAQIKQ | S15 | 0.069 | 85.08 | 316 | 0.054849 | 0.069 | 0.0548 | 0.8333 |
| Q9UJ72 | ANXA10 | RMMIAEAyQSMyGRD | Y59 | 0.086 | 98.61 | 324 | 0.037727 | 0.123 | 0.054 | 1 |
| Q8NE09 | RGS22 | LEKMKKCyLVSNGDy | Y460 | 0.019 | 86.98 | 1264 | 0.008335 | 0.019 | 0.0083 | 1 |
| O00410 | IPO5 | kEGFVEytEQVVKLM | T700 | 0 | 95.87 | 1097 | 0 | 0 | 0 | 1 |
| Q9H013 | ADAM19 | ANYVDKFyRsLNIRI | Y248 | 0 | 93.52 | 955 | 0 | 0.091 | 0.0399 | 1 |
| O00232 | PSMD12 | MVQQCCTyVEEITDL | Y111 | 0.265 | 79.59 | 456 | 0.116252 | 0.265 | 0.1163 | 1 |
| O00232 | PSMD12 | kFFQEENtEkLKLKY | T219 | 0.05 | 77.28 | 456 | 0.034151 | 0.05 | 0.0342 | 1 |
| Q9BV19 | C1orf50 | KREsGQQyFSIISPK | Y131 | 0.277 | 97.18 | 199 | 0.121516 | 0.277 | 0.1215 | 1 |
| Q99595 | TIMM17A | AIkGFRNsPVGVNHR | S40 | 0.175 | 92.99 | 171 | 0.13911 | 0.175 | 0.1391 | 1 |
| P60983 | GMFB | QPRFIVYsYkYQHDD | S72 | 0.036 | 97.77 | 142 | 0.028617 | 0.036 | 0.0286 | 0.8 |
| Q8N239 | KLHL34 | LIIQALNyHTTPSRQ | Y258 | 0.102 | 96.04 | 644 | 0.044746 | 0.102 | 0.0447 | 1 |
| Q8N239 | KLHL34 | VGDRGVVyIsGGKAG | Y477 | 0.196 | 98.49 | 644 | 0.085982 | 0.196 | 0.086 | 1 |
| Q8N239 | KLHL34 | DRGVVyIsGGKAGRG | S479 | 0.006 | 97.83 | 644 | 0.004769 | 0.006 | 0.0048 | 1 |
| Q8N239 | KLHL34 | PFSEIERyDPGADQW | Y544 | 0.046 | 98.48 | 644 | 0.02018 | 0.046 | 0.0202 | 1 |
| P09874 | PARP1 | LFLAHILsPWGAEVk | S479 | 0.076 | 85.26 | 1014 | 0.060414 | 0.076 | 0.0604 | 0.8 |
| P04114 | APOB | RFKHLRKyTYNYEAE | Y52 | 0.023 | 95.49 | 4563 | 0.01009 | 0.023 | 0.0101 | 1 |
| P04114 | APOB | KLKFIIPsPKRPVKL | S928 | 0.168 | 82.72 | 4563 | 0.133546 | 0.186 | 0.1479 | 1 |
| P55199 | ELL | NMSAKDGtCtLQDCM | T258 | 0.191 | 90.4 | 621 | 0.130456 | 0.191 | 0.1305 | 1 |
| Q9ULQ0 | STRIP2 | GFtLGQDtDTLVGLP | T449 | 0.177 | 85.91 | 834 | 0.120893 | 0.177 | 0.1209 | 1 |
| Q07157 | TJP1 | VKEIsQDsLAARDGN | S215 | 0.051 | 87.98 | 1748 | 0.040541 | 0.051 | 0.0405 | 1 |
| Q9Y691 | KCNMB2 | SGEKLLLyHTEETIK | Y130 | 0.125 | 92.63 | 235 | 0.054836 | 0.125 | 0.0548 | 1 |
| Q8TCY9 | URGCP | MkESTTKyYFILSPy | Y383 | 0.026 | 94.67 | 931 | 0.011406 | 0.026 | 0.0114 | 1 |
| Q8TCY9 | URGCP | RAPCRRVsVEDMAHA | S447 | 0.067 | 96.87 | 931 | 0.053259 | 0.067 | 0.0533 | 1 |
| Q9NSA2 | KCND1 | RFLRSVMsLIDVVAI | S265 | 0.005 | 92.17 | 647 | 0.003975 | 0.005 | 0.004 | 1 |

| | | | | | | | | | | |
|---|---|---|---|---|---|---|---|---|---|---|
| Q9UPY5 | SLC7A11 | VTLLRGVsIIGTII | S51 | 0.053 | 95 | 501 | 0.04213 | 0.053 | 0.0421 | 1 |
| Q9H8T0 | AKTIP | IFIRHGLyQDGVFKF | Y117 | 0.009 | 97.74 | 292 | 0.003948 | 0.009 | 0.0039 | 1 |
| Q96J94 | PIWIL1 | DMTAGRRsIAGFVAs | S643 | 0.011 | 87.17 | 861 | 0.008744 | 0.011 | 0.0087 | 1 |
| Q96J94 | PIWIL1 | sIAGFVAsINEGMTR | S650 | 0 | 98.27 | 861 | 0 | 0 | 0 | 1 |
| Q96J94 | PIWIL1 | RGYNPRLtVIVVKKR | T736 | 0.015 | 98.18 | 861 | 0.010245 | 0.015 | 0.0102 | 1 |
| P13804 | ETFA | NDsLAPItLNTItAA | T37 | 0.026 | 98.62 | 333 | 0.017758 | 0.026 | 0.0178 | 1 |
| P13804 | ETFA | tRLGGEVsCLVAGTK | S52 | 0.015 | 98.43 | 333 | 0.011924 | 0.015 | 0.0119 | 1 |
| P13804 | ETFA | LVAQHDVykGLLPEE | Y84 | 0.131 | 96.84 | 333 | 0.057468 | 0.247 | 0.103 | 1 |
| P13804 | ETFA | GLLPEELtPLILATQ | T93 | 0.036 | 98.63 | 333 | 0.024588 | 0.036 | 0.0246 | 1 |
| P17948 | FLT1 | ITKEHSItLNLTIMN | T618 | 0.115 | 91.52 | 1338 | 0.078547 | 0.115 | 0.0785 | 1 |
| P17948 | FLT1 | NQKGsVEssAyLtVQ | S742 | 0.12 | 93.4 | 1338 | 0.09539 | 0.12 | 0.0954 | 1 |
| P16118 | PFKFB1 | FLKRIECyEVNyQPL | Y200 | 0.224 | 93.62 | 471 | 0.098266 | 0.224 | 0.0983 | 1 |
| P16118 | PFKFB1 | YLCRHGEsELNIRGR | S262 | 0.025 | 98.64 | 471 | 0.019873 | 0.025 | 0.0199 | 1 |
| P00747 | PLG | YVNTQGAsLFsVTKK | S32 | 0.03 | 84.61 | 810 | 0.023847 | 0.03 | 0.0238 | 1 |
| P00747 | PLG | KTGNGKNyRGTMSKT | Y111 | 0.069 | 91.63 | 810 | 0.030269 | 0.069 | 0.0303 | 1 |
| Q9HC56 | PCDH9 | KTGVLtAsRVFDREE | S527 | 0.168 | 92.18 | 1237 | 0.133546 | 0.168 | 0.1335 | 1 |
| Q7Z6L1 | TECPR1 | RLSVWAVsLQGKVWY | S215 | 0.033 | 87.98 | 1165 | 0.026232 | 0.033 | 0.0262 | 1 |
| P62955 | CACNG7 | AtPFPMVsLFLVFTA | S107 | 0 | 95.56 | 275 | 0 | 0 | 0 | 1 |
| P49792 | RANBP2 | MMACDRLsQsGHMLL | S352 | 0.002 | 95.17 | 3224 | 0.00159 | 0.002 | 0.0016 | 1 |
| P49792 | RANBP2 | sQsPKDtsFLGsDDI | S399 | 0.05 | 89.43 | 3224 | 0.039746 | 0.0715 | 0.0568 | 1 |
| Q9H3U1 | UNC45A | ItSNPEMtFPGERIy | T724 | 0.123 | 90.09 | 944 | 0.084011 | 0.123 | 0.084 | 1 |
| Q9H3U1 | UNC45A | AVPMIEGyMFEEHEM | Y783 | 0.167 | 95.61 | 944 | 0.07326 | 0.167 | 0.0733 | 1 |
| Q9H3U1 | UNC45A | AGGLAMLtsMRPTLC | T846 | 0.026 | 95.02 | 944 | 0.017758 | 0.026 | 0.0178 | 1 |
| Q9Y2U9 | KLHDC2 | VPPSMSGsCAVCVDR | S94 | 0.155 | 98.85 | 406 | 0.123212 | 0.155 | 0.1232 | 1 |
| P17948 | FLT1 | VNGHLYKtNYLtHRQ | T218 | 0 | 95.78 | 1338 | 0 | 0 | 0 | 1 |
| P17948 | FLT1 | LYKtNYLtHRQTNTI | T222 | 0.045 | 92.62 | 1338 | 0.030736 | 0.045 | 0.0307 | 1 |
| P17948 | FLT1 | ATEKSARyLTRGySL | Y383 | 0.189 | 88.52 | 1338 | 0.082912 | 0.189 | 0.0829 | 1 |
| Q9P212 | PLCE1 | LQLPLSyyyIESSHN | Y1401 | 0.014 | 97.62 | 2302 | 0.006142 | 0.014 | 0.0061 | 1 |
| Q9P212 | PLCE1 | QLPLSyyyIESSHNT | Y1402 | 0.005 | 98.01 | 2302 | 0.002193 | 0.005 | 0.0022 | 1 |
| O94979 | SEC31A | ENGNIILyDPSKIIA | Y101 | 0.043 | 93.82 | 1220 | 0.018863 | 0.043 | 0.0189 | 1 |
| O94979 | SEC31A | QHILASAsPSGRATV | S188 | 0.075 | 93.59 | 1220 | 0.059619 | 0.075 | 0.0596 | 1 |

| | | | | | | | | | | |
|---|---|---|---|---|---|---|---|---|---|---|
| Q9H0C8 | ILKAP | VLGVLEVsRSIGDGQ | S290 | 0 | 98.26 | 392 | 0 | 0 | 0 | 1 |
| Q9H0C8 | ILKAP | ykRCGVtsVPDIRRC | S305 | 0.166 | 98.48 | 392 | 0.131956 | 0.166 | 0.132 | 0.8889 |
| O75310 | UGT2B11 | GYTIERHsGGLIFPP | S184 | 0.017 | 94.16 | 529 | 0.013514 | 0.017 | 0.0135 | 1 |
| O75310 | UGT2B11 | KENIMKLsRIQHDQP | S446 | 0.084 | 98.7 | 529 | 0.066773 | 0.084 | 0.0668 | 1 |
| Q8N3C0 | ASCC3 | RKRWPtMtyRLLNLS | T1106 | 0 | 93.71 | 2202 | 0 | 0 | 0 | 1 |
| P12956 | XRCC6 | VAALCRYtPRRNIPP | T401 | 0.031 | 95.22 | 609 | 0.021173 | 0.062 | 0.0423 | 1 |
| E2RYF6 | MUC22 | VSTAGSEttVstAGS | T496 | 0.02 | 81.21 | 1773 | 0.01366 | 0.02 | 0.0137 | 1 |
| E2RYF6 | MUC22 | GSEttVstAGSETtA | T500 | 0.03 | 82.87 | 1773 | 0.02049 | 0.03 | 0.0205 | 1 |
| E2RYF6 | MUC22 | stAGSETtAAstEDS | T506 | 0.021 | 82.54 | 1773 | 0.014343 | 0.021 | 0.0143 | 1 |
| E2RYF6 | MUC22 | SETtAAstEDSETNT | T510 | 0.018 | 84.8 | 1773 | 0.012294 | 0.018 | 0.0123 | 1 |
| E2RYF6 | MUC22 | TAVSATGsEMttVST | S773 | 0.015 | 79.68 | 1773 | 0.011924 | 0.015 | 0.0119 | 1 |
| E2RYF6 | MUC22 | SATGsEMttVSTEGS | T776 | 0.008 | 91.16 | 1773 | 0.005464 | 0.008 | 0.0055 | 1 |
| E2RYF6 | MUC22 | SETTSASttGSETTT | T870 | 0.025 | 87.7 | 1773 | 0.017075 | 0.025 | 0.0171 | 1 |
| P42679 | MATK | ACENKsWyRVKHHTS | Y87 | 0.078 | 94.19 | 507 | 0.034217 | 0.119 | 0.0522 | 1 |
| Q96C36 | PYCR2 | sAHKIIAssPEMNLP | S33 | 0.012 | 98.26 | 320 | 0.009539 | 0.012 | 0.0095 | 1 |
| Q96C36 | PYCR2 | AHKIIAssPEMNLPt | S34 | 0.089 | 96.55 | 320 | 0.070747 | 0.166 | 0.132 | 1 |
| Q96C36 | PYCR2 | sPEMNLPtVSALRkM | T41 | 0.15 | 96.53 | 320 | 0.102452 | 0.211 | 0.1342 | 1 |
| Q96C36 | PYCR2 | VEAsCIRtRELQSMA | T265 | 0.065 | 95.3 | 320 | 0.044396 | 0.065 | 0.0444 | 1 |
| Q96KR4 | LMLN | tGECAAHtKCGPVIV | T162 | 0.022 | 95.68 | 655 | 0.015026 | 0.022 | 0.015 | 0.9231 |
| P23470 | PTPRG | VCRNDMRsDFsQTML | S433 | 0.035 | 87.38 | 1445 | 0.027822 | 0.035 | 0.0278 | 0.8333 |
| Q9P121 | NTM | TREQSGDyECSASND | Y199 | 0.037 | 96.98 | 344 | 0.016231 | 0.037 | 0.0162 | 1 |
| O95983 | MBD3 | GHRDVFyysPsGKKF | Y36 | 0.152 | 96.92 | 291 | 0.06668 | 0.152 | 0.0667 | 0.9 |
| Q8IY95 | TMEM192 | NRGYNLIyRSTRHLK | Y130 | 0.287 | 87.21 | 271 | 0.125903 | 0.287 | 0.1259 | 0.8889 |
| P05129 | PRKCG | tFCGtPDyIAPEIIA | Y521 | 0.065 | 96.2 | 697 | 0.028515 | 0.14 | 0.0614 | 1 |
| Q969P5 | FBXO32 | APDLHVLsEDRLLWK | S260 | 0.022 | 93.93 | 355 | 0.017488 | 0.022 | 0.0175 | 1 |
| P10644 | PRKAR1A | tLRKRkMyEEFLskV | Y246 | 0.09 | 86.61 | 381 | 0.039482 | 0.09 | 0.0395 | 1 |
| P10644 | PRKAR1A | MNRPRAAtVVArGPL | T338 | 0.013 | 95.82 | 381 | 0.008879 | 0.013 | 0.0089 | 1 |
| Q9Y613 | FHOD1 | EQLVQNAtFRCILAt | T816 | 0.059 | 94.7 | 1164 | 0.040298 | 0.059 | 0.0403 | 0.9091 |
| Q9Y613 | FHOD1 | tFRCILAtLLAVGNF | T823 | 0.001 | 94.45 | 1164 | 0.000683 | 0.149 | 0.1018 | 1 |
| Q9Y613 | FHOD1 | VLQTRPEsSDLySEI | S874 | 0.046 | 90.92 | 1164 | 0.036566 | 0.091 | 0.0724 | 1 |
| Q9Y613 | FHOD1 | AFLLyLGyTPQAARE | Y963 | 0.139 | 88.19 | 1164 | 0.060977 | 0.3 | 0.1316 | 0.9091 |

| | | | | | | | | | | |
|---|---|---|---|---|---|---|---|---|---|---|
| Q6ZU64 | CFAP65 | SQIPPVLtPLRLDFN | T1167 | 0.104 | 78.86 | 1925 | 0.071033 | 0.104 | 0.071 | 1 |
| Q99640 | PKMYT1 | PESFFQQsFQRLSRL | S109 | 0.007 | 97.81 | 499 | 0.005564 | 0.007 | 0.0056 | 1 |
| Q99640 | PKMYT1 | SKEDGRLyAVkRsMs | Y136 | 0.107 | 98.63 | 499 | 0.046939 | 0.107 | 0.0469 | 0.9167 |
| Q99640 | PKMYT1 | RLyAVkRsMsPFRGP | S141 | 0.114 | 94.96 | 499 | 0.09062 | 0.114 | 0.0906 | 1 |
| Q99640 | PKMYT1 | RkLAEVGsHEkVGQH | S160 | 0.113 | 94.91 | 499 | 0.089825 | 0.113 | 0.0898 | 0.8889 |
| Q8TD57 | DNAH3 | LLDFLKKsNDQIAQI | S1314 | 0.115 | 93.1 | 4116 | 0.091415 | 0.095 | 0.1236 | 1 |
| P52630 | STAT2 | LKELkGLsCLVsYQD | S283 | 0.148 | 85.53 | 851 | 0.117647 | 0.148 | 0.1176 | 1 |
| P09874 | PARP1 | GAAVDPDsGLEHsAH | S537 | 0.139 | 89.87 | 1014 | 0.110493 | 0.139 | 0.1105 | 1 |
| O00330 | PDHX | DALCEIEtDKAVVtL | T95 | 0.136 | 85.85 | 501 | 0.09289 | 0.217 | 0.1334 | 1 |
| Q13115 | DUSP4 | ARLRSGLySAVIVyD | Y102 | 0.149 | 87.61 | 394 | 0.065364 | 0.298 | 0.1307 | 1 |
| Q13115 | DUSP4 | LySAVIVyDERsPRA | Y108 | 0 | 96.33 | 394 | 0 | 0 | 0 | 1 |
| Q99614 | TTC1 | sCFQkERsILFSNRA | S155 | 0.004 | 97.98 | 292 | 0.00318 | 0.004 | 0.0032 | 1 |
| Q99614 | TTC1 | LDEALEDykSILEkD | Y212 | 0.027 | 98.58 | 292 | 0.011845 | 0.027 | 0.0118 | 1 |
| O43242 | PSMD3 | NEWARYLyYTGRIkA | Y290 | 0.193 | 90.08 | 534 | 0.084666 | 0.193 | 0.0847 | 1 |
| Q8NBS3 | SLC4A11 | TAMEVARtFATMFsD | T255 | 0 | 90.32 | 891 | 0 | 0 | 0 | 1 |
| Q9Y5F3 | PCDHB1 | GGGLsAHsKVLVEVV | S331 | 0.062 | 95.34 | 818 | 0.049285 | 0.062 | 0.0493 | 1 |
| Q6UXM1 | LRIG3 | KNNEIsWtIEDMNGA | T374 | 0.009 | 85.05 | 1119 | 0.006147 | 0.009 | 0.0061 | 1 |
| Q9P260 | RELCH | AARMFELtLRGMSEA | T991 | 0.021 | 88.33 | 1216 | 0.014343 | 0.021 | 0.0143 | 1 |
| Q7Z6M1 | RABEPK | GDkPRkAtWyTLTVP | T17 | 0.032 | 97.16 | 372 | 0.021856 | 0.032 | 0.0219 | 1 |
| Q7Z6M1 | RABEPK | kPRkAtWyTLTVPGD | Y19 | 0.153 | 97.56 | 372 | 0.067119 | 0.274 | 0.1124 | 1 |
| Q7Z6M1 | RABEPK | GGANPNRsFsDVHTM | S61 | 0.089 | 97.51 | 372 | 0.070747 | 0.089 | 0.0707 | 1 |
| Q13309 | SKP2 | CSKLQNLsLEGLRLS | S212 | 0 | 98.88 | 424 | 0 | 0 | 0 | 1 |
| Q86UW8 | HAPLN4 | QHRAFGsyRGRAELQ | Y112 | 0.137 | 97.43 | 402 | 0.0601 | 0.137 | 0.0601 | 1 |
| P16144 | ITGB4 | TAQGNRDyIPVEGEL | Y1039 | 0.02 | 91.73 | 1822 | 0.008774 | 0.02 | 0.0088 | 1 |
| Q2NL82 | TSR1 | GTLKISGyVrGQTLN | Y277 | 0.021 | 94.14 | 804 | 0.009212 | 0.021 | 0.0092 | 1 |
| Q2NL82 | TSR1 | FRAsPLFsQHtAADK | S630 | 0 | 95.56 | 804 | 0 | 0 | 0 | 1 |
| Q9NVX2 | NLE1 | AFSPTGKyLASGSGD | Y128 | 0.295 | 95.63 | 485 | 0.129412 | 0.295 | 0.1294 | 1 |
| Q7Z7M8 | B3GNT8 | PASARsLyLGEVFTQ | Y272 | 0.006 | 98.17 | 397 | 0.002632 | 0.006 | 0.0026 | 0.8 |
| Q7Z7M8 | B3GNT8 | RPLGPQAsIRLWKQL | S383 | 0.051 | 92.39 | 397 | 0.040541 | 0.051 | 0.0405 | 1 |
| Q13939 | CCIN | TSAGRyIyIsGGTTE | Y330 | 0.095 | 98.47 | 588 | 0.041675 | 0.095 | 0.0417 | 1 |
| Q13939 | CCIN | AGRyIyIsGGTTEQI | S332 | 0 | 97.47 | 588 | 0 | 0 | 0 | 1 |

| | | | | | | | | | | |
|---|---|---|---|---|---|---|---|---|---|---|
| Q13939 | CCIN | GLKTAWRyDMDDNSW | Y348 | 0.024 | 98.12 | 588 | 0.010528 | 0.024 | 0.0105 | 0.8571 |
| Q86UK7 | ZNF598 | DGKVyALyRQLLQHE | Y111 | 0.056 | 94.33 | 904 | 0.024566 | 0.056 | 0.0246 | 0.8333 |
| Q5VZP5 | STYXL2 | SQHFRKAsEFLDEAL | S207 | 0.001 | 98.11 | 1158 | 0.000795 | 0.001 | 0.0008 | 0.8 |
| Q8WVM0 | TFB1M | CRDGPFVyGRTQMtL | Y167 | 0.03 | 98.03 | 346 | 0.013161 | 0.03 | 0.0132 | 1 |
| Q8WVM0 | TFB1M | VyGRTQMtLtFQKEV | T173 | 0.016 | 98.62 | 346 | 0.010928 | 0.016 | 0.0109 | 1 |
| Q8WVM0 | TFB1M | GRTQMtLtFQKEVAE | T175 | 0.026 | 97.17 | 346 | 0.017758 | 0.026 | 0.0178 | 1 |
| Q92673 | SORL1 | VHWAGEKsNVIVALA | S114 | 0.059 | 79.43 | 2214 | 0.0469 | 0.059 | 0.0469 | 0.8333 |
| Q92673 | SORL1 | RPKSSDVyVSyDYGK | Y136 | 0.08 | 94.55 | 2214 | 0.035095 | 0.08 | 0.0351 | 1 |
| Q92673 | SORL1 | NRTNLYIsEAEGLKF | S375 | 0 | 88.75 | 2214 | 0 | 0 | 0 | 0.9167 |
| Q92673 | SORL1 | MPILSKEsAPGLIIA | S499 | 0.15 | 87.81 | 2214 | 0.119237 | 0.142 | 0.1306 | 1 |
| Q7KZF4 | SND1 | HPPVEGsyAPRRGEF | Y728 | 0.082 | 84.24 | 910 | 0.035972 | 0.295 | 0.1294 | 0.8333 |
| P00441 | SOD1 | sIEDsVIsLsGDHCI | S106 | 0.112 | 98.44 | 154 | 0.08903 | 0.112 | 0.089 | 0.8889 |
| P02751 | FN1 | VSGLMVAtKYEVSVY | T1877 | 0.046 | 81.62 | 2477 | 0.031419 | 0.103 | 0.1351 | 1 |
| P04843 | RPN1 | SVIVETVytHVLHPY | Y135 | 0.096 | 98.3 | 607 | 0.042114 | 0.096 | 0.0421 | 0.9231 |
| P04843 | RPN1 | RsEDLLDyGPFRDVP | Y200 | 0.034 | 97.69 | 607 | 0.014915 | 0.034 | 0.0149 | 1 |
| P04843 | RPN1 | GPFSRyDyQRQPDSG | Y266 | 0.333 | 90.42 | 607 | 0.146082 | 0.333 | 0.1461 | 1 |
| P04843 | RPN1 | AAAQDVyyRDEIGNV | Y293 | 0.189 | 94.78 | 607 | 0.082912 | 0.189 | 0.0829 | 1 |
| Q9Y2J8 | PADI2 | FLtAIEIsLDVDADR | S119 | 0.05 | 98.42 | 665 | 0.039746 | 0.05 | 0.0397 | 1 |
| Q9Y2J8 | PADI2 | KEDLKDMsQMILRTK | S182 | 0.056 | 98.41 | 665 | 0.044515 | 0.094 | 0.0747 | 1 |
| Q9Y2J8 | PADI2 | GFPVVLDsPRDGNLK | S371 | 0.018 | 93.48 | 665 | 0.014308 | 0.033 | 0.0266 | 1 |
| Q9Y2J8 | PADI2 | LsGGRRMtKVVRDFL | T444 | 0.003 | 96.99 | 665 | 0.002049 | 0.003 | 0.002 | 0.8 |
| P30304 | CDC25A | GkHQDLkyIsPEIMA | Y359 | 0.054 | 98.25 | 524 | 0.023689 | 0.054 | 0.0237 | 1 |
| P17980 | PSMC3 | HSRkMNVsPDVNyEE | S376 | 0.173 | 89.11 | 439 | 0.13752 | 0.173 | 0.1375 | 1 |
| P17980 | PSMC3 | TELTHEDyMEGILEV | Y420 | 0.004 | 91.92 | 439 | 0.001755 | 0.004 | 0.0018 | 1 |
| P48444 | ARCN1 | kkGVQLQtHPNVDkk | T329 | 0.149 | 97.69 | 511 | 0.101769 | 0.175 | 0.1195 | 1 |
| P48444 | ARCN1 | RHDSRRNtLEWCLPV | T437 | 0.123 | 97.71 | 511 | 0.084011 | 0.123 | 0.084 | 1 |
| Q96DX5 | ASB9 | NQGKGQDsPLHAVAR | S201 | 0.069 | 98.32 | 294 | 0.054849 | 0.069 | 0.0548 | 1 |
| Q96DX5 | ASB9 | LMDFGADtQAKNAEG | T225 | 0.208 | 95.67 | 294 | 0.142067 | 0.208 | 0.1421 | 1 |
| Q96DX5 | ASB9 | VELVPPEsPLAQLFL | S243 | 0.164 | 92.01 | 294 | 0.130366 | 0.164 | 0.1304 | 1 |
| O75077 | ADAM23 | DDDTDVGyVEDGTPC | Y707 | 0.197 | 97.68 | 832 | 0.086421 | 0.197 | 0.0864 | 1 |
| Q5JSL3 | DOCK11 | IYKEPKLtGLSEISL | T1790 | 0.11 | 88.37 | 2073 | 0.075131 | 0.11 | 0.0751 | 1 |

| | | | | | | | | | | |
|---|---|---|---|---|---|---|---|---|---|---|
| Q68DK2 | ZFYVE26 | AERRLNssLERRGRR | S1002 | 0.024 | 80.56 | 2539 | 0.019078 | 0.03 | 0.0442 | 0.875 |
| Q96HS1 | PGAM5 | GREQAELtGLRLASL | T131 | 0.003 | 98.93 | 289 | 0.002049 | 0.003 | 0.002 | 1 |
| Q8NDZ6 | TMEM161B | MTEAtFDtLRLWLII | T313 | 0.14 | 89.13 | 487 | 0.095622 | 0.14 | 0.0956 | 1 |
| Q8IVH8 | MAP4K3 | NPDTRDQyLIFGAEE | Y574 | 0.257 | 93.35 | 894 | 0.112742 | 0.257 | 0.1127 | 1 |
| Q8IVH8 | MAP4K3 | VMNNCLLsIsGKASQ | S616 | 0 | 97.35 | 894 | 0 | 0 | 0 | 1 |
| Q8IVH8 | MAP4K3 | NNCLLsIsGKASQLY | S618 | 0.026 | 95.85 | 894 | 0.020668 | 0.026 | 0.0207 | 1 |
| Q8IZU2 | WDR17 | ATLAIYIyQLDHRYN | Y64 | 0.165 | 92.9 | 1322 | 0.072383 | 0.328 | 0.1439 | 1 |
| Q8IZU2 | WDR17 | DGKVLHKyKHPAAVF | Y518 | 0.002 | 96.95 | 1322 | 0.000877 | 0.002 | 0.0009 | 1 |
| Q8IZU2 | WDR17 | EDTNVRVyYVAtSSD | Y549 | 0.027 | 95.82 | 1322 | 0.011845 | 0.027 | 0.0118 | 1 |
| Q8IZU2 | WDR17 | MWNTEIPyLLISGSW | Y623 | 0.174 | 94.59 | 1322 | 0.076331 | 0.246 | 0.1079 | 1 |
| Q8TC92 | ENOX1 | RGEVNRRsANQFYSM | S299 | 0.074 | 91.42 | 643 | 0.058824 | 0.074 | 0.0588 | 1 |
| O95965 | ITGBL1 | RysDDFCsGHGQCNC | S289 | 0.061 | 89.95 | 494 | 0.04849 | 0.061 | 0.0485 | 0.875 |
| Q5VU57 | AGBL4 | RDAVKAGtGRRFLGG | T382 | 0.08 | 96.95 | 503 | 0.054641 | 0.08 | 0.0546 | 1 |
| Q8TEY7 | USP33 | ELKGDNMySCEKCKK | Y588 | 0.341 | 91.01 | 942 | 0.149592 | 0.341 | 0.1496 | 1 |
| Q8N0X4 | CLYBL | QSRVLPssLMLPKVE | S132 | 0 | 98.6 | 340 | 0 | 0 | 0 | 1 |
| Q8N0X4 | CLYBL | GAGLLRQsREGAAMG | S258 | 0 | 97.88 | 340 | 0 | 0 | 0 | 1 |
| Q6SA08 | TSSK4 | KWENVKIsDFGFAKM | S165 | 0.15 | 96.36 | 328 | 0.119237 | 0.15 | 0.1192 | 1 |
| O75592 | MYCBP2 | LFGGRGEyTAKIKLF | Y1276 | 0.038 | 93.37 | 4678 | 0.01667 | 0.038 | 0.0167 | 0.8 |
| O43314 | PPIP5K2 | FHVELHFsPGAKGCE | S894 | 0 | 91.5 | 1243 | 0 | 0 | 0 | 1 |
| Q6GYQ0 | RALGAPA1 | PHGDVKKstQKVLDt | S13 | 0.063 | 88.17 | 2036 | 0.05008 | 0.127 | 0.101 | 1 |
| P17342 | NPR3 | HWDLPMLsAGALAAG | S156 | 0 | 98.81 | 541 | 0 | 0 | 0 | 0.9091 |
| P17342 | NPR3 | DFEAKQAySSLQTVT | Y316 | 0.055 | 95.99 | 541 | 0.024128 | 0.055 | 0.0241 | 1 |
| Q9NP85 | NPHS2 | EIDAICyyRMENASL | Y195 | 0.061 | 96.58 | 383 | 0.02676 | 0.061 | 0.0268 | 1 |
| Q96PM5 | RCHY1 | HLLHRtCyEEMLkEG | Y173 | 0.181 | 97.35 | 261 | 0.079402 | 0.181 | 0.0794 | 1 |
| P23381 | WARS1 | LRRGIFFsHRDMNQV | S139 | 0 | 98.55 | 471 | 0 | 0 | 0 | 1 |
| Q9H596 | DUSP21 | CMAGVSRsAsLCLAy | S113 | 0.001 | 98.73 | 190 | 0.000795 | 0.001 | 0.0008 | 0.9091 |
| Q9H596 | DUSP21 | AGVSRsAsLCLAyLM | S115 | 0 | 98.9 | 190 | 0 | 0 | 0 | 0.9 |
| Q8WX94 | NLRP7 | PVPVLLGsLLkRKML | S285 | 0.001 | 93.16 | 980 | 0.000795 | 0.001 | 0.0008 | 1 |
| Q9H2K2 | TNKS2 | ALHCAAAsPYPkRkQ | S374 | 0.011 | 97.82 | 1166 | 0.008744 | 0.011 | 0.0087 | 1 |
| Q02388 | COL7A1 | SVPGAtGyRVTWRVL | Y356 | 0.01 | 90.815 | 2944 | 0.004387 | 0.01 | 0.0044 | 1 |
| Q8NER1 | TRPV1 | LTNKKGMtPLALAAG | T336 | 0.126 | 93.78 | 839 | 0.08606 | 0.126 | 0.0861 | 1 |

| | | | | | | | | | | |
|---|---|---|---|---|---|---|---|---|---|---|
| Q12906 | ILF3 | AEQGPILtkHGKNPV | T522 | 0.185 | 88.22 | 894 | 0.126358 | 0.185 | 0.1264 | 1 |
| Q8NE86 | MCU | SRKAEKRttLVLWGG | T232 | 0.124 | 95.69 | 351 | 0.084694 | 0.124 | 0.0847 | 1 |
| Q8NBL1 | POGLUT1 | RRkLGTHyQItkNRL | Y89 | 0 | 98.9 | 392 | 0 | 0 | 0 | 1 |
| Q8NBL1 | POGLUT1 | LGTHyQItkNRLYRE | T92 | 0.198 | 98.56 | 392 | 0.135237 | 0.198 | 0.1352 | 1 |
| Q8NBL1 | POGLUT1 | EYSKFLSyNVTRRKG | Y372 | 0.301 | 98.73 | 392 | 0.132044 | 0.301 | 0.132 | 1 |
| P0C7Q6 | SLC35G6 | FVGGLALsLGLLVyR | S204 | 0.036 | 92.28 | 338 | 0.028617 | 0.036 | 0.0286 | 0.8 |
| Q99504 | EYA3 | YRKVREIyDKHkSNV | Y426 | 0.011 | 98.66 | 573 | 0.004826 | 0.011 | 0.0048 | 0.8571 |
| Q99504 | EYA3 | RFGKkVTyVVIGDGR | Y532 | 0.166 | 98.43 | 573 | 0.072822 | 0.166 | 0.0728 | 1 |
| Q9NRX4 | PHPT1 | DsDGVFKyVLIRVHs | Y22 | 0 | 97.93 | 125 | 0 | 0 | 0 | 0.8571 |
| Q9NRX4 | PHPT1 | KIHVyGysMAyGPAQ | S94 | 0.031 | 92.62 | 125 | 0.024642 | 0.031 | 0.0246 | 1 |
| P61158 | ACTR3 | MEQVIFkyLRAEPED | Y100 | 0.307 | 97.37 | 418 | 0.134676 | 0.307 | 0.1347 | 1 |
| P61158 | ACTR3 | RAEPEDHyFLLtEPP | Y109 | 0.276 | 97.73 | 418 | 0.121077 | 0.276 | 0.1211 | 1 |
| P61158 | ACTR3 | PIAGRDItyFIQQLL | T201 | 0.089 | 97.26 | 418 | 0.060788 | 0.089 | 0.0608 | 0.9 |
| P61158 | ACTR3 | kAVkERysyVCPDLV | S232 | 0.162 | 97.91 | 418 | 0.128776 | 0.162 | 0.1288 | 1 |
| P61158 | ACTR3 | AVkERysyVCPDLVk | Y233 | 0.141 | 97.82 | 418 | 0.061855 | 0.141 | 0.0619 | 1 |
| P61158 | ACTR3 | LYkNIVLsGGstMFR | S322 | 0.016 | 96.01 | 418 | 0.012719 | 0.016 | 0.0127 | 1 |
| P61158 | ACTR3 | IVLsGGstMFRDFGR | T326 | 0.048 | 96.34 | 418 | 0.032785 | 0.048 | 0.0328 | 1 |
| P61158 | ACTR3 | VCHTKkDyEEIGPSI | Y400 | 0.161 | 97.27 | 418 | 0.070628 | 0.161 | 0.0706 | 1 |
| Q8NH41 | OR4K15 | SLLIVADsGFLSLSS | S203 | 0.056 | 85.59 | 348 | 0.044515 | 0.056 | 0.0445 | 0.9091 |
| P31749 | AKT1 | EKATGRYyAMKILKK | Y176 | 0.215 | 95.62 | 480 | 0.094317 | 0.215 | 0.0943 | 1 |
| O75995 | SASH3 | PSPyDHDsLKLQKGD | S193 | 0.133 | 95.34 | 380 | 0.105724 | 0.133 | 0.1057 | 1 |
| Q96PJ5 | FCRL4 | WGEKLtLtPGNtLEV | T70 | 0.175 | 90.52 | 515 | 0.119527 | 0.175 | 0.1195 | 1 |
| Q9BSJ8 | ESYT1 | QVKLTLWyySEERkL | Y981 | 0.245 | 90.47 | 1104 | 0.107478 | 0.245 | 0.1075 | 1 |
| Q9BSJ8 | ESYT1 | GRDPPDPyVsLLLLP | Y1009 | 0.277 | 92.7 | 1104 | 0.121516 | 0.277 | 0.1215 | 0.8182 |
| Q15599 | NHERF2 | IRRVEPGsPAEAAAL | S43 | 0.113 | 88.16 | 337 | 0.089825 | 0.113 | 0.0898 | 1 |
| Q15599 | NHERF2 | LLVVDQEtDEELRRR | T92 | 0.006 | 75.29 | 337 | 0.004098 | 0.006 | 0.0041 | 1 |
| P47897 | QARS1 | kGHNTLPsPWRDRPM | S373 | 0.11 | 90.83 | 775 | 0.087441 | 0.11 | 0.0874 | 1 |
| Q8IY81 | FTSJ3 | LFRRVQAtkPQAsRH | T181 | 0.145 | 93.29 | 847 | 0.099037 | 0.145 | 0.099 | 1 |
| Q8IY81 | FTSJ3 | QAtkPQAsRHESAEI | S186 | 0.049 | 88.67 | 847 | 0.038951 | 0.049 | 0.039 | 1 |
| P21695 | GPD1 | DIGGKKLtEIINtQH | T51 | 0.132 | 98.63 | 349 | 0.090158 | 0.132 | 0.0902 | 1 |
| Q9Y2F9 | BTBD3 | SSCGsAEySAKIELK | Y422 | 0.019 | 94.17 | 522 | 0.008335 | 0.019 | 0.0083 | 0.8667 |

| | | | | | | | | | | |
|---|---|---|---|---|---|---|---|---|---|---|
| Q96J65 | ABCC12 | AMAEANVsLRRMKKI | S407 | 0.01 | 86.94 | 1359 | 0.007949 | 0.01 | 0.0079 | 1 |
| Q9NYU1 | UGGT2 | MLSVLRNtKTPVKFW | T1257 | 0.106 | 93.14 | 1516 | 0.072399 | 0.211 | 0.1444 | 0.9167 |
| Q9P2B2 | PTGFRN | FSKEHtDtFNFRIQR | T497 | 0.097 | 91.43 | 879 | 0.066252 | 0.097 | 0.0663 | 1 |
| P11021 | HSPA5 | kksQIFstAsDNQPt | T453 | 0.106 | 86.47 | 654 | 0.072399 | 0.123 | 0.1058 | 1 |
| Q14008 | CKAP5 | YEKMAKAtGKLKPTs | T1058 | 0.011 | 89.22 | 2032 | 0.007513 | 0.011 | 0.0075 | 1 |
| O15121 | DEGS1 | FVMDDTIsPYSRMKR | S307 | 0.003 | 95.96 | 323 | 0.002385 | 0.003 | 0.0024 | 1 |
| Q8TC07 | TBC1D15 | kYVVLCEsPQDkRTL | S186 | 0.115 | 80.9 | 691 | 0.091415 | 0.115 | 0.0914 | 1 |
| Q8NHM5 | KDM2B | MEGKDFNyEYVQREA | Y94 | 0.143 | 95.17 | 1336 | 0.062732 | 0.247 | 0.1084 | 1 |
| Q9NX02 | NLRP2 | PVPVLLGsLLNRVML | S320 | 0.001 | 90.64 | 1062 | 0.000795 | 0.001 | 0.0008 | 1 |
| Q8WX92 | NELFB | LEDLLEksFSLVKMP | S100 | 0.152 | 90.96 | 580 | 0.120827 | 0.152 | 0.1208 | 1 |
| Q8WX92 | NELFB | GEVVQRLtRMVGKNV | T205 | 0.053 | 98.25 | 580 | 0.0362 | 0.053 | 0.0362 | 1 |
| Q8WX92 | NELFB | QEtLPRDsPDLLLLL | S335 | 0.14 | 95.27 | 580 | 0.111288 | 0.14 | 0.1113 | 1 |
| P37173 | TGFBR2 | NVESFKQtDVYSMAL | T445 | 0.128 | 97.06 | 567 | 0.087426 | 0.206 | 0.1407 | 0.8889 |
| P00736 | C1R | KMLLTFHtDFSNEEN | T118 | 0.051 | 94.95 | 705 | 0.034834 | 0.051 | 0.0348 | 1 |
| Q8NFP9 | NBEA | NSEEQKItEMVyNIF | T913 | 0.079 | 95.65 | 2946 | 0.053958 | 0.079 | 0.054 | 1 |
| P42858 | HTT | VRHVAAAsLIRLVPk | S930 | 0.039 | 97.49 | 3142 | 0.031002 | 0.039 | 0.031 | 1 |
| Q15365 | PCBP1 | VkRIREEsGARINIs | S43 | 0.112 | 90.97 | 356 | 0.08903 | 0.112 | 0.089 | 0.9 |
| Q99704 | DOK1 | ATAFRLDtAQRSHLL | T93 | 0.149 | 93.66 | 481 | 0.101769 | 0.149 | 0.1018 | 1 |
| Q562R1 | ACTBL2 | GMGQkDCyVGDEAQS | Y54 | 0.027 | 95.73 | 376 | 0.011845 | 0.202 | 0.0533 | 1 |
| Q562R1 | ACTBL2 | MDSGDGVtHIVPIYE | T161 | 0.053 | 98.24 | 376 | 0.0362 | 0.092 | 0.0628 | 0.9091 |
| Q562R1 | ACTBL2 | DLAGRDLtDyLMkIL | T187 | 0.014 | 98.1 | 376 | 0.009562 | 0.014 | 0.0096 | 0.9 |
| Q562R1 | ACTBL2 | AGRDLtDyLMkILTE | Y189 | 0.182 | 97.92 | 376 | 0.079841 | 0.182 | 0.0798 | 1 |
| Q562R1 | ACTBL2 | VDIRkDLyANtVLSG | Y295 | 0.004 | 98.18 | 376 | 0.001755 | 0.004 | 0.0018 | 1 |
| Q562R1 | ACTBL2 | RkDLyANtVLSGGst | T298 | 0.006 | 98.42 | 376 | 0.004098 | 0.006 | 0.0041 | 1 |
| Q562R1 | ACTBL2 | tVLSGGstMyPGIAD | T305 | 0 | 98.34 | 376 | 0 | 0 | 0 | 1 |
| Q6ZWH5 | NEK10 | VDKLVNMtYIFQKLA | T189 | 0 | 93.03 | 1172 | 0 | 0.042 | 0.0287 | 0.8 |
| Q9UIV1 | CNOT7 | QHQAGSDsLLtGMAF | S231 | 0 | 98.53 | 285 | 0 | 0 | 0 | 1 |
| O94925 | GLS | IAVNDLGtEyVHRYV | T302 | 0.214 | 98.52 | 669 | 0.146165 | 0.214 | 0.1462 | 1 |
| O94925 | GLS | ITGERVLsPEAVRNT | S449 | 0.169 | 98.48 | 669 | 0.134341 | 0.169 | 0.1343 | 1 |
| P29459 | IL12A | ALCLssIyEDLKMYQ | Y128 | 0.312 | 93.8 | 219 | 0.13687 | 0.312 | 0.1369 | 1 |
| O15379 | HDAC3 | CRFHSEDyIDFLQRV | Y66 | 0 | 98.65 | 428 | 0 | 0 | 0 | 1 |

| | | | | | | | | | | |
|---|---|---|---|---|---|---|---|---|---|---|
| P23468 | PTPRD | QLEKWTEyRITVTAH | Y683 | 0.153 | 92.31 | 1912 | 0.067119 | 0.153 | 0.0671 | 1 |
| P23468 | PTPRD | TKGDGARskPKLVST | S812 | 0.102 | 89.23 | 1912 | 0.081081 | 0.102 | 0.0811 | 1 |
| Q5RL73 | RBM48 | TINLESQyLLIQGVP | Y47 | 0.126 | 96.74 | 367 | 0.055274 | 0.126 | 0.0553 | 0.8182 |
| Q8N1T3 | MYO1H | NNHFILIsGESGAGK | S104 | 0.006 | 89.7 | 1032 | 0.004769 | 0.006 | 0.0048 | 1 |
| Q8N1T3 | MYO1H | IVFVRKNyILNLRyH | Y786 | 0.143 | 83.65 | 1032 | 0.062732 | 0.143 | 0.0627 | 1 |
| Q8N1T3 | MYO1H | YSALKGVsTsNLsDG | S938 | 0.055 | 89.62 | 1032 | 0.04372 | 0.055 | 0.0437 | 1 |
| Q8N1T3 | MYO1H | ALKGVsTsNLsDGIL | S940 | 0 | 88.64 | 1032 | 0 | 0 | 0 | 1 |
| A6NES4 | MROH2A | LIQFLRNsLKKTRGS | S653 | 0.029 | 86.41 | 1674 | 0.023053 | 0.121 | 0.0962 | 1 |
| O00423 | EML1 | VFsAEEGyVKMFLRG | Y186 | 0.225 | 95.97 | 815 | 0.098704 | 0.225 | 0.0987 | 1 |
| O00423 | EML1 | LCMLRDGtLVsGGGK | T473 | 0.153 | 96.74 | 815 | 0.104501 | 0.153 | 0.1045 | 1 |
| O00423 | EML1 | LRDGtLVsGGGKDRK | S476 | 0 | 98.39 | 815 | 0 | 0 | 0 | 1 |
| O00423 | EML1 | GKDRKLIsWsGNyQK | S486 | 0.055 | 97.85 | 815 | 0.04372 | 0.055 | 0.0437 | 1 |
| O00423 | EML1 | LIsWsGNyQKLRKTE | Y491 | 0.288 | 95.89 | 815 | 0.126341 | 0.288 | 0.1263 | 1 |
| O00423 | EML1 | AHEKKLLsTGDDFGK | S755 | 0 | 98.8 | 815 | 0 | 0 | 0 | 1 |
| P51164 | ATP4B | DYQDQLRsPGVtLRP | S79 | 0.123 | 93.81 | 291 | 0.097775 | 0.123 | 0.0978 | 1 |
| P51164 | ATP4B | EKGLEIVyNVSDNRT | Y98 | 0.029 | 94.49 | 291 | 0.012722 | 0.029 | 0.0127 | 1 |
| P62847 | RPS24 | TTGFGMIyDsLDyAk | Y76 | 0.039 | 96.83 | 133 | 0.017109 | 0.039 | 0.0171 | 1 |
| P51530 | DNA2 | LIVGMPGtGKTTTIC | T652 | 0.01 | 93.48 | 1060 | 0.00683 | 0.01 | 0.0068 | 0.8 |
| P51530 | DNA2 | kYQGRDKsIVLVSFV | S975 | 0.183 | 93.93 | 1060 | 0.145469 | 0.183 | 0.1455 | 1 |
| P51530 | DNA2 | VLVSFVRsNKDGtVG | S984 | 0.15 | 90.69 | 1060 | 0.119237 | 0.15 | 0.1192 | 1 |
| Q6ZMR3 | LDHAL6A | SVADLtEsILKNLRR | S262 | 0 | 98.51 | 332 | 0 | 0 | 0 | 0.875 |
| P28039 | AOAH | SDPVPAMtTPEKLYS | T387 | 0.044 | 98.63 | 575 | 0.030053 | 0.044 | 0.0301 | 0.9091 |
| Q6W4X9 | MUC6 | LVAVVYLsRQDKIVI | S458 | 0.126 | 89.78 | 2439 | 0.100159 | 0.126 | 0.1002 | 1 |
| P01111 | NRAS | FAINNSksFADINLY | S89 | 0.019 | 97.96 | 189 | 0.015103 | 0.019 | 0.0151 | 0.9 |
| Q9ULG1 | INO80 | QGHRVLIysQMtRMI | Y1123 | 0 | 95.68 | 1556 | 0 | 0 | 0 | 1 |
| Q9ULG1 | INO80 | GHRVLIysQMtRMID | S1124 | 0.006 | 94.61 | 1556 | 0.004769 | 0.006 | 0.0048 | 1 |
| Q9ULG1 | INO80 | MIDLLEEyMVyRKHT | Y1136 | 0.02 | 94.62 | 1556 | 0.008774 | 0.02 | 0.0088 | 1 |
| P69905 | HBA1 | MFLsFPttktyFPHF | T40 | 0.022 | 98.7 | 142 | 0.015026 | 0.022 | 0.015 | 0.875 |
| P69905 | HBA1 | HFDLsHGsAQVkGHG | S53 | 0.075 | 98.56 | 142 | 0.059619 | 0.075 | 0.0596 | 1 |
| P69905 | HBA1 | LSHCLLVtLAAHLPA | T109 | 0.013 | 98.85 | 142 | 0.008879 | 0.013 | 0.0089 | 0.8182 |
| Q6ZSZ5 | ARHGEF18 | HNEAVSHykLLLQQN | Y564 | 0.162 | 95.81 | 1361 | 0.071067 | 0.162 | 0.0711 | 1 |

| | | | | | | | | | | |
|---|---|---|---|---|---|---|---|---|---|---|
| Q9UDR5 | AASS | VLGSGYIsEPVLEYL | S492 | 0.015 | 94.54 | 926 | 0.011924 | 0.015 | 0.0119 | 1 |
| Q9UDR5 | AASS | PGLNLEGyPNRDSTk | Y700 | 0.291 | 96.81 | 926 | 0.127658 | 0.291 | 0.1277 | 1 |
| Q9UDR5 | AASS | GTLRYKGyMKALNGF | Y730 | 0.026 | 97.65 | 926 | 0.011406 | 0.026 | 0.0114 | 1 |
| Q9UDR5 | AASS | AKTVGLPtAMAAKML | T878 | 0 | 98.51 | 926 | 0 | 0 | 0 | 1 |
| Q9NNW5 | WDR6 | VCGDRRGsVLLFPSR | S514 | 0.047 | 95.29 | 1121 | 0.037361 | 0.047 | 0.0374 | 0.8571 |
| O94832 | MYO1D | VQQPGERsFHsFyQL | S200 | 0 | 93.63 | 1006 | 0 | 0 | 0 | 0.875 |
| Q9BS26 | ERP44 | SKRNIIGyFEQKDSD | Y166 | 0.065 | 98.29 | 406 | 0.028515 | 0.065 | 0.0285 | 1 |
| Q9UBK8 | MTRR | LRALVDYtsDSAEkR | T392 | 0.063 | 98.78 | 698 | 0.04303 | 0.063 | 0.043 | 1 |
| O60336 | MAPKBP1 | GIRSVCVsPNGQHLA | S488 | 0.048 | 95 | 1514 | 0.038156 | 0.088 | 0.07 | 1 |
| O14926 | FSCN2 | FKDCDGHyLAPVGPA | Y228 | 0.047 | 96 | 492 | 0.020618 | 0.047 | 0.0206 | 1 |
| O60725 | ICMT | PGSEARLsLATFLLG | S17 | 0 | 97.86 | 284 | 0 | 0 | 0 | 1 |
| Q14DG7 | TMEM132B | ITTVQVLsPLSDSIL | S636 | 0.096 | 94.12 | 1078 | 0.076312 | 0.152 | 0.1208 | 1 |
| Q9Y2E5 | MAN2B2 | SPKVRDMyAtHLASG | Y435 | 0.017 | 98.69 | 1009 | 0.007458 | 0.288 | 0.1263 | 0.8182 |
| P32297 | CHRNA3 | yNCCEEIyPDITySL | Y228 | 0.263 | 96.75 | 505 | 0.115374 | 0.263 | 0.1154 | 1 |
| P32297 | CHRNA3 | EIyPDITySLYIRRL | Y233 | 0.032 | 97.15 | 505 | 0.014038 | 0.032 | 0.014 | 1 |
| O14745 | NHERF1 | IRsVDPDsPAEAsGL | S186 | 0.122 | 89.94 | 358 | 0.09698 | 0.15 | 0.1192 | 1 |
| Q53SF7 | COBLL1 | EPLDLTKsLNDLGLR | S204 | 0.061 | 93.37 | 1128 | 0.04849 | 0.061 | 0.0485 | 1 |
| P16233 | PNLIP | VNTRFLLytNENPNN | Y58 | 0.23 | 98.77 | 465 | 0.100898 | 0.23 | 0.1009 | 1 |
| P16233 | PNLIP | NTRFLLytNENPNNF | T59 | 0 | 98.77 | 465 | 0 | 0 | 0 | 1 |
| O75715 | GPX5 | KDEKGTIyDYEAIAL | Y41 | 0.238 | 98.08 | 221 | 0.104407 | 0.238 | 0.1044 | 1 |
| Q9BWM7 | SFXN3 | VVLIGRMsAQVPMNM | S93 | 0.004 | 78.82 | 321 | 0.00318 | 0.004 | 0.0032 | 1 |
| P12111 | COL6A3 | GNALEyVsRNIFKRP | S1320 | 0.018 | 88.8 | 3177 | 0.014308 | 0.018 | 0.0143 | 0.8571 |
| Q86UF1 | TSPAN33 | EASKVIYtNGCIDKL | T221 | 0.106 | 97.04 | 283 | 0.072399 | 0.106 | 0.0724 | 1 |
| O75569 | PRKRA | GPAHkREytTICRLE | Y159 | 0.302 | 92.43 | 313 | 0.132483 | 0.302 | 0.1325 | 1 |
| B1ANS9 | WDR64 | LQNQVLDsKNFKSVK | S265 | 0.147 | 86.05 | 1081 | 0.116852 | 0.147 | 0.1169 | 0.8333 |
| Q9UHC6 | CNTNAP2 | VEIDLTEsKVGVHIN | S429 | 0.153 | 95.52 | 1331 | 0.121622 | 0.153 | 0.1216 | 1 |
| Q9UHC6 | CNTNAP2 | MTEDKVWtIVsHDLQ | T638 | 0.158 | 96.89 | 1331 | 0.107916 | 0.158 | 0.1079 | 1 |
| Q9UHC6 | CNTNAP2 | LKsAtEVsFsFDVGN | S857 | 0.009 | 97.58 | 1331 | 0.007154 | 0.009 | 0.0072 | 1 |
| Q9UHC6 | CNTNAP2 | sAtEVsFsFDVGNGP | S859 | 0.163 | 95.89 | 1331 | 0.129571 | 0.163 | 0.1296 | 1 |
| Q8TEL6 | TRPC4AP | TLMTSKKtFLQTATL | T203 | 0.032 | 88.95 | 797 | 0.021856 | 0.032 | 0.0219 | 0.9167 |
| P14210 | HGF | EFKKSAKtTLIKIDP | T48 | 0.201 | 96.41 | 728 | 0.137286 | 0.201 | 0.1373 | 1 |

| | | | | | | | | | | |
|---|---|---|---|---|---|---|---|---|---|---|
| Q13200 | PSMD2 | QIDKyLyssEDyIks | S435 | 0.185 | 75.39 | 908 | 0.147059 | 0.185 | 0.1471 | 0.8 |
| Q96SB8 | SMC6 | SRSyKLksAtGsVVs | S158 | 0.079 | 92.39 | 1091 | 0.062798 | 0.079 | 0.0628 | 1 |
| Q96SB8 | SMC6 | sAtGsVVstRKEELI | S165 | 0.178 | 91.35 | 1091 | 0.141495 | 0.178 | 0.1415 | 1 |
| Q8NFI3 | ENGASE | HGVCVLGtFItEWNE | T190 | 0 | 98.79 | 743 | 0 | 0 | 0 | 1 |
| Q8NFI3 | ENGASE | CVLGtFItEWNEGGR | T193 | 0.03 | 98.09 | 743 | 0.02049 | 0.03 | 0.0205 | 1 |
| Q8NFI3 | ENGASE | VVGGrFDtDKSLELI | T344 | 0.004 | 98.69 | 743 | 0.002732 | 0.004 | 0.0027 | 1 |
| Q9UFP1 | GASK1A | LVRSSDPsHLVYIDN | S478 | 0.178 | 91.49 | 575 | 0.141495 | 0.178 | 0.1415 | 1 |
| Q96HY7 | DHTKD1 | VKTLVFCsGKHFYSL | S809 | 0.105 | 98.94 | 919 | 0.083466 | 0.105 | 0.0835 | 1 |
| P41222 | PTGDS | FRMAtLysRTQTPRA | S150 | 0.007 | 98.49 | 190 | 0.005564 | 0.007 | 0.0056 | 1 |
| P51685 | CCR8 | TLMSVDRyLAVVHAV | Y132 | 0.333 | 93.45 | 355 | 0.146082 | 0.333 | 0.1461 | 1 |
| Q9ULD8 | KCNH3 | RFLyHSRtRDLRDYI | T518 | 0.068 | 91.55 | 1083 | 0.046445 | 0.068 | 0.0464 | 0.875 |
| P07996 | THBS1 | VKGPDPSsPAFRIED | S56 | 0.157 | 83.07 | 1170 | 0.124802 | 0.157 | 0.1248 | 1 |
| P07996 | THBS1 | KGFLLLAsLRQMKKT | S93 | 0.139 | 94.34 | 1170 | 0.110493 | 0.139 | 0.1105 | 0.8889 |
| P36578 | RPL4 | ARPLISVysEkGESS | Y11 | 0.26 | 97.02 | 427 | 0.114058 | 0.26 | 0.1141 | 1 |
| P36578 | RPL4 | QKRyAICsALAAsAL | S126 | 0.001 | 98.14 | 427 | 0.000795 | 0.001 | 0.0008 | 1 |
| P36578 | RPL4 | ICsALAAsALPALVM | S131 | 0.061 | 96.95 | 427 | 0.04849 | 0.061 | 0.0485 | 1 |
| P36578 | RPL4 | FRNIPGItLLNVskL | T228 | 0.121 | 98.3 | 427 | 0.082645 | 0.121 | 0.0826 | 1 |
| P36578 | RPL4 | FRkLDELyGtWRkAA | Y264 | 0.081 | 96.81 | 427 | 0.035534 | 0.081 | 0.0355 | 1 |
| P36578 | RPL4 | AAsLkSNyNLPMHkM | Y277 | 0.17 | 96.7 | 427 | 0.074577 | 0.17 | 0.0746 | 1 |
| Q99259 | GAD1 | NFFRMVIsNPAATQs | S571 | 0.006 | 98.76 | 594 | 0.004769 | 0.006 | 0.0048 | 0.9091 |
| Q13535 | ATR | DTDQLKLyEEPLSkL | Y291 | 0.084 | 81.69 | 2644 | 0.03685 | 0.084 | 0.0369 | 0.9375 |
| Q8NCW6 | GALNT11 | LDEYKEQyFSLRPDL | Y414 | 0.097 | 93.03 | 608 | 0.042553 | 0.097 | 0.0426 | 1 |
| O95427 | PIGN | GHAHRPssRDyKHNI | S224 | 0.174 | 91.06 | 931 | 0.138315 | 0.174 | 0.1383 | 1 |
| O95427 | PIGN | HRPssRDyKHNIKKV | Y227 | 0.011 | 94.59 | 931 | 0.004826 | 0.011 | 0.0048 | 1 |
| Q15646 | OASL | KIRRTRGySGLQRLS | Y389 | 0.219 | 77.71 | 514 | 0.096072 | 0.219 | 0.0961 | 0.875 |
| Q14500 | KCNJ12 | KTYEVPStPRCSAKD | T354 | 0.136 | 94.14 | 433 | 0.09289 | 0.136 | 0.0929 | 1 |
| Q96JB8 | MPP4 | TCAGLVPsNHLLKRK | S304 | 0.008 | 93.8 | 637 | 0.006359 | 0.008 | 0.0064 | 1 |
| Q49A17 | GALNTL6 | AANLCVDsKHGATGT | S469 | 0.02 | 95.92 | 601 | 0.015898 | 0.02 | 0.0159 | 1 |
| Q9NV06 | DCAF13 | YIRALNAtkLERVFA | T48 | 0.046 | 97.15 | 445 | 0.031419 | 0.046 | 0.0314 | 1 |
| Q9NV06 | DCAF13 | kSRSREVyHTKRMQH | Y320 | 0.068 | 97.31 | 445 | 0.029831 | 0.068 | 0.0298 | 1 |
| Q12851 | MAP4K2 | CVNNVLLsLsGKSTH | S542 | 0 | 95.81 | 820 | 0 | 0 | 0 | 1 |

| | | | | | | | | | | |
|---|---|---|---|---|---|---|---|---|---|---|
| Q12851 | MAP4K2 | NNVLLsLsGKSTHIW | S544 | 0.05 | 90.78 | 820 | 0.039746 | 0.05 | 0.0397 | 1 |
| P14868 | DARS1 | IDLrTSTsQAVFRLQ | S192 | 0.08 | 97.99 | 501 | 0.063593 | 0.094 | 0.0747 | 1 |
| Q6PFW1 | PPIP5K1 | FRQKSGRyDISKIPD | Y730 | 0.121 | 93.16 | 1433 | 0.053081 | 0.121 | 0.0531 | 0.8 |
| Q6PFW1 | PPIP5K1 | TAELLRLskALADVV | S764 | 0.024 | 95.85 | 1433 | 0.019078 | 0.024 | 0.0191 | 1 |
| Q9H1Y3 | OPN3 | TVLAYERyIRVVHAR | Y140 | 0.279 | 96.58 | 402 | 0.122393 | 0.279 | 0.1224 | 1 |
| Q92643 | PIGK | RLLsDDRsNILIYMT | S155 | 0.003 | 96.57 | 395 | 0.002385 | 0.003 | 0.0024 | 0.9 |
| P50225 | SULT1A1 | GIPSGMEtLkDTPAP | T95 | 0.142 | 97.55 | 295 | 0.096988 | 0.142 | 0.097 | 1 |
| P24903 | CYP2F1 | SKEYGSMyTVHLGPR | Y66 | 0 | 96.95 | 491 | 0 | 0 | 0 | 1 |
| P24903 | CYP2F1 | ANQSFKKsPAFMPFs | S423 | 0.134 | 95.13 | 491 | 0.106519 | 0.134 | 0.1065 | 1 |
| Q9Y236 | OSGIN2 | kGGALGVtRCLATRQ | T482 | 0.001 | 98.73 | 505 | 0.000683 | 0.001 | 0.0007 | 1 |
| Q14168 | MPP2 | VNSNLERtFRELQTA | T552 | 0.021 | 97.13 | 576 | 0.014343 | 0.021 | 0.0143 | 1 |
| Q11203 | ST3GAL3 | QDFKWLKyIVyKERV | Y241 | 0.281 | 97.08 | 375 | 0.123271 | 0.281 | 0.1233 | 1 |
| Q86UP3 | ZFHX4 | YHAIRAAtMCNLCQR | T1380 | 0.081 | 83.17 | 3567 | 0.055324 | 0.081 | 0.0553 | 1 |
| O75530 | EED | GDPLVFAtVGSNRVT | T115 | 0.013 | 98.93 | 441 | 0.008879 | 0.013 | 0.0089 | 1 |
| O75530 | EED | MMNAIkEsyDyNPNk | S277 | 0 | 98.54 | 441 | 0 | 0 | 0 | 1 |
| Q8NH09 | OR8S1 | YMHLVWGsWGLGFLD | S148 | 0.06 | 97.26 | 359 | 0.047695 | 0.06 | 0.0477 | 1 |
| Q6N022 | TENM4 | GCKMRLSyLSSRTPG | Y1063 | 0.023 | 93.785 | 2769 | 0.01009 | 0.023 | 0.0101 | 1 |
| Q6N022 | TENM4 | LSSRTPGyKSVLRIS | Y1071 | 0.119 | 93.77 | 2769 | 0.052204 | 0.2575 | 0.113 | 1 |
| Q6ZVX9 | PAQR9 | VAAFFNVsKIPERIQ | S293 | 0.176 | 92.5 | 377 | 0.139905 | 0.176 | 0.1399 | 1 |
| O15297 | PPM1D | LDPQKHKyIILGSDG | Y308 | 0.113 | 97.79 | 605 | 0.049571 | 0.206 | 0.0832 | 0.8571 |
| O00203 | AP3B1 | SkQTkLLtQyILNLG | T555 | 0.039 | 94.57 | 1094 | 0.026638 | 0.039 | 0.0266 | 1 |
| Q6UXN9 | WDR82 | DDTFISGsLDkTIrL | S126 | 0 | 98.04 | 313 | 0 | 0 | 0 | 1 |
| Q9H706 | GAREM1 | FIREGHRyKFVNIQT | Y271 | 0.132 | 90.65 | 876 | 0.057907 | 0.359 | 0.1037 | 1 |
| P22059 | OSBP | VHFALLGtWDEKMEC | T645 | 0.178 | 97.92 | 807 | 0.121576 | 0.201 | 0.1373 | 1 |
| O75179 | ANKRD17 | GAEINSRtGSKLGIs | T1213 | 0.054 | 89.12 | 2603 | 0.036883 | 0.054 | 0.0369 | 1 |
| O75179 | ANKRD17 | tGSKLGIsPLMLAAM | S1220 | 0.058 | 94.63 | 2603 | 0.046105 | 0.058 | 0.0461 | 1 |
| Q15678 | PTPN14 | LSkNCFVtRIRLLDS | T23 | 0.003 | 90.87 | 1187 | 0.002049 | 0.003 | 0.002 | 1 |
| Q9H0U3 | MAGT1 | VKAPPRNysVIVMFt | Y72 | 0.004 | 93.49 | 335 | 0.001755 | 0.004 | 0.0018 | 1 |
| Q9H0U3 | MAGT1 | KAPPRNysVIVMFtA | S73 | 0.141 | 93.78 | 335 | 0.112083 | 0.141 | 0.1121 | 1 |
| Q9H0U3 | MAGT1 | ysVIVMFtALQLHRQ | T79 | 0.003 | 95.14 | 335 | 0.002049 | 0.003 | 0.002 | 1 |
| Q66K14 | TBC1D9B | IERDLHRsMPEHPAF | S560 | 0.011 | 86.39 | 1250 | 0.008744 | 0.011 | 0.0087 | 1 |

| | | | | | | | | | | |
|---|---|---|---|---|---|---|---|---|---|---|
| Q96K49 | TMEM87B | YsEYQNIsNTGLSTQ | S274 | 0.15 | 93.09 | 555 | 0.119237 | 0.15 | 0.1192 | 0.8571 |
| Q15286 | RAB35 | VkLQIWDtAGQERFR | T64 | 0.016 | 90.03 | 201 | 0.010928 | 0.016 | 0.0109 | 1 |
| Q15286 | RAB35 | QERFRtItstyYRGT | T74 | 0.019 | 70.68 | 201 | 0.012977 | 0.019 | 0.013 | 1 |
| Q6UWL2 | SUSD1 | ALYPGTNytVNISTA | Y368 | 0.057 | 81.68 | 747 | 0.025005 | 0.057 | 0.025 | 1 |
| Q13627 | DYRK1A | YDDDNyDyIVkNGEK | Y147 | 0.073 | 98.35 | 763 | 0.032024 | 0.073 | 0.032 | 1 |
| Q13627 | DYRK1A | GEKWMDRyEIDSLIG | Y159 | 0.051 | 98.74 | 763 | 0.022373 | 0.051 | 0.0224 | 1 |
| Q8NFJ9 | BBS1 | QMPAAILtMNLLEQH | T332 | 0.077 | 97.57 | 593 | 0.052592 | 0.077 | 0.0526 | 1 |
| Q8NFJ9 | BBS1 | ALLNVIHtPDAVTSL | T369 | 0.204 | 95.2 | 593 | 0.139335 | 0.204 | 0.1393 | 1 |
| Q8NFJ9 | BBS1 | TSLCFGRyGREDNtL | Y381 | 0.198 | 95.84 | 593 | 0.08686 | 0.198 | 0.0869 | 1 |
| Q8NFJ9 | BBS1 | RyGREDNtLIMttRG | T387 | 0.024 | 96.85 | 593 | 0.016392 | 0.024 | 0.0164 | 1 |
| Q8NFJ9 | BBS1 | EDNtLIMttRGGGLI | T391 | 0.021 | 96.96 | 593 | 0.014343 | 0.021 | 0.0143 | 1 |
| Q8NFJ9 | BBS1 | DNtLIMttRGGGLII | T392 | 0.075 | 96.88 | 593 | 0.051226 | 0.075 | 0.0512 | 1 |
| P48048 | KCNJ1 | RPKKRAKtItFSKNA | T191 | 0.205 | 88.28 | 391 | 0.140018 | 0.205 | 0.14 | 1 |
| P48048 | KCNJ1 | FSKNAVIskRGGKLC | S201 | 0.064 | 95.31 | 391 | 0.050875 | 0.064 | 0.0509 | 1 |
| Q02880 | TOP2B | YYkGLGtsTAkEAkE | S640 | 0.183 | 83.46 | 1626 | 0.145469 | 0.183 | 0.1455 | 1 |
| Q02880 | TOP2B | GLPEQFLyGTATkHL | Y707 | 0.211 | 75.08 | 1626 | 0.092563 | 0.211 | 0.0926 | 1 |
| Q02880 | TOP2B | DNERsIPsLVDGFkP | S738 | 0.095 | 95.41 | 1626 | 0.075517 | 0.095 | 0.0755 | 1 |
| Q02880 | TOP2B | WACkLPNyDAREIVN | Y888 | 0.026 | 96.68 | 1626 | 0.011406 | 0.026 | 0.0114 | 1 |
| Q02880 | TOP2B | PHPMLPNyKNFKGTI | Y913 | 0.181 | 95.93 | 1626 | 0.079402 | 0.181 | 0.0794 | 1 |
| Q02880 | TOP2B | QELGQNQyAVSGEIF | Y928 | 0.18 | 93.47 | 1626 | 0.078963 | 0.18 | 0.079 | 1 |
| Q9H2C0 | GAN | kNILAAAsPYIRTkL | S52 | 0.024 | 96.02 | 597 | 0.019078 | 0.024 | 0.0191 | 1 |
| O75147 | OBSL1 | EMRGRVRtVANVtVk | T420 | 0.084 | 86.32 | 1896 | 0.057373 | 0.084 | 0.0574 | 0.9 |
| P13569 | CFTR | QPyIFVAtVPVIVAF | T1019 | 0.014 | 83.13 | 1480 | 0.009562 | 0.014 | 0.0096 | 1 |
| P02461 | COL3A1 | RASQNITyHCKNSIA | Y1370 | 0.013 | 98.7 | 1466 | 0.005703 | 0.013 | 0.0057 | 1 |
| O14917 | PCDH17 | GRSKSGSyRVLENSA | Y64 | 0.24 | 88.28 | 1159 | 0.105285 | 0.277 | 0.1215 | 1 |
| O14917 | PCDH17 | PSsGEIRtLHPFWED | T647 | 0.016 | 87.29 | 1159 | 0.010928 | 0.114 | 0.0779 | 1 |
| Q9BQB6 | VKORC1 | VGTAISCsRVFSSRW | S52 | 0.027 | 92.06 | 163 | 0.021463 | 0.027 | 0.0215 | 1 |
| Q9Y2B1 | RXYLT1 | RTQYSFItGPAVIPG | T146 | 0.169 | 95.83 | 443 | 0.115429 | 0.169 | 0.1154 | 1 |
| Q13823 | GNL2 | NEAQEEIykKGQSKR | Y198 | 0.133 | 89.29 | 731 | 0.058345 | 0.133 | 0.0583 | 1 |
| Q13823 | GNL2 | RDPMGTRsPHIETYL | S234 | 0.151 | 91.35 | 731 | 0.120032 | 0.151 | 0.12 | 1 |
| Q13823 | GNL2 | HTDKKQIsVGFIGyP | S312 | 0.095 | 96.85 | 731 | 0.075517 | 0.095 | 0.0755 | 1 |

| | | | | | | | | | | |
|---|---|---|---|---|---|---|---|---|---|---|
| Q13823 | GNL2 | IsVGFIGyPNVGkSS | Y318 | 0.119 | 94.47 | 731 | 0.052204 | 0.119 | 0.0522 | 1 |
| Q13823 | GNL2 | GkSSVINtLRSKKVC | T329 | 0.045 | 96.86 | 731 | 0.030736 | 0.045 | 0.0307 | 1 |
| Q7Z4N2 | TRPM1 | GKDVTRVyQtMSNPL | Y189 | 0.006 | 87.56 | 1603 | 0.002632 | 0.006 | 0.0026 | 1 |
| Q9P209 | CEP72 | GLkSLDLsRNSLVSL | S62 | 0.134 | 93.15 | 647 | 0.106519 | 0.134 | 0.1065 | 1 |
| Q7L590 | MCM10 | LEEIDWVtFGVILKk | T296 | 0.002 | 93.61 | 875 | 0.001366 | 0.002 | 0.0014 | 1 |
| P40337 | VHL | SVNsREPsQVIFCNr | S72 | 0.01 | 97.48 | 213 | 0.007949 | 0.01 | 0.0079 | 1 |
| P40337 | VHL | QVIFCNrsPRVVLPV | S80 | 0.07 | 98.7 | 213 | 0.055644 | 0.096 | 0.092 | 1 |
| P40337 | VHL | GtGRRIHsyRGHLWL | S111 | 0.04 | 97.37 | 213 | 0.031797 | 0.07 | 0.0556 | 0.875 |
| P40337 | VHL | sLVkPENyRRLDIVR | Y175 | 0.044 | 98.23 | 213 | 0.019302 | 0.044 | 0.0193 | 1 |
| Q92994 | BRF1 | RLTEFEDtPTSQLTI | T270 | 0.069 | 89.56 | 677 | 0.047128 | 0.069 | 0.0471 | 1 |
| Q5BKX5 | ACTMAP | NLQLTDFsPSRATDG | S316 | 0.074 | 96.85 | 351 | 0.058824 | 0.074 | 0.0588 | 1 |
| Q86XL3 | ANKLE2 | FLFGEEPskLDQDVL | S832 | 0.077 | 89.68 | 938 | 0.061208 | 0.077 | 0.0612 | 1 |
| Q9NYU2 | UGGT1 | HDQDVVEyIMNQPNV | Y684 | 0.21 | 90.45 | 1555 | 0.092124 | 0.21 | 0.0921 | 1 |
| Q9Y6Q5 | AP1M2 | _____MsAsAVFILDV | S4 | 0.006 | 96.04 | 423 | 0.004769 | 0.006 | 0.0048 | 1 |
| Q9Y6Q5 | AP1M2 | SNLYLVAtTSKNANA | T70 | 0.024 | 97.77 | 423 | 0.016392 | 0.042 | 0.028 | 0.9091 |
| P62753 | RPS6 | LGEEWkGyVVRIsGG | Y48 | 0.015 | 96.88 | 249 | 0.00658 | 0.015 | 0.0066 | 1 |
| Q13443 | ADAM9 | MNSGAsGsRNFSSCS | S379 | 0.111 | 82.18 | 819 | 0.088236 | 0.111 | 0.0882 | 0.8571 |
| Q13616 | CUL1 | FINNNAVtkMAQSSS | T409 | 0.085 | 89 | 776 | 0.058056 | 0.085 | 0.0581 | 1 |
| P16333 | NCK1 | VQLKETVyCIGQRkF | Y339 | 0.112 | 92.5 | 377 | 0.049133 | 0.112 | 0.0491 | 0.8182 |
| Q9H4I3 | TRABD | SkRDVVktIREVQPD | T94 | 0 | 98.53 | 376 | 0 | 0 | 0 | 1 |
| Q9NZB2 | FAM120A | AFLAQALsPKLYEPD | S745 | 0.014 | 95.54 | 1118 | 0.011129 | 0.014 | 0.0111 | 1 |
| Q9NZB2 | FAM120A | kVEKMRQsVLEGLsF | S841 | 0.01 | 96.87 | 1118 | 0.007949 | 0.01 | 0.0079 | 1 |
| Q9P2D0 | IBTK | VAAGRFHtVLWtREA | T296 | 0 | 98.29 | 1353 | 0 | 0 | 0 | 1 |
| Q9P2D0 | IBTK | RFHtVLWtREAVYTM | T300 | 0.078 | 95.3 | 1353 | 0.053275 | 0.078 | 0.0533 | 1 |
| P22307 | SCP2 | GWKNHKHsVNNPysQ | S192 | 0 | 98.64 | 547 | 0 | 0 | 0 | 1 |
| P22307 | SCP2 | HsVNNPysQFQDEys | S198 | 0 | 98.42 | 547 | 0 | 0 | 0 | 0.8889 |
| P22307 | SCP2 | EMMTDLPssFEEKSI | S265 | 0.166 | 97.75 | 547 | 0.131956 | 0.166 | 0.132 | 1 |
| P22307 | SCP2 | MMTDLPssFEEKSII | S266 | 0.023 | 97.28 | 547 | 0.018283 | 0.023 | 0.0183 | 0.875 |
| Q8N4X5 | AFAP1L2 | RDNHLHFyQDRNRSK | Y383 | 0.292 | 94.35 | 818 | 0.128096 | 0.292 | 0.1281 | 1 |
| O94913 | PCF11 | PVMYLMDsIVKNVGR | S79 | 0.146 | 95.01 | 1555 | 0.116058 | 0.146 | 0.1161 | 1 |
| P07864 | LDHC | CKITIVGtGAVGMAC | T28 | 0.008 | 97.86 | 332 | 0.005464 | 0.008 | 0.0055 | 0.8 |

| | | | | | | | | | | |
|---|---|---|---|---|---|---|---|---|---|---|
| Q6PIF6 | MYO7B | PVDSASTsREMCMHI | S1219 | 0.001 | 88.79 | 2116 | 0.000795 | 0.001 | 0.0008 | 1 |
| Q6PIF6 | MYO7B | ELLARHCyVQLGASA | Y1336 | 0.126 | 88.6 | 2116 | 0.055274 | 0.238 | 0.101 | 1 |
| Q13554 | CAMK2B | sMMHRQEtVECLKKF | T287 | 0.047 | 91.83 | 666 | 0.032102 | 0.047 | 0.0321 | 1 |
| Q8IWW6 | ARHGAP12 | ERGHtLytsDytNEK | T356 | 0.173 | 89.09 | 846 | 0.118161 | 0.173 | 0.1182 | 1 |
| Q8IWW6 | ARHGAP12 | RGHtLytsDytNEKW | S357 | 0.008 | 87.22 | 846 | 0.006359 | 0.089 | 0.0707 | 1 |
| Q00341 | HDLBP | RHEVLLIsAEQDkRA | S216 | 0.172 | 78.52 | 1268 | 0.136725 | 0.172 | 0.1367 | 1 |
| Q00341 | HDLBP | VIGPkGNsLQEILER | S317 | 0.158 | 80.85 | 1268 | 0.125597 | 0.158 | 0.1256 | 1 |
| P04844 | RPN2 | RPFtNLEsAFYSIVG | S50 | 0.082 | 96.19 | 631 | 0.065183 | 0.129 | 0.115 | 1 |
| P04844 | RPN2 | ATVQALQtASHLSQQ | T167 | 0.004 | 97.78 | 631 | 0.002732 | 0.004 | 0.0027 | 1 |
| P04844 | RPN2 | ALFVAATyKLMDHVG | Y216 | 0.035 | 97.55 | 631 | 0.015354 | 0.035 | 0.0154 | 1 |
| O15530 | PDPK1 | LTEGPHLyyVDPVNK | Y485 | 0.075 | 96.7 | 556 | 0.032901 | 0.075 | 0.0329 | 0.8 |
| Q2UY09 | COL28A1 | YYAIsNAtRLLKREG | T141 | 0.087 | 95.14 | 1125 | 0.059422 | 0.087 | 0.0594 | 1 |
| Q9H1D0 | TRPV6 | GPLTSTLyDLTEIDS | Y318 | 0.214 | 93.83 | 765 | 0.093879 | 0.214 | 0.0939 | 1 |
| Q9Y6H5 | SNCAIP | ERNTEKLtPAGLAIK | T387 | 0.003 | 93.6 | 919 | 0.002049 | 0.003 | 0.002 | 0.9 |
| Q75T13 | PGAP1 | GGSLQKQtKFVHECI | T144 | 0 | 98.39 | 922 | 0 | 0 | 0 | 0.9167 |
| Q9UJW3 | DNMT3L | CDRPPSWyLFQFHRL | Y259 | 0.23 | 96.33 | 386 | 0.100898 | 0.23 | 0.1009 | 1 |
| Q8N5Y8 | PARP16 | LAADLRCsLFASALQ | S25 | 0.001 | 98.55 | 322 | 0.000795 | 0.001 | 0.0008 | 0.8 |
| P28289 | TMOD1 | KKFSIVGtRsNDPVA | T234 | 0.004 | 94.32 | 359 | 0.002732 | 0.004 | 0.0027 | 1 |
| P28289 | TMOD1 | FSIVGtRsNDPVAyA | S236 | 0.003 | 93.08 | 359 | 0.002385 | 0.003 | 0.0024 | 1 |
| P31150 | GDI1 | YGksPyLyPLyGLGE | Y226 | 0.004 | 97.31 | 447 | 0.001755 | 0.004 | 0.0018 | 1 |
| P31150 | GDI1 | DIyVCMIsyAHNVAA | S338 | 0 | 97.91 | 447 | 0 | 0 | 0 | 1 |
| P31150 | GDI1 | NVAAQGKyIAIASTT | Y349 | 0.179 | 98.53 | 447 | 0.078525 | 0.179 | 0.0785 | 1 |
| Q13231 | CHIT1 | NRQTFVNsAIRFLRK | S122 | 0.09 | 98.9 | 466 | 0.071542 | 0.09 | 0.0715 | 1 |
| Q9NP77 | SSU72 | CSSNQNRsMEAHNIL | S19 | 0 | 98.74 | 194 | 0 | 0 | 0 | 1 |
| P40426 | PBX3 | FQEEANLyAAKTAVT | Y307 | 0.331 | 90.58 | 434 | 0.145205 | 0.331 | 0.1452 | 0.8182 |
| Q9HCD6 | TANC2 | ISRLVALsCHGtRMR | S387 | 0.136 | 92.7 | 1990 | 0.108108 | 0.136 | 0.1081 | 1 |
| Q92478 | CLEC2B | SEGCAyLsDDGAATA | S127 | 0.096 | 97.67 | 149 | 0.076312 | 0.096 | 0.0763 | 1 |
| O00141 | SGK1 | HkAEEVFyAVkVLQK | Y124 | 0.154 | 95.07 | 431 | 0.067558 | 0.154 | 0.0676 | 1 |
| Q6P179 | ERAP2 | RRNISDIsENLKRYL | S719 | 0.061 | 97.08 | 960 | 0.04849 | 0.061 | 0.0485 | 1 |
| Q2M3M2 | SLC5A9 | IVMPGMIsRALFPDE | S339 | 0 | 97.36 | 681 | 0 | 0 | 0 | 1 |
| Q2M3M2 | SLC5A9 | CGARVGCsNIAyPKL | S366 | 0.089 | 92.51 | 681 | 0.070747 | 0.089 | 0.0707 | 1 |

| | | | | | | | | | | |
|---|---|---|---|---|---|---|---|---|---|---|
| Q2M3M2 | SLC5A9 | VGCsNIAyPKLVMAL | Y370 | 0.007 | 93.61 | 681 | 0.003071 | 0.007 | 0.0031 | 0.9474 |
| Q6P2H8 | TMEM53 | LAKYSAIyHKRGCIV | Y58 | 0.003 | 98 | 277 | 0.001316 | 0.003 | 0.0013 | 0.9333 |
| Q96RL6 | SIGLEC11 | SREVEMstRDRFQLT | T99 | 0.038 | 90.41 | 698 | 0.025955 | 0.038 | 0.026 | 1 |
| Q01968 | OCRL | IATETVGtGIMGKMG | T346 | 0.029 | 91 | 901 | 0.019807 | 0.029 | 0.0198 | 1 |
| Q01968 | OCRL | FERRNQDykDICARM | Y390 | 0.196 | 97.71 | 901 | 0.085982 | 0.196 | 0.086 | 1 |
| Q01968 | OCRL | TSDHKPVsALFHIGV | S528 | 0 | 98.85 | 901 | 0 | 0 | 0 | 1 |
| Q6P9A2 | GALNT18 | QCKTFRWyLVsVYPE | Y453 | 0 | 98.61 | 607 | 0 | 0 | 0 | 0.9333 |
| Q9UJS0 | SLC25A13 | SHQVSFsyFNGFNSL | Y219 | 0.171 | 96.26 | 675 | 0.075015 | 0.171 | 0.075 | 1 |
| Q96J66 | ABCC11 | GIVGRtGsGKSSLGM | S1179 | 0.082 | 80.44 | 1382 | 0.065183 | 0.084 | 0.0953 | 1 |
| Q96J66 | ABCC11 | IWDALERtFLtKAIs | T1254 | 0.004 | 84.98 | 1382 | 0.002732 | 0.004 | 0.0027 | 1 |
| Q96J66 | ABCC11 | VLRKKPGsLFAALMA | S1368 | 0.047 | 77.01 | 1382 | 0.037361 | 0.09 | 0.073 | 1 |
| P42695 | NCAPD3 | MCDLCIRyTIMVDKY | Y972 | 0.306 | 92.35 | 1498 | 0.134238 | 0.306 | 0.1342 | 1 |
| Q53H76 | PLA1A | KGIIAHAtPQCQINQ | T389 | 0.213 | 92.82 | 456 | 0.145482 | 0.213 | 0.1455 | 1 |
| P11766 | ADH5 | TAFGGWksVESVPkL | S324 | 0.005 | 98.65 | 374 | 0.003975 | 0.005 | 0.004 | 0.875 |
| Q9UGK3 | STAP2 | ADGVSVTtRQMHNGT | T191 | 0.019 | 95.17 | 403 | 0.012977 | 0.019 | 0.013 | 1 |
| Q96FB5 | METTL25B | RNLVLVAtKMPLGQA | T459 | 0.055 | 97.65 | 475 | 0.037566 | 0.177 | 0.1209 | 1 |
| Q15477 | SKIC2 | LHLERHDsVFVAAHT | S327 | 0.075 | 96.25 | 1246 | 0.059619 | 0.075 | 0.0596 | 1 |
| Q15477 | SKIC2 | kHMTRTIytsPIkAL | Y358 | 0.003 | 96.86 | 1246 | 0.001316 | 0.003 | 0.0013 | 1 |
| Q15477 | SKIC2 | HMTRTIytsPIkALS | T359 | 0.001 | 95.23 | 1246 | 0.000683 | 0.001 | 0.0007 | 1 |
| Q15477 | SKIC2 | MTRTIytsPIkALSN | S360 | 0 | 90.22 | 1246 | 0 | 0 | 0 | 1 |
| O00571 | DDX3X | RVRPCVVyGGADIGQ | Y301 | 0.303 | 94.43 | 662 | 0.132922 | 0.303 | 0.1329 | 0.8 |
| O00571 | DDX3X | GCHLLVAtPGRLVDM | T323 | 0.123 | 95.71 | 662 | 0.084011 | 0.123 | 0.084 | 0.8182 |
| Q9Y6N7 | ROBO1 | YSVEVAAsTGAGSGV | S855 | 0 | 91.93 | 1651 | 0 | 0 | 0 | 1 |
| Q9UG01 | IFT172 | NYKILHKyVALYATH | Y1445 | 0.015 | 85.88 | 1749 | 0.00658 | 0.015 | 0.0066 | 1 |
| P05556 | ITGB1 | kFKRAEDyPIDLYyL | Y141 | 0.209 | 86.25 | 798 | 0.091685 | 0.209 | 0.0917 | 1 |
| P05556 | ITGB1 | DyPIDLYyLMDLSYS | Y147 | 0.001 | 97.05 | 798 | 0.000439 | 0.001 | 0.0004 | 1 |
| P05556 | ITGB1 | DFRIGFGsFVEkTVM | S186 | 0.001 | 97.2 | 798 | 0.000795 | 0.001 | 0.0008 | 1 |
| P05556 | ITGB1 | LIGWRNVtRLLVFST | T271 | 0.111 | 91.63 | 798 | 0.075815 | 0.111 | 0.0758 | 1 |
| Q9NX78 | TMEM260 | ILAAGVFsFsRLTWQ | S131 | 0.009 | 98.53 | 707 | 0.007154 | 0.009 | 0.0072 | 1 |
| Q9NX78 | TMEM260 | AAGVFsFsRLTWQWS | S133 | 0 | 97.73 | 707 | 0 | 0 | 0 | 1 |
| Q9NX78 | TMEM260 | MWQARMKtPFFIFNL | T596 | 0.127 | 95.98 | 707 | 0.086743 | 0.127 | 0.0867 | 1 |

| | | | | | | | | | | |
|---|---|---|---|---|---|---|---|---|---|---|
| Q8TDX7 | NEK7 | RPDMGYNtLANFRIE | T30 | 0.045 | 96.8 | 302 | 0.030736 | 0.045 | 0.0307 | 1 |
| Q5VU65 | NUP210L | EHAIEPDtKLTFVGD | T1486 | 0.072 | 82.09 | 1888 | 0.049177 | 0.072 | 0.0492 | 1 |
| P49411 | TUFM | DHGKTTLtAAITkIL | T77 | 0.001 | 95.07 | 452 | 0.000683 | 0.001 | 0.0007 | 1 |
| P49411 | TUFM | ETPVIVGsALCALEG | S222 | 0.006 | 96.46 | 452 | 0.004769 | 0.006 | 0.0048 | 1 |
| P19338 | NCL | LkEsFDGsVrARIVt | S595 | 0.145 | 84.2 | 710 | 0.115263 | 0.145 | 0.1153 | 0.875 |
| P21246 | PTN | QFGAECKyQFQAWGE | Y101 | 0.293 | 88.03 | 168 | 0.128535 | 0.293 | 0.1285 | 1 |
| Q13526 | PIN1 | sQFSDCssAkARGDL | S115 | 0.112 | 98.38 | 163 | 0.08903 | 0.112 | 0.089 | 1 |
| Q9BTD8 | RBM42 | RAFSRFPsFLKAKVI | S406 | 0.152 | 93.63 | 480 | 0.120827 | 0.152 | 0.1208 | 1 |
| Q63HM2 | PCNX4 | FLKRFPQtVLGGPRF | T23 | 0.013 | 87.88 | 1172 | 0.008879 | 0.004 | 0.0262 | 1 |
| O75581 | LRP6 | KLNFIHKsNLDGtNR | S209 | 0.013 | 95.02 | 1613 | 0.010334 | 0.013 | 0.0103 | 1 |
| Q13671 | RIN1 | LRERLLLtRPVWLQL | T65 | 0 | 94.51 | 783 | 0 | 0 | 0 | 0.9091 |
| Q8IUH5 | ZDHHC17 | MMDQNGMtPLMWAAY | T192 | 0.023 | 98.32 | 632 | 0.015709 | 0.023 | 0.0157 | 1 |
| Q6HA08 | ASTL | ITRVLKLyGCSPSGP | Y279 | 0.187 | 95.77 | 431 | 0.082034 | 0.187 | 0.082 | 1 |
| Q96BY7 | ATG2B | SVAVKILsDKSESNT | S1179 | 0.132 | 84.19 | 2078 | 0.104929 | 0.132 | 0.1049 | 1 |
| Q5DID0 | UMODL1 | SAPGNAStTVsRLLL | T225 | 0.179 | 85.06 | 1318 | 0.122259 | 0.179 | 0.1223 | 1 |
| O75369 | FLNB | IAQEPGNyEVSIKFN | Y2260 | 0.001 | 78.74 | 2602 | 0.000439 | 0.001 | 0.0004 | 1 |
| Q9H477 | RBKS | KDSFGNDyIENLKQN | Y84 | 0.008 | 98.7 | 322 | 0.003509 | 0.008 | 0.0035 | 0.8889 |
| O43813 | LANCL1 | NkSLAEGyFDAAGRL | Y21 | 0.209 | 98.35 | 399 | 0.091685 | 0.209 | 0.0917 | 1 |
| O43813 | LANCL1 | AHGYVKQsLNCLtkR | S95 | 0.001 | 98.51 | 399 | 0.000795 | 0.001 | 0.0008 | 1 |
| O43813 | LANCL1 | LVkPSVDyVCQLKFP | Y250 | 0.155 | 98.85 | 399 | 0.067996 | 0.213 | 0.0934 | 1 |
| O43813 | LANCL1 | YGLLKKGyGLCHGSA | Y319 | 0.097 | 98.47 | 399 | 0.042553 | 0.181 | 0.0794 | 1 |
| O43813 | LANCL1 | HGSAGNAyAFLTLyN | Y330 | 0.013 | 98.9 | 399 | 0.005703 | 0.013 | 0.0057 | 0.8235 |
| O43813 | LANCL1 | AyAFLTLyNLTQDMK | Y336 | 0.02 | 98.73 | 399 | 0.008774 | 0.02 | 0.0088 | 1 |
| Q86UL3 | GPAT4 | KHVHLMCyRICVRAL | Y218 | 0.181 | 94.94 | 456 | 0.079402 | 0.181 | 0.0794 | 1 |
| Q86UL3 | GPAT4 | AIVCSVWyLPPMtRE | Y387 | 0.026 | 98.69 | 456 | 0.011406 | 0.026 | 0.0114 | 1 |
| Q7L590 | MCM10 | PCGNRSIsLDrLPNk | S824 | 0.123 | 91.32 | 875 | 0.097775 | 0.123 | 0.0978 | 1 |
| Q9Y5Z7 | HCFC2 | GTVPLPRsLHtAsVI | S246 | 0.026 | 97.51 | 792 | 0.020668 | 0.026 | 0.0207 | 1 |
| Q9Y5Z7 | HCFC2 | PLPRsLHtAsVIGNK | T249 | 0.121 | 98.58 | 792 | 0.082645 | 0.121 | 0.0826 | 1 |
| Q9Y5Z7 | HCFC2 | PRsLHtAsVIGNKMY | S251 | 0.138 | 97.8 | 792 | 0.109698 | 0.138 | 0.1097 | 1 |
| Q96PB1 | CASD1 | TNEKIDAyNEAAVSI | Y232 | 0.027 | 96.38 | 797 | 0.011845 | 0.027 | 0.0118 | 1 |
| P00352 | ALDH1A1 | RDRLLLAtMESMNGG | T105 | 0.144 | 98.72 | 501 | 0.098354 | 0.144 | 0.0984 | 1 |

| | | | | | | | | | | |
|---|---|---|---|---|---|---|---|---|---|---|
| P00352 | ALDH1A1 | GKLYSNAyLNDLAGC | Y119 | 0.322 | 98.71 | 501 | 0.141257 | 0.322 | 0.1413 | 1 |
| P00352 | ALDH1A1 | KSNLKRVtLELGGKS | T267 | 0.144 | 98.89 | 501 | 0.098354 | 0.144 | 0.0984 | 1 |
| P00352 | ALDH1A1 | IFVEEsIyDEFVRRS | Y316 | 0.105 | 98.93 | 501 | 0.046062 | 0.105 | 0.0461 | 1 |
| P00352 | ALDH1A1 | kILDLIEsGkKEGAk | S360 | 0.092 | 98.88 | 501 | 0.073132 | 0.092 | 0.0731 | 1 |
| P00352 | ALDH1A1 | NNtFyGLsAGVFTkD | S429 | 0.018 | 98.86 | 501 | 0.014308 | 0.018 | 0.0143 | 0.9091 |
| P00352 | ALDH1A1 | GEyGFHEyTEVkTVt | Y486 | 0.076 | 98.93 | 501 | 0.03334 | 0.076 | 0.0333 | 1 |
| Q5T2S8 | ODAD2 | VTFLREKsPKFSENM | S302 | 0.015 | 88.38 | 1044 | 0.011924 | 0.015 | 0.0119 | 1 |
| P33991 | MCM4 | QMIKLQEsPEDMPAG | S357 | 0.077 | 80.06 | 863 | 0.061208 | 0.077 | 0.0612 | 1 |
| P33991 | MCM4 | rVNVTGIyRAVPIRV | Y394 | 0.22 | 91.32 | 863 | 0.096511 | 0.22 | 0.0965 | 1 |
| Q86X52 | CHSY1 | GFGITCIyKGDLVRV | Y699 | 0.09 | 98.51 | 802 | 0.039482 | 0.09 | 0.0395 | 0.9286 |
| Q99615 | DNAJC7 | YNEAyNyyTkAIDMC | Y51 | 0.042 | 98.3 | 494 | 0.018425 | 0.042 | 0.0184 | 1 |
| Q9BWV7 | TTLL2 | MLGTKHSyWICKPAE | Y208 | 0.327 | 94.51 | 592 | 0.14345 | 0.327 | 0.1434 | 0.8 |
| P51841 | GUCY2F | SYPTFSRtLPsPIRV | T168 | 0.048 | 93.69 | 1108 | 0.032785 | 0.048 | 0.0328 | 1 |
| Q8IWZ8 | SUGP1 | AFLHDKNsREFLYYR | S228 | 0.146 | 86.49 | 645 | 0.116058 | 0.146 | 0.1161 | 1 |
| Q96P53 | WDFY2 | NPETRRLsIGLDNGT | S84 | 0.031 | 98.67 | 400 | 0.024642 | 0.062 | 0.0493 | 1 |
| Q96P53 | WDFY2 | ATRGWLLtSGtDKVI | T383 | 0 | 98.14 | 400 | 0 | 0 | 0 | 1 |
| Q92636 | NSMAF | MPEPKFMyGSHYSSP | Y388 | 0.03 | 95.67 | 917 | 0.013161 | 0.03 | 0.0132 | 1 |
| Q5W041 | ARMC3 | KIVRRNAtMIFGILA | T88 | 0.036 | 96.48 | 872 | 0.024588 | 0.036 | 0.0246 | 1 |
| Q5W041 | ARMC3 | ELKFQLKsNVIPIGH | S782 | 0.092 | 92.52 | 872 | 0.073132 | 0.148 | 0.1217 | 1 |
| Q86VF7 | NRAP | VSHQKKPyCHAHNPK | Y53 | 0.181 | 85.94 | 1730 | 0.079402 | 0.181 | 0.0794 | 1 |
| Q53GL7 | PARP10 | YFENRRRsGGGPVLs | S37 | 0 | 85.72 | 1025 | 0 | 0 | 0 | 1 |
| Q53GL7 | PARP10 | YLENERRsGGGPLED | S199 | 0 | 84.62 | 1025 | 0 | 0 | 0 | 1 |
| Q9Y4C8 | RBM19 | IKNLNFDtTEEKLKE | T741 | 0.09 | 89.93 | 960 | 0.061471 | 0.09 | 0.0615 | 1 |
| P12694 | BCKDHA | KEKVLKLykSMTLLN | Y106 | 0 | 98.9 | 445 | 0 | 0 | 0 | 1 |
| P12694 | BCKDHA | GNDVFAVyNATkEAR | Y307 | 0.197 | 98.94 | 445 | 0.086421 | 0.197 | 0.0864 | 1 |
| P12694 | BCKDHA | TYRIGHHstsDDssA | S337 | 0.088 | 95.77 | 445 | 0.069952 | 0.088 | 0.07 | 1 |
| P12694 | BCKDHA | YRIGHHstsDDssAy | T338 | 0.119 | 95.44 | 445 | 0.081279 | 0.119 | 0.0813 | 1 |
| P12694 | BCKDHA | HHstsDDssAyRsVD | S342 | 0.087 | 92.92 | 445 | 0.069158 | 0.087 | 0.0692 | 1 |
| P23229 | ITGA6 | EVGSLIEyEFRVINL | Y871 | 0.041 | 94.46 | 1130 | 0.017986 | 0.266 | 0.1167 | 1 |
| P49448 | GLUD2 | CKGGIRYstDVsVDE | S153 | 0.152 | 96.17 | 558 | 0.120827 | 0.152 | 0.1208 | 1 |
| P49448 | GLUD2 | EKITRRFtMELAkKG | T206 | 0.041 | 98.77 | 558 | 0.028004 | 0.041 | 0.028 | 1 |

| | | | | | | | | | | |
|---|---|---|---|---|---|---|---|---|---|---|
| P49448 | GLUD2 | AEGANGPttPEADkI | T409 | 0.008 | 98.14 | 558 | 0.005464 | 0.008 | 0.0055 | 1 |
| P49448 | GLUD2 | EGANGPttPEADkIF | T410 | 0 | 97.86 | 558 | 0 | 0 | 0 | 1 |
| P49448 | GLUD2 | SARQIMHtAMkyNLG | T524 | 0.117 | 98.44 | 558 | 0.079913 | 0.117 | 0.0799 | 1 |
| P49448 | GLUD2 | LDLRTAAyVNAIEkV | Y539 | 0.016 | 98.71 | 558 | 0.007019 | 0.016 | 0.007 | 1 |
| P36955 | SERPINF1 | LIEESLtsEFIHDID | S295 | 0.061 | 98.47 | 418 | 0.04849 | 0.061 | 0.0485 | 0.8 |
| P46199 | MTIF2 | EKITFLDtPGHAAFS | T235 | 0.009 | 92.15 | 727 | 0.006147 | 0.009 | 0.0061 | 1 |
| P46199 | MTIF2 | PSMPVGItGWRDLPs | T419 | 0.056 | 96.9 | 727 | 0.038249 | 0.056 | 0.0382 | 1 |
| P48380 | RFX3 | GIRVkPDsPLNRLQE | S260 | 0.091 | 88.75 | 749 | 0.072337 | 0.091 | 0.0723 | 1 |
| P51690 | ARSL | VKLTQHIsAAsLCTP | S81 | 0.011 | 98.66 | 589 | 0.008744 | 0.011 | 0.0087 | 1 |
| P51690 | ARSL | TQHIsAAsLCTPSRA | S84 | 0.023 | 97.84 | 589 | 0.018283 | 0.023 | 0.0183 | 1 |
| P51690 | ARSL | AAFLTGRyPVRSGMV | Y98 | 0.097 | 98.76 | 589 | 0.042553 | 0.097 | 0.0426 | 1 |
| Q9Y2K9 | STXBP5L | LIGYENGtVVFWDLK | T232 | 0.059 | 96.32 | 1186 | 0.040298 | 0.059 | 0.0403 | 1 |
| Q2TB90 | HKDC1 | LPLGLTFsFPCRQTK | S155 | 0.072 | 97.03 | 917 | 0.057234 | 0.072 | 0.0572 | 1 |
| Q2TB90 | HKDC1 | VGMDGTLykIHPQYP | Y417 | 0.181 | 97.14 | 917 | 0.079402 | 0.181 | 0.0794 | 1 |
| Q2TB90 | HKDC1 | EYGLkKksHGLAtVR | S503 | 0.108 | 96.24 | 917 | 0.085851 | 0.108 | 0.0859 | 1 |
| Q2TB90 | HKDC1 | AtVRMLPtYVCGLPD | T514 | 0.054 | 96.44 | 917 | 0.036883 | 0.054 | 0.0369 | 1 |
| Q2TB90 | HKDC1 | EKMTSGMyLGEIVRQ | Y748 | 0.109 | 95.85 | 917 | 0.047817 | 0.109 | 0.0478 | 1 |
| Q2TB90 | HKDC1 | FSRILQEtVKELAPR | T877 | 0.092 | 98.75 | 917 | 0.062837 | 0.092 | 0.0628 | 1 |
| Q2TB90 | HKDC1 | CDVTFMLsEDGSGKG | S892 | 0.026 | 88.97 | 917 | 0.020668 | 0.026 | 0.0207 | 1 |
| Q9Y283 | INVS | KDPsAVHtVRCILDA | T200 | 0.008 | 96.26 | 1065 | 0.005464 | 0.008 | 0.0055 | 1 |
| P56470 | LGALS4 | WDKVVFNtLQGGkWG | T78 | 0.085 | 98.77 | 323 | 0.058056 | 0.085 | 0.0581 | 0.8 |
| P0DMV8 | HSPA1A | GNRttPsyVAFtDtE | Y41 | 0.141 | 98.43 | 641 | 0.061855 | 0.27 | 0.1142 | 0.8 |
| P0DMV8 | HSPA1A | VALNPQNtVFDAkRL | T66 | 0 | 98.04 | 641 | 0 | 0 | 0 | 1 |
| P0DMV8 | HSPA1A | kPkVQVsykGETkAF | Y107 | 0.016 | 94.68 | 641 | 0.007019 | 0.254 | 0.0997 | 1 |
| P0DMV8 | HSPA1A | PVtNAVItVPAyFND | T145 | 0.008 | 97.59 | 641 | 0.005464 | 0.008 | 0.0055 | 0.9167 |
| Q8IXH7 | NELFCD | HKFStRDyIMEPSIF | Y56 | 0.234 | 93.96 | 590 | 0.102652 | 0.234 | 0.1027 | 1 |
| Q8IXH7 | NELFCD | MEPSIFNtLkRYFQA | T65 | 0.126 | 91.97 | 590 | 0.08606 | 0.126 | 0.0861 | 1 |
| Q14376 | GALE | GLKAVGEsVQkPLDy | S97 | 0 | 94.46 | 348 | 0 | 0 | 0 | 1 |
| Q14376 | GALE | sVQkPLDyYRVNLTG | Y104 | 0.019 | 98.38 | 348 | 0.008335 | 0.019 | 0.0083 | 1 |
| Q9BQI7 | PSD2 | ADMDGKRtPRGRRGW | T532 | 0.013 | 89.76 | 771 | 0.008879 | 0.013 | 0.0089 | 1 |
| Q86SR1 | GALNT10 | GTGLCADtkHGALGS | T474 | 0.006 | 89.73 | 603 | 0.004098 | 0.006 | 0.0041 | 1 |

| | | | | | | | | | | |
|---|---|---|---|---|---|---|---|---|---|---|
| Q14160 | SCRIB | EIyRysRsLEELLLD | S37 | 0.104 | 92.09 | 1630 | 0.082671 | 0.104 | 0.0827 | 1 |
| Q6P1A2 | LPCAT3 | QALRLIIsIFLGYPF | S47 | 0.022 | 96.78 | 487 | 0.017488 | 0.022 | 0.0175 | 1 |
| Q6P1A2 | LPCAT3 | LIGLAVDyFDGGkDQ | Y159 | 0.119 | 97.79 | 487 | 0.052204 | 0.184 | 0.0675 | 1 |
| Q6P1A2 | LPCAT3 | TNPRFTGtIASFNIN | T333 | 0.054 | 93.71 | 487 | 0.036883 | 0.054 | 0.0369 | 1 |
| Q9P0K1 | ADAM22 | WGQKVTAsDKYCYEK | S565 | 0.123 | 96.38 | 906 | 0.097775 | 0.123 | 0.0978 | 1 |
| Q9Y4K3 | TRAF6 | VIHSPGFyTGKPGYK | Y381 | 0.026 | 95.18 | 522 | 0.011406 | 0.026 | 0.0114 | 1 |
| Q9Y5Q9 | GTF3C3 | ALVCSERyNLAVVWL | Y453 | 0.323 | 87.35 | 886 | 0.141695 | 0.323 | 0.1417 | 1 |
| Q9Y5Q9 | GTF3C3 | LDARIsLsTLQQQLG | S497 | 0.069 | 94.97 | 886 | 0.054849 | 0.069 | 0.0548 | 0.9 |
| O43148 | RNMT | LNEMAKKyNMKLVYK | Y385 | 0.221 | 98.63 | 476 | 0.09695 | 0.221 | 0.0969 | 0.9 |
| O43148 | RNMT | QVRLPLGtLSKSEWE | T455 | 0.016 | 94.65 | 476 | 0.010928 | 0.13 | 0.0888 | 1 |
| P08133 | ANXA6 | HQAIEGDtsGDFLkA | T657 | 0.158 | 96.54 | 673 | 0.107916 | 0.158 | 0.1079 | 1 |
| P36542 | ATP5F1C | kHLLIGVsSDrGLCG | S97 | 0.075 | 96.09 | 298 | 0.059619 | 0.075 | 0.0596 | 1 |
| Q9NTK5 | OLA1 | CDGIFHLtRAFEDDD | T124 | 0.031 | 96.52 | 396 | 0.021173 | 0.031 | 0.0212 | 1 |
| Q9NTK5 | OLA1 | GALVIPFsGALELKL | S264 | 0.006 | 97.86 | 396 | 0.004769 | 0.006 | 0.0048 | 1 |
| Q1KMD3 | HNRNPUL2 | LVNLDtYtsDLHFQV | T252 | 0.216 | 92.2 | 747 | 0.147531 | 0.216 | 0.1475 | 0.875 |
| Q1KMD3 | HNRNPUL2 | VNLDtYtsDLHFQVS | S253 | 0.1 | 94.16 | 747 | 0.079491 | 0.1 | 0.0795 | 0.875 |
| Q1KMD3 | HNRNPUL2 | YGGQPLFsEkFPTLW | S271 | 0.13 | 85.22 | 747 | 0.103339 | 0.13 | 0.1033 | 1 |
| Q1KMD3 | HNRNPUL2 | PVEERVRtAVPPKTI | T442 | 0.087 | 92.25 | 747 | 0.059422 | 0.087 | 0.0594 | 0.9091 |
| Q1KMD3 | HNRNPUL2 | KENPEKRyNVLGAET | Y481 | 0.202 | 95.21 | 747 | 0.088614 | 0.202 | 0.0886 | 1 |
| P22304 | IDS | PFFLAVGyHKPHIPF | Y225 | 0 | 98.93 | 550 | 0 | 0 | 0 | 0.8571 |
| P22304 | IDS | KPHIPFRyPKEFQKL | Y234 | 0 | 98.86 | 550 | 0 | 0 | 0 | 0.9375 |
| P18433 | PTPRA | sPstNRKyPPLPVDk | Y218 | 0.168 | 96.15 | 802 | 0.073699 | 0.227 | 0.0996 | 1 |
| P18433 | PTPRA | PVEGVPDsDYINASF | S293 | 0.115 | 97.7 | 802 | 0.091415 | 0.115 | 0.0914 | 1 |
| P18433 | PTPRA | IQNDKMRtGNLPANM | T552 | 0.196 | 97 | 802 | 0.133871 | 0.196 | 0.1339 | 1 |
| P18433 | PTPRA | RGEENTDyVNASFID | Y588 | 0.016 | 98.36 | 802 | 0.007019 | 0.016 | 0.007 | 1 |
| P18433 | PTPRA | CYKVVQEyIDAFsDy | Y791 | 0.297 | 95.45 | 802 | 0.13029 | 0.297 | 0.1303 | 1 |
| Q92503 | SEC14L1 | SVQGsHVtRWPGFYI | T626 | 0.025 | 88.58 | 715 | 0.017075 | 0.025 | 0.0171 | 1 |
| Q99707 | MTR | IRVCTRAyHLLVKKL | Y520 | 0.216 | 98.36 | 1265 | 0.094756 | 0.216 | 0.0948 | 1 |
| Q2WGJ9 | FER1L6 | GEVRSQNyQIAITIT | Y83 | 0.063 | 85.82 | 1857 | 0.027637 | 0.063 | 0.0276 | 1 |
| P14635 | CCNB1 | TSKHAkIsTLPQLNS | S413 | 0.019 | 97.94 | 433 | 0.015103 | 0.019 | 0.0151 | 1 |
| Q13423 | NNT | NEkRVALsPAGVQNL | S75 | 0.033 | 97.03 | 1086 | 0.026232 | 0.033 | 0.0262 | 1 |

| | | | | | | | | | | |
|---|---|---|---|---|---|---|---|---|---|---|
| Q13423 | NNT | GAkEVLAsDLVVKVR | S122 | 0.082 | 95.88 | 1086 | 0.065183 | 0.082 | 0.0652 | 1 |
| Q13423 | NNT | LYIHKGItHIGytDL | T369 | 0.15 | 97.48 | 1086 | 0.102452 | 0.15 | 0.1025 | 1 |
| Q9H7F0 | ATP13A3 | ELEASVAsPFTSkTP | S909 | 0.015 | 77.08 | 1226 | 0.011924 | 0.115 | 0.0914 | 1 |
| O95436 | SLC34A2 | ISGILLWyPIPFtRL | Y503 | 0.099 | 94.48 | 690 | 0.04343 | 0.099 | 0.0434 | 1 |
| P23416 | GLRA2 | HYNTGKFtCIEVKFH | T242 | 0.041 | 97.22 | 452 | 0.028004 | 0.041 | 0.028 | 1 |
| P14868 | DARS1 | EkyDtDFyILDkyPL | Y389 | 0.016 | 98.86 | 501 | 0.007019 | 0.016 | 0.007 | 1 |
| P14868 | DARS1 | DFyILDkyPLAVRPF | Y394 | 0.053 | 98.63 | 501 | 0.02325 | 0.053 | 0.0232 | 1 |
| P60900 | PSMA6 | DrHItIFsPEGrLyQ | S17 | 0.086 | 94.66 | 246 | 0.068363 | 0.086 | 0.0684 | 1 |
| P60900 | PSMA6 | FsPEGrLyQVEyAFk | Y23 | 0.059 | 96.11 | 246 | 0.025882 | 0.059 | 0.0259 | 0.9231 |
| P60900 | PSMA6 | VTHLFkItENIGCVM | T73 | 0.194 | 98.65 | 246 | 0.132505 | 0.194 | 0.1325 | 0.8333 |
| Q96Q07 | BTBD9 | FPARVCRyIRIVGTH | Y386 | 0.226 | 98.29 | 612 | 0.099143 | 0.226 | 0.0991 | 1 |
| P22612 | PRKACG | YLyLVMEyVPGGEMF | Y123 | 0.215 | 97.59 | 351 | 0.094317 | 0.215 | 0.0943 | 0.9231 |
| Q8NFF5 | FLAD1 | SLGSRENtVRNPALK | T553 | 0.073 | 93.02 | 587 | 0.04986 | 0.073 | 0.0499 | 1 |
| P48378 | RFX2 | RSsLyNHyLRHCQEH | Y224 | 0.008 | 94.35 | 723 | 0.003509 | 0.008 | 0.0035 | 0.9167 |
| P48378 | RFX2 | GIRLKPDsPLNRLQE | S276 | 0.074 | 89.33 | 723 | 0.058824 | 0.074 | 0.0588 | 1 |
| Q96F63 | CCDC97 | rFRAPLLyEQyIGQy | Y185 | 0.043 | 96.67 | 343 | 0.018863 | 0.189 | 0.0829 | 1 |
| P0CG22 | DHRS4L1 | PLPGFsPyNVsKTAL | Y185 | 0.166 | 89.23 | 281 | 0.072822 | 0.27 | 0.1184 | 0.9167 |
| P0CG22 | DHRS4L1 | GFsPyNVsKTALLGL | S188 | 0.021 | 92.94 | 281 | 0.016693 | 0.021 | 0.0167 | 1 |
| Q7Z7M0 | MEGF8 | DCGCSFHsHCRKRGP | S1169 | 0.007 | 82.385 | 2845 | 0.005564 | 0.007 | 0.0056 | 1 |
| Q7Z7M0 | MEGF8 | CERCRPGsFGNATGS | S1198 | 0.037 | 83.625 | 2845 | 0.029412 | 0.037 | 0.0294 | 0.875 |
| Q8WZ64 | ARAP2 | KHQLEDVtAVLKSFL | T1186 | 0.013 | 96.51 | 1704 | 0.008879 | 0.013 | 0.0089 | 1 |
| Q3KQV9 | UAP1L1 | VTYPkGMyRVGLPSR | Y128 | 0.3 | 97.57 | 507 | 0.131606 | 0.3 | 0.1316 | 1 |
| Q8IY22 | CMIP | RLTLPSkstDADLAR | S647 | 0.097 | 96.22 | 773 | 0.077107 | 0.097 | 0.0771 | 1 |
| Q8IY22 | CMIP | DAGLRLLsEHLTMLQ | S707 | 0.093 | 88.96 | 773 | 0.073927 | 0.093 | 0.0739 | 1 |
| Q8WTS6 | SETD7 | VEADEELtVAYGYDH | T332 | 0.14 | 98.79 | 366 | 0.095622 | 0.14 | 0.0956 | 1 |
| Q5FWF4 | ZRANB3 | QFQkDPDtRVAILSI | T390 | 0 | 94.27 | 1079 | 0 | 0 | 0 | 1 |
| Q9NRD9 | DUOX1 | QKVNFVLsSNRGRRT | S694 | 0.004 | 92.4 | 1551 | 0.00318 | 0.004 | 0.0032 | 1 |
| Q8WXG6 | MADD | TLLHNLIsyMLLMKV | S1380 | 0.004 | 94.69 | 1647 | 0.00318 | -0.005 | 0.0178 | 1 |
| P06737 | PYGL | WIRTQQHyyDKCPKr | Y75 | 0.121 | 91.05 | 847 | 0.053081 | 0.216 | 0.0855 | 0.8667 |
| P06737 | PYGL | AHLCIVGsHAVNGVA | S450 | 0.026 | 98.31 | 847 | 0.020668 | 0.026 | 0.0207 | 1 |
| P06737 | PYGL | QENKLkFsQFLETEY | S547 | 0.096 | 96.77 | 847 | 0.076312 | 0.071 | 0.0995 | 1 |

| | | | | | | | | | | |
|---|---|---|---|---|---|---|---|---|---|---|
| P06737 | PYGL | MIIKLITsVADVVNN | S626 | 0.095 | 98.39 | 847 | 0.075517 | 0.095 | 0.0755 | 0.8 |
| P06737 | PYGL | QIDNGFFsPkQPDLF | S752 | 0.109 | 96.54 | 847 | 0.086646 | 0.151 | 0.1439 | 1 |
| P06737 | PYGL | RFKVFADyEAYVkCQ | Y778 | 0.067 | 98.75 | 847 | 0.029392 | 0.102 | 0.0447 | 1 |
| Q8N135 | LGI4 | CFVVADAsKAGSTTL | S337 | 0.176 | 96.38 | 537 | 0.139905 | 0.176 | 0.1399 | 1 |
| Q8NFW8 | CMAS | FQSVWVStDHDEIEN | T92 | 0 | 98.06 | 434 | 0 | 0 | 0 | 1 |
| Q8NFW8 | CMAS | IREEGyDsVFSVVRR | S168 | 0.003 | 98.57 | 434 | 0.002385 | 0.003 | 0.0024 | 1 |
| Q9NYQ8 | FAT2 | PLFSPPSyRVTISED | Y153 | 0.304 | 93.16 | 4349 | 0.13336 | 0.341 | 0.1496 | 1 |
| Q12923 | PTPN13 | RyKNILPyDATRVPL | Y2249 | 0.026 | 92.93 | 2485 | 0.011406 | 0.026 | 0.0114 | 0.9444 |
| Q12923 | PTPN13 | DDLLTFIsyMRHIHR | S2393 | 0.097 | 93.47 | 2485 | 0.077107 | 0.097 | 0.0771 | 1 |
| Q12923 | PTPN13 | DLLTFIsyMRHIHRS | Y2394 | 0.229 | 94.05 | 2485 | 0.100459 | 0.229 | 0.1005 | 1 |
| O00743 | PPP6C | AFCDLVWsDPEDVDT | S197 | 0 | 98.62 | 305 | 0 | 0 | 0 | 1 |
| O00743 | PPP6C | DVDTWAIsPRGAGWL | S208 | 0.099 | 97.16 | 305 | 0.078697 | 0.099 | 0.0787 | 1 |
| Q9NPH5 | NOX4 | WQENRPDyVNIQLYL | Y491 | 0.097 | 96.19 | 578 | 0.042553 | 0.097 | 0.0426 | 1 |
| Q8NGK0 | OR51G2 | VSATFLLsGIPGLER | S20 | 0.104 | 82.56 | 314 | 0.082671 | 0.104 | 0.0827 | 1 |
| Q9UJM3 | ERRFI1 | IENGKKVsstHyyLL | S390 | 0.187 | 96.71 | 462 | 0.148649 | 0.187 | 0.1486 | 1 |
| O60241 | ADGRB2 | TSRVMtVtVRPPtQP | T845 | 0.186 | 90.78 | 1585 | 0.127041 | 0.186 | 0.127 | 1 |
| O75674 | TOM1L1 | GksHRDPyATSVGHL | Y11 | 0.233 | 96.18 | 476 | 0.102214 | 0.233 | 0.1022 | 1 |
| Q01085 | TIAL1 | HVFVGDLsPEITTED | S105 | 0.029 | 92.38 | 375 | 0.023053 | 0.029 | 0.0231 | 1 |
| P06733 | ENO1 | FDsrGNPtVEVDLFt | T19 | 0.022 | 98.86 | 434 | 0.015026 | 0.044 | 0.0301 | 1 |
| P06733 | ENO1 | tVEVDLFtskGLFRA | T26 | 0.103 | 98.12 | 434 | 0.07035 | 0.103 | 0.0703 | 1 |
| P06733 | ENO1 | LFRAAVPsGAstGIy | S37 | 0.028 | 95.66 | 434 | 0.022258 | 0.028 | 0.0223 | 1 |
| P06733 | ENO1 | ANAILGVsLAVCkAG | S115 | 0 | 98.91 | 434 | 0 | 0 | 0 | 1 |
| P06733 | ENO1 | VEkGVPLyRHIADLA | Y131 | 0.162 | 98.79 | 434 | 0.071067 | 0.162 | 0.0711 | 1 |
| P06733 | ENO1 | MrIGAEVyHNLkNVI | Y189 | 0.13 | 98.91 | 434 | 0.057029 | 0.259 | 0.1136 | 1 |
| P06733 | ENO1 | EFFRsGkyDLDFksP | Y257 | 0.001 | 98.61 | 434 | 0.000439 | 0.001 | 0.0004 | 1 |
| P06733 | ENO1 | PDQLADLyksFIkDy | Y280 | 0.001 | 98.63 | 434 | 0.000439 | 0.001 | 0.0004 | 0.9333 |
| P06733 | ENO1 | AkAVNEksCNCLLLk | S336 | 0 | 98.61 | 434 | 0 | 0 | 0 | 1 |
| Q9Y490 | TLN1 | LVQGVRGsQAQPDsP | S973 | 0 | 81.99 | 2541 | 0 | 0.151 | 0.12 | 1 |
| P08621 | SNRNP70 | RGyAFIEyEHERDMH | Y151 | 0.007 | 97.38 | 437 | 0.003071 | 0.007 | 0.0031 | 1 |
| Q9Y6N8 | CDH10 | KGDGSLKyILsGDGA | Y91 | 0.104 | 92.64 | 788 | 0.045623 | 0.104 | 0.0456 | 1 |
| Q9Y6N8 | CDH10 | DYESRRLyTLKVEAE | Y344 | 0.165 | 96.54 | 788 | 0.072383 | 0.226 | 0.0991 | 1 |

| | | | | | | | | | | |
|---|---|---|---|---|---|---|---|---|---|---|
| Q8NET8 | TRPV3 | EKRLRSLsRKFtDWA | S374 | 0.009 | 87.62 | 790 | 0.007154 | 0.009 | 0.0072 | 1 |
| P14649 | MYL6B | VGDGKILysQCGDVM | Y86 | 0.329 | 95.95 | 208 | 0.144328 | 0.329 | 0.1443 | 1 |
| Q5T440 | IBA57 | RMGWRLLtQDEGPAL | T200 | 0 | 96.37 | 356 | 0 | 0 | 0 | 1 |
| Q9NQT8 | KIF13B | CsGLFERtQKEENEE | T134 | 0.069 | 93.1 | 1826 | 0.047128 | 0.069 | 0.0471 | 1 |
| Q9NQT8 | KIF13B | SLGGNSKtAMVATVs | T324 | 0.08 | 94.78 | 1826 | 0.054641 | 0.08 | 0.0546 | 0.9 |
| Q9NQT8 | KIF13B | tAMVATVsPAADNYD | S331 | 0.002 | 92.46 | 1826 | 0.00159 | 0.002 | 0.0016 | 1 |
| Q02094 | RHAG | VLtAFAFssLVEHRG | S258 | 0 | 98.78 | 409 | 0 | 0 | 0 | 1 |
| A8K2U0 | A2ML1 | QGCVYVQtVLRYNIL | T1312 | 0.051 | 90.96 | 1454 | 0.034834 | 0.092 | 0.0628 | 1 |
| P43405 | SYK | GKISREEsEQIVLIG | S178 | 0 | 96.13 | 635 | 0 | 0 | 0 | 1 |
| P43405 | SYK | LWQLVEHySYKADGL | Y244 | 0.004 | 93.7 | 635 | 0.001755 | 0.004 | 0.0018 | 1 |
| Q9Y6R4 | MAP3K4 | LCQEQTSsQPVIAKA | S979 | 0.068 | 91.16 | 1608 | 0.054054 | 0.068 | 0.0541 | 1 |
| P26639 | TARS1 | LQRIyGIsFPDPkML | S301 | 0 | 97.95 | 723 | 0 | 0 | 0 | 1 |
| P26639 | TARS1 | IGRDQELyFFHELsP | Y333 | 0.143 | 98.14 | 723 | 0.062732 | 0.143 | 0.0627 | 1 |
| P26639 | TARS1 | rGFQEVVtPNIFNsR | T374 | 0.175 | 97.66 | 723 | 0.119527 | 0.175 | 0.1195 | 1 |
| P26639 | TARS1 | VtPNIFNsRLWMTSG | S380 | 0.092 | 96.72 | 723 | 0.073132 | 0.092 | 0.0731 | 1 |
| P26639 | TARS1 | PIRFNLtyVsHDGDD | Y575 | 0.132 | 96.11 | 723 | 0.057907 | 0.132 | 0.0579 | 1 |
| P26639 | TARS1 | RFNLtyVsHDGDDkk | S577 | 0.061 | 89.75 | 723 | 0.04849 | 0.061 | 0.0485 | 1 |
| P26639 | TARS1 | QQLkEFRskQAEEEF | S716 | 0.084 | 93.58 | 723 | 0.066773 | 0.084 | 0.0668 | 1 |
| O14746 | TERT | RPGLLGAsVLGLDDI | S679 | 0.161 | 94.66 | 1132 | 0.127981 | 0.161 | 0.128 | 1 |
| Q96EM0 | L3HYPDH | sPtGSGVtARIALQY | T278 | 0 | 98.79 | 354 | 0 | 0 | 0 | 0.9286 |
| Q9Y5H1 | PCDHGA2 | GYYAQVVyFLEKSPG | Y279 | 0.181 | 93.9 | 932 | 0.079402 | 0.181 | 0.0794 | 1 |
| P05771 | PRKCB | PNGLSDPyVkLkLIP | Y195 | 0.106 | 98.62 | 671 | 0.046501 | 0.178 | 0.0781 | 1 |
| P46060 | RANGAP1 | LLRLGPksSVLIAQQ | S453 | 0.102 | 72.3 | 587 | 0.081081 | 0.102 | 0.0811 | 1 |
| P46060 | RANGAP1 | SVLIAQQtDtsDPEk | T461 | 0.11 | 90.34 | 587 | 0.075131 | 0.11 | 0.0751 | 1 |
| P46060 | RANGAP1 | LIAQQtDtsDPEkVV | T463 | 0.157 | 92.27 | 587 | 0.107233 | 0.157 | 0.1072 | 1 |
| Q9H8M5 | CNNM2 | EAsAFsyyGVMALTA | Y713 | 0.193 | 95.37 | 875 | 0.084666 | 0.193 | 0.0847 | 1 |
| Q5T9C9 | PIP5KL1 | HLQRHPHsLLARLLG | S180 | 0 | 96.41 | 394 | 0 | 0 | 0 | 0.8333 |
| Q16555 | DPYSL2 | PGGIDVHtRFQMPDQ | T74 | 0 | 98.7 | 572 | 0 | 0 | 0 | 0.8 |
| Q16555 | DPYSL2 | MPDQGMTsADDFFQG | S85 | 0.064 | 96.02 | 572 | 0.050875 | 0.125 | 0.0997 | 1 |
| Q14166 | TTLL12 | FkDYRkLsQERPGVL | S348 | 0.073 | 96.43 | 644 | 0.058029 | 0.073 | 0.058 | 1 |
| Q14166 | TTLL12 | tELPQFVsyFQQRER | S402 | 0 | 97.93 | 644 | 0 | 0 | 0 | 1 |

| | | | | | | | | | | |
|---|---|---|---|---|---|---|---|---|---|---|
| Q14166 | TTLL12 | HVTkSLHsIIRHRES | S437 | 0.03 | 96.37 | 644 | 0.023847 | 0.03 | 0.0238 | 0.8333 |
| Q14166 | TTLL12 | DDyEkHFtVMNyDPD | T513 | 0.145 | 95.17 | 644 | 0.099037 | 0.145 | 0.099 | 1 |
| Q7Z6Z7 | HUWE1 | NPTTVLEsPHSLPAk | S1218 | 0.081 | 89.83 | 4374 | 0.064388 | 0.081 | 0.0644 | 1 |
| P49591 | SARS1 | ALANLkVsQIKKVRL | S101 | 0.117 | 94.76 | 514 | 0.093005 | 0.117 | 0.093 | 1 |
| O15021 | MAST4 | KLTDFGLsKVGLMSM | S715 | 0.077 | 76.9 | 2623 | 0.061208 | 0.077 | 0.0612 | 0.8889 |
| Q96QD8 | SLC38A2 | RRRMMNVsKISFFAM | S328 | 0.002 | 97.88 | 506 | 0.00159 | 0.002 | 0.0016 | 0.8889 |
| Q9UN66 | PCDHB8 | DRESRAEyNVTITVT | Y418 | 0.17 | 92.39 | 801 | 0.074577 | 0.17 | 0.0746 | 1 |
| Q9Y4D1 | DAAM1 | HYQQRLQsLYFKKKF | S776 | 0.031 | 97.64 | 1078 | 0.024642 | 0.031 | 0.0246 | 1 |
| Q9Y4D1 | DAAM1 | PKVEAIRsGsEEVFR | S799 | 0.121 | 97.84 | 1078 | 0.096185 | 0.121 | 0.0962 | 1 |
| Q9Y4D1 | DAAM1 | VEAIRsGsEEVFRSG | S801 | 0.018 | 97.78 | 1078 | 0.014308 | 0.018 | 0.0143 | 1 |
| O75487 | GPC4 | QYHFTDEyLECVSKY | Y186 | 0.287 | 92.64 | 556 | 0.125903 | 0.287 | 0.1259 | 1 |
| Q13813 | SPTAN1 | RKAkLGDsHDLQRFL | S1338 | 0.07 | 86.56 | 2472 | 0.055644 | 0.076 | 0.0676 | 0.8 |
| Q07866 | KLC1 | YQRALEIyQtkLGPD | Y369 | 0.123 | 97.62 | 573 | 0.053958 | 0.123 | 0.054 | 1 |
| P14923 | JUP | AMIVNQLskKEASRR | S170 | 0 | 96.77 | 745 | 0 | 0 | 0 | 1 |
| P14923 | JUP | SRRALMGsPQLVAAV | S182 | 0.166 | 96.11 | 745 | 0.131956 | 0.166 | 0.132 | 1 |
| P14923 | JUP | SkNkTLVtQNSGVEA | T430 | 0.157 | 98.6 | 745 | 0.107233 | 0.157 | 0.1072 | 1 |
| P14923 | JUP | VCALRHLtSRHPEAE | T463 | 0.008 | 98.74 | 745 | 0.005464 | 0.008 | 0.0055 | 1 |
| Q9H7D0 | DOCK5 | KDsFQIAtLICSTKL | T624 | 0.117 | 94.86 | 1870 | 0.079913 | 0.117 | 0.0799 | 0.9167 |
| Q8N960 | CEP120 | NKYTKFKsEIQIsIA | S124 | 0.002 | 94.9 | 986 | 0.00159 | 0.002 | 0.0016 | 1 |
| P15311 | EZR | kDNAMLEyLKIAQDL | Y191 | 0.056 | 98.74 | 586 | 0.024566 | 0.056 | 0.0246 | 1 |
| P15311 | EZR | IAQDLEMyGINyFEI | Y201 | 0.131 | 98.39 | 586 | 0.057468 | 0.215 | 0.0943 | 1 |
| P15311 | EZR | CMGNHELyMRRRkPD | Y291 | 0.02 | 93.86 | 586 | 0.008774 | 0.189 | 0.0829 | 1 |
| Q12929 | EPS8 | VINItyDstPEDVKT | S725 | 0.121 | 88.61 | 822 | 0.096185 | 0.121 | 0.0962 | 1 |
| Q9P0M6 | MACROH2A[illegible] | LGQKLsLtQSDISHI | T200 | 0.002 | 93.13 | 372 | 0.001366 | 0.002 | 0.0014 | 0.9091 |
| P09622 | DLD | VIGVELGsVWQRLGA | S230 | 0.006 | 98.84 | 509 | 0.004769 | 0.006 | 0.0048 | 0.9 |
| P09622 | DLD | GFkFkLNtkVTGATK | T276 | 0.173 | 98 | 509 | 0.118161 | 0.173 | 0.1182 | 1 |
| Q9Y6X8 | ZHX2 | LDRLRVEtKLsRREI | T566 | 0.037 | 92.76 | 837 | 0.025272 | 0.037 | 0.0253 | 1 |
| P48449 | LSS | VFGTALNyVSLRILG | Y158 | 0.011 | 98.94 | 732 | 0.004826 | 0.011 | 0.0048 | 0.8 |
| P48449 | LSS | DRFTkSIsIGPISkT | S334 | 0.001 | 98.76 | 732 | 0.000795 | 0.001 | 0.0008 | 0.8 |
| P48449 | LSS | HVSRIPDyLWMGLDG | Y368 | 0.012 | 98.89 | 732 | 0.005264 | 0.012 | 0.0053 | 0.8824 |
| Q9Y5E9 | PCDHB14 | AGTESAHySVAEETE | Y33 | 0.229 | 90.59 | 798 | 0.100459 | 0.229 | 0.1005 | 1 |

| | | | | | | | | | | |
|---|---|---|---|---|---|---|---|---|---|---|
| O75592 | MYCBP2 | EDGSIFFtGsAsKGE | T606 | 0.001 | 94.065 | 4678 | 0.000683 | 0.001 | 0.0007 | 1 |
| P49642 | PRIM1 | IHPFIRksINIIKkY | S215 | 0.002 | 98.7 | 420 | 0.00159 | 0.002 | 0.0016 | 0.8889 |
| P49642 | PRIM1 | PRLDINVskGINHLL | S310 | 0.019 | 94 | 420 | 0.015103 | 0.019 | 0.0151 | 0.8889 |
| P49642 | PRIM1 | HPkTGRIsVPIDLQK | S331 | 0.017 | 98.82 | 420 | 0.013514 | 0.017 | 0.0135 | 1 |
| Q9UQC9 | CLCA2 | MDRNsLQsAVSNIAQ | S876 | 0.087 | 96.89 | 943 | 0.069158 | 0.087 | 0.0692 | 1 |
| Q9BTP7 | FAAP24 | VVEKTRMsEQYFPAL | S97 | 0 | 95.83 | 215 | 0 | 0 | 0 | 1 |
| P16298 | PPP3CB | RYLFLGDyVDRGYFS | Y128 | 0 | 98.9 | 524 | 0 | 0 | 0 | 1 |
| Q9BWG4 | SSBP4 | EKLALyVyEYLLHIG | Y26 | 0.239 | 94.9 | 385 | 0.104846 | 0.239 | 0.1048 | 0.8 |
| P08581 | MET | FGCVyHGtLLDNDGk | T1096 | 0.15 | 93.75 | 1390 | 0.102452 | 0.15 | 0.1025 | 1 |
| Q6UX04 | CWC27 | DELNNKHtIFGKVTG | T128 | 0.003 | 93.38 | 472 | 0.002049 | 0.003 | 0.002 | 1 |
| Q9UHW5 | GPN3 | LVMGPAGsGkSTyCA | S14 | 0.015 | 95.17 | 284 | 0.011924 | 0.015 | 0.0119 | 1 |
| Q8NE18 | NSUN7 | VSHMSILtNNNtsKV | T314 | 0.003 | 92.12 | 718 | 0.002049 | 0.003 | 0.002 | 1 |
| Q8NE18 | NSUN7 | ILtNNNtsKVFVCGV | S319 | 0.017 | 89.27 | 718 | 0.013514 | 0.017 | 0.0135 | 1 |
| Q14191 | WRN | YCPSRkMtQQVTGEL | T777 | 0 | 92.45 | 1432 | 0 | 0 | 0 | 1 |
| Q14807 | KIF22 | LMDLLQLtREEGAEG | T158 | 0.134 | 91.47 | 665 | 0.091524 | 0.199 | 0.1359 | 1 |
| Q9UK32 | RPS6KA6 | EIEILMRyGQHPNII | Y474 | 0.201 | 85.96 | 745 | 0.088176 | 0.201 | 0.0882 | 1 |
| Q6VAB6 | KSR2 | LKSKNVFyDNGKVVI | Y794 | 0.337 | 96.39 | 950 | 0.147837 | 0.337 | 0.1478 | 1 |
| Q6VAB6 | KSR2 | DNGKVVItDFGLFsI | T802 | 0.135 | 93.07 | 950 | 0.092207 | 0.135 | 0.0922 | 1 |
| Q8N7B1 | HORMAD2 | NEDIKKAsVLLIRKL | S157 | 0 | 93.2 | 307 | 0 | 0 | 0 | 1 |
| Q9UER7 | DAXX | VIEQRIPyRGTRyPE | Y259 | 0.063 | 98.15 | 740 | 0.027637 | 0.063 | 0.0276 | 1 |
| P55289 | CDH12 | DREVKEQyQVLIQAK | Y233 | 0.171 | 95.64 | 794 | 0.075015 | 0.171 | 0.075 | 1 |
| Q9UHG0 | DCDC2 | VVKsVLVyRNGDPFY | Y22 | 0.037 | 93.26 | 476 | 0.016231 | 0.082 | 0.036 | 0.9286 |
| Q9UHG0 | DCDC2 | EAFKKLNyLDIGEIK | Y99 | 0.014 | 87.47 | 476 | 0.006142 | 0.014 | 0.0061 | 1 |
| Q9NWS0 | PIH1D1 | FYRRMQNsDFLRELV | S135 | 0.079 | 91.3 | 290 | 0.062798 | 0.079 | 0.0628 | 0.8 |
| Q03154 | ACY1 | kDSEGYIyARGAQDM | Y107 | 0.306 | 98.66 | 408 | 0.134238 | 0.306 | 0.1342 | 1 |
| Q9BUQ8 | DDX23 | EESDQGPyAIILAPt | Y470 | 0.158 | 94.99 | 820 | 0.069312 | 0.158 | 0.0693 | 1 |
| Q9BUQ8 | DDX23 | yAIILAPtRELAQQI | T477 | 0.138 | 88.2 | 820 | 0.094256 | 0.154 | 0.1052 | 1 |
| Q9BUQ8 | DDX23 | GKHKYRQtVMFTATM | T602 | 0.011 | 97.27 | 820 | 0.007513 | 0.011 | 0.0075 | 1 |
| Q9BUQ8 | DDX23 | VERLARSyLRRPAVV | Y620 | 0.056 | 93.94 | 820 | 0.024566 | 0.056 | 0.0246 | 1 |
| Q96S59 | RANBP9 | QNNLRVHyKGHGKTP | Y189 | 0.164 | 98.82 | 729 | 0.071944 | 0.164 | 0.0719 | 1 |
| Q12805 | EFEMP1 | RSDRSVPsDIFQIQA | S408 | 0.169 | 94.02 | 493 | 0.134341 | 0.169 | 0.1343 | 1 |

| | | | | | | | | | | |
|---|---|---|---|---|---|---|---|---|---|---|
| A7KAX9 | ARHGAP32 | LADYCSItNMHAKNL | T514 | 0 | 94.03 | 2087 | 0 | 0 | 0 | 1 |
| P17301 | ITGA2 | KEKNPLMyLTGVQTD | Y1005 | 0.021 | 92.76 | 1181 | 0.009212 | 0.021 | 0.0092 | 1 |
| P17301 | ITGA2 | DVHMKGEyFVNVtTR | Y1071 | 0.091 | 90.29 | 1181 | 0.03992 | 0.266 | 0.0828 | 1 |
| Q15306 | IRF4 | QLDISDPykVyRIVP | Y122 | 0.068 | 94.88 | 451 | 0.029831 | 0.318 | 0.1395 | 0.9 |
| Q9Y2E4 | DIP2C | GGIHLsEtKQLFLEG | T898 | 0 | 92.85 | 1556 | 0 | 0 | 0 | 1 |
| Q9HAS0 | C17orf75 | KLkAIQDtNNLkRFI | T330 | 0.106 | 89.33 | 396 | 0.072399 | 0.106 | 0.0724 | 0.8889 |
| O60645 | EXOC3 | FGSTQGLsDELAKQL | S184 | 0.053 | 92.78 | 745 | 0.04213 | 0.053 | 0.0421 | 1 |
| P43628 | KIR2DL3 | tVILQCWsDVRFQHF | S51 | 0.041 | 95.52 | 341 | 0.032592 | 0.041 | 0.0326 | 1 |
| Q9NRD8 | DUOX2 | QRRVPANyADGVyQA | Y65 | 0.135 | 95.3 | 1548 | 0.059223 | 0.23 | 0.1009 | 1 |
| Q9NRD8 | DUOX2 | LPNPRRLsNAATRGI | S85 | 0 | 95.96 | 1548 | 0 | 0 | 0 | 1 |
| Q01780 | EXOSC10 | DDTHYLLyIyDKMRL | Y446 | 0.248 | 98.16 | 885 | 0.108794 | 0.248 | 0.1088 | 1 |
| Q01780 | EXOSC10 | THYLLyIyDKMRLEM | Y448 | 0.021 | 98 | 885 | 0.009212 | 0.021 | 0.0092 | 1 |
| Q01780 | EXOSC10 | PIFTDEsyLELYRKQ | Y491 | 0.284 | 84.13 | 885 | 0.124587 | 0.284 | 0.1246 | 1 |
| O15020 | SPTBN2 | IETDIVAySGRVQAV | Y477 | 0.193 | 82.07 | 2390 | 0.084666 | 0.193 | 0.0847 | 1 |
| Q6PJF5 | RHBDF2 | IAIIFILsGItGNLA | S700 | 0.001 | 97.98 | 856 | 0.000795 | 0.001 | 0.0008 | 1 |
| Q6PJF5 | RHBDF2 | IFILsGItGNLAsAI | T703 | 0.001 | 97.48 | 856 | 0.000683 | 0.001 | 0.0007 | 1 |
| Q6PJF5 | RHBDF2 | GItGNLAsAIFLPYR | S708 | 0 | 92.15 | 856 | 0 | 0 | 0 | 1 |
| O14744 | PRMT5 | VEKIRRNsEAAMLQE | S103 | 0.019 | 96.7 | 637 | 0.015103 | 0.019 | 0.0151 | 1 |
| O14744 | PRMT5 | SEKEFCSyLQyLEyL | Y280 | 0 | 96.72 | 637 | 0 | 0 | 0 | 0.9167 |
| O14744 | PRMT5 | EFCSyLQyLEyLSQN | Y283 | 0.207 | 95.77 | 637 | 0.090808 | 0.207 | 0.0908 | 1 |
| O14744 | PRMT5 | DNLESQTyEVFEkDP | Y324 | 0.293 | 91.64 | 637 | 0.128535 | 0.293 | 0.1285 | 1 |
| O00757 | FBP2 | IKKKGKIySLNEGyA | Y210 | 0.116 | 98.75 | 339 | 0.050888 | 0.116 | 0.0509 | 1 |
| O00757 | FBP2 | VyGGIFLyPANQKSP | Y265 | 0.327 | 98.19 | 339 | 0.14345 | 0.327 | 0.1434 | 1 |
| Q00325 | SLC25A3 | MQVDPQkykGIFNGF | Y100 | 0.165 | 86.4 | 362 | 0.072383 | 0.165 | 0.0724 | 0.9231 |
| Q00325 | SLC25A3 | IFNGFsVtLkEDGVr | T110 | 0.002 | 89.84 | 362 | 0.001366 | 0.002 | 0.0014 | 1 |
| Q00325 | SLC25A3 | ERTVEALykFVVPkP | Y246 | 0.261 | 87.5 | 362 | 0.114497 | 0.261 | 0.1145 | 0.9167 |
| P21291 | CSRP1 | ADkDGEIyCkGCyAk | Y166 | 0.16 | 88.85 | 193 | 0.07019 | 0.2 | 0.0877 | 1 |
| Q15738 | NSDHL | RILtGLNyEAPKYHI | Y288 | 0.187 | 96.37 | 373 | 0.082034 | 0.187 | 0.082 | 1 |
| P51812 | RPS6KA3 | KLTDFGLskESIDHE | S215 | 0.002 | 65.41 | 740 | 0.00159 | 0.002 | 0.0016 | 1 |
| Q58WW2 | DCAF6 | LAVGCSDssVRIYDR | S214 | 0 | 93.23 | 860 | 0 | 0 | 0 | 0.9091 |
| Q58WW2 | DCAF6 | AVGCSDssVRIYDRR | S215 | 0 | 94.48 | 860 | 0 | 0 | 0 | 1 |

| | | | | | | | | | | |
|---|---|---|---|---|---|---|---|---|---|---|
| P49683 | PRLHR | PLLVILLsYVRVSVK | S244 | 0.138 | 86.5 | 370 | 0.109698 | 0.138 | 0.1097 | 1 |
| Q8NEM2 | SHCBP1 | FYERFRAyQDyILAD | Y85 | 0.018 | 94.88 | 672 | 0.007896 | 0.018 | 0.0079 | 0.8571 |
| Q8NEM2 | SHCBP1 | EEEDEyDyFVRCVEP | Y219 | 0.009 | 90.58 | 672 | 0.003948 | 0.009 | 0.0039 | 1 |
| Q6AI08 | HEATR6 | SIQALIstVLTEAAM | T963 | 0.087 | 95.95 | 1181 | 0.059422 | 0.087 | 0.0594 | 1 |
| P80365 | HSD11B2 | ATVLELNsPGAIELR | S118 | 0.187 | 94.78 | 405 | 0.148649 | 0.187 | 0.1486 | 1 |
| P80365 | HSD11B2 | PQELLQAyGKDYIEH | Y295 | 0.328 | 94.62 | 405 | 0.143889 | 0.328 | 0.1439 | 1 |
| Q6UY18 | LINGO4 | LLQELDLsyNQLSTL | S92 | 0.051 | 98.03 | 593 | 0.040541 | 0.051 | 0.0405 | 1 |
| Q96JG6 | VPS50 | IMsQHNIyVDALLKE | Y815 | 0.08 | 93.1 | 964 | 0.035095 | 0.08 | 0.0351 | 0.8462 |
| Q96JG6 | VPS50 | LMKLEKLtDIRPIPD | T892 | 0.075 | 85.03 | 964 | 0.051226 | 0.075 | 0.0512 | 1 |
| P39687 | ANP32A | kELVLDNsRsNEGKL | S27 | 0.055 | 96.18 | 249 | 0.04372 | 0.055 | 0.0437 | 1 |
| P39687 | ANP32A | NLTHLNLsGNkIkDL | S96 | 0.185 | 98.08 | 249 | 0.147059 | 0.185 | 0.1471 | 1 |
| P52701 | MSH6 | ERVDTCHtPFGkRLL | T767 | 0.009 | 96.82 | 1360 | 0.006147 | 0.009 | 0.0061 | 1 |
| Q9UL62 | TRPC5 | NIYKALAsPsLIALS | S205 | 0.025 | 94.63 | 973 | 0.019873 | 0.025 | 0.0199 | 0.9 |
| Q9UL62 | TRPC5 | YKALAsPsLIALSSE | S207 | 0.006 | 93.7 | 973 | 0.004769 | 0.14 | 0.1113 | 1 |
| Q9UL62 | TRPC5 | GWELKELsKVENEFK | S231 | 0.003 | 93.28 | 973 | 0.002385 | 0.003 | 0.0024 | 1 |
| Q8TB73 | NDNF | SLGRTTVtLAWKPSP | T193 | 0.192 | 97.51 | 568 | 0.131139 | 0.192 | 0.1311 | 0.8571 |
| P22570 | FDXR | HLERTDItkAALGVL | T208 | 0.077 | 98.09 | 491 | 0.052592 | 0.077 | 0.0526 | 1 |
| P22570 | FDXR | PRPRKRLtELLLRTA | T277 | 0.012 | 95.93 | 491 | 0.008196 | 0.012 | 0.0082 | 1 |
| P22570 | FDXR | sPQQVLPsPDGRRAA | S317 | 0.144 | 97.92 | 491 | 0.114468 | 0.144 | 0.1145 | 1 |
| Q9UIF3 | TEKT2 | EVEVIEAtKKALQQK | T147 | 0.023 | 97.11 | 430 | 0.015709 | 0.023 | 0.0157 | 1 |
| Q06124 | PTPN11 | PNEPVsDyINANIIM | Y304 | 0.019 | 98.65 | 593 | 0.008335 | 0.098 | 0.043 | 1 |
| P10155 | RO60 | GLAIVTKyItKGWkE | Y201 | 0.276 | 97.25 | 538 | 0.121077 | 0.276 | 0.1211 | 1 |
| P10155 | RO60 | EKALsVEtEkLLkYL | T222 | 0.075 | 92.47 | 538 | 0.051226 | 0.075 | 0.0512 | 1 |
| P10155 | RO60 | LGSILNAstVAAAMC | S393 | 0.115 | 91.14 | 538 | 0.091415 | 0.115 | 0.0914 | 1 |
| P10155 | RO60 | GSILNAstVAAAMCM | T394 | 0.137 | 89.1 | 538 | 0.093573 | 0.137 | 0.0936 | 0.8571 |
| Q86XI2 | NCAPG2 | DSIERFLyESSSRTL | Y1130 | 0.191 | 88.41 | 1143 | 0.083789 | 0.191 | 0.0838 | 0.8182 |
| Q9H4B4 | PLK3 | ANRKtVHyNPTsTKH | Y506 | 0.304 | 94.47 | 646 | 0.13336 | 0.304 | 0.1334 | 1 |
| O00754 | MAN2B1 | DPTRRFIyVEIAFFS | Y118 | 0.01 | 98.79 | 1011 | 0.004387 | 0.01 | 0.0044 | 1 |
| O00754 | MAN2B1 | TWGKEVIsRFDTPLE | S731 | 0.001 | 98.82 | 1011 | 0.000795 | 0.001 | 0.0008 | 1 |
| O00754 | MAN2B1 | TITRLQEttLVANQL | T954 | 0.098 | 97.09 | 1011 | 0.066935 | 0.098 | 0.0669 | 1 |
| O15269 | SPTLC1 | KTEEAIIySYGFATI | Y164 | 0.04 | 97.27 | 473 | 0.017547 | 0.04 | 0.0175 | 1 |

| | | | | | | | | | | |
|---|---|---|---|---|---|---|---|---|---|---|
| O15269 | SPTLC1 | EKCLPPPsIRVVVTV | S443 | 0.009 | 98.48 | 473 | 0.007154 | 0.009 | 0.0072 | 1 |
| Q9NWM0 | SMOX | KLAMIQQyLKVESCE | Y201 | 0.202 | 95.77 | 555 | 0.088614 | 0.247 | 0.1084 | 1 |
| Q53TQ3 | INO80D | AsKEPECtGQLIQEL | T417 | 0.15 | 86.67 | 1027 | 0.102452 | 0.15 | 0.1025 | 1 |
| P49770 | EIF2B2 | MTIGFSRtVEAFLkE | T173 | 0.172 | 94.4 | 351 | 0.117478 | 0.172 | 0.1175 | 0.8889 |
| P49770 | EIF2B2 | VNkVIIGtktILANG | T236 | 0.092 | 93.06 | 351 | 0.062837 | 0.092 | 0.0628 | 1 |
| P49770 | EIF2B2 | kVIIGtktILANGAL | T238 | 0.006 | 91.79 | 351 | 0.004098 | 0.18 | 0.1229 | 1 |
| Q86WB0 | ZC3HC1 | AGkPFELsPLVCAKY | S98 | 0.035 | 95.49 | 502 | 0.027822 | 0.18 | 0.1431 | 1 |
| Q86WB0 | ZC3HC1 | CFWPDsPsPDRFGML | S163 | 0.028 | 96.14 | 502 | 0.022258 | 0.028 | 0.0223 | 1 |
| Q96Q15 | SMG1 | KLLyLCAtyKALEtV | T256 | 0.005 | 94.63 | 3661 | 0.003415 | 0.005 | 0.0034 | 1 |
| P11245 | NAT2 | AYFERIGyKNSRNKL | Y12 | 0.135 | 98.21 | 290 | 0.059223 | 0.135 | 0.0592 | 1 |
| A6NGE7 | URAD | LAELNAQyRARFGFP | Y111 | 0 | 98.68 | 173 | 0 | 0 | 0 | 1 |
| O43707 | ACTN4 | HRPELIEyDkLRkDD | Y212 | 0.247 | 91.8 | 911 | 0.108355 | 0.247 | 0.1084 | 0.8 |
| O14958 | CASQ2 | GFFKSEDsEyyKAFE | S176 | 0.147 | 97.6 | 399 | 0.116852 | 0.147 | 0.1169 | 1 |
| O14958 | CASQ2 | KSEDsEyyKAFEEAA | Y179 | 0.132 | 98.61 | 399 | 0.057907 | 0.132 | 0.0579 | 1 |
| Q7Z6M2 | FBXO33 | LEIVTPTsLSSLSNA | S268 | 0.006 | 95.08 | 555 | 0.004769 | 0.006 | 0.0048 | 1 |
| P10253 | GAA | MDSRRDFtFNKDGFR | T415 | 0.218 | 98.8 | 952 | 0.148897 | 0.218 | 0.1489 | 1 |
| P10253 | GAA | PFMRNHNsLLsLPQE | S676 | 0.158 | 97.44 | 952 | 0.125597 | 0.158 | 0.1256 | 1 |
| P10253 | GAA | RNHNsLLsLPQEPYS | S679 | 0.119 | 98.32 | 952 | 0.094595 | 0.119 | 0.0946 | 1 |
| Q9BX63 | BRIP1 | AYSGVPMtILSsRDH | T274 | 0 | 95.4 | 1249 | 0 | 0 | 0 | 1 |
| Q9H1D9 | POLR3F | AASkKKVyMLYNLQP | Y150 | 0.296 | 92.98 | 316 | 0.129851 | 0.296 | 0.1299 | 1 |
| O00763 | ACACB | YyLDISLyKEVTDSR | Y1664 | 0.021 | 89.94 | 2458 | 0.009212 | 0.021 | 0.0092 | 1 |
| O00763 | ACACB | AEGIPKIyVAANsGA | Y1826 | 0.033 | 98.1 | 2458 | 0.014477 | 0.033 | 0.0145 | 1 |
| Q16878 | CDO1 | KFDQYRYtRNLVDQG | T59 | 0.04 | 97.24 | 200 | 0.027321 | 0.04 | 0.0273 | 1 |
| Q9UK99 | FBXO3 | RNkNEVFyQCPDQMA | Y223 | 0.13 | 97.94 | 471 | 0.057029 | 0.13 | 0.057 | 1 |
| Q9Y574 | ASB4 | NCRPNGKtPLHVACE | T109 | 0.024 | 97.85 | 426 | 0.016392 | 0.024 | 0.0164 | 1 |
| P62280 | RPS11 | IDkkCPFtGNVsIrG | T63 | 0.067 | 93.9 | 158 | 0.045762 | 0.057 | 0.102 | 1 |
| P08133 | ANXA6 | QGAkyrGsIHDFPGF | S13 | 0.188 | 95.42 | 673 | 0.149444 | 0.188 | 0.1494 | 1 |
| P08133 | ANXA6 | TGkPIEAsIRGELsG | S229 | 0.01 | 89.56 | 673 | 0.007949 | 0.01 | 0.0079 | 1 |
| P08133 | ANXA6 | kNDTsGEykKTLLKL | Y313 | 0.164 | 98.25 | 673 | 0.071944 | 0.164 | 0.0719 | 0.8571 |
| Q16222 | UAP1 | VAYPkGMyDVGLPSR | Y125 | 0.284 | 98.03 | 522 | 0.124587 | 0.284 | 0.1246 | 1 |
| Q16222 | UAP1 | YIMTSGRtMESTkEF | T170 | 0.008 | 95.83 | 522 | 0.005464 | 0.008 | 0.0055 | 1 |

| | | | | | | | | | | |
|---|---|---|---|---|---|---|---|---|---|---|
| Q16222 | UAP1 | VLREDEFsPLkNADS | S404 | 0.103 | 98.54 | 522 | 0.081876 | 0.103 | 0.0819 | 0.9 |
| Q96HL8 | SH3YL1 | KETNRKFyCQDIRAY | Y180 | 0.323 | 96.5 | 342 | 0.141695 | 0.323 | 0.1417 | 1 |
| O95759 | TBC1D8 | LRRVLTAyAHRNPKI | Y580 | 0 | 96.09 | 1140 | 0 | 0 | 0 | 1 |
| Q9Y397 | ZDHHC9 | AAMLFLFsMAtLLRT | S79 | 0.003 | 98.59 | 364 | 0.002385 | 0.003 | 0.0024 | 0.8889 |
| Q9Y397 | ZDHHC9 | LFLFsMAtLLRTSFS | T82 | 0.097 | 98.57 | 364 | 0.066252 | 0.097 | 0.0663 | 1 |
| Q9H5V8 | CDCP1 | LVPKDRLsLVLVPAQ | S458 | 0 | 93.05 | 836 | 0 | 0 | 0 | 1 |
| P14324 | FDPS | DIQDNkCsWLVVQCL | S334 | 0.042 | 98.35 | 419 | 0.033386 | 0.042 | 0.0334 | 1 |
| P10696 | ALPG | ILGGGRKyMFPMGtP | Y225 | 0.126 | 98.88 | 532 | 0.055274 | 0.126 | 0.0553 | 1 |
| P10696 | ALPG | QLtsEEDtLSLVtAD | T369 | 0.02 | 98.76 | 532 | 0.01366 | 0.02 | 0.0137 | 1 |
| P10696 | ALPG | EDtLSLVtADHSHVF | T374 | 0.001 | 98.92 | 532 | 0.000683 | 0.001 | 0.0007 | 0.8333 |
| P10696 | ALPG | ARDRkAytVLLyGNG | T408 | 0.055 | 98.84 | 532 | 0.037566 | 0.055 | 0.0376 | 1 |
| P10696 | ALPG | PDVtEsEsGsPEyRQ | S433 | 0.007 | 98.12 | 532 | 0.005564 | 0.007 | 0.0056 | 1 |
| O60741 | HCN1 | PKVIKMNyLKSWFVV | Y217 | 0.228 | 93.95 | 890 | 0.10002 | 0.228 | 0.1 | 1 |
| Q14164 | IKBKE | GEEGQSIykLTDFGA | Y153 | 0.02 | 98.3 | 716 | 0.008774 | 0.02 | 0.0088 | 1 |
| Q7Z3J2 | VPS35L | HDSVNALtLEDEKRM | T625 | 0.176 | 92.29 | 963 | 0.12021 | 0.176 | 0.1202 | 1 |
| P0C024 | NUDT7 | GKLHLLFtVRSEKLR | T58 | 0.007 | 98.56 | 238 | 0.004781 | 0.007 | 0.0048 | 0.8182 |
| P0C024 | NUDT7 | TDMDDAAtALREAQE | T88 | 0.001 | 98.72 | 238 | 0.000683 | 0.001 | 0.0007 | 0.9167 |
| P42702 | LIFR | VGPRATSyTLVESFS | Y369 | 0.078 | 92.59 | 1097 | 0.034217 | 0.158 | 0.0693 | 1 |
| P42702 | LIFR | KGVENssyLVALDKL | Y492 | 0.122 | 92.72 | 1097 | 0.05352 | 0.122 | 0.0535 | 1 |
| P42702 | LIFR | LRGFLRGyLFyFGKG | Y759 | 0 | 93.47 | 1097 | 0 | 0 | 0 | 1 |
| P42702 | LIFR | DLQGkTSyHLVLRAY | Y807 | 0.153 | 90.03 | 1097 | 0.067119 | 0.153 | 0.0671 | 1 |
| Q9NTJ3 | SMC4 | GPFHkRFsCIIGPNG | S109 | 0 | 91.8 | 1288 | 0 | 0 | 0 | 1 |
| Q9NTJ3 | SMC4 | LsVLIHNsDEHkDIQ | S149 | 0.126 | 83.36 | 1288 | 0.100159 | 0.126 | 0.1002 | 1 |
| Q9NTJ3 | SMC4 | CRDNTSVyHIsGkkK | Y197 | 0.208 | 94.72 | 1288 | 0.091247 | 0.208 | 0.0912 | 1 |
| Q63ZE4 | SLC22A10 | ILSSGALsIGQIILG | S242 | 0.009 | 90.82 | 541 | 0.007154 | 0.009 | 0.0072 | 1 |
| Q96PD6 | MOGAT1 | VFREyVMsVGLVsVs | S158 | 0.097 | 95.51 | 335 | 0.077107 | 0.097 | 0.0771 | 1 |
| Q96PD6 | MOGAT1 | VMsVGLVsVsKKSVS | S163 | 0.037 | 97.06 | 335 | 0.029412 | 0.037 | 0.0294 | 1 |
| P40306 | PSMB10 | HFIAPkIyCCGAGVA | Y81 | 0.291 | 98.59 | 273 | 0.127658 | 0.291 | 0.1277 | 1 |
| Q9HCC0 | MCCC2 | ATVKGGAyYPVTVKK | Y145 | 0.106 | 98.07 | 563 | 0.046501 | 0.106 | 0.0465 | 1 |
| Q9HCC0 | MCCC2 | IARIVDGsRFTEFKA | S342 | 0.001 | 97.5 | 563 | 0.000795 | 0.001 | 0.0008 | 1 |
| Q9HCC0 | MCCC2 | GIIDPADtRLVLGLS | T537 | 0.005 | 98.86 | 563 | 0.003415 | 0.005 | 0.0034 | 1 |

| | | | | | | | | | | |
|---|---|---|---|---|---|---|---|---|---|---|
| P02746 | C1QB | TCKVPGLyYFTYHAS | Y168 | 0 | 97.81 | 253 | 0 | 0 | 0 | 1 |
| P62917 | RPL8 | VGtMPEGtIVCCLEE | T111 | 0.186 | 97.84 | 257 | 0.127041 | 0.186 | 0.127 | 1 |
| P05198 | EIF2S1 | EkGyIDLskRRVsPE | S86 | 0 | 88.35 | 315 | 0 | 0 | 0 | 1 |
| Q9ULL4 | PLXNB3 | IAQTFIDsCTTSEHK | S1793 | 0.01 | 90.02 | 1909 | 0.007949 | 0.01 | 0.0079 | 0.8333 |
| O94779 | CNTN5 | IAILPDGsLRILNAS | S535 | 0.016 | 88.91 | 1100 | 0.012719 | 0.016 | 0.0127 | 1 |
| Q5TGI0 | FAXC | RKQQEIDsKDAIILH | S96 | 0.025 | 70.51 | 409 | 0.019873 | 0.025 | 0.0199 | 1 |
| Q5TGI0 | FAXC | RPNNGVPsLsPFCLK | S114 | 0.023 | 95.44 | 409 | 0.018283 | 0.023 | 0.0183 | 1 |
| Q5TGI0 | FAXC | NNGVPsLsPFCLKME | S116 | 0.15 | 95.93 | 409 | 0.119237 | 0.15 | 0.1192 | 1 |
| P30047 | GCHFR | LGNNFyEyyVDDPPR | Y47 | 0.316 | 98.83 | 84 | 0.138625 | 0.316 | 0.1386 | 1 |
| P13497 | BMP1 | SRGIFLDtIVPKYEV | T285 | 0 | 93.12 | 986 | 0 | 0 | 0 | 1 |
| P13497 | BMP1 | GHIQSPNyPDDYRPS | Y451 | 0.077 | 86.95 | 986 | 0.033779 | 0.077 | 0.0338 | 1 |
| P13497 | BMP1 | DDIKSTSsRLWLKFV | S523 | 0.163 | 93.83 | 986 | 0.129571 | 0.163 | 0.1296 | 1 |
| Q16560 | SNRNP35 | RDLVtGFsKGYAFIE | S91 | 0.148 | 94.04 | 246 | 0.117647 | 0.148 | 0.1176 | 1 |
| O00418 | EEF2K | GkYIkYNsNsGFVRD | S241 | 0.115 | 87.96 | 725 | 0.091415 | 0.115 | 0.0914 | 1 |
| Q460N5 | PARP14 | ADVIVNstSNsFNLK | T1248 | 0.068 | 91.37 | 1801 | 0.046445 | 0.068 | 0.0464 | 0.9091 |
| Q00973 | B4GALNT1 | LVDVLERtPLDLVGG | T377 | 0.122 | 94.29 | 533 | 0.083328 | 0.122 | 0.0833 | 1 |
| P35268 | RPL22 | VtIERsKskItVtsE | S68 | 0.119 | 93.02 | 128 | 0.094595 | 0.119 | 0.0946 | 1 |
| P35268 | RPL22 | KskItVtsEVPFskR | S74 | 0.015 | 93.71 | 128 | 0.011924 | 0.015 | 0.0119 | 1 |
| Q8WWH5 | TRUB1 | FYIRSLVsDIGkELS | S260 | 0.115 | 98.55 | 349 | 0.091415 | 0.115 | 0.0914 | 1 |
| Q63HM2 | PCNX4 | PGHILRVyGGVLPWS | Y999 | 0.007 | 93.12 | 1172 | 0.003071 | 0.007 | 0.0031 | 1 |
| Q63HM2 | PCNX4 | LALKAFRyTLKLMID | Y1028 | 0.096 | 97.02 | 1172 | 0.042114 | 0.179 | 0.0753 | 1 |
| Q63HM2 | PCNX4 | LIKYLEEyERDWYIG | Y1055 | 0.013 | 95.78 | 1172 | 0.005703 | 0.013 | 0.0057 | 1 |
| A6NMX2 | EIF4E1B | GDKIAVWtREAENQA | T192 | 0.024 | 98.73 | 242 | 0.016392 | 0.024 | 0.0164 | 0.8333 |
| Q6ZW61 | BBS12 | TEGINLVtAVLTNPV | T501 | 0.003 | 96.52 | 710 | 0.002049 | 0.003 | 0.002 | 1 |
| O00750 | PIK3C2B | DGNDPDPyVKIYLLP | Y1541 | 0.264 | 94.39 | 1634 | 0.115813 | 0.264 | 0.1158 | 0.8182 |
| Q92736 | RYR2 | NEHtMMFtLNGEILL | T1176 | 0.026 | 96.72 | 4967 | 0.017758 | 0.026 | 0.0178 | 1 |
| Q68DX3 | FRMPD2 | QKKFTItssVtGKKH | S607 | 0.145 | 90.17 | 1309 | 0.115263 | 0.145 | 0.1153 | 1 |
| P00374 | DHFR | PEIDLEkykLLPEYP | Y157 | 0.076 | 97.86 | 187 | 0.03334 | 0.076 | 0.0333 | 1 |
| P00374 | DHFR | IkYkFEVyEkND___ | Y183 | 0.15 | 98.26 | 187 | 0.065803 | 0.15 | 0.0658 | 0.8182 |
| Q9BTY7 | HGH1 | VRHVLALtGCGPGRA | T49 | 0.025 | 97.01 | 390 | 0.017075 | 0.04 | 0.0273 | 1 |
| Q6IQ55 | TTBK2 | GRHDDLWsLFYMLVE | S211 | 0.052 | 98.83 | 1244 | 0.041336 | 0.052 | 0.0413 | 1 |

| | | | | | | | | | | |
|---|---|---|---|---|---|---|---|---|---|---|
| Q7L5D6 | GET4 | LCLKNKssAsVVFTT | S211 | 0.037 | 98.2 | 327 | 0.029412 | 0.037 | 0.0294 | 1 |
| O43741 | PRKAB2 | GGKEVFIsGsFNNWs | S93 | 0.018 | 97.37 | 272 | 0.014308 | 0.018 | 0.0143 | 1 |
| Q16718 | NDUFA5 | HERLRILyTkILDVL | Y28 | 0.093 | 94.65 | 116 | 0.040798 | 0.093 | 0.0408 | 1 |
| Q9UPQ3 | AGAP1 | THLSRVRsLDLDDWP | S663 | 0.006 | 96.11 | 857 | 0.004769 | 0.006 | 0.0048 | 1 |
| Q9UJ99 | CDH22 | IGSVAELsPTGTSVM | S181 | 0.131 | 91.22 | 828 | 0.104134 | 0.131 | 0.1041 | 1 |
| Q9UJ99 | CDH22 | GENTDMTyHLKDESS | Y314 | 0.083 | 93.82 | 828 | 0.036411 | 0.083 | 0.0364 | 1 |
| Q9UJ99 | CDH22 | AANRPVRyAIDRESD | Y430 | 0.236 | 95.27 | 828 | 0.10353 | 0.236 | 0.1035 | 1 |
| Q8NCG5 | CHST4 | DPRAVFRsRERTKGD | S213 | 0.147 | 95.7 | 386 | 0.116852 | 0.147 | 0.1169 | 1 |
| Q9UHP6 | RSPH14 | ARALLNVsISREGKK | S218 | 0 | 98.38 | 348 | 0 | 0 | 0 | 1 |
| P10586 | PTPRF | DGVPGSDyINANyID | Y1405 | 0.003 | 97.67 | 1907 | 0.001316 | 0.003 | 0.0013 | 1 |
| P10586 | PTPRF | RLVNIMPyELTRVCL | Y1676 | 0.095 | 88.9 | 1907 | 0.041675 | 0.3 | 0.1316 | 0.8571 |
| P10586 | PTPRF | RGVEGSDyINASFLD | Y1694 | 0.012 | 96.56 | 1907 | 0.005264 | 0.012 | 0.0053 | 1 |
| P10586 | PTPRF | GREKCHQyWPAERSA | Y1752 | 0 | 95.8 | 1907 | 0 | 0 | 0 | 1 |
| P10586 | PTPRF | AEYNMPQyILREFkV | Y1777 | 0.192 | 94.73 | 1907 | 0.084228 | 0.293 | 0.1063 | 1 |
| P10586 | PTPRF | MVQTEDQyQLCyRAA | Y1888 | 0 | 95.35 | 1907 | 0 | 0 | 0 | 1 |
| P10586 | PTPRF | EDQyQLCyRAALEyL | Y1892 | 0.044 | 94.92 | 1907 | 0.019302 | 0.078 | 0.0318 | 0.9333 |
| P30260 | CDC27 | KSEkALDtLNkAIVI | T691 | 0.182 | 94.49 | 824 | 0.124308 | 0.182 | 0.1243 | 1 |
| Q9UGM6 | WARS2 | RRSAGMNtARyKLAV | T299 | 0.211 | 96.46 | 360 | 0.144116 | 0.211 | 0.1441 | 1 |
| Q9UGM6 | WARS2 | AGMNtARyKLAVADA | Y302 | 0.048 | 98.26 | 360 | 0.021057 | 0.048 | 0.0211 | 1 |
| Q6JQN1 | ACAD10 | MVAPsMAsRVIDRAI | S1005 | 0.168 | 98.56 | 1059 | 0.133546 | 0.168 | 0.1335 | 1 |
| Q86W25 | NLRP13 | KELVPLAtLLITIKT | T358 | 0.077 | 90.44 | 1043 | 0.052592 | 0.119 | 0.0813 | 1 |
| P17612 | PRKACA | DQFERIktLGtGsFG | T49 | 0.001 | 97.35 | 351 | 0.000683 | 0.001 | 0.0007 | 1 |
| P17612 | PRKACA | HKETGNHyAMKILDK | Y70 | 0.253 | 98.73 | 351 | 0.110987 | 0.316 | 0.1386 | 1 |
| Q9H3P2 | NELFA | KLKLLLGtLHLPRRT | T60 | 0.086 | 94.9 | 528 | 0.058739 | 0.086 | 0.0587 | 1 |
| P47895 | ALDH1A3 | RSNLKRVtLELGGKN | T278 | 0.131 | 98.82 | 512 | 0.089475 | 0.131 | 0.0895 | 1 |
| P47895 | ALDH1A3 | VFVEEQVysEFVRRS | Y327 | 0.17 | 98.82 | 512 | 0.074577 | 0.17 | 0.0746 | 1 |
| P47895 | ALDH1A3 | NsTDyGLtAAVFTkN | T440 | 0.007 | 98.74 | 512 | 0.004781 | 0.007 | 0.0048 | 0.9231 |
| P47895 | ALDH1A3 | GEyALAEytEVKTVT | Y497 | 0.083 | 98.86 | 512 | 0.036411 | 0.083 | 0.0364 | 1 |
| O75342 | ALOX12B | GVQDLPGyyYRDDSL | Y496 | 0 | 97.76 | 701 | 0 | 0 | 0 | 0.8667 |
| Q04837 | SSBP1 | kTTWHRIsVFRPGLR | S88 | 0.073 | 97.15 | 148 | 0.058029 | 0.073 | 0.058 | 0.8571 |
| Q9Y3B4 | SF3B6 | RGTAyVVyEDIFDAk | Y64 | 0.018 | 98.05 | 125 | 0.007896 | 0.018 | 0.0079 | 1 |

| | | | | | | | | | | |
|---|---|---|---|---|---|---|---|---|---|---|
| Q8IZF5 | ADGRF3 | LKNLLIAtDKVLDMD | T557 | 0.006 | 95.59 | 1079 | 0.004098 | 0.006 | 0.0041 | 1 |
| P09958 | FURIN | PKFPQQWyLSGVTQR | Y123 | 0.019 | 96.97 | 794 | 0.008335 | 0.019 | 0.0083 | 1 |
| P00918 | CA2 | LkPLsVsyDQATSLR | Y51 | 0.035 | 98.15 | 260 | 0.015354 | 0.035 | 0.0154 | 0.8462 |
| P22309 | UGT1A1 | PQTVLWRyTGTRPSN | Y337 | 0.193 | 92.74 | 533 | 0.084666 | 0.193 | 0.0847 | 1 |
| P07359 | GP1BA | PPGLLTPtPKLEKLS | T163 | 0 | 98.01 | 652 | 0 | 0 | 0 | 1 |
| P69849 | NOMO3 | PLHsDLEyTVTSQkE | Y849 | 0.251 | 81.41 | 1222 | 0.11011 | 0.282 | 0.1237 | 1 |
| P69849 | NOMO3 | LLRDGENyVVLLDST | Y1110 | 0.096 | 93.36 | 1222 | 0.042114 | 0.096 | 0.0421 | 1 |
| P55210 | CASP7 | WRsPGRGsWFVQALC | S239 | 0.003 | 97.61 | 303 | 0.002385 | 0.003 | 0.0024 | 0.8 |
| Q9Y4G8 | RAPGEF2 | LADRIQLsGRYYLkN | S682 | 0.154 | 88.69 | 1499 | 0.122417 | 0.154 | 0.1224 | 1 |
| P13798 | APEH | RFCTNRRsALYYVDL | S280 | 0.003 | 97.51 | 732 | 0.002385 | 0.003 | 0.0024 | 0.8889 |
| P13798 | APEH | RYIPQVktPLLLMLG | T665 | 0.027 | 98.71 | 732 | 0.018441 | 0.027 | 0.0184 | 1 |
| Q7L1S5 | CHST9 | RQLIYDFyyLDyLMF | Y429 | 0.023 | 97.62 | 443 | 0.01009 | 0.023 | 0.0101 | 0.9333 |
| Q7L1S5 | CHST9 | YDFyyLDyLMFNYTT | Y433 | 0.041 | 97.42 | 443 | 0.017986 | 0.041 | 0.018 | 1 |
| P43403 | ZAP70 | AAHLPFFyGSISRAE | Y12 | 0.057 | 96.16 | 619 | 0.025005 | 0.057 | 0.025 | 1 |
| P43403 | ZAP70 | CLRSLGGyVLSLVHD | Y46 | 0.029 | 96.18 | 619 | 0.012722 | 0.029 | 0.0127 | 1 |
| P43403 | ZAP70 | ERQLNGTyAIAGGkA | Y69 | 0.133 | 92.66 | 619 | 0.058345 | 0.133 | 0.0583 | 1 |
| P43403 | ZAP70 | PAELCEFySRDPDGL | Y87 | 0.023 | 97.16 | 619 | 0.01009 | 0.023 | 0.0101 | 1 |
| P43403 | ZAP70 | AHERMPWyHssLTRE | Y164 | 0.145 | 95.48 | 619 | 0.063609 | 0.145 | 0.0636 | 1 |
| P43403 | ZAP70 | RPRkEQGtyALSLIy | T197 | 0.108 | 94.86 | 619 | 0.073765 | 0.161 | 0.1234 | 1 |
| P43403 | ZAP70 | PRkEQGtyALSLIyG | Y198 | 0.258 | 96.85 | 619 | 0.113181 | 0.258 | 0.1132 | 1 |
| Q6UW60 | PCSK4 | HIHIYSAsWGPEDDG | S258 | 0.166 | 97.48 | 755 | 0.131956 | 0.166 | 0.132 | 1 |
| Q9BU70 | TRMO | TGVFSTRsPHRPNAI | S118 | 0.186 | 92.59 | 441 | 0.147854 | 0.186 | 0.1479 | 1 |
| A2RTY3 | HEATR9 | VKyEAyRtLAILGCL | T204 | 0.006 | 95.23 | 570 | 0.004098 | 0.006 | 0.0041 | 0.9091 |
| Q6UXI9 | NPNT | SLGQyQCssFARCYN | S226 | 0.188 | 91.8 | 565 | 0.149444 | 0.188 | 0.1494 | 1 |
| Q9BVN2 | RUSC1 | LLIAVsVsVDKIISH | S494 | 0.056 | 98.25 | 902 | 0.044515 | 0.056 | 0.0445 | 1 |
| Q08J23 | NSUN2 | PEELAWHtNLSRKIL | T129 | 0.088 | 88.41 | 767 | 0.060105 | 0.131 | 0.0844 | 1 |
| Q08J23 | NSUN2 | SRKILRksPHLEkFH | S139 | 0.085 | 73.87 | 767 | 0.067568 | 0.085 | 0.0676 | 1 |
| Q08J23 | NSUN2 | IASLLEKsEGALELA | S339 | 0.013 | 96.52 | 767 | 0.010334 | 0.013 | 0.0103 | 1 |
| Q86WH2 | RASSF3 | TLNSNGIyTGFIkVQ | Y79 | 0.1 | 97.5 | 238 | 0.043869 | 0.1 | 0.0439 | 1 |
| Q9BXF3 | CECR2 | NGESsEYtKMSDNLE | T521 | 0.055 | 96.12 | 1484 | 0.037566 | 0.055 | 0.0376 | 1 |
| O94903 | PLPBP | LADkVNSsWQRKGsP | S129 | 0.074 | 98.37 | 275 | 0.058824 | 0.074 | 0.0588 | 1 |

| | | | | | | | | | | |
|---|---|---|---|---|---|---|---|---|---|---|
| Q9NZ71 | RTEL1 | SKHRVCPyyLSRNLk | Y209 | 0.133 | 91.43 | 1219 | 0.058345 | 0.242 | 0.1003 | 1 |
| Q9Y586 | MAB21L2 | LVyQLNKyyTERCQA | Y15 | 0.045 | 98.62 | 359 | 0.019741 | 0.045 | 0.0197 | 1 |
| Q9Y586 | MAB21L2 | GCRNKCLsVLKTLRD | S252 | 0.184 | 98.5 | 359 | 0.146264 | 0.184 | 0.1463 | 1 |
| Q6ZS30 | NBEAL1 | RRYNLRRsALEIFHV | S1951 | 0.059 | 86.2 | 2694 | 0.0469 | 0.083 | 0.066 | 1 |
| Q9UKS6 | PACSIN3 | EkGPQYGtLEKAWHA | T76 | 0.189 | 98.32 | 424 | 0.12909 | 0.116 | 0.1432 | 1 |
| O94952 | FBXO21 | QHVTAALyGVVNVKK | Y376 | 0.045 | 92.89 | 628 | 0.019741 | 0.045 | 0.0197 | 1 |
| Q9UL17 | TBX21 | SMPGNRLyVHPDSPN | Y220 | 0.125 | 94.7 | 535 | 0.054836 | 0.125 | 0.0548 | 0.8333 |
| P59095 | STARD6 | TSKKITVsSKAsRKF | S36 | 0.075 | 98.73 | 220 | 0.059619 | 0.075 | 0.0596 | 1 |
| O14727 | APAF1 | MLRKHPRsLLILDDV | S238 | 0.047 | 91.32 | 1248 | 0.037361 | 0.094 | 0.0747 | 1 |
| Q8TF71 | SLC16A10 | VLMCIGVtsGVGRLL | T338 | 0.218 | 95.81 | 515 | 0.148897 | 0.218 | 0.1489 | 1 |
| Q8TF71 | SLC16A10 | LMCIGVtsGVGRLLF | S339 | 0.047 | 95.03 | 515 | 0.037361 | 0.047 | 0.0374 | 0.9 |
| A6NDG6 | PGP | KLTKALRyLQQPGCL | Y187 | 0.038 | 98.32 | 321 | 0.01667 | 0.038 | 0.0167 | 1 |
| A6NDG6 | PGP | DVkNNQEsDCVskKK | S295 | 0.18 | 92.48 | 321 | 0.143085 | 0.18 | 0.1431 | 1 |
| Q71RS6 | SLC24A5 | sAPAEVNsRGLTYIT | S435 | 0.038 | 68.35 | 500 | 0.030207 | 0.038 | 0.0302 | 1 |
| P55286 | CDH8 | KNGHYFLySLLPEMV | Y533 | 0.308 | 91.4 | 799 | 0.135115 | 0.308 | 0.1351 | 1 |
| O76000 | OR2B3 | CLRMAAFsWLIGFGN | S148 | 0.015 | 97.98 | 313 | 0.011924 | 0.015 | 0.0119 | 0.8333 |
| O76000 | OR2B3 | WLIGFGNsVLQSSLT | S156 | 0 | 93.85 | 313 | 0 | 0 | 0 | 1 |
| P46459 | NSF | IDsNPyDtDKMAAEF | T114 | 0 | 90.3 | 744 | 0 | 0 | 0 | 1 |
| O60306 | AQR | IkVIEDIyEkEIVKS | Y50 | 0.127 | 92.58 | 1485 | 0.055713 | 0.127 | 0.0557 | 0.9167 |
| O60306 | AQR | LLVHCyLsNLVRREE | S286 | 0.088 | 94.44 | 1485 | 0.069952 | 0.049 | 0.0842 | 1 |
| Q9HB75 | PIDD1 | VIFPPGAtEEPRRVS | T485 | 0.036 | 81.89 | 910 | 0.024588 | 0.036 | 0.0246 | 1 |
| Q9UP95 | SLC12A4 | DAPSLKEsLPLyVVA | S407 | 0.053 | 73.54 | 1085 | 0.04213 | 0.106 | 0.0843 | 1 |
| Q9UP95 | SLC12A4 | LKEsLPLyVVADIAt | Y411 | 0.099 | 89.14 | 1085 | 0.04343 | 0.099 | 0.0434 | 1 |
| Q9UP95 | SLC12A4 | IMAGSNRsGDLRDAQ | S441 | 0.061 | 84.19 | 1085 | 0.04849 | 0.061 | 0.0485 | 1 |
| P16278 | GLB1 | YISGSIHysRVPRFY | Y57 | 0.016 | 98.65 | 677 | 0.007019 | 0.016 | 0.007 | 1 |
| P16278 | GLB1 | LRSSDPDyLAAVDKW | Y154 | 0 | 98.58 | 677 | 0 | 0 | 0 | 0.9375 |
| Q9ULB1 | NRXN1 | AYGILMAtTsRDSAD | T758 | 0.001 | 97.78 | 1477 | 0.000683 | 0.001 | 0.0007 | 1 |
| Q9ULB1 | NRXN1 | GILMAtTsRDSADTL | S760 | 0.05 | 92.53 | 1477 | 0.039746 | 0.05 | 0.0397 | 1 |
| Q9BQS2 | SYT15 | CSPLVKLyLLPDERR | Y188 | 0.28 | 96.92 | 421 | 0.122832 | 0.28 | 0.1228 | 1 |
| Q9BQS2 | SYT15 | ELDTASLsLTVVQNM | S364 | 0.118 | 95.93 | 421 | 0.0938 | 0.118 | 0.0938 | 1 |
| Q96JM7 | L3MBTL3 | EHRRtLItPPGYPNV | T438 | 0.159 | 96.34 | 780 | 0.108599 | 0.159 | 0.1086 | 1 |

| | | | | | | | | | | |
|---|---|---|---|---|---|---|---|---|---|---|
| Q5K651 | SAMD9 | KVSWWNFyFSSESYS | Y695 | 0.093 | 95.4 | 1589 | 0.040798 | 0.093 | 0.0408 | 1 |
| O43598 | DNPH1 | PALYFCGsIrGGRED | S28 | 0 | 96.1 | 174 | 0 | 0 | 0 | 0.8 |
| O43598 | DNPH1 | DVVVAEVtQPsLGVG | T95 | 0.068 | 98.49 | 174 | 0.046445 | 0.068 | 0.0464 | 0.875 |
| Q9Y620 | RAD54B | SGkLQVLsKLLAVIH | S648 | 0.168 | 92.18 | 910 | 0.133546 | 0.168 | 0.1335 | 1 |
| Q9NV66 | TYW1 | YQLIGSHsGVKLCRW | S376 | 0.001 | 97.43 | 732 | 0.000795 | 0.001 | 0.0008 | 1 |
| O94788 | ALDH1A2 | PVDGDyFtFtRHEPI | T170 | 0.047 | 98.78 | 518 | 0.032102 | 0.047 | 0.0321 | 1 |
| O94788 | ALDH1A2 | DGDyFtFtRHEPIGV | T172 | 0.193 | 98.87 | 518 | 0.131822 | 0.193 | 0.1318 | 1 |
| O94788 | ALDH1A2 | RsNLKRVtLELGGKS | T284 | 0.097 | 98.91 | 518 | 0.066252 | 0.097 | 0.0663 | 1 |
| O94788 | ALDH1A2 | IFVEESIyEEFVRRS | Y333 | 0.033 | 98.91 | 518 | 0.014477 | 0.033 | 0.0145 | 1 |
| Q96AC1 | FERMT2 | GWLDSSRsLMEQDVK | S258 | 0.024 | 97.05 | 680 | 0.019078 | 0.024 | 0.0191 | 1 |
| Q96AC1 | FERMT2 | ITPECLVsPRYLkKY | S523 | 0.025 | 88.13 | 680 | 0.019873 | 0.025 | 0.0199 | 1 |
| Q9H0H5 | RACGAP1 | RIKFGkLsLKCRDCR | S313 | 0.048 | 93.34 | 632 | 0.038156 | 0.048 | 0.0382 | 0.9091 |
| Q01094 | E2F1 | YEtSLNLttKRFLEL | T135 | 0.122 | 91.85 | 437 | 0.083328 | 0.122 | 0.0833 | 1 |
| P46100 | ATRX | yYRLDGsttAQsRKk | T2090 | 0.156 | 83.96 | 2492 | 0.10655 | 0.156 | 0.1066 | 0.8333 |
| O75153 | CLUH | RFNPDIFsPGVRFPE | S723 | 0.131 | 85.58 | 1309 | 0.104134 | 0.129 | 0.1174 | 1 |
| O75153 | CLUH | IGIGELItRSAKHIF | T818 | 0 | 98.39 | 1309 | 0 | 0 | 0 | 0.8 |
| O75153 | CLUH | LGEDHEktKESSEYL | T1187 | 0.018 | 94.39 | 1309 | 0.012294 | 0.018 | 0.0123 | 1 |
| Q7Z4G4 | TRMT11 | QRELIESySVkKRHF | Y186 | 0.221 | 96.94 | 463 | 0.09695 | 0.221 | 0.0969 | 1 |
| P31512 | FMO4 | TAAQVLLstRTGTWV | S213 | 0.054 | 98.13 | 558 | 0.042925 | 0.054 | 0.0429 | 1 |
| P31512 | FMO4 | GLKGSILsGTELQAR | S379 | 0.014 | 96.83 | 558 | 0.011129 | 0.014 | 0.0111 | 0.9 |
| P31512 | FMO4 | ELQARWVtRVFKGLC | T389 | 0.003 | 98.5 | 558 | 0.002049 | 0.003 | 0.002 | 1 |
| P31512 | FMO4 | FFGPCTPyQYRLMGP | Y469 | 0.002 | 98.39 | 558 | 0.000877 | 0.002 | 0.0009 | 1 |
| Q9H2V7 | SPNS1 | APAFLLRsRVVLGET | S301 | 0.015 | 94.53 | 528 | 0.011924 | 0.015 | 0.0119 | 1 |
| Q99496 | RNF2 | IYPsRDEyEAHQERV | Y120 | 0.296 | 79.66 | 336 | 0.129851 | 0.296 | 0.1299 | 1 |
| Q15262 | PTPRK | SEETIIQtDEDVPGP | T485 | 0.041 | 92.03 | 1439 | 0.028004 | 0.041 | 0.028 | 1 |
| O96018 | APBA3 | LGStQLVsERNPPTS | S233 | 0.014 | 72.6 | 575 | 0.011129 | 0.125 | 0.0994 | 1 |
| P42892 | ECE1 | RRDEELIyHKVTAAE | Y331 | 0.173 | 98.5 | 770 | 0.075893 | 0.173 | 0.0759 | 0.8182 |
| P42892 | ECE1 | GLITDPHsPsRFRVI | S733 | 0.05 | 98.69 | 770 | 0.039746 | 0.05 | 0.0397 | 1 |
| P42892 | ECE1 | HFRCPPGsPMNPPHK | S759 | 0.184 | 98.47 | 770 | 0.146264 | 0.184 | 0.1463 | 1 |
| P16871 | IL7R | VLMHDVAyRQEKDEN | Y169 | 0.278 | 97.22 | 459 | 0.121955 | 0.278 | 0.122 | 1 |
| Q9Y680 | FKBP7 | DGsKFyCsRTQNEGH | S74 | 0 | 90.63 | 222 | 0 | 0 | 0 | 1 |

| | | | | | | | | | | |
|---|---|---|---|---|---|---|---|---|---|---|
| O43837 | IDH3B | LVIIREQtEGEyssL | T167 | 0.15 | 94.45 | 385 | 0.102452 | 0.15 | 0.1025 | 1 |
| O43837 | IDH3B | TAMLLSAsNMLRHLN | S330 | 0.012 | 98.49 | 385 | 0.009539 | 0.012 | 0.0095 | 1 |
| O43837 | IDH3B | DMGGySttTDFIksV | T369 | 0.014 | 97.55 | 385 | 0.009562 | 0.014 | 0.0096 | 1 |
| Q9NP90 | RAB9B | FsVDDRQsFENLGNW | S94 | 0 | 98.85 | 201 | 0 | 0 | 0 | 1 |
| P06396 | GSN | GRARVHVsEEGtEPE | S261 | 0.068 | 95.03 | 782 | 0.054054 | 0.068 | 0.0541 | 1 |
| Q92499 | DDX1 | DkkGHQFsCVCLHGD | S541 | 0.006 | 98.25 | 740 | 0.004769 | 0.006 | 0.0048 | 1 |
| P82251 | SLC7A9 | GHMLkVLsYIsVRRL | S342 | 0.075 | 83.57 | 487 | 0.059619 | 0.075 | 0.0596 | 1 |
| P82251 | SLC7A9 | LkVLsYIsVRRLTPA | S345 | 0.073 | 82.33 | 487 | 0.058029 | 0.073 | 0.058 | 1 |
| Q9UIK4 | DAPK2 | QAMVRREsVVNLENF | S299 | 0.146 | 94.78 | 370 | 0.116058 | 0.146 | 0.1161 | 1 |
| Q9H3S4 | TPK1 | RFDQIMAsVNTLFQA | S138 | 0.075 | 98.84 | 243 | 0.059619 | 0.075 | 0.0596 | 0.9 |
| Q9UBS0 | RPS6KB2 | GTNLGkIyAMKVLRK | Y96 | 0.074 | 96.77 | 482 | 0.032463 | 0.105 | 0.0461 | 1 |
| Q9UBS0 | RPS6KB2 | PFIVELAyAFQTGGK | Y136 | 0.043 | 92.94 | 482 | 0.018863 | 0.043 | 0.0189 | 1 |
| Q12834 | CDC20 | SYILSSGsRSGHIHH | S285 | 0.001 | 98.94 | 499 | 0.000795 | 0.001 | 0.0008 | 1 |
| Q12834 | CDC20 | TMSPDGAtVASAAAD | T457 | 0.158 | 98.17 | 499 | 0.107916 | 0.158 | 0.1079 | 0.8889 |
| Q12834 | CDC20 | ASAAADEtLRLWRCF | T466 | 0.219 | 98.66 | 499 | 0.14958 | 0.219 | 0.1496 | 0.8571 |
| Q9NPB1 | NT5M | EMAsLQNtDVFICTS | T124 | 0.129 | 98.4 | 228 | 0.088109 | 0.129 | 0.0881 | 1 |
| Q9NPB1 | NT5M | IVLTRDKtVVSADLL | T166 | 0.063 | 98.79 | 228 | 0.04303 | 0.063 | 0.043 | 1 |
| Q96PQ7 | KLHL5 | GGRDGLktLNtVECY | T527 | 0.135 | 96.87 | 755 | 0.092207 | 0.135 | 0.0922 | 1 |
| P06340 | HLA-DOA | VPSAEDVyDCQVEHW | Y187 | 0.071 | 98.16 | 250 | 0.031147 | 0.071 | 0.0311 | 1 |
| Q13435 | SF3B2 | LKIPGLNsPIPESCS | S655 | 0.174 | 85.8 | 895 | 0.138315 | 0.174 | 0.1383 | 1 |
| O75333 | TBX10 | ILNSMHRyQPRFHVV | Y192 | 0.131 | 98.08 | 385 | 0.057468 | 0.131 | 0.0575 | 1 |
| Q5I0X7 | TTC32 | PNFEVPYyNRGLILY | Y96 | 0.327 | 98.89 | 151 | 0.14345 | 0.327 | 0.1434 | 1 |
| P30566 | ADSL | DVMAHVHtFGHCCPK | T94 | 0.037 | 98.69 | 484 | 0.025272 | 0.037 | 0.0253 | 1 |
| P30566 | ADSL | GIIHLGAtSCYVGDN | T111 | 0.167 | 96.94 | 484 | 0.114063 | 0.167 | 0.1141 | 1 |
| Q92888 | ARHGEF1 | WDQEAQIyELVAQTV | Y738 | 0.028 | 96.33 | 912 | 0.012283 | 0.028 | 0.0123 | 1 |
| Q9NVP2 | ASF1B | GyYVNNEyLNPELRE | Y117 | 0.058 | 97.5 | 202 | 0.025444 | 0.058 | 0.0254 | 1 |
| Q13564 | NAE1 | kLLkEQkyDRQLRLW | Y13 | 0.061 | 96.85 | 534 | 0.02676 | 0.061 | 0.0268 | 1 |
| P10746 | UROS | DDCGQDPyIRELGLY | Y19 | 0.255 | 96.07 | 265 | 0.111865 | 0.255 | 0.1119 | 1 |
| Q9UBS5 | GABBR1 | RRDILPDyELKLIHH | Y209 | 0.127 | 91.75 | 961 | 0.055713 | 0.127 | 0.0557 | 1 |
| P23352 | ANOS1 | IsIEPVIyVVQRRWN | Y217 | 0.059 | 94.67 | 680 | 0.025882 | 0.326 | 0.143 | 1 |
| P23352 | ANOS1 | DIRPSRWyQFRVAAV | Y259 | 0.016 | 95.08 | 680 | 0.007019 | 0.016 | 0.007 | 1 |

| | | | | | | | | | | |
|---|---|---|---|---|---|---|---|---|---|---|
| Q7RTY8 | TMPRSS7 | INLVYTTsAFsKFYE | S151 | 0.184 | 91.22 | 843 | 0.146264 | 0.184 | 0.1463 | 1 |
| P22695 | UQCRC2 | ASGNLGHtPFVDEL_ | T447 | 0.082 | 97.67 | 453 | 0.056007 | 0.082 | 0.056 | 1 |
| O75368 | SH3BGRL | GLtAPPGsKEAEVQA | S103 | 0.133 | 91.32 | 114 | 0.105724 | 0.072 | 0.1263 | 0.8333 |
| Q01668 | CACNA1D | GTVMFNAtLFALVRT | T1549 | 0.045 | 88.31 | 2161 | 0.030736 | 0.045 | 0.0307 | 1 |
| P00519 | ABL1 | QGWVPsNyItPVNSL | Y115 | 0.167 | 96.23 | 1130 | 0.07326 | 0.167 | 0.0733 | 1 |
| P00519 | ABL1 | CtREPPFyIITEFMT | Y312 | 0.249 | 93.88 | 1130 | 0.109233 | 0.249 | 0.1092 | 1 |
| Q9HAW8 | UGT1A10 | PQTVLWRyTGTRPSN | Y334 | 0.188 | 95.13 | 530 | 0.082473 | 0.188 | 0.0825 | 1 |
| C9JQI7 | TMEM232 | LSFSGASyHKYPNIF | Y209 | 0.001 | 86.95 | 657 | 0.000439 | 0.2 | 0.0877 | 1 |
| O75449 | KATNA1 | yDSAMVYyQGVLDQM | Y29 | 0.102 | 93.92 | 491 | 0.044746 | 0.102 | 0.0447 | 0.9167 |
| Q13224 | GRIN2B | GVDDALLsLKTGKLD | S720 | 0.146 | 93.01 | 1484 | 0.116058 | 0.146 | 0.1161 | 1 |
| Q13224 | GRIN2B | GIAIQKDsGWKRQVD | S770 | 0.165 | 87.11 | 1484 | 0.131161 | 0.165 | 0.1312 | 1 |
| O95271 | TNKS | HPkRKQVtELLLRKG | T541 | 0.001 | 97.05 | 1327 | 0.000683 | 0.001 | 0.0007 | 1 |
| Q8WW52 | FAM151A | STSPNRTyTQAMVEK | Y223 | 0.033 | 95.76 | 585 | 0.014477 | 0.033 | 0.0145 | 0.8125 |
| Q8WW52 | FAM151A | GGVPQRVtFPVRSSM | T243 | 0.001 | 98.75 | 585 | 0.000683 | 0.001 | 0.0007 | 1 |
| Q12879 | GRIN2A | GVEDALVsLKTGKLD | S719 | 0.142 | 93.37 | 1464 | 0.112878 | 0.142 | 0.1129 | 1 |
| Q12879 | GRIN2A | GKLDAFIyDAAVLNy | Y730 | 0.225 | 93.1 | 1464 | 0.098704 | 0.225 | 0.0987 | 0.8 |
| Q9GZT8 | NIF3L1 | LENRVGIysPHTAyD | Y124 | 0.042 | 98.85 | 377 | 0.018425 | 0.042 | 0.0184 | 1 |
| Q9GZT8 | NIF3L1 | ENRVGIysPHTAyDA | S125 | 0 | 98.76 | 377 | 0 | 0 | 0 | 1 |
| Q9GZT8 | NIF3L1 | IysPHTAyDAAPQGV | Y130 | 0.016 | 98.31 | 377 | 0.007019 | 0.016 | 0.007 | 1 |
| Q14679 | TTLL4 | IKDVVVKtIIsSEPy | T864 | 0 | 98.47 | 1199 | 0 | 0 | 0 | 1 |
| Q14679 | TTLL4 | VVVKtIIsSEPyVTS | S867 | 0.002 | 97.4 | 1199 | 0.00159 | 0.002 | 0.0016 | 1 |
| Q9NY65 | TUBA8 | kEDAANNyARGHYTV | Y103 | 0.024 | 98 | 449 | 0.010528 | 0.024 | 0.0105 | 1 |
| Q9NY65 | TUBA8 | tNLNRLIsQIVssIt | S232 | 0.017 | 97.14 | 449 | 0.013514 | 0.017 | 0.0135 | 0.8889 |
| P50570 | DNM2 | TGTNkGIsPVPINLR | S116 | 0.125 | 90.07 | 870 | 0.099364 | 0.125 | 0.0994 | 1 |
| P50570 | DNM2 | PINLRVysPHVLNLT | S126 | 0.003 | 93.61 | 870 | 0.002385 | 0.003 | 0.0024 | 1 |
| P50570 | DNM2 | LLPLRRGyIGVVNRS | Y231 | 0.129 | 96.93 | 870 | 0.05659 | 0.129 | 0.0566 | 1 |
| P11142 | HSPA8 | VAMNPtNtVFDAkRL | T66 | 0 | 98.03 | 646 | 0 | 0 | 0 | 1 |
| P11142 | HSPA8 | rPkVQVEykGEtksF | Y107 | 0.018 | 94.26 | 646 | 0.007896 | 0.182 | 0.0798 | 1 |
| P11142 | HSPA8 | DGIFEVkstAGDtHL | S221 | 0 | 95.49 | 646 | 0 | 0 | 0 | 1 |
| P11142 | HSPA8 | NADLFRGtLDPVEkA | T313 | 0.023 | 97.64 | 646 | 0.015709 | 0.023 | 0.0157 | 0.9091 |
| P11142 | HSPA8 | IVLVGGStRIPkIQk | T341 | 0.039 | 97.89 | 646 | 0.026638 | 0.039 | 0.0266 | 1 |

| | | | | | | | | | | |
|---|---|---|---|---|---|---|---|---|---|---|
| P11142 | HSPA8 | LLDVTPLsLGIETAG | S400 | 0.082 | 90.13 | 646 | 0.065183 | 0.082 | 0.0652 | 1 |
| Q14008 | CKAP5 | PkLEAGDyADLVkAL | Y317 | 0.044 | 93.16 | 2032 | 0.019302 | 0.077 | 0.0311 | 1 |
| Q5JPE7 | NOMO2 | LLRDGENyVVLLDST | Y1110 | 0.092 | 93.43 | 1267 | 0.040359 | 0.092 | 0.0404 | 1 |
| Q5JPE7 | NOMO2 | STLPRSQyDyILPQV | Y1123 | 0.238 | 87.65 | 1267 | 0.104407 | 0.238 | 0.1044 | 1 |
| Q5JPE7 | NOMO2 | LPRSQyDyILPQVSF | Y1125 | 0.249 | 90.04 | 1267 | 0.109233 | 0.249 | 0.1092 | 1 |
| Q9UBX5 | FBLN5 | QtGPISAtLVMTRPI | T403 | 0.118 | 97.98 | 448 | 0.080596 | 0.118 | 0.0806 | 1 |
| Q86VP6 | CAND1 | QRALKLGtLsALDIL | T670 | 0 | 96.88 | 1230 | 0 | 0 | 0 | 1 |
| Q13426 | XRCC4 | __MERKIsRIHLVSE | S6 | 0.036 | 95.43 | 336 | 0.028617 | 0.036 | 0.0286 | 1 |
| A6NK06 | ACOD1 | LGSAAAAsKFLGLSS | S174 | 0.023 | 98.93 | 481 | 0.018283 | 0.023 | 0.0183 | 1 |
| P43243 | MATR3 | kPVRVHLsQkyKRIk | S471 | 0.032 | 82.78 | 847 | 0.025437 | 0.064 | 0.0753 | 1 |
| Q9P2J3 | KLHL9 | ELATVECyNPRMNEW | Y419 | 0.011 | 98.8 | 617 | 0.004826 | 0.011 | 0.0048 | 0.9231 |
| Q8IXI2 | RHOT1 | TVLRRFGyDDDLDLt | Y276 | 0.157 | 97.47 | 618 | 0.068874 | 0.157 | 0.0689 | 1 |
| Q8IXI2 | RHOT1 | KIPPDCttELNHHAY | T299 | 0.086 | 97.93 | 618 | 0.058739 | 0.086 | 0.0587 | 1 |
| Q07864 | POLE | KEERRGPtLIAVQSs | T1583 | 0.051 | 91.2 | 2286 | 0.034834 | 0.154 | 0.1123 | 1 |
| Q96K76 | USP47 | DPVTSIPyQLQRLFV | Y234 | 0.291 | 93.66 | 1375 | 0.127658 | 0.291 | 0.1277 | 1 |
| Q96K76 | USP47 | LRFLHFPyLLtLQLK | Y384 | 0.042 | 95.92 | 1375 | 0.018425 | 0.042 | 0.0184 | 1 |
| Q96K76 | USP47 | LHFPyLLtLQLKRFD | T387 | 0.064 | 97.32 | 1375 | 0.043713 | 0.064 | 0.0437 | 1 |
| Q96K76 | USP47 | LEKNSLIyELFSVMV | Y487 | 0.003 | 96.82 | 1375 | 0.001316 | 0.003 | 0.0013 | 1 |
| Q96K76 | USP47 | GSAAGGHyyACIKSF | Y504 | 0.133 | 93.59 | 1375 | 0.058345 | 0.133 | 0.0583 | 1 |
| Q96K76 | USP47 | tQEDIKKtHGGsSGS | T535 | 0.06 | 92.54 | 1375 | 0.040981 | 0.06 | 0.041 | 1 |
| Q96K76 | USP47 | FASSTNAyMLIyRLK | Y557 | 0.046 | 93.87 | 1375 | 0.02018 | 0.046 | 0.0202 | 1 |
| Q96K76 | USP47 | TNAyMLIyRLKDPAR | Y561 | 0 | 97.77 | 1375 | 0 | 0 | 0 | 1 |
| P01116 | KRAS | FAINNTKsFEDIHHy | S89 | 0.016 | 98.21 | 189 | 0.012719 | 0.016 | 0.0127 | 0.9 |
| P49736 | MCM2 | QRIRIQEsPGkVAAG | S381 | 0.159 | 82.25 | 904 | 0.126391 | 0.159 | 0.1264 | 1 |
| Q9H903 | MTHFD2L | KKKAGFItPVPGGVG | T320 | 0.016 | 98.64 | 347 | 0.010928 | 0.016 | 0.0109 | 1 |
| O00391 | QSOX1 | IyMADLEsALHYILR | S306 | 0.023 | 97.21 | 747 | 0.018283 | 0.023 | 0.0183 | 1 |
| P23219 | PTGS1 | GKMGPGFtKALGHGV | T220 | 0.016 | 98.75 | 599 | 0.010928 | 0.016 | 0.0109 | 1 |
| Q6PIU2 | NCEH1 | SLLPAsFtKNYkPVV | T297 | 0.211 | 97.63 | 408 | 0.144116 | 0.211 | 0.1441 | 1 |
| P04439 | HLA-A | IEQEGPEyWDQETRN | Y83 | 0.189 | 97.76 | 365 | 0.082912 | 0.23 | 0.1009 | 1 |
| P04439 | HLA-A | DAYDGkDyIALNEDL | Y147 | 0.148 | 98.75 | 365 | 0.064925 | 0.148 | 0.0649 | 1 |
| P04181 | OAT | YNNVLGEyEEyITKL | Y123 | 0.006 | 98.92 | 439 | 0.002632 | 0.006 | 0.0026 | 1 |

| | | | | | | | | | | |
|---|---|---|---|---|---|---|---|---|---|---|
| O75751 | SLC22A3 | LVMLLPEtKGIALPE | T520 | 0.118 | 82.41 | 556 | 0.080596 | 0.118 | 0.0806 | 0.8333 |
| Q9BQI3 | EIF2AK1 | NkLDGQYyAIkKILI | Y193 | 0.288 | 96.97 | 630 | 0.126341 | 0.288 | 0.1263 | 1 |
| P62253 | UBE2G1 | __MtELQsALLLRRQ | S6 | 0.036 | 94.51 | 170 | 0.028617 | 0.036 | 0.0286 | 1 |
| P62253 | UBE2G1 | PPKMkFItEIWHPNV | T76 | 0.183 | 96.86 | 170 | 0.124991 | 0.183 | 0.125 | 1 |
| Q8IUR7 | ARMC8 | kPIEMQLtSAKCLty | T255 | 0.093 | 98.74 | 673 | 0.06352 | 0.093 | 0.0635 | 1 |
| Q8IUR7 | ARMC8 | LtSAKCLtyMCRAGA | T261 | 0.001 | 98.86 | 673 | 0.000683 | 0.001 | 0.0007 | 1 |
| Q9Y6F9 | WNT6 | APNRRTGsPGTRGRA | S302 | 0.073 | 93.1 | 365 | 0.058029 | 0.073 | 0.058 | 1 |
| P35580 | MYH10 | HEMPPHIyAISESAY | Y155 | 0 | 95.16 | 1976 | 0 | 0 | 0 | 1 |
| P35580 | MYH10 | RQRYEILtPNAIPKG | T732 | 0.069 | 88.97 | 1976 | 0.047128 | 0.069 | 0.0471 | 1 |
| Q6UXB8 | PI16 | AQVSPTAsDMLHMRW | S52 | 0.002 | 98.71 | 463 | 0.00159 | 0.002 | 0.0016 | 1 |
| Q15067 | ACOX1 | KEkWLLSsKGLQIIG | S127 | 0 | 94.84 | 660 | 0 | 0 | 0 | 1 |
| Q15067 | ACOX1 | TQEFILNsPTVTSIK | S167 | 0.064 | 98.65 | 660 | 0.050875 | 0.082 | 0.0703 | 1 |
| Q15067 | ACOX1 | PESLTEAyKLRAARL | Y487 | 0.006 | 98.01 | 660 | 0.002632 | 0.006 | 0.0026 | 1 |
| Q6NVY1 | HIBCH | QEVLTMEyRLSQACM | Y330 | 0.068 | 98.85 | 386 | 0.029831 | 0.068 | 0.0298 | 0.8571 |
| P55157 | MTTP | GSSDIREtVMIItGt | T427 | 0.013 | 95.92 | 894 | 0.008879 | 0.013 | 0.0089 | 0.8889 |
| P55157 | MTTP | REtVMIItGtLVRKL | T432 | 0.024 | 98.23 | 894 | 0.016392 | 0.024 | 0.0164 | 1 |
| P55157 | MTTP | tVMIItGtLVRKLCQ | T434 | 0.062 | 96.75 | 894 | 0.042347 | 0.062 | 0.0423 | 0.9167 |
| P55157 | MTTP | ATTALQRyDLPFITD | Y511 | 0.102 | 92.93 | 894 | 0.044746 | 0.102 | 0.0447 | 0.8462 |
| P55157 | MTTP | DVKNILLsIGELPQE | S563 | 0.174 | 92.54 | 894 | 0.138315 | 0.174 | 0.1383 | 1 |
| P21217 | FUT3 | DSARVRyyQSLQAHL | Y210 | 0.194 | 98.28 | 361 | 0.085105 | 0.194 | 0.0851 | 1 |
| P21217 | FUT3 | AHLKVDVyGRSHKPL | Y222 | 0.241 | 98.29 | 361 | 0.105723 | 0.241 | 0.1057 | 1 |
| P12268 | IMPDH2 | ITDPVVLsPkDRVRD | S122 | 0.151 | 97.06 | 514 | 0.120032 | 0.151 | 0.12 | 1 |
| P12268 | IMPDH2 | RPQATAVykVSEYAR | Y348 | 0.209 | 98.09 | 514 | 0.091685 | 0.209 | 0.0917 | 1 |
| P12268 | IMPDH2 | DGIRLKKyrGMGsLD | Y411 | 0.141 | 86.77 | 514 | 0.061855 | 0.141 | 0.0619 | 1 |
| P12268 | IMPDH2 | KKyrGMGsLDAMDkH | S416 | 0.025 | 83.84 | 514 | 0.019873 | 0.025 | 0.0199 | 1 |
| P12268 | IMPDH2 | DAMDkHLssQNryFs | S425 | 0.156 | 75.1 | 514 | 0.124007 | 0.156 | 0.124 | 1 |
| P12268 | IMPDH2 | AMDkHLssQNryFsE | S426 | 0.013 | 75.08 | 514 | 0.010334 | 0.013 | 0.0103 | 1 |
| P12268 | IMPDH2 | HLssQNryFsEADkI | Y430 | 0.296 | 81.68 | 514 | 0.129851 | 0.296 | 0.1299 | 1 |
| P12268 | IMPDH2 | ssQNryFsEADkIkV | S432 | 0.05 | 73.94 | 514 | 0.039746 | 0.05 | 0.0397 | 1 |
| P36954 | POLR2I | KENRILLyACRNCDY | Y37 | 0.279 | 95.99 | 125 | 0.122393 | 0.279 | 0.1224 | 1 |
| P36954 | POLR2I | TQIIADVsQDPTLPR | S73 | 0.103 | 92.05 | 125 | 0.081876 | 0.103 | 0.0819 | 1 |

| | | | | | | | | | | |
|---|---|---|---|---|---|---|---|---|---|---|
| Q9Y6N5 | SQOR | ALGIQLDyEKIkGLP | Y151 | 0.225 | 98.78 | 450 | 0.098704 | 0.225 | 0.0987 | 1 |
| Q9Y6N5 | SQOR | VFGIGDCtNLPtsKT | T338 | 0.026 | 98.93 | 450 | 0.017758 | 0.026 | 0.0178 | 1 |
| Q9Y6N5 | SQOR | ETFPFDQskERLsMy | S408 | 0.184 | 98.57 | 450 | 0.146264 | 0.184 | 0.1463 | 0.8 |
| Q9Y6N5 | SQOR | skERLsMyLMKADLM | Y415 | 0.254 | 98.75 | 450 | 0.111426 | 0.254 | 0.1114 | 1 |
| Q96QU6 | ACCS | RLPDPQRtHVMWAts | T315 | 0.085 | 98.74 | 501 | 0.058056 | 0.085 | 0.0581 | 0.875 |
| Q96QU6 | ACCS | tHVMWAtsKDFGMSG | S322 | 0.123 | 97.69 | 501 | 0.097775 | 0.123 | 0.0978 | 1 |
| P53804 | TTC3 | YRYCDALsMLGEYDW | S310 | 0.025 | 93.92 | 2025 | 0.019873 | 0.025 | 0.0199 | 1 |
| O00116 | AGPS | DETRTIIsLDTsQMN | S257 | 0.016 | 98.94 | 658 | 0.012719 | 0.016 | 0.0127 | 1 |
| O00116 | AGPS | TIIsLDTsQMNRILW | S261 | 0.032 | 98.73 | 658 | 0.025437 | 0.032 | 0.0254 | 1 |
| O00116 | AGPS | SGMKkNIyGNIEDLV | Y326 | 0.242 | 98.72 | 658 | 0.106162 | 0.242 | 0.1062 | 0.8462 |
| Q6PIJ6 | FBXO38 | TMTNCGItDLVLKDC | T938 | 0.012 | 93.84 | 1188 | 0.008196 | 0.012 | 0.0082 | 1 |
| Q9H4G1 | CST9L | VESKTVFsMELLLGR | S88 | 0.092 | 97.85 | 147 | 0.073132 | 0.092 | 0.0731 | 1 |
| P51451 | BLK | VVTkEPIyIVTEyMA | Y309 | 0.248 | 92.61 | 505 | 0.108794 | 0.248 | 0.1088 | 1 |
| P51451 | BLK | QIAEGMAyIERMNsI | Y350 | 0.296 | 95.67 | 505 | 0.129851 | 0.296 | 0.1299 | 1 |
| Q8N6G6 | ADAMTSL1 | PDHLYLEtKTLQGTK | T229 | 0 | 87.37 | 1762 | 0 | 0 | 0 | 1 |
| P54368 | OAZ1 | GGALPEGsKDSFAVL | S152 | 0.038 | 90.26 | 228 | 0.030207 | 0.084 | 0.0668 | 1 |
| P17480 | UBTF | TkILsKKykELPEKK | Y150 | 0.333 | 89.4 | 764 | 0.146082 | 0.333 | 0.1461 | 1 |
| Q9UHD2 | TBK1 | VIGEDGQsVykLTDF | S151 | 0.047 | 98.42 | 729 | 0.037361 | 0.047 | 0.0374 | 1 |
| Q9UHD2 | TBK1 | GEDGQsVykLTDFGA | Y153 | 0.021 | 98.75 | 729 | 0.009212 | 0.021 | 0.0092 | 1 |
| Q9UHD2 | TBK1 | GkPSGAIsGVQkAEN | S247 | 0.074 | 98.18 | 729 | 0.058824 | 0.074 | 0.0588 | 1 |
| Q9UHD2 | TBK1 | SSNQELIyEGRRLVL | Y354 | 0.34 | 98.64 | 729 | 0.149153 | 0.34 | 0.1492 | 1 |
| P37231 | PPARG | CKGFFRRtIRLKLIY | T166 | 0.036 | 98.04 | 505 | 0.024588 | 0.036 | 0.0246 | 1 |
| P12259 | F5 | GIRYSKLsEGAsyLD | S123 | 0.048 | 89.31 | 2224 | 0.038156 | 0.048 | 0.0382 | 1 |
| Q9BRR6 | ADPGK | AGARVAGtQACAtET | T412 | 0.158 | 97.38 | 497 | 0.107916 | 0.158 | 0.1079 | 1 |
| Q504Y2 | PKDCC | MNEKRNLyNAYRFFF | Y341 | 0.121 | 98.21 | 493 | 0.053081 | 0.121 | 0.0531 | 1 |
| Q8IUW3 | SPATA2L | ksFQkMGyVRRDSHR | Y129 | 0.05 | 97.69 | 424 | 0.021934 | 0.05 | 0.0219 | 0.9091 |
| Q8IUW3 | SPATA2L | LLQARRAsGDVASCV | S189 | 0.138 | 92.99 | 424 | 0.109698 | 0.138 | 0.1097 | 1 |
| Q7Z5M8 | ABHD12B | PELKIPHtVNFYLRV | T96 | 0.06 | 98.33 | 362 | 0.040981 | 0.06 | 0.041 | 0.8889 |
| P18124 | RPL7 | FVkLNkAsINMLrIV | S129 | 0.133 | 97.21 | 248 | 0.105724 | 0.133 | 0.1057 | 1 |
| P18124 | RPL7 | WGYPNLksVNELIyk | S149 | 0.017 | 97.65 | 248 | 0.013514 | 0.017 | 0.0135 | 1 |
| Q8NEB9 | PIK3C3 | DGSEPTktPGRtssT | T159 | 0.018 | 92.78 | 887 | 0.012294 | 0.018 | 0.0123 | 1 |

| | | | | | | | | | | |
|---|---|---|---|---|---|---|---|---|---|---|
| Q5JUQ0 | FAM78A | SSSVVLRyRTPHFRA | Y64 | 0.008 | 98.2 | 283 | 0.003509 | 0.008 | 0.0035 | 1 |
| Q9UKU7 | ACAD8 | GPGPkGIsCIVVEkG | S216 | 0 | 98.89 | 415 | 0 | 0 | 0 | 0.9167 |
| Q4W4Y0 | C14orf28 | NKEEHHysAAFQIGG | S267 | 0.026 | 88.91 | 310 | 0.020668 | 0.044 | 0.035 | 1 |
| P05141 | SLC25A5 | GGVAAAIsktAVAPI | S22 | 0.17 | 89.41 | 298 | 0.135135 | 0.17 | 0.1351 | 1 |
| P05141 | SLC25A5 | PkEQGVLsFWRGNLA | S69 | 0.093 | 95.34 | 298 | 0.073927 | 0.18 | 0.1431 | 1 |
| P05141 | SLC25A5 | LTSYPFDtVRRRMMM | T233 | 0.004 | 98.3 | 298 | 0.002732 | 0.004 | 0.0027 | 1 |
| P05141 | SLC25A5 | AFFkGAWsNVLRGMG | S276 | 0.006 | 93.63 | 298 | 0.004769 | 0.006 | 0.0048 | 0.875 |
| Q9H1A4 | ANAPC1 | DPVHNRVtLELsNGS | T596 | 0.004 | 92.24 | 1944 | 0.002732 | 0.116 | 0.0817 | 1 |
| Q8TCU6 | PREX1 | PLRLLVAtKAKEIIK | T704 | 0.091 | 94.33 | 1659 | 0.062154 | 0.091 | 0.0622 | 1 |
| Q86YR7 | MCF2L2 | PSQRLIKyQMLLKGL | Y762 | 0.005 | 95.62 | 1114 | 0.002193 | 0.09 | 0.0395 | 0.9231 |
| Q6ZV29 | PNPLA7 | QGDKSDCtyIMLSGR | T613 | 0.059 | 90.51 | 1317 | 0.040298 | 0.059 | 0.0403 | 1 |
| Q6ZV29 | PNPLA7 | GDKSDCtyIMLSGRL | Y614 | 0.246 | 91.12 | 1317 | 0.107917 | 0.246 | 0.1079 | 1 |
| Q9BYZ6 | RHOBTB2 | PFRAVLKyLytGELD | Y432 | 0.007 | 95.27 | 727 | 0.003071 | 0.007 | 0.0031 | 0.9286 |
| Q7Z388 | DPY19L4 | QGDSAIyySYYKDML | Y100 | 0.076 | 95.47 | 723 | 0.03334 | 0.076 | 0.0333 | 0.9286 |
| Q7Z6K5 | ARPIN | LIDVSRHsILDTHGR | S48 | 0.089 | 93.71 | 226 | 0.070747 | 0.089 | 0.0707 | 1 |
| P16144 | ITGB4 | ESQPYRYtVKARNGA | T1297 | 0.078 | 93.32 | 1822 | 0.053275 | 0.106 | 0.0846 | 1 |
| P35916 | FLT4 | FVEGKNKtVSKLVIQ | T517 | 0.032 | 91.66 | 1363 | 0.021856 | 0.032 | 0.0219 | 1 |
| Q9UPW8 | UNC13A | KTGsSDPyVTVQVGK | Y700 | 0.13 | 95.01 | 1703 | 0.057029 | 0.237 | 0.0958 | 0.9167 |
| Q5VYK3 | ECPAS | EDRFQEFsNIVIPLI | S1655 | 0.033 | 88.6 | 1845 | 0.026232 | 0.143 | 0.1202 | 1 |
| Q8IZ52 | CHPF | PHFRSALtAHPVRDP | T323 | 0.01 | 96.83 | 775 | 0.00683 | 0.01 | 0.0068 | 1 |
| P84085 | ARF5 | AMPVsELtDkLGLQH | T140 | 0.115 | 98.07 | 180 | 0.078547 | 0.115 | 0.0785 | 0.8333 |
| Q8NBP7 | PCSK9 | EEtHLsQsERtARRL | S91 | 0.006 | 98.13 | 692 | 0.004769 | 0.006 | 0.0048 | 1 |
| Q8NBP7 | PCSK9 | HLsQsERtARRLQAQ | T94 | 0.038 | 98.28 | 692 | 0.025955 | 0.07 | 0.0471 | 1 |
| Q8NBP7 | PCSK9 | THLAGVVsGRDAGVA | S235 | 0 | 95.15 | 692 | 0 | 0 | 0 | 1 |
| Q496J9 | SV2C | VGLCLPDtRTQVLM_ | T721 | 0 | 83.13 | 727 | 0 | 0 | 0 | 1 |
| Q13191 | CBLB | KysTKPGsYIFRLSC | S282 | 0 | 98.21 | 982 | 0 | 0 | 0 | 1 |
| Q6PCB7 | SLC27A1 | GMDDRLFyIyTSGTT | Y246 | 0.083 | 97.27 | 646 | 0.036411 | 0.083 | 0.0364 | 1 |
| Q6PCB7 | SLC27A1 | DDRLFyIyTSGTTGL | Y248 | 0.086 | 87.55 | 646 | 0.037727 | 0.086 | 0.0377 | 1 |
| Q6PCB7 | SLC27A1 | AIVVHSRyyRMAAFG | Y266 | 0 | 98 | 646 | 0 | 0 | 0 | 1 |
| Q6PCB7 | SLC27A1 | IVVHSRyyRMAAFGH | Y267 | 0.07 | 97.93 | 646 | 0.030708 | 0.07 | 0.0307 | 1 |
| Q9UF02 | CACNG5 | FNYKyGWsFAFAAIS | S178 | 0.054 | 94.35 | 275 | 0.042925 | 0.054 | 0.0429 | 1 |

| | | | | | | | | | | |
|---|---|---|---|---|---|---|---|---|---|---|
| Q02750 | MAP2K1 | kLCDFGVsGQLIDsM | S212 | 0.048 | 80.47 | 393 | 0.038156 | 0.048 | 0.0382 | 1 |
| O75342 | ALOX12B | ____MATyKVRVAtG | Y4 | 0 | 98.11 | 701 | 0 | 0 | 0 | 1 |
| O75342 | ALOX12B | TyKVRVAtGTDLLSG | T10 | 0.041 | 98.11 | 701 | 0.028004 | 0.063 | 0.043 | 1 |
| O75342 | ALOX12B | GTDLLSGtRDSISLT | T18 | 0.019 | 94.72 | 701 | 0.012977 | 0.019 | 0.013 | 1 |
| Q8IWA0 | WDR75 | SKYLLGAtENGILCC | T560 | 0.027 | 93.99 | 830 | 0.018441 | 0.027 | 0.0184 | 1 |
| Q8IWA0 | WDR75 | PSEPRPLyIQkGISR | Y619 | 0.231 | 89.9 | 830 | 0.101336 | 0.231 | 0.1013 | 1 |
| P52746 | ZNF142 | CPECEyCtNRADALR | T1545 | 0.12 | 81.78 | 1687 | 0.081962 | 0.12 | 0.082 | 1 |
| O75311 | GLRA3 | LMGRTSGyDARIRPN | Y57 | 0.285 | 91.98 | 464 | 0.125025 | 0.285 | 0.125 | 1 |
| Q8IZU8 | DSEL | SNAVSLLsHLWLANt | S1100 | 0 | 97.51 | 1212 | 0 | 0 | 0 | 1 |
| Q8IZU8 | DSEL | sHLWLANtAAALRIN | T1107 | 0.011 | 96.73 | 1212 | 0.007513 | 0.011 | 0.0075 | 0.8182 |
| Q9NQR4 | NIT2 | RAVDNQVyVATAsPA | Y202 | 0.062 | 98.94 | 276 | 0.027199 | 0.121 | 0.052 | 0.8667 |
| Q9NQR4 | NIT2 | QVyVATAsPARDDKA | S207 | 0.003 | 98.91 | 276 | 0.002385 | 0.003 | 0.0024 | 1 |
| Q96D09 | GPRASP2 | LKMLLNLsENPAVAK | S746 | 0 | 93.01 | 838 | 0 | 0 | 0 | 1 |
| Q01970 | PLCB3 | ARLLIEKyEPNQQFL | Y279 | 0.156 | 96.73 | 1234 | 0.068435 | 0.156 | 0.0684 | 1 |
| Q01970 | PLCB3 | SDRKVGIyVEVDMFG | Y746 | 0.157 | 98.17 | 1234 | 0.068874 | 0.157 | 0.0689 | 1 |
| O75792 | RNASEH2A | GyPNDPktKAWLkEH | T216 | 0.004 | 96.34 | 299 | 0.002732 | 0.107 | 0.0731 | 1 |
| P56706 | WNT7B | GQELRVGsREAAFTY | S101 | 0.02 | 96.95 | 349 | 0.015898 | 0.02 | 0.0159 | 0.8889 |
| P56706 | WNT7B | SGSCTTKtCWttLPK | T213 | 0.135 | 93.65 | 349 | 0.092207 | 0.135 | 0.0922 | 1 |
| P56706 | WNT7B | TTKtCWttLPKFREV | T217 | 0.196 | 97.18 | 349 | 0.133871 | 0.196 | 0.1339 | 1 |
| P56706 | WNT7B | GHLLKEKyNAAVQVE | Y232 | 0.289 | 98.4 | 349 | 0.12678 | 0.289 | 0.1268 | 0.9 |
| Q9Y666 | SLC12A7 | HPRLLSFtsQLKAGK | T715 | 0.021 | 94.14 | 1083 | 0.014343 | 0.021 | 0.0143 | 1 |
| Q9Y666 | SLC12A7 | PRLLSFtsQLKAGKG | S716 | 0.113 | 93.83 | 1083 | 0.089825 | 0.113 | 0.0898 | 0.8889 |
| Q9P2R6 | RERE | EFKARVDsFFyILGY | S265 | 0.125 | 94.36 | 1566 | 0.099364 | 0.125 | 0.0994 | 1 |
| P80511 | S100A12 | VRKGHFDtLSKGELK | T27 | 0.174 | 95.05 | 92 | 0.118844 | 0.174 | 0.1188 | 1 |
| Q27J81 | INF2 | NLKkLLEtERkVSAS | T850 | 0.031 | 93.48 | 1249 | 0.021173 | 0.031 | 0.0212 | 1 |
| Q27J81 | INF2 | VAEVQEQytERLQAS | Y865 | 0.038 | 92.99 | 1249 | 0.01667 | 0.038 | 0.0167 | 1 |
| Q5H9U9 | DDX60L | GQFPLsItLVLRLML | T1351 | 0.027 | 90.72 | 1706 | 0.018441 | 0.027 | 0.0184 | 1 |
| P41240 | CSK | AKGSLVDyLRSRGRs | Y277 | 0.203 | 97.76 | 450 | 0.089053 | 0.203 | 0.0891 | 1 |
| Q15024 | EXOSC7 | ENVPCIVtLCkIGYR | T202 | 0.093 | 94.19 | 291 | 0.06352 | 0.093 | 0.0635 | 0.8 |
| O43464 | HTRA2 | MKVtAGIsFAIPSDR | S330 | 0.015 | 92.57 | 458 | 0.011924 | 0.015 | 0.0119 | 1 |
| P27694 | RPA1 | CARVTNKsQIrTWSN | S207 | 0.087 | 88.56 | 616 | 0.069158 | 0.087 | 0.0692 | 0.8571 |

| | | | | | | | | | | |
|---|---|---|---|---|---|---|---|---|---|---|
| P27694 | RPA1 | DFGGRSLsVLSSSTI | S392 | 0.143 | 95.42 | 616 | 0.113673 | 0.143 | 0.1137 | 1 |
| Q9P0K7 | RAI14 | QILCEHKsPINLKDL | S143 | 0.085 | 85.7 | 980 | 0.067568 | 0.085 | 0.0676 | 1 |
| P11217 | PYGM | VDRLRRMsLVEEGAV | S430 | 0.102 | 98.48 | 842 | 0.081081 | 0.102 | 0.0811 | 1 |
| P11217 | PYGM | AHLCIAGsHAVNGVA | S450 | 0 | 98.89 | 842 | 0 | 0 | 0 | 1 |
| P11217 | PYGM | NKFFVPRtVMIGGkA | T603 | 0.016 | 98.81 | 842 | 0.010928 | 0.016 | 0.0109 | 1 |
| Q9UBN4 | TRPC4 | NIYKALAsPSLIALS | S205 | 0.024 | 94.83 | 977 | 0.019078 | 0.024 | 0.0191 | 0.9 |
| Q9BQN1 | FAM83C | LCDLMEAssRRGVPV | S193 | 0.117 | 93.71 | 747 | 0.093005 | 0.117 | 0.093 | 1 |
| Q9HC58 | SLC24A3 | SLVLVLMyLIyIVIM | Y248 | 0.179 | 92.72 | 644 | 0.078525 | 0.179 | 0.0785 | 1 |
| Q9HC58 | SLC24A3 | LVLMyLIyIVIMKYN | Y251 | 0.129 | 88.2 | 644 | 0.05659 | 0.176 | 0.0772 | 1 |
| Q9NUI1 | DECR2 | LSTKVtAsPLQRLGN | S234 | 0.079 | 97.19 | 292 | 0.062798 | 0.079 | 0.0628 | 1 |
| Q16537 | PPP2R5E | LEKDPsLtEPVIRGL | T299 | 0.045 | 97.62 | 467 | 0.030736 | 0.045 | 0.0307 | 1 |
| P49588 | AARS1 | LkLFKMHsPQTsAML | S882 | 0.006 | 93.46 | 968 | 0.004769 | 0.006 | 0.0048 | 1 |
| P49588 | AARS1 | KMHsPQTsAMLFTVD | S886 | 0.006 | 96.31 | 968 | 0.004769 | 0.006 | 0.0048 | 1 |
| P49588 | AARS1 | DNEAGkItCLCQVPQ | T900 | 0.142 | 95.73 | 968 | 0.096988 | 0.142 | 0.097 | 1 |
| Q9HC96 | CAPN10 | CELSPGyyLAVPSTF | Y466 | 0.049 | 95.86 | 672 | 0.021496 | 0.049 | 0.0215 | 1 |
| Q9UPW8 | UNC13A | KKTFLEKsPDLQSLR | S1488 | 0.023 | 89.74 | 1703 | 0.018283 | 0.065 | 0.098 | 1 |
| O60337 | MARCHF6 | NLVHRRIyPFLLMVV | Y850 | 0.122 | 93.93 | 910 | 0.05352 | 0.122 | 0.0535 | 1 |
| O60337 | MARCHF6 | EHIkNDkyLVGQRLV | Y882 | 0.227 | 86.77 | 910 | 0.099582 | 0.227 | 0.0996 | 1 |
| P35590 | TIE1 | GPPTLMttDCPEPLL | T540 | 0.046 | 90.53 | 1138 | 0.031419 | 0.046 | 0.0314 | 1 |
| Q15054 | POLD3 | AKQMLyDyVERKRKE | Y44 | 0.067 | 96.41 | 466 | 0.029392 | 0.067 | 0.0294 | 1 |
| Q14749 | GNMT | NNKAHMVtLDYtVQV | T220 | 0.018 | 94.15 | 295 | 0.012294 | 0.018 | 0.0123 | 1 |
| Q96P20 | NLRP3 | KLVELDLsDNALGDF | S806 | 0.116 | 98.45 | 1036 | 0.09221 | 0.116 | 0.0922 | 1 |
| Q96P20 | NLRP3 | SQSLRkLsLGNNDLG | S975 | 0.163 | 97.59 | 1036 | 0.129571 | 0.163 | 0.1296 | 1 |
| Q8N8A6 | DDX51 | YVHRVGRtARAGkTG | T596 | 0.013 | 89.93 | 666 | 0.008879 | 0.013 | 0.0089 | 0.9091 |
| Q8IVG5 | SAMD9L | IILNCMRsRNPDESA | S842 | 0.073 | 84.5 | 1584 | 0.058029 | 0.073 | 0.058 | 1 |
| P49407 | ARRB1 | yLKERRVyVTLTCAF | Y54 | 0.129 | 95.67 | 418 | 0.05659 | 0.247 | 0.1084 | 1 |
| P49407 | ARRB1 | LFVANVQsFPPAPED | S86 | 0.032 | 92.56 | 418 | 0.025437 | 0.032 | 0.0254 | 1 |
| O43422 | THAP12 | LKALkCLsLVPsVMG | S606 | 0.069 | 92.79 | 761 | 0.054849 | 0.145 | 0.1153 | 1 |
| P61247 | RPS3A | FRkFkLItEDVQGkN | T88 | 0.037 | 94.62 | 264 | 0.025272 | 0.037 | 0.0253 | 0.9091 |
| Q9UBV8 | PEF1 | YDRDRSGsIsYTELQ | S200 | 0.168 | 96.2 | 284 | 0.133546 | 0.168 | 0.1335 | 1 |
| Q9NZ45 | CISD1 | VyCRCWRsKkFPFCD | S77 | 0.029 | 98.26 | 108 | 0.023053 | 0.029 | 0.0231 | 1 |

| | | | | | | | | | | |
|---|---|---|---|---|---|---|---|---|---|---|
| O15245 | SLC22A1 | SGVLMAFsPNYMSML | S196 | 0.009 | 94.52 | 554 | 0.007154 | 0.009 | 0.0072 | 1 |
| Q8IUC4 | RHPN2 | AMAHHEEsVREASLC | S423 | 0.004 | 98.3 | 686 | 0.00318 | 0.004 | 0.0032 | 1 |
| Q16134 | ETFDH | GSVKGIAtNDVGIQk | T216 | 0.06 | 98.46 | 617 | 0.040981 | 0.06 | 0.041 | 0.8 |
| Q16134 | ETFDH | IGLHVtEyEDNLKNS | Y435 | 0 | 98.59 | 617 | 0 | 0 | 0 | 0.8889 |
| P04424 | ASL | LATDLAYyLVRKGMP | Y376 | 0.196 | 98.49 | 464 | 0.085982 | 0.354 | 0.1418 | 1 |
| P04424 | ASL | LQELQTIsPLFSGDV | S417 | 0.038 | 97.8 | 464 | 0.030207 | 0.038 | 0.0302 | 1 |
| P54578 | USP14 | QEEItkQsPTLQRNA | S302 | 0.022 | 84.97 | 494 | 0.017488 | 0.022 | 0.0175 | 1 |
| P54578 | USP14 | SFADDIGsNNCGyYD | S412 | 0.014 | 90.45 | 494 | 0.011129 | 0.014 | 0.0111 | 1 |
| Q15084 | PDIA6 | kGEsPVDyDGGRTRS | Y252 | 0.281 | 96.13 | 440 | 0.123271 | 0.281 | 0.1233 | 1 |
| Q9NU22 | MDN1 | HDTLVRWyTANCLAL | Y186 | 0.086 | 90.1 | 5596 | 0.037727 | 0.086 | 0.0377 | 0.8 |
| P40197 | GP5 | LGKLRQVsLRRNRLR | S369 | 0.06 | 98.43 | 560 | 0.047695 | 0.06 | 0.0477 | 1 |
| P33992 | MCM5 | RVTIMGIySIKkFGL | Y269 | 0.158 | 91.42 | 734 | 0.069312 | 0.158 | 0.0693 | 1 |
| O14841 | OPLAH | IASIRMGttVATNAL | T81 | 0.046 | 82.07 | 1288 | 0.031419 | 0.046 | 0.0314 | 0.875 |
| O14841 | OPLAH | PGENQPLsPEASRKA | S419 | 0.109 | 93.74 | 1288 | 0.086646 | 0.172 | 0.1419 | 1 |
| Q13085 | ACACA | NDPSVMRsPSAGKLI | S756 | 0.039 | 82.69 | 2346 | 0.031002 | 0.039 | 0.031 | 1 |
| Q14790 | CASP8 | MkSKPRGyCLIINNH | Y235 | 0.275 | 98.68 | 479 | 0.120639 | 0.275 | 0.1206 | 1 |
| O14787 | TNPO2 | HMHSIIQyMLQRTQD | Y258 | 0.261 | 97.85 | 897 | 0.114497 | 0.261 | 0.1145 | 1 |
| Q9NTJ3 | SMC4 | EKIRQAFyFALRDTL | Y709 | 0.139 | 91.35 | 1288 | 0.060977 | 0.139 | 0.061 | 0.9 |
| Q9NTJ3 | SMC4 | DRRWRVVtLQGQIIE | T739 | 0.004 | 92.06 | 1288 | 0.002732 | 0.004 | 0.0027 | 1 |
| P29317 | EPHA2 | EVPVAIktLkAGYTE | T647 | 0.104 | 90.58 | 976 | 0.071033 | 0.118 | 0.1026 | 1 |
| P62736 | ACTA2 | APrAVFPsIVGRPRH | S35 | 0.019 | 97.72 | 377 | 0.015103 | 0.019 | 0.0151 | 1 |
| P62736 | ACTA2 | GMGQkDsyVGDEAQs | Y55 | 0.039 | 95.97 | 377 | 0.017109 | 0.039 | 0.0171 | 1 |
| P62736 | ACTA2 | DLAGRDLtDyLMkIL | T188 | 0.017 | 98.24 | 377 | 0.011611 | 0.017 | 0.0116 | 0.9 |
| P62736 | ACTA2 | ERFRCPEtLFQPsFI | T262 | 0.006 | 98.31 | 377 | 0.004098 | 0.006 | 0.0041 | 0.8889 |
| P62736 | ACTA2 | MWISKQEyDEAGPsI | Y364 | 0.122 | 98.31 | 377 | 0.05352 | 0.205 | 0.0899 | 1 |
| Q9UPZ9 | CILK1 | GsVLLGRsIESGELI | S23 | 0.109 | 96.77 | 632 | 0.086646 | 0.109 | 0.0866 | 1 |
| Q9UPZ9 | CILK1 | MNLREVKsLKkLNHA | S53 | 0 | 80.59 | 632 | 0 | 0 | 0 | 1 |
| Q5EB52 | MEST | LKDGGVLsPILTRLM | S201 | 0.023 | 82.24 | 335 | 0.018283 | 0.023 | 0.0183 | 1 |
| P63244 | RACK1 | WVTQIATtPQFPDMI | T24 | 0.057 | 94.17 | 317 | 0.038932 | 0.105 | 0.0707 | 1 |
| P63244 | RACK1 | LSASRDktIIMWkLt | T39 | 0.161 | 97.6 | 317 | 0.109965 | 0.161 | 0.11 | 0.8333 |
| P63244 | RACK1 | LCAATGPsIkIWDLE | S255 | 0.118 | 94.67 | 317 | 0.0938 | 0.118 | 0.0938 | 1 |

| | | | | | | | | | | |
|---|---|---|---|---|---|---|---|---|---|---|
| P54252 | ATXN3 | GGVTsEDyRtFLQQP | Y58 | 0.192 | 87.11 | 361 | 0.084228 | 0.192 | 0.0842 | 1 |
| Q9NWU1 | OXSM | FSRARALstNsDPkL | S255 | 0 | 98.83 | 459 | 0 | 0 | 0 | 0.9091 |
| Q15052 | ARHGEF6 | CEEKEERyLMLFSNV | Y470 | 0.318 | 95.29 | 776 | 0.139502 | 0.318 | 0.1395 | 1 |
| Q15052 | ARHGEF6 | PRMsGFIyQGkIPIA | Y492 | 0.228 | 94.35 | 776 | 0.10002 | 0.228 | 0.1 | 1 |
| A8K7I4 | CLCA1 | ssQTLTLtVtSRASN | T578 | 0.023 | 97.92 | 914 | 0.015709 | 0.023 | 0.0157 | 1 |
| A8K7I4 | CLCA1 | QTLTLtVtSRASNAT | T580 | 0.04 | 97.25 | 914 | 0.027321 | 0.04 | 0.0273 | 1 |
| A8K7I4 | CLCA1 | YSRyFTTyDTNGRyS | Y665 | 0.138 | 94.05 | 914 | 0.060539 | 0.138 | 0.0605 | 0.8571 |
| A8K7I4 | CLCA1 | TyDTNGRySVKVRAL | Y671 | 0.03 | 95.76 | 914 | 0.013161 | 0.03 | 0.0132 | 1 |
| Q86XI6 | PPP1R3B | RMEFAVyyECNGQTY | Y214 | 0 | 97.78 | 285 | 0 | 0 | 0 | 1 |
| P51888 | PRELP | LPGLVFLyMEKNQLE | Y156 | 0.157 | 98.74 | 382 | 0.068874 | 0.157 | 0.0689 | 1 |
| Q9H0B8 | CRISPLD2 | tQVVRCDtKMKDRCK | T292 | 0.084 | 94.6 | 497 | 0.057373 | 0.084 | 0.0574 | 1 |
| Q9H0B8 | CRISPLD2 | SERHGVQsLSKyKPs | S369 | 0.069 | 97.14 | 497 | 0.054849 | 0.069 | 0.0548 | 1 |
| Q9H0B8 | CRISPLD2 | sLSKyKPsSSFMVsK | S376 | 0.184 | 91.64 | 497 | 0.146264 | 0.184 | 0.1463 | 1 |
| Q9H0B8 | CRISPLD2 | PVDKKKTyVGSLRNG | Y468 | 0.065 | 90.91 | 497 | 0.028515 | 0.158 | 0.0693 | 1 |
| Q6L8Q7 | PDE12 | VGVPsSLsPssPsss | S217 | 0.169 | 76.01 | 609 | 0.134341 | 0.169 | 0.1343 | 1 |
| Q6L8Q7 | PDE12 | PsSLsPssPsssWTE | S220 | 0.158 | 85.64 | 609 | 0.125597 | 0.158 | 0.1256 | 1 |
| Q6L8Q7 | PDE12 | sPssPsssWTETDVE | S224 | 0.126 | 94.06 | 609 | 0.100159 | 0.126 | 0.1002 | 1 |
| P51116 | FXR2 | RGsNGAFyKGFVKDV | Y26 | 0.108 | 93.69 | 673 | 0.047378 | 0.108 | 0.0474 | 1 |
| P51116 | FXR2 | VPFIFVGtRENISNA | T350 | 0.161 | 96.04 | 673 | 0.109965 | 0.161 | 0.11 | 1 |
| Q86TU7 | SETD3 | DSFTYEDyRWAVSSV | Y244 | 0.006 | 98.84 | 594 | 0.002632 | 0.006 | 0.0026 | 1 |
| Q9UL18 | AGO1 | HPsRYCAtVRVQRPR | T626 | 0.069 | 97.84 | 857 | 0.047128 | 0.069 | 0.0471 | 1 |
| P98073 | TMPRSS15 | IDEIFLssNLKNEYK | S100 | 0.127 | 86.7 | 1019 | 0.100954 | 0.127 | 0.101 | 1 |
| Q07973 | CYP24A1 | ESICLVLyEKRFGLL | Y220 | 0.124 | 94.31 | 514 | 0.054397 | 0.124 | 0.0544 | 0.8462 |
| Q07973 | CYP24A1 | LKESMRLtPSVPFTt | T388 | 0.137 | 95.87 | 514 | 0.093573 | 0.137 | 0.0936 | 0.875 |
| Q07973 | CYP24A1 | kATVLGEyALPKGTV | Y407 | 0.313 | 96.19 | 514 | 0.137309 | 0.313 | 0.1373 | 1 |
| A5D8V7 | ODAD3 | KVKDAtGtDETHSLV | T368 | 0.09 | 71.82 | 595 | 0.061471 | 0.09 | 0.0615 | 1 |
| Q9H0U4 | RAB1B | IkLQIWDtAGQERFR | T64 | 0.012 | 85.2 | 201 | 0.008196 | 0.012 | 0.0082 | 0.8 |
| Q9NXZ2 | DDX43 | KVRILIAtDLASRGL | T544 | 0.149 | 91.08 | 648 | 0.101769 | 0.149 | 0.1018 | 1 |
| Q9NXZ2 | DDX43 | GVSITTLtRNDWRVA | T592 | 0.038 | 93.55 | 648 | 0.025955 | 0.038 | 0.026 | 0.8 |
| P10809 | HSPD1 | sWGsPkVtkDGVtVA | T74 | 0.015 | 91.8 | 573 | 0.010245 | 0.03 | 0.0205 | 1 |
| P10809 | HSPD1 | sIDLkDKykNIGAkL | Y90 | 0.091 | 94.91 | 573 | 0.03992 | 0.091 | 0.0399 | 1 |

| | | | | | | | | | | |
|---|---|---|---|---|---|---|---|---|---|---|
| P10809 | HSPD1 | VQDVANNtNEEAGDG | T105 | 0 | 91.86 | 573 | 0 | 0 | 0 | 1 |
| P10809 | HSPD1 | NEEAGDGtttATVLA | T113 | 0.088 | 93.54 | 573 | 0.060105 | 0.123 | 0.0781 | 1 |
| P10809 | HSPD1 | EEAGDGtttATVLAR | T114 | 0.172 | 95.75 | 573 | 0.117478 | 0.172 | 0.1175 | 1 |
| P10809 | HSPD1 | kEIGNIIsDAMkkVG | S187 | 0.053 | 93.45 | 573 | 0.04213 | 0.053 | 0.0421 | 1 |
| P10809 | HSPD1 | kCEFQDAyVLLsEkk | Y243 | 0.112 | 97.14 | 573 | 0.049133 | 0.112 | 0.0491 | 1 |
| P10809 | HSPD1 | QDAyVLLsEkkIssI | S247 | 0.025 | 94.35 | 573 | 0.019873 | 0.025 | 0.0199 | 1 |
| P10809 | HSPD1 | VtDALNAtRAAVEEG | T428 | 0.019 | 93.1 | 573 | 0.012977 | 0.019 | 0.013 | 1 |
| P10809 | HSPD1 | GIEIIkRtLkIPAMt | T471 | 0.003 | 97.96 | 573 | 0.002049 | 0.003 | 0.002 | 1 |
| P10809 | HSPD1 | EkGIIDPtkVVRtAL | T522 | 0.026 | 97.47 | 573 | 0.017758 | 0.026 | 0.0178 | 1 |
| P10809 | HSPD1 | DPtkVVRtALLDAAG | T527 | 0.006 | 97.05 | 573 | 0.004098 | 0.006 | 0.0041 | 1 |
| P10809 | HSPD1 | AGVASLLtTAEVVVT | T540 | 0.085 | 94.86 | 573 | 0.058056 | 0.079 | 0.0762 | 1 |
| Q96A11 | GAL3ST3 | LMPPSTVyVTILREP | Y147 | 0.236 | 98.82 | 431 | 0.10353 | 0.236 | 0.1035 | 0.8182 |
| Q96H79 | ZC3HAV1L | HQLIHAAsLKLLQDQ | S218 | 0.029 | 88 | 300 | 0.023053 | 0.029 | 0.0231 | 1 |
| Q86XF0 | DHFR2 | SEIDLEKykLLPEyP | Y157 | 0.099 | 96.4 | 187 | 0.04343 | 0.099 | 0.0434 | 0.9231 |
| P26012 | ITGB8 | CHLKNNVyVKSTTME | Y314 | 0.102 | 97.49 | 769 | 0.044746 | 0.102 | 0.0447 | 1 |
| P26012 | ITGB8 | HPSLGQLsEKLIDNN | S329 | 0.148 | 94.74 | 769 | 0.117647 | 0.148 | 0.1176 | 0.8571 |
| P11498 | PC | kLAKQVGyENAGtVE | Y304 | 0.04 | 96.13 | 1178 | 0.017547 | 0.04 | 0.0175 | 1 |
| P11498 | PC | VGyENAGtVEFLVDR | T309 | 0.002 | 97.77 | 1178 | 0.001366 | 0.002 | 0.0014 | 1 |
| P11498 | PC | VMVNGPttPIPVKAS | T514 | 0.047 | 86.77 | 1178 | 0.032102 | 0.047 | 0.0321 | 1 |
| P11498 | PC | TRGTPLDtEVPMERV | T824 | 0.104 | 96.23 | 1178 | 0.071033 | 0.104 | 0.071 | 1 |
| P11498 | PC | RVFDYsEyWEGARGL | Y837 | 0.005 | 98.21 | 1178 | 0.002193 | 0.005 | 0.0022 | 1 |
| P11498 | PC | FKEVKkAyVEANQML | Y894 | 0 | 96.27 | 1178 | 0 | 0 | 0 | 1 |
| P11498 | PC | VtPEDVLsAAMYPDV | S1009 | 0.022 | 96.03 | 1178 | 0.017488 | 0.044 | 0.035 | 1 |
| Q96N46 | TTC14 | MVLICLGsGIMRDIA | S150 | 0.061 | 84.18 | 770 | 0.04849 | 0.169 | 0.1343 | 1 |
| P98187 | CYP4F8 | YIKIFSKsANIMHAK | S170 | 0.003 | 97.54 | 520 | 0.002385 | 0.003 | 0.0024 | 1 |
| Q5U5Z8 | AGBL2 | MESTFGGsTLGNKRD | S636 | 0 | 96.8 | 902 | 0 | 0 | 0 | 0.9167 |
| O14983 | ATP2A1 | VETLGCtsVICSDKT | S346 | 0.161 | 96.8 | 1001 | 0.127981 | 0.161 | 0.128 | 1 |
| O14983 | ATP2A1 | DRkSMSVyCSPAKSS | Y497 | 0.105 | 96.58 | 1001 | 0.046062 | 0.105 | 0.0461 | 1 |
| O14983 | ATP2A1 | GIRVIMItGDNKGTA | T625 | 0.044 | 84.76 | 1001 | 0.030053 | 0.044 | 0.0301 | 1 |
| O14983 | ATP2A1 | tAVAKTAsEMVLADD | S731 | 0.036 | 91.73 | 1001 | 0.028617 | 0.036 | 0.0286 | 1 |
| Q9H981 | ACTR8 | yVNPLDCyNIHWPIR | Y180 | 0.02 | 98.6 | 624 | 0.008774 | 0.02 | 0.0088 | 1 |

| | | | | | | | | | | |
|---|---|---|---|---|---|---|---|---|---|---|
| P14735 | IDE | YYLRLLMtEVAWtkD | T691 | 0.144 | 98.31 | 1019 | 0.098354 | 0.144 | 0.0984 | 0.8333 |
| P14735 | IDE | APRRHKVsVHVLARE | S955 | 0 | 98.6 | 1019 | 0 | 0 | 0 | 1 |
| Q9Y693 | LHFPL6 | LYPLGWDsEEVRQTC | S145 | 0.103 | 95.16 | 200 | 0.081876 | 0.103 | 0.0819 | 1 |
| P55263 | ADK | GNETEAAtFAREQGF | T246 | 0.103 | 98.22 | 362 | 0.07035 | 0.103 | 0.0703 | 1 |
| Q96BH3 | ELSPBP1 | SNKLWCPttENMDKD | T155 | 0.105 | 96.09 | 223 | 0.071716 | 0.105 | 0.0717 | 1 |
| O95807 | TMEM50A | MWILFGGyVAKEKDI | Y119 | 0.017 | 87.08 | 157 | 0.007458 | 0.017 | 0.0075 | 1 |
| P51788 | CLCN2 | LRLALHRtKGRMLAL | T607 | 0.102 | 77.98 | 898 | 0.069667 | 0.102 | 0.0697 | 1 |
| P51788 | CLCN2 | FQLVERtsLHKTHTI | S802 | 0.174 | 88.9 | 898 | 0.138315 | 0.174 | 0.1383 | 1 |
| Q9H6S3 | EPS8L2 | SQPLTYEsGPDEVRA | S628 | 0.044 | 83.54 | 715 | 0.034976 | 0.044 | 0.035 | 1 |
| P00751 | CFB | TRKVGsQyRLEDSVT | Y181 | 0.331 | 95.05 | 764 | 0.145205 | 0.331 | 0.1452 | 1 |
| Q8N0V5 | GCNT2 | GQDFPLktNREIVQy | T199 | 0.022 | 98.68 | 402 | 0.015026 | 0.022 | 0.015 | 0.8571 |
| Q13972 | RASGRF1 | DIsNLIAsEIIRNED | S1095 | 0 | 95.8 | 1273 | 0 | 0 | 0 | 1 |
| P23142 | FBLN1 | KRyMDGMtVGVVRQV | T656 | 0.023 | 83.89 | 703 | 0.015709 | 0.105 | 0.0717 | 0.8333 |
| Q8TC27 | ADAM32 | LRPHDIAyLLIYMDy | Y275 | 0.011 | 97.13 | 787 | 0.004826 | 0.011 | 0.0048 | 1 |
| Q8TC27 | ADAM32 | TMCITRYsAGVALYP | S300 | 0.014 | 90.22 | 787 | 0.011129 | 0.014 | 0.0111 | 1 |
| Q9HCE1 | MOV10 | DQNPRLLtLEVPGVT | T378 | 0.169 | 93.44 | 1003 | 0.115429 | 0.169 | 0.1154 | 1 |
| Q9HCE1 | MOV10 | LDIPNQLyyEGELQA | Y746 | 0.179 | 96.25 | 1003 | 0.078525 | 0.179 | 0.0785 | 0.9091 |
| O75955 | FLOT1 | YkDRQkFsEQVFKVA | S128 | 0.162 | 94.6 | 427 | 0.128776 | 0.162 | 0.1288 | 1 |
| O75955 | FLOT1 | MGISVVSyTLkDIHD | Y149 | 0.162 | 96.47 | 427 | 0.071067 | 0.162 | 0.0711 | 1 |
| Q9H967 | WDR76 | RRLNSRRsQPLIsLt | S485 | 0.008 | 88.71 | 626 | 0.006359 | 0.008 | 0.0064 | 1 |
| Q99732 | LITAF | SCNKMIVsQLSYNAG | S105 | 0.049 | 94.62 | 161 | 0.038951 | 0.049 | 0.039 | 1 |
| Q86VD1 | MORC1 | QAGMFIYsNNRLIKM | S372 | 0.061 | 94.56 | 984 | 0.04849 | 0.061 | 0.0485 | 1 |
| Q8IYR0 | CFAP206 | ESRLGsVtREITDNR | T113 | 0.214 | 95.89 | 622 | 0.146165 | 0.214 | 0.1462 | 1 |
| Q8IYR0 | CFAP206 | KEELESLyRKIISYV | Y131 | 0.043 | 95.67 | 622 | 0.018863 | 0.043 | 0.0189 | 1 |
| Q96A33 | CCDC47 | NDEDMDTyVFAVGTR | Y259 | 0 | 96.03 | 483 | 0 | 0 | 0 | 1 |
| P09543 | CNP | RGAFSEEykRLDEDL | Y100 | 0.18 | 95.65 | 421 | 0.078963 | 0.18 | 0.079 | 1 |
| P09543 | CNP | TISALFVtPKTTGAR | T292 | 0.051 | 97.2 | 421 | 0.034834 | 0.051 | 0.0348 | 1 |
| Q14315 | FLNC | NDTFTVKytPPGAGR | Y833 | 0.069 | 94.05 | 2725 | 0.030269 | 0.103 | 0.0452 | 1 |
| Q14315 | FLNC | IIDNHDYsYTVKYTA | S927 | 0.075 | 90.96 | 2725 | 0.059619 | 0.053 | 0.0762 | 1 |
| Q8TDB6 | DTX3L | GGPEMyGyPDPSYLk | Y721 | 0.017 | 96.81 | 740 | 0.007458 | 0.017 | 0.0075 | 1 |
| Q9Y3A3 | MOB4 | TFLCHRFtKFVMKyN | T192 | 0.021 | 97.18 | 225 | 0.014343 | 0.021 | 0.0143 | 1 |

| | | | | | | | | | | |
|---|---|---|---|---|---|---|---|---|---|---|
| Q99832 | CCT7 | VDGrGkAtIsNDGAt | T57 | 0.157 | 89.7 | 543 | 0.107233 | 0.205 | 0.14 | 1 |
| Q99832 | CCT7 | GrGkAtIsNDGAtIL | S59 | 0.013 | 89.22 | 543 | 0.010334 | 0.013 | 0.0103 | 1 |
| Q99832 | CCT7 | IRAFRTAtQLAVNkI | T129 | 0.109 | 97.45 | 543 | 0.074448 | 0.172 | 0.1175 | 1 |
| Q99832 | CCT7 | LPIGDVAtQyFADrD | T300 | 0.046 | 93.39 | 543 | 0.031419 | 0.09 | 0.0615 | 1 |
| Q99832 | CCT7 | IGDVAtQyFADrDMF | Y302 | 0.174 | 94.11 | 543 | 0.076331 | 0.34 | 0.1472 | 1 |
| Q99832 | CCT7 | PEEDLkRtMMACGGS | T322 | 0.002 | 92.8 | 543 | 0.001366 | 0.002 | 0.0014 | 1 |
| Q99832 | CCT7 | CPkAkTCtFILrGGA | T371 | 0.008 | 94.11 | 543 | 0.005464 | 0.008 | 0.0055 | 1 |
| Q99832 | CCT7 | INALtAAsEAACLIV | S507 | 0.042 | 96.45 | 543 | 0.033386 | 0.071 | 0.0564 | 1 |
| Q9Y673 | ALG5 | TSkVAFkyCQKyGsD | Y123 | 0.23 | 98.58 | 324 | 0.100898 | 0.23 | 0.1009 | 1 |
| Q9Y673 | ALG5 | CVKGIRDtQCGFKLF | T234 | 0.073 | 93.51 | 324 | 0.04986 | 0.073 | 0.0499 | 1 |
| Q96IJ6 | GMPPA | AGSALyAsRLYLSRY | S258 | 0.019 | 97.67 | 420 | 0.015103 | 0.019 | 0.0151 | 1 |
| Q96IJ6 | GMPPA | AVLGPNVsIGkGVtV | S306 | 0.045 | 98.72 | 420 | 0.035771 | 0.045 | 0.0358 | 1 |
| P49326 | FMO5 | EEGRAsIyKsVIINT | Y56 | 0.21 | 98.57 | 533 | 0.092124 | 0.21 | 0.0921 | 0.8462 |
| P49326 | FMO5 | QyFHSRDyKNPEGFT | Y177 | 0.016 | 98.67 | 533 | 0.007019 | 0.016 | 0.007 | 1 |
| P49326 | FMO5 | RALsQHPtLNDDLPN | T284 | 0.016 | 95.36 | 533 | 0.010928 | 0.016 | 0.0109 | 1 |
| Q8WUA8 | TSKU | PAESFTSsPLSDVNL | S131 | 0.001 | 90.72 | 353 | 0.000795 | 0.001 | 0.0008 | 1 |
| O43929 | ORC4 | QVQYKHLsELLkRtA | S48 | 0.095 | 93.54 | 436 | 0.075517 | 0.095 | 0.0755 | 1 |
| O43929 | ORC4 | LsELLkRtALHGESN | T54 | 0.018 | 89.24 | 436 | 0.012294 | 0.018 | 0.0123 | 1 |
| Q5FYB1 | ARSI | PREYLYRyRTMGNVA | Y252 | 0.063 | 95.29 | 569 | 0.027637 | 0.063 | 0.0276 | 1 |
| Q5FYB1 | ARSI | GNVARRKyAAMVTCM | Y263 | 0.028 | 97.29 | 569 | 0.012283 | 0.028 | 0.0123 | 1 |
| Q5FYB1 | ARSI | SSDNGGQtFsGGsNW | T302 | 0.184 | 91.77 | 569 | 0.125675 | 0.184 | 0.1257 | 0.875 |
| Q5FYB1 | ARSI | GQtFsGGsNWPLRGR | S307 | 0.031 | 96.91 | 569 | 0.024642 | 0.031 | 0.0246 | 1 |
| P49458 | SRP9 | GNLCVkVtDDLVCLV | T43 | 0.115 | 97.98 | 86 | 0.078547 | 0.115 | 0.0785 | 1 |
| Q9NP91 | SLC6A20 | SIYGFKAtFNyENCL | T303 | 0.084 | 95.6 | 592 | 0.057373 | 0.084 | 0.0574 | 0.8889 |
| Q9BRK5 | SDF4 | ELkVDEEtQEVLENL | T193 | 0.128 | 92.25 | 362 | 0.087426 | 0.201 | 0.1311 | 1 |
| O15091 | PRORP | KtLEPRGysLLIRGL | Y213 | 0.252 | 96.06 | 583 | 0.110549 | 0.252 | 0.1105 | 1 |
| Q9NRD0 | FBXO8 | NPDEGVNyFMSKGIL | Y156 | 0.338 | 97.62 | 319 | 0.148276 | 0.338 | 0.1483 | 1 |
| Q8WTX7 | CASTOR1 | EHRVRVLsVARPGLW | S14 | 0.092 | 98.51 | 329 | 0.073132 | 0.092 | 0.0731 | 0.875 |
| Q8WTX7 | CASTOR1 | ARPGLWLyTHPLIKL | Y23 | 0 | 97.17 | 329 | 0 | 0 | 0 | 1 |
| P51451 | BLK | CLDEGGyyISPRITF | Y188 | 0.324 | 91.62 | 505 | 0.142134 | 0.324 | 0.1421 | 1 |
| P51451 | BLK | LQALVQHySKkGDGL | Y205 | 0.01 | 96.67 | 505 | 0.004387 | 0.01 | 0.0044 | 1 |

| | | | | | | | | | | |
|---|---|---|---|---|---|---|---|---|---|---|
| Q7L5L3 | GDPD3 | RyDRNEItIWAsEKS | T175 | 0 | 98.64 | 318 | 0 | 0 | 0 | 1 |
| Q7L5L3 | GDPD3 | NEItIWAsEKSSVMK | S179 | 0.053 | 96.31 | 318 | 0.04213 | 0.053 | 0.0421 | 1 |
| Q8TAD8 | SNIP1 | kQHAVFQyRLVEytR | Y308 | 0 | 96.98 | 396 | 0 | 0 | 0 | 0.8421 |
| P00533 | EGFR | PFGCLLDyVREHKDN | Y801 | 0.079 | 91.14 | 1210 | 0.034656 | 0.209 | 0.0917 | 1 |
| Q14185 | DOCK1 | KLEDAAtyLSLPsTk | Y567 | 0.064 | 86.35 | 1865 | 0.028076 | 0.098 | 0.043 | 1 |
| Q14185 | DOCK1 | AtyLSLPsTkAELEE | S572 | 0.081 | 89.9 | 1865 | 0.064388 | 0.098 | 0.1007 | 1 |
| O94941 | UBOX5 | ATWGRVPsDPFTGVA | S304 | 0.135 | 93.36 | 541 | 0.107313 | 0.135 | 0.1073 | 1 |
| P11717 | IGF2R | DGMIQLNyrGGTPYN | Y709 | 0.006 | 89.59 | 2491 | 0.002632 | 0.006 | 0.0026 | 1 |
| Q9UK73 | FEM1B | RyLLGYVsQQGGQRS | S40 | 0.087 | 94.75 | 627 | 0.069158 | 0.087 | 0.0692 | 1 |
| Q68D85 | NCR3LG1 | QEDPGTVyQCVVRHA | Y226 | 0.006 | 92.76 | 454 | 0.002632 | 0.006 | 0.0026 | 1 |
| P15259 | PGAM2 | WRLNErHyGGLtGLN | Y92 | 0.299 | 96.62 | 253 | 0.131167 | 0.299 | 0.1312 | 1 |
| P15259 | PGAM2 | ErHyGGLtGLNkAET | T96 | 0.036 | 97.11 | 253 | 0.024588 | 0.036 | 0.0246 | 1 |
| P15259 | PGAM2 | MDEKHPyyNsIsKER | Y133 | 0.295 | 98.1 | 253 | 0.129412 | 0.295 | 0.1294 | 1 |
| Q9NQR1 | KMT5A | AAGEELLyDYGDRsk | Y375 | 0.102 | 97.91 | 393 | 0.044746 | 0.102 | 0.0447 | 1 |
| Q5T5C0 | STXBP5 | QKILAVGtQTGALRL | T76 | 0.028 | 98.11 | 1151 | 0.019124 | 0.054 | 0.0374 | 1 |
| Q5T5C0 | STXBP5 | ITGHADGsVKFWDAS | S459 | 0.001 | 95.36 | 1151 | 0.000795 | 0.001 | 0.0008 | 1 |
| Q96DB5 | RMDN1 | NLLLLGktYLkLHNK | T269 | 0 | 98.56 | 314 | 0 | 0 | 0 | 1 |
| Q86UK0 | ABCA12 | GGMKRKLsISIALIG | S1488 | 0.075 | 88.58 | 2595 | 0.059619 | 0.153 | 0.1394 | 1 |
| Q99758 | ABCA3 | GGFLYFFtYIPYFFV | T385 | 0.131 | 90.52 | 1704 | 0.089475 | 0.178 | 0.1122 | 1 |
| O95391 | SLU7 | PTkLELLykSFKVKK | Y354 | 0.327 | 94.51 | 586 | 0.14345 | 0.327 | 0.1434 | 1 |
| Q99518 | FMO2 | AGVsGLIsLKCCVDE | S17 | 0 | 98.93 | 535 | 0 | 0 | 0 | 1 |
| Q99518 | FMO2 | VFHTRFRsMLRNVLP | S241 | 0.086 | 95.1 | 535 | 0.068363 | 0.086 | 0.0684 | 1 |
| P01303 | NPY | PAEDMARyysALRHy | Y48 | 0.33 | 95.46 | 97 | 0.144766 | 0.33 | 0.1448 | 1 |
| Q9BWT6 | MND1 | RIGTSNyyWAFPSKA | Y71 | 0.184 | 95.04 | 205 | 0.080718 | 0.184 | 0.0807 | 1 |
| Q8N436 | CPXM2 | PSLNPDGyEKAYEGG | Y427 | 0.028 | 97.27 | 756 | 0.012283 | 0.028 | 0.0123 | 0.9333 |
| Q6P2Q9 | PRPF8 | RRFTLWWsPTINRAN | S1539 | 0.064 | 93.6 | 2335 | 0.050875 | 0.064 | 0.0509 | 1 |
| Q9H3S5 | PIGM | RDIRHNFsPyFyMLY | S272 | 0.023 | 94.2 | 423 | 0.018283 | 0.023 | 0.0183 | 1 |
| Q9HCU4 | CELSR2 | FEILFNNyVtNRSSS | Y1033 | 0.231 | 81.475 | 2923 | 0.101336 | 0.231 | 0.1013 | 1 |
| Q13144 | EIF2B5 | NITLPEGsVIsLHPP | S447 | 0.084 | 85.89 | 721 | 0.066773 | 0.084 | 0.0668 | 1 |
| Q13144 | EIF2B5 | LPEGsVIsLHPPDAE | S450 | 0 | 86.03 | 721 | 0 | 0.071 | 0.0564 | 1 |
| O75689 | ADAP1 | MDDRRLMyFKDPLDA | Y284 | 0.302 | 98.04 | 374 | 0.132483 | 0.302 | 0.1325 | 1 |

| | | | | | | | | | | |
|---|---|---|---|---|---|---|---|---|---|---|
| Q96NY7 | CLIC6 | STEDVtVsGRkFLDG | S626 | 0.027 | 89.49 | 704 | 0.021463 | 0.027 | 0.0215 | 1 |
| Q99575 | POP1 | KKNIWLEtHIWHAKR | T204 | 0.027 | 94.2 | 1024 | 0.018441 | 0.027 | 0.0184 | 1 |
| P61586 | RHOA | AFGyMECsAKTkDGV | S160 | 0.011 | 98.8 | 193 | 0.008744 | 0.011 | 0.0087 | 1 |
| A1A5B4 | ANO9 | VVYKYRysPCLKEGN | S634 | 0.027 | 85.66 | 782 | 0.021463 | 0.101 | 0.0803 | 1 |
| Q92614 | MYO18A | LMSRQDQsIILLGSS | S493 | 0 | 97.11 | 2054 | 0 | 0 | 0 | 1 |
| O43707 | ACTN4 | NELDyyDsHNVNtRC | S488 | 0.169 | 92.88 | 911 | 0.134341 | 0.169 | 0.1343 | 1 |
| O43707 | ACTN4 | WDALGsLtHsRREAL | T509 | 0.042 | 87.55 | 911 | 0.028687 | 0.098 | 0.0669 | 1 |
| Q6ZS81 | WDFY4 | LIWRLGPtyLFEEAI | T1237 | 0.006 | 96.51 | 3184 | 0.004098 | 0.006 | 0.0041 | 1 |
| Q6ZS81 | WDFY4 | IWRLGPtyLFEEAIS | Y1238 | 0.065 | 96.96 | 3184 | 0.028515 | 0.065 | 0.0285 | 1 |
| Q6ZS81 | WDFY4 | HLSGSLRtIGAVAVG | T1346 | 0.017 | 94.74 | 3184 | 0.011611 | 0.017 | 0.0116 | 1 |
| Q9BTT6 | LRRC1 | EIyRYARsLEELLLD | S37 | 0.13 | 94.81 | 524 | 0.103339 | 0.13 | 0.1033 | 1 |
| Q6UB35 | MTHFD1L | RSRFQFLyDVQVPIV | Y859 | 0.165 | 98.1 | 978 | 0.072383 | 0.165 | 0.0724 | 1 |
| Q15303 | ERBB4 | MARDPQRyLVIQGDD | Y984 | 0.116 | 94.1 | 1308 | 0.050888 | 0.15 | 0.0658 | 0.875 |
| Q9BTU6 | PI4K2A | ERLVVLDyIIRNTDR | Y302 | 0.079 | 98.71 | 479 | 0.034656 | 0.079 | 0.0347 | 1 |
| Q9BTU6 | PI4K2A | RGQILNLtQALkDNK | T431 | 0.172 | 98.54 | 479 | 0.117478 | 0.172 | 0.1175 | 1 |
| Q9H511 | KLHL31 | SVFNGLVyAAGGRNA | Y422 | 0.106 | 98.76 | 634 | 0.046501 | 0.106 | 0.0465 | 1 |
| Q6ZS30 | NBEAL1 | LDMAVsGsRDGTVII | S2511 | 0 | 95.39 | 2694 | 0 | 0.139 | 0.1173 | 1 |
| Q9NZN1 | IL1RAPL1 | KRLIIVMtPNYVVRR | T478 | 0.054 | 86.79 | 696 | 0.036883 | 0.054 | 0.0369 | 1 |
| P54136 | RARS1 | HVGHLRStIIGESIS | T215 | 0.004 | 98.39 | 660 | 0.002732 | 0.004 | 0.0027 | 1 |
| Q8NCT1 | ARRDC4 | AKIERKGyCNGEAIP | Y206 | 0.034 | 95.29 | 418 | 0.014915 | 0.034 | 0.0149 | 1 |
| Q8NCT1 | ARRDC4 | CCIIRVDysLAVyIH | Y291 | 0.069 | 97.06 | 418 | 0.030269 | 0.069 | 0.0303 | 0.8462 |
| Q15185 | PTGES3 | HCIDPNDskHkRTDR | S64 | 0.141 | 98.18 | 160 | 0.112083 | 0.141 | 0.1121 | 1 |
| Q15185 | PTGES3 | kHkRTDRsILCCLRk | S72 | 0.137 | 97.92 | 160 | 0.108903 | 0.137 | 0.1089 | 1 |
| Q8NHC5 | OR14A16 | VMSFDRytAICHPLH | T122 | 0.121 | 96.03 | 309 | 0.082645 | 0.121 | 0.0826 | 1 |
| Q8NEL9 | DDHD1 | VCVRGGLyEVDVTQG | Y264 | 0.156 | 94.74 | 900 | 0.068435 | 0.247 | 0.1084 | 1 |
| Q9NRB3 | CHST12 | RQQLYKLyEADFVLF | Y396 | 0.031 | 98.47 | 414 | 0.013599 | 0.031 | 0.0136 | 0.8571 |
| P61758 | VBP1 | LDEQYQkykFMELNL | Y60 | 0.014 | 95.86 | 197 | 0.006142 | 0.014 | 0.0061 | 1 |
| Q9NRD9 | DUOX1 | LPNPRDLsNTIsRGP | S79 | 0 | 97.69 | 1551 | 0 | 0 | 0 | 1 |
| Q9NRD9 | DUOX1 | RDLsNTIsRGPAGLA | S83 | 0 | 95.98 | 1551 | 0 | 0 | 0 | 1 |
| Q9NRD9 | DUOX1 | RGPAGLAsLRNRTVL | S91 | 0.045 | 97.56 | 1551 | 0.035771 | 0.086 | 0.0684 | 1 |
| Q9NRD9 | DUOX1 | TRVGIILsRGTAASI | S1086 | 0 | 97.64 | 1551 | 0 | 0 | 0 | 1 |

| | | | | | | | | | | |
|---|---|---|---|---|---|---|---|---|---|---|
| Q13206 | DDX10 | PATLEQNyIVCELQQ | Y292 | 0.288 | 95.95 | 875 | 0.126341 | 0.288 | 0.1263 | 1 |
| Q13206 | DDX10 | CELQQKIsVLYSFLR | S302 | 0.045 | 95.31 | 875 | 0.035771 | 0.045 | 0.0358 | 0.8889 |
| Q5VW22 | AGAP6 | SNGMLTyySSLGDyM | Y293 | 0.126 | 94.44 | 663 | 0.055274 | 0.126 | 0.0553 | 0.8462 |
| Q7Z553 | MDGA2 | TLRIVNVsREMSGMy | S506 | 0.169 | 91.98 | 956 | 0.134341 | 0.169 | 0.1343 | 1 |
| Q7Z553 | MDGA2 | sREMSGMyRCQTSQY | Y513 | 0.083 | 93.87 | 956 | 0.036411 | 0.083 | 0.0364 | 1 |
| Q08211 | DHX9 | sQDMkyPsPFFVFGE | S1040 | 0.003 | 92.77 | 1270 | 0.002385 | 0.003 | 0.0024 | 0.8571 |
| Q08211 | DHX9 | AkGMTLVtPLQLLLF | T1063 | 0.021 | 97.05 | 1270 | 0.014343 | 0.021 | 0.0143 | 1 |
| Q08211 | DHX9 | FASkkVQsDGQIVLV | S1077 | 0.006 | 96.62 | 1270 | 0.004769 | 0.006 | 0.0048 | 1 |
| B7ZBB8 | PPP1R3G | NNAGANYtLRYARPA | T348 | 0.188 | 94.77 | 358 | 0.128407 | 0.188 | 0.1284 | 1 |
| P35346 | SSTR5 | AVVHPLssARWRRPR | S147 | 0.023 | 88.43 | 364 | 0.018283 | 0.023 | 0.0183 | 1 |
| O00482 | NR5A2 | CKGFFKRtVQNNKRY | T113 | 0.03 | 96.38 | 541 | 0.02049 | 0.03 | 0.0205 | 1 |
| Q05707 | COL14A1 | NKIGtDGtQVAMVQF | T1070 | 0.04 | 88.65 | 1796 | 0.027321 | 0.04 | 0.0273 | 1 |
| Q05707 | COL14A1 | QVAMVQFtDDPRTEF | T1078 | 0.036 | 85.26 | 1796 | 0.024588 | 0.071 | 0.0484 | 1 |
| Q05707 | COL14A1 | DtLFtAEsGTRRGIP | S1128 | 0.001 | 86.3 | 1796 | 0.000795 | 0.001 | 0.0008 | 1 |
| P04637 | TP53 | DstPPPGtRVRAMAI | T155 | 0.076 | 98.18 | 393 | 0.051909 | 0.076 | 0.0519 | 1 |
| P04637 | TP53 | rHsVVVPyEPPEVGS | Y220 | 0.308 | 97.45 | 393 | 0.135115 | 0.308 | 0.1351 | 1 |
| O60934 | NBN | RLLTGVEyVVGRKNC | Y24 | 0.32 | 95.7 | 754 | 0.140379 | 0.32 | 0.1404 | 0.8182 |
| P08908 | HTR1A | DHGYtIYsTFGAFyI | S199 | 0.091 | 93.82 | 422 | 0.072337 | 0.091 | 0.0723 | 1 |
| O75106 | AOC2 | GRVHAtGyINtAFLK | Y488 | 0.324 | 97.47 | 756 | 0.142134 | 0.324 | 0.1421 | 0.9333 |
| O75106 | AOC2 | HAtGyINtAFLKGGE | T491 | 0.001 | 96.9 | 756 | 0.000683 | 0.001 | 0.0007 | 1 |
| P23528 | CFL1 | yALyDAtyEtkEskk | Y89 | 0.101 | 91.48 | 166 | 0.044307 | 0.202 | 0.0886 | 0.8182 |
| P23528 | CFL1 | LyDAtyEtkEskkED | T91 | 0.174 | 90.64 | 166 | 0.118844 | 0.174 | 0.1188 | 1 |
| P23528 | CFL1 | LGGsAVIsLEGkPL_ | S160 | 0.117 | 89.65 | 166 | 0.093005 | 0.117 | 0.093 | 1 |
| P16144 | ITGB4 | VFRVRAQsQEGWGRE | S1604 | 0.026 | 88.19 | 1822 | 0.020668 | 0.026 | 0.0207 | 1 |
| Q93009 | USP7 | FRGKMVSyIQCkEVD | Y331 | 0.315 | 94.13 | 1102 | 0.138186 | 0.315 | 0.1382 | 1 |
| Q93009 | USP7 | QDtSLILyEEVkPNL | Y735 | 0.109 | 97.49 | 1102 | 0.047817 | 0.109 | 0.0478 | 1 |
| Q93009 | USP7 | EEVkPNLtERIQDYD | T743 | 0.042 | 94.9 | 1102 | 0.028687 | 0.042 | 0.0287 | 1 |
| O15539 | RGS5 | HALMEKDsLPRFVRS | S166 | 0.069 | 98.41 | 181 | 0.054849 | 0.069 | 0.0548 | 0.8333 |
| Q9NZV1 | CRIM1 | CHFSGKAyADEERWD | Y826 | 0.297 | 78.93 | 1036 | 0.13029 | 0.297 | 0.1303 | 1 |
| Q9H7H0 | METTL17 | GSGTGSVtWAAHsIW | T200 | 0.034 | 98.82 | 456 | 0.023222 | 0.034 | 0.0232 | 1 |
| Q8IVL1 | NAV2 | NRFLEAHsSSDVtIG | S2333 | 0.056 | 88.32 | 2488 | 0.044515 | 0.056 | 0.0445 | 1 |

| | | | | | | | | | | |
|---|---|---|---|---|---|---|---|---|---|---|
| Q8IVL1 | NAV2 | AHsSSDVtIGPRLFL | T2338 | 0.06 | 90.52 | 2488 | 0.040981 | 0.06 | 0.041 | 0.875 |
| P46093 | GPR4 | HVLLLSRsAIyLGRP | S248 | 0.002 | 94.15 | 362 | 0.00159 | 0.002 | 0.0016 | 1 |
| P23141 | CES1 | TSyPPMCtQDPKAGQ | T88 | 0.043 | 98.77 | 567 | 0.02937 | 0.043 | 0.0294 | 1 |
| P23141 | CES1 | HRDAGAPtYMYEFQY | T444 | 0.009 | 98.89 | 567 | 0.006147 | 0.009 | 0.0061 | 1 |
| P23141 | CES1 | SEEEIRLsKMVMKFW | S493 | 0 | 98.8 | 567 | 0 | 0 | 0 | 0.9167 |
| P23141 | CES1 | EYNQKEGyLQIGANT | Y526 | 0.153 | 98.89 | 567 | 0.067119 | 0.263 | 0.1049 | 1 |
| Q9HCS5 | EPB41L4A | PSEAELNyLRTAKSL | Y196 | 0.061 | 98.61 | 686 | 0.02676 | 0.061 | 0.0268 | 1 |
| Q96SL1 | SLC49A4 | DKRGLRItVLLTSFL | T119 | 0.008 | 93.85 | 478 | 0.005464 | 0.008 | 0.0055 | 1 |
| Q9Y2G1 | MYRF | GEPKyVKtPEGLKPL | T419 | 0.127 | 94.6 | 1151 | 0.086743 | 0.127 | 0.0867 | 1 |
| P43004 | SLC1A2 | ILPLIISsLITGLSG | S100 | 0 | 93.94 | 574 | 0 | 0 | 0 | 1 |
| Q9Y3D6 | FIS1 | GNyRLKEyEKALKyV | Y87 | 0.163 | 89.52 | 152 | 0.071506 | 0.313 | 0.1373 | 0.8 |
| Q9BXM7 | PINK1 | EGLGHGRtLFLVMkN | T313 | 0.006 | 95.96 | 581 | 0.004098 | 0.006 | 0.0041 | 1 |
| P31025 | LCN1 | KTDEPGKyTADGGKH | Y95 | 0.044 | 97.11 | 176 | 0.019302 | 0.044 | 0.0193 | 1 |
| O95672 | ECEL1 | PHRVLHNyLVWRVVV | Y400 | 0 | 97.82 | 775 | 0 | 0 | 0 | 1 |
| O95672 | ECEL1 | PPQALNAyYLPNKNQ | Y573 | 0.192 | 95.63 | 775 | 0.084228 | 0.192 | 0.0842 | 1 |
| Q86WI1 | PKHD1L1 | STQITCYtRAMPEDS | T102 | 0.083 | 96.35 | 4243 | 0.05669 | 0.083 | 0.0567 | 0.9167 |
| Q86WI1 | PKHD1L1 | yGSNIALssNGKNVR | S180 | 0.052 | 92.53 | 4243 | 0.041336 | 0.041 | 0.055 | 1 |
| Q86WI1 | PKHD1L1 | PHILKTVyPGGRGLK | Y336 | 0.107 | 92.705 | 4243 | 0.046939 | 0.107 | 0.0469 | 0.9167 |
| Q86WI1 | PKHD1L1 | HINRGQKtAETDAYC | T704 | 0.058 | 87.885 | 4243 | 0.039615 | 0.058 | 0.0396 | 0.8333 |
| P23467 | PTPRB | TNGStVPsPVKDIGI | S204 | 0.088 | 80.11 | 1997 | 0.069952 | 0.169 | 0.1351 | 1 |
| P23467 | PTPRB | KyMAtVTsIsGDLKN | S537 | 0.001 | 85.56 | 1997 | 0.000795 | 0.001 | 0.0008 | 1 |
| P23467 | PTPRB | MAtVTsIsGDLKNSS | S539 | 0.035 | 75.25 | 1997 | 0.027822 | 0.035 | 0.0278 | 1 |
| P46778 | RPL21 | HGVVPLAtyMrIYKk | T29 | 0.216 | 95.19 | 160 | 0.147531 | 0.216 | 0.1475 | 1 |
| P04920 | SLC4A2 | KHHPDVTyVkKVRTL | Y1154 | 0.068 | 75.76 | 1241 | 0.029831 | 0.068 | 0.0298 | 1 |
| Q15042 | RAB3GAP1 | CQGPGVRtDFEMVHL | T188 | 0.004 | 97.19 | 981 | 0.002732 | 0.004 | 0.0027 | 1 |
| Q8N201 | INTS1 | KDDyLRAsRALLREI | S495 | 0.008 | 90.81 | 2190 | 0.006359 | 0.008 | 0.0064 | 0.875 |
| Q8N201 | INTS1 | FTEQPEtyYKWDNWP | Y624 | 0.051 | 90.88 | 2190 | 0.022373 | 0.051 | 0.0224 | 1 |
| Q02252 | ALDH6A1 | RVCNLIDsGtKEGAs | S373 | 0.158 | 98.13 | 535 | 0.125597 | 0.158 | 0.1256 | 1 |
| Q8TF64 | GIPC3 | ITDNGAGyAFIKRIk | Y133 | 0.253 | 96.14 | 312 | 0.110987 | 0.253 | 0.111 | 1 |
| Q8TF64 | GIPC3 | IKRIkEGsIINRIEA | S143 | 0.099 | 96.72 | 312 | 0.078697 | 0.099 | 0.0787 | 1 |
| Q9HAU5 | UPF2 | CSLFHQRyADFAPsL | Y233 | 0.111 | 96.08 | 1272 | 0.048694 | 0.111 | 0.0487 | 1 |

| | | | | | | | | | | |
|---|---|---|---|---|---|---|---|---|---|---|
| Q9HAU5 | UPF2 | PNItKLRtDLRFIAE | T264 | 0.088 | 96.36 | 1272 | 0.060105 | 0.088 | 0.0601 | 1 |
| O60518 | RANBP6 | REGFVEytEQVVKLM | T708 | 0 | 96.73 | 1105 | 0 | 0 | 0 | 1 |
| Q9UHI8 | ADAMTS1 | ITIPTGAtNIEVKQR | T749 | 0.062 | 93.04 | 967 | 0.042347 | 0.062 | 0.0423 | 1 |
| Q68EN5 | MATCAP1 | GLRPANPtEEGLASL | T314 | 0.024 | 97.89 | 471 | 0.016392 | 0.024 | 0.0164 | 1 |
| O00341 | SLC1A7 | LGVLTVAyYLWTTFM | Y95 | 0.223 | 96.33 | 560 | 0.097827 | 0.223 | 0.0978 | 1 |
| P40189 | IL6ST | VNLTNDRyLATLTVR | Y396 | 0.154 | 91.77 | 918 | 0.067558 | 0.154 | 0.0676 | 1 |
| Q96RQ9 | IL4I1 | KCMQDPDyEQLLKVV | Y42 | 0.202 | 97.35 | 567 | 0.088614 | 0.202 | 0.0886 | 0.8182 |
| Q96PE1 | ADGRA2 | HPVVyPCtALLLLCL | T777 | 0.043 | 96.17 | 1338 | 0.02937 | 0.169 | 0.1154 | 1 |
| Q86Y07 | VRK2 | SGGFGLIyLAFPTNK | Y44 | 0.046 | 97.42 | 508 | 0.02018 | 0.046 | 0.0202 | 1 |
| Q9BPX1 | HSD17B14 | AGKVVVVtGGGRGIG | T15 | 0.031 | 98.86 | 270 | 0.021173 | 0.031 | 0.0212 | 1 |
| Q8N7X0 | ADGB | YTKNKKGyTFVAEAF | Y1062 | 0.05 | 93.39 | 1667 | 0.021934 | 0.05 | 0.0219 | 1 |
| Q8N7X0 | ADGB | TPLQNYKyIIQCSVL | Y1254 | 0.05 | 92.66 | 1667 | 0.021934 | 0.05 | 0.0219 | 1 |
| Q9Y6K8 | AK5 | LVIGGPGsGKGTQSL | S143 | 0.022 | 94.09 | 562 | 0.017488 | 0.022 | 0.0175 | 0.8889 |
| Q9UBH0 | IL36RN | PVNIMELyLGAKESK | Y89 | 0.04 | 96.24 | 155 | 0.017547 | 0.04 | 0.0175 | 1 |
| Q9Y5M8 | SRPRB | AIVFVVDsAAFQREV | S145 | 0 | 98.49 | 271 | 0 | 0 | 0 | 0.9091 |
| Q9Y277 | VDAC3 | _____MCNtPtyCDLG | T4 | 0.019 | 89.75 | 283 | 0.012977 | 0.019 | 0.013 | 1 |
| Q8NG78 | OR8G5 | PSLTILSsyIFIIAS | S252 | 0.047 | 94.4 | 346 | 0.037361 | 0.047 | 0.0374 | 1 |
| Q5JSJ4 | INTS6L | LKTLPPyyLLTKLES | Y427 | 0.127 | 82.97 | 861 | 0.055713 | 0.127 | 0.0557 | 1 |
| P52789 | HK2 | VEMENQIyAIPEDIM | Y112 | 0.316 | 93.23 | 917 | 0.138625 | 0.316 | 0.1386 | 1 |
| P52789 | HK2 | KkLPLGFtFSFPCHQ | T153 | 0 | 95.25 | 917 | 0 | 0 | 0 | 1 |
| P52789 | HK2 | GTGSNACyMEEMRHI | Y238 | 0 | 98.17 | 917 | 0 | 0 | 0 | 1 |
| P52789 | HK2 | EKMISGMyMGELVRL | Y301 | 0.146 | 95.44 | 917 | 0.064048 | 0.146 | 0.064 | 1 |
| P52789 | HK2 | IVSTRSAsLCAATLA | S384 | 0.135 | 98.62 | 917 | 0.107313 | 0.135 | 0.1073 | 1 |
| P52789 | HK2 | CDVRFLRsEDGsGkG | S445 | 0.006 | 89.57 | 917 | 0.004769 | 0.006 | 0.0048 | 1 |
| P52789 | HK2 | FLRsEDGsGkGAAMV | S449 | 0.055 | 83.08 | 917 | 0.04372 | 0.055 | 0.0437 | 1 |
| P52789 | HK2 | GkGAAMVtAVAyRLA | T457 | 0 | 91.85 | 917 | 0 | 0 | 0 | 1 |
| P52789 | HK2 | QHRARQktLEHLQLS | T473 | 0.158 | 96.09 | 917 | 0.107916 | 0.158 | 0.1079 | 0.9091 |
| P52789 | HK2 | APVkMLPtYVCATPD | T514 | 0.051 | 94.83 | 917 | 0.034834 | 0.051 | 0.0348 | 1 |
| P52789 | HK2 | GTGSNACyMEEMRNV | Y686 | 0.005 | 98.63 | 917 | 0.002193 | 0.005 | 0.0022 | 1 |
| P52789 | HK2 | QRFEkMIsGMyLGEI | S746 | 0 | 94.52 | 917 | 0 | 0 | 0 | 1 |
| P52789 | HK2 | EkMIsGMyLGEIVRN | Y749 | 0.228 | 95.01 | 917 | 0.10002 | 0.228 | 0.1 | 1 |

| | | | | | | | | | | |
|---|---|---|---|---|---|---|---|---|---|---|
| P52789 | HK2 | RNILIDFtkRGLLFR | T762 | 0 | 95.34 | 917 | 0 | 0 | 0 | 1 |
| P52789 | HK2 | FLQSEDGsGkGAALI | S897 | 0.127 | 79.93 | 917 | 0.100954 | 0.127 | 0.101 | 1 |
| P52789 | HK2 | GkGAALItAVACRIR | T905 | 0 | 93.06 | 917 | 0 | 0 | 0 | 1 |
| Q96GQ7 | DDX27 | APDILIAtPGRLIDH | T346 | 0.145 | 91.78 | 796 | 0.099037 | 0.145 | 0.099 | 0.8 |
| Q96GQ7 | DDX27 | VKDLAsVsLKNPVRI | S416 | 0.111 | 84.82 | 796 | 0.088236 | 0.111 | 0.0882 | 1 |
| O75030 | MITF | kGTILKAsVDyIRKL | S357 | 0.106 | 98.25 | 526 | 0.084261 | 0.106 | 0.0843 | 1 |
| O75970 | MPDZ | IKHVLEDsPAGKNGT | S1194 | 0.085 | 88.06 | 2070 | 0.067568 | 0.085 | 0.0676 | 1 |
| Q8N5H7 | SH2D3C | QMRLLWGsQGASSSQ | S826 | 0.175 | 94.08 | 860 | 0.13911 | 0.175 | 0.1391 | 1 |
| Q5GH76 | XKR4 | TIYLGIRsRQSGEND | S280 | 0.037 | 85.72 | 650 | 0.029412 | 0.123 | 0.0978 | 1 |
| Q16769 | QPCT | sFSNIIStLNPtAkR | T126 | 0.066 | 98.8 | 361 | 0.045079 | 0.066 | 0.0451 | 0.875 |
| P78527 | PRKDC | SLLrLQEtLsAADrC | T18 | 0.136 | 83.56 | 4128 | 0.09289 | 0.136 | 0.0929 | 1 |
| P41250 | GARS1 | PEPVLKtsGHVDkFA | S192 | 0.127 | 85.26 | 739 | 0.100954 | 0.144 | 0.1145 | 1 |
| P41250 | GARS1 | DLHLYLYsAkAQVSG | S377 | 0.156 | 91.66 | 739 | 0.124007 | 0.156 | 0.124 | 1 |
| P41250 | GARS1 | GYFIGRIyLYLTkVG | Y414 | 0.312 | 98.86 | 739 | 0.13687 | 0.312 | 0.1369 | 1 |
| P26641 | EEF1G | TQTLVREyFsWEGAF | Y416 | 0 | 96.65 | 437 | 0 | 0 | 0 | 0.8125 |
| O00519 | FAAH | APEAVLFtYVGKAWE | T99 | 0.115 | 98.61 | 579 | 0.078547 | 0.115 | 0.0785 | 1 |
| O00519 | FAAH | kPLLPRLsAFLsNMK | S430 | 0.09 | 95.85 | 579 | 0.071542 | 0.09 | 0.0715 | 1 |
| Q96SQ9 | CYP2S1 | WRQLRkFtMLALRDL | T131 | 0 | 98.29 | 504 | 0 | 0 | 0 | 1 |
| Q96SQ9 | CYP2S1 | TNKNMLMtVIYLLFA | T295 | 0 | 98.11 | 504 | 0 | 0 | 0 | 1 |
| Q96SQ9 | CYP2S1 | PRtLMRttRFRGYTL | T379 | 0.065 | 96.74 | 504 | 0.044396 | 0.065 | 0.0444 | 1 |
| O15511 | ARPC5 | LAAGGVGsIVRVLtA | S140 | 0 | 98.29 | 151 | 0 | 0 | 0 | 1 |
| Q9Y5G1 | PCDHGB3 | DREQIPEyNVtITAT | Y418 | 0.228 | 91.38 | 929 | 0.10002 | 0.228 | 0.1 | 1 |
| Q9Y5G1 | PCDHGB3 | EPRELsSyVSVSARs | Y504 | 0.332 | 92.57 | 929 | 0.145644 | 0.332 | 0.1456 | 1 |
| Q6ZV73 | FGD6 | KGPyLKMystyIKEF | Y966 | 0 | 93.78 | 1430 | 0 | 0 | 0 | 1 |
| Q6ZV73 | FGD6 | GPyLKMystyIKEFD | S967 | 0.043 | 91.27 | 1430 | 0.034181 | 0.043 | 0.0342 | 1 |
| Q6ZV73 | FGD6 | yLKMystyIKEFDKN | Y969 | 0.049 | 93.72 | 1430 | 0.021496 | 0.049 | 0.0215 | 1 |
| P16520 | GNB3 | TDSKLLVsAsQDGkL | S72 | 0 | 98.85 | 340 | 0 | 0 | 0 | 1 |
| P16520 | GNB3 | SKLLVsAsQDGkLIV | S74 | 0 | 98.65 | 340 | 0 | 0 | 0 | 1 |
| Q99829 | CPNE1 | kkDFLGksDPFLEFF | S158 | 0.003 | 96.77 | 537 | 0.002385 | 0.003 | 0.0024 | 1 |
| Q99829 | CPNE1 | QVRLyGPtNFAPIIN | T389 | 0.133 | 92.98 | 537 | 0.090841 | 0.133 | 0.0908 | 1 |
| Q9Y2I1 | NISCH | ATLsVRFsAtSMKEV | S250 | 0.118 | 92.49 | 1504 | 0.0938 | 0.118 | 0.0938 | 1 |

| | | | | | | | | | | |
|---|---|---|---|---|---|---|---|---|---|---|
| P35637 | FUS | VsFDDPPsAkAAIDW | S346 | 0.181 | 90.69 | 526 | 0.14388 | 0.181 | 0.1439 | 0.8571 |
| O00767 | SCD | ILARIKRtGDGNYkS | T351 | 0.175 | 97.38 | 359 | 0.119527 | 0.175 | 0.1195 | 1 |
| P14866 | HNRNPL | DPHktPAsPVVHIrG | S101 | 0.001 | 87.12 | 589 | 0.000795 | 0.001 | 0.0008 | 1 |
| Q8IVN8 | SBSPON | THtEDAGyCMEFKTE | Y168 | 0.059 | 93.38 | 264 | 0.025882 | 0.059 | 0.0259 | 1 |
| P60891 | PRPS1 | sAGAtRVyAILTHGI | Y245 | 0.164 | 98.73 | 318 | 0.071944 | 0.164 | 0.0719 | 1 |
| P35453 | HOXD13 | RRRISAAtNLsERQV | T313 | 0.073 | 96.82 | 343 | 0.04986 | 0.073 | 0.0499 | 1 |
| Q9Y4D1 | DAAM1 | VLQAMLHyQKYASER | Y247 | 0.048 | 95.91 | 1078 | 0.021057 | 0.048 | 0.0211 | 1 |
| Q9Y4D1 | DAAM1 | LDKSTGRyRDEVSLK | Y271 | 0.123 | 77.52 | 1078 | 0.053958 | 0.123 | 0.054 | 1 |
| Q9Y4D1 | DAAM1 | KGQDPDStPLENFNI | T425 | 0.182 | 69.94 | 1078 | 0.124308 | 0.182 | 0.1243 | 1 |
| P54108 | CRISP3 | VRDSCKAsCNCSNsI | S237 | 0.012 | 95.38 | 245 | 0.009539 | 0.012 | 0.0095 | 0.9 |
| Q00610 | CLTC | yDAAkLLyNNVsNFG | Y1218 | 0.005 | 79.55 | 1675 | 0.002193 | 0.005 | 0.0022 | 1 |
| Q00610 | CLTC | GNNRWkQsVELCkKD | S1524 | 0 | 72.91 | 1675 | 0 | 0 | 0 | 0.9 |
| P45974 | USP5 | NkEELLEyEEkKRQA | Y505 | 0.293 | 91.94 | 858 | 0.128535 | 0.293 | 0.1285 | 0.9 |
| Q5T6L9 | ERMARD | ILKIMLPyWEVALVK | Y274 | 0.062 | 98.55 | 678 | 0.027199 | 0.062 | 0.0272 | 1 |
| Q460N5 | PARP14 | WEkGsLVsPGGLQML | S1010 | 0.081 | 85.15 | 1801 | 0.064388 | 0.094 | 0.0823 | 0.875 |
| Q9BV10 | ALG12 | LGLVDRRtHAPtVLA | T288 | 0.001 | 95.82 | 488 | 0.000683 | 0.001 | 0.0007 | 1 |
| Q9BV10 | ALG12 | AAAQTGVsRFLQVNs | S403 | 0.018 | 95.82 | 488 | 0.014308 | 0.018 | 0.0143 | 0.8333 |
| Q9BV10 | ALG12 | QVNsAWRyDKREDVQ | Y414 | 0.3 | 95.49 | 488 | 0.131606 | 0.3 | 0.1316 | 0.8182 |
| P03951 | F11 | TkLDKVVsGFSLKSC | S186 | 0 | 98.58 | 625 | 0 | 0 | 0 | 1 |
| Q14534 | SQLE | SRDGRkVtVIERDLk | T150 | 0.091 | 98.82 | 574 | 0.062154 | 0.091 | 0.0622 | 1 |
| P42285 | MTREX | EKQRVIFtsPIkALS | T188 | 0 | 96.81 | 1042 | 0 | 0 | 0 | 1 |
| P42285 | MTREX | KQRVIFtsPIkALSN | S189 | 0.001 | 91.95 | 1042 | 0.000795 | 0.001 | 0.0008 | 1 |
| P42285 | MTREX | RGVVWEEtIILLPDN | T269 | 0 | 95.99 | 1042 | 0 | 0 | 0 | 1 |
| P42285 | MTREX | RWISSGEyIQMSGRA | Y517 | 0.024 | 93.86 | 1042 | 0.010528 | 0.024 | 0.0105 | 1 |
| Q9BYD6 | MRPL1 | INKVAVFtENASEVK | T161 | 0.021 | 95.98 | 325 | 0.014343 | 0.021 | 0.0143 | 1 |
| Q8NHV1 | GIMAP7 | kHMVILFtRKEELEG | T132 | 0.051 | 97.92 | 300 | 0.034834 | 0.078 | 0.0533 | 1 |
| Q8NHV1 | GIMAP7 | AyFSDDIykDTEERL | Y206 | 0.075 | 98.02 | 300 | 0.032901 | 0.124 | 0.0461 | 0.9286 |
| O15294 | OGT | LPEDAIVyCNFNQLY | Y844 | 0.003 | 98.9 | 1046 | 0.001316 | 0.003 | 0.0013 | 1 |
| O15294 | OGT | IAkNRQEyEDIAVkL | Y976 | 0 | 98.87 | 1046 | 0 | 0 | 0 | 1 |
| O15294 | OGT | VWKQRISsPLFNTkQ | S1004 | 0.084 | 97.34 | 1046 | 0.066773 | 0.168 | 0.1335 | 1 |
| P21108 | PRPS1L1 | sAGAtRVyAILTHGI | Y245 | 0.118 | 98.46 | 318 | 0.051765 | 0.118 | 0.0518 | 1 |

| | | | | | | | | | | |
|---|---|---|---|---|---|---|---|---|---|---|
| Q502W6 | VWA3B | TAFADKEtQAIYLLT | T599 | 0.099 | 87.41 | 1294 | 0.067618 | 0.099 | 0.0676 | 1 |
| Q13510 | ASAH1 | NKtVFKAsSFAGYVG | S202 | 0.008 | 98.69 | 395 | 0.006359 | 0.008 | 0.0064 | 0.8 |
| O76096 | CST7 | QIVKGLKyMLEVEIG | Y88 | 0.143 | 98.81 | 145 | 0.062732 | 0.143 | 0.0627 | 1 |
| O76096 | CST7 | LEVEIGRttCKKNQH | T97 | 0 | 98.68 | 145 | 0 | 0 | 0 | 1 |
| O15244 | SLC22A2 | LVLLLPEtKGKALPE | T517 | 0.075 | 83.69 | 555 | 0.051226 | 0.075 | 0.0512 | 1 |
| P78344 | EIF4G2 | EkAssLIsLLKQEGI | S605 | 0.158 | 95.22 | 907 | 0.125597 | 0.158 | 0.1256 | 0.8571 |
| Q86UB9 | TMEM135 | FGAALPAsYVAILIE | S110 | 0.001 | 98.33 | 458 | 0.000795 | 0.001 | 0.0008 | 1 |
| Q58DX5 | NAALADL2 | LQKNVVAyIsLHsPI | Y524 | 0.006 | 98.28 | 795 | 0.002632 | 0.006 | 0.0026 | 1 |
| Q58DX5 | NAALADL2 | KNVVAyIsLHsPIRG | S526 | 0.011 | 97.95 | 795 | 0.008744 | 0.011 | 0.0087 | 1 |
| Q58DX5 | NAALADL2 | VAyIsLHsPIRGNSS | S529 | 0.032 | 90.39 | 795 | 0.025437 | 0.032 | 0.0254 | 1 |
| Q9Y2X0 | MED16 | CRNLIAFtMDLRSDD | T48 | 0.004 | 95.52 | 877 | 0.002732 | 0.004 | 0.0027 | 1 |
| Q8NEM7 | SUPT20H | AGTQYQktkLtILQs | T350 | 0.163 | 84.81 | 779 | 0.111331 | 0.163 | 0.1113 | 1 |
| Q9NYB0 | TERF2IP | DGSSMSFyVRPsPAK | Y32 | 0.306 | 93.54 | 399 | 0.134238 | 0.329 | 0.1443 | 1 |
| Q9UNZ2 | NSFL1C | KLWksGFsLDNGELr | S192 | 0.009 | 87.59 | 370 | 0.007154 | 0.009 | 0.0072 | 1 |
| Q96N66 | MBOAT7 | LLTLKLVsLASEVQD | S114 | 0 | 97.12 | 472 | 0 | 0 | 0 | 1 |
| Q96N66 | MBOAT7 | GPFFRYRtyLDWLEQ | T172 | 0.029 | 96.41 | 472 | 0.019807 | 0.029 | 0.0198 | 1 |
| Q96N66 | MBOAT7 | PFFRYRtyLDWLEQP | Y173 | 0.001 | 97.39 | 472 | 0.000439 | 0.001 | 0.0004 | 1 |
| Q96N66 | MBOAT7 | PFPGAVPsLRPLLRR | S187 | 0.07 | 97.17 | 472 | 0.055644 | 0.07 | 0.0556 | 1 |
| Q96N66 | MBOAT7 | EkAAsLEyDyEtIRN | Y294 | 0.31 | 97.07 | 472 | 0.135993 | 0.31 | 0.136 | 1 |
| Q96N66 | MBOAT7 | AAsLEyDyEtIRNID | Y296 | 0.219 | 97.95 | 472 | 0.096072 | 0.219 | 0.0961 | 1 |
| Q96N66 | MBOAT7 | sLEyDyEtIRNIDCY | T298 | 0.011 | 97.71 | 472 | 0.007513 | 0.011 | 0.0075 | 1 |
| Q9NP58 | ABCB6 | MNtQENAtRARAVDs | T449 | 0.124 | 89.81 | 842 | 0.084694 | 0.124 | 0.0847 | 1 |
| P18054 | ALOX12 | WKEGLPLtIAADRKD | T151 | 0.042 | 98.34 | 663 | 0.028687 | 0.042 | 0.0287 | 1 |
| P18054 | ALOX12 | NPMLLRRsTsLPsRL | S244 | 0.075 | 98.52 | 663 | 0.059619 | 0.075 | 0.0596 | 1 |
| Q6PCB6 | ABHD17C | NIFSYDYsGyGVSSG | S169 | 0.013 | 98.5 | 329 | 0.010334 | 0.013 | 0.0103 | 1 |
| P20042 | EIF2S2 | RVGTKkTsFVNFTDI | S218 | 0.026 | 93.36 | 333 | 0.020668 | 0.026 | 0.0207 | 1 |
| P20042 | EIF2S2 | LRRYIkEyVTCHtCR | Y278 | 0.043 | 92.65 | 333 | 0.018863 | 0.043 | 0.0189 | 1 |
| P20042 | EIF2S2 | VTCHtCRsPDTILQk | S286 | 0.084 | 84.94 | 333 | 0.066773 | 0.151 | 0.12 | 1 |
| Q8N608 | DPP10 | KASRHSVsPDLKyVL | S133 | 0 | 96.89 | 796 | 0 | 0 | 0 | 1 |
| Q8N608 | DPP10 | SVsPDLKyVLLAyDV | Y138 | 0.293 | 96.76 | 796 | 0.128535 | 0.293 | 0.1285 | 1 |
| Q8N608 | DPP10 | VNYTMQVyPDEGHNV | Y754 | 0.161 | 98.3 | 796 | 0.070628 | 0.161 | 0.0706 | 1 |

| | | | | | | | | | | |
|---|---|---|---|---|---|---|---|---|---|---|
| Q9HC29 | NOD2 | KNARKVVtsRPAAVS | T424 | 0.008 | 95.61 | 1040 | 0.005464 | 0.008 | 0.0055 | 1 |
| Q9HC29 | NOD2 | NARKVVtsRPAAVSA | S425 | 0 | 93.2 | 1040 | 0 | 0 | 0 | 1 |
| Q9HC29 | NOD2 | PVFSWMVsKCHQELL | S493 | 0.058 | 93.41 | 1040 | 0.046105 | 0.058 | 0.0461 | 1 |
| P13569 | CFTR | EENKVRQyDSIQKLL | Y1424 | 0.14 | 87.39 | 1480 | 0.061416 | 0.14 | 0.0614 | 1 |
| Q9BRP7 | FDXACB1 | HPKtQQVsLCyRLTY | S585 | 0.046 | 94.35 | 624 | 0.036566 | 0.046 | 0.0366 | 1 |
| Q9BRP7 | FDXACB1 | tQQVsLCyRLTYQTC | Y588 | 0.063 | 97.77 | 624 | 0.027637 | 0.063 | 0.0276 | 1 |
| P07339 | CTSD | ykLSPEDyTLkVSQA | Y354 | 0.003 | 96.46 | 412 | 0.001316 | 0.003 | 0.0013 | 0.8 |
| P07339 | CTSD | DVFIGRYyTVFDRDN | Y394 | 0.033 | 98.63 | 412 | 0.014477 | 0.033 | 0.0145 | 1 |
| Q9UFH2 | DNAH17 | KNEMFVMsGKIKGKT | S153 | 0.15 | 78.99 | 4462 | 0.119237 | 0.15 | 0.1192 | 1 |
| P62841 | RPS15 | ILPEMVGsMVGVYNG | S92 | 0.033 | 93.49 | 145 | 0.026232 | 0.033 | 0.0262 | 1 |
| P14543 | NID1 | PFLADLDttDGLGKV | T113 | 0.04 | 86.91 | 1247 | 0.027321 | 0.04 | 0.0273 | 0.9091 |
| P50502 | ST13 | QkAIDLFtDAIkLNP | T138 | 0.17 | 98.68 | 369 | 0.116112 | 0.17 | 0.1161 | 1 |
| P50502 | ST13 | ILYAkRAsVFVkLQk | S156 | 0.091 | 98.39 | 369 | 0.072337 | 0.091 | 0.0723 | 1 |
| P50502 | ST13 | LDyDEDAsAMLkEVQ | S218 | 0.185 | 98.01 | 369 | 0.147059 | 0.185 | 0.1471 | 1 |
| Q9BXT8 | RNF17 | HQEVEVKyVDFGNTA | Y760 | 0.036 | 94.81 | 1623 | 0.015793 | 0.036 | 0.0158 | 1 |
| Q9UJA9 | ENPP5 | HYTLIDQsPVAAILP | S267 | 0.09 | 95.76 | 477 | 0.071542 | 0.09 | 0.0715 | 0.8571 |
| Q9NZU0 | FLRT3 | ISTISSPsLQGLTSL | S194 | 0.04 | 94.33 | 649 | 0.031797 | 0.04 | 0.0318 | 1 |
| Q8N4N3 | KLHL36 | FTYGHAGtIYKDFVY | T440 | 0.1 | 98.43 | 616 | 0.068301 | 0.1 | 0.0683 | 1 |
| Q15436 | SEC23A | VPVAALFtPLkERPD | T43 | 0.159 | 98.6 | 765 | 0.108599 | 0.226 | 0.1441 | 1 |
| Q15436 | SEC23A | DLQALKEsMQMSLSL | S150 | 0.092 | 98.07 | 765 | 0.073132 | 0.092 | 0.0731 | 1 |
| P49327 | FASN | GDLVMNVyRDGAWGA | Y1506 | 0.126 | 91.81 | 2511 | 0.055274 | 0.126 | 0.0553 | 1 |
| P49327 | FASN | CkPkYsGtLNLDrVT | T1999 | 0.004 | 95.9 | 2511 | 0.002732 | 0.004 | 0.0027 | 1 |
| P26373 | RPL13 | VRAGrGFsLEELrVA | S77 | 0.013 | 98.19 | 211 | 0.010334 | 0.013 | 0.0103 | 1 |
| P26373 | RPL13 | VARTIGIsVDPrRRN | S97 | 0.061 | 98.04 | 211 | 0.04849 | 0.109 | 0.0866 | 1 |
| Q29980 | MICB | AKtWDtEtEDLTENG | T86 | 0.15 | 96.99 | 383 | 0.102452 | 0.15 | 0.1025 | 1 |
| Q29980 | MICB | NGQDLRRtLtHIkDQ | T99 | 0.068 | 94.86 | 383 | 0.046445 | 0.068 | 0.0464 | 1 |
| O75369 | FLNB | LQsFTVDsskAGLAP | S1447 | 0.076 | 78.99 | 2602 | 0.060414 | 0.175 | 0.1395 | 1 |
| O75369 | FLNB | PYMVSVKyADEEIPR | Y1497 | 0.037 | 80.42 | 2602 | 0.016231 | 0.037 | 0.0162 | 1 |
| O75369 | FLNB | TAAKPGTyVIyVRFG | Y1684 | 0.042 | 84.66 | 2602 | 0.018425 | 0.042 | 0.0184 | 1 |
| P51812 | RPS6KA3 | EIEILLRyGQHPNII | Y470 | 0.225 | 86.44 | 740 | 0.098704 | 0.225 | 0.0987 | 1 |
| P51812 | RPS6KA3 | DESGNPEsIRICDFG | S556 | 0.122 | 93.82 | 740 | 0.09698 | 0.122 | 0.097 | 1 |

| | | | | | | | | | | |
|---|---|---|---|---|---|---|---|---|---|---|
| O14579 | COPE | LVNLIVLsQHLGkPP | S260 | 0.076 | 93.74 | 308 | 0.060414 | 0.093 | 0.0949 | 1 |
| Q53RE8 | ANKRD39 | QPDsAGytALHyASR | T66 | 0.039 | 94.71 | 183 | 0.026638 | 0.039 | 0.0266 | 1 |
| P00558 | PGK1 | rIkAAVPsIkFCLDN | S46 | 0.002 | 98.83 | 417 | 0.00159 | 0.002 | 0.0016 | 1 |
| P00558 | PGK1 | VPMPDkysLEPVAVE | S77 | 0.129 | 98.4 | 417 | 0.102544 | 0.129 | 0.1025 | 1 |
| P00558 | PGK1 | kIEAFRAsLskLGDV | S153 | 0.077 | 98.86 | 417 | 0.061208 | 0.107 | 0.1018 | 1 |
| P00558 | PGK1 | LskLGDVyVNDAFGt | Y161 | 0.047 | 98.93 | 417 | 0.020618 | 0.094 | 0.0412 | 1 |
| P00558 | PGK1 | yVNDAFGtAHRAHSs | T168 | 0.037 | 96.44 | 417 | 0.025272 | 0.055 | 0.0376 | 1 |
| P00558 | PGK1 | tAHRAHSsMVGVNLP | S175 | 0 | 98.84 | 417 | 0 | 0 | 0 | 1 |
| P00558 | PGK1 | IGGGMAFtFLKVLNN | T243 | 0.019 | 98.21 | 417 | 0.012977 | 0.019 | 0.013 | 1 |
| P00558 | PGK1 | NNMEIGTsLFDEEGA | S256 | 0.1 | 98.66 | 417 | 0.079491 | 0.173 | 0.1405 | 1 |
| P00558 | PGK1 | tLPVDFVtADkFDEN | T288 | 0.027 | 98.85 | 417 | 0.018441 | 0.054 | 0.0369 | 1 |
| P00558 | PGK1 | GPESSkkyAEAVTRA | Y324 | 0.025 | 98.87 | 417 | 0.010967 | 0.025 | 0.011 | 1 |
| P48775 | TDO2 | SPASGFQsLQFRLLE | S155 | 0.167 | 97.13 | 406 | 0.132751 | 0.167 | 0.1328 | 0.8571 |
| Q9UFH2 | DNAH17 | LAKDRMNyKMDLEDN | Y952 | 0.001 | 89.13 | 4462 | 0.000439 | 0.001 | 0.0004 | 0.9333 |
| P07741 | APRT | PVLkDPAsFRAAIGL | S38 | 0.007 | 98.63 | 180 | 0.005564 | 0.007 | 0.0056 | 1 |
| O95477 | ABCA1 | RFMtGLDtKNNVKVW | T1586 | 0.107 | 76.13 | 2261 | 0.073082 | 0.135 | 0.0922 | 0.8182 |
| Q05193 | DNM1 | TGTNkGIsPVPINLR | S116 | 0.124 | 90.15 | 864 | 0.098569 | 0.124 | 0.0986 | 1 |
| Q05193 | DNM1 | LLPLRRGyIGVVNRS | Y231 | 0.144 | 96.38 | 864 | 0.063171 | 0.144 | 0.0632 | 1 |
| Q14C86 | GAPVD1 | RTISAYktPRDkVQC | T1389 | 0.031 | 91.55 | 1478 | 0.021173 | 0.031 | 0.0212 | 1 |
| Q9NUQ2 | AGPAT5 | WGVWRLLsAFLPARF | S39 | 0.064 | 95.35 | 364 | 0.050875 | 0.082 | 0.0703 | 1 |
| Q9NUQ2 | AGPAT5 | SkLSIKKtLPsMLIL | T321 | 0 | 98.33 | 364 | 0 | 0 | 0 | 0.9 |
| P49792 | RANBP2 | SFyDQrEyIGRSVHy | Y598 | 0 | 94.76 | 3224 | 0 | 0 | 0 | 1 |
| Q92521 | PIGB | LAFIIRPtAVILWtP | T234 | 0.166 | 94.83 | 554 | 0.11338 | 0.166 | 0.1134 | 1 |
| O15399 | GRIN2D | NYPDMHSyMVRyNQP | Y732 | 0.225 | 92.52 | 1336 | 0.098704 | 0.225 | 0.0987 | 1 |
| O15440 | ABCC5 | SVERINHyIKTLSLE | Y1166 | 0.255 | 89.29 | 1437 | 0.111865 | 0.255 | 0.1119 | 1 |
| Q04206 | RELA | LrLPPVLsHPIFDNr | S180 | 0.004 | 98.52 | 551 | 0.00318 | 0.004 | 0.0032 | 1 |
| Q04206 | RELA | QVAIVFRtPPYADPs | T254 | 0.048 | 98.46 | 551 | 0.032785 | 0.048 | 0.0328 | 1 |
| Q04206 | RELA | RPsDRELsEPMEFQY | S281 | 0.082 | 97.4 | 551 | 0.065183 | 0.082 | 0.0652 | 1 |
| Q9H061 | TMEM126A | AyLGSEQyKLLIkAL | Y178 | 0.338 | 94.67 | 195 | 0.148276 | 0.338 | 0.1483 | 0.8 |
| Q8WUM0 | NUP133 | QGGSFILsSSGSQLI | S223 | 0 | 93.66 | 1156 | 0 | 0.083 | 0.066 | 0.8571 |
| Q13045 | FLII | FIWLGRKsPRLVRAA | S792 | 0.074 | 83.39 | 1269 | 0.058824 | 0.074 | 0.0588 | 1 |

| | | | | | | | | | | |
|---|---|---|---|---|---|---|---|---|---|---|
| Q99747 | NAPG | yDsAAsEyGKAAVAF | Y38 | 0.112 | 98.71 | 312 | 0.049133 | 0.112 | 0.0491 | 1 |
| P50135 | HNMT | IIVVsGssGWDKLWK | S177 | 0 | 98.37 | 292 | 0 | 0 | 0 | 1 |
| P50135 | HNMT | GKVLFNNtLSFIVIE | T284 | 0.116 | 98.63 | 292 | 0.07923 | 0.116 | 0.0792 | 1 |
| Q9BTV4 | TMEM43 | GDLRVSFsYAGLsGD | S244 | 0.184 | 98.2 | 400 | 0.146264 | 0.184 | 0.1463 | 1 |
| Q9BTV4 | TMEM43 | SFsYAGLsGDDPDLG | S249 | 0.05 | 97.37 | 400 | 0.039746 | 0.05 | 0.0397 | 1 |
| Q08881 | ITK | VVIALyDyQtNDPQE | Y182 | 0.213 | 87.92 | 620 | 0.09344 | 0.397 | 0.1118 | 1 |
| Q08881 | ITK | ALRRNEEyCLLDsSE | Y198 | 0.048 | 94.48 | 620 | 0.021057 | 0.048 | 0.0211 | 1 |
| Q08881 | ITK | SRTAGTytVsVFTKA | T274 | 0.013 | 93.42 | 620 | 0.008879 | 0.013 | 0.0089 | 1 |
| Q08881 | ITK | TAGTytVsVFTKAVV | S276 | 0 | 94.23 | 620 | 0 | 0 | 0 | 1 |
| Q9UBW7 | ZMYM2 | SPVALLNtLFyFNtK | T1218 | 0 | 96.24 | 1377 | 0 | 0 | 0 | 1 |
| P98194 | ATP2C1 | AMDVQQLsQIVPKVA | S608 | 0.139 | 89.03 | 919 | 0.110493 | 0.139 | 0.1105 | 1 |
| P98194 | ATP2C1 | FDMFNALssRSQTkS | S825 | 0 | 91.2 | 919 | 0 | 0 | 0 | 1 |
| P98194 | ATP2C1 | DMFNALssRSQTkSV | S826 | 0 | 91.57 | 919 | 0 | 0 | 0 | 1 |
| Q96CG3 | TIFA | tVTCLQMtVYHPGQL | T19 | 0.187 | 97.99 | 184 | 0.127724 | 0.187 | 0.1277 | 1 |
| Q9NVH1 | DNAJC11 | LIIVNAWyGKFVNDk | Y469 | 0 | 92.19 | 559 | 0 | 0 | 0 | 1 |
| P24387 | CRHBP | PHQHRNCsFSIIYPV | S206 | 0.007 | 97.55 | 322 | 0.005564 | 0.007 | 0.0056 | 1 |
| Q8WXF7 | ATL1 | DKEVVAVsVAGAFRK | S71 | 0 | 98.38 | 558 | 0 | 0 | 0 | 1 |
| Q8WXF7 | ATL1 | KMGGEEFsRRYLQQL | S414 | 0.118 | 94.13 | 558 | 0.0938 | 0.118 | 0.0938 | 1 |
| Q8WXF7 | ATL1 | IDELyIQyIKHNDSK | Y432 | 0.142 | 94.52 | 558 | 0.062293 | 0.284 | 0.1246 | 1 |
| Q9BZD2 | SLC29A3 | TALVKVDtSSWTRGF | T157 | 0.029 | 84.21 | 475 | 0.019807 | 0.029 | 0.0198 | 0.8 |
| Q6UX07 | DHRS13 | MGNLRGRtAVVtGAN | T38 | 0.066 | 98.44 | 377 | 0.045079 | 0.107 | 0.0731 | 1 |
| Q6UX07 | DHRS13 | RGRtAVVtGANsGIG | T42 | 0.009 | 98.5 | 377 | 0.006147 | 0.009 | 0.0061 | 1 |
| Q6UX07 | DHRS13 | HIGPFLLtHLLLPCL | T150 | 0.025 | 98.68 | 377 | 0.017075 | 0.025 | 0.0171 | 1 |
| Q6UX07 | DHRS13 | APRGGAQtPLYCALQ | T264 | 0.017 | 94.81 | 377 | 0.011611 | 0.037 | 0.0253 | 0.8 |
| Q6ZT12 | UBR3 | NMLPDQEyKVAFTKT | Y411 | 0.002 | 92.19 | 1888 | 0.000877 | 0.304 | 0.1297 | 1 |
| Q6UXV1 | IZUMO2 | AsFVKNQtQHLMGNS | T93 | 0.052 | 95.94 | 221 | 0.035517 | 0.052 | 0.0355 | 1 |
| Q6ICB4 | PHETA2 | LKGNLLFsFEsREGR | S54 | 0.069 | 95.49 | 259 | 0.054849 | 0.069 | 0.0548 | 1 |
| Q8TAV0 | FAM76A | RFPFEALsQGQQLCK | S20 | 0.174 | 87.79 | 307 | 0.138315 | 0.174 | 0.1383 | 1 |
| Q9NRX3 | NDUFA4L2 | FLAVSTDyKKLKKDR | Y77 | 0.251 | 84.82 | 87 | 0.11011 | 0.251 | 0.1101 | 1 |
| P00338 | LDHA | HGSLFLRtPkIVSGk | T74 | 0.04 | 98.55 | 332 | 0.027321 | 0.06 | 0.0387 | 1 |
| P00338 | LDHA | kIVSGkDyNVTANsk | Y83 | 0.137 | 98.17 | 332 | 0.0601 | 0.137 | 0.0601 | 1 |

| | | | | | | | | | | |
|---|---|---|---|---|---|---|---|---|---|---|
| P00338 | LDHA | DyNVTANskLVIItA | S89 | 0.029 | 98.78 | 332 | 0.023053 | 0.029 | 0.0231 | 1 |
| P00338 | LDHA | IPNVVkysPNCkLLI | S128 | 0.02 | 98.61 | 332 | 0.015898 | 0.02 | 0.0159 | 1 |
| P00338 | LDHA | PkNrVIGsGCNLDsA | S161 | 0.055 | 98.74 | 332 | 0.04372 | 0.07 | 0.0601 | 1 |
| P00338 | LDHA | IkLkGYtsWAIGLSV | S249 | 0.066 | 97.77 | 332 | 0.052464 | 0.066 | 0.0525 | 1 |
| P00338 | LDHA | EEARLkksADtLWGI | S319 | 0.003 | 95.99 | 332 | 0.002385 | 0.003 | 0.0024 | 1 |
| Q5TYX0 | PRAMEF5 | YPAPRESyDADGTLC | Y420 | 0.328 | 93.54 | 476 | 0.143889 | 0.328 | 0.1439 | 1 |
| P07992 | ERCC1 | KSNSIIVsPRQRGNP | S104 | 0.022 | 97.21 | 297 | 0.017488 | 0.022 | 0.0175 | 1 |
| Q02543 | RPL18A | ASGtLrEykVVGRCL | Y10 | 0.155 | 98.17 | 176 | 0.067996 | 0.155 | 0.068 | 1 |
| Q02543 | RPL18A | RHRARAHsIQIMkVE | S123 | 0.166 | 95.57 | 176 | 0.131956 | 0.166 | 0.132 | 1 |
| O60462 | NRP2 | HGDDNGWtPNLDSNK | T319 | 0.183 | 98.52 | 931 | 0.124991 | 0.183 | 0.125 | 1 |
| Q9Y4F5 | CEP170B | KQHAVINyDQDRDEH | Y48 | 0.193 | 93 | 1589 | 0.084666 | 0.193 | 0.0847 | 1 |
| Q92841 | DDX17 | VGIAQTGsGKTLAYL | S219 | 0.157 | 84.26 | 729 | 0.124802 | 0.157 | 0.1248 | 1 |
| Q92841 | DDX17 | VQQVADDyGkCSRLk | Y267 | 0.059 | 90.24 | 729 | 0.025882 | 0.059 | 0.0259 | 1 |
| Q92841 | DDX17 | RLkStCIyGGAPkGP | Y279 | 0.336 | 94.72 | 729 | 0.147398 | 0.336 | 0.1474 | 0.8182 |
| Q92841 | DDX17 | GVEICIAtPGRLIDF | T301 | 0.135 | 93.96 | 729 | 0.092207 | 0.135 | 0.0922 | 1 |
| Q92841 | DDX17 | TNLRRCTyLVLDEAD | Y321 | 0.011 | 97.03 | 729 | 0.004826 | 0.011 | 0.0048 | 1 |
| Q92841 | DDX17 | DrQTLMWsATWPkEV | S356 | 0.017 | 91.39 | 729 | 0.013514 | 0.017 | 0.0135 | 0.8889 |
| Q92841 | DDX17 | NYDYPNssEDyVHrI | S499 | 0.021 | 88.73 | 729 | 0.016693 | 0.021 | 0.0167 | 0.875 |
| Q92841 | DDX17 | YPNssEDyVHrIGRT | Y502 | 0.013 | 92.34 | 729 | 0.005703 | 0.013 | 0.0057 | 1 |
| Q92841 | DDX17 | GTAYTFFtPGNLkQA | T523 | 0.121 | 94.3 | 729 | 0.082645 | 0.121 | 0.0826 | 1 |
| Q96JH7 | VCPIP1 | NYHCkLLsPILARyG | S131 | 0.002 | 95.42 | 1222 | 0.00159 | 0.002 | 0.0016 | 0.8 |
| Q96JH7 | VCPIP1 | YGkDRsGsLLYLHDT | S187 | 0 | 83.22 | 1222 | 0 | 0 | 0 | 1 |
| P01215 | CGA | CFSRAYPtPLRSKKT | T63 | 0.09 | 97.82 | 116 | 0.061471 | 0.09 | 0.0615 | 1 |
| Q96C11 | FGGY | PDFHGNRsPLADLTL | S397 | 0.051 | 97.25 | 551 | 0.040541 | 0.051 | 0.0405 | 1 |
| P21731 | TBXA2R | VMIFFGLsPLLLGAA | S118 | 0 | 98.34 | 343 | 0 | 0 | 0 | 0.9 |
| Q9BWU1 | CDK19 | DIRKMPEyPTLQKDF | Y276 | 0.308 | 96.19 | 502 | 0.135115 | 0.308 | 0.1351 | 1 |
| Q9BR76 | CORO1B | PVKNDQCyEDIrVSR | Y26 | 0.037 | 98.43 | 489 | 0.016231 | 0.037 | 0.0162 | 1 |
| Q9BR76 | CORO1B | CSACKDKsVRIIDPR | S199 | 0.051 | 98.01 | 489 | 0.040541 | 0.065 | 0.0649 | 1 |
| Q14689 | DIP2A | GLWHGVLtSVMNRMH | T569 | 0.003 | 97.03 | 1571 | 0.002049 | 0.003 | 0.002 | 1 |
| Q15645 | TRIP13 | GFIENNVsKLsLLLN | S367 | 0.001 | 94.27 | 432 | 0.000795 | 0.001 | 0.0008 | 1 |
| Q15645 | TRIP13 | ENNVsKLsLLLNDIs | S370 | 0.001 | 97.12 | 432 | 0.000795 | 0.001 | 0.0008 | 1 |

| | | | | | | | | | | |
|---|---|---|---|---|---|---|---|---|---|---|
| Q15645 | TRIP13 | sLLLNDIsRKSEGLs | S377 | 0 | 97.78 | 432 | 0 | 0 | 0 | 1 |
| Q8NEZ3 | WDR19 | YkEAVVAyENAKQWQ | Y918 | 0.084 | 85.81 | 1342 | 0.03685 | 0.084 | 0.0369 | 1 |
| Q96I15 | SCLY | IINAAREsLAKMIGG | S81 | 0.169 | 98.89 | 445 | 0.134341 | 0.169 | 0.1343 | 1 |
| Q5TCX8 | MAP3K21 | FGQVYRAtWQGQEVA | T142 | 0.19 | 97.61 | 1036 | 0.129773 | 0.19 | 0.1298 | 1 |
| Q8TAA3 | PSMA8 | EAVkKGstAVGIRGT | T33 | 0.006 | 89.71 | 256 | 0.004098 | 0.006 | 0.0041 | 1 |
| Q8TAA3 | PSMA8 | PVTVEyItRFIATLK | T116 | 0 | 92.03 | 256 | 0 | 0 | 0 | 1 |
| Q86WZ0 | HEATR4 | VRLEACRsILALKLQ | S820 | 0.047 | 95.44 | 1026 | 0.037361 | 0.047 | 0.0374 | 0.8333 |
| P54277 | PMS1 | EGEAVRLsRQLPMyL | S885 | 0.042 | 86.16 | 932 | 0.033386 | 0.042 | 0.0334 | 1 |
| Q96DI7 | SNRNP40 | NLLRCSWsPDGSkIA | S294 | 0.073 | 94.73 | 357 | 0.058029 | 0.073 | 0.058 | 0.8333 |
| Q96DI7 | SNRNP40 | kLPGHAGsINEVAFH | S329 | 0.024 | 94.92 | 357 | 0.019078 | 0.024 | 0.0191 | 0.8 |
| Q9P2J5 | LARS1 | DIGDTIIyLVH____ | Y1173 | 0.002 | 91.86 | 1176 | 0.000877 | 0.002 | 0.0009 | 1 |
| Q9HCE3 | ZNF532 | IHQHKSPyTCPECGA | Y811 | 0.111 | 78.84 | 1301 | 0.048694 | 0.31 | 0.136 | 1 |
| P54277 | PMS1 | AtVRLLssSQIITSV | S14 | 0 | 92.89 | 932 | 0 | 0 | 0 | 1 |
| P54277 | PMS1 | LKLIRHHyNLkCLKE | Y280 | 0 | 86.77 | 932 | 0 | 0 | 0 | 1 |
| Q14807 | KIF22 | kGLRLkEsGAINTSL | S294 | 0.011 | 83.61 | 665 | 0.008744 | 0.011 | 0.0087 | 1 |
| O00308 | WWP2 | HLDLKVWsCHtLRNE | S87 | 0.019 | 88.41 | 870 | 0.015103 | 0.019 | 0.0151 | 0.8571 |
| P18850 | ATF6 | AIRRRGDtFYVVSFR | T565 | 0.03 | 93.03 | 670 | 0.02049 | 0.053 | 0.0362 | 1 |
| Q96DU3 | SLAMF6 | KMEDTGSyRAQISTK | Y107 | 0.029 | 95.89 | 332 | 0.012722 | 0.029 | 0.0127 | 1 |
| Q6P4Q7 | CNNM4 | yyARHyLyTRNKPAD | Y603 | 0.024 | 94.02 | 775 | 0.010528 | 0.024 | 0.0105 | 0.8571 |
| Q6P4Q7 | CNNM4 | RALVDLQyIKITRQQ | Y708 | 0.172 | 93.93 | 775 | 0.075454 | 0.172 | 0.0755 | 0.8 |
| Q16787 | LAMA3 | HMSRIAMyELLADAD | Y1012 | 0.035 | 89.085 | 3333 | 0.015354 | 0.035 | 0.0154 | 1 |
| Q96BQ5 | CCDC127 | kEFEAVLtERQNIYC | T167 | 0.18 | 96 | 260 | 0.122942 | 0.18 | 0.1229 | 1 |
| Q9BTE3 | MCMBP | HVLHFGkyRDVAECG | Y102 | 0.048 | 89.81 | 642 | 0.021057 | 0.048 | 0.0211 | 1 |
| Q9BTE3 | MCMBP | QQELDLNsPRNTTLE | S118 | 0.12 | 82.09 | 642 | 0.09539 | 0.12 | 0.0954 | 1 |
| Q9BTE3 | MCMBP | STFTEHLyRIIQHLV | Y407 | 0.301 | 95.4 | 642 | 0.132044 | 0.301 | 0.132 | 1 |
| A4D0S4 | LAMB4 | QNDsLDKyyyALyEM | Y252 | 0.171 | 86.67 | 1761 | 0.075015 | 0.171 | 0.075 | 1 |
| A4D0S4 | LAMB4 | NDsLDKyyyALyEMI | Y253 | 0.106 | 91.21 | 1761 | 0.046501 | 0.106 | 0.0465 | 1 |
| A4D0S4 | LAMB4 | DsLDKyyyALyEMIV | Y254 | 0.009 | 93.97 | 1761 | 0.003948 | 0.009 | 0.0039 | 1 |
| Q9NVM4 | PRMT7 | ALVLDIGtGTGLLSM | T73 | 0.088 | 98.17 | 692 | 0.060105 | 0.088 | 0.0601 | 1 |
| Q9H9E3 | COG4 | ELRSLIAyLTTVTTW | Y714 | 0.247 | 94.07 | 785 | 0.108355 | 0.247 | 0.1084 | 1 |
| Q96AP4 | ZUP1 | GNLKHKQyQILAVEG | Y548 | 0.006 | 98.65 | 578 | 0.002632 | 0.006 | 0.0026 | 1 |

| | | | | | | | | | | |
|---|---|---|---|---|---|---|---|---|---|---|
| Q96AP4 | ZUP1 | KLARRQAsQVFTAEK | S569 | 0.037 | 96.96 | 578 | 0.029412 | 0.037 | 0.0294 | 1 |
| P51659 | HSD17B4 | DGRVVLVtGAGAGLG | T15 | 0.027 | 98.89 | 736 | 0.018441 | 0.027 | 0.0184 | 1 |
| P51659 | HSD17B4 | EHMkKQKyGRIIMts | Y143 | 0.121 | 98.85 | 736 | 0.053081 | 0.215 | 0.0877 | 1 |
| P51659 | HSD17B4 | KyGRIIMtssASGIy | T149 | 0.084 | 98.82 | 736 | 0.057373 | 0.141 | 0.0933 | 1 |
| P51659 | HSD17B4 | yGRIIMtssASGIyG | S150 | 0.027 | 98.45 | 736 | 0.021463 | 0.027 | 0.0215 | 1 |
| O60256 | PRPSAP2 | AGLTHLItMDLHQKE | T144 | 0.001 | 98.12 | 369 | 0.000683 | 0.001 | 0.0007 | 1 |
| O60256 | PRPSAP2 | IVAksPAsAkRAQsF | S192 | 0.06 | 90.98 | 369 | 0.047695 | 0.06 | 0.0477 | 1 |
| Q5T0N1 | CFAP70 | AKACYERtIsFVVDA | T989 | 0.017 | 94.77 | 1121 | 0.011611 | 0.017 | 0.0116 | 0.9167 |
| Q14571 | ITPR2 | LLGEIVKysNVIQLL | Y117 | 0.058 | 91.17 | 2701 | 0.025444 | 0.058 | 0.0254 | 1 |
| Q14571 | ITPR2 | QSATSAtssKALWEI | S277 | 0.103 | 84.465 | 2701 | 0.081876 | 0.163 | 0.1296 | 1 |
| Q9BTV5 | FSD1 | EDsKIDHyVLEyRRT | Y198 | 0.137 | 97.55 | 496 | 0.0601 | 0.137 | 0.0601 | 1 |
| Q92824 | PCSK5 | ysKCPEGsyAEDGIC | S1270 | 0.123 | 84.04 | 1860 | 0.097775 | 0.123 | 0.0978 | 0.8571 |
| Q6UXX5 | ITIH6 | MTSYSMRstVVsRYA | S44 | 0.03 | 97.13 | 1313 | 0.023847 | 0.03 | 0.0238 | 1 |
| Q6UXX5 | ITIH6 | SMRstVVsRYAHTLV | S48 | 0 | 93.46 | 1313 | 0 | 0 | 0 | 1 |
| P12110 | COL6A2 | TRVGVVQysHEGTFE | Y665 | 0.003 | 94.36 | 1019 | 0.001316 | 0.003 | 0.0013 | 1 |
| P12110 | COL6A2 | RVGVVQysHEGTFEA | S666 | 0 | 91.34 | 1019 | 0 | 0 | 0 | 1 |
| P12110 | COL6A2 | LEWIAGGtWtPsALK | T701 | 0.184 | 88.68 | 1019 | 0.125675 | 0.184 | 0.1257 | 1 |
| P12110 | COL6A2 | WIAGGtWtPsALKFA | T703 | 0.029 | 91.73 | 1019 | 0.019807 | 0.029 | 0.0198 | 1 |
| Q04917 | YWHAH | rNLLsVAykNVVGAR | Y49 | 0.071 | 98.74 | 246 | 0.031147 | 0.071 | 0.0311 | 0.9333 |
| Q04917 | YWHAH | QYEskVFyLkMkGDy | Y123 | 0.063 | 98.72 | 246 | 0.027637 | 0.126 | 0.0553 | 0.8333 |
| Q04917 | YWHAH | VEASEAAykEAFEIS | Y154 | 0 | 98.85 | 246 | 0 | 0 | 0 | 0.9333 |
| Q04917 | YWHAH | SKEQMQPtHPIRLGL | T168 | 0.066 | 96.55 | 246 | 0.045079 | 0.061 | 0.059 | 1 |
| Q13148 | TARDBP | MsEyIRVtEDENDEP | T8 | 0.045 | 87.52 | 414 | 0.030736 | 0.045 | 0.0307 | 0.8182 |
| Q13148 | TARDBP | IPSEDDGtVLLSTVT | T25 | 0.098 | 86.12 | 414 | 0.066935 | 0.151 | 0.1031 | 1 |
| Q02224 | CENPE | TPVsFDEtLTALQFA | T316 | 0.124 | 90.015 | 2701 | 0.084694 | 0.124 | 0.0847 | 0.9091 |
| Q96SZ6 | CDK5RAP1 | HLPAQSGsSRVLEAM | S406 | 0.019 | 98.5 | 601 | 0.015103 | 0.019 | 0.0151 | 1 |
| Q96SZ6 | CDK5RAP1 | RGYSREAyVELVHHI | Y422 | 0.029 | 97.55 | 601 | 0.012722 | 0.029 | 0.0127 | 0.9412 |
| O43865 | AHCYL1 | RQVDVVItCtGNKNV | T372 | 0.003 | 98.21 | 530 | 0.002049 | 0.003 | 0.002 | 1 |
| Q9UJ83 | HACL1 | VEACRLYtkFsARPs | T135 | 0.202 | 98.04 | 578 | 0.137969 | 0.202 | 0.138 | 1 |
| Q9UJ83 | HACL1 | YkLPFLPtPMGkGVV | T247 | 0.194 | 98.27 | 578 | 0.132505 | 0.194 | 0.1325 | 1 |
| Q9UJ83 | HACL1 | KKSLPMNyYTVFyHV | Y377 | 0.051 | 96.7 | 578 | 0.022373 | 0.051 | 0.0224 | 1 |

| | | | | | | | | | | |
|---|---|---|---|---|---|---|---|---|---|---|
| Q9UJ83 | HACL1 | LQkSLRQsLADttkP | S545 | 0.005 | 98.29 | 578 | 0.003975 | 0.005 | 0.004 | 1 |
| Q9UJ83 | HACL1 | LADttkPsLINIMIE | S553 | 0.009 | 98.79 | 578 | 0.007154 | 0.009 | 0.0072 | 1 |
| Q4ZHG4 | FNDC1 | KVVAsRQytVRYREK | Y297 | 0.044 | 91.68 | 1894 | 0.019302 | 0.044 | 0.0193 | 1 |
| O14974 | PPP1R12A | IKDYDGWtPLHAAAH | T234 | 0.024 | 97.45 | 1030 | 0.016392 | 0.024 | 0.0164 | 1 |
| O60662 | KLHL41 | APMKIPRsMFGVAVH | S486 | 0.026 | 98.51 | 606 | 0.020668 | 0.026 | 0.0207 | 1 |
| P51801 | CLCNKB | AGVVLEDyLDIKNFG | Y138 | 0.082 | 86.44 | 687 | 0.035972 | 0.082 | 0.036 | 1 |
| P51801 | CLCNKB | ILGRNIGsHRVRVEH | S542 | 0.143 | 82.38 | 687 | 0.113673 | 0.143 | 0.1137 | 0.8 |
| Q9NVR7 | TBCCD1 | KIHRCNEsFIyLLSP | S340 | 0.002 | 91.65 | 557 | 0.00159 | 0.002 | 0.0016 | 1 |
| Q9NVR7 | TBCCD1 | RCNEsFIyLLSPLRS | Y343 | 0.016 | 95.69 | 557 | 0.007019 | 0.016 | 0.007 | 0.8462 |
| Q96LI6 | HSFY1 | VLSKLKFyyNPNFKR | Y175 | 0.072 | 96.71 | 401 | 0.031585 | 0.072 | 0.0316 | 1 |
| Q96MU7 | YTHDC1 | QGFARLssEsHHGGs | S417 | 0.143 | 93.26 | 727 | 0.113673 | 0.143 | 0.1137 | 1 |
| Q96MU7 | YTHDC1 | FARLssEsHHGGsPI | S419 | 0.172 | 94.64 | 727 | 0.136725 | 0.172 | 0.1367 | 1 |
| P30419 | NMT1 | EEASKRsyQFWDTQP | Y117 | 0.304 | 90.62 | 496 | 0.13336 | 0.304 | 0.1334 | 1 |
| P30419 | NMT1 | YTLLNENyVEDDDNM | Y180 | 0.3 | 87.7 | 496 | 0.131606 | 0.3 | 0.1316 | 0.8182 |
| O14795 | UNC13B | yMRRFAKtIGKVLMQ | T1144 | 0.003 | 96.05 | 1591 | 0.002049 | 0.003 | 0.002 | 1 |
| O14795 | UNC13B | IGKVLMQyADILSKD | Y1152 | 0 | 97.38 | 1591 | 0 | 0 | 0 | 0.8667 |
| Q15113 | PCOLCE | QVTLRMTtDEGtGGR | T133 | 0.198 | 95.47 | 449 | 0.135237 | 0.198 | 0.1352 | 1 |
| P50750 | CDK9 | FCDEVSkyEkLAkIG | Y19 | 0.141 | 97.73 | 372 | 0.061855 | 0.141 | 0.0619 | 1 |
| Q6ICC9 | RTL6 | IPHMQPDsPLRNNyQ | S160 | 0.161 | 93.58 | 239 | 0.127981 | 0.161 | 0.128 | 1 |
| A8TX70 | COL6A5 | RLGLMSYsNSAKTIS | S280 | 0.057 | 89.55 | 2615 | 0.04531 | 0.114 | 0.0906 | 1 |
| A8TX70 | COL6A5 | LsIQVGKsNtGAAID | S312 | 0 | 87.32 | 2615 | 0 | 0 | 0 | 1 |
| A8TX70 | COL6A5 | IQVGKsNtGAAIDQM | T314 | 0.033 | 93.95 | 2615 | 0.022539 | 0.033 | 0.0225 | 1 |
| A8TX70 | COL6A5 | TQLEEIVsyPPEQTI | S386 | 0.106 | 90.26 | 2615 | 0.084261 | 0.175 | 0.1391 | 1 |
| P48426 | PIP4K2A | SHFkFkEyCPMVFRN | Y93 | 0.178 | 98.82 | 406 | 0.078086 | 0.178 | 0.0781 | 1 |
| Q8WXH2 | JPH3 | GTKEEGKyKQNILVG | Y338 | 0.011 | 92.82 | 748 | 0.004826 | 0.011 | 0.0048 | 1 |
| Q8IWZ3 | ANKHD1 | KNGERMItIrGGTES | T1744 | 0.115 | 85.93 | 2542 | 0.078547 | 0.115 | 0.0785 | 1 |
| Q13332 | PTPRS | RyANVIAyDHsRVIL | Y1428 | 0.095 | 96 | 1948 | 0.041675 | 0.12 | 0.0526 | 1 |
| Q13332 | PTPRS | RMVWEQRsAtIVMMT | S1484 | 0.032 | 96.98 | 1948 | 0.025437 | 0.032 | 0.0254 | 1 |
| Q13332 | PTPRS | VWEQRsAtIVMMTrL | T1486 | 0.003 | 98.01 | 1948 | 0.002049 | 0.003 | 0.002 | 1 |
| Q13332 | PTPRS | RLVNIMPyEStRVCL | Y1717 | 0.094 | 89.73 | 1948 | 0.041236 | 0.23 | 0.0628 | 0.8571 |
| Q13332 | PTPRS | GREKCHQyWPAERSA | Y1793 | 0.026 | 96.36 | 1948 | 0.011406 | 0.026 | 0.0114 | 1 |

| | | | | | | | | | | |
|---|---|---|---|---|---|---|---|---|---|---|
| Q9UPR0 | PLCL2 | LKKIVttyDMMIQSL | Y1004 | 0 | 95.5 | 1127 | 0 | 0 | 0 | 1 |
| P17752 | TPH1 | SNFLKERtGFsIRPV | T225 | 0.119 | 98.22 | 444 | 0.081279 | 0.119 | 0.0813 | 1 |
| P17752 | TPH1 | ITTFQDVyFVSESFE | Y373 | 0.005 | 97.63 | 444 | 0.002193 | 0.005 | 0.0022 | 1 |
| Q9Y6M1 | IGF2BP2 | IKNITkQtQSRVDIH | T223 | 0.105 | 95.39 | 599 | 0.071716 | 0.105 | 0.0717 | 1 |
| Q96E52 | OMA1 | kEQFRLLsELEYEAW | S237 | 0.026 | 88 | 524 | 0.020668 | 0.026 | 0.0207 | 1 |
| Q8WVE0 | EEF1AKMT1 | SETVkyLtRGKILLC | T161 | 0.15 | 98.41 | 214 | 0.102452 | 0.15 | 0.1025 | 0.9 |
| Q8N2U9 | SLC66A2 | EsPLLWQsAIMILTM | S81 | 0.003 | 96.84 | 271 | 0.002385 | 0.003 | 0.0024 | 1 |
| Q7Z4L5 | TTC21B | CAEIAkHsVAQRDyE | S894 | 0.102 | 87.27 | 1316 | 0.081081 | 0.102 | 0.0811 | 1 |
| Q6ZUX3 | TOGARAM2 | LSGTRDstDMLVHNL | T652 | 0.06 | 88.05 | 1019 | 0.040981 | 0.06 | 0.041 | 1 |
| Q12889 | OVGP1 | NREKFIAsVISLLRT | S123 | 0.169 | 98.8 | 678 | 0.134341 | 0.169 | 0.1343 | 1 |
| Q16394 | EXT1 | GEKIAESyQNILAAI | Y131 | 0.028 | 95.94 | 746 | 0.012283 | 0.028 | 0.0123 | 1 |
| Q16394 | EXT1 | SKWTNDYsMVLTGAA | S617 | 0.007 | 97.26 | 746 | 0.005564 | 0.007 | 0.0056 | 1 |
| O75462 | CRLF1 | DFLFQAKyQIRYRVE | Y272 | 0.029 | 94.14 | 422 | 0.012722 | 0.029 | 0.0127 | 1 |
| O60566 | BUB1B | IELGNEDyCIkREYL | Y766 | 0.079 | 86.93 | 1050 | 0.034656 | 0.079 | 0.0347 | 0.8 |
| O43508 | TNFSF12 | LLALRPGsSLRIRTL | S222 | 0.082 | 98.32 | 249 | 0.065183 | 0.082 | 0.0652 | 1 |
| O43314 | PPIP5K2 | ISKIPDIyDCIKYDV | Y727 | 0.114 | 95.04 | 1243 | 0.05001 | 0.298 | 0.1307 | 0.9 |
| Q86WA8 | LONP2 | MPQSMPEyALTRNYL | Y307 | 0.267 | 88.17 | 852 | 0.117129 | 0.295 | 0.1294 | 1 |
| Q6J9G0 | STYK1 | CTEKLPLyMVLEDVA | Y191 | 0.26 | 95.39 | 422 | 0.114058 | 0.26 | 0.1141 | 1 |
| Q6J9G0 | STYK1 | DGLLyDLtEKQVYHI | T224 | 0.07 | 93.02 | 422 | 0.047811 | 0.07 | 0.0478 | 1 |
| Q16836 | HADH | TPMTSQktFEsLVDF | T193 | 0.043 | 97.96 | 314 | 0.02937 | 0.043 | 0.0294 | 1 |
| Q16836 | HADH | TSQktFEsLVDFskA | S196 | 0.029 | 98.24 | 314 | 0.023053 | 0.029 | 0.0231 | 0.875 |
| Q16836 | HADH | FEsLVDFskALGkHP | S201 | 0 | 98.77 | 314 | 0 | 0 | 0 | 1 |
| Q16836 | HADH | DyVGLDttKFIVDGW | T270 | 0.041 | 97.78 | 314 | 0.028004 | 0.041 | 0.028 | 1 |
| Q16836 | HADH | ENPLHQPsPSLNKLV | S290 | 0.019 | 98.74 | 314 | 0.015103 | 0.019 | 0.0151 | 1 |
| Q13492 | PICALM | LNEkAVSyRQVAFDF | Y138 | 0.204 | 98.05 | 652 | 0.089492 | 0.321 | 0.1196 | 0.9091 |
| Q9NUJ1 | ABHD10 | DySGVGSsDGNSEES | S118 | 0.134 | 98.12 | 306 | 0.106519 | 0.134 | 0.1065 | 1 |
| Q9BUL9 | RPP25 | ATRAIVFsGCGrAtT | S68 | 0.079 | 97.06 | 199 | 0.062798 | 0.079 | 0.0628 | 1 |
| O00459 | PIK3R2 | LRKIRDQyLVWLtQK | Y577 | 0.075 | 88.58 | 728 | 0.032901 | 0.075 | 0.0329 | 1 |
| O43323 | DHH | PERTLGAsGPAEGRV | S60 | 0.018 | 93.99 | 396 | 0.014308 | 0.018 | 0.0143 | 0.8182 |
| Q49AG3 | ZBED5 | SWLLRLAyLADIFtK | Y476 | 0.003 | 96.86 | 693 | 0.001316 | 0.116 | 0.0509 | 1 |
| Q49AG3 | ZBED5 | LVARDyEsLIDLTSD | S594 | 0.181 | 95.82 | 693 | 0.14388 | 0.181 | 0.1439 | 1 |

| | | | | | | | | | | |
|---|---|---|---|---|---|---|---|---|---|---|
| P09467 | FBP1 | FDPAVTEyIQRKkFP | Y227 | 0.003 | 98.14 | 338 | 0.001316 | 0.003 | 0.0013 | 1 |
| P09467 | FBP1 | MVADVHRtLVyGGIF | T256 | 0.05 | 98.83 | 338 | 0.034151 | 0.05 | 0.0342 | 1 |
| P09467 | FBP1 | VyGGIFLyPANkKsP | Y265 | 0.264 | 97.93 | 338 | 0.115813 | 0.264 | 0.1158 | 1 |
| P09467 | FBP1 | EKAGGMAttGKEAVL | T297 | 0.08 | 98.6 | 338 | 0.054641 | 0.08 | 0.0546 | 1 |
| P09467 | FBP1 | RAPVILGsPDDVLEF | S321 | 0 | 98.84 | 338 | 0 | 0 | 0 | 1 |
| Q03169 | TNFAIP2 | HLHLVKEyIIQLSkG | Y529 | 0.006 | 96.8 | 654 | 0.002632 | 0.006 | 0.0026 | 1 |
| P29350 | PTPN6 | DPTSERWyHGHMSGG | Y111 | 0.09 | 92.77 | 595 | 0.039482 | 0.09 | 0.0395 | 1 |
| P29350 | PTPN6 | MCEGGRytVGGLETF | T177 | 0.084 | 95.36 | 595 | 0.057373 | 0.084 | 0.0574 | 1 |
| P29350 | PTPN6 | SNIPGsDyINANyIk | Y301 | 0.008 | 98.82 | 595 | 0.003509 | 0.008 | 0.0035 | 1 |
| P29350 | PTPN6 | GRNkCVPyWPEVGMQ | Y364 | 0 | 98.68 | 595 | 0 | 0 | 0 | 1 |
| P29350 | PTPN6 | EVGMQRAyGPysVTN | Y374 | 0.143 | 98.4 | 595 | 0.062732 | 0.143 | 0.0627 | 1 |
| P29350 | PTPN6 | MQRAyGPysVTNCGE | Y377 | 0.16 | 98.7 | 595 | 0.07019 | 0.16 | 0.0702 | 1 |
| P29350 | PTPN6 | QRAyGPysVTNCGEH | S378 | 0.135 | 98.58 | 595 | 0.107313 | 0.135 | 0.1073 | 1 |
| Q9UQ53 | MGAT4B | SGLLEVIsPSPHFYP | S216 | 0.022 | 97.89 | 548 | 0.017488 | 0.022 | 0.0175 | 0.9 |
| P23497 | SP100 | TNkMFEDsQDSCRNL | S92 | 0.097 | 90.78 | 879 | 0.077107 | 0.097 | 0.0771 | 1 |
| P16455 | MGMT | KDCEMkRttLDsPLG | T10 | 0.066 | 95.53 | 207 | 0.045079 | 0.066 | 0.0451 | 1 |
| P16455 | MGMT | MkRttLDsPLGkLEL | S14 | 0.007 | 93.42 | 207 | 0.005564 | 0.007 | 0.0056 | 1 |
| P16455 | MGMT | PLGkLELsGCEQGLH | S22 | 0.13 | 97.78 | 207 | 0.103339 | 0.13 | 0.1033 | 1 |
| Q14185 | DOCK1 | VFDPkELsKMFTEFI | S818 | 0 | 95.9 | 1865 | 0 | 0 | 0 | 1 |
| P35609 | ACTN2 | ILAsDKPyILAEELR | Y844 | 0.317 | 92.22 | 894 | 0.139063 | 0.317 | 0.1391 | 0.9 |
| Q6AI12 | ANKRD40 | GEMPVQLtSRREIRk | T84 | 0.095 | 91.5 | 368 | 0.064886 | 0.095 | 0.0649 | 0.8889 |
| Q8TF05 | PPP4R1 | WISYkLVsEMVKKLH | S806 | 0 | 96.38 | 950 | 0 | 0 | 0 | 1 |
| Q8TF05 | PPP4R1 | LAkTLRQtLLEKDYF | T892 | 0.126 | 91.41 | 950 | 0.08606 | 0.126 | 0.0861 | 1 |
| Q6ZT98 | TTLL7 | KKKRKQKtFIVKPAN | T156 | 0.015 | 92.04 | 887 | 0.010245 | 0.015 | 0.0102 | 0.875 |
| Q6ZT98 | TTLL7 | KRSIKWFtEFLQANQ | T278 | 0.21 | 95.21 | 887 | 0.143433 | 0.21 | 0.1434 | 1 |
| Q13619 | CUL4A | MLSDLQVyKDSFELk | Y227 | 0.08 | 96.25 | 759 | 0.035095 | 0.14 | 0.0614 | 1 |
| Q13619 | CUL4A | QEREVPEyLNHVSkR | Y259 | 0.02 | 97.57 | 759 | 0.008774 | 0.02 | 0.0088 | 1 |
| Q7Z5Q5 | POLN | FEDLVEKyCEKsItV | Y372 | 0.284 | 95.24 | 900 | 0.124587 | 0.284 | 0.1246 | 1 |
| Q969V4 | TEKT1 | VDRILSQtANDLRKQ | T249 | 0.166 | 96.67 | 418 | 0.11338 | 0.166 | 0.1134 | 1 |
| Q8IZY2 | ABCA7 | GGMQRKLsVAIAFVG | S949 | 0.072 | 89.73 | 2146 | 0.057234 | 0.091 | 0.0912 | 1 |
| Q8IZY2 | ABCA7 | AIAFVGGsQVVILDE | S958 | 0.006 | 87.29 | 2146 | 0.004769 | 0.006 | 0.0048 | 1 |

| | | | | | | | | | | |
|---|---|---|---|---|---|---|---|---|---|---|
| P35670 | ATP7B | EAALAVHtLQsMGVD | T1208 | 0.137 | 88.56 | 1465 | 0.093573 | 0.137 | 0.0936 | 1 |
| P35670 | ATP7B | GVDVVLItGDNRKTA | T1220 | 0.027 | 89.36 | 1465 | 0.018441 | 0.027 | 0.0184 | 1 |
| Q9UPW8 | UNC13A | DkkRKFAtKSKNNSW | T1587 | 0.119 | 94.52 | 1703 | 0.081279 | 0.166 | 0.131 | 0.8889 |
| Q96NN9 | AIFM3 | GIGAVPAtGFLRQsG | T433 | 0.056 | 98.52 | 605 | 0.038249 | 0.056 | 0.0382 | 1 |
| Q96NN9 | AIFM3 | AtGFLRQsGIGLDsR | S439 | 0.072 | 97.39 | 605 | 0.057234 | 0.072 | 0.0572 | 1 |
| Q96NN9 | AIFM3 | FAAGDAVtFPLAWRN | T470 | 0.08 | 98.63 | 605 | 0.054641 | 0.08 | 0.0546 | 1 |
| Q15582 | TGFBI | yQLVLQHsRLRGRQH | S37 | 0.025 | 96.21 | 683 | 0.019873 | 0.025 | 0.0199 | 1 |
| Q15582 | TGFBI | YHGQTLEtLGGKKLR | T457 | 0 | 98 | 683 | 0 | 0 | 0 | 1 |
| Q15582 | TGFBI | TLNREGVyTVFAPTN | Y537 | 0.267 | 97.21 | 683 | 0.117129 | 0.267 | 0.1171 | 1 |
| Q15582 | TGFBI | GALVRLksLQGDKLE | S591 | 0.03 | 97.34 | 683 | 0.023847 | 0.03 | 0.0238 | 1 |
| P21439 | ABCB4 | LMKKEGVyFKLVNMQ | Y624 | 0.011 | 88.34 | 1286 | 0.004826 | 0.011 | 0.0048 | 1 |
| Q5JPI3 | C3orf38 | GIVIPPAtEkHNLIQ | T86 | 0.21 | 81.09 | 329 | 0.143433 | 0.21 | 0.1434 | 1 |
| Q49AJ0 | FAM135B | FADFDtMtDRLLDEI | T1191 | 0 | 97.94 | 1406 | 0 | 0 | 0 | 0.8 |
| Q00610 | CLTC | LQkyIEIyVQkVNPS | Y803 | 0.02 | 80.91 | 1675 | 0.008774 | 0.02 | 0.0088 | 1 |
| Q8WYN3 | CSRNP3 | RHNSVRQyTLGEFAR | Y104 | 0.317 | 96.35 | 585 | 0.139063 | 0.317 | 0.1391 | 0.8571 |
| Q9H172 | ABCG4 | SyLSyVRyGFEGVIL | Y573 | 0.27 | 94.3 | 646 | 0.118445 | 0.27 | 0.1184 | 1 |
| Q9H7D0 | DOCK5 | KENRMsCtVNVLNFY | T1224 | 0.023 | 88.58 | 1870 | 0.015709 | 0.172 | 0.1148 | 1 |
| Q9H7D0 | DOCK5 | YERREDFsLRLLTQF | S1395 | 0.024 | 88.61 | 1870 | 0.019078 | 0.024 | 0.0191 | 1 |
| Q92624 | APPBP2 | YSGLEYDyRGLIkLY | Y517 | 0.061 | 97.77 | 585 | 0.02676 | 0.061 | 0.0268 | 1 |
| P16615 | ATP2A2 | VETLGCtsVICSDkT | S346 | 0.144 | 95.14 | 1042 | 0.114468 | 0.144 | 0.1145 | 1 |
| P16615 | ATP2A2 | CNDSALDyNEAkGVY | Y427 | 0.33 | 94.89 | 1042 | 0.144766 | 0.33 | 0.1448 | 1 |
| P16615 | ATP2A2 | kEFtLEFsRDRkSMs | S488 | 0.055 | 80.16 | 1042 | 0.04372 | 0.17 | 0.1351 | 1 |
| P16615 | ATP2A2 | sRDRkSMsVyCtPNk | S495 | 0 | 96.56 | 1042 | 0 | 0 | 0 | 1 |
| P16615 | ATP2A2 | DRkSMsVyCtPNkPs | Y497 | 0.177 | 95.71 | 1042 | 0.077647 | 0.177 | 0.0776 | 1 |
| P16615 | ATP2A2 | GIRVIMItGDNkGTA | T624 | 0.006 | 85.32 | 1042 | 0.004098 | 0.006 | 0.0041 | 1 |
| P16615 | ATP2A2 | tAVAKTAsEMVLADD | S730 | 0.033 | 90.05 | 1042 | 0.026232 | 0.033 | 0.0262 | 1 |
| P16615 | ATP2A2 | sGWLFFRyLAIGCYV | Y836 | 0.173 | 84.48 | 1042 | 0.075893 | 0.173 | 0.0759 | 1 |
| P16615 | ATP2A2 | WLMVLKIsLPVILMD | S973 | 0 | 94.5 | 1042 | 0 | 0 | 0 | 1 |
| Q14032 | BAAT | PEVTDLEyFEEAANF | Y210 | 0.104 | 98.4 | 418 | 0.045623 | 0.104 | 0.0456 | 1 |
| P24534 | EEF1B2 | VKKALGkyGPADVED | Y79 | 0.031 | 73.75 | 225 | 0.013599 | 0.031 | 0.0136 | 0.9167 |
| Q9Y320 | TMX2 | YTDVSTRykVStsPL | Y206 | 0.291 | 94.06 | 296 | 0.127658 | 0.291 | 0.1277 | 1 |

| | | | | | | | | | | |
|---|---|---|---|---|---|---|---|---|---|---|
| Q6PGP7 | SKIC3 | LVEMDSksGPGLIGL | S271 | 0.038 | 86.98 | 1564 | 0.030207 | 0.075 | 0.0596 | 1 |
| Q96NG5 | ZNF558 | ECGKSFSsSFSLTVH | S331 | 0.144 | 92.31 | 402 | 0.114468 | 0.144 | 0.1145 | 0.8 |
| P98164 | LRP2 | LGWPAGVtLDMISKR | T563 | 0.079 | 96.22 | 4655 | 0.053958 | 0.079 | 0.054 | 1 |
| P98164 | LRP2 | DMISKRVyWVDSRFD | Y572 | 0.023 | 95.47 | 4655 | 0.01009 | 0.096 | 0.0421 | 1 |
| P98164 | LRP2 | WVDSRFDyIETVTyD | Y580 | 0.214 | 86.65 | 4655 | 0.093879 | 0.349 | 0.1338 | 1 |
| P98164 | LRP2 | DyIETVTyDGIQRKT | Y586 | 0.13 | 91.37 | 4655 | 0.057029 | 0.237 | 0.104 | 1 |
| Q16620 | NTRK2 | KILVAVKtLKDASDN | T573 | 0.149 | 86.55 | 822 | 0.101769 | 0.149 | 0.1018 | 1 |
| Q6F5E8 | CARMIL2 | QWDVDtIyHRQGCRH | Y185 | 0.008 | 86.99 | 1435 | 0.003509 | 0.121 | 0.0531 | 1 |
| Q6F5E8 | CARMIL2 | RRLsLAQtGLtPRGM | T314 | 0.012 | 88.57 | 1435 | 0.008196 | 0.012 | 0.0082 | 1 |
| Q6F5E8 | CARMIL2 | QRSRPELtARAVHQI | T688 | 0.031 | 91.43 | 1435 | 0.021173 | 0.031 | 0.0212 | 1 |
| Q16719 | KYNU | sLELPADtVQRIAAE | T12 | 0.147 | 95.88 | 465 | 0.100403 | 0.147 | 0.1004 | 1 |
| Q16719 | KYNU | HGLNIEEsMRMIKPR | S186 | 0.018 | 98.66 | 465 | 0.014308 | 0.018 | 0.0143 | 1 |
| P11308 | ERG | KDDFQRLtPSyNADI | T180 | 0.071 | 89.93 | 479 | 0.048494 | 0.071 | 0.0485 | 0.9 |
| P35557 | GCK | IVRRACEsVstRAAH | S373 | 0.168 | 98.67 | 465 | 0.133546 | 0.168 | 0.1335 | 1 |
| P35557 | GCK | tRAAHMCsAGLAGVI | S383 | 0 | 98.84 | 465 | 0 | 0 | 0 | 1 |
| Q9ULH0 | KIDINS220 | NTALMIAsKEGHTEI | S244 | 0 | 94.83 | 1771 | 0 | 0 | 0 | 1 |
| P27448 | MARK3 | SKFRQIVsAVQyCHQ | S164 | 0.14 | 98.64 | 753 | 0.111288 | 0.14 | 0.1113 | 1 |
| P27448 | MARK3 | DIMVGMGysQEEIQE | Y340 | 0.329 | 96.85 | 753 | 0.144328 | 0.329 | 0.1443 | 1 |
| P54132 | BLM | GGGKSLCyQLPACVS | Y699 | 0.004 | 97.63 | 1417 | 0.001755 | 0.004 | 0.0018 | 1 |
| P54132 | BLM | PGVTVVIsPLRSLIV | S714 | 0.012 | 92.91 | 1417 | 0.009539 | 0.012 | 0.0095 | 1 |
| P54132 | BLM | SVPVMALtAtANPRV | T830 | 0 | 91.28 | 1417 | 0 | 0 | 0 | 1 |
| P54132 | BLM | ECDtMADtLQRDGLA | T907 | 0.132 | 96.97 | 1417 | 0.090158 | 0.214 | 0.1462 | 1 |
| Q8N6T3 | ARFGAP1 | sLQEkyNsRAAALFR | S105 | 0.107 | 94.69 | 406 | 0.085056 | 0.139 | 0.1105 | 1 |
| Q17RS7 | GEN1 | EKLLVLLtHyDMIER | T380 | 0.001 | 95.02 | 908 | 0.000683 | 0.182 | 0.1243 | 1 |
| Q11128 | FUT5 | DSARVRyyQSLQAHL | Y223 | 0.21 | 98.21 | 374 | 0.092124 | 0.21 | 0.0921 | 1 |
| Q11128 | FUT5 | AHLKVDVyGRSHKPL | Y235 | 0.238 | 98.25 | 374 | 0.104407 | 0.238 | 0.1044 | 1 |
| O60658 | PDE8A | SIRSSKLsENTVIVG | S153 | 0.011 | 92.14 | 829 | 0.008744 | 0.011 | 0.0087 | 1 |
| O60658 | PDE8A | QGGKIRHyVSIIRVC | Y319 | 0.14 | 93.28 | 829 | 0.061416 | 0.14 | 0.0614 | 1 |
| Q8WVB6 | CHTF18 | SVRDVQAtRVGLKDQ | T575 | 0.073 | 72.08 | 975 | 0.04986 | 0.183 | 0.125 | 1 |
| P62195 | PSMC5 | EkVPDStyEMIGGLD | Y148 | 0.273 | 86.12 | 406 | 0.119761 | 0.273 | 0.1198 | 1 |
| P62195 | PSMC5 | MAREHAPsIIFMDEI | S244 | 0.008 | 87.13 | 406 | 0.006359 | 0.008 | 0.0064 | 1 |

| | | | | | | | | | | |
|---|---|---|---|---|---|---|---|---|---|---|
| Q8TDY2 | RB1CC1 | MEIKVEEsLMMPAVF | S105 | 0.06 | 92.28 | 1594 | 0.047695 | 0.123 | 0.0978 | 1 |
| Q8N9C0 | IGSF22 | DVTKEAVtItWNAPT | T718 | 0.11 | 94.53 | 903 | 0.075131 | 0.11 | 0.0751 | 1 |
| P51784 | USP11 | GQLIIMEtRKKDGTW | T276 | 0.187 | 93.69 | 963 | 0.127724 | 0.187 | 0.1277 | 1 |
| Q8N201 | INTS1 | LsDIDEMsRRRPEIL | S2074 | 0.018 | 86.46 | 2190 | 0.014308 | 0.018 | 0.0143 | 1 |
| Q9BQG1 | SYT3 | AMDLtGFsDPyVKAs | S467 | 0 | 92.67 | 590 | 0 | 0 | 0 | 1 |
| Q9BQG1 | SYT3 | LtGFsDPyVKAsLIs | Y470 | 0.101 | 98.4 | 590 | 0.044307 | 0.101 | 0.0443 | 1 |
| Q9BQG1 | SYT3 | sDPyVKAsLIsEGRR | S474 | 0.017 | 97.17 | 590 | 0.013514 | 0.017 | 0.0135 | 1 |
| Q7L576 | CYFIP1 | SWKLVHPtDkYSNKD | T420 | 0.123 | 94.14 | 1253 | 0.084011 | 0.123 | 0.084 | 1 |
| Q9Y217 | MTMR6 | QLSkQAkyEDLyAFS | Y104 | 0.281 | 96.2 | 621 | 0.123271 | 0.281 | 0.1233 | 1 |
| Q9Y217 | MTMR6 | LLEDQkkyLNPLYSS | Y468 | 0.081 | 95.3 | 621 | 0.035534 | 0.081 | 0.0355 | 1 |
| Q9UHL4 | DPP7 | NQFFRDVtADFEGQs | T206 | 0.101 | 98.92 | 492 | 0.068984 | 0.101 | 0.069 | 1 |
| O95154 | AKR7A3 | rRMDAPtsAAVtRAF | S28 | 0 | 98.52 | 331 | 0 | 0 | 0 | 1 |
| O94910 | ADGRL1 | RTRNIVKyDLRTRIK | Y236 | 0.144 | 95.74 | 1474 | 0.063171 | 0.144 | 0.0632 | 1 |
| Q9NZI7 | UBP1 | HCISTEFtPRKHGGE | T194 | 0.112 | 88.51 | 540 | 0.076498 | 0.173 | 0.1182 | 1 |
| O75911 | DHRS3 | HGkSLMDsDDDALLk | S135 | 0.163 | 93.85 | 302 | 0.129571 | 0.163 | 0.1296 | 1 |
| P41221 | WNT5A | LyQDHMQyIGEGAKt | Y92 | 0.197 | 96.53 | 380 | 0.086421 | 0.31 | 0.1061 | 1 |
| Q14739 | LBR | KAPRNDLsPASSGNA | S357 | 0.06 | 80.22 | 615 | 0.047695 | 0.086 | 0.0684 | 1 |
| Q14739 | LBR | IGtFDLKyFCELRPG | Y385 | 0.001 | 92.76 | 615 | 0.000439 | 0.001 | 0.0004 | 1 |
| Q5VZL5 | ZMYM4 | sPIVLLNtLLFFNTK | T1382 | 0.005 | 97.85 | 1548 | 0.003415 | 0.093 | 0.0635 | 1 |
| Q9HB03 | ELOVL3 | KMLPMLItsLQILQM | T203 | 0.074 | 95.73 | 270 | 0.050543 | 0.074 | 0.0505 | 0.875 |
| P41252 | IARS1 | LIYkAVPsWFVRVEN | S414 | 0.025 | 88.97 | 1262 | 0.019873 | 0.025 | 0.0199 | 1 |
| P41252 | IARS1 | LRLYLINsPVVRAEN | S630 | 0.134 | 93.34 | 1262 | 0.106519 | 0.051 | 0.1178 | 1 |
| P41252 | IARS1 | EEEIEFLyNENTVRE | Y680 | 0.122 | 93.57 | 1262 | 0.05352 | 0.122 | 0.0535 | 1 |
| P41252 | IARS1 | NENTVREsPNItDRW | S688 | 0.106 | 87.05 | 1262 | 0.084261 | 0.187 | 0.1486 | 1 |
| P41252 | IARS1 | VREsPNItDRWILSF | T692 | 0.036 | 93.2 | 1262 | 0.024588 | 0.036 | 0.0246 | 1 |
| P41252 | IARS1 | CRLMAPYtPFLtELM | T773 | 0 | 98.02 | 1262 | 0 | 0 | 0 | 1 |
| P41252 | IARS1 | APYtPFLtELMYQNL | T777 | 0 | 98.36 | 1262 | 0 | 0 | 0 | 1 |
| P41252 | IARS1 | VQDKDtLsIHyLMLP | S800 | 0 | 96.73 | 1262 | 0 | 0 | 0 | 1 |
| P41252 | IARS1 | KDtLsIHyLMLPRVR | Y803 | 0.128 | 95.13 | 1262 | 0.056152 | 0.256 | 0.1123 | 1 |
| A1L0T0 | ILVBL | HGVRFIFtLVGGHIS | T72 | 0 | 98.84 | 632 | 0 | 0 | 0 | 0.9167 |
| A1L0T0 | ILVBL | KNAQMAQsPILLLGG | S141 | 0.014 | 97.83 | 632 | 0.011129 | 0.028 | 0.0223 | 0.8889 |

| | | | | | | | | | | |
|---|---|---|---|---|---|---|---|---|---|---|
| Q9UKY4 | POMT2 | THFGKMGsyyINRTF | S94 | 0.063 | 95.71 | 750 | 0.05008 | 0.126 | 0.1002 | 1 |
| Q9UKY4 | POMT2 | FGKMGsyyINRTFFF | Y96 | 0.187 | 96.67 | 750 | 0.082034 | 0.311 | 0.114 | 0.9286 |
| Q9UKY4 | POMT2 | EtsRNLHsHYHEAPM | S425 | 0 | 96.77 | 750 | 0 | 0 | 0 | 1 |
| Q9H553 | ALG2 | VLENVEHyQELkKMV | Y281 | 0.199 | 97.88 | 416 | 0.087298 | 0.199 | 0.0873 | 1 |
| Q9H553 | ALG2 | ARVKEkFsPEAFTEQ | S399 | 0.092 | 91.99 | 416 | 0.073132 | 0.092 | 0.0731 | 0.8333 |
| Q9UPM8 | AP4E1 | DVGVMAAsLHIyLRM | S225 | 0.001 | 94.03 | 1137 | 0.000795 | 0.001 | 0.0008 | 1 |
| Q9UPM8 | AP4E1 | SSGYKDLtGSFVTIL | T244 | 0.004 | 93.64 | 1137 | 0.002732 | 0.004 | 0.0027 | 1 |
| Q6GYQ0 | RALGAPA1 | ALNVICVsLKFTNKT | S1324 | 0.009 | 96.26 | 2036 | 0.007154 | 0.069 | 0.059 | 1 |
| Q6GYQ0 | RALGAPA1 | IIQILIAtItHLLPS | T1368 | 0.037 | 93.92 | 2036 | 0.025272 | 0.037 | 0.0253 | 1 |
| P68371 | TUBB4B | LERINVyyNEAtGGk | Y51 | 0.021 | 96.91 | 445 | 0.009212 | 0.021 | 0.0092 | 1 |
| P68371 | TUBB4B | NEAtGGkyVPRAVLV | Y59 | 0.268 | 96.69 | 445 | 0.117568 | 0.386 | 0.1327 | 1 |
| P68371 | TUBB4B | sLGGGtGsGMGtLLI | S145 | 0 | 96.22 | 445 | 0 | 0 | 0 | 1 |
| P68371 | TUBB4B | NtFsVVPsPkVSDTV | S172 | 0.02 | 78.3 | 445 | 0.015898 | 0.02 | 0.0159 | 0.8333 |
| P68371 | TUBB4B | PRHGRyLtVAAVFrG | T312 | 0.047 | 96.1 | 445 | 0.032102 | 0.073 | 0.055 | 1 |
| P68371 | TUBB4B | QELFkRIsEQFtAMF | S382 | 0.138 | 94.43 | 445 | 0.109698 | 0.138 | 0.1097 | 1 |
| Q9Y4E6 | WDR7 | RHGsVALyDIRTGKC | Y1380 | 0.167 | 89.69 | 1490 | 0.07326 | 0.167 | 0.0733 | 0.9167 |
| A6NL08 | OR6C75 | YFFLRNFsFLEISFT | S65 | 0.007 | 97.29 | 312 | 0.005564 | 0.007 | 0.0056 | 1 |
| O75694 | NUP155 | LKVDkIItPLNkDHI | T474 | 0.04 | 90.94 | 1391 | 0.027321 | 0.04 | 0.0273 | 1 |
| O75694 | NUP155 | PkKFVLLsAQGsLMF | S504 | 0.007 | 87.59 | 1391 | 0.005564 | 0.007 | 0.0056 | 1 |
| Q6BAA4 | FCRLB | RASDsGRyQCSGtMR | Y166 | 0.049 | 98.28 | 426 | 0.021496 | 0.049 | 0.0215 | 1 |
| Q6BAA4 | FCRLB | VEsAPMFsAKVAVTV | S183 | 0.061 | 97.29 | 426 | 0.04849 | 0.061 | 0.0485 | 1 |
| P55011 | SLC12A2 | MsIDQAkyQRWLIKN | Y842 | 0.196 | 94.32 | 1212 | 0.085982 | 0.299 | 0.1312 | 1 |
| P55011 | SLC12A2 | KNKMKAFyAPVHADD | Y855 | 0.156 | 95.62 | 1212 | 0.068435 | 0.156 | 0.0684 | 1 |
| P55011 | SLC12A2 | DQkLLEAsTQFQkKQ | S1012 | 0.048 | 86.66 | 1212 | 0.038156 | 0.048 | 0.0382 | 1 |
| Q9P2D7 | DNAH1 | IVKVMDDyQVMDEFL | Y892 | 0.033 | 87.61 | 4265 | 0.014477 | 0.033 | 0.0145 | 1 |
| P20591 | MX1 | kWRGKVSyQDyEIEI | Y126 | 0.154 | 91.34 | 662 | 0.067558 | 0.154 | 0.0676 | 1 |
| Q96C10 | DHX58 | LAALQDFyHREHVTK | Y310 | 0.046 | 95.42 | 678 | 0.02018 | 0.046 | 0.0202 | 1 |
| Q96C10 | DHX58 | QKMDQAEyQAKIRDL | Y518 | 0.078 | 98.53 | 678 | 0.034217 | 0.078 | 0.0342 | 1 |
| Q9BSC4 | NOL10 | TPKMGIyyIPVLGPA | Y340 | 0.025 | 92.8 | 688 | 0.010967 | 0.025 | 0.011 | 0.9286 |
| O15239 | NDUFA1 | MERDRRIsGVDryyV | S55 | 0.048 | 97.12 | 70 | 0.038156 | 0.048 | 0.0382 | 1 |
| P55273 | CDKN2D | ARtGFLDtLKVLVEH | T89 | 0.012 | 98.77 | 166 | 0.008196 | 0.012 | 0.0082 | 0.8333 |

| | | | | | | | | | | |
|---|---|---|---|---|---|---|---|---|---|---|
| P45954 | ACADSB | GYkYAIGsLNEGRIG | S289 | 0.025 | 98.03 | 432 | 0.019873 | 0.025 | 0.0199 | 0.8889 |
| O60706 | ABCC9 | GICGRTGsGKSSLSL | S1350 | 0.047 | 78.6 | 1549 | 0.037361 | 0.047 | 0.0374 | 1 |
| Q16543 | CDC37 | HMQRCIDsGLWVPNs | S339 | 0.032 | 82.03 | 378 | 0.025437 | 0.032 | 0.0254 | 1 |
| Q53FZ2 | ACSM3 | DSHTCVKtKHNEIMA | T224 | 0.026 | 98.07 | 586 | 0.017758 | 0.026 | 0.0178 | 0.9 |
| Q53FZ2 | ACSM3 | EIMAIFFtSGTSGYP | T235 | 0.186 | 81.01 | 586 | 0.127041 | 0.186 | 0.127 | 1 |
| Q6P2P2 | PRMT9 | GAGTGILsMFAKkAG | S192 | 0.006 | 98.85 | 845 | 0.004769 | 0.006 | 0.0048 | 1 |
| Q6P2P2 | PRMT9 | PTNVkFQsPAYSSVD | S337 | 0.007 | 91.53 | 845 | 0.005564 | 0.007 | 0.0056 | 0.8 |
| Q6ICB0 | DESI1 | kGLARRLsPIMLGKQ | S25 | 0.058 | 95.47 | 168 | 0.046105 | 0.058 | 0.0461 | 0.8889 |
| Q3LXA3 | TKFC | TAAGGsAskRMALVL | S371 | 0.017 | 86.17 | 575 | 0.013514 | 0.087 | 0.0692 | 1 |
| Q8NI08 | NCOA7 | GRSNSCstFNNDILs | T919 | 0 | 94.13 | 942 | 0 | 0 | 0 | 1 |
| Q8NI08 | NCOA7 | tFNNDILskKEDFIV | S926 | 0.131 | 92.12 | 942 | 0.104134 | 0.131 | 0.1041 | 0.875 |
| Q8NBH2 | KY | WDRSSLKsMsLDLQQ | S144 | 0.106 | 94.13 | 661 | 0.084261 | 0.106 | 0.0843 | 1 |
| Q8NBH2 | KY | QCMtVPGysKGFGYQ | Y249 | 0.232 | 95.38 | 661 | 0.101775 | 0.232 | 0.1018 | 1 |
| P10768 | ESD | PVDPQRMsIFGHsMG | S144 | 0 | 98.89 | 282 | 0 | 0 | 0 | 0.8 |
| Q9BVC4 | MLST8 | RTVQHQDsQVNALEV | S43 | 0.083 | 88.73 | 326 | 0.065978 | 0.083 | 0.066 | 1 |
| A8MV23 | SERPINE3 | stMSFQGtWRKRFSS | T197 | 0.179 | 88.87 | 424 | 0.122259 | 0.179 | 0.1223 | 1 |
| A8MV23 | SERPINE3 | KADRPFIyFLREPNT | Y383 | 0.001 | 95.23 | 424 | 0.000439 | 0.054 | 0.0237 | 1 |
| P14866 | HNRNPL | EyGPHADsPVLMVYG | S381 | 0.038 | 88.23 | 589 | 0.030207 | 0.116 | 0.0922 | 1 |
| P14866 | HNRNPL | sGkSERSssGLLEWE | S543 | 0.116 | 91.11 | 589 | 0.09221 | 0.116 | 0.0922 | 1 |
| P19474 | TRIM21 | GPLRPFFsPGFNDGG | S447 | 0.027 | 98.17 | 475 | 0.021463 | 0.027 | 0.0215 | 1 |
| Q96GQ7 | DDX27 | TDHVMLFtQtKKQAH | T470 | 0.001 | 93.8 | 796 | 0.000683 | 0.001 | 0.0007 | 1 |
| P53004 | BLVRA | DLRNPHPsSAFLNLI | S33 | 0.005 | 94.6 | 296 | 0.003975 | 0.005 | 0.004 | 1 |
| P53004 | BLVRA | SQEVEVAyICSESSS | Y72 | 0 | 98.89 | 296 | 0 | 0 | 0 | 1 |
| P53004 | BLVRA | FSGISRLtWLVsLFG | T174 | 0.008 | 98.63 | 296 | 0.005464 | 0.008 | 0.0055 | 0.8889 |
| P53004 | BLVRA | CLEtEKksPLSWIEE | S211 | 0.093 | 97.93 | 296 | 0.073927 | 0.093 | 0.0739 | 1 |
| Q5D0E6 | DALRD3 | LATLFESyKCSMEQG | Y417 | 0.174 | 97.74 | 543 | 0.076331 | 0.174 | 0.0763 | 0.9286 |
| O60469 | DSCAM | VRVEDQKtMRGNVAV | T135 | 0.046 | 79.43 | 2012 | 0.031419 | 0.158 | 0.1079 | 1 |
| Q9ULC5 | ACSL5 | KGIIRHDsFWDKLIF | S395 | 0.045 | 90.29 | 683 | 0.035771 | 0.045 | 0.0358 | 1 |
| P00450 | CP | SKYKKVVyRQyTDST | Y784 | 0 | 98.74 | 1065 | 0 | 0 | 0 | 1 |
| P00450 | CP | KKVVyRQyTDSTFRV | Y787 | 0.245 | 98.21 | 1065 | 0.107478 | 0.245 | 0.1075 | 1 |
| Q9HCM2 | PLXNA4 | PYHKKEHyLSGVNEs | Y159 | 0.055 | 93.56 | 1894 | 0.024128 | 0.098 | 0.043 | 1 |

| | | | | | | | | | | |
|---|---|---|---|---|---|---|---|---|---|---|
| Q9HCM2 | PLXNA4 | VLRDMAFsKDHEQLY | S488 | 0.116 | 94.21 | 1894 | 0.09221 | 0.116 | 0.0922 | 1 |
| P19525 | EIF2AK2 | DMkEtKytVDKRFGM | T258 | 0.052 | 87.1 | 551 | 0.035517 | 0.073 | 0.0499 | 1 |
| P19525 | EIF2AK2 | HRIDGKTyVIKRVkY | Y293 | 0.166 | 97.75 | 551 | 0.072822 | 0.166 | 0.0728 | 0.8 |
| Q9BVC3 | DSCC1 | NPYEGPDsQkEkDSN | S152 | 0.161 | 87.66 | 393 | 0.127981 | 0.161 | 0.128 | 1 |
| Q969V3 | NCLN | RIIAEALtRVIYNLT | T423 | 0.02 | 93.36 | 563 | 0.01366 | 0.16 | 0.1093 | 1 |
| Q9UIJ7 | AK3 | HPASGRVyNIEFNPP | Y138 | 0.166 | 93.33 | 227 | 0.072822 | 0.166 | 0.0728 | 1 |
| Q9Y6Y9 | LY96 | IKFSKGKyKCVVEAI | Y131 | 0.121 | 93.54 | 160 | 0.053081 | 0.121 | 0.0531 | 1 |
| Q96GQ5 | RUSF1 | GATRAALtVHQARRN | T215 | 0.072 | 96.16 | 468 | 0.049177 | 0.072 | 0.0492 | 1 |
| Q9NS15 | LTBP3 | CMNtGGsyNCHCNRG | Y639 | 0.242 | 87.9 | 1303 | 0.106162 | 0.242 | 0.1062 | 1 |
| Q2PZI1 | DPY19L1 | RtEMGLYySYFKTIV | Y68 | 0.094 | 96.4 | 675 | 0.041236 | 0.094 | 0.0412 | 1 |
| Q2PZI1 | DPY19L1 | VFAGAMPtMAsVkLS | T558 | 0.055 | 94.47 | 675 | 0.037566 | 0.082 | 0.0529 | 0.8889 |
| Q2PZI1 | DPY19L1 | RARTKIVysMysRKA | Y589 | 0.037 | 95.55 | 675 | 0.016231 | 0.23 | 0.0685 | 1 |
| Q2PZI1 | DPY19L1 | ARTKIVysMysRKAA | S590 | 0.091 | 94.68 | 675 | 0.072337 | 0.173 | 0.1407 | 1 |
| Q2PZI1 | DPY19L1 | TKIVysMysRKAAEE | Y592 | 0.009 | 94.65 | 675 | 0.003948 | 0.009 | 0.0039 | 1 |
| Q2PZI1 | DPY19L1 | KIVysMysRKAAEEV | S593 | 0.016 | 94.4 | 675 | 0.012719 | 0.016 | 0.0127 | 1 |
| O75762 | TRPA1 | ETMLHRAsLFDHHEL | S317 | 0 | 91.01 | 1119 | 0 | 0 | 0 | 1 |
| O75762 | TRPA1 | ILATASAsWNIVNLL | S354 | 0.045 | 88.39 | 1119 | 0.035771 | 0.045 | 0.0358 | 1 |
| O75762 | TRPA1 | SKSKDKKsPLHFAAS | S448 | 0.001 | 88.3 | 1119 | 0.000795 | 0.001 | 0.0008 | 1 |
| A4D161 | FAM221A | TGCQCRAyLyVPLNG | Y111 | 0.074 | 96.06 | 298 | 0.032463 | 0.074 | 0.0325 | 1 |
| A4D161 | FAM221A | GYMRLDDsGIGVPSV | S210 | 0.003 | 93.62 | 298 | 0.002385 | 0.003 | 0.0024 | 1 |
| P20930 | FLG | IINLFKQySKKDKNT | Y18 | 0.073 | 90.64 | 4061 | 0.032024 | 0.073 | 0.032 | 0.9 |
| Q8TDR2 | STK35 | kVADFGLskVCAGLA | S385 | 0.038 | 74.7 | 534 | 0.030207 | 0.038 | 0.0302 | 1 |
| O00629 | KPNA4 | ITFLRNVtWVMVNLC | T221 | 0.018 | 96.19 | 521 | 0.012294 | 0.018 | 0.0123 | 1 |
| Q92900 | UPF1 | VTSATIVyHLARQGN | Y518 | 0.137 | 97.37 | 1129 | 0.0601 | 0.137 | 0.0601 | 1 |
| Q9Y4L1 | HYOU1 | DNPHVALyQARFPEH | Y116 | 0.053 | 98.32 | 999 | 0.02325 | 0.053 | 0.0232 | 1 |
| Q9Y4L1 | HYOU1 | NTATALSyGVFRRKD | Y213 | 0.236 | 97.34 | 999 | 0.10353 | 0.236 | 0.1035 | 1 |
| Q9Y4L1 | HYOU1 | AAAMGAVyQAAALSK | Y421 | 0.184 | 97.4 | 999 | 0.080718 | 0.184 | 0.0807 | 1 |
| Q9Y4L1 | HYOU1 | VVRDAVVyPILVEFT | Y443 | 0.181 | 97.42 | 999 | 0.079402 | 0.181 | 0.0794 | 1 |
| Q9Y4L1 | HYOU1 | KLkGVGDsFkkYPDY | S527 | 0.05 | 93.8 | 999 | 0.039746 | 0.05 | 0.0397 | 1 |
| Q9Y4L1 | HYOU1 | DkLYQPEyQEVsTEE | Y759 | 0.276 | 93.76 | 999 | 0.121077 | 0.276 | 0.1211 | 1 |
| Q9Y4L1 | HYOU1 | QPEyQEVsTEEQREE | S763 | 0.118 | 92.7 | 999 | 0.0938 | 0.118 | 0.0938 | 1 |

| | | | | | | | | | | |
|---|---|---|---|---|---|---|---|---|---|---|
| Q8IVV2 | LOXHD1 | TYIVtVFtGDVRGAG | T304 | 0.03 | 89.14 | 2067 | 0.02049 | 0.03 | 0.0205 | 1 |
| P49915 | GMPS | GVDSTVCtALLNRAL | T253 | 0 | 95.08 | 693 | 0 | 0 | 0 | 1 |
| P49915 | GMPS | SQMPVILtPLHFDrD | T616 | 0 | 98.57 | 693 | 0 | 0 | 0 | 1 |
| P49915 | GMPS | FMTGIPAtPGNEIPV | T652 | 0.142 | 96.49 | 693 | 0.096988 | 0.142 | 0.097 | 1 |
| P49915 | GMPS | IKKIPGIsRIMYDLT | S676 | 0.069 | 97.67 | 693 | 0.054849 | 0.069 | 0.0548 | 1 |
| Q92608 | DOCK2 | ISSsPTHsLyVFVRN | S222 | 0.001 | 91.8 | 1830 | 0.000795 | 0.001 | 0.0008 | 1 |
| Q92608 | DOCK2 | SsPTHsLyVFVRNFV | Y224 | 0.071 | 94.03 | 1830 | 0.031147 | 0.071 | 0.0311 | 1 |
| Q92608 | DOCK2 | NNLkVVFtDLGNkDL | T281 | 0.057 | 93.64 | 1830 | 0.038932 | 0.113 | 0.0771 | 1 |
| Q92608 | DOCK2 | GVAVMDItDIIKGkA | T331 | 0.108 | 90.93 | 1830 | 0.073765 | 0.145 | 0.1164 | 1 |
| Q92608 | DOCK2 | LVCSTkLtQNVGLLG | T612 | 0.044 | 88.66 | 1830 | 0.030053 | 0.044 | 0.0301 | 0.8571 |
| Q92608 | DOCK2 | FNTVLEAyIQQHFSA | Y699 | 0.043 | 93.25 | 1830 | 0.018863 | 0.043 | 0.0189 | 1 |
| Q92608 | DOCK2 | LQKQkVQsMNEIVQS | S838 | 0.019 | 94.24 | 1830 | 0.015103 | 0.019 | 0.0151 | 1 |
| Q8WUJ3 | CEMIP | SWTFLNKtLHPGGMA | T167 | 0.121 | 93 | 1361 | 0.082645 | 0.17 | 0.1161 | 1 |
| Q8WUJ3 | CEMIP | KDKRPFLsIIsARYS | S758 | 0.004 | 96.77 | 1361 | 0.00318 | 0.004 | 0.0032 | 1 |
| Q99986 | VRK1 | AkRFSRktVLQLsLR | T153 | 0.025 | 98.34 | 396 | 0.017075 | 0.025 | 0.0171 | 1 |
| Q99986 | VRK1 | RktVLQLsLRILDIL | S158 | 0 | 98.86 | 396 | 0 | 0 | 0 | 1 |
| O60603 | TLR2 | LQSYEPKsLKsIQNV | S193 | 0 | 93.5 | 784 | 0 | 0 | 0 | 1 |
| Q10567 | AP1B1 | AVkKVIAsMtVGKDV | S40 | 0.122 | 83.9 | 949 | 0.09698 | 0.122 | 0.097 | 1 |
| Q92889 | ERCC4 | NLFVRKLyLWPRFHV | Y191 | 0.235 | 94.77 | 916 | 0.103091 | 0.235 | 0.1031 | 1 |
| Q15303 | ERBB4 | DQNKFLCyADTIHWQ | Y157 | 0.254 | 94.8 | 1308 | 0.111426 | 0.254 | 0.1114 | 1 |
| Q8N0X7 | SPART | kEEAKNyykQGIGHL | Y46 | 0.032 | 97.22 | 666 | 0.014038 | 0.032 | 0.014 | 1 |
| Q99728 | BARD1 | PKPDSDVtQTINtVA | T714 | 0.118 | 72.93 | 777 | 0.080596 | 0.118 | 0.0806 | 1 |
| Q99728 | BARD1 | DSDQRFCtQYIIYED | T734 | 0.004 | 92.21 | 777 | 0.002732 | 0.004 | 0.0027 | 1 |
| Q02548 | PAX5 | VVEKIAEyKRQNPTM | Y102 | 0.042 | 95.18 | 391 | 0.018425 | 0.042 | 0.0184 | 1 |
| P09104 | ENO2 | TVEVDLytAKGLFRA | T26 | 0.096 | 98.12 | 434 | 0.065569 | 0.12 | 0.0996 | 1 |
| P09104 | ENO2 | LFRAAVPsGAstGly | S37 | 0.004 | 95.02 | 434 | 0.00318 | 0.004 | 0.0032 | 1 |
| P09104 | ENO2 | AERELPLyRHIAQLA | Y131 | 0.072 | 98.61 | 434 | 0.031585 | 0.072 | 0.0316 | 1 |
| P09104 | ENO2 | LkVNQIGsVTEAIQA | S349 | 0.001 | 98.52 | 434 | 0.000795 | 0.001 | 0.0008 | 1 |
| P35241 | RDX | TAVLLASyAVQAKyG | Y128 | 0.18 | 98.65 | 583 | 0.078963 | 0.278 | 0.122 | 1 |
| P35241 | RDX | VQAKyGDyNkEIHkP | Y137 | 0.324 | 96.42 | 583 | 0.142134 | 0.324 | 0.1421 | 1 |
| P35241 | RDX | CMGNHELyMRRRkPD | Y291 | 0.052 | 93.65 | 583 | 0.022812 | 0.267 | 0.1171 | 1 |

| | | | | | | | | | | |
|---|---|---|---|---|---|---|---|---|---|---|
| Q7Z6L1 | TECPR1 | TEKGGWTyAIDFPAT | Y140 | 0.136 | 93.91 | 1165 | 0.059661 | 0.25 | 0.1018 | 1 |
| P62879 | GNB2 | TDsRLLVsAsQDGkL | S72 | 0 | 98.9 | 340 | 0 | 0 | 0 | 1 |
| P62879 | GNB2 | sRLLVsAsQDGkLII | S74 | 0 | 98.73 | 340 | 0 | 0 | 0 | 1 |
| O43432 | EIF4G3 | RELTIEFsKPLLPVG | S1343 | 0.084 | 86.96 | 1585 | 0.066773 | 0.084 | 0.0668 | 0.875 |
| Q96GZ6 | SLC41A3 | WVLIAKQsPPIVKIL | S275 | 0.081 | 90.12 | 507 | 0.064388 | 0.081 | 0.0644 | 1 |
| Q86TB3 | ALPK2 | LtDVGIAtLAkGYKG | T2095 | 0.092 | 92.3 | 2170 | 0.062837 | 0.092 | 0.0628 | 1 |
| Q96LA6 | FCRL1 | MWKEDTGsyWCEAQT | S83 | 0.149 | 97.37 | 429 | 0.118442 | 0.149 | 0.1184 | 1 |
| Q96LA6 | FCRL1 | WKEDTGsyWCEAQTM | Y84 | 0.021 | 97.46 | 429 | 0.009212 | 0.021 | 0.0092 | 1 |
| P57740 | NUP107 | LQTKEEVsIEVLKTY | S569 | 0 | 96.16 | 925 | 0 | 0 | 0 | 1 |
| O43193 | MLNR | YLGSMAVsDLLILLG | S83 | 0 | 97.48 | 412 | 0 | 0 | 0 | 0.8 |
| P27987 | ITPKB | PSLRKDMyQKMIEVD | Y770 | 0.253 | 95.49 | 946 | 0.110987 | 0.253 | 0.111 | 1 |
| P27987 | ITPKB | NHNILIAyRDRLKAI | Y854 | 0 | 98.58 | 946 | 0 | 0 | 0 | 1 |
| Q96PZ0 | PUS7 | RLMyIHSyQSyVWNN | Y479 | 0.006 | 97.42 | 661 | 0.002632 | 0.006 | 0.0026 | 0.9412 |
| C9JRZ8 | AKR1B15 | TAYSPLGsPDRPWAK | S215 | 0.096 | 97.32 | 316 | 0.076312 | 0.096 | 0.0763 | 0.8 |
| Q9BT88 | SYT11 | QTQGSDPyIKMTILP | Y197 | 0.116 | 96.74 | 431 | 0.050888 | 0.116 | 0.0509 | 0.8333 |
| Q14644 | RASA3 | EGLRVFQsVkIKIGE | S13 | 0.098 | 90.69 | 834 | 0.077902 | 0.167 | 0.1328 | 1 |
| Q14644 | RASA3 | EkSLCPFyGEDFyCE | Y61 | 0.206 | 92.68 | 834 | 0.090369 | 0.206 | 0.0904 | 1 |
| Q14644 | RASA3 | PRSFRHLsFyIFDRD | S77 | 0.031 | 97.03 | 834 | 0.024642 | 0.061 | 0.0486 | 1 |
| Q14644 | RASA3 | SFRHLsFyIFDRDVF | Y79 | 0.15 | 97.17 | 834 | 0.065803 | 0.15 | 0.0658 | 1 |
| P42345 | MTOR | KTEQNQGtRREAIRV | T880 | 0.009 | 84.08 | 2549 | 0.006147 | 0.009 | 0.0061 | 1 |
| O60508 | CDC40 | RTFATyGyALDPSLD | Y137 | 0.013 | 94.9 | 579 | 0.005703 | 0.013 | 0.0057 | 1 |
| P26927 | MST1 | VWCNGEEyRGAVDRT | Y199 | 0.068 | 91.16 | 711 | 0.029831 | 0.068 | 0.0298 | 1 |
| P23786 | CPT2 | RATNMTVsAIRFLKT | S158 | 0.018 | 98.7 | 658 | 0.014308 | 0.018 | 0.0143 | 1 |
| P23786 | CPT2 | QyFRLFNstRLPKPS | S223 | 0.045 | 98.75 | 658 | 0.035771 | 0.045 | 0.0358 | 0.8889 |
| P23786 | CPT2 | yFRLFNstRLPKPSR | T224 | 0.032 | 98.81 | 658 | 0.021856 | 0.032 | 0.0219 | 0.8889 |
| P23786 | CPT2 | APEFPLAyLTSENRD | Y290 | 0.095 | 98.7 | 658 | 0.041675 | 0.095 | 0.0417 | 1 |
| P23786 | CPT2 | SVyTkRCsEAFVREP | S513 | 0.035 | 98.79 | 658 | 0.027822 | 0.035 | 0.0278 | 1 |
| Q9UHP7 | CLEC2D | DTKNWtssQRFCDSQ | S99 | 0 | 98.92 | 191 | 0 | 0 | 0 | 1 |
| Q8IWT1 | SCN4B | EDLHFRWtyNssDAF | T69 | 0.065 | 97.97 | 228 | 0.044396 | 0.065 | 0.0444 | 1 |
| Q8IWT1 | SCN4B | DLHFRWtyNssDAFK | Y70 | 0.205 | 97.52 | 228 | 0.089931 | 0.205 | 0.0899 | 1 |
| P10321 | HLA-C | VEQEGPEyWDRETQk | Y83 | 0.107 | 97.6 | 366 | 0.046939 | 0.107 | 0.0469 | 1 |

| | | | | | | | | | | |
|---|---|---|---|---|---|---|---|---|---|---|
| P10321 | HLA-C | SAYDGkDyIALNEDL | Y147 | 0.116 | 98.75 | 366 | 0.050888 | 0.116 | 0.0509 | 1 |
| O95460 | MATN4 | RVGLVQFsSRVRTEF | S431 | 0.002 | 93.38 | 622 | 0.00159 | 0.002 | 0.0016 | 1 |
| P15880 | RPS2 | TFDAIsktysyLtPD | T247 | 0.008 | 92.59 | 293 | 0.005464 | 0.008 | 0.0055 | 1 |
| P15880 | RPS2 | sktysyLtPDLWkET | T252 | 0.117 | 88.68 | 293 | 0.079913 | 0.117 | 0.0799 | 1 |
| Q8TC12 | RDH11 | AGLAYCHskLANILF | S205 | 0.003 | 97.77 | 318 | 0.002385 | 0.003 | 0.0024 | 0.9 |
| Q9GZV9 | FGF23 | APHQTIysALMIRSE | S71 | 0 | 97 | 251 | 0 | 0 | 0 | 0.8 |
| Q9GZV9 | FGF23 | GMNPPPysQFLsRRN | S155 | 0 | 94.44 | 251 | 0 | 0 | 0 | 0.8333 |
| Q08188 | TGM3 | ASAPIGRytMALQIF | Y111 | 0.042 | 98.88 | 693 | 0.018425 | 0.042 | 0.0184 | 1 |
| Q08188 | TGM3 | tMALQIFsQGGISsV | S119 | 0.109 | 96.95 | 693 | 0.086646 | 0.109 | 0.0866 | 1 |
| Q08188 | TGM3 | KYVGRVLsAMINSND | S221 | 0 | 98.76 | 693 | 0 | 0 | 0 | 0.9 |
| Q9Y2L1 | DIS3 | SGTYLQGtFRASREN | T250 | 0.211 | 95.26 | 958 | 0.144116 | 0.211 | 0.1441 | 1 |
| Q9Y2L1 | DIS3 | QTkELREtNSMVEEF | T659 | 0.001 | 93.18 | 958 | 0.000683 | 0.001 | 0.0007 | 1 |
| Q9Y2L1 | DIS3 | FMLLANIsVAKkIHE | S673 | 0.168 | 98.74 | 958 | 0.133546 | 0.168 | 0.1335 | 1 |
| Q96KN2 | CNDP1 | ANIDDtQyLAAKRAI | Y427 | 0 | 97.16 | 507 | 0 | 0 | 0 | 0.9412 |
| Q9UPZ3 | HPS5 | MASSNPVyMEMEKGD | Y822 | 0.174 | 76.47 | 1129 | 0.076331 | 0.174 | 0.0763 | 1 |
| P08237 | PFKM | VSKsGsHtVAVMNVG | T402 | 0.113 | 95.09 | 780 | 0.077181 | 0.113 | 0.0772 | 1 |
| P52788 | SMS | TYTNKNGsFANLRIY | S57 | 0.103 | 94.93 | 366 | 0.081876 | 0.127 | 0.1291 | 1 |
| P52788 | SMS | ADGRLVEyDIDEVVy | Y134 | 0.079 | 98.78 | 366 | 0.034656 | 0.079 | 0.0347 | 1 |
| P52788 | SMS | MGsGkEDytGkDVLI | Y189 | 0.058 | 98.68 | 366 | 0.025444 | 0.058 | 0.0254 | 1 |
| P52788 | SMS | CkKYMRktCGDVLDN | T236 | 0.168 | 97.27 | 366 | 0.114746 | 0.233 | 0.1476 | 1 |
| Q96P48 | ARAP1 | RRsFDLTtPyRIFSF | T502 | 0.122 | 87.92 | 1450 | 0.083328 | 0.207 | 0.1303 | 1 |
| Q8TEU7 | RAPGEF6 | SVARLRGtWEKLPSK | T968 | 0.008 | 95.1 | 1601 | 0.005464 | 0.008 | 0.0055 | 1 |
| Q15349 | RPS6KA2 | EIEILLRyGQHPNII | Y463 | 0.202 | 87.47 | 733 | 0.088614 | 0.202 | 0.0886 | 1 |
| Q96M32 | AK7 | RKLFILLsTVMtWAR | S153 | 0 | 95.78 | 723 | 0 | 0 | 0 | 1 |
| Q96M32 | AK7 | ILLsTVMtWARsKAL | T157 | 0 | 94.12 | 723 | 0 | 0 | 0 | 1 |
| Q9NYU1 | UGGT2 | SAFHKVLsEKAQNEE | S203 | 0.133 | 93.29 | 1516 | 0.105724 | 0.16 | 0.139 | 1 |
| Q9NYU1 | UGGT2 | AQNEEILyVLRHYIQ | Y213 | 0.009 | 94.96 | 1516 | 0.003948 | 0.009 | 0.0039 | 1 |
| Q9NYU1 | UGGT2 | RKMyLSGyGVELAIK | Y232 | 0.001 | 94.81 | 1516 | 0.000439 | 0.001 | 0.0004 | 0.9333 |
| C9JPN9 | USP17L12 | APASKMLtLLTSAKV | T268 | 0.148 | 97.19 | 530 | 0.101086 | 0.148 | 0.1011 | 1 |
| Q8TAP6 | CEP76 | GINRPVCsyVkPLRA | S322 | 0.09 | 94.61 | 659 | 0.071542 | 0.09 | 0.0715 | 1 |
| Q8TAP6 | CEP76 | INRPVCsyVkPLRAG | Y323 | 0.188 | 95.46 | 659 | 0.082473 | 0.188 | 0.0825 | 0.9412 |

| | | | | | | | | | | |
|---|---|---|---|---|---|---|---|---|---|---|
| P56373 | P2RX3 | FTRLDSVsEKSSVSP | S269 | 0.144 | 86.52 | 397 | 0.114468 | 0.144 | 0.1145 | 1 |
| Q9BRP0 | OVOL2 | GVQQQyAyKQRRDKL | Y206 | 0.278 | 87.22 | 275 | 0.121955 | 0.278 | 0.122 | 0.8462 |
| P78348 | ASIC1 | IQAFASSsTLHGLAH | S25 | 0.179 | 84.44 | 528 | 0.14229 | 0.179 | 0.1423 | 1 |
| P78348 | ASIC1 | ACRIDCEtRYLVENC | T316 | 0.023 | 94 | 528 | 0.015709 | 0.023 | 0.0157 | 1 |
| Q9Y5E8 | PCDHB15 | ENFYRLVtEGALDRE | T405 | 0.006 | 92.68 | 787 | 0.004098 | 0.006 | 0.0041 | 1 |
| Q99873 | PRMT1 | DrATLyVtAIEDrQY | T201 | 0.028 | 98.89 | 371 | 0.019124 | 0.028 | 0.0191 | 1 |
| Q99873 | PRMT1 | KRTGFSTsPEsPytH | S304 | 0.096 | 98.77 | 371 | 0.076312 | 0.096 | 0.0763 | 1 |
| Q99873 | PRMT1 | PytHWkQtVFYMEDY | T315 | 0.016 | 98.66 | 371 | 0.010928 | 0.016 | 0.0109 | 1 |
| Q13185 | CBX3 | FLMKWkDsDEADLVL | S145 | 0.179 | 86.45 | 183 | 0.14229 | 0.179 | 0.1423 | 1 |
| Q9NRG9 | AAAS | GSkILATtPSAVFRV | T306 | 0.093 | 88.83 | 546 | 0.06352 | 0.219 | 0.1496 | 1 |
| Q9UQ80 | PA2G4 | DAMIMEEtGkIFKKE | T60 | 0.017 | 98.09 | 394 | 0.011611 | 0.017 | 0.0116 | 1 |
| Q9UQ80 | PA2G4 | IAFPTSIsVNNCVCH | S81 | 0.038 | 98.81 | 394 | 0.030207 | 0.038 | 0.0302 | 0.875 |
| Q9UQ80 | PA2G4 | NNCVCHFsPLksDQD | S90 | 0.021 | 98.56 | 394 | 0.016693 | 0.021 | 0.0167 | 1 |
| Q13451 | FKBP5 | EPNAELIyEVTLkSF | Y243 | 0.015 | 98.82 | 457 | 0.00658 | 0.015 | 0.0066 | 1 |
| Q53H54 | TRMT12 | RNRVAPGsPCMLTQL | S82 | 0.091 | 89.07 | 448 | 0.072337 | 0.091 | 0.0723 | 1 |
| Q86VP6 | CAND1 | GSsyARSsVVTAVKF | S984 | 0.034 | 92.61 | 1230 | 0.027027 | 0.034 | 0.027 | 1 |
| Q86VP6 | CAND1 | VVTAVKFtISDHPQP | T992 | 0.02 | 91.48 | 1230 | 0.01366 | 0.02 | 0.0137 | 1 |
| Q86VP6 | CAND1 | kQDELkRsAMRAVAA | S1175 | 0.075 | 90.89 | 1230 | 0.059619 | 0.13 | 0.1056 | 1 |
| P26006 | ITGA3 | KEWDLSEysYKDPED | Y237 | 0.305 | 94.1 | 1051 | 0.133799 | 0.305 | 0.1338 | 1 |
| P26006 | ITGA3 | EWDLSEysYKDPEDQ | S238 | 0.128 | 93.94 | 1051 | 0.101749 | 0.128 | 0.1017 | 1 |
| P26006 | ITGA3 | PIIISMNysLPLRMP | Y574 | 0.039 | 92.37 | 1051 | 0.017109 | 0.039 | 0.0171 | 1 |
| Q10567 | AP1B1 | LEISGTFtRQVGSIS | T739 | 0.063 | 91.22 | 949 | 0.04303 | 0.094 | 0.0606 | 1 |
| Q9Y616 | IRAK3 | FQNIIEGtRNFHKDF | T162 | 0 | 94.36 | 596 | 0 | 0 | 0 | 0.8 |
| Q63HQ2 | EGFLAM | QAGKGRLssPRHVTT | S126 | 0.111 | 91.88 | 1017 | 0.088236 | 0.111 | 0.0882 | 1 |
| Q63HQ2 | EGFLAM | AMNSHGPsPRSWPSD | S225 | 0.017 | 92.08 | 1017 | 0.013514 | 0.017 | 0.0135 | 1 |
| Q8N5M1 | ATPAF2 | LQELRARtAAGtLFI | T266 | 0 | 98.61 | 289 | 0 | 0 | 0 | 0.9091 |
| Q8N5M1 | ATPAF2 | RARtAAGtLFIHLCS | T270 | 0.15 | 97.72 | 289 | 0.102452 | 0.15 | 0.1025 | 0.9 |
| Q96F25 | ALG14 | VTPRESLsILVVAGS | S38 | 0.17 | 93.45 | 216 | 0.135135 | 0.17 | 0.1351 | 1 |
| Q96F25 | ALG14 | IADtDEMsANKINSF | S77 | 0.01 | 91.85 | 216 | 0.007949 | 0.01 | 0.0079 | 1 |
| Q9HC35 | EML4 | KLKLEWAyGyRGKDC | Y265 | 0.081 | 98.37 | 981 | 0.035534 | 0.081 | 0.0355 | 0.8667 |
| Q9HC35 | EML4 | EERTQRHyLGHTDCV | Y305 | 0.042 | 98.76 | 981 | 0.018425 | 0.042 | 0.0184 | 1 |

| | | | | | | | | | | |
|---|---|---|---|---|---|---|---|---|---|---|
| Q9HC35 | EML4 | PDKIRIAtGQIAGVD | T326 | 0 | 98.9 | 981 | 0 | 0 | 0 | 1 |
| Q9HC35 | EML4 | GkGPKGVyQIsKQIK | Y499 | 0.129 | 93.91 | 981 | 0.05659 | 0.129 | 0.0566 | 1 |
| Q5U651 | RASIP1 | TEFFRKLsMAVNLLC | S833 | 0.186 | 95.32 | 963 | 0.147854 | 0.186 | 0.1479 | 1 |
| P21953 | BCKDHB | DVDTICKsVIKTGRL | S318 | 0.032 | 98.88 | 392 | 0.025437 | 0.032 | 0.0254 | 1 |
| P21953 | BCKDHB | IPDKWKCyDALRKMI | Y383 | 0.306 | 98.85 | 392 | 0.134238 | 0.306 | 0.1342 | 0.8889 |
| Q9Y6P5 | SESN1 | LPLHYRHyIGIMAAA | Y119 | 0.001 | 98.63 | 492 | 0.000439 | 0.106 | 0.0465 | 0.8462 |
| Q6UX73 | C16orf89 | tQGPLQQsQDyINLF | S220 | 0.041 | 92.94 | 402 | 0.032592 | 0.041 | 0.0326 | 1 |
| Q9UBF2 | COPG2 | RTPVPSkyIRFIFNR | Y472 | 0.048 | 94.67 | 871 | 0.021057 | 0.048 | 0.0211 | 1 |
| P29803 | PDHA2 | TRAEGLKyyRMMLTV | Y62 | 0.144 | 98.85 | 388 | 0.063171 | 0.144 | 0.0632 | 1 |
| P29803 | PDHA2 | RAEGLKyyRMMLTVR | Y63 | 0.144 | 98.87 | 388 | 0.063171 | 0.144 | 0.0632 | 1 |
| P29803 | PDHA2 | MELQTYRyHGHsMsD | Y287 | 0.134 | 98.58 | 388 | 0.058784 | 0.134 | 0.0588 | 1 |
| P29803 | PDHA2 | TYRyHGHsMsDPGVs | S291 | 0.131 | 97.94 | 388 | 0.104134 | 0.118 | 0.1451 | 1 |
| Q53GT1 | KLHL22 | DVHQVACySCTSGQW | Y517 | 0.119 | 98.35 | 634 | 0.052204 | 0.119 | 0.0522 | 1 |
| Q7L7V1 | DHX32 | RkQILGSssSGKFFC | S395 | 0.139 | 67.61 | 743 | 0.110493 | 0.139 | 0.1105 | 1 |
| O94826 | TOMM70 | TEkNVDLsTFyQNRA | S153 | 0.002 | 97.28 | 608 | 0.00159 | 0.002 | 0.0016 | 1 |
| Q96DR7 | ARHGEF26 | TFDPYVkyCTNEVyQ | Y532 | 0.008 | 95.71 | 871 | 0.003509 | 0.008 | 0.0035 | 1 |
| Q9H244 | P2RY12 | ARIPYTLsQtRDVFD | S262 | 0.056 | 92.61 | 342 | 0.044515 | 0.056 | 0.0445 | 1 |
| Q12882 | DPYD | VKIICGksLsVNEMT | S260 | 0.171 | 95.53 | 1025 | 0.13593 | 0.171 | 0.1359 | 1 |
| Q12882 | DPYD | IICGksLsVNEMTLS | S262 | 0.169 | 95.68 | 1025 | 0.134341 | 0.169 | 0.1343 | 1 |
| Q12882 | DPYD | LKADVVIsAFGsVLS | S436 | 0.043 | 98.3 | 1025 | 0.034181 | 0.043 | 0.0342 | 1 |
| Q12882 | DPYD | AGWGFALtKtFsLDk | T573 | 0.009 | 98.86 | 1025 | 0.006147 | 0.009 | 0.0061 | 1 |
| Q12882 | DPYD | WGFALtKtFsLDkDI | T575 | 0.045 | 98.07 | 1025 | 0.030736 | 0.045 | 0.0307 | 1 |
| Q12882 | DPYD | FALtKtFsLDkDIVT | S577 | 0.09 | 96.61 | 1025 | 0.071542 | 0.09 | 0.0715 | 1 |
| Q12882 | DPYD | RTTyGGVsGtAIRPI | S766 | 0.15 | 98.73 | 1025 | 0.119237 | 0.15 | 0.1192 | 1 |
| Q8NBN7 | RDH13 | KYNTkAAyCQSKLAI | Y200 | 0.162 | 95.92 | 331 | 0.071067 | 0.162 | 0.0711 | 1 |
| O75923 | DYSF | LAStTQYsRAVFDGC | S638 | 0.011 | 81.44 | 2080 | 0.008744 | 0.011 | 0.0087 | 0.8333 |
| O75923 | DYSF | GLTYPKFsDVTGkIK | S905 | 0.003 | 84.96 | 2080 | 0.002385 | 0.003 | 0.0024 | 1 |
| Q13402 | MYO7A | TLLTDSAtTAKELCN | T1278 | 0.047 | 88.7 | 2215 | 0.032102 | 0.032 | 0.0561 | 1 |
| Q9UK17 | KCND3 | GDCCYEEyKDRKREN | Y136 | 0.113 | 91.02 | 655 | 0.049571 | 0.113 | 0.0496 | 0.8182 |
| Q9BXT2 | CACNG6 | SGLLLLVsLEVFRHs | S188 | 0.008 | 92.81 | 260 | 0.006359 | 0.17 | 0.1351 | 1 |
| P52888 | THOP1 | GEVVGKFyLDLyPRE | Y412 | 0.015 | 98.51 | 689 | 0.00658 | 0.015 | 0.0066 | 0.8 |

| | | | | | | | | | | |
|---|---|---|---|---|---|---|---|---|---|---|
| P52888 | THOP1 | NSkVGMDyRsCILRP | Y641 | 0.018 | 98.52 | 689 | 0.007896 | 0.018 | 0.0079 | 0.9286 |
| Q8ND71 | GIMAP8 | AIFGADFtKYAIMLF | T557 | 0.059 | 90.83 | 665 | 0.040298 | 0.059 | 0.0403 | 1 |
| P61353 | RPL27 | IKSFVkVyNyNHLMP | Y75 | 0.081 | 96.86 | 136 | 0.035534 | 0.081 | 0.0355 | 0.8182 |
| P00325 | ADH1B | TSTFSQytVVDENAV | T151 | 0 | 98.95 | 375 | 0 | 0 | 0 | 1 |
| P28066 | PSMA5 | DrGVNtFsPEGrLFQ | S16 | 0.091 | 89.75 | 241 | 0.072337 | 0.091 | 0.0723 | 1 |
| Q15418 | RPS6KA1 | VKGAMAAtysALNSS | T701 | 0.015 | 80.94 | 735 | 0.010245 | 0.015 | 0.0102 | 0.8333 |
| O43791 | SPOP | EETKAMEsQRAYRFV | S119 | 0.043 | 98.05 | 374 | 0.034181 | 0.043 | 0.0342 | 0.8571 |
| Q9NZC9 | SMARCAL1 | AGLINIVsFDLLsKL | S528 | 0.018 | 91.68 | 954 | 0.014308 | 0.018 | 0.0143 | 1 |
| O43913 | ORC5 | AVLNFPkyCEPVVkG | Y236 | 0.02 | 86.41 | 435 | 0.008774 | 0.02 | 0.0088 | 1 |
| P43243 | MATR3 | sNLPHsGysDsAVLk | Y508 | 0.211 | 82.34 | 847 | 0.092563 | 0.211 | 0.0926 | 1 |
| P49748 | ACADVL | VVVLSRAsRSLSEGH | S586 | 0.043 | 98.87 | 655 | 0.034181 | 0.043 | 0.0342 | 1 |
| P38432 | COIL | VEtPKkDysLLPLLA | Y461 | 0.312 | 87.91 | 576 | 0.13687 | 0.312 | 0.1369 | 1 |
| Q96AY4 | TTC28 | LGNMGAVyIAMGDFE | Y364 | 0.236 | 90.69 | 2481 | 0.10353 | 0.236 | 0.1035 | 0.8 |
| Q96AY4 | TTC28 | HGNLAVAyQALGAHD | Y564 | 0.132 | 91.59 | 2481 | 0.057907 | 0.132 | 0.0579 | 0.8 |
| Q96AY4 | TTC28 | YGNLGLTyESLGTFE | Y1047 | 0.15 | 90.99 | 2481 | 0.065803 | 0.15 | 0.0658 | 0.8 |
| P07686 | HEXB | HRGILIDtsRHYLPV | T209 | 0 | 98.87 | 556 | 0 | 0 | 0 | 0.9 |
| P07686 | HEXB | RGILIDtsRHYLPVK | S210 | 0 | 98.84 | 556 | 0 | 0 | 0 | 1 |
| Q7Z628 | NET1 | QEAIyEMsRGEQDLI | S183 | 0.172 | 95.85 | 596 | 0.136725 | 0.172 | 0.1367 | 1 |
| Q9Y639 | NPTN | RAEDSGEyHCVyHFV | Y216 | 0.031 | 97.83 | 398 | 0.013599 | 0.031 | 0.0136 | 0.9286 |
| Q9Y639 | NPTN | SGEyHCVyHFVSAPK | Y220 | 0 | 96.69 | 398 | 0 | 0 | 0 | 1 |
| Q9UBB4 | ATXN10 | ASsLQLItECFRCLR | T82 | 0.095 | 97.94 | 475 | 0.064886 | 0.095 | 0.0649 | 1 |
| Q9UBB4 | ATXN10 | EWPFLIItDLFLKSP | T221 | 0.057 | 95.92 | 475 | 0.038932 | 0.057 | 0.0389 | 1 |
| Q9P2H3 | IFT80 | SAGEDCKyKVWDsYG | Y212 | 0.038 | 98.61 | 777 | 0.01667 | 0.038 | 0.0167 | 1 |
| Q9P2H3 | IFT80 | PKFPGMRtDILNAQT | T414 | 0.05 | 89.78 | 777 | 0.034151 | 0.082 | 0.056 | 1 |
| Q9P2H3 | IFT80 | TVSLSNDtIAIRDKA | T428 | 0.021 | 96.55 | 777 | 0.014343 | 0.021 | 0.0143 | 1 |
| O15143 | ARPC1B | DFKCRIFsAyIkEVE | S170 | 0 | 98.78 | 372 | 0 | 0 | 0 | 1 |
| O15143 | ARPC1B | TGMDGGMsIWDVKsL | S355 | 0.001 | 97.23 | 372 | 0.000795 | 0.001 | 0.0008 | 1 |
| P45880 | VDAC2 | ICQGLKLtFDttFsP | T109 | 0.145 | 97.41 | 294 | 0.099037 | 0.145 | 0.099 | 0.875 |
| P45880 | VDAC2 | sLIGVGytQtLRPGV | T259 | 0.163 | 98.16 | 294 | 0.111331 | 0.163 | 0.1113 | 1 |
| Q9BQA1 | WDR77 | GALLLGAssLsGRCW | S47 | 0.014 | 97.02 | 342 | 0.011129 | 0.014 | 0.0111 | 1 |
| Q8NHU6 | TDRD7 | NFPDPHRsINRQITN | S831 | 0 | 90.89 | 1098 | 0 | 0 | 0 | 0.875 |

| | | | | | | | | | | |
|---|---|---|---|---|---|---|---|---|---|---|
| Q5THR3 | EFCAB6 | KKRFLDFsKEPNGKI | S647 | 0.149 | 78 | 1501 | 0.118442 | 0.149 | 0.1184 | 1 |
| Q9ULC5 | ACSL5 | HGESLRssLVGVVVP | S581 | 0.133 | 84.44 | 683 | 0.105724 | 0.172 | 0.1367 | 1 |
| Q9ULC5 | ACSL5 | RTQIDSLyEHIQD__ | Y678 | 0.087 | 87.75 | 683 | 0.038166 | 0.087 | 0.0382 | 1 |
| P19404 | NDUFV2 | GIKVGETtPDKLFTL | T164 | 0.062 | 95.57 | 249 | 0.042347 | 0.062 | 0.0423 | 1 |
| P19404 | NDUFV2 | VQINDNYyEDLTAKD | Y193 | 0.03 | 97.77 | 249 | 0.013161 | 0.03 | 0.0132 | 1 |
| Q9UHC7 | MKRN1 | LSNCNHTyCLKCIRK | Y309 | 0.005 | 95.27 | 482 | 0.002193 | 0.005 | 0.0022 | 1 |
| Q8NHU6 | TDRD7 | EDEMKEyySKNPKIT | Y502 | 0.151 | 91.14 | 1098 | 0.066242 | 0.151 | 0.0662 | 1 |
| P26358 | DNMT1 | WDGFFsttVTNPEPM | T1526 | 0.085 | 80.69 | 1616 | 0.058056 | 0.085 | 0.0581 | 1 |
| Q96FZ2 | HMCES | NksPQSNsPVLLSRL | S50 | 0.01 | 98.74 | 354 | 0.007949 | 0.01 | 0.0079 | 1 |
| Q6ZNA4 | RNF111 | NrGAsQGtIERCTYP | T901 | 0.14 | 92.56 | 994 | 0.095622 | 0.14 | 0.0956 | 1 |
| O14734 | ACOT8 | DPkLPVLyQVERTRT | Y96 | 0 | 98.67 | 319 | 0 | 0 | 0 | 1 |
| Q9NZC7 | WWOX | RQRYDGStTAMEILQ | T111 | 0.128 | 88.75 | 414 | 0.087426 | 0.128 | 0.0874 | 1 |
| Q92616 | GCN1 | DEkkQVFtSEKFLVM | T526 | 0.052 | 87.98 | 2671 | 0.035517 | 0.052 | 0.0355 | 1 |
| Q92616 | GCN1 | LCTVLHLtERLFLDH | T546 | 0.028 | 90.82 | 2671 | 0.019124 | 0.028 | 0.0191 | 1 |
| Q6ZMQ8 | AATK | HCKYREDyFVtADQL | Y283 | 0.131 | 92.72 | 1374 | 0.057468 | 0.131 | 0.0575 | 1 |
| Q6ZMQ8 | AATK | YREDyFVtADQLWVP | T286 | 0.088 | 86.69 | 1374 | 0.060105 | 0.088 | 0.0601 | 1 |
| Q6ZMT4 | KDM7A | SKPVQAGtRTFIKEL | T119 | 0.026 | 98.53 | 941 | 0.017758 | 0.026 | 0.0178 | 1 |
| Q6ZMT4 | KDM7A | SKMTLHNyVKYFMNP | Y210 | 0.003 | 98.32 | 941 | 0.001316 | 0.003 | 0.0013 | 1 |
| O60231 | DHX16 | ELKVLVAsATMDTAR | S552 | 0.006 | 88.56 | 1041 | 0.004769 | 0.187 | 0.1486 | 1 |
| P16885 | PLCG2 | FRLWLCGsQMVALNF | S1007 | 0.006 | 96.67 | 1265 | 0.004769 | 0.006 | 0.0048 | 1 |
| P16885 | PLCG2 | LFsLNGRtGyVLQPE | T1034 | 0.042 | 96.03 | 1265 | 0.028687 | 0.042 | 0.0287 | 1 |
| P16885 | PLCG2 | sLNGRtGyVLQPESM | Y1036 | 0.003 | 97.99 | 1265 | 0.001316 | 0.003 | 0.0013 | 1 |
| P16885 | PLCG2 | AVKSGFRsVPLKNGY | S1164 | 0.004 | 98.34 | 1265 | 0.00318 | 0.004 | 0.0032 | 0.8182 |
| P15882 | CHN1 | ADIsVNMyEDINIIt | Y333 | 0.065 | 96.82 | 459 | 0.028515 | 0.13 | 0.057 | 1 |
| P15882 | CHN1 | yEDINIItGALKLyF | T340 | 0 | 98.46 | 459 | 0 | 0 | 0 | 1 |
| P15882 | CHN1 | ItGALKLyFRDLPIP | Y346 | 0.006 | 98.13 | 459 | 0.002632 | 0.006 | 0.0026 | 1 |
| O95388 | CCN4 | TSCGLGVstRISNVN | S235 | 0.02 | 92.66 | 367 | 0.015898 | 0.02 | 0.0159 | 1 |
| Q92729 | PTPRU | ITQYEISyQSIESSD | Y524 | 0.293 | 92.73 | 1446 | 0.128535 | 0.293 | 0.1285 | 1 |
| Q92729 | PTPRU | NLHPGTTyLFSVRAR | Y563 | 0.136 | 92.24 | 1446 | 0.059661 | 0.136 | 0.0597 | 1 |
| O00567 | NOP56 | IINDNATyCRLAQFI | Y210 | 0.001 | 90.84 | 594 | 0.000439 | 0.001 | 0.0004 | 1 |
| O00567 | NOP56 | LINIESFsSRVVSLS | S268 | 0.001 | 91.99 | 594 | 0.000795 | 0.001 | 0.0008 | 1 |

| | | | | | | | | | | |
|---|---|---|---|---|---|---|---|---|---|---|
| O00567 | NOP56 | AKNKGRIsRYLANkC | S369 | 0 | 92.25 | 594 | 0 | 0 | 0 | 1 |
| Q9NXE8 | CWC25 | KKsWHPQtLRNVEKV | T16 | 0.186 | 92.04 | 425 | 0.127041 | 0.186 | 0.127 | 1 |
| Q13557 | CAMK2D | sMMHRQEtVDCLKKF | T287 | 0.049 | 92.66 | 499 | 0.033468 | 0.049 | 0.0335 | 1 |
| Q53HC5 | KLHL26 | RSVSSkVyQLPEPGA | Y331 | 0.198 | 97.73 | 615 | 0.08686 | 0.198 | 0.0869 | 1 |
| O15037 | KHNYN | HGLQHyFssRGIAIA | S458 | 0 | 94.71 | 678 | 0 | 0 | 0 | 0.8 |
| O15037 | KHNYN | GLQHyFssRGIAIAV | S459 | 0 | 94.4 | 678 | 0 | 0 | 0 | 1 |
| Q9BQS7 | HEPH | GLIGPLItCKRGALD | T205 | 0 | 96.73 | 1158 | 0 | 0 | 0 | 1 |
| Q9NY15 | STAB1 | LPADRRVtALVPsEA | T1022 | 0 | 83.57 | 2570 | 0 | 0 | 0 | 0.9 |
| O14921 | RGS13 | tKYGPVVyAAyLKME | Y48 | 0 | 97.55 | 159 | 0 | 0 | 0 | 0.9333 |
| O14921 | RGS13 | GPVVyAAyLKMEHSD | Y51 | 0.16 | 95.92 | 159 | 0.07019 | 0.32 | 0.1404 | 1 |
| O14921 | RGS13 | RFLkSEMyQkLLKTM | Y146 | 0.077 | 95.44 | 159 | 0.033779 | 0.077 | 0.0338 | 1 |
| Q8TD43 | TRPM4 | FLRLSDRtDPAAVYs | T97 | 0.17 | 93.86 | 1214 | 0.116112 | 0.17 | 0.1161 | 1 |
| Q9NS86 | LANCL2 | IESGkTLsREERKTE | S239 | 0 | 98.26 | 450 | 0 | 0 | 0 | 1 |
| Q92620 | DHX38 | VRKCIVAtNIAETSL | T805 | 0.144 | 91.37 | 1227 | 0.098354 | 0.144 | 0.0984 | 1 |
| O75159 | SOCS5 | EQWNHNFsFDAHDPC | S438 | 0.074 | 94.97 | 536 | 0.058824 | 0.074 | 0.0588 | 1 |
| Q96GX1 | TCTN2 | CVTNLELyQERDGII | Y327 | 0.201 | 79.73 | 697 | 0.088176 | 0.201 | 0.0882 | 1 |
| P00488 | F13A1 | GILDTCLyVMDRAQM | Y241 | 0.32 | 98.51 | 732 | 0.140379 | 0.32 | 0.1404 | 0.875 |
| Q96KG9 | SCYL1 | sTRHRVLtSAFSRAT | T469 | 0.152 | 94.08 | 808 | 0.103818 | 0.152 | 0.1038 | 1 |
| P05089 | ARG1 | sLAIGsIsGHARVHP | S109 | 0.102 | 98.93 | 322 | 0.081081 | 0.161 | 0.128 | 0.9091 |
| P78509 | RELN | DPsIIVLyAKNNSAD | Y286 | 0.114 | 95.595 | 3460 | 0.05001 | 0.114 | 0.05 | 1 |
| P78509 | RELN | LCTPsMDtTGyGNLR | T468 | 0.004 | 94.22 | 3460 | 0.002732 | 0.004 | 0.0027 | 1 |
| P78509 | RELN | LITSFLDsSQSRFLQ | S764 | 0 | 89.105 | 3460 | 0 | 0 | 0 | 1 |
| P78509 | RELN | QSRFLQFtLRLGSkS | T773 | 0.06 | 83.515 | 3460 | 0.040981 | 0.116 | 0.0792 | 1 |
| Q7Z6J6 | FRMD5 | SGQtPAtsELNFLRk | S191 | 0 | 97.8 | 570 | 0 | 0 | 0 | 1 |
| Q14562 | DHX8 | LCKMLIMsVHLGCSE | S996 | 0 | 89.99 | 1220 | 0 | 0 | 0 | 1 |
| P30874 | SSTR2 | IFCLTVMsIDRYLAV | S137 | 0.001 | 94.85 | 369 | 0.000795 | 0.067 | 0.0533 | 0.8571 |
| Q86Y38 | XYLT1 | VLQVSRQysNVRVTP | Y378 | 0.127 | 98.56 | 959 | 0.055713 | 0.127 | 0.0557 | 1 |
| P34059 | GALNS | LSFASAEyQEALSRI | Y461 | 0.057 | 98.83 | 522 | 0.025005 | 0.057 | 0.025 | 1 |
| P34931 | HSPA1L | GNRttPsyVAFtDtE | Y43 | 0.07 | 98.29 | 641 | 0.030708 | 0.133 | 0.0566 | 0.8 |
| P34931 | HSPA1L | VAMNPQNtVFDAkRL | T68 | 0 | 97.74 | 641 | 0 | 0 | 0 | 0.9091 |
| P34931 | HSPA1L | VLtKLKEtAEAFLGH | T132 | 0.131 | 97.7 | 641 | 0.089475 | 0.142 | 0.1263 | 1 |

| | | | | | | | | | | |
|---|---|---|---|---|---|---|---|---|---|---|
| P34931 | HSPA1L | PVTNAVItVPAyFND | T147 | 0.007 | 97.51 | 641 | 0.004781 | 0.007 | 0.0048 | 0.9167 |
| P34931 | HSPA1L | PtAAAIAyGLDkGGQ | Y185 | 0.12 | 92.22 | 641 | 0.052642 | 0.17 | 0.0746 | 1 |
| Q9H0S4 | DDX47 | ARsILLAtDVASRGL | T320 | 0.109 | 92.16 | 455 | 0.074448 | 0.218 | 0.1489 | 1 |
| P82932 | MRPS6 | ____MPRyELALILK | Y4 | 0.212 | 95.91 | 125 | 0.093001 | 0.212 | 0.093 | 1 |
| P43115 | PTGER3 | LVGQLLTtPVVIVVY | T107 | 0.187 | 92.89 | 390 | 0.127724 | 0.187 | 0.1277 | 1 |
| O14657 | TOR1B | NAGGDLItKtALDFW | T222 | 0.133 | 93.58 | 336 | 0.090841 | 0.133 | 0.0908 | 0.8 |
| O14657 | TOR1B | GGDLItKtALDFWRA | T224 | 0.196 | 93.89 | 336 | 0.133871 | 0.196 | 0.1339 | 1 |
| Q14139 | UBE4A | LLTFNPTyCALkELN | Y464 | 0.05 | 97.25 | 1066 | 0.021934 | 0.05 | 0.0219 | 1 |
| Q86Y56 | DNAAF5 | SkMTRLIsCRIINtF | S721 | 0 | 98.18 | 855 | 0 | 0 | 0 | 1 |
| P04062 | GBA1 | VGFLETIsPGYSIHT | S523 | 0.003 | 98.38 | 536 | 0.002385 | 0.003 | 0.0024 | 1 |
| P26640 | VARS1 | QLkkLGSsLDWDRAC | S437 | 0.002 | 98.18 | 1264 | 0.00159 | 0.002 | 0.0016 | 1 |
| P26640 | VARS1 | WGHRIPAyFVtVSDP | Y734 | 0.224 | 97.45 | 1264 | 0.098266 | 0.224 | 0.0983 | 1 |
| P26640 | VARS1 | LRFGLCAyMSQGRDI | Y929 | 0.074 | 88.94 | 1264 | 0.032463 | 0.074 | 0.0325 | 1 |
| P26640 | VARS1 | DVGLRLLsPFMPFVT | S1061 | 0.031 | 98.51 | 1264 | 0.024642 | 0.031 | 0.0246 | 0.8 |
| Q5SWX8 | ODR4 | TKKIFCRtYDIHDPk | T151 | 0.035 | 95.42 | 454 | 0.023905 | 0.035 | 0.0239 | 0.8 |
| Q9BWV3 | CDADC1 | LLLATVAssVPNFKH | S288 | 0.009 | 96.6 | 514 | 0.007154 | 0.009 | 0.0072 | 0.9 |
| Q58FF7 | HSP90AB3P | FIHGVVDsEDLPLNI | S325 | 0.029 | 77.62 | 597 | 0.023053 | 0.029 | 0.0231 | 1 |
| O75716 | STK16 | AAQRCTIsyRAPELF | S197 | 0.11 | 96.83 | 305 | 0.087441 | 0.11 | 0.0874 | 1 |
| O75716 | STK16 | AQRCTIsyRAPELFS | Y198 | 0.172 | 98.19 | 305 | 0.075454 | 0.172 | 0.0755 | 1 |
| O00206 | TLR4 | YGRGENIyDAFVIyS | Y674 | 0.269 | 89.25 | 839 | 0.118006 | 0.269 | 0.118 | 0.8333 |
| O00206 | TLR4 | IHEGFHKsrKVIVVV | S730 | 0.065 | 87.7 | 839 | 0.051669 | 0.114 | 0.0906 | 0.8 |
| P52333 | JAK3 | LLYRIRFyFPNWFGL | Y105 | 0.258 | 92.18 | 1124 | 0.113181 | 0.301 | 0.132 | 0.9167 |
| P52333 | JAK3 | ELLkTVSyKACLPPS | Y190 | 0.16 | 91.03 | 1124 | 0.07019 | 0.16 | 0.0702 | 1 |
| P52333 | JAK3 | RHSLMAkyIMDLERL | Y238 | 0.014 | 94.54 | 1124 | 0.006142 | 0.014 | 0.0061 | 1 |
| P52333 | JAK3 | tGGSRPGsyVLRRSP | S398 | 0.043 | 94.83 | 1124 | 0.034181 | 0.043 | 0.0342 | 1 |
| P52333 | JAK3 | GGSRPGsyVLRRSPQ | Y399 | 0.123 | 94.66 | 1124 | 0.053958 | 0.123 | 0.054 | 1 |
| Q4KMQ2 | ANO6 | DCKFRRQsEDPsCPN | S256 | 0.156 | 85.17 | 910 | 0.124007 | 0.156 | 0.124 | 1 |
| Q9Y3C8 | UFC1 | EyQsLIRyVENNkNA | Y42 | 0.114 | 97.29 | 167 | 0.05001 | 0.114 | 0.05 | 0.9231 |
| P15085 | CPA1 | ALASLyGtKFNYGsI | T346 | 0.127 | 98.77 | 419 | 0.086743 | 0.127 | 0.0867 | 1 |
| P15085 | CPA1 | GtKFNYGsIIKAIYQ | S352 | 0.142 | 98.6 | 419 | 0.112878 | 0.142 | 0.1129 | 1 |
| P78527 | PRKDC | QTECRHksIELFYkF | S1187 | 0 | 93.69 | 4128 | 0 | 0 | 0 | 1 |

| | | | | | | | | | | |
|---|---|---|---|---|---|---|---|---|---|---|
| Q6RI45 | BRWD3 | QMGDELIyFRQGHEA | Y948 | 0 | 93.63 | 1802 | 0 | 0 | 0 | 1 |
| Q6RI45 | BRWD3 | GESFSIKyHDMPDVI | Y1024 | 0.152 | 89.03 | 1802 | 0.06668 | 0.152 | 0.0667 | 1 |
| Q6RI45 | BRWD3 | VKAAKIVtDVLLRFI | T1235 | 0.014 | 91.47 | 1802 | 0.009562 | 0.014 | 0.0096 | 1 |
| Q14315 | FLNC | LNPSGAktDtyVtDN | T1300 | 0.127 | 88.77 | 2725 | 0.086743 | 0.127 | 0.0867 | 0.9167 |
| P01009 | SERPINA1 | QHCkkLSsWVLLMKY | S261 | 0 | 98.66 | 418 | 0 | 0 | 0 | 1 |
| P62851 | RPS25 | kHRAQVIyTRNTkGG | Y109 | 0.139 | 92.54 | 125 | 0.060977 | 0.139 | 0.061 | 0.8182 |
| P25391 | LAMA1 | SVAKQKKsLNtAGQF | S348 | 0 | 92.25 | 3075 | 0 | 0 | 0 | 1 |
| Q7L1I2 | SV2B | FLFCRLIsGIGIGGA | S208 | 0 | 95.91 | 683 | 0 | 0 | 0 | 0.9 |
| Q9UJU6 | DBNL | VRVVtEksPtDWALF | S24 | 0.169 | 96 | 430 | 0.134341 | 0.169 | 0.1343 | 1 |
| Q9UJU6 | DBNL | VVtEksPtDWALFTY | T26 | 0.075 | 98.03 | 430 | 0.051226 | 0.075 | 0.0512 | 1 |
| Q9P2E9 | RRBP1 | KKVEGAPtQGRKAEG | T287 | 0.041 | 78.41 | 1410 | 0.028004 | 0.041 | 0.028 | 1 |
| Q8IX01 | SUGP2 | GPEIEQFsIENstDN | S806 | 0.122 | 81.75 | 1082 | 0.09698 | 0.122 | 0.097 | 0.8333 |
| Q8IX01 | SUGP2 | EQFsIENstDNPDLW | S810 | 0.042 | 69.48 | 1082 | 0.033386 | 0.042 | 0.0334 | 1 |
| Q14C87 | TMEM132D | DVLTFPVsIsRNsTE | S310 | 0.091 | 91.99 | 1099 | 0.072337 | 0.122 | 0.097 | 0.9 |
| O60308 | CEP104 | LKNTLRAsVFLVRRA | S481 | 0 | 97.89 | 925 | 0 | 0 | 0 | 1 |
| O60308 | CEP104 | TAHCVERtIPVLLtR | T529 | 0.002 | 97.94 | 925 | 0.001366 | 0.002 | 0.0014 | 1 |
| O60308 | CEP104 | KDLGtGssGFtIDNV | S603 | 0.038 | 91.58 | 925 | 0.030207 | 0.038 | 0.0302 | 1 |
| Q9Y5R2 | MMP24 | SKEGyYTyFYKGRDY | Y537 | 0.295 | 92.92 | 645 | 0.129412 | 0.295 | 0.1294 | 0.9 |
| P35232 | PHB1 | NVPVITGskDLQNVN | S82 | 0.07 | 92.25 | 272 | 0.055644 | 0.07 | 0.0556 | 1 |
| P35232 | PHB1 | ERVLPsItTEILkSV | T123 | 0.016 | 94.17 | 272 | 0.010928 | 0.016 | 0.0109 | 1 |
| P51993 | FUT6 | NSARVRyyQSLQAHL | Y209 | 0.214 | 98.17 | 359 | 0.093879 | 0.214 | 0.0939 | 1 |
| P51993 | FUT6 | AHLKVDVyGRSHKPL | Y221 | 0.204 | 98.22 | 359 | 0.089492 | 0.204 | 0.0895 | 1 |
| Q9UJX2 | CDC23 | LLLIAGLtRERGLLH | T43 | 0.008 | 95.81 | 597 | 0.005464 | 0.156 | 0.1066 | 0.8889 |
| Q9UJX2 | CDC23 | AyFLYMysRYLSGEK | S121 | 0 | 97.34 | 597 | 0 | 0 | 0 | 1 |
| Q9HCG8 | CWC22 | VLECIKLsEETTTss | S571 | 0.135 | 82.02 | 908 | 0.107313 | 0.135 | 0.1073 | 1 |
| Q86VI1 | EXOC3L1 | GQQFLEAyVSLRELE | Y163 | 0.198 | 93.04 | 746 | 0.08686 | 0.222 | 0.0974 | 1 |
| P17066 | HSPA6 | GNRttPsyVAFtDtE | Y43 | 0.166 | 98.25 | 643 | 0.072822 | 0.293 | 0.1146 | 0.8 |
| P17066 | HSPA6 | PTAAAIAyGLDRRGA | Y185 | 0.124 | 92.92 | 643 | 0.054397 | 0.124 | 0.0544 | 1 |
| P17066 | HSPA6 | LLDVAPLsLGLEtAG | S402 | 0.039 | 90.52 | 643 | 0.031002 | 0.039 | 0.031 | 1 |
| Q9NSD9 | FARSB | GASDVVLykIDVPAN | Y63 | 0.002 | 95.72 | 589 | 0.000877 | 0.002 | 0.0009 | 1 |
| Q9UHP3 | USP25 | QVLYLDRyMHRNREI | Y404 | 0.005 | 96.96 | 1055 | 0.002193 | 0.005 | 0.0022 | 0.9231 |

| | | | | | | | | | | |
|---|---|---|---|---|---|---|---|---|---|---|
| P16615 | ATP2A2 | AMGVVVAtGVNtEIG | T226 | 0.044 | 94.8 | 1042 | 0.030053 | 0.044 | 0.0301 | 1 |
| P16615 | ATP2A2 | VVAtGVNtEIGKIRD | T230 | 0.023 | 93.1 | 1042 | 0.015709 | 0.023 | 0.0157 | 1 |
| Q9P2K1 | CC2D2A | GPTAYVLtWEQGRYL | T1401 | 0 | 92.68 | 1620 | 0 | 0 | 0 | 1 |
| Q14997 | PSME4 | VkCCPEEsLKLFVPH | S642 | 0 | 94.77 | 1843 | 0 | 0 | 0 | 1 |
| O95861 | BPNT1 | MRLVASAySIAQKAG | Y16 | 0.263 | 98.87 | 308 | 0.115374 | 0.263 | 0.1154 | 1 |
| O95861 | BPNT1 | ADRLAQMsICSsLAR | S57 | 0 | 98.05 | 308 | 0 | 0 | 0 | 1 |
| O95861 | BPNT1 | AQMsICSsLARKFPK | S61 | 0.002 | 97.96 | 308 | 0.00159 | 0.002 | 0.0016 | 1 |
| O95861 | BPNT1 | ASAYVFAsPGCkKWD | S240 | 0.06 | 97.87 | 308 | 0.047695 | 0.12 | 0.0954 | 1 |
| O43422 | THAP12 | EYCRGQAyIVSSGFS | Y330 | 0.041 | 93.24 | 761 | 0.017986 | 0.041 | 0.018 | 0.9167 |
| O95573 | ACSL3 | kNWEEGGyFNTDkPH | Y516 | 0.291 | 91.62 | 720 | 0.127658 | 0.291 | 0.1277 | 1 |
| O95715 | CXCL14 | EKMVIITtKSVSRYR | T72 | 0.016 | 95.38 | 111 | 0.010928 | 0.016 | 0.0109 | 1 |
| P61601 | NCALD | LEIVQAIyKMVSSVM | Y129 | 0.26 | 87.42 | 193 | 0.114058 | 0.26 | 0.1141 | 0.9 |
| O00222 | GRM8 | NGDAPGRyDIFQYQI | Y471 | 0.025 | 94.05 | 908 | 0.010967 | 0.025 | 0.011 | 0.8571 |
| Q5TCH4 | CYP4A22 | GKVRVQLyDPDyMkV | Y100 | 0.038 | 97.6 | 519 | 0.01667 | 0.038 | 0.0167 | 0.9333 |
| P22676 | CALB2 | NELKGFLsDLLKKAN | S137 | 0.153 | 84.31 | 271 | 0.121622 | 0.153 | 0.1216 | 1 |
| Q9BWV1 | BOC | MLGESEPsAPsRPyV | S693 | 0.027 | 92.99 | 1114 | 0.021463 | 0.027 | 0.0215 | 0.8 |
| Q7L523 | RRAGA | DtRRLGAtIDVEHSH | T42 | 0.154 | 94.34 | 313 | 0.105184 | 0.154 | 0.1052 | 1 |
| P06681 | C2 | VtDQFLCsGtQEDEs | S666 | 0.001 | 98.28 | 752 | 0.000795 | 0.001 | 0.0008 | 1 |
| P06681 | C2 | DQFLCsGtQEDEsPC | T668 | 0.201 | 93.58 | 752 | 0.137286 | 0.201 | 0.1373 | 1 |
| P06681 | C2 | sGtQEDEsPCKGESG | S673 | 0.077 | 96.07 | 752 | 0.061208 | 0.077 | 0.0612 | 1 |
| P01031 | C5 | SMDIDVSyKHKGALH | Y1317 | 0.169 | 91.2 | 1676 | 0.074138 | 0.169 | 0.0741 | 1 |
| P49327 | FASN | ASRLsIPtyGLQCTR | T2268 | 0 | 93.92 | 2511 | 0 | 0 | 0 | 1 |
| O15460 | P4HA2 | AEKELVQsLkEYILV | S43 | 0.144 | 93.73 | 535 | 0.114468 | 0.071 | 0.1337 | 1 |
| P84022 | SMAD3 | EVFAECLsDSAIFVQ | S309 | 0.057 | 98.38 | 425 | 0.04531 | 0.057 | 0.0453 | 1 |
| Q9Y5Y9 | SCN10A | IAFDPYyyFQKKWNI | Y724 | 0.001 | 85.52 | 1956 | 0.000439 | 0.001 | 0.0004 | 0.8182 |
| O75027 | ABCB7 | VLIGyGVsRAGAAFF | S199 | 0.119 | 94.07 | 752 | 0.094595 | 0.119 | 0.0946 | 1 |
| O75027 | ABCB7 | SkAIDRGtRGIsFVL | T257 | 0.142 | 89.48 | 752 | 0.096988 | 0.142 | 0.097 | 1 |
| O75027 | ABCB7 | YETASLKstSTLAML | S370 | 0.001 | 93.3 | 752 | 0.000795 | 0.001 | 0.0008 | 1 |
| O75027 | ABCB7 | DVVKHRTsIFIAHRL | S660 | 0.001 | 93.89 | 752 | 0.000795 | 0.001 | 0.0008 | 1 |
| Q8IY81 | FTSJ3 | LTLYHRTsVTDFLRA | S253 | 0.173 | 91.5 | 847 | 0.13752 | 0.173 | 0.1375 | 1 |
| Q02338 | BDH1 | KFGVEAFsDCLRYEM | S219 | 0.02 | 98.84 | 343 | 0.015898 | 0.02 | 0.0159 | 1 |

| | | | | | | | | | | |
|---|---|---|---|---|---|---|---|---|---|---|
| Q02338 | BDH1 | FIAATsLySPESIQA | Y248 | 0.142 | 87.77 | 343 | 0.062293 | 0.142 | 0.0623 | 1 |
| P51957 | NEK4 | ELIRTMLsKRPEERP | S243 | 0.159 | 96.58 | 841 | 0.126391 | 0.159 | 0.1264 | 1 |
| Q15428 | SF3A2 | LLMAAEPyETIAFKV | Y176 | 0.236 | 93.69 | 464 | 0.10353 | 0.236 | 0.1035 | 1 |
| Q9NVX7 | KBTBD4 | LSNAVIyyRVGDNVW | Y351 | 0.065 | 97.56 | 518 | 0.028515 | 0.207 | 0.0908 | 1 |
| Q9P126 | CLEC1B | HNLtWEEsKQYCtDM | S126 | 0 | 98.82 | 229 | 0 | 0 | 0 | 1 |
| A2RTX5 | TARS3 | EVLsPNMyNsKLWEA | Y457 | 0.294 | 97.01 | 802 | 0.128974 | 0.294 | 0.129 | 1 |
| A2RTX5 | TARS3 | LsPNMyNsKLWEASG | S459 | 0.129 | 95.45 | 802 | 0.102544 | 0.129 | 0.1025 | 1 |
| P61916 | NPC2 | PIQKDktysYLNkLP | Y107 | 0.163 | 98.74 | 151 | 0.071506 | 0.163 | 0.0715 | 0.9167 |
| Q4LE39 | ARID4B | KLTDASWytVVFDDG | Y89 | 0.007 | 96.21 | 1312 | 0.003071 | 0.007 | 0.0031 | 0.9333 |
| O95352 | ATG7 | LPAVIAAsKRKLVIN | S490 | 0.067 | 97.08 | 703 | 0.053259 | 0.067 | 0.0533 | 1 |
| Q96CT2 | KLHL29 | TKAEELVyEtVIKWI | Y498 | 0.062 | 94.2 | 875 | 0.027199 | 0.062 | 0.0272 | 0.9286 |
| Q96CT2 | KLHL29 | AEELVyEtVIKWIKK | T500 | 0.026 | 94.64 | 875 | 0.017758 | 0.026 | 0.0178 | 1 |
| Q9BYK8 | HELZ2 | LRILGLEySLRVPPS | Y1551 | 0.241 | 86.53 | 2649 | 0.105723 | 0.241 | 0.1057 | 1 |
| Q9BYK8 | HELZ2 | RLRsDCFyEQPDEDG | Y1754 | 0.262 | 90.57 | 2649 | 0.114936 | 0.262 | 0.1149 | 1 |
| Q9HC62 | SENP2 | LRKKCLKyLDsMGQK | Y493 | 0.012 | 98.55 | 589 | 0.005264 | 0.012 | 0.0053 | 0.8667 |
| Q9HC62 | SENP2 | KCLKyLDsMGQKGHR | S496 | 0.019 | 97.71 | 589 | 0.015103 | 0.019 | 0.0151 | 1 |
| Q12846 | STX4 | kVRTIRQtIVkLGNk | T51 | 0.116 | 97.91 | 297 | 0.07923 | 0.116 | 0.0792 | 1 |
| P00568 | AK1 | RVDDNEEtIKKRLET | T145 | 0.052 | 96.16 | 194 | 0.035517 | 0.052 | 0.0355 | 0.9 |
| Q13733 | ATP1A4 | EAVETLGsTSTICSD | S377 | 0.043 | 94.76 | 1029 | 0.034181 | 0.043 | 0.0342 | 0.8571 |
| Q13733 | ATP1A4 | GIkVIMVtGDHPItA | T623 | 0.01 | 91.95 | 1029 | 0.00683 | 0.01 | 0.0068 | 0.8571 |
| Q9UIF8 | BAZ2B | RLQGEVAyyAPCGKK | Y772 | 0.004 | 97.11 | 2168 | 0.001755 | 0.004 | 0.0018 | 0.8462 |
| P05109 | S100A8 | LNsIIDVyHkYSLIK | Y16 | 0.038 | 98.83 | 93 | 0.01667 | 0.038 | 0.0167 | 0.8462 |
| P78371 | CCT2 | AEtARLTsFIGAIAI | S28 | 0.037 | 92.92 | 535 | 0.029412 | 0.037 | 0.0294 | 1 |
| P78371 | CCT2 | LGGSLADsyLDEGFL | S212 | 0.018 | 93.95 | 535 | 0.014308 | 0.018 | 0.0143 | 0.8182 |
| P78371 | CCT2 | AkILIANtGMDtDkI | T242 | 0.103 | 89.66 | 535 | 0.07035 | 0.169 | 0.1113 | 1 |
| P78371 | CCT2 | ILDEAERsLHDALCV | S389 | 0.006 | 91.28 | 535 | 0.004769 | 0.006 | 0.0048 | 1 |
| P78371 | CCT2 | TVYGGGCsEMLMAHA | S413 | 0.061 | 95.63 | 535 | 0.04849 | 0.061 | 0.0485 | 1 |
| P78371 | CCT2 | kEAVAMEsyAkALRM | S438 | 0.033 | 95.99 | 535 | 0.026232 | 0.033 | 0.0262 | 1 |
| P78371 | CCT2 | EAVAMEsyAkALRML | Y439 | 0.011 | 96.78 | 535 | 0.004826 | 0.011 | 0.0048 | 1 |
| P78371 | CCT2 | ADNAGyDsADLVAQL | S458 | 0.152 | 93.35 | 535 | 0.120827 | 0.152 | 0.1208 | 1 |
| P21941 | MATN1 | MSYMEKGtMtGAALK | T352 | 0.074 | 88.54 | 496 | 0.050543 | 0.074 | 0.0505 | 1 |

| | | | | | | | | | | |
|---|---|---|---|---|---|---|---|---|---|---|
| P21941 | MATN1 | YMEKGtMtGAALKYL | T354 | 0.028 | 93.63 | 496 | 0.019124 | 0.028 | 0.0191 | 1 |
| A8K4G0 | CD300LB | KDNQKDRtFtVTMEG | T86 | 0.145 | 98.63 | 201 | 0.099037 | 0.145 | 0.099 | 1 |
| O00443 | PIK3C2A | DGADPNPyVkTYLLP | Y1595 | 0.204 | 93.74 | 1686 | 0.089492 | 0.204 | 0.0895 | 1 |
| Q8N8U2 | CDYL2 | SSDRRKEstRIAEAI | S328 | 0.011 | 95.77 | 506 | 0.008744 | 0.011 | 0.0087 | 1 |
| P05181 | CYP2E1 | ELKNIPKsFtRLAQR | S56 | 0.033 | 98.41 | 493 | 0.026232 | 0.05 | 0.0397 | 0.875 |
| Q8WW24 | TEKT4 | MEALAAEtNLLLAQK | T116 | 0.116 | 98.66 | 435 | 0.07923 | 0.116 | 0.0792 | 1 |
| Q8IY17 | PNPLA6 | GPTLLLNsDIIRARL | S818 | 0.068 | 90.74 | 1375 | 0.054054 | 0.135 | 0.1074 | 1 |
| Q8IY17 | PNPLA6 | DAsLtPWtVRCLRQA | T864 | 0 | 89.92 | 1375 | 0 | 0 | 0 | 1 |
| Q8IY17 | PNPLA6 | PAKLHELyEkVFSRR | Y953 | 0.033 | 88.73 | 1375 | 0.014477 | 0.066 | 0.029 | 1 |
| Q8IY17 | PNPLA6 | SRLARVLtGNtIALV | T975 | 0.042 | 86.58 | 1375 | 0.028687 | 0.155 | 0.1139 | 1 |
| Q8IY17 | PNPLA6 | ARVLtGNtIALVLGG | T978 | 0.104 | 91.9 | 1375 | 0.071033 | 0.169 | 0.1154 | 1 |
| Q14181 | POLA2 | EGTRSSGsHLVFVPs | S418 | 0.031 | 95.28 | 598 | 0.024642 | 0.031 | 0.0246 | 1 |
| Q8WWY8 | LIPH | LDKVAAIsLMFSTGS | S393 | 0.086 | 96.72 | 451 | 0.068363 | 0.086 | 0.0684 | 1 |
| P18858 | LIG1 | ARLRMVEtLsNLLRS | T311 | 0.002 | 98.09 | 919 | 0.001366 | 0.002 | 0.0014 | 1 |
| P18858 | LIG1 | LRMVEtLsNLLRSVV | S313 | 0.001 | 98.27 | 919 | 0.000795 | 0.001 | 0.0008 | 1 |
| P18858 | LIG1 | QGMILKQtFCEVPDL | T506 | 0.053 | 96.57 | 919 | 0.0362 | 0.053 | 0.0362 | 0.8889 |
| P18858 | LIG1 | kPMLAHPtRGIsEVL | T548 | 0.029 | 90.25 | 919 | 0.019807 | 0.029 | 0.0198 | 1 |
| P18858 | LIG1 | IAEFLEQsVKDSCEG | S714 | 0 | 93.87 | 919 | 0 | 0 | 0 | 1 |
| P18858 | LIG1 | SAQVACLyRKQSQIQ | Y897 | 0.266 | 91.89 | 919 | 0.11669 | 0.266 | 0.1167 | 0.8 |
| Q8N1M1 | BEST3 | LLLKWRGsIyKLLYR | S27 | 0.114 | 94.91 | 668 | 0.09062 | 0.114 | 0.0906 | 0.8 |
| Q96S65 | CSRNP1 | VPSrGGCtLGMALRH | T110 | 0.004 | 92.85 | 589 | 0.002732 | 0.182 | 0.1243 | 1 |
| Q9UBC3 | DNMT3B | VGDKRDIsRFLECNP | S711 | 0.033 | 94.59 | 853 | 0.026232 | 0.033 | 0.0262 | 0.875 |
| Q9UBC3 | DNMT3B | FGFPVHytDVsNMGr | T816 | 0.001 | 96.59 | 853 | 0.000683 | 0.001 | 0.0007 | 0.875 |
| Q9UBC3 | DNMT3B | KLLGRSWsVPVIRHL | S835 | 0.101 | 97.04 | 853 | 0.080286 | 0.101 | 0.0803 | 1 |
| Q02750 | MAP2K1 | VGtRSYMsPERLQGT | S231 | 0.008 | 94.93 | 393 | 0.006359 | 0.008 | 0.0064 | 1 |
| Q13085 | ACACA | RTIQVENsHLILTGA | S1844 | 0.001 | 90.68 | 2346 | 0.000795 | 0.001 | 0.0008 | 1 |
| Q13085 | ACACA | NPRHMEMyADRESRG | Y2108 | 0.078 | 97.85 | 2346 | 0.034217 | 0.078 | 0.0342 | 1 |
| Q13085 | ACACA | MRRVDPVyIHLAERL | Y2144 | 0.132 | 91.39 | 2346 | 0.057907 | 0.263 | 0.1154 | 1 |
| O43781 | DYRK3 | KHHGRSstKVIDFGS | T351 | 0.011 | 97.54 | 588 | 0.007513 | 0.011 | 0.0075 | 1 |
| O43781 | DYRK3 | VVLVGGRsRRGKKRG | S469 | 0.154 | 95.98 | 588 | 0.122417 | 0.154 | 0.1224 | 1 |
| P26010 | ITGB7 | APQRVRVtLRPGEPQ | T131 | 0.167 | 96.59 | 798 | 0.114063 | 0.167 | 0.1141 | 1 |

| | | | | | | | | | | |
|---|---|---|---|---|---|---|---|---|---|---|
| P26010 | ITGB7 | RFLRAEGyPVDLYYL | Y150 | 0.221 | 82.32 | 798 | 0.09695 | 0.221 | 0.0969 | 1 |
| P11277 | SPTB | QELEKENyHDQkRIT | Y493 | 0.071 | 90.46 | 2137 | 0.031147 | 0.071 | 0.0311 | 0.8 |
| Q14397 | GCKR | LLLAAHKtVDQGIAA | T280 | 0.148 | 95.08 | 625 | 0.101086 | 0.148 | 0.1011 | 1 |
| Q14397 | GCKR | FERAHQVtYSQSPKI | T306 | 0.001 | 98.65 | 625 | 0.000683 | 0.001 | 0.0007 | 1 |
| Q14397 | GCKR | AtLMKSVsTSLEKKG | S321 | 0.111 | 98.63 | 625 | 0.088236 | 0.111 | 0.0882 | 0.8889 |
| Q2KHR3 | QSER1 | AMKETFksyMELLVs | S1551 | 0.127 | 94.98 | 1735 | 0.100954 | 0.127 | 0.101 | 1 |
| Q2KHR3 | QSER1 | MKETFksyMELLVsI | Y1552 | 0.001 | 96.17 | 1735 | 0.000439 | 0.313 | 0.1373 | 1 |
| P01579 | IFNG | DDQSIQKsVETIKED | S92 | 0.08 | 95.29 | 166 | 0.063593 | 0.08 | 0.0636 | 1 |
| P07320 | CRYGD | _MGKITLyEDrGFQG | Y7 | 0.235 | 98.8 | 174 | 0.103091 | 0.235 | 0.1031 | 0.9167 |
| Q969H0 | FBXW7 | WRIRAsNtKLVCAVG | T680 | 0.182 | 96.99 | 707 | 0.124308 | 0.182 | 0.1243 | 0.8571 |
| Q969H0 | FBXW7 | KLVCAVGsRNGtEEt | S688 | 0.115 | 97.02 | 707 | 0.091415 | 0.115 | 0.0914 | 0.8889 |
| O00445 | SYT5 | VGGLSDPyVKVHLLQ | Y278 | 0.097 | 98.56 | 386 | 0.042553 | 0.097 | 0.0426 | 1 |
| Q9NWW9 | PLAAT2 | GYAHWAIyVGDGYVV | Y27 | 0.31 | 98.69 | 162 | 0.135993 | 0.31 | 0.136 | 0.8333 |
| O94985 | CLSTN1 | KKETILCssDKTDMN | S407 | 0.063 | 92.31 | 981 | 0.05008 | 0.063 | 0.0501 | 1 |
| Q5VU97 | CACHD1 | HDKISVLtVADTVRT | T268 | 0.037 | 96.62 | 1274 | 0.025272 | 0.037 | 0.0253 | 1 |
| Q5VU97 | CACHD1 | VkssDsPtQHAVGFQ | T311 | 0.209 | 90.03 | 1274 | 0.14275 | 0.209 | 0.1427 | 1 |
| Q9BYT1 | SLC17A9 | AsAWGsItAVtPLLA | T104 | 0.024 | 97.64 | 436 | 0.016392 | 0.024 | 0.0164 | 1 |
| Q9BYT1 | SLC17A9 | WGsItAVtPLLAHLS | T107 | 0.06 | 95.68 | 436 | 0.040981 | 0.06 | 0.041 | 1 |
| P49917 | LIG4 | MHkDGDVykYFSRNG | Y288 | 0.017 | 98.61 | 911 | 0.007458 | 0.017 | 0.0075 | 1 |
| Q8WYK1 | CNTNAP5 | NKDGLLLsTELSEGS | S406 | 0 | 93.13 | 1306 | 0 | 0 | 0 | 1 |
| Q8WYK1 | CNTNAP5 | AERNLKEtsLQVDNL | T886 | 0.046 | 92.94 | 1306 | 0.031419 | 0.046 | 0.0314 | 1 |
| Q8WYK1 | CNTNAP5 | ERNLKEtsLQVDNLP | S887 | 0.059 | 95.12 | 1306 | 0.0469 | 0.118 | 0.0938 | 1 |
| Q8WYK1 | CNTNAP5 | NIALSFVttQAPSLL | T1046 | 0.035 | 94.58 | 1306 | 0.023905 | 0.035 | 0.0239 | 1 |
| Q8WYK1 | CNTNAP5 | VLLCKNGsLQVRYHL | S1073 | 0.045 | 93.43 | 1306 | 0.035771 | 0.045 | 0.0358 | 1 |
| P85298 | ARHGAP8 | DQyVENDyTIVYFHY | Y70 | 0.045 | 88.91 | 464 | 0.019741 | 0.045 | 0.0197 | 0.8571 |
| Q01973 | ROR1 | EDGFCQPyRGIACAR | Y173 | 0.155 | 89.18 | 937 | 0.067996 | 0.255 | 0.1119 | 0.8571 |
| P09668 | CTSH | GIMGEDtyPYQGKDG | Y205 | 0.088 | 98.7 | 335 | 0.038604 | 0.088 | 0.0386 | 1 |
| Q9UKQ2 | ADAM28 | RVFEMANyVNMLYKK | Y237 | 0 | 95.14 | 775 | 0 | 0.307 | 0.1347 | 1 |
| Q86UR5 | RIMS1 | DGRPRNPyVKMyFLP | Y781 | 0.05 | 96.73 | 1692 | 0.021934 | 0.05 | 0.0219 | 0.8182 |
| Q9P0N9 | TBC1D7 | TEkLCTFsQRFPLPS | S46 | 0.006 | 96.47 | 293 | 0.004769 | 0.006 | 0.0048 | 0.8889 |
| Q8NB16 | MLKL | FYQYDVKsEIYSFGI | S393 | 0.006 | 97.23 | 471 | 0.004769 | 0.006 | 0.0048 | 1 |

| | | | | | | | | | | |
|---|---|---|---|---|---|---|---|---|---|---|
| Q9Y5R8 | TRAPPC1 | GIKVVMNtDLGVGPI | T86 | 0 | 98.11 | 145 | 0 | 0 | 0 | 1 |
| P00367 | GLUD1 | sWEVIEGyRAQHSQH | Y135 | 0.284 | 98.61 | 558 | 0.124587 | 0.284 | 0.1246 | 1 |
| P00367 | GLUD1 | CKGGIRYstDVsVDE | S153 | 0.182 | 96.3 | 558 | 0.144674 | 0.182 | 0.1447 | 1 |
| P00367 | GLUD1 | EkITRRFtMELAkkG | T206 | 0.043 | 98.77 | 558 | 0.02937 | 0.043 | 0.0294 | 1 |
| P00367 | GLUD1 | VPAPDMstGEREMSW | T228 | 0.105 | 97.32 | 558 | 0.071716 | 0.105 | 0.0717 | 1 |
| P00367 | GLUD1 | AVGEsDGsIWNPDGI | S336 | 0.111 | 97.07 | 558 | 0.088236 | 0.149 | 0.1184 | 1 |
| P00367 | GLUD1 | AEGANGPttPEADkI | T409 | 0.004 | 98.15 | 558 | 0.002732 | 0.004 | 0.0027 | 1 |
| P00367 | GLUD1 | EGANGPttPEADkIF | T410 | 0 | 97.93 | 558 | 0 | 0 | 0 | 1 |
| P00367 | GLUD1 | VHSGLAytMERSARQ | T513 | 0.203 | 98.21 | 558 | 0.138652 | 0.203 | 0.1387 | 1 |
| P00367 | GLUD1 | LDLRTAAyVNAIEkV | Y539 | 0.014 | 98.74 | 558 | 0.006142 | 0.014 | 0.0061 | 1 |
| P00367 | GLUD1 | IEkVFKVyNEAGVTF | Y550 | 0.042 | 95.27 | 558 | 0.018425 | 0.212 | 0.093 | 1 |
| Q9Y4K4 | MAP4K5 | RRNPQQDyELVQRVG | Y20 | 0.169 | 95.11 | 846 | 0.074138 | 0.169 | 0.0741 | 1 |
| Q149M9 | NWD1 | VtGFSNGsIsLVssK | S1068 | 0.037 | 87.28 | 1564 | 0.029412 | 0.143 | 0.1137 | 0.8333 |
| Q149M9 | NWD1 | GFSNGsIsLVssKGD | S1070 | 0 | 82.59 | 1564 | 0 | 0.119 | 0.0946 | 0.8333 |
| Q149M9 | NWD1 | GALVASAsPQSSSFK | S1188 | 0.006 | 91.34 | 1564 | 0.004769 | 0.006 | 0.0048 | 1 |
| O60687 | SRPX2 | PKGGNyHssLGtRCE | S81 | 0.037 | 84.34 | 465 | 0.029412 | 0.037 | 0.0294 | 1 |
| O60687 | SRPX2 | KGGNyHssLGtRCEL | S82 | 0.034 | 87.96 | 465 | 0.027027 | 0.034 | 0.027 | 1 |
| O60687 | SRPX2 | NyHssLGtRCELSCD | T85 | 0 | 90.46 | 465 | 0 | 0 | 0 | 1 |
| P09429 | HMGB1 | AADDkQPyEkkAAKL | Y144 | 0.243 | 95.4 | 215 | 0.106601 | 0.243 | 0.1066 | 0.8182 |
| P46060 | RANGAP1 | KLkELNLsFCEIkRD | S301 | 0.126 | 98.25 | 587 | 0.100159 | 0.126 | 0.1002 | 1 |
| P12755 | SKI | RSTERCEtVLEGETI | T103 | 0.091 | 92.99 | 728 | 0.062154 | 0.091 | 0.0622 | 1 |
| P10911 | MCF2 | LIQTERVyVRELYTV | Y510 | 0.084 | 98.26 | 925 | 0.03685 | 0.084 | 0.0369 | 1 |
| P10911 | MCF2 | YGEKEEVyIVQAsNV | Y787 | 0.152 | 92.64 | 925 | 0.06668 | 0.152 | 0.0667 | 1 |
| O00178 | GTPBP1 | GNVDAGKstLLGVLt | S174 | 0.118 | 93.76 | 669 | 0.0938 | 0.118 | 0.0938 | 1 |
| O00178 | GTPBP1 | NVDAGKstLLGVLtH | T175 | 0.193 | 94.42 | 669 | 0.131822 | 0.193 | 0.1318 | 1 |
| O00178 | GTPBP1 | stLLGVLtHGELDNG | T181 | 0.037 | 92.51 | 669 | 0.025272 | 0.037 | 0.0253 | 0.9 |
| O00178 | GTPBP1 | HEkyLKTtVFGMTGH | T263 | 0.007 | 91.24 | 669 | 0.004781 | 0.007 | 0.0048 | 1 |
| O00178 | GTPBP1 | RKGMVMVsPRLNPQA | S481 | 0.142 | 96.7 | 669 | 0.112878 | 0.142 | 0.1129 | 1 |
| O00178 | GTPBP1 | QAMVHCGsIRQTAtI | S517 | 0.031 | 92.27 | 669 | 0.024642 | 0.031 | 0.0246 | 1 |
| Q9Y676 | MRPS18B | sLSSVPIsPYkDEPW | S49 | 0.115 | 78.67 | 258 | 0.091415 | 0.137 | 0.1089 | 1 |
| Q9Y676 | MRPS18B | KYLEsEEyQERYGSR | Y64 | 0.131 | 96.56 | 258 | 0.057468 | 0.131 | 0.0575 | 1 |

| | | | | | | | | | | |
|---|---|---|---|---|---|---|---|---|---|---|
| Q14703 | MBTPS1 | FkNVKERtNWTNERT | T235 | 0.073 | 92.45 | 1052 | 0.04986 | 0.216 | 0.1475 | 1 |
| Q14703 | MBTPS1 | LSGTSVAsPVVAGAV | S417 | 0 | 96.78 | 1052 | 0 | 0 | 0 | 1 |
| O43143 | DHX15 | LQLPVWEykDRFTDI | Y142 | 0.315 | 96.25 | 795 | 0.138186 | 0.315 | 0.1382 | 1 |
| O43143 | DHX15 | DILVRHQsFVLVGEt | S155 | 0.186 | 97.98 | 795 | 0.147854 | 0.186 | 0.1479 | 1 |
| O43143 | DHX15 | VLVGEtGsGkTTQIP | S164 | 0.056 | 86.07 | 795 | 0.044515 | 0.056 | 0.0445 | 1 |
| O43143 | DHX15 | NDPLLERyGVIILDE | Y254 | 0.121 | 98.1 | 795 | 0.053081 | 0.121 | 0.0531 | 1 |
| O43143 | DHX15 | LDEAHERtLATDILM | T266 | 0.21 | 91.2 | 795 | 0.143433 | 0.21 | 0.1434 | 0.8 |
| O43143 | DHX15 | VTAISkAsAQQRAGR | S465 | 0.157 | 91.46 | 795 | 0.124802 | 0.157 | 0.1248 | 1 |
| O43143 | DHX15 | RLYTEkAykTEMQDN | Y490 | 0.312 | 97.59 | 795 | 0.13687 | 0.312 | 0.1369 | 1 |
| O43143 | DHX15 | HLTLLNVyHAFKQNH | Y631 | 0.099 | 97.83 | 795 | 0.04343 | 0.099 | 0.0434 | 1 |
| O43143 | DHX15 | DNFINYRsLMSADNV | S653 | 0.044 | 87.22 | 795 | 0.034976 | 0.044 | 0.035 | 1 |
| O43143 | DHX15 | NIRKALVtGYFMQVA | T695 | 0.089 | 97.97 | 795 | 0.060788 | 0.089 | 0.0608 | 1 |
| O43143 | DHX15 | HLERTGHyLTVKDNQ | Y710 | 0.018 | 97.16 | 795 | 0.007896 | 0.018 | 0.0079 | 1 |
| O14727 | APAF1 | GSSFLTSsDDQtIRL | S901 | 0 | 89.8 | 1248 | 0 | 0 | 0 | 0.9 |
| O75131 | CPNE3 | NkDLFGksDPYLEFH | S159 | 0.003 | 97.07 | 537 | 0.002385 | 0.003 | 0.0024 | 1 |
| O75131 | CPNE3 | YkNSGVIsVKQCEIT | S268 | 0.052 | 98.4 | 537 | 0.041336 | 0.052 | 0.0413 | 1 |
| O75131 | CPNE3 | GDGGSLRsPLGEVAI | S476 | 0.031 | 94.47 | 537 | 0.024642 | 0.031 | 0.0246 | 1 |
| P27708 | CAD | QYRVSLRyVAPPSLR | Y2103 | 0.052 | 94.85 | 2225 | 0.022812 | 0.052 | 0.0228 | 1 |
| P27708 | CAD | RFGstQEyEACFGQF | Y2158 | 0.268 | 82.09 | 2225 | 0.117568 | 0.268 | 0.1176 | 1 |
| P27708 | CAD | DSDPRAAyFRQAENG | Y2203 | 0.033 | 91.63 | 2225 | 0.014477 | 0.061 | 0.0268 | 0.9286 |
| P27708 | CAD | RQAENGMyIRMALLA | Y2212 | 0.077 | 94.03 | 2225 | 0.033779 | 0.134 | 0.0588 | 1 |
| P27708 | CAD | IRMALLAtVLGRF__ | T2220 | 0.009 | 89.96 | 2225 | 0.006147 | 0.129 | 0.0841 | 1 |
| Q06278 | AOX1 | FGGKVLKtGIIAAVt | T814 | 0 | 98.51 | 1338 | 0 | 0 | 0 | 1 |
| Q06278 | AOX1 | tGIIAAVtAFAANKH | T821 | 0 | 98.87 | 1338 | 0 | 0 | 0 | 1 |
| Q06278 | AOX1 | ITGGRHPyLGKYKAG | Y851 | 0.04 | 98.49 | 1338 | 0.017547 | 0.08 | 0.0351 | 0.8 |
| Q06278 | AOX1 | WRECMAMsSYSLRKV | S984 | 0.004 | 98.31 | 1338 | 0.00318 | 0.004 | 0.0032 | 1 |
| Q06278 | AOX1 | LTLNSPLtPEKIRMA | T1307 | 0 | 98.65 | 1338 | 0 | 0.092 | 0.0624 | 1 |
| Q6ZN44 | UNC5A | NIVARRRsASAAVIV | S232 | 0.125 | 95.99 | 842 | 0.099364 | 0.125 | 0.0994 | 1 |
| Q8N3G9 | TMEM130 | KMEKGLSstIRVVGH | S91 | 0.001 | 94.73 | 435 | 0.000795 | 0.001 | 0.0008 | 1 |
| P32119 | PRDX2 | GCEVLGVsVDSQFTH | S76 | 0 | 98.87 | 198 | 0 | 0 | 0 | 1 |
| P32119 | PRDX2 | TRrLsEDyGVLktDE | Y115 | 0.005 | 98.66 | 198 | 0.002193 | 0.005 | 0.0022 | 1 |

| | | | | | | | | | | |
|---|---|---|---|---|---|---|---|---|---|---|
| P12111 | COL6A3 | VRVAVVQySDRTRPE | Y1072 | 0 | 96.69 | 3177 | 0 | 0 | 0 | 1 |
| Q9NVG8 | TBC1D13 | PLNPNPDsRWNtyFk | S108 | 0.034 | 86.66 | 400 | 0.027027 | 0.034 | 0.027 | 1 |
| P00751 | CFB | IYLVLDGsDSIGASN | S278 | 0.005 | 88.97 | 764 | 0.003975 | 0.005 | 0.004 | 0.8 |
| O43900 | PRICKLE3 | DKVPYVNsPGEkYRI | S122 | 0.171 | 89.95 | 615 | 0.13593 | 0.171 | 0.1359 | 1 |
| Q9H257 | CARD9 | FLESLELyYPQLYKK | Y86 | 0.23 | 92.21 | 536 | 0.100898 | 0.23 | 0.1009 | 1 |
| Q8WWA0 | ITLN1 | CTVGDRWssQQGsKA | S101 | 0.01 | 98.83 | 313 | 0.007949 | 0.01 | 0.0079 | 1 |
| Q8WWA0 | ITLN1 | TVGDRWssQQGsKAV | S102 | 0 | 98.82 | 313 | 0 | 0 | 0 | 1 |
| P21796 | VDAC1 | LARGLkLtFDssFsP | T98 | 0.098 | 96.78 | 283 | 0.066935 | 0.098 | 0.0669 | 1 |
| P21796 | VDAC1 | DGtEFGGsIyQkVNK | S193 | 0.109 | 97.22 | 283 | 0.086646 | 0.109 | 0.0866 | 1 |
| P21796 | VDAC1 | IDPDACFsAkVNNss | S234 | 0.046 | 97.53 | 283 | 0.036566 | 0.046 | 0.0366 | 1 |
| P21796 | VDAC1 | PGIKLTLsALLDGkN | S260 | 0.095 | 96.97 | 283 | 0.075517 | 0.095 | 0.0755 | 1 |
| P08865 | RPSA | EPRLLVVtDPRADHQ | T125 | 0.019 | 95.75 | 295 | 0.012977 | 0.038 | 0.026 | 1 |
| P08865 | RPSA | PTIALCNtDSPLRyV | T150 | 0.011 | 95.39 | 295 | 0.007513 | 0.011 | 0.0075 | 1 |
| P08865 | RPSA | CNNKGAHsVGLMWWM | S170 | 0 | 95.67 | 295 | 0 | 0 | 0 | 1 |
| Q6XPS3 | TPTE2 | CKGGKGRtGTMVCAL | T327 | 0 | 98.23 | 522 | 0 | 0 | 0 | 1 |
| Q9NYR8 | RDH8 | KRyQVVAtMRDLGKk | T38 | 0.003 | 98.85 | 311 | 0.002049 | 0.003 | 0.002 | 1 |
| Q9ULZ2 | STAP1 | FQERLKItALPLyFE | T22 | 0.165 | 84.98 | 295 | 0.112697 | 0.165 | 0.1127 | 1 |
| Q9ULZ2 | STAP1 | KItALPLyFEGFLLI | Y27 | 0.193 | 92.52 | 295 | 0.084666 | 0.193 | 0.0847 | 0.8333 |
| Q9ULZ2 | STAP1 | RGTTLFFyTDKKSII | Y57 | 0.001 | 92.29 | 295 | 0.000439 | 0.001 | 0.0004 | 1 |
| Q9NTI2 | ATP8A2 | EELGQVkyLFsDKTG | Y424 | 0.101 | 94.76 | 1188 | 0.044307 | 0.101 | 0.0443 | 1 |
| Q9NTI2 | ATP8A2 | IMNFKKCsIAGVtyG | S444 | 0.004 | 92.67 | 1188 | 0.00318 | 0.004 | 0.0032 | 1 |
| Q9NTI2 | ATP8A2 | CsIAGVtyGHFPELA | Y450 | 0.062 | 88.07 | 1188 | 0.027199 | 0.062 | 0.0272 | 1 |
| Q9NTI2 | ATP8A2 | NIEDRHPtAPCIQEF | T493 | 0.121 | 85.26 | 1188 | 0.082645 | 0.121 | 0.0826 | 1 |
| Q9NTI2 | ATP8A2 | FTARtPFsVIIEAMG | S549 | 0.183 | 90.6 | 1188 | 0.145469 | 0.183 | 0.1455 | 1 |
| Q9NTI2 | ATP8A2 | LQAGVPEtIATLLKA | T690 | 0.03 | 94.25 | 1188 | 0.02049 | 0.03 | 0.0205 | 0.8 |
| Q9UNH5 | CDC14A | RKKIVHytCFDQRKR | T85 | 0.003 | 96.74 | 594 | 0.002049 | 0.003 | 0.002 | 0.8333 |
| Q9UER7 | DAXX | RPAkLyVyINELCTV | Y126 | 0.181 | 95.74 | 740 | 0.079402 | 0.181 | 0.0794 | 0.9091 |
| Q7Z5J1 | HSD11B1L | LTSRALPsLTDSKGs | S152 | 0.188 | 98.41 | 315 | 0.149444 | 0.188 | 0.1494 | 0.8 |
| Q7Z5J1 | HSD11B1L | KGsLVVVsSLLGRVP | S164 | 0.095 | 93.41 | 315 | 0.075517 | 0.095 | 0.0755 | 1 |
| Q9H3T2 | SEMA6C | AGPHSNItVMFLGSN | T439 | 0.113 | 94.01 | 930 | 0.077181 | 0.113 | 0.0772 | 1 |
| Q05823 | RNASEL | HLADFDksIkWAGDP | S507 | 0.039 | 97.76 | 741 | 0.031002 | 0.039 | 0.031 | 1 |

| | | | | | | | | | | |
|---|---|---|---|---|---|---|---|---|---|---|
| Q9UKT7 | FBXL3 | WDAFIRHsPKVNLVM | S284 | 0.009 | 98.32 | 428 | 0.007154 | 0.009 | 0.0072 | 1 |
| Q6VY07 | PACS1 | LGSVDSKySSSFLDs | Y680 | 0 | 96.89 | 963 | 0 | 0 | 0 | 1 |
| Q9ULT0 | TTC7A | RGQyVMLsECLERAM | S400 | 0 | 92.9 | 858 | 0 | 0 | 0 | 1 |
| Q9UKZ9 | PCOLCE2 | TTITRDGsLHATVsI | S331 | 0.175 | 83.9 | 415 | 0.13911 | 0.175 | 0.1391 | 1 |
| Q6ISB3 | GRHL2 | LGLLyDyyKVPRDKR | Y71 | 0.032 | 83.91 | 625 | 0.014038 | 0.22 | 0.0965 | 1 |
| Q12955 | ANK3 | AAttsGytPLHLsAR | T534 | 0.013 | 97.72 | 4377 | 0.008879 | 0.013 | 0.0089 | 1 |
| Q12955 | ANK3 | GytPLHLsAREGHED | S539 | 0 | 97.21 | 4377 | 0 | 0 | 0 | 1 |
| Q12955 | ANK3 | AQTKMGytPLHVGCH | T732 | 0.014 | 97.04 | 4377 | 0.009562 | 0.014 | 0.0096 | 1 |
| Q9NQC7 | CYLD | yWEERIFyLLLQECs | Y22 | 0.315 | 92.16 | 956 | 0.138186 | 0.315 | 0.1382 | 1 |
| Q9NQC7 | CYLD | LLkVPkGsIGQYIQD | S45 | 0.057 | 91.64 | 956 | 0.04531 | 0.057 | 0.0453 | 1 |
| Q9H488 | POFUT1 | FTNLHVSyQKYFkLE | Y87 | 0.026 | 98.8 | 388 | 0.011406 | 0.026 | 0.0114 | 1 |
| Q9H488 | POFUT1 | QAYHRVIsLEDFMEK | S104 | 0.067 | 98.64 | 388 | 0.053259 | 0.108 | 0.0951 | 1 |
| Q96AE7 | TTC17 | QTsWVLSsMAALyWR | S1019 | 0.091 | 94.62 | 1141 | 0.072337 | 0.142 | 0.1364 | 0.8571 |
| O94855 | SEC24D | LDPDSIPsPIQVIEN | S266 | 0.153 | 93.4 | 1032 | 0.121622 | 0.153 | 0.1216 | 0.8 |
| O94855 | SEC24D | ASrGGQVyATNTRGQ | Y283 | 0.224 | 98.3 | 1032 | 0.098266 | 0.224 | 0.0983 | 1 |
| O94855 | SEC24D | QLADLYKsCETDALI | S779 | 0.142 | 98.5 | 1032 | 0.112878 | 0.142 | 0.1129 | 1 |
| P00450 | CP | KKLVYREytDAsFtN | Y428 | 0.092 | 98.34 | 1065 | 0.040359 | 0.183 | 0.0803 | 1 |
| P33032 | MC5R | tIFYALRyHHIMtAR | Y150 | 0.227 | 91.61 | 325 | 0.099582 | 0.227 | 0.0996 | 1 |
| P43088 | PTGFR | PILGHRDyKIQASRT | Y177 | 0.009 | 97.27 | 359 | 0.003948 | 0.009 | 0.0039 | 0.8824 |
| Q9Y496 | KIF3A | SLGGNSKtMMCANIG | T316 | 0.055 | 96.49 | 699 | 0.037566 | 0.055 | 0.0376 | 0.8 |
| Q99623 | PHB2 | WFQyPIIyDIRARPR | Y81 | 0.062 | 91.02 | 299 | 0.027199 | 0.062 | 0.0272 | 1 |
| Q99623 | PHB2 | ARPRkIssPTGSkDL | S92 | 0.062 | 91.39 | 299 | 0.049285 | 0.062 | 0.0493 | 1 |
| Q99623 | PHB2 | DLQMVNIsLRVLSRP | S105 | 0.07 | 94.95 | 299 | 0.055644 | 0.07 | 0.0556 | 1 |
| Q99623 | PHB2 | yQrLGLDyEERVLPs | Y128 | 0.163 | 87.07 | 299 | 0.071506 | 0.283 | 0.1088 | 1 |
| Q99623 | PHB2 | TERAKDFsLILDDVA | S176 | 0.043 | 92.06 | 299 | 0.034181 | 0.043 | 0.0342 | 1 |
| Q9UDY2 | TJP2 | EAIyGPNtKMVRFKK | T510 | 0.181 | 86.49 | 1190 | 0.123625 | 0.181 | 0.1236 | 1 |
| Q08AH1 | ACSM1 | FREMGDLtRRVANVF | T95 | 0 | 98.82 | 577 | 0 | 0 | 0 | 1 |
| Q08AH1 | ACSM1 | SQCPSLKtKLLVsDH | T184 | 0.152 | 98.26 | 577 | 0.103818 | 0.152 | 0.1038 | 1 |
| Q08AH1 | ACSM1 | LKtKLLVsDHSREGW | S189 | 0.12 | 96.82 | 577 | 0.09539 | 0.12 | 0.0954 | 1 |
| O43639 | NCK2 | VQLVDNVyCIGQRRF | Y342 | 0.162 | 92.78 | 380 | 0.071067 | 0.162 | 0.0711 | 0.8333 |
| Q9UQN3 | CHMP2B | KTRTFAVsSkVTSMs | S79 | 0.169 | 93.93 | 213 | 0.134341 | 0.169 | 0.1343 | 1 |

| | | | | | | | | | | |
|---|---|---|---|---|---|---|---|---|---|---|
| Q9UQN3 | CHMP2B | VTSMsTQtkVMNSQM | T89 | 0.174 | 91.65 | 213 | 0.118844 | 0.174 | 0.1188 | 1 |
| Q9Y5E5 | PCDHB4 | ASDEIKQtFSINEVT | T290 | 0.01 | 94.72 | 795 | 0.00683 | 0.01 | 0.0068 | 1 |
| Q8IZJ1 | UNC5B | LERHsLAstELtCKI | S803 | 0.14 | 89.75 | 945 | 0.111288 | 0.14 | 0.1113 | 1 |
| O75369 | FLNB | kAFVGQKssFLVDCS | S2531 | 0.122 | 85.97 | 2602 | 0.09698 | 0.157 | 0.1345 | 1 |
| O75369 | FLNB | VVkERGDyVLAVKWG | Y2581 | 0.004 | 85.31 | 2602 | 0.001755 | 0.004 | 0.0018 | 1 |
| Q8NHS4 | CLHC1 | CLCKQGQtHRVMEYI | T446 | 0.133 | 83.26 | 586 | 0.090841 | 0.133 | 0.0908 | 1 |
| O43280 | TREH | LSHPERFsLIYSEHP | S154 | 0.014 | 98.32 | 583 | 0.011129 | 0.014 | 0.0111 | 1 |
| O43280 | TREH | KNYLLNRyYVPyGGP | Y278 | 0 | 98.83 | 583 | 0 | 0 | 0 | 1 |
| O43280 | TREH | NDsQAtKyRILRSQR | Y376 | 0.076 | 98.86 | 583 | 0.03334 | 0.076 | 0.0333 | 1 |
| Q5TEJ8 | THEMIS2 | VMHLGIRsARCVLGM | S143 | 0.058 | 91.05 | 643 | 0.046105 | 0.058 | 0.0461 | 1 |
| Q8NI35 | PATJ | IKQVLEDsPAGKTNA | S1111 | 0.07 | 87.47 | 1801 | 0.055644 | 0.105 | 0.0835 | 1 |
| Q96CJ1 | EAF2 | RVLkLGEsFEkQPRC | S22 | 0.087 | 95.94 | 260 | 0.069158 | 0.153 | 0.1216 | 1 |
| Q00653 | NFKB2 | kGRKTyPtVKICNYE | T79 | 0.181 | 96.28 | 900 | 0.123625 | 0.181 | 0.1236 | 1 |
| Q00653 | NFKB2 | kIEVDLVtHsDPPRA | T97 | 0.065 | 94.56 | 900 | 0.044396 | 0.13 | 0.0888 | 1 |
| Q00653 | NFKB2 | PPRAHAHsLVGkQCs | S108 | 0.166 | 91.75 | 900 | 0.131956 | 0.166 | 0.132 | 1 |
| Q00653 | NFKB2 | SDSkQFTyyPLVEDk | Y325 | 0 | 95.58 | 900 | 0 | 0 | 0 | 1 |
| Q13247 | SRSF6 | RLIVENLssRCsWQD | S118 | 0.105 | 79.95 | 344 | 0.083466 | 0.105 | 0.0835 | 1 |
| Q86Y37 | CACUL1 | QASPPDLyIERFNIA | Y206 | 0.198 | 96.67 | 369 | 0.08686 | 0.198 | 0.0869 | 1 |
| P43116 | PTGER2 | IRHGRtAyLQLYATL | Y196 | 0.038 | 93.2 | 358 | 0.01667 | 0.038 | 0.0167 | 0.9231 |
| Q8IYR6 | TMEFF1 | CGSNGDtyQNECFLR | Y118 | 0.265 | 95.26 | 380 | 0.116252 | 0.265 | 0.1163 | 0.8462 |
| O95484 | CLDN9 | GWLGtLVsCALPLWK | S24 | 0.046 | 93.95 | 217 | 0.036566 | 0.046 | 0.0366 | 0.8182 |
| Q8WZ60 | KLHL6 | LLQEARMyHLsGNEI | Y297 | 0.152 | 88.12 | 621 | 0.06668 | 0.152 | 0.0667 | 0.8333 |
| P22234 | PAICS | WVCRRIAtGsFLkRN | T105 | 0.039 | 96.81 | 425 | 0.026638 | 0.039 | 0.0266 | 1 |
| P22234 | PAICS | IFEILEKsWLPQNCT | S179 | 0.01 | 97.81 | 425 | 0.007949 | 0.01 | 0.0079 | 1 |
| P22234 | PAICS | QQKDkQsyRDLkEVt | Y231 | 0 | 94.28 | 425 | 0 | 0 | 0 | 1 |
| P22234 | PAICS | AHkGPDEtLrIkAEY | T309 | 0.003 | 98.49 | 425 | 0.002049 | 0.003 | 0.002 | 1 |
| P22234 | PAICS | LsNHLVWsKLRAsIL | S400 | 0.01 | 98.49 | 425 | 0.007949 | 0.01 | 0.0079 | 1 |
| Q8WUJ3 | CEMIP | VAVNDEGsRNLDDMA | S241 | 0.134 | 93.68 | 1361 | 0.106519 | 0.134 | 0.1065 | 1 |
| Q02153 | GUCY1B1 | IGQRMPRyCLFGNTV | Y540 | 0.144 | 88.77 | 619 | 0.063171 | 0.144 | 0.0632 | 1 |
| Q9UGP8 | SEC63 | EYQEYNPyEVLNLDP | Y106 | 0.129 | 88.96 | 760 | 0.05659 | 0.129 | 0.0566 | 1 |
| Q9UGP8 | SEC63 | AALtDEEsRKNWEEF | S157 | 0.144 | 86.42 | 760 | 0.114468 | 0.144 | 0.1145 | 1 |

| | | | | | | | | | | |
|---|---|---|---|---|---|---|---|---|---|---|
| Q9UGP8 | SEC63 | kARVLLLsHLARMkI | S307 | 0 | 95.88 | 760 | 0 | 0 | 0 | 1 |
| Q5VWK5 | IL23R | ICGKDIssGYPPDIP | S121 | 0 | 96.99 | 629 | 0 | 0 | 0 | 1 |
| Q8NB46 | ANKRD52 | kERKRKWtPLHAAAA | T654 | 0.041 | 95.35 | 1076 | 0.028004 | 0.041 | 0.028 | 1 |
| Q8NB46 | ANKRD52 | SGERADItDVMDAYG | T681 | 0.055 | 91.4 | 1076 | 0.037566 | 0.055 | 0.0376 | 1 |
| Q16270 | IGFBP7 | GCQLRAAsQRAEsRG | S137 | 0.121 | 95.67 | 282 | 0.096185 | 0.121 | 0.0962 | 1 |
| Q07021 | C1QBP | EQEPELtstPNFVVE | S164 | 0.013 | 94.94 | 282 | 0.010334 | 0.013 | 0.0103 | 1 |
| Q07021 | C1QBP | IFsIREVsFQstGEs | S210 | 0.023 | 96.18 | 282 | 0.018283 | 0.023 | 0.0183 | 1 |
| Q9NQW7 | XPNPEP1 | AQYkDGttDVTRTMH | T425 | 0.008 | 98.95 | 623 | 0.005464 | 0.008 | 0.0055 | 1 |
| Q9Y487 | ATP6V0A2 | EsGAtIPsFMNIIPT | S355 | 0.117 | 81.61 | 856 | 0.093005 | 0.117 | 0.093 | 1 |
| P07910 | HNRNPC | YGkIVGCsVHkGFAF | S47 | 0.104 | 95.23 | 306 | 0.082671 | 0.104 | 0.0827 | 0.875 |
| P07910 | HNRNPC | kGFAFVQyVNErNAR | Y57 | 0.039 | 96.33 | 306 | 0.017109 | 0.039 | 0.0171 | 1 |
| Q96FW1 | OTUB1 | YAEDDNIyQQkIkDL | Y68 | 0.232 | 93.48 | 271 | 0.101775 | 0.232 | 0.1018 | 1 |
| P07384 | CAPN1 | ANAsRARsEQFINLR | S474 | 0.044 | 93.22 | 714 | 0.034976 | 0.044 | 0.035 | 1 |
| P07384 | CAPN1 | FRLPPGEyVVVPSTF | Y494 | 0.038 | 96.1 | 714 | 0.01667 | 0.038 | 0.0167 | 1 |
| Q8N5I2 | ARRDC1 | QVRAAIHtPRFsKDH | T117 | 0.091 | 91.75 | 433 | 0.062154 | 0.091 | 0.0622 | 1 |
| P22033 | MMUT | PKEkLTGtIQNDILK | T216 | 0.2 | 93.32 | 750 | 0.136603 | 0.2 | 0.1366 | 1 |
| P22033 | MMUT | YIFPPEPsMKIIADI | S238 | 0.059 | 96.74 | 750 | 0.0469 | 0.059 | 0.0469 | 1 |
| P22033 | MMUT | IIADIFEyTAKHMPK | Y248 | 0.17 | 95.85 | 750 | 0.074577 | 0.17 | 0.0746 | 1 |
| P22033 | MMUT | TLADGLEySRTGLQA | Y287 | 0 | 97.7 | 750 | 0 | 0 | 0 | 1 |
| Q9Y5G7 | PCDHGA6 | DREGEAVyRLVLTAM | Y208 | 0.109 | 92.02 | 932 | 0.047817 | 0.109 | 0.0478 | 1 |
| Q9Y3T9 | NOC2L | kLTREEGtPLtLyYs | T613 | 0.027 | 93.53 | 749 | 0.018441 | 0.027 | 0.0184 | 0.9 |
| Q9H2E6 | SEMA6A | CKNDMGGsQRVLEKQ | S265 | 0.104 | 97.23 | 1030 | 0.082671 | 0.104 | 0.0827 | 1 |
| Q96RP7 | GAL3ST4 | QRLVFLKtHKsGSSS | T74 | 0.021 | 98.52 | 486 | 0.014343 | 0.021 | 0.0143 | 0.8889 |
| Q96HD9 | ACY3 | VLANPAAtSGCRRYV | T58 | 0.108 | 98.48 | 319 | 0.073765 | 0.108 | 0.0738 | 1 |
| Q9BWT1 | CDCA7 | GFGNVHAyLkSLkQE | Y359 | 0.264 | 94.99 | 371 | 0.115813 | 0.264 | 0.1158 | 1 |
| Q1L5Z9 | LONRF2 | VRtFPDGsSVVDAIG | S606 | 0.094 | 86.32 | 754 | 0.074722 | 0.094 | 0.0747 | 1 |
| P13497 | BMP1 | RMFLRFYsDNSVQRK | S843 | 0.025 | 92.41 | 986 | 0.019873 | 0.025 | 0.0199 | 1 |
| Q92974 | ARHGEF2 | FPTLDkPsVVsLQNL | S519 | 0.03 | 92.89 | 986 | 0.023847 | 0.184 | 0.1463 | 1 |
| Q8N7C7 | RNF148 | GANGVIIyNYQGTGS | Y130 | 0.023 | 98.12 | 305 | 0.01009 | 0.023 | 0.0101 | 1 |
| P61081 | UBE2M | NELNLPktCDIsFSD | T46 | 0.167 | 98 | 183 | 0.114063 | 0.167 | 0.1141 | 1 |
| P61081 | UBE2M | SFkVGQGyPHDPPkV | Y86 | 0.043 | 98.22 | 183 | 0.018863 | 0.043 | 0.0189 | 1 |

| | | | | | | | | | | |
|---|---|---|---|---|---|---|---|---|---|---|
| Q9P2R3 | ANKFY1 | AFNLRGQsPLHILGQ | S1008 | 0.041 | 96.17 | 1169 | 0.032592 | 0.082 | 0.0652 | 0.8889 |
| Q9P2R3 | ANKFY1 | RLLCHKCsTKEIPII | S1138 | 0 | 91.03 | 1169 | 0 | 0 | 0 | 1 |
| Q2TBF2 | WSCD2 | sYFIVYQtQVQDNRC | T323 | 0.099 | 89.95 | 565 | 0.067618 | 0.099 | 0.0676 | 1 |
| P04632 | CAPNS1 | IkRWQAIykQFDTDR | Y178 | 0.008 | 97.79 | 268 | 0.003509 | 0.008 | 0.0035 | 1 |
| O43572 | AKAP10 | AMILYDKyFSLQAtH | Y442 | 0.101 | 95.17 | 662 | 0.044307 | 0.101 | 0.0443 | 1 |
| Q15645 | TRIP13 | kkEDINLsVRKLLNR | S41 | 0.097 | 96.48 | 432 | 0.077107 | 0.097 | 0.0771 | 1 |
| P51617 | IRAK1 | LCEISRGtHNFSEEL | T209 | 0 | 96.55 | 712 | 0 | 0 | 0 | 1 |
| P17252 | PRKCA | PNGLSDPyVkLkLIP | Y195 | 0.1 | 98.61 | 672 | 0.043869 | 0.1 | 0.0439 | 1 |
| P17252 | PRKCA | sDKDRRLsVEIWDWD | S241 | 0.037 | 98.15 | 672 | 0.029412 | 0.037 | 0.0294 | 1 |
| P51571 | SSR4 | VRFFDEEsySLLRKA | S111 | 0.152 | 88.96 | 173 | 0.120827 | 0.152 | 0.1208 | 1 |
| P48448 | ALDH3B2 | tAATKHLtPVtLELG | T124 | 0.159 | 98.76 | 385 | 0.108599 | 0.159 | 0.1086 | 1 |
| P48448 | ALDH3B2 | TKHLtPVtLELGGkN | T127 | 0.003 | 98.88 | 385 | 0.002049 | 0.003 | 0.002 | 1 |
| Q96BP3 | PPWD1 | YPTSVCFsPDGKkIA | S289 | 0.048 | 97.39 | 646 | 0.038156 | 0.048 | 0.0382 | 1 |
| Q9Y597 | KCTD3 | RFSTSRQtLMWIPDS | T35 | 0.205 | 94.35 | 815 | 0.140018 | 0.205 | 0.14 | 0.8333 |
| O75529 | TAF5L | RTYPLRIyAGHLADV | Y422 | 0.009 | 97.09 | 589 | 0.003948 | 0.009 | 0.0039 | 0.8 |
| O75529 | TAF5L | DktVRLWsAQQGNsV | S454 | 0.167 | 95.92 | 589 | 0.132751 | 0.167 | 0.1328 | 1 |
| Q8TAS1 | UHMK1 | kYIQTDGyRAPEAEL | Y197 | 0.245 | 97.89 | 419 | 0.107478 | 0.245 | 0.1075 | 1 |
| Q7Z7G8 | VPS13B | LLQGPSDtKDLHSTK | T901 | 0.085 | 84.56 | 4022 | 0.058056 | 0.085 | 0.0581 | 0.8889 |
| Q6NUT2 | DPY19L2 | VFAGAMPtMAsIKLS | T643 | 0.065 | 94.38 | 758 | 0.044396 | 0.065 | 0.0444 | 1 |
| Q6NUT2 | DPY19L2 | GAMPtMAsIKLSTLH | S646 | 0.035 | 94.85 | 758 | 0.027822 | 0.035 | 0.0278 | 1 |
| Q6NUT2 | DPY19L2 | KIVYSTYsRKSAKEV | S678 | 0.021 | 94.07 | 758 | 0.016693 | 0.021 | 0.0167 | 1 |
| P07205 | PGK2 | NNQRIKAsIPSIKYC | S43 | 0.036 | 97.16 | 417 | 0.028617 | 0.036 | 0.0286 | 1 |
| P07205 | PGK2 | AKAVVLMsHLGRPDG | S62 | 0 | 98.87 | 417 | 0 | 0 | 0 | 1 |
| P07205 | PGK2 | KIEAFRAsLskLGDV | S153 | 0.074 | 98.84 | 417 | 0.058824 | 0.069 | 0.083 | 1 |
| P07205 | PGK2 | LskLGDVyVNDAFGt | Y161 | 0.045 | 98.91 | 417 | 0.019741 | 0.09 | 0.0395 | 1 |
| P07205 | PGK2 | yVNDAFGtAHRAHSS | T168 | 0.036 | 96.46 | 417 | 0.024588 | 0.036 | 0.0246 | 1 |
| P07205 | PGK2 | IIGGGMAyTFLKVLN | Y242 | 0.079 | 98.54 | 417 | 0.034656 | 0.158 | 0.0693 | 1 |
| P26641 | EEF1G | ALIAAQysGAQVRVL | S25 | 0.169 | 98.44 | 437 | 0.134341 | 0.169 | 0.1343 | 0.8571 |
| Q8N9W5 | DNAAF3 | RHYLGSRyDARrGVS | Y202 | 0.285 | 95.11 | 541 | 0.125025 | 0.285 | 0.125 | 1 |
| P14616 | INSRR | AARNCMVsQDFtVKI | S1124 | 0.114 | 91.45 | 1297 | 0.09062 | 0.114 | 0.0906 | 1 |
| Q9H3Z4 | DNAJC5 | TSDDIKKsYRKLALk | S34 | 0.138 | 94.88 | 198 | 0.109698 | 0.138 | 0.1097 | 0.8571 |

| | | | | | | | | | | |
|---|---|---|---|---|---|---|---|---|---|---|
| Q6P1M3 | LLGL2 | DLFQFNktVEHGFPH | T26 | 0.035 | 96.08 | 1020 | 0.023905 | 0.172 | 0.1119 | 0.8889 |
| Q6P1M3 | LLGL2 | TVRFWDAsGVCLRLL | S453 | 0.161 | 98.11 | 1020 | 0.127981 | 0.161 | 0.128 | 1 |
| Q6P1M3 | LLGL2 | WAGtNGGtIYAFSLR | T741 | 0.183 | 96.51 | 1020 | 0.124991 | 0.183 | 0.125 | 1 |
| Q6UY14 | ADAMTSL4 | sLsAEGPtTQPVDVY | T576 | 0.053 | 88.82 | 1074 | 0.0362 | 0.053 | 0.0362 | 1 |
| Q9NRF9 | POLE3 | kEARsAIsRAASVFV | S38 | 0.093 | 92.68 | 147 | 0.073927 | 0.093 | 0.0739 | 1 |
| P11388 | TOP2A | NDAGGRNsTECTLIL | S452 | 0.015 | 92.22 | 1531 | 0.011924 | 0.015 | 0.0119 | 1 |
| P11388 | TOP2A | GLQykkNyEDEDSLk | Y522 | 0 | 90.81 | 1531 | 0 | 0 | 0 | 1 |
| P11388 | TOP2A | NKELILFsNsDNERs | S707 | 0.055 | 90.09 | 1531 | 0.04372 | 0.11 | 0.0874 | 0.875 |
| P11388 | TOP2A | DNERsIPsMVDGLkP | S717 | 0.095 | 95.59 | 1531 | 0.075517 | 0.095 | 0.0755 | 1 |
| P30679 | GNA15 | KRFILDMyTRMYTGC | Y316 | 0.022 | 98.64 | 374 | 0.009651 | 0.022 | 0.0097 | 1 |
| P07585 | DCN | KASYSGVsLFsNPVQ | S328 | 0.006 | 97.63 | 359 | 0.004769 | 0.006 | 0.0048 | 1 |
| O14776 | TCERG1 | SDVsWSDtRRtLRkD | T921 | 0.102 | 94.28 | 1098 | 0.069667 | 0.103 | 0.095 | 1 |
| Q8NFM7 | IL17RD | QNVSPGDyIIELVDD | Y268 | 0.153 | 92.89 | 739 | 0.067119 | 0.153 | 0.0671 | 1 |
| P57735 | RAB25 | ALLVFDLtKHQTYAV | T95 | 0.191 | 98.19 | 213 | 0.130456 | 0.191 | 0.1305 | 1 |
| Q9NYY8 | FASTKD2 | RISSRkLsEECNSLs | S160 | 0.067 | 85.72 | 710 | 0.053259 | 0.067 | 0.0533 | 1 |
| Q9NYY8 | FASTKD2 | sKAPTFPsSNYFTAM | S181 | 0.008 | 92.99 | 710 | 0.006359 | 0.008 | 0.0064 | 1 |
| Q9NYY8 | FASTKD2 | LFKGLADyVAATFDI | Y406 | 0.322 | 97.67 | 710 | 0.141257 | 0.322 | 0.1413 | 1 |
| Q8WZ74 | CTTNBP2 | LLDLFISsACLIPVK | S1178 | 0.152 | 81.57 | 1663 | 0.120827 | 0.152 | 0.1208 | 1 |
| Q92830 | KAT2A | EEIYGANsPIWESGF | S372 | 0.092 | 91.22 | 837 | 0.073132 | 0.092 | 0.0731 | 1 |
| Q9H4A4 | RNPEP | PGVDPDDtyNEtPyE | T408 | 0.203 | 93.04 | 650 | 0.138652 | 0.203 | 0.1387 | 1 |
| Q9H4A4 | RNPEP | PDDtyNEtPyEKGFC | T412 | 0 | 97.19 | 650 | 0 | 0 | 0 | 1 |
| Q9H4A4 | RNPEP | DtyNEtPyEKGFCFV | Y414 | 0.313 | 97.91 | 650 | 0.137309 | 0.313 | 0.1373 | 0.9 |
| Q9H4A4 | RNPEP | NQGkQKYtLPLyHAM | T604 | 0.017 | 96.07 | 650 | 0.011611 | 0.017 | 0.0116 | 1 |
| Q9H4A4 | RNPEP | QKYtLPLyHAMMGGS | Y608 | 0.019 | 97.97 | 650 | 0.008335 | 0.019 | 0.0083 | 1 |
| P50454 | SERPINH1 | AATLAERsAGLAFSL | S47 | 0.007 | 96.39 | 418 | 0.005564 | 0.007 | 0.0056 | 1 |
| P50454 | SERPINH1 | ADDFVRsskQHyNCE | S150 | 0 | 98.42 | 418 | 0 | 0 | 0 | 1 |
| P50454 | SERPINH1 | VRsskQHyNCEHskI | Y154 | 0.089 | 97.79 | 418 | 0.039043 | 0.089 | 0.039 | 1 |
| P50454 | SERPINH1 | NEWAAQTtDGkLPEV | T181 | 0.006 | 97.24 | 418 | 0.004098 | 0.006 | 0.0041 | 1 |
| P50454 | SERPINH1 | HRTGLYNyyDDEkEk | Y245 | 0.08 | 98.82 | 418 | 0.035095 | 0.153 | 0.0646 | 1 |
| Q9NVH2 | INTS7 | LGDLMELyKVIGRSA | Y475 | 0.001 | 91.76 | 962 | 0.000439 | 0.001 | 0.0004 | 1 |
| Q96QU1 | PCDH15 | PVFSKRIyKGMVAPD | Y931 | 0.097 | 88.73 | 1955 | 0.042553 | 0.161 | 0.0706 | 1 |

| | | | | | | | | | | |
|---|---|---|---|---|---|---|---|---|---|---|
| Q96QU1 | PCDH15 | PVFQKKFyIGGVSED | Y1149 | 0.033 | 87.68 | 1955 | 0.014477 | 0.299 | 0.1312 | 1 |
| Q96QU1 | PCDH15 | GNYSVMAyRLIIPPI | Y1181 | 0.136 | 93.58 | 1955 | 0.059661 | 0.175 | 0.0768 | 1 |
| Q16696 | CYP2A13 | AKQLRRFsIATLRGF | S131 | 0 | 98.6 | 494 | 0 | 0 | 0 | 1 |
| Q3MIT2 | PUS10 | VSFPPQLsVREHAAW | S153 | 0.09 | 96.99 | 529 | 0.071542 | 0.09 | 0.0715 | 1 |
| Q86US8 | SMG6 | APKNGRPyNQLALLA | Y769 | 0.016 | 97.84 | 1419 | 0.007019 | 0.016 | 0.007 | 0.8571 |
| Q86US8 | SMG6 | KLDAVYyyMRSLAAs | Y789 | 0.082 | 98.54 | 1419 | 0.035972 | 0.082 | 0.036 | 1 |
| Q9H4B4 | PLK3 | FEDADNIyIFLELCS | Y136 | 0.169 | 96.97 | 646 | 0.074138 | 0.169 | 0.0741 | 1 |
| P08708 | RPS17 | LRNkIAGyVtHLMkR | Y53 | 0.255 | 96.73 | 135 | 0.111865 | 0.255 | 0.1119 | 1 |
| P08708 | RPS17 | NkIAGyVtHLMkRIQ | T55 | 0.146 | 96.96 | 135 | 0.09972 | 0.146 | 0.0997 | 1 |
| Q07001 | CHRND | HLFQEKGyNKELRPV | Y38 | 0.302 | 93.58 | 517 | 0.132483 | 0.302 | 0.1325 | 1 |
| O95475 | SIX6 | ALWLEAHyQEAEKLR | Y96 | 0.239 | 97.21 | 246 | 0.104846 | 0.239 | 0.1048 | 1 |
| Q6ZXV5 | TMTC3 | LEEADQLyRQAISMR | Y552 | 0 | 95.98 | 915 | 0 | 0 | 0 | 1 |
| Q32P44 | EML3 | GGHGsHVtsVRFTHD | T843 | 0.062 | 98.45 | 896 | 0.042347 | 0.062 | 0.0423 | 1 |
| Q32P44 | EML3 | GHGsHVtsVRFTHDD | S844 | 0.065 | 98.54 | 896 | 0.051669 | 0.065 | 0.0517 | 1 |
| O75146 | HIP1R | GLPLPSSsILsWKFC | S80 | 0.132 | 96.18 | 1068 | 0.104929 | 0.132 | 0.1049 | 1 |
| O75146 | HIP1R | LPSSsILsWKFCHVL | S83 | 0 | 97.07 | 1068 | 0 | 0 | 0 | 1 |
| Q9H2U9 | ADAM7 | VNFVNMIyKTLNIHV | Y237 | 0.009 | 96.11 | 754 | 0.003948 | 0.075 | 0.0329 | 1 |
| Q9H2U9 | ADAM7 | ETTLLRFsFWQEKIL | S272 | 0 | 91.96 | 754 | 0 | 0 | 0 | 1 |
| Q9H4G4 | GLIPR2 | stRILkHsPESSRGQ | S55 | 0.001 | 96.23 | 154 | 0.000795 | 0.001 | 0.0008 | 0.8333 |
| P40763 | STAT3 | TEELHLItFETEVyH | T440 | 0.12 | 97.08 | 770 | 0.081962 | 0.12 | 0.082 | 1 |
| P40763 | STAT3 | LLGPGVNysGCQITW | Y539 | 0.214 | 90.86 | 770 | 0.093879 | 0.214 | 0.0939 | 1 |
| P40763 | STAT3 | LSTkPPGtFLLRFSE | T605 | 0.006 | 95.75 | 770 | 0.004098 | 0.006 | 0.0041 | 1 |
| P40763 | STAT3 | kEGGVTFtWVEkDIS | T622 | 0.005 | 94.9 | 770 | 0.003415 | 0.005 | 0.0034 | 0.875 |
| P40763 | STAT3 | IQsVEPytkQQLNNM | T641 | 0.212 | 93.65 | 770 | 0.144799 | 0.212 | 0.1448 | 1 |
| Q9ULH4 | LRFN2 | MVNLHQLsLDHNLLD | S179 | 0.13 | 98.16 | 789 | 0.103339 | 0.13 | 0.1033 | 1 |
| P05771 | PRKCB | RkGtDELyAVKILKK | Y368 | 0.034 | 93.96 | 671 | 0.014915 | 0.15 | 0.0336 | 1 |
| P05771 | PRKCB | NIWDGVttktFCGtP | T498 | 0.149 | 96.16 | 671 | 0.101769 | 0.149 | 0.1018 | 1 |
| P05771 | PRKCB | tFCGtPDyIAPEIIA | Y507 | 0.196 | 97.97 | 671 | 0.085982 | 0.196 | 0.086 | 1 |
| P29322 | EPHA8 | NQNNWLRtsWVPRDG | T86 | 0.008 | 95.04 | 1005 | 0.005464 | 0.008 | 0.0055 | 1 |
| P29322 | EPHA8 | LEYEIKYyEKDKEMQ | Y478 | 0.152 | 93.05 | 1005 | 0.06668 | 0.296 | 0.1279 | 1 |
| Q7Z5Q1 | CPEB2 | AFSNQQSyIAAISAR | Y495 | 0.063 | 97.07 | 589 | 0.027637 | 0.063 | 0.0276 | 0.8462 |

| | | | | | | | | | | |
|---|---|---|---|---|---|---|---|---|---|---|
| Q9NPF4 | OSGEP | AVLGFEGsANkIGVG | S10 | 0.058 | 96.81 | 335 | 0.046105 | 0.058 | 0.0461 | 1 |
| Q9NPF4 | OSGEP | ITGATSPtVLyVsGG | T127 | 0.01 | 98.34 | 335 | 0.00683 | 0.01 | 0.0068 | 1 |
| Q9NPF4 | OSGEP | ATSPtVLyVsGGNTQ | Y130 | 0 | 98.19 | 335 | 0 | 0 | 0 | 0.8 |
| Q9NPF4 | OSGEP | SPtVLyVsGGNTQVI | S132 | 0.09 | 94.82 | 335 | 0.071542 | 0.09 | 0.0715 | 0.8 |
| Q9NPF4 | OSGEP | HRTPLSDsGVTQRYR | S318 | 0.016 | 97.24 | 335 | 0.012719 | 0.016 | 0.0127 | 1 |
| Q9BWX1 | PHF7 | IQKYAHTsAKHFFKC | S197 | 0.026 | 96.21 | 381 | 0.020668 | 0.026 | 0.0207 | 1 |
| Q969M7 | UBE2F | GGKFQFEtEVPDAYN | T85 | 0.009 | 98.57 | 185 | 0.006147 | 0.009 | 0.0061 | 0.9091 |
| O15067 | PFAS | GNVLkELsDPAGAII | S530 | 0 | 95.49 | 1338 | 0 | 0 | 0 | 1 |
| O15067 | PFAS | PAGAIIytSRFQLGD | T539 | 0.013 | 94.67 | 1338 | 0.008879 | 0.013 | 0.0089 | 1 |
| O15067 | PFAS | VASkRYLtNkVDRSV | T681 | 0.081 | 96.26 | 1338 | 0.055324 | 0.149 | 0.1018 | 1 |
| O15067 | PFAS | HLLYVALsPGQHRLG | S857 | 0.086 | 95.52 | 1338 | 0.068363 | 0.091 | 0.0814 | 1 |
| Q8IUD2 | ERC1 | RGLtPPAsyNLDDDQ | S1050 | 0.084 | 83.4 | 1116 | 0.066773 | 0.084 | 0.0668 | 1 |
| Q6UWX4 | HHIPL2 | AGSHGKRyRVPSDNP | Y395 | 0.123 | 95.58 | 724 | 0.053958 | 0.123 | 0.054 | 1 |
| Q6UWX4 | HHIPL2 | PGAHPAIyAyGIRNM | Y414 | 0.249 | 98.7 | 724 | 0.109233 | 0.249 | 0.1092 | 1 |
| Q6UWX4 | HHIPL2 | AHPAIyAyGIRNMWR | Y416 | 0.246 | 98.42 | 724 | 0.107917 | 0.246 | 0.1079 | 1 |
| Q6UWX4 | HHIPL2 | THSKFIIsFAEDEAG | S568 | 0.003 | 98.33 | 724 | 0.002385 | 0.003 | 0.0024 | 1 |
| P17655 | CAPN2 | IkYLNQDyEALRNEC | Y32 | 0.176 | 98.13 | 700 | 0.077209 | 0.176 | 0.0772 | 0.8889 |
| P62263 | RPS14 | VkADRDEssPyAAML | S69 | 0.118 | 95.65 | 151 | 0.0938 | 0.118 | 0.0938 | 0.875 |
| P62263 | RPS14 | IGRIEDVtPIPsDst | T133 | 0.158 | 96.43 | 151 | 0.107916 | 0.198 | 0.122 | 0.9 |
| Q9NYV6 | RRN3 | IARYVPstPWFLMPI | T200 | 0 | 95.42 | 651 | 0 | 0.162 | 0.1106 | 1 |
| Q9NYV6 | RRN3 | VHNLLRIsVYFPTLR | S233 | 0.05 | 96.68 | 651 | 0.039746 | 0.178 | 0.1415 | 0.875 |
| Q9UET6 | FTSJ1 | SKDKRDVyYRLAKEN | Y12 | 0.078 | 95.5 | 329 | 0.034217 | 0.078 | 0.0342 | 1 |
| Q11206 | ST3GAL4 | MDFHWIEtILSDKKR | T202 | 0.052 | 95.54 | 333 | 0.035517 | 0.052 | 0.0355 | 1 |
| Q9Y664 | KPTN | LWQRsFssPLLAMAH | S364 | 0.038 | 97.21 | 436 | 0.030207 | 0.038 | 0.0302 | 0.8333 |
| Q9UIG0 | BAZ1B | PkFLPHkyDVkLQNE | Y222 | 0 | 89.63 | 1483 | 0 | 0 | 0 | 0.9375 |
| Q86WV6 | STING1 | YRGSYWRtVRACLGC | T84 | 0.013 | 93.14 | 379 | 0.008879 | 0.013 | 0.0089 | 1 |
| Q14558 | PRPSAP1 | AGLTHIItMDLHQKE | T132 | 0.001 | 97.74 | 356 | 0.000683 | 0.001 | 0.0007 | 0.8 |
| Q14558 | PRPSAP1 | DAAKRAQsYAERLRL | S186 | 0.052 | 93.6 | 356 | 0.041336 | 0.141 | 0.1121 | 0.8333 |
| P46059 | SLC15A1 | LDWAKEKyDERLISQ | Y269 | 0.305 | 90.14 | 708 | 0.133799 | 0.305 | 0.1338 | 1 |
| O14757 | CHEK1 | KRMtRFFtkLDADks | T382 | 0.06 | 89 | 476 | 0.040981 | 0.06 | 0.041 | 0.8889 |
| O14757 | CHEK1 | tkLDADksYQCLKET | S389 | 0.029 | 89.43 | 476 | 0.023053 | 0.029 | 0.0231 | 0.8889 |

| | | | | | | | | | | |
|---|---|---|---|---|---|---|---|---|---|---|
| Q9H2P0 | ADNP | RksFLtkyFNkQPYP | Y774 | 0 | 92.85 | 1102 | 0 | 0 | 0 | 1 |
| Q99714 | HSD17B10 | KGGIVGMtLPIARDL | T179 | 0.014 | 98.94 | 261 | 0.009562 | 0.014 | 0.0096 | 0.8 |
| Q99714 | HSD17B10 | IAPGLFGtPLLtsLP | T203 | 0.193 | 90.08 | 261 | 0.131822 | 0.193 | 0.1318 | 0.8 |
| P16157 | ANK1 | VNFAtLLyKELTAVP | Y1230 | 0.091 | 89.63 | 1881 | 0.03992 | 0.143 | 0.0532 | 0.8333 |
| P16157 | ANK1 | REGRLRCyCMTDDKV | Y1259 | 0.038 | 85.79 | 1881 | 0.01667 | 0.159 | 0.0698 | 1 |
| Q9H4A5 | GOLPH3L | GDVLLDEtLkHIKAT | T122 | 0 | 98.66 | 285 | 0 | 0 | 0 | 1 |
| P04196 | HRG | RVENTTVyyLVLDVQ | Y67 | 0.15 | 91.98 | 525 | 0.065803 | 0.15 | 0.0658 | 1 |
| P04196 | HRG | VIDFNCTtSSVSSAL | T128 | 0.025 | 95.5 | 525 | 0.017075 | 0.025 | 0.0171 | 0.8333 |
| Q96QK1 | VPS35 | YEsRKsMsCYVLSNV | S437 | 0 | 96.12 | 796 | 0 | 0 | 0 | 0.8 |
| Q12899 | TRIM26 | GFTWGkVyWEVEVER | Y369 | 0.092 | 97.36 | 539 | 0.040359 | 0.092 | 0.0404 | 1 |
| Q8IW45 | NAXD | VGGCQEytGAPYFAA | T86 | 0.165 | 98.14 | 347 | 0.112697 | 0.165 | 0.1127 | 1 |
| Q9NPB9 | ACKR4 | VVIVFIVtQLPyNIV | T252 | 0.087 | 90.59 | 350 | 0.059422 | 0.087 | 0.0594 | 1 |
| Q14690 | PDCD11 | GKKSVkLsLNPkNVN | S169 | 0.005 | 95.68 | 1871 | 0.003975 | 0.005 | 0.004 | 1 |
| Q14690 | PDCD11 | EAKKLMMtLKKTLIE | T520 | 0 | 95.78 | 1871 | 0 | 0 | 0 | 1 |
| P60484 | PTEN | HKNHYkIyNLCAERH | Y68 | 0.001 | 98.74 | 403 | 0.000439 | 0.001 | 0.0004 | 1 |
| P60484 | PTEN | TGVMICAyLLHRGKF | Y138 | 0.141 | 98.9 | 403 | 0.061855 | 0.282 | 0.1237 | 1 |
| P60484 | PTEN | AQEALDFyGEVRTRD | Y155 | 0 | 98.42 | 403 | 0 | 0 | 0 | 1 |
| P60484 | PTEN | QRRyVyyysyLLKNH | Y178 | 0.107 | 98.26 | 403 | 0.046939 | 0.107 | 0.0469 | 1 |
| P60484 | PTEN | LKVkIYSsNsGPtRR | S227 | 0.116 | 97.34 | 403 | 0.09221 | 0.116 | 0.0922 | 1 |
| P60484 | PTEN | RREDKFMyFEFPQPL | Y240 | 0.286 | 98.21 | 403 | 0.125464 | 0.286 | 0.1255 | 1 |
| O00408 | PDE2A | EHIAMPIyKLLQDLF | Y871 | 0.005 | 98.19 | 941 | 0.002193 | 0.005 | 0.0022 | 1 |
| Q6UXZ0 | TMIGD1 | VNGKtENyILDTTPG | Y40 | 0.335 | 98.12 | 262 | 0.14696 | 0.335 | 0.147 | 1 |
| Q3KR16 | PLEKHG6 | LLTTELIyVRKLKIM | Y176 | 0.133 | 94.72 | 790 | 0.058345 | 0.133 | 0.0583 | 1 |
| Q3KR16 | PLEKHG6 | WCEKHKRsGRQMLCD | S297 | 0.035 | 79.78 | 790 | 0.027822 | 0.035 | 0.0278 | 1 |
| Q3KR16 | PLEKHG6 | PHQRITKyPLLLHAV | Y316 | 0.004 | 97.82 | 790 | 0.001755 | 0.004 | 0.0018 | 1 |
| Q13395 | TARBP1 | DLDRCHLyLMVLtEL | Y754 | 0 | 94.62 | 1621 | 0 | 0.337 | 0.1478 | 1 |
| Q8WV16 | DCAF4 | DEQYLMAsDMAGKIK | S388 | 0.007 | 98.2 | 495 | 0.005564 | 0.007 | 0.0056 | 1 |
| Q9BUB5 | MKNK1 | AKVQGAVsLQNGKEY | S68 | 0.116 | 96.58 | 465 | 0.09221 | 0.187 | 0.1486 | 1 |
| Q14331 | FRG1 | DSRIALksGYGkYLG | S111 | 0.084 | 91.94 | 258 | 0.066773 | 0.084 | 0.0668 | 0.8 |
| Q9ULI1 | NWD2 | HEDTGPEsDPVVIVR | S445 | 0.097 | 82.67 | 1742 | 0.077107 | 0.097 | 0.0771 | 1 |
| P22670 | RFX1 | GLRIKAssPLLRLME | S515 | 0.097 | 92.14 | 979 | 0.077107 | 0.097 | 0.0771 | 1 |

| | | | | | | | | | | |
|---|---|---|---|---|---|---|---|---|---|---|
| P42858 | HTT | SSPTIRRtAAGSAVs | T269 | 0.054 | 96.92 | 3142 | 0.036883 | 0.054 | 0.0369 | 1 |
| P42858 | HTT | tAAGSAVsICQHSRR | S276 | 0.097 | 94.755 | 3142 | 0.077107 | 0.097 | 0.0771 | 1 |
| Q86WU2 | LDHD | VDPGADAsLCGMAAT | S163 | 0.034 | 98.15 | 507 | 0.027027 | 0.034 | 0.027 | 1 |
| O75787 | ATP6AP2 | SILKsPGsVVFRNGN | S27 | 0.183 | 85.5 | 350 | 0.145469 | 0.183 | 0.1455 | 0.8333 |
| Q8TE73 | DNAH5 | GLRHLMFyyQDVEEA | Y83 | 0.141 | 90.58 | 4624 | 0.061855 | 0.226 | 0.0792 | 1 |
| O95243 | MBD4 | RFDVyFIsPQGLkFR | S112 | 0.057 | 91.47 | 580 | 0.04531 | 0.057 | 0.0453 | 0.8333 |
| Q02127 | DHODH | HRLAVRFtsLGLLPR | T62 | 0.017 | 97.57 | 395 | 0.011611 | 0.034 | 0.0232 | 1 |
| Q02127 | DHODH | FVEIGSVtPKPQEGN | T121 | 0 | 98.61 | 395 | 0 | 0 | 0 | 1 |
| Q02127 | DHODH | SVDAAEDyAEGVRVL | Y194 | 0.003 | 98.76 | 395 | 0.001316 | 0.003 | 0.0013 | 1 |
| Q02127 | DHODH | YLVVNVSsPNtAGLR | S214 | 0.101 | 96.41 | 395 | 0.080286 | 0.101 | 0.0803 | 1 |
| Q02127 | DHODH | VNVSsPNtAGLRSLQ | T217 | 0.114 | 91.93 | 395 | 0.077864 | 0.114 | 0.0779 | 1 |
| P08567 | PLEK | DPAYLHyyDPAGAED | Y278 | 0.234 | 92.47 | 350 | 0.102652 | 0.234 | 0.1027 | 0.8 |
| P28827 | PTPRM | RyGNIIAyDHSRVRL | Y935 | 0.002 | 95.93 | 1452 | 0.000877 | 0.002 | 0.0009 | 1 |
| P53350 | PLK1 | LELCRRRsLLELHkR | S137 | 0.044 | 96.35 | 603 | 0.034976 | 0.044 | 0.035 | 1 |
| P53350 | PLK1 | tLCGtPNyIAPEVLs | Y217 | 0.212 | 93.86 | 603 | 0.093001 | 0.212 | 0.093 | 1 |
| Q6P996 | PDXDC1 | LVEDELSsPVVVFRF | S387 | 0.062 | 91.76 | 788 | 0.049285 | 0.062 | 0.0493 | 0.875 |
| Q9GZN1 | ACTR6 | ENTVMIDyVLPDFST | Y236 | 0.009 | 95.2 | 396 | 0.003948 | 0.009 | 0.0039 | 1 |
| Q5HYJ3 | FAM76B | RYPFEELsQGQQLCK | S22 | 0.171 | 88.12 | 339 | 0.13593 | 0.171 | 0.1359 | 1 |
| Q8TDX6 | CSGALNACT1 | LNtCRLNtQPGKKVF | T375 | 0.109 | 97.89 | 532 | 0.074448 | 0.109 | 0.0744 | 1 |
| Q9NRZ5 | AGPAT4 | TEKKHEIsMQVARAK | S188 | 0.147 | 97.14 | 378 | 0.116852 | 0.147 | 0.1169 | 1 |
| Q9NRZ5 | AGPAT4 | RNVVsAVyDCTLNFR | Y226 | 0.058 | 98.61 | 378 | 0.025444 | 0.058 | 0.0254 | 0.8125 |
| Q9H857 | NT5DC2 | RTFHNPTyFSRRLVR | Y457 | 0.121 | 93.8 | 520 | 0.053081 | 0.121 | 0.0531 | 0.9375 |
| Q12996 | CSTF3 | IDkARKTyERLVAQF | Y54 | 0.058 | 98.46 | 717 | 0.025444 | 0.058 | 0.0254 | 1 |
| Q12996 | CSTF3 | LVAQFPssGRFWKLY | S64 | 0.054 | 97.67 | 717 | 0.042925 | 0.107 | 0.0851 | 1 |
| Q12996 | CSTF3 | LWKCyLSyVRETKGk | Y107 | 0.169 | 96.96 | 717 | 0.074138 | 0.335 | 0.1462 | 1 |
| Q12996 | CSTF3 | KEKMAQAyDFALDKI | Y126 | 0.022 | 98.64 | 717 | 0.009651 | 0.022 | 0.0097 | 1 |
| Q12996 | CSTF3 | KIGMEIMsyQIWVDY | S139 | 0.001 | 97.86 | 717 | 0.000795 | 0.001 | 0.0008 | 1 |
| Q12996 | CSTF3 | tEDQTLItkRVMFAY | T273 | 0.003 | 98.56 | 717 | 0.002049 | 0.003 | 0.002 | 1 |
| Q12996 | CSTF3 | SELkALGykDVSRAK | Y546 | 0.125 | 84.27 | 717 | 0.054836 | 0.367 | 0.1418 | 1 |
| O15068 | MCF2L | VERARALsLDGEQLI | S412 | 0.09 | 97.59 | 1137 | 0.071542 | 0.179 | 0.1423 | 1 |
| P08397 | HMBS | sVPVAVHtAMKDGQL | T269 | 0.031 | 98.51 | 361 | 0.021173 | 0.031 | 0.0212 | 1 |

| | | | | | | | | | | |
|---|---|---|---|---|---|---|---|---|---|---|
| Q15399 | TLR1 | KTtILNIsQNyISEL | S53 | 0.105 | 98.06 | 786 | 0.083466 | 0.16 | 0.1272 | 1 |
| Q15399 | TLR1 | KkGDCsWtKSLLSLN | T422 | 0.098 | 94.74 | 786 | 0.066935 | 0.098 | 0.0669 | 1 |
| Q15399 | TLR1 | SLLSLNMsSNILTDT | S431 | 0.05 | 98.28 | 786 | 0.039746 | 0.05 | 0.0397 | 1 |
| Q9NQC7 | CYLD | LAFGGYLsEVVEENT | S568 | 0.14 | 79.65 | 956 | 0.111288 | 0.14 | 0.1113 | 0.8571 |
| O00214 | LGALS8 | tPMGPGRtVVVkGEV | T200 | 0.111 | 95.87 | 317 | 0.075815 | 0.111 | 0.0758 | 0.8 |
| Q5VV42 | CDKAL1 | VVEVVEEtIkGHsVR | T170 | 0.09 | 95.67 | 579 | 0.061471 | 0.09 | 0.0615 | 1 |
| Q5VV42 | CDKAL1 | ILNHPRVyAFLHIPV | Y315 | 0.025 | 98.71 | 579 | 0.010967 | 0.025 | 0.011 | 1 |
| Q2TAY7 | SMU1 | kQTQPERyIHLENLL | Y114 | 0.151 | 86.81 | 513 | 0.066242 | 0.151 | 0.0662 | 1 |
| Q5T6F0 | DCAF12 | NKELGAVsLDGyFHL | S271 | 0.003 | 94.45 | 453 | 0.002385 | 0.068 | 0.0541 | 1 |
| P13797 | PLS3 | NLNLALNsAsAIGCH | S203 | 0.002 | 98.53 | 630 | 0.00159 | 0.127 | 0.101 | 1 |
| P13797 | PLS3 | DIkDSkAyFHLLNQI | Y302 | 0.043 | 98.47 | 630 | 0.018863 | 0.043 | 0.0189 | 1 |
| Q4VC44 | FLYWCH1 | QARLGCRsRAITQGH | S641 | 0.001 | 89.71 | 716 | 0.000795 | 0.001 | 0.0008 | 0.875 |
| P10515 | DLAT | kVPLPsLsPtMQAGT | S100 | 0.117 | 80 | 647 | 0.093005 | 0.117 | 0.093 | 1 |
| Q96EY7 | PTCD3 | YEQALNLytELLNNR | Y279 | 0.118 | 96.74 | 689 | 0.051765 | 0.118 | 0.0518 | 1 |
| Q14CZ7 | FASTKD3 | RHLQLLGyQVVQIPY | Y624 | 0.326 | 96.99 | 662 | 0.143012 | 0.326 | 0.143 | 0.9167 |
| P0CG48 | UBC | DyNIQkEstLHLVLr | S65 | 0.177 | 96.16 | 685 | 0.1407 | 0.177 | 0.1407 | 1 |
| Q9Y5A9 | YTHDF2 | IFDDFsHyEkRQEEE | Y560 | 0.218 | 94.67 | 579 | 0.095633 | 0.218 | 0.0956 | 1 |
| Q9BT04 | FUZ | KsFHDSItLIVLsSE | T80 | 0.041 | 95.99 | 418 | 0.028004 | 0.041 | 0.028 | 1 |
| Q9BT04 | FUZ | SItLIVLsSEVGISE | S85 | 0.023 | 93.76 | 418 | 0.018283 | 0.023 | 0.0183 | 1 |
| P28072 | PSMB6 | AMERDGssGGVIRLA | S204 | 0.015 | 94.04 | 239 | 0.011924 | 0.015 | 0.0119 | 0.9167 |
| Q03001 | DST | VNLVLKNtQAAEALV | T1181 | 0.024 | 92.825 | 7570 | 0.016392 | 0.024 | 0.0164 | 1 |
| Q562E7 | WDR81 | GRISNFHyLMQLNRL | Y362 | 0 | 92.53 | 1941 | 0 | 0 | 0 | 1 |
| P20908 | COL5A1 | WPkENPGsWFSEFKR | S1702 | 0.124 | 92.99 | 1838 | 0.098569 | 0.124 | 0.0986 | 0.8 |
| Q96BR9 | ZBTB8A | QDVFCDCsILVEGKV | S27 | 0.186 | 91.18 | 441 | 0.147854 | 0.186 | 0.1479 | 1 |
| P49441 | INPP1 | DTTFKWDsCAAHAIL | S317 | 0.02 | 98.5 | 399 | 0.015898 | 0.02 | 0.0159 | 0.8 |
| P29323 | EPHB2 | KYLADMNyVHRDLAA | Y742 | 0.073 | 84.91 | 1055 | 0.032024 | 0.263 | 0.1154 | 0.8571 |
| Q7L8C5 | SYT13 | HDGGCDCyVQGsVAN | Y194 | 0.1 | 92.8 | 426 | 0.043869 | 0.1 | 0.0439 | 1 |
| Q86VD9 | PIGZ | LPSTPTHyTLLFTHT | Y458 | 0.06 | 91.52 | 579 | 0.026321 | 0.06 | 0.0263 | 0.9091 |
| Q86VD9 | PIGZ | FVVTPGttRRAVEKC | T527 | 0.165 | 93.42 | 579 | 0.112697 | 0.165 | 0.1127 | 1 |
| P29144 | TPP2 | KLDSsDIyNELkETY | Y1042 | 0.283 | 86.23 | 1249 | 0.124148 | 0.283 | 0.1241 | 0.8182 |
| Q8IX18 | DHX40 | kSEALQRsGRAGRTS | S394 | 0.06 | 91.71 | 779 | 0.047695 | 0.06 | 0.0477 | 1 |

| | | | | | | | | | | |
|---|---|---|---|---|---|---|---|---|---|---|
| Q8IX18 | DHX40 | GkCFRIYsKDFWNQC | S410 | 0.142 | 94.24 | 779 | 0.112878 | 0.142 | 0.1129 | 1 |
| Q8IX18 | DHX40 | CTMDGRGsPVHIHPS | S648 | 0.165 | 88.86 | 779 | 0.131161 | 0.165 | 0.1312 | 1 |
| Q9C0B1 | FTO | DTWLPYLtPKDDEFY | T32 | 0.051 | 97.62 | 505 | 0.034834 | 0.054 | 0.0486 | 1 |
| P15498 | VAV1 | GKVIyTLsALSWTPI | S113 | 0.15 | 95.56 | 845 | 0.119237 | 0.15 | 0.1192 | 0.8571 |
| Q12851 | MAP4K2 | NVVAYIGsyLRNDRL | S79 | 0.055 | 94.16 | 820 | 0.04372 | 0.055 | 0.0437 | 1 |
| Q96KB5 | PBK | MEYGGEKsLNDLIEE | S122 | 0.161 | 96 | 322 | 0.127981 | 0.161 | 0.128 | 1 |
| Q9HA64 | FN3KRP | LDAPGGGsVLVMEHM | S84 | 0.025 | 95.76 | 309 | 0.019873 | 0.05 | 0.0397 | 1 |
| Q9HA64 | FN3KRP | GSGyRGSsLNIMRNL | S300 | 0 | 98.4 | 309 | 0 | 0 | 0 | 1 |
| Q06033 | ITIH3 | VAFVIDIsGsMAGRK | S292 | 0.009 | 93.86 | 890 | 0.007154 | 0.009 | 0.0072 | 1 |
| Q06033 | ITIH3 | SMEDKGMtNINDGLL | T360 | 0.079 | 89.64 | 890 | 0.053958 | 0.079 | 0.054 | 1 |
| Q06033 | ITIH3 | EHRIPERstsIVIML | S386 | 0.163 | 95.51 | 890 | 0.129571 | 0.163 | 0.1296 | 0.875 |
| Q06033 | ITIH3 | RIPERstsIVIMLTD | S388 | 0.125 | 97.88 | 890 | 0.099364 | 0.125 | 0.0994 | 1 |
| Q06033 | ITIH3 | ALDLSLKyHFVTPLT | Y592 | 0.132 | 90.77 | 890 | 0.057907 | 0.132 | 0.0579 | 0.9 |
| P06703 | S100A6 | VAIFHKysGREGDkH | S20 | 0.123 | 91.91 | 90 | 0.097775 | 0.123 | 0.0978 | 1 |
| Q01344 | IL5RA | QRNVNLEyQVKINAP | Y65 | 0.128 | 95.89 | 420 | 0.056152 | 0.128 | 0.0562 | 1 |
| P49641 | MAN2A2 | GLTGLLKsIRRVDEE | S783 | 0.032 | 97.98 | 1150 | 0.025437 | 0.032 | 0.0254 | 1 |
| P49641 | MAN2A2 | LPLQANFyPMPVMAY | Y923 | 0.19 | 98.64 | 1150 | 0.08335 | 0.19 | 0.0833 | 1 |
| Q9UBT2 | UBA2 | TTIkkGVtECyECHP | T156 | 0.083 | 98.47 | 640 | 0.05669 | 0.083 | 0.0567 | 1 |
| P49589 | CARS1 | CRLHLyNsLtRNkEV | S34 | 0.045 | 96.57 | 748 | 0.035771 | 0.045 | 0.0358 | 1 |
| P49589 | CARS1 | DGkkVTWyCCGPTVy | Y53 | 0.004 | 98.72 | 748 | 0.001755 | 0.004 | 0.0018 | 1 |
| P49589 | CARS1 | GPTVyDAsHMGHARS | S63 | 0.084 | 94.84 | 748 | 0.066773 | 0.084 | 0.0668 | 0.8889 |
| P49589 | CARS1 | VLPPDVLtRVSEYVP | T239 | 0.006 | 98.13 | 748 | 0.004098 | 0.006 | 0.0041 | 1 |
| P49589 | CARS1 | QKIVDNGyGyVSNGS | Y260 | 0.146 | 97.52 | 748 | 0.064048 | 0.247 | 0.0923 | 1 |
| P49589 | CARS1 | IVDNGyGyVSNGSVY | Y262 | 0.15 | 97.28 | 748 | 0.065803 | 0.15 | 0.0658 | 1 |
| P49589 | CARS1 | LVSQCNLyMAARKAV | Y528 | 0.165 | 97.18 | 748 | 0.072383 | 0.165 | 0.0724 | 1 |
| Q9UHQ9 | CYB5R1 | VVEFRGPsGLLTYTG | S150 | 0.187 | 97.5 | 305 | 0.148649 | 0.187 | 0.1486 | 0.8 |
| Q8NAT2 | TDRD5 | MIEMRRCysNQLVsD | Y510 | 0.026 | 92.7 | 981 | 0.011406 | 0.026 | 0.0114 | 1 |
| Q8NAT2 | TDRD5 | CysNQLVsDRYVMPE | S516 | 0.058 | 91.41 | 981 | 0.046105 | 0.058 | 0.0461 | 1 |
| P09382 | LGALS1 | ANtIVCNskDGGAWG | S63 | 0.088 | 98.79 | 135 | 0.069952 | 0.088 | 0.07 | 1 |
| Q10469 | MGAT2 | YLRLLLDsLRKAQGI | S137 | 0.009 | 98.5 | 447 | 0.007154 | 0.009 | 0.0072 | 1 |
| P53673 | CRYBA4 | EFTAECPsVLELGFE | S35 | 0.161 | 97.87 | 196 | 0.127981 | 0.161 | 0.128 | 1 |

| | | | | | | | | | | |
|---|---|---|---|---|---|---|---|---|---|---|
| P51575 | P2RX1 | VRHCRPIyEFHGLYE | Y274 | 0.165 | 96.24 | 399 | 0.072383 | 0.165 | 0.0724 | 1 |
| Q86UW1 | SLC51A | LWTLGIIsRQARLHL | S240 | 0.067 | 88.63 | 340 | 0.053259 | 0.163 | 0.1296 | 1 |
| Q709C8 | VPS13C | EDsPADQtLIVQSQP | T624 | 0.124 | 90.45 | 3753 | 0.084694 | 0.124 | 0.0847 | 1 |
| Q00796 | SORD | AAkPNNLsLVVHGPG | S11 | 0.016 | 98.88 | 357 | 0.012719 | 0.016 | 0.0127 | 1 |
| Q00796 | SORD | ICGSDVHyWEyGRIG | Y51 | 0.07 | 98.49 | 357 | 0.030708 | 0.07 | 0.0307 | 1 |
| Q15031 | LARS2 | DKSKPKFyVLsMFPY | Y85 | 0 | 97.28 | 903 | 0 | 0 | 0 | 1 |
| Q15031 | LARS2 | KPKFyVLsMFPYPSG | S88 | 0.032 | 96.05 | 903 | 0.025437 | 0.064 | 0.0509 | 1 |
| Q15031 | LARS2 | HVRVYTIsDtIARFQ | S108 | 0 | 97.37 | 903 | 0 | 0 | 0 | 1 |
| Q15031 | LARS2 | RVYTIsDtIARFQKM | T110 | 0 | 98.18 | 903 | 0 | 0 | 0 | 1 |
| Q15031 | LARS2 | GTSHVAIsPSHRLLH | S315 | 0.03 | 93.02 | 903 | 0.023847 | 0.06 | 0.0477 | 1 |
| Q15031 | LARS2 | VPGKDCLtPVMAVNM | T343 | 0.103 | 96.43 | 903 | 0.07035 | 0.174 | 0.1153 | 1 |
| Q6YHK3 | CD109 | LNSADEIyELRVTGR | Y102 | 0.019 | 89.88 | 1445 | 0.008335 | 0.019 | 0.0083 | 1 |
| O15195 | VILL | GASSDLHyWVGKQAG | Y62 | 0.138 | 94.76 | 856 | 0.060539 | 0.256 | 0.1123 | 0.8333 |
| O95450 | ADAMTS2 | RCHLyCEsREtGEVV | S661 | 0.034 | 89 | 1211 | 0.027027 | 0.034 | 0.027 | 1 |
| Q92974 | ARHGEF2 | IDEAEVIySELMSDF | Y198 | 0.161 | 75.41 | 986 | 0.070628 | 0.161 | 0.0706 | 1 |
| Q92974 | ARHGEF2 | VTQRItkyPLLISRI | Y394 | 0.016 | 98.32 | 986 | 0.007019 | 0.016 | 0.007 | 1 |
| Q8WVC6 | DCAKD | YFLRGYRyVILDIPL | Y109 | 0.063 | 98.17 | 231 | 0.027637 | 0.063 | 0.0276 | 1 |
| Q96KN7 | RPGRIP1 | VTSDLDHyLRREALS | Y864 | 0.246 | 91.05 | 1286 | 0.107917 | 0.246 | 0.1079 | 1 |
| Q96PZ2 | FAM111A | KLtHsENsSLYMALN | S102 | 0.147 | 84.64 | 611 | 0.116852 | 0.147 | 0.1169 | 1 |
| Q8N612 | FHIP1B | VEEMIAStAYLELFL | T351 | 0 | 96.36 | 972 | 0 | 0 | 0 | 1 |
| P63241 | EIF5A | GIQDGyLsLLQDSGE | S100 | 0.119 | 95.95 | 154 | 0.094595 | 0.119 | 0.0946 | 1 |
| O95157 | NXPH3 | CsRDHAQsSAtWSCS | S206 | 0.02 | 95.39 | 252 | 0.015898 | 0.02 | 0.0159 | 1 |
| Q5GFL6 | VWA2 | RAELEEItGsPKHVM | T491 | 0.094 | 84.78 | 755 | 0.064203 | 0.094 | 0.0642 | 1 |
| Q6NSJ0 | MYORG | DVTSIHKyMVRRyFN | Y295 | 0.291 | 96.94 | 714 | 0.127658 | 0.291 | 0.1277 | 1 |
| P11387 | TOP1 | AkMLDHEyTTkEIFR | Y268 | 0.34 | 95.51 | 765 | 0.149153 | 0.34 | 0.1492 | 1 |
| P11387 | TOP1 | GEKDWQkyETARRLK | Y444 | 0 | 98.02 | 765 | 0 | 0 | 0 | 1 |
| P11387 | TOP1 | RQRAVALyFIDkLAL | Y480 | 0.157 | 98.47 | 765 | 0.068874 | 0.157 | 0.0689 | 1 |
| P11387 | TOP1 | ADTVGCCsLRVEHIN | S506 | 0.155 | 97.8 | 765 | 0.123212 | 0.155 | 0.1232 | 1 |
| P11387 | TOP1 | LTAkVFRtyNASITL | T591 | 0.085 | 97.83 | 765 | 0.058056 | 0.085 | 0.0581 | 0.9167 |
| Q9H0D6 | XRN2 | VRPRRLLyMAIDGVA | Y99 | 0.025 | 98.69 | 950 | 0.010967 | 0.025 | 0.011 | 1 |
| Q9H0D6 | XRN2 | NLAkCLRyyIADRLN | Y176 | 0.128 | 98.02 | 950 | 0.056152 | 0.241 | 0.1057 | 1 |

| | | | | | | | | | | |
|---|---|---|---|---|---|---|---|---|---|---|
| Q9H0D6 | XRN2 | LAkCLRyyIADRLNN | Y177 | 0.015 | 98.12 | 950 | 0.00658 | 0.015 | 0.0066 | 1 |
| Q9H0D6 | XRN2 | HktGGYLtESGYVNL | T379 | 0.051 | 96.62 | 950 | 0.034834 | 0.077 | 0.0587 | 1 |
| Q5UIP0 | RIF1 | MVSSIIDsLEILFNK | S143 | 0.035 | 90.34 | 2472 | 0.027822 | 0.035 | 0.0278 | 0.8571 |
| Q96PU5 | NEDD4L | ILRVkVVsGIDLAKK | S28 | 0.121 | 97.38 | 975 | 0.096185 | 0.121 | 0.0962 | 1 |
| Q96DN5 | TBC1D31 | QYPVFNQyPkFIVDY | Y680 | 0.321 | 82.94 | 1066 | 0.140818 | 0.321 | 0.1408 | 1 |
| P27352 | CBLIF | SNITVIYtINNQLRG | T317 | 0.091 | 98.35 | 417 | 0.062154 | 0.091 | 0.0622 | 0.875 |
| O60341 | KDM1A | QsFGMDVtLLEARDR | T305 | 0.086 | 98.9 | 852 | 0.058739 | 0.086 | 0.0587 | 1 |
| O60341 | KDM1A | GLDIkLNtAVRQVRY | T588 | 0.079 | 98.85 | 852 | 0.053958 | 0.079 | 0.054 | 1 |
| Q8TCT9 | HM13 | EVtEMFSyEESNPkD | Y346 | 0.303 | 92.79 | 377 | 0.132922 | 0.303 | 0.1329 | 1 |
| O94759 | TRPM2 | MLALAEEyEHRAIGV | Y682 | 0.013 | 95.47 | 1503 | 0.005703 | 0.013 | 0.0057 | 1 |
| Q92545 | TMEM131 | DLQKNIIsKITAELS | S818 | 0 | 83.43 | 1883 | 0 | 0 | 0 | 1 |
| Q92545 | TMEM131 | IIVRNNLtVMDAVMV | T961 | 0.019 | 88.75 | 1883 | 0.012977 | 0.019 | 0.013 | 1 |
| Q92545 | TMEM131 | GkLPGPGssLRFkIT | S988 | 0.008 | 84.85 | 1883 | 0.006359 | 0.008 | 0.0064 | 1 |
| O94808 | GFPT2 | AVATRRGsPLLIGVR | S202 | 0.062 | 88.82 | 682 | 0.049285 | 0.144 | 0.1145 | 0.8571 |
| Q9Y4I1 | MYO5A | VEEKVLAsNPIMEsI | S199 | 0.007 | 87.97 | 1855 | 0.005564 | 0.007 | 0.0056 | 1 |
| Q9Y4I1 | MYO5A | AsNPIMEsIGNAKTT | S205 | 0.022 | 94.24 | 1855 | 0.017488 | 0.022 | 0.0175 | 1 |
| Q14520 | HABP2 | EIGSDDCyVGDGySy | Y195 | 0.289 | 85.38 | 560 | 0.12678 | 0.289 | 0.1268 | 1 |
| Q14520 | HABP2 | yVGDGySyRGKMNRT | Y202 | 0.059 | 86.76 | 560 | 0.025882 | 0.059 | 0.0259 | 1 |
| Q15291 | RBBP5 | LEsFGQNyPEEADGt | Y13 | 0.303 | 87.93 | 538 | 0.132922 | 0.303 | 0.1329 | 1 |
| Q15291 | RBBP5 | LQDLVNRtPWKkCCF | T252 | 0.054 | 94.64 | 538 | 0.036883 | 0.054 | 0.0369 | 1 |
| Q99466 | NOTCH4 | QPDRAGRtPLHAAVA | T1669 | 0.066 | 94.58 | 2003 | 0.045079 | 0.066 | 0.0451 | 1 |
| Q8NI35 | PATJ | NEYCLDNtSLAEAVE | T748 | 0.145 | 79.92 | 1801 | 0.099037 | 0.191 | 0.1305 | 1 |
| O94910 | ADGRL1 | RERTCKDyIKAVVET | Y616 | 0.029 | 94.78 | 1474 | 0.012722 | 0.029 | 0.0127 | 0.875 |
| P08134 | RHOC | AFGyLECsAKTKEGV | S160 | 0.002 | 98.79 | 193 | 0.00159 | 0.002 | 0.0016 | 1 |
| Q969X5 | ERGIC1 | NPLASHDyILkIVPT | Y187 | 0.033 | 98.12 | 290 | 0.014477 | 0.033 | 0.0145 | 1 |
| Q969X5 | ERGIC1 | LkIVPTVyEDkSGKQ | Y196 | 0.199 | 92.29 | 290 | 0.087298 | 0.199 | 0.0873 | 0.9091 |
| Q969X5 | ERGIC1 | KQRYSYQyTVANKEy | Y209 | 0.029 | 95.49 | 290 | 0.012722 | 0.029 | 0.0127 | 1 |
| Q96J02 | ITCH | FTVDGIPyFVDHNRR | Y495 | 0.25 | 86.76 | 903 | 0.109671 | 0.25 | 0.1097 | 1 |
| Q96J02 | ITCH | HNRRtttyIDPRTGk | Y506 | 0.231 | 90.74 | 903 | 0.101336 | 0.231 | 0.1013 | 1 |
| P10966 | CD8B | TNKMVMLsCEAKISL | S40 | 0.109 | 97.14 | 210 | 0.086646 | 0.109 | 0.0866 | 1 |
| Q13576 | IQGAP2 | IPECADKyyDALVkA | Y579 | 0.014 | 87.55 | 1575 | 0.006142 | 0.014 | 0.0061 | 1 |

| | | | | | | | | | | |
|---|---|---|---|---|---|---|---|---|---|---|
| Q13576 | IQGAP2 | LHKkyDyyyNTDsKE | Y612 | 0.077 | 86.7 | 1575 | 0.033779 | 0.131 | 0.0575 | 1 |
| O75864 | PPP1R37 | LALKVNHsLLRLDLD | S418 | 0.171 | 92.7 | 691 | 0.13593 | 0.171 | 0.1359 | 1 |
| Q86XA9 | HEATR5A | sPsIQLAsLEVVRQI | S1626 | 0.001 | 93.04 | 2040 | 0.000795 | 0.001 | 0.0008 | 1 |
| P48735 | IDH2 | DQTDDQVtIDsALAT | T97 | 0 | 98.43 | 452 | 0 | 0 | 0 | 1 |
| P48735 | IDH2 | DDQVtIDsALATQky | S100 | 0 | 98.69 | 452 | 0 | 0 | 0 | 1 |
| P48735 | IDH2 | ALATQkysVAVkCAt | S108 | 0.038 | 98.09 | 452 | 0.030207 | 0.038 | 0.0302 | 1 |
| P48735 | IDH2 | sVAVkCAtItPDEAR | T115 | 0.164 | 98.15 | 452 | 0.112014 | 0.151 | 0.1464 | 1 |
| P48735 | IDH2 | IQkkWPLyMSTKNTI | Y247 | 0.177 | 98.87 | 452 | 0.077647 | 0.177 | 0.0776 | 1 |
| Q13075 | NAIP | sPDLStGyWKLSPKQ | Y989 | 0.013 | 90.08 | 1403 | 0.005703 | 0.013 | 0.0057 | 1 |
| Q13075 | NAIP | IRPALELsKASVtKC | S1051 | 0.113 | 87.45 | 1403 | 0.089825 | 0.113 | 0.1025 | 1 |
| Q13075 | NAIP | ELsKASVtKCSISKL | T1056 | 0.113 | 92.73 | 1403 | 0.077181 | 0.213 | 0.1487 | 1 |
| Q9NRY4 | ARHGAP35 | NENWLSVsRKMQASP | S296 | 0.174 | 93.49 | 1499 | 0.138315 | 0.174 | 0.1383 | 1 |
| Q9NRY4 | ARHGAP35 | ASPEYQDyVYLEGTQ | Y308 | 0.088 | 97.26 | 1499 | 0.038604 | 0.088 | 0.0386 | 1 |
| Q9BXN1 | ASPN | RCVLSRMsVQLGNFG | S372 | 0.053 | 96.19 | 380 | 0.04213 | 0.053 | 0.0421 | 1 |
| Q32P41 | TRMT5 | VRENNyTyEFDFSKV | Y272 | 0.007 | 98.58 | 509 | 0.003071 | 0.007 | 0.0031 | 1 |
| Q32P41 | TRMT5 | ANDLNPEsHkWLLYN | S332 | 0.05 | 97.46 | 509 | 0.039746 | 0.05 | 0.0397 | 1 |
| Q32P41 | TRMT5 | QIPASVLyKNQTRNP | Y474 | 0.058 | 89.17 | 509 | 0.025444 | 0.148 | 0.0649 | 0.9 |
| O75343 | GUCY1B2 | ESMWCMVyLCSPKLR | Y291 | 0.003 | 96.33 | 617 | 0.001316 | 0.003 | 0.0013 | 1 |
| Q8TE73 | DNAH5 | TSLFVKVtNQIIsAC | T401 | 0.013 | 92.01 | 4624 | 0.008879 | 0.013 | 0.0089 | 1 |
| Q8TE73 | DNAH5 | YQLILENyGADIDMI | Y596 | 0.031 | 91.75 | 4624 | 0.013599 | 0.031 | 0.0136 | 1 |
| Q8TE73 | DNAH5 | MKMMLAEyQRVKSKI | Y759 | 0.255 | 90.34 | 4624 | 0.111865 | 0.255 | 0.1119 | 1 |
| Q5VU65 | NUP210L | KMGRKytstPRHIEV | S1074 | 0.076 | 87.15 | 1888 | 0.060414 | 0.076 | 0.0604 | 1 |
| O95498 | VNN2 | DPHRFGHtPVQARLS | T119 | 0.036 | 97.33 | 520 | 0.024588 | 0.036 | 0.0246 | 1 |
| O75533 | SF3B1 | kDLLPRLtPILkNRH | T1021 | 0.104 | 90.09 | 1304 | 0.071033 | 0.104 | 0.071 | 1 |
| O75533 | SF3B1 | LPALMNEyRVPELNV | Y1136 | 0.211 | 90.3 | 1304 | 0.092563 | 0.211 | 0.0926 | 0.8333 |
| O75533 | SF3B1 | KDYIYAVtPLLEDAL | T1170 | 0.159 | 94.4 | 1304 | 0.108599 | 0.159 | 0.1086 | 1 |
| Q6ZWK6 | TMPRSS11F | EDDKSFYyLASFKVT | Y61 | 0.008 | 97.04 | 438 | 0.003509 | 0.008 | 0.0035 | 0.9286 |
| Q9NTI5 | PDS5B | KDIFMFItRQLKGLE | T111 | 0.027 | 97.18 | 1447 | 0.018441 | 0.027 | 0.0184 | 1 |
| Q9NTI5 | PDS5B | KNLNkQAyDLAkALL | Y215 | 0.171 | 96.92 | 1447 | 0.075015 | 0.337 | 0.1466 | 1 |
| Q9NTI5 | PDS5B | LAQIYKkyALQSAAG | Y419 | 0.042 | 96.93 | 1447 | 0.018425 | 0.042 | 0.0184 | 1 |
| Q9NTI5 | PDS5B | FSKVMVItRNLPDPG | T533 | 0.066 | 93.33 | 1447 | 0.045079 | 0.066 | 0.0451 | 1 |

| | | | | | | | | | | |
|---|---|---|---|---|---|---|---|---|---|---|
| Q5T7M9 | DIPK1A | TSAKNLGyNDKyDLK | Y302 | 0.071 | 93.8 | 428 | 0.031147 | 0.117 | 0.0513 | 1 |
| Q6DKI7 | PVRIG | ATELSsFtIRCGFLG | T59 | 0.126 | 89.39 | 326 | 0.08606 | 0.126 | 0.0861 | 0.875 |
| O60741 | HCN1 | GAVGKKMyFIQHGVA | Y513 | 0.088 | 97.65 | 890 | 0.038604 | 0.088 | 0.0386 | 1 |
| Q9HAW7 | UGT1A7 | PQTVLWRyTGTRPSN | Y334 | 0.166 | 95.89 | 530 | 0.072822 | 0.166 | 0.0728 | 1 |
| P23634 | ATP2B4 | LVKGIIDsTVGEHRQ | S771 | 0.079 | 84.89 | 1241 | 0.062798 | 0.079 | 0.0628 | 1 |
| Q9ULW6 | NAP1L2 | WNEGkNVtLktIKKK | T342 | 0.214 | 94.71 | 460 | 0.146165 | 0.214 | 0.1462 | 1 |
| Q5BKT4 | ALG10 | AGWSIADsLKSKSIF | S387 | 0.186 | 95.28 | 473 | 0.147854 | 0.186 | 0.1479 | 1 |
| Q8TCY9 | URGCP | RsRLVVLsTVGVPGT | S696 | 0.01 | 93.83 | 931 | 0.007949 | 0.01 | 0.0079 | 0.9091 |
| Q8N6P7 | IL22RA1 | LGGKQREyEFFGLTP | Y186 | 0.07 | 96.97 | 574 | 0.030708 | 0.07 | 0.0307 | 1 |
| Q32MH5 | ATOSA | PSGTIQVtLFNPNKT | T960 | 0.058 | 97.35 | 1076 | 0.039615 | 0.058 | 0.0396 | 1 |
| Q14324 | MYBPC2 | AVNAIGVsQPSMNTK | S722 | 0.086 | 94.85 | 1141 | 0.068363 | 0.124 | 0.0986 | 1 |
| Q14324 | MYBPC2 | GVNIAGRsEPATLAQ | S820 | 0.134 | 89.86 | 1141 | 0.106519 | 0.182 | 0.1447 | 1 |
| Q14324 | MYBPC2 | GNsEIMGyFVQKADK | Y967 | 0.109 | 94.74 | 1141 | 0.047817 | 0.198 | 0.0797 | 1 |
| Q14324 | MYBPC2 | TENICGLsDSPGVSK | S1015 | 0.048 | 95.32 | 1141 | 0.038156 | 0.048 | 0.0382 | 1 |
| O14976 | GAK | YALKRLLsNEEEkNR | S73 | 0.065 | 77.66 | 1311 | 0.051669 | 0.072 | 0.0779 | 1 |
| O14976 | GAK | IPPHDTQyTVFHsLI | Y285 | 0.151 | 95.02 | 1311 | 0.066242 | 0.151 | 0.0662 | 1 |
| Q9H239 | MMP28 | SGVLDRAtLRQMtRP | T82 | 0.006 | 93.05 | 520 | 0.004098 | 0.006 | 0.0041 | 1 |
| Q86UG4 | SLCO6A1 | FIAFIFstLIFSGFs | T588 | 0.097 | 94.57 | 719 | 0.066252 | 0.097 | 0.0663 | 1 |
| Q86UG4 | SLCO6A1 | tLIFSGFsGVPIVLA | S595 | 0.01 | 96.98 | 719 | 0.007949 | 0.01 | 0.0079 | 1 |
| Q86UG4 | SLCO6A1 | FKMSGETsCILRDVN | S642 | 0.012 | 91.25 | 719 | 0.009539 | 0.012 | 0.0095 | 0.875 |
| P36406 | TRIM23 | RPLWKHyyLNTQAVV | Y469 | 0.13 | 90.62 | 574 | 0.057029 | 0.13 | 0.057 | 1 |
| P57737 | CORO7 | SERQLLLyEAEALAG | Y712 | 0.003 | 98.66 | 925 | 0.001316 | 0.003 | 0.0013 | 0.9286 |
| P57737 | CORO7 | CLRLRQssLEPVAFR | S807 | 0.107 | 95.82 | 925 | 0.085056 | 0.107 | 0.0851 | 1 |
| P52790 | HK3 | EkMISGMyLGEIVRH | Y755 | 0.179 | 93.93 | 923 | 0.078525 | 0.179 | 0.0785 | 1 |
| P35270 | SPR | ASLLsPGsVLVLsAR | S35 | 0.007 | 97.02 | 261 | 0.005564 | 0.007 | 0.0056 | 1 |
| P35270 | SPR | PGsVLVLsARNDEAL | S40 | 0.014 | 98.78 | 261 | 0.011129 | 0.014 | 0.0111 | 1 |
| P35270 | SPR | QQLARETsVDPDMRK | S213 | 0.01 | 98.41 | 261 | 0.007949 | 0.01 | 0.0079 | 1 |
| P56192 | MARS1 | HIIHADIyRWFNISF | Y344 | 0.085 | 98.74 | 900 | 0.037288 | 0.159 | 0.0671 | 1 |
| P56192 | MARS1 | TPNAQFItRSWLRDG | T482 | 0.011 | 91.61 | 900 | 0.007513 | 0.011 | 0.0075 | 1 |
| P56192 | MARS1 | NPEQVDLyQFMAKDN | Y549 | 0.012 | 98.93 | 900 | 0.005264 | 0.012 | 0.0053 | 0.9231 |
| Q96L21 | RPL10L | PLGGHMVsDEyEQLs | S54 | 0.021 | 97.42 | 214 | 0.016693 | 0.021 | 0.0167 | 0.9091 |

| | | | | | | | | | | |
|---|---|---|---|---|---|---|---|---|---|---|
| P29320 | EPHA3 | CAKCPPHsSTQEDGS | S294 | 0.115 | 96.59 | 983 | 0.091415 | 0.115 | 0.0914 | 1 |
| P29320 | EPHA3 | FAAVSITtNQAAPSP | T432 | 0.047 | 88.06 | 983 | 0.032102 | 0.047 | 0.0321 | 1 |
| Q14289 | PTK2B | TLRKLIGsVDDLLPS | S923 | 0.177 | 94.33 | 1009 | 0.1407 | 0.177 | 0.1407 | 1 |
| O60841 | EIF5B | PDsDVAAtLKKQKKN | T775 | 0 | 89.73 | 1220 | 0 | 0 | 0 | 1 |
| O60841 | EIF5B | EATDILVskISRQSI | S1184 | 0 | 91.88 | 1220 | 0 | 0 | 0 | 1 |
| O94819 | KBTBD11 | PGAAQRAtDAVGPQL | T224 | 0.09 | 95.88 | 623 | 0.061471 | 0.09 | 0.0615 | 1 |
| P07195 | LDHB | HGSLFLQtPkIVADk | T75 | 0.036 | 98.12 | 334 | 0.024588 | 0.072 | 0.0492 | 0.8889 |
| P07195 | LDHB | EVAQLkksADtLWDI | S320 | 0.001 | 96.88 | 334 | 0.000795 | 0.001 | 0.0008 | 1 |
| Q8N4P6 | LRRC71 | DGLRLNRsLLWLsLA | S281 | 0.145 | 94.63 | 559 | 0.115263 | 0.145 | 0.1153 | 1 |
| O60443 | GSDME | __MFAkAtRNFLREV | T6 | 0.002 | 87.44 | 496 | 0.001366 | 0.002 | 0.0014 | 0.8182 |
| P51854 | TKTL1 | SQVKHKPtAVVAKTF | T210 | 0.002 | 98.77 | 596 | 0.001366 | 0.002 | 0.0014 | 0.9286 |
| O75165 | DNAJC13 | GkItGRRyNCyKHHW | Y114 | 0.118 | 95.8 | 2243 | 0.051765 | 0.118 | 0.0518 | 1 |
| O75165 | DNAJC13 | GQVRKYSsTERDsLL | S308 | 0.042 | 93.7 | 2243 | 0.033386 | 0.042 | 0.0334 | 1 |
| Q969Z0 | TBRG4 | KSQkDQNtFQKLLHI | T450 | 0.073 | 92.77 | 631 | 0.04986 | 0.073 | 0.0499 | 1 |
| P0C7M7 | ACSM4 | LTTYPITtLCSPPTV | T316 | 0 | 98.61 | 580 | 0 | 0 | 0 | 0.8182 |
| Q9UQC2 | GAB2 | DPDVLEyyKNDHSKk | Y49 | 0.215 | 93.96 | 676 | 0.094317 | 0.215 | 0.0943 | 0.9 |
| Q8IXL7 | MSRB3 | KRLTPLQyHVTQEKG | Y59 | 0.147 | 98.7 | 192 | 0.064487 | 0.147 | 0.0645 | 1 |
| Q9H2T7 | RANBP17 | DTYAPEItkAFITSR | T409 | 0.023 | 95.38 | 1088 | 0.015709 | 0.023 | 0.0157 | 0.9091 |
| Q9ULZ3 | PYCARD | TDKLVSFyLETYGAE | Y60 | 0.044 | 79.59 | 195 | 0.019302 | 0.31 | 0.136 | 0.8 |
| Q9UKF6 | CPSF3 | GPSVVMAsPGMMQSG | S328 | 0.112 | 89.28 | 684 | 0.08903 | 0.112 | 0.089 | 1 |
| O94808 | GFPT2 | CKDRGALtVGVTNTV | T444 | 0 | 98.71 | 682 | 0 | 0 | 0 | 1 |
| O94808 | GFPT2 | GRPIILCsKDDTESS | S622 | 0.033 | 95.45 | 682 | 0.026232 | 0.033 | 0.0262 | 0.875 |
| Q8N766 | EMC1 | tGRIIHssVQkKAKG | S773 | 0.112 | 95.87 | 993 | 0.08903 | 0.112 | 0.089 | 1 |
| Q8TDG2 | ACTRT1 | ASPDRCFsAWIGASI | S339 | 0.08 | 97.79 | 376 | 0.063593 | 0.08 | 0.0636 | 1 |
| P31948 | STIP1 | LdPTNMTyITNQAAV | Y261 | 0.014 | 96.18 | 543 | 0.006142 | 0.014 | 0.0061 | 1 |
| P31948 | STIP1 | YPQAMkHytEAIkRN | Y383 | 0.046 | 98.32 | 543 | 0.02018 | 0.046 | 0.0202 | 1 |
| Q9NW82 | WDR70 | VYYDPNksQRGAkLC | S504 | 0.017 | 93.72 | 654 | 0.013514 | 0.017 | 0.0135 | 0.8889 |
| O96024 | B3GALT4 | QRNAIRAsWGGLREA | S93 | 0.029 | 96.76 | 378 | 0.023053 | 0.04 | 0.0358 | 0.8571 |
| O96024 | B3GALT4 | CYGKFLLtSHRLDPW | T330 | 0.036 | 97.03 | 378 | 0.024588 | 0.036 | 0.0246 | 0.8462 |
| Q3MJ13 | WDR72 | LLYPHGLssKLDQSW | S474 | 0.023 | 80.54 | 1102 | 0.018283 | 0.023 | 0.0183 | 0.8333 |
| P22830 | FECH | VKLLDELsPNTAPHK | S151 | 0.067 | 96.31 | 423 | 0.053259 | 0.067 | 0.0533 | 1 |

| | | | | | | | | | | |
|---|---|---|---|---|---|---|---|---|---|---|
| P22830 | FECH | LNAIYRYyNQVGRKP | Y210 | 0.136 | 98.04 | 423 | 0.059661 | 0.136 | 0.0597 | 1 |
| P22830 | FECH | MPWLGPQtDEsIkGL | T315 | 0.002 | 98.67 | 423 | 0.001366 | 0.002 | 0.0014 | 1 |
| O75339 | CILP | QQHQAGEyFCKAQSD | Y374 | 0.056 | 94.93 | 1184 | 0.024566 | 0.056 | 0.0246 | 1 |
| O75339 | CILP | RLQKFVNttKVLPFN | T551 | 0.017 | 93.52 | 1184 | 0.011611 | 0.017 | 0.0116 | 1 |
| O75339 | CILP | LQKFVNttKVLPFNK | T552 | 0.055 | 94.22 | 1184 | 0.037566 | 0.055 | 0.0376 | 1 |
| O15392 | BIRC5 | FLEGCACtPERMAEA | T34 | 0.213 | 98.2 | 142 | 0.145482 | 0.213 | 0.1455 | 1 |
| Q9H6X2 | ANTXR1 | QLAHKFIsPQLRMSF | S75 | 0.045 | 89.38 | 564 | 0.035771 | 0.045 | 0.0358 | 1 |
| P03956 | MMP1 | TYRIENytPDLPRAD | T122 | 0.047 | 98.7 | 469 | 0.032102 | 0.047 | 0.0321 | 1 |
| P03956 | MMP1 | KLTFDAIttIRGEVM | T288 | 0.038 | 98.27 | 469 | 0.025955 | 0.038 | 0.026 | 1 |
| P05165 | PCCA | IkTVAIHsDVDASSV | S94 | 0 | 97.63 | 728 | 0 | 0 | 0 | 0.9167 |
| P05165 | PCCA | HPGyGFLsENkEFAR | S147 | 0.032 | 95.48 | 728 | 0.025437 | 0.032 | 0.0254 | 1 |
| P05165 | PCCA | WAVECRVyAEDPyKS | Y401 | 0.128 | 95.82 | 728 | 0.056152 | 0.246 | 0.1079 | 1 |
| P05165 | PCCA | MADALDNyVIRGVTH | Y473 | 0.01 | 98.49 | 728 | 0.004387 | 0.175 | 0.0768 | 1 |
| Q9UDR5 | AASS | DLRKVYGtVLSRHHH | T263 | 0.006 | 98.1 | 926 | 0.004098 | 0.006 | 0.0041 | 1 |
| Q6TGC4 | PADI6 | TVKMTSPsPsVDADK | S91 | 0.072 | 93.71 | 694 | 0.057234 | 0.072 | 0.0572 | 1 |
| Q9NZM1 | MYOF | LASTTQysRAVFDGN | S620 | 0.017 | 80.44 | 2061 | 0.013514 | 0.017 | 0.0135 | 0.8 |
| Q9NZM1 | MYOF | AHtKPVVtLTSYWED | T642 | 0.064 | 91.29 | 2061 | 0.043713 | 0.064 | 0.0437 | 1 |
| Q9NZM1 | MYOF | KDSFSDPyAHICFLH | Y1163 | 0.165 | 95.54 | 2061 | 0.072383 | 0.165 | 0.0724 | 0.9167 |
| Q2Q1W2 | TRIM71 | SGAFAPLtKATGDGL | T487 | 0 | 96.96 | 868 | 0 | 0 | 0 | 1 |
| Q00536 | CDK16 | LGEGtyAtVyKGKSK | T178 | 0.063 | 86.07 | 496 | 0.04303 | 0.063 | 0.043 | 1 |
| O15068 | MCF2L | QDGSPVItFPDYPAF | T96 | 0.039 | 96.28 | 1137 | 0.026638 | 0.039 | 0.0266 | 1 |
| O43390 | HNRNPR | ALLERTGyTLDVtTG | Y136 | 0.236 | 94.99 | 633 | 0.10353 | 0.236 | 0.1035 | 1 |
| O43390 | HNRNPR | GkIPRDLyEDELVPL | Y177 | 0.111 | 95.63 | 633 | 0.048694 | 0.111 | 0.0487 | 1 |
| O43390 | HNRNPR | rGFCFLEyEDHkSAA | Y296 | 0.037 | 93.58 | 633 | 0.016231 | 0.037 | 0.0162 | 1 |
| Q92547 | TOPBP1 | VTNTWLVtCIDYQTL | T618 | 0.178 | 90.63 | 1522 | 0.121576 | 0.178 | 0.1216 | 1 |
| P29401 | TKT | LNLRkIssDLDGHPV | S105 | 0.149 | 94.31 | 623 | 0.118442 | 0.149 | 0.1184 | 1 |
| P29401 | TKT | LQHQMDIyQkRCEAF | Y202 | 0.063 | 98.44 | 623 | 0.027637 | 0.063 | 0.0276 | 1 |
| P29401 | TKT | kIATRkAyGQALAkL | Y321 | 0.001 | 98.82 | 623 | 0.000439 | 0.001 | 0.0004 | 1 |
| P29401 | TKT | IIALDGDtkNstFsE | T342 | 0.081 | 98.1 | 623 | 0.055324 | 0.081 | 0.0553 | 1 |
| P29401 | TKT | DGDtkNstFsEIFkK | T346 | 0.007 | 98.27 | 623 | 0.004781 | 0.007 | 0.0048 | 1 |
| P29401 | TKT | DtkNstFsEIFkKEH | S348 | 0 | 98.63 | 623 | 0 | 0 | 0 | 1 |

| | | | | | | | | | | |
|---|---|---|---|---|---|---|---|---|---|---|
| P29401 | TKT | NRtVPFCsTFAAFFT | S387 | 0 | 98.83 | 623 | 0 | 0 | 0 | 0.9167 |
| P29401 | TKT | SNINLCGsHCGVSIG | S415 | 0.004 | 98.11 | 623 | 0.00318 | 0.004 | 0.0032 | 1 |
| P29401 | TKT | tSTVFYPsDGVATEk | S449 | 0 | 98.55 | 623 | 0 | 0 | 0 | 1 |
| P29401 | TKT | kGICFIRtSRPENAI | T472 | 0.003 | 98.68 | 623 | 0.002049 | 0.003 | 0.002 | 1 |
| P29401 | TKT | RPENAIIyNNNEDFQ | Y481 | 0.157 | 98.49 | 623 | 0.068874 | 0.157 | 0.0689 | 1 |
| P29401 | TKT | DRkLILDsARATKGR | S548 | 0.002 | 98.37 | 623 | 0.00159 | 0.002 | 0.0016 | 1 |
| Q58FF7 | HSP90AB3P | SKEQVANsAFVERVR | S370 | 0.154 | 95.86 | 597 | 0.122417 | 0.173 | 0.1375 | 1 |
| O95831 | AIFM1 | PYMRPPLsKELWFsD | S176 | 0.084 | 98.75 | 613 | 0.066773 | 0.084 | 0.0668 | 1 |
| O95831 | AIFM1 | GAEVksRtTLFRKIG | T281 | 0.214 | 98.78 | 613 | 0.146165 | 0.214 | 0.1462 | 1 |
| O95831 | AIFM1 | QDNPksAtEQsGtGI | T521 | 0 | 98.09 | 613 | 0 | 0 | 0 | 1 |
| O95831 | AIFM1 | sAtEQsGtGIRsEsE | T526 | 0.138 | 96.92 | 613 | 0.094256 | 0.138 | 0.0943 | 0.8333 |
| P36959 | GMPR | KLFyGMssDtAMNKH | S271 | 0.015 | 90.49 | 345 | 0.011924 | 0.015 | 0.0119 | 1 |
| P27361 | MAPK3 | GAYGMVSsAYDHVRK | S58 | 0.07 | 97.42 | 379 | 0.055644 | 0.07 | 0.0556 | 1 |
| P27361 | MAPK3 | EyVAtRWyrAPEIML | Y210 | 0.19 | 96.75 | 379 | 0.08335 | 0.31 | 0.1111 | 1 |
| Q9ULB5 | CDH7 | DREEQAYyTLRAQAL | Y118 | 0.307 | 93.17 | 785 | 0.134676 | 0.307 | 0.1347 | 1 |
| Q9ULB5 | CDH7 | DREAKDQyLLVIQAK | Y226 | 0.152 | 95.81 | 785 | 0.06668 | 0.152 | 0.0667 | 1 |
| Q9ULB5 | CDH7 | PRFPRRSyQYNVPES | Y267 | 0.312 | 96.4 | 785 | 0.13687 | 0.312 | 0.1369 | 1 |
| Q9ULB5 | CDH7 | ANsGVIttAKSLDRE | T436 | 0.031 | 95.7 | 785 | 0.021173 | 0.031 | 0.0212 | 1 |
| O00180 | KCNK1 | TAVVQRItVHVTRRP | T157 | 0.048 | 93.53 | 336 | 0.032785 | 0.048 | 0.0328 | 0.9091 |
| Q7Z4F1 | LRP10 | KLHLACGsERLTLRS | S82 | 0.116 | 86.32 | 713 | 0.09221 | 0.116 | 0.0922 | 1 |
| P51570 | GALK1 | PMALELMtVLVGsPR | T61 | 0.021 | 98.58 | 392 | 0.014343 | 0.021 | 0.0143 | 1 |
| O96028 | NSD2 | DKDHLLKyNVGDLVW | Y220 | 0.133 | 92.54 | 1365 | 0.058345 | 0.133 | 0.0583 | 0.8889 |
| Q14623 | IHH | PAARFRAtFAsHVQP | T293 | 0.151 | 89.3 | 411 | 0.103135 | 0.151 | 0.1031 | 1 |
| O00159 | MYO1C | FAVADTVyRALRTER | Y124 | 0.126 | 89.85 | 1063 | 0.055274 | 0.336 | 0.1474 | 1 |
| O00159 | MYO1C | RDQAVMIsGEsGAGK | S139 | 0.089 | 89.23 | 1063 | 0.070747 | 0.089 | 0.0707 | 1 |
| O00159 | MYO1C | GAGKtEAtKRLLQFY | T150 | 0.015 | 90.72 | 1063 | 0.010245 | 0.207 | 0.1414 | 1 |
| O00159 | MYO1C | FEQFCINyCNEKLQQ | Y440 | 0.059 | 91.12 | 1063 | 0.025882 | 0.059 | 0.0259 | 1 |
| O00159 | MYO1C | LQSkEPAyVRCIKPN | Y622 | 0.113 | 92.75 | 1063 | 0.049571 | 0.113 | 0.0496 | 1 |
| O00159 | MYO1C | LIRHQVkyLGLLENL | Y648 | 0.267 | 82.32 | 1063 | 0.117129 | 0.267 | 0.1171 | 1 |
| O00159 | MYO1C | RSRQLLLtPNAVVIV | T943 | 0 | 88.59 | 1063 | 0 | 0 | 0 | 1 |
| A8MTJ3 | GNAT3 | LLLGAGEsGkSTIVk | S44 | 0.027 | 97.55 | 354 | 0.021463 | 0.027 | 0.0215 | 1 |

| | | | | | | | | | | |
|---|---|---|---|---|---|---|---|---|---|---|
| Q9P0L2 | MARK1 | AKFRQIVsAVQyCHQ | S168 | 0.139 | 98.51 | 795 | 0.110493 | 0.139 | 0.1105 | 1 |
| Q92523 | CPT1B | CQAVIESsyQVAKAL | S533 | 0.068 | 98.67 | 772 | 0.054054 | 0.068 | 0.0541 | 1 |
| Q92523 | CPT1B | QAVIESsyQVAKALA | Y534 | 0.328 | 98.72 | 772 | 0.143889 | 0.328 | 0.1439 | 1 |
| Q9H972 | C14orf93 | LKGACVAyFLtKRRE | Y367 | 0.181 | 95.29 | 538 | 0.079402 | 0.181 | 0.0794 | 1 |
| Q9H972 | C14orf93 | DVTEELMsDEEDSLN | S428 | 0.042 | 95.39 | 538 | 0.033386 | 0.042 | 0.0334 | 1 |
| A6NMZ7 | COL6A6 | IRQMGGNtNtGAALN | T513 | 0.088 | 87.66 | 2263 | 0.060105 | 0.088 | 0.0601 | 1 |
| A6NMZ7 | COL6A6 | QMGGNtNtGAALNFT | T515 | 0.033 | 92.75 | 2263 | 0.022539 | 0.033 | 0.0225 | 1 |
| A6NMZ7 | COL6A6 | DIMFLVDsSGsIGPE | S629 | 0 | 92.33 | 2263 | 0 | 0 | 0 | 1 |
| A6NMZ7 | COL6A6 | RKFLILItDGEAQDI | T730 | 0.039 | 91.77 | 2263 | 0.026638 | 0.039 | 0.0266 | 1 |
| Q5JTZ9 | AARS2 | LRRAVRFsMEILKAP | S361 | 0 | 91.36 | 985 | 0 | 0 | 0 | 1 |
| Q01638 | IL1RL1 | MTEDAGDyTCKFIHN | Y179 | 0.018 | 95.71 | 556 | 0.007896 | 0.018 | 0.0079 | 1 |
| Q01638 | IL1RL1 | ANYsVtAtRSFTVKD | T197 | 0.099 | 94.86 | 556 | 0.067618 | 0.099 | 0.0676 | 1 |
| P33151 | CDH5 | DREKQARyEIVVEAR | Y223 | 0.184 | 96.11 | 784 | 0.080718 | 0.184 | 0.0807 | 1 |
| Q12788 | TBL3 | ACEkLEAtMLRLRRD | T702 | 0.122 | 81.07 | 808 | 0.083328 | 0.122 | 0.0833 | 1 |
| Q6IEE8 | SLFN12L | AEVENKGySYKKDGI | Y86 | 0.185 | 95.83 | 588 | 0.081157 | 0.185 | 0.0812 | 1 |
| P49459 | UBE2A | VsAIVEQsWRDC___ | S148 | 0.027 | 95.13 | 152 | 0.021463 | 0.076 | 0.0604 | 1 |
| Q8IV77 | CNGA4 | SFFLDLAsLMPTDVV | S121 | 0 | 94.11 | 575 | 0 | 0 | 0 | 1 |
| Q96F07 | CYFIP2 | PANVQPYyLYGSkPL | Y911 | 0.303 | 96.51 | 1278 | 0.132922 | 0.303 | 0.1329 | 1 |
| O60478 | GPR137B | HKRLsyQsVFLFLCL | S79 | 0.037 | 90.83 | 399 | 0.029412 | 0.037 | 0.0294 | 0.8889 |
| Q6P996 | PDXDC1 | PEykSMKsCLYVGMA | S572 | 0.121 | 78.38 | 788 | 0.096185 | 0.161 | 0.128 | 1 |
| Q14847 | LASP1 | kGyEKkPyCNAHyPk | Y52 | 0.269 | 93.2 | 261 | 0.118006 | 0.269 | 0.118 | 1 |
| O75334 | PPFIA2 | LERRREAsQHEIKDV | S1098 | 0.028 | 94.11 | 1257 | 0.022258 | 0.028 | 0.0223 | 1 |
| P50993 | ATP1A2 | ARGIVIAtGDRTVMG | T259 | 0.076 | 93.45 | 1020 | 0.051909 | 0.076 | 0.0519 | 1 |
| P50993 | ATP1A2 | EAVETLGsTSTICSD | S367 | 0.037 | 94.79 | 1020 | 0.029412 | 0.037 | 0.0294 | 1 |
| P50993 | ATP1A2 | LCFVGLMsMIDPPRA | S587 | 0 | 94.98 | 1020 | 0 | 0 | 0 | 1 |
| P50993 | ATP1A2 | GIkVIMVtGDHPItA | T614 | 0.016 | 92.15 | 1020 | 0.010928 | 0.016 | 0.0109 | 0.8571 |
| P50993 | ATP1A2 | IISEGNEtVEDIAAR | T637 | 0.012 | 91 | 1020 | 0.008196 | 0.012 | 0.0082 | 1 |
| P50993 | ATP1A2 | ICKTRRNsVFQQGMk | S940 | 0.077 | 95.99 | 1020 | 0.061208 | 0.077 | 0.0612 | 1 |
| P17987 | TCP1 | PTSVIsGyRLACkEA | Y121 | 0.003 | 96 | 556 | 0.001316 | 0.003 | 0.0013 | 1 |
| P17987 | TCP1 | HGRSQMEsMLISGYA | S208 | 0.071 | 92.33 | 556 | 0.056439 | 0.071 | 0.0564 | 0.9091 |
| P17987 | TCP1 | AIAEFArsLLVIPNT | S444 | 0 | 95.74 | 556 | 0 | 0 | 0 | 1 |

| | | | | | | | | | | |
|---|---|---|---|---|---|---|---|---|---|---|
| P17987 | TCP1 | PTIVkVKsLkFATEA | S513 | 0.076 | 94.76 | 556 | 0.060414 | 0.076 | 0.0604 | 1 |
| O95477 | ABCA1 | GGMQRKLsVALAFVG | S1042 | 0.083 | 87.58 | 2261 | 0.065978 | 0.083 | 0.066 | 1 |
| P18206 | VCL | kLEAMtNsKQsIAkk | S380 | 0.014 | 93.33 | 1134 | 0.011129 | 0.014 | 0.0111 | 1 |
| P18206 | VCL | LGEIsALtskLADLR | T442 | 0.081 | 93.18 | 1134 | 0.055324 | 0.081 | 0.0553 | 1 |
| P18206 | VCL | EAMTQEVsDVFsDTT | S596 | 0.017 | 92.67 | 1134 | 0.013514 | 0.017 | 0.0135 | 0.8889 |
| P18206 | VCL | AANFENHsGkLGAtA | S637 | 0.086 | 96.59 | 1134 | 0.068363 | 0.155 | 0.1232 | 1 |
| P18206 | VCL | tVEGIQAsVkTAREL | S664 | 0.029 | 94.91 | 1134 | 0.023053 | 0.029 | 0.0231 | 1 |
| P18206 | VCL | VkTARELtPQVVsAA | T672 | 0.107 | 95.66 | 1134 | 0.073082 | 0.185 | 0.1231 | 1 |
| P18206 | VCL | tksLLDAsEEAIkKD | S726 | 0.062 | 94.59 | 1134 | 0.049285 | 0.062 | 0.0493 | 1 |
| P18206 | VCL | LQksFLDsGyRILGA | S820 | 0.018 | 96.78 | 1134 | 0.014308 | 0.018 | 0.0143 | 1 |
| P37088 | SCNN1A | NLRPGVEtsIsMRKE | T375 | 0.023 | 97.31 | 669 | 0.015709 | 0.023 | 0.0157 | 1 |
| Q8WVX9 | FAR1 | _MVSIPEyYEGkNVL | Y7 | 0.158 | 98.6 | 515 | 0.069312 | 0.305 | 0.1311 | 1 |
| Q8WVX9 | FAR1 | EkIIAINsELtQPkL | S84 | 0.04 | 96.77 | 515 | 0.031797 | 0.04 | 0.0318 | 1 |
| Q8WVX9 | FAR1 | VQLNVIAtRQLILLA | T132 | 0.011 | 98.01 | 515 | 0.007513 | 0.011 | 0.0075 | 1 |
| P33076 | CIITA | RGCTLLLtARPRGRL | T546 | 0.009 | 90.35 | 1130 | 0.006147 | 0.009 | 0.0061 | 1 |
| P15924 | DSP | IAELCANsIKDyELQ | S957 | 0.081 | 89.63 | 2871 | 0.064388 | 0.162 | 0.1288 | 1 |
| Q9NYW0 | TAS2R10 | STIGFILtGLAIsRI | T48 | 0.173 | 96.07 | 307 | 0.118161 | 0.173 | 0.1182 | 1 |
| Q9NYW0 | TAS2R10 | ILtGLAIsRIFLIWI | S53 | 0.101 | 97 | 307 | 0.080286 | 0.101 | 0.0803 | 0.875 |
| Q8NBX0 | SCCPDH | KPNIkICtQVKGPEA | T362 | 0.012 | 98.78 | 429 | 0.008196 | 0.012 | 0.0082 | 1 |
| Q8NBX0 | SCCPDH | PkAGGVFtPGAAFSk | T400 | 0.05 | 98.72 | 429 | 0.034151 | 0.05 | 0.0342 | 0.8182 |
| Q4J6C6 | PREPL | TKDsRFLtINIMNKT | T266 | 0.005 | 96.97 | 727 | 0.003415 | 0.005 | 0.0034 | 1 |
| Q4J6C6 | PREPL | TEFKLMRtAADTPAI | T327 | 0.074 | 97.86 | 727 | 0.050543 | 0.074 | 0.0505 | 1 |
| Q4J6C6 | PREPL | NCPFQLCsPIRPPKY | S410 | 0.038 | 93.85 | 727 | 0.030207 | 0.038 | 0.0302 | 1 |
| Q4J6C6 | PREPL | KPLLVHVyGAYGMDL | Y479 | 0.285 | 98.07 | 727 | 0.125025 | 0.285 | 0.125 | 1 |
| Q16363 | LAMA4 | RFDIEVRtPADNGLI | T1081 | 0 | 87.52 | 1823 | 0 | 0 | 0 | 1 |
| Q16363 | LAMA4 | YHEISIIyHNDKKMI | Y1143 | 0.049 | 91.15 | 1823 | 0.021496 | 0.049 | 0.0215 | 1 |
| Q16363 | LAMA4 | SSVSPTRyELIVDKS | Y1324 | 0.133 | 89.23 | 1823 | 0.058345 | 0.133 | 0.0583 | 1 |
| Q9Y4B6 | DCAF1 | EDHyVEFskHSQDRV | S1185 | 0.007 | 97.14 | 1507 | 0.005564 | 0.007 | 0.0056 | 1 |
| Q9Y4B6 | DCAF1 | MDTVCRLyEVGRQRL | Y1387 | 0.085 | 94.59 | 1507 | 0.037288 | 0.17 | 0.0746 | 0.9333 |
| O14981 | BTAF1 | IIKPLMEtIKkEENT | T884 | 0.004 | 95.01 | 1849 | 0.002732 | 0.004 | 0.0027 | 1 |
| O14981 | BTAF1 | EYLNPLHyTGPPTER | Y1359 | 0.004 | 91.13 | 1849 | 0.001755 | 0.004 | 0.0018 | 1 |

| | | | | | | | | | | |
|---|---|---|---|---|---|---|---|---|---|---|
| O14981 | BTAF1 | FRNIKFNyCILDEGH | Y1400 | 0.039 | 92.79 | 1849 | 0.017109 | 0.039 | 0.0171 | 1 |
| O60264 | SMARCA5 | GFDKENVyDELrQCI | Y970 | 0.257 | 89.54 | 1052 | 0.112742 | 0.257 | 0.1127 | 0.8333 |
| P16410 | CTLA4 | IAsFVCEyASPGKAT | Y60 | 0.075 | 94.45 | 223 | 0.032901 | 0.075 | 0.0329 | 1 |
| Q9NZ01 | TECR | QWYPARQsLRLDPKG | S49 | 0.029 | 95.67 | 308 | 0.023053 | 0.029 | 0.0231 | 1 |
| Q9NZ01 | TECR | YIKRLLEtLFVHRFS | T145 | 0.035 | 95.91 | 308 | 0.023905 | 0.035 | 0.0239 | 1 |
| P08238 | HSP90AB1 | IPNPQERtLtLVDtG | T83 | 0.047 | 97.42 | 724 | 0.032102 | 0.047 | 0.0321 | 1 |
| P08238 | HSP90AB1 | NPQERtLtLVDtGIG | T85 | 0.129 | 97.88 | 724 | 0.088109 | 0.129 | 0.0881 | 1 |
| P08238 | HSP90AB1 | LkEDQtEyLEErRVK | Y192 | 0.059 | 96.07 | 724 | 0.025882 | 0.205 | 0.0899 | 1 |
| P08238 | HSP90AB1 | KEVVKkHsQFIGYPI | S206 | 0.039 | 87.49 | 724 | 0.031002 | 0.078 | 0.062 | 1 |
| P08238 | HSP90AB1 | DDItQEEyGEFyksL | Y301 | 0.106 | 96.53 | 724 | 0.046501 | 0.106 | 0.0465 | 1 |
| P08238 | HSP90AB1 | QEEyGEFyksLtNDW | Y305 | 0.029 | 95.15 | 724 | 0.012722 | 0.029 | 0.0127 | 1 |
| P08238 | HSP90AB1 | FIRGVVDsEDLPLNI | S383 | 0 | 89.43 | 724 | 0 | 0 | 0 | 1 |
| P08238 | HSP90AB1 | kkFyEAFskNLkLGI | S434 | 0.08 | 96.7 | 724 | 0.063593 | 0.097 | 0.0842 | 0.8 |
| P08238 | HSP90AB1 | stNRRRLsELLRyHt | S452 | 0.031 | 97.32 | 724 | 0.024642 | 0.031 | 0.0246 | 1 |
| P08238 | HSP90AB1 | RLsELLRyHtsQsGD | Y457 | 0.101 | 97.65 | 724 | 0.044307 | 0.101 | 0.0443 | 1 |
| P08238 | HSP90AB1 | sELLRyHtsQsGDEM | T459 | 0.063 | 97.46 | 724 | 0.04303 | 0.063 | 0.043 | 1 |
| P08238 | HSP90AB1 | sQsGDEMtSLSEYVS | T467 | 0.104 | 95.89 | 724 | 0.071033 | 0.164 | 0.1228 | 1 |
| P08238 | HSP90AB1 | kEtQksIyyItGEsk | Y484 | 0.044 | 97.46 | 724 | 0.019302 | 0.044 | 0.0193 | 1 |
| P08238 | HSP90AB1 | EtQksIyyItGEskE | Y485 | 0.229 | 97.47 | 724 | 0.100459 | 0.229 | 0.1005 | 1 |
| P08238 | HSP90AB1 | krGFEVVyMTEPIDE | Y512 | 0.102 | 97.73 | 724 | 0.044746 | 0.178 | 0.0717 | 1 |
| P08238 | HSP90AB1 | VtstyGWtANMERIM | T599 | 0.109 | 91.4 | 724 | 0.074448 | 0.109 | 0.0744 | 1 |
| P28330 | ACADL | KHFIPQMtAGKCIGA | T162 | 0.178 | 98.56 | 430 | 0.121576 | 0.178 | 0.1216 | 1 |
| O15455 | TLR3 | SLKKLELssNQIKEF | S205 | 0.112 | 98.47 | 904 | 0.08903 | 0.112 | 0.089 | 1 |
| P42765 | ACAA2 | VLCGGtEsMSQAPyC | S121 | 0 | 98.31 | 397 | 0 | 0 | 0 | 0.9091 |
| Q9Y4F1 | FARP1 | IGQTPAEsDFQLLEI | S211 | 0 | 96.51 | 1045 | 0 | 0 | 0 | 1 |
| Q86VP6 | CAND1 | YCIQAFEsFVRRCPK | S281 | 0.073 | 90.11 | 1230 | 0.058029 | 0.073 | 0.058 | 0.8 |
| O00231 | PSMD11 | EARLVSLyFDTkRYQ | Y137 | 0.213 | 89.86 | 422 | 0.09344 | 0.213 | 0.0934 | 1 |
| O00231 | PSMD11 | QAtLDMQsGIIHAAE | S214 | 0.006 | 92.48 | 422 | 0.004769 | 0.006 | 0.0048 | 1 |
| Q6IN85 | PPP4R3A | AQLtDEAtDEEKRQE | T323 | 0.202 | 92.14 | 833 | 0.137969 | 0.202 | 0.138 | 1 |
| Q8IVI9 | NOSTRIN | AKKKLMVstKKHEAL | S143 | 0.135 | 96.2 | 506 | 0.107313 | 0.135 | 0.1073 | 1 |
| Q8IVI9 | NOSTRIN | LLNKLtKstEKLEKE | S176 | 0.174 | 96.18 | 506 | 0.138315 | 0.174 | 0.1383 | 1 |

| | | | | | | | | | | |
|---|---|---|---|---|---|---|---|---|---|---|
| P13637 | ATP1A3 | EAVETLGsTSTICSD | S359 | 0.032 | 95.35 | 1013 | 0.025437 | 0.032 | 0.0254 | 1 |
| P13637 | ATP1A3 | PFNSTNKyQLsIHEt | Y478 | 0.175 | 92.14 | 1013 | 0.07677 | 0.253 | 0.0873 | 1 |
| P13637 | ATP1A3 | STNKyQLsIHEtEDP | S481 | 0.01 | 94.57 | 1013 | 0.007949 | 0.01 | 0.0079 | 1 |
| P13637 | ATP1A3 | GIkVIMVtGDHPItA | T607 | 0.02 | 92.43 | 1013 | 0.01366 | 0.02 | 0.0137 | 1 |
| P13637 | ATP1A3 | IISEGNEtVEDIAAR | T630 | 0.017 | 91.95 | 1013 | 0.011611 | 0.017 | 0.0116 | 1 |
| P13637 | ATP1A3 | TGDGVNDsPALkkAD | S712 | 0.002 | 86.38 | 1013 | 0.00159 | 0.002 | 0.0016 | 1 |
| P13637 | ATP1A3 | ICKTRRNsVFQQGMk | S933 | 0.077 | 96.11 | 1013 | 0.061208 | 0.152 | 0.1208 | 1 |
| P13637 | ATP1A3 | GGWVEKEtYY_____ | T1011 | 0.088 | 92.5 | 1013 | 0.060105 | 0.088 | 0.0601 | 1 |
| Q9UHI8 | ADAMTS1 | CGGGVQytMRECDNP | T582 | 0.039 | 94.98 | 967 | 0.026638 | 0.039 | 0.0266 | 0.875 |
| Q8NCT3 | MATCAP2 | MLRHEIGtHYFRGIN | T335 | 0.005 | 91.31 | 520 | 0.003415 | 0.005 | 0.0034 | 0.8 |
| Q96PN6 | ADCY10 | WDNFYKFsNRAKNLL | S1443 | 0.102 | 92.78 | 1610 | 0.081081 | 0.069 | 0.1029 | 0.875 |
| Q96PN6 | ADCY10 | RRTMTLTyYDGISRY | Y1459 | 0.073 | 88.31 | 1610 | 0.032024 | 0.073 | 0.032 | 0.9333 |
| P54802 | NAGLU | AWRLLLRsVYNCSGE | S500 | 0.02 | 98.89 | 743 | 0.015898 | 0.02 | 0.0159 | 1 |
| P54802 | NAGLU | SAPSLATsPAFRYDL | S553 | 0.16 | 98.56 | 743 | 0.127186 | 0.16 | 0.1272 | 1 |
| P54802 | NAGLU | YEEARSAyLSkELAs | Y583 | 0.281 | 98.71 | 743 | 0.123271 | 0.281 | 0.1233 | 1 |
| P54802 | NAGLU | FVLSKQRyPsQPRGD | Y715 | 0.211 | 98.8 | 743 | 0.092563 | 0.211 | 0.0926 | 1 |
| P40394 | ADH7 | VGAtECIsPKDSTKP | S254 | 0.066 | 97.87 | 386 | 0.052464 | 0.066 | 0.0525 | 1 |
| P46020 | PHKA1 | LPHMWGQsLyILGSL | S405 | 0 | 98.28 | 1223 | 0 | 0 | 0 | 1 |
| P46020 | PHKA1 | VLGTSKLyDIRKTIF | Y511 | 0.007 | 95.4 | 1223 | 0.003071 | 0.007 | 0.0031 | 0.9333 |
| P08590 | MYL3 | DsNGCINyEAFVKHI | Y185 | 0.118 | 96.8 | 195 | 0.051765 | 0.118 | 0.0518 | 1 |
| Q99666 | RGPD5 | GFYFAkLyYEAkEyD | Y36 | 0.089 | 93.11 | 1765 | 0.039043 | 0.089 | 0.039 | 1 |
| Q9Y2G9 | SBNO2 | AGVGkGRtVAGVILE | T287 | 0.004 | 95.95 | 1366 | 0.002732 | 0.004 | 0.0027 | 1 |
| Q9Y2G9 | SBNO2 | DMKVSGMyIARQLSF | Y476 | 0.222 | 92.12 | 1366 | 0.097388 | 0.222 | 0.0974 | 1 |
| P50440 | GATM | YRSIIKDyFHRGAKW | Y203 | 0.003 | 98.42 | 423 | 0.001316 | 0.003 | 0.0013 | 1 |
| P50440 | GATM | DELYNQDyPIHsVED | Y227 | 0.132 | 97.51 | 423 | 0.057907 | 0.132 | 0.0579 | 1 |
| P50440 | GATM | RRHLAPDyRVHIISF | Y289 | 0.307 | 98.69 | 423 | 0.134676 | 0.307 | 0.1347 | 1 |
| P49721 | PSMB2 | ILNLPTFsVRIIDkN | S179 | 0.089 | 98.54 | 201 | 0.070747 | 0.169 | 0.1375 | 1 |
| P43403 | ZAP70 | FRKFSSRsDVWSYGV | S520 | 0.024 | 97.23 | 619 | 0.019078 | 0.024 | 0.0191 | 1 |
| P43403 | ZAP70 | VTMWEALsyGQkPYK | S534 | 0.05 | 97.61 | 619 | 0.039746 | 0.114 | 0.0999 | 1 |
| P31946 | YWHAB | AAAMkAVtEQGHELs | T32 | 0.013 | 98.24 | 246 | 0.008879 | 0.013 | 0.0089 | 1 |
| P31946 | YWHAB | GDyFRyLsEVAsGDN | S132 | 0.021 | 97.53 | 246 | 0.016693 | 0.021 | 0.0167 | 0.9167 |

| | | | | | | | | | | |
|---|---|---|---|---|---|---|---|---|---|---|
| P31946 | YWHAB | VSNSQQAyQEAFEIS | Y151 | 0 | 98.86 | 246 | 0 | 0 | 0 | 0.9333 |
| P04440 | HLA-DPB1 | TQRFLERyIyNREEF | Y57 | 0.311 | 98.03 | 258 | 0.136431 | 0.311 | 0.1364 | 1 |
| Q9UKF6 | CPSF3 | NFNyHILsPCDLSNy | S496 | 0.146 | 87.28 | 684 | 0.116058 | 0.146 | 0.1161 | 1 |
| Q12840 | KIF5A | MNEHSSRsHSIFLIN | S205 | 0.004 | 90.03 | 1032 | 0.00318 | 0.06 | 0.0477 | 1 |
| Q12840 | KIF5A | SLGGNCRttMFICCS | T298 | 0.06 | 95.79 | 1032 | 0.040981 | 0.06 | 0.041 | 1 |
| Q12840 | KIF5A | LGGNCRttMFICCSP | T299 | 0.113 | 95.92 | 1032 | 0.077181 | 0.113 | 0.0772 | 1 |
| Q8N8D1 | PDCD7 | VRKKQADtKRMVDIL | T312 | 0.085 | 96.9 | 485 | 0.058056 | 0.085 | 0.0581 | 1 |
| P13861 | PRKAR2A | GDDGDNFyVIERGTY | Y177 | 0.131 | 97.85 | 404 | 0.057468 | 0.131 | 0.0575 | 1 |
| P13861 | PRKAR2A | tNkPRAAsAyAVGDV | S350 | 0.051 | 96.47 | 404 | 0.040541 | 0.051 | 0.0405 | 1 |
| Q9HD67 | MYO10 | RYKRNQIyTYIGSIL | Y93 | 0.073 | 96.85 | 2058 | 0.032024 | 0.073 | 0.032 | 1 |
| Q9HD67 | MYO10 | kLILKFLsVIsQQSL | S175 | 0.049 | 90.21 | 2058 | 0.038951 | 0.049 | 0.039 | 1 |
| Q9HD67 | MYO10 | LKFLsVIsQQSLELS | S178 | 0.005 | 86.49 | 2058 | 0.003975 | 0.005 | 0.004 | 1 |
| Q9HD67 | MYO10 | yLSTPENyHYLNQSG | Y286 | 0.041 | 95.67 | 2058 | 0.017986 | 0.041 | 0.018 | 1 |
| Q9HD67 | MYO10 | NVQQAVDsRDSLAMA | S397 | 0.051 | 90.1 | 2058 | 0.040541 | 0.051 | 0.0405 | 1 |
| Q9HD67 | MYO10 | HyAGEVQyDVRGILE | Y555 | 0.005 | 95.37 | 2058 | 0.002193 | 0.005 | 0.0022 | 1 |
| P54709 | ATP1B3 | SRSDPtsyAGyIEDL | Y103 | 0.148 | 91.88 | 279 | 0.064925 | 0.148 | 0.0649 | 1 |
| P54709 | ATP1B3 | DPtsyAGyIEDLkkF | Y106 | 0.308 | 93.85 | 279 | 0.135115 | 0.308 | 0.1351 | 1 |
| P61088 | UBE2N | IEtArAWtrLyAMNN | T144 | 0.036 | 98.67 | 152 | 0.024588 | 0.036 | 0.0246 | 1 |
| Q8IZP7 | HS6ST3 | DLSLVGCyNLTFMNE | Y323 | 0.209 | 97.27 | 471 | 0.091685 | 0.209 | 0.0917 | 0.9167 |
| Q5JTW2 | CEP78 | DWAPLLstLKINkDL | T61 | 0.039 | 86.03 | 689 | 0.026638 | 0.039 | 0.0266 | 1 |
| Q5JTW2 | CEP78 | LAKGLNKsAsLVHLS | S147 | 0.104 | 90.04 | 689 | 0.082671 | 0.104 | 0.0827 | 0.875 |
| P51878 | CASP5 | RLLQGLGytVVDEKN | Y228 | 0.078 | 98.42 | 434 | 0.034217 | 0.078 | 0.0342 | 0.8333 |
| Q9BXP8 | PAPPA2 | PDSPKRAyMSVKELK | Y666 | 0.036 | 88.56 | 1791 | 0.015793 | 0.036 | 0.0158 | 0.9412 |
| Q7Z4J2 | GLT6D1 | GPLVHVKsLGEHIAS | S153 | 0.135 | 92.99 | 276 | 0.107313 | 0.135 | 0.1073 | 0.875 |
| Q969R5 | L3MBTL2 | AVLPsRVyWIASVIQ | Y230 | 0.151 | 96.74 | 705 | 0.066242 | 0.151 | 0.0662 | 1 |
| Q9Y223 | GNE | ATCNRADySkLAPIM | Y22 | 0.186 | 97.95 | 722 | 0.081596 | 0.186 | 0.0816 | 1 |
| Q7RTP6 | MICAL3 | KtHPVsEyEFEVIIG | Y214 | 0.323 | 94.92 | 2002 | 0.141695 | 0.323 | 0.1417 | 1 |
| Q7RTP6 | MICAL3 | VSKNFsQysIDPVTR | Y463 | 0.308 | 98.04 | 2002 | 0.135115 | 0.308 | 0.1351 | 1 |
| Q7RTP6 | MICAL3 | SKNFsQysIDPVTRY | S464 | 0.06 | 98.18 | 2002 | 0.047695 | 0.06 | 0.0477 | 1 |
| Q96QV1 | HHIP | SLAFHPNyKKNGKLY | Y293 | 0.11 | 97.88 | 700 | 0.048255 | 0.11 | 0.0483 | 1 |
| Q15021 | NCAPD2 | yNHMLsAtVKIIQML | T241 | 0.075 | 95.65 | 1401 | 0.051226 | 0.075 | 0.0512 | 0.875 |

| | | | | | | | | | | |
|---|---|---|---|---|---|---|---|---|---|---|
| P36952 | SERPINB5 | QVGAKGDtANEIGQV | T50 | 0.018 | 98.5 | 375 | 0.012294 | 0.018 | 0.0123 | 1 |
| P36952 | SERPINB5 | LKLIKRLyVDKsLNL | Y93 | 0.026 | 98.56 | 375 | 0.011406 | 0.026 | 0.0114 | 1 |
| P36952 | SERPINB5 | DNSVNDQtKILVVNA | T157 | 0.07 | 96.83 | 375 | 0.047811 | 0.122 | 0.0833 | 1 |
| P36952 | SERPINB5 | KDVEDEstGLEkIEk | T241 | 0.048 | 88.44 | 375 | 0.032785 | 0.048 | 0.0328 | 1 |
| P36952 | SERPINB5 | NADHPFIyIIRHNKT | Y356 | 0.005 | 98.69 | 375 | 0.002193 | 0.005 | 0.0022 | 1 |
| Q9UKU0 | ACSL6 | INtADIstVIVDKPQ | T197 | 0.031 | 98.67 | 697 | 0.021173 | 0.031 | 0.0212 | 1 |
| Q9UKU0 | ACSL6 | GKWLPAGtLkIIDRK | T549 | 0.032 | 97.04 | 697 | 0.021856 | 0.032 | 0.0219 | 1 |
| Q9Y6M0 | PRSS21 | NIYLSPRyLGNSPyD | Y130 | 0.248 | 95.38 | 314 | 0.108794 | 0.248 | 0.1088 | 1 |
| P19174 | PLCG1 | PDGMPVIyHGHTLTT | Y379 | 0.306 | 95.71 | 1290 | 0.134238 | 0.306 | 0.1342 | 0.8182 |
| P19174 | PLCG1 | ISADGLPsPNQLkRK | S451 | 0.05 | 98 | 1290 | 0.039746 | 0.05 | 0.0397 | 1 |
| P19174 | PLCG1 | KKIALELsELVVyCR | S954 | 0.044 | 89.04 | 1290 | 0.034976 | 0.044 | 0.035 | 1 |
| P19174 | PLCG1 | RACyRDMssFPETkA | S981 | 0.042 | 97.22 | 1290 | 0.033386 | 0.042 | 0.0334 | 1 |
| P26374 | CHML | YEQHPDEyQAFRQCS | Y314 | 0.195 | 93.59 | 656 | 0.085544 | 0.195 | 0.0855 | 1 |
| Q92519 | TRIB2 | DKHGCPAyVSPEILN | Y218 | 0.091 | 95.72 | 343 | 0.03992 | 0.091 | 0.0399 | 1 |
| Q92519 | TRIB2 | EPSERLtsQEILDHP | S299 | 0.059 | 98.27 | 343 | 0.0469 | 0.059 | 0.0469 | 1 |
| Q7Z4Q2 | HEATR3 | SPVSSMEsLLLKTLV | S246 | 0.065 | 96.55 | 680 | 0.051669 | 0.065 | 0.0517 | 1 |
| Q99758 | ABCA3 | PLIIKELsKVYEQRV | S1387 | 0 | 87.97 | 1704 | 0 | -0.003 | 0.0123 | 1 |
| Q99758 | ABCA3 | ANKLVRtysGGNKRK | Y1515 | 0.161 | 79.5 | 1704 | 0.070628 | 0.28 | 0.1126 | 1 |
| O60303 | KATNIP | LNGIEIFssKGEPVQ | S1000 | 0.154 | 93.7 | 1618 | 0.122417 | 0.154 | 0.1224 | 1 |
| Q8NFG4 | FLCN | CVRIIPYsSQYEEAY | S406 | 0.141 | 96.12 | 579 | 0.112083 | 0.141 | 0.1121 | 1 |
| Q9UJY1 | HSPB8 | kDGyVEVsGKHEEkQ | S122 | 0.175 | 98.08 | 196 | 0.13911 | 0.175 | 0.1391 | 0.8571 |
| P05177 | CYP1A2 | FLWFLQKtVQEHyQD | T267 | 0.038 | 95.32 | 516 | 0.025955 | 0.038 | 0.026 | 1 |
| Q8NEB9 | PIK3C3 | PAQLFFKtEDGGkyP | T625 | 0.164 | 95.06 | 887 | 0.112014 | 0.164 | 0.112 | 1 |
| Q8NEB9 | PIK3C3 | ENLDLkLtPYKVLAT | T668 | 0.184 | 97.29 | 887 | 0.125675 | 0.184 | 0.1257 | 1 |
| Q96JB5 | CDK5RAP3 | WQEIIALyEKDNTyL | Y104 | 0 | 96.38 | 506 | 0 | 0 | 0 | 1 |
| Q96JB5 | CDK5RAP3 | LyEKDNTyLVELssL | Y110 | 0.193 | 95.26 | 506 | 0.084666 | 0.193 | 0.0847 | 1 |
| Q96JB5 | CDK5RAP3 | NTyLVELssLLVRNV | S115 | 0 | 94.96 | 506 | 0 | 0 | 0 | 1 |
| Q96JB5 | CDK5RAP3 | KrGNSTVyEWRTGTE | Y239 | 0.188 | 95.32 | 506 | 0.082473 | 0.188 | 0.0825 | 1 |
| A1L390 | PLEKHG3 | LRSIVEDyLLKIIDT | Y119 | 0.025 | 91.24 | 1219 | 0.010967 | 0.025 | 0.011 | 1 |
| A1L390 | PLEKHG3 | VEEKRNWtHHIKRLI | T386 | 0.042 | 92.9 | 1219 | 0.028687 | 0.07 | 0.0478 | 1 |
| Q9H5Z1 | DHX35 | ITISGIVyVIDCGFV | Y356 | 0.11 | 97.22 | 703 | 0.048255 | 0.11 | 0.0483 | 1 |

| | | | | | | | | | | |
|---|---|---|---|---|---|---|---|---|---|---|
| Q9H5Z1 | DHX35 | RFHSTGAyRTIRDDH | Y627 | 0.096 | 95.32 | 703 | 0.042114 | 0.096 | 0.0421 | 1 |
| Q5T1V6 | DDX59 | LPPQLLNsPYLHDQK | S578 | 0.058 | 93.51 | 619 | 0.046105 | 0.058 | 0.0461 | 0.875 |
| O00410 | IPO5 | RkQAEETyENIPGQS | Y33 | 0.176 | 91.17 | 1097 | 0.077209 | 0.274 | 0.0924 | 1 |
| O00410 | IPO5 | ssAFDEVyPALPsDV | Y78 | 0.056 | 91.7 | 1097 | 0.024566 | 0.106 | 0.045 | 1 |
| O00410 | IPO5 | NDDSVLksLVEIADT | S239 | 0.078 | 96.44 | 1097 | 0.062003 | 0.078 | 0.062 | 1 |
| O60312 | ATP10A | ERLHDQVsVELPHLG | S723 | 0.041 | 91.22 | 1499 | 0.032592 | 0.058 | 0.0547 | 1 |
| O60312 | ATP10A | DSVRKRMsVVIRHPL | S750 | 0.01 | 94.99 | 1499 | 0.007949 | 0.01 | 0.0079 | 1 |
| O60312 | ATP10A | TQNyLNVyAAEGLRT | Y803 | 0 | 87.68 | 1499 | 0 | 0 | 0 | 1 |
| O60312 | ATP10A | GMQAVMAsDFAVPKF | S1060 | 0.028 | 86.34 | 1499 | 0.022258 | 0.109 | 0.0866 | 1 |
| O43396 | TXNL1 | APAFSsMsNKYPQAV | S48 | 0.009 | 97.66 | 289 | 0.007154 | 0.009 | 0.0072 | 1 |
| Q96BW5 | PTER | KYGGHGysHILtNVV | S314 | 0.024 | 98.6 | 349 | 0.019078 | 0.024 | 0.0191 | 0.8889 |
| P78413 | IRX4 | KNATREttSTLKAWL | T154 | 0.174 | 93.4 | 519 | 0.118844 | 0.174 | 0.1188 | 0.8571 |
| P82914 | MRPS15 | EARIIALsVKIRSYE | S133 | 0.06 | 96.11 | 257 | 0.047695 | 0.06 | 0.0477 | 1 |
| Q9NTJ4 | MAN2C1 | RLKRLsNtDGLPRVQ | T482 | 0.003 | 95.92 | 1040 | 0.002049 | 0.003 | 0.002 | 1 |
| Q9NTJ4 | MAN2C1 | GLPRVQLssPRQLFS | S491 | 0.006 | 98.36 | 1040 | 0.004769 | 0.006 | 0.0048 | 1 |
| Q9NTJ4 | MAN2C1 | IRVKLDPtGRLTSLV | T701 | 0.18 | 94.55 | 1040 | 0.122942 | 0.18 | 0.1229 | 1 |
| P17655 | CAPN2 | TNRARERsDtFINLR | S462 | 0.041 | 92.13 | 700 | 0.032592 | 0.041 | 0.0326 | 1 |
| P17655 | CAPN2 | IQKYQKIyREIDVDR | Y611 | 0.014 | 89.45 | 700 | 0.006142 | 0.014 | 0.0061 | 1 |
| O43795 | MYO1B | GAGktEAsKLVMSYV | S118 | 0.047 | 90.47 | 1136 | 0.037361 | 0.047 | 0.0374 | 0.8889 |
| O43795 | MYO1B | RDFSRYNyLSLDsAk | Y234 | 0.038 | 95.38 | 1136 | 0.01667 | 0.038 | 0.0167 | 0.8125 |
| O43795 | MYO1B | YNyLSLDsAkVNGVD | S239 | 0.067 | 81.11 | 1136 | 0.053259 | 0.067 | 0.0533 | 1 |
| O43795 | MYO1B | SVLERAFsFRTVEAK | S325 | 0.085 | 90.57 | 1136 | 0.067568 | 0.085 | 0.0676 | 1 |
| O43795 | MYO1B | INLkRPPtAGsQFkA | T569 | 0.064 | 85.78 | 1136 | 0.043713 | 0.064 | 0.0437 | 0.8182 |
| Q53R41 | FASTKD1 | TQDLSSLsVLMVNIS | S194 | 0.018 | 94.82 | 847 | 0.014308 | 0.018 | 0.0143 | 1 |
| Q15047 | SETDB1 | KMGFHVIyKTPCGLC | Y627 | 0 | 95.93 | 1291 | 0 | 0 | 0 | 1 |
| Q15047 | SETDB1 | ECLPTGVyECNKRCk | Y779 | 0.297 | 96.97 | 1291 | 0.13029 | 0.297 | 0.1303 | 1 |
| Q08209 | PPP3CA | RYLFLGDyVDRGYFS | Y119 | 0 | 98.9 | 521 | 0 | 0 | 0 | 1 |
| Q14118 | DAG1 | LHRMRsFsEVELHNM | S218 | 0.15 | 91.62 | 895 | 0.119237 | 0.15 | 0.1192 | 0.8571 |
| Q9Y2E5 | MAN2B2 | RAFVVPHsHMDVGWV | S35 | 0.082 | 97.34 | 1009 | 0.065183 | 0.082 | 0.0652 | 1 |
| Q9Y2E5 | MAN2B2 | FVWRGSPsLSERQEI | S199 | 0.142 | 96.25 | 1009 | 0.112878 | 0.165 | 0.1312 | 0.8333 |
| Q9Y2E5 | MAN2B2 | TPSHIPFsNRSGFYW | S225 | 0.119 | 92.34 | 1009 | 0.094595 | 0.156 | 0.124 | 1 |

| | | | | | | | | | | |
|---|---|---|---|---|---|---|---|---|---|---|
| P13995 | MTHFD2 | RQKAGYItPVPGGVG | T306 | 0.018 | 98.76 | 350 | 0.012294 | 0.018 | 0.0123 | 1 |
| P13995 | MTHFD2 | VAMLMKNtIIAAkKV | T324 | 0 | 98.5 | 350 | 0 | 0 | 0 | 1 |
| P67775 | PPP2CA | DRGGWGIsPRGAGYT | S212 | 0.122 | 97.69 | 309 | 0.09698 | 0.122 | 0.097 | 1 |
| O14936 | CASK | LPAFkRKtLVLLGAH | T741 | 0.015 | 96.7 | 926 | 0.010245 | 0.015 | 0.0102 | 1 |
| P16885 | PLCG2 | RKREGsDsyAItFRA | S679 | 0.081 | 85.77 | 1265 | 0.064388 | 0.081 | 0.0644 | 1 |
| P16885 | PLCG2 | KREGsDsyAItFRAR | Y680 | 0.075 | 92.74 | 1265 | 0.032901 | 0.075 | 0.0329 | 1 |
| P16885 | PLCG2 | GsDsyAItFRARGKV | T683 | 0 | 93.17 | 1265 | 0 | 0 | 0 | 1 |
| Q6P2Q9 | PRPF8 | KGYLPSHyERVQMLL | Y2256 | 0.267 | 86.61 | 2335 | 0.117129 | 0.301 | 0.132 | 0.9091 |
| P02675 | FGB | GGETSEMyLIQPDss | Y255 | 0.041 | 98.05 | 491 | 0.017986 | 0.041 | 0.018 | 1 |
| P02675 | FGB | PDssVKPyRVyCDMN | Y266 | 0.209 | 97.66 | 491 | 0.091685 | 0.209 | 0.0917 | 1 |
| P02675 | FGB | sVKPyRVyCDMNTEN | Y269 | 0.176 | 98.55 | 491 | 0.077209 | 0.176 | 0.0772 | 1 |
| P02675 | FGB | NTENGGWtVIQNRQD | T280 | 0.016 | 98.67 | 491 | 0.010928 | 0.016 | 0.0109 | 1 |
| P02675 | FGB | FGRKWDPykQGFGNV | Y299 | 0.001 | 98.8 | 491 | 0.000439 | 0.001 | 0.0004 | 1 |
| P02675 | FGB | NDKISQLtRMGPtEL | T333 | 0.032 | 97.05 | 491 | 0.021856 | 0.032 | 0.0219 | 1 |
| P02675 | FGB | QLtRMGPtELLIEME | T338 | 0.103 | 97.93 | 491 | 0.07035 | 0.103 | 0.0703 | 1 |
| P02675 | FGB | GDKVKAHyGGFTVQN | Y356 | 0 | 98.26 | 491 | 0 | 0 | 0 | 1 |
| P02675 | FGB | VQNEANKyQIsVNKY | Y368 | 0.001 | 98.72 | 491 | 0.000439 | 0.001 | 0.0004 | 1 |
| P02675 | FGB | SDPRKQCsKEDGGGW | S425 | 0.061 | 97.86 | 491 | 0.04849 | 0.061 | 0.0485 | 1 |
| P02675 | FGB | RYYWGGQyTWDMAKH | Y452 | 0.05 | 98.15 | 491 | 0.021934 | 0.05 | 0.0219 | 1 |
| P02675 | FGB | MNWkGSWySMRKMSM | Y475 | 0.286 | 98.52 | 491 | 0.125464 | 0.286 | 0.1255 | 1 |
| Q02978 | SLC25A11 | PADQRRGykNVFNAL | Y161 | 0.251 | 91.57 | 314 | 0.11011 | 0.251 | 0.1101 | 1 |
| Q9UKM9 | RALY | kGyAFVQysNERHAR | Y62 | 0.049 | 94.23 | 306 | 0.021496 | 0.049 | 0.0215 | 1 |
| P50579 | METAP2 | PKGQECEyPPTQDGR | Y137 | 0.121 | 98.47 | 478 | 0.053081 | 0.121 | 0.0531 | 0.9333 |
| P50579 | METAP2 | GEVYAIEtFGStGKG | T365 | 0 | 98.88 | 478 | 0 | 0 | 0 | 0.9091 |
| Q68CR1 | SEL1L3 | GQGSERLsSMNLGYk | S608 | 0 | 98.36 | 1132 | 0 | 0 | 0 | 1 |
| Q01740 | FMO1 | AGVSGLAsIKCCLEE | S17 | 0 | 98.93 | 532 | 0 | 0 | 0 | 1 |
| Q01740 | FMO1 | FKGQyFHsRQYKHPD | S173 | 0.084 | 98.56 | 532 | 0.066773 | 0.084 | 0.0668 | 1 |
| Q01740 | FMO1 | PAHLQKPtLAIIGLI | T365 | 0.129 | 98.04 | 532 | 0.088109 | 0.129 | 0.0881 | 1 |
| O75845 | SC5D | LWDRIGGsFkNPSSF | S253 | 0.024 | 98.29 | 299 | 0.019078 | 0.024 | 0.0191 | 1 |
| P50914 | RPL14 | VEVGRVAyVSFGPHA | Y14 | 0.272 | 97.65 | 215 | 0.119322 | 0.272 | 0.1193 | 1 |
| P25089 | FPR3 | ILPFRMVsVAMREKW | S84 | 0.082 | 94.54 | 353 | 0.065183 | 0.082 | 0.0652 | 1 |

| | | | | | | | | | | |
|---|---|---|---|---|---|---|---|---|---|---|
| Q641Q3 | METRNL | HEPERQDsAIHLRVs | S223 | 0.002 | 95.19 | 311 | 0.00159 | 0.002 | 0.0016 | 1 |
| Q9UKR5 | ERG28 | RSWLVMVsIIAMGNT | S16 | 0.026 | 98.19 | 140 | 0.020668 | 0.026 | 0.0207 | 0.8333 |
| Q29RF7 | PDS5A | EKNGkLWsPDEEVSP | S823 | 0.031 | 89.88 | 1337 | 0.024642 | 0.031 | 0.0246 | 1 |
| Q29RF7 | PDS5A | NSTLRLLsAMLVsEG | S865 | 0.149 | 97.36 | 1337 | 0.118442 | 0.149 | 0.1184 | 1 |
| Q29RF7 | PDS5A | EQKRISksDMSRLRL | S883 | 0.069 | 93.09 | 1337 | 0.054849 | 0.069 | 0.0548 | 1 |
| Q29RF7 | PDS5A | KLLLPLEyMAIFALC | Y949 | 0 | 97.58 | 1337 | 0 | 0 | 0 | 1 |
| Q9BSM1 | PCGF1 | QNkYVRCsVRAEVRH | S195 | 0.114 | 96.75 | 259 | 0.09062 | 0.114 | 0.0906 | 0.8333 |
| Q9UPN3 | MACF1 | LEDLLQDsMDEKEQL | S831 | 0.039 | 92.74 | 7388 | 0.031002 | 0.039 | 0.031 | 1 |
| Q9UPN3 | MACF1 | IVQLkPRsPDHVLKN | S862 | 0.099 | 88.98 | 7388 | 0.078697 | 0.099 | 0.0787 | 1 |
| Q6YHU6 | THADA | LFCRIQEssQVLykL | S696 | 0.008 | 91.52 | 1953 | 0.006359 | 0.088 | 0.07 | 1 |
| P49759 | CLK1 | FLHSNKLtHTDLkPE | T285 | 0.005 | 98.58 | 484 | 0.003415 | 0.005 | 0.0034 | 1 |
| P49759 | CLK1 | EHHstLVstRHyRAP | S341 | 0.083 | 92.23 | 484 | 0.065978 | 0.125 | 0.0994 | 0.8889 |
| O60711 | LPXN | REQNDkTyCQPCFNk | Y375 | 0.22 | 94.33 | 386 | 0.096511 | 0.22 | 0.0965 | 1 |
| Q96HA8 | NTAQ1 | AVHYQPAsPPRDACV | S15 | 0.152 | 97.87 | 205 | 0.120827 | 0.152 | 0.1208 | 1 |
| Q9Y5N6 | ORC6 | SGLNkEtyQsCLKSF | Y79 | 0.077 | 95.56 | 252 | 0.033779 | 0.077 | 0.0338 | 1 |
| O43916 | CHST1 | AsRDLLRsLyDCDLY | S126 | 0.139 | 98 | 411 | 0.110493 | 0.139 | 0.1105 | 0.8 |
| O43916 | CHST1 | RDLLRsLyDCDLYFL | Y128 | 0.01 | 98.19 | 411 | 0.004387 | 0.01 | 0.0044 | 0.9286 |
| O43916 | CHST1 | PPWLKGKyMLVRYED | Y297 | 0.092 | 98.58 | 411 | 0.040359 | 0.092 | 0.0404 | 1 |
| P16989 | YBX3 | NPRKyLrsVGDGETV | S134 | 0.001 | 93.66 | 372 | 0.000795 | 0.001 | 0.0008 | 1 |
| P08575 | PTPRC | NLkPYTKyVLsLHAY | Y455 | 0.159 | 92.9 | 1306 | 0.069751 | 0.318 | 0.1395 | 1 |
| P08575 | PTPRC | DLQySTDyTFKAYFH | Y550 | 0.063 | 94.53 | 1306 | 0.027637 | 0.063 | 0.0276 | 1 |
| O00746 | NME4 | ESVLAEHyQDLRRkP | Y85 | 0.189 | 96.74 | 187 | 0.082912 | 0.216 | 0.0948 | 0.8 |
| O00746 | NME4 | SRNVIHAsDSVEGAQ | S153 | 0.001 | 98.19 | 187 | 0.000795 | 0.001 | 0.0008 | 1 |
| Q9UGI8 | TES | NQALARQyMQMLPKE | Y126 | 0.042 | 95.5 | 421 | 0.018425 | 0.042 | 0.0184 | 1 |
| Q9UGI8 | TES | kEGDPAIyAERAGyD | Y251 | 0.049 | 97.1 | 421 | 0.021496 | 0.049 | 0.0215 | 1 |
| Q9UGI8 | TES | kLyCGRHyCDsEkPR | Y293 | 0.133 | 98.67 | 421 | 0.058345 | 0.133 | 0.0583 | 1 |
| Q9UGI8 | TES | KPVCKPCyVKNHAVV | Y353 | 0.121 | 95.34 | 421 | 0.053081 | 0.121 | 0.0531 | 1 |
| P00533 | EGFR | MARDPQRyLVIQGDE | Y978 | 0.245 | 93.89 | 1210 | 0.107478 | 0.278 | 0.122 | 1 |
| Q9NPQ8 | RIC8A | GVRLLtDtLELtLGV | T195 | 0.086 | 96.98 | 531 | 0.058739 | 0.212 | 0.1499 | 0.875 |
| Q13724 | MOGS | YFGMKTRsPKPLLTG | S111 | 0.012 | 96.99 | 837 | 0.009539 | 0.161 | 0.1291 | 1 |
| Q5SW96 | LDLRAP1 | QKVTLKVsPRGIILT | S93 | 0.102 | 96.7 | 308 | 0.081081 | 0.174 | 0.1383 | 0.875 |

| | | | | | | | | | | |
|---|---|---|---|---|---|---|---|---|---|---|
| Q9NTG1 | PKDREJ | SFTVGPSsLLstLIQ | S692 | 0.013 | 86.32 | 2253 | 0.010334 | 0.013 | 0.0103 | 1 |
| Q9NTG1 | PKDREJ | KDFLPAGyLLyIVAs | Y708 | 0.181 | 90.39 | 2253 | 0.079402 | 0.349 | 0.1485 | 1 |
| Q9NTG1 | PKDREJ | LPAGyLLyIVAsVLN | Y711 | 0.028 | 91.4 | 2253 | 0.012283 | 0.028 | 0.0123 | 1 |
| Q9NTG1 | PKDREJ | yLLyIVAsVLNNMKt | S715 | 0.001 | 90.09 | 2253 | 0.000795 | 0.001 | 0.0008 | 1 |
| P08185 | SERPINA6 | SMALAMLsLGTCGHT | S78 | 0.007 | 97.7 | 405 | 0.005564 | 0.007 | 0.0056 | 1 |
| P07814 | EPRS1 | ILESKsLyDEVAAQG | Y827 | 0.157 | 89.72 | 1512 | 0.068874 | 0.327 | 0.1406 | 1 |
| P07814 | EPRS1 | LLsLkAQyKEKTGKE | Y864 | 0.069 | 86.44 | 1512 | 0.030269 | 0.121 | 0.0531 | 1 |
| Q6ZTR5 | CFAP47 | GPSyRQDyALFLRFE | Y366 | 0.088 | 88.64 | 3187 | 0.038604 | 0.088 | 0.0386 | 1 |
| Q6ZTR5 | CFAP47 | HVYLAFNsICKASTK | S518 | 0.068 | 89.44 | 3187 | 0.054054 | 0.068 | 0.0541 | 1 |
| Q70EK8 | USP53 | SREKALPsDNIRHAL | S95 | 0.012 | 96.46 | 1073 | 0.009539 | 0.012 | 0.0095 | 1 |
| P60520 | GABARAPL2 | IVEkVSGsQIVDIDk | S39 | 0.162 | 92.3 | 117 | 0.128776 | 0.162 | 0.1288 | 1 |
| P09211 | GSTP1 | AFLAsPEyVNLPING | Y199 | 0.231 | 98.39 | 210 | 0.101336 | 0.257 | 0.1127 | 0.8889 |
| Q9Y678 | COPG1 | LtEsETEyVIRCTKH | Y646 | 0.15 | 93.68 | 874 | 0.065803 | 0.294 | 0.1275 | 1 |
| P24928 | POLR2A | ELkRMsVtEGGIkYP | T37 | 0.003 | 92.47 | 1970 | 0.002049 | 0.003 | 0.002 | 1 |
| P24928 | POLR2A | KkRLTHVyDLCkGkN | Y145 | 0.289 | 93.9 | 1970 | 0.12678 | 0.289 | 0.1268 | 1 |
| P24928 | POLR2A | ALLCDTMtCRGHLMA | T1406 | 0.103 | 95.2 | 1970 | 0.07035 | 0.132 | 0.0819 | 1 |
| Q7Z443 | PKD1L3 | AKVVITLyGsEGRSE | Y768 | 0.023 | 90.36 | 1732 | 0.01009 | 0.434 | 0.1362 | 1 |
| Q7Z443 | PKD1L3 | tQDyLWLsIAtRHPW | S890 | 0 | 82.47 | 1732 | 0 | 0 | 0 | 1 |
| Q9UQP3 | TNN | GLRPGMEyMVHVWAQ | Y949 | 0.101 | 90.39 | 1299 | 0.044307 | 0.101 | 0.0443 | 1 |
| O15355 | PPM1G | EGKEEPGsDSGTTAV | S325 | 0.109 | 93.72 | 546 | 0.086646 | 0.109 | 0.0866 | 1 |
| O15355 | PPM1G | VNGGLNLsRAIGDHF | S397 | 0 | 98.08 | 546 | 0 | 0 | 0 | 0.8182 |
| Q9Y2Q0 | ATP8A1 | ALIIDGKtLKYALtF | T748 | 0.001 | 92.35 | 1164 | 0.000683 | 0.001 | 0.0007 | 1 |
| Q9Y2Q0 | ATP8A1 | YSIAQFKyLKNLLMI | Y839 | 0.165 | 95.37 | 1164 | 0.072383 | 0.329 | 0.1441 | 0.9167 |
| Q13541 | EIF4EBP1 | PGGtRIIyDRkFLME | Y54 | 0.081 | 87.61 | 118 | 0.035534 | 0.221 | 0.0969 | 1 |
| P54756 | EPHA5 | IVMWEVVsYGERPYW | S873 | 0.148 | 91.2 | 1037 | 0.117647 | 0.148 | 0.1176 | 1 |
| Q96RQ3 | MCCC1 | VQTVAVYsEADRNSM | S80 | 0 | 97.75 | 725 | 0 | 0 | 0 | 0.9091 |
| Q96RQ3 | MCCC1 | GPAPSQQsYLSMEKI | S106 | 0 | 96.43 | 725 | 0 | 0 | 0 | 0.9 |
| P61254 | RPL26 | IrRkIMssPLskELR | S32 | 0.001 | 97.1 | 145 | 0.000795 | 0.001 | 0.0008 | 0.9 |
| Q68DU8 | KCTD16 | RDGFLFRyILDYLRD | Y84 | 0.183 | 94.84 | 428 | 0.080279 | 0.183 | 0.0803 | 0.8889 |
| Q13895 | BYSL | LQPHPQLsPEIRREL | S414 | 0.117 | 95.48 | 437 | 0.093005 | 0.117 | 0.093 | 1 |
| P0CG38 | POTEI | APrAVFPsIVGRPRQ | S733 | 0.037 | 95.93 | 1075 | 0.029412 | 0.037 | 0.0294 | 1 |

| | | | | | | | | | | |
|---|---|---|---|---|---|---|---|---|---|---|
| P0CG38 | POTEI | AGRELTDyLMkILTE | Y888 | 0.173 | 96.79 | 1075 | 0.075893 | 0.232 | 0.1018 | 1 |
| P0CG38 | POTEI | MWISkQEyDEsGPsI | Y1062 | 0.114 | 97.39 | 1075 | 0.05001 | 0.114 | 0.05 | 1 |
| Q8N2G8 | GHDC | DHCLQEAsPRyKSLR | S467 | 0.109 | 95.04 | 530 | 0.086646 | 0.109 | 0.0866 | 1 |
| Q8N2G8 | GHDC | LQEAsPRyKSLRFWG | Y470 | 0.004 | 95.9 | 530 | 0.001755 | 0.004 | 0.0018 | 1 |
| Q9Y4W6 | AFG3L2 | MTLGGRVsEEIFFGR | S634 | 0.014 | 97.22 | 797 | 0.011129 | 0.014 | 0.0111 | 1 |
| Q9Y6X5 | ENPP4 | HVKNVFItKtFPNHY | T68 | 0 | 98.83 | 453 | 0 | 0 | 0 | 1 |
| Q9Y6X5 | ENPP4 | KNVFItKtFPNHYsI | T70 | 0.159 | 98.75 | 453 | 0.108599 | 0.159 | 0.1086 | 1 |
| Q9Y6X5 | ENPP4 | KtFPNHYsIVTGLYE | S76 | 0 | 98.95 | 453 | 0 | 0 | 0 | 1 |
| Q9UGM3 | DMBT1 | SPSyPAYyPNNAKCV | Y1786 | 0.021 | 82.08 | 2413 | 0.009212 | 0.021 | 0.0092 | 1 |
| Q9UGM3 | DMBT1 | RMTIHFRsDISFQNT | S1861 | 0.041 | 89.31 | 2413 | 0.032592 | 0.041 | 0.0326 | 1 |
| Q96MF6 | COQ10A | GIPAYPRtCTVDFSI | T186 | 0.068 | 96.5 | 247 | 0.046445 | 0.068 | 0.0464 | 1 |
| Q8NHY0 | B4GALNT2 | LAALEKTyNTyRSNt | Y538 | 0.264 | 93.72 | 566 | 0.115813 | 0.264 | 0.1158 | 1 |
| Q9HB89 | NMUR1 | VLNVTALsVERyVAV | S156 | 0.007 | 92.26 | 426 | 0.005564 | 0.007 | 0.0056 | 1 |
| P28340 | POLD1 | PSPFLRItVALPRLV | T213 | 0.163 | 97.86 | 1107 | 0.111331 | 0.163 | 0.1113 | 0.8571 |
| P28340 | POLD1 | IAPLRVLsFDIECAG | S314 | 0 | 98.57 | 1107 | 0 | 0 | 0 | 1 |
| P28340 | POLD1 | TRRRLAVyCLkDAYL | Y511 | 0.316 | 91.25 | 1107 | 0.138625 | 0.316 | 0.1386 | 1 |
| P28340 | POLD1 | REAADWVsGHFPsPI | S783 | 0.163 | 96.4 | 1107 | 0.129571 | 0.163 | 0.1296 | 0.875 |
| P28340 | POLD1 | LLISkkRyAGLLFSS | Y809 | 0.237 | 94.31 | 1107 | 0.103968 | 0.237 | 0.104 | 1 |
| P28340 | POLD1 | VANLVtAsLRRLLID | S847 | 0.004 | 95.89 | 1107 | 0.00318 | 0.004 | 0.0032 | 0.8 |
| Q6TDU7 | DNAI7 | KEEtEFHsTLYHMVK | S674 | 0.035 | 94.33 | 716 | 0.027822 | 0.035 | 0.0278 | 1 |
| Q92673 | SORL1 | KLEPGGKyHIIVQLG | Y2002 | 0.149 | 86.49 | 2214 | 0.065364 | 0.298 | 0.1307 | 1 |
| Q8N1P7 | CRYBG2 | GFNNHVLsVRIKGGI | S1478 | 0 | 97.33 | 1661 | 0 | 0 | 0 | 1 |
| Q96G97 | BSCL2 | QLLEVELyADYRENS | Y184 | 0.281 | 94.92 | 398 | 0.123271 | 0.281 | 0.1233 | 1 |
| Q9UKX3 | MYH13 | NTKRVIQyFATIAVT | Y195 | 0.013 | 88.55 | 1938 | 0.005703 | 0.013 | 0.0057 | 1 |
| P49327 | FASN | LQLSYTAtDRHPQAL | T1273 | 0 | 91.61 | 2511 | 0 | 0 | 0 | 0.9091 |
| Q96S38 | RPS6KC1 | AEDLLQFsANIPALY | S109 | 0.003 | 90.79 | 1066 | 0.002385 | 0.129 | 0.1025 | 0.8333 |
| Q96F07 | CYFIP2 | SWKLVHPtDkFCNkD | T419 | 0.116 | 92.05 | 1278 | 0.07923 | 0.116 | 0.0792 | 1 |
| Q9UGP8 | SEC63 | GkPGNyQytVFLRSD | Y694 | 0.207 | 95.32 | 760 | 0.090808 | 0.207 | 0.0908 | 1 |
| Q9UGP8 | SEC63 | kPGNyQytVFLRSDS | T695 | 0.147 | 96.48 | 760 | 0.100403 | 0.147 | 0.1004 | 0.8571 |
| Q96T83 | SLC9A7 | PIIFHAGySLKKRHF | Y197 | 0.194 | 87.53 | 725 | 0.085105 | 0.194 | 0.0851 | 1 |
| P50991 | CCT4 | ARMLVELsKAQDIEA | S95 | 0.006 | 95.78 | 539 | 0.004769 | 0.006 | 0.0048 | 1 |

| | | | | | | | | | | |
|---|---|---|---|---|---|---|---|---|---|---|
| P50991 | CCT4 | LLNSATTsLNSKVVS | S170 | 0.074 | 91.91 | 539 | 0.058824 | 0.074 | 0.0588 | 1 |
| P50991 | CCT4 | SQYsSLLsPMSVNAV | S184 | 0.008 | 94.96 | 539 | 0.006359 | 0.008 | 0.0064 | 1 |
| P50991 | CCT4 | GLIQFCLsAPKTDMD | S254 | 0.128 | 86.69 | 539 | 0.101749 | 0.144 | 0.127 | 1 |
| P50991 | CCT4 | AMEVIPstLAENAGL | T464 | 0.069 | 97.12 | 539 | 0.047128 | 0.132 | 0.0895 | 1 |
| Q08752 | PPID | ___MSHPsPQAkPSN | S5 | 0.056 | 97.49 | 370 | 0.044515 | 0.112 | 0.089 | 1 |
| Q08752 | PPID | VDkILLItEDLkNIG | T223 | 0 | 97.9 | 370 | 0 | 0 | 0 | 1 |
| Q08752 | PPID | WEMAIKKyAEVLryV | Y246 | 0.004 | 98.53 | 370 | 0.001755 | 0.004 | 0.0018 | 1 |
| Q08752 | PPID | KyAEVLryVDSSkAV | Y252 | 0.003 | 97.51 | 370 | 0.001316 | 0.003 | 0.0013 | 0.8125 |
| Q8TDY4 | ASAP3 | RDyIMAkyVEHRFAR | Y535 | 0.105 | 96.12 | 903 | 0.046062 | 0.105 | 0.0461 | 0.875 |
| P29317 | EPHA2 | NDMPIYMySVCNVMS | Y67 | 0.09 | 94.77 | 976 | 0.039482 | 0.09 | 0.0395 | 1 |
| P46940 | IQGAP1 | FLsAIVssVDkIPyG | S1152 | 0.016 | 91.27 | 1657 | 0.012719 | 0.089 | 0.0707 | 0.8 |
| P53621 | COPA | ICRVttVtEIGkDVI | T1207 | 0 | 96.99 | 1224 | 0 | 0 | 0 | 1 |
| P53621 | COPA | DVIGLrIsPLQFR__ | S1219 | 0.008 | 92.28 | 1224 | 0.006359 | 0.008 | 0.0064 | 0.8571 |
| Q9NP66 | HMG20A | PPEEKQRyLDEADRD | Y152 | 0.157 | 96.06 | 347 | 0.068874 | 0.157 | 0.0689 | 1 |
| Q9NP66 | HMG20A | yMkELEQyQktEAyk | Y170 | 0.191 | 89.63 | 347 | 0.083789 | 0.191 | 0.0838 | 1 |
| Q7L014 | DDX46 | EPFRkNFyVEVPELA | Y336 | 0.314 | 91.74 | 1031 | 0.137747 | 0.314 | 0.1377 | 1 |
| Q7L014 | DDX46 | GPIAVIMtPtRELAL | T452 | 0 | 93.37 | 1031 | 0 | 0 | 0 | 1 |
| Q7L014 | DDX46 | RELALQItKECKKFS | T462 | 0.084 | 95.04 | 1031 | 0.057373 | 0.084 | 0.0574 | 1 |
| Q7L014 | DDX46 | VTNLRRVtyVVLDEA | T524 | 0.021 | 96.79 | 1031 | 0.014343 | 0.021 | 0.0143 | 1 |
| Q7L014 | DDX46 | TNLRRVtyVVLDEAD | Y525 | 0.011 | 97.53 | 1031 | 0.004826 | 0.011 | 0.0048 | 1 |
| Q7L014 | DDX46 | AGNkGYAytFITEDQ | Y720 | 0.339 | 95.91 | 1031 | 0.148714 | 0.339 | 0.1487 | 1 |
| Q7L014 | DDX46 | GNkGYAytFITEDQA | T721 | 0.003 | 96.05 | 1031 | 0.002049 | 0.003 | 0.002 | 1 |
| Q9P2R7 | SUCLA2 | PEEAkAVsSQMIGkK | S132 | 0 | 98.08 | 463 | 0 | 0 | 0 | 1 |
| Q9P2R7 | SUCLA2 | KINFDsNsAYRQKkI | S281 | 0.063 | 96.56 | 463 | 0.05008 | 0.063 | 0.0501 | 1 |
| Q9P2R7 | SUCLA2 | ARMVVKLsEIVtLAK | S444 | 0.049 | 96.47 | 463 | 0.038951 | 0.049 | 0.039 | 1 |
| Q13480 | GAB1 | DPDVLEyyKNDHAKK | Y48 | 0.261 | 93.52 | 694 | 0.114497 | 0.261 | 0.1145 | 0.9 |
| Q8N8V2 | GBP7 | VPGGHNIyLEAKKKI | Y435 | 0 | 94.03 | 638 | 0 | 0 | 0 | 1 |
| Q8IYB1 | MB21D2 | EKNGTIIsIILGVGs | S221 | 0.023 | 97.91 | 491 | 0.018283 | 0.023 | 0.0183 | 1 |
| Q8IYB1 | MB21D2 | LKKCISSsLMQAyQA | S305 | 0.018 | 96.65 | 491 | 0.014308 | 0.018 | 0.0143 | 1 |
| Q8IYB1 | MB21D2 | SSsLMQAyQACKAII | Y310 | 0.066 | 97.5 | 491 | 0.028953 | 0.066 | 0.029 | 0.9231 |
| Q13950 | RUNX2 | AELVRTDsPNFLCSV | S118 | 0.166 | 94.27 | 521 | 0.131956 | 0.166 | 0.132 | 1 |

| | | | | | | | | | | |
|---|---|---|---|---|---|---|---|---|---|---|
| Q14554 | PDIA5 | KPLLIMFyAPWCSMC | Y178 | 0.235 | 96.96 | 519 | 0.103091 | 0.235 | 0.1031 | 1 |
| Q14008 | CKAP5 | QKNPKNQsETLNWLS | S733 | 0.086 | 94.71 | 2032 | 0.068363 | 0.086 | 0.0684 | 1 |
| P11413 | G6PD | DFFARNsyVAGQyDD | Y107 | 0.336 | 98.84 | 515 | 0.147398 | 0.336 | 0.1474 | 1 |
| P11413 | G6PD | NsyVAGQyDDAASYQ | Y112 | 0.263 | 98.1 | 515 | 0.115374 | 0.263 | 0.1154 | 1 |
| P11413 | G6PD | sQANRLFyLALPPtV | Y139 | 0 | 98.85 | 515 | 0 | 0 | 0 | 1 |
| P11413 | G6PD | LALPPtVyEAVTKNI | Y147 | 0.009 | 96.85 | 515 | 0.003948 | 0.009 | 0.0039 | 1 |
| P11413 | G6PD | EGEATkGyLDDPTVP | Y322 | 0.002 | 98.43 | 515 | 0.000877 | 0.002 | 0.0009 | 1 |
| P11413 | G6PD | VQPNEAVytkMMTKk | Y401 | 0.294 | 98.61 | 515 | 0.128974 | 0.294 | 0.129 | 1 |
| P11413 | G6PD | EEsELDLtyGNRyKN | T423 | 0.162 | 96.62 | 515 | 0.110648 | 0.162 | 0.1106 | 1 |
| P11413 | G6PD | EsELDLtyGNRyKNV | Y424 | 0.28 | 95.95 | 515 | 0.122832 | 0.28 | 0.1228 | 1 |
| P11413 | G6PD | NVkLPDAyERLILDV | Y437 | 0.237 | 94.95 | 515 | 0.103968 | 0.237 | 0.104 | 1 |
| Q9UBL0 | ARPP21 | LHEFLINtLkNNSRD | T156 | 0.117 | 94.27 | 812 | 0.079913 | 0.117 | 0.0799 | 0.8889 |
| Q9UBL0 | ARPP21 | NVDQtGksVIINkTS | S218 | 0.039 | 94.87 | 812 | 0.031002 | 0.039 | 0.031 | 1 |
| Q9HCK5 | AGO4 | PKIDVYHyDVDIKPE | Y47 | 0.022 | 98.43 | 861 | 0.009651 | 0.022 | 0.0097 | 1 |
| Q9HCK5 | AGO4 | TYVRCTRsVSIPAPA | S798 | 0.063 | 98.48 | 861 | 0.05008 | 0.063 | 0.0501 | 1 |
| Q14669 | TRIP12 | RSLNPPLtIVRktFE | T1937 | 0.131 | 93.53 | 1992 | 0.089475 | 0.131 | 0.0895 | 0.8889 |
| Q9BYX4 | IFIH1 | QPFLKKWyRVIGLsG | Y385 | 0.228 | 95.05 | 1025 | 0.10002 | 0.228 | 0.1 | 1 |
| Q9BYX4 | IFIH1 | GRARADEsTYVLVAH | S828 | 0.081 | 92.53 | 1025 | 0.064388 | 0.114 | 0.0906 | 1 |
| Q8WUF8 | ARB2A | HELTSWKsFPSIFKF | S381 | 0 | 98.7 | 416 | 0 | 0 | 0 | 0.9167 |
| Q8IVU1 | IGDCC3 | QHGDGNAtVRFVsLR | T606 | 0.196 | 90.5 | 814 | 0.133871 | 0.196 | 0.1339 | 1 |
| Q9UI15 | TAGLN3 | DMAAVQRtLMALGsV | T128 | 0.003 | 97.75 | 199 | 0.002049 | 0.003 | 0.002 | 1 |
| Q9UI15 | TAGLN3 | RtLMALGsVAVTKDD | S134 | 0.121 | 94.73 | 199 | 0.096185 | 0.121 | 0.0962 | 1 |
| Q12965 | MYO1E | MDDyIFTyIGSVLIS | Y51 | 0.18 | 91.26 | 1108 | 0.078963 | 0.18 | 0.079 | 1 |
| Q9Y4F3 | MARF1 | SNRLRRLsDNCGGkV | S536 | 0 | 91.92 | 1742 | 0 | 0 | 0 | 0.9091 |
| P08575 | PTPRC | LFLAEFQsIPRVFsk | S659 | 0.109 | 95.69 | 1306 | 0.086646 | 0.109 | 0.0866 | 1 |
| P08575 | PTPRC | RyVDILPyDyNRVEL | Y689 | 0.005 | 98.31 | 1306 | 0.002193 | 0.005 | 0.0022 | 1 |
| P08575 | PTPRC | NGDAGSNyINASyID | Y707 | 0.008 | 98.33 | 1306 | 0.003509 | 0.008 | 0.0035 | 1 |
| P08575 | PTPRC | NRNkCAEyWPSMEEG | Y765 | 0.005 | 97.64 | 1306 | 0.002193 | 0.005 | 0.0022 | 1 |
| P08575 | PTPRC | QHKRCPDyIIQkLNI | Y792 | 0.046 | 94 | 1306 | 0.02018 | 0.135 | 0.0592 | 1 |
| P08575 | PTPRC | CSAGVGRtGtyIGID | T860 | 0 | 98.06 | 1306 | 0 | 0 | 0 | 1 |
| P08575 | PTPRC | AGVGRtGtyIGIDAM | T862 | 0 | 98.03 | 1306 | 0 | 0 | 0 | 1 |

| | | | | | | | | | | |
|---|---|---|---|---|---|---|---|---|---|---|
| P08575 | PTPRC | GVGRtGtyIGIDAML | Y863 | 0.041 | 98.21 | 1306 | 0.017986 | 0.082 | 0.036 | 1 |
| P08575 | PTPRC | AENKVDVyGYVVKLR | Y882 | 0.129 | 98.13 | 1306 | 0.05659 | 0.258 | 0.1132 | 1 |
| P08575 | PTPRC | MVQVEAQyILIHQAL | Y902 | 0.008 | 97.76 | 1306 | 0.003509 | 0.008 | 0.0035 | 1 |
| P11362 | FGFR1 | EEVEVQDsVPADSGL | S91 | 0.016 | 89.3 | 822 | 0.012719 | 0.016 | 0.0127 | 1 |
| Q7L0Y3 | TRMT10C | AKRLNLAtECLPLDk | T333 | 0.133 | 97.98 | 403 | 0.090841 | 0.133 | 0.0908 | 1 |
| Q13425 | SNTB2 | RLGDAILsVNGtDLR | S165 | 0.12 | 93.96 | 540 | 0.09539 | 0.12 | 0.0954 | 1 |
| O15033 | AREL1 | KkVyCyVsPKQFSVK | S358 | 0.158 | 88.46 | 823 | 0.125597 | 0.158 | 0.1256 | 1 |
| O15033 | AREL1 | LHkNIGGsETFQDkV | S436 | 0.031 | 80.51 | 823 | 0.024642 | 0.031 | 0.0246 | 1 |
| P52895 | AKR1C2 | VkREDIFytskLWSN | Y81 | 0.001 | 98.92 | 323 | 0.000439 | 0.001 | 0.0004 | 0.9412 |
| P52895 | AKR1C2 | NQVECHPyFNQRkLL | Y196 | 0.067 | 98.7 | 323 | 0.029392 | 0.067 | 0.0294 | 1 |
| P52895 | AKR1C2 | GLNRNVRyLtLDIFA | Y305 | 0.061 | 97.26 | 323 | 0.02676 | 0.061 | 0.0268 | 1 |
| Q9NWZ5 | UCKL1 | DMKIFVDtDsDIRLV | T228 | 0.166 | 92.65 | 548 | 0.11338 | 0.166 | 0.1134 | 1 |
| Q9NWZ5 | UCKL1 | IGNFGDRyFGTDAVP | Y529 | 0.291 | 94.28 | 548 | 0.127658 | 0.291 | 0.1277 | 1 |
| Q14517 | FAT1 | DRETTDKytLNItVy | Y788 | 0.154 | 88.41 | 4588 | 0.067558 | 0.154 | 0.0676 | 1 |
| Q14517 | FAT1 | DKytLNItVyDLGIP | T793 | 0.14 | 89.835 | 4588 | 0.095622 | 0.1625 | 0.1401 | 1 |
| Q14517 | FAT1 | SSNDkLMyKITSGNP | Y1176 | 0.194 | 91.995 | 4588 | 0.085105 | 0.2315 | 0.1016 | 1 |
| Q8N3S3 | PHTF2 | GSETSKKysNtSILL | Y695 | 0.151 | 83.11 | 785 | 0.066242 | 0.151 | 0.0662 | 1 |
| P22105 | TNXB | RKYKMNLyGLRGRQR | Y1330 | 0.279 | 92.84 | 4244 | 0.122393 | 0.279 | 0.1224 | 0.8 |
| Q9Y4P3 | TBL2 | ATCADDrtIRIWSTk | T113 | 0.06 | 97.82 | 447 | 0.040981 | 0.093 | 0.0605 | 0.8571 |
| Q9Y4P3 | TBL2 | GTWkLWDtDVEYkkk | T308 | 0.114 | 97 | 447 | 0.077864 | 0.216 | 0.1462 | 1 |
| Q9Y4P3 | TBL2 | APCRLALsPNAQVLA | S340 | 0.012 | 97.71 | 447 | 0.009539 | 0.012 | 0.0095 | 1 |
| Q02742 | GCNT1 | MKDLyAMsANWKYLI | S207 | 0.145 | 96.98 | 428 | 0.115263 | 0.145 | 0.1153 | 1 |
| Q02742 | GCNT1 | EERWKKRyEVVNGKL | Y259 | 0.259 | 98.27 | 428 | 0.11362 | 0.259 | 0.1136 | 1 |
| P03891 | MT-ND2 | LGGLPPLtGFLPKWA | T258 | 0.019 | 96.04 | 347 | 0.012977 | 0.019 | 0.013 | 1 |
| Q5SY80 | CATSPERE | YGSALWRySTNSPNy | Y23 | 0.044 | 94.07 | 951 | 0.019302 | 0.044 | 0.0193 | 0.9167 |
| Q5SY80 | CATSPERE | ySTNSPNyRIFSTRs | Y30 | 0.327 | 87.98 | 951 | 0.14345 | 0.327 | 0.1434 | 0.9 |
| Q5SY80 | CATSPERE | yRIFSTRsTIKLEyE | S37 | 0.045 | 92.75 | 951 | 0.035771 | 0.045 | 0.0358 | 1 |
| Q5SY80 | CATSPERE | RsTIKLEyEGtLFTE | Y43 | 0.147 | 92.72 | 951 | 0.064487 | 0.147 | 0.0645 | 1 |
| Q5SY80 | CATSPERE | SKNLRVRyIWGEyGC | Y745 | 0.059 | 93.24 | 951 | 0.025882 | 0.059 | 0.0259 | 1 |
| Q5SY80 | CATSPERE | VRyIWGEyGCPLRLD | Y750 | 0.175 | 90.24 | 951 | 0.07677 | 0.175 | 0.0768 | 1 |
| P24385 | CCND1 | YRLTRFLsRVIkCDP | S234 | 0.072 | 93.54 | 295 | 0.057234 | 0.072 | 0.0572 | 1 |

| | | | | | | | | | | |
|---|---|---|---|---|---|---|---|---|---|---|
| Q8N2A8 | PLD6 | GVRVRVVtDCDYMAL | T124 | 0.017 | 96.23 | 252 | 0.011611 | 0.017 | 0.0116 | 1 |
| Q96GX5 | MASTL | IAVPKPPsIEEFSIV | S31 | 0.18 | 86.58 | 879 | 0.143085 | 0.18 | 0.1431 | 1 |
| Q9H4G0 | EPB41L1 | IMELHKTyRGMTPGE | Y261 | 0.233 | 97.27 | 881 | 0.102214 | 0.233 | 0.1022 | 1 |
| Q6GYQ0 | RALGAPA1 | LSPELFEsPNIQFFV | S1554 | 0.018 | 87.73 | 2036 | 0.014308 | 0.018 | 0.0143 | 1 |
| Q6GYQ0 | RALGAPA1 | LNNtTLVsCIQIRsE | S1569 | 0.015 | 93.46 | 2036 | 0.011924 | 0.015 | 0.0119 | 1 |
| Q6GYQ0 | RALGAPA1 | AGLAsANsNVRIIVR | S1593 | 0.179 | 85.93 | 2036 | 0.14229 | 0.179 | 0.1423 | 1 |
| P55895 | RAG2 | GDVPEARyGHSINVV | Y138 | 0.138 | 98.56 | 527 | 0.060539 | 0.138 | 0.0605 | 0.9167 |
| P55895 | RAG2 | RMICNIIsLEDNKIE | S291 | 0.137 | 95.75 | 527 | 0.108903 | 0.137 | 0.1089 | 1 |
| Q96NG3 | ODAD4 | KFQQAIDtWEEKIPL | T382 | 0.167 | 97.24 | 672 | 0.114063 | 0.167 | 0.1141 | 1 |
| O75369 | FLNB | GPNHIVGsPFkAKVt | S2465 | 0.041 | 80.18 | 2602 | 0.032592 | 0.078 | 0.0817 | 0.8333 |
| Q9Y2I2 | NTNG1 | DELHLARyFyAIsDI | Y284 | 0.096 | 91.46 | 539 | 0.042114 | 0.096 | 0.0421 | 1 |
| Q8WWI5 | SLC44A1 | PKLPVPAsAPIPFFH | S167 | 0.058 | 89.83 | 657 | 0.046105 | 0.058 | 0.0461 | 1 |
| O75369 | FLNB | ALGLEAVsDsGTKAE | S1172 | 0.113 | 80.07 | 2602 | 0.089825 | 0.113 | 0.0898 | 1 |
| Q9UHE8 | STEAP1 | WLDKWMLtRKQFGLL | T160 | 0.07 | 97.23 | 339 | 0.047811 | 0.07 | 0.0478 | 0.8571 |
| Q13287 | NMI | MRDkLELsFskSRNG | S180 | 0.076 | 92.78 | 307 | 0.060414 | 0.076 | 0.0604 | 1 |
| Q13287 | NMI | QsGsAVItFVEIGVA | T206 | 0.199 | 95.52 | 307 | 0.13592 | 0.199 | 0.1359 | 1 |
| Q6PL18 | ATAD2 | RIFLRNVtHRLAIDK | T997 | 0.001 | 91.15 | 1390 | 0.000683 | 0.001 | 0.0007 | 1 |
| Q6PL18 | ATAD2 | SkIDLHkyLTVkDyL | Y1043 | 0.167 | 87.84 | 1390 | 0.07326 | 0.167 | 0.0733 | 0.9167 |
| Q6PL18 | ATAD2 | kyLTVkDyLRDIDLI | Y1049 | 0 | 92.77 | 1390 | 0 | 0 | 0 | 0.8571 |
| P98198 | ATP8B2 | sMEyPKLyEPGQLNL | Y959 | 0.021 | 92.61 | 1209 | 0.009212 | 0.021 | 0.0092 | 1 |
| P51991 | HNRNPA3 | IGGLsFEttDDsLrE | T46 | 0.002 | 96.23 | 378 | 0.001366 | 0.002 | 0.0014 | 1 |
| P27338 | MAOB | LRNQKVkyVDLGGSy | Y53 | 0.202 | 98.5 | 520 | 0.088614 | 0.202 | 0.0886 | 0.9091 |
| P27338 | MAOB | LYAkVLGsLEALEPV | S374 | 0.133 | 97.73 | 520 | 0.105724 | 0.064 | 0.1228 | 1 |
| P27338 | MAOB | EALEPVHyEEKNWCE | Y383 | 0.162 | 98.67 | 520 | 0.071067 | 0.162 | 0.0711 | 1 |
| Q16706 | MAN2A1 | NDYFRDKtQyIFNNM | T192 | 0 | 98.05 | 1144 | 0 | 0 | 0 | 1 |
| Q16706 | MAN2A1 | FIWSEISyLSKWWDI | Y218 | 0.005 | 98.64 | 1144 | 0.002193 | 0.005 | 0.0022 | 1 |
| Q16706 | MAN2A1 | ARMLLDQyRKKSKLF | Y407 | 0.02 | 98.27 | 1144 | 0.008774 | 0.02 | 0.0088 | 1 |
| Q16706 | MAN2A1 | WDLQFKNyQQLFDYM | Y442 | 0.051 | 98.64 | 1144 | 0.022373 | 0.051 | 0.0224 | 1 |
| Q16706 | MAN2A1 | HYWSGYFtSRPFyKR | T509 | 0.004 | 98.61 | 1144 | 0.002732 | 0.004 | 0.0027 | 1 |
| Q16706 | MAN2A1 | LRAAEILyyFALRQA | Y532 | 0.04 | 97.54 | 1144 | 0.017547 | 0.04 | 0.0175 | 1 |
| Q16706 | MAN2A1 | RAAEILyyFALRQAH | Y533 | 0 | 97.61 | 1144 | 0 | 0 | 0 | 1 |

| | | | | | | | | | | |
|---|---|---|---|---|---|---|---|---|---|---|
| Q16706 | MAN2A1 | kVYKILEsASSNSHL | S723 | 0.187 | 88.99 | 1144 | 0.148649 | 0.187 | 0.1486 | 1 |
| Q16706 | MAN2A1 | SNSHLADyVLYkNkV | Y733 | 0.046 | 94.94 | 1144 | 0.02018 | 0.046 | 0.0202 | 1 |
| Q16706 | MAN2A1 | EGQSVEVsNIVDIRK | S865 | 0.123 | 98.51 | 1144 | 0.097775 | 0.123 | 0.0978 | 1 |
| Q13618 | CUL3 | DMsISNTtMDEFRQH | T483 | 0.137 | 96.1 | 768 | 0.093573 | 0.137 | 0.0936 | 1 |
| Q6P6C2 | ALKBH5 | SLPVRrGsVTVLSGY | S252 | 0.024 | 98.17 | 394 | 0.019078 | 0.024 | 0.0191 | 1 |
| O15068 | MCF2L | LKEMLKysRNCEGAE | S785 | 0.001 | 94.55 | 1137 | 0.000795 | 0.001 | 0.0008 | 1 |
| O60383 | GDF9 | PRAVGHRyGSPVHTM | Y394 | 0.213 | 84.1 | 454 | 0.09344 | 0.213 | 0.0934 | 1 |
| Q13017 | ARHGAP5 | RPPsYTIsIVGLsGt | S16 | 0 | 97.76 | 1502 | 0 | 0 | 0 | 1 |
| Q13017 | ARHGAP5 | TIsIVGLsGtEKDKG | S21 | 0.08 | 93.33 | 1502 | 0.063593 | 0.113 | 0.0951 | 1 |
| Q13017 | ARHGAP5 | KPVIIAAtKCDECVD | T199 | 0.047 | 96.93 | 1502 | 0.032102 | 0.047 | 0.0321 | 1 |
| Q13017 | ARHGAP5 | NHPDYEEyINLEGTR | Y306 | 0.066 | 96.12 | 1502 | 0.028953 | 0.066 | 0.029 | 0.9231 |
| P01730 | CD4 | sFLtKGPsKLNDRAD | S74 | 0.18 | 93.53 | 458 | 0.143085 | 0.18 | 0.1431 | 1 |
| Q9BUB7 | TMEM70 | sEDGRLIyTGNMARA | Y96 | 0.02 | 92.71 | 260 | 0.008774 | 0.02 | 0.0088 | 1 |
| Q9BUB7 | TMEM70 | HEATTDTyKAITyNA | Y179 | 0.02 | 94.49 | 260 | 0.008774 | 0.02 | 0.0088 | 1 |
| Q9H0A0 | NAT10 | TPNLLARtVETVEGG | T137 | 0.018 | 90.27 | 1025 | 0.012294 | 0.018 | 0.0123 | 1 |
| P08253 | MMP2 | GNKyEsCtsAGRsDG | T364 | 0.188 | 93.43 | 660 | 0.128407 | 0.188 | 0.1284 | 1 |
| P08253 | MMP2 | DGKMWCAttANYDDD | T377 | 0.219 | 94.54 | 660 | 0.14958 | 0.219 | 0.1496 | 1 |
| Q58DX5 | NAALADL2 | AEVIDVsyGMADDLK | Y281 | 0.265 | 93.5 | 795 | 0.116252 | 0.265 | 0.1163 | 1 |
| Q9HB63 | NTN4 | VQEQLKItNLRVQLL | T222 | 0.006 | 98.25 | 628 | 0.004098 | 0.006 | 0.0041 | 0.8889 |
| Q12774 | ARHGEF5 | LIVSEAsyLRSLNIA | Y1189 | 0 | 95.99 | 1597 | 0 | 0.15 | 0.0658 | 1 |
| Q12774 | ARHGEF5 | NQTyQERtFQsLMNs | T1277 | 0.165 | 94.75 | 1597 | 0.112697 | 0.165 | 0.1127 | 1 |
| O43854 | EDIL3 | SPEYIkSyKIAySND | Y246 | 0.01 | 98.13 | 480 | 0.004387 | 0.01 | 0.0044 | 1 |
| O43854 | EDIL3 | DGkTWAMyKVKGTNE | Y260 | 0.165 | 97.37 | 480 | 0.072383 | 0.165 | 0.0724 | 1 |
| Q8TD08 | MAPK15 | APEVLLSsHRYTLGV | S192 | 0.087 | 71.18 | 544 | 0.069158 | 0.106 | 0.0843 | 1 |
| Q9BUB5 | MKNK1 | HFNEREAsRVVRDVA | S148 | 0.045 | 98.64 | 465 | 0.035771 | 0.086 | 0.0688 | 1 |
| Q9BUB5 | MKNK1 | APEKGLPtPQVLQRN | T385 | 0.004 | 93.92 | 465 | 0.002732 | 0.004 | 0.0027 | 1 |
| Q9UKU9 | ANGPTL2 | SRyQDGVyWAEFRGG | Y467 | 0.077 | 98.52 | 493 | 0.033779 | 0.077 | 0.0338 | 0.8 |
| Q9UKU9 | ANGPTL2 | AEFRGGSySLKKVVM | Y476 | 0.077 | 98.31 | 493 | 0.033779 | 0.077 | 0.0338 | 1 |
| Q9BS92 | NIPSNAP3B | FYEFRTYyLKPSNMN | Y43 | 0.341 | 96.93 | 247 | 0.149592 | 0.341 | 0.1496 | 1 |
| Q08426 | EHHADH | PAALDLItSGRRILA | T153 | 0.148 | 98.41 | 723 | 0.101086 | 0.148 | 0.1011 | 1 |
| Q08426 | EHHADH | QKTGkGWyQyDkPLG | Y580 | 0.072 | 98.76 | 723 | 0.031585 | 0.072 | 0.0316 | 1 |

| | | | | | | | | | | |
|---|---|---|---|---|---|---|---|---|---|---|
| Q08426 | EHHADH | TGkGWyQyDkPLGRI | Y582 | 0.283 | 98.5 | 723 | 0.124148 | 0.283 | 0.1241 | 1 |
| Q08426 | EHHADH | VLEKLQKyyRQNPDI | Y682 | 0.078 | 98.69 | 723 | 0.034217 | 0.078 | 0.0342 | 1 |
| P42345 | MTOR | APEAPLPsRkAALET | S1131 | 0 | 86.5 | 2549 | 0 | 0 | 0 | 1 |
| P42345 | MTOR | IIHPIVRtLDQsPEL | T1162 | 0.007 | 84.96 | 2549 | 0.004781 | 0.007 | 0.0048 | 1 |
| P42345 | MTOR | IVRtLDQsPELRstA | S1166 | 0.062 | 80.93 | 2549 | 0.049285 | 0.062 | 0.0493 | 1 |
| P42345 | MTOR | VFQLGKkyQIFIPMV | Y1188 | 0 | 84.22 | 2549 | 0 | 0 | 0 | 1 |
| Q86UK0 | ABCA12 | MLLGRILyAPyNPVT | Y811 | 0.104 | 93.04 | 2595 | 0.045623 | 0.199 | 0.0841 | 1 |
| O75390 | CS | RGMkGLVyEtsVLDP | Y80 | 0.274 | 98.83 | 466 | 0.1202 | 0.274 | 0.1202 | 1 |
| O75390 | CS | MkGLVyEtsVLDPDE | T82 | 0.004 | 98.78 | 466 | 0.002732 | 0.004 | 0.0027 | 1 |
| O75390 | CS | kGLVyEtsVLDPDEG | S83 | 0.039 | 98.78 | 466 | 0.031002 | 0.039 | 0.031 | 0.9 |
| Q9UQ49 | NEU3 | QEDDRGItyRIPALL | T23 | 0.009 | 97.63 | 428 | 0.006147 | 0.009 | 0.0061 | 0.8182 |
| Q9UQ49 | NEU3 | EDDRGItyRIPALLy | Y24 | 0.276 | 98.44 | 428 | 0.121077 | 0.276 | 0.1211 | 0.8182 |
| Q13057 | COASY | GLTGISGsGKSSIAQ | S369 | 0.02 | 97.01 | 564 | 0.015898 | 0.02 | 0.0159 | 1 |
| P15924 | DSP | CANsIKDyELQLASY | Y961 | 0.041 | 89.8 | 2871 | 0.017986 | 0.349 | 0.1378 | 0.6 |
| A6NNM8 | TTLL13 | VRDGAVGsKRKLSTL | S310 | 0.124 | 89.69 | 815 | 0.098569 | 0.124 | 0.0986 | 1 |
| Q5T653 | MRPL2 | LLRKVNGtAIIQLPs | T223 | 0.07 | 96.85 | 305 | 0.047811 | 0.07 | 0.0478 | 1 |
| Q5T653 | MRPL2 | VATVGRVsNVDHNKR | S248 | 0.068 | 97.95 | 305 | 0.054054 | 0.068 | 0.0541 | 1 |
| Q9H7D0 | DOCK5 | EVFALLVsSLLENLL | S1198 | 0.123 | 91.06 | 1870 | 0.097775 | 0.123 | 0.0978 | 1 |
| Q8N6M0 | OTUD6B | VVALRSQtAEYMQsH | T183 | 0 | 98.38 | 293 | 0 | 0 | 0 | 1 |
| Q8IVV2 | LOXHD1 | NIGVKVLyEMTVWTG | Y1554 | 0.095 | 91.69 | 2067 | 0.041675 | 0.095 | 0.0417 | 1 |
| Q8IVV2 | LOXHD1 | GDVVGGGtDsNIFMT | T1568 | 0.026 | 88.06 | 2067 | 0.017758 | 0.052 | 0.0355 | 1 |
| Q8IVV2 | LOXHD1 | VVGGGtDsNIFMTLY | S1570 | 0.003 | 90.07 | 2067 | 0.002385 | 0.003 | 0.0024 | 1 |
| P17516 | AKR1C4 | MPVLGFGtyAPPEVP | T23 | 0.063 | 98.03 | 323 | 0.04303 | 0.063 | 0.043 | 1 |
| P17516 | AKR1C4 | VkREDIFytskLWCT | Y81 | 0.003 | 98.89 | 323 | 0.001316 | 0.003 | 0.0013 | 0.9375 |
| P17516 | AKR1C4 | NQVECHPyLNQskLL | Y196 | 0.084 | 98.8 | 323 | 0.03685 | 0.084 | 0.0369 | 1 |
| P17516 | AKR1C4 | GLNRNYRyVVMDFLM | Y305 | 0.07 | 98.2 | 323 | 0.030708 | 0.07 | 0.0307 | 1 |
| O94973 | AP2A2 | FHLCSVPtRALLLST | T536 | 0.006 | 94.07 | 939 | 0.004098 | 0.006 | 0.0041 | 1 |
| O94953 | KDM4B | EFkDFNkyVAyIEsQ | Y31 | 0.101 | 97.63 | 1096 | 0.044307 | 0.101 | 0.0443 | 1 |
| Q08J23 | NSUN2 | EkMkVINtGIkVWCR | T583 | 0.001 | 94.06 | 767 | 0.000683 | 0.11 | 0.0751 | 0.8333 |
| Q8TB24 | RIN3 | QQARtLAsRADTQAQ | S896 | 0.056 | 85.31 | 985 | 0.044515 | 0.116 | 0.0922 | 0.8333 |
| O95996 | APC2 | EELHQVVssILRNLs | S510 | 0 | 98 | 2303 | 0 | 0 | 0 | 1 |

| | | | | | | | | | | |
|---|---|---|---|---|---|---|---|---|---|---|
| O95996 | APC2 | ssILRNLsWRADINS | S517 | 0.047 | 97.68 | 2303 | 0.037361 | 0.047 | 0.0374 | 1 |
| Q6UWP2 | DHRS11 | HIRAtCIsPGVVETQ | S197 | 0 | 98.84 | 260 | 0 | 0 | 0 | 0.8889 |
| Q7L576 | CYFIP1 | NLSkIDKyFKQLQVV | Y298 | 0.191 | 95.26 | 1253 | 0.083789 | 0.191 | 0.0838 | 1 |
| Q9NRN7 | AASDHPPT | kEWAFEEsKIDEHHF | S233 | 0.085 | 98.17 | 309 | 0.067568 | 0.085 | 0.0676 | 0.8889 |
| Q8NA54 | IQUB | ARELQNIyKCIMLKN | Y502 | 0.221 | 93.55 | 791 | 0.09695 | 0.221 | 0.0969 | 1 |
| Q5HYC2 | KIAA2026 | SRHLREKtTIAVTSR | T204 | 0.045 | 91.05 | 2103 | 0.030736 | 0.045 | 0.0307 | 0.8889 |
| P17540 | CKMT2 | ECLTPAIyAKLRNKV | Y73 | 0.025 | 98.61 | 419 | 0.010967 | 0.025 | 0.011 | 1 |
| P17540 | CKMT2 | NKVTPNGyTLDQCIQ | Y85 | 0.107 | 98.33 | 419 | 0.046939 | 0.2 | 0.0827 | 1 |
| P17540 | CKMT2 | DEHyVLssRVRtGRs | S163 | 0.014 | 97.78 | 419 | 0.011129 | 0.014 | 0.0111 | 1 |
| P17540 | CKMT2 | VADVyDIsNIDRIGR | S371 | 0.004 | 97.25 | 419 | 0.00318 | 0.004 | 0.0032 | 1 |
| Q07617 | SPAG1 | LQALAPEsRALRKDK | S72 | 0.107 | 85.68 | 926 | 0.085056 | 0.107 | 0.0851 | 1 |
| Q9H082 | RAB33B | IKIQLWDtAGQERFR | T89 | 0.008 | 90.75 | 229 | 0.005464 | 0.008 | 0.0055 | 0.8 |
| Q9BZM2 | PLA2G2F | VLstAHGsLLNLKAM | S21 | 0 | 91.66 | 168 | 0 | 0 | 0 | 0.9167 |
| O95497 | VNN1 | HYPSKKMtGSGIYAP | T275 | 0.001 | 98.79 | 513 | 0.000683 | 0.001 | 0.0007 | 1 |
| Q7Z5L7 | PODN | PANAFDstPNLKGIF | T545 | 0.022 | 95.98 | 613 | 0.015026 | 0.022 | 0.015 | 1 |
| Q8N0S6 | CENPL | RLVRVSTsVASAHTD | S310 | 0.147 | 93.74 | 344 | 0.116852 | 0.147 | 0.1169 | 1 |
| O75132 | ZBED4 | REMSTQMsTLSQVIP | S952 | 0.01 | 91.82 | 1171 | 0.007949 | 0.01 | 0.0079 | 1 |
| O75132 | ZBED4 | TEEEAEQykQDLIRE | Y1025 | 0.274 | 95.89 | 1171 | 0.1202 | 0.396 | 0.1366 | 1 |
| O95405 | ZFYVE9 | LYPCPLFsVRFRKPL | S1091 | 0.003 | 98.36 | 1425 | 0.002385 | 0.003 | 0.0024 | 1 |
| P52732 | KIF11 | NINQsLLtLGRVITA | T294 | 0.056 | 90.24 | 1056 | 0.038249 | 0.056 | 0.0382 | 1 |
| P52732 | KIF11 | LTRILQDsLGGRTRT | S323 | 0.055 | 90.12 | 1056 | 0.04372 | 0.055 | 0.0437 | 1 |
| P27635 | RPL10 | PLCGHMVsDEyEQLs | S54 | 0.02 | 97.45 | 214 | 0.015898 | 0.02 | 0.0159 | 1 |
| Q5T5J6 | SWT1 | IsGVKSLsKEELSAE | S526 | 0.077 | 95.9 | 900 | 0.061208 | 0.077 | 0.0612 | 1 |
| Q5JZY3 | EPHA10 | SDLVCKIsGFGRGPR | S787 | 0.174 | 93.76 | 1008 | 0.138315 | 0.174 | 0.1383 | 0.8 |
| Q9HAC7 | SUGCT | DAVASAVsGLMHITG | S183 | 0.125 | 97.08 | 445 | 0.099364 | 0.187 | 0.1486 | 1 |
| P21333 | FLNA | kAyVGQkssFTVDCS | S2576 | 0.137 | 88.5 | 2647 | 0.108903 | 0.137 | 0.1089 | 1 |
| P21333 | FLNA | LLkDkGEytLVVkWG | Y2626 | 0.008 | 88.75 | 2647 | 0.003509 | 0.008 | 0.0035 | 1 |
| Q08257 | CRYZ | CGVNPVEtyIRSGTY | T52 | 0.065 | 97.81 | 329 | 0.044396 | 0.126 | 0.087 | 1 |
| Q08257 | CRYZ | kkGDRVFtsstISGG | T95 | 0.004 | 98.84 | 329 | 0.002732 | 0.004 | 0.0027 | 1 |
| Q08257 | CRYZ | kGDRVFtsstISGGy | S96 | 0.088 | 98.04 | 329 | 0.069952 | 0.127 | 0.1184 | 0.8182 |
| Q08257 | CRYZ | sstISGGyAEyALAA | Y103 | 0.004 | 98.77 | 329 | 0.001755 | 0.004 | 0.0018 | 0.8125 |

| | | | | | | | | | | |
|---|---|---|---|---|---|---|---|---|---|---|
| Q08257 | CRYZ | RDTMAKEsSIIGVTL | S264 | 0.126 | 98.09 | 329 | 0.100159 | 0.126 | 0.1002 | 1 |
| P10523 | SAG | RDKSVTIyLGNRDYI | Y29 | 0.16 | 98.19 | 405 | 0.07019 | 0.16 | 0.0702 | 1 |
| P10523 | SAG | LVKGKKVyVtLtCAF | Y62 | 0.172 | 95.62 | 405 | 0.075454 | 0.172 | 0.0755 | 1 |
| P10523 | SAG | KGKKVyVtLtCAFRY | T64 | 0.055 | 96.16 | 405 | 0.037566 | 0.055 | 0.0376 | 0.8889 |
| P10523 | SAG | LTssEVAtEVPFRLM | T353 | 0.15 | 97.32 | 405 | 0.102452 | 0.15 | 0.1025 | 1 |
| Q02241 | KIF23 | LkDPVGVyCRVRPLG | Y29 | 0.041 | 98.11 | 960 | 0.017986 | 0.041 | 0.018 | 1 |
| Q02241 | KIF23 | KNGLLFTyGVTGsGK | Y111 | 0.019 | 95.87 | 960 | 0.008335 | 0.019 | 0.0083 | 1 |
| Q02241 | KIF23 | FTyGVTGsGKTHTMT | S116 | 0.041 | 93.32 | 960 | 0.032592 | 0.041 | 0.0472 | 1 |
| Q02241 | KIF23 | GTNkMVPyRDSKLTH | Y388 | 0.127 | 95.99 | 960 | 0.055713 | 0.127 | 0.0557 | 1 |
| Q68CZ2 | TNS3 | GsFIVRDsHsFRGAy | S1199 | 0.141 | 91.01 | 1445 | 0.112083 | 0.141 | 0.1121 | 1 |
| Q9C0J8 | WDR33 | tLHAHkNtVMEVKLN | T290 | 0.197 | 97.12 | 1336 | 0.134554 | 0.197 | 0.1346 | 1 |
| Q6UWY5 | OLFML1 | MKDAVYNsPKVYLLI | S172 | 0.109 | 89.86 | 402 | 0.086646 | 0.109 | 0.0866 | 1 |
| Q6UWY5 | OLFML1 | QGTGQVIyKGFLFFH | Y221 | 0.231 | 96.59 | 402 | 0.101336 | 0.231 | 0.1013 | 0.9091 |
| P25205 | MCM3 | VRPkVVRsVHYCPAT | S144 | 0.025 | 89.32 | 808 | 0.019873 | 0.025 | 0.0199 | 1 |
| P25205 | MCM3 | DrVQVVGtYRCLPGK | T240 | 0.011 | 89.55 | 808 | 0.007513 | 0.011 | 0.0075 | 0.9091 |
| Q7L0X0 | TRIL | SGDIFAAsPALyRLD | S348 | 0.014 | 91.68 | 811 | 0.011129 | 0.014 | 0.0111 | 1 |
| Q0IIM8 | TBC1D8B | DNVTNkDsPLPSNVQ | S718 | 0.076 | 74.98 | 1120 | 0.060414 | 0.076 | 0.0604 | 1 |
| O43405 | COCH | MLSRWSAsFtVtKGK | S120 | 0.04 | 95.82 | 550 | 0.031797 | 0.04 | 0.0318 | 1 |
| Q7RTY8 | TMPRSS7 | SPHIRRLsGIRAYFE | S351 | 0.054 | 88.24 | 843 | 0.042925 | 0.054 | 0.0429 | 1 |
| Q7RTY8 | TMPRSS7 | YPPKCKCtWKFQTSL | T397 | 0.17 | 95.08 | 843 | 0.116112 | 0.17 | 0.1161 | 1 |
| A7MBM2 | DISP2 | FFDLWRLsKSQLPSG | S569 | 0.081 | 87.78 | 1401 | 0.064388 | 0.081 | 0.0644 | 1 |
| P40879 | SLC26A3 | NIGRTNIyKNKKDYY | Y520 | 0.272 | 91.1 | 764 | 0.119322 | 0.272 | 0.1193 | 1 |
| Q9BY15 | ADGRE3 | TEGRQEIssTATTIL | S158 | 0.032 | 90.17 | 652 | 0.025437 | 0.032 | 0.0254 | 0.9 |
| Q9BY15 | ADGRE3 | VESKVLEtALKDPEQ | T175 | 0.116 | 93.44 | 652 | 0.07923 | 0.116 | 0.0792 | 1 |
| Q9BY15 | ADGRE3 | TGQGSQWsRDGCFLI | S317 | 0.093 | 93.86 | 652 | 0.073927 | 0.093 | 0.0739 | 1 |
| Q93062 | RBPMS | LQEEEVRtLFVsGLP | T25 | 0.003 | 95.42 | 196 | 0.002049 | 0.003 | 0.002 | 0.875 |
| O75323 | NIPSNAP2 | PRsGPNIyELrSYQL | Y187 | 0.005 | 97.82 | 286 | 0.002193 | 0.005 | 0.0022 | 0.9167 |
| Q15788 | NCOA1 | GKIIsIDtSSLRAAG | T277 | 0.159 | 93.91 | 1441 | 0.108599 | 0.159 | 0.1086 | 1 |
| Q15788 | NCOA1 | QPQGREPsYARQLFQ | S308 | 0.147 | 92.87 | 1441 | 0.116852 | 0.147 | 0.1169 | 1 |
| Q13868 | EXOSC2 | DSVLLLssMNLPGGE | S112 | 0.038 | 83.58 | 293 | 0.030207 | 0.038 | 0.0302 | 1 |
| O60239 | SH3BP5 | ELEkLNQsTDDINRR | S55 | 0.101 | 97.64 | 455 | 0.080286 | 0.047 | 0.0926 | 1 |

| | | | | | | | | | | |
|---|---|---|---|---|---|---|---|---|---|---|
| O60239 | SH3BP5 | AVEDSkPyWEARRVA | Y101 | 0.155 | 98.53 | 455 | 0.067996 | 0.31 | 0.136 | 1 |
| O60239 | SH3BP5 | LKRAINKskPYFELk | S208 | 0 | 98.03 | 455 | 0 | 0 | 0 | 1 |
| P15104 | GLUL | GVANRsAsIRIPRTV | S322 | 0.003 | 97.71 | 373 | 0.002385 | 0.003 | 0.0024 | 1 |
| P15104 | GLUL | yFEDRRPsANCDPFS | S343 | 0 | 98.78 | 373 | 0 | 0 | 0 | 1 |
| Q9H0F6 | SHARPIN | GPIRLQVtLEDAASA | T224 | 0.184 | 94.85 | 387 | 0.125675 | 0.184 | 0.1257 | 1 |
| Q9H0F6 | SHARPIN | CLCVPERsLASYGVR | S282 | 0.172 | 95.52 | 387 | 0.136725 | 0.172 | 0.1367 | 1 |
| P32321 | DCTD | TAENkLDtKyPYVCH | T77 | 0.048 | 97.62 | 178 | 0.032785 | 0.048 | 0.0328 | 1 |
| P08514 | ITGA2B | LLVGAPLyMESRADR | Y344 | 0.206 | 98.64 | 1039 | 0.090369 | 0.33 | 0.1448 | 1 |
| P08514 | ITGA2B | LAEVGRVyLFLQPRG | Y360 | 0.082 | 98.76 | 1039 | 0.035972 | 0.082 | 0.036 | 1 |
| Q9UPU3 | SORCS3 | LYDFNLGsVTEsSLW | S224 | 0.152 | 87.36 | 1222 | 0.120827 | 0.152 | 0.1208 | 1 |
| Q9UPU3 | SORCS3 | NLGsVTEsSLWRSTD | S228 | 0.14 | 92.37 | 1222 | 0.111288 | 0.14 | 0.1113 | 1 |
| Q9UPU3 | SORCS3 | NDKVGLKtVLSyLyV | T251 | 0.141 | 88.83 | 1222 | 0.096305 | 0.141 | 0.0963 | 1 |
| Q9UPU3 | SORCS3 | SLDQKLYssMDFGRR | S321 | 0.048 | 95.03 | 1222 | 0.038156 | 0.048 | 0.0382 | 1 |
| Q9UPU3 | SORCS3 | LDQKLYssMDFGRRW | S322 | 0.023 | 95.13 | 1222 | 0.018283 | 0.023 | 0.0183 | 1 |
| P09848 | LCT | ADRMLQFsLGWFAHP | S1150 | 0.052 | 95.27 | 1927 | 0.041336 | 0.071 | 0.0682 | 1 |
| P09848 | LCT | PIFRNGDyPDtMKWK | Y1164 | 0.014 | 96.27 | 1927 | 0.006142 | 0.014 | 0.0061 | 1 |
| P09848 | LCT | DGIDLRGyVAWSLMD | Y1313 | 0.033 | 98.68 | 1927 | 0.014477 | 0.066 | 0.029 | 1 |
| P09848 | LCT | GSFWLKMtPFGFRRI | T1725 | 0.009 | 94.81 | 1927 | 0.006147 | 0.009 | 0.0061 | 1 |
| Q15208 | STK38 | FTFQNMNskRKAEtW | S264 | 0.075 | 90.24 | 465 | 0.059619 | 0.075 | 0.0596 | 1 |
| P07355 | ANXA2 | RQDIAFAyQRRTKkE | Y75 | 0.103 | 98.7 | 339 | 0.045185 | 0.185 | 0.0737 | 1 |
| P07355 | ANXA2 | DAsELkAsMkGLGtD | S117 | 0.021 | 97.53 | 339 | 0.016693 | 0.034 | 0.027 | 0.8889 |
| P07355 | ANXA2 | LQEINRVykEMYkTD | Y147 | 0 | 98.67 | 339 | 0 | 0 | 0 | 1 |
| P07355 | ANXA2 | EkDIIsDtsGDFRkL | T163 | 0.085 | 97.22 | 339 | 0.058056 | 0.085 | 0.0581 | 1 |
| P07355 | ANXA2 | IsIMtERsVPHLQkV | S221 | 0.117 | 98.15 | 339 | 0.093005 | 0.134 | 0.1065 | 0.8571 |
| P07355 | ANXA2 | LQkVFDRyksysPyD | Y232 | 0 | 98.7 | 339 | 0 | 0 | 0 | 1 |
| P07355 | ANXA2 | FDRyksysPyDMLEs | S236 | 0.1 | 97.77 | 339 | 0.079491 | 0.147 | 0.1357 | 1 |
| P07355 | ANXA2 | sPyDMLEsIRkEVkG | S243 | 0 | 98.44 | 339 | 0 | 0 | 0 | 1 |
| P07355 | ANXA2 | FADRLyDsMkGkGTR | S277 | 0.02 | 97.75 | 339 | 0.015898 | 0.02 | 0.0159 | 1 |
| P07355 | ANXA2 | FkRkYGksLyyyIQQ | S314 | 0.099 | 98.13 | 339 | 0.078697 | 0.176 | 0.1424 | 1 |
| P07355 | ANXA2 | yyyIQQDtkGDYQkA | T323 | 0.03 | 97.22 | 339 | 0.02049 | 0.03 | 0.0205 | 1 |
| Q15796 | SMAD2 | RMCTIRMsFVkGWGA | S417 | 0.005 | 98.68 | 467 | 0.003975 | 0.005 | 0.004 | 1 |

| | | | | | | | | | | |
|---|---|---|---|---|---|---|---|---|---|---|
| A0AVT1 | UBA6 | VEIDDALySRQRYVL | Y44 | 0.142 | 92.19 | 1052 | 0.062293 | 0.142 | 0.0623 | 1 |
| A0AVT1 | UBA6 | FPAAIEHtIQWARDk | T637 | 0 | 98.03 | 1052 | 0 | 0 | 0 | 1 |
| A0AVT1 | UBA6 | VQGVKMLyVPVMPGH | Y994 | 0.336 | 94.68 | 1052 | 0.147398 | 0.336 | 0.1474 | 1 |
| P28329 | CHAT | DQLRFAAsLISGVLs | S237 | 0.075 | 96.86 | 748 | 0.059619 | 0.088 | 0.07 | 1 |
| P28329 | CHAT | KDSTNRDsLDMIERC | S377 | 0.056 | 98.39 | 748 | 0.044515 | 0.056 | 0.0445 | 1 |
| P28329 | CHAT | LHRRLVPtyESASIR | T553 | 0.028 | 98.93 | 748 | 0.019124 | 0.028 | 0.0191 | 1 |
| Q9Y232 | CDYL | VRKQDGFtHILLstk | T353 | 0.051 | 98.09 | 598 | 0.034834 | 0.051 | 0.0348 | 1 |
| P36544 | CHRNA7 | YECCKEPyPDVtFTV | Y217 | 0.235 | 94.36 | 502 | 0.103091 | 0.235 | 0.1031 | 1 |
| P36544 | CHRNA7 | PDVtFTVtMRRRTLY | T225 | 0.204 | 94.73 | 502 | 0.139335 | 0.204 | 0.1393 | 1 |
| Q92610 | ZNF592 | AFKTASstADHSATQ | T842 | 0.059 | 88.44 | 1267 | 0.040298 | 0.155 | 0.1059 | 1 |
| P37268 | FDFT1 | LEDFPtIsLEFRNLA | S129 | 0 | 98.65 | 417 | 0 | 0 | 0 | 1 |
| Q92859 | NEO1 | TFELVPtsPPKDVTV | S956 | 0.139 | 83.61 | 1461 | 0.110493 | 0.139 | 0.1105 | 1 |
| Q9ULB4 | CDH9 | PKFTKDLyTASVPEM | Y164 | 0.141 | 95.89 | 789 | 0.061855 | 0.141 | 0.0619 | 1 |
| Q9ULB4 | CDH9 | DFENQMLyTLRVDAS | Y343 | 0.094 | 95.84 | 789 | 0.041236 | 0.094 | 0.0412 | 1 |
| Q9ULB4 | CDH9 | PVFTKVsyLIEVDED | Y388 | 0.175 | 96.62 | 789 | 0.07677 | 0.175 | 0.0768 | 1 |
| Q9ULB4 | CDH9 | ARNNLIKysVDRHTD | Y419 | 0.179 | 96.68 | 789 | 0.078525 | 0.179 | 0.0785 | 1 |
| P30837 | ALDH1B1 | VAEQTPLsALYLASL | S217 | 0.012 | 98.76 | 517 | 0.009539 | 0.012 | 0.0095 | 1 |
| P30837 | ALDH1B1 | DSNLKRVtLELGGKS | T283 | 0.184 | 98.9 | 517 | 0.125675 | 0.184 | 0.1257 | 1 |
| P30837 | ALDH1B1 | TFVEEsIyNEFLERT | Y332 | 0.138 | 98.73 | 517 | 0.060539 | 0.138 | 0.0605 | 1 |
| P30837 | ALDH1B1 | QFERVLGyIQLGQkE | Y373 | 0.249 | 98.82 | 517 | 0.109233 | 0.249 | 0.1092 | 1 |
| P30837 | ALDH1B1 | RGFFIkPtVFGGVQD | T401 | 0 | 98.93 | 517 | 0 | 0 | 0 | 1 |
| P21397 | MAOA | RDRVGGRtyTIRNEH | T52 | 0.068 | 98.89 | 527 | 0.046445 | 0.068 | 0.0464 | 0.8182 |
| P21397 | MAOA | LYAKVLGsQEALHPV | S383 | 0.075 | 98.46 | 527 | 0.059619 | 0.082 | 0.0728 | 1 |
| Q8IWA4 | MFN1 | AFFGRTSsGKSSVIN | S86 | 0.048 | 91.03 | 741 | 0.038156 | 0.079 | 0.0628 | 1 |
| Q8IWA4 | MFN1 | QNRIFFVsAkEVLsA | S284 | 0.017 | 95.64 | 741 | 0.013514 | 0.062 | 0.0582 | 1 |
| O75534 | CSDE1 | EEADPTIySGkVIRP | Y597 | 0.224 | 89.72 | 798 | 0.098266 | 0.224 | 0.0983 | 1 |
| O75165 | DNAJC13 | TLyALTSsTKIIKEA | S1843 | 0.157 | 87.4 | 2243 | 0.124802 | 0.157 | 0.1248 | 1 |
| Q13671 | RIN1 | PTtGyLVyRRAEWPE | Y701 | 0.004 | 91.09 | 783 | 0.001755 | 0.004 | 0.0018 | 0.8824 |
| Q8N0U8 | VKORC1L1 | kCSAALAsRWGRGFG | S64 | 0.155 | 94.01 | 176 | 0.123212 | 0.155 | 0.1232 | 1 |
| O95147 | DUSP14 | FWRQLIDyERQLFGk | Y161 | 0.094 | 98.86 | 198 | 0.041236 | 0.094 | 0.0412 | 1 |
| P10275 | AR | CVRMRHLsQEFGWLQ | S792 | 0.001 | 97.57 | 920 | 0.000795 | 0.001 | 0.0008 | 1 |

| | | | | | | | | | | |
|---|---|---|---|---|---|---|---|---|---|---|
| P19525 | EIF2AK2 | KGtLRyMsPEQISSQ | S456 | 0 | 93.14 | 551 | 0 | 0 | 0 | 0.875 |
| Q99497 | PARK7 | VVICPDAsLEDAkkE | S57 | 0.107 | 98.82 | 189 | 0.085056 | 0.106 | 0.1227 | 1 |
| Q99497 | PARK7 | GFGSKVTtHPLAkDk | T125 | 0 | 98.91 | 189 | 0 | 0 | 0 | 0.8333 |
| Q99497 | PARK7 | EkDGLILtSRGPGTS | T154 | 0 | 98.96 | 189 | 0 | 0 | 0 | 0.9231 |
| Q8NI37 | PPTC7 | RTSHRLHtANLGDSG | T153 | 0.019 | 98.7 | 304 | 0.012977 | 0.019 | 0.013 | 0.8182 |
| Q8TDY4 | ASAP3 | FAAKMPRyRGAALAR | Y34 | 0.047 | 87.93 | 903 | 0.020618 | 0.308 | 0.1351 | 0.9167 |
| Q9UF56 | FBXL17 | CKQLSDTsIIAVASH | S429 | 0.001 | 98.26 | 701 | 0.000795 | 0.001 | 0.0008 | 1 |
| Q9UF56 | FBXL17 | ITELDNEtVMEIVKR | T557 | 0.125 | 98.49 | 701 | 0.085377 | 0.125 | 0.0854 | 0.8889 |
| Q8IZ73 | RPUSD2 | FAKTAAVsERIHEQV | S290 | 0.128 | 94.65 | 545 | 0.101749 | 0.128 | 0.1017 | 1 |
| P55291 | CDH15 | SIVKALDyESCEHyE | Y330 | 0.151 | 92.29 | 814 | 0.066242 | 0.151 | 0.0662 | 0.8 |
| P55291 | CDH15 | DyESCEHyELKVSVQ | Y336 | 0.231 | 94.74 | 814 | 0.101336 | 0.231 | 0.1013 | 1 |
| Q58EX2 | SDK2 | RNGLVLGyKVMYkEK | Y1235 | 0.021 | 89.72 | 2172 | 0.009212 | 0.069 | 0.0303 | 1 |
| Q58EX2 | SDK2 | GLGKyVLyEVQVLAF | Y1272 | 0.221 | 89.72 | 2172 | 0.09695 | 0.221 | 0.0969 | 1 |
| P22314 | UBA1 | ADIDEGLysRQLyVL | Y55 | 0.05 | 96.37 | 1058 | 0.021934 | 0.05 | 0.0219 | 1 |
| P22314 | UBA1 | MkRLQTSsVLVSGLr | S74 | 0.14 | 98.66 | 1058 | 0.111288 | 0.14 | 0.1113 | 1 |
| P22314 | UBA1 | IkLVVADtRGLFGQL | T191 | 0.092 | 98.73 | 1058 | 0.062837 | 0.092 | 0.0628 | 1 |
| P22314 | UBA1 | SFksLVAsLAEPDFV | S309 | 0 | 97.87 | 1058 | 0 | 0 | 0 | 1 |
| P22314 | UBA1 | AEPDFVVtDFAkFSR | T318 | 0.095 | 97.65 | 1058 | 0.064886 | 0.095 | 0.0649 | 1 |
| P22314 | UBA1 | GQVAVFGsDLQEkLG | S460 | 0.127 | 96.02 | 1058 | 0.100954 | 0.127 | 0.101 | 1 |
| P22314 | UBA1 | GPDTErIyDDDFFQN | Y560 | 0.068 | 96.96 | 1058 | 0.029831 | 0.068 | 0.0298 | 1 |
| P22314 | UBA1 | MDFIVAAsNLRAENy | S866 | 0.001 | 98.51 | 1058 | 0.000795 | 0.001 | 0.0008 | 1 |
| P22314 | UBA1 | sNLRAENyDIPSADr | Y873 | 0.029 | 97.24 | 1058 | 0.012722 | 0.029 | 0.0127 | 1 |
| Q86SE5 | RALYL | KTQtSNVtNKNDPkS | T11 | 0.158 | 92.43 | 291 | 0.107916 | 0.158 | 0.1079 | 0.8889 |
| Q9NPZ5 | B3GAT2 | MRTTRKVsVWPVGLV | S205 | 0 | 98.82 | 323 | 0 | 0 | 0 | 1 |
| Q13951 | CBFB | RQtPSREyVDLEREA | Y85 | 0.256 | 93.28 | 182 | 0.112304 | 0.256 | 0.1123 | 1 |
| P25100 | ADRA1D | WVVALVVsVGPLLGW | S228 | 0.076 | 85.65 | 572 | 0.060414 | 0.076 | 0.0604 | 1 |
| P60174 | TPI1 | GAFTGEIsPGMIkDC | S80 | 0.068 | 98.86 | 249 | 0.054054 | 0.068 | 0.0541 | 1 |
| Q8TEY7 | USP33 | ELRARGLtGLKNIGN | T185 | 0 | 95.57 | 942 | 0 | 0.053 | 0.0362 | 0.9167 |
| O15034 | RIMBP2 | CVTSRGssDELQCTL | S377 | 0.037 | 92.44 | 1052 | 0.029412 | 0.037 | 0.0294 | 1 |
| Q9H2D1 | SLC25A32 | GAMTLCItNPLWVtK | T138 | 0.024 | 95.33 | 315 | 0.016392 | 0.024 | 0.0164 | 1 |
| Q05639 | EEF1A2 | VDSGkstttGHLIyk | T23 | 0.055 | 92.39 | 463 | 0.037566 | 0.055 | 0.0376 | 0.8571 |

| | | | | | | | | | | |
|---|---|---|---|---|---|---|---|---|---|---|
| Q05639 | EEF1A2 | WkFETTkyyltIIDA | Y85 | 0.195 | 90.91 | 463 | 0.085544 | 0.195 | 0.0855 | 1 |
| Q05639 | EEF1A2 | EHALLAytLGVkQLI | T142 | 0.14 | 82.69 | 463 | 0.095622 | 0.173 | 0.1182 | 0.8571 |
| Q05639 | EEF1A2 | ykIGGIGtVPVGrVE | T261 | 0.107 | 93.93 | 463 | 0.073082 | 0.122 | 0.0833 | 1 |
| Q05639 | EEF1A2 | FAPVNITtEVKsVEM | T287 | 0 | 96.02 | 463 | 0 | 0 | 0 | 1 |
| Q6VVX0 | CYP2R1 | GYSIPKGtTVItNLY | T398 | 0.086 | 98.54 | 501 | 0.058739 | 0.086 | 0.0587 | 1 |
| Q6VVX0 | CYP2R1 | PKGtTVItNLYSVHF | T402 | 0.004 | 98.89 | 501 | 0.002732 | 0.004 | 0.0027 | 1 |
| P25963 | NFKBIA | IHEEkALtMEVIRQV | T90 | 0 | 97.65 | 317 | 0 | 0 | 0 | 1 |
| Q8N6Y1 | PCDH20 | RRGPRPEsFRIDPKT | S794 | 0.011 | 88.28 | 951 | 0.008744 | 0.011 | 0.0087 | 1 |
| P62256 | UBE2H | PDkYPFksPSIGFMN | S65 | 0.151 | 97.1 | 183 | 0.120032 | 0.151 | 0.12 | 1 |
| Q96G03 | PGM2 | DARLDQEtAQWLRWD | T19 | 0.091 | 94.9 | 612 | 0.062154 | 0.145 | 0.099 | 1 |
| Q96G03 | PGM2 | MLSSTVSsKILRAIA | S381 | 0.037 | 97.96 | 612 | 0.029412 | 0.037 | 0.0294 | 0.8571 |
| Q96G03 | PGM2 | SQQLKAIyVEyGYHI | Y469 | 0.231 | 97.62 | 612 | 0.101336 | 0.231 | 0.1013 | 0.8333 |
| Q96G03 | PGM2 | LKAIyVEyGYHITKA | Y472 | 0.2 | 97.41 | 612 | 0.087737 | 0.2 | 0.0877 | 0.9286 |
| Q96A35 | MRPL24 | AGERVRVstRsGRII | S147 | 0.017 | 96.61 | 216 | 0.013514 | 0.017 | 0.0135 | 1 |
| Q8IWV8 | UBR2 | LGSIIGYsDGLRRIL | S328 | 0.039 | 90.29 | 1755 | 0.031002 | 0.181 | 0.1439 | 1 |
| P14384 | CPM | TRIHIMPsMNPDGFE | S124 | 0.028 | 98.85 | 443 | 0.022258 | 0.028 | 0.0223 | 0.8889 |
| O95977 | S1PR4 | LCAFDRCssLLPLYS | S194 | 0.026 | 92.74 | 384 | 0.020668 | 0.066 | 0.0525 | 1 |
| Q92508 | PIEZO1 | FKLVALytVWVALKE | T822 | 0.077 | 93.41 | 2521 | 0.052592 | 0.077 | 0.0526 | 1 |
| Q6ZW49 | PAXIP1 | ESQVSSAyAQAIRER | Y661 | 0.234 | 94.35 | 1069 | 0.102652 | 0.234 | 0.1027 | 1 |
| Q14416 | GRM2 | IGRYNIFtYLRAGSG | T457 | 0.003 | 95.76 | 872 | 0.002049 | 0.003 | 0.002 | 1 |
| Q14416 | GRM2 | PLPASRCsEPCLQNE | S501 | 0.187 | 90.07 | 872 | 0.148649 | 0.187 | 0.1486 | 1 |
| P57088 | TMEM33 | MMTNKLDtAMWLSRL | T25 | 0.148 | 94.4 | 247 | 0.101086 | 0.148 | 0.1011 | 0.875 |
| Q9Y279 | VSIG4 | DHIQQAKyQGRLHVs | Y82 | 0.011 | 95.47 | 399 | 0.004826 | 0.011 | 0.0048 | 1 |
| Q06830 | PRDX1 | NCQVIGAsVDsHFCH | S77 | 0.015 | 98.8 | 199 | 0.011924 | 0.015 | 0.0119 | 1 |
| Q06830 | PRDX1 | PMNIPLVsDPkRtIA | S106 | 0.092 | 98.86 | 199 | 0.073132 | 0.112 | 0.0971 | 1 |
| Q06830 | PRDX1 | kRtIAQDyGVLkADE | Y116 | 0.097 | 98.48 | 199 | 0.042553 | 0.097 | 0.0426 | 1 |
| Q06830 | PRDX1 | LkADEGIsFRGLFII | S126 | 0.004 | 97.77 | 199 | 0.00318 | 0.004 | 0.0032 | 1 |
| Q15020 | SART3 | VPDLWIRysQyLDrQ | Y350 | 0.012 | 98.55 | 963 | 0.005264 | 0.012 | 0.0053 | 1 |
| Q15020 | SART3 | PDLWIRysQyLDrQL | S351 | 0.028 | 98.09 | 963 | 0.022258 | 0.028 | 0.0223 | 1 |
| Q15020 | SART3 | LWIRysQyLDrQLKV | Y353 | 0.235 | 98.29 | 963 | 0.103091 | 0.235 | 0.1031 | 1 |
| Q5SYC1 | CLVS2 | ANWDQSRyTLVDILR | Y131 | 0.064 | 96.4 | 327 | 0.028076 | 0.064 | 0.0281 | 1 |

| | | | | | | | | | | |
|---|---|---|---|---|---|---|---|---|---|---|
| Q8WWV3 | RTN4IP1 | RVLILGAsGGVGTFA | S214 | 0.153 | 97.74 | 396 | 0.121622 | 0.153 | 0.1216 | 1 |
| Q9NWA0 | MED9 | IKCMDKDsPEVHQDL | S80 | 0.183 | 88.92 | 146 | 0.145469 | 0.183 | 0.1455 | 1 |
| Q9H9A6 | LRRC40 | MVsDVDLsFNKLSFI | S457 | 0.06 | 93.65 | 602 | 0.047695 | 0.06 | 0.0477 | 1 |
| O95886 | DLGAP3 | IRSAVGstQLLLSQK | T840 | 0 | 97.94 | 979 | 0 | 0 | 0 | 1 |
| Q8N4L2 | PIP4P2 | CTVCNEAtPIKNPPT | T85 | 0.039 | 83.37 | 257 | 0.026638 | 0.039 | 0.0266 | 0.9 |
| P63261 | ACTG1 | APrAVFPsIVGRPRH | S33 | 0.024 | 97.61 | 375 | 0.019078 | 0.024 | 0.0191 | 0.9091 |
| P63261 | ACTG1 | MEkIWHHtFyNELRV | T89 | 0 | 98.67 | 375 | 0 | 0 | 0 | 0.8889 |
| P63261 | ACTG1 | TTGIVMDsGdGVtHt | S155 | 0.006 | 98.34 | 375 | 0.004769 | 0.006 | 0.0048 | 0.8333 |
| P63261 | ACTG1 | sGdGVtHtVPIyEGy | T162 | 0.016 | 98.5 | 375 | 0.010928 | 0.016 | 0.0109 | 1 |
| P63261 | ACTG1 | DLAGRDLtDyLMkIL | T186 | 0.02 | 98.15 | 375 | 0.01366 | 0.034 | 0.0247 | 0.9 |
| P63261 | ACTG1 | VDIRkDLyANtVLsG | Y294 | 0.006 | 98.35 | 375 | 0.002632 | 0.006 | 0.0026 | 1 |
| P63261 | ACTG1 | RkDLyANtVLsGGtt | T297 | 0.003 | 98.61 | 375 | 0.002049 | 0.003 | 0.002 | 1 |
| P63261 | ACTG1 | LyANtVLsGGttMyP | S300 | 0.005 | 98.46 | 375 | 0.003975 | 0.005 | 0.004 | 1 |
| P63261 | ACTG1 | LsGGttMyPGIADRM | Y306 | 0.163 | 98.68 | 375 | 0.071506 | 0.268 | 0.1176 | 0.8667 |
| P63261 | ACTG1 | MWISkQEyDEsGPsI | Y362 | 0.134 | 98.3 | 375 | 0.058784 | 0.268 | 0.1176 | 1 |
| P09923 | ALPI | TRGNEVIsVMNRAKQ | S150 | 0.044 | 98.92 | 528 | 0.034976 | 0.044 | 0.035 | 1 |
| P09923 | ALPI | TTRVQHAsPAGTyAH | S174 | 0 | 98.78 | 528 | 0 | 0 | 0 | 0.9 |
| P09923 | ALPI | HAsPAGTyAHTVNRN | Y179 | 0.047 | 98.92 | 528 | 0.020618 | 0.047 | 0.0206 | 1 |
| Q3B7T1 | EDRF1 | kKSkPFSsFGMAYDF | S106 | 0 | 88.85 | 1238 | 0 | 0 | 0 | 1 |
| P12107 | COL11A1 | SRKGYEKtVIEINtP | T1766 | 0 | 97.93 | 1806 | 0 | 0 | 0 | 1 |
| P12107 | COL11A1 | KtVIEINtPKIDQVP | T1772 | 0.008 | 94.84 | 1806 | 0.005464 | 0.008 | 0.0055 | 1 |
| Q9NPD5 | SLCO1B3 | LAALSFSyIAKALGG | Y38 | 0.106 | 96.28 | 702 | 0.046501 | 0.106 | 0.0465 | 1 |
| Q9NPD5 | SLCO1B3 | GTTFILLtVKIVQPE | T559 | 0.063 | 85.26 | 702 | 0.04303 | 0.156 | 0.1066 | 1 |
| Q9NPD5 | SLCO1B3 | FGALIDKtCMKWstN | T598 | 0.03 | 94.75 | 702 | 0.02049 | 0.03 | 0.0205 | 1 |
| Q9NPD5 | SLCO1B3 | DKtCMKWstNsCGAQ | S603 | 0.084 | 86.34 | 702 | 0.066773 | 0.084 | 0.0668 | 1 |
| O75970 | MPDZ | SSLGMTVsANKDGLG | S1023 | 0.097 | 88.88 | 2070 | 0.077107 | 0.171 | 0.1359 | 1 |
| P54760 | EPHB4 | GGREDLTyALRCREC | Y357 | 0.132 | 96.2 | 987 | 0.057907 | 0.132 | 0.0579 | 1 |
| P14625 | HSP90B1 | IFLRELIsNAsDALD | S106 | 0.08 | 92.56 | 803 | 0.063593 | 0.098 | 0.0779 | 0.8889 |
| P14625 | HSP90B1 | RELIsNAsDALDkIR | S109 | 0.062 | 93.38 | 803 | 0.049285 | 0.062 | 0.0493 | 1 |
| P14625 | HSP90B1 | LsGNEELtVkIKCDk | T133 | 0.118 | 96.76 | 803 | 0.080596 | 0.118 | 0.0806 | 1 |
| P14625 | HSP90B1 | GQFGVGFysAFLVAD | Y200 | 0.091 | 92.72 | 803 | 0.03992 | 0.178 | 0.0767 | 1 |

| | | | | | | | | | | |
|---|---|---|---|---|---|---|---|---|---|---|
| P14625 | HSP90B1 | QFGVGFysAFLVADK | S201 | 0.004 | 92.34 | 803 | 0.00318 | 0.004 | 0.0032 | 1 |
| P14625 | HSP90B1 | EGEVTFKsILFVPts | S385 | 0 | 95.89 | 803 | 0 | 0 | 0 | 1 |
| P14625 | HSP90B1 | FHDMMPKyLNFVkGV | Y429 | 0.011 | 95.31 | 803 | 0.004826 | 0.011 | 0.0048 | 1 |
| P14625 | HSP90B1 | FVkGVVDsDDLPLNV | S439 | 0.018 | 91.38 | 803 | 0.014308 | 0.018 | 0.0143 | 1 |
| P14625 | HSP90B1 | DITSLDQyVERMKEK | Y527 | 0.018 | 98.14 | 803 | 0.007896 | 0.018 | 0.0079 | 1 |
| P14625 | HSP90B1 | KEKQDKIyFMAGssR | Y539 | 0.118 | 97.62 | 803 | 0.051765 | 0.196 | 0.0717 | 1 |
| P14625 | HSP90B1 | sRKEAEssPFVERLL | S552 | 0.04 | 96.01 | 803 | 0.031797 | 0.04 | 0.0318 | 1 |
| P14625 | HSP90B1 | VSQRLtEsPCALVAS | S643 | 0.158 | 95.98 | 803 | 0.125597 | 0.158 | 0.1256 | 1 |
| P14625 | HSP90B1 | yyAsQkktFEINPrH | T684 | 0.083 | 95.48 | 803 | 0.05669 | 0.083 | 0.0567 | 1 |
| P14625 | HSP90B1 | TATLRSGyLLPDTkA | Y727 | 0.147 | 92.32 | 803 | 0.064487 | 0.147 | 0.0645 | 1 |
| Q08211 | DHX9 | VtQPRRIsAVsVAER | S449 | 0.001 | 88.56 | 1270 | 0.000795 | 0.158 | 0.1427 | 0.8333 |
| P36896 | ACVR1B | QRVGTKRyMAPEVLD | Y380 | 0.162 | 98.57 | 505 | 0.071067 | 0.162 | 0.0711 | 1 |
| Q6V702 | CFAP299 | KTEDFEVyFTGKKRL | Y144 | 0.089 | 93.48 | 233 | 0.039043 | 0.089 | 0.039 | 0.8182 |
| Q14997 | PSME4 | WTRKLStyIRLYGRK | Y82 | 0.028 | 94.59 | 1843 | 0.012283 | 0.028 | 0.0123 | 1 |
| Q14997 | PSME4 | TMQKAITyFEIFLPT | Y213 | 0.008 | 97.71 | 1843 | 0.003509 | 0.008 | 0.0035 | 1 |
| Q14997 | PSME4 | AEkIHRQyETIGLDF | Y1087 | 0.045 | 93.43 | 1843 | 0.019741 | 0.045 | 0.0197 | 0.9091 |
| Q92979 | EMG1 | RPDITHQsLLMLMDS | S91 | 0 | 98.88 | 244 | 0 | 0 | 0 | 0.8571 |
| P05026 | ATP1B1 | DPkSYEAyVLNIVRF | Y101 | 0.168 | 94.35 | 303 | 0.073699 | 0.168 | 0.0737 | 1 |
| P05026 | ATP1B1 | EyFGLGNsPGFPLQY | S235 | 0.173 | 87.26 | 303 | 0.13752 | 0.173 | 0.1375 | 1 |
| P05026 | ATP1B1 | GkLLQPKyLQPLLAV | Y254 | 0.091 | 97.02 | 303 | 0.03992 | 0.091 | 0.0399 | 1 |
| P20648 | ATP4A | ETALLKFsELTLGNA | S471 | 0.03 | 94.06 | 1035 | 0.023847 | 0.03 | 0.0238 | 1 |
| P20648 | ATP4A | GIRVIMVtGDHPItA | T628 | 0.015 | 92.61 | 1035 | 0.010245 | 0.015 | 0.0102 | 1 |
| Q6UX71 | PLXDC2 | VKIHGILsNtHRQAA | S150 | 0.074 | 92.73 | 529 | 0.058824 | 0.074 | 0.0588 | 1 |
| Q68EM7 | ARHGAP17 | AQNMQEAstQLEDSL | S82 | 0.102 | 95.33 | 881 | 0.081081 | 0.148 | 0.1239 | 1 |
| Q9Y2H1 | STK38L | KkDTGHIyAMkILRk | Y116 | 0.009 | 95.53 | 464 | 0.003948 | 0.159 | 0.0698 | 1 |
| Q05397 | PTK2 | KPTLNFFyQQVkSDy | Y148 | 0.008 | 98.43 | 1052 | 0.003509 | 0.008 | 0.0035 | 1 |
| Q05397 | PTK2 | yQQVkSDyMLEIADQ | Y155 | 0.048 | 97.3 | 1052 | 0.021057 | 0.048 | 0.0211 | 1 |
| Q05397 | PTK2 | ALEKKSNyEVLEkDV | Y194 | 0.042 | 92.8 | 1052 | 0.018425 | 0.042 | 0.0184 | 1 |
| P24928 | POLR2A | GIVQDTLtAVRKFtK | T543 | 0.049 | 95.97 | 1970 | 0.033468 | 0.095 | 0.0649 | 1 |
| P24928 | POLR2A | LtAVRKFtKRDVFLE | T549 | 0 | 97.02 | 1970 | 0 | 0 | 0 | 1 |
| P24928 | POLR2A | PGNtLRQtFENQVNR | T736 | 0.104 | 80.77 | 1970 | 0.071033 | 0.208 | 0.1421 | 1 |

| | | | | | | | | | | |
|---|---|---|---|---|---|---|---|---|---|---|
| P24928 | POLR2A | SEyNNFksMVVsGAk | S768 | 0.074 | 92.54 | 1970 | 0.058824 | 0.074 | 0.0588 | 1 |
| P24928 | POLR2A | NFksMVVsGAkGskI | S772 | 0 | 93.22 | 1970 | 0 | 0 | 0 | 1 |
| Q8IY82 | DRC7 | EVEPMEHtKKLLYQY | T665 | 0.062 | 80.86 | 874 | 0.042347 | 0.149 | 0.1018 | 1 |
| P28370 | SMARCA1 | GFDRENVyEELRQCV | Y985 | 0.247 | 91.4 | 1054 | 0.108355 | 0.247 | 0.1084 | 0.9231 |
| Q6DD88 | ATL3 | HSCFSDVtCFLLPHP | T261 | 0.06 | 97.6 | 541 | 0.040981 | 0.06 | 0.041 | 1 |
| Q0VDF9 | HSPA14 | EKNGKLRyEIDTGEE | Y100 | 0 | 97 | 509 | 0 | 0 | 0 | 1 |
| O95932 | TGM6 | tLtVSLAsPPsAVIG | S103 | 0.096 | 96.7 | 706 | 0.076312 | 0.096 | 0.0763 | 0.875 |
| O95932 | TGM6 | VIGRYLLsIRLSSHR | S115 | 0.123 | 96.02 | 706 | 0.097775 | 0.123 | 0.0978 | 1 |
| O95932 | TGM6 | SDSRVDItDLYKYPE | T438 | 0.183 | 98.38 | 706 | 0.124991 | 0.183 | 0.125 | 1 |
| Q9HCH3 | CPNE5 | GQSQFNIyEVVNPKK | Y284 | 0.188 | 97.85 | 593 | 0.082473 | 0.188 | 0.0825 | 1 |
| Q9UJX2 | CDC23 | kCCAFNDtREEGkAL | T542 | 0.078 | 83.42 | 597 | 0.053275 | 0.078 | 0.0533 | 1 |
| P53041 | PPP5C | GDFVDRGsFsVEVIL | S277 | 0.06 | 95.36 | 499 | 0.047695 | 0.06 | 0.0477 | 1 |
| P53041 | PPP5C | FVDRGsFsVEVILtL | S279 | 0.002 | 98.59 | 499 | 0.00159 | 0.002 | 0.0016 | 1 |
| P53041 | PPP5C | PMCDLLWsDPQPQNG | S387 | 0 | 98.08 | 499 | 0 | 0 | 0 | 0.9167 |
| Q9NY30 | BTG4 | ATTVFFVtRLVKKHD | T13 | 0.127 | 98.32 | 223 | 0.086743 | 0.127 | 0.0867 | 0.8889 |
| Q969R8 | ITFG2 | TSGKVSVykNDDsRP | Y48 | 0.017 | 97.48 | 447 | 0.007458 | 0.017 | 0.0075 | 1 |
| Q969R8 | ITFG2 | VSQPGCAyAILLCTW | Y207 | 0.017 | 95.72 | 447 | 0.007458 | 0.017 | 0.0075 | 1 |
| Q969R8 | ITFG2 | RDVVLHQtsGRIHNk | T245 | 0.1 | 86.1 | 447 | 0.068301 | 0.183 | 0.1291 | 1 |
| P21281 | ATP6V1B2 | SAIDGMNsIArGQkI | S182 | 0 | 92.53 | 511 | 0 | 0 | 0 | 0.9167 |
| P15144 | ANPEP | ELVEPTEyLVVHLKG | Y161 | 0.02 | 97.73 | 967 | 0.008774 | 0.02 | 0.0088 | 0.9375 |
| P15144 | ANPEP | AVDALAssHPLStPA | S452 | 0.102 | 95.14 | 967 | 0.081081 | 0.13 | 0.1033 | 1 |
| P15144 | ANPEP | AssHPLStPAsEINT | T457 | 0.1 | 93.26 | 967 | 0.068301 | 0.1 | 0.0683 | 1 |
| P15144 | ANPEP | VWIVPItsIRDGRQQ | S590 | 0.006 | 98.2 | 967 | 0.004769 | 0.006 | 0.0048 | 1 |
| O15457 | MSH4 | GVLNyTKtPGGsRRL | T344 | 0.061 | 87.36 | 936 | 0.041664 | 0.061 | 0.0417 | 1 |
| O15457 | MSH4 | QIMAQIGsYVPAEYS | S704 | 0.002 | 97.19 | 936 | 0.00159 | 0.002 | 0.0016 | 1 |
| O15457 | MSH4 | YVPAEYSsFRIAKQI | S712 | 0.014 | 96.63 | 936 | 0.011129 | 0.014 | 0.0111 | 1 |
| Q8WXA8 | HTR3C | MSMGNNLyDQIMFYV | Y234 | 0.327 | 94.28 | 447 | 0.14345 | 0.327 | 0.1434 | 1 |
| Q8N807 | PDILT | SCKGVVEsAALVVWL | S141 | 0.106 | 97.97 | 584 | 0.084261 | 0.106 | 0.0843 | 1 |
| Q8N807 | PDILT | RVIKQHLtDFVIEyN | T255 | 0.049 | 94.19 | 584 | 0.033468 | 0.049 | 0.0335 | 0.8571 |
| Q8N807 | PDILT | IsELHIMsHMLLFVS | S276 | 0.124 | 96.74 | 584 | 0.098569 | 0.124 | 0.0986 | 1 |
| Q8N807 | PDILT | MPSDDITyESLKKFG | Y356 | 0.289 | 97.85 | 584 | 0.12678 | 0.289 | 0.1268 | 1 |

| | | | | | | | | | | |
|---|---|---|---|---|---|---|---|---|---|---|
| Q9H251 | CDH23 | DYETTQAyQLTVNAt | Y201 | 0.269 | 88.15 | 3354 | 0.118006 | 0.3275 | 0.1437 | 1 |
| Q96EB6 | SIRT1 | FRSRDGIyARLAVDF | Y280 | 0.335 | 74.06 | 747 | 0.14696 | 0.335 | 0.147 | 1 |
| Q96EB6 | SIRT1 | QAMFDIEyFRKDPRP | Y301 | 0.294 | 91.5 | 747 | 0.128974 | 0.294 | 0.129 | 1 |
| Q96EB6 | SIRT1 | CHkFIALsDkEGkLL | S333 | 0.098 | 98.51 | 747 | 0.077902 | 0.098 | 0.0779 | 1 |
| Q96EB6 | SIRT1 | GkLLRNYtQNIDTLE | T344 | 0 | 98.28 | 747 | 0 | 0 | 0 | 1 |
| Q96EB6 | SIRT1 | DLLIVIGssLkVRPV | S441 | 0.073 | 95.83 | 747 | 0.058029 | 0.073 | 0.058 | 1 |
| P07814 | EPRS1 | AALLNQHyQVNFkGk | Y224 | 0.039 | 89.08 | 1512 | 0.017109 | 0.269 | 0.1131 | 1 |
| P07814 | EPRS1 | CMRDPtLyRCkIQPH | Y357 | 0.029 | 97.23 | 1512 | 0.012722 | 0.029 | 0.0127 | 1 |
| P07814 | EPRS1 | RkPYIWEySRLNLNN | Y423 | 0.113 | 87.4 | 1512 | 0.049571 | 0.113 | 0.0496 | 0.9286 |
| P07814 | EPRS1 | GLkPVWysPKVFIEG | S547 | 0.008 | 90.07 | 1512 | 0.006359 | 0.008 | 0.0064 | 1 |
| P01591 | JCHAIN | ENIsDPtsPLRTRFV | S77 | 0.102 | 90.77 | 159 | 0.081081 | 0.102 | 0.0811 | 1 |
| Q9UHB7 | AFF4 | TAADKRLtVLCLRCE | T999 | 0.059 | 97.89 | 1163 | 0.040298 | 0.085 | 0.0581 | 1 |
| Q92621 | NUP205 | NVVNIARyLyHGNtN | Y902 | 0.005 | 93.1 | 2012 | 0.002193 | 0.005 | 0.0022 | 0.9333 |
| P61421 | ATP6V0D1 | DRRAFIItINsFGtE | T224 | 0 | 92.25 | 351 | 0 | 0 | 0 | 1 |
| P61421 | ATP6V0D1 | AFIItINsFGtELsk | S227 | 0.133 | 89.01 | 351 | 0.105724 | 0.133 | 0.1057 | 1 |
| Q9NR30 | DDX21 | ATSVDQRsLINsNVG | S628 | 0 | 70.17 | 783 | 0 | 0.079 | 0.0628 | 1 |
| P00450 | CP | RLYKKALyLQYTDET | Y72 | 0.001 | 98.67 | 1065 | 0.000439 | 0.001 | 0.0004 | 1 |
| P29372 | MPG | PEDEAAHsRGGRQTP | S137 | 0 | 98.28 | 298 | 0 | 0 | 0 | 1 |
| P09471 | GNAO1 | LLLGAGEsGkSTIVk | S44 | 0.028 | 97.53 | 354 | 0.022258 | 0.028 | 0.0223 | 1 |
| Q9UHB4 | NDOR1 | MISNQRVtGPSHFQD | T220 | 0.089 | 97.94 | 597 | 0.060788 | 0.089 | 0.0608 | 1 |
| P35236 | PTPN7 | WPtEEEtyGPFQIRI | Y215 | 0.27 | 95.38 | 360 | 0.118445 | 0.27 | 0.1184 | 1 |
| Q9P243 | ZFAT | TDGCDYStPDKYKLQ | T809 | 0.066 | 82.31 | 1243 | 0.045079 | 0.066 | 0.0451 | 1 |
| Q9UN37 | VPS4A | LARKTEGySGADISI | Y329 | 0.192 | 97.65 | 437 | 0.084228 | 0.192 | 0.0842 | 1 |
| Q96G42 | KLHDC7B | FYRLLRYsPVKDAWD | S467 | 0.155 | 97.68 | 594 | 0.123212 | 0.155 | 0.1232 | 1 |
| O15446 | POLR1G | LAGKRHRyRVLSSCP | Y80 | 0.132 | 94.03 | 510 | 0.057907 | 0.132 | 0.0579 | 1 |
| Q9UIC8 | LCMT1 | MDGHILDsKRYAVIG | S162 | 0.063 | 97.53 | 334 | 0.05008 | 0.063 | 0.0501 | 1 |
| Q6ZTQ4 | CDHR3 | SRNIPLSyFLISPPK | Y172 | 0.187 | 92.76 | 885 | 0.082034 | 0.187 | 0.082 | 1 |
| O15439 | ABCC4 | YRKALRLsNMAMGKT | S182 | 0.177 | 84.95 | 1325 | 0.1407 | 0.177 | 0.1407 | 1 |
| Q12931 | TRAP1 | RELIsNAsDALEkLR | S121 | 0.089 | 94.13 | 704 | 0.070747 | 0.089 | 0.0707 | 1 |
| Q12931 | TRAP1 | DAPLNIRsIFyVPDM | S341 | 0 | 97.11 | 704 | 0 | 0.144 | 0.1145 | 1 |
| P09327 | VIL1 | PSGRDPEtPIIVVKQ | T693 | 0.176 | 77.98 | 827 | 0.12021 | 0.114 | 0.136 | 0.875 |

| | | | | | | | | | | |
|---|---|---|---|---|---|---|---|---|---|---|
| P42081 | CD86 | QIKDKGLyQCIIHHK | Y108 | 0.089 | 98.17 | 329 | 0.039043 | 0.089 | 0.039 | 1 |
| P42081 | CD86 | KMSVLLRtKNSTIEY | T175 | 0.211 | 86.67 | 329 | 0.144116 | 0.211 | 0.1441 | 1 |
| Q8NE28 | STKLD1 | CKLQDLssNVLMtDK | S178 | 0.026 | 83.32 | 680 | 0.020668 | 0.026 | 0.0207 | 0.875 |
| P04183 | TK1 | ykCLVIKyAKDTRYs | Y55 | 0.134 | 95.64 | 234 | 0.058784 | 0.261 | 0.112 | 0.8 |
| P04183 | TK1 | VIGGADkyHsVCRLC | Y181 | 0.133 | 96.95 | 234 | 0.058345 | 0.133 | 0.0583 | 0.9375 |
| P04183 | TK1 | GGADkyHsVCRLCYF | S183 | 0.007 | 98.49 | 234 | 0.005564 | 0.007 | 0.0056 | 1 |
| P55822 | SH3BGR | KVFVATSsGSIAIRK | S74 | 0.118 | 90.08 | 239 | 0.0938 | 0.118 | 0.0938 | 1 |
| Q5ST30 | VARS2 | VHPDDSRyTHLHGRQ | Y357 | 0.236 | 96.24 | 1063 | 0.10353 | 0.236 | 0.1035 | 1 |
| Q9NRU3 | CNNM1 | GDCFMLRsDAVLDFA | S444 | 0.184 | 93.67 | 951 | 0.146264 | 0.184 | 0.1463 | 1 |
| P23470 | PTPRG | AILLKDDyFVSGAGL | Y132 | 0.028 | 95.43 | 1445 | 0.012283 | 0.028 | 0.0123 | 0.8235 |
| P23470 | PTPRG | HDRVVSKsAVRDsWN | S320 | 0.029 | 93.96 | 1445 | 0.023053 | 0.029 | 0.0231 | 1 |
| P23470 | PTPRG | SKsAVRDsWNHDMTD | S325 | 0.097 | 77.46 | 1445 | 0.077107 | 0.166 | 0.1419 | 1 |
| Q9Y216 | MTMR7 | DCHDVYIsLIRLARP | S96 | 0.004 | 96.4 | 660 | 0.00318 | 0.004 | 0.0032 | 1 |
| Q9Y216 | MTMR7 | IRKANPGsDFVyVVD | S233 | 0.073 | 97.29 | 660 | 0.058029 | 0.073 | 0.058 | 1 |
| Q13867 | BLMH | QHkyNkLyTVEyLSN | Y284 | 0.105 | 98.84 | 455 | 0.046062 | 0.105 | 0.0461 | 1 |
| P58335 | ANTXR2 | VSPEMRLsFIVFSsQ | S81 | 0 | 96.17 | 489 | 0 | 0 | 0 | 0.8889 |
| Q12860 | CNTN1 | HYGNNFGyIVAFKPF | Y742 | 0 | 96.7 | 1018 | 0 | 0 | 0 | 1 |
| Q8N5U6 | RNF10 | VHKKDLKsVVATESH | S277 | 0.03 | 97.15 | 811 | 0.023847 | 0.03 | 0.0238 | 1 |
| Q96D15 | RCN3 | DAHGNFQyDHEAFLG | Y54 | 0.307 | 92.72 | 328 | 0.134676 | 0.307 | 0.1347 | 1 |
| Q9HBG6 | IFT122 | AVRCLDMsAsRKkLA | S523 | 0.045 | 92.93 | 1241 | 0.035771 | 0.045 | 0.0358 | 1 |
| P48729 | CSNK1A1 | EFIVGGkyKLVRKIG | Y17 | 0.098 | 98.02 | 337 | 0.042991 | 0.098 | 0.043 | 1 |
| P48729 | CSNK1A1 | AHLGIEQsRRDDMES | S199 | 0.011 | 98.78 | 337 | 0.008744 | 0.011 | 0.0087 | 1 |
| P48729 | CSNK1A1 | RFEEAPDyMYLRQLF | Y274 | 0.248 | 98.76 | 337 | 0.108794 | 0.248 | 0.1088 | 1 |
| Q86UA1 | PRPF39 | IQKNLPKsRKVLLEA | S512 | 0 | 96.47 | 669 | 0 | 0 | 0 | 1 |
| P62857 | RPS28 | tGsQGQCtQVRVEFM | T28 | 0.168 | 95.65 | 69 | 0.114746 | 0.168 | 0.1147 | 1 |
| P18887 | XRCC1 | LSFVRFHsPPDKDEA | S151 | 0.163 | 92.18 | 633 | 0.129571 | 0.163 | 0.1296 | 1 |
| Q9UI12 | ATP6V1H | kSGRLEWsPVHkSEk | S367 | 0.174 | 89.65 | 483 | 0.138315 | 0.174 | 0.1383 | 1 |
| P42695 | NCAPD3 | KIPHRVFtLDVVLAL | T412 | 0.01 | 96.7 | 1498 | 0.00683 | 0.01 | 0.0068 | 0.8 |
| P33121 | ACSL1 | PEKIENIyMRSEPVA | Y577 | 0 | 93.99 | 698 | 0 | 0 | 0 | 0.9333 |
| P52294 | KPNA1 | PIDEVIStPGVVARF | T124 | 0.198 | 95.61 | 538 | 0.135237 | 0.198 | 0.1352 | 1 |
| P52294 | KPNA1 | LQTAEFRtRkEAAWA | T396 | 0.007 | 96.71 | 538 | 0.004781 | 0.007 | 0.0048 | 1 |

| | | | | | | | | | | |
|---|---|---|---|---|---|---|---|---|---|---|
| P05771 | PRKCB | EVKNHKFtARFFKQP | T40 | 0.132 | 95.62 | 671 | 0.090158 | 0.158 | 0.1338 | 0.875 |
| Q8WXI4 | ACOT11 | GEGYRNPtEVQMSQL | T45 | 0.091 | 96.29 | 607 | 0.062154 | 0.131 | 0.0838 | 1 |
| P00813 | ADA | MRHQPNWsPkVVELC | S162 | 0.002 | 98.85 | 363 | 0.00159 | 0.002 | 0.0016 | 1 |
| P00813 | ADA | TLEDQALyNRLRQEN | Y249 | 0.011 | 98.84 | 363 | 0.004826 | 0.011 | 0.0048 | 0.9091 |
| P00813 | ADA | RELLDLLyKAyGMPP | Y348 | 0.219 | 98.38 | 363 | 0.096072 | 0.311 | 0.1364 | 1 |
| P11215 | ITGAM | QIQSVVTyDLALDSG | Y676 | 0.057 | 94.44 | 1152 | 0.025005 | 0.057 | 0.025 | 1 |
| P50416 | CPT1A | SQTKNVVsGVLFGTG | S106 | 0 | 89.66 | 773 | 0 | 0 | 0 | 1 |
| P50416 | CPT1A | MGMVKIFsGRkPMLy | S158 | 0.062 | 89.92 | 773 | 0.049285 | 0.062 | 0.0493 | 1 |
| P50416 | CPT1A | SKYLAVEsPFLkEVL | S671 | 0.098 | 96.09 | 773 | 0.077902 | 0.098 | 0.0779 | 1 |
| P00738 | HP | ICLPSKDyAEVGRVG | Y272 | 0.203 | 95.58 | 406 | 0.089053 | 0.203 | 0.0891 | 0.9 |
| Q9Y4F9 | RIPOR2 | RLCPGDQyEIFMKyG | Y223 | 0.054 | 93.79 | 1068 | 0.023689 | 0.3 | 0.1316 | 1 |
| Q9Y4F9 | RIPOR2 | QyEIFMKyGRQRWKL | Y229 | 0.004 | 94.06 | 1068 | 0.001755 | 0.247 | 0.1084 | 1 |
| Q07812 | BAX | tPTWQtVtIFVAGVL | T174 | 0.037 | 92.33 | 192 | 0.025272 | 0.037 | 0.0253 | 1 |
| Q07812 | BAX | VAGVLTAsLtIWKKM | S184 | 0.037 | 94.26 | 192 | 0.029412 | 0.037 | 0.0294 | 1 |
| Q13232 | NME3 | GKNLIHGsDSVEsAR | S137 | 0.011 | 98.54 | 169 | 0.008744 | 0.011 | 0.0087 | 1 |
| Q13232 | NME3 | HGsDSVEsARREIAL | S142 | 0.041 | 97.63 | 169 | 0.032592 | 0.041 | 0.0326 | 0.8571 |
| P80303 | NUCB2 | TkELEKVyDPKNEED | Y278 | 0.043 | 93.68 | 420 | 0.018863 | 0.043 | 0.0189 | 1 |
| Q04637 | EIF4G1 | QKELQALyALQALVV | Y1526 | 0.106 | 94 | 1599 | 0.046501 | 0.106 | 0.0465 | 0.8182 |
| P54577 | YARS1 | HPDADsLyVEkIDVG | Y388 | 0.097 | 97.97 | 528 | 0.042553 | 0.097 | 0.0426 | 1 |
| P54577 | YARS1 | QkMrGVEsQGMLLCA | S436 | 0.077 | 97.53 | 528 | 0.061208 | 0.077 | 0.0612 | 1 |
| P54577 | YARS1 | QGMLLCAsIEGINRQ | S444 | 0.096 | 95.25 | 528 | 0.076312 | 0.121 | 0.1041 | 1 |
| P06732 | CKM | KVLtLELykkLRDKE | Y39 | 0.028 | 98.62 | 381 | 0.012283 | 0.028 | 0.0123 | 1 |
| P06732 | CKM | DPNyVLssRVRTGRs | S129 | 0.01 | 98.03 | 381 | 0.007949 | 0.01 | 0.0079 | 1 |
| P06732 | CKM | GEFkGKyyPLKsMtE | Y174 | 0.315 | 94.07 | 381 | 0.138186 | 0.315 | 0.1382 | 1 |
| O95453 | PARN | YQKLKKHsMDFLLFQ | S61 | 0.01 | 95.38 | 639 | 0.007949 | 0.01 | 0.0079 | 1 |
| O95453 | PARN | KERyIVIskVDEEER | S242 | 0.051 | 90.4 | 639 | 0.040541 | 0.051 | 0.0405 | 1 |
| O75369 | FLNB | EGRtFEMsDFIVDTR | S2021 | 0.063 | 78.24 | 2602 | 0.05008 | 0.063 | 0.0501 | 0.9 |
| O95232 | LUC7L3 | HESVCkYyLCGFCPA | Y38 | 0.334 | 90.01 | 432 | 0.146521 | 0.334 | 0.1465 | 1 |
| P21399 | ACO1 | DFNRRADsLQkNQDL | S138 | 0.001 | 97.16 | 889 | 0.000795 | 0.001 | 0.0008 | 1 |
| P21399 | ACO1 | PHPLVTStDIVLTIT | T258 | 0.04 | 95.16 | 889 | 0.027321 | 0.04 | 0.0273 | 0.9167 |
| P21399 | ACO1 | KLKYIKkyLQAVGMF | Y336 | 0.007 | 98.38 | 889 | 0.003071 | 0.007 | 0.0031 | 1 |

| | | | | | | | | | | |
|---|---|---|---|---|---|---|---|---|---|---|
| P21399 | ACO1 | AGLNVMPyIKTSLSP | Y466 | 0.038 | 97.45 | 889 | 0.01667 | 0.038 | 0.0167 | 1 |
| P21399 | ACO1 | REFNSYGsRrGNDAV | S711 | 0.033 | 91.69 | 889 | 0.026232 | 0.033 | 0.0262 | 1 |
| P21399 | ACO1 | KAVLAEsyERIHRsN | Y800 | 0.3 | 93.34 | 889 | 0.131606 | 0.3 | 0.1316 | 1 |
| P31321 | PRKAR1B | tLRKRkMyEEFLsKV | Y246 | 0.284 | 91.87 | 381 | 0.124587 | 0.284 | 0.1246 | 1 |
| P31321 | PRKAR1B | LNRPRAAtVVArGPL | T338 | 0.039 | 96.78 | 381 | 0.026638 | 0.039 | 0.0266 | 1 |
| Q9NZI8 | IGF2BP1 | VNELQNLtAAEVVVP | T517 | 0.053 | 89.17 | 577 | 0.0362 | 0.053 | 0.0362 | 1 |
| Q52LD8 | RFTN2 | SLEGFFIyEEEGSGV | Y323 | 0 | 97.13 | 501 | 0 | 0 | 0 | 1 |
| Q14108 | SCARB2 | RVEEVGPytYRELRN | Y89 | 0.022 | 98.8 | 478 | 0.009651 | 0.022 | 0.0097 | 1 |
| Q9NW13 | RBM28 | RLIIRNLsFKCSEDD | S122 | 0.115 | 92.14 | 759 | 0.091415 | 0.115 | 0.0914 | 1 |
| P48167 | GLRB | RCkMQLEsFGyTTDD | S203 | 0.03 | 96.75 | 497 | 0.023847 | 0.03 | 0.0238 | 1 |
| P48167 | GLRB | MMGVyAPtLLIVVLS | T277 | 0.02 | 96.61 | 497 | 0.01366 | 0.02 | 0.0137 | 1 |
| O14828 | SCAMP3 | LKRIHsLyRRTGASF | Y304 | 0.243 | 89.13 | 347 | 0.106601 | 0.243 | 0.1066 | 1 |
| P09327 | VIL1 | SFFDGDCyIILAIHK | Y46 | 0.006 | 94.94 | 827 | 0.002632 | 0.006 | 0.0026 | 0.8333 |
| Q9H0R5 | GBP3 | VQKAIAHyDQQMGQK | Y333 | 0.001 | 98.07 | 595 | 0.000439 | 0.001 | 0.0004 | 0.9231 |
| Q9UNK4 | PLA2G2D | GKMPILSyWPYGCHC | Y41 | 0.151 | 98.22 | 145 | 0.066242 | 0.151 | 0.0662 | 0.9286 |
| Q9UNK4 | PLA2G2D | CSIyKDyyRYNFSQG | Y86 | 0.051 | 97.03 | 145 | 0.022373 | 0.051 | 0.0224 | 1 |
| Q16698 | DECR1 | QGKVAFItGGGtGLG | T65 | 0.042 | 98.84 | 335 | 0.028687 | 0.042 | 0.0287 | 1 |
| Q13151 | HNRNPA0 | sRCFGFVtySNVEEA | T54 | 0.124 | 96 | 305 | 0.084694 | 0.124 | 0.0847 | 1 |
| Q13151 | HNRNPA0 | RCFGFVtySNVEEAD | Y55 | 0.038 | 96.71 | 305 | 0.01667 | 0.038 | 0.0167 | 1 |
| P55072 | VCP | NVFIIGAtNRPDIID | T623 | 0.021 | 91.87 | 806 | 0.014343 | 0.021 | 0.0143 | 1 |
| P55072 | VCP | IPLPDEksRVAILkA | S652 | 0.168 | 94.44 | 806 | 0.133546 | 0.168 | 0.1335 | 1 |
| P55072 | VCP | LEFLAkMtNGFsGAD | T679 | 0.018 | 93.43 | 806 | 0.012294 | 0.096 | 0.0656 | 1 |
| P55072 | VCP | AkMtNGFsGADLtEI | S683 | 0.013 | 94.51 | 806 | 0.010334 | 0.099 | 0.0787 | 0.875 |
| Q9UKN7 | MYO15A | AtKLILRyLAAMNQK | Y1331 | 0.007 | 91.11 | 3530 | 0.003071 | 0.007 | 0.0031 | 0.9333 |
| P51884 | LUM | sLEDLQLtHNKITKL | T145 | 0.118 | 98.78 | 338 | 0.080596 | 0.118 | 0.0806 | 1 |
| P51884 | LUM | GLPVSLLtLyLDNNK | T209 | 0.107 | 98.93 | 338 | 0.073082 | 0.164 | 0.112 | 1 |
| P51884 | LUM | ISNIPDEyFkRFNAL | Y224 | 0.22 | 96.95 | 338 | 0.096511 | 0.22 | 0.0965 | 0.9091 |
| P51884 | LUM | ILGPLSYsKIKHLRL | S304 | 0.001 | 96.62 | 338 | 0.000795 | 0.001 | 0.0008 | 1 |
| Q12906 | ILF3 | LELLCEksIGtANRP | S258 | 0.038 | 97.77 | 894 | 0.030207 | 0.038 | 0.0302 | 0.875 |
| Q12906 | ILF3 | RQQREDItQSAQHAL | T315 | 0.072 | 98.61 | 894 | 0.049177 | 0.072 | 0.0492 | 1 |
| P50461 | CSRP3 | TDKDGELyCKVCyAK | Y167 | 0.195 | 90.55 | 194 | 0.085544 | 0.195 | 0.0855 | 1 |

| | | | | | | | | | | |
|---|---|---|---|---|---|---|---|---|---|---|
| P32004 | L1CAM | NLKVkDAtQItQGPR | T518 | 0.08 | 91.87 | 1257 | 0.054641 | 0.08 | 0.0546 | 0.9167 |
| Q96Q91 | SLC4A9 | RKGLSDFsSVLAILL | S666 | 0.036 | 91.09 | 983 | 0.028617 | 0.036 | 0.0286 | 1 |
| Q15166 | PON3 | GTSVASVyHGKILIG | Y336 | 0.245 | 97.38 | 354 | 0.107478 | 0.245 | 0.1075 | 1 |
| P60510 | PPP4C | GDVPETNyLFMGDFV | Y77 | 0.023 | 98.91 | 307 | 0.01009 | 0.023 | 0.0101 | 1 |
| Q8IYD2 | KLHDC8A | AAMGISVtAKDyRVY | T123 | 0.101 | 98.1 | 350 | 0.068984 | 0.101 | 0.069 | 1 |
| P17405 | SMPD1 | PLCCRRGsGLPPAsR | S233 | 0.152 | 97.59 | 631 | 0.120827 | 0.152 | 0.1208 | 0.8889 |
| P17405 | SMPD1 | DGNYSGSsHVVLDHE | S510 | 0.147 | 98.39 | 631 | 0.116852 | 0.147 | 0.1169 | 1 |
| Q96P31 | FCRL3 | KNTHQKVyyKDGKQL | Y133 | 0.011 | 95.08 | 734 | 0.004826 | 0.011 | 0.0048 | 1 |
| Q96P31 | FCRL3 | VSRDNSKyHCTAyRK | Y161 | 0.055 | 93.87 | 734 | 0.024128 | 0.055 | 0.0241 | 1 |
| P63267 | ACTG2 | APrAVFPsIVGRPRH | S34 | 0.016 | 97.66 | 376 | 0.012719 | 0.016 | 0.0127 | 1 |
| P63267 | ACTG2 | GMGQkDsyVGDEAQs | Y54 | 0.037 | 96.13 | 376 | 0.016231 | 0.103 | 0.0452 | 1 |
| P63267 | ACTG2 | DLAGRDLtDyLMkIL | T187 | 0.012 | 98.36 | 376 | 0.008196 | 0.012 | 0.0082 | 0.9 |
| P63267 | ACTG2 | ERFRCPEtLFQPsFI | T261 | 0.003 | 98.43 | 376 | 0.002049 | 0.003 | 0.002 | 0.8889 |
| P63267 | ACTG2 | IDIRkDLyANNVLSG | Y295 | 0.006 | 98.36 | 376 | 0.002632 | 0.006 | 0.0026 | 1 |
| P63267 | ACTG2 | LSGGttMyPGIADRM | Y307 | 0.185 | 98.66 | 376 | 0.081157 | 0.255 | 0.1119 | 0.8571 |
| P63267 | ACTG2 | MWISKPEyDEAGPSI | Y363 | 0.115 | 98.36 | 376 | 0.050449 | 0.193 | 0.0847 | 1 |
| Q6PH85 | DCUN1D2 | KKKLERLyGRYKDPQ | Y69 | 0.078 | 98.13 | 259 | 0.034217 | 0.078 | 0.0342 | 1 |
| Q9BT88 | SYT11 | ELQVSLSyQPVAQRM | Y301 | 0.073 | 97.51 | 431 | 0.032024 | 0.073 | 0.032 | 0.8462 |
| Q9BT88 | SYT11 | TGLSGNPyVKVNVyy | Y331 | 0.209 | 96.78 | 431 | 0.091685 | 0.209 | 0.0917 | 1 |
| Q9BXP5 | SRRT | ETEAVKRyNDyKLDF | Y172 | 0.28 | 88.29 | 876 | 0.122832 | 0.28 | 0.1228 | 1 |
| Q9UHA3 | RSL24D1 | kELTVDNsFEFEkRR | S68 | 0.065 | 92.14 | 163 | 0.051669 | 0.065 | 0.0517 | 1 |
| Q96P11 | NSUN5 | HLAKVLVyELLLGkG | Y76 | 0.211 | 95.9 | 429 | 0.092563 | 0.211 | 0.0926 | 0.8182 |
| Q8IZY2 | ABCA7 | ANPRAGLyLQQMPYP | Y528 | 0.053 | 89.34 | 2146 | 0.02325 | 0.274 | 0.1202 | 1 |
| Q16630 | CPSF6 | HGQNPVVtPCNkQFL | T157 | 0.114 | 93.04 | 551 | 0.077864 | 0.114 | 0.0779 | 1 |
| Q9HCS7 | XAB2 | FEKARDVyEEAIRTV | Y279 | 0.049 | 89.36 | 855 | 0.021496 | 0.049 | 0.0215 | 0.9231 |
| Q8IWZ5 | TRIM42 | PSSRIIEyCRNDNKL | Y301 | 0.26 | 90.07 | 723 | 0.114058 | 0.26 | 0.1141 | 1 |
| Q96T58 | SPEN | FHPKATRtLFIGNLE | T439 | 0.132 | 96.83 | 3664 | 0.090158 | 0.132 | 0.0902 | 0.8 |
| Q96T58 | SPEN | KGMALVLyNEIEYAQ | Y559 | 0.026 | 96.46 | 3664 | 0.011406 | 0.026 | 0.0114 | 1 |
| Q96T58 | SPEN | AQAAVKEtKGRKIGG | T572 | 0.017 | 93.3 | 3664 | 0.011611 | 0.017 | 0.0116 | 1 |
| Q8N1W1 | ARHGEF28 | sCsEAPLyGQMMIYA | Y11 | 0.175 | 88.43 | 1705 | 0.07677 | 0.334 | 0.1465 | 1 |
| P43034 | PAFAH1B1 | GTLIAsCsNDQtVRV | S253 | 0 | 98.36 | 410 | 0 | 0 | 0 | 1 |

| | | | | | | | | | | |
|---|---|---|---|---|---|---|---|---|---|---|
| P43034 | PAFAH1B1 | AsCsNDQtVRVWVVA | T257 | 0.001 | 97.31 | 410 | 0.000683 | 0.001 | 0.0007 | 0.8333 |
| Q5IJ48 | CRB2 | ERPAATtsRWLLWLD | S971 | 0.059 | 87.51 | 1285 | 0.0469 | 0.059 | 0.0469 | 1 |
| Q9NR97 | TLR8 | RLEILDLsFNyIKGS | S345 | 0 | 93.49 | 1041 | 0 | 0 | 0 | 1 |
| Q96MK3 | FAM20A | VPNPWIRsYTLAGKE | S362 | 0.111 | 86.1 | 541 | 0.088236 | 0.111 | 0.0882 | 0.8333 |
| P35658 | NUP214 | AVTSVCWsPKGkQLA | S201 | 0.116 | 95.84 | 2090 | 0.09221 | 0.116 | 0.0922 | 1 |
| Q9NR64 | KLHL1 | GGRDGLktLNtVECY | T519 | 0.145 | 97.01 | 748 | 0.099037 | 0.145 | 0.099 | 1 |
| Q9NR64 | KLHL1 | TVLEGPIyAVGGHDG | Y556 | 0.102 | 98.56 | 748 | 0.044746 | 0.102 | 0.0447 | 1 |
| Q9Y5S1 | TRPV2 | FQHTGIysVMIQKVI | S524 | 0.162 | 85.99 | 764 | 0.128776 | 0.181 | 0.1439 | 1 |
| Q96J66 | ABCC11 | LMQKKGKyAQLIQKM | Y726 | 0.004 | 81.31 | 1382 | 0.001755 | 0.004 | 0.0018 | 1 |
| Q7KZF4 | SND1 | IGkEVCFtIENktPQ | T98 | 0.079 | 94.37 | 910 | 0.053958 | 0.079 | 0.054 | 1 |
| Q7KZF4 | SND1 | GrEyGMIyLGkDTNG | Y113 | 0.19 | 96.32 | 910 | 0.08335 | 0.19 | 0.0833 | 1 |
| Q7KZF4 | SND1 | sEGNGsHtIRDLkyT | T172 | 0.129 | 95.35 | 910 | 0.088109 | 0.129 | 0.0881 | 1 |
| Q7KZF4 | SND1 | WRDyVAPtANLDQkD | T333 | 0.189 | 78.27 | 910 | 0.12909 | 0.189 | 0.1291 | 1 |
| Q7KZF4 | SND1 | kVNVtVDyIRPAsPA | Y421 | 0.019 | 91.08 | 910 | 0.008335 | 0.019 | 0.0083 | 1 |
| Q7KZF4 | SND1 | DDQRSsHyDELLAAE | Y476 | 0.14 | 90.3 | 910 | 0.061416 | 0.14 | 0.0614 | 0.8182 |
| Q7KZF4 | SND1 | FLQrAGRsEAVVEyV | S527 | 0.107 | 96.54 | 910 | 0.085056 | 0.107 | 0.0851 | 1 |
| O94805 | ACTL6B | KEKLPQVsKSWHNYM | S244 | 0.127 | 95.14 | 426 | 0.100954 | 0.127 | 0.101 | 1 |
| P31948 | STIP1 | PHNHVLysNrSAAyA | S42 | 0.002 | 98.21 | 543 | 0.00159 | 0.002 | 0.0016 | 0.875 |
| P31948 | STIP1 | FEEAkrTyEEGLkHE | Y95 | 0.148 | 97.74 | 543 | 0.064925 | 0.25 | 0.1097 | 0.8125 |
| P24821 | TNC | FDRYRLNysLPtGQW | Y1019 | 0.313 | 85.92 | 2201 | 0.137309 | 0.313 | 0.1373 | 1 |
| P24821 | TNC | DRYRLNysLPtGQWV | S1020 | 0.112 | 81.4 | 2201 | 0.08903 | 0.156 | 0.124 | 1 |
| O94829 | IPO13 | CEMLGRMySTIPQAS | Y735 | 0.019 | 96.74 | 963 | 0.008335 | 0.019 | 0.0083 | 1 |
| O60762 | DPM1 | sPRQNkysVLLPTYN | S28 | 0 | 98.83 | 260 | 0 | 0 | 0 | 0.9 |
| Q8NCW5 | NAXE | sLTAPkKsATQFtGR | S247 | 0.015 | 97.76 | 288 | 0.011924 | 0.015 | 0.0119 | 1 |
| Q8NCW5 | NAXE | QFtGRyHyLGGrFVP | Y257 | 0.205 | 98.83 | 288 | 0.089931 | 0.205 | 0.0899 | 1 |
| P13674 | P4HA1 | TEKDLVtsLkDyIkA | S41 | 0.143 | 96.14 | 534 | 0.113673 | 0.143 | 0.1137 | 1 |
| P54920 | NAPA | YEQSADyykGEESNS | Y152 | 0.155 | 97.21 | 295 | 0.067996 | 0.155 | 0.068 | 1 |
| Q6KCM7 | SLC25A25 | ALRktGQysGMLDCA | Y316 | 0.265 | 87.86 | 469 | 0.116252 | 0.336 | 0.1474 | 1 |
| P35221 | CTNNA1 | GVkLVRMsAsQLEAL | S453 | 0.045 | 91.51 | 906 | 0.035771 | 0.045 | 0.0358 | 1 |
| Q5VTT5 | MYOM3 | ASLKVSCtyKEDEGL | T335 | 0 | 85.3 | 1437 | 0 | 0 | 0 | 0.8462 |
| Q14573 | ITPR3 | THGRHVQyLDFLHTV | Y1299 | 0.056 | 88.25 | 2671 | 0.024566 | 0.112 | 0.0491 | 0.9231 |

| | | | | | | | | | | |
|---|---|---|---|---|---|---|---|---|---|---|
| P33176 | KIF5B | EHSSRSHsIFLINVK | S206 | 0.054 | 93.33 | 963 | 0.042925 | 0.054 | 0.0429 | 1 |
| A6NCI4 | VWA3A | tLQPDGGsNLLQALK | S247 | 0.015 | 85.27 | 1184 | 0.011924 | 0.015 | 0.0119 | 1 |
| Q96K12 | FAR2 | NIAIIRPsIVGAtWQ | S230 | 0.066 | 97.22 | 515 | 0.052464 | 0.066 | 0.0525 | 1 |
| Q96K12 | FAR2 | RPsIVGAtWQEPFPG | T235 | 0.012 | 98.26 | 515 | 0.008196 | 0.012 | 0.0082 | 1 |
| Q96K12 | FAR2 | PNGIIIAtGKGFLRA | T257 | 0.077 | 96.64 | 515 | 0.052592 | 0.077 | 0.0526 | 1 |
| Q9H9G7 | AGO3 | DYQPGITyIVVQKRH | Y705 | 0.004 | 98.63 | 860 | 0.001755 | 0.004 | 0.0018 | 1 |
| P60866 | RPS20 | LtSRNVksLEkVCAD | S31 | 0.173 | 91.84 | 119 | 0.13752 | 0.173 | 0.1375 | 1 |
| P60866 | RPS20 | KRLIDLHsPsEIVkQ | S93 | 0.014 | 84.03 | 119 | 0.011129 | 0.014 | 0.0111 | 1 |
| Q96L08 | SUSD3 | GNGAsVGtVLMFRCP | T55 | 0.008 | 93.66 | 255 | 0.005464 | 0.008 | 0.0055 | 1 |
| P08686 | CYP21A2 | VVLNSKRtIEEAMVK | T77 | 0.047 | 98.28 | 494 | 0.032102 | 0.047 | 0.0321 | 1 |
| Q96P15 | SERPINB11 | HKSYLDVsEEGTEAA | S340 | 0.158 | 90.57 | 392 | 0.125597 | 0.158 | 0.1256 | 1 |
| Q15750 | TAB1 | VLLNNKLyVANVGtN | Y179 | 0.09 | 97.94 | 504 | 0.039482 | 0.09 | 0.0395 | 1 |
| Q96PE2 | ARHGEF17 | FLKFLEQsMRENKEK | S1192 | 0.11 | 84.03 | 2063 | 0.087441 | 0.11 | 0.0874 | 1 |
| P53634 | CTSC | TDECAIEsIAVAATP | S452 | 0.172 | 92.24 | 463 | 0.136725 | 0.172 | 0.1367 | 0.8333 |
| Q9BQG0 | MYBBP1A | LLSIIRRsLRssSSk | S859 | 0.003 | 87.36 | 1328 | 0.002385 | 0.003 | 0.0024 | 1 |
| Q5T7W0 | ZNF618 | ANDAGTVsKLCHLFL | S782 | 0.172 | 92.74 | 954 | 0.136725 | 0.172 | 0.1367 | 1 |
| O15230 | LAMA5 | EVGVAVHtPQRAPQQ | T1130 | 0.082 | 89.965 | 3695 | 0.056007 | 0.082 | 0.056 | 1 |
| Q9H1B5 | XYLT2 | KRLLKAVyHEQHFFY | Y256 | 0.07 | 98.69 | 865 | 0.030708 | 0.07 | 0.0307 | 1 |
| Q9H1B5 | XYLT2 | VVELAQGyDNVRVTP | Y284 | 0.217 | 97.3 | 865 | 0.095195 | 0.217 | 0.0952 | 1 |
| Q9H1B5 | XYLT2 | GTPALKAyWENTYDA | Y536 | 0.084 | 98.62 | 865 | 0.03685 | 0.084 | 0.0369 | 1 |
| P61981 | YWHAG | AAAMkNVtELNEPLs | T31 | 0.017 | 97.7 | 247 | 0.011611 | 0.017 | 0.0116 | 1 |
| P61981 | YWHAG | RNLLsVAykNVVGAR | Y49 | 0.055 | 98.67 | 247 | 0.024128 | 0.055 | 0.0241 | 0.8571 |
| P61981 | YWHAG | VESSEkAysEAHEIS | Y154 | 0 | 98.85 | 247 | 0 | 0 | 0 | 1 |
| P61981 | YWHAG | SkEHMQPtHPIRLGL | T168 | 0.067 | 96.5 | 247 | 0.045762 | 0.132 | 0.0899 | 1 |
| P61981 | YWHAG | RLGLALNysVFYYEI | Y179 | 0.061 | 98.8 | 247 | 0.02676 | 0.105 | 0.0419 | 1 |
| P61981 | YWHAG | LGLALNysVFYYEIQ | S180 | 0.002 | 98.64 | 247 | 0.00159 | 0.002 | 0.0016 | 1 |
| Q7Z7E8 | UBE2Q1 | FGANksQysLTRAQQ | Y393 | 0.205 | 93.99 | 422 | 0.089931 | 0.263 | 0.1154 | 0.9375 |
| Q8WV93 | AFG1L | SKPPRGLyVyGDVGt | Y133 | 0.011 | 98.24 | 481 | 0.004826 | 0.011 | 0.0048 | 0.9286 |
| Q8WV93 | AFG1L | yVyGDVGtGKtMVMD | T140 | 0.02 | 90.2 | 481 | 0.01366 | 0.102 | 0.0697 | 0.9 |
| Q5JVL4 | EFHC1 | IyDCDPFtRRYYKEk | T351 | 0.001 | 94.63 | 640 | 0.000683 | 0.001 | 0.0007 | 1 |
| O75643 | SNRNP200 | VDDFKIIyIAPMRsL | Y540 | 0.002 | 96.86 | 2136 | 0.000877 | 0.192 | 0.0842 | 1 |

| | | | | | | | | | | |
|---|---|---|---|---|---|---|---|---|---|---|
| O75643 | SNRNP200 | FVHSRkEtGKTARAI | T731 | 0 | 89.34 | 2136 | 0 | 0 | 0 | 1 |
| O75643 | SNRNP200 | VNWLGyAyLyIRMLR | Y924 | 0 | 96.38 | 2136 | 0 | 0 | 0 | 1 |
| O75643 | SNRNP200 | NSAFESLyQDKFPFF | Y1321 | 0.023 | 93.21 | 2136 | 0.01009 | 0.023 | 0.0101 | 1 |
| O75643 | SNRNP200 | kGNIIIStPEkWDIL | T1428 | 0.088 | 94.16 | 2136 | 0.060105 | 0.088 | 0.0601 | 1 |
| O75643 | SNRNP200 | SRKQTRLtAIDILTT | T1572 | 0.007 | 93.64 | 2136 | 0.004781 | 0.007 | 0.0048 | 1 |
| O75643 | SNRNP200 | KIHAYVDyPIyDVLQ | Y1679 | 0.081 | 92.66 | 2136 | 0.035534 | 0.162 | 0.0711 | 1 |
| O75643 | SNRNP200 | NkQDAVDyLTWTFLY | Y1754 | 0.282 | 93.66 | 2136 | 0.123709 | 0.282 | 0.1237 | 1 |
| O75643 | SNRNP200 | IISNAAEyENIPIRH | Y1855 | 0 | 94.75 | 2136 | 0 | 0 | 0 | 1 |
| O75643 | SNRNP200 | FNDPHVKtNLLLQAH | T1891 | 0 | 96.46 | 2136 | 0 | 0 | 0 | 1 |
| Q96JP0 | FEM1C | DLLGALkyWKKAMNM | Y268 | 0.16 | 98.11 | 617 | 0.07019 | 0.16 | 0.0702 | 1 |
| Q96JP0 | FEM1C | HLAVDkNtTCVGRYP | T494 | 0.043 | 95.88 | 617 | 0.02937 | 0.043 | 0.0294 | 1 |
| P0CG48 | UBC | DyNIQkEstLHLVLr | S521 | 0.18 | 95.32 | 685 | 0.143085 | 0.18 | 0.1431 | 1 |
| P08195 | SLC3A2 | GLkGRLDyLSSLkVk | Y250 | 0.256 | 97.52 | 630 | 0.112304 | 0.256 | 0.1123 | 1 |
| P08195 | SLC3A2 | IRVILDLtPNyRGEN | T309 | 0.001 | 97.31 | 630 | 0.000683 | 0.001 | 0.0007 | 1 |
| P08195 | SLC3A2 | PNyRGENsWFstQVD | S317 | 0.173 | 95.24 | 630 | 0.13752 | 0.173 | 0.1375 | 1 |
| P08195 | SLC3A2 | NKDLLLTssyLsDsG | S402 | 0.108 | 95.45 | 630 | 0.085851 | 0.108 | 0.0859 | 1 |
| P08195 | SLC3A2 | KDLLLTssyLsDsGs | S403 | 0.155 | 94.59 | 630 | 0.123212 | 0.155 | 0.1232 | 1 |
| P08195 | SLC3A2 | DLLLTssyLsDsGsT | Y404 | 0.172 | 94.65 | 630 | 0.075454 | 0.172 | 0.0755 | 1 |
| P08195 | SLC3A2 | GQsEDPGsLLsLFRR | S518 | 0.048 | 96.94 | 630 | 0.038156 | 0.048 | 0.0382 | 1 |
| P08195 | SLC3A2 | LsLFRRLsDQRsKER | S527 | 0.027 | 98.22 | 630 | 0.021463 | 0.027 | 0.0215 | 1 |
| P08195 | SLC3A2 | AKADLLLsTQPGREE | S598 | 0.118 | 97.04 | 630 | 0.0938 | 0.118 | 0.0938 | 1 |
| O60909 | B4GALT2 | GVLMGGRyTPPDCTP | Y134 | 0.235 | 98.51 | 372 | 0.103091 | 0.235 | 0.1031 | 1 |
| Q9Y6N6 | LAMC3 | ADGPWEPyQFYSASC | Y160 | 0 | 94.94 | 1575 | 0 | 0 | 0 | 1 |
| O14908 | GIPC1 | RPRLVFHtQLAHGsP | T62 | 0 | 97.6 | 333 | 0 | 0 | 0 | 1 |
| Q9HD40 | SEPSECS | SSLLNKItNsLVLDI | T118 | 0.013 | 98.71 | 501 | 0.008879 | 0.013 | 0.0089 | 1 |
| P30876 | POLR2B | LQTQHQktFIGkIPI | T147 | 0.021 | 93.2 | 1174 | 0.014343 | 0.021 | 0.0143 | 1 |
| P30876 | POLR2B | kDSkyAytGECRSCL | T218 | 0.031 | 94.65 | 1174 | 0.021173 | 0.031 | 0.0212 | 1 |
| P30876 | POLR2B | SHLRRLNsPIGRDGk | S487 | 0.077 | 90.84 | 1174 | 0.061208 | 0.077 | 0.0612 | 1 |
| P30876 | POLR2B | QGMRHAIyDkLDDDG | Y845 | 0.133 | 93.68 | 1174 | 0.058345 | 0.224 | 0.0833 | 1 |
| P30876 | POLR2B | DDVIIGktVtLPENE | T870 | 0.03 | 94.55 | 1174 | 0.02049 | 0.03 | 0.0205 | 1 |
| P30876 | POLR2B | QIGDkFAsRHGQkGT | S937 | 0 | 95.3 | 1174 | 0 | 0 | 0 | 1 |

| | | | | | | | | | | |
|---|---|---|---|---|---|---|---|---|---|---|
| P30876 | POLR2B | AVNVQkIsNLLsDyG | S1012 | 0.006 | 94.76 | 1174 | 0.004769 | 0.006 | 0.0048 | 1 |
| Q7L576 | CYFIP1 | QPNAQPQyLHGSKAL | Y887 | 0.265 | 97.58 | 1253 | 0.116252 | 0.265 | 0.1163 | 1 |
| Q8N4S9 | MARVELD2 | NVALFILyMAAAIVY | Y304 | 0.028 | 95.73 | 558 | 0.012283 | 0.238 | 0.1044 | 0.9286 |
| P54646 | PRKAA2 | tsCGsPNyAAPEVIS | Y179 | 0.207 | 97.31 | 552 | 0.090808 | 0.207 | 0.0908 | 1 |
| Q15437 | SEC23B | MKKATkHyEMLANRT | Y333 | 0 | 98.82 | 767 | 0 | 0 | 0 | 1 |
| P42704 | LRPPRC | LQkLGAVyDVSHyNA | Y160 | 0.269 | 83.93 | 1394 | 0.118006 | 0.269 | 0.118 | 0.8333 |
| P42704 | LRPPRC | GIEPGPDtyLALLNA | T269 | 0.001 | 86.32 | 1394 | 0.000683 | 0.001 | 0.0007 | 0.8333 |
| P30307 | CDC25C | LYILkGGyRDFFPEY | Y416 | 0 | 98.75 | 473 | 0 | 0 | 0 | 0.8947 |
| O75691 | UTP20 | LCMTATVsHILDQRE | S1131 | 0.033 | 94.43 | 2785 | 0.026232 | 0.033 | 0.0262 | 1 |
| Q9Y2C9 | TLR6 | FHAFIsysEHDsAWV | S649 | 0.007 | 90.3 | 796 | 0.005564 | 0.007 | 0.0056 | 1 |
| Q8N9I5 | FADS6 | NVVGLGDsstWRLPC | S155 | 0.096 | 81.55 | 356 | 0.076312 | 0.096 | 0.0763 | 1 |
| Q8N9I5 | FADS6 | PCLNRyVyMFLAPFL | Y168 | 0.013 | 96.6 | 356 | 0.005703 | 0.013 | 0.0057 | 1 |
| Q8N8M0 | NAT16 | NVIDAGEtVLVEGLR | T118 | 0 | 98.65 | 369 | 0 | 0 | 0 | 0.8 |
| Q96EN8 | MOCOS | VVGMRNVtMAINVIs | T162 | 0.001 | 95.55 | 888 | 0.000683 | 0.001 | 0.0007 | 1 |
| Q96QB1 | DLC1 | TSEQAELsYKKVSEG | S1355 | 0.01 | 93.3 | 1528 | 0.007949 | 0.01 | 0.0079 | 1 |
| P26951 | IL3RA | FDDISRLssGSQSSH | S174 | 0.027 | 92.13 | 378 | 0.021463 | 0.027 | 0.0215 | 1 |
| P26951 | IL3RA | HFNRKFRyELQIQKR | Y238 | 0.109 | 97.6 | 378 | 0.047817 | 0.109 | 0.0478 | 1 |
| Q9BRP4 | PAAF1 | HkGGILDtAIVDrGR | T182 | 0 | 98.84 | 392 | 0 | 0 | 0 | 1 |
| Q9BRP4 | PAAF1 | DrGRNVVsAsRDGtA | S193 | 0 | 98.89 | 392 | 0 | 0 | 0 | 1 |
| Q9BRP4 | PAAF1 | GRNVVsAsRDGtARL | S195 | 0 | 98.9 | 392 | 0 | 0 | 0 | 1 |
| Q9BRP4 | PAAF1 | VsAsRDGtARLWDCG | T199 | 0.127 | 98.85 | 392 | 0.086743 | 0.127 | 0.0867 | 1 |
| Q8N2K0 | ABHD12 | FDWIkARsGDNPVyI | S235 | 0.009 | 96.51 | 398 | 0.007154 | 0.009 | 0.0072 | 1 |
| Q8N2K0 | ABHD12 | sLGtGVAtNLVRRLC | T253 | 0 | 98.62 | 398 | 0 | 0 | 0 | 1 |
| Q8N2K0 | ABHD12 | PDALILEsPFTNIRE | S273 | 0 | 98.25 | 398 | 0 | 0 | 0 | 1 |
| Q2TBA0 | KLHL40 | APMQTARsLFGATVH | S501 | 0.034 | 98.46 | 621 | 0.027027 | 0.034 | 0.027 | 1 |
| Q2TBA0 | KLHL40 | EAFPQERsSLSLVSL | S548 | 0.164 | 97.13 | 621 | 0.130366 | 0.164 | 0.1304 | 1 |
| Q13242 | SRSF9 | DGVGMVEyLrkEDME | Y152 | 0 | 85.47 | 221 | 0 | 0.047 | 0.0206 | 1 |
| Q9NR20 | DYRK4 | LSIVRRFtLsVLKCL | T211 | 0.017 | 97.92 | 520 | 0.011611 | 0.017 | 0.0116 | 1 |
| Q9NR20 | DYRK4 | IVRRFtLsVLKCLQM | S213 | 0.156 | 95.91 | 520 | 0.124007 | 0.156 | 0.124 | 1 |
| Q9UQ26 | RIMS2 | VGHQLIVtILGAKDL | T825 | 0.158 | 96.7 | 1411 | 0.107916 | 0.158 | 0.1079 | 1 |
| Q9HAW0 | BRF2 | SPSVPAkyVEDKEkM | Y193 | 0.203 | 95.6 | 419 | 0.089053 | 0.203 | 0.0891 | 1 |

| | | | | | | | | | | |
|---|---|---|---|---|---|---|---|---|---|---|
| P12109 | COL6A1 | LSLVKENyAELLEDA | Y793 | 0.038 | 92.24 | 1028 | 0.01667 | 0.038 | 0.0167 | 0.8571 |
| Q12809 | KCNH2 | GDLLTALyFISRGSI | Y780 | 0.199 | 94.19 | 1159 | 0.087298 | 0.199 | 0.0873 | 1 |
| P08069 | IGF1R | DGVFTTYsDVWSFGV | S1193 | 0.034 | 87.8 | 1367 | 0.027027 | 0.034 | 0.027 | 1 |
| P08069 | IGF1R | EPGFREVsFyySEEN | S1278 | 0.061 | 92.09 | 1367 | 0.04849 | 0.061 | 0.0485 | 1 |
| P0CG48 | UBC | DyNIQkEstLHLVLr | S445 | 0.185 | 96.16 | 685 | 0.147059 | 0.185 | 0.1471 | 0.8571 |
| P49419 | ALDH7A1 | RQAsVADyEETVkkA | Y88 | 0.079 | 98.85 | 539 | 0.034656 | 0.079 | 0.0347 | 1 |
| P49419 | ALDH7A1 | QVLGsLVsLEMGKIL | S133 | 0.002 | 98.84 | 539 | 0.00159 | 0.002 | 0.0016 | 1 |
| P49419 | ALDH7A1 | EVkQGLSsSIFTkDL | S457 | 0 | 98.92 | 539 | 0 | 0 | 0 | 1 |
| C9J1S8 | TRIM49D1 | FGDILHRsEsVLLHM | S255 | 0.015 | 92.56 | 452 | 0.011924 | 0.015 | 0.0119 | 1 |
| P24386 | CHM | FLTFCMEyEKYPDEY | Y305 | 0.123 | 94.15 | 653 | 0.053958 | 0.123 | 0.054 | 1 |
| O94844 | RHOBTB1 | SMQAVLDyLyTKQLS | Y542 | 0.068 | 93.48 | 696 | 0.029831 | 0.183 | 0.0803 | 0.9091 |
| P34897 | SHMT2 | ERVLELVsITANkNT | S404 | 0.178 | 98.73 | 504 | 0.141495 | 0.178 | 0.1415 | 0.8889 |
| Q8IWB7 | WDFY1 | PkEDGVItASEDrTI | T41 | 0 | 98.56 | 410 | 0 | 0 | 0 | 1 |
| Q8TEY7 | USP33 | sLLQFyIsRQWLNKF | S743 | 0.013 | 93.4 | 942 | 0.010334 | 0.013 | 0.0103 | 1 |
| Q8TEY7 | USP33 | WNFLQsIyGGGPEVI | Y909 | 0.244 | 94.46 | 942 | 0.107039 | 0.244 | 0.107 | 1 |
| Q68CR1 | SEL1L3 | GFFGPLKyYRLRSLH | Y420 | 0 | 96.74 | 1132 | 0 | 0 | 0 | 0.8667 |
| O15344 | MID1 | RFTSQGSyGVAGNVF | Y528 | 0.228 | 92.21 | 667 | 0.10002 | 0.228 | 0.1 | 1 |
| Q96P69 | GPR78 | GVLLVNLsLGHLLLA | S49 | 0.05 | 98.49 | 363 | 0.039746 | 0.05 | 0.0397 | 1 |
| Q96P69 | GPR78 | LASNAALsVAALsAD | S96 | 0.022 | 96.29 | 363 | 0.017488 | 0.022 | 0.0175 | 1 |
| Q96P69 | GPR78 | ALsVAALsADQWLAV | S101 | 0.073 | 90.63 | 363 | 0.058029 | 0.124 | 0.0986 | 0.8889 |
| Q9NUZ1 | ACOXL | TKNFVSRsLVIGEVL | S36 | 0.169 | 90.33 | 547 | 0.134341 | 0.169 | 0.1343 | 1 |
| Q9NUZ1 | ACOXL | LVIGEVLsMADMATG | S44 | 0.004 | 94.25 | 547 | 0.00318 | 0.004 | 0.0032 | 1 |
| Q9H2M9 | RAB3GAP2 | QDCVLSLsPTNDLMV | S83 | 0.003 | 94.08 | 1393 | 0.002385 | 0.003 | 0.0024 | 0.8889 |
| A2PYH4 | HFM1 | VICAPTGsGKTVVFE | S307 | 0.024 | 79.36 | 1435 | 0.019078 | 0.024 | 0.0191 | 1 |
| A2PYH4 | HFM1 | TATAVIMtRLSTRDK | T670 | 0.003 | 94.61 | 1435 | 0.002049 | 0.003 | 0.002 | 1 |
| O43765 | SGTA | DPAyskAyGRMGLAL | Y162 | 0.216 | 98.64 | 313 | 0.094756 | 0.266 | 0.1167 | 1 |
| Q9UHX1 | PUF60 | RkVVAEVyDQERFDN | Y547 | 0.098 | 88.55 | 559 | 0.042991 | 0.234 | 0.1027 | 0.8182 |
| P11509 | CYP2A6 | LKNLVMTtLNLFIGG | T295 | 0 | 98.88 | 494 | 0 | 0 | 0 | 1 |
| P19838 | NFKB1 | MRNDLyQtPLHLAVI | T584 | 0.016 | 94.29 | 968 | 0.010928 | 0.016 | 0.0109 | 1 |
| P33261 | CYP2C19 | WKEIRRFsLMTLRNF | S127 | 0.004 | 97.51 | 490 | 0.00318 | 0.004 | 0.0032 | 0.9 |
| P98175 | RBM10 | QKVSMHYsDPKPKIN | S207 | 0.072 | 82.79 | 930 | 0.057234 | 0.072 | 0.0572 | 1 |

| | | | | | | | | | | |
|---|---|---|---|---|---|---|---|---|---|---|
| Q96RW7 | HMCN1 | PKPKIAWtVNDMFIV | T646 | 0.083 | 89.465 | 5635 | 0.05669 | 0.128 | 0.0967 | 1 |
| Q96RW7 | HMCN1 | NQFGKIQsETTVtVT | S878 | 0.116 | 89.75 | 5635 | 0.09221 | 0.116 | 0.0922 | 1 |
| Q712K3 | UBE2R2 | VPTTLAEyCIkTkVP | Y190 | 0.273 | 90.65 | 238 | 0.119761 | 0.273 | 0.1198 | 1 |
| Q00013 | MPP1 | ESAGLIPsPELQEWR | S220 | 0 | 94.37 | 466 | 0 | 0 | 0 | 1 |
| Q8N884 | CGAS | EYSNTRAyYFVKFKR | Y248 | 0.262 | 95.86 | 522 | 0.114936 | 0.262 | 0.1149 | 0.8182 |
| Q8N884 | CGAS | DCLkLMkyLLEQLkE | Y415 | 0.128 | 96.17 | 522 | 0.056152 | 0.263 | 0.1154 | 1 |
| Q8IY17 | PNPLA6 | VLNGRLRsVIQRGSG | S676 | 0.009 | 85.18 | 1375 | 0.007154 | 0.009 | 0.0072 | 1 |
| Q8IY17 | PNPLA6 | LAkLPEGtLGHIKRR | T727 | 0.001 | 89.42 | 1375 | 0.000683 | 0.03 | 0.0205 | 1 |
| P0C672 | TSPAN19 | IDFVISEyGSKDKPE | Y129 | 0.011 | 94.11 | 248 | 0.004826 | 0.011 | 0.0048 | 1 |
| O00187 | MASP2 | YDFLKIQtDREEHGP | T250 | 0.189 | 95.25 | 686 | 0.12909 | 0.189 | 0.1291 | 1 |
| O00187 | MASP2 | TyKAVIQySCEETFy | Y394 | 0.075 | 95.1 | 686 | 0.032901 | 0.075 | 0.0329 | 1 |
| Q86UX2 | ITIH5 | EIRTPLLsDIRIDYP | S484 | 0.146 | 91.81 | 942 | 0.116058 | 0.146 | 0.1161 | 1 |
| Q9NYG2 | ZDHHC3 | MFGTQVHsICTDETG | S241 | 0.127 | 95.89 | 299 | 0.100954 | 0.127 | 0.101 | 0.8889 |
| Q9UGI6 | KCNN3 | ADVDIILsIPMFLRL | S408 | 0.004 | 91.58 | 731 | 0.00318 | 0.004 | 0.0032 | 1 |
| Q96C45 | ULK4 | TEHNPtFtRLVEESK | T1023 | 0.023 | 84.63 | 1275 | 0.015709 | 0.038 | 0.026 | 1 |
| Q9NTJ5 | SACM1L | NQEGVFRsNCMDCLD | S387 | 0 | 96.37 | 587 | 0 | 0 | 0 | 0.8 |
| Q9UJ04 | TSPYL4 | RLHMQRRsFIIQNIP | S244 | 0.114 | 96.72 | 414 | 0.09062 | 0.186 | 0.1479 | 1 |
| Q14410 | GK2 | GREKPAyyALEGSVA | Y316 | 0.189 | 97.66 | 553 | 0.082912 | 0.189 | 0.0829 | 1 |
| Q14410 | GK2 | ATESEIRyATWKKAV | Y500 | 0.162 | 96.77 | 553 | 0.071067 | 0.162 | 0.0711 | 1 |
| Q93063 | EXT2 | VVRLPADsPIPERGD | S75 | 0.116 | 92.5 | 718 | 0.09221 | 0.116 | 0.0922 | 1 |
| Q86T03 | PIP4P1 | CGVCNEAtPIKNAPP | T111 | 0.043 | 87.4 | 277 | 0.02937 | 0.043 | 0.0294 | 0.9 |
| P10745 | RBP3 | AIVVGERtGGGALDL | T259 | 0.081 | 94.22 | 1247 | 0.055324 | 0.081 | 0.0553 | 1 |
| Q9BXT6 | MOV10L1 | NPEFEQAyNFEPMDV | Y624 | 0.187 | 87.23 | 1211 | 0.082034 | 0.187 | 0.082 | 1 |
| Q9Y450 | HBS1L | EERERGVtMDVGMTk | T325 | 0.117 | 89.6 | 684 | 0.079913 | 0.117 | 0.0799 | 1 |
| Q4KMP7 | TBC1D10B | sCQGMyEtMEQLRNL | T579 | 0 | 95.49 | 808 | 0 | 0 | 0 | 1 |
| Q96Q11 | TRNT1 | LFTMKLQsPEFQSLF | S34 | 0.16 | 94.47 | 434 | 0.127186 | 0.16 | 0.1272 | 1 |
| P23396 | RPS3 | rELAEDGysGVEVRV | Y34 | 0.058 | 95.56 | 243 | 0.025444 | 0.058 | 0.0254 | 0.8462 |
| P23396 | RPS3 | RFGFPEGsVELyAEk | S83 | 0.139 | 91.98 | 243 | 0.110493 | 0.139 | 0.1105 | 1 |
| P23396 | RPS3 | sGDPVNyyVDtAVRH | Y167 | 0.09 | 95.98 | 243 | 0.039482 | 0.09 | 0.0395 | 1 |
| P47211 | GALR1 | AIVHSRRssSLRVSR | S143 | 0.163 | 89.6 | 349 | 0.129571 | 0.163 | 0.1296 | 1 |
| O43294 | TGFB1I1 | HFHARRGsLCAtCGL | S403 | 0.096 | 89.31 | 461 | 0.076312 | 0.096 | 0.0763 | 0.8333 |

| | | | | | | | | | | |
|---|---|---|---|---|---|---|---|---|---|---|
| Q16762 | TST | RVLDASWysPGTREA | Y37 | 0.284 | 97.65 | 297 | 0.124587 | 0.284 | 0.1246 | 1 |
| Q16762 | TST | PDAVGLDsGHIRGAV | S202 | 0.159 | 97.34 | 297 | 0.126391 | 0.159 | 0.1264 | 1 |
| P42229 | STAT5A | LWNLkPFttRDFSIR | T647 | 0.172 | 97.09 | 794 | 0.117478 | 0.172 | 0.1175 | 1 |
| Q8N543 | OGFOD1 | RSWLsDIsKIDLEST | S131 | 0.169 | 96.08 | 542 | 0.134341 | 0.169 | 0.1343 | 1 |
| P16157 | ANK1 | VLSKTGFtPLHIAAH | T208 | 0 | 92.29 | 1881 | 0 | 0 | 0 | 1 |
| P16157 | ANK1 | AKAKDDQtPLHCAAR | T472 | 0.002 | 97.46 | 1881 | 0.001366 | 0.002 | 0.0014 | 1 |
| P16157 | ANK1 | NAKTKLGysPLHQAA | Y735 | 0.108 | 90.21 | 1881 | 0.047378 | 0.315 | 0.1103 | 1 |
| P16157 | ANK1 | AKTKLGysPLHQAAQ | S736 | 0.018 | 93.58 | 1881 | 0.014308 | 0.018 | 0.0143 | 1 |
| Q9NY56 | OBP2A | GNLEAtFtFMREDRC | T67 | 0.121 | 97.8 | 170 | 0.082645 | 0.121 | 0.0826 | 0.8889 |
| P49368 | CCT3 | ALDDMIstLkkISIP | T136 | 0.197 | 94.43 | 545 | 0.134554 | 0.197 | 0.1346 | 1 |
| P49368 | CCT3 | KEIDIKKyARVEkIP | Y198 | 0.294 | 91.26 | 545 | 0.128974 | 0.294 | 0.129 | 1 |
| P49368 | CCT3 | PRIVLLDssLEykkG | S243 | 0.113 | 93.14 | 545 | 0.089825 | 0.113 | 0.0898 | 1 |
| P49368 | CCT3 | yLMRANItAIrRVRk | T310 | 0.003 | 96.53 | 545 | 0.002049 | 0.003 | 0.002 | 1 |
| P49368 | CCT3 | VkLQTYktAVEtAVL | T508 | 0.181 | 96.62 | 545 | 0.123625 | 0.181 | 0.1236 | 1 |
| P49368 | CCT3 | TYktAVEtAVLLLRI | T512 | 0.118 | 95.75 | 545 | 0.080596 | 0.118 | 0.0806 | 1 |
| Q9BYC2 | OXCT2 | VKDLQVVssNVGVED | S89 | 0.063 | 98.39 | 517 | 0.05008 | 0.063 | 0.0501 | 1 |
| Q9BYC2 | OXCT2 | KDLQVVssNVGVEDF | S90 | 0 | 97.22 | 517 | 0 | 0 | 0 | 1 |
| Q96IK5 | GMCL1 | TVCGYyTsAGTYGLD | S219 | 0.132 | 98.22 | 515 | 0.104929 | 0.132 | 0.1049 | 1 |
| Q96IK5 | GMCL1 | MQVEMDIytALKKWM | Y276 | 0 | 98.47 | 515 | 0 | 0 | 0 | 1 |
| P61163 | ACTR1A | LFSNIVLsGGSTLFk | S301 | 0.014 | 97.89 | 376 | 0.011129 | 0.014 | 0.0111 | 1 |
| Q8N0W3 | FCSK | DGRVPLVsGVVFFsV | S223 | 0.009 | 94.2 | 1084 | 0.007154 | 0.009 | 0.0072 | 1 |
| Q8N0W3 | FCSK | VsGVVFFsVEtAERL | S229 | 0.107 | 97.84 | 1084 | 0.085056 | 0.107 | 0.0851 | 1 |
| Q8N0W3 | FCSK | VVFFsVEtAERLLAT | T232 | 0.003 | 97.36 | 1084 | 0.002049 | 0.003 | 0.002 | 1 |
| O15327 | INPP4B | VRDRKLNtLVQIsVI | T61 | 0 | 92.22 | 924 | 0 | 0 | 0 | 0.8182 |
| O15327 | INPP4B | LNtLVQIsVIHPVEQ | S66 | 0.044 | 95.96 | 924 | 0.034976 | 0.044 | 0.035 | 0.8571 |
| Q9H5Y7 | SLITRK6 | IEVLEEGsFMNLTRL | S406 | 0.039 | 93.53 | 841 | 0.031002 | 0.039 | 0.031 | 1 |
| O60880 | SH2D1A | LAtGLDGsyLLRDSE | S28 | 0.006 | 96.94 | 128 | 0.004769 | 0.006 | 0.0048 | 1 |
| Q13421 | MSLN | PGRFVAEsAEVLLPR | S200 | 0.064 | 95.63 | 630 | 0.050875 | 0.064 | 0.0509 | 1 |
| Q9BPW8 | NIPSNAP1 | KPEyLDAyNsLTEAV | Y87 | 0.022 | 98.45 | 284 | 0.009651 | 0.022 | 0.0097 | 1 |
| Q9BPW8 | NIPSNAP1 | PRMGPNIyELrTYkL | Y185 | 0.008 | 97.84 | 284 | 0.003509 | 0.008 | 0.0035 | 0.8571 |
| Q14674 | ESPL1 | REAAPQDyEAVARGS | Y140 | 0.105 | 87.48 | 2120 | 0.046062 | 0.105 | 0.0461 | 1 |

| | | | | | | | | | | |
|---|---|---|---|---|---|---|---|---|---|---|
| Q9HBA9 | FOLH1B | YTLRVDCtPLMYSLV | T159 | 0.004 | 97.97 | 442 | 0.002732 | 0.004 | 0.0027 | 1 |
| Q9HBA9 | FOLH1B | YSLVYNLtKELKSPD | T170 | 0.004 | 98.71 | 442 | 0.002732 | 0.004 | 0.0027 | 1 |
| Q9HBA9 | FOLH1B | YAVVLRKyADKIYNI | Y299 | 0.066 | 95.94 | 442 | 0.028953 | 0.066 | 0.029 | 1 |
| Q8TC07 | TBC1D15 | QKQkTDEyFRMkLQW | Y383 | 0 | 98.84 | 691 | 0 | 0 | 0 | 0.9231 |
| Q8TC07 | TBC1D15 | LCkAEAIsLQMVkCK | S611 | 0.029 | 96.49 | 691 | 0.023053 | 0.029 | 0.0231 | 1 |
| Q5T7B8 | KIF24 | ITVEDkEtLLVHEkK | T253 | 0.073 | 92.57 | 1368 | 0.04986 | 0.097 | 0.0663 | 0.8 |
| Q9UKX5 | ITGA11 | AYLGYTVtsVVsSRQ | T430 | 0.114 | 96.85 | 1188 | 0.077864 | 0.114 | 0.0779 | 1 |
| Q9UKX5 | ITGA11 | YLGYTVtsVVsSRQG | S431 | 0.07 | 96.01 | 1188 | 0.055644 | 0.07 | 0.0556 | 1 |
| Q9UKX5 | ITGA11 | YTVtsVVsSRQGRVY | S434 | 0.133 | 84.57 | 1188 | 0.105724 | 0.133 | 0.1057 | 1 |
| Q04759 | PRKCQ | RMLMNARyFLEMSDT | Y121 | 0.328 | 87.83 | 706 | 0.143889 | 0.328 | 0.1439 | 1 |
| Q96PN6 | ADCY10 | AARMMMYyPGIVTCD | Y421 | 0.135 | 93.49 | 1610 | 0.059223 | 0.158 | 0.0693 | 1 |
| Q9Y5S2 | CDC42BPB | TSSLLILtENENEkR | T1196 | 0.05 | 94.46 | 1711 | 0.034151 | 0.201 | 0.1373 | 1 |
| Q9UQV4 | LAMP3 | FNIDPNAtQAsGNCG | T268 | 0.186 | 98.21 | 416 | 0.127041 | 0.186 | 0.127 | 1 |
| Q5JSJ4 | INTS6L | RTDLGTSyLDIAKGA | Y24 | 0.135 | 95.3 | 861 | 0.059223 | 0.135 | 0.0592 | 0.8 |
| Q6ZUM4 | ARHGAP27 | LPAPNHDtLRMLFQH | T821 | 0 | 97.67 | 889 | 0 | 0 | 0 | 1 |
| Q96PL1 | SCGB3A2 | NELGPEAsEAVKKLL | S79 | 0.119 | 90.87 | 93 | 0.094595 | 0.119 | 0.0946 | 1 |
| Q8WYQ5 | DGCR8 | PSEPFGAsVtIDGVT | S545 | 0.146 | 93.29 | 773 | 0.116058 | 0.146 | 0.1161 | 0.8889 |
| P62424 | RPL7A | RAGVNtVttLVENkK | T144 | 0.191 | 97.82 | 266 | 0.130456 | 0.191 | 0.1305 | 1 |
| P62424 | RPL7A | CRkMGVPyCIIkGKA | Y181 | 0.012 | 97.86 | 266 | 0.005264 | 0.012 | 0.0053 | 1 |
| P62424 | RPL7A | rTNYNDryDEIrrHW | Y230 | 0.262 | 93.8 | 266 | 0.114936 | 0.262 | 0.1149 | 1 |
| Q99619 | SPSB2 | GWNPKDCsENIEVKE | S54 | 0.052 | 97.5 | 263 | 0.041336 | 0.052 | 0.0413 | 1 |
| Q99619 | SPSB2 | EVKEGGLyFERRPVA | Y65 | 0.261 | 98.57 | 263 | 0.114497 | 0.261 | 0.1145 | 1 |
| P12259 | F5 | KSLSSEMyVKSYTIH | Y2149 | 0.234 | 92.04 | 2224 | 0.102652 | 0.234 | 0.1027 | 1 |
| P12259 | F5 | KSYTIHYsEQGVEWK | S2158 | 0.03 | 92.26 | 2224 | 0.023847 | 0.03 | 0.0238 | 1 |
| P12259 | F5 | FFNPPIIsRFIRVIP | S2198 | 0.001 | 88.55 | 2224 | 0.000795 | 0.001 | 0.0008 | 1 |
| P12259 | F5 | IPKTWNQsIALRLEL | S2211 | 0.157 | 89.42 | 2224 | 0.124802 | 0.157 | 0.1248 | 1 |
| Q6UXH1 | CRELD2 | TLSKyEssEIRLLEI | S71 | 0.094 | 90.09 | 353 | 0.074722 | 0.095 | 0.1393 | 1 |
| Q6UXH1 | CRELD2 | GGsQRPCsGNGHCSG | S149 | 0.062 | 94.77 | 353 | 0.049285 | 0.062 | 0.0493 | 1 |
| P35222 | CTNNB1 | AVMVHQLskKEAsRH | S179 | 0 | 96.95 | 781 | 0 | 0 | 0 | 1 |
| P35222 | CTNNB1 | QLskKEAsRHAIMRs | S184 | 0.011 | 94.61 | 781 | 0.008744 | 0.011 | 0.0087 | 1 |
| P35222 | CTNNB1 | sRHAIMRsPQMVsAI | S191 | 0.17 | 96.36 | 781 | 0.135135 | 0.17 | 0.1351 | 1 |

| | | | | | | | | | | |
|---|---|---|---|---|---|---|---|---|---|---|
| P35222 | CTNNB1 | AGTLHNLsHHREGLL | S222 | 0.031 | 98.5 | 781 | 0.024642 | 0.031 | 0.0246 | 1 |
| P35222 | CTNNB1 | LAYGNQEsKLIILAS | S311 | 0.011 | 98.81 | 781 | 0.008744 | 0.011 | 0.0087 | 1 |
| P35222 | CTNNB1 | LVQNCLWtLRNLSDA | T384 | 0 | 98.89 | 781 | 0 | 0 | 0 | 1 |
| P35222 | CTNNB1 | AGDREDItEPAICAL | T461 | 0.004 | 98.74 | 781 | 0.002732 | 0.004 | 0.0027 | 1 |
| P35222 | CTNNB1 | ICALRHLtsRHQEAE | T472 | 0.001 | 98.85 | 781 | 0.000683 | 0.001 | 0.0007 | 1 |
| P35222 | CTNNB1 | HWPLIkAtVGLIRNL | T510 | 0 | 98.87 | 781 | 0 | 0 | 0 | 1 |
| Q6ZS81 | WDFY4 | CTLMALQsILSIAGG | S489 | 0.025 | 95.72 | 3184 | 0.019873 | 0.025 | 0.0199 | 1 |
| Q96SD1 | DCLRE1C | SSYRACFsFHSSySE | S317 | 0.026 | 94.54 | 692 | 0.020668 | 0.026 | 0.0207 | 1 |
| Q96SD1 | DCLRE1C | CFsFHSSySEIKDFL | Y322 | 0.265 | 96.66 | 692 | 0.116252 | 0.265 | 0.1163 | 1 |
| P05129 | PRKCG | PNGLSDPyVkLKLIP | Y195 | 0.104 | 98.3 | 697 | 0.045623 | 0.104 | 0.0456 | 1 |
| Q15746 | MYLK | EVyEDAGsHyLCLLK | S469 | 0.033 | 82.07 | 1914 | 0.026232 | 0.033 | 0.0262 | 1 |
| Q15746 | MYLK | yEDAGsHyLCLLKAR | Y471 | 0.255 | 89.3 | 1914 | 0.111865 | 0.255 | 0.1119 | 0.9091 |
| Q8N4A0 | GALNT4 | AWSTLLRtIHsVLET | T154 | 0.004 | 98.85 | 578 | 0.002732 | 0.004 | 0.0027 | 1 |
| Q8N4A0 | GALNT4 | TLLRtIHsVLETSPA | S157 | 0.077 | 98.74 | 578 | 0.061208 | 0.077 | 0.0612 | 1 |
| Q8N4A0 | GALNT4 | IILVDDLsDRVyLKT | S177 | 0.118 | 96.65 | 578 | 0.0938 | 0.118 | 0.0938 | 1 |
| Q8N4A0 | GALNT4 | SRIDPIRsPtMAGGL | S306 | 0.009 | 98.64 | 578 | 0.007154 | 0.009 | 0.0072 | 1 |
| Q8N4A0 | GALNT4 | IDPIRsPtMAGGLFA | T308 | 0.022 | 98.67 | 578 | 0.015026 | 0.022 | 0.015 | 1 |
| Q8IVU3 | HERC6 | KRDYVSKyIDYIFNV | Y859 | 0.172 | 95.3 | 1022 | 0.075454 | 0.172 | 0.0755 | 0.9167 |
| Q8N5N7 | MRPL50 | PPLRSRAyTPPEDLQ | Y60 | 0.112 | 97.25 | 158 | 0.049133 | 0.112 | 0.0491 | 1 |
| P02786 | TFRC | VkDsAQNsVIIVDkN | S199 | 0.157 | 97.06 | 760 | 0.124802 | 0.157 | 0.1248 | 1 |
| P02786 | TFRC | LVENPGGyVAYSkAA | Y219 | 0.204 | 96.75 | 760 | 0.089492 | 0.204 | 0.0895 | 0.8333 |
| P02786 | TFRC | NAIGVLIyMDQtKFP | Y282 | 0.031 | 97.19 | 760 | 0.013599 | 0.031 | 0.0136 | 1 |
| P02786 | TFRC | NIPVQTIsRAAAEkL | S338 | 0.017 | 97.22 | 760 | 0.013514 | 0.017 | 0.0135 | 1 |
| P02786 | TFRC | GFVEPDHyVVVGAQR | Y402 | 0.253 | 97.85 | 760 | 0.110987 | 0.253 | 0.111 | 0.8182 |
| P02786 | TFRC | WGPGAAksGVGTALL | S419 | 0.005 | 97.87 | 760 | 0.003975 | 0.005 | 0.004 | 1 |
| P02786 | TFRC | LHLKAFTyINLDkAV | Y481 | 0 | 98.57 | 760 | 0 | 0 | 0 | 0.8947 |
| P02786 | TFRC | DVELNLDyERYNsQL | Y611 | 0.161 | 94.42 | 760 | 0.070628 | 0.161 | 0.0706 | 1 |
| P02786 | TFRC | RVEyHFLsPYVsPkE | S687 | 0 | 96.08 | 760 | 0 | 0 | 0 | 1 |
| Q5VYK3 | ECPAS | PsEGsGFtkLHLVES | T836 | 0.058 | 93.43 | 1845 | 0.039615 | 0.083 | 0.053 | 1 |
| Q9NYR9 | NKIRAS2 | KVKLWEVsVADRRsL | S150 | 0.063 | 97.48 | 191 | 0.05008 | 0.063 | 0.0501 | 0.8333 |
| Q02817 | MUC2 | STWGNFHyKTFDGDV | Y45 | 0.024 | 94.95 | 5179 | 0.010528 | 0.024 | 0.0105 | 1 |

| | | | | | | | | | | |
|---|---|---|---|---|---|---|---|---|---|---|
| Q8TF66 | LRRC15 | LANLQNIsLQNNRLR | S371 | 0.109 | 98.52 | 581 | 0.086646 | 0.109 | 0.0866 | 0.8333 |
| Q92973 | TNPO1 | HHLNKPEyIQMLMPP | Y569 | 0.196 | 98.18 | 898 | 0.085982 | 0.196 | 0.086 | 1 |
| Q92973 | TNPO1 | DENWRRFsDQFPLPL | S881 | 0.119 | 87.83 | 898 | 0.094595 | 0.119 | 0.0946 | 1 |
| Q96MG7 | NSMCE3 | EFQWGPRtNLETSkM | T254 | 0.008 | 91.67 | 304 | 0.005464 | 0.008 | 0.0055 | 0.8889 |
| P63151 | PPP2R2A | GSDsVVMtGSYNNFF | T359 | 0.004 | 98.82 | 447 | 0.002732 | 0.004 | 0.0027 | 1 |
| P63151 | PPP2R2A | RNTKRDItLEASREN | T378 | 0.163 | 96.79 | 447 | 0.111331 | 0.163 | 0.1113 | 1 |
| P63151 | PPP2R2A | VATTNNLyIFQDKVN | Y440 | 0.057 | 96.15 | 447 | 0.025005 | 0.057 | 0.025 | 1 |
| Q9HB07 | MYG1 | HQRSFTEtMSSLsPG | T115 | 0.172 | 98.44 | 376 | 0.117478 | 0.172 | 0.1175 | 1 |
| Q9HB07 | MYG1 | PWQTKLSsAGLIYLH | S131 | 0 | 98.78 | 376 | 0 | 0 | 0 | 1 |
| Q8TE68 | EPS8L1 | QKIKYAFsLLARLRG | S328 | 0 | 98.14 | 723 | 0 | 0 | 0 | 1 |
| Q13609 | DNASE1L3 | KEQYAFLyKEKLVSV | Y106 | 0.198 | 98.38 | 305 | 0.08686 | 0.198 | 0.0869 | 0.8 |
| Q86XR8 | CEP57 | LAKLIQEsPtVELkD | S388 | 0.107 | 91.54 | 500 | 0.085056 | 0.107 | 0.0851 | 1 |
| P22413 | ENPP1 | RPVYPTKtFPNHYSI | T256 | 0.171 | 98.59 | 925 | 0.116795 | 0.171 | 0.1168 | 1 |
| P22413 | ENPP1 | GIIDNKMyDPKMNAS | Y280 | 0.067 | 98.62 | 925 | 0.029392 | 0.067 | 0.0294 | 1 |
| Q8NGY3 | OR6K3 | AICNPLRyQMIMtPR | Y148 | 0.282 | 94.48 | 331 | 0.123709 | 0.282 | 0.1237 | 0.8182 |
| P78559 | MAP1A | DWVKNLIsPELGVVF | S129 | 0.153 | 91.61 | 2803 | 0.121622 | 0.153 | 0.1216 | 1 |
| Q8IWR0 | ZC3H7A | RRAMRVMsIERKkWM | S750 | 0.024 | 77.92 | 971 | 0.019078 | 0.164 | 0.1304 | 1 |
| Q9UQQ1 | NAALADL1 | GTIALTRyGGVGRGA | Y193 | 0.096 | 95.91 | 740 | 0.042114 | 0.096 | 0.0421 | 1 |
| Q9UQQ1 | NAALADL1 | QGALGCHyRLGPGFR | Y315 | 0 | 98.29 | 740 | 0 | 0 | 0 | 1 |
| Q9UQQ1 | NAALADL1 | FPEErYYsHVLWAPR | S679 | 0.107 | 95.6 | 740 | 0.085056 | 0.107 | 0.0851 | 1 |
| Q96EX3 | DYNC2I2 | GAtAVAFssFDPRLF | S343 | 0.165 | 96.66 | 536 | 0.131161 | 0.165 | 0.1312 | 0.875 |
| Q008S8 | ECT2L | PSRRFEEyLNLLyAV | Y716 | 0.043 | 91.31 | 904 | 0.018863 | 0.043 | 0.0189 | 0.9167 |
| Q008S8 | ECT2L | LyAVRLHtPAEHVDR | T727 | 0.023 | 81.95 | 904 | 0.015709 | 0.023 | 0.0157 | 1 |
| P42229 | STAT5A | LkIkLGHyAtQLQKT | Y90 | 0.281 | 95.83 | 794 | 0.123271 | 0.281 | 0.1233 | 1 |
| P54753 | EPHB3 | EVFVAIKtLKVGYTE | T666 | 0.179 | 87.35 | 998 | 0.122259 | 0.179 | 0.1223 | 1 |
| Q9Y4E1 | WASHC2C | sVEEIRRssQsWSLA | S31 | 0.126 | 86.36 | 1320 | 0.100159 | 0.126 | 0.1002 | 1 |
| A4D1P6 | WDR91 | DGVIKVWsFNPIMQT | S436 | 0.09 | 93.06 | 747 | 0.071542 | 0.09 | 0.0715 | 1 |
| Q86TI2 | DPP9 | DSYRyPRtGSKNPkI | T308 | 0.032 | 97.92 | 863 | 0.021856 | 0.032 | 0.0219 | 1 |
| Q86TI2 | DPP9 | YIARAGWtRDGKyAW | T358 | 0.054 | 97.1 | 863 | 0.036883 | 0.054 | 0.0369 | 1 |
| Q86TI2 | DPP9 | GWtRDGKyAWAMFLD | Y363 | 0.227 | 96.76 | 863 | 0.099582 | 0.227 | 0.0996 | 1 |
| Q12913 | PTPRJ | VyAIIMLtKCVEQGR | T1139 | 0.014 | 93.47 | 1337 | 0.009562 | 0.014 | 0.0096 | 0.875 |

| | | | | | | | | | | |
|---|---|---|---|---|---|---|---|---|---|---|
| P05187 | ALPP | HVPDsGAtATAYLCG | T117 | 0.001 | 98.91 | 535 | 0.000683 | 0.001 | 0.0007 | 0.9167 |
| P05187 | ALPP | TTRVQHAsPAGTyAH | S177 | 0 | 98.66 | 535 | 0 | 0 | 0 | 0.9091 |
| P05187 | ALPP | HAsPAGTyAHTVNRN | Y182 | 0.04 | 98.91 | 535 | 0.017547 | 0.04 | 0.0175 | 1 |
| P05187 | ALPP | DPSLMEMtEAALRLL | T314 | 0.001 | 98.9 | 535 | 0.000683 | 0.001 | 0.0007 | 1 |
| P05187 | ALPP | QLtsEEDtLSLVtAD | T372 | 0.022 | 98.88 | 535 | 0.015026 | 0.022 | 0.015 | 1 |
| P05187 | ALPP | EDtLSLVtADHSHVF | T377 | 0.001 | 98.91 | 535 | 0.000683 | 0.001 | 0.0007 | 0.8333 |
| P05187 | ALPP | ARDRkAytVLLyGNG | T411 | 0.062 | 98.86 | 535 | 0.042347 | 0.062 | 0.0423 | 1 |
| P05187 | ALPP | PDVtEsEsGsPEyRQ | S436 | 0.006 | 98.16 | 535 | 0.004769 | 0.006 | 0.0048 | 1 |
| P05187 | ALPP | sPEyRQQsAVPLDEE | S445 | 0.166 | 98.83 | 535 | 0.131956 | 0.166 | 0.132 | 1 |
| Q8IU57 | IFNLR1 | TFsVPKysKFSKPTC | S210 | 0.111 | 97.81 | 520 | 0.088236 | 0.111 | 0.0882 | 1 |
| P36507 | MAP2K2 | kLCDFGVsGQLIDsM | S216 | 0.035 | 83.35 | 400 | 0.027822 | 0.035 | 0.0278 | 1 |
| Q05BV3 | EML5 | WKRGKMLsMAPGHtD | S144 | 0.025 | 92.18 | 1969 | 0.019873 | 0.032 | 0.0318 | 1 |
| P32455 | GBP1 | IEVFIRssFkDVDHL | S372 | 0.08 | 94.87 | 592 | 0.063593 | 0.08 | 0.0636 | 1 |
| P32455 | GBP1 | EEVKAGIySKPGGYR | Y427 | 0.198 | 93.99 | 592 | 0.08686 | 0.198 | 0.0869 | 1 |
| P32455 | GBP1 | MMEQkERsYQEHLKQ | S523 | 0.131 | 94.01 | 592 | 0.104134 | 0.131 | 0.1041 | 1 |
| Q8TE76 | MORC4 | NKSCLTFtDDGCGMt | T87 | 0.075 | 98.68 | 937 | 0.051226 | 0.075 | 0.0512 | 1 |
| Q9UJX4 | ANAPC5 | VAkCQVAsAASYDQP | S674 | 0.077 | 91.96 | 755 | 0.061208 | 0.077 | 0.0612 | 1 |
| P61086 | UBE2K | EIAGPPDtPYEGGRY | T49 | 0.155 | 98.31 | 200 | 0.105867 | 0.155 | 0.1059 | 0.8 |
| P48544 | KCNJ5 | QPKKRAEtLMFsNNA | T199 | 0.216 | 91.82 | 419 | 0.147531 | 0.216 | 0.1475 | 1 |
| P48544 | KCNJ5 | RAEtLMFsNNAVISM | S203 | 0.015 | 97.53 | 419 | 0.011924 | 0.015 | 0.0119 | 1 |
| P78536 | ADAM17 | KkNIyLNsGLTSTkN | S382 | 0.019 | 95.89 | 824 | 0.015103 | 0.019 | 0.0151 | 1 |
| O95376 | ARIH2 | RYTLQYTyPYAyyME | Y428 | 0.001 | 95.99 | 493 | 0.000439 | 0.001 | 0.0004 | 0.9286 |
| Q9Y3R0 | GRIP1 | MAINGIPtEDSTFEE | T532 | 0.033 | 74.37 | 1128 | 0.022539 | 0.094 | 0.0642 | 1 |
| O43679 | LDB2 | LTNFTLNyLRLCVIL | Y203 | 0.301 | 94.6 | 373 | 0.132044 | 0.301 | 0.132 | 0.9231 |
| Q8N2R8 | FAM43A | TKMKLtVsAQGIRMV | S114 | 0.148 | 89.13 | 423 | 0.117647 | 0.148 | 0.1176 | 1 |
| Q5TIE3 | VWA5B1 | FIFLIDRsssMsGIs | S369 | 0.019 | 93.54 | 1220 | 0.015103 | 0.019 | 0.0151 | 1 |
| Q5TIE3 | VWA5B1 | EQGSNRRyQVSALHT | Y881 | 0.077 | 91.58 | 1220 | 0.033779 | 0.35 | 0.1468 | 1 |
| P48637 | GSS | IAKAWELyGSPNALV | Y199 | 0.311 | 98.45 | 474 | 0.136431 | 0.311 | 0.1364 | 1 |
| O00442 | RTCA | NGVSRIKtGPVTLHT | T314 | 0.004 | 97.46 | 366 | 0.002732 | 0.004 | 0.0027 | 1 |
| P48200 | IREB2 | AGLRVKPyIRTSLSP | Y541 | 0.085 | 92.91 | 963 | 0.037288 | 0.085 | 0.0373 | 1 |
| P48200 | IREB2 | TPREFNSyGARRGND | Y784 | 0.059 | 89.32 | 963 | 0.025882 | 0.059 | 0.0259 | 1 |

| | | | | | | | | | | |
|---|---|---|---|---|---|---|---|---|---|---|
| Q15389 | ANGPT1 | DVNGGGWtVIQHRED | T325 | 0.01 | 98.61 | 498 | 0.00683 | 0.01 | 0.0068 | 1 |
| Q15389 | ANGPT1 | IGNEKQNyRLyLkGH | Y401 | 0.004 | 98.17 | 498 | 0.001755 | 0.004 | 0.0018 | 1 |
| Q9Y2C2 | UST | NRVGKCGsRtVVLLL | S115 | 0.129 | 90.46 | 406 | 0.102544 | 0.129 | 0.1025 | 0.875 |
| Q5TAX3 | TUT4 | RVQyLGytMKVFAKR | T1101 | 0.001 | 98.11 | 1644 | 0.000683 | 0.001 | 0.0007 | 1 |
| Q8N1B4 | VPS52 | SKIyLSYyRSyLGRL | Y300 | 0.02 | 96.58 | 723 | 0.008774 | 0.02 | 0.0088 | 0.9333 |
| Q8N1B4 | VPS52 | yLSYyRSyLGRLMKV | Y303 | 0.19 | 96.62 | 723 | 0.08335 | 0.19 | 0.0833 | 1 |
| Q14CN2 | CLCA4 | KVQTEKAsIMFMQSI | S236 | 0.006 | 97.24 | 919 | 0.004769 | 0.006 | 0.0048 | 1 |
| P22736 | NR4A1 | IPGFAELsPADQDLL | S431 | 0.187 | 96.22 | 598 | 0.148649 | 0.187 | 0.1486 | 1 |
| Q5JRX3 | PITRM1 | KQFHATHyHPSNARF | Y258 | 0.05 | 98.75 | 1037 | 0.021934 | 0.05 | 0.0219 | 1 |
| Q5JRX3 | PITRM1 | NGYTREAyFSVGLQG | Y388 | 0.12 | 98.85 | 1037 | 0.052642 | 0.12 | 0.0526 | 1 |
| Q5JRX3 | PITRM1 | DsGHLyAsIRAGRTL | S723 | 0.153 | 98.32 | 1037 | 0.121622 | 0.153 | 0.1216 | 1 |
| Q5JRX3 | PITRM1 | IRAGRTLtPAGDLQE | T731 | 0.006 | 98.7 | 1037 | 0.004098 | 0.006 | 0.0041 | 1 |
| P04628 | WNT1 | AFIFAITsAGVtHSV | S132 | 0 | 98.69 | 370 | 0 | 0 | 0 | 1 |
| P04628 | WNT1 | AITsAGVtHSVARSC | T136 | 0 | 98.71 | 370 | 0 | 0 | 0 | 1 |
| Q7RTX1 | TAS1R1 | NDKYQVEtMVLLLQK | T198 | 0.016 | 96.04 | 841 | 0.010928 | 0.016 | 0.0109 | 1 |
| Q7RTX1 | TAS1R1 | WALSRHItGVPGIQR | T310 | 0.176 | 94.89 | 841 | 0.12021 | 0.176 | 0.1202 | 1 |
| Q7Z4L5 | TTC21B | DREAILEsDARVKEQ | S95 | 0.126 | 84.77 | 1316 | 0.100159 | 0.126 | 0.1002 | 1 |
| P43627 | KIR2DL2 | SVTLSCssRSSYDMY | S151 | 0.089 | 97.48 | 348 | 0.070747 | 0.089 | 0.0707 | 1 |
| Q5TBC7 | BCL2L15 | AKNVIAEtIKGQTGA | T79 | 0.039 | 91.33 | 163 | 0.026638 | 0.039 | 0.0266 | 1 |
| A6NJI9 | LRRC72 | DVFELFLsKKELTEV | S53 | 0.088 | 90.67 | 287 | 0.069952 | 0.088 | 0.07 | 1 |
| Q99661 | KIF2C | EGAEINksLLALkEC | S519 | 0.102 | 90.92 | 725 | 0.081081 | 0.102 | 0.0811 | 0.8571 |
| Q99661 | KIF2C | LGQNkAHtPFRESkL | T537 | 0.216 | 77.33 | 725 | 0.147531 | 0.216 | 0.1475 | 1 |
| Q9UP79 | ADAMTS8 | MSVAARIyKHPsIKN | Y254 | 0.014 | 93.42 | 889 | 0.006142 | 0.014 | 0.0061 | 1 |
| Q9UP79 | ADAMTS8 | ARIyKHPsIKNSINL | S258 | 0.135 | 94.57 | 889 | 0.107313 | 0.135 | 0.1073 | 1 |
| Q9UP79 | ADAMTS8 | VTIPAGAtNIDVKQR | T714 | 0.016 | 93.88 | 889 | 0.010928 | 0.016 | 0.0109 | 1 |
| Q92466 | DDB2 | PSTVAVGsKGGDIML | S131 | 0 | 96.13 | 427 | 0 | 0 | 0 | 1 |
| Q92466 | DDB2 | PVNAACFsPDGARLL | S297 | 0.056 | 94.56 | 427 | 0.044515 | 0.056 | 0.0445 | 0.8 |
| Q92466 | DDB2 | HRHFQHLtPIkAAWH | T338 | 0.08 | 94.9 | 427 | 0.054641 | 0.08 | 0.0546 | 1 |
| O95239 | KIF4A | DKNSSFRskLHLVDL | S234 | 0.025 | 92.21 | 1232 | 0.019873 | 0.025 | 0.0199 | 0.9 |
| Q2NL67 | PARP6 | IVLRLRFsLsQyLDG | S101 | 0.115 | 92.15 | 630 | 0.091415 | 0.115 | 0.0914 | 1 |
| P13762 | HLA-DRB4 | RVWNLIRyIyNQEEy | Y59 | 0.302 | 98.2 | 266 | 0.132483 | 0.302 | 0.1325 | 1 |

| | | | | | | | | | | |
|---|---|---|---|---|---|---|---|---|---|---|
| P13762 | HLA-DRB4 | yIyNQEEyARYNSDL | Y66 | 0.167 | 98.26 | 266 | 0.07326 | 0.167 | 0.0733 | 1 |
| Q8NFF2 | SLC24A4 | sEDVAGAtFMAAGSS | T143 | 0.059 | 77.67 | 622 | 0.040298 | 0.059 | 0.0403 | 1 |
| Q86VS3 | IQCH | RIRTSRRtIIHIPSL | T450 | 0.057 | 97.68 | 1027 | 0.038932 | 0.057 | 0.0389 | 1 |
| Q86VS3 | IQCH | LATHLMysPKAIKRI | S557 | 0 | 97.51 | 1027 | 0 | 0 | 0 | 1 |
| Q8WZA2 | RAPGEF4 | IAAHCKEyKNLNSFF | Y857 | 0.27 | 96.29 | 1011 | 0.118445 | 0.336 | 0.1474 | 1 |
| Q9Y3I0 | RTCB | LACARIAsPEGQDyL | S300 | 0.129 | 97.76 | 505 | 0.102544 | 0.129 | 0.1025 | 1 |
| Q9Y3I0 | RTCB | GMAAAGNyAWVNRSS | Y316 | 0.006 | 98.61 | 505 | 0.002632 | 0.006 | 0.0026 | 1 |
| Q9Y3I0 | RTCB | RALSRAKsRRNLDFQ | S439 | 0 | 85.83 | 505 | 0 | 0 | 0 | 1 |
| Q9NRZ7 | AGPAT3 | LPRTkGFttAVKCLR | T212 | 0.133 | 97.91 | 376 | 0.090841 | 0.133 | 0.0908 | 1 |
| P38405 | GNAL | LLLGAGEsGkSTIVk | S53 | 0.033 | 97.5 | 381 | 0.026232 | 0.033 | 0.0262 | 1 |
| P38405 | GNAL | NTNRLREsLDLFESI | S256 | 0.056 | 98.66 | 381 | 0.044515 | 0.079 | 0.0628 | 1 |
| Q9Y5A7 | NUB1 | kDLAkQYsDRLECCE | S46 | 0.08 | 93.64 | 615 | 0.063593 | 0.08 | 0.0636 | 1 |
| Q9Y5A7 | NUB1 | DRKNLLEtRLHITGR | T103 | 0.079 | 94.19 | 615 | 0.053958 | 0.06 | 0.0649 | 1 |
| P43626 | KIR2DL1 | MTQDLAGtYRCYGSV | T97 | 0.135 | 97.99 | 348 | 0.092207 | 0.135 | 0.0922 | 1 |
| O00506 | STK25 | kGsFGEVykGIDNHT | Y35 | 0.327 | 97.25 | 426 | 0.14345 | 0.327 | 0.1434 | 1 |
| P54819 | AK2 | HPksGRsyHEEFNPP | Y152 | 0.202 | 91.98 | 239 | 0.088614 | 0.202 | 0.0886 | 1 |
| Q4VNC1 | ATP13A4 | ENRLKEEtKPVLEEL | T699 | 0.015 | 93.75 | 1196 | 0.010245 | 0.015 | 0.0102 | 1 |
| Q4VNC1 | ATP13A4 | LIsARIRtVMItGDN | T713 | 0.099 | 94.93 | 1196 | 0.067618 | 0.099 | 0.0676 | 1 |
| Q4VNC1 | ATP13A4 | RIRtVMItGDNLQTA | T717 | 0.025 | 89.3 | 1196 | 0.017075 | 0.025 | 0.0171 | 1 |
| Q4VNC1 | ATP13A4 | HFALTGKsFHVIsQH | S799 | 0.045 | 89.51 | 1196 | 0.035771 | 0.045 | 0.0358 | 1 |
| A8TX70 | COL6A5 | VVFLIDNsRNIAKDE | S1971 | 0 | 75.5 | 2615 | 0 | 0.137 | 0.1089 | 1 |
| O95298 | NDUFC2 | YLVKREDyLYAVRDR | Y80 | 0.146 | 93.16 | 119 | 0.064048 | 0.341 | 0.1496 | 1 |
| Q09161 | NCBP1 | VWEILHstIRKMNKH | T644 | 0.014 | 98.57 | 790 | 0.009562 | 0.055 | 0.0376 | 0.9 |
| Q9NQI0 | DDX4 | KAGYTKLtPVQKYSI | T312 | 0.107 | 90.66 | 724 | 0.073082 | 0.107 | 0.0731 | 1 |
| Q9NQI0 | DDX4 | CVRAVVIyGGTQLGH | Y400 | 0.122 | 95.09 | 724 | 0.05352 | 0.279 | 0.1224 | 0.8182 |
| Q9NQI0 | DDX4 | GGTQLGHsIRQIVQG | S408 | 0.041 | 94.32 | 724 | 0.032592 | 0.041 | 0.0326 | 1 |
| Q9NQI0 | DDX4 | IGLKQIKyLVLDEAD | Y442 | 0.157 | 97.6 | 724 | 0.068874 | 0.157 | 0.0689 | 1 |
| P43686 | PSMC4 | DQKPDVMyADIGGMD | Y164 | 0.229 | 87.78 | 418 | 0.100459 | 0.229 | 0.1005 | 1 |
| Q8WXS8 | ADAMTS14 | GCDKEVGsMKADDKC | S696 | 0.188 | 82.39 | 1223 | 0.149444 | 0.188 | 0.1494 | 0.8333 |
| P36871 | PGM1 | GGIILtAsHNPGGPN | S117 | 0.079 | 95.6 | 562 | 0.062798 | 0.079 | 0.0628 | 0.8571 |
| P36871 | PGM1 | MHGVVGPyVKKILCE | Y232 | 0.019 | 98.94 | 562 | 0.008335 | 0.019 | 0.0083 | 1 |

| | | | | | | | | | | |
|---|---|---|---|---|---|---|---|---|---|---|
| P36871 | PGM1 | ELGAPANsAVNCVPL | S247 | 0.055 | 98.79 | 562 | 0.04372 | 0.106 | 0.0843 | 1 |
| P36871 | PGM1 | tkIALyEtPtGWKFF | T355 | 0 | 98.3 | 562 | 0 | 0 | 0 | 0.9091 |
| P36871 | PGM1 | NFFTRYDyEEVEAEG | Y430 | 0.052 | 95.03 | 562 | 0.022812 | 0.052 | 0.0228 | 1 |
| Q13620 | CUL4B | KDVFEAFykKDLAKR | Y607 | 0.022 | 94.68 | 913 | 0.009651 | 0.322 | 0.1413 | 1 |
| Q96JC1 | VPS39 | SIKELFPtGkQLEPL | T202 | 0.01 | 86.2 | 886 | 0.00683 | 0.01 | 0.0068 | 1 |
| Q6ZUX3 | TOGARAM2 | MLLSIIItVADNLNS | T851 | 0.101 | 92.73 | 1019 | 0.068984 | 0.101 | 0.069 | 0.8571 |
| Q14201 | BTG3 | AAVVFFFtRLVRKHD | T13 | 0.181 | 96.44 | 252 | 0.123625 | 0.181 | 0.1236 | 1 |
| P98073 | TMPRSS15 | FSTTNRMtVLLItND | T613 | 0.105 | 90.47 | 1019 | 0.071716 | 0.105 | 0.0717 | 1 |
| P98073 | TMPRSS15 | RMtVLLItNDVLARG | T618 | 0.131 | 87.09 | 1019 | 0.089475 | 0.131 | 0.0895 | 1 |
| Q8N987 | NECAB1 | KLSNESRyMIyEFWE | Y299 | 0.088 | 95.06 | 351 | 0.038604 | 0.088 | 0.0386 | 1 |
| Q9ULC6 | PADI1 | LNVPFsDsKRVRVFC | S208 | 0.048 | 97.24 | 663 | 0.038156 | 0.048 | 0.0382 | 1 |
| Q9ULC6 | PADI1 | GFRLLLAsPSACLKL | S494 | 0.039 | 98.88 | 663 | 0.031002 | 0.039 | 0.031 | 1 |
| Q9ULC6 | PADI1 | yLGIPKPyGPIINGR | Y602 | 0.287 | 96.68 | 663 | 0.125903 | 0.287 | 0.1259 | 1 |
| P52948 | NUP98 | kLySDrHyDLNQLLE | Y1504 | 0.305 | 82.71 | 1817 | 0.133799 | 0.431 | 0.1453 | 1 |
| Q13007 | IL24 | NRTVEVRtLKSFSTL | T133 | 0.03 | 84.8 | 206 | 0.02049 | 0.03 | 0.0205 | 1 |
| P02511 | CRYAB | RLEKDrFsVNLDVkH | S76 | 0.094 | 94.57 | 175 | 0.074722 | 0.094 | 0.0747 | 1 |
| Q10713 | PMPCA | TSRDTTMyAVSADSK | Y151 | 0.094 | 98.34 | 525 | 0.041236 | 0.094 | 0.0412 | 1 |
| Q10713 | PMPCA | GPGKGMFsRLYLNVL | S361 | 0.089 | 94.9 | 525 | 0.070747 | 0.089 | 0.0707 | 1 |
| Q8NFD2 | ANKK1 | GASLELPtHQGWTPL | T590 | 0.107 | 94.71 | 765 | 0.073082 | 0.107 | 0.0731 | 1 |
| Q8NFD2 | ANKK1 | ARNKVGWtPAHLAAL | T694 | 0.061 | 95.87 | 765 | 0.041664 | 0.061 | 0.0417 | 1 |
| Q96RW7 | HMCN1 | AGTGRAQtFFDVSEP | T512 | 0.009 | 92.74 | 5635 | 0.006147 | 0.009 | 0.0061 | 1 |
| Q9ULK0 | GRID1 | GGRsMLDtIKKGHIT | T368 | 0.187 | 90.7 | 1009 | 0.127724 | 0.187 | 0.1277 | 1 |
| O94832 | MYO1D | DQLVVFHtKDNKDLI | T921 | 0.07 | 94.06 | 1006 | 0.047811 | 0.07 | 0.0478 | 0.8 |
| Q9BQP7 | MGME1 | DAELCsQyWtKWLLR | Y317 | 0.331 | 98.04 | 344 | 0.145205 | 0.331 | 0.1452 | 1 |
| Q12756 | KIF1A | VVRRPYAyMyNSDKD | Y1605 | 0.006 | 89.13 | 1690 | 0.002632 | 0.383 | 0.1107 | 1 |
| Q12756 | KIF1A | RRPYAyMyNSDKDTV | Y1607 | 0.216 | 87.98 | 1690 | 0.094756 | 0.216 | 0.0948 | 1 |
| P05981 | HPN | IDACQGDsGGPFVCE | S353 | 0.174 | 97.28 | 417 | 0.138315 | 0.174 | 0.1383 | 1 |
| Q9P2K2 | TXNDC16 | NIHLFIKtMKAPLLT | T246 | 0.071 | 90.78 | 825 | 0.048494 | 0.071 | 0.0485 | 0.9 |
| Q6P047 | C8orf74 | LGNKLRDyRGHFNTT | Y98 | 0.134 | 94.79 | 294 | 0.058784 | 0.165 | 0.0724 | 1 |
| Q6P047 | C8orf74 | HHTFIRHyKLYQYVL | Y122 | 0.103 | 94.8 | 294 | 0.045185 | 0.103 | 0.0452 | 1 |
| Q9NY33 | DPP3 | LERVILGsEAAQQHP | S130 | 0.009 | 98.38 | 737 | 0.007154 | 0.009 | 0.0072 | 1 |

| | | | | | | | | | | |
|---|---|---|---|---|---|---|---|---|---|---|
| Q9NY33 | DPP3 | YYEVRLAsVLGSEPS | S218 | 0 | 98.75 | 737 | 0 | 0 | 0 | 0.8 |
| Q9NY33 | DPP3 | IEAHkRGsRFWIQDK | S297 | 0 | 98.57 | 737 | 0 | 0 | 0 | 1 |
| Q9NY33 | DPP3 | TEGFkNVsLGNVLAV | S408 | 0.025 | 92.38 | 737 | 0.019873 | 0.025 | 0.0199 | 0.9 |
| Q9NY33 | DPP3 | RRLQVLkstGDVAGG | S630 | 0 | 98.48 | 737 | 0 | 0 | 0 | 1 |
| Q9NY33 | DPP3 | RLQVLkstGDVAGGR | T631 | 0.061 | 98.43 | 737 | 0.041664 | 0.061 | 0.0417 | 1 |
| Q9NY33 | DPP3 | kLIVQPNtRLEGsDV | T677 | 0.181 | 98.46 | 737 | 0.123625 | 0.181 | 0.1236 | 1 |
| Q86X76 | NIT1 | WEQTQkIyNCHVLLN | Y143 | 0.177 | 98.67 | 327 | 0.077647 | 0.177 | 0.0776 | 1 |
| Q86X76 | NIT1 | GPMCEsNsTMPGPSL | S180 | 0.142 | 94.66 | 327 | 0.112878 | 0.142 | 0.1129 | 1 |
| Q86X76 | NIT1 | RHHEKRAsyGHSMVV | S268 | 0.081 | 97.3 | 327 | 0.064388 | 0.081 | 0.0644 | 1 |
| P52848 | NDST1 | DCSINPKsPLLyVTR | S204 | 0.109 | 93.71 | 882 | 0.086646 | 0.109 | 0.0866 | 1 |
| P52848 | NDST1 | yPEMDLDsRAFLKDY | S844 | 0 | 97.84 | 882 | 0 | 0 | 0 | 1 |
| P54652 | HSPA2 | VAMNPtNtIFDAkRL | T67 | 0 | 98.29 | 639 | 0 | 0 | 0 | 1 |
| P54652 | HSPA2 | kPkVQVEykGEtkTF | Y108 | 0.016 | 93.96 | 639 | 0.007019 | 0.271 | 0.1128 | 1 |
| P54652 | HSPA2 | NDSQRQAtKDAGtIt | T159 | 0.04 | 93.74 | 639 | 0.027321 | 0.04 | 0.0273 | 1 |
| Q8IWV8 | UBR2 | LTTFGTAtYKVGLKV | T1312 | 0.001 | 96.12 | 1755 | 0.000683 | 0.166 | 0.1134 | 0.8889 |
| Q9Y620 | RAD54B | FYSVLIIsYEMLLRS | S409 | 0.166 | 91.85 | 910 | 0.131956 | 0.166 | 0.132 | 1 |
| Q9Y620 | RAD54B | LSSYRkIyEEPIILS | Y492 | 0.163 | 88.31 | 910 | 0.071506 | 0.163 | 0.0715 | 1 |
| Q5S007 | LRRK2 | RKtIINEsLNFKIRD | S1508 | 0.007 | 85.61 | 2527 | 0.005564 | 0.007 | 0.0056 | 1 |
| Q9HCN4 | GPN1 | TTFVQRLtGHLHAQG | T40 | 0.043 | 95.42 | 374 | 0.02937 | 0.043 | 0.0294 | 1 |
| Q9Y5H0 | PCDHGA3 | SLNPQSGsLVTAERI | S83 | 0.064 | 90.08 | 932 | 0.050875 | 0.064 | 0.0509 | 1 |
| Q9Y5H0 | PCDHGA3 | DyEDAMFyEIKIEAQ | Y315 | 0.217 | 90.68 | 932 | 0.095195 | 0.37 | 0.1395 | 1 |
| Q16518 | RPE65 | IVItEFGtCAFPDPc | T105 | 0.023 | 95.39 | 533 | 0.015709 | 0.045 | 0.031 | 0.8889 |
| Q149N8 | SHPRH | QHAKCVNyDEKNLKI | Y691 | 0.166 | 79.16 | 1683 | 0.072822 | 0.263 | 0.1154 | 0.9167 |
| P15924 | DSP | IyQLEEEyENLLKAS | Y255 | 0.086 | 94.125 | 2871 | 0.037727 | 0.086 | 0.0377 | 1 |
| P15924 | DSP | EELLYDWsDkNTNIA | S299 | 0.018 | 96.25 | 2871 | 0.014308 | 0.018 | 0.0143 | 1 |
| P15924 | DSP | VQNLVNksKKIVQLk | S442 | 0.011 | 96.76 | 2871 | 0.008744 | 0.011 | 0.0087 | 1 |
| Q9Y6Y0 | IVNS1ABP | RAVLACCsPYLFEIF | S54 | 0.016 | 95.74 | 642 | 0.012719 | 0.016 | 0.0127 | 1 |
| Q9Y6Y0 | IVNS1ABP | DSDPHGIsHVKFDDL | S71 | 0.025 | 83.36 | 642 | 0.019873 | 0.025 | 0.0199 | 1 |
| P31939 | ATIC | LGLNLVAsGGtAkAL | S34 | 0.062 | 94.24 | 592 | 0.049285 | 0.062 | 0.0493 | 1 |
| P31939 | ATIC | NLVAsGGtAkALrDA | T37 | 0.027 | 96.71 | 592 | 0.018441 | 0.027 | 0.0184 | 1 |
| P31939 | ATIC | QSSESKDtSLETRRQ | T166 | 0.113 | 98.16 | 592 | 0.077181 | 0.113 | 0.0772 | 1 |

| | | | | | | | | | | |
|---|---|---|---|---|---|---|---|---|---|---|
| P31939 | ATIC | ALkAFtHtAQYDEAI | T182 | 0.088 | 98.67 | 592 | 0.060105 | 0.088 | 0.0601 | 1 |
| P31939 | ATIC | KQYSkGVsQMPLRyG | S202 | 0.069 | 98.29 | 592 | 0.054849 | 0.069 | 0.0548 | 0.8889 |
| P31939 | ATIC | LGIPAAAsFkHVsPA | S264 | 0 | 98.94 | 592 | 0 | 0 | 0 | 1 |
| P31939 | ATIC | yktLtPIsAAyArAR | S300 | 0 | 98.83 | 592 | 0 | 0 | 0 | 1 |
| P31939 | ATIC | LtPIsAAyArARGAD | Y303 | 0 | 98.93 | 592 | 0 | 0 | 0 | 1 |
| P31939 | ATIC | kIISREVsDGIIAPG | S338 | 0.035 | 98.43 | 592 | 0.027822 | 0.035 | 0.0278 | 1 |
| P31939 | ATIC | sLFsNVVtkNkDLPE | T405 | 0.013 | 98.67 | 592 | 0.008879 | 0.013 | 0.0089 | 1 |
| P31939 | ATIC | LRDLIVAtIAVKYTQ | T422 | 0.04 | 98.94 | 592 | 0.027321 | 0.04 | 0.0273 | 1 |
| P31939 | ATIC | KLTEVSIsSDAFFPF | S537 | 0.026 | 98.92 | 592 | 0.020668 | 0.026 | 0.0207 | 1 |
| Q9BW92 | TARS2 | LPRGTRVyNALVAFI | Y332 | 0.228 | 98.5 | 718 | 0.10002 | 0.228 | 0.1 | 1 |
| Q9BW92 | TARS2 | NsGDGAFyGPKIDVH | Y536 | 0.183 | 92.26 | 718 | 0.080279 | 0.183 | 0.0803 | 1 |
| Q9BW92 | TARS2 | LRAAGLVsDLDADSG | S643 | 0.042 | 97.85 | 718 | 0.033386 | 0.042 | 0.0334 | 1 |
| Q6PD62 | CTR9 | ITQATLLyTMADKII | Y117 | 0.088 | 91.08 | 1173 | 0.038604 | 0.088 | 0.0386 | 0.8571 |
| Q6PD62 | CTR9 | RGDkENAsQCFEkVL | S361 | 0.062 | 91.98 | 1173 | 0.049285 | 0.062 | 0.0493 | 1 |
| Q9NZQ3 | NCKIPSD | AIVRTTPyLQHRHRL | Y673 | 0.005 | 96.4 | 722 | 0.002193 | 0.005 | 0.0022 | 1 |
| P20916 | MAG | QCLCVVKsNPEPsVA | S436 | 0.001 | 93.26 | 626 | 0.000795 | 0.001 | 0.0008 | 1 |
| Q7Z401 | DENND4A | LNNsMWGsAVYLCYK | S190 | 0.17 | 88.35 | 1863 | 0.135135 | 0.17 | 0.1351 | 1 |
| Q00266 | MAT1A | VIPVRIHtIVISVQH | T202 | 0.186 | 98.8 | 395 | 0.127041 | 0.186 | 0.127 | 1 |
| P23743 | DGKA | ILPPSSIyPSVLASG | Y335 | 0.136 | 85.28 | 735 | 0.059661 | 0.136 | 0.0597 | 1 |
| O00458 | IFRD1 | QLGPGIEsEEILKTL | S162 | 0.003 | 95.36 | 451 | 0.002385 | 0.003 | 0.0024 | 1 |
| O15523 | DDX3Y | RVRPCVVyGGADIGQ | Y299 | 0.312 | 94.62 | 660 | 0.13687 | 0.312 | 0.1369 | 0.8333 |
| O15523 | DDX3Y | GCHLLVAtPGRLVDM | T321 | 0.121 | 96.04 | 660 | 0.082645 | 0.121 | 0.0826 | 1 |
| O15523 | DDX3Y | IGLDFCkyLVLDEAD | Y341 | 0.061 | 97.94 | 660 | 0.02676 | 0.061 | 0.0268 | 1 |
| O15523 | DDX3Y | LGAtGsDsLTLVFVE | S440 | 0.111 | 85.01 | 660 | 0.088236 | 0.111 | 0.0882 | 1 |
| P52179 | MYOM1 | sGNAKVNyIFNEKEI | Y1277 | 0.127 | 92.3 | 1685 | 0.055713 | 0.164 | 0.0719 | 1 |
| P52179 | MYOM1 | SKKDAGIyEVILKDD | Y1426 | 0.003 | 90.1 | 1685 | 0.001316 | 0.003 | 0.0013 | 1 |
| Q13535 | ATR | QDTLAVAsFRSKAYT | S1645 | 0 | 81.82 | 2644 | 0 | 0 | 0 | 1 |
| Q14692 | BMS1 | IKCDRKVsLyGYLRG | S281 | 0.003 | 94.16 | 1282 | 0.002385 | 0.003 | 0.0024 | 1 |
| Q14692 | BMS1 | FEGAVIRtVSGIRGQ | T1063 | 0.039 | 96.61 | 1282 | 0.026638 | 0.039 | 0.0266 | 1 |
| Q70CQ2 | USP34 | LAFKyFMsPtLtMRL | S324 | 0.028 | 89.535 | 3546 | 0.022258 | 0.119 | 0.0946 | 1 |
| Q8WXE0 | CASKIN2 | IRNtyNQtALDIVNQ | T256 | 0.216 | 89.16 | 1202 | 0.147531 | 0.216 | 0.1475 | 1 |

| | | | | | | | | | | |
|---|---|---|---|---|---|---|---|---|---|---|
| Q9H300 | PARL | GASGAIMtVLAAVCT | T282 | 0.015 | 98.56 | 379 | 0.010245 | 0.015 | 0.0102 | 1 |
| P0CG39 | POTEJ | AGRELTDyLMkILTE | Y851 | 0.188 | 96.66 | 1038 | 0.082473 | 0.267 | 0.1171 | 1 |
| P0CG39 | POTEJ | MWISkQEyDEsGPsI | Y1025 | 0.109 | 96.9 | 1038 | 0.047817 | 0.109 | 0.0478 | 1 |
| Q53GS7 | GLE1 | LILIKEDyFPRIEAI | Y652 | 0.079 | 95.02 | 698 | 0.034656 | 0.079 | 0.0347 | 0.9333 |
| P02771 | AFP | TkLSQKFtKVNFTEI | T248 | 0 | 95.57 | 609 | 0 | 0 | 0 | 1 |
| P02771 | AFP | ILRVAKGyQELLEKC | Y377 | 0.028 | 95.65 | 609 | 0.012283 | 0.028 | 0.0123 | 1 |
| P02771 | AFP | LQKyIQEsQALAkRs | S408 | 0.058 | 93.95 | 609 | 0.046105 | 0.058 | 0.0461 | 0.8 |
| Q5W0V3 | FHIP2A | SSyHVEGtGyDtyLR | T589 | 0.198 | 66.25 | 765 | 0.135237 | 0.198 | 0.1352 | 0.8889 |
| Q86UR5 | RIMS1 | ItKVKKGsLADVVGH | S642 | 0.041 | 85.45 | 1692 | 0.032592 | -0.012 | 0.0824 | 1 |
| Q16720 | ATP2B3 | VtAVGVNsQTGIIFT | S283 | 0.019 | 90.58 | 1220 | 0.015103 | 0.045 | 0.0358 | 1 |
| Q8N335 | GPD1L | PQTSAEVyRILKQKG | Y309 | 0.191 | 98.42 | 351 | 0.083789 | 0.191 | 0.0838 | 1 |
| P11216 | PYGB | VDRLRRMsVIEEGDC | S430 | 0.087 | 97.71 | 843 | 0.069158 | 0.17 | 0.1351 | 1 |
| P11216 | PYGB | KVkINPSsMFDVHVK | S562 | 0.004 | 98.16 | 843 | 0.00318 | 0.004 | 0.0032 | 1 |
| P11216 | PYGB | GyNAREyyDHLPELk | Y733 | 0.2 | 97.85 | 843 | 0.087737 | 0.243 | 0.1066 | 1 |
| P11216 | PYGB | QISSGFFsPkEPDCF | S752 | 0.108 | 96.02 | 843 | 0.085851 | 0.108 | 0.0859 | 1 |
| P11216 | PYGB | SDRTITEyAREIWGV | Y821 | 0.012 | 98.47 | 843 | 0.005264 | 0.012 | 0.0053 | 0.8667 |
| Q9UBN7 | HDAC6 | SSNFDSIyICPSTFA | Y570 | 0.047 | 96.16 | 1215 | 0.020618 | 0.047 | 0.0206 | 1 |
| P06213 | INSR | DGVFTTssDMWsFGV | S1217 | 0.038 | 90.12 | 1382 | 0.030207 | 0.077 | 0.0612 | 1 |
| P35250 | RFC2 | AQQALRRtMEIyskt | T157 | 0.174 | 90.15 | 354 | 0.118844 | 0.174 | 0.1188 | 1 |
| P35250 | RFC2 | MEIyskttRFALACN | T165 | 0.003 | 93.98 | 354 | 0.002049 | 0.003 | 0.002 | 1 |
| P35250 | RFC2 | FALACNAsDkIIEPI | S174 | 0.184 | 91.35 | 354 | 0.146264 | 0.184 | 0.1463 | 1 |
| Q5SRH9 | TTC39A | AIRKSRRyFSSNPIS | Y445 | 0.023 | 97.95 | 613 | 0.01009 | 0.023 | 0.0101 | 1 |
| Q5SRH9 | TTC39A | MMyIWNGyAVIGKQP | Y467 | 0.06 | 96.12 | 613 | 0.026321 | 0.06 | 0.0263 | 1 |
| Q9Y2Z2 | MTO1 | TAAARCGsRtLLLTH | S60 | 0.032 | 98.21 | 717 | 0.025437 | 0.032 | 0.0254 | 1 |
| Q9Y2Z2 | MTO1 | AARCGsRtLLLTHRV | T62 | 0.002 | 98.75 | 717 | 0.001366 | 0.002 | 0.0014 | 1 |
| Q9Y2Z2 | MTO1 | VVSRTEGyIGVLIDD | Y471 | 0.15 | 97.25 | 717 | 0.065803 | 0.15 | 0.0658 | 1 |
| Q9Y2Z2 | MTO1 | VLIDDLttLGtSEPy | T481 | 0.099 | 89.65 | 717 | 0.067618 | 0.166 | 0.1098 | 0.8 |
| Q9Y2Z2 | MTO1 | tLGtSEPyRMFTSRV | Y488 | 0.248 | 89.89 | 717 | 0.108794 | 0.248 | 0.1088 | 0.8571 |
| Q9Y2Z2 | MTO1 | VREkLHFsRPQTIGA | S660 | 0.082 | 84.19 | 717 | 0.065183 | 0.082 | 0.0652 | 1 |
| Q9HCN3 | PGAP6 | APQRLSFySWYGSAR | Y59 | 0.184 | 90.55 | 771 | 0.080718 | 0.184 | 0.0807 | 1 |
| Q9HCN3 | PGAP6 | RVCsDtPsVMRLRLN | S384 | 0.021 | 94.1 | 771 | 0.016693 | 0.065 | 0.1172 | 1 |

| | | | | | | | | | | |
|---|---|---|---|---|---|---|---|---|---|---|
| P35237 | SERPINB6 | LVLVNAVyFRGNWDE | Y168 | 0.102 | 91.92 | 376 | 0.044746 | 0.177 | 0.068 | 0.8 |
| P35237 | SERPINB6 | KADFSGMsQTDLSLS | S306 | 0.027 | 96.06 | 376 | 0.021463 | 0.027 | 0.0215 | 1 |
| Q96RL7 | VPS13A | HRQKVKQyKELykkK | Y365 | 0 | 92.89 | 3174 | 0 | 0 | 0 | 1 |
| Q96RL7 | VPS13A | KEIMDRAyDsFDIQL | Y706 | 0.186 | 79.12 | 3174 | 0.081596 | 0.3565 | 0.135 | 1 |
| Q14697 | GANAB | RsIRPGLsPyRALLD | S58 | 0.072 | 97.41 | 944 | 0.057234 | 0.107 | 0.0982 | 1 |
| Q14697 | GANAB | IRPGLsPyRALLDSL | Y60 | 0.044 | 98.55 | 944 | 0.019302 | 0.044 | 0.0193 | 1 |
| Q14697 | GANAB | VTEGGEPyRLYNLDV | Y277 | 0.037 | 98.87 | 944 | 0.016231 | 0.037 | 0.0162 | 1 |
| Q14697 | GANAB | GsGEtPQtDVRWMSE | T351 | 0.047 | 96.15 | 944 | 0.032102 | 0.094 | 0.0642 | 1 |
| Q14697 | GANAB | ISDVFRQyASLTGTQ | Y379 | 0.133 | 98.78 | 944 | 0.058345 | 0.191 | 0.0838 | 0.8235 |
| Q14697 | GANAB | ELrNLGLyVkTRDGS | Y489 | 0.075 | 98.88 | 944 | 0.032901 | 0.075 | 0.0329 | 1 |
| Q14697 | GANAB | FSGNTLVsSsADPEG | S874 | 0.021 | 98.72 | 944 | 0.016693 | 0.021 | 0.0167 | 1 |
| Q64LD2 | WDR25 | EGsEFLssTDASTRD | S394 | 0 | 92.32 | 544 | 0 | 0 | 0 | 0.8 |
| P19338 | NCL | RsIsLyytGEkGQNQ | T464 | 0.143 | 86.95 | 710 | 0.097671 | 0.143 | 0.0977 | 1 |
| Q9HCG7 | GBA2 | LGGIGGGtItRGWRG | T161 | 0 | 98.19 | 927 | 0 | 0 | 0 | 1 |
| Q9HCG7 | GBA2 | GIGGGtItRGWRGQF | T163 | 0 | 98.29 | 927 | 0 | 0 | 0 | 0.9 |
| Q9HCG7 | GBA2 | GQCRLEFsLAWDMPR | S409 | 0.006 | 98.71 | 927 | 0.004769 | 0.006 | 0.0048 | 1 |
| Q9HCG7 | GBA2 | EDLTRRRyLMsGVMA | Y576 | 0.013 | 95.58 | 927 | 0.005703 | 0.013 | 0.0057 | 1 |
| O43581 | SYT7 | IGGTSDPyVKVWLMy | Y305 | 0.112 | 98.49 | 403 | 0.049133 | 0.112 | 0.0491 | 0.9167 |
| Q92902 | HPS1 | EGDFYCSyFLWFEND | Y611 | 0.003 | 96.76 | 700 | 0.001316 | 0.003 | 0.0013 | 1 |
| P62979 | RPS27A | DyNIQkEstLHLVLr | S65 | 0.161 | 94.68 | 156 | 0.127981 | 0.161 | 0.128 | 1 |
| Q5JTH9 | RRP12 | LkKTLLDsLRSTSsP | S824 | 0.018 | 84.78 | 1297 | 0.014308 | 0.018 | 0.0143 | 1 |
| Q05655 | PRKCD | WGLNkQGyKCRQCNA | Y187 | 0.136 | 86.76 | 676 | 0.059661 | 0.136 | 0.0597 | 0.9091 |
| P07358 | C8B | DFGTHYItEAVLGGI | T346 | 0.184 | 96.46 | 591 | 0.125675 | 0.184 | 0.1257 | 1 |
| Q9BPX3 | NCAPG | sPEIRVYtkALSSLE | T851 | 0.033 | 95.47 | 1015 | 0.022539 | 0.033 | 0.0225 | 0.8333 |
| Q16875 | PFKFB3 | QGRIGGDsGLSsRGK | S269 | 0.157 | 98.06 | 520 | 0.124802 | 0.157 | 0.1248 | 1 |
| Q16875 | PFKFB3 | KkFASALskFVEEQN | S283 | 0.076 | 98.29 | 520 | 0.060414 | 0.076 | 0.0604 | 1 |
| Q16875 | PFKFB3 | RYPTGESyQDLVQRL | Y363 | 0.063 | 98.5 | 520 | 0.027637 | 0.063 | 0.0276 | 1 |
| P29474 | NOS3 | DPRLPPCtLRQALTF | T854 | 0.119 | 95.65 | 1203 | 0.081279 | 0.119 | 0.0813 | 1 |
| P29323 | EPHB2 | CRGCPSGtFKANQGD | T276 | 0.099 | 95.72 | 1055 | 0.067618 | 0.099 | 0.0676 | 1 |
| P29323 | EPHB2 | GLKAGAIyVFQVRAR | Y504 | 0.167 | 94.04 | 1055 | 0.07326 | 0.167 | 0.0733 | 1 |
| Q7L1Q6 | BZW1 | NLVKEGVsAAFAVkL | S182 | 0 | 97.9 | 419 | 0 | 0 | 0 | 1 |

| | | | | | | | | | | |
|---|---|---|---|---|---|---|---|---|---|---|
| Q7L1Q6 | BZW1 | GLkELSEyVRNQQTI | Y244 | 0.235 | 94.24 | 419 | 0.103091 | 0.235 | 0.1031 | 1 |
| O60542 | PSPN | QLWSLTLsVAELGLG | S74 | 0.042 | 96.4 | 156 | 0.033386 | 0.042 | 0.0334 | 1 |
| Q9UJF2 | RASAL2 | ATkSIEEyLKLVGQQ | Y381 | 0.078 | 96.89 | 1139 | 0.034217 | 0.078 | 0.0342 | 1 |
| Q9UJF2 | RASAL2 | LKLVGQQyLHDALGE | Y389 | 0.075 | 92.22 | 1139 | 0.032901 | 0.075 | 0.0329 | 1 |
| P40926 | MDH2 | LVsRLTLyDIAHtPG | Y56 | 0.126 | 98.9 | 338 | 0.055274 | 0.248 | 0.1078 | 0.9286 |
| P40926 | MDH2 | TLyDIAHtPGVAADL | T61 | 0 | 98.34 | 338 | 0 | 0 | 0 | 1 |
| P40926 | MDH2 | VECsFVKsQEtECty | S280 | 0.052 | 98.6 | 338 | 0.041336 | 0.104 | 0.0827 | 1 |
| Q15034 | HERC3 | SDRGQLFsWGAGSDG | S107 | 0 | 97.19 | 1050 | 0 | 0 | 0 | 1 |
| Q9NQX3 | GPHN | NLPPFPAsVKDGYAV | S371 | 0.025 | 98.29 | 736 | 0.019873 | 0.025 | 0.0199 | 1 |
| Q9NQX3 | GPHN | DRFIIGEsQAGEQPT | S393 | 0.02 | 96.5 | 736 | 0.015898 | 0.035 | 0.0284 | 1 |
| Q9NQX3 | GPHN | SRADVIItsGGVsMG | T570 | 0.005 | 98.55 | 736 | 0.003415 | 0.005 | 0.0034 | 1 |
| Q9NQX3 | GPHN | RADVIItsGGVsMGE | S571 | 0.017 | 97.44 | 736 | 0.013514 | 0.017 | 0.0135 | 1 |
| Q9NQX3 | GPHN | KLDPRPEyHRCILTW | Y673 | 0.147 | 96.45 | 736 | 0.064487 | 0.293 | 0.1283 | 1 |
| P43364 | MAGEA11 | VHLLLRkyRVkGLIT | Y237 | 0.072 | 94.74 | 429 | 0.031585 | 0.103 | 0.0452 | 1 |
| Q9Y6Q1 | CAPN6 | KYLKKGNyVLVPTMF | Y467 | 0.079 | 97.74 | 641 | 0.034656 | 0.079 | 0.0347 | 0.9091 |
| Q9Y6Q1 | CAPN6 | ANETVNPyLVIKCGK | Y540 | 0.22 | 97.42 | 641 | 0.096511 | 0.22 | 0.0965 | 1 |
| Q5VYK3 | ECPAS | LSMMVGAySTLEGAQ | Y470 | 0 | 90.83 | 1845 | 0 | 0 | 0 | 1 |
| Q14721 | KCNB1 | AEKDEDDtKFKSIPA | T359 | 0.143 | 90.72 | 858 | 0.097671 | 0.143 | 0.0977 | 1 |
| Q14721 | KCNB1 | SFWWATItMTTVGYG | T374 | 0.173 | 97.14 | 858 | 0.118161 | 0.173 | 0.1182 | 1 |
| Q96PP8 | GBP5 | GLYRTGKsYLMNKLA | S52 | 0.15 | 91.03 | 586 | 0.119237 | 0.15 | 0.1192 | 1 |
| Q8NF91 | SYNE1 | LKsWIIKyGRRESVE | Y530 | 0.148 | 95.15 | 8797 | 0.064925 | 0.148 | 0.0649 | 1 |
| O75937 | DNAJC8 | kkRFRQLsILVHPDk | S81 | 0.153 | 94.2 | 253 | 0.121622 | 0.153 | 0.1216 | 1 |
| Q9NS28 | RGS18 | QLMEQDSyTRFLKSD | Y189 | 0.207 | 96.52 | 235 | 0.090808 | 0.207 | 0.0908 | 1 |
| Q9NS28 | RGS18 | RFLKSDIyLDLMEGR | Y198 | 0.12 | 91.45 | 235 | 0.052642 | 0.12 | 0.0526 | 1 |
| Q8N6S4 | ANKRD13C | YGDRQMItALLRKLK | T194 | 0.082 | 96.98 | 541 | 0.056007 | 0.082 | 0.056 | 1 |
| Q9ULU4 | ZMYND8 | kFGVFNysPFRTPyT | S363 | 0.089 | 95.97 | 1186 | 0.070747 | 0.089 | 0.0707 | 1 |
| Q9ULU4 | ZMYND8 | ysPFRTPyTPNSQYQ | Y369 | 0.16 | 96.73 | 1186 | 0.07019 | 0.16 | 0.0702 | 1 |
| Q9NXS2 | QPCTL | IPHsPGPtRIQAIEL | T257 | 0.201 | 98.15 | 382 | 0.137286 | 0.201 | 0.1373 | 1 |
| Q2M2H8 | MGAM2 | QLSFRLPsANVYGLG | S215 | 0.126 | 94.98 | 2515 | 0.100159 | 0.126 | 0.1002 | 1 |
| P38117 | ETFB | VKEVIAVsCGPAQCQ | S65 | 0.022 | 98.68 | 255 | 0.017488 | 0.022 | 0.0175 | 1 |
| P38117 | ETFB | LkLPAVVtADLrLNE | T182 | 0.024 | 98.88 | 255 | 0.016392 | 0.024 | 0.0164 | 1 |

| | | | | | | | | | | |
|---|---|---|---|---|---|---|---|---|---|---|
| Q9Y305 | ACOT9 | EPELREkyLTVQNTV | Y104 | 0.015 | 96.27 | 439 | 0.00658 | 0.015 | 0.0066 | 1 |
| Q9Y305 | ACOT9 | YIQVRVHsEVASLQE | S375 | 0.017 | 95.92 | 439 | 0.013514 | 0.017 | 0.0135 | 1 |
| Q99459 | CDC5L | SKKDRIEsLEkRLEI | S691 | 0.093 | 86.59 | 802 | 0.073927 | 0.093 | 0.0739 | 1 |
| Q14517 | FAT1 | PEFtQTAyKAAFDEN | Y468 | 0.211 | 87.4 | 4588 | 0.092563 | 0.211 | 0.0926 | 1 |
| Q14517 | FAT1 | YELMPRVytLRIRAS | Y534 | 0.208 | 86.04 | 4588 | 0.091247 | 0.208 | 0.0912 | 1 |
| O75602 | SPAG6 | ALKRIAAsALSDIAK | S189 | 0.049 | 98.3 | 509 | 0.038951 | 0.049 | 0.039 | 1 |
| Q9UIW2 | PLXNA1 | KSWVERYyADIAKMP | Y1817 | 0.175 | 93.43 | 1896 | 0.07677 | 0.175 | 0.0768 | 1 |
| Q9HB71 | CACYBP | kPISVEGsSkkVkTD | S141 | 0.148 | 98.15 | 228 | 0.117647 | 0.148 | 0.1176 | 1 |
| Q9NQU5 | PAK6 | KsLVGtPyWMAPEVI | Y566 | 0.261 | 97.73 | 681 | 0.114497 | 0.261 | 0.1145 | 1 |
| Q92854 | SEMA4D | SLNLPDKtLQFVKDH | T390 | 0.046 | 98.49 | 862 | 0.031419 | 0.046 | 0.0314 | 1 |
| P11511 | CYP19A1 | MVtVCAEsLKTHLDR | S167 | 0.076 | 97.4 | 503 | 0.060414 | 0.076 | 0.0604 | 0.8889 |
| Q96BX8 | MOB3A | FRkPtALsAPRYMDL | S114 | 0.12 | 98.57 | 217 | 0.09539 | 0.12 | 0.0954 | 1 |
| O94822 | LTN1 | SSQLRALySMyLRKT | Y1499 | 0 | 90.19 | 1766 | 0 | 0.211 | 0.0723 | 1 |
| Q16678 | CYP1B1 | GLINKDLtSRVMIFS | T457 | 0.191 | 96.57 | 543 | 0.130456 | 0.191 | 0.1305 | 1 |
| Q6WRI0 | IGSF10 | QTSDSGIyKCTAKNP | Y2603 | 0.031 | 89.6 | 2623 | 0.013599 | 0.031 | 0.0136 | 1 |
| Q9BUI4 | POLR3C | GDsGGGMyVINLHkA | Y334 | 0.114 | 96.54 | 534 | 0.05001 | 0.114 | 0.05 | 1 |
| Q9HCS4 | TCF7L1 | SREEQAkyyELARKE | Y395 | 0.179 | 97.46 | 588 | 0.078525 | 0.179 | 0.0785 | 0.8333 |
| P27105 | STOM | SVDGVVyyRVQNATL | Y124 | 0.057 | 97.14 | 288 | 0.025005 | 0.057 | 0.025 | 1 |
| Q9BX79 | STRA6 | TLDPGYytYRNFLKI | T565 | 0.049 | 92.7 | 667 | 0.033468 | 0.049 | 0.0335 | 1 |
| P48507 | GCLM | KkIVAIGtsDLDkTQ | T164 | 0.026 | 97.6 | 274 | 0.017758 | 0.026 | 0.0178 | 1 |
| P48507 | GCLM | kIVAIGtsDLDkTQL | S165 | 0.018 | 95.28 | 274 | 0.014308 | 0.018 | 0.0143 | 0.9091 |
| P14616 | INSRR | GLRHFtEyRIDIHAC | Y783 | 0.174 | 88.63 | 1297 | 0.076331 | 0.174 | 0.0763 | 1 |
| Q13371 | PDCL | ALPALLIyKGGELIG | Y239 | 0.305 | 97.49 | 301 | 0.133799 | 0.305 | 0.1338 | 1 |
| P40939 | HADHA | CKTLQEVtQLsQEAQ | T117 | 0.171 | 98.24 | 763 | 0.116795 | 0.171 | 0.1168 | 1 |
| P40939 | HADHA | LQEVtQLsQEAQRIV | S120 | 0.028 | 97.7 | 763 | 0.022258 | 0.028 | 0.0223 | 1 |
| P40939 | HADHA | IATkDRKtVLGtPEV | T167 | 0.006 | 96.25 | 763 | 0.004098 | 0.006 | 0.0041 | 1 |
| P40939 | HADHA | DRKtVLGtPEVLLGA | T171 | 0.013 | 98.05 | 763 | 0.008879 | 0.013 | 0.0089 | 1 |
| P40939 | HADHA | ALPGAGGtQRLPKMV | T185 | 0.016 | 98.25 | 763 | 0.010928 | 0.016 | 0.0109 | 1 |
| P40939 | HADHA | EEkVRKQtkGLyPAP | T294 | 0.216 | 93.72 | 763 | 0.147531 | 0.216 | 0.1475 | 0.8571 |
| P40939 | HADHA | SVDkGLKtILkDAtL | T387 | 0.025 | 98.76 | 763 | 0.017075 | 0.025 | 0.0171 | 1 |
| P40939 | HADHA | EKVIGMHyFsPVDkM | Y499 | 0.05 | 98.06 | 763 | 0.021934 | 0.05 | 0.0219 | 1 |

| | | | | | | | | | | |
|---|---|---|---|---|---|---|---|---|---|---|
| P40939 | HADHA | TEKTSkDtsAsAVAV | T521 | 0 | 98.79 | 763 | 0 | 0 | 0 | 1 |
| P40939 | HADHA | EKTSkDtsAsAVAVG | S522 | 0.038 | 98.63 | 763 | 0.030207 | 0.038 | 0.0302 | 1 |
| P40939 | HADHA | TSkDtsAsAVAVGLk | S524 | 0.057 | 98.73 | 763 | 0.04531 | 0.057 | 0.0453 | 1 |
| P40939 | HADHA | RKSGKGFyIyQEGVk | Y637 | 0.025 | 97.09 | 763 | 0.010967 | 0.025 | 0.011 | 1 |
| P40939 | HADHA | LKkYEAAyGkQFtPC | Y740 | 0.318 | 95.85 | 763 | 0.139502 | 0.318 | 0.1395 | 1 |
| Q969Y0 | NXPE3 | sVSLVHPsEGIRVLQ | S191 | 0.032 | 96.52 | 559 | 0.025437 | 0.032 | 0.0254 | 1 |
| Q969Y0 | NXPE3 | VVRLLDRsPKTVVVI | S474 | 0.169 | 98.45 | 559 | 0.134341 | 0.169 | 0.1343 | 1 |
| Q9H8E8 | KAT14 | GDNFFRFtCsDCsAD | T105 | 0.19 | 89.47 | 782 | 0.129773 | 0.19 | 0.1298 | 1 |
| Q9H8E8 | KAT14 | RFtCsDCsADGKEQy | S110 | 0.137 | 87.15 | 782 | 0.108903 | 0.137 | 0.1089 | 1 |
| O95163 | ELP1 | YCLSILTsHVKkTTP | S842 | 0 | 95.26 | 1332 | 0 | 0.089 | 0.0707 | 1 |
| O15040 | TECPR2 | SIGMLyLyCRHLNQM | Y60 | 0.144 | 93.15 | 1411 | 0.063171 | 0.176 | 0.0772 | 1 |
| Q14690 | PDCD11 | LARYSHVsQHsPskk | S1357 | 0.026 | 78.49 | 1871 | 0.020668 | 0.099 | 0.0787 | 0.8333 |
| P49590 | HARS2 | EKVRttEtQVFVATP | T410 | 0.139 | 97.44 | 506 | 0.094939 | 0.139 | 0.0949 | 1 |
| Q6UWU2 | GLB1L | GSLRGLYttVDFGPA | T234 | 0.047 | 98.63 | 654 | 0.032102 | 0.047 | 0.0321 | 1 |
| Q6UWU2 | GLB1L | SLRGLYttVDFGPAD | T235 | 0.023 | 98.71 | 654 | 0.015709 | 0.023 | 0.0157 | 1 |
| Q6UWU2 | GLB1L | FLTGKLGsKLDILVE | S469 | 0.067 | 97.12 | 654 | 0.053259 | 0.067 | 0.0533 | 1 |
| O75771 | RAD51D | KLLDAGLytGEVTEI | Y98 | 0.324 | 96.56 | 328 | 0.142134 | 0.324 | 0.1421 | 1 |
| O75771 | RAD51D | LLDAGLytGEVTEIV | T99 | 0.131 | 96.12 | 328 | 0.089475 | 0.131 | 0.0895 | 1 |
| Q7Z7H5 | TMED4 | QkEVFLPstPGLGMH | S71 | 0.115 | 80.79 | 227 | 0.091415 | 0.115 | 0.0914 | 1 |
| Q5JSH3 | WDR44 | HCGRLLAsAGQDNVV | S528 | 0 | 98.22 | 913 | 0 | 0 | 0 | 0.8 |
| Q9Y606 | PUS1 | ARTDKGVsAAGQVVS | S150 | 0.023 | 98.61 | 427 | 0.018283 | 0.188 | 0.1494 | 1 |
| Q9UHC6 | CNTNAP2 | KQISAIAtQGRYSSS | T100 | 0.006 | 98.41 | 1331 | 0.004098 | 0.006 | 0.0041 | 1 |
| Q9UHC6 | CNTNAP2 | NYNGVNItDLARRKK | T348 | 0.026 | 92.17 | 1331 | 0.017758 | 0.026 | 0.0178 | 1 |
| P35609 | ACTN2 | HRPDLIDysKLNKDD | Y200 | 0.286 | 91.98 | 894 | 0.125464 | 0.286 | 0.1255 | 1 |
| P35609 | ACTN2 | LEQAEKGyEEWLLNE | Y385 | 0.067 | 95.39 | 894 | 0.029392 | 0.067 | 0.0294 | 1 |
| Q9H1C4 | UNC93B1 | TLVAAAVsYLRMEQK | S510 | 0.144 | 93.79 | 597 | 0.114468 | 0.144 | 0.1145 | 1 |
| Q9UK97 | FBXO9 | LRSLEQLsLVCRGFY | S217 | 0.055 | 94.99 | 447 | 0.04372 | 0.055 | 0.0437 | 1 |
| Q9UK97 | FBXO9 | AWHQVEyyRYIRFFP | Y297 | 0.225 | 97.19 | 447 | 0.098704 | 0.225 | 0.0987 | 1 |
| P13196 | ALAS1 | LLAGALEsVRILKSA | S488 | 0 | 98.87 | 640 | 0 | 0 | 0 | 1 |
| P36957 | DLST | DISVAVAtPRGLVVP | T323 | 0.202 | 86.96 | 453 | 0.137969 | 0.202 | 0.138 | 1 |
| Q9NY93 | DDX56 | LEQFSIPtCVLNGEL | T285 | 0.157 | 88.49 | 547 | 0.107233 | 0.157 | 0.1072 | 0.8889 |

| | | | | | | | | | | |
|---|---|---|---|---|---|---|---|---|---|---|
| P42684 | ABL2 | AkFPIKWtAPESLAy | T452 | 0.084 | 96.24 | 1182 | 0.057373 | 0.084 | 0.0574 | 0.8333 |
| Q8WUT4 | LRRN4 | VHGYQIRySAEGWAG | Y614 | 0.299 | 91.51 | 740 | 0.131167 | 0.299 | 0.1312 | 1 |
| P98187 | CYP4F8 | LTMCLKEsLRLHPPI | S388 | 0.018 | 98.79 | 520 | 0.014308 | 0.018 | 0.0143 | 1 |
| Q6N069 | NAA16 | LGkkEEAyEFVRKGL | Y66 | 0.282 | 93.58 | 864 | 0.123709 | 0.282 | 0.1237 | 1 |
| Q32P51 | HNRNPA1L2 | IGGLsFEtTDEsLrS | T25 | 0.001 | 94.23 | 320 | 0.000683 | 0.001 | 0.0007 | 1 |
| Q32P51 | HNRNPA1L2 | VTFDDHDsVDkIVIQ | S158 | 0.074 | 92.9 | 320 | 0.058824 | 0.147 | 0.117 | 1 |
| P83876 | TXNL4A | KGRGLVVsPkDYsTk | S132 | 0.125 | 85.45 | 142 | 0.099364 | 0.125 | 0.0994 | 1 |
| P08174 | CD55 | GyRQSVTyACNKGFT | Y251 | 0.062 | 98.32 | 381 | 0.027199 | 0.062 | 0.0272 | 0.8462 |
| Q9P2M1 | LRP2BP | QSMLGLyysTKEPKE | Y183 | 0.074 | 98.37 | 347 | 0.032463 | 0.074 | 0.0325 | 0.8 |
| Q68E01 | INTS3 | LDLLSELyQKQPKIG | Y681 | 0.125 | 95.78 | 1043 | 0.054836 | 0.125 | 0.0548 | 0.9286 |
| Q6VY07 | PACS1 | IkIysLSsQPIDHEG | S255 | 0.156 | 88.41 | 963 | 0.124007 | 0.156 | 0.124 | 0.8889 |
| Q96MM6 | HSPA12B | PSLPEKDtVRWVLtV | T200 | 0.103 | 85.43 | 686 | 0.07035 | 0.204 | 0.1393 | 1 |
| Q96MM6 | HSPA12B | DtVRWVLtVPAIWKQ | T206 | 0.016 | 95.92 | 686 | 0.010928 | 0.016 | 0.0109 | 1 |
| P51659 | HSD17B4 | DIVISNAyVDLAPTS | Y602 | 0.136 | 98.66 | 736 | 0.059661 | 0.136 | 0.0597 | 0.8333 |
| Q14568 | HSP90AA2P | LRELISNssDALDKI | S52 | 0.059 | 97.16 | 343 | 0.0469 | 0.059 | 0.0469 | 1 |
| Q14568 | HSP90AA2P | RELISNssDALDKIW | S53 | 0.127 | 95.54 | 343 | 0.100954 | 0.127 | 0.101 | 1 |
| Q14568 | HSP90AA2P | LkEDQTEyLEEQRIK | Y197 | 0.084 | 97.38 | 343 | 0.03685 | 0.084 | 0.0369 | 1 |
| Q9UM54 | MYO6 | YSKDRIYtYVANILI | T88 | 0.031 | 93.74 | 1294 | 0.021173 | 0.031 | 0.0212 | 1 |
| Q9UM54 | MYO6 | NTKFVLRyLTESYGT | Y167 | 0 | 91.75 | 1294 | 0 | 0 | 0 | 1 |
| Q9UM54 | MYO6 | KLRstGAsFIRCIKP | S664 | 0.051 | 93.29 | 1294 | 0.040541 | 0.051 | 0.0405 | 1 |
| Q9UM54 | MYO6 | LGLNENDykFGLTkV | Y749 | 0.108 | 86.95 | 1294 | 0.047378 | 0.108 | 0.0474 | 1 |
| A6NHR9 | SMCHD1 | KPPtPAVsNVRsVAS | S1503 | 0 | 79.01 | 2005 | 0 | 0 | 0 | 1 |
| A6NHR9 | SMCHD1 | QQQMAALtkEKDQLS | T1627 | 0.058 | 75.81 | 2005 | 0.039615 | 0.058 | 0.0396 | 1 |
| Q9Y5F0 | PCDHB13 | LKsAENFyTLLTERP | Y401 | 0.184 | 92.69 | 798 | 0.080718 | 0.184 | 0.0807 | 1 |
| P50990 | CCT8 | GMVFkkEtEGDVtsV | T227 | 0.203 | 86.96 | 548 | 0.138652 | 0.203 | 0.1387 | 1 |
| P50990 | CCT8 | tGANVVVtGGkVADM | T293 | 0 | 93.61 | 548 | 0 | 0 | 0 | 1 |
| P50990 | CCT8 | VADMALHyANkYNIM | Y304 | 0.246 | 92.91 | 548 | 0.107917 | 0.246 | 0.1079 | 1 |
| P50990 | CCT8 | DLRRLCktVGAtALP | T327 | 0.043 | 89.44 | 548 | 0.02937 | 0.043 | 0.0294 | 1 |
| P50990 | CCT8 | LVPGGGAtEIELAkQ | T415 | 0.031 | 93.79 | 548 | 0.021173 | 0.031 | 0.0212 | 1 |
| P50990 | CCT8 | EAGILDTyLGKyWAI | Y501 | 0.146 | 93.32 | 548 | 0.064048 | 0.146 | 0.064 | 1 |
| P50990 | CCT8 | yWAIKLAtNAAVtVL | T512 | 0.025 | 96.26 | 548 | 0.017075 | 0.025 | 0.0171 | 1 |

| | | | | | | | | | | |
|---|---|---|---|---|---|---|---|---|---|---|
| P21980 | TGM2 | RKLVAEVsLQNPLPV | S608 | 0.11 | 97.66 | 687 | 0.087441 | 0.11 | 0.0874 | 1 |
| Q9HCX4 | TRPC7 | YAIAVVLsFSRIAyI | S551 | 0 | 93.45 | 862 | 0 | 0 | 0 | 0.8889 |
| Q86YT5 | SLC13A5 | GFALAKGsEAsGLsV | S424 | 0.088 | 90.25 | 568 | 0.069952 | 0.088 | 0.07 | 0.875 |
| Q86YT5 | SLC13A5 | LAKGsEAsGLsVWMG | S427 | 0.138 | 91.48 | 568 | 0.109698 | 0.138 | 0.1097 | 1 |
| Q9BQI6 | SLF1 | DDLGSYVsLSCDDFS | S654 | 0.129 | 88.8 | 1058 | 0.102544 | 0.129 | 0.1025 | 1 |
| P43146 | DCC | NRERALNtTQPGSLQ | T479 | 0.149 | 84.8 | 1447 | 0.101769 | 0.149 | 0.1018 | 0.9167 |
| P43146 | DCC | SQPIKVAtQPELQVP | T521 | 0.12 | 87.21 | 1447 | 0.081962 | 0.12 | 0.082 | 0.9231 |
| P62913 | RPL11 | AkVLEQLtGQtPVFs | T44 | 0.001 | 93.88 | 178 | 0.000683 | 0.001 | 0.0007 | 1 |
| Q9P1U1 | ACTR3B | PRKWIkQyTGINAIN | Y256 | 0.272 | 97.32 | 418 | 0.119322 | 0.272 | 0.1193 | 1 |
| Q86XA9 | HEATR5A | LsVIsSAsKLFGVVC | S783 | 0.027 | 91.62 | 2040 | 0.021463 | 0.153 | 0.1216 | 1 |
| Q86XA9 | HEATR5A | GHSLALGsLHRYLGG | S916 | 0.004 | 92.24 | 2040 | 0.00318 | 0.004 | 0.0032 | 1 |
| Q6IN85 | PPP4R3A | kDEFYNRyIMkSFLF | Y573 | 0.221 | 97.96 | 833 | 0.09695 | 0.221 | 0.0969 | 0.9231 |
| P28749 | RBL1 | LKLAEILyyKILETV | Y468 | 0.001 | 98.48 | 1068 | 0.000439 | 0.117 | 0.0513 | 1 |
| P28749 | RBL1 | TGSLALFyRKVYHLA | Y792 | 0 | 95.85 | 1068 | 0 | 0 | 0 | 0.9333 |
| Q2M2I8 | AAK1 | EDEIKkYttLSYRAP | T221 | 0.142 | 97.54 | 961 | 0.096988 | 0.142 | 0.097 | 1 |
| Q2M2I8 | AAK1 | DMHCLIRyMLEPDPD | Y292 | 0.221 | 98.7 | 961 | 0.09695 | 0.221 | 0.0969 | 1 |
| P05362 | ICAM1 | FENTSAPyQLQtFVL | Y207 | 0.26 | 96.48 | 532 | 0.114058 | 0.26 | 0.1141 | 1 |
| P05362 | ICAM1 | SAPyQLQtFVLPAtP | T211 | 0 | 94.04 | 532 | 0 | 0 | 0 | 0.8333 |
| P42262 | GRIA2 | StLEEKEyPGAHTtt | Y285 | 0.264 | 92.26 | 883 | 0.115813 | 0.264 | 0.1158 | 1 |
| Q9BZC7 | ABCA2 | GGMkRKLsVAIAFVG | S1132 | 0.071 | 89.77 | 2435 | 0.056439 | 0.071 | 0.0564 | 1 |
| Q9Y5H4 | PCDHGA1 | DFEEYKMysMEVQAQ | Y315 | 0.123 | 91.95 | 931 | 0.053958 | 0.123 | 0.054 | 1 |
| Q99594 | TEAD3 | RLRLLEYsAFMEVQR | S231 | 0.017 | 97.64 | 435 | 0.013514 | 0.017 | 0.0135 | 1 |
| Q9UBL3 | ASH2L | GYDKFSYsWRSKKGT | S459 | 0 | 98.58 | 628 | 0 | 0 | 0 | 1 |
| Q9UBL3 | ASH2L | IKFKsyLyFEEKDFV | Y519 | 0 | 92.66 | 628 | 0 | 0 | 0 | 0.9231 |
| P23469 | PTPRE | NILPNDHsRVILSQL | S174 | 0.096 | 96.8 | 700 | 0.076312 | 0.096 | 0.0763 | 1 |
| P46952 | HAAO | AWERTQGsVALsVtQ | S270 | 0 | 97.6 | 286 | 0 | 0 | 0 | 1 |
| P46952 | HAAO | GsVALsVtQDPACKK | T276 | 0.208 | 97.69 | 286 | 0.142067 | 0.208 | 0.1421 | 1 |
| Q9Y5P8 | PPP2R3B | RHNDHALstkMIDRI | S348 | 0.001 | 97.95 | 575 | 0.000795 | 0.001 | 0.0008 | 0.8889 |
| P35503 | UGT1A3 | PQTVLWRyTGTRPSN | Y338 | 0.15 | 96.29 | 534 | 0.065803 | 0.268 | 0.1062 | 1 |
| Q9UNI6 | DUSP12 | VDTSSAIyKQYRLQK | Y179 | 0.246 | 94.59 | 340 | 0.107917 | 0.246 | 0.1079 | 0.8182 |
| P23193 | TCEA1 | VNAIRKQstDEEVTS | S57 | 0.147 | 83.22 | 301 | 0.116852 | 0.147 | 0.1169 | 1 |

| | | | | | | | | | | |
|---|---|---|---|---|---|---|---|---|---|---|
| Q9UBR4 | LHX3 | LETLKsAyNTSPKPA | Y176 | 0.115 | 96.05 | 397 | 0.050449 | 0.115 | 0.0504 | 1 |
| Q9H7Z7 | PTGES2 | AAAMyLIsKRLKSRH | S290 | 0.062 | 92.02 | 377 | 0.049285 | 0.062 | 0.0493 | 1 |
| P02787 | TF | DNLKGKKsCHTAVGR | S468 | 0 | 98.75 | 698 | 0 | 0 | 0 | 1 |
| P02787 | TF | EPNNKEGyyGytGAF | Y533 | 0.011 | 98.69 | 698 | 0.004826 | 0.011 | 0.0048 | 1 |
| P02787 | TF | DLLFRDDtVCLAKLH | T654 | 0.073 | 97.76 | 698 | 0.04986 | 0.073 | 0.0499 | 1 |
| P02787 | TF | KLHDRNTyEKyLGEE | Y666 | 0.244 | 96.37 | 698 | 0.107039 | 0.244 | 0.107 | 1 |
| P02787 | TF | DRNTyEKyLGEEyVK | Y669 | 0.063 | 97.43 | 698 | 0.027637 | 0.063 | 0.0276 | 1 |
| Q9BZZ2 | SIGLEC1 | LLLCRVDsDPPAQLR | S1009 | 0.106 | 82.84 | 1709 | 0.084261 | 0.106 | 0.0843 | 1 |
| Q9BZZ2 | SIGLEC1 | MLEDEGVyICEASNT | Y1065 | 0.143 | 90.41 | 1709 | 0.062732 | 0.143 | 0.0627 | 1 |
| O75469 | NR1I2 | GYHFNVMtCEGCKGF | T57 | 0 | 97.42 | 434 | 0 | 0 | 0 | 1 |
| Q86X55 | CARM1 | QDYVRTGtyQRAILQ | T171 | 0.073 | 97.8 | 608 | 0.04986 | 0.073 | 0.0499 | 0.9 |
| Q86X55 | CARM1 | DYVRTGtyQRAILQN | Y172 | 0.054 | 98.5 | 608 | 0.023689 | 0.054 | 0.0237 | 0.9412 |
| Q86X55 | CARM1 | KIYAVEAsTMAQHAE | S216 | 0.099 | 89.98 | 608 | 0.078697 | 0.099 | 0.0787 | 1 |
| Q86X55 | CARM1 | LIANkRQsYDISIVA | S447 | 0 | 97.24 | 608 | 0 | 0 | 0 | 0.875 |
| Q3YEC7 | RABL6 | VIRGDRNtGKTALWH | T54 | 0.01 | 95.64 | 729 | 0.00683 | 0.01 | 0.0068 | 1 |
| Q9H0A0 | NAT10 | SRQsILNsLSRGKkA | S593 | 0.143 | 78.17 | 1025 | 0.113673 | 0.143 | 0.1137 | 1 |
| Q9NPD7 | NRN1 | LGDSMANyPQGLDDK | Y52 | 0.28 | 84.65 | 142 | 0.122832 | 0.28 | 0.1228 | 1 |
| O43488 | AKR7A2 | RRMDAPAsAAAVrAF | S56 | 0 | 98.46 | 359 | 0 | 0 | 0 | 1 |
| P29466 | CASP1 | HRPEHKtsDSTFLVF | S227 | 0 | 98.61 | 404 | 0 | 0 | 0 | 1 |
| P46779 | RPL28 | kRNkQTYsTEPNNLk | S26 | 0.103 | 96.62 | 137 | 0.081876 | 0.103 | 0.0819 | 0.8889 |
| P46779 | RPL28 | INkNARAtLssIRHM | T89 | 0.001 | 97.05 | 137 | 0.000683 | 0.001 | 0.0007 | 0.8 |
| Q9H6A9 | PCNX3 | LITYYVKsIIYYVSR | S1491 | 0.002 | 95.15 | 2034 | 0.00159 | 0.002 | 0.0016 | 1 |
| Q8WWN8 | ARAP3 | sPGELALyKSEQAFS | Y422 | 0.208 | 90.32 | 1544 | 0.091247 | 0.208 | 0.0912 | 1 |
| Q6P3W7 | SCYL2 | QRDNLQKsQFFKGLP | S350 | 0.088 | 90.24 | 929 | 0.069952 | 0.088 | 0.07 | 1 |
| Q6P3W7 | SCYL2 | MDLLLtKtPPDEIKN | T440 | 0.032 | 85.66 | 929 | 0.021856 | 0.032 | 0.0219 | 1 |
| P23229 | ITGA6 | TFTKDFHyIVFGAPG | Y202 | 0.041 | 87.71 | 1130 | 0.017986 | 0.259 | 0.1115 | 1 |
| P23229 | ITGA6 | kDEITFVsGAPRANH | S317 | 0.006 | 97.92 | 1130 | 0.004769 | 0.006 | 0.0048 | 1 |
| P23229 | ITGA6 | IVIGAPQyFDRDGEV | Y379 | 0.238 | 94.36 | 1130 | 0.104407 | 0.238 | 0.1044 | 1 |
| Q8WWP7 | GIMAP1 | GkSATGNsILGQRRF | S46 | 0.031 | 98 | 306 | 0.024642 | 0.031 | 0.0246 | 1 |
| Q5H9S7 | DCAF17 | HGILIVMySSGLVRL | Y228 | 0.21 | 95 | 520 | 0.092124 | 0.21 | 0.0921 | 1 |
| O15230 | LAMA5 | DLWVLERsMDFGRty | S175 | 0.005 | 95.43 | 3695 | 0.003975 | 0.005 | 0.004 | 1 |

| | | | | | | | | | | |
|---|---|---|---|---|---|---|---|---|---|---|
| P14618 | PKM | MCRLDIDsPPItARN | S37 | 0.036 | 97.78 | 531 | 0.028617 | 0.036 | 0.0286 | 1 |
| P14618 | PKM | RNtGIICtIGPAsRs | T50 | 0.06 | 98.88 | 531 | 0.040981 | 0.061 | 0.0561 | 1 |
| P14618 | PKM | ICtIGPAsRsVEtLk | S55 | 0 | 98.62 | 531 | 0 | 0 | 0 | 1 |
| P14618 | PKM | tHEyHAEtIkNVrtA | T87 | 0.011 | 98.83 | 531 | 0.007513 | 0.011 | 0.0075 | 1 |
| P14618 | PKM | IkNVrtAtEsFAsDP | T95 | 0.073 | 98.84 | 531 | 0.04986 | 0.12 | 0.0791 | 0.8889 |
| P14618 | PKM | kGATLkItLDNAyME | T143 | 0.077 | 98.32 | 531 | 0.052592 | 0.102 | 0.0638 | 1 |
| P14618 | PKM | ENILWLDykNICkVV | Y161 | 0.059 | 97.05 | 531 | 0.025882 | 0.059 | 0.0259 | 0.9 |
| P14618 | PKM | yVDDGLIsLQVkQkG | S182 | 0.05 | 98.08 | 531 | 0.039746 | 0.05 | 0.0397 | 1 |
| P14618 | PKM | kGADFLVtEVENGGs | T195 | 0.015 | 98.81 | 531 | 0.010245 | 0.015 | 0.0102 | 1 |
| P14618 | PKM | DVDMVFAsFIRkAsD | S243 | 0.026 | 98.09 | 531 | 0.020668 | 0.026 | 0.0207 | 1 |
| P14618 | PKM | GKNIkIIskIENHEG | S269 | 0.001 | 98.91 | 531 | 0.000795 | 0.001 | 0.0008 | 1 |
| P14618 | PKM | FDEILEAsDGIMVAR | S287 | 0.052 | 98.88 | 531 | 0.041336 | 0.052 | 0.0413 | 1 |
| P14618 | PKM | REAEAAIyHLQLFEE | Y390 | 0.21 | 98.79 | 531 | 0.092124 | 0.21 | 0.0921 | 1 |
| P14618 | PKM | APItsDPtEAtAVGA | T409 | 0.096 | 97.12 | 531 | 0.065569 | 0.08 | 0.082 | 1 |
| P14618 | PKM | AVGAVEAsFkCCsGA | S420 | 0.061 | 98.82 | 531 | 0.04849 | 0.08 | 0.0683 | 1 |
| P14618 | PKM | EAsFkCCsGAIIVLt | S425 | 0.115 | 98.34 | 531 | 0.091415 | 0.127 | 0.1125 | 1 |
| P14618 | PKM | sGAIIVLtksGRSAH | T432 | 0.146 | 98.17 | 531 | 0.09972 | 0.146 | 0.0997 | 1 |
| P14618 | PKM | SAHQVARyRPRAPII | Y444 | 0.112 | 98.89 | 531 | 0.049133 | 0.112 | 0.0491 | 0.9091 |
| P14618 | PKM | RAPIIAVtRNPQtAR | T454 | 0.042 | 98.56 | 531 | 0.028687 | 0.042 | 0.0287 | 1 |
| P14618 | PKM | AVtRNPQtARQAHLy | T459 | 0.103 | 98.74 | 531 | 0.07035 | 0.206 | 0.1407 | 1 |
| P14618 | PKM | GDVVIVLtGWRPGsG | T513 | 0.06 | 96.95 | 531 | 0.040981 | 0.06 | 0.041 | 1 |
| Q8N398 | VWA5B2 | GSLsSAEyEARVRAR | Y292 | 0.068 | 90.53 | 1242 | 0.029831 | 0.068 | 0.0298 | 1 |
| Q8N398 | VWA5B2 | KALEPALsDIsVDWF | S525 | 0.057 | 92.43 | 1242 | 0.04531 | 0.057 | 0.0453 | 1 |
| Q15573 | TAF1A | KNIGVMNyLKISLQH | Y133 | 0.221 | 91.93 | 450 | 0.09695 | 0.221 | 0.0969 | 1 |
| O95185 | UNC5C | GLIVREVsIEIsRQQ | S123 | 0.156 | 96.09 | 931 | 0.124007 | 0.156 | 0.124 | 1 |
| Q8TEX9 | IPO4 | DPDLRRCtYSLFAAL | T586 | 0 | 98.03 | 1081 | 0 | 0 | 0 | 1 |
| Q8TEX9 | IPO4 | LARVVPsyMQAVNRE | Y755 | 0.035 | 98.71 | 1081 | 0.015354 | 0.035 | 0.0154 | 1 |
| P09651 | HNRNPA1 | IGGLsFEtTDEsLrS | T25 | 0 | 95.54 | 372 | 0 | 0 | 0 | 1 |
| P09651 | HNRNPA1 | SrGFGFVtyATVEEV | T61 | 0.013 | 97.22 | 372 | 0.008879 | 0.013 | 0.0089 | 1 |
| P09651 | HNRNPA1 | rGFGFVtyATVEEVD | Y62 | 0 | 97.69 | 372 | 0 | 0 | 0 | 1 |
| P09651 | HNRNPA1 | VTFDDHDsVDkIVIQ | S158 | 0.058 | 95.54 | 372 | 0.046105 | 0.116 | 0.0922 | 1 |

| | | | | | | | | | | |
|---|---|---|---|---|---|---|---|---|---|---|
| O75077 | ADAM23 | VVLVAVEtWtEKDQI | T352 | 0 | 97.42 | 832 | 0 | 0 | 0 | 1 |
| O75077 | ADAM23 | LVAVEtWtEKDQIDI | T354 | 0.178 | 97.07 | 832 | 0.121576 | 0.178 | 0.1216 | 1 |
| Q6ZTA4 | TRIM67 | GVFMPALsLNRNVQV | S756 | 0.008 | 95.42 | 783 | 0.006359 | 0.008 | 0.0064 | 1 |
| Q12904 | AIMP1 | HPDADsLyVEEVDVG | Y175 | 0.057 | 97.33 | 312 | 0.025005 | 0.057 | 0.025 | 1 |
| Q12904 | AIMP1 | AKMRGVLsQAMVMCA | S223 | 0.11 | 97.1 | 312 | 0.087441 | 0.11 | 0.0874 | 1 |
| P16152 | CBR1 | FsGDVVLtARDVTrG | T36 | 0.001 | 98.95 | 277 | 0.000683 | 0.001 | 0.0007 | 0.9 |
| Q96AA3 | RFT1 | NVRLTLLystTLFLA | Y55 | 0.28 | 98.34 | 541 | 0.122832 | 0.28 | 0.1228 | 1 |
| Q96AA3 | RFT1 | RLTLLystTLFLARE | T57 | 0.099 | 96.75 | 541 | 0.067618 | 0.099 | 0.0676 | 0.9 |
| Q96AA3 | RFT1 | VAAAVLEsLLKLALL | S331 | 0.157 | 93.75 | 541 | 0.124802 | 0.157 | 0.1248 | 1 |
| Q96AA3 | RFT1 | SKEEVDRyNFVMLAL | Y404 | 0.222 | 89.99 | 541 | 0.097388 | 0.345 | 0.1272 | 1 |
| A5A3E0 | POTEF | APrAVFPsIVGRPRQ | S733 | 0.043 | 96.49 | 1075 | 0.034181 | 0.043 | 0.0342 | 1 |
| A5A3E0 | POTEF | SSSLEksyELPDGQV | Y940 | 0.269 | 96.41 | 1075 | 0.118006 | 0.269 | 0.118 | 1 |
| A5A3E0 | POTEF | MWISkQEyDEsGPsI | Y1062 | 0.109 | 97.1 | 1075 | 0.047817 | 0.109 | 0.0478 | 1 |
| O60232 | ZNRD2 | IsRLMGDyLLrGYRM | Y41 | 0.242 | 96.22 | 199 | 0.106162 | 0.242 | 0.1062 | 1 |
| O43293 | DAPK3 | LGETKQEtLTNISAV | T225 | 0.061 | 90.29 | 454 | 0.041664 | 0.061 | 0.0417 | 1 |
| Q92995 | USP13 | DPtQDFNtQMtkLGH | T385 | 0 | 94.47 | 863 | 0 | 0.117 | 0.0799 | 1 |
| Q92995 | USP13 | QDFNtQMtkLGHGLL | T388 | 0 | 95.76 | 863 | 0 | 0 | 0 | 0.9 |
| Q9Y2T7 | YBX2 | NPRKFLRsVGDGETV | S137 | 0.006 | 92.7 | 364 | 0.004769 | 0.006 | 0.0048 | 1 |
| Q9BXJ9 | NAA15 | LGkkEEAyELVRRGL | Y66 | 0.23 | 94.49 | 866 | 0.100898 | 0.284 | 0.1246 | 1 |
| Q9BXJ9 | NAA15 | LEDAADVyRGLQERN | Y247 | 0.176 | 98.49 | 866 | 0.077209 | 0.3 | 0.1316 | 1 |
| Q9BXJ9 | NAA15 | MLERLkIyEEAWTkY | Y280 | 0.052 | 96.49 | 866 | 0.022812 | 0.09 | 0.0395 | 1 |
| Q9BXJ9 | NAA15 | RKITLRsyVDLLkLE | Y538 | 0.014 | 97.67 | 866 | 0.006142 | 0.014 | 0.0061 | 1 |
| B2RTY4 | MYO9A | NPkYVKMyDNHQLGK | Y203 | 0.091 | 82.76 | 2548 | 0.03992 | 0.18 | 0.0785 | 1 |
| P23468 | PTPRD | MLGAEDLtPEDDMPI | T277 | 0.149 | 86.78 | 1912 | 0.101769 | 0.149 | 0.1018 | 0.8889 |
| Q13107 | USP4 | SETRRLyyDEQEsEA | Y751 | 0.214 | 86.93 | 963 | 0.093879 | 0.214 | 0.0939 | 1 |
| Q13107 | USP4 | LyyDEQEsEAyEKHV | S756 | 0.077 | 84.26 | 963 | 0.061208 | 0.077 | 0.0612 | 1 |
| O15327 | INPP4B | LQRMQVHsPHLKDAL | S353 | 0.071 | 87.21 | 924 | 0.056439 | 0.098 | 0.0779 | 1 |
| P20592 | MX2 | KYIQRQQtINLVVVP | T258 | 0.046 | 90.85 | 715 | 0.031419 | 0.046 | 0.0314 | 1 |
| P20592 | MX2 | PRLAERLttELIMHI | T375 | 0.13 | 87.29 | 715 | 0.088792 | 0.202 | 0.138 | 1 |
| Q96P20 | NLRP3 | RGDILLSsLIRKKLL | S334 | 0 | 91.02 | 1036 | 0 | 0 | 0 | 1 |
| O15123 | ANGPT2 | YYWKGSGySLKATTM | Y482 | 0.226 | 97.27 | 496 | 0.099143 | 0.226 | 0.0991 | 1 |

| | | | | | | | | | | |
|---|---|---|---|---|---|---|---|---|---|---|
| Q9H9Y6 | POLR1B | DIFIGVVyyQRLRHM | Y998 | 0.059 | 98.01 | 1135 | 0.025882 | 0.059 | 0.0259 | 1 |
| Q9H9Y6 | POLR1B | IFIGVVyyQRLRHMV | Y999 | 0.042 | 97.12 | 1135 | 0.018425 | 0.042 | 0.0184 | 1 |
| O75112 | LDB3 | HMEDGEPyCEKDyIN | Y656 | 0.298 | 94.31 | 727 | 0.130728 | 0.298 | 0.1307 | 1 |
| Q9BQG0 | MYBBP1A | LLKSPAEsCDLLGDI | S613 | 0.063 | 91.25 | 1328 | 0.05008 | 0.063 | 0.0501 | 1 |
| P09884 | POLA1 | DVPEkSEyLEVKYSA | Y452 | 0 | 95.87 | 1462 | 0 | 0 | 0 | 0.9333 |
| Q13617 | CUL2 | TSKLHRMyTDMsVSA | Y477 | 0.333 | 92.32 | 745 | 0.146082 | 0.333 | 0.1461 | 1 |
| P24298 | GPT | NPVQGAMysFPRVQL | Y413 | 0.079 | 96.73 | 496 | 0.034656 | 0.079 | 0.0347 | 0.9375 |
| P0C1S8 | WEE2 | IGVGEFGtVYKCIKR | T225 | 0.179 | 93.24 | 567 | 0.122259 | 0.179 | 0.1223 | 1 |
| Q9UKI8 | TLK1 | RINNEDNsQFKDHPT | S444 | 0.068 | 93.9 | 766 | 0.054054 | 0.068 | 0.0541 | 1 |
| Q7Z5R6 | APBB1IP | GFKDLNEsLNALEDQ | S55 | 0 | 89.22 | 666 | 0 | 0.09 | 0.0915 | 1 |
| Q9GZV3 | SLC5A7 | IVWVMRItVFVFGAS | T384 | 0.122 | 95.45 | 580 | 0.083328 | 0.122 | 0.0833 | 1 |
| Q9NR30 | DDX21 | EEILsVAyKkDSEDN | Y361 | 0.272 | 92.94 | 783 | 0.119322 | 0.272 | 0.1193 | 0.8333 |
| Q13509 | TUBB3 | GTGSGMGtLLISKVr | T149 | 0.001 | 98.13 | 450 | 0.000683 | 0.001 | 0.0007 | 1 |
| Q13509 | TUBB3 | NtFsVVPsPKVSDTV | S172 | 0.007 | 82.76 | 450 | 0.005564 | 0.007 | 0.0056 | 0.8333 |
| Q13509 | TUBB3 | PRHGRyLtVAtVFRG | T312 | 0.047 | 96.1 | 450 | 0.032102 | 0.078 | 0.0533 | 1 |
| Q13509 | TUBB3 | PPRGLkMssTFIGNS | S364 | 0.002 | 95.05 | 450 | 0.00159 | 0.002 | 0.0016 | 1 |
| Q13509 | TUBB3 | PRGLkMssTFIGNST | S365 | 0 | 95.14 | 450 | 0 | 0 | 0 | 1 |
| Q13509 | TUBB3 | QELFkrIsEQFtAMF | S382 | 0.139 | 95.3 | 450 | 0.110493 | 0.139 | 0.1105 | 1 |
| P49959 | MRE11 | DALDDENtFKILVAT | T12 | 0.098 | 96.82 | 708 | 0.066935 | 0.098 | 0.0669 | 1 |
| P49959 | MRE11 | EQQLFYIsQPGSSVV | S264 | 0 | 98.44 | 708 | 0 | 0 | 0 | 1 |
| P49959 | MRE11 | PFSVLRFsQkFVDrV | S382 | 0.125 | 90.92 | 708 | 0.099364 | 0.125 | 0.0994 | 1 |
| Q92835 | INPP5D | ETQLPIytPLTHHGE | T835 | 0.079 | 98.43 | 1189 | 0.053958 | 0.079 | 0.054 | 0.875 |
| Q9H8X9 | ZDHHC11 | NVRLMKNySQPMPLF | Y106 | 0.331 | 92.51 | 412 | 0.145205 | 0.331 | 0.1452 | 1 |
| O43692 | PI15 | NWIGEAPyKVGVPCS | Y223 | 0.119 | 98.6 | 258 | 0.052204 | 0.119 | 0.0522 | 0.8462 |
| Q9BVA1 | TUBB2B | QLERINVyYNEATGN | Y50 | 0.029 | 96.51 | 445 | 0.012722 | 0.058 | 0.0254 | 0.8 |
| Q9BVA1 | TUBB2B | MPGFAPLtsrGsQQy | T274 | 0.168 | 88.99 | 445 | 0.114746 | 0.168 | 0.1147 | 0.8571 |
| Q9BVA1 | TUBB2B | QELFkRIsEQFtAMF | S382 | 0.133 | 94.65 | 445 | 0.105724 | 0.133 | 0.1057 | 1 |
| O60500 | NPHS1 | EDNGLtLtCEAFsEA | T416 | 0.214 | 90.15 | 1241 | 0.146165 | 0.214 | 0.1462 | 1 |
| O60500 | NPHS1 | tLtCEAFsEAFTKET | S421 | 0.18 | 81.19 | 1241 | 0.143085 | 0.18 | 0.1431 | 1 |
| A8MZ26 | EFCAB9 | NyQEFKLyTIIyTDK | Y164 | 0.161 | 94.56 | 197 | 0.070628 | 0.161 | 0.0706 | 1 |
| Q14139 | UBE4A | FNYRRPMyPILRYMW | Y749 | 0.013 | 95.72 | 1066 | 0.005703 | 0.093 | 0.0408 | 0.8182 |

| | | | | | | | | | | |
|---|---|---|---|---|---|---|---|---|---|---|
| P10586 | PTPRF | RGYRVyytPDsRRPP | T452 | 0.048 | 91.57 | 1907 | 0.032785 | 0.048 | 0.0328 | 1 |
| P10586 | PTPRF | DMGVGVFtPtIEART | T594 | 0.091 | 85.51 | 1907 | 0.062154 | 0.091 | 0.0622 | 0.8889 |
| P10586 | PTPRF | SSPVLVRtDEDVPsG | T703 | 0.01 | 89.4 | 1907 | 0.00683 | 0.01 | 0.0068 | 1 |
| P10586 | PTPRF | SKPKIVTttGAVPGR | T816 | 0.01 | 65.22 | 1907 | 0.00683 | 0.153 | 0.1277 | 1 |
| Q92544 | TM9SF4 | FTDVNKIyLHNHLSF | Y178 | 0.278 | 94.31 | 642 | 0.121955 | 0.305 | 0.1338 | 1 |
| Q92544 | TM9SF4 | NHLSFILyyHREDME | Y188 | 0.251 | 94.95 | 642 | 0.11011 | 0.277 | 0.1215 | 1 |
| Q6ZMW3 | EML6 | ALGMKDGsFIVLRVR | S389 | 0.02 | 96.54 | 1958 | 0.015898 | 0.02 | 0.0159 | 1 |
| Q6ZMW3 | EML6 | GSYLAVGsNDGPVDV | S427 | 0.005 | 93.15 | 1958 | 0.003975 | 0.005 | 0.004 | 1 |
| Q92616 | GCN1 | LGLVIRLtsADALRP | T2294 | 0 | 88.77 | 2671 | 0 | 0 | 0 | 1 |
| Q92616 | GCN1 | NTVVRAYsDQAIVNL | S2612 | 0 | 89.51 | 2671 | 0 | 0 | 0 | 1 |
| Q7Z5G4 | GOLGA7 | VRTLNNLyAEAEKLG | Y54 | 0.013 | 96.39 | 137 | 0.005703 | 0.013 | 0.0057 | 0.8 |
| Q9P0V3 | SH3BP4 | KkVTVGLyGPKHIHP | Y479 | 0 | 91.36 | 963 | 0 | 0 | 0 | 1 |
| Q14677 | CLINT1 | NWRRVYKsLLLLAYL | S82 | 0 | 96.31 | 625 | 0 | 0 | 0 | 1 |
| Q9BY64 | UGT2B28 | DLLGLPKtRAFITHG | T368 | 0.065 | 98.82 | 529 | 0.044396 | 0.065 | 0.0444 | 1 |
| Q53GQ0 | HSD17B12 | DFAsEDIyDkIkTGL | Y115 | 0.191 | 98.49 | 312 | 0.083789 | 0.339 | 0.1382 | 0.9167 |
| Q5XUX0 | FBXO31 | QYDNCLtyRRIYLPP | Y286 | 0.001 | 98.67 | 539 | 0.000439 | 0.001 | 0.0004 | 1 |
| Q9P2J2 | IGSF9 | SPRIDPDyVGRVRLQ | Y81 | 0.019 | 92.08 | 1179 | 0.008335 | 0.019 | 0.0083 | 1 |
| Q9P2J2 | IGSF9 | VRLQKGAsLQIEGLR | S92 | 0.144 | 90.08 | 1179 | 0.114468 | 0.144 | 0.1145 | 1 |
| P29033 | GJB2 | NtVDCFVsRPtEKTV | S183 | 0.125 | 93.42 | 226 | 0.099364 | 0.125 | 0.0994 | 1 |
| P41227 | NAA10 | VPHGHItsLAVkRSH | S74 | 0.061 | 98.32 | 235 | 0.04849 | 0.061 | 0.0485 | 1 |
| Q9H0X4 | FAM234A | EVSGHLYsGSTGHQI | S243 | 0.013 | 90.96 | 552 | 0.010334 | 0.013 | 0.0103 | 1 |
| P06731 | CEACAM5 | tRNDTASyKCETQNP | Y213 | 0.051 | 92.89 | 702 | 0.022373 | 0.051 | 0.0224 | 1 |
| P06731 | CEACAM5 | NPVSARRsDSVILNV | S226 | 0.081 | 95.46 | 702 | 0.064388 | 0.081 | 0.0644 | 1 |
| P48058 | GRIA4 | REEVIDFsKPFMsLG | S514 | 0.09 | 96.02 | 902 | 0.071542 | 0.09 | 0.0715 | 1 |
| P48058 | GRIA4 | TKEFFRRsKIAVYEK | S684 | 0.175 | 92.26 | 902 | 0.13911 | 0.175 | 0.1391 | 1 |
| P13796 | LCP1 | DIkDSkAyyHLLEQV | Y299 | 0.048 | 97.68 | 627 | 0.021057 | 0.086 | 0.0377 | 1 |
| P13796 | LCP1 | IkDSkAyyHLLEQVA | Y300 | 0.01 | 97.39 | 627 | 0.004387 | 0.01 | 0.0044 | 1 |
| Q7Z7M9 | GALNT5 | KCKSFKWyLENVFPD | Y795 | 0 | 98.52 | 940 | 0 | 0 | 0 | 1 |
| P62873 | GNB1 | TDsRLLVsAsQDGkL | S72 | 0 | 98.76 | 340 | 0 | 0 | 0 | 1 |
| P62873 | GNB1 | sRLLVsAsQDGkLII | S74 | 0 | 97.56 | 340 | 0 | 0 | 0 | 1 |
| Q13610 | PWP1 | YDKSVALyDCRSPDE | Y327 | 0.068 | 96.74 | 501 | 0.029831 | 0.068 | 0.0298 | 1 |

| | | | | | | | | | | |
|---|---|---|---|---|---|---|---|---|---|---|
| Q6ZMY6 | WDR88 | IKDHHTRsITSCCFD | S232 | 0.114 | 94.55 | 472 | 0.09062 | 0.145 | 0.1153 | 0.8571 |
| P04080 | CSTB | VHLrVFQsLPHENkP | S72 | 0.17 | 97.43 | 98 | 0.135135 | 0.17 | 0.1351 | 1 |
| P20853 | CYP2A7 | EDRTKMPyMEAVIHE | Y351 | 0.294 | 98.87 | 494 | 0.128974 | 0.294 | 0.129 | 1 |
| Q5VTT5 | MYOM3 | IFSKVKyyNVEYMKt | Y1245 | 0.273 | 85.73 | 1437 | 0.119761 | 0.425 | 0.1433 | 1 |
| Q5VTT5 | MYOM3 | KDSDKGKyTLEIAAG | Y1291 | 0.009 | 96.98 | 1437 | 0.003948 | 0.009 | 0.0039 | 1 |
| Q68CQ1 | MROH7 | DPIMKVLsIRGLVIL | S1022 | 0.007 | 94.89 | 1323 | 0.005564 | 0.007 | 0.0056 | 0.9091 |
| Q8N6G5 | CSGALNACT2 | QYRMCIQsKAMNEAS | S505 | 0.022 | 93.28 | 542 | 0.017488 | 0.022 | 0.0175 | 1 |
| P68036 | UBE2L3 | RADLAEEySkDRKKF | Y129 | 0.247 | 96.59 | 154 | 0.108355 | 0.247 | 0.1084 | 0.9091 |
| Q9H845 | ACAD9 | IFTVFAKtEVVDsDG | T229 | 0.005 | 98.19 | 621 | 0.003415 | 0.005 | 0.0034 | 1 |
| Q9H845 | ACAD9 | ILNSGRFsMGsVVAG | S306 | 0.12 | 91.36 | 621 | 0.09539 | 0.12 | 0.0954 | 0.8889 |
| Q9UBS8 | RNF14 | FLECRHVyCKACLKD | Y242 | 0.032 | 96.45 | 474 | 0.014038 | 0.032 | 0.014 | 1 |
| Q8TDX9 | PKD1L1 | DItFTAVtKETIPLE | T530 | 0.203 | 89.11 | 2849 | 0.138652 | 0.203 | 0.1387 | 1 |
| O95922 | TTLL1 | sQsNKEAyVIsLyIN | Y178 | 0.184 | 91.42 | 423 | 0.080718 | 0.184 | 0.0807 | 1 |
| O95922 | TTLL1 | NKEAyVIsLyINNPL | S181 | 0.132 | 95.01 | 423 | 0.104929 | 0.132 | 0.1049 | 0.875 |
| O95922 | TTLL1 | RKFDLRLyVLVSTyR | Y200 | 0.013 | 97.92 | 423 | 0.005703 | 0.013 | 0.0057 | 0.9286 |
| O95922 | TTLL1 | LyVLVSTyRPLRCyM | Y206 | 0.042 | 96.24 | 423 | 0.018425 | 0.042 | 0.0184 | 0.8333 |
| O95922 | TTLL1 | TyRPLRCyMyKLGFC | Y212 | 0.028 | 97.16 | 423 | 0.012283 | 0.028 | 0.0123 | 1 |
| O95922 | TTLL1 | RPLRCyMyKLGFCRF | Y214 | 0.016 | 97.06 | 423 | 0.007019 | 0.016 | 0.007 | 1 |
| Q9BV36 | MLPH | GSAkVIRsLHGRLQG | S140 | 0.178 | 90.14 | 600 | 0.141495 | 0.178 | 0.1415 | 1 |
| O75925 | PIAS1 | YLVKQLsstVLLQRL | S290 | 0.154 | 97.26 | 651 | 0.122417 | 0.154 | 0.1224 | 0.8571 |
| O75925 | PIAS1 | GIrNPDHsRALIkEk | S308 | 0.007 | 94.79 | 651 | 0.005564 | 0.007 | 0.0056 | 1 |
| Q9H6R3 | ACSS3 | LIFGGFAsKELssRI | S204 | 0.118 | 83.85 | 686 | 0.0938 | 0.182 | 0.1447 | 1 |
| Q9H6R3 | ACSS3 | GFAsKELssRIDHVK | S208 | 0.054 | 88.98 | 686 | 0.042925 | 0.054 | 0.0429 | 1 |
| P78368 | CSNK1G2 | DRTFtLKtVLMIAIQ | T141 | 0.003 | 98.73 | 415 | 0.002049 | 0.003 | 0.002 | 1 |
| P78368 | CSNK1G2 | DFGLAKEyIDPETKK | Y195 | 0.123 | 98.49 | 415 | 0.053958 | 0.123 | 0.054 | 1 |
| P12830 | CDH1 | DCTGRQRtAyFsLDT | T66 | 0.118 | 74.96 | 882 | 0.080596 | 0.118 | 0.0806 | 0.8571 |
| P12830 | CDH1 | RQRtAyFsLDTRFKV | S70 | 0.165 | 86.49 | 882 | 0.131161 | 0.165 | 0.1312 | 1 |
| Q8N9U0 | TC2N | SSTPLTLsFFVKVGM | S388 | 0.157 | 88.92 | 490 | 0.124802 | 0.157 | 0.1248 | 1 |
| Q8N9U0 | TC2N | AVNQWKEtVINPEKV | T473 | 0.045 | 91.33 | 490 | 0.030736 | 0.045 | 0.0307 | 1 |
| Q8IYW5 | RNF168 | PFCRRRVsSWTRYHT | S59 | 0.035 | 95.92 | 571 | 0.027822 | 0.035 | 0.0278 | 1 |
| P48960 | ADGRE5 | NVtMGQssARMKLNW | S378 | 0.123 | 85.87 | 835 | 0.097775 | 0.123 | 0.0978 | 0.8571 |

| | | | | | | | | | | |
|---|---|---|---|---|---|---|---|---|---|---|
| Q8IXK2 | GALNT12 | AGGLFAVsKKYFEYL | S318 | 0.06 | 98.91 | 581 | 0.047695 | 0.06 | 0.0477 | 1 |
| Q9BYT3 | STK33 | GAAIEEIyTFGRILG | Y116 | 0.059 | 97.53 | 514 | 0.025882 | 0.059 | 0.0259 | 1 |
| P02787 | TF | NQLRGKKsCHtGLGR | S136 | 0 | 98.69 | 698 | 0 | 0 | 0 | 1 |
| P02787 | TF | RGKKsCHtGLGRsAG | T139 | 0.003 | 97.07 | 698 | 0.002049 | 0.003 | 0.002 | 1 |
| P02787 | TF | NIPIGLLyCDLPEPR | Y155 | 0.095 | 90.73 | 698 | 0.041675 | 0.095 | 0.0417 | 1 |
| Q8WU67 | ABHD3 | HLSEELGyRCVVFNN | Y169 | 0.022 | 98.56 | 409 | 0.009651 | 0.022 | 0.0097 | 0.8667 |
| P78346 | RPP30 | DSTMRRytIssALNL | T169 | 0.062 | 92.66 | 268 | 0.042347 | 0.062 | 0.0423 | 0.9091 |
| P78346 | RPP30 | MRRytIssALNLMQI | S172 | 0.082 | 91.15 | 268 | 0.065183 | 0.082 | 0.0652 | 0.9 |
| Q8IYR2 | SMYD4 | RLQEASQtISDLERN | T165 | 0.093 | 94.03 | 804 | 0.06352 | 0.093 | 0.0635 | 1 |
| P35580 | MYH10 | EsFGNAktVkNDNSS | T233 | 0.009 | 85.58 | 1976 | 0.006147 | 0.146 | 0.1019 | 1 |
| P35580 | MYH10 | ERTFHIFyQLLSGAG | Y285 | 0.009 | 95.31 | 1976 | 0.003948 | 0.009 | 0.0039 | 0.9375 |
| Q9BQS8 | FYCO1 | IPTTRCNsHKENIQG | S1427 | 0.019 | 94.35 | 1478 | 0.015103 | 0.019 | 0.0151 | 1 |
| Q96Q15 | SMG1 | LAGQPAVtVRHGFDL | T1047 | 0 | 98.13 | 3661 | 0 | -0.0435 | 0.0299 | 1 |
| O00469 | PLOD2 | DILLtLDyPKEALkL | Y323 | 0.111 | 96.15 | 737 | 0.048694 | 0.111 | 0.0487 | 1 |
| O00469 | PLOD2 | RQDEKCDyYFsVDAD | Y387 | 0.238 | 98.43 | 737 | 0.104407 | 0.238 | 0.1044 | 1 |
| O00469 | PLOD2 | EKCDyYFsVDADVVL | S390 | 0.002 | 98.53 | 737 | 0.00159 | 0.002 | 0.0016 | 1 |
| O00469 | PLOD2 | YYARSEDyVDIVQGN | Y444 | 0.051 | 97.55 | 737 | 0.022373 | 0.051 | 0.0224 | 1 |
| O00469 | PLOD2 | MEHYGkWsGGkHHDS | S590 | 0.132 | 87.86 | 737 | 0.104929 | 0.132 | 0.1049 | 1 |
| O00469 | PLOD2 | GGYENVPtDDIHMKQ | T608 | 0.097 | 85.36 | 737 | 0.066252 | 0.097 | 0.0663 | 1 |
| O00469 | PLOD2 | REFIAPVtLKVFAGY | T634 | 0.004 | 96.35 | 737 | 0.002732 | 0.004 | 0.0027 | 1 |
| O00469 | PLOD2 | LLNFVVKysPERQRS | Y655 | 0.093 | 95.07 | 737 | 0.040798 | 0.093 | 0.0408 | 1 |
| O00469 | PLOD2 | LNFVVKysPERQRSL | S656 | 0.082 | 93.73 | 737 | 0.065183 | 0.082 | 0.0652 | 1 |
| Q15139 | PRKD1 | DTLRKRHyWRLDsKC | Y443 | 0.302 | 92.59 | 912 | 0.132483 | 0.302 | 0.1325 | 1 |
| Q15149 | PLEC | EEVGFDWsDRNTNMT | S782 | 0.009 | 92.265 | 4684 | 0.007154 | 0.009 | 0.0072 | 1 |
| Q15149 | PLEC | HLKENAAyFQFFSDV | Y861 | 0.135 | 96.01 | 4684 | 0.059223 | 0.234 | 0.1027 | 1 |
| Q15149 | PLEC | QLHVDMKsLLAWQSL | S1029 | 0.005 | 90.53 | 4684 | 0.003975 | 0.005 | 0.004 | 1 |
| Q15149 | PLEC | yQAFLRDsQDAGGFG | S1077 | 0 | 86.89 | 4684 | 0 | 0 | 0 | 0.7222 |
| Q9NZM6 | PKD2L2 | TMsQLSstLSRCVKD | T399 | 0.086 | 85.79 | 624 | 0.058739 | 0.086 | 0.0587 | 1 |
| Q8IU85 | CAMK1D | LKPENLLyysQDEES | Y152 | 0.16 | 93.82 | 385 | 0.07019 | 0.16 | 0.0702 | 1 |
| Q8WVV9 | HNRNPLL | sHHkVsVsPVVHVRG | S75 | 0 | 91.82 | 542 | 0 | 0 | 0 | 1 |
| P06702 | S100A9 | INTFHQYsVKLGHPD | S23 | 0.004 | 98.65 | 114 | 0.00318 | 0.004 | 0.0032 | 1 |

| | | | | | | | | | | |
|---|---|---|---|---|---|---|---|---|---|---|
| P06702 | S100A9 | VKLGHPDtLNQGEFk | T31 | 0.182 | 98.76 | 114 | 0.124308 | 0.182 | 0.1243 | 1 |
| P54296 | MYOM2 | SANRHGLsEPsEIts | S594 | 0.096 | 91.29 | 1465 | 0.076312 | 0.096 | 0.0763 | 0.9 |
| P07858 | CTSB | WTRKGLVsGGLyESH | S169 | 0.049 | 98.87 | 339 | 0.038951 | 0.049 | 0.039 | 0.8182 |
| Q9UHW9 | SLC12A6 | VkHPRLLtFAsQLkA | T778 | 0.005 | 95.7 | 1150 | 0.003415 | 0.005 | 0.0034 | 1 |
| Q9UHW9 | SLC12A6 | MEFLEVLtEGLERVL | T1129 | 0.042 | 93.08 | 1150 | 0.028687 | 0.042 | 0.0287 | 0.8889 |
| P83111 | LACTB | AsISKsLtMVALAKL | T170 | 0 | 98.65 | 547 | 0 | 0 | 0 | 1 |
| P83111 | LACTB | VTTRLLIsHLSGIRH | S215 | 0.017 | 97.31 | 547 | 0.013514 | 0.017 | 0.0135 | 1 |
| P83111 | LACTB | ERASGCkyLDyMQKI | Y343 | 0.043 | 97.4 | 547 | 0.018863 | 0.043 | 0.0189 | 1 |
| P83111 | LACTB | TPyVDNSyKWAGGGF | Y393 | 0.047 | 94.83 | 547 | 0.020618 | 0.047 | 0.0206 | 0.9231 |
| Q9NUP9 | LIN7C | ANATAkAtVAAFAAS | T78 | 0.07 | 88.07 | 197 | 0.047811 | 0.07 | 0.0478 | 1 |
| O95379 | TNFAIP8 | IIKNLIKtVIKLAIL | T78 | 0.217 | 96.13 | 198 | 0.148214 | 0.217 | 0.1482 | 1 |
| Q6P158 | DHX57 | TKEGVRAsYNYCRQN | S1142 | 0.149 | 93.17 | 1386 | 0.118442 | 0.058 | 0.1316 | 1 |
| Q92743 | HTRA1 | IKKFLtEsHDRQAKG | S367 | 0.007 | 88.86 | 480 | 0.005564 | 0.007 | 0.0056 | 1 |
| Q9Y572 | RIPK3 | KLADFGLstFQGGSQ | S164 | 0.041 | 82.14 | 518 | 0.032592 | 0.041 | 0.0326 | 0.8889 |
| Q6STE5 | SMARCD3 | DsDGsIAsWELRVEG | S181 | 0.115 | 92.18 | 483 | 0.091415 | 0.115 | 0.0914 | 1 |
| P08559 | PDHA1 | NPTDHLItAyRAHGF | T116 | 0.033 | 98.9 | 390 | 0.022539 | 0.033 | 0.0225 | 1 |
| P08559 | PDHA1 | REILAELtGRKGGCA | T139 | 0.13 | 98.74 | 390 | 0.088792 | 0.13 | 0.0888 | 1 |
| P08559 | PDHA1 | FICENNRyGMGtsVE | Y227 | 0.226 | 98.73 | 390 | 0.099143 | 0.226 | 0.0991 | 1 |
| P08559 | PDHA1 | RAAAstDyykrGDFI | Y242 | 0.026 | 98.84 | 390 | 0.011406 | 0.026 | 0.0114 | 1 |
| P08559 | PDHA1 | AAAstDyykrGDFIP | Y243 | 0.106 | 98.82 | 390 | 0.046501 | 0.106 | 0.0465 | 1 |
| P08559 | PDHA1 | MELQTYRyHGHsMsD | Y289 | 0.133 | 98.63 | 390 | 0.058345 | 0.133 | 0.0583 | 1 |
| P08559 | PDHA1 | TYRyHGHsMsDPGVs | S293 | 0.131 | 97.84 | 390 | 0.104134 | 0.131 | 0.1041 | 1 |
| Q8IZL9 | CDK20 | ____MDQyCILGRIG | Y4 | 0.298 | 93.04 | 346 | 0.130728 | 0.298 | 0.1307 | 1 |
| Q969N2 | PIGT | NYFVRLytEPLLVNL | T510 | 0.012 | 96.48 | 578 | 0.008196 | 0.124 | 0.0847 | 1 |
| Q9NUX5 | POT1 | GSFLRIysLHTKLQS | S243 | 0.081 | 92.53 | 634 | 0.064388 | 0.081 | 0.0644 | 1 |
| P54727 | RAD23B | _MQVtLktLQQQtFk | T7 | 0.129 | 90.95 | 409 | 0.088109 | 0.129 | 0.0881 | 0.8333 |
| Q86TL0 | ATG4D | IHLCGRRyRFEGEGD | Y101 | 0.095 | 95 | 474 | 0.041675 | 0.095 | 0.0417 | 1 |
| Q86TL0 | ATG4D | PSCTVGFyAGDRkEF | Y398 | 0.127 | 98.31 | 474 | 0.055713 | 0.168 | 0.0737 | 0.9091 |
| A1L4K1 | FSD2 | PSSERAVyMtAPSPP | Y465 | 0.064 | 95.89 | 749 | 0.028076 | 0.064 | 0.0281 | 1 |
| A1L4K1 | FSD2 | QLQPGRSyIIyVRAL | Y538 | 0.124 | 92.39 | 749 | 0.054397 | 0.124 | 0.0544 | 1 |
| Q9BSU3 | NAA11 | SEVEPkyyADGEDAy | Y138 | 0.278 | 96.56 | 229 | 0.121955 | 0.278 | 0.122 | 0.9 |

| | | | | | | | | | | |
|---|---|---|---|---|---|---|---|---|---|---|
| Q7Z5P4 | HSD17B13 | YEFAKRQsILVLWDI | S61 | 0.018 | 98.67 | 300 | 0.014308 | 0.018 | 0.0143 | 1 |
| Q99081 | TCF12 | MCQLHLksEkPQTKL | S610 | 0.181 | 91.42 | 682 | 0.14388 | 0.181 | 0.1439 | 1 |
| Q7Z388 | DPY19L4 | VKINYSPyVNYFTRV | Y697 | 0.123 | 92.68 | 723 | 0.053958 | 0.257 | 0.1127 | 0.8889 |
| Q9Y4G8 | RAPGEF2 | IEDFLLtyRTFLSSP | Y302 | 0.111 | 93 | 1499 | 0.048694 | 0.111 | 0.0487 | 1 |
| O60312 | ATP10A | AVVGIVIyAGHETKA | Y281 | 0.024 | 93.09 | 1499 | 0.010528 | 0.091 | 0.0299 | 1 |
| Q9UBL6 | CPNE7 | EESTGKAsQYYILLI | S509 | 0.127 | 88.05 | 633 | 0.100954 | 0.127 | 0.101 | 1 |
| P08574 | CYC1 | CTFLRWAsEPEHDHR | S278 | 0.02 | 98.24 | 325 | 0.015898 | 0.02 | 0.0159 | 0.8889 |
| Q8NGH3 | OR2D3 | LIVVVLFyGSGIFTy | Y268 | 0.065 | 91.73 | 330 | 0.028515 | 0.065 | 0.0285 | 0.875 |
| Q16850 | CYP51A1 | LDrCIkEtLRLRPPI | T376 | 0 | 98.93 | 503 | 0 | 0.102 | 0.0697 | 0.8889 |
| Q8IWX7 | UNC45B | KKDLKQItSHLLDML | T279 | 0.215 | 95.27 | 931 | 0.146848 | 0.215 | 0.1468 | 1 |
| Q9BYX7 | POTEKP | APQAVFPsIVGRPRH | S33 | 0.007 | 97.18 | 375 | 0.005564 | 0.007 | 0.0056 | 1 |
| Q9BYX7 | POTEKP | AGRELTDyLMkILTE | Y188 | 0.198 | 97.87 | 375 | 0.08686 | 0.198 | 0.0869 | 1 |
| Q9BYX7 | POTEKP | MWISkQEyDEsGPsI | Y362 | 0.109 | 98.36 | 375 | 0.047817 | 0.109 | 0.0478 | 1 |
| Q13002 | GRIK2 | NRSLIVTtILEEPYV | T437 | 0.019 | 96.5 | 908 | 0.012977 | 0.019 | 0.013 | 1 |
| Q13002 | GRIK2 | AKQTKIEyGAVEDGA | Y682 | 0.029 | 94.34 | 908 | 0.012722 | 0.029 | 0.0127 | 1 |
| Q13002 | GRIK2 | TMTFFKKsKIsTyDK | S697 | 0.132 | 90.69 | 908 | 0.104929 | 0.132 | 0.1049 | 1 |
| Q9BUA3 | SPINDOC | ILEQHPHtLDLsPSE | T117 | 0.114 | 91.15 | 381 | 0.077864 | 0.114 | 0.0779 | 1 |
| A8MPY1 | GABRR3 | PDGNVLLsLRITVSA | S175 | 0.184 | 96.28 | 467 | 0.146264 | 0.184 | 0.1463 | 1 |
| Q8TC44 | POC1B | GRLIVsCsEDKTIKI | S163 | 0.001 | 97.5 | 478 | 0.000795 | 0.001 | 0.0008 | 1 |
| Q01484 | ANK2 | ATFSPIVtLEPRRRK | T1221 | 0.087 | 95.8 | 3957 | 0.059422 | 0.087 | 0.0594 | 1 |
| P63096 | GNAI1 | LLLGAGEsGkSTIVk | S44 | 0.034 | 97.7 | 354 | 0.027027 | 0.054 | 0.0479 | 1 |
| P63096 | GNAI1 | LNDSAAyyLNDLDRI | Y155 | 0.014 | 98.13 | 354 | 0.006142 | 0.014 | 0.0061 | 0.9444 |
| P54687 | BCAT1 | RVREMFGsGtACVVC | S331 | 0.021 | 98.23 | 386 | 0.016693 | 0.021 | 0.0167 | 1 |
| P78549 | NTHL1 | RRYQVLLsLMLSsQT | S135 | 0.006 | 98.09 | 312 | 0.004769 | 0.006 | 0.0048 | 1 |
| P78549 | NTHL1 | sQTKDQVtAGAMQRL | T147 | 0.054 | 97.58 | 312 | 0.036883 | 0.054 | 0.0369 | 1 |
| Q96NR8 | RDH12 | LQGTGVTtYAVHPGV | T225 | 0.003 | 98.81 | 316 | 0.002049 | 0.003 | 0.002 | 0.9231 |
| Q96BR5 | COA7 | NCEENQHsDsCYkLG | S68 | 0.026 | 98.19 | 231 | 0.020668 | 0.026 | 0.0207 | 1 |
| Q96BR5 | COA7 | IWACANAsRMYkLGD | S191 | 0.146 | 98.6 | 231 | 0.116058 | 0.146 | 0.1161 | 0.875 |
| Q9HCJ5 | ZSWIM6 | LLEAGPysGLGEIIH | S753 | 0.021 | 93.5 | 1215 | 0.016693 | 0.021 | 0.0167 | 1 |
| Q9NR09 | BIRC6 | LLAGGLLtyKsPAts | T574 | 0.164 | 88.08 | 4857 | 0.112014 | 0.2 | 0.1366 | 1 |
| Q7Z304 | MAMDC2 | GQKARLLsRPLRGVS | S571 | 0.032 | 94.77 | 686 | 0.025437 | 0.032 | 0.0254 | 0.8889 |

| | | | | | | | | | | |
|---|---|---|---|---|---|---|---|---|---|---|
| Q2TBF2 | WSCD2 | RAKYIGCyLDDTQSR | Y134 | 0.269 | 97.4 | 565 | 0.118006 | 0.269 | 0.118 | 0.8 |
| Q8IWZ6 | BBS7 | KGEGVFKsDNISTIS | S567 | 0.001 | 98.16 | 715 | 0.000795 | 0.001 | 0.0008 | 1 |
| P05186 | ALPL | QVPDsAGtATAyLCG | T113 | 0.001 | 98.89 | 524 | 0.000683 | 0.001 | 0.0007 | 0.9167 |
| P05186 | ALPL | sAGtATAyLCGVKAN | Y117 | 0.006 | 98.78 | 524 | 0.002632 | 0.006 | 0.0026 | 0.9444 |
| P05186 | ALPL | WAKDAGKsVGIVTTT | S160 | 0.117 | 98.8 | 524 | 0.093005 | 0.117 | 0.093 | 1 |
| P21580 | TNFAIP3 | VLRkALFstLkETDT | S128 | 0.11 | 96.7 | 790 | 0.087441 | 0.11 | 0.0874 | 1 |
| P21580 | TNFAIP3 | LRkALFstLkETDTR | T129 | 0.001 | 96.81 | 790 | 0.000683 | 0.132 | 0.0902 | 0.8571 |
| P21580 | TNFAIP3 | kEKLLKEyLMVIEIP | Y306 | 0.173 | 98.79 | 790 | 0.075893 | 0.235 | 0.1031 | 1 |
| O15550 | KDM6A | HTILGMNtVQLYMKV | T1131 | 0.133 | 97.55 | 1401 | 0.090841 | 0.133 | 0.0908 | 1 |
| Q8N752 | CSNK1A1L | SRRDDMEsLGyVFMy | S206 | 0.112 | 98.89 | 337 | 0.08903 | 0.112 | 0.089 | 1 |
| Q8N752 | CSNK1A1L | DDMEsLGyVFMyFNR | Y209 | 0 | 98.86 | 337 | 0 | 0 | 0 | 1 |
| Q8N752 | CSNK1A1L | sLGyVFMyFNRTSLP | Y213 | 0.022 | 98.83 | 337 | 0.009651 | 0.022 | 0.0097 | 1 |
| Q13505 | MTX1 | kISNPWQsPSGTLPA | S197 | 0.094 | 90.54 | 466 | 0.074722 | 0.094 | 0.0747 | 1 |
| O75051 | PLXNA2 | VKVLNCDtITQVKEK | T1524 | 0.05 | 93.56 | 1894 | 0.034151 | 0.05 | 0.0342 | 1 |
| Q5T764 | IFIT1B | QNVRVVEsVSLLGLI | S439 | 0.064 | 94.43 | 474 | 0.050875 | 0.064 | 0.0509 | 1 |
| O00148 | DDX39A | CPHVVVGtPGRILAL | T171 | 0.158 | 97.02 | 427 | 0.107916 | 0.158 | 0.1079 | 0.8 |
| O00148 | DDX39A | VQEIFRLtPHEkQCM | T217 | 0.121 | 98.34 | 427 | 0.082645 | 0.121 | 0.0826 | 1 |
| O95104 | SCAF4 | IDSIVRQsRHQFGTD | S73 | 0 | 96.79 | 1147 | 0 | 0 | 0 | 0.8182 |
| O94823 | ATP10B | NCsNIRIyERKEQTY | Y155 | 0.192 | 91.22 | 1461 | 0.084228 | 0.192 | 0.0842 | 1 |
| Q9UK23 | NAGPA | RAVEPLRtFsVLEPG | T103 | 0.118 | 93.53 | 515 | 0.080596 | 0.118 | 0.0806 | 0.8889 |
| Q9UK23 | NAGPA | VEPLRtFsVLEPGGP | S105 | 0.042 | 97.96 | 515 | 0.033386 | 0.042 | 0.0334 | 0.8 |
| Q9UK23 | NAGPA | RRATVEEtARAADCR | T126 | 0.043 | 98.17 | 515 | 0.02937 | 0.043 | 0.0294 | 1 |
| Q5SZQ8 | CELF3 | LTYCARDsALKAQSA | S60 | 0.164 | 94.35 | 465 | 0.130366 | 0.164 | 0.1304 | 1 |
| Q96BK5 | PINX1 | sFSLEEKsKISKNRV | S117 | 0.028 | 85.65 | 328 | 0.022258 | 0.028 | 0.0223 | 1 |
| Q8WXG1 | RSAD2 | FNVEEDMtEQIkALN | T234 | 0.138 | 98.37 | 361 | 0.094256 | 0.138 | 0.0943 | 1 |
| P63165 | SUMO1 | QrIADNHtPkELGME | T76 | 0.141 | 89.94 | 101 | 0.096305 | 0.141 | 0.0963 | 1 |
| O43189 | PHF1 | DVAHLVLyHLSVCCK | Y260 | 0.087 | 98.12 | 567 | 0.038166 | 0.087 | 0.0382 | 0.8125 |
| Q9H649 | NSUN3 | LINVIKVsELDGRKM | S190 | 0.107 | 98.27 | 340 | 0.085056 | 0.107 | 0.0851 | 0.8 |
| Q96F07 | CYFIP2 | YLMDGNVsNIYKLDA | S279 | 0.076 | 91.48 | 1278 | 0.060414 | 0.076 | 0.0604 | 1 |
| Q8NHS3 | MFSD8 | FLSSVGFsVVMMSIW | S54 | 0.131 | 89.28 | 518 | 0.104134 | 0.131 | 0.1041 | 1 |
| Q15124 | PGM5 | GGGLRRPtGLFEGQR | T31 | 0.009 | 96.93 | 567 | 0.006147 | 0.009 | 0.0061 | 1 |

| | | | | | | | | | | |
|---|---|---|---|---|---|---|---|---|---|---|
| Q15124 | PGM5 | GCTMVVGsDGRYFSR | S66 | 0.021 | 98.9 | 567 | 0.016693 | 0.021 | 0.0167 | 1 |
| Q15124 | PGM5 | SDKIyQIsKTIEEYA | S156 | 0 | 97.85 | 567 | 0 | 0 | 0 | 1 |
| Q15124 | PGM5 | FDADGDRyMILGQNG | Y299 | 0.073 | 98.75 | 567 | 0.032024 | 0.146 | 0.064 | 1 |
| Q15124 | PGM5 | VRATLRLyAEsyERD | Y522 | 0.022 | 98.56 | 567 | 0.009651 | 0.022 | 0.0097 | 0.9231 |
| Q6UX98 | ZDHHC24 | WLWPFLAsPLPGDGI | S264 | 0.071 | 96.22 | 284 | 0.056439 | 0.071 | 0.0564 | 1 |
| Q6P2Q9 | PRPF8 | PNIVIKGsELQLPFQ | S1887 | 0.125 | 82.12 | 2335 | 0.099364 | 0.125 | 0.0994 | 0.8889 |
| Q9HB96 | FANCE | AATTSMAyAKLMLTV | Y489 | 0.01 | 95.27 | 536 | 0.004387 | 0.01 | 0.0044 | 1 |
| Q9NUL7 | DDX28 | ALKSRLIsLEQLsFL | S276 | 0.17 | 95.51 | 540 | 0.135135 | 0.17 | 0.1351 | 1 |
| Q9NUL7 | DDX28 | LIsLEQLsFLVLDEA | S281 | 0.073 | 94.23 | 540 | 0.058029 | 0.073 | 0.058 | 1 |
| Q6P656 | CFAP161 | ATGQVLRyGQDFCLG | Y143 | 0.129 | 96.75 | 301 | 0.05659 | 0.129 | 0.0566 | 1 |
| Q969R2 | OSBP2 | NHYVWRKststVHNI | S682 | 0.04 | 94.19 | 916 | 0.031797 | 0.04 | 0.0318 | 1 |
| Q8IYK4 | COLGALT2 | QLDWELIyIGRKRMQ | Y467 | 0.011 | 98.68 | 626 | 0.004826 | 0.011 | 0.0048 | 1 |
| Q8IYK4 | COLGALT2 | NKHPVAEyKEyyEsR | Y538 | 0.186 | 93.83 | 626 | 0.081596 | 0.186 | 0.0816 | 1 |
| Q8IYK4 | COLGALT2 | VAEyKEyyEsRDLKA | Y542 | 0.042 | 95.39 | 626 | 0.018425 | 0.042 | 0.0184 | 1 |
| P56945 | BCAR1 | VRSQVTHySNLLCDL | Y815 | 0.146 | 97.4 | 870 | 0.064048 | 0.146 | 0.064 | 1 |
| Q15149 | PLEC | KNRSkGIyQsLEGAV | Y480 | 0.21 | 92.22 | 4684 | 0.092124 | 0.358 | 0.1419 | 1 |
| A3QJZ6 | PRAMEF22 | NLRKLFIsDGCRYLL | S242 | 0.017 | 88.72 | 481 | 0.013514 | 0.017 | 0.0135 | 0.8333 |
| Q8TET4 | GANC | TLSItGIsFCGADIG | S611 | 0.049 | 97.79 | 914 | 0.038951 | 0.049 | 0.039 | 1 |
| Q9BY41 | HDAC8 | VHsLIEAyALHKQMR | Y48 | 0.31 | 98.61 | 377 | 0.135993 | 0.31 | 0.136 | 1 |
| P51610 | HCFC1 | GKYSNDLyELQASRW | Y111 | 0.23 | 95.46 | 2035 | 0.100898 | 0.23 | 0.1009 | 1 |
| P51610 | HCFC1 | PRLGHSFsLVGNkCY | S143 | 0.08 | 97.09 | 2035 | 0.063593 | 0.08 | 0.0636 | 1 |
| P51610 | HCFC1 | ESHTAVVyTEKDNKK | Y208 | 0.099 | 95.14 | 2035 | 0.04343 | 0.156 | 0.0535 | 1 |
| P51610 | HCFC1 | RGPFSEIsAFkTCLP | S1881 | 0.042 | 95.84 | 2035 | 0.033386 | 0.042 | 0.0334 | 1 |
| P51610 | HCFC1 | TSGKIIEySVYLAIQ | Y1923 | 0.076 | 95.01 | 2035 | 0.03334 | 0.076 | 0.0333 | 1 |
| P35579 | MYH9 | sEEIVEMykGKKRHE | Y138 | 0.014 | 91.03 | 1960 | 0.006142 | 0.137 | 0.0601 | 1 |
| P35579 | MYH9 | HEMPPHIyAITDTAy | Y151 | 0 | 94.4 | 1960 | 0 | 0 | 0 | 1 |
| P35579 | MYH9 | MQDrEDQsILCtGES | S169 | 0 | 94.15 | 1960 | 0 | 0 | 0 | 1 |
| P35579 | MYH9 | EDQsILCtGESGAGk | T173 | 0.135 | 87.89 | 1960 | 0.092207 | 0.135 | 0.0922 | 1 |
| P35579 | MYH9 | NTKkVIQyLAyVAsS | Y190 | 0.009 | 92.53 | 1960 | 0.003948 | 0.009 | 0.0039 | 1 |
| P35579 | MYH9 | FQEFRQRyEILtPNs | Y721 | 0.054 | 89.82 | 1960 | 0.023689 | 0.054 | 0.0237 | 1 |
| P35579 | MYH9 | RQRyEILtPNsIPkG | T725 | 0.069 | 87.57 | 1960 | 0.047128 | 0.069 | 0.0471 | 1 |

| | | | | | | | | | | |
|---|---|---|---|---|---|---|---|---|---|---|
| P46531 | NOTCH1 | QTDRtGEtALHLAAR | T1930 | 0.011 | 88.39 | 2555 | 0.007513 | 0.011 | 0.0075 | 1 |
| P46531 | NOTCH1 | ILIRNRAtDLDARMH | T1986 | 0.168 | 84.92 | 2555 | 0.114746 | 0.168 | 0.1147 | 1 |
| P30613 | PKLR | GDLVIVVtGWRPGsG | T556 | 0.017 | 97.49 | 574 | 0.011611 | 0.017 | 0.0116 | 1 |
| Q8N6Q8 | METTL25 | LKyGLKVyGIDSSNT | Y201 | 0.215 | 97.77 | 603 | 0.094317 | 0.215 | 0.0943 | 1 |
| O75074 | LRP3 | SPAWPLNyPPGTNCS | Y66 | 0.033 | 89.51 | 770 | 0.014477 | 0.033 | 0.0145 | 1 |
| Q92882 | OSTF1 | TNWWKGTsKGRtGLI | S54 | 0.015 | 96.93 | 214 | 0.011924 | 0.015 | 0.0119 | 1 |
| Q92882 | OSTF1 | GRtGLIPsNYVAEQA | S63 | 0.166 | 95.38 | 214 | 0.131956 | 0.166 | 0.132 | 1 |
| O60343 | TBC1D4 | RFEINLIsPDTKSVV | S370 | 0.124 | 81.75 | 1298 | 0.098569 | 0.124 | 0.0986 | 1 |
| O75147 | OBSL1 | PSDAGtYsCAVGtAR | S1699 | 0.122 | 91.18 | 1896 | 0.09698 | 0.16 | 0.1366 | 1 |
| Q9NPI0 | TMEM138 | DPHFYQDsLWLRkEF | S150 | 0.176 | 71.85 | 162 | 0.139905 | 0.176 | 0.1399 | 1 |
| Q08170 | SRSF4 | RLIVENLssRCsWQD | S112 | 0.079 | 81.86 | 494 | 0.062798 | 0.079 | 0.0628 | 1 |
| P55039 | DRG2 | KPNSVVIsCGMKLNL | S269 | 0.003 | 98.18 | 364 | 0.002385 | 0.003 | 0.0024 | 1 |
| P55039 | DRG2 | LWEYLALtCIYtKKR | T290 | 0.104 | 91.82 | 364 | 0.071033 | 0.104 | 0.071 | 0.9167 |
| P29374 | ARID4A | GTDVSAkyRGAFCEA | Y20 | 0.263 | 84.09 | 1257 | 0.115374 | 0.263 | 0.1154 | 1 |
| P24752 | ACAT1 | EVVIVSAtRTPIGSF | T48 | 0.041 | 98.87 | 427 | 0.028004 | 0.082 | 0.056 | 0.9167 |
| P24752 | ACAT1 | AINSytRsKAAWEAG | S222 | 0 | 98.8 | 427 | 0 | 0 | 0 | 0.9091 |
| P24752 | ACAT1 | QkENGTVtAANAStL | T279 | 0 | 98.79 | 427 | 0 | 0 | 0 | 1 |
| P35398 | RORA | CKGFFRRsQQSNATY | S100 | 0.028 | 94.24 | 523 | 0.022258 | 0.028 | 0.0223 | 0.8 |
| Q92561 | PHYHIP | GVFFSCNtEFNtGQP | T188 | 0.21 | 97.34 | 330 | 0.143433 | 0.21 | 0.1434 | 1 |
| Q53H82 | LACTB2 | AEAkIQQyISHRNIR | Y213 | 0.088 | 98.45 | 288 | 0.038604 | 0.088 | 0.0386 | 0.8667 |
| O00148 | DDX39A | QRrILVAtNLFGrGM | T343 | 0.129 | 90.77 | 427 | 0.088109 | 0.129 | 0.0881 | 0.8889 |
| Q92622 | RUBCN | DFSKYyVsNFskDLL | S774 | 0 | 96.59 | 972 | 0 | 0 | 0 | 1 |
| Q92622 | RUBCN | TEDLHLYsLNDLtAT | S851 | 0.02 | 97.45 | 972 | 0.015898 | 0.02 | 0.0159 | 1 |
| Q92622 | RUBCN | LYsLNDLtATRkGEL | T856 | 0.197 | 97.63 | 972 | 0.134554 | 0.197 | 0.1346 | 1 |
| Q10567 | AP1B1 | MLSkDLDyyGtLLkK | Y276 | 0.269 | 89.82 | 949 | 0.118006 | 0.416 | 0.139 | 1 |
| Q10567 | AP1B1 | LSkDLDyyGtLLkKL | Y277 | 0.194 | 90.59 | 949 | 0.085105 | 0.325 | 0.1272 | 0.8182 |
| Q96H22 | CENPN | PLLGQALtIAsKHHQ | T180 | 0.108 | 94.22 | 339 | 0.073765 | 0.108 | 0.0738 | 1 |
| Q96H22 | CENPN | GQALtIAsKHHQIVk | S183 | 0.094 | 91.78 | 339 | 0.074722 | 0.094 | 0.0747 | 1 |
| Q9H6Z4 | RANBP3 | SLRLILNtKLWAQMQ | T457 | 0.005 | 97.5 | 567 | 0.003415 | 0.005 | 0.0034 | 1 |
| A4D1U4 | DENND11 | FKsMAsGsHKIQSDF | S115 | 0.031 | 94.53 | 455 | 0.024642 | 0.072 | 0.0572 | 0.8571 |
| A4D1U4 | DENND11 | MkSVGILsPSYTLLY | S158 | 0.097 | 93.11 | 455 | 0.077107 | 0.097 | 0.0771 | 1 |

| | | | | | | | | | | |
|---|---|---|---|---|---|---|---|---|---|---|
| O15084 | ANKRD28 | VSDRAGRtALHHAAF | T142 | 0.073 | 97.48 | 1053 | 0.04986 | 0.073 | 0.0499 | 1 |
| Q9UHD1 | CHORDC1 | WSCCRRktsDFNtFL | T199 | 0.207 | 95.93 | 332 | 0.141384 | 0.207 | 0.1414 | 1 |
| Q9NSY2 | STARD5 | PALAAQMsEAVAEKM | S10 | 0.043 | 98.34 | 213 | 0.034181 | 0.043 | 0.0342 | 1 |
| Q9P2S2 | NRXN2 | HTEAEDVsLRFMsQR | S757 | 0.086 | 97.2 | 1712 | 0.068363 | 0.086 | 0.0684 | 1 |
| Q9P2S2 | NRXN2 | DVsLRFMsQRAyGLM | S762 | 0.025 | 93.93 | 1712 | 0.019873 | 0.025 | 0.0199 | 1 |
| Q9P2S2 | NRXN2 | AyGLMMAtTsREsAD | T772 | 0 | 97.13 | 1712 | 0 | 0 | 0 | 1 |
| Q9P2S2 | NRXN2 | GLMMAtTsREsADtL | S774 | 0.056 | 92.38 | 1712 | 0.044515 | 0.056 | 0.0445 | 1 |
| Q9P2S2 | NRXN2 | MAtTsREsADtLRLE | S777 | 0.06 | 88.12 | 1712 | 0.047695 | 0.152 | 0.1208 | 1 |
| Q9P2S2 | NRXN2 | TsREsADtLRLELDG | T780 | 0.043 | 93.79 | 1712 | 0.02937 | 0.043 | 0.0294 | 1 |
| Q13043 | STK4 | LGEGsyGsVykAIHk | S43 | 0.168 | 90.43 | 487 | 0.133546 | 0.183 | 0.1455 | 1 |
| Q13043 | STK4 | SIMQQCDsPHVVKYY | S82 | 0.078 | 94.89 | 487 | 0.062003 | 0.078 | 0.062 | 1 |
| Q9UHK6 | AMACR | LAGHDINyLALSGVL | Y126 | 0.154 | 98.38 | 382 | 0.067558 | 0.154 | 0.0676 | 1 |
| Q96KV7 | WDR90 | SAQARAPsVMRLWDF | S433 | 0.028 | 82.61 | 1748 | 0.022258 | 0.028 | 0.0223 | 1 |
| P42685 | FRK | yFVALFDyQARtAED | Y53 | 0.111 | 93.21 | 505 | 0.048694 | 0.111 | 0.0487 | 0.9231 |
| P42685 | FRK | QGyIPsNyVAEDRSL | Y104 | 0.116 | 93.56 | 505 | 0.050888 | 0.116 | 0.0509 | 1 |
| P42685 | FRK | ysENKTGsFLIRESE | S139 | 0.06 | 98.03 | 505 | 0.047695 | 0.083 | 0.0792 | 1 |
| P42685 | FRK | LNEFVSHyTKTSDGL | Y193 | 0.037 | 97.02 | 505 | 0.016231 | 0.037 | 0.0162 | 1 |
| P42685 | FRK | LVGEHNIyKVADFGL | Y368 | 0.023 | 96.31 | 505 | 0.01009 | 0.023 | 0.0101 | 1 |
| Q9NQA5 | TRPV5 | QQkRILEsPLLRAsK | S47 | 0.057 | 92.72 | 729 | 0.04531 | 0.057 | 0.0453 | 1 |
| Q9NQA5 | TRPV5 | EsPLLRAsKENDLSV | S53 | 0 | 94.76 | 729 | 0 | 0 | 0 | 1 |
| Q92552 | MRPS27 | PVssLtIsRLIDNIs | S73 | 0.044 | 92.62 | 414 | 0.034976 | 0.041 | 0.0459 | 1 |
| Q9NZL9 | MAT2B | PNRRVLVtGAtGLLG | T34 | 0.01 | 98.73 | 334 | 0.00683 | 0.01 | 0.0068 | 1 |
| Q9NZL9 | MAT2B | VLRIPILyGEVEkLE | Y186 | 0.007 | 98.81 | 334 | 0.003071 | 0.007 | 0.0031 | 0.8889 |
| Q9NZL9 | MAT2B | FRIGIkEsLWPFLID | S318 | 0.041 | 98.13 | 334 | 0.032592 | 0.041 | 0.0326 | 1 |
| Q9Y3Z3 | SAMHD1 | LLVRIIDtPQFQRLR | T138 | 0.038 | 98.42 | 626 | 0.025955 | 0.038 | 0.026 | 1 |
| Q9Y3Z3 | SAMHD1 | NIIDtMItDAFLKAD | T387 | 0.034 | 98.75 | 626 | 0.023222 | 0.034 | 0.0232 | 1 |
| Q9Y3Z3 | SAMHD1 | VDVINMDyGMQEKNP | Y507 | 0.193 | 91.91 | 626 | 0.084666 | 0.193 | 0.0847 | 1 |
| Q9Y3Z3 | SAMHD1 | KNPIDHVsFYCKTAP | S519 | 0.064 | 98.18 | 626 | 0.050875 | 0.064 | 0.0509 | 1 |
| Q9Y3Z3 | SAMHD1 | DVIAPLItPQkkEWN | T592 | 0.032 | 93.62 | 626 | 0.021856 | 0.032 | 0.0219 | 1 |
| Q96BJ3 | AIDA | LPsEPGMtLLtIRIE | T167 | 0.076 | 97.04 | 306 | 0.051909 | 0.076 | 0.0519 | 0.8333 |
| Q96BJ3 | AIDA | GPIVIELyKKPTDFK | Y276 | 0.219 | 98.14 | 306 | 0.096072 | 0.219 | 0.0961 | 1 |

| | | | | | | | | | | |
|---|---|---|---|---|---|---|---|---|---|---|
| Q8NF50 | DOCK8 | IGDCAEPytVIKESD | Y365 | 0.03 | 88.91 | 2099 | 0.013161 | 0.03 | 0.0132 | 1 |
| Q9NXZ2 | DDX43 | KKNFYKEstATSAMS | S202 | 0.119 | 93.31 | 648 | 0.094595 | 0.119 | 0.0946 | 1 |
| Q9NXZ2 | DDX43 | RPGMLVLtPTRELAL | T323 | 0 | 95.67 | 648 | 0 | 0 | 0 | 1 |
| Q9NXZ2 | DDX43 | VEGECCKySYKGLRs | Y339 | 0.028 | 88.33 | 648 | 0.012283 | 0.028 | 0.0123 | 1 |
| Q9NXZ2 | DDX43 | GVDIIIAtPGRLNDL | T372 | 0.134 | 96.13 | 648 | 0.091524 | 0.134 | 0.0915 | 1 |
| Q53S08 | RAB6D | ITLVGNKtDLADKRQ | T128 | 0.156 | 96.37 | 254 | 0.10655 | 0.156 | 0.1066 | 0.8571 |
| P23246 | SFPQ | FkRLFAkyGEPGEVF | Y320 | 0.068 | 90.37 | 707 | 0.029831 | 0.115 | 0.0504 | 1 |
| P23246 | SFPQ | ALsVRNLsPyVsNEL | S379 | 0.132 | 95.7 | 707 | 0.104929 | 0.132 | 0.1049 | 1 |
| Q9Y234 | LIPT1 | NINLTFFttKKKYDR | T119 | 0.191 | 97.97 | 373 | 0.130456 | 0.191 | 0.1305 | 1 |
| Q9Y234 | LIPT1 | INLTFFttKKKYDRM | T120 | 0.185 | 97.52 | 373 | 0.126358 | 0.185 | 0.1264 | 1 |
| Q16763 | UBE2S | PPkGyFLtKIFHPNV | T81 | 0.075 | 97.96 | 222 | 0.051226 | 0.075 | 0.0512 | 0.8571 |
| Q16763 | UBE2S | AARARLLtEIHGGAG | T152 | 0.112 | 98.1 | 222 | 0.076498 | 0.149 | 0.1018 | 1 |
| P30511 | HLA-F | RGYHQHAyDGkDYIS | Y139 | 0.131 | 98.56 | 346 | 0.057468 | 0.131 | 0.0575 | 1 |
| O60218 | AKR1B10 | VGLGTWKsPLGkVKE | S23 | 0.079 | 96.36 | 316 | 0.062798 | 0.079 | 0.0628 | 1 |
| P86791 | CCZ1 | SsVLRQCysMyKLFN | Y142 | 0.214 | 98.63 | 482 | 0.093879 | 0.214 | 0.0939 | 1 |
| P86791 | CCZ1 | LRQCysMyKLFNGTF | Y145 | 0 | 98.56 | 482 | 0 | 0 | 0 | 1 |
| O43156 | TTI1 | LVtQAPIsARAGPVY | S967 | 0.009 | 92.89 | 1089 | 0.007154 | 0.009 | 0.0072 | 1 |
| Q14624 | ITIH4 | VVFVIDKsGsMsGRK | S282 | 0.013 | 97.19 | 930 | 0.010334 | 0.013 | 0.0103 | 1 |
| Q15782 | CHI3L2 | SEVMLyQtINSLkTK | T84 | 0.147 | 95.04 | 390 | 0.100403 | 0.229 | 0.1491 | 1 |
| Q15782 | CHI3L2 | LLLTAGVsAGRQMID | S185 | 0.065 | 98.85 | 390 | 0.051669 | 0.13 | 0.1033 | 0.875 |
| Q5XPI4 | RNF123 | SNFGTIRsttCVyKG | S126 | 0.008 | 96.75 | 1314 | 0.006359 | 0.008 | 0.0064 | 1 |
| Q5XPI4 | RNF123 | FGTIRsttCVyKGKW | T128 | 0 | 95.45 | 1314 | 0 | 0 | 0 | 1 |
| Q6UXG2 | ELAPOR1 | GNQGRKMsVCTDNVT | S712 | 0 | 91.91 | 1013 | 0 | 0 | 0 | 1 |
| Q6UXG2 | ELAPOR1 | TGYKAGVssQPVsLA | S756 | 0.126 | 82.12 | 1013 | 0.100159 | 0.126 | 0.1002 | 1 |
| Q7Z6G8 | ANKS1B | KELRGtEstQDACAK | S1080 | 0.091 | 89.34 | 1248 | 0.072337 | 0.168 | 0.1351 | 0.8571 |
| P49916 | LIG3 | FSKSEAHtADGISIR | T796 | 0.122 | 94.4 | 1009 | 0.083328 | 0.122 | 0.0833 | 1 |
| Q6P1M0 | SLC27A4 | QCLQERRtVPILFAs | T75 | 0.119 | 97.81 | 643 | 0.081279 | 0.188 | 0.1406 | 1 |
| Q6P1M0 | SLC27A4 | GFTDKLFyIyTSGTT | Y243 | 0.05 | 97.68 | 643 | 0.021934 | 0.05 | 0.0219 | 1 |
| Q6P1M0 | SLC27A4 | TDKLFyIyTSGTTGL | Y245 | 0.107 | 90.43 | 643 | 0.046939 | 0.214 | 0.0939 | 1 |
| Q6P1M0 | SLC27A4 | AIVVHSRyyRMAALV | Y263 | 0 | 97.62 | 643 | 0 | 0 | 0 | 1 |
| Q6P1M0 | SLC27A4 | IVVHSRyyRMAALVY | Y264 | 0.085 | 97.15 | 643 | 0.037288 | 0.085 | 0.0373 | 1 |

| | | | | | | | | | | |
|---|---|---|---|---|---|---|---|---|---|---|
| Q6P1M0 | SLC27A4 | PLRRFDGyLNQGANN | Y463 | 0.109 | 92.23 | 643 | 0.047817 | 0.109 | 0.0478 | 1 |
| Q8NGY1 | OR10Z1 | TPMyLFLsFLSFSET | S64 | 0.165 | 93.81 | 313 | 0.131161 | 0.165 | 0.1312 | 1 |
| Q0VAM2 | RASGEF1B | RLsyLVAtEICMPVK | T271 | 0.077 | 95.66 | 473 | 0.052592 | 0.077 | 0.0526 | 0.875 |
| Q9NQG7 | HPS4 | GLVRMNLytHCVKGL | Y544 | 0.029 | 92.3 | 708 | 0.012722 | 0.029 | 0.0127 | 1 |
| Q96PY5 | FMNL2 | TKLGLDEyLDkLKHT | Y352 | 0.302 | 94.81 | 1086 | 0.132483 | 0.302 | 0.1325 | 1 |
| Q8NDW8 | TTC21A | KKEAVIEtLNKALDL | T858 | 0.112 | 87.88 | 1320 | 0.076498 | 0.112 | 0.0765 | 0.8571 |
| P15531 | NME1 | GEtNPADskPGtIRG | S99 | 0.131 | 98.48 | 152 | 0.104134 | 0.131 | 0.1041 | 1 |
| P15531 | NME1 | GRNIIhGsDsVEsAE | S120 | 0.008 | 98.42 | 152 | 0.006359 | 0.008 | 0.0064 | 0.9 |
| P50402 | EMD | VGsTRRLyEKkIFEy | Y34 | 0.159 | 94.84 | 254 | 0.069751 | 0.159 | 0.0698 | 0.8462 |
| Q96H55 | MYO19 | NkLSREPsFIVVHYA | S526 | 0.019 | 92.3 | 970 | 0.015103 | 0.019 | 0.0151 | 1 |
| Q96H55 | MYO19 | HYAGPVRyHTAGLVE | Y538 | 0.025 | 94.39 | 970 | 0.010967 | 0.025 | 0.011 | 1 |
| Q9NVH6 | TMLHE | tWPDGHVtKYDLNWL | T126 | 0 | 98.36 | 421 | 0 | 0 | 0 | 1 |
| O43490 | PROM1 | MATAIKEtKEALENM | T266 | 0.008 | 96.02 | 865 | 0.005464 | 0.008 | 0.0055 | 1 |
| O43490 | PROM1 | ALENMNstLKSLHQQ | T276 | 0.129 | 94.84 | 865 | 0.088109 | 0.129 | 0.0881 | 1 |
| O43490 | PROM1 | SsVIIEEtKKYGRTI | T739 | 0.142 | 91.9 | 865 | 0.096988 | 0.142 | 0.097 | 1 |
| Q5TAA0 | TTC22 | AKIHIRAyLHDLKRA | Y346 | 0.013 | 96.63 | 569 | 0.005703 | 0.013 | 0.0057 | 1 |
| Q5TAA0 | TTC22 | LVRLLFEtMEREGEG | T550 | 0.056 | 69.69 | 569 | 0.038249 | 0.056 | 0.0382 | 1 |
| Q99538 | LGMN | MYRKMVFyIEACESG | Y185 | 0.047 | 98.94 | 433 | 0.020618 | 0.047 | 0.0206 | 0.8571 |
| Q99538 | LGMN | QYGNKtIstMKVMQF | S276 | 0.045 | 98.37 | 433 | 0.035771 | 0.045 | 0.0358 | 1 |
| O95460 | MATN4 | LVFVIDSsRsVRPFE | S42 | 0.033 | 91.67 | 622 | 0.026232 | 0.033 | 0.0262 | 0.875 |
| O95460 | MATN4 | RVGVIQYsSQVQSVF | S79 | 0.016 | 92.9 | 622 | 0.012719 | 0.03 | 0.0238 | 1 |
| Q8IXY8 | PPIL6 | CTGKAGFsQRGIRLH | S185 | 0.168 | 97.17 | 311 | 0.133546 | 0.168 | 0.1335 | 1 |
| Q8IXY8 | PPIL6 | QRGIRLHyKNSIFHR | Y193 | 0 | 98.78 | 311 | 0 | 0 | 0 | 1 |
| Q9GZU1 | MCOLN1 | LKTINLQsLINNEIP | S244 | 0.141 | 79.28 | 580 | 0.112083 | 0.177 | 0.1407 | 1 |
| Q9Y265 | RUVBL1 | LLAGPPGtGktALAL | T74 | 0.03 | 92.14 | 456 | 0.02049 | 0.049 | 0.0322 | 1 |
| Q9Y265 | RUVBL1 | APIVIFAsNRGNCVI | S331 | 0.011 | 96.44 | 456 | 0.008744 | 0.011 | 0.0087 | 1 |
| Q9Y265 | RUVBL1 | RGTEDITsPHGIPLD | S346 | 0.036 | 95.18 | 456 | 0.028617 | 0.036 | 0.0286 | 1 |
| O95025 | SEMA3D | FKRINVDyRLTQIVV | Y447 | 0.115 | 92.61 | 777 | 0.050449 | 0.199 | 0.0873 | 1 |
| Q9Y6E7 | SIRT4 | KRVKEADsLLVVGss | S255 | 0.03 | 98.68 | 314 | 0.023847 | 0.03 | 0.0238 | 1 |
| Q9Y6E7 | SIRT4 | DsLLVVGssLQVYSG | S261 | 0.059 | 98.33 | 314 | 0.0469 | 0.059 | 0.0469 | 1 |
| P07954 | FH | kVPNDkyyGAQTVRs | Y68 | 0.28 | 98.69 | 510 | 0.122832 | 0.28 | 0.1228 | 1 |

| | | | | | | | | | | |
|---|---|---|---|---|---|---|---|---|---|---|
| P07954 | FH | yGAQTVRsTMNFkIG | S75 | 0.068 | 98.25 | 510 | 0.054054 | 0.11 | 0.0967 | 1 |
| P07954 | FH | AAEVNQDyGLDPkIA | Y110 | 0.301 | 98.39 | 510 | 0.132044 | 0.301 | 0.132 | 1 |
| P07954 | FH | LGQEFSGyVQQVkYA | Y251 | 0.026 | 98.9 | 510 | 0.011406 | 0.026 | 0.0114 | 1 |
| Q13011 | ECH1 | LALAAEIsSksPVAV | S265 | 0.062 | 98 | 328 | 0.049285 | 0.062 | 0.0493 | 1 |
| Q13011 | ECH1 | AAEIsSksPVAVQST | S268 | 0.152 | 98.12 | 328 | 0.120827 | 0.152 | 0.1208 | 0.8571 |
| Q9NVH0 | EXD2 | LKVVQCHsQGGLRSL | S568 | 0.188 | 91.81 | 621 | 0.149444 | 0.188 | 0.1494 | 1 |
| Q5VYY2 | LIPM | EVATEDGyILSVNRI | Y71 | 0.081 | 98.81 | 423 | 0.035534 | 0.162 | 0.0711 | 0.9231 |
| Q5VYY2 | LIPM | LAQKIKMyFALAPIA | Y209 | 0 | 98.88 | 423 | 0 | 0 | 0 | 0.8947 |
| P37173 | TGFBR2 | AKGNLQEyLTRHVIS | Y336 | 0.26 | 98.63 | 567 | 0.114058 | 0.26 | 0.1141 | 1 |
| Q76FK4 | NOL8 | VFAyINIsVAEADLK | S59 | 0.188 | 92.75 | 1167 | 0.149444 | 0.188 | 0.1494 | 1 |
| Q9H3U1 | UNC45A | LMLAALRtLVGICsE | T218 | 0.016 | 94.87 | 944 | 0.010928 | 0.016 | 0.0109 | 1 |
| Q13217 | DNAJC3 | SKLkNDNtEAFYKIS | T222 | 0.124 | 91.97 | 504 | 0.084694 | 0.124 | 0.0847 | 1 |
| Q13217 | DNAJC3 | FYKISTLyYQLGDHE | Y232 | 0.256 | 98.42 | 504 | 0.112304 | 0.256 | 0.1123 | 0.8 |
| Q14993 | COL19A1 | SIQSQVIsLYMDCNL | S184 | 0.032 | 96.66 | 1142 | 0.025437 | 0.032 | 0.0254 | 0.8333 |
| O14920 | IKBKB | MVTGTIHtyPVTEDE | T324 | 0.184 | 97.24 | 756 | 0.125675 | 0.184 | 0.1257 | 0.8889 |
| O14920 | IKBKB | VTGTIHtyPVTEDEs | Y325 | 0.087 | 97.46 | 756 | 0.038166 | 0.087 | 0.0382 | 1 |
| O14920 | IKBKB | KNsMAsMsQQLKAKL | S476 | 0.177 | 96.49 | 756 | 0.1407 | 0.177 | 0.1407 | 1 |
| O14920 | IKBKB | DQRtEGDsQEMVRLL | S587 | 0.02 | 96.47 | 756 | 0.015898 | 0.02 | 0.0159 | 1 |
| O14920 | IKBKB | LLLQAIQsFEkKVRV | S600 | 0.126 | 93.35 | 756 | 0.100159 | 0.126 | 0.1002 | 1 |
| Q8IV77 | CNGA4 | AVSVHLstLsRVQIF | T342 | 0.163 | 79.31 | 575 | 0.111331 | 0.163 | 0.1113 | 1 |
| Q9HCK4 | ROBO2 | RKSDAGMyTCVGTNM | Y201 | 0.047 | 97.06 | 1378 | 0.020618 | 0.047 | 0.0206 | 1 |
| A8MYV0 | DCDC2C | ETFKYFPyWKsPRVP | Y216 | 0 | 86.22 | 364 | 0 | 0 | 0 | 1 |
| Q9H5Z1 | DHX35 | EGRVVGVtQPRRVAA | T106 | 0.004 | 97.42 | 703 | 0.002732 | 0.004 | 0.0027 | 0.9 |
| Q96CW1 | AP2M1 | SkFDsERsISFIPPD | S262 | 0.172 | 82.02 | 435 | 0.136725 | 0.172 | 0.1367 | 0.8 |
| Q96CW1 | AP2M1 | RMAGMKEsQISAEIE | S364 | 0.115 | 96.2 | 435 | 0.091415 | 0.115 | 0.0914 | 1 |
| Q96CW1 | AP2M1 | FEVPFAPsGLkVRYL | S397 | 0.072 | 94.4 | 435 | 0.057234 | 0.072 | 0.0572 | 1 |
| Q96CW1 | AP2M1 | YIGRSGIyEtRC___ | Y431 | 0.056 | 97.08 | 435 | 0.024566 | 0.056 | 0.0246 | 0.8462 |
| Q96CW1 | AP2M1 | GRSGIyEtRC_____ | T433 | 0.01 | 95.36 | 435 | 0.00683 | 0.01 | 0.0068 | 1 |
| O43896 | KIF1C | SRSHAVFtIVFTQRC | T222 | 0.153 | 97.23 | 1103 | 0.104501 | 0.153 | 0.1045 | 1 |
| O43896 | KIF1C | TAMIAALsPADINyE | S326 | 0.009 | 94.93 | 1103 | 0.007154 | 0.009 | 0.0072 | 1 |
| O43896 | KIF1C | LsPADINyEETLSTL | Y332 | 0.104 | 93.29 | 1103 | 0.045623 | 0.104 | 0.0456 | 1 |

| | | | | | | | | | | |
|---|---|---|---|---|---|---|---|---|---|---|
| Q8NFU5 | IPMK | RkDAVAAsIQkIEKI | S229 | 0.025 | 98.43 | 416 | 0.019873 | 0.025 | 0.0199 | 1 |
| Q9Y2U5 | MAP3K2 | LRDPQEKtLSIFMEY | T432 | 0.186 | 92.62 | 619 | 0.127041 | 0.186 | 0.127 | 0.8 |
| Q5VVW2 | GARNL3 | EKSPFFLsVtLsDQN | S124 | 0.002 | 97.74 | 1013 | 0.00159 | 0.002 | 0.0016 | 1 |
| Q5VVW2 | GARNL3 | SPFFLsVtLsDQNNQ | T126 | 0.027 | 94.86 | 1013 | 0.018441 | 0.066 | 0.0451 | 1 |
| Q5VVW2 | GARNL3 | FFLsVtLsDQNNQRV | S128 | 0.074 | 86.25 | 1013 | 0.058824 | 0.074 | 0.0588 | 1 |
| Q9UIG8 | SLCO3A1 | NVVYRyLyVSIAIAL | Y626 | 0.021 | 93.82 | 710 | 0.009212 | 0.021 | 0.0092 | 1 |
| Q8N3U4 | STAG2 | VIHALQCtHYVILWQ | T737 | 0.016 | 98.11 | 1231 | 0.010928 | 0.016 | 0.0109 | 0.8 |
| Q92903 | CDS1 | IkLSPKKtWEGFIGG | T276 | 0.2 | 93.47 | 461 | 0.136603 | 0.2 | 0.1366 | 1 |
| Q5JQC9 | AKAP4 | DDIDWLRsHRGVCKV | S19 | 0.116 | 87.51 | 854 | 0.09221 | 0.116 | 0.0922 | 1 |
| Q8IWJ2 | GCC2 | INTMLQLsPEEKGKL | S1649 | 0.177 | 81.54 | 1684 | 0.1407 | 0.177 | 0.1407 | 1 |
| Q9UGT4 | SUSD2 | VAAGDRVsIMLASGA | S560 | 0.137 | 97.2 | 822 | 0.108903 | 0.137 | 0.1089 | 0.8 |
| Q8NBJ9 | SIDT2 | SVIVKVTsNKAFPCS | S191 | 0.158 | 94.65 | 832 | 0.125597 | 0.158 | 0.1256 | 1 |
| Q8NBJ9 | SIDT2 | AFPCSVIsIQDVLCP | S201 | 0 | 96.49 | 832 | 0 | 0 | 0 | 1 |
| Q96DM1 | PGBD4 | WCDGKEVtMLsTFHN | T394 | 0.087 | 94.97 | 585 | 0.059422 | 0.087 | 0.0594 | 0.8 |
| Q96J01 | THOC3 | GNKDDVVtFIDAkTH | T166 | 0.078 | 98.62 | 351 | 0.053275 | 0.078 | 0.0533 | 1 |
| Q96J01 | THOC3 | DWPVRTLsFSHDGKM | S273 | 0.15 | 98.82 | 351 | 0.119237 | 0.15 | 0.1192 | 1 |
| Q96J01 | THOC3 | LWEVQCEsPTFTVAW | S308 | 0.092 | 92.19 | 351 | 0.073132 | 0.184 | 0.1463 | 0.8 |
| P11388 | TOP2A | EKGFQQIsFVNSIAT | S312 | 0.045 | 96.59 | 1531 | 0.035771 | 0.045 | 0.0358 | 0.9 |
| Q68CQ4 | UTP25 | SSDILIAsPLGLRTI | S451 | 0.01 | 98.26 | 756 | 0.007949 | 0.081 | 0.0644 | 1 |
| Q68CQ4 | UTP25 | GIRNLIFyELPTYPH | Y688 | 0.009 | 98.08 | 756 | 0.003948 | 0.009 | 0.0039 | 1 |
| P40938 | RFC3 | IEHQTIttPSkKKIE | T76 | 0.132 | 71.5 | 356 | 0.090158 | 0.132 | 0.0902 | 1 |
| P40938 | RFC3 | VAKFMALyKKFMEDG | Y343 | 0.313 | 88.02 | 356 | 0.137309 | 0.337 | 0.1478 | 1 |
| O14522 | PTPRT | THCIDFHyyFssRDR | Y107 | 0.215 | 93.2 | 1441 | 0.094317 | 0.215 | 0.0943 | 1 |
| O14522 | PTPRT | IDFHyyFssRDRSSP | S110 | 0.048 | 88.43 | 1441 | 0.038156 | 0.048 | 0.0382 | 1 |
| O14522 | PTPRT | DFHyyFssRDRSSPG | S111 | 0.157 | 85.06 | 1441 | 0.124802 | 0.157 | 0.1248 | 1 |
| O14522 | PTPRT | DIRARQLtLQWEPFG | T404 | 0.19 | 92.22 | 1441 | 0.129773 | 0.19 | 0.1298 | 1 |
| O14522 | PTPRT | EPFGyAVtRCHSYNL | T415 | 0.023 | 91.19 | 1441 | 0.015709 | 0.023 | 0.0157 | 1 |
| Q9BRA2 | TXNDC17 | IFAyFtGsKDAGGkS | S34 | 0.091 | 97.14 | 123 | 0.072337 | 0.091 | 0.0723 | 0.8571 |
| Q9BRA2 | TXNDC17 | LkVtAVPtLLkyGTP | T95 | 0.042 | 98.72 | 123 | 0.028687 | 0.042 | 0.0287 | 0.8 |
| Q9BX63 | BRIP1 | FDELLQVyYDAIkYK | Y742 | 0.001 | 92.79 | 1249 | 0.000439 | 0.001 | 0.0004 | 1 |
| P02008 | HBZ | ADtIGTEtLERLFLs | T29 | 0.063 | 98.75 | 142 | 0.04303 | 0.063 | 0.043 | 1 |

| | | | | | | | | | | |
|---|---|---|---|---|---|---|---|---|---|---|
| P02008 | HBZ | LFLsHPQtkTyFPHF | T40 | 0.042 | 98.76 | 142 | 0.028687 | 0.042 | 0.0287 | 0.8571 |
| P02008 | HBZ | HFDLHPGsAQLRAHG | S53 | 0.066 | 97.36 | 142 | 0.052464 | 0.066 | 0.0525 | 1 |
| Q9UGL1 | KDM5B | CSCPPYkyKLRyRyT | Y730 | 0.065 | 96.3 | 1544 | 0.028515 | 0.065 | 0.0285 | 1 |
| Q8WUX9 | CHMP7 | KEKAEEVyRLYQNSP | Y164 | 0.025 | 93.09 | 453 | 0.010967 | 0.025 | 0.011 | 1 |
| P14314 | PRKCSH | VKRPRGVsLTNHHFY | S24 | 0.051 | 94.55 | 528 | 0.040541 | 0.051 | 0.0405 | 1 |
| P14314 | PRKCSH | LCPFKLVsQkPkLGG | S434 | 0.183 | 97.87 | 528 | 0.145469 | 0.183 | 0.1455 | 1 |
| Q15019 | SEPTIN2 | DEDFkEQtRLLkASI | T228 | 0.084 | 84.86 | 361 | 0.057373 | 0.084 | 0.0574 | 1 |
| Q15019 | SEPTIN2 | MQDLQEVtQDLHyEN | T291 | 0.001 | 91.6 | 361 | 0.000683 | 0.001 | 0.0007 | 1 |
| Q15019 | SEPTIN2 | EVtQDLHyENFRSER | Y296 | 0.04 | 92.07 | 361 | 0.017547 | 0.04 | 0.0175 | 1 |
| Q9Y490 | TLN1 | LGDLIsAtKAAAGKV | T2098 | 0 | 85.43 | 2541 | 0 | 0 | 0 | 1 |
| Q969Y2 | GTPBP3 | PAPGSGAtIFALSSG | T35 | 0.006 | 97.45 | 492 | 0.004098 | 0.006 | 0.0041 | 1 |
| Q9NP92 | MRPS30 | KYMVYPQtFALNADR | T97 | 0.127 | 95.23 | 439 | 0.086743 | 0.127 | 0.0867 | 1 |
| P62910 | RPL32 | KKFIrHQsDrYVKIK | S25 | 0.152 | 95.95 | 135 | 0.120827 | 0.152 | 0.1208 | 1 |
| Q15717 | ELAVL1 | LGyGFVNyVTAkDAE | Y68 | 0.019 | 96.73 | 326 | 0.008335 | 0.019 | 0.0083 | 1 |
| Q86XL3 | ANKLE2 | skEPPLyyGVCPVyE | Y200 | 0.276 | 92.49 | 938 | 0.121077 | 0.276 | 0.1211 | 0.8 |
| Q86XL3 | ANKLE2 | AVKMIKGsRFkAFST | S235 | 0.173 | 83.01 | 938 | 0.13752 | 0.173 | 0.1375 | 1 |
| Q9Y5F1 | PCDHB12 | PEFEQAFyEVKILEN | Y247 | 0.181 | 94.41 | 795 | 0.079402 | 0.181 | 0.0794 | 1 |
| Q8IX18 | DHX40 | QHGMIGVtQPRkVAA | T104 | 0.032 | 97.54 | 779 | 0.021856 | 0.032 | 0.0219 | 0.8 |
| O00305 | CACNB4 | PVAFAVKtNVsyCGA | T99 | 0.008 | 97.96 | 520 | 0.005464 | 0.008 | 0.0055 | 1 |
| P05155 | SERPING1 | LVLLNAIyLsAKWKT | Y294 | 0.251 | 93.54 | 500 | 0.11011 | 0.251 | 0.1101 | 0.8333 |
| Q9NQQ7 | SLC35C2 | TALDVGLsNWsFLyV | S101 | 0.048 | 97.75 | 365 | 0.038156 | 0.048 | 0.0382 | 0.9091 |
| Q9NQQ7 | SLC35C2 | DVGLsNWsFLyVTVS | S104 | 0 | 97.13 | 365 | 0 | 0 | 0 | 1 |
| Q6ZMW3 | EML6 | HHVDKLVtVGIKHIK | T834 | 0 | 96.25 | 1958 | 0 | 0 | 0 | 1 |
| Q8N9N7 | LRRC57 | NLRtIDLsNNkIEsL | S46 | 0.071 | 98.68 | 239 | 0.056439 | 0.071 | 0.0564 | 1 |
| Q70E73 | RAPH1 | LFKNPQNyLLGKKET | Y366 | 0.303 | 95.48 | 1250 | 0.132922 | 0.303 | 0.1329 | 1 |
| Q70E73 | RAPH1 | NVYYGQDyRNKYKAP | Y456 | 0.004 | 98.44 | 1250 | 0.001755 | 0.004 | 0.0018 | 1 |
| Q70E73 | RAPH1 | AKYGKQLyMNyQEAL | Y510 | 0.265 | 98.11 | 1250 | 0.116252 | 0.265 | 0.1163 | 1 |
| Q70E73 | RAPH1 | GKQLyMNyQEALKRT | Y513 | 0.278 | 97.87 | 1250 | 0.121955 | 0.278 | 0.122 | 1 |
| Q9NXC5 | MIOS | TRtGLLAtLTRDSNI | T285 | 0.006 | 97.57 | 875 | 0.004098 | 0.006 | 0.0041 | 1 |
| P35228 | NOS2 | QLIRyAGyQMPDGSI | Y270 | 0.06 | 97.31 | 1153 | 0.026321 | 0.111 | 0.0487 | 1 |
| P35228 | NOS2 | INIAVLHsFQKQNVT | S425 | 0.009 | 98.71 | 1153 | 0.007154 | 0.009 | 0.0072 | 1 |

| | | | | | | | | | | |
|---|---|---|---|---|---|---|---|---|---|---|
| P35228 | NOS2 | AAESFMKyMQNEyRs | Y446 | 0.079 | 96.21 | 1153 | 0.034656 | 0.149 | 0.0622 | 1 |
| Q9UJX0 | OSGIN1 | DALLRPDtDFGGNMK | T97 | 0.179 | 85.73 | 477 | 0.122259 | 0.179 | 0.1223 | 1 |
| Q15149 | PLEC | NVQIALDyLRHRQVk | Y246 | 0.3 | 95.415 | 4684 | 0.131606 | 0.3275 | 0.1437 | 1 |
| Q9NPF8 | ADAP2 | PERRFVLtCPsEKEQ | T342 | 0 | 95.97 | 381 | 0 | 0 | 0 | 1 |
| P17643 | TYRP1 | FLGVGQEsFGEVDFS | S207 | 0.117 | 98.05 | 537 | 0.093005 | 0.117 | 0.093 | 1 |
| P17643 | TYRP1 | DDLMGSRsNFDSTLI | S270 | 0.115 | 98.54 | 537 | 0.091415 | 0.115 | 0.0914 | 1 |
| P16499 | PDE6A | KTKNILAsPIMNGKD | S180 | 0.044 | 95.23 | 860 | 0.034976 | 0.044 | 0.035 | 1 |
| P16499 | PDE6A | AFLNCDRySVGLLDM | Y275 | 0.018 | 95.04 | 860 | 0.007896 | 0.018 | 0.0079 | 1 |
| P55283 | CDH4 | DNDIPIRySITGVGA | Y205 | 0.162 | 94.81 | 916 | 0.071067 | 0.162 | 0.0711 | 1 |
| Q96GR2 | ACSBG1 | DKWEHISySQyyLLA | Y135 | 0.038 | 98.56 | 724 | 0.01667 | 0.038 | 0.0167 | 1 |
| Q6ZSI9 | CAPN12 | FHLNNQLtQTLTSRy | T661 | 0.071 | 80.01 | 719 | 0.048494 | 0.071 | 0.0485 | 0.9 |
| Q6ZSI9 | CAPN12 | tQTLTSRyRDSRLRV | Y668 | 0.143 | 87.89 | 719 | 0.062732 | 0.143 | 0.0627 | 0.9167 |
| Q96QT4 | TRPM7 | QRAAQKLtFAFNQMK | T1685 | 0.044 | 90.45 | 1865 | 0.030053 | 0.044 | 0.0301 | 1 |
| P51959 | CCNG1 | HDNGLRMtARLRDFE | T48 | 0.052 | 95.66 | 295 | 0.035517 | 0.052 | 0.0355 | 1 |
| O60674 | JAK2 | MEyLPyGsLRDyLQK | S936 | 0.149 | 95.49 | 1132 | 0.118442 | 0.149 | 0.1184 | 1 |
| O60674 | JAK2 | PyGsLRDyLQKHKER | Y940 | 0.16 | 95.01 | 1132 | 0.07019 | 0.16 | 0.0702 | 1 |
| O60674 | JAK2 | DHIKLLQyTSQICKG | Y956 | 0.003 | 97.74 | 1132 | 0.001316 | 0.003 | 0.0013 | 1 |
| P38570 | ITGAE | EEFKSARtARELNLI | T350 | 0.06 | 85.88 | 1179 | 0.040981 | 0.06 | 0.041 | 0.8 |
| P38570 | ITGAE | RELNLIAsDPDEtHA | S359 | 0.18 | 93.15 | 1179 | 0.143085 | 0.18 | 0.1431 | 0.8 |
| Q9ULT8 | HECTD1 | EPTYTIMyREMKDSD | Y1876 | 0.005 | 92.29 | 2610 | 0.002193 | 0.131 | 0.0397 | 1 |
| Q5T0N1 | CFAP70 | FKGDtPVtFIRAEFN | T31 | 0.001 | 91.65 | 1121 | 0.000683 | 0.001 | 0.0007 | 1 |
| Q7Z3V4 | UBE3B | PELQRLIsGDNAEID | S925 | 0.064 | 83.41 | 1068 | 0.050875 | 0.064 | 0.0509 | 0.8 |
| Q13045 | FLII | KLEHLSVsHNNLTTL | S64 | 0.101 | 91.87 | 1269 | 0.080286 | 0.171 | 0.147 | 1 |
| Q9Y5B9 | SUPT16H | NFDMVIVyKDysKkV | Y861 | 0.116 | 93.36 | 1047 | 0.050888 | 0.116 | 0.0509 | 1 |
| Q9BSU1 | PHAF1 | LEVVPERsLGNEQWE | S11 | 0.002 | 98.14 | 422 | 0.00159 | 0.002 | 0.0016 | 1 |
| Q9BSU1 | PHAF1 | GATVKRMyIYSGNSL | Y172 | 0.222 | 98.12 | 422 | 0.097388 | 0.222 | 0.0974 | 1 |
| Q04656 | ATP7A | NRSAIVKyNASsVtP | Y321 | 0.221 | 84.11 | 1500 | 0.09695 | 0.414 | 0.1499 | 0.8182 |
| O14544 | SOCS6 | SDDRyLLsLSFRSHG | S419 | 0.056 | 94.76 | 535 | 0.044515 | 0.056 | 0.0445 | 0.8889 |
| Q687X5 | STEAP4 | GTGDFGRsLGLKMLQ | S33 | 0.01 | 92.66 | 459 | 0.007949 | 0.01 | 0.0079 | 1 |
| P09417 | QDPR | EADFSSWtPLEFLVE | T203 | 0 | 98.38 | 244 | 0 | 0 | 0 | 1 |
| Q9C026 | TRIM9 | DDKAWAMyVDNNRsW | Y619 | 0.279 | 96.68 | 710 | 0.122393 | 0.279 | 0.1224 | 1 |

| | | | | | | | | | | |
|---|---|---|---|---|---|---|---|---|---|---|
| Q9C026 | TRIM9 | MyVDNNRsWFMHNNS | S625 | 0.007 | 96.47 | 710 | 0.005564 | 0.007 | 0.0056 | 1 |
| P52565 | ARHGDIA | GMkYIQHtyRKGVkI | T132 | 0.168 | 96.3 | 204 | 0.114746 | 0.168 | 0.1147 | 1 |
| P52565 | ARHGDIA | YGPRAEEyEFLtPVE | Y156 | 0.31 | 97.08 | 204 | 0.135993 | 0.31 | 0.136 | 0.875 |
| Q96HW7 | INTS4 | SQQFLQQsLERVYSL | S617 | 0.004 | 89.14 | 963 | 0.00318 | 0.004 | 0.0032 | 0.8333 |
| Q13432 | UNC119 | HNKADySySGTP___ | Y236 | 0.108 | 95.62 | 240 | 0.047378 | 0.108 | 0.0474 | 0.9 |
| Q9UJQ1 | LAMP5 | VFWVDRAyALKMLFV | Y115 | 0.01 | 97.74 | 280 | 0.004387 | 0.01 | 0.0044 | 1 |
| P50895 | BCAM | DGsPSPEytLFRLQD | Y301 | 0.106 | 94.03 | 628 | 0.046501 | 0.106 | 0.0465 | 1 |
| P50895 | BCAM | GsPSPEytLFRLQDE | T302 | 0.16 | 93.94 | 628 | 0.109282 | 0.16 | 0.1093 | 1 |
| P19087 | GNAT2 | LLLGAGEsGkSTIVk | S44 | 0.031 | 97.3 | 354 | 0.024642 | 0.031 | 0.0246 | 1 |
| Q9H4L4 | SENP3 | RtITYFDsQRTLNRR | S483 | 0.027 | 93.58 | 574 | 0.021463 | 0.027 | 0.0215 | 1 |
| Q9H4L4 | SENP3 | CPKHIAkyLQAEAVk | Y498 | 0.035 | 98.16 | 574 | 0.015354 | 0.035 | 0.0154 | 0.8571 |
| Q9Y2M5 | KLHL20 | LVDEAKNyLLLPQER | Y293 | 0.248 | 91.95 | 609 | 0.108794 | 0.248 | 0.1088 | 1 |
| Q9Y2M5 | KLHL20 | PLMQGPRtRPRKPIR | T308 | 0.09 | 85.65 | 609 | 0.061471 | 0.09 | 0.0615 | 1 |
| O95816 | BAG2 | NAKsHLMsLysACss | S137 | 0.163 | 94.27 | 211 | 0.129571 | 0.163 | 0.1296 | 1 |
| Q9BUI4 | POLR3C | DYAEVsNtFVRLADT | T147 | 0.065 | 96.54 | 534 | 0.044396 | 0.065 | 0.0444 | 1 |
| Q5JU69 | TOR2A | PHHGFsNsGIMEERL | S246 | 0 | 87.89 | 321 | 0 | 0 | 0 | 1 |
| Q5TAH2 | SLC9C2 | NyyFMFLyVLESTLK | Y651 | 0.242 | 86.85 | 1124 | 0.106162 | 0.242 | 0.1062 | 1 |
| P17948 | FLT1 | KYGNLSNyLKSKRDL | Y920 | 0.241 | 91.02 | 1338 | 0.105723 | 0.315 | 0.1382 | 1 |
| Q9GZM7 | TINAGL1 | IYSHTPVsLGRPERY | S387 | 0.036 | 94.75 | 467 | 0.028617 | 0.036 | 0.0286 | 1 |
| Q9HC21 | SLC25A19 | AQGEPkVyNTLRHAV | Y155 | 0.078 | 84.02 | 320 | 0.034217 | 0.288 | 0.1263 | 1 |
| Q16401 | PSMD5 | LVkALANskTIAEIF | S466 | 0.159 | 96.24 | 504 | 0.126391 | 0.159 | 0.1264 | 0.8889 |
| P46939 | UTRN | DQKELSVsVRRLAIL | S569 | 0.029 | 84.16 | 3433 | 0.023053 | 0.029 | 0.0231 | 1 |
| P20142 | PGC | DVFLRSyySVyDLGN | Y372 | 0.009 | 98.42 | 388 | 0.003948 | 0.009 | 0.0039 | 1 |
| Q9BSA4 | TTYH2 | LPVWREVtMELTKLS | T195 | 0.165 | 93.06 | 534 | 0.112697 | 0.165 | 0.1127 | 1 |
| Q8IVS8 | GLYCTK | CLHILNRyGLRAALP | Y274 | 0.268 | 95.97 | 523 | 0.117568 | 0.268 | 0.1176 | 1 |
| Q8IVS8 | GLYCTK | VKsMAQFyGLLAHVA | Y348 | 0.056 | 98.01 | 523 | 0.024566 | 0.056 | 0.0246 | 1 |
| Q8NEM0 | MCPH1 | PTRTLVMtSMPsEKQ | T653 | 0.123 | 97.39 | 835 | 0.084011 | 0.123 | 0.084 | 1 |
| Q8NEM0 | MCPH1 | HLCGGRVsQVPRQAs | S787 | 0.16 | 95.07 | 835 | 0.127186 | 0.16 | 0.1272 | 1 |
| P00738 | HP | RTEGDGVyTLNNEKQ | Y125 | 0.025 | 92.57 | 406 | 0.010967 | 0.025 | 0.011 | 1 |
| O60701 | UGDH | LVFIsVNtPTKTYGM | T91 | 0.046 | 91.29 | 494 | 0.031419 | 0.089 | 0.0605 | 1 |
| O60701 | UGDH | VQALCAVyEHWVPRE | Y199 | 0.112 | 98.5 | 494 | 0.049133 | 0.112 | 0.0491 | 1 |

| | | | | | | | | | | |
|---|---|---|---|---|---|---|---|---|---|---|
| O60701 | UGDH | ESSSIyIskYLMDEG | S354 | 0.003 | 98.45 | 494 | 0.002385 | 0.003 | 0.0024 | 1 |
| O60844 | ZG16 | AIQARSssYSGEyGG | S25 | 0.17 | 98.24 | 167 | 0.135135 | 0.186 | 0.1479 | 1 |
| O14638 | ENPP3 | RAMYPTKtFPNHYTI | T205 | 0.143 | 98.68 | 875 | 0.097671 | 0.143 | 0.0977 | 1 |
| O14638 | ENPP3 | KMEYMTDyFPRINFF | Y389 | 0.147 | 98.3 | 875 | 0.064487 | 0.147 | 0.0645 | 1 |
| O14638 | ENPP3 | NItHGFLyPPASNRT | Y694 | 0.021 | 98.71 | 875 | 0.009212 | 0.021 | 0.0092 | 1 |
| O14638 | ENPP3 | VRDVELLtGLDFyQD | T846 | 0.09 | 98.86 | 875 | 0.061471 | 0.09 | 0.0615 | 1 |
| O14638 | ENPP3 | LLtGLDFyQDKVQPV | Y851 | 0.109 | 98.7 | 875 | 0.047817 | 0.109 | 0.0478 | 1 |
| Q66K79 | CPZ | ELLVIEFssRPGQHE | S228 | 0 | 98.4 | 652 | 0 | 0 | 0 | 1 |
| Q96CT7 | CCDC124 | LKKEWLRsPDNPMNQ | S207 | 0.084 | 82.74 | 223 | 0.066773 | 0.084 | 0.0668 | 1 |
| Q9Y231 | FUT9 | NLESPTHtPQKSGIE | T142 | 0.024 | 96.1 | 359 | 0.016392 | 0.024 | 0.0164 | 1 |
| Q13508 | ART3 | IALMAyIsEAQEQTP | S103 | 0.157 | 95.16 | 389 | 0.124802 | 0.157 | 0.1248 | 1 |
| Q13508 | ART3 | QSINKtCsHyECAFL | S252 | 0.047 | 97.04 | 389 | 0.037361 | 0.047 | 0.0374 | 0.875 |
| Q13508 | ART3 | INKtCsHyECAFLGG | Y254 | 0.062 | 97.54 | 389 | 0.027199 | 0.062 | 0.0272 | 1 |
| Q6P158 | DHX57 | ESVKSSAtRLLYCTT | T639 | 0.173 | 96.14 | 1386 | 0.118161 | 0.173 | 0.1182 | 1 |
| P35998 | PSMC2 | ADSEDPkyIINVkQF | Y111 | 0.018 | 87.39 | 433 | 0.007896 | 0.018 | 0.0079 | 1 |
| P46063 | RECQL | FRPLQLEtINVTMAG | T99 | 0 | 97.72 | 649 | 0 | 0 | 0 | 1 |
| O00238 | BMPR1B | YIMADMYsFGLILWE | S402 | 0.096 | 98.79 | 502 | 0.076312 | 0.096 | 0.0763 | 0.9091 |
| P58107 | EPPK1 | GLLDTQTsQVLTAVD | S1390 | 0.011 | 83.72 | 5088 | 0.008744 | 0.011 | 0.0087 | 1 |
| Q14194 | CRMP1 | RAALVGGttMIIDHV | T101 | 0.003 | 98.89 | 572 | 0.002049 | 0.003 | 0.002 | 0.8182 |
| Q14194 | CRMP1 | AALVGGttMIIDHVV | T102 | 0.022 | 98.88 | 572 | 0.015026 | 0.022 | 0.015 | 1 |
| Q14194 | CRMP1 | GSPLVVIsQGKIVFE | S448 | 0.046 | 98.14 | 572 | 0.036566 | 0.046 | 0.0366 | 1 |
| Q9NVU7 | SDAD1 | MMLMNLIsRLVGIHE | S322 | 0 | 92.77 | 687 | 0 | 0 | 0 | 1 |
| Q6U841 | SLC4A10 | LAEVPIPtRFLFILL | T368 | 0.018 | 91.96 | 1118 | 0.012294 | 0.018 | 0.0123 | 1 |
| O14786 | NRP1 | NyPENGWtPGEDsyR | T316 | 0.197 | 98.61 | 923 | 0.134554 | 0.197 | 0.1346 | 1 |
| O14786 | NRP1 | GWtPGEDsyREWIQV | S321 | 0.186 | 97.55 | 923 | 0.147854 | 0.186 | 0.1479 | 1 |
| O14786 | NRP1 | KETKKKyyVKTyKID | Y354 | 0.312 | 98.59 | 923 | 0.13687 | 0.312 | 0.1369 | 1 |
| O14786 | NRP1 | KKyyVKTyKIDVSsN | Y358 | 0.011 | 98.53 | 923 | 0.004826 | 0.011 | 0.0048 | 1 |
| O14786 | NRP1 | KITDYPCsGMLGMVS | S432 | 0.095 | 88.9 | 923 | 0.075517 | 0.095 | 0.0755 | 1 |
| Q7Z6J0 | SH3RF1 | CAKALyNyEGkEPGD | Y145 | 0.173 | 95.02 | 888 | 0.075893 | 0.173 | 0.0759 | 1 |
| Q8WVF5 | KCTD4 | TLtKYPDtFLEGIVN | T57 | 0.079 | 96.61 | 259 | 0.053958 | 0.079 | 0.054 | 0.8 |
| P98196 | ATP11A | GRQAARNsDYAIPKF | S854 | 0.006 | 85.9 | 1134 | 0.004769 | 0.185 | 0.1471 | 1 |

| | | | | | | | | | | |
|---|---|---|---|---|---|---|---|---|---|---|
| P35913 | PDE6B | LLQADRCsLFMYRQR | S93 | 0.054 | 97.18 | 854 | 0.042925 | 0.054 | 0.0429 | 1 |
| P35913 | PDE6B | EIQLILPtRARLGKE | T469 | 0 | 97.22 | 854 | 0 | 0 | 0 | 1 |
| Q5VVH5 | IRAK1BP1 | AASKVFItFEVKGKE | T247 | 0.093 | 96.22 | 260 | 0.06352 | 0.093 | 0.0635 | 1 |
| Q8IZS8 | CACNA2D3 | DCRNRKWyIQAATsP | Y247 | 0.032 | 94.04 | 1091 | 0.014038 | 0.032 | 0.014 | 0.8667 |
| Q99952 | PTPN18 | AVQTEEQyRFLYHTV | Y281 | 0 | 98.02 | 460 | 0 | 0 | 0 | 1 |
| Q8IUH3 | RBM45 | KNVGyAkyADRIsAN | Y434 | 0 | 92.39 | 476 | 0 | 0 | 0 | 0.9412 |
| Q8IUH3 | RBM45 | AkyADRIsANDAIAT | S439 | 0.075 | 91.96 | 476 | 0.059619 | 0.075 | 0.0596 | 1 |
| Q7Z553 | MDGA2 | NTKYTPNtGPNADRS | T784 | 0.092 | 82.32 | 956 | 0.062837 | 0.092 | 0.0628 | 0.9 |
| Q7Z553 | MDGA2 | SGSKEGFyMyIETSR | Y798 | 0.006 | 93.08 | 956 | 0.002632 | 0.006 | 0.0026 | 0.9375 |
| Q5FVE4 | ACSBG2 | RHFVAQKyKKQIDHM | Y657 | 0.228 | 95.98 | 666 | 0.10002 | 0.228 | 0.1 | 1 |
| Q9BYX2 | TBC1D2 | EGIFEIktPSRVITL | T115 | 0.191 | 85.52 | 928 | 0.130456 | 0.191 | 0.1305 | 0.8889 |
| P06127 | CD5 | GSLGGTIsYEAQDKT | S186 | 0.139 | 90.95 | 495 | 0.110493 | 0.139 | 0.1105 | 1 |
| Q9H9T3 | ELP3 | RPDGLkLyPTLVIRG | Y318 | 0.187 | 96.58 | 547 | 0.082034 | 0.187 | 0.082 | 1 |
| Q9H9T3 | ELP3 | LALVPPWtRVYRVQR | T360 | 0.007 | 98.44 | 547 | 0.004781 | 0.007 | 0.0048 | 1 |
| Q9H9T3 | ELP3 | VELVrRDyVANGGWE | Y427 | 0.137 | 98.17 | 547 | 0.0601 | 0.137 | 0.0601 | 1 |
| Q92835 | INPP5D | SRTGkDGsFLVRAsE | S27 | 0 | 95.74 | 1189 | 0 | 0 | 0 | 1 |
| Q9Y2V2 | CARHSP1 | APGTKHEtWsGHVIs | T139 | 0.038 | 95.91 | 147 | 0.025955 | 0.038 | 0.026 | 1 |
| Q8WU66 | TSPEAR | LLLGLRLsPAQLHFL | S124 | 0.131 | 89.92 | 669 | 0.104134 | 0.131 | 0.1041 | 1 |
| P37235 | HPCAL1 | LEIVQAIyKMVSSVM | Y129 | 0.261 | 88.38 | 193 | 0.114497 | 0.261 | 0.1145 | 0.9 |
| Q8NHC7 | OR14C36 | QFFCDIPsLLKLSCS | S181 | 0.176 | 87.83 | 312 | 0.139905 | 0.176 | 0.1399 | 1 |
| Q9UL33 | TRAPPC2L | PTEDYKVyGYVTNSk | Y70 | 0.221 | 98.29 | 140 | 0.09695 | 0.221 | 0.0969 | 0.8333 |
| Q96GW9 | MARS2 | TAATRFStGtDEHGL | T90 | 0.017 | 98.17 | 593 | 0.011611 | 0.017 | 0.0116 | 1 |
| Q96GW9 | MARS2 | ATRFStGtDEHGLKI | T92 | 0 | 98.48 | 593 | 0 | 0 | 0 | 1 |
| Q6P4A8 | PLBD1 | SLDKGTLyIVEQIPT | Y375 | 0.324 | 98.91 | 553 | 0.142134 | 0.324 | 0.1421 | 0.9091 |
| Q6P4A8 | PLBD1 | SMKyIMRyNNYKKDP | Y456 | 0.267 | 98.81 | 553 | 0.117129 | 0.267 | 0.1171 | 1 |
| O75509 | TNFRSF21 | QCARGTFsDVPssVM | S177 | 0.037 | 98.5 | 655 | 0.029412 | 0.037 | 0.0294 | 1 |
| O75509 | TNFRSF21 | GTFsDVPssVMKCKA | S181 | 0.037 | 97.73 | 655 | 0.029412 | 0.157 | 0.1248 | 1 |
| Q9Y5X1 | SNX9 | AkPGtEQyLLAkQLA | Y219 | 0.02 | 92.31 | 595 | 0.008774 | 0.02 | 0.0088 | 1 |
| Q9Y5X1 | SNX9 | GLkSYIEyQLtPTNT | Y272 | 0.03 | 97.03 | 595 | 0.013161 | 0.03 | 0.0132 | 1 |
| Q14353 | GAMT | PPADCRyyAFPQMIT | Y223 | 0.042 | 97.13 | 236 | 0.018425 | 0.042 | 0.0184 | 0.8824 |
| Q6ZMW3 | EML6 | KGKILVGtKDGEIIE | T1663 | 0 | 91.96 | 1958 | 0 | 0 | 0 | 1 |

| | | | | | | | | | | |
|---|---|---|---|---|---|---|---|---|---|---|
| Q6ZMW3 | EML6 | FVIQMDFsADGKyIQ | S1824 | 0.008 | 94.64 | 1958 | 0.006359 | 0.008 | 0.0064 | 1 |
| Q9NRW7 | VPS45 | SAENDEFyANNMYLN | Y272 | 0.035 | 94.8 | 570 | 0.015354 | 0.035 | 0.0154 | 1 |
| Q9NRW7 | VPS45 | GRLkENLyPYLGPST | Y487 | 0.153 | 97.73 | 570 | 0.067119 | 0.153 | 0.0671 | 1 |
| Q9NXE4 | SMPD4 | ILPDsPLyHNKVQFT | Y172 | 0.031 | 84.42 | 866 | 0.013599 | 0.244 | 0.107 | 0.9167 |
| P39656 | DDOST | VCASGPRtLVLLDNL | T47 | 0.002 | 96.58 | 456 | 0.001366 | 0.002 | 0.0014 | 1 |
| P42356 | PI4KA | SCFLSNRsRtyDMIQ | S2087 | 0.086 | 81.75 | 2102 | 0.068363 | 0.086 | 0.0684 | 0.8 |
| P54868 | HMGCS2 | GGLKLEDtYTNKDLD | T346 | 0.088 | 86.4 | 508 | 0.060105 | 0.088 | 0.0601 | 1 |
| P54868 | HMGCS2 | IGAFSYGsGLAAsFF | S414 | 0.177 | 98.67 | 508 | 0.1407 | 0.177 | 0.1407 | 1 |
| P54868 | HMGCS2 | YGsGLAAsFFsFRVs | S419 | 0 | 98.94 | 508 | 0 | 0 | 0 | 0.8 |
| P54868 | HMGCS2 | sQDAAPGsPLDKLVs | S433 | 0.064 | 98.46 | 508 | 0.050875 | 0.064 | 0.0509 | 1 |
| P35659 | DEK | kkNVGQFsGFPFEkG | S131 | 0.069 | 80.25 | 375 | 0.054849 | 0.093 | 0.0739 | 0.875 |
| Q9NRG7 | SDR39U1 | KPGPGRItWDELAAs | T40 | 0.148 | 98.44 | 293 | 0.101086 | 0.148 | 0.1011 | 1 |
| Q9BXR0 | QTRT1 | DGNETLLsPEKSVQI | S139 | 0.166 | 96.15 | 403 | 0.131956 | 0.166 | 0.132 | 1 |
| O14983 | ATP2A1 | VATTGVGtEIGKIRD | T230 | 0.022 | 95.13 | 1001 | 0.015026 | 0.022 | 0.015 | 1 |
| Q15418 | RPS6KA1 | EIEILLRyGQHPNII | Y466 | 0.214 | 87.3 | 735 | 0.093879 | 0.214 | 0.0939 | 1 |
| Q15418 | RPS6KA1 | VLHTIGKtVEYLHSQ | T522 | 0.01 | 96.65 | 735 | 0.00683 | 0.01 | 0.0068 | 1 |
| P06744 | GPI | FQkLQQWyREHRsEL | Y17 | 0.256 | 98.5 | 558 | 0.112304 | 0.256 | 0.1123 | 1 |
| P06744 | GPI | HGHILVDysKNLVtE | Y55 | 0.118 | 98.89 | 558 | 0.051765 | 0.236 | 0.1035 | 1 |
| P06744 | GPI | GHILVDysKNLVtED | S56 | 0.013 | 98.87 | 558 | 0.010334 | 0.013 | 0.0103 | 1 |
| P06744 | GPI | DLGPLMVtEALkPyS | T168 | 0 | 98.92 | 558 | 0 | 0 | 0 | 1 |
| P06744 | GPI | DGTHIAktLAQLNPE | T195 | 0.035 | 98.82 | 558 | 0.023905 | 0.07 | 0.0478 | 1 |
| P06744 | GPI | SSLFIIAskTFttQE | S210 | 0.019 | 96.47 | 558 | 0.015103 | 0.038 | 0.0302 | 1 |
| P06744 | GPI | AkHFVALstNttkVK | S247 | 0 | 95.38 | 558 | 0 | 0 | 0 | 1 |
| P06744 | GPI | FQQGDMEsNGkYITk | S359 | 0 | 98.85 | 558 | 0 | 0 | 0 | 1 |
| Q96N96 | SPATA13 | GLDILDRssELIHSG | S453 | 0.037 | 93.26 | 652 | 0.029412 | 0.037 | 0.0294 | 1 |
| Q96N96 | SPATA13 | LDILDRssELIHSGE | S454 | 0.112 | 90.71 | 652 | 0.08903 | 0.112 | 0.089 | 1 |
| Q96N96 | SPATA13 | LRRDMLyyKGRLDMD | Y498 | 0.249 | 97.56 | 652 | 0.109233 | 0.249 | 0.1092 | 1 |
| Q9NPA0 | EMC7 | SYVVEVVsPAYRFDP | S100 | 0.083 | 92.82 | 242 | 0.065978 | 0.083 | 0.066 | 1 |
| Q5W5X9 | TTC23 | LtKAERLsKELLQCG | S167 | 0 | 94.8 | 447 | 0 | 0 | 0 | 1 |
| Q86VX9 | MON1A | PDLRHFLyKSkSSGL | Y547 | 0.231 | 97.91 | 652 | 0.101336 | 0.231 | 0.1013 | 0.9167 |
| Q9H115 | NAPB | SKHDSAtsFVDAGNA | S84 | 0.107 | 98 | 298 | 0.085056 | 0.058 | 0.1017 | 1 |

| | | | | | | | | | | |
|---|---|---|---|---|---|---|---|---|---|---|
| Q9H115 | NAPB | YEQSADyykGEESNS | Y152 | 0.156 | 97.4 | 298 | 0.068435 | 0.156 | 0.0684 | 1 |
| Q14184 | DOC2B | ANGySDPyVKTyLRP | Y305 | 0.011 | 98.71 | 412 | 0.004826 | 0.011 | 0.0048 | 0.8333 |
| Q8NDZ2 | SIMC1 | LEDDFQQtLRRQRQH | T584 | 0.18 | 86.56 | 872 | 0.122942 | 0.18 | 0.1229 | 1 |
| P17812 | CTPS1 | GKGIIASsVGTILKS | S22 | 0 | 96.3 | 591 | 0 | 0 | 0 | 1 |
| P17812 | CTPS1 | TSVkEKIsMFCHVEP | S231 | 0.145 | 97.22 | 591 | 0.115263 | 0.145 | 0.1153 | 1 |
| P17812 | CTPS1 | EEQGVVDyFLRRLDL | Y265 | 0.314 | 95.61 | 591 | 0.137747 | 0.314 | 0.1377 | 1 |
| P17812 | CTPS1 | SADLEPItsQEEPVR | T346 | 0.206 | 93.76 | 591 | 0.140701 | 0.206 | 0.1407 | 1 |
| P17812 | CTPS1 | STEFDPTtSHPVVVD | T428 | 0.174 | 95.91 | 591 | 0.118844 | 0.174 | 0.1188 | 1 |
| P17812 | CTPS1 | MRLGKRRtLFQTkNs | T455 | 0.012 | 97.9 | 591 | 0.008196 | 0.012 | 0.0082 | 1 |
| P17812 | CTPS1 | tLFQTkNsVMRkLYG | S462 | 0.019 | 95.31 | 591 | 0.015103 | 0.019 | 0.0151 | 1 |
| P56705 | WNT4 | RNLEVMDsVRRGAQL | S66 | 0.001 | 95.62 | 351 | 0.000795 | 0.001 | 0.0008 | 1 |
| P56705 | WNT4 | GKVVTQGtREAAFVy | T106 | 0.024 | 97.63 | 351 | 0.016392 | 0.024 | 0.0164 | 0.8889 |
| O14966 | RAB29 | GFtGWTEtsVkENkN | T154 | 0.001 | 98.69 | 203 | 0.000683 | 0.001 | 0.0007 | 0.8182 |
| O14966 | RAB29 | FtGWTEtsVkENkNI | S155 | 0.013 | 98.41 | 203 | 0.010334 | 0.013 | 0.0103 | 0.8571 |
| P21333 | FLNA | tADFkVytkGAGsGE | T507 | 0.161 | 84.53 | 2647 | 0.109965 | 0.161 | 0.11 | 1 |
| P21333 | FLNA | WPQEAGEyAVHVLCN | Y643 | 0 | 88.81 | 2647 | 0 | 0 | 0 | 1 |
| P21333 | FLNA | KDNGNGtysCsyVPR | Y731 | 0.337 | 92.21 | 2647 | 0.147837 | 0.337 | 0.1478 | 1 |
| P21333 | FLNA | NGtysCsyVPRKPVK | Y735 | 0.016 | 92.55 | 2647 | 0.007019 | 0.016 | 0.007 | 1 |
| Q6NUI2 | GPAT2 | GLPLVLLstHKTLLD | S203 | 0.006 | 94.28 | 795 | 0.004769 | 0.006 | 0.0048 | 1 |
| Q6NUI2 | GPAT2 | LPLVLLstHKTLLDG | T204 | 0.037 | 90.81 | 795 | 0.025272 | 0.037 | 0.0253 | 0.9 |
| O75165 | DNAJC13 | QPyKyAGyPMLIRTI | Y1395 | 0.021 | 89.34 | 2243 | 0.009212 | 0.021 | 0.0092 | 1 |
| O75165 | DNAJC13 | ENPTIRKsLAGMLTP | S1608 | 0.011 | 94.4 | 2243 | 0.008744 | 0.011 | 0.0087 | 1 |
| P07947 | YES1 | NGyIPsNyVAPADSI | Y146 | 0.133 | 94.63 | 543 | 0.058345 | 0.133 | 0.0583 | 1 |
| P07947 | YES1 | SIQAEEWyFGKMGRK | Y159 | 0.282 | 95.09 | 543 | 0.123709 | 0.282 | 0.1237 | 0.8 |
| P07947 | YES1 | ETtkGAysLsIRDWD | S195 | 0 | 97.19 | 543 | 0 | 0.117 | 0.093 | 1 |
| P07947 | YES1 | VVSEEPIyIVTEFMS | Y345 | 0.295 | 97.06 | 543 | 0.129412 | 0.295 | 0.1294 | 1 |
| Q6XPS3 | TPTE2 | FVEGRQQyFSDLFNI | Y135 | 0.015 | 86.95 | 522 | 0.00658 | 0.04 | 0.0175 | 1 |
| Q75N90 | FBN3 | THVRstCyGAIEKGS | Y621 | 0.223 | 86.905 | 2809 | 0.097827 | 0.223 | 0.0978 | 1 |
| Q13402 | MYO7A | AFIAVLKyMGDyPsK | Y1780 | 0.14 | 91.77 | 2215 | 0.061416 | 0.14 | 0.0614 | 1 |
| P33993 | MCM7 | IREVRADsVGkLVTV | S156 | 0.037 | 85.39 | 719 | 0.029412 | 0.037 | 0.0294 | 1 |
| P33993 | MCM7 | QGLLSETyLEAHRIV | Y297 | 0.309 | 91.06 | 719 | 0.135554 | 0.309 | 0.1356 | 0.875 |

| | | | | | | | | | | |
|---|---|---|---|---|---|---|---|---|---|---|
| Q9UPU5 | USP24 | FEIVKLVtGSLIACH | T673 | 0.009 | 87.8 | 2620 | 0.006147 | 0.009 | 0.0061 | 0.8 |
| P16144 | ITGB4 | RVDKDCAyCTDEMFR | Y50 | 0.05 | 92.35 | 1822 | 0.021934 | 0.05 | 0.0219 | 1 |
| P16144 | ITGB4 | ARVLSQLtsDyTIGF | T164 | 0.053 | 83.47 | 1822 | 0.0362 | 0.105 | 0.0717 | 1 |
| P16144 | ITGB4 | FQKTRTGsFHIRRGE | S397 | 0.027 | 78.36 | 1822 | 0.021463 | 0.027 | 0.0215 | 1 |
| P16144 | ITGB4 | RRGEVGIyQVQLRAL | Y408 | 0.021 | 91.92 | 1822 | 0.009212 | 0.021 | 0.0092 | 1 |
| O94827 | PLEKHG5 | EEEGCMEyMRGLLRD | Y448 | 0.307 | 93.8 | 1006 | 0.134676 | 0.307 | 0.1347 | 1 |
| Q96SI9 | STRBP | TLKVILTsPLIRDEL | S166 | 0 | 94.1 | 672 | 0 | 0.057 | 0.0453 | 1 |
| Q7Z7C8 | TAF8 | KASVETLtEMLQSYI | T62 | 0 | 96.62 | 310 | 0 | 0 | 0 | 0.875 |
| Q9HA92 | RSAD1 | TREARIQtLEPDNWM | T323 | 0.131 | 98.49 | 442 | 0.089475 | 0.131 | 0.0895 | 1 |
| Q86WV1 | SKAP1 | SRGLFyyyANEKSKQ | Y143 | 0 | 94.95 | 359 | 0 | 0 | 0 | 1 |
| Q9Y6W6 | DUSP10 | IIVyDENtNEPSRVM | T240 | 0.178 | 93.86 | 482 | 0.121576 | 0.178 | 0.1216 | 1 |
| Q9NYQ6 | CELSR1 | EGEEERVsyyMEGLF | S281 | 0.126 | 87.795 | 3014 | 0.100159 | 0.143 | 0.1137 | 1 |
| Q9NYQ6 | CELSR1 | GEEERVsyyMEGLFD | Y282 | 0.088 | 89.96 | 3014 | 0.038604 | 0.088 | 0.0386 | 1 |
| P10398 | ARAF | LIDVARQtAQGMDyL | T413 | 0.007 | 98.38 | 606 | 0.004781 | 0.208 | 0.1421 | 1 |
| O14802 | POLR3A | EALVLMGtkANLVTP | T525 | 0.015 | 94.4 | 1390 | 0.010245 | 0.015 | 0.0102 | 1 |
| O14802 | POLR3A | NEAADAMsRLARLAP | S681 | 0.039 | 93.26 | 1390 | 0.031002 | 0.039 | 0.031 | 1 |
| O14802 | POLR3A | KNELILttESIMKkS | T949 | 0.034 | 91.17 | 1390 | 0.023222 | 0.053 | 0.0362 | 1 |
| O14802 | POLR3A | kKFIkGVsEKIKKTR | S977 | 0 | 91.99 | 1390 | 0 | 0 | 0 | 1 |
| P46940 | IQGAP1 | DEGDAQktLQALQIP | T557 | 0.005 | 93.11 | 1657 | 0.003415 | 0.005 | 0.0034 | 1 |
| P46940 | IQGAP1 | LAEVAQHyQDTLIRA | Y579 | 0.009 | 94.25 | 1657 | 0.003948 | 0.009 | 0.0039 | 1 |
| P24462 | CYP3A7 | YRKVWGIyDCQQPML | Y75 | 0.053 | 97.4 | 503 | 0.02325 | 0.053 | 0.0232 | 0.8 |
| Q9BUJ2 | HNRNPUL1 | AsNPSkkyNILGtNA | Y450 | 0.253 | 95.67 | 856 | 0.110987 | 0.253 | 0.111 | 1 |
| O75351 | VPS4B | LGRkTDGysGADIsI | Y336 | 0.212 | 97.51 | 444 | 0.093001 | 0.212 | 0.093 | 1 |
| P38646 | HSPA9 | PNNTFyAtkRLIGRR | T120 | 0.125 | 96.78 | 679 | 0.085377 | 0.215 | 0.1429 | 1 |
| P38646 | HSPA9 | VEAHGkLysPsQIGA | Y161 | 0.159 | 96.24 | 679 | 0.069751 | 0.26 | 0.0967 | 1 |
| P38646 | HSPA9 | EAHGkLysPsQIGAF | S162 | 0.024 | 96.34 | 679 | 0.019078 | 0.188 | 0.1494 | 0.8 |
| P38646 | HSPA9 | VLMKMkEtAENyLGH | T177 | 0.071 | 97.6 | 679 | 0.048494 | 0.125 | 0.0854 | 1 |
| P38646 | HSPA9 | GGGTFDIsILEIQKG | S253 | 0.019 | 97.07 | 679 | 0.015103 | 0.019 | 0.0151 | 1 |
| P38646 | HSPA9 | KGVFEVksTNGDtFL | S266 | 0 | 95.4 | 679 | 0 | 0 | 0 | 1 |
| P38646 | HSPA9 | VtDLIrRtIAPCQkA | T362 | 0.031 | 97.87 | 679 | 0.021173 | 0.031 | 0.0212 | 1 |
| P38646 | HSPA9 | VILVGGMtRMPkVQQ | T390 | 0.032 | 98.11 | 679 | 0.021856 | 0.032 | 0.0219 | 1 |

| | | | | | | | | | | |
|---|---|---|---|---|---|---|---|---|---|---|
| P63010 | AP2B1 | ANSAVVLsAVKVLMk | S258 | 0.101 | 96.92 | 937 | 0.080286 | 0.119 | 0.0946 | 1 |
| P63010 | AP2B1 | LLPkDsDyyNMLLkK | Y276 | 0.267 | 93.02 | 937 | 0.117129 | 0.267 | 0.1171 | 1 |
| P63010 | AP2B1 | LPkDsDyyNMLLkKL | Y277 | 0.094 | 94 | 937 | 0.041236 | 0.094 | 0.0412 | 0.8182 |
| Q6DT37 | CDC42BPG | VGtPDYIsPEILQAM | S239 | 0.003 | 93.94 | 1551 | 0.002385 | 0.003 | 0.0024 | 1 |
| Q9BXM9 | FSD1L | GLkFDSkyMNFRVRA | Y273 | 0.327 | 91.2 | 530 | 0.14345 | 0.327 | 0.1434 | 1 |
| Q9BXM9 | FSD1L | AIESGQHyWEVKAQK | Y386 | 0.005 | 96.35 | 530 | 0.002193 | 0.005 | 0.0022 | 1 |
| Q9BXM9 | FSD1L | CGGLsLstGMQVPSA | T499 | 0.106 | 95.12 | 530 | 0.072399 | 0.106 | 0.0724 | 1 |
| Q01813 | PFKP | VYFIYEGyQGMVDGG | Y64 | 0.137 | 98.08 | 784 | 0.0601 | 0.137 | 0.0601 | 1 |
| Q01813 | PFKP | TDFIyQLySEEGKGV | Y654 | 0.006 | 98.2 | 784 | 0.002632 | 0.006 | 0.0026 | 1 |
| Q8IVF4 | DNAH10 | WKKAYFDtRAKIEAS | T440 | 0.091 | 91.43 | 4471 | 0.062154 | 0.091 | 0.0622 | 1 |
| O75891 | ALDH1L1 | PRHGSIIyHPSLLPr | Y105 | 0.214 | 97.53 | 902 | 0.093879 | 0.214 | 0.0939 | 0.9375 |
| P04001 | OPN1MW | RWVYHLTsVWMIFVV | S57 | 0.072 | 94.41 | 364 | 0.057234 | 0.072 | 0.0572 | 1 |
| P09327 | VIL1 | DQERGGRtyVGVVDG | T206 | 0.077 | 76.83 | 827 | 0.052592 | 0.107 | 0.109 | 1 |
| P53621 | COPA | SARVkGLsFHPKRPW | S16 | 0.084 | 97.65 | 1224 | 0.066773 | 0.168 | 0.1335 | 1 |
| P53621 | COPA | PTMPLIVsGADDRQV | S222 | 0 | 98.26 | 1224 | 0 | 0 | 0 | 1 |
| P53621 | COPA | SkAWEVDtCrGHyNN | T244 | 0.077 | 93.36 | 1224 | 0.052592 | 0.077 | 0.0526 | 1 |
| P53621 | COPA | PRQELILsNSEDksI | S266 | 0 | 97.77 | 1224 | 0 | 0 | 0 | 1 |
| P53621 | COPA | LsNSEDksIRVWDMS | S272 | 0.03 | 95.39 | 1224 | 0.023847 | 0.03 | 0.0238 | 1 |
| Q9BUX1 | CHAC1 | VGFVRGYsRRFWQGD | S61 | 0.163 | 96.56 | 222 | 0.129571 | 0.163 | 0.1296 | 1 |
| O75881 | CYP7B1 | LQKQHGDtFTVLLGG | T80 | 0.073 | 95.8 | 506 | 0.04986 | 0.122 | 0.0806 | 1 |
| O75881 | CYP7B1 | SLICLESsIFEALRL | S355 | 0.024 | 98.71 | 506 | 0.019078 | 0.024 | 0.0191 | 1 |
| Q9Y4C2 | TCAF1 | NLWVKMFsHQVQKNL | S874 | 0 | 98.01 | 921 | 0 | 0 | 0 | 1 |
| P22891 | PROZ | TRTYCERssVAAMHW | S344 | 0.045 | 89.79 | 400 | 0.035771 | 0.045 | 0.0358 | 0.8889 |
| Q9BQE3 | TUBA1C | CDPRHGkyMACCLLy | Y312 | 0.043 | 94.42 | 449 | 0.018863 | 0.043 | 0.0189 | 0.8667 |
| Q9BQE3 | TUBA1C | yMACCLLyRGDVVPk | Y319 | 0.023 | 95.51 | 449 | 0.01009 | 0.023 | 0.0101 | 1 |
| Q9BQE3 | TUBA1C | QRAVCMLsNTTAVAE | S379 | 0 | 95 | 449 | 0 | 0 | 0 | 0.8889 |
| Q9BQE3 | TUBA1C | MAALEkDyEEVGADs | Y432 | 0.021 | 94.85 | 449 | 0.009212 | 0.021 | 0.0092 | 0.9375 |
| Q9C0D5 | TANC1 | LIkRRDKtRMFCHPS | T809 | 0.094 | 91.41 | 1861 | 0.064203 | 0.148 | 0.1011 | 1 |
| P43250 | GRK6 | KACGRQLtQNFLsHt | T106 | 0.156 | 94.9 | 576 | 0.10655 | 0.156 | 0.1066 | 1 |
| O43657 | TSPAN6 | rDWtDTNyysEkGFP | Y167 | 0.282 | 92.29 | 245 | 0.123709 | 0.306 | 0.1342 | 1 |
| Q9Y2M0 | FAN1 | NRLKNHPsLRCHEDL | S646 | 0.09 | 93.3 | 1017 | 0.071542 | 0.09 | 0.0715 | 1 |

| | | | | | | | | | | |
|---|---|---|---|---|---|---|---|---|---|---|
| Q96HA7 | TONSL | GAsVtLRtRKGLSPL | T628 | 0.182 | 92.16 | 1378 | 0.124308 | 0.182 | 0.1243 | 1 |
| Q8WVJ2 | NUDCD2 | DQMQRkLtLERFQkE | T123 | 0.006 | 84.91 | 157 | 0.004098 | 0.006 | 0.0041 | 1 |
| Q9BYJ1 | ALOXE3 | LPLCHPIyKLLLPHT | Y434 | 0.123 | 98.35 | 711 | 0.053958 | 0.244 | 0.107 | 1 |
| Q9NXL6 | SIDT1 | FHFTASPsQPQYFLY | S184 | 0.038 | 92.57 | 827 | 0.030207 | 0.038 | 0.0302 | 1 |
| O14818 | PSMA7 | DRAItVFsPDGHLFQ | S11 | 0.093 | 90.39 | 248 | 0.073927 | 0.186 | 0.1479 | 1 |
| O14818 | PSMA7 | PVTVEyItRyIASLk | T108 | 0 | 98.7 | 248 | 0 | 0 | 0 | 0.8 |
| O00471 | EXOC5 | FLDGELksDVFTNSE | S147 | 0.184 | 69.52 | 708 | 0.146264 | 0.184 | 0.1463 | 1 |
| Q9HC36 | MRM3 | WEESGLRyDKAYPGD | Y94 | 0.107 | 88.68 | 420 | 0.046939 | 0.107 | 0.0469 | 1 |
| Q9C0A0 | CNTNAP4 | GPNGDHItLQLRRAR | T233 | 0.071 | 90.22 | 1308 | 0.048494 | 0.111 | 0.0758 | 1 |
| Q3SY77 | UGT3A2 | yLLMDRVsQILQDHG | S43 | 0 | 97.03 | 523 | 0 | 0 | 0 | 1 |
| Q3SY77 | UGT3A2 | KsAAVAAsVILRsHP | S441 | 0.127 | 97.49 | 523 | 0.100954 | 0.127 | 0.101 | 1 |
| Q4G0Z9 | MCMDC2 | AERMTHGyYLASRRI | Y565 | 0.039 | 92.7 | 681 | 0.017109 | 0.039 | 0.0171 | 1 |
| Q9H0K4 | RSPH6A | CSLCPQYsVAVVRsN | S620 | 0.064 | 92.93 | 717 | 0.050875 | 0.064 | 0.0509 | 1 |
| Q9H0K4 | RSPH6A | YsVAVVRsNLWPGAY | S626 | 0.078 | 97.35 | 717 | 0.062003 | 0.078 | 0.062 | 0.875 |
| Q9UBF2 | COPG2 | DGVtMQVtVRsKERT | T854 | 0.026 | 97.87 | 871 | 0.017758 | 0.045 | 0.03 | 0.8333 |
| Q9UBF2 | COPG2 | tMQVtVRsKERTPVD | S857 | 0.007 | 95.24 | 871 | 0.005564 | 0.007 | 0.0056 | 1 |
| Q49AN0 | CRY2 | SGDYIRRyLPKLkAF | Y451 | 0.108 | 97.93 | 593 | 0.047378 | 0.108 | 0.0474 | 0.8333 |
| Q49AN0 | CRY2 | AFPSRyIyEPWNAPE | Y464 | 0.001 | 98.45 | 593 | 0.000439 | 0.001 | 0.0004 | 1 |
| Q96F10 | SAT2 | TWKGRTIyLEDIYVM | Y90 | 0.31 | 98.44 | 170 | 0.135993 | 0.31 | 0.136 | 1 |
| Q96J84 | KIRREL1 | TQADAGTytCRAIVP | Y369 | 0.027 | 97.04 | 757 | 0.011845 | 0.027 | 0.0118 | 1 |
| Q96J84 | KIRREL1 | QADAGTytCRAIVPR | T370 | 0.178 | 97.15 | 757 | 0.121576 | 0.178 | 0.1216 | 1 |
| Q5T0W9 | FAM83B | FKEIVEAstRGVsVY | S170 | 0.105 | 95.44 | 1011 | 0.083466 | 0.105 | 0.0835 | 1 |
| P63000 | RAC1 | VrHHCPNtPIILVGT | T108 | 0.188 | 97.73 | 192 | 0.128407 | 0.24 | 0.1487 | 0.8571 |
| Q99798 | ACO2 | QEIERGKsYLRLRPD | S87 | 0.086 | 97.04 | 780 | 0.068363 | 0.086 | 0.0684 | 1 |
| Q99798 | ACO2 | kDINQEVyNFLATAG | Y151 | 0.14 | 98.35 | 780 | 0.061416 | 0.241 | 0.0918 | 1 |
| Q99798 | ACO2 | GsLSGWssPkDVILK | S243 | 0.039 | 98.47 | 780 | 0.031002 | 0.039 | 0.031 | 0.875 |
| Q99798 | ACO2 | GSCTNSSyEDMGRSA | Y390 | 0.088 | 98.73 | 780 | 0.038604 | 0.176 | 0.0772 | 1 |
| Q99798 | ACO2 | IKKGEKNtIVtsYNr | T467 | 0 | 98.86 | 780 | 0 | 0 | 0 | 1 |
| Q99798 | ACO2 | GEKNtIVtsYNrNFt | T470 | 0.017 | 98.91 | 780 | 0.011611 | 0.034 | 0.0232 | 1 |
| Q99798 | ACO2 | EKNtIVtsYNrNFtG | S471 | 0 | 98.84 | 780 | 0 | 0 | 0 | 1 |
| Q99798 | ACO2 | EFDPGQDtyQHPPkD | T543 | 0.15 | 95.83 | 780 | 0.102452 | 0.106 | 0.1198 | 0.875 |

| | | | | | | | | | | |
|---|---|---|---|---|---|---|---|---|---|---|
| Q99798 | ACO2 | EFGPVPDtARYYKKH | T646 | 0.07 | 98.84 | 780 | 0.047811 | 0.115 | 0.0847 | 1 |
| Q99798 | ACO2 | VVIGDENyGEGssRE | Y665 | 0.025 | 98.48 | 780 | 0.010967 | 0.025 | 0.011 | 1 |
| Q86UB2 | BIVM | KGCSYVLyKPHGKNK | Y288 | 0.07 | 97.84 | 503 | 0.030708 | 0.07 | 0.0307 | 0.9167 |
| Q86UB2 | BIVM | DIVTDLNtQNPEYLD | T384 | 0.173 | 94.9 | 503 | 0.118161 | 0.173 | 0.1182 | 1 |
| P22223 | CDH3 | QRLNQLKsNKDRDtK | S133 | 0.063 | 88.9 | 829 | 0.05008 | 0.063 | 0.0501 | 1 |
| P22223 | CDH3 | DRDtKIFySITGPGA | Y143 | 0.215 | 94.12 | 829 | 0.094317 | 0.215 | 0.0943 | 1 |
| Q86V15 | CASZ1 | MHKKRDNsLQHGFMR | S521 | 0.077 | 76.03 | 1759 | 0.061208 | 0.077 | 0.0612 | 1 |
| Q9P2J5 | LARS1 | TTDVNPYyDSFVRWQ | Y214 | 0 | 98.56 | 1176 | 0 | 0 | 0 | 1 |
| Q9P2J5 | LARS1 | FVRWQFLtLRERNKI | T224 | 0.013 | 98.34 | 1176 | 0.008879 | 0.013 | 0.0089 | 1 |
| Q9P2J5 | LARS1 | EGVGPQEyTLLkLkV | Y264 | 0.277 | 94.91 | 1176 | 0.121516 | 0.277 | 0.1215 | 1 |
| Q9P2J5 | LARS1 | NIFLVAAtLRPETMF | T293 | 0.152 | 94.79 | 1176 | 0.103818 | 0.152 | 0.1038 | 1 |
| Q9P2J5 | LARS1 | QkAARNMsYQGFTkD | S335 | 0.002 | 96.6 | 1176 | 0.00159 | 0.002 | 0.0016 | 1 |
| Q9P2J5 | LARS1 | ASLSAPLtSYKVIYV | T367 | 0.203 | 95.35 | 1176 | 0.138652 | 0.203 | 0.1387 | 1 |
| Q9P2J5 | LARS1 | GVVTSVPsDsPDDIA | S394 | 0.002 | 95.24 | 1176 | 0.00159 | 0.002 | 0.0016 | 1 |
| Q9P2J5 | LARS1 | DAGDALIyMEPEkQV | Y507 | 0.009 | 96.09 | 1176 | 0.003948 | 0.009 | 0.0039 | 1 |
| Q9P2J5 | LARS1 | MTKEVWDyVFFkEAP | Y637 | 0.21 | 98.52 | 1176 | 0.092124 | 0.21 | 0.0921 | 1 |
| Q9P2J5 | LARS1 | kQEFEFWyPVDLRVS | Y666 | 0.147 | 97.92 | 1176 | 0.064487 | 0.147 | 0.0645 | 1 |
| Q02246 | CNTN2 | TDGRHFVsQTTGNLY | S186 | 0.023 | 93.11 | 1040 | 0.018283 | 0.023 | 0.0183 | 1 |
| Q02246 | CNTN2 | NAsDLGNySCLAtSH | Y205 | 0.02 | 94.75 | 1040 | 0.008774 | 0.02 | 0.0088 | 1 |
| Q02246 | CNTN2 | GNySCLAtSHMDFST | T210 | 0.175 | 94 | 1040 | 0.119527 | 0.175 | 0.1195 | 0.875 |
| Q02246 | CNTN2 | PNTKYHVtVRAyNRA | T887 | 0.023 | 95.2 | 1040 | 0.015709 | 0.023 | 0.0157 | 1 |
| Q02246 | CNTN2 | YHVtVRAyNRAGTGP | Y891 | 0.336 | 90.04 | 1040 | 0.147398 | 0.336 | 0.1474 | 1 |
| Q6P1X5 | TAF2 | MIDALANsVTPAVSV | S805 | 0.092 | 80.57 | 1199 | 0.073132 | 0.139 | 0.1105 | 1 |
| O95237 | LRAT | PRTHLTHyGIYLGDN | Y61 | 0.11 | 97.43 | 230 | 0.048255 | 0.11 | 0.0483 | 1 |
| Q9HCJ1 | ANKH | VGLVFVNsKRDRTKA | S79 | 0 | 88.99 | 492 | 0 | 0 | 0 | 1 |
| P40429 | RPL13A | PYHFrAPsRIFWRTV | S77 | 0.116 | 98.21 | 203 | 0.09221 | 0.116 | 0.0922 | 1 |
| Q6PI73 | LILRA6 | LAPGQSLtLQCGsDV | T242 | 0.176 | 93.98 | 481 | 0.12021 | 0.176 | 0.1202 | 1 |
| Q6PI73 | LILRA6 | SLtLQCGsDVGYDRF | S247 | 0.052 | 95.58 | 481 | 0.041336 | 0.052 | 0.0413 | 1 |
| Q6WRX3 | ZYG11A | LHRLFTEtFsMEVTM | T371 | 0 | 93.08 | 759 | 0 | 0 | 0 | 0.8889 |
| Q6WRX3 | ZYG11A | VRLLsEVtCLLFKAL | T427 | 0.059 | 95.49 | 759 | 0.040298 | 0.059 | 0.0403 | 1 |
| Q6WRX3 | ZYG11A | LQKNCLLsLTNSRIL | S450 | 0.081 | 92.85 | 759 | 0.064388 | 0.081 | 0.0644 | 1 |

| | | | | | | | | | | |
|---|---|---|---|---|---|---|---|---|---|---|
| P41732 | TSPAN7 | kDTFLRTyTDAMQTy | Y124 | 0.147 | 91.55 | 249 | 0.064487 | 0.147 | 0.0645 | 1 |
| P41732 | TSPAN7 | yTDAMQTyNGNDERS | Y131 | 0.17 | 93.84 | 249 | 0.074577 | 0.17 | 0.0746 | 1 |
| Q504T8 | MIDN | RPRRDIGtILQILND | T305 | 0.006 | 92.41 | 468 | 0.004098 | 0.072 | 0.0492 | 1 |
| Q5TCS8 | AK9 | NTEEAAKyREKIYYF | Y957 | 0.016 | 87.43 | 1911 | 0.007019 | 0.29 | 0.1272 | 1 |
| Q9Y5V3 | MAGED1 | EFLWGLRsYHEtskM | S638 | 0.014 | 92.37 | 778 | 0.011129 | 0.014 | 0.0111 | 0.875 |
| Q70J99 | UNC13D | PEQRALLyEDALytV | Y56 | 0.07 | 95.59 | 1090 | 0.030708 | 0.07 | 0.0307 | 1 |
| Q70J99 | UNC13D | LLyEDALytVLHRLG | Y61 | 0 | 96.2 | 1090 | 0 | 0 | 0 | 1 |
| Q70J99 | UNC13D | VSGFSDPyCLLGIEQ | Y135 | 0.155 | 91.55 | 1090 | 0.067996 | 0.31 | 0.136 | 1 |
| Q5R372 | RABGAP1L | FLALNEEtPKDKQVY | T383 | 0.095 | 91.14 | 815 | 0.064886 | 0.095 | 0.0649 | 1 |
| P53992 | SEC24C | LDPDAIPsPIQVIED | S328 | 0.174 | 95.6 | 1094 | 0.138315 | 0.174 | 0.1383 | 0.8 |
| Q13077 | TRAF1 | YKLCLRLyLNGDGtG | Y310 | 0.257 | 98.17 | 416 | 0.112742 | 0.257 | 0.1127 | 1 |
| Q8NET6 | CHST13 | SAAFQRRyGARIVQR | Y200 | 0.124 | 96.85 | 341 | 0.054397 | 0.124 | 0.0544 | 1 |
| O75019 | LILRA1 | GILETQEyRLYREKK | Y58 | 0.138 | 91.8 | 489 | 0.060539 | 0.186 | 0.0816 | 1 |
| Q7Z5U6 | WDR53 | KWTGGHSsPVLCLNA | S11 | 0.182 | 97 | 358 | 0.144674 | 0.182 | 0.1447 | 0.8 |
| Q7Z5U6 | WDR53 | QTENLLAsADDSGAI | S111 | 0.021 | 98.78 | 358 | 0.016693 | 0.021 | 0.0167 | 0.8889 |
| Q495T6 | MMEL1 | KARTLYRsCMNQsVI | S174 | 0.022 | 98.23 | 779 | 0.017488 | 0.022 | 0.0175 | 1 |
| Q495T6 | MMEL1 | QPTLGMPsREyYFNG | S260 | 0.12 | 93.32 | 779 | 0.09539 | 0.12 | 0.0954 | 1 |
| Q495T6 | MMEL1 | LGMPsREyYFNGGSN | Y263 | 0.023 | 94.98 | 779 | 0.01009 | 0.023 | 0.0101 | 1 |
| Q495T6 | MMEL1 | SIKTDVHsPLKYRVL | S742 | 0.039 | 98.54 | 779 | 0.031002 | 0.039 | 0.031 | 0.875 |
| P35626 | GRK3 | AsVGtHGyMAPEVLQ | Y356 | 0.264 | 96.8 | 688 | 0.115813 | 0.264 | 0.1158 | 1 |
| Q6XQN6 | NAPRT | TEDPEkQtLPGSkAA | T416 | 0.043 | 92.38 | 538 | 0.02937 | 0.043 | 0.0294 | 1 |
| P0DJD1 | RGPD2 | LIKIIDDsDsNLsVV | S724 | 0.055 | 86.91 | 1756 | 0.04372 | 0.055 | 0.0437 | 1 |
| P53618 | COPB1 | NMVDLNDyLQHILkS | Y865 | 0 | 87.89 | 953 | 0 | 0 | 0 | 1 |
| P53618 | COPB1 | KsQGMALsLGDkINL | S939 | 0.102 | 81.58 | 953 | 0.081081 | 0.102 | 0.0811 | 1 |
| Q6PIY7 | TENT2 | IkDQFLKsWHRLKNK | S461 | 0.02 | 98.73 | 484 | 0.015898 | 0.02 | 0.0159 | 1 |
| Q7L5N7 | LPCAT2 | VQPVLLRyPNKLDTV | Y250 | 0.238 | 96.66 | 544 | 0.104407 | 0.238 | 0.1044 | 0.9333 |
| Q7L5N7 | LPCAT2 | VRkHLDEyASIASSS | Y367 | 0.012 | 91.03 | 544 | 0.005264 | 0.012 | 0.0053 | 1 |
| P05091 | ALDH2 | VLKCLryyAGWADky | Y149 | 0.001 | 98.93 | 517 | 0.000439 | 0.001 | 0.0004 | 1 |
| P05091 | ALDH2 | PIDGDFFsYTrHEPV | S169 | 0.032 | 98.77 | 517 | 0.025437 | 0.032 | 0.0254 | 1 |
| P05091 | ALDH2 | VAEQTPLtALyVANL | T217 | 0.011 | 98.91 | 517 | 0.007513 | 0.011 | 0.0075 | 1 |
| P05091 | ALDH2 | QTPLtALyVANLIKE | Y220 | 0.297 | 98.91 | 517 | 0.13029 | 0.297 | 0.1303 | 1 |

| | | | | | | | | | | |
|---|---|---|---|---|---|---|---|---|---|---|
| P05091 | ALDH2 | TFVQEDIyDEFVERS | Y332 | 0.084 | 98.87 | 517 | 0.03685 | 0.084 | 0.0369 | 1 |
| P05091 | ALDH2 | GLAAAVFtKDLDKAN | T450 | 0.046 | 98.86 | 517 | 0.031419 | 0.046 | 0.0314 | 1 |
| P05091 | ALDH2 | GEyGLQAytEVKTVT | Y502 | 0.079 | 98.92 | 517 | 0.034656 | 0.079 | 0.0347 | 1 |
| Q8N394 | TMTC2 | VIAERVLyIPSMGFC | Y425 | 0.005 | 98.66 | 836 | 0.002193 | 0.005 | 0.0022 | 1 |
| Q8N394 | TMTC2 | YNMMGEAyMRLSKLP | Y653 | 0.238 | 98.5 | 836 | 0.104407 | 0.238 | 0.1044 | 1 |
| Q96L93 | KIF16B | ERtKTFTyDFSFYSA | Y58 | 0.085 | 96.57 | 1317 | 0.037288 | 0.151 | 0.0595 | 1 |
| Q96L93 | KIF16B | TGSGKsytMMGNSGD | T111 | 0.021 | 96.17 | 1317 | 0.014343 | 0.042 | 0.0287 | 1 |
| Q86TD4 | SRL | LQRLRKIyHsSIKPL | Y64 | 0.011 | 97.12 | 473 | 0.004826 | 0.011 | 0.0048 | 1 |
| Q86X29 | LSR | AVtLGDyyQGRRITI | Y190 | 0.001 | 93.12 | 649 | 0.000439 | 0.001 | 0.0004 | 1 |
| Q03013 | GSTM4 | LDFPNLPyLIDGAHK | Y62 | 0.047 | 98.92 | 218 | 0.020618 | 0.047 | 0.0206 | 1 |
| Q96ND0 | FAM210A | YTVTLGGtsVtVKyL | T209 | 0 | 88.86 | 272 | 0 | 0.015 | 0.0102 | 1 |
| P24347 | MMP11 | FRYPLSLsPDDCRGV | S246 | 0.176 | 92.83 | 488 | 0.139905 | 0.176 | 0.1399 | 1 |
| O14641 | DVL2 | GGGGVGEtKVIyHLD | T14 | 0 | 89.74 | 736 | 0 | 0 | 0 | 1 |
| O14641 | DVL2 | GAkYFFksMDQDFGV | S59 | 0.111 | 91.57 | 736 | 0.088236 | 0.111 | 0.0882 | 1 |
| P04350 | TUBB4A | QLERINVyYNEAtGG | Y50 | 0.019 | 96.55 | 444 | 0.008335 | 0.019 | 0.0083 | 0.8 |
| P04350 | TUBB4A | NEAtGGNyVPRAVLV | Y59 | 0.266 | 96.56 | 444 | 0.11669 | 0.266 | 0.1167 | 1 |
| P04350 | TUBB4A | sLGGGtGsGMGtLLI | S145 | 0 | 96.51 | 444 | 0 | 0 | 0 | 1 |
| P04350 | TUBB4A | NtFsVVPsPkVSDTV | S172 | 0.01 | 84.76 | 444 | 0.007949 | 0.01 | 0.0079 | 0.8333 |
| P04350 | TUBB4A | ALtVPELtQQMFDAk | T290 | 0.008 | 96.22 | 444 | 0.005464 | 0.008 | 0.0055 | 1 |
| P04350 | TUBB4A | PRHGRyLtVAAVFrG | T312 | 0.042 | 96.33 | 444 | 0.028687 | 0.083 | 0.0566 | 1 |
| P04350 | TUBB4A | AAtFIGNsTAIQELF | S371 | 0.001 | 97.13 | 444 | 0.000795 | 0.001 | 0.0008 | 1 |
| P04350 | TUBB4A | QELFkRIsEQFtAMF | S382 | 0.14 | 95.07 | 444 | 0.111288 | 0.14 | 0.1113 | 1 |
| Q6ZWJ1 | STXBP4 | KESMIGVsFEEAKSI | S82 | 0.184 | 95.17 | 553 | 0.146264 | 0.184 | 0.1463 | 1 |
| Q5HYA8 | TMEM67 | LKFVAASyDIRGNFL | Y337 | 0.029 | 94.62 | 995 | 0.012722 | 0.317 | 0.1391 | 1 |
| Q5HYA8 | TMEM67 | IFLVDAVsGRENDLG | S449 | 0.026 | 91.68 | 995 | 0.020668 | 0.026 | 0.0207 | 1 |
| Q5HYA8 | TMEM67 | VLASLLKtAGWKRRI | T543 | 0.117 | 92.37 | 995 | 0.079913 | 0.117 | 0.0799 | 1 |
| P43320 | CRYBB2 | RRtDsLssLRPIKVD | S96 | 0 | 98.47 | 205 | 0 | 0 | 0 | 1 |
| P13667 | PDIA4 | CKQFAPEyEkIANIL | Y101 | 0.031 | 97.5 | 645 | 0.013599 | 0.031 | 0.0136 | 1 |
| P13667 | PDIA4 | RKGRPyDyNGPrEky | Y266 | 0.228 | 95.54 | 645 | 0.10002 | 0.228 | 0.1 | 1 |
| P13667 | PDIA4 | EkyGIVDyMIEQSGP | Y278 | 0.337 | 96.71 | 645 | 0.147837 | 0.337 | 0.1478 | 0.9091 |
| P13667 | PDIA4 | IEQSGPPsKEILTLk | S287 | 0.021 | 97.32 | 645 | 0.016693 | 0.021 | 0.0167 | 1 |

| | | | | | | | | | | |
|---|---|---|---|---|---|---|---|---|---|---|
| P13667 | PDIA4 | KGESDPAyQQyQDAA | Y320 | 0.226 | 96.9 | 645 | 0.099143 | 0.226 | 0.0991 | 1 |
| P13667 | PDIA4 | SDPAyQQyQDAANNL | Y323 | 0 | 96.1 | 645 | 0 | 0 | 0 | 1 |
| P13667 | PDIA4 | VsNDAkRytRRPLVV | Y409 | 0.078 | 95.58 | 645 | 0.034217 | 0.078 | 0.0342 | 1 |
| P13667 | PDIA4 | AKDFPEytFAIADEE | T449 | 0.125 | 97.76 | 645 | 0.085377 | 0.125 | 0.0854 | 1 |
| P13667 | PDIA4 | AIADEEDyAGEVKDL | Y458 | 0.186 | 95.56 | 645 | 0.081596 | 0.186 | 0.0816 | 1 |
| P13667 | PDIA4 | MEPEEFDsDTLREFV | S495 | 0.164 | 97.87 | 645 | 0.130366 | 0.164 | 0.1304 | 1 |
| P13667 | PDIA4 | CKQLEPVyNsLAkKY | Y565 | 0.031 | 97.77 | 645 | 0.013599 | 0.031 | 0.0136 | 1 |
| P13667 | PDIA4 | AtANDVPsDrYkVEG | S592 | 0.169 | 88.14 | 645 | 0.134341 | 0.169 | 0.1343 | 1 |
| P13667 | PDIA4 | VEGFPTIyFAPsGDk | Y604 | 0.143 | 98.47 | 645 | 0.062732 | 0.143 | 0.0627 | 1 |
| O15530 | PDPK1 | FQDDEKLyFGLsyAK | Y156 | 0.141 | 97.96 | 556 | 0.061855 | 0.141 | 0.0619 | 1 |
| Q13045 | FLII | LDRGLDIyVWRGAQA | Y657 | 0.183 | 94.92 | 1269 | 0.080279 | 0.321 | 0.1408 | 1 |
| Q9NU19 | TBC1D22B | PREVRPItWRLLSGY | T219 | 0 | 98.62 | 505 | 0 | 0 | 0 | 1 |
| P42263 | GRIA3 | WEKFVYLyDTERGFS | Y165 | 0.226 | 92.52 | 894 | 0.099143 | 0.226 | 0.0991 | 1 |
| Q7L1T6 | CYB5R4 | IYPTGLFtPELDRLQ | T361 | 0.104 | 97.19 | 521 | 0.071033 | 0.104 | 0.071 | 1 |
| O15075 | DCLK1 | KYTERDAsGMLyNLA | S489 | 0.083 | 95.59 | 740 | 0.065978 | 0.071 | 0.0903 | 1 |
| Q3ZCM7 | TUBB8 | _MREIVLtQIGQCGN | T7 | 0 | 98.23 | 444 | 0 | 0 | 0 | 1 |
| Q3ZCM7 | TUBB8 | NTFsILPsPKVSDTV | S172 | 0.009 | 86.08 | 444 | 0.007154 | 0.009 | 0.0072 | 0.8333 |
| Q3ZCM7 | TUBB8 | MPGFAPLtsrGsQQy | T274 | 0.186 | 89.88 | 444 | 0.127041 | 0.24 | 0.1492 | 0.8571 |
| Q9UJV9 | DDX41 | GVHMMVAtPGRLMDL | T320 | 0.152 | 94.51 | 622 | 0.103818 | 0.152 | 0.1038 | 0.8 |
| O75469 | NR1I2 | PHMADMStyMFKGII | T248 | 0.199 | 98.37 | 434 | 0.13592 | 0.199 | 0.1359 | 1 |
| Q7RTR0 | NLRP9 | QKKMLPEssLLIALG | S269 | 0.003 | 93.45 | 991 | 0.002385 | 0.003 | 0.0024 | 1 |
| Q7RTR0 | NLRP9 | KKMLPEssLLIALGK | S270 | 0.001 | 95.32 | 991 | 0.000795 | 0.065 | 0.0517 | 1 |
| P14598 | NCF1 | KRGWIPAsFLEPLDS | S208 | 0.06 | 96.16 | 390 | 0.047695 | 0.06 | 0.0477 | 1 |
| P14598 | NCF1 | TGYFPSMyLQKsGQD | Y279 | 0.184 | 96.96 | 390 | 0.080718 | 0.184 | 0.0807 | 0.9091 |
| P14598 | NCF1 | QDAYRRNsVRFLQQR | S328 | 0.028 | 89.62 | 390 | 0.022258 | 0.028 | 0.0223 | 1 |
| P49792 | RANBP2 | LIKIIDDsDsNLsVV | S725 | 0.058 | 88.665 | 3224 | 0.046105 | 0.058 | 0.0461 | 1 |
| O75578 | ITGA10 | GGRRLFLsGAPRFRH | S449 | 0.005 | 98.46 | 1167 | 0.003975 | 0.087 | 0.0692 | 1 |
| O75578 | ITGA10 | RRKVLVStTLENRKE | T828 | 0.009 | 91.79 | 1167 | 0.006147 | 0.009 | 0.0061 | 1 |
| P02788 | LTF | RQPRTHyyAVAVVKK | Y112 | 0.261 | 98.54 | 710 | 0.114497 | 0.261 | 0.1145 | 0.8889 |
| P02788 | LTF | NSVKGKKsCHtAVDR | S477 | 0 | 98.72 | 710 | 0 | 0 | 0 | 1 |
| P02788 | LTF | KGKKsCHtAVDRtAG | T480 | 0.007 | 98.55 | 710 | 0.004781 | 0.007 | 0.0048 | 1 |

| | | | | | | | | | | |
|---|---|---|---|---|---|---|---|---|---|---|
| P02788 | LTF | APNHAVVsRMDKVER | S620 | 0.001 | 98.77 | 710 | 0.000795 | 0.001 | 0.0008 | 1 |
| Q9UBJ2 | ABCD2 | HRPSLWKyHTHLLQF | Y670 | 0.237 | 90.25 | 740 | 0.103968 | 0.237 | 0.104 | 1 |
| Q9UI10 | EIF2B4 | MVRLGLQySQGLVSG | Y214 | 0.062 | 93.06 | 523 | 0.027199 | 0.062 | 0.0272 | 1 |
| Q9UIV8 | SERPINB13 | AKILGIPyKNNDLSM | Y237 | 0.015 | 94.31 | 391 | 0.00658 | 0.015 | 0.0066 | 1 |
| Q99439 | CNN2 | RTWIEGLtGLsIGPD | T42 | 0.167 | 92.33 | 309 | 0.114063 | 0.167 | 0.1141 | 0.875 |
| Q9NVM9 | INTS13 | KLVNFIVsDsGAHVL | S81 | 0 | 97.79 | 706 | 0 | 0 | 0 | 1 |
| Q6UX41 | BTNL8 | tVLDAGLyGCRISSQ | Y112 | 0.028 | 97.77 | 500 | 0.012283 | 0.028 | 0.0123 | 1 |
| Q9BXS5 | AP1M1 | GAPRPPAtVtNAVsW | T152 | 0.085 | 91.97 | 423 | 0.058056 | 0.17 | 0.1161 | 1 |
| Q9BXS5 | AP1M1 | EYMIKAKsQFKRRsT | S299 | 0.067 | 97.71 | 423 | 0.053259 | 0.067 | 0.0533 | 1 |
| Q9BXS5 | AP1M1 | KsQFKRRsTANNVEI | S305 | 0.116 | 97.04 | 423 | 0.09221 | 0.116 | 0.0922 | 1 |
| Q5VT66 | MTARC1 | DLLLPIktPTTNAVH | T140 | 0.195 | 93.47 | 337 | 0.133188 | 0.195 | 0.1332 | 1 |
| Q5VT66 | MTARC1 | kSPLFGQyFVLENPG | Y317 | 0.276 | 98.86 | 337 | 0.121077 | 0.276 | 0.1211 | 0.9167 |
| Q8IVF2 | AHNAK2 | VKQVLKDsSAAkLFN | S145 | 0.152 | 84.84 | 5795 | 0.120827 | 0.152 | 0.1208 | 1 |
| Q9P2I0 | CPSF2 | LNCAIYAtIPVYKMG | T87 | 0 | 98.72 | 782 | 0 | 0 | 0 | 1 |
| Q9P2I0 | CPSF2 | LLtNVLEtLRGDGNV | T226 | 0.096 | 98.12 | 782 | 0.065569 | 0.177 | 0.1209 | 1 |
| Q9P2I0 | CPSF2 | NVLIAVDtAGRVLEL | T239 | 0.015 | 98.23 | 782 | 0.010245 | 0.015 | 0.0102 | 1 |
| Q9P2I0 | CPSF2 | KSQVEWMsDkLMRCF | S288 | 0.003 | 91.95 | 782 | 0.002385 | 0.003 | 0.0024 | 1 |
| Q12830 | BPTF | TISTLRLtItQLENN | T909 | 0.1 | 95.11 | 3046 | 0.068301 | 0.171 | 0.1168 | 1 |
| Q9BQD7 | ANTKMT | GKtVDLGsGDGRIVL | S84 | 0.09 | 94.47 | 235 | 0.071542 | 0.115 | 0.0987 | 1 |
| Q3MJ16 | PLA2G4E | DLIIHLNyCAGSQTK | Y727 | 0 | 92.47 | 868 | 0 | 0 | 0 | 1 |
| Q13761 | RUNX3 | ksFtLtItVFTNPTQ | T155 | 0.169 | 98.45 | 415 | 0.115429 | 0.169 | 0.1154 | 0.8571 |
| O76083 | PDE9A | ANSERTPykVRPVAI | Y76 | 0.095 | 94.27 | 593 | 0.041675 | 0.14 | 0.0614 | 0.8462 |
| Q9Y6M7 | SLC4A7 | SDFAVFLtIVIMVTI | T867 | 0 | 91.55 | 1214 | 0 | 0 | 0 | 1 |
| Q86X27 | RALGPS2 | PIFRLTKtWALLSRK | T158 | 0.008 | 98.04 | 583 | 0.005464 | 0.008 | 0.0055 | 0.8571 |
| Q86Z14 | KLB | LRWVRRNyGDMDIyI | Y880 | 0.228 | 97.67 | 1044 | 0.10002 | 0.228 | 0.1 | 1 |
| Q86Z14 | KLB | NyGDMDIyITASGID | Y886 | 0.305 | 98.75 | 1044 | 0.133799 | 0.305 | 0.1338 | 1 |
| Q86Z14 | KLB | DDRLRKYyLGKyLQE | Y906 | 0.01 | 96.57 | 1044 | 0.004387 | 0.01 | 0.0044 | 1 |
| Q86Z14 | KLB | RKYyLGKyLQEVLKA | Y910 | 0.018 | 98 | 1044 | 0.007896 | 0.018 | 0.0079 | 1 |
| Q86V21 | AACS | VPEWFkGsRLNYAEN | S97 | 0.009 | 98.68 | 672 | 0.007154 | 0.009 | 0.0072 | 1 |
| Q8NBW4 | SLC38A9 | FNILGTVsVLyLIFL | S321 | 0.04 | 89.05 | 561 | 0.031797 | 0.04 | 0.0318 | 1 |
| Q08499 | PDE4D | SRsGNQVsEFIsNTF | S330 | 0.093 | 85.56 | 809 | 0.073927 | 0.093 | 0.0739 | 1 |

| | | | | | | | | | | |
|---|---|---|---|---|---|---|---|---|---|---|
| P16591 | FER | PLAEQDWyHGAIPRI | Y461 | 0.117 | 94.66 | 822 | 0.051326 | 0.117 | 0.0513 | 0.8182 |
| P16591 | FER | EyVLSVysDGQRRHF | S498 | 0.06 | 94.3 | 822 | 0.047695 | 0.06 | 0.0477 | 1 |
| P16591 | FER | KtsVAVktCkEDLPQ | T592 | 0.105 | 91.83 | 822 | 0.071716 | 0.105 | 0.0717 | 1 |
| P16591 | FER | EAkILkQyDHPNIVK | Y615 | 0.162 | 93.15 | 822 | 0.071067 | 0.162 | 0.0711 | 0.8333 |
| P20742 | PZP | GMGLKVFtNsKIRKP | T675 | 0.113 | 91.2 | 1482 | 0.077181 | 0.113 | 0.0772 | 0.9 |
| Q7Z3Z3 | PIWIL3 | SIAGFVAsTNAELTK | S672 | 0 | 98.34 | 882 | 0 | 0 | 0 | 1 |
| Q7Z3Z3 | PIWIL3 | DHEAKKMsTYLKTIS | S743 | 0.133 | 92.76 | 882 | 0.105724 | 0.133 | 0.1057 | 1 |
| P0DPH7 | TUBA3C | kEDAANNyARGHytI | Y103 | 0.018 | 98.05 | 450 | 0.007896 | 0.018 | 0.0079 | 0.9474 |
| P0DPH7 | TUBA3C | MVDNEAIyDICRRNL | Y210 | 0.324 | 95.38 | 450 | 0.142134 | 0.324 | 0.1421 | 1 |
| O43772 | SLC25A20 | GMLSGVFtTGIMTPG | T124 | 0.091 | 91.71 | 301 | 0.062154 | 0.091 | 0.0622 | 0.9 |
| Q13835 | PKP1 | AVsLLRRtGNAEIQK | T341 | 0.168 | 93.61 | 747 | 0.114746 | 0.168 | 0.1147 | 1 |
| Q99797 | MIPEP | SSTFLMGtNFPNKIE | T223 | 0.043 | 93.59 | 713 | 0.02937 | 0.043 | 0.0294 | 1 |
| Q99797 | MIPEP | MQFLEKLsDKLsErT | S325 | 0.161 | 98.36 | 713 | 0.127981 | 0.161 | 0.128 | 1 |
| Q99797 | MIPEP | SMLGRTRyQHVTGTR | Y510 | 0.024 | 98.16 | 713 | 0.010528 | 0.024 | 0.0105 | 1 |
| Q99797 | MIPEP | LVGyGARyySyLMSR | Y632 | 0.255 | 97.22 | 713 | 0.111865 | 0.255 | 0.1119 | 1 |
| Q99797 | MIPEP | VGyGARyySyLMSRA | Y633 | 0.019 | 97.98 | 713 | 0.008335 | 0.019 | 0.0083 | 1 |
| O75828 | CBR3 | FsGDVVLtARDVARG | T36 | 0 | 98.92 | 277 | 0 | 0 | 0 | 0.9 |
| Q99685 | MGLL | EESSPRRtPQSIPYQ | T10 | 0.075 | 98.31 | 303 | 0.051226 | 0.075 | 0.0512 | 1 |
| Q8N8Q8 | COX18 | VALRGAVtLPLAAyQ | T98 | 0.013 | 93.4 | 333 | 0.008879 | 0.013 | 0.0089 | 0.8182 |
| Q8NBP5 | MFSD9 | STLVGCWsDVVGRRS | S103 | 0.018 | 95.02 | 474 | 0.014308 | 0.018 | 0.0143 | 1 |
| Q9HBH5 | RDH14 | RLEGtNVtVNVLHPG | T241 | 0.174 | 98.81 | 336 | 0.118844 | 0.174 | 0.1188 | 1 |
| Q9NYQ7 | CELSR3 | DVsDHLFysFERGNE | Y1208 | 0.231 | 94 | 3312 | 0.101336 | 0.231 | 0.1013 | 1 |
| P10915 | HAPLN1 | HKKTyGGyQGRVFLK | Y108 | 0.208 | 97.45 | 354 | 0.091247 | 0.208 | 0.0912 | 0.9231 |
| Q07020 | RPL18 | FLARRtNstFNQVVL | S41 | 0.166 | 94.73 | 188 | 0.131956 | 0.166 | 0.132 | 1 |
| Q07020 | RPL18 | LPGRENktAVVVGtI | T79 | 0.069 | 97.47 | 188 | 0.047128 | 0.069 | 0.0471 | 1 |
| Q9BZQ4 | NMNAT2 | CGSFNPItKGHIQMF | T21 | 0.044 | 98.11 | 307 | 0.030053 | 0.044 | 0.0301 | 0.875 |
| P40925 | MDH1 | GLPINDFsREkMDLT | S309 | 0.033 | 98.56 | 334 | 0.026232 | 0.033 | 0.0262 | 0.875 |
| P19338 | NCL | tLVLSNLsYsAtEET | S494 | 0.125 | 82.86 | 710 | 0.099364 | 0.125 | 0.0994 | 1 |
| P84090 | ERH | _____MsHtILLVQPt | T4 | 0.01 | 98.53 | 104 | 0.00683 | 0.01 | 0.0068 | 0.9091 |
| Q9Y6M4 | CSNK1G3 | LHLEyRFykQLGSGD | Y90 | 0.096 | 97.92 | 447 | 0.042114 | 0.189 | 0.0829 | 0.9167 |
| Q9Y6M4 | CSNK1G3 | DFGLAKEyIDPETKK | Y192 | 0.094 | 98.47 | 447 | 0.041236 | 0.094 | 0.0412 | 1 |

| | | | | | | | | | | |
|---|---|---|---|---|---|---|---|---|---|---|
| P53041 | PPP5C | GYYRRAAsNMALGkF | S105 | 0 | 94.45 | 499 | 0 | 0 | 0 | 1 |
| A2RUS2 | DENND3 | FLLRRIPtLKIRVAS | T781 | 0.016 | 91.25 | 1198 | 0.010928 | 0.016 | 0.0109 | 0.8571 |
| P0DTE5 | UGT2A2 | SKILGRPtTLCETMG | T251 | 0.132 | 96.78 | 536 | 0.090158 | 0.132 | 0.0902 | 1 |
| O94832 | MYO1D | RFEKGRIyTFIGEVV | Y39 | 0.113 | 97.17 | 1006 | 0.049571 | 0.113 | 0.0496 | 1 |
| O94832 | MYO1D | LKQEQEEyQREGIPW | Y423 | 0.092 | 88.58 | 1006 | 0.040359 | 0.183 | 0.0803 | 1 |
| O94832 | MYO1D | EVTkRPLtAATLFKN | T563 | 0.082 | 85.58 | 1006 | 0.056007 | 0.044 | 0.1031 | 1 |
| O94832 | MYO1D | NLASKEPyyVRCIKP | Y585 | 0.12 | 92.93 | 1006 | 0.052642 | 0.12 | 0.0526 | 1 |
| O94832 | MYO1D | LASKEPyyVRCIKPN | Y586 | 0.119 | 93.69 | 1006 | 0.052204 | 0.119 | 0.0522 | 1 |
| Q7Z3S7 | CACNA2D4 | ILLGVVGsDVALREL | S533 | 0.009 | 94.72 | 1137 | 0.007154 | 0.009 | 0.0072 | 1 |
| P49795 | RGS19 | SFDKLMHsPAGRSVF | S97 | 0.069 | 95.58 | 217 | 0.054849 | 0.069 | 0.0548 | 1 |
| P49795 | RGS19 | ARLIyEDyVsILsPk | Y146 | 0.122 | 98.12 | 217 | 0.05352 | 0.122 | 0.0535 | 1 |
| P49795 | RGS19 | SYPRFLSsPtyRALL | S199 | 0.086 | 96.36 | 217 | 0.068363 | 0.086 | 0.099 | 1 |
| P49795 | RGS19 | RFLSsPtyRALLLQG | Y202 | 0.086 | 94.73 | 217 | 0.037727 | 0.086 | 0.0377 | 1 |
| Q15654 | TRIP6 | RIVALDrsFHIGCYK | S424 | 0.148 | 90.01 | 476 | 0.117647 | 0.148 | 0.1176 | 1 |
| Q14653 | IRF3 | EEWEFEVtAFYRGRQ | T207 | 0.029 | 98.15 | 427 | 0.019807 | 0.029 | 0.0198 | 1 |
| Q14653 | IRF3 | sHPLsLtsDQYKAYL | S405 | 0.164 | 91.21 | 427 | 0.130366 | 0.164 | 0.1304 | 1 |
| Q13496 | MTM1 | YITNYRLyLRSLETD | Y71 | 0.104 | 96.2 | 603 | 0.045623 | 0.104 | 0.0456 | 1 |
| Q13496 | MTM1 | RNKDDEkyLDVIRET | Y260 | 0.06 | 98.71 | 603 | 0.026321 | 0.06 | 0.0263 | 1 |
| Q13496 | MTM1 | QISkLTIyDARPSVN | Y277 | 0.145 | 98.89 | 603 | 0.063609 | 0.145 | 0.0636 | 1 |
| Q13496 | MTM1 | GyEsDDAyHNAELFF | Y301 | 0.157 | 97.85 | 603 | 0.068874 | 0.157 | 0.0689 | 1 |
| Q13496 | MTM1 | RQKVTERtVSLWSLI | T495 | 0.119 | 98.64 | 603 | 0.081279 | 0.134 | 0.1169 | 1 |
| P25092 | GUCY2C | FKTYSWStSYVyKNG | T189 | 0.111 | 93.71 | 1073 | 0.075815 | 0.111 | 0.0758 | 1 |
| P25092 | GUCY2C | SWStSYVyKNGTETE | Y193 | 0.033 | 94.62 | 1073 | 0.014477 | 0.033 | 0.0145 | 1 |
| P53350 | PLK1 | SDkYGLGyQLCDNSV | Y425 | 0.022 | 95.57 | 603 | 0.009651 | 0.022 | 0.0097 | 0.9412 |
| P53350 | PLK1 | ILHLsNGsVQINFFQ | S529 | 0.084 | 92.68 | 603 | 0.066773 | 0.084 | 0.0668 | 0.8 |
| P53350 | PLK1 | INFFQDHtkLILCPL | T539 | 0.002 | 94.53 | 603 | 0.001366 | 0.002 | 0.0014 | 1 |
| Q9HDB9 | ERVK-5 | MGQTKSKtKsKYASY | T8 | 0.057 | 67.46 | 667 | 0.038932 | 0.129 | 0.0881 | 1 |
| P17301 | ITGA2 | KLNLQTStSIPNVTE | T101 | 0.003 | 93.11 | 1181 | 0.002049 | 0.003 | 0.002 | 1 |
| P17301 | ITGA2 | HLQYFGRsLDGYGDL | S621 | 0.131 | 98.37 | 1181 | 0.104134 | 0.131 | 0.1041 | 1 |
| P24821 | TNC | KLIPGVEyLVsIIAM | Y1775 | 0.095 | 86.37 | 2201 | 0.041675 | 0.095 | 0.0417 | 1 |
| P24821 | TNC | PGVEyLVsIIAMKGF | S1778 | 0.176 | 85.92 | 2201 | 0.139905 | 0.176 | 0.1399 | 1 |

| | | | | | | | | | | |
|---|---|---|---|---|---|---|---|---|---|---|
| P24821 | TNC | tITAkFTtDLDsPRD | T1885 | 0.002 | 86.71 | 2201 | 0.001366 | 0.002 | 0.0014 | 1 |
| Q96S55 | WRNIP1 | VDTLAYLsDGDARAG | S436 | 0.036 | 85.35 | 665 | 0.028617 | 0.084 | 0.0668 | 1 |
| O00170 | AIP | EGTVLDDsRARGkPM | S53 | 0.029 | 96.18 | 330 | 0.023053 | 0.029 | 0.0231 | 1 |
| O00170 | AIP | VWETIVCtMREGEIA | T79 | 0.084 | 96.61 | 330 | 0.057373 | 0.084 | 0.0574 | 1 |
| O15409 | FOXP2 | WTVDEVEyQKRRSQK | Y580 | 0.189 | 89.53 | 715 | 0.082912 | 0.189 | 0.0829 | 0.8182 |
| Q96PY5 | FMNL2 | RKPLENLsDEDrFMM | S763 | 0.101 | 89.35 | 1086 | 0.080286 | 0.101 | 0.0803 | 1 |
| Q7LBE3 | SLC26A9 | QWKDMIGtAFSLAIV | T336 | 0.209 | 92.79 | 791 | 0.14275 | 0.209 | 0.1427 | 1 |
| Q7LBE3 | SLC26A9 | YLYPLPKsVLGALIA | S432 | 0.024 | 93.96 | 791 | 0.019078 | 0.024 | 0.0191 | 1 |
| P61313 | RPL15 | kFARsLQsVAEErAG | S100 | 0.031 | 97.48 | 204 | 0.024642 | 0.031 | 0.0246 | 0.9 |
| Q6ZR08 | DNAH12 | EKIMNTWyPKVINLF | Y264 | 0.116 | 91.155 | 3092 | 0.050888 | 0.2125 | 0.0932 | 1 |
| Q8IYM9 | TRIM22 | AWILGVHsKISsLNk | S373 | 0.005 | 92.37 | 498 | 0.003975 | 0.005 | 0.004 | 1 |
| Q8NI36 | WDR36 | ISAAMDCsIRTWDLP | S645 | 0.016 | 94.78 | 951 | 0.012719 | 0.016 | 0.0127 | 1 |
| Q9Y375 | NDUFAF1 | MYSYFMFtRGGPYWQ | T243 | 0.115 | 92 | 327 | 0.078547 | 0.115 | 0.0785 | 1 |
| Q8IX12 | CCAR1 | MAIGGHWsPSLDGPD | S505 | 0 | 94.83 | 1150 | 0 | 0 | 0 | 1 |
| Q00059 | TFAM | VssYLRFskEQLPIF | S61 | 0.119 | 97.99 | 246 | 0.094595 | 0.138 | 0.1453 | 1 |
| Q00059 | TFAM | PRsAYNVyVAERFQE | Y165 | 0.1 | 98.05 | 246 | 0.043869 | 0.1 | 0.0439 | 1 |
| Q00059 | TFAM | sDsEkELyIQHAkED | Y200 | 0.122 | 97.69 | 246 | 0.05352 | 0.122 | 0.0535 | 1 |
| O75143 | ATG13 | VTPAYRLsRkQGHEY | S141 | 0.008 | 97.21 | 517 | 0.006359 | 0.008 | 0.0064 | 1 |
| P80217 | IFI35 | VRELLPGsVMLGFAR | S178 | 0.154 | 92.24 | 286 | 0.122417 | 0.154 | 0.1224 | 1 |
| P36915 | GNL1 | rGyDPNRyrLHFERD | Y81 | 0.162 | 74.55 | 607 | 0.071067 | 0.311 | 0.1364 | 1 |
| P36915 | GNL1 | EERSFQDyLGkIHGA | Y155 | 0.21 | 95.69 | 607 | 0.092124 | 0.238 | 0.1044 | 1 |
| Q9BT09 | CNPY3 | sGVKYtksDLRLIEV | S92 | 0.083 | 88.4 | 278 | 0.065978 | 0.082 | 0.1225 | 1 |
| Q9NRC8 | SIRT7 | TLTHMSItRLHEQKL | T153 | 0.054 | 98.69 | 400 | 0.036883 | 0.054 | 0.0369 | 1 |
| Q9NYK1 | TLR7 | PERLSNCsRsLKNLI | S722 | 0.141 | 95.86 | 1049 | 0.112083 | 0.141 | 0.1121 | 1 |
| A8MY62 | LACTBL1 | sGAPNEYtMyRISSI | T74 | 0.036 | 98.61 | 500 | 0.024588 | 0.068 | 0.046 | 1 |
| A8MY62 | LACTBL1 | APNEYtMyRISSISK | Y76 | 0.005 | 98.62 | 500 | 0.002193 | 0.005 | 0.0022 | 1 |
| A8MY62 | LACTBL1 | LILRRMAsQLsGLPR | S136 | 0.004 | 97.32 | 500 | 0.00318 | 0.004 | 0.0032 | 1 |
| A8MY62 | LACTBL1 | RRMAsQLsGLPRRLR | S139 | 0.009 | 94.59 | 500 | 0.007154 | 0.009 | 0.0072 | 1 |
| P43304 | GPD2 | LVERDDFssGTSSRS | S104 | 0.018 | 97.39 | 727 | 0.014308 | 0.018 | 0.0143 | 0.9231 |
| P43304 | GPD2 | KLVGAIVyYDGQHND | Y218 | 0.061 | 93.9 | 727 | 0.02676 | 0.061 | 0.0268 | 1 |
| P43304 | GPD2 | INAtGPFtDsVRKMD | T294 | 0.003 | 96.71 | 727 | 0.002049 | 0.003 | 0.002 | 1 |

| | | | | | | | | | | |
|---|---|---|---|---|---|---|---|---|---|---|
| P43304 | GPD2 | KMTIAGTtDtPtDVt | T355 | 0.012 | 88.7 | 727 | 0.008196 | 0.012 | 0.0082 | 1 |
| P43304 | GPD2 | NEVRNYLsCDVEVRR | S384 | 0.137 | 83.85 | 727 | 0.108903 | 0.137 | 0.1089 | 1 |
| P43304 | GPD2 | RLVSEFPyIEAEVKY | Y532 | 0.031 | 97.9 | 727 | 0.013599 | 0.031 | 0.0136 | 0.9091 |
| P43304 | GPD2 | CtAVDMIsRRTRLAF | S553 | 0.01 | 96.93 | 727 | 0.007949 | 0.01 | 0.0079 | 1 |
| P41279 | MAP3K8 | MRELIEAsLERNPNH | S368 | 0.002 | 98.72 | 467 | 0.00159 | 0.002 | 0.0016 | 1 |
| Q6XYQ8 | SYT10 | KAMDItGssDPyVKV | S398 | 0.114 | 91.32 | 523 | 0.09062 | 0.114 | 0.0906 | 1 |
| Q6XYQ8 | SYT10 | AMDItGssDPyVKVS | S399 | 0 | 93.59 | 523 | 0 | 0 | 0 | 1 |
| Q6XYQ8 | SYT10 | ItGssDPyVKVSLMC | Y402 | 0.137 | 97.95 | 523 | 0.0601 | 0.137 | 0.0601 | 1 |
| Q8TEQ6 | GEMIN5 | RVkDLIVsGDEKGVV | S127 | 0.012 | 98.01 | 1508 | 0.009539 | 0.012 | 0.0095 | 1 |
| Q8TEQ6 | GEMIN5 | FNLCPLQtEDDKQLL | T345 | 0.076 | 92.91 | 1508 | 0.051909 | 0.076 | 0.0519 | 1 |
| Q92889 | ERCC4 | CISMSRYykRPVLLI | Y752 | 0.101 | 86.3 | 916 | 0.044307 | 0.279 | 0.1224 | 0.8 |
| Q92696 | RABGGTA | PLLYEkEtLQYFQTL | T404 | 0.074 | 98.57 | 567 | 0.050543 | 0.074 | 0.0505 | 1 |
| Q06187 | BTK | HNsAGLIsRLKyPVs | S371 | 0.089 | 90.75 | 659 | 0.070747 | 0.089 | 0.0707 | 1 |
| P48052 | CPA2 | SQLLMFPyGyKCTKL | Y316 | 0.2 | 98.59 | 419 | 0.087737 | 0.2 | 0.0877 | 1 |
| P48052 | CPA2 | GGSIDWSyDyGIKyS | Y369 | 0.231 | 98.77 | 419 | 0.101336 | 0.231 | 0.1013 | 0.8571 |
| P48052 | CPA2 | SyDyGIKySFAFELR | Y375 | 0.019 | 98.79 | 419 | 0.008335 | 0.019 | 0.0083 | 1 |
| Q5FYB0 | ARSJ | PLRGSKGtYWEGGIR | T347 | 0.042 | 97.52 | 599 | 0.028687 | 0.042 | 0.0287 | 0.9 |
| Q96PE1 | ADGRA2 | ASYLGNDtRIRWYHN | T277 | 0.083 | 87.41 | 1338 | 0.05669 | 0.083 | 0.0567 | 1 |
| Q5T9A4 | ATAD3B | VRDIAIAtRNtKKNR | T335 | 0.092 | 91.17 | 648 | 0.062837 | 0.092 | 0.0628 | 1 |
| Q8TD26 | CHD6 | LRVRMLyyLKAEILG | Y1238 | 0.053 | 87.95 | 2715 | 0.02325 | 0.053 | 0.0232 | 0.8 |
| O15146 | MUSK | NKHGEKFstAKAAAT | S293 | 0.143 | 75.25 | 869 | 0.113673 | 0.143 | 0.1137 | 1 |
| Q8IW93 | ARHGEF19 | VENILKRtAQGsEDE | T533 | 0.019 | 94.6 | 802 | 0.012977 | 0.068 | 0.0464 | 1 |
| P00439 | PAH | FTHLDKRsLPALtNI | S87 | 0.034 | 78.01 | 452 | 0.027027 | 0.034 | 0.027 | 0.8889 |
| Q16849 | PTPRN | RHPDFLPyDHARIKL | Y746 | 0.145 | 98.53 | 979 | 0.063609 | 0.145 | 0.0636 | 0.8667 |
| P05107 | ITGB2 | NPRRVECsGRGRCRC | S591 | 0.016 | 92.67 | 769 | 0.012719 | 0.016 | 0.0127 | 0.8889 |
| P32247 | BRS3 | AGCVWIVsMIFALPE | S175 | 0 | 98.66 | 399 | 0 | 0 | 0 | 0.9167 |
| P32247 | BRS3 | NMTFEsCtSYPVSKK | T204 | 0.204 | 91.48 | 399 | 0.139335 | 0.204 | 0.1393 | 0.875 |
| Q6P2Q9 | PRPF8 | LFFMRGItPLLERWL | T645 | 0.069 | 93.47 | 2335 | 0.047128 | 0.069 | 0.0471 | 1 |
| Q15018 | ABRAXAS2 | ASkVNEEsLDRILKD | S79 | 0.14 | 93.17 | 415 | 0.111288 | 0.14 | 0.1113 | 1 |
| Q4KMG0 | CDON | GDVKyVtyMVNVLEH | Y304 | 0.171 | 91.06 | 1287 | 0.075015 | 0.171 | 0.075 | 1 |
| Q8WTV0 | SCARB1 | QVRERGPyVYREFRH | Y92 | 0.005 | 98.34 | 552 | 0.002193 | 0.005 | 0.0022 | 1 |

| | | | | | | | | | | |
|---|---|---|---|---|---|---|---|---|---|---|
| P20813 | CYP2B6 | WKVLRRFsVTTMRDF | S128 | 0 | 95.21 | 491 | 0 | 0 | 0 | 0.8889 |
| Q9Y458 | TBX22 | GLDPGKQyHVAIDVV | Y138 | 0.087 | 93.99 | 520 | 0.038166 | 0.087 | 0.0382 | 1 |
| A2RU49 | HYKK | TEYVLKIsNTKASKN | S67 | 0.023 | 97.45 | 373 | 0.018283 | 0.023 | 0.0183 | 1 |
| Q9BVK2 | ALG8 | QWYYEATsEWTLDYP | S61 | 0.123 | 94.03 | 526 | 0.097775 | 0.123 | 0.0978 | 1 |
| Q8ND90 | PNMA1 | YKRLTLFsGRDIPGP | S165 | 0.031 | 92.6 | 353 | 0.024642 | 0.031 | 0.0246 | 1 |
| Q8N136 | DAW1 | PPGIMLEyEKHGELK | Y19 | 0.296 | 94.38 | 415 | 0.129851 | 0.296 | 0.1299 | 1 |
| Q8N136 | DAW1 | YKGNIVItGSKDNTC | T404 | 0 | 98.48 | 415 | 0 | 0 | 0 | 1 |
| O95865 | DDAH2 | GGGDLPNsQEALQKL | S245 | 0.005 | 95.77 | 285 | 0.003975 | 0.005 | 0.004 | 0.875 |
| Q8NEC7 | GSTCD | FVTCPCCyGFIQNTs | Y541 | 0.011 | 96.39 | 633 | 0.004826 | 0.011 | 0.0048 | 1 |
| Q92615 | LARP4B | SQMDSDQyVPITTVA | Y188 | 0.284 | 92.04 | 738 | 0.124587 | 0.284 | 0.1246 | 1 |
| Q92615 | LARP4B | LREIsEStPVEEVEA | T247 | 0.093 | 88.82 | 738 | 0.06352 | 0.093 | 0.0635 | 1 |
| Q6P2C8 | MED27 | kLDIWsksNyQVFQK | S234 | 0.057 | 97.14 | 311 | 0.04531 | 0.057 | 0.0453 | 1 |
| P58872 | RHBDL3 | RLIRHVAyEtLPREI | Y141 | 0.264 | 85.6 | 404 | 0.115813 | 0.264 | 0.1158 | 1 |
| Q13336 | SLC14A1 | PEENRIFyLQAKKRM | Y377 | 0.109 | 97.42 | 389 | 0.047817 | 0.109 | 0.0478 | 1 |
| Q9Y5J1 | UTP18 | HVHTMDFsPRSGyFA | S530 | 0 | 94.72 | 556 | 0 | 0.04 | 0.0318 | 1 |
| Q9Y5J1 | UTP18 | DFsPRSGyFALGNEk | Y535 | 0.19 | 94.2 | 556 | 0.08335 | 0.19 | 0.0833 | 1 |
| Q6YN16 | HSDL2 | PHPkLLGtIytAAEE | T53 | 0.09 | 94.26 | 418 | 0.061471 | 0.171 | 0.1158 | 1 |
| O60784 | TOM1 | LLGNPFssPVGQRIE | S11 | 0.154 | 96.2 | 492 | 0.122417 | 0.154 | 0.1224 | 1 |
| O75360 | PROP1 | RESLARDtGLsEARI | T106 | 0.128 | 97.95 | 226 | 0.087426 | 0.128 | 0.0874 | 1 |
| Q8WY07 | SLC7A3 | VDLMsIGtLLAYSLV | T414 | 0.026 | 93.78 | 619 | 0.017758 | 0.026 | 0.0178 | 0.8 |
| P62241 | RPS8 | LGRPAANtkIGPRRI | T36 | 0.032 | 97.97 | 208 | 0.021856 | 0.032 | 0.0219 | 1 |
| P62241 | RPS8 | sECCTRKtRIIDVVy | T76 | 0.028 | 97.31 | 208 | 0.019124 | 0.052 | 0.0365 | 1 |
| P62241 | RPS8 | NCIVLIDstPyrQWy | S106 | 0.031 | 96.27 | 208 | 0.024642 | 0.031 | 0.0246 | 1 |
| P62241 | RPS8 | VLIDstPyrQWyESH | Y109 | 0.004 | 95.85 | 208 | 0.001755 | 0.004 | 0.0018 | 1 |
| P62241 | RPS8 | stPyrQWyESHyALP | Y113 | 0.075 | 94 | 208 | 0.032901 | 0.118 | 0.0439 | 1 |
| P62241 | RPS8 | rQWyESHyALPLGRK | Y117 | 0.154 | 92.4 | 208 | 0.067558 | 0.28 | 0.116 | 1 |
| P62241 | RPS8 | EGkELEFyLrKIKAR | Y198 | 0.208 | 96.19 | 208 | 0.091247 | 0.208 | 0.0912 | 1 |
| Q9UKK3 | PARP4 | DQMTRLCsPSCHSVS | S1048 | 0.145 | 83.05 | 1724 | 0.115263 | 0.145 | 0.1153 | 1 |
| Q9UKK3 | PARP4 | NSLITQFtsFVAVEK | T1173 | 0.004 | 91.49 | 1724 | 0.002732 | 0.21 | 0.1434 | 1 |
| P52333 | JAK3 | DASRLLLySsQICkG | Y929 | 0 | 97.78 | 1124 | 0 | 0 | 0 | 1 |
| Q9Y2H6 | FNDC3A | IANRTkNsLtLQWKA | S386 | 0.064 | 90.34 | 1198 | 0.050875 | 0.064 | 0.0509 | 1 |

| | | | | | | | | | | |
|---|---|---|---|---|---|---|---|---|---|---|
| Q9Y2H6 | FNDC3A | LLPGKtYsFRLRAAN | S732 | 0.101 | 94.75 | 1198 | 0.080286 | 0.101 | 0.0803 | 1 |
| Q9Y2H6 | FNDC3A | KMGFGPFsEKCDITT | S747 | 0.1 | 95.53 | 1198 | 0.079491 | 0.1 | 0.0795 | 1 |
| Q9Y2H6 | FNDC3A | sLGAGPFsHMIKLKT | S940 | 0.12 | 93.37 | 1198 | 0.09539 | 0.12 | 0.0954 | 1 |
| Q9UBQ7 | GRHPR | ANLKVIStMSVGIDH | T80 | 0 | 98.66 | 328 | 0 | 0 | 0 | 1 |
| Q9UBQ7 | GRHPR | HRTRNtMsLLAANNL | S306 | 0.005 | 98.4 | 328 | 0.003975 | 0.005 | 0.004 | 1 |
| P22102 | GART | WRIPRVFsWLQQEGH | S714 | 0.066 | 93.09 | 1010 | 0.052464 | 0.1 | 0.0795 | 0.875 |
| Q96G46 | DUS3L | LDIRGkLyLAPLTTC | Y308 | 0.044 | 98.33 | 650 | 0.019302 | 0.044 | 0.0193 | 1 |
| Q96G46 | DUS3L | QRHWDIsssERLDIL | S543 | 0.064 | 97.6 | 650 | 0.050875 | 0.064 | 0.0509 | 1 |
| Q9BVI0 | PHF20 | AVNkDGTyTVKFYDG | Y116 | 0.1 | 95.52 | 1012 | 0.043869 | 0.1 | 0.0439 | 0.9091 |
| Q92520 | FAM3C | LEDNVLMsGVKNNVG | S94 | 0.125 | 98.74 | 227 | 0.099364 | 0.125 | 0.0994 | 1 |
| Q15561 | TEAD4 | GVSSQYEsPENMIIT | S322 | 0.016 | 96 | 434 | 0.012719 | 0.016 | 0.0127 | 1 |
| P16870 | CPE | YWEDNKNsLISyLEQ | S364 | 0 | 98.75 | 476 | 0 | 0 | 0 | 1 |
| P16870 | CPE | GIDHDVtsAkDGDYW | S404 | 0.037 | 98.64 | 476 | 0.029412 | 0.037 | 0.0294 | 1 |
| O94822 | LTN1 | AFCkDEItSVLQDHL | T191 | 0.001 | 93.38 | 1766 | 0.000683 | 0.001 | 0.0007 | 1 |
| Q6ZMP0 | THSD4 | GKyEGGGtMFTYkRP | T480 | 0.111 | 91.6 | 1018 | 0.075815 | 0.111 | 0.0758 | 1 |
| Q8IUG5 | MYO18B | MAHSRSAtRFSMVMS | T715 | 0.164 | 91.4 | 2567 | 0.112014 | 0.194 | 0.1325 | 0.875 |
| Q9UIQ6 | LNPEP | TAVVPLRyELSLHPN | Y177 | 0.009 | 98.92 | 1025 | 0.003948 | 0.009 | 0.0039 | 1 |
| Q9UIQ6 | LNPEP | DEQYTALsNMPkkSS | S329 | 0.016 | 98.91 | 1025 | 0.012719 | 0.016 | 0.0127 | 1 |
| Q9UHV5 | RAPGEFL1 | HSYVTIRsRLSASVQ | S332 | 0.102 | 91.11 | 662 | 0.081081 | 0.102 | 0.0811 | 0.875 |
| Q9UHV5 | RAPGEFL1 | GERGRREtANLELLL | T462 | 0.015 | 88.3 | 662 | 0.010245 | 0.015 | 0.0102 | 1 |
| Q12836 | ZP4 | QIAKDKNyGSyyGVG | Y327 | 0.143 | 90.86 | 540 | 0.062732 | 0.143 | 0.0627 | 0.9167 |
| Q8WYR4 | RSPH1 | YLYAETGsKYVGTWV | S135 | 0.126 | 96.84 | 309 | 0.100159 | 0.148 | 0.1176 | 0.8 |
| Q8NF50 | DOCK8 | FFELLVKsMAQHVHN | S993 | 0 | 91.87 | 2099 | 0 | 0.013 | 0.0755 | 1 |
| P15498 | VAV1 | KMEVFQEyYGLPPPP | Y603 | 0.024 | 93.62 | 845 | 0.010528 | 0.024 | 0.0105 | 1 |
| P15498 | VAV1 | LTELVEFyQQNSLkD | Y745 | 0.053 | 96.37 | 845 | 0.02325 | 0.053 | 0.0232 | 0.9231 |
| Q86SJ6 | DSG4 | SVVDREItPLFLIYC | T120 | 0.162 | 92.62 | 1040 | 0.110648 | 0.162 | 0.1106 | 1 |
| O00522 | KRIT1 | TNRILHEykNLSTSE | Y605 | 0 | 96.75 | 736 | 0 | 0 | 0 | 1 |
| Q16531 | DDB1 | kDLLFILtAKYNACI | T81 | 0.017 | 96.99 | 1140 | 0.011611 | 0.017 | 0.0116 | 1 |
| Q16531 | DDB1 | APTICFVyQDPQGRH | Y182 | 0.238 | 96.53 | 1140 | 0.104407 | 0.238 | 0.1044 | 1 |
| Q16531 | DDB1 | QGRHVkTyEVsLREk | Y193 | 0.241 | 98.03 | 1140 | 0.105723 | 0.241 | 0.1057 | 1 |
| Q16531 | DDB1 | VGSRLGDsQLVKLNV | S331 | 0.031 | 98.12 | 1140 | 0.024642 | 0.041 | 0.0349 | 1 |

| | | | | | | | | | | |
|---|---|---|---|---|---|---|---|---|---|---|
| Q16531 | DDB1 | VTLGTQPtVLRtFRS | T636 | 0.021 | 92.83 | 1140 | 0.014343 | 0.021 | 0.0143 | 1 |
| Q16531 | DDB1 | SNHKLVFsNVNLKEV | S669 | 0.137 | 80.68 | 1140 | 0.108903 | 0.159 | 0.1264 | 0.8333 |
| Q16531 | DDB1 | GTTALRPsASTQALS | S755 | 0.016 | 96.78 | 1140 | 0.012719 | 0.016 | 0.0127 | 1 |
| Q16531 | DDB1 | LGkDPNTyFIVGTAM | Y828 | 0 | 98.17 | 1140 | 0 | 0 | 0 | 1 |
| Q6P9H5 | GIMAP6 | TRPVTKTsQRRSREW | S79 | 0.097 | 96.07 | 292 | 0.077107 | 0.097 | 0.0771 | 0.8 |
| Q9NZ32 | ACTR10 | NTEELySyLKEFIHI | Y67 | 0.1 | 94.13 | 417 | 0.043869 | 0.1 | 0.0439 | 1 |
| Q9H6E5 | TUT1 | WAQGRGLsGSGPLLS | S423 | 0.147 | 94.81 | 874 | 0.116852 | 0.147 | 0.1169 | 1 |
| Q9NSB8 | HOMER2 | KKNWMPAsKQAVTVS | S28 | 0.083 | 95.93 | 354 | 0.065978 | 0.083 | 0.066 | 0.875 |
| Q6P587 | FAHD1 | EEGDIILtGtPkGVG | T187 | 0.013 | 98.75 | 224 | 0.008879 | 0.013 | 0.0089 | 1 |
| O60882 | MMP20 | FVGDEYysyDERKRK | S416 | 0.01 | 97.99 | 483 | 0.007949 | 0.01 | 0.0079 | 0.8333 |
| O60882 | MMP20 | YFFSGPKtyKyDtEK | T461 | 0.01 | 94.97 | 483 | 0.00683 | 0.01 | 0.0068 | 0.9 |
| O60882 | MMP20 | SGPKtyKyDtEKEDV | Y464 | 0.077 | 97.63 | 483 | 0.033779 | 0.077 | 0.0338 | 0.8571 |
| Q9H1H9 | KIF13A | GEKVSkVsLVDLAGS | S249 | 0.042 | 95.07 | 1805 | 0.033386 | 0.042 | 0.0334 | 1 |
| Q6Q0C0 | TRAF7 | VFKDPVItTCGHtFC | T144 | 0.035 | 90.91 | 670 | 0.023905 | 0.035 | 0.0239 | 1 |
| Q9H4L7 | SMARCAD1 | AEkLkEMsQLMLKEP | S801 | 0.044 | 88.36 | 1026 | 0.034976 | 0.044 | 0.035 | 1 |
| Q6P461 | ACSM6 | ACVRLGItFVPGSPQ | T137 | 0.057 | 96.52 | 480 | 0.038932 | 0.057 | 0.0389 | 1 |
| Q6P461 | ACSM6 | GMGFSQAsRRWMDLQ | S251 | 0 | 93.1 | 480 | 0 | 0 | 0 | 1 |
| Q96Q42 | ALS2 | ALLPSLRtEVWtWGk | T526 | 0.045 | 93.46 | 1657 | 0.030736 | 0.09 | 0.0615 | 1 |
| Q96Q42 | ALS2 | SLRtEVWtWGkGKEG | T530 | 0 | 97.84 | 1657 | 0 | 0 | 0 | 1 |
| Q9NZ08 | ERAP1 | GELRILAsTQFEPtA | S179 | 0 | 98.85 | 941 | 0 | 0 | 0 | 1 |
| Q9NZ08 | ERAP1 | AsTQFEPtAARMAFP | T185 | 0.132 | 98.65 | 941 | 0.090158 | 0.132 | 0.0902 | 0.9 |
| Q9NZ08 | ERAP1 | TKSGVKVsVyAVPDK | S268 | 0.021 | 98.92 | 941 | 0.016693 | 0.021 | 0.0167 | 1 |
| Q9NZ08 | ERAP1 | SGVKVsVyAVPDKIN | Y270 | 0.288 | 98.92 | 941 | 0.126341 | 0.288 | 0.1263 | 1 |
| Q9NZ08 | ERAP1 | FkSGIVQyLQKHSYK | Y464 | 0.03 | 98.48 | 941 | 0.013161 | 0.03 | 0.0132 | 1 |
| Q9NZ08 | ERAP1 | QKHSYKNtKNEDLWD | T473 | 0.01 | 98.77 | 941 | 0.00683 | 0.01 | 0.0068 | 1 |
| Q9NZ08 | ERAP1 | NVGMNGYyIVHYEDD | Y608 | 0.007 | 98.83 | 941 | 0.003071 | 0.007 | 0.0031 | 0.875 |
| O75078 | ADAM11 | VQDDLLEtLARLMVY | T308 | 0.005 | 95.08 | 769 | 0.003415 | 0.005 | 0.0034 | 1 |
| Q9HAR2 | ADGRL3 | AVVRRELsCESYPIE | S35 | 0.062 | 94.48 | 1447 | 0.049285 | 0.062 | 0.0493 | 1 |
| P49863 | GZMK | LCNSQSyyNGDPFIT | Y187 | 0.048 | 93.72 | 264 | 0.021057 | 0.048 | 0.0211 | 0.8125 |
| O14905 | WNT9B | QLSPFREtGQVLKLR | T234 | 0.048 | 98.52 | 357 | 0.032785 | 0.095 | 0.0649 | 1 |
| O95299 | NDUFA10 | SKRLTERsRVItVDG | S57 | 0.092 | 89.17 | 355 | 0.073132 | 0.066 | 0.0945 | 1 |

| | | | | | | | | | | |
|---|---|---|---|---|---|---|---|---|---|---|
| O95299 | NDUFA10 | TERsRVItVDGNICT | T61 | 0 | 95.65 | 355 | 0 | 0 | 0 | 0.9 |
| O95299 | NDUFA10 | EMKItsAyLQDIENA | Y233 | 0.118 | 92.17 | 355 | 0.051765 | 0.118 | 0.0518 | 1 |
| Q504Q3 | PAN2 | GIPFIDDyISTQEQV | Y1028 | 0.16 | 92.72 | 1202 | 0.07019 | 0.261 | 0.1001 | 0.9167 |
| Q6UXZ3 | CD300LD | SLTVQCAyGSGWETY | Y41 | 0.171 | 97.66 | 194 | 0.075015 | 0.171 | 0.075 | 0.9375 |
| Q04912 | MST1R | GVVYHGEyIDQAQNR | Y1101 | 0.309 | 93.71 | 1400 | 0.135554 | 0.309 | 0.1356 | 1 |
| Q04912 | MST1R | QVARGMEyLAEQKFV | Y1198 | 0.328 | 96.19 | 1400 | 0.143889 | 0.328 | 0.1439 | 0.8889 |
| P43630 | KIR3DL2 | SWSSYDIyHLsREGE | Y253 | 0 | 95.51 | 455 | 0 | 0 | 0 | 1 |
| P43630 | KIR3DL2 | SYDIyHLsREGEAHE | S256 | 0.181 | 93.03 | 455 | 0.14388 | 0.181 | 0.1439 | 0.875 |
| O43314 | PPIP5K2 | QIVVGICsMAKKsKS | S50 | 0.005 | 97.47 | 1243 | 0.003975 | 0.005 | 0.004 | 1 |
| O43314 | PPIP5K2 | ICsMAKKsKSKPMkE | S55 | 0.066 | 95.74 | 1243 | 0.052464 | 0.126 | 0.1023 | 1 |
| O43314 | PPIP5K2 | FVKNSMkyyDDCAKI | Y334 | 0.006 | 85.12 | 1243 | 0.002632 | 0.006 | 0.0026 | 1 |
| O43314 | PPIP5K2 | VKNSMkyyDDCAKIL | Y335 | 0.031 | 88.62 | 1243 | 0.013599 | 0.062 | 0.0272 | 1 |
| Q16658 | FSCN1 | HPQVNIysVtRKRYA | S146 | 0.001 | 97.1 | 493 | 0.000795 | 0.001 | 0.0008 | 1 |
| Q16658 | FSCN1 | AANERNVsTRQGMDL | S274 | 0 | 98.18 | 493 | 0 | 0 | 0 | 1 |
| Q16658 | FSCN1 | FRTHTGkyWTLTATG | Y314 | 0.202 | 97.67 | 493 | 0.088614 | 0.202 | 0.0886 | 1 |
| Q16658 | FSCN1 | IkDSTGKyWTVGSDS | Y432 | 0.269 | 98.38 | 493 | 0.118006 | 0.324 | 0.1421 | 1 |
| Q7L591 | DOK3 | RKVWALLyAGGPSGV | Y89 | 0.237 | 95.08 | 496 | 0.103968 | 0.298 | 0.1307 | 1 |
| Q08211 | DHX9 | LDANDELtPLGRILA | T849 | 0.023 | 95.66 | 1270 | 0.015709 | 0.085 | 0.0388 | 1 |
| Q08211 | DHX9 | NALIHkSsVNCPFss | S1026 | 0.023 | 93.61 | 1270 | 0.018283 | 0.023 | 0.0183 | 1 |
| Q86U38 | NOP9 | VLDAILTsPsVTRKL | S533 | 0.024 | 96.28 | 636 | 0.019078 | 0.024 | 0.0191 | 1 |
| Q8NGA6 | OR10H5 | yLFLCALsItEILYT | S67 | 0.015 | 96.09 | 315 | 0.011924 | 0.015 | 0.0119 | 1 |
| Q8NGA6 | OR10H5 | FLCALsItEILYTVA | T69 | 0.009 | 96.25 | 315 | 0.006147 | 0.009 | 0.0061 | 0.8182 |
| Q00537 | CDK17 | LGEGtyAtVykGRSk | T205 | 0.1 | 85.87 | 523 | 0.068301 | 0.1 | 0.0683 | 1 |
| Q6UXG2 | ELAPOR1 | PCAEGRYsLGTGIRF | S117 | 0.003 | 87.68 | 1013 | 0.002385 | 0.003 | 0.0024 | 1 |
| Q9UJ70 | NAGK | DPLVPLRsLGLsLsG | S70 | 0.008 | 98.42 | 344 | 0.006359 | 0.008 | 0.0064 | 1 |
| Q9UJ70 | NAGK | PLRsLGLsLsGGDQE | S74 | 0 | 96.06 | 344 | 0 | 0 | 0 | 0.9333 |
| Q9UJ70 | NAGK | RsLGLsLsGGDQEDA | S76 | 0.148 | 88.16 | 344 | 0.117647 | 0.148 | 0.1176 | 1 |
| Q09472 | EP300 | SAKRLPstRLGTFLE | T1296 | 0.098 | 97.53 | 2414 | 0.066935 | 0.098 | 0.0669 | 1 |
| Q8IV35 | WDR49 | ETRLLTGsTDGTVKI | S211 | 0 | 94.95 | 697 | 0 | 0 | 0 | 1 |
| Q01064 | PDE1B | IDKPKALsLLLHAAD | S363 | 0.05 | 98.4 | 536 | 0.039746 | 0.05 | 0.0397 | 0.8889 |
| O60245 | PCDH7 | GRNGIERyELLQEPG | Y180 | 0.168 | 93.03 | 1069 | 0.073699 | 0.168 | 0.0737 | 1 |

| | | | | | | | | | | |
|---|---|---|---|---|---|---|---|---|---|---|
| Q9Y6M7 | SLC4A7 | HIPHDLFtEMDELCy | T114 | 0.001 | 91.18 | 1214 | 0.000683 | 0.001 | 0.0007 | 0.9091 |
| Q9Y6M7 | SLC4A7 | PAGKAPQyHEIGRsI | Y481 | 0.129 | 91.18 | 1214 | 0.05659 | 0.129 | 0.0566 | 0.9231 |
| Q9UBF8 | PI4KB | CRQsINFsLQCALLL | S209 | 0.037 | 95.28 | 816 | 0.029412 | 0.188 | 0.1494 | 1 |
| Q9UKJ1 | PILRA | QWQSIEGtKLSITQA | T145 | 0.003 | 98.62 | 303 | 0.002049 | 0.003 | 0.002 | 0.9 |
| Q8TCB0 | IFI44 | SFFNSVRsVFQGHVt | S208 | 0.001 | 96.41 | 444 | 0.000795 | 0.001 | 0.0008 | 1 |
| Q8TCB0 | IFI44 | sVFQGHVtHQALVGT | T215 | 0.106 | 91.75 | 444 | 0.072399 | 0.106 | 0.0724 | 1 |
| Q8TCB0 | IFI44 | DISVVSNySSEWELD | Y392 | 0.088 | 93.97 | 444 | 0.038604 | 0.088 | 0.0386 | 1 |
| P13010 | XRCC5 | EEQMkYksEGkCFsV | S335 | 0.166 | 88.67 | 732 | 0.131956 | 0.166 | 0.132 | 1 |
| P13010 | XRCC5 | PAEVttksQIPLskI | S526 | 0.036 | 94.27 | 732 | 0.028617 | 0.036 | 0.0286 | 1 |
| Q16348 | SLC15A2 | EFCERFSyyGMKAVL | Y60 | 0.003 | 94.91 | 729 | 0.001316 | 0.003 | 0.0013 | 1 |
| Q16348 | SLC15A2 | HAEERTRyFSVFyLS | Y183 | 0.034 | 81.73 | 729 | 0.014915 | 0.034 | 0.0149 | 1 |
| Q16348 | SLC15A2 | LTRVLFLyIPLPMFW | Y306 | 0.131 | 94.7 | 729 | 0.057468 | 0.262 | 0.1149 | 1 |
| Q16348 | SLC15A2 | ALLDQQGsRWTLQAI | S321 | 0 | 95.64 | 729 | 0 | 0 | 0 | 1 |
| Q16348 | SLC15A2 | SIKSFQKtPHySKLH | T449 | 0.052 | 85.32 | 729 | 0.035517 | 0.052 | 0.0355 | 1 |
| Q16348 | SLC15A2 | DFHFHLKyHNLSLyT | Y470 | 0.122 | 89.53 | 729 | 0.05352 | 0.241 | 0.1057 | 1 |
| Q01518 | CAP1 | MGkVPtIsINkTDGC | S409 | 0.098 | 98.19 | 475 | 0.077902 | 0.098 | 0.0779 | 1 |
| Q01518 | CAP1 | DGCHAyLskNsLDCE | S421 | 0.104 | 95.65 | 475 | 0.082671 | 0.104 | 0.0827 | 1 |
| Q01518 | CAP1 | HAyLskNsLDCEIVs | S424 | 0 | 94.17 | 475 | 0 | 0 | 0 | 1 |
| Q9UHN6 | CEMIP2 | AKSLLQGtIQMIQER | T326 | 0 | 96.2 | 1383 | 0 | 0 | 0 | 1 |
| Q9UHN6 | CEMIP2 | TRSTFKKyVPTPDRy | Y888 | 0.042 | 97.97 | 1383 | 0.018425 | 0.042 | 0.0184 | 1 |
| Q9UHN6 | CEMIP2 | MKNSWQItPRNNISL | T910 | 0.011 | 91.26 | 1383 | 0.007513 | 0.011 | 0.0075 | 1 |
| Q9UHN6 | CEMIP2 | VVMLEKGytIHWNGP | Y1044 | 0.005 | 98.46 | 1383 | 0.002193 | 0.005 | 0.0022 | 1 |
| Q9UHN6 | CEMIP2 | VMLEKGytIHWNGPA | T1045 | 0 | 98.26 | 1383 | 0 | 0 | 0 | 1 |
| Q9UHN6 | CEMIP2 | KFYFDsStGLLFLyL | T1125 | 0.042 | 94.94 | 1383 | 0.028687 | 0.084 | 0.0574 | 1 |
| Q9UHN6 | CEMIP2 | StGLLFLyLKAKSHR | Y1131 | 0.168 | 96.89 | 1383 | 0.073699 | 0.294 | 0.129 | 1 |
| Q9UHN6 | CEMIP2 | KtGIPPRsIVLLSTR | S1285 | 0.054 | 95.91 | 1383 | 0.042925 | 0.097 | 0.0783 | 1 |
| Q86YV0 | RASAL3 | LPPARRLsLRLRGLG | S367 | 0.162 | 95.69 | 1011 | 0.128776 | 0.162 | 0.1288 | 1 |
| Q86YV0 | RASAL3 | ATkAIDEyMKLVAQD | Y512 | 0.132 | 94.3 | 1011 | 0.057907 | 0.132 | 0.0579 | 1 |
| Q16637 | SMN1 | RETCVVVyTGyGNRE | Y127 | 0.077 | 96.65 | 294 | 0.033779 | 0.077 | 0.0338 | 0.9286 |
| Q86YR7 | MCF2L2 | KWDILGKsLEFHRQL | S426 | 0.029 | 89.2 | 1114 | 0.023053 | 0.102 | 0.0811 | 1 |
| P22626 | HNRNPA2B1 | kLFIGGLsFEtTEEs | S29 | 0.169 | 95.92 | 353 | 0.134341 | 0.169 | 0.1343 | 1 |

| | | | | | | | | | | |
|---|---|---|---|---|---|---|---|---|---|---|
| P22626 | HNRNPA2B1 | IGGLsFEtTEEsLrN | T32 | 0.002 | 95.66 | 353 | 0.001366 | 0.002 | 0.0014 | 1 |
| Q5UE93 | PIK3R6 | VERLEEIyCSLLGPA | Y255 | 0.079 | 92.79 | 754 | 0.034656 | 0.276 | 0.1211 | 1 |
| O60506 | SYNCRIP | VtTGQRKyGGPPPDS | Y144 | 0.047 | 96.12 | 623 | 0.020618 | 0.047 | 0.0206 | 1 |
| O60506 | SYNCRIP | rGFCFLEyEDHkTAA | Y293 | 0.055 | 94.03 | 623 | 0.024128 | 0.055 | 0.0241 | 1 |
| P47972 | NPTX2 | KVSLPLRtNYLYGKI | T233 | 0.131 | 96.96 | 431 | 0.089475 | 0.131 | 0.0895 | 1 |
| Q6ZW49 | PAXIP1 | TRTVKFLtAIsVVKH | T916 | 0.067 | 95.33 | 1069 | 0.045762 | 0.124 | 0.0847 | 0.9 |
| Q6ZW49 | PAXIP1 | VKFLtAIsVVKHIVT | S919 | 0.142 | 95.51 | 1069 | 0.112878 | 0.142 | 0.1129 | 1 |
| Q06187 | BTK | HEKLVQLyGVCTkQR | Y461 | 0.216 | 90.76 | 659 | 0.094756 | 0.216 | 0.0948 | 1 |
| P49961 | ENTPD1 | ALPENVKyGIVLDAG | Y49 | 0.257 | 97.37 | 510 | 0.112742 | 0.257 | 0.1127 | 1 |
| P49961 | ENTPD1 | SHTSLYIyKWPAEKE | Y65 | 0.018 | 97.73 | 510 | 0.007896 | 0.018 | 0.0079 | 1 |
| P22897 | MRC1 | LECLRMGsSLVSIES | S1268 | 0.11 | 90.2 | 1456 | 0.087441 | 0.127 | 0.1114 | 1 |
| P60893 | GPR85 | VVGNLLIsILLVKDK | S42 | 0.161 | 96.45 | 370 | 0.127981 | 0.161 | 0.128 | 1 |
| P23109 | AMPD1 | LLRFIKksyQIDADR | S322 | 0.079 | 94.63 | 780 | 0.062798 | 0.141 | 0.1121 | 1 |
| Q15223 | NECTIN1 | SANGKPPsVVSWETR | S181 | 0.082 | 97.36 | 517 | 0.065183 | 0.082 | 0.0652 | 1 |
| Q15223 | NECTIN1 | TVTVISRyRLVPSRE | Y211 | 0.122 | 97.8 | 517 | 0.05352 | 0.122 | 0.0535 | 1 |
| P23415 | GLRA1 | HyNTGKFtCIEARFH | T236 | 0.037 | 97.4 | 457 | 0.025272 | 0.037 | 0.0253 | 1 |
| Q92769 | HDAC2 | TkYHSDEyIkFLRSI | Y73 | 0 | 98.3 | 488 | 0 | 0 | 0 | 1 |
| Q8NI36 | WDR36 | ETELRSLsPDCGGSI | S863 | 0.076 | 87.83 | 951 | 0.060414 | 0.076 | 0.0604 | 1 |
| O14569 | CYB561D2 | GYLLGsAsLLLGMCS | S173 | 0.1 | 98.61 | 222 | 0.079491 | 0.1 | 0.0795 | 1 |
| Q9NR30 | DDX21 | KKEAQELsQNsAIkQ | S455 | 0.021 | 93.72 | 783 | 0.016693 | 0.021 | 0.0167 | 1 |
| Q9NR30 | DDX21 | AIkQDAQsLHGDIPQ | S466 | 0.158 | 93.52 | 783 | 0.125597 | 0.158 | 0.1256 | 1 |
| Q9NR30 | DDX21 | PPKDVESyIHRSGRT | Y521 | 0.001 | 88.65 | 783 | 0.000439 | 0.001 | 0.0004 | 1 |
| Q9NR30 | DDX21 | TGVCICFyQHKEEyQ | Y541 | 0.106 | 94.88 | 783 | 0.046501 | 0.106 | 0.0465 | 1 |
| O43683 | BUB1 | MKLFPkGtIFTAKCE | T960 | 0.026 | 96.49 | 1085 | 0.017758 | 0.157 | 0.1072 | 1 |
| Q9NXS3 | KLHL28 | CVLDQKVyVIGGIAT | Y334 | 0.216 | 98.17 | 571 | 0.094756 | 0.216 | 0.0948 | 1 |
| Q9NXS3 | KLHL28 | VVLAGELyALGGYDG | Y389 | 0.088 | 98.65 | 571 | 0.038604 | 0.088 | 0.0386 | 1 |
| Q9NXS3 | KLHL28 | YLQSVEKyIPKIRkW | Y406 | 0.135 | 98.17 | 571 | 0.059223 | 0.135 | 0.0592 | 1 |
| Q96L92 | SNX27 | RVKKNsttDQVYQAI | T297 | 0.074 | 97.26 | 541 | 0.050543 | 0.074 | 0.0505 | 1 |
| Q96L92 | SNX27 | PRWVKIFtPYFNYMH | T504 | 0.022 | 92.37 | 541 | 0.015026 | 0.022 | 0.015 | 1 |
| Q15858 | SCN9A | EYADKIFtYIFILEM | T1234 | 0.05 | 87.28 | 1988 | 0.034151 | 0.05 | 0.0342 | 1 |
| Q15858 | SCN9A | IMyAAVDsVNVDKQP | S1418 | 0.049 | 92.29 | 1988 | 0.038951 | 0.049 | 0.039 | 1 |

| | | | | | | | | | | |
|---|---|---|---|---|---|---|---|---|---|---|
| Q96TA1 | NIBAN2 | HNYGLVLyENkAAYE | Y101 | 0.327 | 97.87 | 746 | 0.14345 | 0.327 | 0.1434 | 1 |
| Q96TA1 | NIBAN2 | PPRAVINsAGYKILt | S119 | 0.057 | 97.18 | 746 | 0.04531 | 0.057 | 0.0453 | 1 |
| Q96TA1 | NIBAN2 | ILtSVDQyLELIGNs | Y131 | 0.081 | 93.21 | 746 | 0.035534 | 0.081 | 0.0355 | 1 |
| Q96JA1 | LRIG1 | LAELssLsVLRLSHN | S334 | 0.184 | 98.26 | 1093 | 0.146264 | 0.184 | 0.1463 | 1 |
| Q9UBR1 | UPB1 | QEVKRVLyGkELRKL | Y30 | 0.155 | 97.28 | 384 | 0.067996 | 0.155 | 0.068 | 1 |
| Q09666 | AHNAK | VQEVTQNsPAARTGV | S41 | 0.107 | 90.39 | 5890 | 0.085056 | 0.107 | 0.0851 | 1 |
| Q96GG9 | DCUN1D1 | LEQLYNRyKDPQDEN | Y72 | 0.04 | 98.05 | 259 | 0.017547 | 0.04 | 0.0175 | 1 |
| Q8N3E9 | PLCD3 | PGGEPVIyHGHTLTS | Y396 | 0.277 | 94.3 | 789 | 0.121516 | 0.277 | 0.1215 | 1 |
| Q8N3E9 | PLCD3 | RHNARQLtRVYPLGL | T583 | 0 | 98.32 | 789 | 0 | 0 | 0 | 1 |
| O43597 | SPRY2 | CDKQCLCsAQNVIDy | S208 | 0.183 | 90.86 | 315 | 0.145469 | 0.183 | 0.1455 | 1 |
| O43597 | SPRY2 | PGCRCKNsNtVCCKV | S296 | 0.157 | 77.8 | 315 | 0.124802 | 0.157 | 0.1248 | 1 |
| Q53EU6 | GPAT3 | SKyNMVSyLLRMMTS | Y352 | 0.019 | 92.2 | 434 | 0.008335 | 0.019 | 0.0083 | 1 |
| P0DPD8 | F1AKMT4-E( | VLGTLSNsRDFLRHF | S859 | 0.145 | 97.46 | 883 | 0.115263 | 0.145 | 0.1153 | 1 |
| Q9UHJ3 | SFMBT1 | INAAYKPsRVLRELQ | S553 | 0.088 | 91.57 | 866 | 0.069952 | 0.088 | 0.07 | 1 |
| Q6DN72 | FCRL6 | RGSPPILySFYLDEK | Y239 | 0 | 95.87 | 434 | 0 | 0 | 0 | 1 |
| P35542 | SAA4 | MIsNHQNsNRYLyAR | S50 | 0.003 | 91.68 | 130 | 0.002385 | 0.154 | 0.1224 | 1 |
| Q9Y512 | SAMM50 | GNFERNFsVNLYKVT | S189 | 0.125 | 96.93 | 469 | 0.099364 | 0.125 | 0.0994 | 1 |
| Q9Y512 | SAMM50 | RGMSAEysFPIWkTS | S216 | 0.166 | 96.58 | 469 | 0.131956 | 0.166 | 0.132 | 1 |
| Q9Y512 | SAMM50 | GCLsRTAsFAVRKES | S243 | 0 | 91.79 | 469 | 0 | 0 | 0 | 1 |
| Q9BXI3 | NT5C1A | AVTIAVssRALFRMD | S58 | 0.071 | 94.98 | 368 | 0.056439 | 0.071 | 0.0564 | 1 |
| P06748 | NPM1 | DENEHQLsLrTVsLG | S43 | 0.021 | 97.54 | 294 | 0.016693 | 0.021 | 0.0167 | 0.875 |
| Q9HC29 | NOD2 | GFIWLIRsLyEMQEE | S745 | 0 | 95.26 | 1040 | 0 | 0 | 0 | 0.8889 |
| Q9HC29 | NOD2 | EGLKKNssLKILKLS | S984 | 0.135 | 97.36 | 1040 | 0.107313 | 0.135 | 0.1073 | 1 |
| O60216 | RAD21 | FECNLESsVESIIsP | S40 | 0.058 | 88.04 | 631 | 0.046105 | 0.058 | 0.0461 | 1 |
| Q04912 | MST1R | AASFSPRsVSIRRLk | S215 | 0.018 | 96.16 | 1400 | 0.014308 | 0.018 | 0.0143 | 1 |
| P00505 | GOT2 | MNLGVGAyRDDNGkP | Y67 | 0.138 | 97.02 | 430 | 0.060539 | 0.138 | 0.0605 | 0.8462 |
| P00505 | GOT2 | GkPyVLPsVRkAEAQ | S79 | 0 | 98.8 | 430 | 0 | 0 | 0 | 1 |
| P00505 | GOT2 | GALRIGAsFLQRFFk | S143 | 0.142 | 98.87 | 430 | 0.112878 | 0.053 | 0.1244 | 1 |
| P00505 | GOT2 | ERLIkEFsIyMTkDG | S399 | 0.095 | 98.5 | 430 | 0.075517 | 0.052 | 0.0905 | 1 |
| Q92905 | COPS5 | YFkyCkIsALALLKM | S58 | 0.009 | 96.9 | 334 | 0.007154 | 0.009 | 0.0072 | 1 |
| Q8WZA1 | POMGNT1 | CTVKDEGsFHLKDtA | S182 | 0.142 | 97.94 | 660 | 0.112878 | 0.142 | 0.1129 | 1 |

| | | | | | | | | | | |
|---|---|---|---|---|---|---|---|---|---|---|
| Q96IY4 | CPB2 | LGIKySFtIELRDTG | T383 | 0.095 | 98.6 | 423 | 0.064886 | 0.095 | 0.0649 | 0.9 |
| Q96CS7 | PLEKHB2 | MDCINIRtGQECRDT | T58 | 0.172 | 94.14 | 222 | 0.117478 | 0.172 | 0.1175 | 1 |
| A6NFN9 | ANKUB1 | DVGIsFCsTLKCFVK | S71 | 0.092 | 92.95 | 502 | 0.073132 | 0.092 | 0.0731 | 1 |
| A6NFN9 | ANKUB1 | NTLKDYQtDIGTTLR | T148 | 0.113 | 88.71 | 502 | 0.077181 | 0.113 | 0.0772 | 1 |
| Q9Y585 | OR1A2 | PALNPFIySLRNWDM | Y288 | 0.087 | 84.76 | 309 | 0.038166 | 0.087 | 0.0382 | 1 |
| Q96DU7 | ITPKC | LRPFVPAyyGMVLQD | Y456 | 0.311 | 97.6 | 683 | 0.136431 | 0.311 | 0.1364 | 1 |
| Q5T089 | MORN1 | AAAHLGEyVLMIRDV | Y435 | 0.029 | 90.54 | 497 | 0.012722 | 0.071 | 0.0311 | 1 |
| Q96Q04 | LMTK3 | HSNYKEDyyLTPERL | Y296 | 0.162 | 92.9 | 1460 | 0.071067 | 0.162 | 0.0711 | 0.8333 |
| O95251 | KAT7 | EIsIKEIsQETAVNP | S545 | 0.053 | 96.52 | 611 | 0.04213 | 0.053 | 0.0421 | 1 |
| O75912 | DGKI | VCDGtDLtPKIQELK | T594 | 0.164 | 95.15 | 1065 | 0.112014 | 0.164 | 0.112 | 1 |
| P42356 | PI4KA | KMGYVREyILWAAsK | Y1672 | 0.041 | 94.85 | 2102 | 0.017986 | 0.041 | 0.018 | 1 |
| P42356 | PI4KA | QSAAkAPyLAkFKVK | Y1803 | 0 | 89.57 | 2102 | 0 | 0 | 0 | 1 |
| P42356 | PI4KA | LDLFVFPyRVVATAP | Y1884 | 0.129 | 90.61 | 2102 | 0.05659 | 0.129 | 0.0566 | 0.8333 |
| P18669 | PGAM1 | YkLVLIRhGEsAWNL | H11 | 0.075 | 98.3 | 254 | 0.038943 | 0.075 | 0.0389 | 0.9167 |
| P18669 | PGAM1 | VLIRhGEsAWNLENr | S14 | 0.006 | 96.38 | 254 | 0.004769 | 0.006 | 0.0048 | 0.8889 |
| P18669 | PGAM1 | GWyDADLsPAGHEEA | S31 | 0.028 | 97.76 | 254 | 0.022258 | 0.05 | 0.0397 | 1 |
| P18669 | PGAM1 | ErHyGGLtGLNkAET | T96 | 0.026 | 97.04 | 254 | 0.017758 | 0.045 | 0.03 | 1 |
| P18669 | PGAM1 | DQLPsCEsLkDTIAR | S155 | 0.017 | 94.76 | 254 | 0.013514 | 0.017 | 0.0135 | 1 |
| Q86VI3 | IQGAP3 | KEGKAAQtERVLRNP | T628 | 0.001 | 91.9 | 1631 | 0.000683 | 0.001 | 0.0007 | 1 |
| P42694 | HELZ | VTVSTKKsHQTWTFA | S277 | 0.097 | 77.57 | 1942 | 0.077107 | 0.119 | 0.0946 | 0.8571 |
| P08913 | ADRA2A | ISLDRyWsItQAIEY | S149 | 0.168 | 92.16 | 465 | 0.133546 | 0.168 | 0.1335 | 1 |
| O00411 | POLRMT | LTPDLLSyAAALQCM | Y299 | 0.058 | 94.51 | 1230 | 0.025444 | 0.212 | 0.093 | 1 |
| O00411 | POLRMT | TRKVVKQtVMTVVyG | T993 | 0.001 | 89.92 | 1230 | 0.000683 | 0.001 | 0.0007 | 1 |
| Q8WU10 | PYROXD1 | HTSLPDIyAAGDICT | Y354 | 0.014 | 98.9 | 500 | 0.006142 | 0.014 | 0.0061 | 1 |
| Q8WU10 | PYROXD1 | RCTKGREyIKVVMQN | Y446 | 0.081 | 95.5 | 500 | 0.035534 | 0.081 | 0.0355 | 0.8333 |
| Q9Y281 | CFL2 | yALyDAtyEtkEskk | Y89 | 0.091 | 92.15 | 166 | 0.03992 | 0.181 | 0.0794 | 1 |
| Q9Y281 | CFL2 | LyDAtyEtkEskkED | T91 | 0.176 | 90.38 | 166 | 0.12021 | 0.176 | 0.1202 | 1 |
| Q9Y281 | CFL2 | LGGNVVVsLEGKPL_ | S160 | 0.116 | 90.14 | 166 | 0.09221 | 0.116 | 0.0922 | 1 |
| P13796 | LCP1 | ALVIFQLyEkIkVPV | Y430 | 0 | 98.62 | 627 | 0 | 0 | 0 | 1 |
| P13796 | LCP1 | SFkDPKIstsLPVLD | S547 | 0.158 | 97.97 | 627 | 0.125597 | 0.158 | 0.1256 | 1 |
| P13796 | LCP1 | kDPKIstsLPVLDLI | S549 | 0 | 98.4 | 627 | 0 | 0 | 0 | 1 |

| | | | | | | | | | | |
|---|---|---|---|---|---|---|---|---|---|---|
| B0I1T2 | MYO1G | HHRHHLHytsRQLCP | Y497 | 0.083 | 96.65 | 1018 | 0.036411 | 0.23 | 0.1009 | 1 |
| B0I1T2 | MYO1G | RHHLHytsRQLCPTD | S499 | 0.009 | 95 | 1018 | 0.007154 | 0.009 | 0.0072 | 1 |
| P48444 | ARCN1 | VLITTKNsNILEDLE | S74 | 0.109 | 90.01 | 511 | 0.086646 | 0.144 | 0.1311 | 1 |
| P13533 | MYH6 | KAGVYKLtGAIMHyG | T354 | 0 | 94.78 | 1939 | 0 | 0 | 0 | 1 |
| Q96T76 | MMS19 | KFLNLSSsPsMAVRI | S965 | 0.092 | 93.27 | 1030 | 0.073132 | 0.092 | 0.0731 | 1 |
| Q9BTW9 | TBCD | DLHMRHGsILACAEV | S621 | 0 | 98.33 | 1192 | 0 | 0 | 0 | 1 |
| Q9BTW9 | TBCD | QLyDRQLyRGLGGQL | Y668 | 0.169 | 94.06 | 1192 | 0.074138 | 0.169 | 0.0741 | 0.9167 |
| Q01105 | SET | FyFDENPyFENkVLS | Y146 | 0.054 | 97.63 | 290 | 0.023689 | 0.054 | 0.0237 | 0.8571 |
| Q9Y5H9 | PCDHA2 | KSYEIQVtAtDKGTP | T321 | 0.215 | 94.42 | 948 | 0.146848 | 0.215 | 0.1468 | 1 |
| Q08828 | ADCY1 | EALFRGytLRLEQAA | T53 | 0.032 | 91.34 | 1119 | 0.021856 | 0.106 | 0.1098 | 1 |
| Q96M69 | LRGUK | YSYFEPRyILVVPMN | Y529 | 0.079 | 95.29 | 825 | 0.034656 | 0.079 | 0.0347 | 1 |
| Q96M69 | LRGUK | VPMNKEKyEGyLRRK | Y540 | 0.12 | 95.83 | 825 | 0.052642 | 0.12 | 0.0526 | 0.8571 |
| P49802 | RGS7 | YKLMKSDsYPRFIRS | S434 | 0.066 | 97.03 | 495 | 0.052464 | 0.066 | 0.0525 | 0.9 |
| Q00341 | HDLBP | IkKIREEsNtkIDLP | S611 | 0.081 | 79.38 | 1268 | 0.064388 | 0.081 | 0.0644 | 1 |
| Q00341 | HDLBP | KIREEsNtkIDLPAE | T613 | 0.082 | 81.48 | 1268 | 0.056007 | 0.082 | 0.056 | 1 |
| Q8TD47 | RPS4Y2 | AGFIDVIsIEKTGEH | S91 | 0.172 | 97.18 | 263 | 0.136725 | 0.172 | 0.1367 | 0.8571 |
| Q7Z4H8 | POGLUT3 | GDEHVAQsPYILKGP | S128 | 0.136 | 97.6 | 507 | 0.108108 | 0.136 | 0.1081 | 1 |
| Q7Z4H8 | POGLUT3 | GFFDFFKyKYQVNVD | Y367 | 0.234 | 98.72 | 507 | 0.102652 | 0.234 | 0.1027 | 1 |
| Q7Z4H8 | POGLUT3 | ALEPWKHyVPIkRNL | Y413 | 0.336 | 98.87 | 507 | 0.147398 | 0.336 | 0.1474 | 0.9167 |
| Q7Z4H8 | POGLUT3 | YYQVLQKyAERQssK | Y468 | 0 | 98.61 | 507 | 0 | 0 | 0 | 1 |
| Q14289 | PTK2B | EGAPQALsIKTSSLA | S332 | 0.147 | 94.36 | 1009 | 0.116852 | 0.147 | 0.1169 | 1 |
| Q13595 | TRA2A | GrRIRVDysItkRAH | Y194 | 0.317 | 89.14 | 282 | 0.139063 | 0.317 | 0.1391 | 1 |
| Q13595 | TRA2A | rRIRVDysItkRAHt | S195 | 0.08 | 80.97 | 282 | 0.063593 | 0.107 | 0.0851 | 1 |
| Q9BQ39 | DDX50 | PPQDVESyIHRSGRT | Y472 | 0.001 | 89.11 | 737 | 0.000439 | 0.001 | 0.0004 | 1 |
| Q8NHU2 | CFAP61 | EKANLAVtLCNDKEE | T56 | 0 | 96.69 | 1237 | 0 | 0 | 0 | 1 |
| Q8NHU2 | CFAP61 | ILRLGFKsCLYYRVY | S585 | 0.044 | 94.09 | 1237 | 0.034976 | 0.044 | 0.035 | 0.875 |
| P56556 | NDUFA6 | GKIELEEtIKVWKQR | T93 | 0.063 | 95.43 | 128 | 0.04303 | 0.063 | 0.043 | 1 |
| Q9H2X9 | SLC12A5 | RYyHWTLsFLGMSLC | S622 | 0.013 | 95.98 | 1139 | 0.010334 | 0.013 | 0.0103 | 1 |
| O60423 | ATP8B3 | TIKCLKKsNIKIWVL | S755 | 0 | 89.18 | 1300 | 0 | 0 | 0 | 1 |
| O60423 | ATP8B3 | NIKIWVLtGDKQEtA | T763 | 0.009 | 88.31 | 1300 | 0.006147 | 0.009 | 0.0061 | 1 |
| Q9NX46 | ADPRS | PDPGtPGsERTEALY | S67 | 0.166 | 96.5 | 363 | 0.131956 | 0.166 | 0.132 | 1 |

| | | | | | | | | | | |
|---|---|---|---|---|---|---|---|---|---|---|
| Q9H0V1 | TMEM168 | VSKRWSDytLHLPtG | Y625 | 0.29 | 82.54 | 697 | 0.127219 | 0.336 | 0.1474 | 0.8182 |
| Q9H799 | CPLANE1 | DVIVLTTsSNDAWLA | S70 | 0.002 | 89.87 | 3197 | 0.00159 | 0.02 | 0.0159 | 1 |
| Q16774 | GUK1 | AAQADMEssKEPGLF | S158 | 0.133 | 94.63 | 197 | 0.105724 | 0.133 | 0.1057 | 0.8571 |
| Q16774 | GUK1 | AQADMEssKEPGLFD | S159 | 0.179 | 93.82 | 197 | 0.14229 | 0.179 | 0.1423 | 1 |
| Q9H3E2 | SNX25 | IMLAEILttKVLKPV | T135 | 0.015 | 96.16 | 840 | 0.010245 | 0.015 | 0.0102 | 1 |
| P54707 | ATP12A | GIkVIMVtGDHPItA | T632 | 0.011 | 91.08 | 1039 | 0.007513 | 0.011 | 0.0075 | 0.8571 |
| O43511 | SLC26A4 | FGTSRHIsVGPFPVV | S137 | 0 | 91.82 | 780 | 0 | 0 | 0 | 1 |
| O43511 | SLC26A4 | VGPFPVVsLMVGSVV | S145 | 0 | 90.43 | 780 | 0 | 0 | 0 | 1 |
| O43511 | SLC26A4 | LLEPLQKsVLAAVVI | S448 | 0.008 | 92.4 | 780 | 0.006359 | 0.008 | 0.0064 | 1 |
| P46379 | BAG6 | SLEVLVktLDsQTRT | T23 | 0.163 | 92.11 | 1132 | 0.111331 | 0.163 | 0.1113 | 1 |
| Q53F39 | MPPE1 | INFVMVNsVALNGDG | S196 | 0.018 | 98.57 | 396 | 0.014308 | 0.018 | 0.0143 | 1 |
| Q14684 | RRP1B | VLCPESQsPNGVRFH | S160 | 0.137 | 83.01 | 758 | 0.108903 | 0.137 | 0.1089 | 1 |
| P63279 | UBE2I | FkDDYPssPPkCKFE | S71 | 0.005 | 97.8 | 158 | 0.003975 | 0.005 | 0.004 | 1 |
| O60503 | ADCY9 | RIQVSEEsYRVLskM | S1212 | 0.006 | 95.92 | 1353 | 0.004769 | 0.006 | 0.0048 | 1 |
| O14772 | FPGT | LNSYKLLsIEEMLIy | S576 | 0 | 97.19 | 607 | 0 | 0 | 0 | 1 |
| Q9NZU5 | LMCD1 | tQKLGLQyMELIPKE | Y133 | 0.112 | 97.93 | 365 | 0.049133 | 0.112 | 0.0491 | 1 |
| Q13404 | UBE2V1 | QPPEGQCysN_____ | Y145 | 0.211 | 96.19 | 147 | 0.092563 | 0.211 | 0.0926 | 1 |
| Q16819 | MEP1A | YGFGVTLyPNSRESS | Y472 | 0.027 | 97.49 | 746 | 0.011845 | 0.174 | 0.0736 | 1 |
| Q16819 | MEP1A | SSSMVFTtsKSHTSP | T529 | 0.011 | 98.19 | 746 | 0.007513 | 0.011 | 0.0075 | 1 |
| Q8WUJ0 | STYX | VHGNAGIsRSAAFVI | S125 | 0.106 | 98.66 | 223 | 0.084261 | 0.106 | 0.0843 | 0.8333 |
| Q9UQ84 | EXO1 | NITVPEDyINGFIRA | Y271 | 0.163 | 96.86 | 846 | 0.071506 | 0.163 | 0.0715 | 1 |
| Q9UQ84 | EXO1 | RANNTFLyQLVFDPI | Y284 | 0.072 | 98.17 | 846 | 0.031585 | 0.072 | 0.0316 | 0.9231 |
| Q9HC10 | OTOF | LRVIIWNtDEVVLED | T1741 | 0.037 | 93.42 | 1997 | 0.025272 | 0.071 | 0.0482 | 1 |
| Q9HC10 | OTOF | DFFtGEKssDIFVRG | S1757 | 0.038 | 84.88 | 1997 | 0.030207 | 0.038 | 0.0302 | 1 |
| Q9HC10 | OTOF | FFtGEKssDIFVRGW | S1758 | 0.003 | 86.35 | 1997 | 0.002385 | 0.003 | 0.0024 | 1 |
| Q9UNN5 | FAF1 | VSkLRIRtPsGEFLE | T580 | 0.097 | 88 | 650 | 0.066252 | 0.097 | 0.0663 | 1 |
| Q9Y2T1 | AXIN2 | IENNSIVsKQLKPAT | S146 | 0.032 | 91.56 | 843 | 0.025437 | 0.032 | 0.0254 | 1 |
| Q9HAU8 | RNPEPL1 | SKWRtFQtALFLDRL | T565 | 0.003 | 97.99 | 725 | 0.002049 | 0.003 | 0.002 | 1 |
| Q9UM54 | MYO6 | REKLHLSsPDNFRYL | S267 | 0.149 | 93.7 | 1294 | 0.118442 | 0.149 | 0.1184 | 1 |
| Q9UM54 | MYO6 | QILQNRKsPEyLKAG | S297 | 0.001 | 92.94 | 1294 | 0.000795 | 0.001 | 0.0008 | 1 |
| Q9UM54 | MYO6 | QNRKsPEyLKAGSMK | Y300 | 0.085 | 91.31 | 1294 | 0.037288 | 0.085 | 0.0373 | 1 |

| | | | | | | | | | | |
|---|---|---|---|---|---|---|---|---|---|---|
| P46019 | PHKA2 | HIGVLGTsKLYVIRN | S508 | 0.016 | 97.03 | 1235 | 0.012719 | 0.016 | 0.0127 | 1 |
| Q9H8Y5 | ANKZF1 | DSRARPPyTVAADKs | Y572 | 0.32 | 91.93 | 726 | 0.140379 | 0.32 | 0.1404 | 1 |
| Q9H8Y5 | ANKZF1 | TVAADKstRNEFRRF | T580 | 0.001 | 91.51 | 726 | 0.000683 | 0.001 | 0.0007 | 1 |
| Q9Y253 | POLH | QTLVSHGsVPQLFSQ | S242 | 0.138 | 97.96 | 713 | 0.109698 | 0.138 | 0.1097 | 1 |
| Q9Y253 | POLH | VATQLVVsIRVQGDk | S369 | 0.025 | 98.67 | 713 | 0.019873 | 0.025 | 0.0199 | 1 |
| Q9Y253 | POLH | TMLFLCAtKFSASAP | T427 | 0.187 | 98.48 | 713 | 0.127724 | 0.187 | 0.1277 | 1 |
| P22083 | FUT4 | rLLTDrAsyGEAQAV | S228 | 0.15 | 93.53 | 530 | 0.119237 | 0.15 | 0.1192 | 1 |
| P22083 | FUT4 | LLTDrAsyGEAQAVL | Y229 | 0.117 | 94.1 | 530 | 0.051326 | 0.117 | 0.0513 | 1 |
| P22083 | FUT4 | TVARYKFyLAFENSQ | Y410 | 0.008 | 98.82 | 530 | 0.003509 | 0.008 | 0.0035 | 1 |
| Q8TDD1 | DDX54 | NPDIIIAtPGRLVHV | T223 | 0.155 | 95.22 | 881 | 0.105867 | 0.155 | 0.1059 | 1 |
| Q8N531 | FBXL6 | LTLAkEGsPFLTQkW | S431 | 0.014 | 96.41 | 539 | 0.011129 | 0.014 | 0.0111 | 0.875 |
| Q16666 | IFI16 | kKIMFHAtVATQtQF | T232 | 0.148 | 94.95 | 785 | 0.101086 | 0.148 | 0.1011 | 0.875 |
| Q16666 | IFI16 | KTVNQkTtIyEIQDD | T332 | 0 | 94.47 | 785 | 0 | 0 | 0 | 1 |
| Q16666 | IFI16 | VNQkTtIyEIQDDRG | Y334 | 0.009 | 97.08 | 785 | 0.003948 | 0.009 | 0.0039 | 1 |
| Q9Y210 | TRPC6 | NAYKGLAsPAYLSLS | S282 | 0.049 | 94.74 | 931 | 0.038951 | 0.049 | 0.039 | 1 |
| Q9Y210 | TRPC6 | KNDYKKLsMQCKDFV | S322 | 0.067 | 95.63 | 931 | 0.053259 | 0.067 | 0.0533 | 1 |
| P54803 | GALC | VSGGGATsRLLVNYP | S68 | 0.016 | 98.82 | 685 | 0.012719 | 0.032 | 0.0254 | 1 |
| P54803 | GALC | TDGLGNLtIIIETMS | T381 | 0 | 98.88 | 685 | 0 | 0 | 0 | 1 |
| P54803 | GALC | ANGSYRVtGDLAGWI | T608 | 0.024 | 98.5 | 685 | 0.016392 | 0.024 | 0.0164 | 1 |
| Q9Y614 | ACTL7B | AMHVtSQsLLSIYSY | S182 | 0.001 | 97.88 | 415 | 0.000795 | 0.001 | 0.0008 | 1 |
| Q9BXR6 | CFHR5 | AQPKKESyKVGDVLK | Y165 | 0.055 | 89.68 | 569 | 0.024128 | 0.055 | 0.0241 | 0.8571 |
| O75116 | ROCK2 | LQMKAEDyDVVkVIG | Y92 | 0.09 | 97.77 | 1388 | 0.039482 | 0.143 | 0.0496 | 1 |
| Q9UJC5 | SH3BGRL2 | RVFIAsssGFVAIKK | S11 | 0.172 | 83.18 | 107 | 0.136725 | 0.172 | 0.1367 | 1 |
| O75891 | ALDH1L1 | ALKFAELtLKAGIPK | T613 | 0.028 | 97.49 | 902 | 0.019124 | 0.028 | 0.0191 | 1 |
| Q8NE35 | CPEB3 | AFSNQQSyIAAISAR | Y604 | 0.14 | 96.18 | 698 | 0.061416 | 0.14 | 0.0614 | 0.8571 |
| Q76LX8 | ADAMTS13 | LVPLLDGtECGVEKW | T358 | 0.011 | 91.87 | 1427 | 0.007513 | 0.011 | 0.0075 | 1 |
| O43304 | SEC14L5 | LIYSGSNyQGPGGLV | Y455 | 0.137 | 88.92 | 696 | 0.0601 | 0.137 | 0.0601 | 1 |
| Q13683 | ITGA7 | RKEGEPGsLFGFSVA | S50 | 0.036 | 97.1 | 1181 | 0.028617 | 0.036 | 0.0286 | 1 |
| Q13683 | ITGA7 | yLGFSIDsGKGLVRA | S313 | 0.058 | 96.7 | 1181 | 0.046105 | 0.08 | 0.0636 | 1 |
| Q5VWZ2 | LYPLAL1 | PTAPPRSyTPMkGGI | Y66 | 0.167 | 98.69 | 237 | 0.07326 | 0.167 | 0.0733 | 1 |
| Q14566 | MCM6 | PVHPELVsGTFLCLD | S153 | 0.141 | 86.23 | 821 | 0.112083 | 0.141 | 0.1121 | 1 |

| | | | | | | | | | | |
|---|---|---|---|---|---|---|---|---|---|---|
| Q8NA66 | CNBD1 | LIRMCPyyEEWPTLS | Y323 | 0.174 | 89.75 | 436 | 0.076331 | 0.174 | 0.0763 | 1 |
| Q5JTH9 | RRP12 | EGLVKsCsEtLLRVM | S327 | 0.038 | 94 | 1297 | 0.030207 | 0.038 | 0.0302 | 1 |
| Q5JTH9 | RRP12 | SASGPAQsVAKMFRA | S467 | 0.133 | 93.41 | 1297 | 0.105724 | 0.133 | 0.1057 | 1 |
| P55084 | HADHB | LTGLLHRtSVPKEVV | T91 | 0.115 | 98.3 | 474 | 0.078547 | 0.127 | 0.1119 | 1 |
| P55084 | HADHB | SRLEQDEyALrSHsL | Y244 | 0.103 | 98.74 | 474 | 0.045185 | 0.201 | 0.0864 | 1 |
| P55084 | HADHB | SFLTDGAsAMLIMAE | S316 | 0.005 | 98.88 | 474 | 0.003975 | 0.005 | 0.004 | 1 |
| P55084 | HADHB | MGYkPKAyLRDFMyV | Y336 | 0.24 | 98.52 | 474 | 0.105285 | 0.24 | 0.1053 | 1 |
| P55084 | HADHB | LLGPTyAtPkVLEKA | T359 | 0 | 98.79 | 474 | 0 | 0 | 0 | 1 |
| P39023 | RPL3 | LGYKAGMtHIVREVD | T54 | 0.095 | 98.23 | 403 | 0.064886 | 0.095 | 0.0649 | 1 |
| P39023 | RPL3 | LIkNNAstDyDLsDk | T305 | 0.166 | 94.63 | 403 | 0.11338 | 0.166 | 0.1134 | 1 |
| P42336 | PIK3CA | YCRACGMyLKHLNRQ | Y698 | 0.081 | 97.97 | 1068 | 0.035534 | 0.081 | 0.0355 | 1 |
| P42336 | PIK3CA | ALQFNsHtLHQWLKD | T876 | 0.134 | 93.91 | 1068 | 0.091524 | 0.188 | 0.1482 | 1 |
| Q9NQX4 | MYO5C | GDAIIHAysGQNMGD | Y125 | 0.189 | 91.82 | 1742 | 0.082912 | 0.189 | 0.0829 | 1 |
| Q9NQX4 | MYO5C | ARNNRNQsIIVSGES | S156 | 0.001 | 95.39 | 1742 | 0.000795 | 0.026 | 0.0207 | 1 |
| Q96N76 | UROC1 | TSGLGGMsGAQAKAA | S253 | 0.099 | 90.19 | 676 | 0.078697 | 0.099 | 0.0787 | 1 |
| P08183 | ABCB1 | VQLLERFyDPLAGkV | Y1087 | 0.072 | 95.06 | 1280 | 0.031585 | 0.141 | 0.0619 | 1 |
| P08183 | ABCB1 | LLAQKGIyFsMVSVQ | Y1267 | 0.003 | 88.12 | 1280 | 0.001316 | 0.003 | 0.0013 | 1 |
| Q15047 | SETDB1 | TVGPGKKyKVKFDNK | Y230 | 0.075 | 96.55 | 1291 | 0.032901 | 0.075 | 0.0329 | 1 |
| P33763 | S100A5 | ALttMVTtFHKYSGR | T15 | 0.215 | 97.64 | 92 | 0.146848 | 0.215 | 0.1468 | 1 |
| P18074 | ERCC2 | TsVALEKyQEACENG | Y584 | 0.026 | 94.41 | 760 | 0.011406 | 0.026 | 0.0114 | 1 |
| Q5VU97 | CACHD1 | VMAtSHVtDEWMTQM | T712 | 0.004 | 91.17 | 1274 | 0.002732 | 0.004 | 0.0027 | 1 |
| O43175 | PHGDH | QIPQAtAsMkDGKWE | S127 | 0.08 | 98.12 | 533 | 0.063593 | 0.08 | 0.0636 | 1 |
| O43175 | PHGDH | kGtIQVItQGTsLkN | T358 | 0.015 | 94.24 | 533 | 0.010245 | 0.015 | 0.0102 | 1 |
| O43175 | PHGDH | kNAGNCLsPAVIVGL | S371 | 0.087 | 93.25 | 533 | 0.069158 | 0.087 | 0.0692 | 1 |
| Q14696 | MESD | LMMFVTVsGsPtEKE | S121 | 0.145 | 91.83 | 234 | 0.115263 | 0.145 | 0.1153 | 1 |
| Q14571 | ITPR2 | GDQRyAPyKyMLRLC | Y549 | 0.225 | 86.05 | 2701 | 0.098704 | 0.225 | 0.0987 | 1 |
| Q00722 | PLCB2 | VQGLIDKyEPsGINA | Y274 | 0.246 | 94.84 | 1185 | 0.107917 | 0.246 | 0.1079 | 1 |
| Q00722 | PLCB2 | KPGVPLPsPEDLRGK | S450 | 0.043 | 98.74 | 1185 | 0.034181 | 0.043 | 0.0342 | 1 |
| Q00722 | PLCB2 | EMSSLVNyIQPTKFV | Y552 | 0.069 | 98.26 | 1185 | 0.030269 | 0.069 | 0.0303 | 1 |
| Q00722 | PLCB2 | DYNKRQMsRIyPKGT | S601 | 0 | 98.7 | 1185 | 0 | 0 | 0 | 1 |
| Q00722 | PLCB2 | KRQMsRIyPKGTRMD | Y604 | 0.264 | 98.32 | 1185 | 0.115813 | 0.264 | 0.1158 | 1 |

| | | | | | | | | | | |
|---|---|---|---|---|---|---|---|---|---|---|
| P11274 | BCR | NGIEVKLsVKFNsRE | S1030 | 0.092 | 92.81 | 1271 | 0.073132 | 0.106 | 0.112 | 0.8571 |
| Q9Y276 | BCS1L | LsVAPQQsLVLLEDV | S277 | 0.1 | 93.29 | 419 | 0.079491 | 0.1 | 0.0795 | 1 |
| Q8IU80 | TMPRSS6 | LKELITStRLGTyYN | T129 | 0.183 | 91.17 | 811 | 0.124991 | 0.183 | 0.125 | 1 |
| Q8IU80 | TMPRSS6 | PLEKRLItsVyGCSR | T287 | 0.115 | 94.97 | 811 | 0.078547 | 0.115 | 0.0785 | 0.8571 |
| O43557 | TNFSF14 | tHGLyKRtPRYPEEL | T170 | 0.154 | 98.26 | 240 | 0.105184 | 0.154 | 0.1052 | 1 |
| Q8IVF4 | DNAH10 | PVRGLSVtYyLEAKM | T1318 | 0.078 | 85.19 | 4471 | 0.053275 | 0.102 | 0.0697 | 1 |
| Q9NTG7 | SIRT3 | KGLLLRLyTQNIDGL | Y226 | 0 | 98.82 | 399 | 0 | 0 | 0 | 1 |
| Q5JVF3 | PCID2 | SSNLkDDySTAQRVT | Y204 | 0.064 | 95.59 | 399 | 0.028076 | 0.064 | 0.0281 | 1 |
| Q5JVF3 | PCID2 | TVELLKKyHLMQFAE | Y274 | 0.14 | 96.87 | 399 | 0.061416 | 0.14 | 0.0614 | 1 |
| Q5JVF3 | PCID2 | LMQFAEVtRAVSEGN | T283 | 0.122 | 98.25 | 399 | 0.083328 | 0.122 | 0.0833 | 1 |
| Q6ZQQ6 | WDR87 | LFVATGssEVLVFDT | S392 | 0.099 | 90.695 | 2873 | 0.078697 | 0.099 | 0.0787 | 1 |
| Q6V1X1 | DPP8 | DsFRYPktGtANPkV | T334 | 0.041 | 97.83 | 898 | 0.028004 | 0.041 | 0.028 | 0.8462 |
| Q6V1X1 | DPP8 | RLVYFEGtKDSPLEH | T543 | 0.061 | 98.3 | 898 | 0.041664 | 0.061 | 0.0417 | 0.9167 |
| P25325 | MPST | PLQLLDAsWYLPkLG | S35 | 0.06 | 98.75 | 297 | 0.047695 | 0.06 | 0.0477 | 1 |
| P25325 | MPST | DPAFIktyEDIkENL | Y166 | 0.168 | 98.67 | 297 | 0.073699 | 0.276 | 0.0997 | 1 |
| P25325 | MPST | GKPDVPIyDGSWVEW | Y272 | 0.005 | 98.88 | 297 | 0.002193 | 0.005 | 0.0022 | 0.8571 |
| Q96GD3 | SCMH1 | QDPRNttstClAtVV | S74 | 0.129 | 96.56 | 660 | 0.102544 | 0.129 | 0.1025 | 1 |
| O75309 | CDH16 | RLGALALsPKGSTSL | S192 | 0.112 | 93.48 | 829 | 0.08903 | 0.112 | 0.089 | 1 |
| O60229 | KALRN | DVCKRGFtVIIDMRG | T95 | 0.036 | 93.35 | 2986 | 0.024588 | 0.036 | 0.0246 | 1 |
| Q8IYW2 | CFAP46 | KMRQPEVsEDCIQMy | S65 | 0 | 92.73 | 2715 | 0 | 0 | 0 | 1 |
| Q8IYW2 | CFAP46 | sEDCIQMyFKVKAPI | Y72 | 0.073 | 92.01 | 2715 | 0.032024 | 0.073 | 0.032 | 0.8571 |
| Q96LJ8 | UBXN10 | RLLLAVRsPtGQRFV | S205 | 0.05 | 95.06 | 280 | 0.039746 | 0.05 | 0.0397 | 1 |
| Q96LJ8 | UBXN10 | KTSYRHCsIEtMEVP | S240 | 0.022 | 95.84 | 280 | 0.017488 | 0.022 | 0.0175 | 1 |
| Q86TM6 | SYVN1 | AVtDAIMsRRAIRNM | S265 | 0.069 | 89.53 | 617 | 0.054849 | 0.069 | 0.0548 | 1 |
| P13693 | TPT1 | IsHDEMFsDIykIRE | S15 | 0.026 | 98.02 | 172 | 0.020668 | 0.026 | 0.0207 | 0.8571 |
| Q9Y623 | MYH4 | GGKVTAKtEAGAtVT | T64 | 0.185 | 86.25 | 1939 | 0.126358 | 0.185 | 0.1264 | 1 |
| P30038 | ALDH4A1 | VEsVVSGtLRSAFEY | T336 | 0.007 | 98.92 | 563 | 0.004781 | 0.007 | 0.0048 | 1 |
| P30038 | ALDH4A1 | rNAAGNFyINDKSTG | Y505 | 0.249 | 98.85 | 563 | 0.109233 | 0.249 | 0.1092 | 1 |
| Q9P218 | COL20A1 | SLRGPEGsEARGIRA | S545 | 0.098 | 87.09 | 1284 | 0.077902 | 0.181 | 0.1456 | 0.9091 |
| Q9P218 | COL20A1 | ARGIRARtPtLAPPR | T554 | 0.015 | 82.71 | 1284 | 0.010245 | 0.015 | 0.0102 | 1 |
| Q9C0C7 | AMBRA1 | GLGLAyGtNKGDLVI | T1028 | 0.001 | 90.05 | 1298 | 0.000683 | 0.001 | 0.0007 | 0.8 |

| | | | | | | | | | | |
|---|---|---|---|---|---|---|---|---|---|---|
| Q96PP9 | GBP4 | QAELKRLsEHLtEsI | S429 | 0.086 | 94.54 | 640 | 0.068363 | 0.086 | 0.0684 | 1 |
| Q96PP9 | GBP4 | LsEHLtEsILRGIFS | S435 | 0.036 | 92.08 | 640 | 0.028617 | 0.036 | 0.0286 | 1 |
| P57764 | GSDMD | SSTGFQPyCLVVRkP | Y37 | 0.085 | 95.13 | 484 | 0.037288 | 0.085 | 0.0373 | 0.8571 |
| Q8TBX8 | PIP4K2C | LVMRNMFsHRLPVHR | S208 | 0.022 | 93.13 | 421 | 0.017488 | 0.022 | 0.0175 | 1 |
| Q8TBX8 | PIP4K2C | GAPQKEVyFMGLIDI | Y368 | 0 | 98.09 | 421 | 0 | 0 | 0 | 0.8235 |
| P23276 | KEL | LRLLMSQyGHFPFFR | Y201 | 0.164 | 92.95 | 732 | 0.071944 | 0.164 | 0.0719 | 1 |
| P23276 | KEL | PSKVQEHsSLSISIT | S272 | 0.003 | 96.62 | 732 | 0.002385 | 0.003 | 0.0024 | 1 |
| O75843 | AP1G2 | VLFETVLtIMDIRSA | T296 | 0 | 96.84 | 785 | 0 | 0 | 0 | 1 |
| Q8IYA7 | MKX | WALRIKLyNKYVQGN | Y146 | 0.19 | 93.62 | 352 | 0.08335 | 0.19 | 0.0833 | 1 |
| Q9HCJ5 | ZSWIM6 | IIPLVVKsVKCATVL | S1097 | 0.03 | 92.64 | 1215 | 0.023847 | 0.03 | 0.0238 | 1 |
| P19447 | ERCC3 | KDKPIGCsVAISTYS | S404 | 0.118 | 88.81 | 782 | 0.0938 | 0.134 | 0.1065 | 1 |
| Q9H2Y9 | SLCO5A1 | ASNASIytGVIIVPs | T474 | 0.011 | 94.58 | 848 | 0.007513 | 0.189 | 0.1291 | 1 |
| Q96RU8 | TRIB1 | RLESLEDtHIMkGED | T229 | 0.217 | 91.1 | 372 | 0.148214 | 0.217 | 0.1482 | 1 |
| P23409 | MYF6 | FEALKRRtVANPNQR | T120 | 0.192 | 97.36 | 242 | 0.131139 | 0.192 | 0.1311 | 1 |
| P16581 | SELE | KESGAWSyNTSTEAM | Y24 | 0.312 | 91.41 | 610 | 0.13687 | 0.312 | 0.1369 | 1 |
| P16581 | SELE | HSPIGEFtyKSSCAF | T452 | 0.013 | 89.71 | 610 | 0.008879 | 0.013 | 0.0089 | 1 |
| Q86W24 | NLRP14 | RLIKMNLtQNTLGYE | T1022 | 0.061 | 92.58 | 1093 | 0.041664 | 0.061 | 0.0417 | 1 |
| Q6ZS30 | NBEAL1 | LEHVLsQsIKEQTEI | S1625 | 0.044 | 82.15 | 2694 | 0.034976 | 0.044 | 0.035 | 1 |
| Q9H3R1 | NDST4 | IYENILKyVSMDSWN | Y139 | 0.012 | 91.91 | 872 | 0.005264 | 0.012 | 0.0053 | 1 |
| Q9H3R1 | NDST4 | DCFVNPQsPLLHITK | S199 | 0.124 | 93.03 | 872 | 0.098569 | 0.124 | 0.0986 | 1 |
| Q9H3R1 | NDST4 | SIISNLPsPKTFEEV | S626 | 0.151 | 93.86 | 872 | 0.120032 | 0.151 | 0.12 | 1 |
| Q9H3R1 | NDST4 | SDFLFEKsANyFHSE | S667 | 0.002 | 98.41 | 872 | 0.00159 | 0.002 | 0.0016 | 1 |
| Q9H3R1 | NDST4 | LFEKsANyFHSEEAP | Y670 | 0.008 | 98.65 | 872 | 0.003509 | 0.008 | 0.0035 | 1 |
| Q9H3R1 | NDST4 | yPPMDPEsRTFLSNY | S834 | 0 | 97.66 | 872 | 0 | 0 | 0 | 1 |
| Q96JB1 | DNAH8 | ALQDFQKyKtLWtED | Y1010 | 0.005 | 86.615 | 4490 | 0.002193 | 0.005 | 0.0022 | 0.8 |
| O75110 | ATP9A | PGtVVRSstIPEQLG | S376 | 0.176 | 83.05 | 1047 | 0.139905 | 0.176 | 0.1399 | 0.8571 |
| O75110 | ATP9A | PEQLGRIsYLLtDkT | S386 | 0.169 | 94.97 | 1047 | 0.134341 | 0.169 | 0.1343 | 1 |
| O75110 | ATP9A | GRIsYLLtDkTGTLT | T390 | 0.011 | 95.65 | 1047 | 0.007513 | 0.011 | 0.0075 | 1 |
| Q7Z6J2 | TAMALIN | GNVLRLEtLYGTSIR | T186 | 0.012 | 91.8 | 395 | 0.008196 | 0.012 | 0.0082 | 1 |
| Q9UHR4 | BAIAP2L1 | NRLTESTyRNVMEQF | Y16 | 0.093 | 97.58 | 511 | 0.040798 | 0.093 | 0.0408 | 1 |
| Q9UHR4 | BAIAP2L1 | DVKyMNAtLKRYQTE | T118 | 0.049 | 97.32 | 511 | 0.033468 | 0.072 | 0.0457 | 1 |

| | | | | | | | | | | |
|---|---|---|---|---|---|---|---|---|---|---|
| Q9UHR4 | BAIAP2L1 | KLEsLEKsQAELKKI | S136 | 0.031 | 96.34 | 511 | 0.024642 | 0.031 | 0.0246 | 1 |
| Q9NR16 | CD163L1 | VGGSNRCsGRLEVYF | S593 | 0.063 | 76.07 | 1453 | 0.05008 | 0.083 | 0.066 | 1 |
| Q9BZF3 | OSBPL6 | KNKFWGKsMEILPVG | S704 | 0.183 | 72.76 | 934 | 0.145469 | 0.183 | 0.1455 | 1 |
| Q8IWV8 | UBR2 | FkVGEPTySCRDCAV | Y110 | 0.089 | 90.72 | 1755 | 0.039043 | 0.235 | 0.1031 | 1 |
| P10586 | PTPRF | SKGsGPLsPSIQSRT | S1000 | 0.089 | 90.34 | 1907 | 0.070747 | 0.166 | 0.1362 | 1 |
| Q9Y2A7 | NCKAP1 | SDISEMRsLsELLGP | S838 | 0.127 | 97.85 | 1128 | 0.100954 | 0.127 | 0.101 | 1 |
| Q9Y2A7 | NCKAP1 | ISEMRsLsELLGPYG | S840 | 0.013 | 97.92 | 1128 | 0.010334 | 0.013 | 0.0103 | 1 |
| Q9Y2A7 | NCKAP1 | FLALASSsLLKIGQE | S1065 | 0.102 | 89.37 | 1128 | 0.081081 | 0.184 | 0.1485 | 1 |
| P35228 | NOS2 | KVVCMDKyRLSCLEE | Y575 | 0.204 | 95.42 | 1153 | 0.089492 | 0.204 | 0.0895 | 1 |
| Q9P0M2 | AKAP7 | LYRIDLCsMLKKKQS | S264 | 0.037 | 97.9 | 348 | 0.029412 | 0.037 | 0.0294 | 1 |
| P84074 | HPCA | LEIVQAIyKMVSSVM | Y129 | 0.26 | 87.69 | 193 | 0.114058 | 0.26 | 0.1141 | 1 |
| Q99650 | OSMR | GTNVTICyVSRNIQN | Y168 | 0.023 | 86.59 | 979 | 0.01009 | 0.023 | 0.0101 | 1 |
| Q9HCM3 | KIAA1549 | FLNGSEVsELLRNLS | S1129 | 0.084 | 90.97 | 1950 | 0.066773 | 0.084 | 0.0668 | 1 |
| Q8TF27 | AGAP11 | LKTWKKKyVTLCSNG | Y143 | 0.254 | 95.54 | 550 | 0.111426 | 0.254 | 0.1114 | 1 |
| P51817 | PRKX | AKKLVDRtWtLCGtP | T201 | 0.032 | 90.57 | 358 | 0.021856 | 0.032 | 0.0219 | 1 |
| Q96HA7 | TONSL | FPRAAEAyQkQLRFA | Y334 | 0.134 | 98.07 | 1378 | 0.058784 | 0.134 | 0.0588 | 1 |
| Q9C0H5 | ARHGAP39 | NTSRFYYyNASTQRT | Y83 | 0.15 | 89.56 | 1083 | 0.065803 | 0.316 | 0.1386 | 0.8 |
| Q8N323 | NXPE1 | NGTSHVFtECGLTLN | T218 | 0.18 | 97.09 | 547 | 0.122942 | 0.18 | 0.1229 | 1 |
| Q8N323 | NXPE1 | PPVPGGYtLQGKWIT | T320 | 0.025 | 98.28 | 547 | 0.017075 | 0.025 | 0.0171 | 1 |
| Q9GZX3 | CHST6 | DLFQWAVsRALCSPP | S126 | 0.024 | 98.11 | 395 | 0.019078 | 0.024 | 0.0191 | 1 |
| Q5T9G4 | ARMC12 | LVFAERLsEGRNAPH | S259 | 0.01 | 96.35 | 340 | 0.007949 | 0.01 | 0.0079 | 1 |
| Q9H3R0 | KDM4C | AVtSQtFyEVMFDDG | Y908 | 0.094 | 91.64 | 1056 | 0.041236 | 0.094 | 0.0412 | 1 |
| Q9H3R0 | KDM4C | MFDDGsFsRDTFPED | S918 | 0.053 | 86.37 | 1056 | 0.04213 | 0.053 | 0.0421 | 1 |
| Q9H3R0 | KDM4C | MKREDIYtLDEELPk | T985 | 0.021 | 89.51 | 1056 | 0.014343 | 0.114 | 0.0779 | 1 |
| Q49A26 | GLYR1 | DQsDNDMsAVYRAYI | S545 | 0 | 98.73 | 553 | 0 | 0 | 0 | 1 |
| Q6NUI2 | GPAT2 | LFLCRLLsPLLKAFA | S695 | 0.056 | 90.36 | 795 | 0.044515 | 0.056 | 0.0445 | 0.8571 |
| Q9HD20 | ATP13A1 | VLDLQADsRLHVIFG | S379 | 0.155 | 78.4 | 1204 | 0.123212 | 0.155 | 0.1232 | 0.8 |
| P32929 | CTH | ESLGGFEsLAELPAI | S346 | 0.003 | 98.91 | 405 | 0.002385 | 0.003 | 0.0024 | 1 |
| P32929 | CTH | SDTLIRLsVGLEDEE | S377 | 0.023 | 98.89 | 405 | 0.018283 | 0.023 | 0.0183 | 0.9167 |
| Q8NEA9 | GMCL2 | GFDLLVIyTNGYIIF | Y424 | 0 | 96 | 526 | 0 | 0 | 0 | 1 |
| Q9UNE2 | RPH3AL | TKCGIEAsPGQKRPL | S128 | 0.18 | 84.08 | 315 | 0.143085 | 0.18 | 0.1431 | 1 |

| | | | | | | | | | | |
|---|---|---|---|---|---|---|---|---|---|---|
| Q8NCN5 | PDPR | ADySNKLyyQLEQET | Y109 | 0.002 | 98.05 | 879 | 0.000877 | 0.002 | 0.0009 | 1 |
| Q8NCN5 | PDPR | LNVIGIPsEIIsPkk | S154 | 0.157 | 98.28 | 879 | 0.124802 | 0.157 | 0.1248 | 1 |
| Q8NCN5 | PDPR | NWQGGILsGGFEKNP | S302 | 0.018 | 98.09 | 879 | 0.014308 | 0.018 | 0.0143 | 1 |
| Q8NCN5 | PDPR | MEVMPLMyDLKVPrW | Y442 | 0.259 | 93.69 | 879 | 0.11362 | 0.259 | 0.1136 | 1 |
| Q8NCN5 | PDPR | DFINRGEyEIDIAGY | Y840 | 0.02 | 98.28 | 879 | 0.008774 | 0.02 | 0.0088 | 1 |
| P49279 | SLC11A1 | RSCAILPtVLVAVFR | T412 | 0.01 | 84.3 | 550 | 0.00683 | 0.129 | 0.0881 | 1 |
| P09871 | C1S | PNDKTKFyAAGLVSW | Y648 | 0.059 | 97.27 | 688 | 0.025882 | 0.059 | 0.0259 | 1 |
| Q6P1J6 | PLB1 | DGNLETHttLPNILK | T1139 | 0.045 | 96.48 | 1458 | 0.030736 | 0.045 | 0.0307 | 1 |
| P19801 | AOC1 | EEPPLFSsHKPRGDF | S284 | 0.005 | 97.64 | 751 | 0.003975 | 0.124 | 0.1109 | 1 |
| Q9Y5S8 | NOX1 | VRGQtEEsMNEsHPR | S234 | 0.019 | 91.42 | 564 | 0.015103 | 0.019 | 0.0151 | 1 |
| Q9Y5S8 | NOX1 | tEEsMNEsHPRKCAE | S238 | 0.018 | 90.6 | 564 | 0.014308 | 0.055 | 0.0437 | 1 |
| Q9Y5S8 | NOX1 | GKVGFLNyRLFLTGW | Y470 | 0.108 | 91.67 | 564 | 0.047378 | 0.108 | 0.0474 | 1 |
| O95342 | ABCB11 | ILLLDMAtsALDNEs | T586 | 0.092 | 76.34 | 1321 | 0.062837 | 0.092 | 0.0628 | 1 |
| Q99569 | PKP4 | DNDRVVssVAtALRN | S891 | 0.172 | 97.41 | 1192 | 0.136725 | 0.172 | 0.1367 | 0.875 |
| Q86UV5 | USP48 | QQRDLREyTQATIyV | Y859 | 0.22 | 85.16 | 1035 | 0.096511 | 0.22 | 0.0965 | 0.8333 |
| Q9NUT2 | ABCB8 | SSFKLVIsQGLRsCT | S271 | 0.135 | 89.67 | 735 | 0.107313 | 0.135 | 0.1073 | 1 |
| Q9NUT2 | ABCB8 | VIsQGLRsCTQVAGC | S276 | 0.022 | 91.65 | 735 | 0.017488 | 0.022 | 0.0175 | 1 |
| Q9NUT2 | ABCB8 | SMLSTRLtLLLMVAT | T295 | 0.136 | 86.08 | 735 | 0.09289 | 0.136 | 0.0929 | 1 |
| P61006 | RAB8A | RtITTAyyRGAMGIM | Y78 | 0.029 | 88.55 | 207 | 0.012722 | 0.029 | 0.0127 | 0.8333 |
| Q8TAT5 | NEIL3 | MkMIRDFsILFyRCR | S229 | 0 | 96.98 | 605 | 0 | 0 | 0 | 1 |
| O43823 | AKAP8 | CSVCKFRsFDDEEIQ | S399 | 0.051 | 79.96 | 692 | 0.040541 | 0.051 | 0.0405 | 0.8 |
| Q8WVY7 | UBLCP1 | EFYSkkNtIMFDDIG | T248 | 0.001 | 98.87 | 318 | 0.000683 | 0.001 | 0.0007 | 0.8 |
| P23497 | SP100 | AAADKQFyEKKAAKL | Y818 | 0.198 | 93.25 | 879 | 0.08686 | 0.198 | 0.0869 | 0.8333 |
| Q9HBH0 | RHOF | FPEHyAPsVFEKYTA | S51 | 0.161 | 87.87 | 211 | 0.127981 | 0.161 | 0.128 | 0.8333 |
| Q9ULS6 | KCNS2 | RHSTGLRsLGATLKY | S317 | 0.014 | 86.82 | 477 | 0.011129 | 0.014 | 0.0111 | 1 |
| P49754 | VPS41 | IGQLKAIsPYLPRGD | S446 | 0.048 | 91.86 | 854 | 0.038156 | 0.048 | 0.0382 | 1 |
| Q9ULM3 | YEATS2 | tIVVGNVsKYIPPDK | S216 | 0.018 | 95.81 | 1422 | 0.014308 | 0.018 | 0.0143 | 0.9 |
| Q5VU92 | DCAF12L1 | NQELGAVsLDGYFHL | S281 | 0.003 | 93.55 | 463 | 0.002385 | 0.047 | 0.0374 | 0.8889 |
| P49841 | GSK3B | DRPQEVSyTDTKVIG | Y56 | 0.196 | 97.18 | 420 | 0.085982 | 0.258 | 0.1132 | 1 |
| Q9UI30 | TRMT112 | PkGPVEGyEENEEFL | Y74 | 0.286 | 96.41 | 125 | 0.125464 | 0.286 | 0.1255 | 1 |
| P48454 | PPP3CC | RYLFLGDyVDRGYFS | Y115 | 0 | 98.9 | 512 | 0 | 0 | 0 | 1 |

| | | | | | | | | | | |
|---|---|---|---|---|---|---|---|---|---|---|
| Q9BQG0 | MYBBP1A | WRVVRFLsPPALQGY | S399 | 0.153 | 93.82 | 1328 | 0.121622 | 0.153 | 0.1216 | 1 |
| Q6MZP7 | LIN54 | GCKNFEEsPERktLM | S635 | 0 | 94.73 | 749 | 0 | 0 | 0 | 1 |
| Q9UKM7 | MAN1B1 | QMNRQMEtGLSPEIV | T565 | 0.001 | 98.59 | 699 | 0.000683 | 0.001 | 0.0007 | 1 |
| Q9H2S1 | KCNN2 | FEKRKRLsDYALIFG | S135 | 0.019 | 94.31 | 579 | 0.015103 | 0.019 | 0.0151 | 1 |
| P18887 | XRCC1 | RPDWTRDstHLICAF | S357 | 0.131 | 92 | 633 | 0.104134 | 0.131 | 0.1041 | 1 |
| P18887 | XRCC1 | LICAFANtPKysQVL | T367 | 0.166 | 84.26 | 633 | 0.11338 | 0.166 | 0.1134 | 1 |
| P18887 | XRCC1 | AFANtPKysQVLGLG | Y370 | 0.23 | 90.09 | 633 | 0.100898 | 0.23 | 0.1009 | 1 |
| Q8N6M5 | ALLC | EAIAELKsDDWSYLV | S129 | 0.052 | 97.66 | 391 | 0.041336 | 0.052 | 0.0413 | 1 |
| Q8ND71 | GIMAP8 | GKSAAGNsILGRQAF | S266 | 0.186 | 91.53 | 665 | 0.147854 | 0.186 | 0.1479 | 1 |
| Q8ND71 | GIMAP8 | SWRKKKVsIIDAPDI | S299 | 0.09 | 94.78 | 665 | 0.071542 | 0.09 | 0.0715 | 1 |
| Q96RR1 | TWNK | LKRFSVRyLRPARSL | Y197 | 0.06 | 96.63 | 684 | 0.026321 | 0.06 | 0.0263 | 1 |
| P31040 | SDHA | QRAFGGQsLkFGKGG | S177 | 0.16 | 98.59 | 664 | 0.127186 | 0.16 | 0.1272 | 1 |
| P31040 | SDHA | DQIVPGLyACGEAAC | Y436 | 0.082 | 98.93 | 664 | 0.035972 | 0.082 | 0.036 | 1 |
| P31040 | SDHA | HWRkHTLsyVDVGTG | S628 | 0.023 | 98.89 | 664 | 0.018283 | 0.023 | 0.0183 | 1 |
| A5PLN9 | TRAPPC13 | ILVCAVSyTTQAGEK | Y150 | 0.037 | 95.61 | 417 | 0.016231 | 0.037 | 0.0162 | 1 |
| O60733 | PLA2G6 | MAyMRGMyFRMKDEV | Y541 | 0 | 96.27 | 806 | 0 | 0 | 0 | 1 |
| P13866 | SLC5A1 | MVMPGMIsRILYTEK | S335 | 0 | 97.69 | 664 | 0 | 0 | 0 | 0.8182 |
| P13866 | SLC5A1 | ILVLIGIsIAWVPIV | S437 | 0 | 88.09 | 664 | 0 | 0 | 0 | 0.8 |
| Q53GD3 | SLC44A4 | KFLNRNAyIMIAIYG | Y560 | 0.05 | 98.17 | 710 | 0.021934 | 0.05 | 0.0219 | 1 |
| Q96CC6 | RHBDF1 | LGMVGRLtNRtYRKR | T380 | 0.169 | 78.13 | 855 | 0.115429 | 0.169 | 0.1154 | 1 |
| Q8NAA4 | ATG16L2 | RHQAVTGsRDRTVKE | S441 | 0 | 97.63 | 619 | 0 | 0 | 0 | 1 |
| Q8NAA4 | ATG16L2 | FKCGSDWtKAVFsPD | T552 | 0.006 | 95.43 | 619 | 0.004098 | 0.006 | 0.0041 | 0.8462 |
| Q68DD2 | PLA2G4F | DRRKQVVsKALQQVL | S336 | 0.063 | 96.72 | 849 | 0.05008 | 0.063 | 0.0501 | 1 |
| Q68DD2 | PLA2G4F | PVVAVLGsGGGtRAM | S362 | 0.004 | 94.94 | 849 | 0.00318 | 0.004 | 0.0032 | 1 |
| Q68DD2 | PLA2G4F | VLGsGGGtRAMSSLY | T366 | 0.058 | 90.32 | 849 | 0.039615 | 0.058 | 0.0396 | 1 |
| P22694 | PRKACB | EDFERkktLGtGsFG | T49 | 0.001 | 97.21 | 351 | 0.000683 | 0.001 | 0.0007 | 1 |
| P22694 | PRKACB | LLIDHQGyIQVtDFG | Y180 | 0.035 | 98.77 | 351 | 0.015354 | 0.035 | 0.0154 | 0.9286 |
| P22694 | PRKACB | AKRVKGRtWtLCGtP | T196 | 0.072 | 97.16 | 351 | 0.049177 | 0.143 | 0.0977 | 1 |
| P22694 | PRKACB | tLCGtPEyLAPEIIL | Y205 | 0.189 | 98.53 | 351 | 0.082912 | 0.189 | 0.0829 | 1 |
| O94856 | NFASC | YHVyENGsLEIKMIR | S485 | 0.004 | 87.8 | 1347 | 0.00318 | 0.004 | 0.0032 | 1 |
| Q3SYG4 | BBS9 | SHKItIDtNKSPVSL | T552 | 0.145 | 92.69 | 887 | 0.099037 | 0.145 | 0.099 | 1 |

| | | | | | | | | | | |
|---|---|---|---|---|---|---|---|---|---|---|
| P18754 | RCC1 | DSHtAALtDDGRVFL | T135 | 0.011 | 98.34 | 421 | 0.007513 | 0.011 | 0.0075 | 1 |
| P18754 | RCC1 | SNYHQLGtPGTESCF | T274 | 0.124 | 94.4 | 421 | 0.084694 | 0.124 | 0.0847 | 0.8571 |
| Q86SF2 | GALNT7 | LWKRVPLtPQEkRLR | T368 | 0.108 | 98.12 | 657 | 0.073765 | 0.108 | 0.0738 | 1 |
| P41235 | HNF4A | SIADVCEsMKEQLLV | S190 | 0.065 | 95.06 | 474 | 0.051669 | 0.065 | 0.0517 | 1 |
| P41235 | HNF4A | LAEMsRVsIRILDEL | S265 | 0.085 | 96.52 | 474 | 0.067568 | 0.085 | 0.0676 | 1 |
| Q9Y2V7 | COG6 | FGTARRstVVRGFID | T276 | 0.094 | 95.09 | 657 | 0.064203 | 0.094 | 0.0642 | 0.8889 |
| Q9Y2V7 | COG6 | TLALFEFtDRRLEML | T518 | 0.018 | 89.53 | 657 | 0.012294 | 0.018 | 0.0123 | 1 |
| Q9Y2V7 | COG6 | VMNPINEyKDPENIL | Y638 | 0.274 | 84.25 | 657 | 0.1202 | 0.274 | 0.1202 | 1 |
| Q6V0I7 | FAT4 | APPGSYVsGIsATDG | S495 | 0.134 | 89.15 | 4981 | 0.106519 | 0.134 | 0.1065 | 1 |
| Q92187 | ST8SIA4 | KPGDVIHyVLDRRRT | Y110 | 0.068 | 94.77 | 359 | 0.029831 | 0.068 | 0.0298 | 1 |
| Q92187 | ST8SIA4 | KNKLKVRtAYPSLRL | T253 | 0.061 | 98.33 | 359 | 0.041664 | 0.061 | 0.0417 | 1 |
| Q96MG8 | PCMTD1 | EGyRDNAykDLAWKH | Y51 | 0.259 | 96.44 | 357 | 0.11362 | 0.259 | 0.1136 | 0.9167 |
| Q5BKZ1 | ZNF326 | VNkFKKTsIRkQQTN | S369 | 0.128 | 80.19 | 582 | 0.101749 | 0.128 | 0.1017 | 1 |
| O14732 | IMPA2 | GPkRDPAtLkLFLSN | T182 | 0.152 | 95.54 | 288 | 0.103818 | 0.152 | 0.1038 | 1 |
| O60763 | USO1 | VRALksLsKKYRLEV | S50 | 0.002 | 93.55 | 962 | 0.00159 | 0.002 | 0.0016 | 1 |
| P40227 | CCT6A | GLHPRIItEGFEAAk | T120 | 0.123 | 94.21 | 531 | 0.084011 | 0.123 | 0.084 | 1 |
| P40227 | CCT6A | KHksETDtsLIRGLV | T204 | 0.139 | 90.69 | 531 | 0.094939 | 0.139 | 0.0949 | 1 |
| P40227 | CCT6A | PFsLDALskEGIVAL | S306 | 0.158 | 91.47 | 531 | 0.125597 | 0.158 | 0.1256 | 1 |
| P40227 | CCT6A | IkGPNkHtLTQIkDA | T383 | 0.188 | 91.34 | 531 | 0.128407 | 0.188 | 0.1284 | 1 |
| Q03426 | MVK | AVSLNLRtFLRLQPH | T37 | 0.029 | 97.82 | 396 | 0.019807 | 0.029 | 0.0198 | 1 |
| Q9Y2S7 | POLDIP2 | ERHWRIFsLsGtLEt | S288 | 0.091 | 95.25 | 368 | 0.072337 | 0.091 | 0.0723 | 1 |
| Q05329 | GAD2 | ALGIGtDsVILIKCD | S298 | 0.033 | 98.42 | 585 | 0.026232 | 0.033 | 0.0262 | 1 |
| Q05329 | GAD2 | EYGTTMVsyQPLGDK | S546 | 0.122 | 97.95 | 585 | 0.09698 | 0.122 | 0.097 | 1 |
| Q9UQM7 | CAMK2A | CTRFTEEyQLFEELG | Y13 | 0.17 | 96.62 | 478 | 0.074577 | 0.17 | 0.0746 | 1 |
| Q9UQM7 | CAMK2A | sCMHRQEtVDCLKKF | T286 | 0.054 | 91.78 | 478 | 0.036883 | 0.054 | 0.0369 | 1 |
| P51160 | PDE6C | KVPVEVLtRWMYTVR | T546 | 0.004 | 96.32 | 858 | 0.002732 | 0.004 | 0.0027 | 1 |
| Q08881 | ITK | ENQVIkVsDFGMtRF | S499 | 0.101 | 92.84 | 620 | 0.080286 | 0.089 | 0.111 | 1 |
| P31947 | SFN | LssIEQksNEEGsEE | S69 | 0.103 | 87.34 | 248 | 0.081876 | 0.103 | 0.0819 | 1 |
| P31947 | SFN | IDsARsAyQEAMDIS | Y151 | 0 | 98.83 | 248 | 0 | 0 | 0 | 1 |
| P31947 | SFN | sEDSYkDstLIMQLL | S216 | 0 | 96.54 | 248 | 0 | 0 | 0 | 1 |
| P49406 | MRPL19 | EPERRFLsPEFIPRR | S77 | 0.095 | 94.73 | 292 | 0.075517 | 0.095 | 0.0755 | 0.8333 |

| | | | | | | | | | | |
|---|---|---|---|---|---|---|---|---|---|---|
| Q96AE4 | FUBP1 | kIQIAPDsGGLPERs | S140 | 0.131 | 72.94 | 644 | 0.104134 | 0.131 | 0.1041 | 0.8571 |
| Q96AE4 | FUBP1 | MLtGtPEsVQsAkRL | S156 | 0.187 | 94.62 | 644 | 0.148649 | 0.187 | 0.1486 | 1 |
| Q5VYJ5 | MALRD1 | GPTPTLNtGPMKDNT | T896 | 0.056 | 81.63 | 2156 | 0.038249 | 0.056 | 0.0382 | 1 |
| P47900 | P2RY1 | LVLILGCyGLIVRAL | Y237 | 0.083 | 91.44 | 373 | 0.036411 | 0.083 | 0.0364 | 1 |
| P52948 | NUP98 | EGRLEAVsRkQGAQF | S856 | 0.009 | 89.52 | 1817 | 0.007154 | 0.009 | 0.0072 | 1 |
| Q9BV73 | CEP250 | LHHsLsHsLLAVAQA | S2394 | 0.025 | 81.35 | 2442 | 0.019873 | 0.025 | 0.0199 | 1 |
| O60674 | JAK2 | kFGSLDTyLKKNKNC | Y637 | 0.255 | 93.14 | 1132 | 0.111865 | 0.255 | 0.1119 | 1 |
| Q9P265 | DIP2B | KRAERIAsVLGDKGH | S1035 | 0.035 | 90.63 | 1576 | 0.027822 | 0.035 | 0.0278 | 1 |
| Q9HCJ6 | VAT1L | CTPVEFVyKIPDDMS | Y147 | 0.087 | 98.82 | 419 | 0.038166 | 0.087 | 0.0382 | 0.9167 |
| Q9HCJ6 | VAT1L | PMNFVTAyVMLFEVA | Y169 | 0.274 | 98.9 | 419 | 0.1202 | 0.274 | 0.1202 | 1 |
| Q9H7E9 | C8orf33 | MHSLFGDyRAQMEAE | Y166 | 0.194 | 96.35 | 229 | 0.085105 | 0.194 | 0.0851 | 0.8 |
| Q49AJ0 | FAM135B | DLFQRGYyQIRVTLK | Y28 | 0.029 | 93.42 | 1406 | 0.012722 | 0.058 | 0.0254 | 1 |
| Q8J025 | APCDD1 | FKAYQFYyGSNRCTN | Y110 | 0.141 | 98.55 | 514 | 0.061855 | 0.141 | 0.0619 | 1 |
| P52292 | KPNA2 | FESAWALtNIASGtS | T145 | 0.013 | 97.61 | 529 | 0.008879 | 0.013 | 0.0089 | 1 |
| P52292 | KPNA2 | MSSLACGyLRNLTWT | Y225 | 0.238 | 96.41 | 529 | 0.104407 | 0.238 | 0.1044 | 1 |
| P52292 | KPNA2 | kEATWTMsNItAGRQ | S360 | 0 | 96.79 | 529 | 0 | 0.058 | 0.0461 | 1 |
| P52292 | KPNA2 | TWTMsNItAGRQDQI | T363 | 0.005 | 96.22 | 529 | 0.003415 | 0.005 | 0.0034 | 1 |
| Q9Y5X3 | SNX5 | EVFLQRLssHPVLsK | S151 | 0.061 | 95.39 | 404 | 0.04849 | 0.117 | 0.093 | 1 |
| Q9Y5X3 | SNX5 | LssHPVLsKDRNFHV | S157 | 0.161 | 95.6 | 404 | 0.127981 | 0.161 | 0.128 | 1 |
| Q8WVM8 | SCFD1 | FWQKHkGsPFPEVAE | S332 | 0.143 | 95.05 | 642 | 0.113673 | 0.143 | 0.1137 | 1 |
| P16401 | H1-5 | kGtGAsGsFkLNkkA | S107 | 0.161 | 96.21 | 226 | 0.127981 | 0.161 | 0.128 | 0.8571 |
| P07202 | TPO | QLMNEELtERLFVLS | T561 | 0.015 | 94.18 | 933 | 0.010245 | 0.015 | 0.0102 | 1 |
| P15088 | CPA3 | sWNsIPNtNDPCADN | T264 | 0.069 | 94.63 | 417 | 0.047128 | 0.069 | 0.0471 | 0.875 |
| P15088 | CPA3 | NDPCADNyRGSAPES | Y272 | 0.226 | 97.12 | 417 | 0.099143 | 0.226 | 0.0991 | 1 |
| P15088 | CPA3 | STRYETRyIYGPIES | Y346 | 0.166 | 98.68 | 417 | 0.072822 | 0.166 | 0.0728 | 1 |
| Q29983 | MICA | CLQELRRyLKsGVVL | Y194 | 0.001 | 93.21 | 383 | 0.000439 | 0.001 | 0.0004 | 1 |
| Q9NYQ8 | FAT2 | DREsEPRyILKVEAR | Y889 | 0.121 | 85.84 | 4349 | 0.053081 | 0.121 | 0.0531 | 1 |
| Q15111 | PLCL1 | YPEDFVNyNKKFLSR | Y634 | 0.008 | 96.43 | 1095 | 0.003509 | 0.008 | 0.0035 | 1 |
| Q8N1V2 | CFAP52 | QLKRIVMsIGVDDDD | S210 | 0.035 | 94.83 | 620 | 0.027822 | 0.035 | 0.0278 | 1 |
| Q8N1V2 | CFAP52 | VFCKsPGyKPIKKIQ | Y281 | 0.003 | 90.09 | 620 | 0.001316 | 0.003 | 0.0013 | 1 |
| Q8N1V2 | CFAP52 | QGGItSItLRGEGHQ | T297 | 0.119 | 94.01 | 620 | 0.081279 | 0.119 | 0.0813 | 1 |

| | | | | | | | | | | |
|---|---|---|---|---|---|---|---|---|---|---|
| P43351 | RAD52 | EDVGYGVsEGLkSkA | S129 | 0.007 | 97.87 | 418 | 0.005564 | 0.007 | 0.0056 | 1 |
| P43351 | RAD52 | GLkSkALsLEkARKE | S138 | 0 | 98.35 | 418 | 0 | 0 | 0 | 1 |
| P43351 | RAD52 | NCILDKDyLRsLNkL | Y171 | 0.109 | 97.95 | 418 | 0.047817 | 0.109 | 0.0478 | 1 |
| P35749 | MYH11 | SEKIVDMykGKkRHE | Y142 | 0.018 | 92.05 | 1972 | 0.007896 | 0.174 | 0.0763 | 1 |
| P35749 | MYH11 | HEMPPHIyAIADTAy | Y155 | 0 | 94.93 | 1972 | 0 | 0 | 0 | 1 |
| P35749 | MYH11 | NTKKVIQyLAVVASS | Y194 | 0.003 | 92.63 | 1972 | 0.001316 | 0.003 | 0.0013 | 1 |
| P35749 | MYH11 | DPLNDNVtSLLNASS | T604 | 0.019 | 86.63 | 1972 | 0.012977 | 0.14 | 0.0956 | 1 |
| O95487 | SEC24B | EFIASSDyMLRPPQP | Y670 | 0.305 | 92.98 | 1268 | 0.133799 | 0.305 | 0.1338 | 1 |
| Q7Z7B1 | PIGW | LIVPMVAtLTIWASF | T71 | 0.024 | 95.13 | 504 | 0.016392 | 0.024 | 0.0164 | 1 |
| Q8IYB7 | DIS3L2 | SAGALNKsLTQTFGD | S643 | 0.095 | 93.09 | 885 | 0.075517 | 0.095 | 0.0755 | 1 |
| Q8IYB7 | DIS3L2 | LARKEVLtNMCSRPM | T662 | 0.077 | 92.94 | 885 | 0.052592 | 0.077 | 0.0526 | 1 |
| Q8IYB7 | DIS3L2 | FDVLVLRyGVQKRIY | Y789 | 0.303 | 93.05 | 885 | 0.132922 | 0.303 | 0.1329 | 1 |
| Q96KN1 | LRATD2 | NSFLTDAsQGRRGRV | S159 | 0.008 | 96.42 | 310 | 0.006359 | 0.008 | 0.0064 | 1 |
| O94889 | KLHL18 | LVDEAkDyHLMPERR | Y263 | 0.203 | 89.54 | 574 | 0.089053 | 0.203 | 0.0891 | 0.8 |
| O94889 | KLHL18 | HtRRSRVsLVAsCGR | S518 | 0.164 | 97.86 | 574 | 0.130366 | 0.164 | 0.1304 | 1 |
| O94889 | KLHL18 | SRVsLVAsCGRLYAV | S522 | 0.021 | 91.09 | 574 | 0.016693 | 0.021 | 0.0167 | 0.8 |
| P07332 | FES | PLHEQLWyHGAIPRA | Y461 | 0.15 | 97.67 | 822 | 0.065803 | 0.15 | 0.0658 | 0.9091 |
| P07332 | FES | EARILKQySHPNIVR | Y614 | 0.168 | 95.1 | 822 | 0.073699 | 0.265 | 0.091 | 1 |
| Q96CM3 | RPUSD4 | VKKyWAItVHVPMPs | T189 | 0.009 | 98.56 | 377 | 0.006147 | 0.009 | 0.0061 | 1 |
| Q96CM3 | RPUSD4 | QHHkMTLsPSYRMDD | S221 | 0.174 | 94.22 | 377 | 0.138315 | 0.174 | 0.1383 | 1 |
| O14530 | TXNDC9 | LHIkVIPtLALLkDG | T145 | 0.048 | 96.34 | 226 | 0.032785 | 0.048 | 0.0328 | 1 |
| P21291 | CSRP1 | AVHGEEIyCkSCyGK | Y57 | 0.12 | 90.66 | 193 | 0.052642 | 0.12 | 0.0526 | 1 |
| P11586 | MTHFD1 | kERAsFItPVPGGVG | T269 | 0.01 | 96.9 | 935 | 0.00683 | 0.01 | 0.0068 | 1 |
| P11586 | MTHFD1 | PGGVGPMtVAMLMQS | T279 | 0.165 | 94.76 | 935 | 0.112697 | 0.165 | 0.1127 | 1 |
| P11586 | MTHFD1 | FACVRQPsQGPTFGI | S413 | 0.042 | 91.49 | 935 | 0.033386 | 0.042 | 0.0334 | 1 |
| P11586 | MTHFD1 | SKKGEPVsAEDLGVs | S601 | 0.044 | 96.76 | 935 | 0.034976 | 0.044 | 0.035 | 1 |
| P11586 | MTHFD1 | sAEDLGVsGALtVLM | S608 | 0.001 | 95.48 | 935 | 0.000795 | 0.001 | 0.0008 | 1 |
| P11586 | MTHFD1 | LGVsGALtVLMKDAI | T612 | 0.19 | 98.13 | 935 | 0.129773 | 0.19 | 0.1298 | 1 |
| P11586 | MTHFD1 | EsELDLIsRLSREHG | S771 | 0.111 | 98.33 | 935 | 0.088236 | 0.111 | 0.0882 | 1 |
| P11586 | MTHFD1 | RIIAQkIyGADDIEL | Y834 | 0 | 97.69 | 935 | 0 | 0 | 0 | 1 |
| P11586 | MTHFD1 | CMAkTHLsLsHNPEQ | S870 | 0.014 | 94.83 | 935 | 0.011129 | 0.014 | 0.0111 | 1 |

| | | | | | | | | | | |
|---|---|---|---|---|---|---|---|---|---|---|
| P11586 | MTHFD1 | AkTHLsLsHNPEQKG | S872 | 0 | 95.3 | 935 | 0 | 0 | 0 | 1 |
| P09105 | HBQ1 | HLDLsPGsSQVRAHG | S53 | 0.067 | 97.54 | 142 | 0.053259 | 0.067 | 0.0533 | 1 |
| Q4G0N4 | NADK2 | kPVIGVNtDPERsEG | T183 | 0.002 | 97.85 | 442 | 0.001366 | 0.002 | 0.0014 | 1 |
| Q4G0N4 | NADK2 | ESLSSRAsYYEIsVD | S289 | 0.007 | 97.21 | 442 | 0.005564 | 0.007 | 0.0056 | 1 |
| Q4G0N4 | NADK2 | RAsYYEIsVDDGPWE | S294 | 0.067 | 95.87 | 442 | 0.053259 | 0.067 | 0.0533 | 1 |
| Q4G0N4 | NADK2 | RELVEkVtNEYNEsL | T357 | 0.059 | 98.43 | 442 | 0.040298 | 0.059 | 0.0403 | 1 |
| Q4G0N4 | NADK2 | EEPkILFsIREPIAN | S376 | 0.028 | 97.09 | 442 | 0.022258 | 0.028 | 0.0223 | 1 |
| Q12864 | CDH17 | YNLTIEVsDKDFKTL | S425 | 0.145 | 95.2 | 832 | 0.115263 | 0.145 | 0.1153 | 1 |
| Q12864 | CDH17 | TGSSKILyHIIKGDs | Y489 | 0.164 | 96.31 | 832 | 0.071944 | 0.164 | 0.0719 | 1 |
| P08754 | GNAI3 | LLLGAGEsGkSTIVk | S44 | 0.031 | 97.68 | 354 | 0.024642 | 0.031 | 0.0246 | 1 |
| P08754 | GNAI3 | DrIsQSNyIPTQQDV | Y167 | 0.122 | 96.05 | 354 | 0.05352 | 0.122 | 0.0535 | 1 |
| Q8NI17 | IL31RA | RLENIAKtEPPKIFR | T122 | 0.097 | 87.1 | 732 | 0.066252 | 0.097 | 0.0663 | 0.9167 |
| Q9UPW5 | AGTPBP1 | GILQSLksVtNIkLG | S278 | 0.139 | 94.36 | 1226 | 0.110493 | 0.139 | 0.1105 | 1 |
| Q9UPW5 | AGTPBP1 | LQSLksVtNIkLGRK | T280 | 0.019 | 93.76 | 1226 | 0.012977 | 0.019 | 0.013 | 1 |
| O95479 | H6PD | QLKTAEDyQALNkDI | Y117 | 0.062 | 97.86 | 791 | 0.027199 | 0.062 | 0.0272 | 1 |
| O95479 | H6PD | VVGQYQSySEQVRRE | Y316 | 0 | 89.35 | 791 | 0 | 0 | 0 | 1 |
| O95479 | H6PD | IGHGDLGsPAVLVSR | S409 | 0.133 | 91.77 | 791 | 0.105724 | 0.133 | 0.1057 | 1 |
| O95479 | H6PD | PLSDYYAysPVRERD | Y449 | 0.242 | 98.08 | 791 | 0.106162 | 0.242 | 0.1062 | 1 |
| O95479 | H6PD | LSDYYAysPVRERDA | S450 | 0.109 | 96.29 | 791 | 0.086646 | 0.109 | 0.0866 | 1 |
| Q3ZCQ8 | TIMM50 | DRVLLDLsAFLKTIA | S280 | 0.025 | 96.27 | 353 | 0.019873 | 0.025 | 0.0199 | 1 |
| Q3ZCQ8 | TIMM50 | VRTVLEHyALEDDPL | Y301 | 0.284 | 89.46 | 353 | 0.124587 | 0.284 | 0.1246 | 1 |
| O00299 | CLIC1 | LAALNPEsNTAGLDI | S103 | 0.019 | 97.91 | 241 | 0.015103 | 0.019 | 0.0151 | 0.9091 |
| O00299 | CLIC1 | sAEDEGVsQRkFLDG | S163 | 0.016 | 93.78 | 241 | 0.012719 | 0.016 | 0.0127 | 1 |
| O00299 | CLIC1 | AFrGVHRyLsNAyAR | Y209 | 0.013 | 98.61 | 241 | 0.005703 | 0.013 | 0.0057 | 0.875 |
| A4FU01 | MTMR11 | SWHHPGGsDLLRCGG | S267 | 0.009 | 92.31 | 709 | 0.007154 | 0.009 | 0.0072 | 1 |
| Q8TCJ2 | STT3B | ALVGKAMssNEtAAy | S639 | 0.031 | 93.29 | 826 | 0.024642 | 0.031 | 0.0246 | 1 |
| Q8TCJ2 | STT3B | LVGKAMssNEtAAyk | S640 | 0.181 | 93.62 | 826 | 0.14388 | 0.181 | 0.1439 | 1 |
| Q8TCJ2 | STT3B | ssNEtAAykIMRTLD | Y646 | 0.317 | 94.96 | 826 | 0.139063 | 0.317 | 0.1391 | 1 |
| Q8TCJ2 | STT3B | kDIREsDyFTPQGEF | Y695 | 0.091 | 93.08 | 826 | 0.03992 | 0.091 | 0.0399 | 1 |
| Q8TCJ2 | STT3B | TLLNCLMykMsYYRF | Y718 | 0.045 | 94.39 | 826 | 0.019741 | 0.045 | 0.0197 | 1 |
| Q8TCJ2 | STT3B | NCLMykMsYYRFGEM | S721 | 0 | 93.15 | 826 | 0 | 0 | 0 | 1 |

| | | | | | | | | | | |
|---|---|---|---|---|---|---|---|---|---|---|
| Q16644 | MAPKAPK3 | kYAVTDDyQLSkQVL | Y43 | 0.069 | 98.44 | 382 | 0.030269 | 0.069 | 0.0303 | 1 |
| Q5T5U3 | ARHGAP21 | KKPtAtGtFGVRLDD | T1144 | 0.015 | 87.93 | 1958 | 0.010245 | 0.134 | 0.0915 | 1 |
| Q96PC2 | IP6K3 | MRKCAQstsACLGVR | T243 | 0.015 | 96.64 | 410 | 0.010245 | 0.015 | 0.0102 | 1 |
| Q13490 | BIRC2 | VCMDKEVsVVFIPCG | S580 | 0.14 | 90.42 | 618 | 0.111288 | 0.14 | 0.1113 | 1 |
| Q9BRS8 | LARP6 | YVsVKLLtSFKKVKH | T133 | 0.14 | 93.33 | 491 | 0.095622 | 0.14 | 0.0956 | 1 |
| Q7Z589 | EMSY | RKLELEAyAGVISAL | Y28 | 0 | 97.9 | 1322 | 0 | 0 | 0 | 1 |
| Q969N2 | PIGT | GQGKKDWsLFRMFSR | S259 | 0.164 | 92.52 | 578 | 0.130366 | 0.164 | 0.1304 | 1 |
| Q53G59 | KLHL12 | MPLLTPRyITDVIDA | Y231 | 0.213 | 90.23 | 568 | 0.09344 | 0.213 | 0.0934 | 1 |
| Q9NYH9 | UTP6 | LAVIGADsVTLkNkY | S473 | 0.067 | 75.66 | 597 | 0.053259 | 0.067 | 0.0533 | 1 |
| Q14149 | MORC3 | DHyGIMMyHRNRLIk | Y318 | 0 | 97.87 | 939 | 0 | 0 | 0 | 1 |
| Q16665 | HIF1A | DGDMIYIsDNVNKYM | S113 | 0.008 | 95.6 | 826 | 0.006359 | 0.008 | 0.0064 | 0.875 |
| Q5T8I9 | HENMT1 | EkAkIENsPTPFCVG | S329 | 0.112 | 95.46 | 393 | 0.08903 | 0.112 | 0.089 | 1 |
| O75635 | SERPINB7 | QERKFNLsVIEDPsM | S217 | 0.057 | 96.08 | 380 | 0.04531 | 0.057 | 0.0453 | 1 |
| O75635 | SERPINB7 | FQNLMEWtNPRRMTS | T262 | 0.007 | 94.99 | 380 | 0.004781 | 0.136 | 0.0929 | 1 |
| Q05940 | SLC18A2 | QPEsQKGtPLttLLK | T283 | 0.211 | 80.95 | 514 | 0.144116 | 0.211 | 0.1441 | 1 |
| Q15569 | TESK1 | KIGAGFFsEVYKVRH | S69 | 0.127 | 83.24 | 626 | 0.100954 | 0.127 | 0.101 | 1 |
| Q6SPF0 | SAMD1 | VsykGsIsyRNAARV | S92 | 0.007 | 95.44 | 538 | 0.005564 | 0.007 | 0.0056 | 1 |
| Q6S8J7 | POTEA | SKNkGGLtPLLLAVH | T200 | 0.015 | 98.03 | 498 | 0.010245 | 0.015 | 0.0102 | 1 |
| Q17RY0 | CPEB4 | AFSNQQSyIAAISAR | Y635 | 0.059 | 97.05 | 729 | 0.025882 | 0.059 | 0.0259 | 0.8462 |
| Q16281 | CNGA3 | VDPSSNLyyRWLTAI | Y168 | 0.099 | 91.78 | 694 | 0.04343 | 0.099 | 0.0434 | 1 |
| Q69YN4 | VIRMA | EkLVLEHskDDDNLD | S1464 | 0.118 | 83.39 | 1812 | 0.0938 | 0.118 | 0.0938 | 1 |
| Q6UXI7 | VIT | kVAGQSGyKGSYSNG | Y113 | 0.124 | 93.5 | 678 | 0.054397 | 0.124 | 0.0544 | 1 |
| P23975 | SLC6A2 | LWKGVKtsGKVVWIT | S259 | 0.082 | 93.84 | 617 | 0.065183 | 0.082 | 0.0652 | 0.8182 |
| P17028 | ZNF24 | rQFGYQDsPGPREAV | S63 | 0.169 | 71.19 | 368 | 0.134341 | 0.169 | 0.1343 | 1 |
| O75443 | TECTA | SFPKFVVtAKNEDRD | T1144 | 0.041 | 94.46 | 2155 | 0.028004 | 0.041 | 0.028 | 1 |
| Q9HCM9 | TRIM39 | RLWNGDKyAATTTPF | Y439 | 0.177 | 98.13 | 518 | 0.077647 | 0.177 | 0.0776 | 1 |
| Q9H583 | HEATR1 | HQLASISsPVVTSLL | S917 | 0.086 | 83.23 | 2144 | 0.068363 | 0.157 | 0.1248 | 1 |
| Q9H583 | HEATR1 | SLLINLGsPVKEVRR | S929 | 0.178 | 88.29 | 2144 | 0.141495 | 0.178 | 0.1415 | 0.8333 |
| Q14457 | BECN1 | LVPyGNHsYLESLTD | S337 | 0.005 | 92.38 | 450 | 0.003975 | 0.005 | 0.004 | 1 |
| P61764 | STXBP1 | LPYESQVySLDSADS | Y145 | 0.009 | 96.81 | 594 | 0.003948 | 0.009 | 0.0039 | 1 |
| Q969Z3 | MTARC2 | DPSERELyKLSPLFG | Y307 | 0.133 | 96.09 | 335 | 0.058345 | 0.133 | 0.0583 | 0.8182 |

| | | | | | | | | | | |
|---|---|---|---|---|---|---|---|---|---|---|
| Q7Z3Q1 | SLC46A3 | GLIPGLVsTFILLsI | S87 | 0.092 | 94.27 | 461 | 0.073132 | 0.148 | 0.1217 | 1 |
| P21964 | COMT | VLEAIDtyCEQkEWA | Y82 | 0.163 | 98.2 | 271 | 0.071506 | 0.163 | 0.0715 | 1 |
| P21964 | COMT | DGLEkAIykGPGsEA | Y262 | 0.234 | 98.57 | 271 | 0.102652 | 0.305 | 0.1338 | 1 |
| P52849 | NDST2 | TNPADRAysWyQHQR | Y710 | 0.211 | 97.72 | 883 | 0.092563 | 0.211 | 0.0926 | 1 |
| P52849 | NDST2 | ADRAysWyQHQRAHG | Y713 | 0.113 | 97.55 | 883 | 0.049571 | 0.113 | 0.0496 | 1 |
| P52849 | NDST2 | YPDMDTEsRLFLTDF | S843 | 0 | 97.9 | 883 | 0 | 0 | 0 | 1 |
| P98160 | HSPG2 | PQLEDAGsREFREVS | S105 | 0.13 | 85.41 | 4391 | 0.103339 | 0.13 | 0.1033 | 1 |
| Q9H341 | OR51M1 | IFRGPVAtIPIVLLL | T168 | 0.056 | 91.72 | 326 | 0.038249 | 0.056 | 0.0382 | 1 |
| Q9NR63 | CYP26B1 | AVELASTsRFELATR | S462 | 0.011 | 85.96 | 512 | 0.008744 | 0.113 | 0.0898 | 1 |
| O00154 | ACOT7 | TNkAtLWyVPLSLkN | Y162 | 0.006 | 98.18 | 380 | 0.002632 | 0.006 | 0.0026 | 0.875 |
| O00154 | ACOT7 | MTFTSNKsMEIEVLV | S303 | 0.164 | 97.97 | 380 | 0.130366 | 0.164 | 0.1304 | 1 |
| O60437 | PPL | ALPGSGTtPLLRTRV | T583 | 0.05 | 79.97 | 1756 | 0.034151 | 0.05 | 0.0342 | 1 |
| O14512 | SOCS7 | DSSSFAAsLRELEKC | S391 | 0.025 | 83.87 | 581 | 0.019873 | 0.071 | 0.0564 | 1 |
| Q9Y2P5 | SLC27A5 | HERVLQMsKMLSLSG | S316 | 0.004 | 94.24 | 690 | 0.00318 | 0.004 | 0.0032 | 0.8889 |
| Q9Y2P5 | SLC27A5 | TFDGEKLyQHVRAWL | Y613 | 0.189 | 96.04 | 690 | 0.082912 | 0.189 | 0.0829 | 1 |
| Q6ZXV5 | TMTC3 | FDNPAAVsPTPTRQL | S274 | 0.118 | 91.75 | 915 | 0.0938 | 0.118 | 0.0938 | 1 |
| Q6ZXV5 | TMTC3 | NFERALKyFLQAtHV | Y468 | 0.311 | 98.18 | 915 | 0.136431 | 0.311 | 0.1364 | 1 |
| Q6ZXV5 | TMTC3 | LKyFLQAtHVQPDDI | T473 | 0.214 | 96.3 | 915 | 0.146165 | 0.214 | 0.1462 | 1 |
| O15031 | PLXNB2 | DGRILKVyLTPDGTS | Y419 | 0.096 | 91.02 | 1838 | 0.042114 | 0.096 | 0.0421 | 1 |
| Q5T230 | UTF1 | LPTYRRVsAALAQQQ | S99 | 0.18 | 95.06 | 341 | 0.143085 | 0.18 | 0.1431 | 0.8571 |
| Q969S3 | ZNF622 | ALEFADFyDFRssyP | Y347 | 0.306 | 88.98 | 477 | 0.134238 | 0.306 | 0.1342 | 0.8333 |
| Q8WWQ2 | HPSE2 | LRFGGKRtDFLQFQN | T120 | 0.001 | 93.43 | 592 | 0.000683 | 0.001 | 0.0007 | 1 |
| Q8WWQ2 | HPSE2 | IYSRAsLyGPNIGRP | Y302 | 0.122 | 98.49 | 592 | 0.05352 | 0.323 | 0.1417 | 1 |
| Q8IU60 | DCP2 | yKMGVPTyGAIILDE | Y100 | 0.209 | 98.02 | 420 | 0.091685 | 0.209 | 0.0917 | 1 |
| P55000 | SLURP1 | TCKEPMTsAsCRTIt | S34 | 0.134 | 88.86 | 103 | 0.106519 | 0.134 | 0.1065 | 0.8571 |
| Q9Y2P4 | SLC27A6 | EKRGELVtVLDKFLS | T55 | 0.052 | 94.69 | 619 | 0.035517 | 0.052 | 0.0355 | 0.8571 |
| Q9Y2P4 | SLC27A6 | GEPGLLIsRVNAKNP | S426 | 0.043 | 93.44 | 619 | 0.034181 | 0.043 | 0.0342 | 1 |
| Q8N3D4 | EHBP1L1 | VDISVTLyRDPHVDQ | Y85 | 0.119 | 90.91 | 1523 | 0.052204 | 0.119 | 0.0522 | 0.9 |
| Q5K4L6 | SLC27A3 | IHGSRRFsYsEAERE | S92 | 0.132 | 98.61 | 683 | 0.104929 | 0.132 | 0.1049 | 1 |
| Q96QP1 | ALPK1 | FLyGLDVsGKLLQVA | S124 | 0.033 | 89.49 | 1244 | 0.026232 | 0.033 | 0.0262 | 0.875 |
| Q8N2G6 | ZCCHC24 | KKRCFGEyKCPKCKR | Y170 | 0.028 | 97.99 | 241 | 0.012283 | 0.028 | 0.0123 | 1 |

| | | | | | | | | | | |
|---|---|---|---|---|---|---|---|---|---|---|
| Q9NR96 | TLR9 | KLNLSFNyQKRVSFA | Y345 | 0.098 | 95.15 | 1032 | 0.042991 | 0.098 | 0.043 | 1 |
| P25774 | CTSS | IDNKGIDsDASYPYK | S200 | 0.177 | 98.78 | 331 | 0.1407 | 0.177 | 0.1407 | 0.875 |
| P34932 | HSPA4 | AEksNLAyDIVQLPt | Y89 | 0.11 | 95.34 | 840 | 0.048255 | 0.11 | 0.0483 | 1 |
| P34932 | HSPA4 | VQLPtGLtGIkVTYM | T99 | 0.041 | 93.02 | 840 | 0.028004 | 0.063 | 0.043 | 1 |
| P34932 | HSPA4 | MEEERNFttEQVtAM | T113 | 0.074 | 97.1 | 840 | 0.050543 | 0.14 | 0.0956 | 1 |
| P34932 | HSPA4 | EEERNFttEQVtAML | T114 | 0.019 | 97.04 | 840 | 0.012977 | 0.019 | 0.013 | 1 |
| P34932 | HSPA4 | NFttEQVtAMLLskL | T118 | 0.008 | 98.46 | 840 | 0.005464 | 0.008 | 0.0055 | 1 |
| P34932 | HSPA4 | YTDAERRsVMDATQI | S155 | 0.028 | 96.78 | 840 | 0.022258 | 0.028 | 0.0223 | 1 |
| P34932 | HSPA4 | ALQCAILsPAFkVRE | S384 | 0.065 | 88.72 | 840 | 0.051669 | 0.098 | 0.0815 | 1 |
| P34932 | HSPA4 | PQSDGSsskVKVKVR | S476 | 0.004 | 90.6 | 840 | 0.00318 | 0.004 | 0.0032 | 1 |
| P34932 | HSPA4 | RDkLsGEyEkFVsED | Y636 | 0.074 | 95.2 | 840 | 0.032463 | 0.074 | 0.0325 | 1 |
| P34932 | HSPA4 | EDQPkQVyVDkLAEL | Y671 | 0.005 | 97.99 | 840 | 0.002193 | 0.005 | 0.0022 | 1 |
| Q99805 | TM9SF2 | HVDIKIyyHVVETGS | Y218 | 0.275 | 91.77 | 663 | 0.120639 | 0.275 | 0.1206 | 1 |
| P08603 | CFH | PDRkKDQykVGEVLK | Y587 | 0.069 | 79.6 | 1231 | 0.030269 | 0.069 | 0.0303 | 0.8333 |
| Q7L211 | ABHD13 | ISQCRMPsLFISGLs | S260 | 0.001 | 98.57 | 337 | 0.000795 | 0.001 | 0.0008 | 1 |
| Q92954 | PRG4 | YDKCCPDyEsFCAEV | Y100 | 0.123 | 94.13 | 1404 | 0.053958 | 0.123 | 0.054 | 1 |
| P17706 | PTPN2 | ENDyINAsLVDIEEA | S72 | 0 | 98.93 | 415 | 0 | 0 | 0 | 0.8 |
| P31513 | FMO3 | HNSKIQEyIIAFAKE | Y90 | 0.05 | 98.83 | 532 | 0.021934 | 0.05 | 0.0219 | 0.8667 |
| P59190 | RAB15 | VRIQIWDtAGQERYQ | T64 | 0.017 | 84.84 | 212 | 0.011611 | 0.017 | 0.0116 | 1 |
| Q8IWS0 | PHF6 | sQGIyMVyCRKHKKt | Y127 | 0.277 | 95.01 | 365 | 0.121516 | 0.277 | 0.1215 | 1 |
| Q6NUM9 | RETSAT | GPNGRkEyPMySGEk | Y167 | 0.114 | 98.41 | 610 | 0.05001 | 0.114 | 0.05 | 1 |
| Q6NUM9 | RETSAT | RAGGAVLtkATVQSV | T316 | 0.018 | 97.7 | 610 | 0.012294 | 0.018 | 0.0123 | 1 |
| Q6NUM9 | RETSAT | LHLPSTNyYVYyDTD | Y408 | 0.238 | 98.44 | 610 | 0.104407 | 0.238 | 0.1044 | 1 |
| Q6NUM9 | RETSAT | MDQAMERyVSMPREE | Y423 | 0.019 | 96.65 | 610 | 0.008335 | 0.019 | 0.0083 | 1 |
| Q6NUM9 | RETSAT | KGkRGsDyETFKNSF | Y486 | 0.043 | 96.03 | 610 | 0.018863 | 0.043 | 0.0189 | 1 |
| Q9HCM4 | EPB41L5 | kAKWLEMyGVDMHVV | Y230 | 0.154 | 95.48 | 733 | 0.067558 | 0.154 | 0.0676 | 0.8462 |
| Q99677 | LPAR4 | RTLRKPAtLsQIGTN | T236 | 0.184 | 86.7 | 370 | 0.125675 | 0.184 | 0.1257 | 1 |
| P20020 | ATP2B1 | DkDPLLLsGTHVMEG | S262 | 0.109 | 90.29 | 1220 | 0.086646 | 0.109 | 0.0866 | 0.8889 |
| P20020 | ATP2B1 | VtAVGVNsQTGIIFT | S282 | 0.022 | 88.78 | 1220 | 0.017488 | 0.039 | 0.031 | 1 |
| P52701 | MSH6 | RTLLEEEyFREkLSD | Y642 | 0.013 | 93.71 | 1360 | 0.005703 | 0.099 | 0.0434 | 1 |
| P52701 | MSH6 | sMANFEEyIPLDSDT | Y709 | 0.275 | 95.29 | 1360 | 0.120639 | 0.275 | 0.1206 | 1 |

| | | | | | | | | | | |
|---|---|---|---|---|---|---|---|---|---|---|
| Q86WG5 | SBF2 | FENKILEtEktLPAA | T568 | 0.015 | 83.95 | 1849 | 0.010245 | 0.015 | 0.0102 | 1 |
| P54289 | CACNA2D1 | VTLAKTAsGVNQLVD | S52 | 0.035 | 94.33 | 1103 | 0.027822 | 0.035 | 0.0278 | 1 |
| P54289 | CACNA2D1 | SATGLARyyPASPWV | Y217 | 0.096 | 97.32 | 1103 | 0.042114 | 0.228 | 0.1 | 1 |
| P54289 | CACNA2D1 | ATGLARyyPASPWVD | Y218 | 0.078 | 97.75 | 1103 | 0.034217 | 0.078 | 0.0342 | 1 |
| Q13884 | SNTB1 | PQAGVDLsFAtRTGT | S397 | 0.111 | 90.56 | 538 | 0.088236 | 0.177 | 0.1407 | 1 |
| Q13884 | SNTB1 | KtIIQSPyEKLKMSS | Y483 | 0.146 | 91.72 | 538 | 0.064048 | 0.146 | 0.064 | 0.8462 |
| P60981 | DSTN | PLkSkMIyAsskDAI | Y117 | 0.102 | 90.34 | 165 | 0.044746 | 0.102 | 0.0447 | 1 |
| Q4G176 | ACSF3 | yTKLMEyyDrHFTQP | Y304 | 0.212 | 97.15 | 576 | 0.093001 | 0.212 | 0.093 | 1 |
| Q4G176 | ACSF3 | VVFkDGQyWIRGRTS | Y466 | 0.004 | 95.53 | 576 | 0.001755 | 0.004 | 0.0018 | 0.95 |
| O75251 | NDUFS7 | FRASPRQsDVMIVAG | S117 | 0 | 95.1 | 213 | 0 | 0 | 0 | 1 |
| P46459 | NSF | AVETkNFsGAELEGL | S437 | 0.044 | 86.66 | 744 | 0.034976 | 0.044 | 0.035 | 0.8571 |
| Q92887 | ABCC2 | sMLQASVsTERLEky | S609 | 0 | 88.87 | 1545 | 0 | 0 | 0 | 1 |
| Q92887 | ABCC2 | sTERLEkyLGGDDLD | Y616 | 0.036 | 89.36 | 1545 | 0.015793 | 0.036 | 0.0158 | 1 |
| Q86W34 | AMZ2 | GITMIDLyPRDSWNF | Y179 | 0.078 | 94.48 | 360 | 0.034217 | 0.078 | 0.0342 | 1 |
| Q86W34 | AMZ2 | SDFySMHyKGKVKKL | Y214 | 0.101 | 84.6 | 360 | 0.044307 | 0.101 | 0.0443 | 1 |
| P48051 | KCNJ6 | FSTHAVIsMRDGKLC | S212 | 0.004 | 97.11 | 423 | 0.00318 | 0.004 | 0.0032 | 1 |
| P26718 | KLRK1 | NCStPNTyICMQRTV | Y209 | 0.002 | 98.76 | 216 | 0.000877 | 0.002 | 0.0009 | 0.9375 |
| Q6P2C0 | WDR93 | PGMVLIFsKNGsVCL | S541 | 0.11 | 89.91 | 686 | 0.087441 | 0.163 | 0.1296 | 1 |
| Q08AD1 | CAMSAP2 | QFRSLyTyCPETEEI | Y1415 | 0.249 | 91.54 | 1489 | 0.109233 | 0.249 | 0.1092 | 0.8 |
| P43307 | SSR1 | VSGEPEAsPSADTTI | S80 | 0.109 | 93.8 | 286 | 0.086646 | 0.085 | 0.1149 | 1 |
| P43307 | SSR1 | QRQATFEySFIPAEP | Y156 | 0.133 | 96.99 | 286 | 0.058345 | 0.133 | 0.0583 | 1 |
| P51172 | SCNN1D | GAIRLVCsRGNRLKT | S242 | 0.156 | 65.83 | 802 | 0.124007 | 0.156 | 0.124 | 1 |
| P51172 | SCNN1D | RKLLPLVtLCDGNPR | T296 | 0 | 92.62 | 802 | 0 | 0 | 0 | 1 |
| P51172 | SCNN1D | sFsVRPGtEATISIR | T513 | 0.184 | 93.27 | 802 | 0.125675 | 0.184 | 0.1257 | 1 |
| A0AV96 | RBM47 | ALMERtGysMVQENG | Y44 | 0.295 | 89.62 | 593 | 0.129412 | 0.295 | 0.1294 | 1 |
| A0AV96 | RBM47 | QENGQRKyGGPPPGW | Y55 | 0.03 | 92.01 | 593 | 0.013161 | 0.03 | 0.0132 | 1 |
| P39019 | RPS19 | GAGVGsMtkIYGGRQ | T76 | 0.022 | 96.13 | 145 | 0.015026 | 0.022 | 0.015 | 1 |
| Q9BVA0 | KATNB1 | kyEsyVQtGCTSLKL | T577 | 0.072 | 94.6 | 655 | 0.049177 | 0.072 | 0.0492 | 1 |
| O00244 | ATOX1 | NkKVCIEsEHsMDtL | S44 | 0.006 | 98.47 | 68 | 0.004769 | 0.006 | 0.0048 | 1 |
| P48736 | PIK3CG | LWHFRYEsLKHPKAY | S582 | 0.003 | 92.48 | 1102 | 0.002385 | 0.003 | 0.0024 | 1 |
| O60568 | PLOD3 | YYARSEDyVELVQRK | Y444 | 0.034 | 96.98 | 738 | 0.014915 | 0.034 | 0.0149 | 1 |

| | | | | | | | | | | |
|---|---|---|---|---|---|---|---|---|---|---|
| O60568 | PLOD3 | VGVWNVPyISQAYVI | Y460 | 0.165 | 97.7 | 738 | 0.072383 | 0.23 | 0.1009 | 1 |
| O60568 | PLOD3 | PTTWGTRyIMVSFVD | Y730 | 0.234 | 97.65 | 738 | 0.102652 | 0.234 | 0.1027 | 1 |
| P14868 | DARS1 | AANINkEsIVDVEGV | S112 | 0 | 98.77 | 501 | 0 | 0 | 0 | 1 |
| O15357 | INPPL1 | HLLLTVKsMDGyEsY | S827 | 0.012 | 91.8 | 1258 | 0.009539 | 0.012 | 0.0095 | 1 |
| Q92729 | PTPRU | LIYFQAAsHLKGETR | S710 | 0 | 87.26 | 1446 | 0 | 0.152 | 0.1208 | 1 |
| Q9UGC6 | RGS17 | LDPNPHMyEDAQLQI | Y171 | 0.046 | 98.41 | 210 | 0.02018 | 0.046 | 0.0202 | 1 |
| P50395 | GDI2 | ___MNEEyDVIVLGT | Y5 | 0.103 | 96.51 | 445 | 0.045185 | 0.103 | 0.0452 | 1 |
| P50395 | GDI2 | ITPLEDLykRFkIPG | Y53 | 0.07 | 98.44 | 445 | 0.030708 | 0.07 | 0.0307 | 1 |
| P50395 | GDI2 | GKIyKVPstEAEALA | S121 | 0.079 | 83.45 | 445 | 0.062798 | 0.152 | 0.1208 | 1 |
| P50395 | GDI2 | LDQPCyEtINRIkLy | T205 | 0.011 | 98.43 | 445 | 0.007513 | 0.011 | 0.0075 | 1 |
| P50395 | GDI2 | YGksPyLyPLyGLGE | Y226 | 0 | 97.42 | 445 | 0 | 0 | 0 | 1 |
| P50395 | GDI2 | KyIAIVStTVETkEP | T355 | 0 | 98.46 | 445 | 0 | 0 | 0 | 1 |
| P50395 | GDI2 | PkDLGtEsQIFIsRT | S396 | 0.091 | 98.26 | 445 | 0.072337 | 0.079 | 0.0995 | 1 |
| P50395 | GDI2 | tEsQIFIsRTYDATT | S401 | 0.03 | 98.49 | 445 | 0.023847 | 0.03 | 0.0238 | 1 |
| O95347 | SMC2 | sNLQDLVykNGQAGI | Y67 | 0.207 | 90.89 | 1197 | 0.090808 | 0.336 | 0.1283 | 1 |
| O95347 | SMC2 | NANVLFktkFVDGVS | T1159 | 0.078 | 90.84 | 1197 | 0.053275 | 0.078 | 0.0533 | 1 |
| P14410 | SI | PTVSDTLyDVKVAQN | Y190 | 0.039 | 97.93 | 1827 | 0.017109 | 0.076 | 0.0333 | 1 |
| P14410 | SI | KVAQNPFsIQVIRKS | S200 | 0.078 | 97.94 | 1827 | 0.062003 | 0.072 | 0.0872 | 1 |
| P14410 | SI | FPRQDNSysVNsARY | Y983 | 0.024 | 90.32 | 1827 | 0.010528 | 0.024 | 0.0105 | 1 |
| P14410 | SI | RNILNIRyTLLPyFY | Y1587 | 0.002 | 98.36 | 1827 | 0.000877 | 0.002 | 0.0009 | 1 |
| P14410 | SI | IRyTLLPyFYTQMHE | Y1592 | 0.084 | 98.16 | 1827 | 0.03685 | 0.144 | 0.0632 | 1 |
| Q9UBP0 | SPAST | VARELQPsIIFIDEV | S436 | 0.03 | 93.79 | 616 | 0.023847 | 0.03 | 0.0238 | 1 |
| Q9UBP0 | SPAST | sPQtLEAyIRWNKDF | Y604 | 0.138 | 95.87 | 616 | 0.060539 | 0.138 | 0.0605 | 1 |
| O43895 | XPNPEP2 | DAVAVIRyLVWLEKN | Y379 | 0 | 98.74 | 674 | 0 | 0 | 0 | 1 |
| O43895 | XPNPEP2 | SFETISAsGLNAALA | S422 | 0.007 | 96.87 | 674 | 0.005564 | 0.007 | 0.0056 | 1 |
| O43895 | XPNPEP2 | LNAALAHysPTKELN | Y431 | 0.318 | 84.14 | 674 | 0.139502 | 0.318 | 0.1395 | 1 |
| O43895 | XPNPEP2 | HLQYLNRyyQTIREK | Y617 | 0.145 | 98.55 | 674 | 0.063609 | 0.145 | 0.0636 | 1 |
| O43895 | XPNPEP2 | LQYLNRyyQTIREKV | Y618 | 0 | 98.6 | 674 | 0 | 0 | 0 | 1 |
| Q14667 | BLTP2 | WSVTQMVsDLSQVTV | S1384 | 0.155 | 92.45 | 2235 | 0.123212 | 0.155 | 0.1232 | 0.8333 |
| Q15831 | STK11 | GFKVDIWsAGVTLYN | S240 | 0.02 | 98.77 | 433 | 0.015898 | 0.02 | 0.0159 | 1 |
| P17931 | LGALS3 | RRVIVCNtkLDNNWG | T175 | 0.054 | 98.9 | 250 | 0.036883 | 0.054 | 0.0369 | 1 |

| | | | | | | | | | | |
|---|---|---|---|---|---|---|---|---|---|---|
| P17931 | LGALS3 | NDAHLLQyNHRVKkL | Y221 | 0.011 | 98.86 | 250 | 0.004826 | 0.011 | 0.0048 | 1 |
| Q9NYU2 | UGGT1 | KGQRAVIsNGRIIGP | S895 | 0.029 | 94.73 | 1555 | 0.023053 | 0.029 | 0.0231 | 1 |
| Q9BXI6 | TBC1D10A | KyLPGyysEKLEAIQ | S232 | 0.167 | 91.63 | 508 | 0.132751 | 0.167 | 0.1328 | 0.8333 |
| Q9BXI6 | TBC1D10A | ACQGQyEtIERLRsL | T330 | 0 | 96.29 | 508 | 0 | 0 | 0 | 1 |
| Q9H892 | TTC12 | LKDMKVLyTNRAQAy | Y143 | 0.176 | 98.64 | 705 | 0.077209 | 0.176 | 0.0772 | 1 |
| Q9H892 | TTC12 | yTNRAQAyMKLEDyE | Y150 | 0.34 | 98.72 | 705 | 0.149153 | 0.34 | 0.1492 | 1 |
| P15735 | PHKG2 | AkEFYQKyDPkDVIG | Y24 | 0.041 | 97.46 | 406 | 0.017986 | 0.041 | 0.018 | 1 |
| P04839 | CYBB | DIPPKFFytRkLLGS | Y41 | 0.02 | 95.94 | 570 | 0.008774 | 0.02 | 0.0088 | 1 |
| P04839 | CYBB | IPPKFFytRkLLGSA | T42 | 0.051 | 95.93 | 570 | 0.034834 | 0.082 | 0.0609 | 1 |
| Q8NE00 | TMEM104 | TKyLQILtsLMRWIA | T237 | 0.101 | 78.23 | 496 | 0.068984 | 0.101 | 0.069 | 1 |
| O95954 | FTCD | EELDVPVyLYGEAAR | Y124 | 0.009 | 98.44 | 541 | 0.003948 | 0.009 | 0.0039 | 1 |
| O95954 | FTCD | PAIRAGEyEALPKKL | Y146 | 0.238 | 97.71 | 541 | 0.104407 | 0.238 | 0.1044 | 1 |
| O95954 | FTCD | VTALHTVyEETCREA | Y250 | 0.225 | 98.77 | 541 | 0.098704 | 0.225 | 0.0987 | 1 |
| Q15283 | RASA2 | TkVkkKTsNPQFNEI | S217 | 0.173 | 93.27 | 850 | 0.13752 | 0.173 | 0.1375 | 0.875 |
| P11831 | SRF | MKKAYELstLTGTQV | S176 | 0.178 | 98.15 | 508 | 0.141495 | 0.178 | 0.1415 | 1 |
| P11831 | SRF | GHVYTFAtRKLQPMI | T199 | 0.171 | 98.37 | 508 | 0.116795 | 0.171 | 0.1168 | 1 |
| P82673 | MRPS35 | DQDWPsVyPVAAPFK | Y70 | 0.072 | 95.96 | 323 | 0.031585 | 0.072 | 0.0316 | 1 |
| P82673 | MRPS35 | EEyIWENsSSERNIL | S256 | 0.082 | 88.79 | 323 | 0.065183 | 0.092 | 0.0988 | 1 |
| O43747 | AP1G1 | RMRIKLtyNHKGSAM | Y800 | 0.057 | 92.69 | 822 | 0.025005 | 0.057 | 0.025 | 1 |
| Q6ZUA9 | MROH5 | LLSTCFRsVFALPLL | S503 | 0.082 | 90.67 | 1318 | 0.065183 | 0.082 | 0.0652 | 1 |
| P62906 | RPL10A | VKKLAKkyDAFLASE | Y107 | 0.312 | 81.39 | 217 | 0.13687 | 0.312 | 0.1369 | 1 |
| Q5JXA9 | SIRPB2 | SEPLNCDySIyIHNV | Y110 | 0.231 | 91.18 | 342 | 0.101336 | 0.231 | 0.1013 | 1 |
| Q5JXA9 | SIRPB2 | LNCDySIyIHNVTRE | Y113 | 0.045 | 94.87 | 342 | 0.019741 | 0.045 | 0.0197 | 0.9091 |
| Q9UDY2 | TJP2 | KSRANEEyGLRLGsQ | Y319 | 0.228 | 87.44 | 1190 | 0.10002 | 0.228 | 0.1 | 1 |
| P09172 | DBH | GGPGSSRyLRLEVHy | Y327 | 0.18 | 95.06 | 617 | 0.078963 | 0.18 | 0.079 | 1 |
| P09172 | DBH | yLRLEVHyHNPLVIE | Y334 | 0.015 | 92.01 | 617 | 0.00658 | 0.015 | 0.0066 | 1 |
| P09172 | DBH | SSGIRLYyTAKLRRF | Y353 | 0.249 | 98.28 | 617 | 0.109233 | 0.249 | 0.1092 | 1 |
| P09172 | DBH | LTGRKVVtVLVRDGR | T422 | 0 | 98.77 | 617 | 0 | 0 | 0 | 1 |
| P08294 | SOD3 | GRGGNQAsVENGNAG | S195 | 0.005 | 98.38 | 240 | 0.003975 | 0.005 | 0.004 | 1 |
| P46459 | NSF | FPFIkICsPDkMIGF | S569 | 0.068 | 91.68 | 744 | 0.054054 | 0.068 | 0.0541 | 1 |
| P46459 | NSF | RKLLIIGtTSRKDVL | T645 | 0 | 96.76 | 744 | 0 | 0 | 0 | 1 |

| | | | | | | | | | | |
|---|---|---|---|---|---|---|---|---|---|---|
| P46459 | NSF | LLMLIEMsLQMDPEy | S717 | 0 | 92.6 | 744 | 0 | 0 | 0 | 1 |
| A6NH11 | GLTPD2 | MMRRFHAsLKPEGDV | S92 | 0 | 98.6 | 291 | 0 | 0.048 | 0.0382 | 1 |
| Q9NXH9 | TRMT1 | ADFRVSLsHACkNAV | S479 | 0.011 | 97.06 | 659 | 0.008744 | 0.011 | 0.0087 | 0.875 |
| Q9NXH9 | TRMT1 | RERLsETsPAFrILS | S517 | 0.166 | 94.99 | 659 | 0.131956 | 0.166 | 0.132 | 1 |
| P51589 | CYP2J2 | WKEQRRFtLtALRNF | T141 | 0 | 98.45 | 502 | 0 | 0 | 0 | 1 |
| P51589 | CYP2J2 | tRDFIDAyLKEMSKH | Y281 | 0 | 97.77 | 502 | 0 | 0 | 0 | 1 |
| P07311 | ACYP1 | EWLEtRGsPkSHIDk | S71 | 0.024 | 98.02 | 99 | 0.019078 | 0.024 | 0.0191 | 0.875 |
| O14745 | NHERF1 | IRLVEPGsPAEkAGL | S46 | 0.094 | 86.77 | 358 | 0.074722 | 0.094 | 0.0747 | 1 |
| O15042 | U2SURP | YYRWkLYsILQGDsP | S479 | 0.048 | 93.16 | 1029 | 0.038156 | 0.048 | 0.0382 | 1 |
| Q13445 | TMED1 | EPTEAGDyKLCFDNS | Y103 | 0.028 | 94.42 | 227 | 0.012283 | 0.028 | 0.0123 | 1 |
| Q9NYI0 | PSD3 | ADMDGKKtPRGKRGW | T805 | 0.022 | 87.64 | 1048 | 0.015026 | 0.022 | 0.015 | 1 |
| P62899 | RPL31 | YRIRVRLsRKRNEDE | S89 | 0.091 | 94.1 | 125 | 0.072337 | 0.091 | 0.0723 | 1 |
| P62899 | RPL31 | DsPNkLytLVtyVPV | T104 | 0 | 94.63 | 125 | 0 | 0 | 0 | 0.8333 |
| Q86W24 | NLRP14 | KVMLPEAsLLVTTRL | S302 | 0.065 | 95.36 | 1093 | 0.051669 | 0.106 | 0.0911 | 1 |
| Q86W24 | NLRP14 | LVTTRLttsKrLKQL | T311 | 0.025 | 83.96 | 1093 | 0.017075 | 0.211 | 0.1368 | 1 |
| O43379 | WDR62 | sLSALAFsPDGKYIV | S120 | 0.013 | 94.75 | 1518 | 0.010334 | 0.013 | 0.0103 | 1 |
| O43379 | WDR62 | GVACVAFsPNMKHIV | S164 | 0.024 | 94.14 | 1518 | 0.019078 | 0.024 | 0.0191 | 1 |
| O43379 | WDR62 | GVRVMQVsPDGQHLA | S501 | 0.045 | 94.67 | 1518 | 0.035771 | 0.045 | 0.0358 | 1 |
| O43379 | WDR62 | NRDIQMIsCGADkSI | S598 | 0 | 97.28 | 1518 | 0 | 0 | 0 | 1 |
| O43379 | WDR62 | NGkQKKCykGSQGDE | Y666 | 0.215 | 89.58 | 1518 | 0.094317 | 0.215 | 0.0943 | 1 |
| O43379 | WDR62 | LkVHVDPsGTFLATS | S684 | 0 | 92.73 | 1518 | 0 | 0 | 0 | 1 |
| Q6XZF7 | DNMBP | IQkHLQDsLADLKSL | S896 | 0.161 | 82.71 | 1577 | 0.127981 | 0.161 | 0.128 | 1 |
| Q6UW63 | POGLUT2 | DGSFIVRyRMyASyK | Y101 | 0.007 | 98.39 | 502 | 0.003071 | 0.007 | 0.0031 | 1 |
| Q6UW63 | POGLUT2 | RyRMyASyKNLkVEI | Y107 | 0.118 | 94.42 | 502 | 0.051765 | 0.118 | 0.0518 | 1 |
| Q6UW63 | POGLUT2 | ELQPWKHyIPVKSNL | Y408 | 0.266 | 98.89 | 502 | 0.11669 | 0.266 | 0.1167 | 1 |
| Q969V3 | NCLN | YNLLFFAsGGGKFNY | S275 | 0.006 | 96.36 | 563 | 0.004769 | 0.023 | 0.0183 | 1 |
| Q8NGN0 | OR4D5 | SVLyTIVtPMLNPAI | T279 | 0.015 | 93.12 | 318 | 0.010245 | 0.015 | 0.0102 | 0.8889 |
| Q16739 | UGCG | ISWRTGRyRLRCGGT | Y380 | 0.057 | 88.85 | 394 | 0.025005 | 0.057 | 0.025 | 0.9231 |
| O75417 | POLQ | PKAVLEKyHSFGVKK | Y84 | 0.01 | 86.96 | 2590 | 0.004387 | 0.01 | 0.0044 | 1 |
| O75417 | POLQ | VGIKVDGyMGSTsPS | Y171 | 0.201 | 84.64 | 2590 | 0.088176 | 0.299 | 0.1048 | 1 |
| P11940 | PABPC1 | MtDESGkskGFGFVs | S230 | 0.16 | 91.55 | 636 | 0.127186 | 0.16 | 0.1272 | 1 |

| | | | | | | | | | | |
|---|---|---|---|---|---|---|---|---|---|---|
| P20248 | CCNA2 | NYIDRFLsSMSVLRG | S244 | 0.161 | 97.93 | 432 | 0.127981 | 0.161 | 0.128 | 1 |
| Q8TE59 | ADAMTS19 | HGEKMLEsFCKWQHE | S399 | 0.163 | 93.32 | 1207 | 0.129571 | 0.163 | 0.1296 | 1 |
| P82664 | MRPS10 | QYEMRTLyRCLELEH | Y140 | 0.305 | 98.27 | 201 | 0.133799 | 0.305 | 0.1338 | 1 |
| P82664 | MRPS10 | TGSTADVyLEyIQRN | Y156 | 0.033 | 97.83 | 201 | 0.014477 | 0.033 | 0.0145 | 1 |
| P27708 | CAD | YGVRVLGtPVETIEL | T504 | 0.022 | 93.77 | 2225 | 0.015026 | 0.022 | 0.015 | 1 |
| P27708 | CAD | KGWkEIEyEVVRDAY | Y602 | 0.005 | 97.06 | 2225 | 0.002193 | 0.005 | 0.0022 | 1 |
| P27708 | CAD | AFEPSVDyCVVkIPR | Y735 | 0.02 | 97.64 | 2225 | 0.008774 | 0.02 | 0.0088 | 1 |
| P27708 | CAD | kIGSCMksVGEVMGI | S761 | 0.036 | 91.83 | 2225 | 0.028617 | 0.036 | 0.0286 | 1 |
| P27708 | CAD | IECNVRVsRsFPFVS | S1220 | 0.103 | 93.52 | 2225 | 0.081876 | 0.103 | 0.0819 | 1 |
| P27708 | CAD | CNVRVsRsFPFVSkT | S1222 | 0.026 | 94.4 | 2225 | 0.020668 | 0.026 | 0.0207 | 1 |
| P27708 | CAD | GVkVPQFsFSRLAGA | S1267 | 0.073 | 86.95 | 2225 | 0.058029 | 0.069 | 0.0823 | 1 |
| O96008 | TOMM40 | GQAGMHAtYYHKASD | T273 | 0.086 | 96.64 | 361 | 0.058739 | 0.086 | 0.0587 | 0.8333 |
| P48745 | CCN3 | KSCGMGFstRVTNRN | S225 | 0.009 | 91.74 | 357 | 0.007154 | 0.03 | 0.0238 | 1 |
| Q92542 | NCSTN | QSQPLPPssLQRFLR | S425 | 0.042 | 95.78 | 709 | 0.033386 | 0.042 | 0.0334 | 1 |
| Q92542 | NCSTN | SQPLPPssLQRFLRA | S426 | 0.025 | 96.79 | 709 | 0.019873 | 0.025 | 0.0199 | 1 |
| Q92542 | NCSTN | FLRArNIsGVVLADH | S437 | 0.018 | 91.69 | 709 | 0.014308 | 0.018 | 0.0143 | 1 |
| Q92542 | NCSTN | ARLARALsPAFELsQ | S632 | 0.015 | 95.47 | 709 | 0.011924 | 0.015 | 0.0119 | 0.8571 |
| Q92542 | NCSTN | STEYStWtESRWKDI | T649 | 0.001 | 96.1 | 709 | 0.000683 | 0.001 | 0.0007 | 0.8333 |
| Q9NZK5 | ADA2 | AKHLIERsQVFNILR | S96 | 0.074 | 98.72 | 511 | 0.058824 | 0.074 | 0.0588 | 1 |
| Q9NZK5 | ADA2 | tGHsLHDyKEALMIP | Y337 | 0.018 | 98.82 | 511 | 0.007896 | 0.018 | 0.0079 | 1 |
| Q9NZK5 | ADA2 | HPAVRTysWKkDIPI | S398 | 0.002 | 98.86 | 511 | 0.00159 | 0.002 | 0.0016 | 0.8889 |
| Q00722 | PLCB2 | LSERsVRtYVEVELF | T701 | 0 | 98.27 | 1185 | 0 | 0 | 0 | 1 |
| Q00722 | PLCB2 | KLSPSTNsINPVWKE | S728 | 0.012 | 96.64 | 1185 | 0.009539 | 0.044 | 0.035 | 1 |
| Q8TCU3 | SLC7A13 | TPREILssDAVAITW | S272 | 0.07 | 84.7 | 470 | 0.055644 | 0.07 | 0.0556 | 0.8889 |
| Q6PGP7 | SKIC3 | HyQLGLTyWFMGEET | Y464 | 0.12 | 93.71 | 1564 | 0.052642 | 0.202 | 0.0751 | 0.8182 |
| Q6PGP7 | SKIC3 | LNPESIYsVFkVAAI | S634 | 0 | 92.54 | 1564 | 0 | 0 | 0 | 1 |
| Q6PGP7 | SKIC3 | AMQDAtLsKAALNEL | S1141 | 0.001 | 89.12 | 1564 | 0.000795 | 0.088 | 0.07 | 1 |
| Q4KWH8 | PLCH1 | SQSKVDMyARVLQEG | Y332 | 0 | 96.83 | 1693 | 0 | 0 | 0 | 1 |
| O15195 | VILL | PAGRSPAtPIVLVKQ | T684 | 0.171 | 96.96 | 856 | 0.116795 | 0.171 | 0.1168 | 1 |
| Q8TC59 | PIWIL2 | LKDDRIEtyVRTIQS | T651 | 0.101 | 95.66 | 973 | 0.068984 | 0.101 | 0.069 | 1 |
| Q8TC59 | PIWIL2 | KDDRIEtyVRTIQST | Y652 | 0.002 | 96.43 | 973 | 0.000877 | 0.002 | 0.0009 | 1 |

| | | | | | | | | | | |
|---|---|---|---|---|---|---|---|---|---|---|
| Q8TC59 | PIWIL2 | NLTLTKWySRVVFQM | Y772 | 0.22 | 97.76 | 973 | 0.096511 | 0.22 | 0.0965 | 1 |
| Q9Y5F8 | PCDHGB7 | DFEEVERytINIEAk | Y315 | 0.191 | 90.17 | 929 | 0.083789 | 0.191 | 0.0838 | 1 |
| Q96LD1 | SGCZ | EDGRVLFsADEDEIT | S153 | 0.065 | 94.35 | 299 | 0.051669 | 0.065 | 0.0517 | 1 |
| Q9Y2J2 | EPB41L3 | tCDVEkRsRGQVLFD | S130 | 0.175 | 94.74 | 1087 | 0.13911 | 0.175 | 0.1391 | 1 |
| O00472 | ELL2 | SAIELPDyLIKYIAI | Y528 | 0.231 | 95.55 | 640 | 0.101336 | 0.231 | 0.1013 | 1 |
| O00472 | ELL2 | YQkIkQSsPNYHEEK | S604 | 0.166 | 70.22 | 640 | 0.131956 | 0.166 | 0.132 | 1 |
| Q8N0Z6 | TTC5 | THCRNkVsLQNLSMV | S138 | 0 | 98.66 | 440 | 0 | 0 | 0 | 0.8182 |
| Q8N0Z6 | TTC5 | GNSYLSLyFSTGQNP | Y193 | 0.094 | 91.5 | 440 | 0.041236 | 0.094 | 0.0412 | 0.8667 |
| O15553 | MEFV | RVGSISFyNVTARSH | Y732 | 0.043 | 97.53 | 781 | 0.018863 | 0.043 | 0.0189 | 1 |
| O15553 | MEFV | VTARSHIyTFASCSF | Y741 | 0.293 | 94.86 | 781 | 0.128535 | 0.293 | 0.1285 | 1 |
| Q5T5U3 | ARHGAP21 | PLFTNDkyADFIEAN | Y1240 | 0.145 | 93.5 | 1958 | 0.063609 | 0.145 | 0.0636 | 1 |
| P21462 | FPR1 | PMsIVAVsYGLIATK | S220 | 0.032 | 96.12 | 350 | 0.025437 | 0.032 | 0.0254 | 1 |
| O95461 | LARGE1 | MKHISTPyMFLSDID | Y558 | 0.123 | 98.18 | 756 | 0.053958 | 0.123 | 0.054 | 1 |
| O95461 | LARGE1 | FLPMyGLyEYLRKSV | Y573 | 0.206 | 97.93 | 756 | 0.090369 | 0.206 | 0.0904 | 1 |
| O95461 | LARGE1 | MIVPAFEtLRyRLSF | T599 | 0.041 | 95.23 | 756 | 0.028004 | 0.041 | 0.028 | 1 |
| Q13308 | PTK7 | AtVFANGsLLLTQVR | S285 | 0.036 | 94.1 | 1070 | 0.028617 | 0.036 | 0.0286 | 1 |
| Q9HBT6 | CDH20 | GNSARVVySILQGQP | Y205 | 0.188 | 95.35 | 801 | 0.082473 | 0.188 | 0.0825 | 1 |
| O95396 | MOCS3 | kAFSAAAsLRRLNsA | S144 | 0.107 | 97.61 | 460 | 0.085056 | 0.107 | 0.0851 | 1 |
| Q9NQ86 | TRIM36 | PPQESTKsCMDCsAs | S170 | 0 | 88.64 | 728 | 0 | 0 | 0 | 1 |
| Q9NQ86 | TRIM36 | sCMDCsAsyCNECFK | S177 | 0.037 | 91.39 | 728 | 0.029412 | 0.037 | 0.0294 | 1 |
| Q9NQ86 | TRIM36 | CMDCsAsyCNECFKI | Y178 | 0.075 | 92.77 | 728 | 0.032901 | 0.075 | 0.0329 | 1 |
| P21589 | NT5E | AsQIsGLyLPYKVLP | Y158 | 0.336 | 98.09 | 574 | 0.147398 | 0.336 | 0.1474 | 1 |
| Q9Y2X3 | NOP58 | HAsLVGQtsPKHKGk | T350 | 0.039 | 91.07 | 529 | 0.026638 | 0.039 | 0.0266 | 1 |
| Q969P6 | TOP1MT | AEEVATFyGRMLDHE | Y96 | 0.066 | 98.21 | 601 | 0.028953 | 0.066 | 0.029 | 1 |
| Q969P6 | TOP1MT | RQRAVALyFIDkLAL | Y316 | 0.16 | 98.34 | 601 | 0.07019 | 0.21 | 0.0921 | 1 |
| Q969P6 | TOP1MT | VPVEKPVyKNLQLFM | Y384 | 0.11 | 98.39 | 601 | 0.048255 | 0.11 | 0.0483 | 1 |
| P23945 | FSHR | GPVILDIsRtRIHSL | S226 | 0.046 | 98.46 | 695 | 0.036566 | 0.046 | 0.0366 | 1 |
| P23945 | FSHR | VILDIsRtRIHSLPS | T228 | 0.073 | 98.27 | 695 | 0.04986 | 0.073 | 0.0499 | 1 |
| O95782 | AP2A1 | LLRLYkAsPDLVPMG | S179 | 0.177 | 89.2 | 977 | 0.1407 | 0.177 | 0.1407 | 1 |
| Q3B8N5 | PROX2 | SSNLLKVyFPDVQFN | Y466 | 0.187 | 95.39 | 592 | 0.082034 | 0.187 | 0.082 | 0.9091 |
| Q8IV08 | PLD3 | NMDWRsLtQVKELGV | T225 | 0.04 | 97.88 | 490 | 0.027321 | 0.04 | 0.0273 | 0.8571 |

| | | | | | | | | | | |
|---|---|---|---|---|---|---|---|---|---|---|
| Q8IV08 | PLD3 | GVVMYNCsCLARDLT | S238 | 0.182 | 97.55 | 490 | 0.144674 | 0.182 | 0.1447 | 1 |
| Q9H5V8 | CDCP1 | RKEERVEyyIPGSTT | Y283 | 0 | 90.99 | 836 | 0 | 0 | 0 | 0.8333 |
| Q13243 | SRSF5 | RLIVENLssRVsWQD | S116 | 0.049 | 68.73 | 272 | 0.038951 | 0.049 | 0.039 | 1 |
| Q9Y5Y2 | NUBP2 | FIQEFPGsPAFAALT | S249 | 0.152 | 91.73 | 271 | 0.120827 | 0.152 | 0.1208 | 1 |
| Q8IXR5 | FAM178B | FPDMTSRsRRLRSQL | S520 | 0.005 | 94.36 | 679 | 0.003975 | 0.005 | 0.004 | 0.9 |
| Q9GZW8 | MS4A7 | GLFLLADsMVALRTA | S138 | 0.001 | 90.74 | 240 | 0.000795 | 0.109 | 0.0866 | 1 |
| Q8N126 | CADM3 | QEDPNGKtFTVSsSV | T187 | 0.081 | 97.18 | 398 | 0.055324 | 0.081 | 0.0553 | 0.875 |
| Q8N126 | CADM3 | GKtFTVSsSVtFQVT | S192 | 0.018 | 97.94 | 398 | 0.014308 | 0.018 | 0.0143 | 1 |
| Q8N126 | CADM3 | NKsDSGTyGCTATSN | Y297 | 0.08 | 91.46 | 398 | 0.035095 | 0.157 | 0.0689 | 1 |
| P51681 | CCR5 | QFWKNFQtLkIVILG | T195 | 0.068 | 95.56 | 352 | 0.046445 | 0.068 | 0.0464 | 1 |
| Q9UBM8 | MGAT4C | IKKIKVNtGtEDRQN | T396 | 0.146 | 96.93 | 478 | 0.09972 | 0.146 | 0.0997 | 1 |
| Q9UBM8 | MGAT4C | KIKVNtGtEDRQNDI | T398 | 0.16 | 90.22 | 478 | 0.109282 | 0.16 | 0.1093 | 0.8333 |
| P16112 | ACAN | LRDQTVRyPIVsPRT | Y536 | 0.115 | 94.01 | 2530 | 0.050449 | 0.115 | 0.0504 | 1 |
| P12814 | ACTN1 | EVENQILtRDAKGIs | T737 | 0.005 | 76.83 | 892 | 0.003415 | 0.193 | 0.1318 | 0.9 |
| Q6EMK4 | VASN | NLHDLDVsDNQLERV | S224 | 0.147 | 97.18 | 673 | 0.116852 | 0.147 | 0.1169 | 1 |
| Q6UWE0 | LRSAM1 | PKSCSLLsLATIKVL | S79 | 0.148 | 77.15 | 723 | 0.117647 | 0.177 | 0.1407 | 0.8 |
| Q9UJA5 | TRMT6 | IKKKKKKyEAIItVV | Y154 | 0.246 | 87.36 | 497 | 0.107917 | 0.246 | 0.1079 | 1 |
| Q9UJA5 | TRMT6 | ItVVKPStRILSIMy | T165 | 0.18 | 98.5 | 497 | 0.122942 | 0.18 | 0.1229 | 1 |
| Q9UJA5 | TRMT6 | tRILSIMyyAREPGK | Y172 | 0.061 | 98.12 | 497 | 0.02676 | 0.225 | 0.0987 | 1 |
| Q9UJA5 | TRMT6 | INHMRyDtLAQMLtL | T187 | 0.01 | 98.08 | 497 | 0.00683 | 0.01 | 0.0068 | 1 |
| Q9UJA5 | TRMT6 | ERMGGFGsIIQLyPG | S226 | 0.179 | 98.39 | 497 | 0.14229 | 0.179 | 0.1423 | 1 |
| Q9UJA5 | TRMT6 | FGsIIQLyPGGGPVR | Y231 | 0.119 | 97.33 | 497 | 0.052204 | 0.119 | 0.0522 | 1 |
| Q8IY21 | DDX60 | SGkTyASyYCMEKVL | Y796 | 0.146 | 89.91 | 1712 | 0.064048 | 0.281 | 0.1206 | 1 |
| Q8IY21 | DDX60 | LKLYFLFsLQFLVKE | S1416 | 0 | 92.67 | 1712 | 0 | 0 | 0 | 1 |
| Q8IY21 | DDX60 | NAYALDFyKHGSLIG | Y1646 | 0.11 | 93.42 | 1712 | 0.048255 | 0.11 | 0.0483 | 1 |
| Q05513 | PRKCZ | EILRGEEyGFSVDWW | Y428 | 0.048 | 93.43 | 592 | 0.021057 | 0.048 | 0.0211 | 1 |
| P11021 | HSPA5 | GNrItPsyVAFtPEG | Y65 | 0.089 | 98.75 | 654 | 0.039043 | 0.089 | 0.039 | 0.8 |
| P11021 | HSPA5 | LtsNPENtVFDAkRL | T91 | 0 | 98.75 | 654 | 0 | 0 | 0 | 0.9091 |
| P11021 | HSPA5 | IVLVGGstRIPkIQQ | T366 | 0.038 | 98.18 | 654 | 0.025955 | 0.038 | 0.026 | 1 |
| P54577 | YARS1 | LKAGCEVtILFADLH | T70 | 0.071 | 98.41 | 528 | 0.048494 | 0.122 | 0.0882 | 1 |
| Q9NVD7 | PARVA | VLEKINEtLKLPPRs | T167 | 0.171 | 92.97 | 372 | 0.116795 | 0.171 | 0.1168 | 0.8333 |

| | | | | | | | | | | |
|---|---|---|---|---|---|---|---|---|---|---|
| Q04828 | AKR1C1 | kskALEAtkLAIEAG | T38 | 0.006 | 98.77 | 323 | 0.004098 | 0.006 | 0.0041 | 1 |
| Q04828 | AKR1C1 | AGFRHIDsAHLyNNE | S51 | 0 | 98.72 | 323 | 0 | 0 | 0 | 0.9 |
| Q04828 | AKR1C1 | VkREDIFytskLWCN | Y81 | 0.001 | 98.93 | 323 | 0.000439 | 0.001 | 0.0004 | 0.9412 |
| Q04828 | AKR1C1 | NQVECHPyFNQRkLL | Y196 | 0.089 | 98.87 | 323 | 0.039043 | 0.178 | 0.0781 | 1 |
| Q04828 | AKR1C1 | GLNRNVRyLtLDIFA | Y305 | 0.066 | 98.61 | 323 | 0.028953 | 0.066 | 0.029 | 1 |
| Q96JY6 | PDLIM2 | AQSKIRQsPsPLRLQ | S72 | 0.156 | 91.58 | 352 | 0.124007 | 0.156 | 0.124 | 0.875 |
| Q8N142 | ADSS1 | AtDADIIsRCQGGNN | S61 | 0 | 97.46 | 457 | 0 | 0 | 0 | 1 |
| Q8N142 | ADSS1 | GVGKsREsMIQLF__ | S452 | 0.082 | 93.07 | 457 | 0.065183 | 0.164 | 0.1304 | 1 |
| Q2TAZ0 | ATG2A | QSRTLALsRCsLEVI | S809 | 0.001 | 88.76 | 1938 | 0.000795 | 0.001 | 0.0008 | 1 |
| Q2TAZ0 | ATG2A | TLALsRCsLEVILPS | S812 | 0.052 | 92.01 | 1938 | 0.041336 | 0.052 | 0.0413 | 1 |
| Q00887 | PSG9 | TRKDAGTytLHIIKR | Y120 | 0.037 | 97.35 | 426 | 0.016231 | 0.037 | 0.0162 | 1 |
| Q00887 | PSG9 | RKDAGTytLHIIKRG | T121 | 0 | 97.67 | 426 | 0 | 0 | 0 | 1 |
| Q00887 | PSG9 | TKyIAGPyECEIRNP | Y215 | 0 | 97.51 | 426 | 0 | 0 | 0 | 1 |
| Q8TF17 | SH3TC2 | LRHRHLKsQLQATKS | S958 | 0.008 | 85.32 | 1288 | 0.006359 | 0.008 | 0.0064 | 1 |
| Q8TF17 | SH3TC2 | LESLGQLyRNLNTAR | Y1011 | 0.129 | 93.63 | 1288 | 0.05659 | 0.25 | 0.1068 | 0.8 |
| Q16512 | PKN1 | GMGyGDRtstFCGtP | T772 | 0.062 | 88.21 | 942 | 0.042347 | 0.062 | 0.0423 | 1 |
| P50995 | ANXA11 | YKRMYGksLyHDIsG | S480 | 0.08 | 97.71 | 505 | 0.063593 | 0.08 | 0.0636 | 1 |
| P50995 | ANXA11 | yHDIsGDtsGDyRKI | T489 | 0.06 | 97.72 | 505 | 0.040981 | 0.06 | 0.041 | 1 |
| P50995 | ANXA11 | sGDtsGDyRKILLkI | Y493 | 0.196 | 98.69 | 505 | 0.085982 | 0.196 | 0.086 | 1 |
| Q8IW70 | TMEM151B | GLEGAPAtRLRFtKC | T230 | 0.002 | 93.67 | 566 | 0.001366 | 0.08 | 0.0546 | 1 |
| O75344 | FKBP6 | DFLDCAEsDKFCALS | S148 | 0.034 | 88.93 | 327 | 0.027027 | 0.034 | 0.027 | 0.8182 |
| Q9H159 | CDH19 | DREERsLyILRAQVI | Y113 | 0.228 | 93.01 | 772 | 0.10002 | 0.228 | 0.1 | 1 |
| Q9H159 | CDH19 | TDPDNRKsPIRYsIt | S402 | 0.115 | 91.63 | 772 | 0.091415 | 0.115 | 0.0914 | 1 |
| Q9NYK5 | MRPL39 | AMHLsEWyCRKSILA | Y99 | 0.211 | 93.04 | 338 | 0.092563 | 0.211 | 0.0926 | 1 |
| Q16527 | CSRP2 | AIHDEEIyCKSCyGK | Y57 | 0.097 | 89.67 | 193 | 0.042553 | 0.097 | 0.0426 | 1 |
| Q9UMQ6 | CAPN11 | GSWRRGSsAGGCRNH | S404 | 0.001 | 93.89 | 739 | 0.000795 | 0.182 | 0.1447 | 1 |
| Q9UMQ6 | CAPN11 | KKEFFTKyQDHGFsE | Y493 | 0.183 | 92.64 | 739 | 0.080279 | 0.183 | 0.0803 | 1 |
| P11172 | UMPS | GDNLGQQyNSPQEVI | Y432 | 0.315 | 91.16 | 480 | 0.138186 | 0.315 | 0.1382 | 0.8 |
| P11172 | UMPS | EVIGkRGsDIIIVGR | S444 | 0.001 | 97.66 | 480 | 0.000795 | 0.001 | 0.0008 | 0.8182 |
| P19174 | PLCG1 | AHESkEWyHASLTRA | Y669 | 0.242 | 96.83 | 1290 | 0.106162 | 0.242 | 0.1062 | 1 |
| P19174 | PLCG1 | KRNEPNSyAISFRAE | Y702 | 0.128 | 95.55 | 1290 | 0.056152 | 0.128 | 0.0562 | 1 |

| | | | | | | | | | | |
|---|---|---|---|---|---|---|---|---|---|---|
| O60566 | BUB1B | LNANDEAtVSVLGEL | T1008 | 0.09 | 90.52 | 1050 | 0.061471 | 0.09 | 0.0615 | 1 |
| O60674 | JAK2 | kVPVTHEtQEECLGM | T174 | 0.117 | 90.09 | 1132 | 0.079913 | 0.117 | 0.0799 | 1 |
| O60674 | JAK2 | DQTPLAIyNSISYkT | Y201 | 0.228 | 90.62 | 1132 | 0.10002 | 0.228 | 0.1 | 1 |
| Q8N6N6 | NATD1 | HTEVPDAyRGRGIAK | Y64 | 0.308 | 91.77 | 113 | 0.135115 | 0.308 | 0.1351 | 1 |
| P29074 | PTPN4 | MsGGILIyKNRVRMN | Y244 | 0.039 | 91.47 | 926 | 0.017109 | 0.039 | 0.0171 | 0.8462 |
| Q495M3 | SLC36A2 | IGMAALGyLRFGDDI | Y318 | 0.02 | 96.41 | 483 | 0.008774 | 0.02 | 0.0088 | 1 |
| Q4ZHG4 | FNDC1 | VTGYLVYsASyEDFI | S1694 | 0.123 | 94.11 | 1894 | 0.097775 | 0.123 | 0.0978 | 0.875 |
| P67809 | YBX1 | NPRKyLrsVGDGEtV | S102 | 0.002 | 93.79 | 324 | 0.00159 | 0.002 | 0.0016 | 1 |
| P67809 | YBX1 | GGVPVQGskYAADrN | S136 | 0.007 | 92.57 | 324 | 0.005564 | 0.007 | 0.0056 | 0.875 |
| Q8IZD9 | DOCK3 | FRPVMDTyIQKHFAG | Y689 | 0.054 | 92.79 | 2030 | 0.023689 | 0.054 | 0.0237 | 1 |
| Q8IZD9 | DOCK3 | HFAGALAyKELIRCL | Y700 | 0.069 | 93.6 | 2030 | 0.030269 | 0.069 | 0.0303 | 1 |
| Q8IZD9 | DOCK3 | VAEFVRGtLGSMPsT | T815 | 0.017 | 92.01 | 2030 | 0.011611 | 0.017 | 0.0116 | 1 |
| Q13003 | GRIK3 | TLGVSILyRKPNGTN | Y544 | 0.1 | 92.93 | 919 | 0.043869 | 0.1 | 0.0439 | 1 |
| P55769 | SNU13 | LLDLVQQsCNykQLR | S29 | 0 | 97.84 | 128 | 0 | 0 | 0 | 1 |
| P46940 | IQGAP1 | AKLGNFFsPkVVsLK | S86 | 0.095 | 85.49 | 1657 | 0.075517 | 0.095 | 0.0755 | 1 |
| P59044 | NLRP6 | DAVCRDLsEALRAAP | S749 | 0.047 | 95.5 | 892 | 0.037361 | 0.047 | 0.0374 | 1 |
| Q9Y490 | TLN1 | CsPSDAytkkELIEC | T1946 | 0.116 | 74.83 | 2541 | 0.07923 | 0.116 | 0.0792 | 1 |
| Q6ZRH7 | CATSPERG | VYFHSNGsEYIMALT | S357 | 0.1 | 93.04 | 1159 | 0.079491 | 0.1 | 0.0795 | 1 |
| Q6ZRH7 | CATSPERG | PKELSIKyMARSFRG | Y519 | 0.094 | 95.68 | 1159 | 0.041236 | 0.094 | 0.0412 | 0.8462 |
| Q5TCQ9 | MAGI3 | REPIRLKtVKPGKVI | T102 | 0.019 | 93.45 | 1481 | 0.012977 | 0.085 | 0.0581 | 1 |
| Q5TCQ9 | MAGI3 | INKDLRHyLsLQFQK | Y116 | 0 | 94.68 | 1481 | 0 | 0.091 | 0.0399 | 1 |
| O14617 | AP3D1 | VFLkYPEsLRPAFPR | S180 | 0.026 | 94.47 | 1153 | 0.020668 | 0.026 | 0.0207 | 1 |
| P27986 | PIK3R1 | LRKtRDQyLMWLTQK | Y580 | 0.085 | 90.06 | 724 | 0.037288 | 0.085 | 0.0373 | 1 |
| O75223 | GGCT | NYESAPPsPQYkkII | S136 | 0.001 | 98.56 | 188 | 0.000795 | 0.001 | 0.0008 | 0.875 |
| P11310 | ACADM | AEKkGDEyIINGQKM | Y183 | 0.114 | 98.71 | 421 | 0.05001 | 0.114 | 0.05 | 1 |
| P11310 | ACADM | VDSGRrNtyyASIAK | T351 | 0.127 | 98.26 | 421 | 0.086743 | 0.127 | 0.0867 | 1 |
| Q7Z3G6 | PRICKLE2 | FYQDGKIyCGRHHAE | Y182 | 0.308 | 97.88 | 844 | 0.135115 | 0.308 | 0.1351 | 1 |
| Q9Y2Z4 | YARS2 | VWLNRDktsPFELyQ | T297 | 0.089 | 93.42 | 477 | 0.060788 | 0.089 | 0.0608 | 0.9 |
| Q9Y2Z4 | YARS2 | WLNRDktsPFELyQF | S298 | 0.003 | 93.37 | 477 | 0.002385 | 0.003 | 0.0024 | 0.8333 |
| Q9Y2Z4 | YARS2 | ktsPFELyQFFVrQP | Y303 | 0.179 | 96.48 | 477 | 0.078525 | 0.179 | 0.0785 | 1 |
| Q9Y2Z4 | YARS2 | MITEGGVsINHQQVT | S435 | 0.169 | 93.57 | 477 | 0.134341 | 0.169 | 0.1343 | 1 |

| | | | | | | | | | | |
|---|---|---|---|---|---|---|---|---|---|---|
| Q5VYK3 | ECPAS | LMHLLIPtLFHMKYP | T144 | 0.026 | 92.05 | 1845 | 0.017758 | 0.047 | 0.0321 | 0.9091 |
| P41218 | MNDA | TAPFKYEsPENGKST | S227 | 0.162 | 79.7 | 407 | 0.128776 | 0.162 | 0.1288 | 1 |
| P41218 | MNDA | KsVHKKNtIyEIQDN | T337 | 0 | 97.67 | 407 | 0 | 0 | 0 | 1 |
| P41218 | MNDA | VHKKNtIyEIQDNTG | Y339 | 0 | 98.36 | 407 | 0 | 0 | 0 | 1 |
| Q14574 | DSC3 | TMHtRLKysILQQtP | Y282 | 0.232 | 97.79 | 896 | 0.101775 | 0.232 | 0.1018 | 1 |
| P11717 | IGF2R | DQVLQLVyKDGSPCP | Y1592 | 0.001 | 91.43 | 2491 | 0.000439 | 0.001 | 0.0004 | 0.8947 |
| Q14934 | NFATC4 | LTLTVPEysNKRVSR | Y653 | 0.14 | 95.96 | 902 | 0.061416 | 0.14 | 0.0614 | 1 |
| Q14934 | NFATC4 | SNGRRKRsPTQSFRF | S676 | 0.117 | 94.47 | 902 | 0.093005 | 0.117 | 0.093 | 1 |
| Q9HAQ2 | KIF9 | DAsQDLVyEtVAKDV | Y70 | 0.199 | 97.9 | 790 | 0.087298 | 0.327 | 0.1261 | 1 |
| Q8N3R9 | PALS1 | tVRNEMDsVIIsRIV | S277 | 0.157 | 94.37 | 675 | 0.124802 | 0.157 | 0.1248 | 1 |
| O95619 | YEATS4 | MAEFGPDsGGRVkGV | S12 | 0.059 | 92.63 | 227 | 0.0469 | 0.059 | 0.0469 | 1 |
| Q9H6W3 | RIOX1 | AELNRIPssRRRAAR | S197 | 0.17 | 98.54 | 641 | 0.135135 | 0.17 | 0.1351 | 1 |
| Q9H6W3 | RIOX1 | ELNRIPssRRRAARL | S198 | 0.019 | 98.72 | 641 | 0.015103 | 0.019 | 0.0151 | 1 |
| P07900 | HSP90AA1 | LRELISNssDALDkI | S52 | 0.057 | 95.8 | 732 | 0.04531 | 0.057 | 0.0453 | 1 |
| P07900 | HSP90AA1 | RELISNssDALDkIR | S53 | 0.067 | 93.59 | 732 | 0.053259 | 0.067 | 0.0533 | 1 |
| P07900 | HSP90AA1 | IPNkQDRtLtIVDtG | T88 | 0.106 | 97.61 | 732 | 0.072399 | 0.127 | 0.1075 | 1 |
| P07900 | HSP90AA1 | GtIAksGtkAFMEAL | T115 | 0.132 | 83.45 | 732 | 0.090158 | 0.132 | 0.0902 | 1 |
| P07900 | HSP90AA1 | kHNDDEQyAWEsSAG | Y160 | 0.222 | 97.93 | 732 | 0.097388 | 0.222 | 0.0974 | 1 |
| P07900 | HSP90AA1 | DEQyAWEsSAGGsFT | S164 | 0.123 | 95.58 | 732 | 0.097775 | 0.072 | 0.1192 | 1 |
| P07900 | HSP90AA1 | LkEDQtEyLEErRIk | Y197 | 0.085 | 96.61 | 732 | 0.037288 | 0.133 | 0.0583 | 1 |
| P07900 | HSP90AA1 | kEIVKkHsQFIGyPI | S211 | 0.028 | 90.27 | 732 | 0.022258 | 0.028 | 0.0223 | 1 |
| P07900 | HSP90AA1 | DDItNEEyGEFyksL | Y309 | 0.174 | 96.64 | 732 | 0.076331 | 0.257 | 0.1127 | 1 |
| P07900 | HSP90AA1 | NEEyGEFyksLtNDW | Y313 | 0.037 | 95.68 | 732 | 0.016231 | 0.037 | 0.0162 | 1 |
| P07900 | HSP90AA1 | FIrGVVDsEDLPLNI | S391 | 0.008 | 91.28 | 732 | 0.006359 | 0.008 | 0.0064 | 1 |
| P07900 | HSP90AA1 | LAEDkENykkFyEQF | Y434 | 0.091 | 97.81 | 732 | 0.03992 | 0.091 | 0.0399 | 0.9286 |
| P07900 | HSP90AA1 | kkFyEQFskNIkLGI | S442 | 0.106 | 97.78 | 732 | 0.084261 | 0.106 | 0.0843 | 0.8 |
| P07900 | HSP90AA1 | sELLryytsAsGDEM | T467 | 0.056 | 97.71 | 732 | 0.038249 | 0.056 | 0.0382 | 1 |
| P07900 | HSP90AA1 | EMVsLkDyCtRMkEN | Y480 | 0.037 | 97.29 | 732 | 0.016231 | 0.037 | 0.0162 | 1 |
| P07900 | HSP90AA1 | kENQkHIyyITGEtk | Y492 | 0.032 | 97.7 | 732 | 0.014038 | 0.032 | 0.014 | 1 |
| P07900 | HSP90AA1 | ENQkHIyyITGEtkD | Y493 | 0.224 | 97.91 | 732 | 0.098266 | 0.372 | 0.1361 | 1 |
| P07900 | HSP90AA1 | FEGktLVsVTkEGLE | S543 | 0.076 | 96.36 | 732 | 0.060414 | 0.121 | 0.0997 | 1 |

| | | | | | | | | | | |
|---|---|---|---|---|---|---|---|---|---|---|
| P07900 | HSP90AA1 | kVEkVVVsNRLVTsP | S589 | 0.06 | 90.94 | 732 | 0.047695 | 0.06 | 0.0477 | 1 |
| P07900 | HSP90AA1 | TsPCCIVtstyGWtA | T601 | 0.031 | 92.92 | 732 | 0.021173 | 0.031 | 0.0212 | 1 |
| P07900 | HSP90AA1 | VtstyGWtANMERIM | T607 | 0.135 | 92.75 | 732 | 0.092207 | 0.135 | 0.0922 | 1 |
| Q9Y6U3 | SCIN | MAKLyMVsDASGSMR | S255 | 0.15 | 79.62 | 715 | 0.119237 | 0.15 | 0.1192 | 0.8 |
| Q9Y6U3 | SCIN | QMNYSkNtQIQVLPE | T326 | 0.004 | 79.52 | 715 | 0.002732 | 0.108 | 0.0738 | 1 |
| P06132 | UROD | AWGEETDyTPVWCMR | Y30 | 0.264 | 98.86 | 367 | 0.115813 | 0.306 | 0.1342 | 0.9 |
| P06132 | UROD | VFQAItLtRQRLAGR | T141 | 0 | 98.81 | 367 | 0 | 0 | 0 | 1 |
| P51812 | RPS6KA3 | VKGAMAAtysALNRN | T706 | 0.013 | 75.13 | 740 | 0.008879 | 0.013 | 0.0089 | 0.8182 |
| Q14344 | GNA13 | DkLGEPDyIPSQQDI | Y189 | 0.078 | 96.83 | 377 | 0.034217 | 0.078 | 0.0342 | 1 |
| Q14344 | GNA13 | ILLARRPtkGIHEYD | T203 | 0.19 | 94.64 | 377 | 0.129773 | 0.19 | 0.1298 | 0.8889 |
| Q14344 | GNA13 | NRVFSNVsIILFLNK | S285 | 0.163 | 89.98 | 377 | 0.129571 | 0.163 | 0.1296 | 1 |
| Q02809 | PLOD1 | RSAQFFNyKIQALGL | Y53 | 0.192 | 96.01 | 727 | 0.084228 | 0.255 | 0.1119 | 1 |
| Q02809 | PLOD1 | LAQHGSEyQSVKLVG | Y344 | 0.129 | 97.07 | 727 | 0.05659 | 0.254 | 0.11 | 1 |
| Q02809 | PLOD1 | VALTEPNsLRLLIQQ | S392 | 0.004 | 97.2 | 727 | 0.00318 | 0.004 | 0.0032 | 1 |
| Q02809 | PLOD1 | YYARsEDyVDIVQGR | Y434 | 0.063 | 97.62 | 727 | 0.027637 | 0.063 | 0.0276 | 1 |
| Q02809 | PLOD1 | WHKFLLEyIAPMTEK | Y619 | 0.153 | 97.96 | 727 | 0.067119 | 0.153 | 0.0671 | 1 |
| Q02809 | PLOD1 | LNRVGVDyEGGGCRF | Y675 | 0.054 | 97.5 | 727 | 0.023689 | 0.054 | 0.0237 | 1 |
| O94851 | MICAL2 | LAERESLyRLLPQTT | Y445 | 0.21 | 97.83 | 1124 | 0.092124 | 0.21 | 0.0921 | 1 |
| O94851 | MICAL2 | INkNFEQyTLDPGTR | Y463 | 0.22 | 97.29 | 1124 | 0.096511 | 0.22 | 0.0965 | 1 |
| O15244 | SLC22A2 | RRtVGIFyQVAYTVG | Y241 | 0.01 | 90.15 | 555 | 0.004387 | 0.01 | 0.0044 | 1 |
| Q93084 | ATP2A3 | VETLGCtsVICSDkT | S346 | 0.174 | 94.82 | 999 | 0.138315 | 0.174 | 0.1383 | 1 |
| Q93084 | ATP2A3 | CNDSALDyNEAkGVY | Y427 | 0.309 | 92.95 | 999 | 0.135554 | 0.309 | 0.1356 | 1 |
| Q93084 | ATP2A3 | YEkVGEAtETALTCL | T441 | 0.108 | 88.71 | 999 | 0.073765 | 0.176 | 0.1202 | 1 |
| Q99536 | VAT1 | APGPGQLtLRLRACG | T80 | 0.031 | 98.8 | 393 | 0.021173 | 0.031 | 0.0212 | 1 |
| Q99536 | VAT1 | ENVTVFGtASASKHE | T220 | 0.046 | 98.77 | 393 | 0.031419 | 0.046 | 0.0314 | 1 |
| Q99536 | VAT1 | IDyHTTDyVDEIkKI | Y245 | 0.132 | 98.53 | 393 | 0.057907 | 0.233 | 0.0946 | 1 |
| O75051 | PLXNA2 | DDPKFHSyVsLPFGC | Y293 | 0.061 | 96.6 | 1894 | 0.02676 | 0.122 | 0.0535 | 1 |
| O75051 | PLXNA2 | PKFHSyVsLPFGCTR | S295 | 0 | 95.97 | 1894 | 0 | 0 | 0 | 1 |
| O75051 | PLXNA2 | TSRDRMTsVASYVYN | S434 | 0 | 97.23 | 1894 | 0 | 0 | 0 | 1 |
| O75051 | PLXNA2 | SYVYNGYsVVFVGTK | S444 | 0.011 | 94.73 | 1894 | 0.008744 | 0.011 | 0.0087 | 1 |
| Q4G0P3 | HYDIN | EEVNRMLtPsEFLKE | T47 | 0.181 | 88.63 | 5121 | 0.123625 | 0.181 | 0.1236 | 1 |

| | | | | | | | | | | |
|---|---|---|---|---|---|---|---|---|---|---|
| Q4G0P3 | HYDIN | CSNTVQKyELALVVD | Y669 | 0.079 | 88.905 | 5121 | 0.034656 | 0.079 | 0.0347 | 1 |
| Q6NXR4 | TTI2 | VNRLGILtVRHLKRL | T377 | 0.001 | 95.46 | 508 | 0.000683 | 0.001 | 0.0007 | 1 |
| O75443 | TECTA | NEsELCDsVARYASA | S956 | 0.057 | 87.08 | 2155 | 0.04531 | 0.073 | 0.0626 | 1 |
| Q13905 | RAPGEF1 | RLEWQKQtsEGLAEY | T950 | 0.071 | 92.58 | 1077 | 0.048494 | 0.071 | 0.0485 | 0.8333 |
| P41594 | GRM5 | LGCEIRDsCWHSAVA | S98 | 0.004 | 94.64 | 1212 | 0.00318 | 0.004 | 0.0032 | 1 |
| P41594 | GRM5 | ACFCEGMtVRGLLMA | T282 | 0 | 92.07 | 1212 | 0 | 0 | 0 | 1 |
| Q6P1R4 | DUS1L | RHGAQLCyTPMLHAQ | Y45 | 0.001 | 98.48 | 473 | 0.000439 | 0.001 | 0.0004 | 0.9412 |
| Q9Y2X0 | MED16 | TVTRVCDyHtKLFLI | Y547 | 0.339 | 89.88 | 877 | 0.148714 | 0.339 | 0.1487 | 0.8 |
| Q15257 | PTPA | VFKVFNRyLEVMRKL | Y223 | 0.016 | 98.9 | 358 | 0.007019 | 0.016 | 0.007 | 0.9231 |
| Q14249 | ENDOG | VHAYHRAtNADYRGS | T128 | 0.154 | 98.81 | 297 | 0.105184 | 0.154 | 0.1052 | 1 |
| Q6PL24 | TMED8 | KLGKEKNsRLVVKRG | S181 | 0.113 | 95.25 | 325 | 0.089825 | 0.113 | 0.0898 | 1 |
| Q6PL24 | TMED8 | VVtIRVPtHPEGKRV | T197 | 0.102 | 97.52 | 325 | 0.069667 | 0.102 | 0.0697 | 1 |
| Q6PL24 | TMED8 | MPVYRRDsHRDVQAG | S284 | 0 | 97.91 | 325 | 0 | 0 | 0 | 0.8 |
| Q9UKG9 | CROT | LEESLKKyLEsVkPF | Y35 | 0 | 98.88 | 612 | 0 | 0 | 0 | 0.9333 |
| Q9UKG9 | CROT | INQAKAQyLREASDL | Y387 | 0.025 | 98.06 | 612 | 0.010967 | 0.025 | 0.011 | 1 |
| Q9UKG9 | CROT | TSFGKKLtKNKMLHP | T409 | 0 | 97.95 | 612 | 0 | 0 | 0 | 1 |
| Q9UKG9 | CROT | QLALQLAyyRLHGHP | Y428 | 0.047 | 98.92 | 612 | 0.020618 | 0.047 | 0.0206 | 1 |
| Q9H222 | ABCG5 | GtFLGEVyVNGRALR | Y114 | 0.287 | 89.71 | 651 | 0.125903 | 0.287 | 0.1259 | 1 |
| O95626 | ANP32D | NLIHLNLsGNKIkDL | S96 | 0.152 | 97.9 | 131 | 0.120827 | 0.152 | 0.1208 | 1 |
| Q15397 | PUM3 | NMLTEELyGNtFQLy | Y276 | 0.126 | 98.31 | 648 | 0.055274 | 0.126 | 0.0553 | 0.9091 |
| Q15397 | PUM3 | TEELyGNtFQLyksA | T279 | 0.158 | 97.54 | 648 | 0.107916 | 0.158 | 0.1079 | 1 |
| P61011 | SRP54 | EIIIVDTsGRHKQED | S192 | 0.032 | 85.59 | 504 | 0.025437 | 0.032 | 0.0254 | 1 |
| Q9NZQ9 | TMOD4 | DMLRENRsLQSLNIE | S253 | 0.038 | 97.96 | 345 | 0.030207 | 0.038 | 0.0302 | 1 |
| Q8NEE8 | TTC16 | YYLYRAKsRQLLQNI | S449 | 0 | 92.81 | 873 | 0 | 0 | 0 | 1 |
| Q8NEE8 | TTC16 | GARQDVAtVLLLNPK | T465 | 0.075 | 91.47 | 873 | 0.051226 | 0.075 | 0.0512 | 1 |
| Q14642 | INPP5A | YsDTLEStPMLEKEK | T143 | 0.148 | 93.13 | 412 | 0.101086 | 0.148 | 0.1011 | 0.8 |
| P41231 | P2RY2 | TWNASTTyMFHLAVS | Y71 | 0.146 | 93.44 | 377 | 0.064048 | 0.228 | 0.0772 | 0.8462 |
| P29692 | EEF1D | PALVAKSsILLDVkP | S197 | 0.161 | 90.19 | 281 | 0.127981 | 0.161 | 0.128 | 1 |
| Q9NQ66 | PLCB1 | LMEPLEKyPLEsGVP | Y444 | 0.24 | 97.95 | 1216 | 0.105285 | 0.303 | 0.1329 | 1 |
| P51398 | DAP3 | VWNKREstEkGsPLG | T216 | 0.126 | 90.2 | 398 | 0.08606 | 0.126 | 0.0861 | 1 |
| P51398 | DAP3 | ItRVRNAtDAVGIVL | T237 | 0.011 | 93.73 | 398 | 0.007513 | 0.011 | 0.0075 | 1 |

| | | | | | | | | | | |
|---|---|---|---|---|---|---|---|---|---|---|
| P51398 | DAP3 | LkELKRQssLGMFHL | S251 | 0.026 | 87.44 | 398 | 0.020668 | 0.026 | 0.0207 | 0.9 |
| P51398 | DAP3 | INALWGRttLKREDk | T272 | 0.132 | 92.47 | 398 | 0.090158 | 0.132 | 0.0902 | 0.9 |
| Q8TDZ2 | MICAL1 | GKLAIGItANFVNGR | T241 | 0.022 | 97.22 | 1067 | 0.015026 | 0.022 | 0.015 | 1 |
| Q8TDZ2 | MICAL1 | LYQLLsQtsPENMHR | T446 | 0.079 | 90.43 | 1067 | 0.053958 | 0.079 | 0.054 | 1 |
| Q15058 | KIF14 | FAYGQtGsGKSYTMM | S451 | 0.015 | 92.57 | 1648 | 0.011924 | 0.015 | 0.0119 | 1 |
| Q15393 | SF3B3 | EVFGVIRsLMAFRLt | S64 | 0.055 | 97.35 | 1217 | 0.04372 | 0.076 | 0.0642 | 1 |
| Q15393 | SF3B3 | GRIVILEyQPskNMF | Y92 | 0.106 | 96.81 | 1217 | 0.046501 | 0.106 | 0.0465 | 1 |
| Q15393 | SF3B3 | kGRAVMIsAIEkQkL | S133 | 0.004 | 97.66 | 1217 | 0.00318 | 0.004 | 0.0032 | 1 |
| Q15393 | SF3B3 | VAFPLQYtPRkFVIH | T784 | 0.053 | 98.18 | 1217 | 0.0362 | 0.053 | 0.0362 | 1 |
| Q15393 | SF3B3 | IETDHNAyTEATkAQ | Y807 | 0.081 | 95.44 | 1217 | 0.035534 | 0.133 | 0.048 | 1 |
| Q15393 | SF3B3 | SNTGEDWyVLVGVAK | Y904 | 0.094 | 98.07 | 1217 | 0.041236 | 0.094 | 0.0412 | 1 |
| Q15393 | SF3B3 | SVAGGFVyTYKLVNN | Y926 | 0.196 | 98.62 | 1217 | 0.085982 | 0.298 | 0.1307 | 1 |
| Q15393 | SF3B3 | PNkQkNVsEELDRtP | S1194 | 0.003 | 95.23 | 1217 | 0.002385 | 0.003 | 0.0024 | 1 |
| P55265 | ADAR | DLSGkLGtPKKEINR | T167 | 0.121 | 86.04 | 1226 | 0.082645 | 0.141 | 0.0963 | 0.8333 |
| Q9BWD1 | ACAT2 | AVCLAVQsIGIGDSS | S105 | 0.15 | 98.11 | 397 | 0.119237 | 0.15 | 0.1192 | 0.875 |
| Q15056 | EIF4H | VEFDEVDsLKEALty | S94 | 0.111 | 82.65 | 248 | 0.088236 | 0.111 | 0.0882 | 1 |
| Q6ZQN7 | SLCO4C1 | TMKFALFtsGVALTL | T460 | 0.004 | 97.19 | 724 | 0.002732 | 0.004 | 0.0027 | 1 |
| Q6ZQN7 | SLCO4C1 | MKFALFtsGVALTLS | S461 | 0.012 | 96.15 | 724 | 0.009539 | 0.012 | 0.0095 | 1 |
| O60437 | PPL | HLSSLQDyMQRCTNE | Y218 | 0.028 | 94.7 | 1756 | 0.012283 | 0.056 | 0.0246 | 1 |
| Q96E11 | MRRF | ISQMADDtVAELDRH | T245 | 0.137 | 97.13 | 262 | 0.093573 | 0.137 | 0.0936 | 1 |
| P62495 | ETF1 | RMEKRHNyVRKVAEt | Y201 | 0.066 | 93.65 | 437 | 0.028953 | 0.066 | 0.029 | 1 |
| P62495 | ETF1 | ENLDIMRyVLHCQGT | Y331 | 0.116 | 89.07 | 437 | 0.050888 | 0.23 | 0.1002 | 1 |
| O95267 | RASGRP1 | HLEPEELsEHLTyLE | S210 | 0.009 | 94.33 | 797 | 0.007154 | 0.009 | 0.0072 | 1 |
| Q96P47 | AGAP3 | RHLGAHLsRVRSLDL | S676 | 0.061 | 96.35 | 875 | 0.04849 | 0.061 | 0.0485 | 1 |
| Q96P47 | AGAP3 | RGLTPLAyARRAGSQ | Y825 | 0.104 | 96.97 | 875 | 0.045623 | 0.104 | 0.0456 | 1 |
| P10586 | PTPRF | EESDQGKyECVATNS | Y205 | 0.032 | 93.05 | 1907 | 0.014038 | 0.032 | 0.014 | 1 |
| Q49AG3 | ZBED5 | AVQLSNStVARRIkD | T247 | 0.172 | 95.04 | 693 | 0.117478 | 0.172 | 0.1175 | 1 |
| Q3KP31 | ZNF791 | ECAKTFIsLENFRRH | S423 | 0.155 | 88.18 | 576 | 0.123212 | 0.155 | 0.1232 | 0.8 |
| Q6NTF7 | APOBEC3H | LEELDKNsRAIKRRL | S170 | 0.004 | 98.18 | 200 | 0.00318 | 0.004 | 0.0032 | 0.9091 |
| Q8NG68 | TTL | SSVYAEVsRLLLAtG | S18 | 0.081 | 98.39 | 377 | 0.064388 | 0.081 | 0.0644 | 1 |
| Q16877 | PFKFB4 | IVQVKLGsPDYVNRD | S176 | 0.172 | 80.99 | 469 | 0.136725 | 0.172 | 0.1367 | 1 |

| | | | | | | | | | | |
|---|---|---|---|---|---|---|---|---|---|---|
| Q16877 | PFKFB4 | EDLDRDLsYIKIMDV | S214 | 0.173 | 97.86 | 469 | 0.13752 | 0.173 | 0.1375 | 1 |
| Q16877 | PFKFB4 | IGGDPGLsPRGREFA | S275 | 0.006 | 98.49 | 469 | 0.004769 | 0.006 | 0.0048 | 1 |
| Q9H6R3 | ACSS3 | SAGAIEEsILSHGTV | S574 | 0.067 | 80.68 | 686 | 0.053259 | 0.067 | 0.0533 | 0.8889 |
| Q15008 | PSMD6 | DYKTFVTyTVyVsMI | Y222 | 0.282 | 90.07 | 389 | 0.123709 | 0.282 | 0.1237 | 0.9167 |
| Q15008 | PSMD6 | TFVTyTVyVsMIALE | Y225 | 0.092 | 91.32 | 389 | 0.040359 | 0.092 | 0.0404 | 1 |
| Q15008 | PSMD6 | VTyTVyVsMIALERP | S227 | 0.001 | 91.93 | 389 | 0.000795 | 0.001 | 0.0008 | 1 |
| Q96SM3 | CPXM1 | FRWLstVyAGSNLAM | Y536 | 0 | 98.78 | 734 | 0 | 0 | 0 | 0.8 |
| Q9ULQ1 | TPCN1 | SARERYLtFKALNQN | T382 | 0.004 | 93.52 | 816 | 0.002732 | 0.004 | 0.0027 | 1 |
| Q16851 | UGP2 | DVkGGTLtQyEGkLR | T296 | 0.016 | 87.09 | 508 | 0.010928 | 0.016 | 0.0109 | 1 |
| Q16851 | UGP2 | SKFKIFNtNNLWISL | T329 | 0.038 | 91.8 | 508 | 0.025955 | 0.038 | 0.026 | 1 |
| Q16851 | UGP2 | RFLPVKTtSDLLLVM | T398 | 0.078 | 90.88 | 508 | 0.053275 | 0.078 | 0.0533 | 1 |
| Q16851 | UGP2 | sFtkVQDyLRRFEsI | Y442 | 0.008 | 91.56 | 508 | 0.003509 | 0.008 | 0.0035 | 1 |
| O75083 | WDR1 | PkGNNFLyTNGkCVI | Y34 | 0.016 | 98.57 | 606 | 0.007019 | 0.016 | 0.007 | 1 |
| O75083 | WDR1 | NPALADIytEHAHQV | Y54 | 0.052 | 97.41 | 606 | 0.022812 | 0.104 | 0.0456 | 1 |
| O75083 | WDR1 | PSGFyIAsGDVsGkL | S75 | 0 | 98.82 | 606 | 0 | 0 | 0 | 1 |
| O75083 | WDR1 | GHNKVINsVDIkQSR | S151 | 0.013 | 98.82 | 606 | 0.010334 | 0.013 | 0.0103 | 1 |
| O75083 | WDR1 | SRPYRLAtGsDDNCA | T164 | 0 | 98.85 | 606 | 0 | 0 | 0 | 1 |
| O75083 | WDR1 | PYRLAtGsDDNCAAF | S166 | 0 | 98.76 | 606 | 0 | 0 | 0 | 1 |
| O75083 | WDR1 | kAHDGGIyAIsWSPD | Y238 | 0.287 | 98.72 | 606 | 0.125903 | 0.337 | 0.1478 | 1 |
| O75083 | WDR1 | DGGIyAIsWSPDSTH | S241 | 0.14 | 98.7 | 606 | 0.111288 | 0.181 | 0.1439 | 1 |
| O75083 | WDR1 | SHDGHINyWDsETGE | Y348 | 0.237 | 95.69 | 606 | 0.103968 | 0.237 | 0.104 | 1 |
| O75083 | WDR1 | GHINyWDsETGENDs | S351 | 0.099 | 94.4 | 606 | 0.078697 | 0.142 | 0.1328 | 1 |
| O75083 | WDR1 | SGQLISCsMDDTVRY | S383 | 0 | 98.52 | 606 | 0 | 0 | 0 | 1 |
| O75083 | WDR1 | TsLMLrDysGQGVVk | Y398 | 0.235 | 90.17 | 606 | 0.103091 | 0.345 | 0.1208 | 1 |
| O75083 | WDR1 | VDGNVRLysILGTTL | Y472 | 0.056 | 98.22 | 606 | 0.024566 | 0.056 | 0.0246 | 1 |
| O75083 | WDR1 | HRLHHVssLAWLDEH | S581 | 0.06 | 98.34 | 606 | 0.047695 | 0.12 | 0.0954 | 1 |
| P34972 | CNR2 | CLRyPPSyKALLTRG | Y141 | 0.253 | 92.17 | 360 | 0.110987 | 0.341 | 0.1496 | 0.8182 |
| Q6XZB0 | LIPI | KRPCLEFsQLSVKDS | S25 | 0.177 | 93.78 | 460 | 0.1407 | 0.177 | 0.1407 | 1 |
| Q6XZB0 | LIPI | IEtILMMytRNNLNC | Y48 | 0.1 | 98.07 | 460 | 0.043869 | 0.1 | 0.0439 | 1 |
| Q6XZB0 | LIPI | EtILMMytRNNLNCA | T49 | 0.055 | 97.74 | 460 | 0.037566 | 0.055 | 0.0376 | 1 |
| P51946 | CCNH | ___MYHNsSQkRHWT | S5 | 0.158 | 95.78 | 323 | 0.125597 | 0.158 | 0.1256 | 1 |

| | | | | | | | | | | |
|---|---|---|---|---|---|---|---|---|---|---|
| Q9BZL4 | PPP1R12C | HVAAAKGyIEVMRLL | Y239 | 0.277 | 96.06 | 782 | 0.121516 | 0.277 | 0.1215 | 0.8889 |
| Q9H0H0 | INTS2 | CPPLMDItLHMLNGY | T877 | 0.012 | 94.91 | 1204 | 0.008196 | 0.097 | 0.0663 | 1 |
| Q12797 | ASPH | LVRkYPQsPRARYGK | S374 | 0.008 | 96.16 | 758 | 0.006359 | 0.008 | 0.0064 | 1 |
| Q12797 | ASPH | LRGAIETyQEVASLP | Y405 | 0.044 | 98.02 | 758 | 0.019302 | 0.044 | 0.0193 | 1 |
| Q12797 | ASPH | KNDLGVGyLLIGDND | Y464 | 0.311 | 95.62 | 758 | 0.136431 | 0.311 | 0.1364 | 1 |
| Q12797 | ASPH | NDNAkKVyEEVLSVt | Y477 | 0 | 97.64 | 758 | 0 | 0 | 0 | 1 |
| Q12797 | ASPH | IEsGDPGtDDGRFYF | T522 | 0.01 | 95.95 | 758 | 0.00683 | 0.01 | 0.0068 | 1 |
| Q12797 | ASPH | WtPkETGyTELVkSL | Y583 | 0.278 | 97.93 | 758 | 0.121955 | 0.278 | 0.122 | 1 |
| Q92819 | HAS2 | GkREVMYtAFRALGR | T196 | 0.002 | 97.15 | 552 | 0.001366 | 0.002 | 0.0014 | 1 |
| Q92819 | HAS2 | DTMLDPAsSVEMVKV | S221 | 0 | 95.93 | 552 | 0 | 0 | 0 | 1 |
| Q92819 | HAS2 | VLSLGYAtKYTARSK | T328 | 0 | 98.64 | 552 | 0 | 0 | 0 | 1 |
| Q9UGI0 | ZRANB1 | RGETLGytRFQGVYL | T556 | 0.091 | 77.02 | 708 | 0.062154 | 0.091 | 0.0622 | 1 |
| Q9H6T3 | RPAP3 | LEEAKKDyERVLELE | Y224 | 0.131 | 96.61 | 665 | 0.057468 | 0.131 | 0.0575 | 1 |
| Q8NBJ7 | SUMF2 | AIDIFPVtNKDFRDF | T70 | 0.073 | 98.9 | 301 | 0.04986 | 0.073 | 0.0499 | 1 |
| Q5SW79 | CEP170 | ___MSLtsWFLVsSG | S5 | 0.056 | 83.08 | 1584 | 0.044515 | 0.135 | 0.1073 | 1 |
| Q5SW79 | CEP170 | LtsWFLVsSGGTRHR | S10 | 0.07 | 92.86 | 1584 | 0.055644 | 0.084 | 0.0668 | 0.875 |
| Q5SW79 | CEP170 | HLVkDLGsLNGTFVN | S62 | 0.039 | 95.65 | 1584 | 0.031002 | 0.039 | 0.031 | 1 |
| Q15652 | JMJD1C | DHKHLMPtQIIPGsV | T1905 | 0.003 | 94.53 | 2540 | 0.002049 | 0.06 | 0.041 | 1 |
| Q9UJT9 | FBXL7 | ESRLRYLsIAHCGRV | S356 | 0.059 | 98.86 | 491 | 0.0469 | 0.059 | 0.0469 | 1 |
| Q8N4J0 | CARNMT1 | kQFVRDWsETGkAER | S170 | 0.027 | 98.86 | 409 | 0.021463 | 0.027 | 0.0215 | 0.9 |
| Q14185 | DOCK1 | GKELAEQyENEMFDY | Y1304 | 0.006 | 93.92 | 1865 | 0.002632 | 0.006 | 0.0026 | 1 |
| P82675 | MRPS5 | ISLRFKRtHIKMKKQ | T301 | 0.007 | 97.24 | 430 | 0.004781 | 0.007 | 0.0048 | 1 |
| P82675 | MRPS5 | SGSINMLsLtQGLFR | S344 | 0.124 | 97.51 | 430 | 0.098569 | 0.124 | 0.0986 | 1 |
| P82675 | MRPS5 | SINMLsLtQGLFRGL | T346 | 0.006 | 97.86 | 430 | 0.004098 | 0.006 | 0.0041 | 1 |
| A9UHW6 | MIF4GD | IQSFDAEtQQLLkTA | T18 | 0.199 | 94.37 | 222 | 0.13592 | 0.199 | 0.1359 | 1 |
| Q99638 | RAD9A | SRIGDELyLEPLEDG | Y28 | 0.329 | 96.11 | 391 | 0.144328 | 0.329 | 0.1443 | 0.9091 |
| Q13501 | SQSTM1 | PGGFQAHyRDEDGDL | Y67 | 0.211 | 96.06 | 440 | 0.092563 | 0.211 | 0.0926 | 0.9167 |
| Q15155 | NOMO1 | LLRDGENyVVLLDST | Y1110 | 0.082 | 93.34 | 1222 | 0.035972 | 0.082 | 0.036 | 1 |
| Q15155 | NOMO1 | STLPRSQyDyILPQV | Y1123 | 0.214 | 89.41 | 1222 | 0.093879 | 0.214 | 0.0939 | 1 |
| Q15155 | NOMO1 | LPRSQyDyILPQVSF | Y1125 | 0.204 | 90.04 | 1222 | 0.089492 | 0.204 | 0.0895 | 1 |
| Q8NGP6 | OR5M8 | PMLNLIIysLRNKNV | Y289 | 0.052 | 82.5 | 311 | 0.022812 | 0.052 | 0.0228 | 1 |

| | | | | | | | | | | |
|---|---|---|---|---|---|---|---|---|---|---|
| Q8NGP6 | OR5M8 | MLNLIIysLRNKNVK | S290 | 0.093 | 75.95 | 311 | 0.073927 | 0.093 | 0.0739 | 1 |
| Q13698 | CACNA1S | FKGKMHKtCYFIGtD | T225 | 0.031 | 93.07 | 1873 | 0.021173 | 0.031 | 0.0212 | 1 |
| Q9UBI4 | STOML1 | SKDGAVLsVGADVQF | S140 | 0.121 | 92.8 | 398 | 0.096185 | 0.121 | 0.0962 | 0.8571 |
| Q9NRW4 | DUSP22 | NKVTHILsVHDSARP | S36 | 0.022 | 98.87 | 184 | 0.017488 | 0.022 | 0.0175 | 1 |
| P14416 | DRD2 | AISIDRYtAVAMPML | T134 | 0.124 | 90.18 | 443 | 0.084694 | 0.124 | 0.0847 | 1 |
| P54922 | ADPRH | AAGDALGyyNGKWEF | Y19 | 0.068 | 98.77 | 357 | 0.029831 | 0.068 | 0.0298 | 1 |
| Q9Y247 | FAM50B | PAsRWEAyDPEKKWD | Y313 | 0.324 | 89.98 | 325 | 0.142134 | 0.324 | 0.1421 | 1 |
| Q14152 | EIF3A | LHRLyHLsREMRkNL | S303 | 0.135 | 86.63 | 1382 | 0.107313 | 0.135 | 0.1073 | 1 |
| P18077 | RPL35A | SkAIFAGykRGLRNQ | Y14 | 0.156 | 98.37 | 110 | 0.068435 | 0.156 | 0.0684 | 1 |
| P18077 | RPL35A | LRNQREHtALLkIEG | T25 | 0.093 | 98.33 | 110 | 0.06352 | 0.093 | 0.0635 | 1 |
| P18077 | RPL35A | ARDETEFyLGkRCAY | Y42 | 0.296 | 97.34 | 110 | 0.129851 | 0.296 | 0.1299 | 1 |
| Q15366 | PCBP2 | TIAGIPQsIIECVkQ | S154 | 0.134 | 89.49 | 365 | 0.106519 | 0.134 | 0.1065 | 1 |
| P23443 | RPS6KB1 | sIHDGtVtHtFCGtI | T250 | 0.189 | 80.79 | 525 | 0.12909 | 0.189 | 0.1291 | 1 |
| P23443 | RPS6KB1 | QVFLGFtyVAPsVLE | Y413 | 0.223 | 89.18 | 525 | 0.097827 | 0.223 | 0.0978 | 1 |
| Q6NXG1 | ESRP1 | MELIDDNtVVRARGL | T225 | 0.123 | 97.33 | 681 | 0.084011 | 0.123 | 0.084 | 1 |
| Q12866 | MERTK | QVHNATCtVRIAAVT | T458 | 0.163 | 92.21 | 999 | 0.111331 | 0.163 | 0.1113 | 1 |
| Q9HAV0 | GNB4 | DATLVQItSNMDSVG | T34 | 0.069 | 96.73 | 340 | 0.047128 | 0.069 | 0.0471 | 1 |
| Q9HAV0 | GNB4 | YDSRLLVsAsQDGkL | S72 | 0 | 98.86 | 340 | 0 | 0 | 0 | 1 |
| Q9HAV0 | GNB4 | SRLLVsAsQDGkLII | S74 | 0 | 98.64 | 340 | 0 | 0 | 0 | 1 |
| Q99437 | ATP6V0B | GIyITGSsIIGGGVK | S75 | 0.155 | 94.92 | 205 | 0.123212 | 0.155 | 0.1232 | 1 |
| Q9BT22 | ALG1 | VLGDVGRsPRMQYHA | S46 | 0.11 | 93.82 | 464 | 0.087441 | 0.11 | 0.0874 | 1 |
| O14975 | SLC27A2 | GAKVLLVsPELQAAV | S158 | 0.055 | 97.58 | 620 | 0.04372 | 0.055 | 0.0437 | 1 |
| O14975 | SLC27A2 | FWDDCRkyNVTVIQY | Y304 | 0.293 | 98.45 | 620 | 0.128535 | 0.293 | 0.1285 | 1 |
| O14975 | SLC27A2 | EGRIGMAsIKMKENH | S528 | 0 | 94.56 | 620 | 0 | 0 | 0 | 0.8889 |
| Q96FS4 | SIPA1 | LVsRGCEtRELALPR | T685 | 0.153 | 95.69 | 1042 | 0.104501 | 0.153 | 0.1045 | 0.8182 |
| Q13310 | PABPC4 | RDMITRRsLGyAyVN | S51 | 0.164 | 92.02 | 644 | 0.130366 | 0.164 | 0.1304 | 1 |
| Q9H3Q3 | GAL3ST2 | PWLFLARyVEGVGSQ | Y102 | 0.227 | 97.83 | 398 | 0.099582 | 0.227 | 0.0996 | 1 |
| Q02218 | OGDH | RVTDRNItLsLVANP | T367 | 0.051 | 96.18 | 1023 | 0.034834 | 0.051 | 0.0348 | 1 |
| Q02218 | OGDH | TDRNItLsLVANPsH | S369 | 0.003 | 95.97 | 1023 | 0.002385 | 0.003 | 0.0024 | 1 |
| Q02218 | OGDH | MFTQPLMyKQIRkQk | Y527 | 0.089 | 96.8 | 1023 | 0.039043 | 0.089 | 0.039 | 1 |
| Q02218 | OGDH | MVGQVAItRIEQLsP | T928 | 0 | 98.4 | 1023 | 0 | 0 | 0 | 1 |

| | | | | | | | | | | |
|---|---|---|---|---|---|---|---|---|---|---|
| Q02218 | OGDH | EHKNQGYyDyVkPRL | Y966 | 0.151 | 97.77 | 1023 | 0.066242 | 0.151 | 0.0662 | 1 |
| Q02218 | OGDH | KNQGYyDyVkPRLRT | Y968 | 0.158 | 97.77 | 1023 | 0.069312 | 0.158 | 0.0693 | 1 |
| Q9NVP1 | DDX18 | GTGVLILsPtRELAM | S258 | 0 | 92.63 | 670 | 0 | 0 | 0 | 1 |
| Q9NVP1 | DDX18 | LLDHMQNtPGFMyKN | T319 | 0.164 | 91.35 | 670 | 0.112014 | 0.164 | 0.112 | 0.8 |
| Q9NVP1 | DDX18 | LLPTRRQtMLFSATQ | T360 | 0.016 | 94.35 | 670 | 0.010928 | 0.016 | 0.0109 | 0.8571 |
| P62826 | RAN | NLQyyDIsAkSNyNF | S150 | 0.006 | 97.58 | 216 | 0.004769 | 0.006 | 0.0048 | 0.8571 |
| P18206 | VCL | ALLMAEMsrLVRGGS | S1002 | 0 | 91.54 | 1134 | 0 | 0 | 0 | 1 |
| P23458 | JAK1 | ELPkDIsyKRyIPET | Y217 | 0.072 | 91.72 | 1154 | 0.031585 | 0.072 | 0.0316 | 1 |
| O95817 | BAG3 | GkKTDKkyLMIEEyL | Y451 | 0.146 | 97.77 | 575 | 0.064048 | 0.146 | 0.064 | 0.9 |
| P48728 | AMT | GAPMRAHsPILNMEG | S334 | 0.155 | 98.44 | 403 | 0.123212 | 0.155 | 0.1232 | 1 |
| P48728 | AMT | TKIGTVTsGCPsPSL | S349 | 0.048 | 98.69 | 403 | 0.038156 | 0.048 | 0.0382 | 0.875 |
| P48728 | AMT | TVTsGCPsPSLkKNV | S353 | 0.054 | 98.74 | 403 | 0.042925 | 0.054 | 0.0429 | 0.875 |
| Q9UIF7 | MUTYH | MKKVFRVyQGQQPGT | Y494 | 0.121 | 94.63 | 546 | 0.053081 | 0.121 | 0.0531 | 1 |
| P35367 | HRH1 | VGNLYIVsLsVADLI | S68 | 0.083 | 97.91 | 487 | 0.065978 | 0.112 | 0.0951 | 0.8889 |
| Q12884 | FAP | PCSQSVRsVFAVNWI | S551 | 0.107 | 98.65 | 760 | 0.085056 | 0.107 | 0.0851 | 0.8889 |
| Q12884 | FAP | KDDNLEHyKNstVMA | Y677 | 0.002 | 98.75 | 760 | 0.000877 | 0.002 | 0.0009 | 1 |
| Q9NWU5 | MRPL22 | PRRPAEIyHCRRQIK | Y68 | 0.255 | 98.09 | 206 | 0.111865 | 0.255 | 0.1119 | 1 |
| Q9BXK5 | BCL2L13 | RQMLLELtRRGQEPL | T168 | 0.121 | 93.22 | 485 | 0.082645 | 0.121 | 0.0826 | 0.9091 |
| Q92556 | ELMO1 | VLNSHDLyQKVAQEI | Y216 | 0.049 | 93.65 | 727 | 0.021496 | 0.049 | 0.0215 | 1 |
| Q92556 | ELMO1 | KsMyTRDyKKLGFIN | Y356 | 0.27 | 94.91 | 727 | 0.118445 | 0.27 | 0.1184 | 1 |
| Q92556 | ELMO1 | AKHHQDAyIRIVLEN | Y395 | 0.002 | 97.91 | 727 | 0.000877 | 0.002 | 0.0009 | 1 |
| Q92556 | ELMO1 | HECPFGRssIELTkM | S416 | 0.006 | 98.28 | 727 | 0.004769 | 0.006 | 0.0048 | 1 |
| Q92556 | ELMO1 | ECPFGRssIELTkML | S417 | 0 | 98.73 | 727 | 0 | 0 | 0 | 1 |
| Q92556 | ELMO1 | CIQLLNKtWkEMRAt | T466 | 0.006 | 98.2 | 727 | 0.004098 | 0.006 | 0.0041 | 1 |
| O00476 | SLC17A3 | SSFLMGAsRGFSSIA | S355 | 0.022 | 91.14 | 420 | 0.017488 | 0.022 | 0.0175 | 0.9 |
| O43306 | ADCY6 | QDsMWVVsYVVLGIL | S212 | 0 | 90.58 | 1168 | 0 | 0.031 | 0.0246 | 1 |
| Q9P109 | GCNT4 | EKSFPIAySLVVHKD | Y135 | 0.03 | 98.94 | 453 | 0.013161 | 0.03 | 0.0132 | 0.9 |
| Q9P109 | GCNT4 | NYYEGFFyPsCtGsH | Y379 | 0.148 | 90.99 | 453 | 0.064925 | 0.148 | 0.0649 | 0.8462 |
| Q9HD20 | ATP13A1 | LRDGkEVtPVSSIPV | T558 | 0.11 | 88.38 | 1204 | 0.075131 | 0.11 | 0.0751 | 1 |
| Q9HD20 | ATP13A1 | VSSIPVEtHRALASC | T567 | 0.019 | 91.12 | 1204 | 0.012977 | 0.019 | 0.013 | 1 |
| Q9HD20 | ATP13A1 | LGstDLCyIAAVKGA | Y649 | 0.085 | 92.19 | 1204 | 0.037288 | 0.085 | 0.0373 | 1 |

| | | | | | | | | | | |
|---|---|---|---|---|---|---|---|---|---|---|
| Q9HD20 | ATP13A1 | FSQCPPDyHHIHTEI | Y671 | 0.034 | 87.49 | 1204 | 0.014915 | 0.034 | 0.0149 | 1 |
| Q9HD20 | ATP13A1 | ARVLALGykELGHLT | Y690 | 0.129 | 94.19 | 1204 | 0.05659 | 0.129 | 0.0566 | 1 |
| Q9HD20 | ATP13A1 | FVGFIVVsCPLKADS | S722 | 0.184 | 86.44 | 1204 | 0.146264 | 0.184 | 0.1463 | 0.8333 |
| Q9HD20 | ATP13A1 | QKEFVItsLKELGyV | S851 | 0.021 | 91.38 | 1204 | 0.016693 | 0.021 | 0.0167 | 1 |
| P30101 | PDIA3 | CkrLAPEyEAAATrL | Y67 | 0.039 | 98 | 505 | 0.017109 | 0.039 | 0.0171 | 1 |
| P30101 | PDIA3 | DGEEAGAyDGPrTAD | Y115 | 0.176 | 93.28 | 505 | 0.077209 | 0.176 | 0.0772 | 1 |
| P30101 | PDIA3 | kKQAGPAsVPLrtEE | S136 | 0.044 | 95.16 | 505 | 0.034976 | 0.044 | 0.035 | 1 |
| P30101 | PDIA3 | ANDVPsPyEVrGFPt | Y445 | 0.143 | 95.15 | 505 | 0.062732 | 0.143 | 0.0627 | 1 |
| P30101 | PDIA3 | VrGFPtIyFsPANkK | Y454 | 0.185 | 98.5 | 505 | 0.081157 | 0.331 | 0.1452 | 1 |
| P30101 | PDIA3 | ELsDFIsyLQREAtN | Y479 | 0.218 | 98.14 | 505 | 0.095633 | 0.218 | 0.0956 | 1 |
| Q9Y334 | VWA7 | AKLALLAsIQAFSLL | S288 | 0 | 97.98 | 891 | 0 | 0 | 0 | 0.9 |
| Q9Y334 | VWA7 | YKAVALAsGGEVIFt | S475 | 0.013 | 94.68 | 891 | 0.010334 | 0.013 | 0.0103 | 0.9091 |
| P45985 | MAP2K4 | LATGRFPyPKWNSVF | Y307 | 0.247 | 93.81 | 399 | 0.108355 | 0.281 | 0.1233 | 1 |
| O95544 | NADK | ARNTAWVsFDGRKRQ | S381 | 0.103 | 98.38 | 446 | 0.081876 | 0.103 | 0.0819 | 1 |
| Q7Z494 | NPHP3 | DHPRVAQsLHQLASV | S987 | 0.009 | 89.56 | 1330 | 0.007154 | 0.009 | 0.0072 | 1 |
| Q7Z494 | NPHP3 | FGNAEQLyKQALEIS | Y1008 | 0.104 | 93.71 | 1330 | 0.045623 | 0.104 | 0.0456 | 1 |
| Q8TEM1 | NUP210 | KAAkQGDtILVsGMk | T205 | 0.062 | 89.97 | 1887 | 0.042347 | 0.062 | 0.0423 | 1 |
| Q8N0W3 | FCSK | SDTPPLAyELGGAVL | Y729 | 0.24 | 95.06 | 1084 | 0.105285 | 0.24 | 0.1053 | 1 |
| Q8IXT1 | DDIAS | IILVSKRsNCPKCGs | S39 | 0.103 | 81.91 | 998 | 0.081876 | 0.103 | 0.0819 | 1 |
| Q8NDG6 | TDRD9 | TRLIYMTtGVLLQKI | T232 | 0.066 | 93.55 | 1382 | 0.045079 | 0.066 | 0.0451 | 0.8 |
| Q96S38 | RPS6KC1 | RPLWNLRsPAEELKA | S332 | 0.154 | 86.19 | 1066 | 0.122417 | 0.154 | 0.1224 | 1 |
| Q6YHK3 | CD109 | LKAyFLGsKSSMAVH | S450 | 0.017 | 87.53 | 1445 | 0.013514 | 0.03 | 0.0253 | 1 |
| Q6YHK3 | CD109 | ISVtQPDsIVGIVAV | S585 | 0 | 93.57 | 1445 | 0 | 0 | 0 | 1 |
| Q9H4A6 | GOLPH3 | ALkHVKEtQPPEtVQ | T143 | 0.096 | 97.4 | 298 | 0.065569 | 0.096 | 0.0656 | 1 |
| Q9H4A6 | GOLPH3 | KEtQPPEtVQNWIEL | T148 | 0.13 | 98.41 | 298 | 0.088792 | 0.13 | 0.0888 | 1 |
| Q9H4A6 | GOLPH3 | WNPLKLHyQLRNVRE | Y168 | 0.255 | 97.84 | 298 | 0.111865 | 0.255 | 0.1119 | 1 |
| Q8N9R8 | SCAI | LVVKKLRyYARFIVV | Y181 | 0.042 | 97.74 | 606 | 0.018425 | 0.042 | 0.0184 | 1 |
| O60641 | SNAP91 | LDyLIQAtNEtNVNI | T49 | 0.004 | 92.06 | 907 | 0.002732 | 0.004 | 0.0027 | 1 |
| Q01995 | TAGLN | LKNGVILsKLVNsLy | S63 | 0 | 96.46 | 201 | 0 | 0 | 0 | 1 |
| Q01995 | TAGLN | FLKAAEDyGVIktDM | Y104 | 0.113 | 95.38 | 201 | 0.049571 | 0.223 | 0.0971 | 1 |
| Q01995 | TAGLN | DMAAVQRtLMALGsL | T129 | 0.003 | 97.95 | 201 | 0.002049 | 0.003 | 0.002 | 1 |

| | | | | | | | | | | |
|---|---|---|---|---|---|---|---|---|---|---|
| Q01995 | TAGLN | RtLMALGsLAVtKND | S135 | 0.116 | 95.83 | 201 | 0.09221 | 0.149 | 0.1277 | 1 |
| Q01995 | TAGLN | ALGsLAVtKNDGHyR | T139 | 0.083 | 92.97 | 201 | 0.05669 | 0.166 | 0.1134 | 1 |
| Q8NB66 | UNC13C | KTGsSDPyVTVQVGK | Y1244 | 0.17 | 94.57 | 2214 | 0.074577 | 0.17 | 0.0746 | 0.9167 |
| Q6MZM0 | HEPHL1 | SIYKKAVyRRFTDGT | Y79 | 0.009 | 98.48 | 1159 | 0.003948 | 0.009 | 0.0039 | 1 |
| O75146 | HIP1R | KVKANKHsPHLSRLQ | S929 | 0.021 | 90.39 | 1068 | 0.016693 | 0.021 | 0.0167 | 1 |
| P16402 | H1-3 | kGtGAsGsFkLNkkA | S105 | 0.155 | 95.9 | 221 | 0.123212 | 0.155 | 0.1232 | 0.875 |
| P50851 | LRBA | TVLLIMHtLKYYYWA | T620 | 0.071 | 97.215 | 2863 | 0.048494 | 0.111 | 0.0723 | 1 |
| P50851 | LRBA | QALKAMGyFLKHLAP | Y745 | 0.137 | 97.545 | 2863 | 0.0601 | 0.137 | 0.0601 | 1 |
| Q8IVV2 | LOXHD1 | TDANVFItLFGTQDD | T992 | 0.063 | 90.46 | 2067 | 0.04303 | 0.063 | 0.043 | 1 |
| P04075 | ALDOA | VLkIGEHtPsALAIM | T158 | 0.113 | 98.68 | 364 | 0.077181 | 0.113 | 0.0772 | 0.8333 |
| P04075 | ALDOA | HHIyLEGtLLkPNMV | T227 | 0.033 | 98.96 | 364 | 0.022539 | 0.033 | 0.0225 | 0.9231 |
| P04075 | ALDOA | VTGItFLsGGQsEEE | S272 | 0.086 | 96.94 | 364 | 0.068363 | 0.142 | 0.1162 | 1 |
| P04075 | ALDOA | LLkPWALtFsYGRAL | T299 | 0.03 | 98.92 | 364 | 0.02049 | 0.05 | 0.033 | 1 |
| P04075 | ALDOA | VkRALANsLACQGky | S336 | 0.128 | 98.71 | 364 | 0.101749 | 0.128 | 0.1017 | 1 |
| P04075 | ALDOA | sLACQGkytPsGQAG | Y343 | 0.227 | 97.3 | 364 | 0.099582 | 0.227 | 0.0996 | 1 |
| Q96CM8 | ACSF2 | StPIGGLsyVQGCtk | S59 | 0 | 97.35 | 615 | 0 | 0 | 0 | 1 |
| Q96CM8 | ACSF2 | GyCVMLGyWGEPQKT | Y472 | 0.151 | 97.08 | 615 | 0.066242 | 0.302 | 0.1325 | 0.9231 |
| P09619 | PDGFRB | CTKGGPIyIITEyCR | Y678 | 0.275 | 93.55 | 1106 | 0.120639 | 0.275 | 0.1206 | 1 |
| Q8N5T2 | TBC1D19 | YHEPSKLyQIFREMY | Y387 | 0.029 | 98.53 | 526 | 0.012722 | 0.029 | 0.0127 | 0.875 |
| Q96KP4 | CNDP2 | TFFKDVDyVCISDNY | Y190 | 0.156 | 98.85 | 475 | 0.068435 | 0.156 | 0.0684 | 1 |
| Q96KP4 | CNDP2 | GKkkPCItYGLRGIC | T207 | 0 | 98.79 | 475 | 0 | 0 | 0 | 1 |
| Q96KP4 | CNDP2 | VMLLPVGsADDGAHs | S439 | 0.095 | 98.54 | 475 | 0.075517 | 0.095 | 0.0755 | 1 |
| O96014 | WNT11 | KLMRLHNsEVGRQAL | S194 | 0.038 | 98.01 | 354 | 0.030207 | 0.038 | 0.0302 | 1 |
| O96014 | WNT11 | GRQALRAsLEMKCKC | S204 | 0.06 | 97.99 | 354 | 0.047695 | 0.06 | 0.0477 | 1 |
| O96014 | WNT11 | AADLKTRyLSAtKVV | Y241 | 0.136 | 97.97 | 354 | 0.059661 | 0.136 | 0.0597 | 0.8462 |
| O96014 | WNT11 | KTSNGsDsCDLMCCG | S308 | 0.075 | 97.08 | 354 | 0.059619 | 0.075 | 0.0596 | 1 |
| P14867 | GABRA1 | VSDHDMEytIDVFFR | Y87 | 0.006 | 96.95 | 456 | 0.002632 | 0.006 | 0.0026 | 1 |
| P14867 | GABRA1 | TEDGTLLyTMRLTVR | Y156 | 0.167 | 96.81 | 456 | 0.07326 | 0.328 | 0.1439 | 1 |
| Q86X24 | HORMAD1 | KIsTEHQsLVLVKRL | S26 | 0 | 93.51 | 394 | 0 | 0 | 0 | 1 |
| Q86X24 | HORMAD1 | VPKRRKFsEPKEHI_ | S388 | 0 | 83.62 | 394 | 0 | 0.057 | 0.0453 | 1 |
| Q9Y2L5 | TRAPPC8 | KSPMVRKyAFHMILA | Y560 | 0.018 | 96.08 | 1435 | 0.007896 | 0.018 | 0.0079 | 1 |

| | | | | | | | | | | |
|---|---|---|---|---|---|---|---|---|---|---|
| Q9Y2L5 | TRAPPC8 | LHILGVVyNLGTIQG | Y841 | 0 | 96.11 | 1435 | 0 | 0 | 0 | 1 |
| Q9C0C4 | SEMA4C | kKLLFAGsRSQLVQL | S488 | 0.001 | 98.12 | 833 | 0.000795 | 0.001 | 0.0008 | 0.875 |
| Q16798 | ME3 | QQSDLDKyIILMtLQ | Y107 | 0.136 | 96.16 | 604 | 0.059661 | 0.136 | 0.0597 | 0.8667 |
| Q16798 | ME3 | EGRGIFAsGsPFKSV | S467 | 0.016 | 95.08 | 604 | 0.012719 | 0.016 | 0.0127 | 1 |
| Q16798 | ME3 | RGIFAsGsPFKSVTL | S469 | 0.086 | 84.79 | 604 | 0.068363 | 0.086 | 0.0684 | 0.875 |
| Q16798 | ME3 | LStIRDVsLRIAIKV | S545 | 0 | 98.74 | 604 | 0 | 0 | 0 | 1 |
| Q8IZJ3 | CPAMD8 | AQPQAPGyLIAAPsV | Y33 | 0.15 | 90.46 | 1885 | 0.065803 | 0.15 | 0.0658 | 1 |
| Q8IZJ3 | CPAMD8 | QFKPGLAyVGKVELS | Y362 | 0.122 | 87.9 | 1885 | 0.05352 | 0.122 | 0.0535 | 1 |
| Q8IZJ3 | CPAMD8 | IYTsEVVsQRGLVGF | S400 | 0.047 | 90.59 | 1885 | 0.037361 | 0.047 | 0.0374 | 1 |
| Q01433 | AMPD2 | RKAPVLQyLYYLAQI | Y667 | 0 | 98.35 | 879 | 0 | 0.145 | 0.0636 | 1 |
| P43354 | NR4A2 | CQHyGVRtCEGCKGF | T279 | 0.016 | 97.24 | 598 | 0.010928 | 0.016 | 0.0109 | 0.8333 |
| Q14767 | LTBP2 | APTRMDCySGQKGHA | Y1414 | 0.289 | 84.83 | 1821 | 0.12678 | 0.289 | 0.1268 | 1 |
| Q92685 | ALG3 | CASYRVHsIFVLRLF | S166 | 0.03 | 95.17 | 438 | 0.023847 | 0.03 | 0.0238 | 1 |
| Q8N9B8 | RASGEF1A | KAKLKsFsAKIVQLL | S114 | 0.037 | 92.26 | 481 | 0.029412 | 0.133 | 0.1057 | 1 |
| Q96DM1 | PGBD4 | VVPCFEIyHTKKNy_ | Y579 | 0.236 | 90.07 | 585 | 0.10353 | 0.283 | 0.1241 | 0.8 |
| O75618 | DEDD | HRDVRVLsFLFVDVI | S46 | 0.185 | 92.59 | 318 | 0.147059 | 0.185 | 0.1471 | 1 |
| Q7L2E3 | DHX30 | MYPVkEHyLEDILAk | Y622 | 0.216 | 95.68 | 1194 | 0.094756 | 0.346 | 0.1308 | 1 |
| Q7L2E3 | DHX30 | sVFVRDSsQVHPLAV | S1092 | 0.001 | 95.88 | 1194 | 0.000795 | 0.001 | 0.0008 | 1 |
| Q7L2E3 | DHX30 | PLAVLLLtDGDVHIR | T1103 | 0.086 | 89.03 | 1194 | 0.058739 | 0.163 | 0.1103 | 1 |
| Q7L2E3 | DHX30 | LGRMVERsLRsELAA | S1152 | 0.01 | 92.78 | 1194 | 0.007949 | 0.01 | 0.0079 | 1 |
| P02144 | MB | RKDMAsNyKELGFQG | Y147 | 0.014 | 97.8 | 154 | 0.006142 | 0.014 | 0.0061 | 1 |
| Q9Y566 | SHANK1 | DLGGSPNyKDRRGLT | Y275 | 0.228 | 92.92 | 2161 | 0.10002 | 0.228 | 0.1 | 0.8333 |
| P99999 | CYCS | HGLFGRktGQAPGys | T41 | 0.011 | 98.61 | 105 | 0.007513 | 0.011 | 0.0075 | 0.8 |
| Q8TBK2 | SETD6 | HELQAPAsRWRPYFA | S138 | 0.117 | 96.89 | 473 | 0.093005 | 0.117 | 0.093 | 1 |
| Q14894 | CRYM | PVTKHRGyLGVMPAY | Y58 | 0.191 | 95.37 | 314 | 0.083789 | 0.191 | 0.0838 | 0.8333 |
| Q495B1 | ANKDD1A | GMNALLLsAWFGHLR | S98 | 0 | 97.37 | 522 | 0 | 0.037 | 0.0294 | 1 |
| P54764 | EPHA4 | GEFGEVCsGRLKVPG | S637 | 0.164 | 92.62 | 986 | 0.130366 | 0.164 | 0.1304 | 1 |
| Q08945 | SSRP1 | VYQEVkGsMNDGrLR | S17 | 0.097 | 78.07 | 709 | 0.077107 | 0.097 | 0.0771 | 0.8 |
| P98160 | HSPG2 | EsDQGAytCEAMNAR | T478 | 0.021 | 90.97 | 4391 | 0.014343 | 0.042 | 0.0287 | 1 |
| Q460N5 | PARP14 | VkINHRHyTVNLNTy | Y1576 | 0.157 | 92.15 | 1801 | 0.068874 | 0.265 | 0.0988 | 0.9091 |
| Q13275 | SEMA3F | yEVLFLGtDRGTVQK | T485 | 0 | 98.2 | 785 | 0 | 0 | 0 | 0.9 |

| | | | | | | | | | | |
|---|---|---|---|---|---|---|---|---|---|---|
| Q5W0U4 | BSPRY | FNEkCTRsPLLTQLW | S211 | 0.05 | 93.83 | 402 | 0.039746 | 0.05 | 0.0397 | 1 |
| Q9Y4B4 | RAD54L2 | GGVLLMGyEMyRLLt | Y404 | 0.153 | 92.44 | 1467 | 0.067119 | 0.153 | 0.0671 | 0.9286 |
| Q9Y4B4 | RAD54L2 | LLMGyEMyRLLtLKk | Y407 | 0.01 | 93.48 | 1467 | 0.004387 | 0.01 | 0.0044 | 1 |
| Q9Y4B4 | RAD54L2 | YTLEKKIyDRQIskQ | Y875 | 0.029 | 89.95 | 1467 | 0.012722 | 0.029 | 0.0127 | 1 |
| Q9Y4B4 | RAD54L2 | QIskQGMsDRVVDDL | S885 | 0.063 | 84.34 | 1467 | 0.05008 | 0.063 | 0.0501 | 0.8889 |
| Q9BUB4 | ADAT1 | IGSLQARsQIsKVEL | S431 | 0.003 | 96.59 | 502 | 0.002385 | 0.003 | 0.0024 | 1 |
| Q9BUB4 | ADAT1 | LQARsQIsKVELFRS | S434 | 0.003 | 97.55 | 502 | 0.002385 | 0.003 | 0.0024 | 0.8889 |
| Q92698 | RAD54L | PDFVFMLsSKAGGCG | S572 | 0.133 | 88.36 | 747 | 0.105724 | 0.133 | 0.1057 | 1 |
| Q7Z6L1 | TECPR1 | GHDGVRDsILFIYYV | S645 | 0.021 | 90.54 | 1165 | 0.016693 | 0.021 | 0.0167 | 1 |
| Q7Z6L1 | TECPR1 | PIALRDVsIIPEsPG | S938 | 0.025 | 97.83 | 1165 | 0.019873 | 0.025 | 0.0199 | 1 |
| Q7Z6L1 | TECPR1 | SNNVCRVsVGPLDQV | S1082 | 0.022 | 97.76 | 1165 | 0.017488 | 0.022 | 0.0175 | 1 |
| Q9Y5I4 | PCDHAC2 | LsAsVAItVAVVDRV | T677 | 0.157 | 87.78 | 1007 | 0.107233 | 0.157 | 0.1072 | 1 |
| Q6IMI4 | SULT6B1 | ITMCTSEtFQALDTF | T44 | 0.018 | 93.94 | 303 | 0.012294 | 0.018 | 0.0123 | 1 |
| O00400 | SLC33A1 | FsAADAVtGLKLVEE | T331 | 0.087 | 96.43 | 549 | 0.059422 | 0.195 | 0.1332 | 1 |
| P19086 | GNAZ | IKLLLLGtSNSGKST | T41 | 0.084 | 96.32 | 355 | 0.057373 | 0.123 | 0.079 | 1 |
| P19086 | GNAZ | IIFCVELsGyDLKLY | S229 | 0.045 | 97.42 | 355 | 0.035771 | 0.048 | 0.0531 | 0.8571 |
| P19086 | GNAZ | FCVELsGyDLKLYED | Y231 | 0.042 | 97.63 | 355 | 0.018425 | 0.042 | 0.0184 | 1 |
| P19086 | GNAZ | NNNWFINtsLILFLN | T263 | 0.127 | 92.03 | 355 | 0.086743 | 0.127 | 0.0867 | 1 |
| P19086 | GNAZ | QFVFDAVtDVIIQNN | T341 | 0.007 | 92.04 | 355 | 0.004781 | 0.007 | 0.0048 | 1 |
| P15924 | DSP | TEDMLKVyEARLTEE | Y787 | 0.089 | 92.78 | 2871 | 0.039043 | 0.2885 | 0.1148 | 1 |
| Q04864 | REL | GkDCRDGyyEAEFGQ | Y88 | 0.262 | 97.17 | 619 | 0.114936 | 0.319 | 0.1399 | 1 |
| Q04864 | REL | kDCRDGyyEAEFGQE | Y89 | 0.225 | 98.03 | 619 | 0.098704 | 0.225 | 0.0987 | 1 |
| Q86XM0 | CATSPERD | TREDNLYyGNLGIVP | Y317 | 0.012 | 94.88 | 798 | 0.005264 | 0.012 | 0.0053 | 1 |
| P55884 | EIF3B | RMTLDTLsIyETPSM | S447 | 0.095 | 90.55 | 814 | 0.075517 | 0.095 | 0.0755 | 1 |
| P55884 | EIF3B | GIKDFSWsPGGNIIA | S473 | 0.049 | 93.17 | 814 | 0.038951 | 0.049 | 0.039 | 1 |
| P55884 | EIF3B | kGtQGVVtNFEIFRM | T542 | 0.018 | 88.82 | 814 | 0.012294 | 0.018 | 0.0123 | 1 |
| P12956 | XRCC6 | ykYsGRDsLIFLVDA | S37 | 0.034 | 89.4 | 609 | 0.027027 | 0.034 | 0.027 | 0.875 |
| P12956 | XRCC6 | LAVVFYGtEkDkNSV | T90 | 0 | 93.47 | 609 | 0 | 0 | 0 | 0.9 |
| Q86X83 | COMMD2 | TYLLTESsKLMISEL | S73 | 0.015 | 92.45 | 199 | 0.011924 | 0.015 | 0.0119 | 1 |
| A6NH52 | TVP23A | LLSLDFWsVKNVTGR | S76 | 0.078 | 95.96 | 213 | 0.062003 | 0.078 | 0.062 | 0.8571 |
| P43146 | DCC | HKPDEGLyQCEASLG | Y115 | 0 | 95.2 | 1447 | 0 | 0 | 0 | 1 |

| | | | | | | | | | | |
|---|---|---|---|---|---|---|---|---|---|---|
| P43146 | DCC | GDSGSIIsRTAKVAV | S129 | 0.031 | 91.99 | 1447 | 0.024642 | 0.088 | 0.07 | 1 |
| P06213 | INSR | GCtVINGsLIINIRG | S366 | 0.01 | 97.05 | 1382 | 0.007949 | 0.01 | 0.0079 | 1 |
| P06213 | INSR | GYLKIRRsyALVSLs | S400 | 0.005 | 90.35 | 1382 | 0.003975 | 0.005 | 0.004 | 1 |
| P06213 | INSR | GLRHFTGyRIELQAC | Y818 | 0.176 | 92.64 | 1382 | 0.077209 | 0.176 | 0.0772 | 1 |
| P06213 | INSR | SVRIRATsLAGNGSW | S929 | 0.003 | 92.76 | 1382 | 0.002385 | 0.068 | 0.0563 | 0.9091 |
| A6NES4 | MROH2A | sLEKGFLyRALGFTL | Y688 | 0.073 | 92.72 | 1674 | 0.032024 | 0.073 | 0.032 | 0.9333 |
| O43520 | ATP8B1 | CCINGQIyGDHRDAS | Y476 | 0.001 | 89.47 | 1251 | 0.000439 | 0.001 | 0.0004 | 1 |
| Q4AC94 | C2CD3 | AARTVSIsVQIIRAC | S1200 | 0.104 | 90.19 | 2353 | 0.082671 | 0.104 | 0.0827 | 1 |
| Q13474 | DRP2 | MRALSFKtGIACLCG | T498 | 0.03 | 97.25 | 957 | 0.02049 | 0.03 | 0.0205 | 1 |
| Q14028 | CNGB1 | CYYFAVKtLITIGGL | T842 | 0 | 91.07 | 1251 | 0 | 0 | 0 | 1 |
| P49327 | FASN | LWKDNWVsFMDtMLQ | S1028 | 0.091 | 93.93 | 2511 | 0.072337 | 0.178 | 0.1419 | 0.875 |
| Q5T4T1 | TMEM170B | HRQGRVIsVIAVSIG | S71 | 0.033 | 92.3 | 132 | 0.026232 | 0.033 | 0.0262 | 1 |
| Q12931 | TRAP1 | QLTSLSEyASRMRAG | Y486 | 0.052 | 97.96 | 704 | 0.022812 | 0.052 | 0.0228 | 0.8571 |
| Q12931 | TRAP1 | RAGtRNIyyLCAPNr | Y498 | 0.296 | 97.52 | 704 | 0.129851 | 0.296 | 0.1299 | 1 |
| Q12931 | TRAP1 | AGtRNIyyLCAPNrH | Y499 | 0.081 | 98.15 | 704 | 0.035534 | 0.081 | 0.0355 | 0.8462 |
| Q12931 | TRAP1 | NrHLAEHsPyyEAMk | S511 | 0.105 | 96.04 | 704 | 0.083466 | 0.105 | 0.0835 | 1 |
| Q12931 | TRAP1 | LAEHsPyyEAMkKKD | Y514 | 0.286 | 95.35 | 704 | 0.125464 | 0.286 | 0.1255 | 1 |
| Q8NCE0 | TSEN2 | QPTFRTTyMAyHyFR | Y343 | 0 | 98.16 | 465 | 0 | 0 | 0 | 1 |
| Q8NCE0 | TSEN2 | FRTTyMAyHyFRSKG | Y346 | 0.066 | 98.37 | 465 | 0.028953 | 0.066 | 0.029 | 1 |
| P51857 | AKR1D1 | NQVECHPyFTQPKLL | Y199 | 0.086 | 98.03 | 326 | 0.037727 | 0.086 | 0.0377 | 1 |
| Q6NXE6 | ARMC6 | LkVLIEAtkAFLDNP | T283 | 0.04 | 97.88 | 501 | 0.027321 | 0.04 | 0.0273 | 1 |
| Q9BX10 | GTPBP2 | SDMPRkItEVLVRKV | T157 | 0 | 90.61 | 602 | 0 | 0 | 0 | 1 |
| Q96FJ0 | STAMBPL1 | kNDLLKkyNVEYQEY | Y129 | 0.002 | 97.95 | 436 | 0.000877 | 0.002 | 0.0009 | 1 |
| Q9P241 | ATP10D | VNNRIRttKYTLLNF | T79 | 0.219 | 76.88 | 1426 | 0.14958 | 0.219 | 0.1496 | 0.8 |
| Q14683 | SMC1A | IGPFQRFtAIIGPNG | T28 | 0 | 93.71 | 1233 | 0 | 0 | 0 | 1 |
| Q14683 | SMC1A | AFVSMVYsEEGAEDr | S85 | 0.137 | 90.45 | 1233 | 0.108903 | 0.137 | 0.1089 | 1 |
| Q14683 | SMC1A | kVVQLHEySEELEkL | Y117 | 0.003 | 92.82 | 1233 | 0.001316 | 0.003 | 0.0013 | 1 |
| Q14683 | SMC1A | MKNPkERtALFEEIS | T152 | 0.049 | 91.97 | 1233 | 0.033468 | 0.098 | 0.0669 | 1 |
| Q14683 | SMC1A | LFEEISrsGELAQEy | S161 | 0 | 91.13 | 1233 | 0 | 0 | 0 | 1 |
| Q14683 | SMC1A | sGELAQEyDkRKKEM | Y168 | 0.011 | 95.46 | 1233 | 0.004826 | 0.011 | 0.0048 | 1 |
| Q14683 | SMC1A | VRDkFQEtSDEFEAA | T1041 | 0.006 | 89.61 | 1233 | 0.004098 | 0.006 | 0.0041 | 1 |

| | | | | | | | | | | |
|---|---|---|---|---|---|---|---|---|---|---|
| P68871 | HBB | kGtFAtLsELHCDkL | S90 | 0.003 | 97.54 | 147 | 0.002385 | 0.003 | 0.0024 | 1 |
| P68871 | HBB | tPPVQAAyQkVVAGV | Y131 | 0.003 | 98.43 | 147 | 0.001316 | 0.003 | 0.0013 | 0.8824 |
| P40692 | MLH1 | KNtHPFLyLsLEISP | Y293 | 0.102 | 96.67 | 756 | 0.044746 | 0.102 | 0.0447 | 1 |
| P40692 | MLH1 | tHPFLyLsLEISPQN | S295 | 0.023 | 95.49 | 756 | 0.018283 | 0.023 | 0.0183 | 1 |
| P29279 | CCN2 | MGIStRVtNDNASCR | T222 | 0.011 | 85.62 | 349 | 0.007513 | 0.011 | 0.0075 | 1 |
| P58397 | ADAMTS12 | GNYKLAGtVFQyDRK | T770 | 0.101 | 83.81 | 1594 | 0.068984 | 0.119 | 0.0813 | 1 |
| P58397 | ADAMTS12 | LAGtVFQyDRKGDLE | Y774 | 0.048 | 92.02 | 1594 | 0.021057 | 0.048 | 0.0211 | 1 |
| Q96B02 | UBE2W | GMTLNEKsVQNsITQ | S29 | 0.176 | 73.4 | 151 | 0.139905 | 0.176 | 0.1399 | 1 |
| Q6UXI7 | VIT | RkLMILItDGRSYDD | T601 | 0.077 | 96.44 | 678 | 0.052592 | 0.077 | 0.0526 | 1 |
| Q9NQY0 | BIN3 | QKDMKkstDADLAMs | T49 | 0.038 | 97.93 | 253 | 0.025955 | 0.038 | 0.026 | 1 |
| P23458 | JAK1 | IEVQkGRySLHGSDR | Y507 | 0.239 | 97.76 | 1154 | 0.104846 | 0.239 | 0.1048 | 1 |
| Q969I6 | SLC38A4 | ILVPtIKyIFGFIGA | Y476 | 0.292 | 90.97 | 547 | 0.128096 | 0.292 | 0.1281 | 1 |
| P22105 | TNXB | PLTADGttAPERKEE | T1245 | 0.047 | 87.78 | 4244 | 0.032102 | 0.047 | 0.0321 | 0.9231 |
| P42345 | MTOR | ICTVGRLsSMNPAFV | S719 | 0.097 | 85.02 | 2549 | 0.077107 | 0.187 | 0.1494 | 1 |
| P12081 | HARS1 | ETQVLVAsAQkKLLE | S415 | 0 | 98.07 | 509 | 0 | 0 | 0 | 1 |
| A6NHR9 | SMCHD1 | RIDFLPHyDtLVKSG | Y121 | 0.194 | 90.68 | 2005 | 0.085105 | 0.194 | 0.0851 | 1 |
| P25788 | PSMA3 | DLSASTFsPDGRVFQ | S16 | 0.088 | 93.13 | 255 | 0.069952 | 0.088 | 0.07 | 1 |
| P25788 | PSMA3 | kAVENSStAIGIRCk | T36 | 0.003 | 98.16 | 255 | 0.002049 | 0.003 | 0.002 | 1 |
| Q92636 | NSMAF | GSSVFTTsQDSTLKM | S649 | 0 | 98.32 | 917 | 0 | 0 | 0 | 1 |
| Q92636 | NSMAF | DSTLKMFsKESKMLQ | S658 | 0.102 | 90.56 | 917 | 0.081081 | 0.102 | 0.0811 | 1 |
| Q8NFW1 | COL22A1 | DLVFLLDtSSSVGKE | T45 | 0.03 | 95.83 | 1626 | 0.02049 | 0.03 | 0.0205 | 1 |
| Q8NFW1 | COL22A1 | yHGGNTNtGDALRYI | T117 | 0.031 | 96.37 | 1626 | 0.021173 | 0.031 | 0.0212 | 1 |
| Q9P107 | GMIP | PFGKWtLsSAAQTHQ | S488 | 0.175 | 92.84 | 970 | 0.13911 | 0.175 | 0.1391 | 0.8889 |
| Q00688 | FKBP3 | DEALLtMsKGEkARL | S183 | 0.004 | 91.4 | 224 | 0.00318 | 0.004 | 0.0032 | 1 |
| P01042 | KNG1 | IDFVAREttCsKEsN | T326 | 0 | 92.3 | 644 | 0 | 0 | 0 | 1 |
| Q9BRJ2 | MRPL45 | RMKKtMAsQVsIRRI | S118 | 0.025 | 88.92 | 306 | 0.019873 | 0.171 | 0.1359 | 1 |
| Q9BRJ2 | MRPL45 | KtMAsQVsIRRIkDY | S121 | 0.101 | 92.68 | 306 | 0.080286 | 0.101 | 0.0803 | 1 |
| Q9UNW1 | MINPP1 | VALIRHGtRYPTVKQ | T91 | 0.032 | 98.59 | 487 | 0.021856 | 0.064 | 0.0437 | 0.8 |
| Q9UNW1 | MINPP1 | DDAKVLEyLNDLKQY | Y315 | 0 | 98.77 | 487 | 0 | 0 | 0 | 0.8571 |
| Q93008 | USP9X | AFRGKHLsFVVRFPN | S887 | 0.016 | 87.1 | 2554 | 0.012719 | 0.016 | 0.0127 | 1 |
| Q9UHL0 | DDX25 | GHQVsLLsGELTVEQ | S369 | 0.077 | 85.81 | 483 | 0.061208 | 0.114 | 0.0906 | 0.8 |

| | | | | | | | | | | |
|---|---|---|---|---|---|---|---|---|---|---|
| P05455 | SSB | ICHQIEyyFGDFNLP | Y24 | 0.199 | 93.31 | 408 | 0.087298 | 0.199 | 0.0873 | 1 |
| P05455 | SSB | DkTKIRRsPskPLPE | S92 | 0.1 | 92.25 | 408 | 0.079491 | 0.1 | 0.0795 | 1 |
| O60832 | DKC1 | SYLRRVVyPLEKLLT | Y287 | 0.301 | 96.38 | 514 | 0.132044 | 0.301 | 0.132 | 1 |
| Q8WXI4 | ACOT11 | PSNQVYLsYNNVSSL | S388 | 0.051 | 90.34 | 607 | 0.040541 | 0.051 | 0.0405 | 0.9 |
| Q8WXI4 | ACOT11 | AQAFLLLsDLRQRPE | S444 | 0.109 | 92.15 | 607 | 0.086646 | 0.127 | 0.1111 | 0.8333 |
| Q15149 | PLEC | ISLVIRGtQGAEEVL | T1235 | 0.062 | 94.54 | 4684 | 0.042347 | 0.062 | 0.0423 | 1 |
| Q8IZ81 | ELMOD2 | HPKLGYSyAIVGINL | Y204 | 0.115 | 98.36 | 293 | 0.050449 | 0.115 | 0.0504 | 1 |
| Q96DE0 | NUDT16 | PRVVAHFyAKRLtLE | Y111 | 0.002 | 98.71 | 195 | 0.000877 | 0.002 | 0.0009 | 0.8667 |
| Q14511 | NEDD9 | RNkVMNSsNQLCEQL | S780 | 0.007 | 93.33 | 834 | 0.005564 | 0.007 | 0.0056 | 1 |
| Q5HY92 | FIGN | GLTALPRsILLFGPR | S521 | 0.008 | 94.54 | 759 | 0.006359 | 0.008 | 0.0064 | 1 |
| Q92887 | ABCC2 | RLHTIMDsDKVMVLD | S1501 | 0.133 | 86.45 | 1545 | 0.105724 | 0.133 | 0.1057 | 1 |
| Q9BTC0 | DIDO1 | FQVTDNRyKsKYRSI | Y721 | 0 | 94.24 | 2240 | 0 | 0 | 0 | 0.9231 |
| Q9Y6R7 | FCGBP | HRGSRRVsYVGLVTV | S526 | 0 | 87.28 | 5405 | 0 | 0 | 0 | 0.9 |
| Q7Z3Z4 | PIWIL4 | EITYVDyyKQQYDIT | Y340 | 0.097 | 94.33 | 852 | 0.042553 | 0.097 | 0.0426 | 1 |
| Q7Z3Z4 | PIWIL4 | LNQPMLVsLLKKKRN | S358 | 0.017 | 93.78 | 852 | 0.013514 | 0.017 | 0.0135 | 1 |
| Q6VVB1 | NHLRC1 | LTFTKENsLLVLDTA | S376 | 0.092 | 94.4 | 395 | 0.073132 | 0.092 | 0.0731 | 1 |
| Q03113 | GNA12 | LLLGAGEsGkSTFLk | S68 | 0.017 | 97 | 381 | 0.013514 | 0.017 | 0.0135 | 1 |
| P07948 | LYN | ILLYEIVtyGKIPYP | T438 | 0.133 | 97.25 | 512 | 0.090841 | 0.133 | 0.0908 | 1 |
| P54296 | MYOM2 | FCEVRDTsLVMLWKA | S828 | 0.135 | 84.58 | 1465 | 0.107313 | 0.135 | 0.1073 | 1 |
| P54296 | MYOM2 | GSSPVSGyFVDFREE | Y847 | 0.195 | 94.75 | 1465 | 0.085544 | 0.195 | 0.0855 | 0.9333 |
| P63010 | AP2B1 | kLQNNNVyTIAkRNV | Y874 | 0.318 | 96.92 | 937 | 0.139502 | 0.318 | 0.1395 | 1 |
| Q8NFY9 | KBTBD8 | GNHQRMFtVEAyDIE | T498 | 0.163 | 93.42 | 601 | 0.111331 | 0.163 | 0.1113 | 1 |
| Q8NFY9 | KBTBD8 | RMFtVEAyDIELNKW | Y502 | 0.014 | 96.64 | 601 | 0.006142 | 0.014 | 0.0061 | 1 |
| Q8NE71 | ABCF1 | YIVAGRRyGLVGPNG | Y332 | 0.074 | 96.88 | 845 | 0.032463 | 0.074 | 0.0325 | 1 |
| P42330 | AKR1C3 | AGFRHIDsAHLyNNE | S51 | 0 | 98.67 | 323 | 0 | 0 | 0 | 0.9 |
| P42330 | AKR1C3 | VkREDIFytskLWST | Y81 | 0.013 | 98.93 | 323 | 0.005703 | 0.013 | 0.0057 | 1 |
| P42330 | AKR1C3 | VDLYLIHsPMSLKPG | S118 | 0.103 | 97.74 | 323 | 0.081876 | 0.153 | 0.1405 | 1 |
| P42330 | AKR1C3 | DIVDLCTtWEAMEKC | T147 | 0 | 98.91 | 323 | 0 | 0 | 0 | 1 |
| P42330 | AKR1C3 | NQVECHPyFNRSkLL | Y196 | 0.088 | 98.78 | 323 | 0.038604 | 0.088 | 0.0386 | 1 |
| P42330 | AKR1C3 | GLDRNLHyFNSDSFA | Y305 | 0.053 | 96.25 | 323 | 0.02325 | 0.093 | 0.0408 | 1 |
| P78357 | CNTNAP1 | LHGIsGWsPRIGDPN | S62 | 0.108 | 96.82 | 1384 | 0.085851 | 0.108 | 0.0859 | 1 |

| | | | | | | | | | | |
|---|---|---|---|---|---|---|---|---|---|---|
| Q9BSJ8 | ESYT1 | kGLIEGksDPyALVR | S356 | 0.004 | 86.44 | 1104 | 0.00318 | 0.004 | 0.0032 | 1 |
| P01569 | IFNA5 | LPQtHSLsNRRtLMI | S33 | 0.024 | 90.09 | 189 | 0.019078 | 0.024 | 0.0191 | 1 |
| P01569 | IFNA5 | HSLsNRRtLMIMAQM | T37 | 0 | 95.19 | 189 | 0 | 0.101 | 0.069 | 1 |
| Q9Y2K3 | MYH15 | LHNRENQsILFTGES | S171 | 0 | 93.57 | 1946 | 0 | 0 | 0 | 1 |
| Q9UQ52 | CNTN6 | QQEDEGFyECIASNL | Y295 | 0.005 | 95.86 | 1028 | 0.002193 | 0.005 | 0.0022 | 1 |
| Q86YT9 | JAML | KDEyVLYyySNLSVP | Y73 | 0.18 | 94.13 | 394 | 0.078963 | 0.18 | 0.079 | 1 |
| Q7Z745 | MROH2B | HKQLMLEsIIRGLYH | S1348 | 0.026 | 90.97 | 1585 | 0.020668 | 0.048 | 0.0382 | 0.9 |
| Q8TD84 | DSCAML1 | SEIIEAKtHGREPSF | T1474 | 0 | 82.85 | 2053 | 0 | 0 | 0 | 0.9231 |
| Q8TCB7 | METTL6 | MLRFKASsKLGENFY | S204 | 0.166 | 97.06 | 284 | 0.131956 | 0.166 | 0.132 | 1 |
| Q5TIA1 | MEI1 | PHMKEKFsKKLASSs | S594 | 0 | 94.25 | 1274 | 0 | 0 | 0 | 1 |
| Q14952 | KIR2DS3 | SCSSWSSyDMYHLST | Y155 | 0.028 | 97.87 | 304 | 0.012283 | 0.028 | 0.0123 | 1 |
| P06756 | ITGAV | ARtLGMVyIYDGKNM | Y284 | 0.328 | 98.6 | 1048 | 0.143889 | 0.328 | 0.1439 | 1 |
| P06756 | ITGAV | EDkKGIVyIFNGRST | Y404 | 0.11 | 98.65 | 1048 | 0.048255 | 0.11 | 0.0483 | 1 |
| P06756 | ITGAV | RDkLTPItIFMEyRL | T586 | 0.099 | 92.22 | 1048 | 0.067618 | 0.099 | 0.0676 | 1 |
| P06756 | ITGAV | PItIFMEyRLDYRTA | Y591 | 0.056 | 95.26 | 1048 | 0.024566 | 0.056 | 0.0246 | 1 |
| P06756 | ITGAV | KFDLQIQsSNLFDKV | S749 | 0.167 | 92.91 | 1048 | 0.132751 | 0.167 | 0.1328 | 1 |
| P16591 | FER | YVSNVSKsWLLMIQQ | S70 | 0.117 | 92.12 | 822 | 0.093005 | 0.145 | 0.147 | 0.875 |
| Q9H5I5 | PIEZO2 | KsSPLLVyLRNNLLM | Y1056 | 0.125 | 91.895 | 2752 | 0.054836 | 0.247 | 0.1073 | 1 |
| Q13822 | ENPP2 | NLCKsytsCCHDFDE | S85 | 0.075 | 92.94 | 863 | 0.059619 | 0.075 | 0.0596 | 1 |
| Q13822 | ENPP2 | PHERRILtILQWLTL | T288 | 0.016 | 96.27 | 863 | 0.010928 | 0.016 | 0.0109 | 1 |
| Q13822 | ENPP2 | PsVYAFYsEQPDFSG | S308 | 0.016 | 98.33 | 863 | 0.012719 | 0.016 | 0.0127 | 1 |
| Q13822 | ENPP2 | EDVTCDRtEFLSNYL | T370 | 0.165 | 98.06 | 863 | 0.112697 | 0.156 | 0.1491 | 1 |
| Q13822 | ENPP2 | TEERHLLyGRPAVLy | Y599 | 0.174 | 96.97 | 863 | 0.076331 | 0.174 | 0.0763 | 1 |
| Q96FE5 | LINGO1 | YLRFLNLsyNPIstI | S295 | 0.054 | 98.73 | 620 | 0.042925 | 0.054 | 0.0429 | 1 |
| O14979 | HNRNPDL | kMFIGGLsWDTskkD | S156 | 0.077 | 92.48 | 420 | 0.061208 | 0.077 | 0.0612 | 1 |
| Q86XP0 | PLA2G4D | APGMKFKtKTLTDTS | T61 | 0.048 | 98.02 | 818 | 0.032785 | 0.048 | 0.0328 | 1 |
| Q9Y5F2 | PCDHB11 | PEFEQAFyEVKIREN | Y247 | 0.198 | 94.71 | 797 | 0.08686 | 0.198 | 0.0869 | 1 |
| Q07617 | SPAG1 | ykDALSkysECLKIN | Y646 | 0.034 | 95.99 | 926 | 0.014915 | 0.034 | 0.0149 | 0.8 |
| Q07617 | SPAG1 | NkECAIytNRALCYL | T661 | 0.061 | 95.51 | 926 | 0.041664 | 0.061 | 0.0417 | 1 |
| P09601 | HMOX1 | GFKLVMAsLyHIyVA | S53 | 0 | 98.71 | 288 | 0 | 0 | 0 | 1 |
| P47897 | QARS1 | QLsAALEyVRSHPLD | Y93 | 0.174 | 94.59 | 775 | 0.076331 | 0.291 | 0.1277 | 1 |

| | | | | | | | | | | |
|---|---|---|---|---|---|---|---|---|---|---|
| Q9Y2X7 | GIT1 | RSLGRHIsIVkHLRH | S46 | 0.041 | 97.91 | 761 | 0.032592 | 0.086 | 0.0684 | 1 |
| P00480 | OTC | QAATPKGyEPDASVT | Y223 | 0.229 | 97.55 | 354 | 0.100459 | 0.32 | 0.1404 | 1 |
| P00480 | OTC | RLQAFQGyQVTMKTA | Y284 | 0.138 | 98.6 | 354 | 0.060539 | 0.138 | 0.0605 | 1 |
| Q8TDJ6 | DMXL2 | GTPSLPVsLSWVRDG | S1251 | 0.159 | 85.43 | 3036 | 0.126391 | 0.159 | 0.1264 | 1 |
| O96000 | NDUFB10 | AkAYQDryQDLGAyS | Y137 | 0.259 | 94.67 | 172 | 0.11362 | 0.259 | 0.1136 | 1 |
| P62714 | PPP2CB | DRGGWGIsPRGAGYT | S212 | 0.088 | 97.32 | 309 | 0.069952 | 0.05 | 0.0846 | 1 |
| P07437 | TUBB | AkFWEVIsDEHGIDP | S25 | 0.006 | 95.21 | 444 | 0.004769 | 0.006 | 0.0048 | 1 |
| P07437 | TUBB | LDrIsVyyNEAtGGk | Y51 | 0.059 | 96.75 | 444 | 0.025882 | 0.118 | 0.0518 | 0.9231 |
| P07437 | TUBB | NtFsVVPsPkVSDTV | S172 | 0.026 | 93.16 | 444 | 0.020668 | 0.026 | 0.0207 | 0.8333 |
| P07437 | TUBB | ALtVPELtQQVFDAk | T290 | 0.021 | 93.42 | 444 | 0.014343 | 0.042 | 0.0287 | 1 |
| P07437 | TUBB | PRHGRyLtVAAVFrG | T312 | 0.055 | 96.74 | 444 | 0.037566 | 0.11 | 0.0751 | 1 |
| P07437 | TUBB | AVtFIGNsTAIQELF | S371 | 0 | 97.21 | 444 | 0 | 0 | 0 | 1 |
| P07437 | TUBB | QELFkRIsEQFtAMF | S382 | 0.13 | 95.6 | 444 | 0.103339 | 0.13 | 0.1033 | 0.8889 |
| O94817 | ATG12 | LLkAVGDtPIMktKK | T65 | 0.065 | 88.41 | 140 | 0.044396 | 0.065 | 0.0444 | 1 |
| Q9ULV8 | CBLC | RLSVsPPsLRDLLPR | S43 | 0.091 | 86.61 | 474 | 0.072337 | 0.091 | 0.0723 | 1 |
| Q8NGR2 | OR1L6 | MLNPFIysLRNKDMK | S328 | 0.124 | 82.16 | 347 | 0.098569 | 0.174 | 0.1383 | 0.8571 |
| A2RRD8 | ZNF320 | DCGKAFNsPsHLIRH | S452 | 0.037 | 88.44 | 509 | 0.029412 | 0.037 | 0.0294 | 0.8 |
| O95155 | UBE4B | LEtVDPtyIFHPRCR | Y710 | 0.039 | 94.86 | 1302 | 0.017109 | 0.039 | 0.0171 | 1 |
| Q9UBV4 | WNT16 | GHLLKDKyENSIQIs | Y253 | 0.27 | 98.55 | 365 | 0.118445 | 0.27 | 0.1184 | 0.8333 |
| Q6J4K2 | SLC8B1 | TLAWALGyLGLYVFy | Y233 | 0.09 | 95.16 | 584 | 0.039482 | 0.09 | 0.0395 | 1 |
| Q6J4K2 | SLC8B1 | yLGLYVFyVVTVILC | Y240 | 0.03 | 96.23 | 584 | 0.013161 | 0.03 | 0.0132 | 1 |
| O15169 | AXIN1 | ARAIYRKyILDNNGI | Y148 | 0.214 | 96.58 | 862 | 0.093879 | 0.214 | 0.0939 | 1 |
| O15169 | AXIN1 | NGIVSRQtKPATKSF | T160 | 0.052 | 96.47 | 862 | 0.035517 | 0.052 | 0.0355 | 1 |
| P55196 | AFDN | EKENPKDyCIARVML | Y295 | 0.04 | 92.88 | 1824 | 0.017547 | 0.04 | 0.0175 | 0.8182 |
| Q9UL26 | RAB22A | DtGVGKssIVWRFVE | S20 | 0.161 | 98.63 | 194 | 0.127981 | 0.161 | 0.128 | 0.8889 |
| P30041 | PRDX6 | FHDFLGDsWGILFSH | S32 | 0.03 | 98.69 | 224 | 0.023847 | 0.052 | 0.0422 | 1 |
| P30041 | PRDX6 | VEDHLAWskDINAyN | S83 | 0.018 | 98.72 | 224 | 0.014308 | 0.018 | 0.0143 | 1 |
| P30041 | PRDX6 | DEkGMPVtARVVFVF | T130 | 0.008 | 97.43 | 224 | 0.005464 | 0.008 | 0.0055 | 1 |
| Q9GZL7 | WDR12 | FYTDNKkyAVDDVPF | Y15 | 0.078 | 86.8 | 423 | 0.034217 | 0.103 | 0.0452 | 1 |
| Q9GZL7 | WDR12 | VFNCISYsPLCkRLA | S306 | 0.001 | 95.23 | 423 | 0.000795 | 0.001 | 0.0008 | 1 |
| P62993 | GRB2 | EECDQNWykAELNGk | Y37 | 0.138 | 87.78 | 217 | 0.060539 | 0.138 | 0.0605 | 1 |

| | | | | | | | | | | |
|---|---|---|---|---|---|---|---|---|---|---|
| P62993 | GRB2 | GAFLIREsEsAPGDF | S88 | 0.139 | 88.94 | 217 | 0.110493 | 0.139 | 0.1105 | 1 |
| Q9ULT8 | HECTD1 | LPSIRKAsLALIRKM | S544 | 0 | 92.99 | 2610 | 0 | 0.044 | 0.0368 | 1 |
| P16452 | EPB42 | RVVTtFAsAQGTGGR | S294 | 0.021 | 93.97 | 691 | 0.016693 | 0.021 | 0.0167 | 1 |
| Q16864 | ATP6V1F | PAVLEIPskEHPYDA | S93 | 0.172 | 94.98 | 119 | 0.136725 | 0.172 | 0.1367 | 1 |
| Q9NZD2 | GLTP | TNPAkFRtLQNILEV | T69 | 0.114 | 98.65 | 209 | 0.077864 | 0.114 | 0.0779 | 1 |
| Q9NSI6 | BRWD1 | ISDMAVNyENTMIAA | Y238 | 0 | 84.73 | 2320 | 0 | 0 | 0 | 1 |
| Q9NSI6 | BRWD1 | GFGCSKPyEKIPDQM | Y551 | 0 | 87.19 | 2320 | 0 | 0 | 0 | 1 |
| P22079 | LPO | ILLATSHtLFLREHN | T384 | 0 | 98.94 | 712 | 0 | 0 | 0 | 0.9091 |
| P22079 | LPO | RGLLAKKsKLMKQNK | S521 | 0.008 | 98.78 | 712 | 0.006359 | 0.008 | 0.0064 | 1 |
| Q9P2E8 | MARCHF4 | CELCYYKyHVIAIst | Y212 | 0.093 | 89.43 | 410 | 0.040798 | 0.126 | 0.0553 | 1 |
| Q8TEV9 | SMCR8 | SRDFILIsEFSEQVG | S61 | 0 | 94.83 | 937 | 0 | 0 | 0 | 1 |
| P08648 | ITGA5 | GYCQGGFsAEFTKTG | S210 | 0.117 | 98.68 | 1049 | 0.093005 | 0.117 | 0.093 | 1 |
| P08648 | ITGA5 | DEFGRFGssLTPLGD | S394 | 0.173 | 98.38 | 1049 | 0.13752 | 0.173 | 0.1375 | 1 |
| P08648 | ITGA5 | EFGRFGssLTPLGDL | S395 | 0.082 | 98.41 | 1049 | 0.065183 | 0.082 | 0.0652 | 1 |
| Q6ZR08 | DNAH12 | TTLGALVtIDVHARD | T1166 | 0.082 | 91.375 | 3092 | 0.056007 | 0.082 | 0.056 | 1 |
| Q6ZNB7 | AGMO | QGFRMLFyTMKPSET | Y21 | 0.205 | 96.23 | 445 | 0.089931 | 0.205 | 0.0899 | 0.8667 |
| Q6ZNB7 | AGMO | MLLELVVsWILKGKP | S58 | 0.036 | 94.94 | 445 | 0.028617 | 0.036 | 0.0286 | 1 |
| Q86WV6 | STING1 | RVysNSIyELLENGQ | Y245 | 0.237 | 94.9 | 379 | 0.103968 | 0.237 | 0.104 | 1 |
| P01011 | SERPINA3 | IKDLDsQtMMVLVNY | T203 | 0.027 | 95.51 | 423 | 0.018441 | 0.027 | 0.0184 | 1 |
| P01011 | SERPINA3 | LKYTGNAsALFILPD | S273 | 0.021 | 94.7 | 423 | 0.016693 | 0.021 | 0.0167 | 1 |
| P01011 | SERPINA3 | EAMLLPEtLKRWRDs | T295 | 0.11 | 95.83 | 423 | 0.075131 | 0.11 | 0.0751 | 0.8889 |
| O95620 | DUS4L | FRtLVRKysCDLCYT | Y49 | 0.199 | 98.68 | 317 | 0.087298 | 0.199 | 0.0873 | 1 |
| Q53T59 | HS1BP3 | RRLQNAHtGLDLTVP | T18 | 0.162 | 95.1 | 392 | 0.110648 | 0.162 | 0.1106 | 1 |
| Q9Y5E2 | PCDHB7 | QINPTSGsLHLKAQL | S300 | 0.083 | 96.33 | 793 | 0.065978 | 0.083 | 0.066 | 1 |
| Q01082 | SPTBN1 | RTQILAAsyELHkFy | S1804 | 0.11 | 88.03 | 2364 | 0.087441 | 0.085 | 0.1157 | 0.875 |
| Q9NX74 | DUS2 | MGCPkQYsTkGGMGA | S121 | 0 | 79.21 | 493 | 0 | 0 | 0 | 1 |
| Q9NX74 | DUS2 | LVkRIERtGIAAIAV | T175 | 0.146 | 98.67 | 493 | 0.09972 | 0.146 | 0.0997 | 1 |
| Q9NX74 | DUS2 | IEDFRQAtAASSVMV | T234 | 0 | 98.62 | 493 | 0 | 0 | 0 | 1 |
| Q9ULX3 | NOB1 | tEIRDkAtRRRLAVL | T42 | 0.014 | 92.37 | 412 | 0.009562 | 0.014 | 0.0096 | 1 |
| Q14155 | ARHGEF7 | IYQGKLPtTGMTITK | T528 | 0.071 | 86.53 | 803 | 0.048494 | 0.071 | 0.0485 | 1 |
| Q8NDV3 | SMC1B | SGPERQKtVALDGtL | T638 | 0.004 | 93.02 | 1235 | 0.002732 | 0.004 | 0.0027 | 1 |

| | | | | | | | | | | |
|---|---|---|---|---|---|---|---|---|---|---|
| Q8NDV3 | SMC1B | KtVALDGtLFLKSGV | T644 | 0.154 | 90.33 | 1235 | 0.105184 | 0.154 | 0.1052 | 1 |
| Q9BT23 | LIMD2 | AALHGEFyCKPHFQQ | Y87 | 0.094 | 89.79 | 127 | 0.041236 | 0.242 | 0.1062 | 1 |
| Q9UN76 | SLC6A14 | NWSKKSDyLLSMIGy | Y45 | 0.029 | 98.3 | 642 | 0.012722 | 0.029 | 0.0127 | 1 |
| Q9UN76 | SLC6A14 | yLLSMIGyAVGLGNV | Y52 | 0.103 | 97.9 | 642 | 0.045185 | 0.103 | 0.0452 | 1 |
| Q9Y490 | TLN1 | rNGDtMEyRkKQRPL | Y81 | 0.108 | 86.2 | 2541 | 0.047378 | 0.108 | 0.0474 | 1 |
| Q9Y490 | TLN1 | ItNHDEysLVRELME | S128 | 0 | 88.69 | 2541 | 0 | 0 | 0 | 1 |
| Q9NXJ5 | PGPEP1 | RyLCDFTyYTSLyQs | Y153 | 0 | 98.61 | 209 | 0 | 0 | 0 | 0.8333 |
| P20702 | ITGAX | LGsRDLQssVTLDLA | S673 | 0.001 | 94.25 | 1163 | 0.000795 | 0.001 | 0.0008 | 1 |
| Q16795 | NDUFA9 | NHLGrMGsQVIIPYR | S78 | 0.003 | 96.47 | 377 | 0.002385 | 0.003 | 0.0024 | 0.8889 |
| O60507 | TPST1 | KANKTFAyHKDMPLI | Y65 | 0.038 | 95.55 | 370 | 0.01667 | 0.076 | 0.0333 | 1 |
| O60507 | TPST1 | MVRDGRAsVHSMISR | S189 | 0 | 98.68 | 370 | 0 | 0 | 0 | 0.9 |
| Q9UG01 | IFT172 | GQTDNIIyVyKIGED | Y91 | 0.111 | 96.31 | 1749 | 0.048694 | 0.222 | 0.0974 | 1 |
| Q9UG01 | IFT172 | TDNIIyVyKIGEDWG | Y93 | 0.075 | 95.69 | 1749 | 0.032901 | 0.135 | 0.0592 | 1 |
| Q9UG01 | IFT172 | CDRKIVAyGKEGHML | Y222 | 0.134 | 95.01 | 1749 | 0.058784 | 0.262 | 0.1135 | 1 |
| Q9UG01 | IFT172 | GQsVVLGsyDRLRVF | S259 | 0.015 | 91.92 | 1749 | 0.011924 | 0.015 | 0.0119 | 1 |
| Q8NEV1 | CSNK2A3 | REAMEHPyFYTVVkD | Y323 | 0.001 | 98.69 | 391 | 0.000439 | 0.001 | 0.0004 | 1 |
| P43121 | MCAM | SGNHMKEsREVTVPV | S238 | 0.159 | 98.17 | 646 | 0.126391 | 0.159 | 0.1264 | 1 |
| Q8NEZ5 | FBXO22 | ILPHTVLyMADSETF | Y104 | 0 | 97.45 | 403 | 0 | 0 | 0 | 1 |
| Q6P3X3 | TTC27 | VERAMRQtQALADQF | T410 | 0 | 96.72 | 843 | 0 | 0 | 0 | 1 |
| Q6P3X3 | TTC27 | DGEIWRLyAHVYGNG | Y715 | 0.278 | 96.17 | 843 | 0.121955 | 0.278 | 0.122 | 1 |
| Q9H7B2 | RPF2 | DFDVTEDyRRLksLL | Y156 | 0.262 | 94.91 | 306 | 0.114936 | 0.262 | 0.1149 | 1 |
| Q9H7B2 | RPF2 | ELEEMGPsLDLVLRR | S220 | 0.063 | 96.29 | 306 | 0.05008 | 0.063 | 0.0501 | 1 |
| O95602 | POLR1A | PLMASGksLPCFEPy | S934 | 0.079 | 90.24 | 1720 | 0.062798 | 0.114 | 0.1063 | 1 |
| O95602 | POLR1A | sLPCFEPyEFTPRAG | Y941 | 0.143 | 88.49 | 1720 | 0.062732 | 0.143 | 0.0627 | 1 |
| P51788 | CLCN2 | TITALFKtRFRLDFP | T308 | 0.106 | 84.41 | 898 | 0.072399 | 0.106 | 0.0724 | 1 |
| P05166 | PCCB | NMVDIIHsVVDEREF | S322 | 0.07 | 97.69 | 539 | 0.055644 | 0.07 | 0.0556 | 1 |
| P05166 | PCCB | VEAAQAEyIEkFANP | Y486 | 0.268 | 90.68 | 539 | 0.117568 | 0.268 | 0.1176 | 1 |
| Q15233 | NONO | RFACHsAsLtVRNLP | S149 | 0.143 | 97.3 | 471 | 0.113673 | 0.143 | 0.1137 | 1 |
| Q15233 | NONO | ACHsAsLtVRNLPQY | T151 | 0.206 | 98.27 | 471 | 0.140701 | 0.206 | 0.1407 | 1 |
| Q86UY8 | NT5DC3 | KLQKGQIyKQGNLyE | Y365 | 0.014 | 97.42 | 548 | 0.006142 | 0.014 | 0.0061 | 1 |
| Q9UBB5 | MBD2 | KSDVYyFsPsGKKFR | S181 | 0.056 | 90.62 | 411 | 0.044515 | 0.056 | 0.0445 | 0.8333 |

| | | | | | | | | | | |
|---|---|---|---|---|---|---|---|---|---|---|
| O00481 | BTN3A1 | GLTDGNKyRTLTEPR | Y441 | 0.226 | 97.81 | 513 | 0.099143 | 0.226 | 0.0991 | 0.8182 |
| O75436 | VPS26A | RDVNKKFsVRYFLNL | S269 | 0.112 | 97.52 | 327 | 0.08903 | 0.112 | 0.089 | 1 |
| Q9UIR0 | BTNL2 | tEMQMEEyRGWVEWI | Y91 | 0.147 | 83.1 | 455 | 0.064487 | 0.147 | 0.0645 | 1 |
| P58499 | FAM3B | TYAYRLLsGGGRSKY | S80 | 0.008 | 98.53 | 235 | 0.006359 | 0.008 | 0.0064 | 1 |
| Q8N1F8 | STK11IP | AATAQYLsPRARDAA | S306 | 0.046 | 88.61 | 1088 | 0.036566 | 0.046 | 0.0366 | 1 |
| A5D8W1 | CFAP69 | EEVIQQLsNLECLLA | S281 | 0.171 | 96.33 | 941 | 0.13593 | 0.171 | 0.1359 | 1 |
| O60493 | SNX3 | VGrGrFttyEIRVkt | T48 | 0.135 | 98.56 | 162 | 0.092207 | 0.135 | 0.0922 | 1 |
| O60493 | SNX3 | GrGrFttyEIRVktN | Y49 | 0.031 | 98.8 | 162 | 0.013599 | 0.031 | 0.0136 | 1 |
| O60493 | SNX3 | tyEIRVktNLPIFkL | T55 | 0.046 | 96.97 | 162 | 0.031419 | 0.061 | 0.0492 | 1 |
| O60493 | SNX3 | DEIIDksytPSkIRH | Y154 | 0.183 | 92.49 | 162 | 0.080279 | 0.183 | 0.0803 | 1 |
| Q15181 | PPA1 | EERAAPFsLEyRVFL | S14 | 0.186 | 95.63 | 289 | 0.147854 | 0.186 | 0.1479 | 0.8333 |
| Q15181 | PPA1 | MNTtLsEsPFkCDPD | S250 | 0.113 | 95.56 | 289 | 0.089825 | 0.113 | 0.0898 | 0.8333 |
| O95294 | RASAL1 | ___MAKssSLNVRVV | S5 | 0.051 | 88.65 | 804 | 0.040541 | 0.096 | 0.077 | 1 |
| O95294 | RASAL1 | EVVARTAtVWRSLGP | T44 | 0.006 | 94.42 | 804 | 0.004098 | 0.208 | 0.1364 | 1 |
| Q96DH6 | MSI2 | VTFADPAsVDkVLGQ | S74 | 0.059 | 88.79 | 328 | 0.0469 | 0.059 | 0.0469 | 1 |
| P21333 | FLNA | dGtQTVNyVPsREGP | Y1511 | 0.075 | 89.79 | 2647 | 0.032901 | 0.075 | 0.0329 | 1 |
| P21333 | FLNA | RYtILIkyGGDEIPF | Y1622 | 0.1 | 85.37 | 2647 | 0.043869 | 0.1 | 0.0439 | 1 |
| P21333 | FLNA | DEIPFsPyRVRAVPt | Y1632 | 0.207 | 86.68 | 2647 | 0.090808 | 0.207 | 0.0908 | 1 |
| P21333 | FLNA | TAPQPGKyVICVRFG | Y1720 | 0 | 78.36 | 2647 | 0 | 0 | 0 | 1 |
| Q99460 | PSMD1 | GtARQDVyDLLKTNL | Y494 | 0.027 | 92.52 | 953 | 0.011845 | 0.027 | 0.0118 | 1 |
| Q99460 | PSMD1 | VNQFRQLySkVINDk | Y713 | 0.033 | 90.79 | 953 | 0.014477 | 0.033 | 0.0145 | 1 |
| Q99460 | PSMD1 | TMPETCRyQPFkPLS | Y900 | 0.136 | 80.97 | 953 | 0.059661 | 0.136 | 0.0597 | 1 |
| O43462 | MBTPS2 | ITEVAEDsPAIGPRG | S265 | 0.106 | 87.15 | 519 | 0.084261 | 0.106 | 0.0843 | 1 |
| P02751 | FN1 | ALKNNQKsEPLIGRK | S2073 | 0.114 | 87.63 | 2477 | 0.09062 | 0.121 | 0.1343 | 1 |
| Q5SY16 | NOL9 | CkNNYENyIDIVKyV | Y386 | 0 | 96.82 | 702 | 0 | 0 | 0 | 1 |
| Q5SY16 | NOL9 | VKyVFsAyKRESPLI | Y397 | 0.242 | 95.69 | 702 | 0.106162 | 0.242 | 0.1062 | 1 |
| Q05BV3 | EML5 | ADGKRLAsVGIDDSH | S789 | 0 | 95.8 | 1969 | 0 | 0 | 0 | 1 |
| Q6NZ67 | MZT2B | stEEMELyELAQAAG | Y41 | 0.309 | 91.77 | 158 | 0.135554 | 0.309 | 0.1356 | 1 |
| Q9H5I5 | PIEZO2 | ARLFGIKsVIQtDCS | S312 | 0.105 | 85.41 | 2752 | 0.083466 | 0.085 | 0.1121 | 1 |
| O60353 | FZD6 | SIFCLCAtLFTFLTF | T213 | 0 | 95.19 | 706 | 0 | 0.156 | 0.1066 | 0.8 |
| O95602 | POLR1A | GSHDELRsPSACLVV | S1695 | 0.016 | 91.1 | 1720 | 0.012719 | 0.019 | 0.066 | 1 |

| | | | | | | | | | | |
|---|---|---|---|---|---|---|---|---|---|---|
| Q14232 | EIF2B1 | SAVAAIRtLLEFLkR | T29 | 0 | 94.4 | 305 | 0 | 0 | 0 | 1 |
| P50443 | SLC26A2 | KLHENYNssIAGHIP | S358 | 0.011 | 89.74 | 739 | 0.008744 | 0.011 | 0.0087 | 1 |
| P50443 | SLC26A2 | FRFVAPLyyINKECF | Y588 | 0.287 | 94.63 | 739 | 0.125903 | 0.287 | 0.1259 | 1 |
| P50443 | SLC26A2 | ECFkSALykQtVNPI | Y600 | 0.187 | 94.7 | 739 | 0.082034 | 0.187 | 0.082 | 1 |
| P50443 | SLC26A2 | kSALykQtVNPILIK | T603 | 0.128 | 90.05 | 739 | 0.087426 | 0.128 | 0.0874 | 1 |
| Q9HAT2 | SIAE | SVKAHsDtWMVVLDP | T81 | 0.127 | 97.63 | 523 | 0.086743 | 0.127 | 0.0867 | 0.8571 |
| Q9HAT2 | SIAE | ATRELsNtAAYQSVR | T146 | 0.154 | 95.97 | 523 | 0.105184 | 0.181 | 0.1236 | 1 |
| P07237 | P4HB | CKALAPEyAkAAGKL | Y63 | 0.035 | 98.28 | 508 | 0.015354 | 0.035 | 0.0154 | 1 |
| P07237 | P4HB | EsDLAQQyGVrGyPT | Y94 | 0.26 | 95.41 | 508 | 0.114058 | 0.26 | 0.1141 | 1 |
| P07237 | P4HB | ILFIFIDsDHTDNQR | S293 | 0.019 | 95.83 | 508 | 0.015103 | 0.019 | 0.0151 | 1 |
| P07237 | P4HB | ADrTVIDyNGERtLD | Y457 | 0.178 | 94.65 | 508 | 0.078086 | 0.178 | 0.0781 | 1 |
| P15121 | AKR1B1 | MPILGLGtWksPPGQ | T20 | 0.007 | 98.73 | 316 | 0.004781 | 0.007 | 0.0048 | 1 |
| P15121 | AKR1B1 | LGLGtWksPPGQVTE | S23 | 0.051 | 97.92 | 316 | 0.040541 | 0.1 | 0.0797 | 1 |
| P15121 | AKR1B1 | REELFIVsKLWCTyH | S77 | 0 | 98.92 | 316 | 0 | 0 | 0 | 0.8889 |
| P15121 | AKR1B1 | NQIECHPyLtQEkLI | Y190 | 0.109 | 98.85 | 316 | 0.047817 | 0.109 | 0.0478 | 1 |
| P15121 | AKR1B1 | IECHPyLtQEkLIQy | T192 | 0.002 | 98.79 | 316 | 0.001366 | 0.002 | 0.0014 | 1 |
| Q9Y6X3 | MAU2 | FTTALRLtNHQELWA | T409 | 0.173 | 95.94 | 613 | 0.118161 | 0.173 | 0.1182 | 1 |
| Q9Y6X3 | MAU2 | LRETLKMsNAEDLNR | S489 | 0.002 | 96.5 | 613 | 0.00159 | 0.002 | 0.0016 | 1 |
| O15394 | NCAM2 | NIEDAGIyRCQATDA | Y91 | 0.097 | 94.88 | 837 | 0.042553 | 0.097 | 0.0426 | 1 |
| Q9Y2D2 | SLC35A3 | LGVyQWLsLVILMTG | S143 | 0 | 96.46 | 325 | 0 | 0 | 0 | 0.9091 |
| P24385 | CCND1 | ATCMFVAskMkETIP | S111 | 0 | 98.43 | 295 | 0 | 0 | 0 | 1 |
| Q8N653 | LZTR1 | GtPPAPRyHHSAVVY | Y119 | 0.33 | 95.61 | 840 | 0.144766 | 0.33 | 0.1448 | 0.9091 |
| Q8WVM7 | STAG1 | FDEDsGDyPLtMPGP | Y162 | 0.079 | 95.16 | 1258 | 0.034656 | 0.079 | 0.0347 | 1 |
| Q8WVM7 | STAG1 | IDDRNKLtEHFIItL | T568 | 0 | 96.92 | 1258 | 0 | 0 | 0 | 1 |
| Q8WVM7 | STAG1 | LtEHFIItLPMLLSK | T574 | 0.17 | 97.13 | 1258 | 0.116112 | 0.17 | 0.1161 | 1 |
| Q8WVM7 | STAG1 | DIyNVLStLKRLTSF | T694 | 0.04 | 97.12 | 1258 | 0.027321 | 0.04 | 0.0273 | 1 |
| Q9BZE4 | GTPBP4 | LskTQRktPTVIHkH | T30 | 0.086 | 95.39 | 634 | 0.058739 | 0.086 | 0.0587 | 1 |
| Q15036 | SNX17 | VLHCRVRysQLLGLH | Y37 | 0.288 | 95.21 | 470 | 0.126341 | 0.288 | 0.1263 | 1 |
| Q15036 | SNX17 | RREQLEkyMQAVRQD | Y82 | 0.029 | 96.8 | 470 | 0.012722 | 0.029 | 0.0127 | 1 |
| P33992 | MCM5 | PNVyEVIsksIAPsI | S336 | 0.065 | 87.91 | 734 | 0.051669 | 0.065 | 0.0517 | 1 |
| Q9Y5Z0 | BACE2 | LNLDCREyNADkAIV | Y295 | 0.018 | 98.08 | 518 | 0.007896 | 0.018 | 0.0079 | 1 |

| | | | | | | | | | | |
|---|---|---|---|---|---|---|---|---|---|---|
| Q00839 | HNRNPU | WAGGRASyGVskGkV | Y327 | 0.145 | 97.51 | 825 | 0.063609 | 0.145 | 0.0636 | 1 |
| Q00839 | HNRNPU | AAENPGkyNILGTNt | Y525 | 0.305 | 92.77 | 825 | 0.133799 | 0.305 | 0.1338 | 1 |
| Q00839 | HNRNPU | VCPkDEDykQRTQkK | Y613 | 0.151 | 92.63 | 825 | 0.066242 | 0.151 | 0.0662 | 1 |
| Q00839 | HNRNPU | AQkLLEQykEEskkA | Y669 | 0.033 | 93.12 | 825 | 0.014477 | 0.033 | 0.0145 | 1 |
| Q00839 | HNRNPU | LEQykEEskkALPPE | S673 | 0.133 | 89.6 | 825 | 0.105724 | 0.133 | 0.1057 | 1 |
| Q15262 | PTPRK | RyGNIIAyDHSRVIL | Y922 | 0.007 | 95.54 | 1439 | 0.003071 | 0.007 | 0.0031 | 1 |
| Q15262 | PTPRK | EDDPSSDyINANyID | Y940 | 0.003 | 97.21 | 1439 | 0.001316 | 0.003 | 0.0013 | 1 |
| P13747 | HLA-E | RGYEQFAyDGkDYLT | Y139 | 0.128 | 98.32 | 358 | 0.056152 | 0.128 | 0.0562 | 1 |
| P32298 | GRK4 | RILEKVQsRFVVsLA | S244 | 0.117 | 95.14 | 578 | 0.093005 | 0.117 | 0.093 | 1 |
| Q92698 | RAD54L | LkNSENQtyQALDsL | T308 | 0.18 | 88.09 | 747 | 0.122942 | 0.18 | 0.1229 | 1 |
| Q92698 | RAD54L | kNSENQtyQALDsLN | Y309 | 0.147 | 90.05 | 747 | 0.064487 | 0.147 | 0.0645 | 1 |
| Q92698 | RAD54L | LFILDEAsLSDTHDR | S671 | 0.13 | 86.7 | 747 | 0.103339 | 0.13 | 0.1033 | 1 |
| Q68CJ6 | NUGGC | RDVALVVtKMDKLHL | T354 | 0.079 | 93.4 | 796 | 0.053958 | 0.076 | 0.0719 | 1 |
| P98082 | DAB2 | QRIWVNIsLsGIKII | S98 | 0.013 | 97.9 | 770 | 0.010334 | 0.013 | 0.0103 | 0.875 |
| O95490 | ADGRL2 | ESETGKNsIDyIYNT | S339 | 0.092 | 88.37 | 1459 | 0.073132 | 0.092 | 0.0731 | 1 |
| Q9UQL6 | HDAC5 | FAAGLGRsLREAQAG | S1071 | 0.036 | 95.77 | 1122 | 0.028617 | 0.036 | 0.0286 | 0.8889 |
| P98196 | ATP11A | QSKSQkRsAVEKsMN | S287 | 0.026 | 81.99 | 1134 | 0.020668 | 0.026 | 0.0207 | 1 |
| P98196 | ATP11A | CkLFRRNtQLLELtt | T708 | 0.124 | 83.93 | 1134 | 0.084694 | 0.142 | 0.097 | 1 |
| P98196 | ATP11A | NtQLLELttkRIEEQ | T714 | 0.152 | 82.54 | 1134 | 0.103818 | 0.152 | 0.1038 | 1 |
| P55072 | VCP | EAINEDNsVVsLSQP | S37 | 0.092 | 90.77 | 806 | 0.073132 | 0.092 | 0.0731 | 1 |
| P55072 | VCP | VEFkVVEtDPSPyCI | T168 | 0.064 | 94.55 | 806 | 0.043713 | 0.064 | 0.0437 | 1 |
| P55072 | VCP | yCIVAPDtVIHCEGE | T180 | 0.139 | 92.67 | 806 | 0.094939 | 0.139 | 0.0949 | 1 |
| P55072 | VCP | MskLAGEsEsNLRkA | S282 | 0.008 | 87.84 | 806 | 0.006359 | 0.008 | 0.0064 | 1 |
| P55072 | VCP | IAPkREktHGEVERR | T316 | 0.063 | 74.84 | 806 | 0.04303 | 0.063 | 0.043 | 1 |
| P55072 | VCP | HVIVMAAtNRPNsID | T347 | 0.014 | 92.39 | 806 | 0.009562 | 0.014 | 0.0096 | 1 |
| P55072 | VCP | LEILQIHtkNMkLAD | T385 | 0.111 | 91.94 | 806 | 0.075815 | 0.111 | 0.0758 | 1 |
| P55072 | VCP | LEQVANEtHGHVGAD | T403 | 0 | 85.09 | 806 | 0 | 0 | 0 | 1 |
| Q6IBW4 | NCAPH2 | RQGEVLAsRKDFRMN | S175 | 0.066 | 87.81 | 605 | 0.052464 | 0.066 | 0.0525 | 1 |
| P98164 | LRP2 | YFDKIEHstFDGLDR | S897 | 0.001 | 94.49 | 4655 | 0.000795 | 0.001 | 0.0008 | 1 |
| P61978 | HNRNPK | FQECCPHsTDrVVLI | S188 | 0.165 | 84.21 | 463 | 0.131161 | 0.165 | 0.1312 | 1 |
| P61978 | HNRNPK | IkGRAQPyDPNFyDE | Y225 | 0.013 | 81.05 | 463 | 0.005703 | 0.327 | 0.1434 | 1 |

| | | | | | | | | | | |
|---|---|---|---|---|---|---|---|---|---|---|
| P61978 | HNRNPK | FyDETyDyGGFTMMF | Y236 | 0.02 | 77.03 | 463 | 0.008774 | 0.146 | 0.0601 | 1 |
| P60709 | ACTB | APrAVFPsIVGRPRH | S33 | 0.018 | 97.47 | 375 | 0.014308 | 0.018 | 0.0143 | 1 |
| P60709 | ACTB | GMGQkDsyVGDEAQs | Y53 | 0.051 | 95.87 | 375 | 0.022373 | 0.345 | 0.1421 | 1 |
| P60709 | ACTB | MEkIWHHtFyNELRV | T89 | 0.001 | 98.6 | 375 | 0.000683 | 0.001 | 0.0007 | 0.9 |
| P60709 | ACTB | TTGIVMdsGdGVtHt | S155 | 0.026 | 98.51 | 375 | 0.020668 | 0.026 | 0.0207 | 0.8333 |
| P60709 | ACTB | sGdGVtHtVPIyEGy | T162 | 0.016 | 98.53 | 375 | 0.010928 | 0.016 | 0.0109 | 1 |
| P60709 | ACTB | DLAGRDLtDyLMkIL | T186 | 0.013 | 98.23 | 375 | 0.008879 | 0.013 | 0.0089 | 0.9 |
| P60709 | ACTB | AGRDLtDyLMkILtE | Y188 | 0.15 | 98.29 | 375 | 0.065803 | 0.3 | 0.1316 | 1 |
| P60709 | ACTB | VDIRkDLyANtVLsG | Y294 | 0.004 | 98.21 | 375 | 0.001755 | 0.004 | 0.0018 | 1 |
| P60709 | ACTB | RkDLyANtVLsGGtt | T297 | 0 | 98.63 | 375 | 0 | 0 | 0 | 1 |
| P60709 | ACTB | LyANtVLsGGttMyP | S300 | 0.005 | 98.44 | 375 | 0.003975 | 0.005 | 0.004 | 1 |
| P60709 | ACTB | MWISkQEyDEsGPsI | Y362 | 0.121 | 98.1 | 375 | 0.053081 | 0.241 | 0.1055 | 1 |
| Q09161 | NCBP1 | NNYNEAVyLVRFLSD | Y126 | 0.221 | 98.6 | 790 | 0.09695 | 0.221 | 0.0969 | 1 |
| Q09161 | NCBP1 | WYVYAFLsSLPWVGK | S175 | 0 | 98.73 | 790 | 0 | 0 | 0 | 1 |
| Q09161 | NCBP1 | PQVLAQAtEMLYMRL | T395 | 0.036 | 98.7 | 790 | 0.024588 | 0.036 | 0.0246 | 1 |
| O95757 | HSPA4L | ARsGGIEtIANEYSD | T25 | 0.066 | 97.17 | 839 | 0.045079 | 0.066 | 0.0451 | 1 |
| O95757 | HSPA4L | FkKLHGRsFDDPIVQ | S74 | 0.12 | 95.01 | 839 | 0.09539 | 0.164 | 0.1304 | 1 |
| O95757 | HSPA4L | ALQCAILsPAFkVRE | S384 | 0.076 | 89.27 | 839 | 0.060414 | 0.076 | 0.0604 | 1 |
| O95757 | HSPA4L | GQDLLNsyIENEGKM | Y600 | 0.193 | 97.59 | 839 | 0.084666 | 0.193 | 0.0847 | 1 |
| Q9BXF9 | TEKT3 | QFNKVNLsFTNRIAE | S346 | 0.155 | 98.38 | 490 | 0.123212 | 0.155 | 0.1232 | 1 |
| Q9BXF9 | TEKT3 | DLAVKANsLyIDQEK | S465 | 0.025 | 98.01 | 490 | 0.019873 | 0.025 | 0.0199 | 1 |
| Q9H3G5 | CPVL | DWkGSQEyKKAEKKV | Y411 | 0.025 | 97.85 | 476 | 0.010967 | 0.025 | 0.011 | 0.9167 |
| Q8IXI1 | RHOT2 | tPCLFVSskADLPEG | S524 | 0.045 | 93.5 | 618 | 0.035771 | 0.132 | 0.1049 | 1 |
| Q8IVG5 | SAMD9L | GRNIERVsFYLGFsI | S1566 | 0.125 | 89.71 | 1584 | 0.099364 | 0.177 | 0.1407 | 0.875 |
| Q01432 | AMPD3 | KYVtGFDsVDDEsKH | S525 | 0 | 97.5 | 767 | 0 | 0 | 0 | 0.9 |
| Q01432 | AMPD3 | EGNDIRktNVAQIRM | T732 | 0.043 | 96.61 | 767 | 0.02937 | 0.043 | 0.0294 | 1 |
| Q3SXY7 | LRIT3 | sSKDAGDyKCKAKNL | Y326 | 0.023 | 94.37 | 679 | 0.01009 | 0.023 | 0.0101 | 1 |
| Q8NFA0 | USP32 | DSCLRAFtSEEELGE | T1243 | 0.189 | 93.59 | 1604 | 0.12909 | 0.189 | 0.1291 | 1 |
| Q8NFA0 | USP32 | KNPNCkWyCyNDssC | Y1539 | 0.112 | 96.91 | 1604 | 0.049133 | 0.259 | 0.1136 | 1 |
| Q8NFA0 | USP32 | PNCkWyCyNDssCKE | Y1541 | 0.058 | 96.89 | 1604 | 0.025444 | 0.058 | 0.0254 | 1 |
| Q9HBW1 | LRRC4 | YAFNRVPsLMRLDLG | S172 | 0.115 | 96.86 | 653 | 0.091415 | 0.115 | 0.0914 | 1 |

| | | | | | | | | | | |
|---|---|---|---|---|---|---|---|---|---|---|
| Q05397 | PTK2 | ALRtLLAtVDEtIPL | T967 | 0.139 | 95.23 | 1052 | 0.094939 | 0.139 | 0.0949 | 0.8571 |
| Q9Y6E2 | BZW2 | EKDANsVtSSLRKAN | T204 | 0.082 | 94.28 | 419 | 0.056007 | 0.082 | 0.056 | 1 |
| Q9Y6E2 | BZW2 | DAGLkELsDFLRVQQ | S243 | 0.026 | 92.63 | 419 | 0.020668 | 0.026 | 0.0207 | 1 |
| O43300 | LRRTM2 | SLEFLDLsTNRLRSL | S189 | 0.026 | 98.75 | 516 | 0.020668 | 0.026 | 0.0207 | 1 |
| Q8IUQ4 | SIAH1 | SVLFPCKyASSGCEI | Y100 | 0.327 | 92.96 | 282 | 0.14345 | 0.327 | 0.1434 | 0.9 |
| Q8IUQ4 | SIAH1 | ELCEFRPySCPCPGA | Y126 | 0.161 | 92.17 | 282 | 0.070628 | 0.161 | 0.0706 | 0.9167 |
| Q9NY28 | GALNT8 | DLFRKFGyNAYLSNQ | Y149 | 0.001 | 96.78 | 637 | 0.000439 | 0.001 | 0.0004 | 1 |
| Q9NY28 | GALNT8 | SIIQRAItsIINRtP | T202 | 0.162 | 98.3 | 637 | 0.110648 | 0.162 | 0.1106 | 1 |
| Q9NY28 | GALNT8 | IIQRAItsIINRtPs | S203 | 0.036 | 98.6 | 637 | 0.028617 | 0.036 | 0.0286 | 1 |
| Q9NY28 | GALNT8 | ItsIINRtPsRLLKE | T208 | 0.099 | 98.42 | 637 | 0.067618 | 0.099 | 0.0676 | 1 |
| P22735 | TGM1 | HEDWRQEyVLNESGR | Y267 | 0.041 | 98.65 | 817 | 0.017986 | 0.041 | 0.018 | 0.8571 |
| P22735 | TGM1 | SVEILLSyLRTGySV | Y365 | 0.006 | 98.52 | 817 | 0.002632 | 0.006 | 0.0026 | 1 |
| P25787 | PSMA2 | sFsLTTFsPsGKLVQ | S14 | 0.183 | 89.16 | 234 | 0.145469 | 0.183 | 0.1455 | 1 |
| Q9UNQ0 | ABCG2 | VRENLQFsAALRLAT | S143 | 0 | 95.62 | 655 | 0 | 0.118 | 0.0938 | 1 |
| Q9UNQ0 | ABCG2 | sGGERKRtsIGMELI | T194 | 0.003 | 94.14 | 655 | 0.002049 | 0.003 | 0.002 | 1 |
| Q9UNQ0 | ABCG2 | GGERKRtsIGMELIT | S195 | 0.041 | 93.56 | 655 | 0.032592 | 0.081 | 0.0644 | 1 |
| Q12891 | HYAL2 | AQQFMLEtLRYVKAV | T190 | 0 | 98.64 | 473 | 0 | 0 | 0 | 1 |
| Q12891 | HYAL2 | PDCYNHDyVQNWESY | Y216 | 0.208 | 95.62 | 473 | 0.091247 | 0.208 | 0.0912 | 1 |
| Q12891 | HYAL2 | ANHALPVyVFTRPty | Y291 | 0.086 | 98.75 | 473 | 0.037727 | 0.086 | 0.0377 | 1 |
| Q12891 | HYAL2 | VyVFTRPtySRRLTG | T297 | 0.026 | 98.42 | 473 | 0.017758 | 0.026 | 0.0178 | 1 |
| Q12891 | HYAL2 | yVFTRPtySRRLTGL | Y298 | 0.143 | 98.23 | 473 | 0.062732 | 0.143 | 0.0627 | 1 |
| P12111 | COL6A3 | LFTSsAGyRAAEGIP | Y543 | 0.248 | 89.26 | 3177 | 0.108794 | 0.248 | 0.1088 | 1 |
| Q99678 | GPR20 | AIVRPEGsRRCRQPA | S158 | 0.065 | 88.45 | 358 | 0.051669 | 0.065 | 0.0517 | 1 |
| P82987 | ADAMTSL3 | SLENEGTyVCIATNA | Y1365 | 0.028 | 88.52 | 1691 | 0.012283 | 0.094 | 0.0412 | 1 |
| P54098 | POLG | QLLSGERyKEDPWLW | Y479 | 0.229 | 89 | 1239 | 0.100459 | 0.304 | 0.1334 | 1 |
| Q6W5P4 | NPSR1 | NSVVLFStWRRKKKS | T74 | 0.084 | 92.16 | 371 | 0.057373 | 0.185 | 0.1264 | 1 |
| Q13976 | PRKG1 | GLIKHTEyMEFLKsV | Y212 | 0.167 | 89.14 | 671 | 0.07326 | 0.167 | 0.0733 | 1 |
| Q9Y3D7 | PAM16 | RGRAGHRsAAASNLs | S45 | 0.153 | 90.94 | 125 | 0.121622 | 0.153 | 0.1216 | 1 |
| Q9P1W9 | PIM2 | REAFEAEyRLGPLLG | Y32 | 0.127 | 98.38 | 311 | 0.055713 | 0.254 | 0.1114 | 1 |
| Q9P1W9 | PIM2 | FDGTRVYsPPEWISR | S204 | 0 | 98.35 | 311 | 0 | 0 | 0 | 1 |
| P10828 | THRB | FLLAFEHyINYRKHH | Y406 | 0.185 | 98.17 | 461 | 0.081157 | 0.185 | 0.0812 | 1 |

| | | | | | | | | | | |
|---|---|---|---|---|---|---|---|---|---|---|
| P98171 | ARHGAP4 | ERGLQAEyETQVkEM | Y17 | 0.338 | 84.87 | 946 | 0.148276 | 0.338 | 0.1483 | 1 |
| P27708 | CAD | VLRGEVAyIDGQVLV | Y1794 | 0.2 | 93.75 | 2225 | 0.087737 | 0.2 | 0.0877 | 1 |
| Q8TDN2 | KCNV2 | WGVRLKytPRCCRIC | T195 | 0.079 | 94.32 | 545 | 0.053958 | 0.079 | 0.054 | 1 |
| O60547 | GMDS | TGRIEHLykNPQAHI | Y69 | 0.304 | 87.39 | 372 | 0.13336 | 0.304 | 0.1334 | 1 |
| Q5GAN3 | RNASE13 | yPKGFRGyCNGLMSY | Y51 | 0.177 | 96.78 | 156 | 0.077647 | 0.177 | 0.0776 | 1 |
| Q9P218 | COL20A1 | TLtGRVttKKAPSPS | T644 | 0.078 | 88.13 | 1284 | 0.053275 | 0.078 | 0.0533 | 1 |
| Q12768 | WASHC5 | RITDPKMtCHIDQLN | T837 | 0.058 | 91.15 | 1159 | 0.039615 | 0.058 | 0.0396 | 1 |
| Q12768 | WASHC5 | CHIDQLNtWyDMKTH | T845 | 0.092 | 94.29 | 1159 | 0.062837 | 0.092 | 0.0628 | 0.8333 |
| Q8WV41 | SNX33 | LHKFtVIsVPHLPEk | S285 | 0.025 | 97.46 | 574 | 0.019873 | 0.025 | 0.0199 | 1 |
| O75054 | IGSF3 | SSSFRTRtAIEKAES | T617 | 0.207 | 85.41 | 1194 | 0.141384 | 0.207 | 0.1414 | 1 |
| Q496F6 | CD300E | sCESIVEtKGEEKVE | T64 | 0.038 | 97.43 | 205 | 0.025955 | 0.038 | 0.026 | 1 |
| Q14126 | DSG2 | NPIAKIHsDLAEErG | S75 | 0.013 | 87.66 | 1118 | 0.010334 | 0.013 | 0.0103 | 1 |
| Q14126 | DSG2 | DREEHSSyTLTVEAR | Y235 | 0.094 | 97.39 | 1118 | 0.041236 | 0.094 | 0.0412 | 1 |
| Q96S82 | UBL7 | LDQIIVAtPGLSSDP | T147 | 0.017 | 76.44 | 380 | 0.011611 | 0.017 | 0.0116 | 1 |
| O60674 | JAK2 | AGNQTGLyVLRCSPK | Y423 | 0.024 | 94.79 | 1132 | 0.010528 | 0.024 | 0.0105 | 1 |
| O60674 | JAK2 | SPKDFNKyFLTFAVE | Y435 | 0.181 | 97.84 | 1132 | 0.079402 | 0.181 | 0.0794 | 1 |
| Q03001 | DST | RDKVQKKtFtKWINQ | T40 | 0.01 | 96.25 | 7570 | 0.00683 | 0.01 | 0.0068 | 0.9167 |
| Q03001 | DST | NVQIALDyLKRRQVk | Y102 | 0.184 | 96.8 | 7570 | 0.080718 | 0.344 | 0.145 | 1 |
| Q9Y230 | RUVBL2 | IrGtsyQsPHGIPID | S342 | 0.052 | 93.86 | 463 | 0.041336 | 0.063 | 0.0547 | 1 |
| O43405 | COCH | HMPNWFGttKYVKPL | T339 | 0.175 | 92.35 | 550 | 0.119527 | 0.175 | 0.1195 | 1 |
| Q9Y2K9 | STXBP5L | ITGHADGsIKFWDAS | S472 | 0.02 | 97.07 | 1186 | 0.015898 | 0.02 | 0.0159 | 1 |
| Q2M3D2 | EXOC3L2 | HRRALVEyVRPLLRG | Y235 | 0.031 | 96.65 | 409 | 0.013599 | 0.031 | 0.0136 | 1 |
| Q9NQH7 | XPNPEP3 | LQPGMVItIEPGIyI | T449 | 0 | 98.95 | 507 | 0 | 0 | 0 | 1 |
| Q9NQH7 | XPNPEP3 | ItIEPGIyIPEDDKD | Y455 | 0.101 | 98.91 | 507 | 0.044307 | 0.159 | 0.0592 | 1 |
| Q15198 | PDGFRL | GQLtLVNsTSADTGE | S133 | 0.049 | 95.45 | 375 | 0.038951 | 0.049 | 0.039 | 0.9091 |
| Q15198 | PDGFRL | DMKRGFVyLQPHSEH | Y233 | 0.007 | 97.01 | 375 | 0.003071 | 0.007 | 0.0031 | 1 |
| Q15198 | PDGFRL | AGGRSQIsVKYQLLY | S257 | 0.158 | 96.9 | 375 | 0.125597 | 0.158 | 0.1256 | 1 |
| P13929 | ENO3 | RFRAAVPsGAstGIy | S37 | 0.028 | 95.91 | 434 | 0.022258 | 0.028 | 0.0223 | 1 |
| P13929 | ENO3 | AAVPsGAstGIyEAL | S40 | 0.122 | 74.95 | 434 | 0.09698 | 0.127 | 0.1426 | 1 |
| P13929 | ENO3 | MRIGAEVyHHLKGVI | Y189 | 0.117 | 98.9 | 434 | 0.051326 | 0.117 | 0.0513 | 1 |
| P13929 | ENO3 | EFyRNGkyDLDFksP | Y257 | 0.002 | 98.56 | 434 | 0.000877 | 0.002 | 0.0009 | 1 |

| | | | | | | | | | | |
|---|---|---|---|---|---|---|---|---|---|---|
| P13929 | ENO3 | LKVNQIGsVTEsIQA | S349 | 0 | 98.77 | 434 | 0 | 0 | 0 | 1 |
| P54278 | PMS2 | ASRACRKsVMIGTAL | S815 | 0.086 | 90.9 | 862 | 0.068363 | 0.086 | 0.0684 | 1 |
| Q53EL6 | PDCD4 | LPELALDtPRAPQLV | T254 | 0.159 | 93.88 | 469 | 0.108599 | 0.159 | 0.1086 | 1 |
| Q9Y312 | AAR2 | LLNLLCRsEAAMMKH | S284 | 0 | 95.92 | 384 | 0 | 0 | 0 | 1 |
| P35606 | COPB2 | MQIRVFNyNTLERVH | Y86 | 0.123 | 95.16 | 906 | 0.053958 | 0.123 | 0.054 | 1 |
| P35606 | COPB2 | PTQPFILtSSDDMLI | T116 | 0 | 98.82 | 906 | 0 | 0 | 0 | 1 |
| P35606 | COPB2 | WAHDSsEyAIRESNS | Y403 | 0.008 | 97.27 | 906 | 0.003509 | 0.008 | 0.0035 | 1 |
| P35606 | COPB2 | kNNVAFMsyFLQGKV | S731 | 0 | 92.39 | 906 | 0 | 0 | 0 | 1 |
| P35606 | COPB2 | NNVAFMsyFLQGKVD | Y732 | 0.194 | 92.74 | 906 | 0.085105 | 0.265 | 0.1163 | 1 |
| Q9NXW9 | ALKBH4 | VEQCNLDyCPERGSA | Y158 | 0.008 | 98.66 | 302 | 0.003509 | 0.008 | 0.0035 | 1 |
| Q9BUF5 | TUBB6 | QLERINVyYNESssQ | Y50 | 0.028 | 96.4 | 446 | 0.012283 | 0.056 | 0.0246 | 0.8 |
| Q9BUF5 | TUBB6 | PRGLkMAsTFIGNST | S365 | 0.004 | 95.69 | 446 | 0.00318 | 0.004 | 0.0032 | 1 |
| Q9BYE2 | TMPRSS13 | YNSTIQEsLHRsECP | S298 | 0.082 | 92.48 | 586 | 0.065183 | 0.082 | 0.0652 | 1 |
| P10586 | PTPRF | MktsVLLsWEVPDSY | S1031 | 0.123 | 80.11 | 1907 | 0.097775 | 0.072 | 0.1256 | 1 |
| Q9BXL7 | CARD11 | SGPVTLHyKVNHEGY | Y751 | 0.081 | 93.32 | 1154 | 0.035534 | 0.081 | 0.0355 | 1 |
| Q96S38 | RPS6KC1 | EDDyEAAsDFyRkGV | S263 | 0.147 | 90.15 | 1066 | 0.116852 | 0.147 | 0.1169 | 1 |
| Q96S38 | RPS6KC1 | yEAAsDFyRkGVDLL | Y266 | 0.007 | 95.39 | 1066 | 0.003071 | 0.007 | 0.0031 | 0.9231 |
| Q9NWX6 | THG1L | FsEFNINyNNELPMY | Y228 | 0.09 | 96.22 | 298 | 0.039482 | 0.09 | 0.0395 | 1 |
| Q9NWX6 | THG1L | LPMYRKGtVLIWQKV | T239 | 0.061 | 98.75 | 298 | 0.041664 | 0.061 | 0.0417 | 1 |
| A6NE02 | BTBD17 | LHRLATKyGVSSLQR | Y145 | 0.14 | 96.49 | 478 | 0.061416 | 0.14 | 0.0614 | 1 |
| P58753 | TIRAP | SGLSRAAyPPELRFM | Y187 | 0.094 | 92.73 | 221 | 0.041236 | 0.094 | 0.0412 | 0.9167 |
| P10912 | GHR | SLKVDkEyEVRVRSK | Y226 | 0.14 | 97.37 | 638 | 0.061416 | 0.14 | 0.0614 | 1 |
| P53677 | AP3M2 | ItVGPKQtMGkTIEG | T300 | 0.125 | 92.46 | 418 | 0.085377 | 0.125 | 0.0854 | 1 |
| Q9BXS9 | SLC26A6 | VSCSMSRsLVQESTG | S405 | 0 | 93.71 | 759 | 0 | 0 | 0 | 1 |
| Q9BXS9 | SLC26A6 | QVAGAISsLFILLII | S423 | 0 | 96.82 | 759 | 0 | 0 | 0 | 1 |
| A0JNW5 | BLTP3B | GPGAVIHsLLAEKNG | S1269 | 0.102 | 87.49 | 1464 | 0.081081 | 0.129 | 0.1109 | 0.8571 |
| P43353 | ALDH3B1 | LLEHRFDyIFFtGsP | Y183 | 0.167 | 98.9 | 468 | 0.07326 | 0.167 | 0.0733 | 1 |
| P43353 | ALDH3B1 | RFDyIFFtGsPRVGk | T187 | 0.116 | 98.83 | 468 | 0.07923 | 0.116 | 0.0792 | 0.9 |
| P43353 | ALDH3B1 | TAAAkHLtPVtLELG | T205 | 0.151 | 98.74 | 468 | 0.103135 | 0.151 | 0.1031 | 1 |
| P43353 | ALDH3B1 | PLALYAFsNSSQVVk | S368 | 0.027 | 98.25 | 468 | 0.021463 | 0.027 | 0.0215 | 1 |
| Q05193 | DNM1 | IstVRQCtKKLQQYP | T443 | 0 | 94.92 | 864 | 0 | 0.047 | 0.0321 | 1 |

| | | | | | | | | | | |
|---|---|---|---|---|---|---|---|---|---|---|
| P51955 | NEK2 | VGtPYYMsPEQMNRM | S184 | 0 | 90.24 | 445 | 0 | 0 | 0 | 1 |
| Q9P1W8 | SIRPG | EFKSGPGtEMALGAK | T138 | 0.009 | 98.25 | 387 | 0.006147 | 0.009 | 0.0061 | 1 |
| Q99624 | SLC38A3 | QNIGAMSsyLYIIKS | S159 | 0 | 95.47 | 504 | 0 | 0 | 0 | 0.8 |
| Q16627 | CCL14 | SDKWVQDyIKDMKEN | Y86 | 0.218 | 97.72 | 93 | 0.095633 | 0.218 | 0.0956 | 1 |
| Q9Y2S2 | CRYL1 | GGFQVKLyDIEQQQI | Y35 | 0.294 | 98.84 | 319 | 0.128974 | 0.294 | 0.129 | 1 |
| Q8IUS5 | EPHX4 | APIHRQNyKLDCLIt | Y142 | 0.02 | 96.33 | 362 | 0.008774 | 0.02 | 0.0088 | 0.8 |
| Q96T76 | MMS19 | DMkLVWPsAkLLQAA | S371 | 0 | 97.06 | 1030 | 0 | 0.056 | 0.0445 | 0.8889 |
| Q13555 | CAMK2G | CTRFTDDyQLFEELG | Y14 | 0.077 | 96.37 | 558 | 0.033779 | 0.154 | 0.0676 | 1 |
| Q13555 | CAMK2G | YYSEADAsHCIHQIL | S114 | 0 | 98.35 | 558 | 0 | 0 | 0 | 1 |
| Q13555 | CAMK2G | sMMHRQEtVECLRKF | T287 | 0.046 | 92.17 | 558 | 0.031419 | 0.046 | 0.0314 | 1 |
| Q9Y6M9 | NDUFB9 | WKKLRREsWEREVkQ | S132 | 0.034 | 96.84 | 179 | 0.027027 | 0.034 | 0.027 | 1 |
| P04083 | ANXA1 | RQQIkAAyLQEtGkP | Y84 | 0.015 | 98.78 | 346 | 0.00658 | 0.015 | 0.0066 | 1 |
| P04083 | ANXA1 | IRDINRVyREELkRD | Y156 | 0.009 | 98.8 | 346 | 0.003948 | 0.009 | 0.0039 | 1 |
| P04083 | ANXA1 | AkDItsDtsGDFRNA | T172 | 0.118 | 97.38 | 346 | 0.080596 | 0.118 | 0.0806 | 1 |
| Q13547 | HDAC1 | TkYHSDDyIkFLRSI | Y72 | 0 | 98.35 | 482 | 0 | 0 | 0 | 1 |
| P31751 | AKT2 | EKATGRyyAMKILRk | Y178 | 0.192 | 95.35 | 481 | 0.084228 | 0.192 | 0.0842 | 1 |
| P31751 | AKT2 | EVLEDNDyGRAVDWW | Y327 | 0.072 | 85.53 | 481 | 0.031585 | 0.072 | 0.0316 | 1 |
| Q8NB90 | AFG2A | VPLPDAAtRREIFKL | T801 | 0.092 | 91.22 | 893 | 0.062837 | 0.092 | 0.0628 | 0.8571 |
| O00541 | PES1 | STAARTFyLIkDIRF | Y65 | 0.234 | 94.7 | 588 | 0.102652 | 0.234 | 0.1027 | 1 |
| P51532 | SMARCA4 | sPNPPNLtKKMKKIV | T1459 | 0.214 | 88.33 | 1647 | 0.146165 | 0.214 | 0.1462 | 1 |
| Q96P50 | ACAP3 | VREDLELsLVRNAQA | S135 | 0.134 | 95.07 | 834 | 0.106519 | 0.134 | 0.1065 | 1 |
| Q9GZV4 | EIF5A2 | PNIkRNDyQLICIQD | Y89 | 0.229 | 94.44 | 153 | 0.100459 | 0.229 | 0.1005 | 1 |
| Q9HD33 | MRPL47 | RLDKVVDsMDALDkV | S138 | 0.038 | 98.15 | 250 | 0.030207 | 0.038 | 0.0302 | 1 |
| P11473 | VDR | CKGFFRRsMKRKALF | S51 | 0.052 | 96.99 | 427 | 0.041336 | 0.052 | 0.0413 | 1 |
| Q7Z406 | MYH14 | YSGLIYTySGLFCVV | Y137 | 0.143 | 90.48 | 1995 | 0.062732 | 0.143 | 0.0627 | 1 |
| Q7Z406 | MYH14 | TEAIVEMyRGKKRHE | Y162 | 0.149 | 92.95 | 1995 | 0.065364 | 0.292 | 0.1281 | 1 |
| Q7Z406 | MYH14 | NTKKVIQyLAHVAss | Y214 | 0.006 | 92.88 | 1995 | 0.002632 | 0.006 | 0.0026 | 1 |
| Q7Z406 | MYH14 | LRDQADFsVLHYAGK | S595 | 0.003 | 90.92 | 1995 | 0.002385 | 0.003 | 0.0024 | 1 |
| Q7Z406 | MYH14 | RQRYEILtPNAIPKG | T749 | 0.069 | 88.59 | 1995 | 0.047128 | 0.069 | 0.0471 | 1 |
| Q8IXT5 | RBM12B | NCGsGRVtPIKIMNL | T925 | 0.137 | 85.42 | 1001 | 0.093573 | 0.21 | 0.1434 | 1 |
| P0C0L5 | C4B | GAHAAAItAYALTLT | T1196 | 0.011 | 95.05 | 1744 | 0.007513 | 0.011 | 0.0075 | 1 |

| | | | | | | | | | | |
|---|---|---|---|---|---|---|---|---|---|---|
| P06576 | ATP5F1B | kIPVGPEtLGRIMNV | T140 | 0.01 | 94.37 | 529 | 0.00683 | 0.01 | 0.0068 | 0.9 |
| P06576 | ATP5F1B | IDNIFRFtQAGsEVs | T312 | 0.033 | 93.44 | 529 | 0.022539 | 0.033 | 0.0225 | 1 |
| P06576 | ATP5F1B | tQAGsEVsALLGRIP | S319 | 0.039 | 94.24 | 529 | 0.031002 | 0.039 | 0.031 | 1 |
| Q13523 | PRPF4B | NWTDAEGyYRVNIGE | Y674 | 0.225 | 87.24 | 1007 | 0.098704 | 0.225 | 0.0987 | 1 |
| Q13523 | PRPF4B | PyLVsRFyRAPEIII | Y855 | 0.048 | 97.07 | 1007 | 0.021057 | 0.127 | 0.0557 | 0.8462 |
| Q14258 | TRIM25 | CPQCRAVyQARPQLH | Y57 | 0.327 | 90.4 | 630 | 0.14345 | 0.327 | 0.1434 | 0.8182 |
| P23368 | ME2 | GLGDLGVyGMGIPVG | Y175 | 0.008 | 98.21 | 584 | 0.003509 | 0.008 | 0.0035 | 1 |
| Q4G0M1 | ERFE | RAPVAGFyALAAtLH | Y257 | 0.003 | 96.23 | 354 | 0.001316 | 0.003 | 0.0013 | 1 |
| P30838 | ALDH3A1 | QYLDKDLyPVINGGV | Y163 | 0.071 | 98.91 | 453 | 0.031147 | 0.071 | 0.0311 | 1 |
| P30838 | ALDH3A1 | ERFDHILyTGSTGVG | Y186 | 0.165 | 98.93 | 453 | 0.072383 | 0.165 | 0.0724 | 0.9091 |
| P30838 | ALDH3A1 | tAAAkHLtPVTLELG | T205 | 0.156 | 98.93 | 453 | 0.10655 | 0.156 | 0.1066 | 1 |
| P30838 | ALDH3A1 | KKSLkEFyGEDAkKS | Y272 | 0.261 | 98.85 | 453 | 0.114497 | 0.261 | 0.1145 | 1 |
| Q96FV9 | THOC1 | LINALNKsGLSDLAE | S640 | 0.186 | 86.6 | 657 | 0.147854 | 0.186 | 0.1479 | 1 |
| Q53EP0 | FNDC3B | DLRPATDyHVRVYAM | Y351 | 0.143 | 93.74 | 1204 | 0.062732 | 0.143 | 0.0627 | 0.9 |
| Q53EP0 | FNDC3B | KAPIDNGskITNYLL | S407 | 0.118 | 91.56 | 1204 | 0.0938 | 0.087 | 0.1222 | 1 |
| Q53EP0 | FNDC3B | RAGITWVtLQWsKPE | T492 | 0.099 | 95.78 | 1204 | 0.067618 | 0.099 | 0.0676 | 1 |
| Q9Y2W2 | WBP11 | LRkLEVEyEQkRAQL | Y114 | 0.072 | 96.89 | 641 | 0.031585 | 0.072 | 0.0316 | 1 |
| Q9BZ29 | DOCK9 | YKFPSTFyEGRADMC | Y1468 | 0.116 | 88.05 | 2069 | 0.050888 | 0.331 | 0.1452 | 0.8 |
| Q14192 | FHL2 | ILREESPyCVVCFET | Y27 | 0.252 | 94.61 | 279 | 0.110549 | 0.252 | 0.1105 | 0.9 |
| Q9HCC8 | GDPD2 | PMLAPENtLMsLRKT | T240 | 0.005 | 98.46 | 539 | 0.003415 | 0.005 | 0.0034 | 1 |
| Q9HCC8 | GDPD2 | APENtLMsLRKTAEC | S243 | 0 | 97.98 | 539 | 0 | 0 | 0 | 1 |
| Q9HCC8 | GDPD2 | PTRITAHsSDFSWtE | S293 | 0 | 97.71 | 539 | 0 | 0 | 0 | 1 |
| Q99715 | COL12A1 | VLITDGksQDDVEAP | S1312 | 0.148 | 82.07 | 3063 | 0.117647 | 0.148 | 0.1176 | 0.8571 |
| P38919 | EIF4A3 | ATQVVLIsATLPHEI | S218 | 0.001 | 92.3 | 411 | 0.000795 | 0.001 | 0.0008 | 0.9 |
| Q9HCS2 | CYP4F12 | LDMFEHIsLMTLDSL | S196 | 0.059 | 97.23 | 524 | 0.0469 | 0.059 | 0.0469 | 0.8182 |
| Q9BSJ8 | ESYT1 | VkGKSDPyVkLKLAG | Y677 | 0.126 | 94.57 | 1104 | 0.055274 | 0.248 | 0.1074 | 1 |
| P12955 | PEPD | VECRVFKtDMELEVL | T188 | 0.135 | 98.84 | 493 | 0.092207 | 0.135 | 0.0922 | 0.8333 |
| P41226 | UBA7 | DPTTEHIyGDNFFSR | Y524 | 0.025 | 92.11 | 1012 | 0.010967 | 0.025 | 0.011 | 1 |
| P41226 | UBA7 | PRsAFRHsYLHLAEN | S884 | 0.029 | 97.9 | 1012 | 0.023053 | 0.029 | 0.0231 | 1 |
| Q14693 | LPIN1 | HkVsQNGyKFLYCSA | Y717 | 0.047 | 97.71 | 890 | 0.020618 | 0.047 | 0.0206 | 1 |
| P21266 | GSTM3 | LDFPNLPyLLDGkNk | Y66 | 0.058 | 98.7 | 225 | 0.025444 | 0.058 | 0.0254 | 1 |

| | | | | | | | | | | |
|---|---|---|---|---|---|---|---|---|---|---|
| O95563 | MPC2 | GFIWSRysLVIIPKN | S86 | 0.069 | 93.14 | 127 | 0.054849 | 0.069 | 0.0548 | 1 |
| Q1EHB4 | SLC5A12 | ILVCAVFsGLIMYSH | S293 | 0.005 | 97.55 | 618 | 0.003975 | 0.005 | 0.004 | 0.9 |
| Q92537 | SUSD6 | WKPAMEIsCRLNEDK | S101 | 0.168 | 91.5 | 303 | 0.133546 | 0.168 | 0.1335 | 1 |
| P46109 | CRKL | KIHyLDtttLIEPAP | T96 | 0.112 | 79.32 | 303 | 0.076498 | 0.195 | 0.1332 | 1 |
| Q04446 | GBE1 | REGDNVNyDWIHWDP | Y173 | 0.07 | 98.08 | 702 | 0.030708 | 0.07 | 0.0307 | 1 |
| Q04446 | GBE1 | SRPKKPRsLRIyEsH | S196 | 0.035 | 98.38 | 702 | 0.027822 | 0.068 | 0.0548 | 1 |
| Q04446 | GBE1 | KPRsLRIyEsHVGIS | Y200 | 0.002 | 98.92 | 702 | 0.000877 | 0.002 | 0.0009 | 0.8889 |
| Q04446 | GBE1 | yLEkCIAyAESHDQA | Y476 | 0.112 | 98.67 | 702 | 0.049133 | 0.218 | 0.0935 | 0.9091 |
| Q04446 | GBE1 | kFkIVLDsDAAEyGG | S652 | 0.006 | 98.83 | 702 | 0.004769 | 0.006 | 0.0048 | 1 |
| Q04446 | GBE1 | LDsDAAEyGGHQRLD | Y657 | 0.155 | 98.83 | 702 | 0.067996 | 0.155 | 0.068 | 1 |
| O75414 | NME6 | ARHVAPDsIRGSFGL | S114 | 0.095 | 98.59 | 186 | 0.075517 | 0.095 | 0.0755 | 1 |
| P23467 | PTPRB | KYKIQILtVsGGLFS | T1067 | 0.007 | 87.17 | 1997 | 0.004781 | 0.007 | 0.0048 | 1 |
| P23467 | PTPRB | KIQILtVsGGLFSKE | S1069 | 0 | 80.99 | 1997 | 0 | 0 | 0 | 1 |
| P23467 | PTPRB | GLRPGRSyQFNVKTV | Y1328 | 0.1 | 87.21 | 1997 | 0.043869 | 0.1 | 0.0439 | 1 |
| Q8N3C0 | ASCC3 | PVRIVGLstALANAR | S1490 | 0.001 | 88.78 | 2202 | 0.000795 | 0.001 | 0.0008 | 1 |
| Q8N3C0 | ASCC3 | AMLPCPDyDTDTKtV | Y1909 | 0.003 | 87.15 | 2202 | 0.001316 | 0.003 | 0.0013 | 1 |
| P50748 | KNTC1 | HKLNTEEyLRVIGKP | Y1762 | 0.171 | 79.39 | 2209 | 0.075015 | 0.171 | 0.075 | 1 |
| Q96Q91 | SLC4A9 | PVVLGsLtEVSLPSR | T257 | 0.039 | 76.32 | 983 | 0.026638 | 0.18 | 0.1229 | 1 |
| Q7Z6M3 | MILR1 | EAHEsGPyKCKAQVt | Y101 | 0.04 | 98.6 | 343 | 0.017547 | 0.04 | 0.0175 | 1 |
| Q13643 | FHL3 | VPDkGAHyCVPCYEN | Y149 | 0.17 | 96.38 | 280 | 0.074577 | 0.17 | 0.0746 | 0.9 |
| Q6ZPD9 | DPY19L3 | ALCIGLLtsVYLATL | T61 | 0.071 | 96.11 | 716 | 0.048494 | 0.071 | 0.0485 | 1 |
| Q6ZPD9 | DPY19L3 | LCIGLLtsVYLATLH | S62 | 0.157 | 96.82 | 716 | 0.124802 | 0.157 | 0.1248 | 1 |
| P38570 | ITGAE | HGEEGRVyVyRLSEQ | Y549 | 0.106 | 97.89 | 1179 | 0.046501 | 0.106 | 0.0465 | 1 |
| P38570 | ITGAE | EEGRVyVyRLSEQDG | Y551 | 0.228 | 96.61 | 1179 | 0.10002 | 0.228 | 0.1 | 1 |
| Q9BQI3 | EIF2AK1 | VGTCLYAsPEQLEGS | S498 | 0 | 92.74 | 630 | 0 | 0 | 0 | 0.9 |
| Q6K0P9 | PYHIN1 | ENMsKLMsEMHSFIQ | S390 | 0.177 | 96.78 | 492 | 0.1407 | 0.177 | 0.1407 | 1 |
| Q9HBG4 | ATP6V0A4 | FNRTNKFtAGFQNIV | T373 | 0.158 | 88.63 | 840 | 0.107916 | 0.158 | 0.1079 | 1 |
| Q9HBG4 | ATP6V0A4 | NSYKMKMsVILGIVQ | S544 | 0.02 | 94.73 | 840 | 0.015898 | 0.02 | 0.0159 | 1 |
| Q9UPZ3 | HPS5 | LPPLPVItLRSEPQY | T273 | 0.189 | 89.89 | 1129 | 0.12909 | 0.189 | 0.1291 | 1 |
| O15120 | AGPAT2 | QVLEAIPtsGLtAAD | T233 | 0.015 | 98.24 | 278 | 0.010245 | 0.015 | 0.0102 | 1 |
| Q9NSE4 | IARS2 | PDQRADLyLEGKDQL | Y621 | 0 | 98.64 | 1012 | 0 | 0 | 0 | 1 |

| | | | | | | | | | | |
|---|---|---|---|---|---|---|---|---|---|---|
| Q9NSE4 | IARS2 | TRELSNFyFSIIkDR | Y798 | 0.004 | 97.71 | 1012 | 0.001755 | 0.004 | 0.0018 | 1 |
| P13716 | ALAD | AKFASCFyGPFRDAA | Y205 | 0.117 | 94.31 | 330 | 0.051326 | 0.117 | 0.0513 | 0.9167 |
| P13716 | ALAD | PFRDAAkssPAFGDR | S214 | 0.029 | 80.37 | 330 | 0.023053 | 0.029 | 0.0231 | 0.8571 |
| Q12794 | HYAL1 | GTGKSQMyVQHRVAE | Y261 | 0.023 | 98.77 | 435 | 0.01009 | 0.023 | 0.0101 | 1 |
| P26038 | MSN | EELIQDItQRLFFLQ | T98 | 0 | 97.93 | 577 | 0 | 0 | 0 | 1 |
| P26038 | MSN | ETAVLLAsYAVQsky | S127 | 0 | 97.95 | 577 | 0 | 0 | 0 | 1 |
| P26038 | MSN | LAsYAVQskyGDFNk | S132 | 0.16 | 97.03 | 577 | 0.127186 | 0.16 | 0.1272 | 1 |
| P26038 | MSN | REDAVLEyLKIAQDL | Y191 | 0.079 | 98.47 | 577 | 0.034656 | 0.119 | 0.0522 | 1 |
| P26038 | MSN | DALGLNIyEQNDRLT | Y228 | 0.039 | 97.81 | 577 | 0.017109 | 0.075 | 0.0329 | 1 |
| P26038 | MSN | CMGNHELyMRRRkPD | Y291 | 0.018 | 93.08 | 577 | 0.007896 | 0.101 | 0.0443 | 1 |
| Q96GA7 | SDSL | CLPPSLtsVVVIVCG | S303 | 0.106 | 97.29 | 329 | 0.084261 | 0.106 | 0.0843 | 1 |
| Q00G26 | PLIN5 | KVAEVRRsVDALQTA | S322 | 0.128 | 92.5 | 463 | 0.101749 | 0.128 | 0.1017 | 1 |
| O14607 | UTY | NkLKSVKsPVPMVHL | S1213 | 0.069 | 96.51 | 1347 | 0.054849 | 0.069 | 0.0548 | 1 |
| Q5PRF9 | SAMD4B | NTFQEDGsGMKDVPS | S299 | 0.146 | 89.9 | 694 | 0.116058 | 0.146 | 0.1161 | 1 |
| Q92630 | DYRK2 | sQKLLDAsKRAKNFV | S449 | 0.101 | 97.84 | 601 | 0.080286 | 0.101 | 0.0803 | 1 |
| Q70J99 | UNC13D | ELTkHSTsAVDLSTC | S629 | 0.001 | 95.15 | 1090 | 0.000795 | 0.001 | 0.0008 | 1 |
| Q14974 | KPNB1 | PTFLVELsRVLANPG | S41 | 0.003 | 97.83 | 876 | 0.002385 | 0.003 | 0.0024 | 1 |
| Q14974 | KPNB1 | PEQLQDKsNEILtAI | S170 | 0.032 | 94.92 | 876 | 0.025437 | 0.032 | 0.0254 | 1 |
| Q14974 | KPNB1 | MQVVCEAtQCPDTRV | T226 | 0.106 | 98.02 | 876 | 0.072399 | 0.209 | 0.1424 | 1 |
| Q14974 | KPNB1 | NNLRssAyEsLMEIV | Y529 | 0.036 | 98.17 | 876 | 0.015793 | 0.036 | 0.0158 | 1 |
| Q14974 | KPNB1 | LRssAyEsLMEIVkN | S531 | 0.006 | 98.12 | 876 | 0.004769 | 0.006 | 0.0048 | 1 |
| Q14974 | KPNB1 | LMEIVkNsAkDCYPA | S539 | 0.006 | 96.05 | 876 | 0.004769 | 0.006 | 0.0048 | 1 |
| Q14974 | KPNB1 | LQSLLCAtLQNVLRk | T587 | 0.021 | 96.44 | 876 | 0.014343 | 0.021 | 0.0143 | 1 |
| Q14974 | KPNB1 | LGGEFLKyMEAFkPF | Y646 | 0.239 | 97.64 | 876 | 0.104846 | 0.239 | 0.1048 | 0.8462 |
| Q14974 | KPNB1 | LNtLQQAsQAQVDKs | S743 | 0.118 | 95 | 876 | 0.0938 | 0.118 | 0.0938 | 1 |
| Q14974 | KPNB1 | LLTEGRRsKtNKAkT | S853 | 0.155 | 90.1 | 876 | 0.123212 | 0.155 | 0.1232 | 1 |
| Q8TCG5 | CPT1C | LRGAKILsENVDCHV | S538 | 0.075 | 95.78 | 803 | 0.059619 | 0.075 | 0.0596 | 1 |
| Q8TEM1 | NUP210 | EAEQILMsPNsyIKL | S1301 | 0.051 | 93.27 | 1887 | 0.040541 | 0.051 | 0.0405 | 1 |
| Q8TEM1 | NUP210 | QILMsPNsyIKLQtN | S1304 | 0.02 | 92.66 | 1887 | 0.015898 | 0.02 | 0.0159 | 1 |
| Q8TEM1 | NUP210 | PVSYLRVsMSPVLHT | S1380 | 0.105 | 91.19 | 1887 | 0.083466 | 0.148 | 0.1246 | 1 |
| P07099 | EPHX1 | LLTNVMLyWTTGTII | Y361 | 0.052 | 98.78 | 455 | 0.022812 | 0.052 | 0.0228 | 0.8125 |

| | | | | | | | | | | |
|---|---|---|---|---|---|---|---|---|---|---|
| A2A3L6 | TTC24 | yDQALKYyKEALAQC | Y376 | 0.1 | 97.33 | 582 | 0.043869 | 0.1 | 0.0439 | 1 |
| Q641Q2 | WASHC2A | sVEEIRRssQsWSLA | S31 | 0.119 | 85.97 | 1341 | 0.094595 | 0.119 | 0.0946 | 1 |
| P11171 | EPB41 | VMELHkSyRSMtPAQ | Y374 | 0.234 | 96.58 | 864 | 0.102652 | 0.234 | 0.1027 | 1 |
| O95864 | FADS2 | LVIDRkVyNITkWSI | Y44 | 0.158 | 98.43 | 444 | 0.069312 | 0.158 | 0.0693 | 1 |
| P20701 | ITGAL | ELSSSGIsADLSRGH | S355 | 0.123 | 98.49 | 1170 | 0.097775 | 0.123 | 0.0978 | 1 |
| O75746 | SLC25A12 | DFsDIMVtIRSHMLT | T188 | 0.219 | 88.48 | 678 | 0.14958 | 0.219 | 0.1496 | 1 |
| P06213 | INSR | KLIMItDyLLLFRVy | Y87 | 0.208 | 94.65 | 1382 | 0.091247 | 0.364 | 0.1355 | 0.9167 |
| Q9UIA0 | CYTH4 | NKtAIGTyLGERDPI | Y114 | 0.18 | 97.39 | 394 | 0.078963 | 0.18 | 0.079 | 1 |
| Q76N89 | HECW1 | SRRLIsFsLsDFQAM | S208 | 0.012 | 92.08 | 1606 | 0.009539 | 0.012 | 0.0095 | 1 |
| Q76N89 | HECW1 | RLIsFsLsDFQAMGL | S210 | 0.153 | 93.36 | 1606 | 0.121622 | 0.153 | 0.1216 | 0.8333 |
| P07942 | LAMB1 | IICDSRYsDIEPSTE | S195 | 0.005 | 89.78 | 1786 | 0.003975 | 0.005 | 0.004 | 1 |
| P07942 | LAMB1 | IQNLLkItNLRIKFV | T231 | 0.009 | 91.78 | 1786 | 0.006147 | 0.009 | 0.0061 | 1 |
| P07942 | LAMB1 | CDDCQHNtMGRNCEQ | T369 | 0.004 | 92.29 | 1786 | 0.002732 | 0.004 | 0.0027 | 1 |
| P57768 | SNX16 | DPPGPFDsLEESRAF | S222 | 0.042 | 80.02 | 344 | 0.033386 | 0.042 | 0.0334 | 1 |
| Q13356 | PPIL2 | KGyVRLHtNKGDLNL | T286 | 0.131 | 95.52 | 520 | 0.089475 | 0.131 | 0.0895 | 1 |
| Q13356 | PPIL2 | AYLDKkHtIFGRVVG | T395 | 0.001 | 95.68 | 520 | 0.000683 | 0.001 | 0.0007 | 1 |
| P26583 | HMGB2 | sAkDkQPyEQkAAkL | Y144 | 0.306 | 95.31 | 209 | 0.134238 | 0.306 | 0.1342 | 0.8 |
| O94989 | ARHGEF15 | VVtSEASyLRsLRLL | Y432 | 0 | 96.03 | 841 | 0 | 0.12 | 0.0526 | 1 |
| O94989 | ARHGEF15 | LRsLRLLtDTFVLSQ | T440 | 0.07 | 97.11 | 841 | 0.047811 | 0.07 | 0.0478 | 1 |
| Q9H7T0 | CATSPERB | PRNMFHMsKKtGKFK | S902 | 0.002 | 89.24 | 1116 | 0.00159 | 0.002 | 0.0016 | 1 |
| Q9H7T0 | CATSPERB | FKFPItQyPVsLEII | Y951 | 0.166 | 93.72 | 1116 | 0.072822 | 0.166 | 0.0728 | 1 |
| Q969T9 | WBP2 | PyRVIFLskGkDAMQ | S61 | 0.14 | 97.19 | 261 | 0.111288 | 0.14 | 0.1113 | 1 |
| Q969T9 | WBP2 | kGkDAMQsFMMPFYL | S69 | 0.007 | 96.95 | 261 | 0.005564 | 0.007 | 0.0056 | 0.8 |
| P05013 | IFNA6 | HsLGHRRtMMLLAQM | T37 | 0 | 96.17 | 189 | 0 | 0.145 | 0.099 | 1 |
| P56597 | NME5 | PEQCSNFyVEKYGKM | Y58 | 0.231 | 94.35 | 212 | 0.101336 | 0.231 | 0.1013 | 0.8 |
| P25789 | PSMA4 | DSRTTIFsPEGRLYQ | S13 | 0.122 | 92.63 | 261 | 0.09698 | 0.122 | 0.097 | 1 |
| P25789 | PSMA4 | LNEDMACsVAGItsD | S75 | 0 | 97.37 | 261 | 0 | 0 | 0 | 0.8333 |
| Q8WXG9 | ADGRV1 | DNAFGIIsFNMLPsI | S148 | 0 | 93.32 | 6306 | 0 | 0 | 0 | 1 |
| P05543 | SERPINA7 | NsHVEMQtKGKVVGL | T181 | 0.018 | 91.34 | 415 | 0.012294 | 0.018 | 0.0123 | 1 |
| P32322 | PYCR1 | VREGATVyATGTHAQ | Y135 | 0.028 | 98.81 | 319 | 0.012283 | 0.028 | 0.0123 | 1 |
| P19438 | TNFRSF1A | ECEsGSFtASENHLR | T90 | 0.039 | 98.48 | 455 | 0.026638 | 0.039 | 0.0266 | 1 |

| | | | | | | | | | | |
|---|---|---|---|---|---|---|---|---|---|---|
| A2RRH5 | WDR27 | KVLCLLAsLFGGKIA | S376 | 0.134 | 96.58 | 827 | 0.106519 | 0.134 | 0.1065 | 1 |
| Q9Y4G6 | TLN2 | GNPKAQHtHDAITEA | T1793 | 0.04 | 85.65 | 2542 | 0.027321 | 0.04 | 0.0273 | 1 |
| Q9UQ16 | DNM3 | PINLRVysPHVLNLT | S126 | 0.003 | 93.57 | 869 | 0.002385 | 0.003 | 0.0024 | 1 |
| Q9UQ16 | DNM3 | HIADRMGtPHLQKVL | T274 | 0.042 | 94.21 | 869 | 0.028687 | 0.042 | 0.0287 | 0.9 |
| Q9UK61 | TASOR | YAVVSFTyKDDIQtP | Y327 | 0.271 | 93.49 | 1670 | 0.118884 | 0.271 | 0.1189 | 0.8182 |
| Q9UHC3 | ASIC3 | TVEQKKAyEMSELLG | Y429 | 0.242 | 94.26 | 531 | 0.106162 | 0.242 | 0.1062 | 1 |
| Q9NS23 | RASSF1 | VkHLHVLsRtRAREV | S222 | 0.122 | 93.78 | 344 | 0.09698 | 0.122 | 0.097 | 0.875 |
| P47989 | XDH | RKLGLSGtKLGCGEG | T39 | 0.034 | 98.78 | 1333 | 0.023222 | 0.034 | 0.0232 | 1 |
| P47989 | XDH | AKLPAQKtEVFRGVL | T324 | 0.012 | 98.07 | 1333 | 0.008196 | 0.012 | 0.0082 | 1 |
| P47989 | XDH | ETRSTVVsTAVALAA | S810 | 0.004 | 98.9 | 1333 | 0.00318 | 0.004 | 0.0032 | 1 |
| P47989 | XDH | TKMVQVAsRALKIPT | S1052 | 0.022 | 98.84 | 1333 | 0.017488 | 0.04 | 0.0318 | 1 |
| Q9NPQ8 | RIC8A | VIQPMGMsPRGHLTS | S502 | 0.074 | 81.14 | 531 | 0.058824 | 0.116 | 0.0922 | 1 |
| Q12913 | PTPRJ | GVRKAALsWSNGNGt | S226 | 0.004 | 91.5 | 1337 | 0.00318 | 0.004 | 0.0032 | 1 |
| Q9H3D4 | TP63 | VIRAMPVykKAEHVT | Y231 | 0.051 | 97.09 | 680 | 0.022373 | 0.051 | 0.0224 | 1 |
| Q9H3D4 | TP63 | RPILIIVtLEtRDGQ | T326 | 0.111 | 97.97 | 680 | 0.075815 | 0.111 | 0.0758 | 1 |
| Q5VT25 | CDC42BPA | LEtIRLIyFKNKMAE | Y1510 | 0.049 | 92.18 | 1732 | 0.021496 | 0.049 | 0.0215 | 1 |
| P28331 | NDUFS1 | GsPVDLtytYDHLGD | Y440 | 0.141 | 96.93 | 727 | 0.061855 | 0.141 | 0.0619 | 0.875 |
| P28331 | NDUFS1 | kPMVVLGssALQRND | S478 | 0.018 | 96.79 | 727 | 0.014308 | 0.018 | 0.0143 | 0.8 |
| P28331 | NDUFS1 | PMVVLGssALQRNDG | S479 | 0.133 | 96.18 | 727 | 0.105724 | 0.133 | 0.1057 | 0.9 |
| Q8WUM9 | SLC20A1 | TTHCKVGsVVSVGWL | S629 | 0 | 95.68 | 679 | 0 | 0 | 0 | 1 |
| O00591 | GABRP | ISEsNMDyTATIYLR | Y82 | 0.032 | 95.89 | 440 | 0.014038 | 0.032 | 0.014 | 1 |
| Q8NGM1 | OR4C15 | TPKMIVDsLYVTKTI | S84 | 0.123 | 94.61 | 316 | 0.097775 | 0.123 | 0.0978 | 1 |
| Q6TDP4 | KLHL17 | CLNSAERyDPLtGTW | Y456 | 0.061 | 98.36 | 642 | 0.02676 | 0.061 | 0.0268 | 0.8 |
| Q6TDP4 | KLHL17 | SLNsIEKyNPRTNKW | Y597 | 0.048 | 98.76 | 642 | 0.021057 | 0.048 | 0.0211 | 1 |
| P11277 | SPTB | VDALEREsQQLMDSH | S1018 | 0.014 | 86.48 | 2137 | 0.011129 | 0.091 | 0.0766 | 1 |
| Q9ULJ7 | ANKRD50 | IEDAHGHtPLTLAAR | T580 | 0.022 | 95.43 | 1429 | 0.015026 | 0.022 | 0.015 | 1 |
| Q5I0G3 | MDH1B | RtFVNLKtVLLMRYA | T265 | 0 | 98.59 | 518 | 0 | 0 | 0 | 1 |
| P17480 | UBTF | PPNSYsLyCAELMAN | Y308 | 0.209 | 93.1 | 764 | 0.091685 | 0.209 | 0.0917 | 1 |
| P16452 | EPB42 | ERNPPENtFLRLtAM | T560 | 0.101 | 88.56 | 691 | 0.068984 | 0.101 | 0.069 | 1 |
| O75962 | TRIO | VTAVDKRyRDFSLRM | Y1228 | 0.006 | 89.66 | 3097 | 0.002632 | 0.278 | 0.1068 | 1 |
| Q96LI5 | CNOT6L | GVIDyIFysKTHMNV | Y493 | 0.128 | 98.25 | 555 | 0.056152 | 0.167 | 0.0733 | 1 |

| | | | | | | | | | | |
|---|---|---|---|---|---|---|---|---|---|---|
| P00747 | PLG | YDGKISKtMSGLECQ | T200 | 0.089 | 94.84 | 810 | 0.060788 | 0.141 | 0.0922 | 1 |
| O15360 | FANCA | VQMQREWsFARtHPL | S347 | 0.178 | 85.06 | 1455 | 0.141495 | 0.178 | 0.1415 | 1 |
| O15360 | FANCA | REWsFARtHPLLTsL | T351 | 0.132 | 83.64 | 1455 | 0.090158 | 0.132 | 0.0902 | 1 |
| Q9UID3 | VPS51 | MKRVVEDtTAIDVQV | T620 | 0.028 | 94.03 | 782 | 0.019124 | 0.028 | 0.0191 | 1 |
| Q9BZJ0 | CRNKL1 | ARRALGtsIGkCPKN | S599 | 0.002 | 90.36 | 848 | 0.00159 | 0.002 | 0.0016 | 1 |
| Q99436 | PSMB7 | DVTGPHLySIYPHGS | Y154 | 0.177 | 98.23 | 277 | 0.077647 | 0.259 | 0.1136 | 0.8889 |
| Q09428 | ABCC8 | SPFNKQRyKMVIEAC | Y798 | 0.178 | 93.58 | 1581 | 0.078086 | 0.286 | 0.1255 | 1 |
| Q8N1C3 | GABRG1 | YPKNEIEyKWKKPSV | Y218 | 0.053 | 96.86 | 465 | 0.02325 | 0.053 | 0.0232 | 0.875 |
| Q96JT2 | SLC45A3 | VPLLGsAsDHWRGRY | S75 | 0 | 90.32 | 553 | 0 | 0.032 | 0.0254 | 0.875 |
| Q14012 | CAMK1 | AEDIRDIyDFRDVLG | Y20 | 0.151 | 96.93 | 370 | 0.066242 | 0.151 | 0.0662 | 1 |
| Q96PU5 | NEDD4L | IRLLQFVtGTSRVPM | T903 | 0.028 | 82.8 | 975 | 0.019124 | 0.028 | 0.0191 | 0.9167 |
| Q8IY18 | SMC5 | kIELLEAtEKSIGPP | T199 | 0.024 | 88.27 | 1101 | 0.016392 | 0.024 | 0.0164 | 1 |
| Q8IY18 | SMC5 | NTACKENtsQYFFIT | T1046 | 0.099 | 78.3 | 1101 | 0.067618 | 0.174 | 0.1188 | 1 |
| Q8IY18 | SMC5 | TACKENtsQYFFITP | S1047 | 0.033 | 83.84 | 1101 | 0.026232 | 0.033 | 0.0262 | 1 |
| Q8N1B3 | CCNQ | MRsIPIAtACTIYHK | T50 | 0 | 98.17 | 248 | 0 | 0 | 0 | 0.9091 |
| P19823 | ITIH2 | ICPNCREtAVDGELV | T267 | 0.175 | 92.38 | 946 | 0.119527 | 0.175 | 0.1195 | 1 |
| P19823 | ITIH2 | DGELVVLyDVKREEK | Y277 | 0 | 98.14 | 946 | 0 | 0 | 0 | 0.8571 |
| Q9BWS9 | CHID1 | KVFGSkFtQIsPVWL | T104 | 0.206 | 97.91 | 393 | 0.140701 | 0.206 | 0.1407 | 1 |
| Q9BWS9 | CHID1 | GSkFtQIsPVWLQLk | S107 | 0 | 98.84 | 393 | 0 | 0 | 0 | 0.9 |
| Q13740 | ALCAM | QYDDVPEykDRLNLS | Y86 | 0.063 | 94.67 | 583 | 0.027637 | 0.063 | 0.0276 | 1 |
| Q9UDY8 | MALT1 | GTPEETGsYLVskDL | S656 | 0 | 88.92 | 824 | 0 | 0 | 0 | 1 |
| P32019 | INPP5B | VGIMLLLyVKQEHAA | Y432 | 0.071 | 98.69 | 993 | 0.031147 | 0.071 | 0.0311 | 1 |
| P32019 | INPP5B | NDELYRktLEEIVRS | T651 | 0.122 | 90.1 | 993 | 0.083328 | 0.122 | 0.0833 | 1 |
| P38606 | ATP6V1A | MRALDEyyDkHFTEF | Y465 | 0.022 | 93.82 | 617 | 0.009651 | 0.022 | 0.0097 | 0.9375 |
| P38606 | ATP6V1A | EAkIksDyAQLLEDM | Y601 | 0.149 | 92.89 | 617 | 0.065364 | 0.149 | 0.0654 | 1 |
| Q86SX6 | GLRX5 | LKKLGIHsALLDEKK | S145 | 0.013 | 96.21 | 157 | 0.010334 | 0.013 | 0.0103 | 1 |
| Q9ULV0 | MYO5B | HNDDFEMtsFWLsNT | T1546 | 0.006 | 93.18 | 1848 | 0.004098 | 0.006 | 0.0041 | 0.9167 |
| O15031 | PLXNB2 | EkIIDQVyRGQPCSC | Y1499 | 0.134 | 82.38 | 1838 | 0.058784 | 0.134 | 0.0588 | 1 |
| O15031 | PLXNB2 | RVNTLMHyNVRDGAT | Y1548 | 0.168 | 90.79 | 1838 | 0.073699 | 0.168 | 0.0737 | 1 |
| Q9P286 | PAK5 | HRDIKSDsILLTSDG | S573 | 0.002 | 96.43 | 719 | 0.00159 | 0.002 | 0.0016 | 1 |
| Q96EK7 | FAM120B | SLLVRGLtTLVLVNs | T781 | 0.03 | 97.18 | 910 | 0.02049 | 0.212 | 0.1448 | 1 |

| | | | | | | | | | | |
|---|---|---|---|---|---|---|---|---|---|---|
| Q96L93 | KIF16B | FFSVMLQsAtsPLHI | S1278 | 0.019 | 86.37 | 1317 | 0.015103 | 0.029 | 0.1128 | 1 |
| Q96L93 | KIF16B | VMLQsAtsPLHINKV | S1281 | 0 | 85.77 | 1317 | 0 | 0 | 0 | 1 |
| P29375 | KDM5A | IRPLAEktGICkIRP | T46 | 0.109 | 96.25 | 1690 | 0.074448 | 0.217 | 0.1485 | 1 |
| P29375 | KDM5A | GSLLksHyERILYPY | Y162 | 0.001 | 90.56 | 1690 | 0.000439 | 0.001 | 0.0004 | 1 |
| Q9C0H9 | SRCIN1 | GMLKSPNtAILIKDE | T228 | 0.163 | 90.96 | 1183 | 0.111331 | 0.163 | 0.1113 | 0.8889 |
| P52566 | ARHGDIB | GLkYVQHtyRTGVkV | T129 | 0.147 | 95.73 | 201 | 0.100403 | 0.147 | 0.1004 | 1 |
| P52566 | ARHGDIB | YHNkSFFtDDDKQDH | T179 | 0.167 | 97.3 | 201 | 0.114063 | 0.167 | 0.1141 | 1 |
| Q15149 | PLEC | NVQIALDyLRHRQVk | Y246 | 0.3 | 95.415 | 4684 | 0.131606 | 0.3275 | 0.1437 | 1 |
| Q15149 | PLEC | KEkLLLWsQRMVEGy | S305 | 0.005 | 93.21 | 4684 | 0.003975 | 0.005 | 0.004 | 1 |
| Q9UNE7 | STUB1 | YPEAAACyGRAITRN | Y49 | 0.05 | 96.92 | 303 | 0.021934 | 0.05 | 0.0219 | 1 |
| O60883 | GPR37L1 | LGLINQFstFFKGAI | S404 | 0.173 | 88.23 | 481 | 0.13752 | 0.173 | 0.1375 | 0.875 |
| Q6EMB2 | TTLL5 | FKFDVRLyVLVTSyD | Y227 | 0.032 | 98.17 | 1281 | 0.014038 | 0.032 | 0.014 | 0.9375 |
| P20132 | SDS | KMDSAQPsGSFkIRG | S37 | 0.072 | 97.17 | 328 | 0.057234 | 0.072 | 0.0572 | 1 |
| O14511 | NRG2 | PLERNQRyIFFLEPT | Y194 | 0.195 | 88.56 | 850 | 0.085544 | 0.195 | 0.0855 | 0.9 |
| Q9H4M9 | EHD1 | AEGLRQLyAQkLLPL | Y29 | 0.053 | 97.3 | 534 | 0.02325 | 0.053 | 0.0232 | 1 |
| Q96SE0 | ABHD1 | IRQFDERytsVAFGY | Y291 | 0.031 | 98.14 | 405 | 0.013599 | 0.031 | 0.0136 | 1 |
| Q96SE0 | ABHD1 | RQFDERytsVAFGYQ | T292 | 0.038 | 97.59 | 405 | 0.025955 | 0.038 | 0.026 | 0.8889 |
| Q96SE0 | ABHD1 | QFDERytsVAFGYQD | S293 | 0.029 | 97.32 | 405 | 0.023053 | 0.029 | 0.0231 | 1 |
| Q96SE0 | ABHD1 | GYQDCVTyyKAASPR | Y304 | 0.113 | 98.23 | 405 | 0.049571 | 0.113 | 0.0496 | 0.8571 |
| Q96SE0 | ABHD1 | YQDCVTyyKAASPRT | Y305 | 0.065 | 98.18 | 405 | 0.028515 | 0.065 | 0.0285 | 0.8333 |
| P35052 | GPC1 | CGEVRQIyGAKGFSL | Y39 | 0.037 | 95.81 | 558 | 0.016231 | 0.068 | 0.0298 | 1 |
| P35052 | GPC1 | ARAFRDLySELRLYY | Y145 | 0.006 | 97.84 | 558 | 0.002632 | 0.006 | 0.0026 | 1 |
| P62750 | RPL23A | kkIEDNNtLVFIVDV | T95 | 0.148 | 97.4 | 156 | 0.101086 | 0.148 | 0.1011 | 1 |
| Q9HBJ7 | USP29 | RHTFSRLsRVLIIHL | S504 | 0.031 | 97.24 | 922 | 0.024642 | 0.031 | 0.0246 | 0.8 |
| Q13472 | TOP3A | kGIADLLsNGRMRRR | S54 | 0.016 | 97.06 | 1001 | 0.012719 | 0.016 | 0.0127 | 1 |
| P07948 | LYN | EGFIPsNyVAKLNTL | Y117 | 0.109 | 92.87 | 512 | 0.047817 | 0.109 | 0.0478 | 1 |
| P07948 | LYN | GAFLIREsETLKGsF | S158 | 0.15 | 91.29 | 512 | 0.119237 | 0.168 | 0.1335 | 1 |
| P07948 | LYN | ETLKGsFsLSVRDFD | S166 | 0 | 95.94 | 512 | 0 | 0.096 | 0.0763 | 1 |
| P07948 | LYN | AWEIPREsIKLVKRL | S246 | 0.146 | 95.11 | 512 | 0.116058 | 0.146 | 0.1161 | 1 |
| P07948 | LYN | VTREEPIyIItEyMA | Y316 | 0.242 | 94.98 | 512 | 0.106162 | 0.242 | 0.1062 | 1 |
| Q6UVK1 | CSPG4 | VEQGRVRyLSTDPQH | Y743 | 0.006 | 92.28 | 2322 | 0.002632 | 0.294 | 0.108 | 1 |

| | | | | | | | | | | |
|---|---|---|---|---|---|---|---|---|---|---|
| Q86UT6 | NLRX1 | KYMLPQAsILVTTRP | S285 | 0.046 | 96.79 | 975 | 0.036566 | 0.046 | 0.0366 | 0.8333 |
| P00491 | PNP | MRQRALstWkQMGEQ | T177 | 0.084 | 93.06 | 289 | 0.057373 | 0.084 | 0.0574 | 1 |
| P00491 | PNP | VFGFSLItNKVIMDy | T242 | 0.064 | 95.99 | 289 | 0.043713 | 0.064 | 0.0437 | 0.9091 |
| Q9BXR5 | TLR10 | IQQLDLKtFEFNKEL | T91 | 0.106 | 94.36 | 811 | 0.072399 | 0.212 | 0.1448 | 1 |
| Q9BXR5 | TLR10 | LPHyEEGsLPILNTT | S184 | 0 | 95.07 | 811 | 0 | 0 | 0 | 1 |
| Q9BXR5 | TLR10 | YIQQDKIyLLLTKMD | Y320 | 0.05 | 87.75 | 811 | 0.021934 | 0.05 | 0.0219 | 1 |
| P49354 | FNTA | VISNttGyNDrAVLE | Y250 | 0.209 | 96.59 | 379 | 0.091685 | 0.209 | 0.0917 | 1 |
| Q15375 | EPHA7 | NQNNWLRtNWIsKGN | T87 | 0.003 | 94.46 | 998 | 0.002049 | 0.003 | 0.002 | 1 |
| Q15375 | EPHA7 | ALVSVKVyyKKCWSI | Y201 | 0.116 | 94.17 | 998 | 0.050888 | 0.116 | 0.0509 | 1 |
| Q15375 | EPHA7 | LVSVKVyyKKCWSII | Y202 | 0.27 | 93.05 | 998 | 0.118445 | 0.27 | 0.1184 | 0.8462 |
| Q15842 | KCNJ8 | RAETLIFsRHAVIAV | S194 | 0.023 | 96.35 | 424 | 0.018283 | 0.042 | 0.0334 | 1 |
| Q15842 | KCNJ8 | CHVIDKRsPLyDIsA | S275 | 0.088 | 90.72 | 424 | 0.069952 | 0.088 | 0.07 | 1 |
| Q15842 | KCNJ8 | IDKRsPLyDIsAtDL | Y278 | 0.266 | 92.88 | 424 | 0.11669 | 0.266 | 0.1167 | 1 |
| Q15842 | KCNJ8 | RsPLyDIsAtDLANQ | S281 | 0.13 | 92.09 | 424 | 0.103339 | 0.13 | 0.1033 | 1 |
| Q9NQZ7 | ENTPD7 | PVKKHKEtPLyILCt | T167 | 0.006 | 97.85 | 604 | 0.004098 | 0.006 | 0.0041 | 1 |
| Q9NQZ7 | ENTPD7 | tPLyILCtAGMRLLP | T174 | 0.083 | 96.14 | 604 | 0.05669 | 0.152 | 0.1038 | 0.875 |
| Q5VYK3 | ECPAS | VRALSINtLVKISkS | T1277 | 0.005 | 90.97 | 1845 | 0.003415 | 0.005 | 0.0034 | 0.9091 |
| Q5VYK3 | ECPAS | EEPIyKTsCALtIHA | S1458 | 0.001 | 89.63 | 1845 | 0.000795 | 0.001 | 0.0008 | 1 |
| Q5VYK3 | ECPAS | yKTsCALtIHAIGRY | T1462 | 0 | 93.7 | 1845 | 0 | 0 | 0 | 0.8333 |
| Q9H227 | GBA3 | PARPRVPyTSAKEYA | Y450 | 0 | 98.74 | 469 | 0 | 0 | 0 | 1 |
| Q6R327 | RICTOR | TDFHYRHsPDTAEGQ | S265 | 0.184 | 75.59 | 1708 | 0.146264 | 0.184 | 0.1463 | 1 |
| Q6R327 | RICTOR | REARFLAsKMGIIAT | S284 | 0.009 | 94.07 | 1708 | 0.007154 | 0.009 | 0.0072 | 1 |
| Q6R327 | RICTOR | DHISVRAtILLGELL | T424 | 0.075 | 94.99 | 1708 | 0.051226 | 0.075 | 0.0512 | 1 |
| Q16647 | PTGIS | QAQDRVHsADVFHTF | S200 | 0 | 98.35 | 500 | 0 | 0 | 0 | 1 |
| Q16647 | PTGIS | PQRDPEIytDPEVFK | Y398 | 0.034 | 98.22 | 500 | 0.014915 | 0.034 | 0.0149 | 1 |
| O95671 | ASMTL | SFAtPyGyAMEtAkQ | Y57 | 0.014 | 95.69 | 621 | 0.006142 | 0.014 | 0.0061 | 0.9231 |
| O95671 | ASMTL | PyGyAMEtAkQkALE | T61 | 0.021 | 96.34 | 621 | 0.014343 | 0.021 | 0.0143 | 1 |
| Q2VWP7 | PRTG | SVWTSHRtPKAtSVK | T610 | 0.025 | 89.48 | 1150 | 0.017075 | 0.025 | 0.0171 | 1 |
| Q13751 | LAMB3 | AGMLIERsSDFGkTW | S136 | 0.01 | 95.79 | 1172 | 0.007949 | 0.01 | 0.0079 | 1 |
| P43243 | MATR3 | HCSSLPHyQkLkKFL | Y827 | 0.283 | 89.48 | 847 | 0.124148 | 0.283 | 0.1241 | 1 |
| O60894 | RAMP1 | FLAVHGRyFRsCPIs | Y100 | 0.233 | 98.36 | 148 | 0.102214 | 0.233 | 0.1022 | 1 |

| | | | | | | | | | | |
|---|---|---|---|---|---|---|---|---|---|---|
| O75116 | ROCK2 | PRIFQILyANEGEsK | Y1232 | 0.07 | 87.17 | 1388 | 0.030708 | 0.07 | 0.0307 | 0.8333 |
| Q9UG01 | IFT172 | WMKAAELsIKFLPPQ | S1322 | 0 | 83.89 | 1749 | 0 | 0 | 0 | 1 |
| Q5TEJ8 | THEMIS2 | EMSDSRRysLADLTA | Y440 | 0.169 | 94.84 | 643 | 0.074138 | 0.169 | 0.0741 | 1 |
| O94906 | PRPF6 | AVRHLPQsVRIYIRA | S372 | 0.016 | 90.22 | 941 | 0.012719 | 0.016 | 0.0127 | 1 |
| O95433 | AHSA1 | TLkEtFLtsPEELyR | T217 | 0.148 | 94.12 | 338 | 0.101086 | 0.148 | 0.1011 | 0.8 |
| P40937 | RFC5 | FGKVtEEtVytCTGH | T234 | 0.21 | 95.97 | 340 | 0.143433 | 0.21 | 0.1434 | 1 |
| P40937 | RFC5 | KVtEEtVytCTGHPL | Y236 | 0.283 | 93.27 | 340 | 0.124148 | 0.283 | 0.1241 | 1 |
| Q6U841 | SLC4A10 | FLYMGAssLKGIQFF | S949 | 0.038 | 85.05 | 1118 | 0.030207 | 0.038 | 0.0302 | 1 |
| Q8IY67 | RAVER1 | AEYMKKDsAARAkSD | S185 | 0.107 | 98.15 | 606 | 0.085056 | 0.107 | 0.0851 | 1 |
| Q9UBC5 | MYO1A | GYAHRQGyGPFLERY | Y631 | 0.129 | 93.07 | 1043 | 0.05659 | 0.167 | 0.0733 | 1 |
| Q8IVW8 | SPNS2 | WHWALRVsPVLGMIT | S266 | 0.053 | 92.84 | 549 | 0.04213 | 0.053 | 0.0421 | 1 |
| Q8TEQ8 | PIGO | RRYGNLKsPEPPMLF | S696 | 0 | 80.6 | 1089 | 0 | 0.07 | 0.0592 | 1 |
| Q86WI3 | NLRC5 | MLKRLDLsHLLLNsS | S1558 | 0.078 | 88.42 | 1866 | 0.062003 | 0.078 | 0.062 | 0.875 |
| Q7LDG7 | RASGRP2 | QEGNRRHssLIDIDS | S116 | 0.097 | 85.08 | 609 | 0.077107 | 0.097 | 0.0771 | 1 |
| Q99551 | MTERF1 | IASIISRyPRAITRt | Y128 | 0.154 | 98.69 | 399 | 0.067558 | 0.154 | 0.0676 | 1 |
| Q99551 | MTERF1 | ENPRVLDsSISTLkS | S354 | 0.132 | 97.96 | 399 | 0.104929 | 0.132 | 0.1049 | 1 |
| Q9H223 | EHD4 | FLNRFMCsQLPNQVL | S142 | 0 | 97.17 | 541 | 0 | 0 | 0 | 1 |
| Q6ZQY3 | GADL1 | SLVARFMtEALNPSV | T136 | 0.093 | 98.73 | 521 | 0.06352 | 0.093 | 0.0635 | 1 |
| Q6ZQY3 | GADL1 | SVYTYEVsPVFLLVE | S149 | 0.013 | 98.37 | 521 | 0.010334 | 0.013 | 0.0103 | 1 |
| Q6ZQY3 | GADL1 | VNRALALsRYLVDEI | S416 | 0.011 | 98.69 | 521 | 0.008744 | 0.011 | 0.0087 | 1 |
| Q6ZQY3 | GADL1 | KLLMEPEyANICFWY | Y437 | 0.047 | 98.36 | 521 | 0.020618 | 0.047 | 0.0206 | 1 |
| Q6ZQY3 | GADL1 | ERMMKKGsLMLGYQP | S478 | 0.177 | 98.29 | 521 | 0.1407 | 0.177 | 0.1407 | 1 |
| Q3MHD2 | LSM12 | FSVGSQVsCRTCQEQ | S16 | 0.024 | 95.79 | 195 | 0.019078 | 0.024 | 0.0191 | 1 |
| Q6VUC0 | TFAP2E | DLMAQDRsPLGNsRP | S364 | 0.126 | 94.61 | 442 | 0.100159 | 0.126 | 0.1002 | 1 |
| Q86TI0 | TBC1D1 | EGMNSsKtKLELQKH | T410 | 0.046 | 88.74 | 1168 | 0.031419 | 0.046 | 0.0314 | 1 |
| Q8WV93 | AFG1L | PAAGKLyyLTSEADV | Y296 | 0.146 | 95.41 | 481 | 0.064048 | 0.146 | 0.064 | 0.9 |
| Q9NR80 | ARHGEF4 | ssKDQMRtNVINEIL | T286 | 0.152 | 97.13 | 690 | 0.103818 | 0.152 | 0.1038 | 1 |
| P20073 | ANXA7 | AGDTsGDyRRLLLAI | Y478 | 0.182 | 97.94 | 488 | 0.079841 | 0.182 | 0.0798 | 0.9231 |
| Q6P2E9 | EDC4 | EIVASsDssISskAR | S106 | 0 | 90.14 | 1401 | 0 | 0 | 0 | 1 |
| Q9NVI1 | FANCI | EDADVSVtQRTAFQI | T952 | 0.167 | 90.92 | 1328 | 0.114063 | 0.167 | 0.1141 | 1 |
| Q9NVI1 | FANCI | SKLLEPSsPQFVQML | S1000 | 0.094 | 93.39 | 1328 | 0.074722 | 0.094 | 0.0747 | 1 |

| | | | | | | | | | | |
|---|---|---|---|---|---|---|---|---|---|---|
| P35244 | RPA3 | tGkMFILsDGEGKNG | S44 | 0.094 | 98.7 | 121 | 0.074722 | 0.094 | 0.0747 | 1 |
| Q13042 | CDC16 | SAIGYIHsLMGNFEN | S490 | 0.182 | 94.28 | 620 | 0.144674 | 0.182 | 0.1447 | 1 |
| Q9Y4G6 | TLN2 | LLGItKDsVMRVDEK | S339 | 0.083 | 83.03 | 2542 | 0.065978 | 0.083 | 0.066 | 1 |
| Q96BY6 | DOCK10 | IASGAEPyIkNPDSN | Y526 | 0.087 | 86.79 | 2186 | 0.038166 | 0.087 | 0.0382 | 1 |
| P20594 | NPR2 | ENLTNGDyVFFYLDV | Y250 | 0.184 | 92.46 | 1047 | 0.080718 | 0.184 | 0.0807 | 1 |
| Q9ULA0 | DNPEP | EPHVRMVtLYDNEEV | T306 | 0.001 | 98.95 | 485 | 0.000683 | 0.001 | 0.0007 | 1 |
| Q9ULA0 | DNPEP | NSKQRYAsNAVSEAL | S393 | 0.067 | 98.8 | 485 | 0.053259 | 0.091 | 0.0723 | 0.8 |
| Q9ULA0 | DNPEP | LRVLDLGsPQLAMHS | S444 | 0.044 | 98.88 | 485 | 0.034976 | 0.044 | 0.035 | 1 |
| Q9Y478 | PRKAB1 | GVMVLSAtHRYKKKy | T254 | 0.037 | 98.14 | 270 | 0.025272 | 0.071 | 0.0482 | 1 |
| Q3T906 | GNPTAB | GDITLKGyNLsKsAL | Y728 | 0.303 | 89.02 | 1256 | 0.132922 | 0.303 | 0.1329 | 0.875 |
| Q8NA19 | L3MBTL4 | DKFVWMDyLKACKLQ | Y165 | 0.208 | 91.32 | 623 | 0.091247 | 0.208 | 0.0912 | 1 |
| P21333 | FLNA | REATTEFsVDARALt | S1279 | 0.151 | 89.73 | 2647 | 0.120032 | 0.151 | 0.12 | 1 |
| Q15413 | RYR3 | sQHRRIFyEAGGAGT | Y251 | 0.035 | 90.915 | 4870 | 0.015354 | 0.035 | 0.0154 | 1 |
| Q15413 | RYR3 | RHLTTGHyLALTEDQ | Y294 | 0.055 | 94.65 | 4870 | 0.024128 | 0.055 | 0.0241 | 0.8462 |
| P10745 | RBP3 | QRGVYLLtSHRTATA | T541 | 0.002 | 98.33 | 1247 | 0.001366 | 0.002 | 0.0014 | 1 |
| Q9H3P7 | ACBD3 | QYPGRGVyLLkFDNS | Y504 | 0.006 | 98.13 | 528 | 0.002632 | 0.006 | 0.0026 | 1 |
| Q9H3P7 | ACBD3 | LWRSKSVyYRVYYTR | Y521 | 0.173 | 98.44 | 528 | 0.075893 | 0.173 | 0.0759 | 1 |
| P00387 | CYB5R3 | VGQHIyLsARIDGNL | S82 | 0.1 | 98.47 | 301 | 0.079491 | 0.139 | 0.1105 | 1 |
| P00387 | CYB5R3 | PIIRtVksVGMIAGG | S174 | 0.061 | 98.76 | 301 | 0.04849 | 0.061 | 0.0485 | 1 |
| O95251 | KAT7 | NFNMkCPtPGCNsLG | T187 | 0.065 | 81.9 | 611 | 0.044396 | 0.188 | 0.1284 | 1 |
| O95251 | KAT7 | CPtPGCNsLGHLTGk | S192 | 0.016 | 87.27 | 611 | 0.012719 | 0.016 | 0.0127 | 1 |
| Q9H7B4 | SMYD3 | PGELLFRsDPLAYTV | S33 | 0.036 | 98.52 | 428 | 0.028617 | 0.036 | 0.0286 | 0.9 |
| Q9BZZ5 | API5 | VGQHKDAyQVILDGV | Y28 | 0.003 | 97.88 | 524 | 0.001316 | 0.003 | 0.0013 | 1 |
| Q8TCG2 | PI4K2B | ERLVILDyIIRNTDR | Y298 | 0.075 | 98.67 | 481 | 0.032901 | 0.075 | 0.0329 | 1 |
| O94898 | LRIG2 | ISEIKTSsFPRMQLK | S163 | 0.028 | 94.88 | 1065 | 0.022258 | 0.028 | 0.0223 | 1 |
| O94898 | LRIG2 | QLKyLNLsNNRITTL | S175 | 0.162 | 98.38 | 1065 | 0.128776 | 0.162 | 0.1288 | 1 |
| O94898 | LRIG2 | MLQQLYVsQNAIERI | S295 | 0.071 | 98.17 | 1065 | 0.056439 | 0.071 | 0.0564 | 1 |
| O95758 | PTBP3 | DRPPCsPsRVLHLRk | S58 | 0.046 | 80.74 | 552 | 0.036566 | 0.046 | 0.0366 | 1 |
| O95758 | PTBP3 | QPVyIQysNHRELkt | S131 | 0.086 | 81.76 | 552 | 0.068363 | 0.086 | 0.0684 | 1 |
| Q9NV35 | NUDT15 | KGSVGAGsFQLPGGH | S42 | 0.121 | 96.83 | 164 | 0.096185 | 0.121 | 0.0962 | 1 |
| Q8N3C0 | ASCC3 | SkSEILRtQMLVTtP | T578 | 0.041 | 88.07 | 2202 | 0.028004 | 0.1 | 0.0682 | 1 |

| | | | | | | | | | | |
|---|---|---|---|---|---|---|---|---|---|---|
| Q8N3C0 | ASCC3 | RtQMLVTtPEkWDVV | T584 | 0.12 | 89.71 | 2202 | 0.081962 | 0.12 | 0.082 | 1 |
| P55290 | CDH13 | IGHVMEGsPTGTTVM | S258 | 0.151 | 94.72 | 713 | 0.120032 | 0.151 | 0.12 | 1 |
| O00339 | MATN2 | TRVGLLQyGSTVKNE | Y101 | 0.003 | 92.45 | 956 | 0.001316 | 0.003 | 0.0013 | 1 |
| O00339 | MATN2 | MRHLSTGtMtGLAIQ | T134 | 0.13 | 82.29 | 956 | 0.088792 | 0.13 | 0.0888 | 1 |
| O00339 | MATN2 | HLSTGtMtGLAIQYA | T136 | 0.004 | 88.79 | 956 | 0.002732 | 0.004 | 0.0027 | 1 |
| O00339 | MATN2 | PRVIMIVtDGRPQDS | T169 | 0.048 | 92.3 | 956 | 0.032785 | 0.048 | 0.0328 | 1 |
| Q8TF21 | ANKRD24 | CEGASPEtVEVLLQG | T229 | 0 | 97.07 | 1146 | 0 | 0 | 0 | 1 |
| Q8TF21 | ANKRD24 | LGQDAAHyGALAGDK | Y253 | 0.202 | 95.12 | 1146 | 0.088614 | 0.202 | 0.0886 | 1 |
| Q9Y5B9 | SUPT16H | KkAAsITsEVFNkFF | S191 | 0.153 | 97.56 | 1047 | 0.121622 | 0.153 | 0.1216 | 1 |
| Q96N87 | SLC6A18 | LFFGMLFtLGLsTMF | T412 | 0.069 | 97.66 | 628 | 0.047128 | 0.069 | 0.0471 | 1 |
| Q96N87 | SLC6A18 | MLFtLGLsTMFGTVE | S416 | 0.025 | 96.74 | 628 | 0.019873 | 0.025 | 0.0199 | 1 |
| P61978 | HNRNPK | AQyLLQNsVkQySGk | S454 | 0.011 | 85.86 | 463 | 0.008744 | 0.011 | 0.0087 | 1 |
| O75879 | GATB | FQNVIKEtRAEPKKV | T415 | 0.174 | 94.4 | 557 | 0.118844 | 0.174 | 0.1188 | 1 |
| O75879 | GATB | IGLVRKAtQSRADPV | T538 | 0.058 | 83.21 | 557 | 0.039615 | 0.058 | 0.0396 | 1 |
| Q9UBZ9 | REV1 | DQTKCAAsVGIGSNI | S607 | 0 | 98.59 | 1251 | 0 | 0 | 0 | 0.9091 |
| Q9BXF6 | RAB11FIP5 | VLTTMHRsLIGVDKF | S116 | 0.179 | 91.59 | 653 | 0.14229 | 0.179 | 0.1423 | 1 |
| Q658N2 | WSCD1 | REIEMFDsAILLIRN | S423 | 0.058 | 98.02 | 575 | 0.046105 | 0.058 | 0.0461 | 1 |
| Q658N2 | WSCD1 | KRLLVVHyEELRRSL | Y490 | 0.177 | 96.37 | 575 | 0.077647 | 0.177 | 0.0776 | 1 |
| P0C7P3 | SLFN14 | FANtQGGyVLIGVDD | Y242 | 0.156 | 95.9 | 912 | 0.068435 | 0.156 | 0.0684 | 0.9231 |
| O95084 | PRSS23 | VKLSTGCtGTLVAEK | T161 | 0 | 97 | 383 | 0 | 0 | 0 | 0.8667 |
| Q8NEU8 | APPL2 | LEEALQDsPQtRSLL | S16 | 0.16 | 97.19 | 664 | 0.127186 | 0.16 | 0.1272 | 1 |
| Q8NEU8 | APPL2 | ALQDsPQtRSLLSVF | T19 | 0.17 | 96.07 | 664 | 0.116112 | 0.17 | 0.1161 | 1 |
| Q8NEU8 | APPL2 | GIILQAEsRkENEEW | S359 | 0.095 | 96.67 | 664 | 0.075517 | 0.095 | 0.0755 | 1 |
| P54762 | EPHB1 | IYTEMRFtVRDCSSL | T92 | 0.038 | 89.71 | 984 | 0.025955 | 0.105 | 0.0922 | 1 |
| Q06278 | AOX1 | ISFLFKFyLEVsQIL | Y527 | 0.293 | 98.15 | 1338 | 0.128535 | 0.293 | 0.1285 | 1 |
| Q06278 | AOX1 | KKMDPVHyPSLADKy | Y542 | 0.307 | 90.57 | 1338 | 0.134676 | 0.307 | 0.1347 | 1 |
| P19623 | SRM | LDGVIQCtERDEFSY | T72 | 0.005 | 98.78 | 302 | 0.003415 | 0.005 | 0.0034 | 1 |
| Q9ULL8 | SHROOM4 | ALILIKGsFRILKLI | S80 | 0.08 | 89.63 | 1493 | 0.063593 | 0.08 | 0.0636 | 0.8333 |
| Q9Y6K1 | DNMT3A | RKPIRVLsLFDGIAt | S638 | 0.02 | 98.65 | 912 | 0.015898 | 0.02 | 0.0159 | 0.9 |
| Q9Y6K1 | DNMT3A | RLLGRSWsVPVIRHL | S894 | 0.088 | 97.45 | 912 | 0.069952 | 0.088 | 0.07 | 1 |
| Q3SY69 | ALDH1L2 | IDSPKHGsIIyHPSI | S124 | 0.035 | 96.26 | 923 | 0.027822 | 0.035 | 0.0278 | 1 |

| | | | | | | | | | | |
|---|---|---|---|---|---|---|---|---|---|---|
| Q3SY69 | ALDH1L2 | PKHGsIIyHPSILPR | Y127 | 0.218 | 96.15 | 923 | 0.095633 | 0.218 | 0.0956 | 0.9412 |
| Q29RF7 | PDS5A | LADIFRIyAPEAPyt | Y101 | 0.298 | 91.19 | 1337 | 0.130728 | 0.298 | 0.1307 | 1 |
| Q29RF7 | PDS5A | IyAPEAPytsHDKLK | Y107 | 0.126 | 90.68 | 1337 | 0.055274 | 0.126 | 0.0553 | 1 |
| Q29RF7 | PDS5A | MHMLDLMssIIMEGD | S192 | 0.031 | 96 | 1337 | 0.024642 | 0.031 | 0.0246 | 1 |
| Q29RF7 | PDS5A | HMLDLMssIIMEGDG | S193 | 0.054 | 95.34 | 1337 | 0.042925 | 0.054 | 0.0429 | 1 |
| Q29RF7 | PDS5A | MMGLAQLykkYCLHG | Y426 | 0.234 | 97.65 | 1337 | 0.102652 | 0.234 | 0.1027 | 1 |
| Q8IY82 | DRC7 | QALRVHsyKSMQPEM | Y516 | 0.049 | 96.43 | 874 | 0.021496 | 0.113 | 0.0496 | 1 |
| Q8IY82 | DRC7 | PRTMTEyyQGRPDFL | Y552 | 0.141 | 96.99 | 874 | 0.061855 | 0.21 | 0.0921 | 1 |
| Q08345 | DDR1 | DDPLCMItDyMENGD | T701 | 0.046 | 93.05 | 913 | 0.031419 | 0.181 | 0.1236 | 0.875 |
| P15090 | FABP4 | GVTSTRVyERA_____ | Y129 | 0.119 | 98.18 | 132 | 0.052204 | 0.119 | 0.0522 | 1 |
| Q9BQ24 | ZFYVE21 | KKLVRsPsGLRMVPE | S19 | 0.131 | 90.56 | 234 | 0.104134 | 0.131 | 0.1041 | 1 |
| P03952 | KLKB1 | SSTPQENtISGYSLL | T275 | 0.003 | 97 | 638 | 0.002049 | 0.003 | 0.002 | 1 |
| P33908 | MAN1A1 | CHESYNRtFMkLGPE | T515 | 0.114 | 98.38 | 653 | 0.077864 | 0.114 | 0.0779 | 1 |
| Q9NQN1 | OR2S2 | NGVLILVtILDsRLH | T50 | 0.18 | 91.88 | 319 | 0.122942 | 0.18 | 0.1229 | 1 |
| O14773 | TPP1 | VSGTSAStPVFGGIL | T478 | 0 | 98.67 | 563 | 0 | 0 | 0 | 0.8571 |
| Q16280 | CNGA2 | GDIGKEMyIIKEGKL | Y492 | 0.136 | 92.13 | 664 | 0.059661 | 0.136 | 0.0597 | 1 |
| Q16280 | CNGA2 | NIRSLGYsDLFCLsK | S546 | 0 | 92.97 | 664 | 0 | 0 | 0 | 1 |
| Q16280 | CNGA2 | YsDLFCLsKDDLMEA | S552 | 0.169 | 91.29 | 664 | 0.134341 | 0.169 | 0.1343 | 1 |
| P32970 | CD70 | GGPALGRsFLHGPEL | S84 | 0.046 | 94.21 | 193 | 0.036566 | 0.046 | 0.0366 | 1 |
| Q53EQ6 | TIGD5 | RWRGDRVtVLLAANL | T276 | 0.057 | 92.2 | 642 | 0.038932 | 0.057 | 0.0389 | 1 |
| P27216 | ANXA13 | IIAIKEAyQRLFDRS | Y128 | 0.042 | 98.82 | 316 | 0.018425 | 0.042 | 0.0184 | 1 |
| P35222 | CTNNB1 | LFVQLLYsPIENIQR | S605 | 0.172 | 95.49 | 781 | 0.136725 | 0.172 | 0.1367 | 1 |
| Q15392 | DHCR24 | FQHICtAyELVLADG | Y190 | 0.013 | 98.81 | 516 | 0.005703 | 0.013 | 0.0057 | 0.8 |
| Q15392 | DHCR24 | TNREGLEyIPLRHYY | Y321 | 0.134 | 98.75 | 516 | 0.058784 | 0.134 | 0.0588 | 1 |
| Q15392 | DHCR24 | QDAFPEVyDkICkAA | Y507 | 0.299 | 96.27 | 516 | 0.131167 | 0.299 | 0.1312 | 1 |
| Q13045 | FLII | EDNQGIVyAWVGRAS | Y1102 | 0.031 | 94.98 | 1269 | 0.013599 | 0.031 | 0.0136 | 1 |
| Q13045 | FLII | SLKACQVyIQHMRSK | Y1227 | 0.046 | 86.36 | 1269 | 0.02018 | 0.38 | 0.1443 | 1 |
| P42357 | HAL | NVLAKGysGIsLETL | S223 | 0.042 | 97.08 | 657 | 0.033386 | 0.042 | 0.0334 | 1 |
| P42357 | HAL | AKGysGIsLETLKQV | S226 | 0.083 | 98.25 | 657 | 0.065978 | 0.083 | 0.066 | 1 |
| P42357 | HAL | ADIVAALtLEVLKGT | T337 | 0 | 98.65 | 657 | 0 | 0 | 0 | 1 |
| P42357 | HAL | VHGVVNDtIAFVKNI | T410 | 0.055 | 98.8 | 657 | 0.037566 | 0.055 | 0.0376 | 0.8 |

| | | | | | | | | | | |
|---|---|---|---|---|---|---|---|---|---|---|
| P42357 | HAL | IPESRPLsPTAFSLQ | S635 | 0.134 | 80.87 | 657 | 0.106519 | 0.165 | 0.1312 | 1 |
| P83436 | COG7 | VDRCVRFtNGLGTCG | T413 | 0.033 | 91.51 | 770 | 0.022539 | 0.115 | 0.0785 | 1 |
| Q92598 | HSPH1 | ARAGGIEtIANEFsD | T25 | 0.066 | 97.41 | 858 | 0.045079 | 0.066 | 0.0451 | 1 |
| Q92598 | HSPH1 | kEkENLSyDLVPLkN | Y89 | 0.071 | 95.73 | 858 | 0.031147 | 0.071 | 0.0311 | 1 |
| Q92598 | HSPH1 | ISVPSFFtDAERRSV | T149 | 0.021 | 96.64 | 858 | 0.014343 | 0.021 | 0.0143 | 1 |
| Q92598 | HSPH1 | ALQCAILsPAFkVRE | S384 | 0.076 | 88.99 | 858 | 0.060414 | 0.076 | 0.0604 | 1 |
| Q92598 | HSPH1 | PFELEAFySDPQGVP | Y446 | 0.118 | 97.18 | 858 | 0.051765 | 0.118 | 0.0518 | 1 |
| Q92598 | HSPH1 | GKDLLNMyIETEGKM | Y614 | 0.217 | 97.93 | 858 | 0.095195 | 0.217 | 0.0952 | 1 |
| Q92598 | HSPH1 | RDkLCGPyEkFICEQ | Y653 | 0.099 | 94.51 | 858 | 0.04343 | 0.189 | 0.0807 | 1 |
| Q92598 | HSPH1 | FLRLLtEtEDWLyEE | T672 | 0.009 | 96.8 | 858 | 0.006147 | 0.009 | 0.0061 | 1 |
| Q92598 | HSPH1 | EDQAKQAyVDkLEEL | Y688 | 0.006 | 97.78 | 858 | 0.002632 | 0.006 | 0.0026 | 1 |
| Q13635 | PTCH1 | QTMFQLMtPKQMYEH | T368 | 0.008 | 91.21 | 1447 | 0.005464 | 0.075 | 0.0512 | 1 |
| Q9H0M0 | WWP1 | RQtLFEDsFQQIMAL | S580 | 0 | 98.28 | 922 | 0 | 0.042 | 0.0334 | 0.875 |
| Q10471 | GALNT2 | EDRTRVVsPIIDVIN | S250 | 0.06 | 98.84 | 571 | 0.047695 | 0.06 | 0.0477 | 1 |
| Q10471 | GALNT2 | LFVMDKFyFEELGKY | Y317 | 0.158 | 98.86 | 571 | 0.069312 | 0.158 | 0.0693 | 1 |
| Q10471 | GALNT2 | KWYLENVyPELRVPD | Y434 | 0.042 | 98.54 | 571 | 0.018425 | 0.042 | 0.0184 | 0.9091 |
| Q09328 | MGAT5 | RQNQSLVyGKVDSFW | Y452 | 0.085 | 94.62 | 741 | 0.037288 | 0.085 | 0.0373 | 1 |
| Q09328 | MGAT5 | PKFNPPKssKNtDFF | S544 | 0.053 | 95.08 | 741 | 0.04213 | 0.053 | 0.0421 | 1 |
| Q09328 | MGAT5 | PPKssKNtDFFIGkP | T548 | 0.213 | 94.4 | 741 | 0.145482 | 0.213 | 0.1455 | 1 |
| P98155 | VLDLR | AIAVDWVyKTIYWTD | Y524 | 0.229 | 93.47 | 873 | 0.100459 | 0.229 | 0.1005 | 1 |
| P50747 | HLCS | LYQKFMAyLsQGGKV | Y236 | 0.009 | 93.97 | 726 | 0.003948 | 0.063 | 0.0276 | 1 |
| Q8NEN0 | ARMC2 | GHLLVQVtATLRNLV | T452 | 0.011 | 98.71 | 867 | 0.007513 | 0.011 | 0.0075 | 1 |
| P51790 | CLCN3 | CCCGNIFsYLFPKYS | S297 | 0.011 | 93.05 | 818 | 0.008744 | 0.011 | 0.0087 | 1 |
| Q86TI0 | TBC1D1 | LYNILKAySLLDQEV | Y880 | 0.005 | 97.95 | 1168 | 0.002193 | 0.005 | 0.0022 | 1 |
| O14764 | GABRD | RVsLGIttVLTMTTL | T286 | 0.07 | 93.62 | 452 | 0.047811 | 0.193 | 0.1318 | 1 |
| P13639 | EEF2 | KANIRNMsVIAHVDH | S23 | 0.001 | 94.97 | 858 | 0.000795 | 0.001 | 0.0008 | 1 |
| P13639 | EEF2 | kkLWGDryFDPANGk | Y265 | 0.209 | 94.27 | 858 | 0.091685 | 0.209 | 0.0917 | 1 |
| P13639 | EEF2 | EGkKLPRtFCQLILD | T288 | 0 | 95.92 | 858 | 0 | 0 | 0 | 1 |
| P13639 | EEF2 | DALLQMItIHLPsPV | T355 | 0 | 95.5 | 858 | 0 | 0 | 0 | 1 |
| P13639 | EEF2 | kYRCELLyEGPPDDE | Y373 | 0.088 | 95.64 | 858 | 0.038604 | 0.088 | 0.0386 | 1 |
| P13639 | EEF2 | yAFGRVFsGLVSTGL | S418 | 0 | 97.96 | 858 | 0 | 0 | 0 | 1 |

| | | | | | | | | | | |
|---|---|---|---|---|---|---|---|---|---|---|
| P13639 | EEF2 | VrIMGPNytPGkkED | Y434 | 0.173 | 89.23 | 858 | 0.075893 | 0.173 | 0.0759 | 1 |
| P13639 | EEF2 | yLkPIQRtILMMGRY | T450 | 0.001 | 95.42 | 858 | 0.000683 | 0.001 | 0.0007 | 1 |
| O43707 | ACTN4 | EVENQILtRDAkGIs | T756 | 0 | 79.09 | 911 | 0 | 0.167 | 0.1141 | 1 |
| Q3KNS1 | PTCHD3 | KIEVVHFtsLsRQLE | T368 | 0.021 | 92.59 | 767 | 0.014343 | 0.155 | 0.1059 | 1 |
| Q3KNS1 | PTCHD3 | MISAWHKtNLADDIR | T469 | 0.099 | 86.99 | 767 | 0.067618 | 0.099 | 0.0676 | 1 |
| O96013 | PAK4 | VIHRDIKsDsILLTH | S443 | 0.005 | 96.62 | 591 | 0.003975 | 0.005 | 0.004 | 1 |
| O96013 | PAK4 | HRDIKsDsILLTHDG | S445 | 0.002 | 96.48 | 591 | 0.00159 | 0.002 | 0.0016 | 1 |
| O96013 | PAK4 | ksLVGtPyWMAPELI | Y480 | 0.104 | 97.55 | 591 | 0.045623 | 0.238 | 0.1044 | 1 |
| Q9HCB6 | SPON1 | FYKPGTSyRVTLSAA | Y78 | 0.269 | 90.16 | 807 | 0.118006 | 0.269 | 0.118 | 1 |
| P08473 | MME | VSSLSRTyKESRNAF | Y387 | 0.079 | 98.77 | 750 | 0.034656 | 0.079 | 0.0347 | 1 |
| P08473 | MME | DNKLNNEyLELNyKE | Y497 | 0.052 | 98.24 | 750 | 0.022812 | 0.052 | 0.0228 | 1 |
| P08473 | MME | NEyLELNyKEDEyFE | Y502 | 0.333 | 98.72 | 750 | 0.146082 | 0.333 | 0.1461 | 1 |
| P08473 | MME | LNyKEDEyFENIIQN | Y507 | 0.196 | 98.78 | 750 | 0.085982 | 0.196 | 0.086 | 1 |
| P08473 | MME | SIKTDVHsPGNFRII | S713 | 0.05 | 98.78 | 750 | 0.039746 | 0.05 | 0.0397 | 1 |
| P25786 | PSMA1 | DNDVTVWsPQGRIHQ | S14 | 0.086 | 95.21 | 263 | 0.068363 | 0.086 | 0.0684 | 1 |
| P25786 | PSMA1 | SATVGLksktHAVLV | S40 | 0.024 | 98.48 | 263 | 0.019078 | 0.024 | 0.0191 | 0.8 |
| P25786 | PSMA1 | VsLIGsktQIPtQRY | T116 | 0.101 | 98.35 | 263 | 0.068984 | 0.101 | 0.069 | 0.8182 |
| P25786 | PSMA1 | yFDCRAMsIGARSQS | S160 | 0.008 | 98.73 | 263 | 0.006359 | 0.008 | 0.0064 | 0.8462 |
| Q9HAB3 | SLC52A2 | QSFSCLPyGRLAYHL | Y305 | 0.216 | 90.72 | 445 | 0.094756 | 0.216 | 0.0948 | 1 |
| Q9NW08 | POLR3B | RkGAVGAsVTSsTHE | S202 | 0.13 | 92.34 | 1133 | 0.103339 | 0.13 | 0.1033 | 1 |
| Q9NW08 | POLR3B | NRIDtLMyLLAyPQK | Y710 | 0.163 | 94.76 | 1133 | 0.071506 | 0.163 | 0.0715 | 0.8333 |
| Q9NW08 | POLR3B | PQKPMVKtkTIELIE | T722 | 0.025 | 94.09 | 1133 | 0.017075 | 0.025 | 0.0171 | 1 |
| Q96P11 | NSUN5 | AEHCLRAsPETTLSS | S408 | 0.116 | 90.69 | 429 | 0.09221 | 0.116 | 0.0922 | 1 |
| Q8NG97 | OR2Z1 | TPMYFLLsQLSLFDI | S64 | 0.153 | 95.81 | 314 | 0.121622 | 0.153 | 0.1216 | 1 |
| Q8N7M0 | DYNLT5 | KDLMIPRyKLIVIVH | Y126 | 0.308 | 94.77 | 179 | 0.135115 | 0.308 | 0.1351 | 1 |
| Q99720 | SIGMAR1 | YWAEISDtIISGTFH | T127 | 0.032 | 98.48 | 223 | 0.021856 | 0.032 | 0.0219 | 1 |
| Q14694 | USP10 | TkQEVEIsRRVtLEk | S670 | 0.172 | 89.3 | 798 | 0.136725 | 0.172 | 0.1367 | 1 |
| Q14694 | USP10 | VEIsRRVtLEkLPPV | T674 | 0.094 | 95.58 | 798 | 0.064203 | 0.094 | 0.0642 | 0.8 |
| Q14694 | USP10 | EIskELLsPGVkNKN | S718 | 0.01 | 92.7 | 798 | 0.007949 | 0.01 | 0.0079 | 0.8889 |
| Q8WWN8 | ARAP3 | kNQRLEKyKDVIGCL | Y1016 | 0 | 97.51 | 1544 | 0 | 0 | 0 | 0.9333 |
| Q05084 | ICA1 | TGKALCFsSQQRLAL | S123 | 0.027 | 95.92 | 483 | 0.021463 | 0.027 | 0.0215 | 1 |

| | | | | | | | | | | |
|---|---|---|---|---|---|---|---|---|---|---|
| O14980 | XPO1 | VPMFRNVsLKCLTEI | S264 | 0 | 98.15 | 1071 | 0 | 0 | 0 | 1 |
| O14980 | XPO1 | QLGsILktNVRACkA | T694 | 0.089 | 98.3 | 1071 | 0.060788 | 0.089 | 0.0608 | 1 |
| O14980 | XPO1 | AREPEVLsTMAIIVN | S802 | 0.156 | 97.15 | 1071 | 0.124007 | 0.156 | 0.124 | 1 |
| P46821 | MAP1B | DWMKNLIsPDLGVVF | S351 | 0.035 | 88.22 | 2468 | 0.027822 | 0.035 | 0.0278 | 1 |
| O43852 | CALU | ARHLVyEsDQNkDGK | S277 | 0.007 | 88.05 | 315 | 0.005564 | 0.007 | 0.0056 | 1 |
| Q13315 | ATM | RLIRDPEtIkHLDRH | T39 | 0.004 | 87.65 | 3056 | 0.002732 | 0.004 | 0.0027 | 0.875 |
| Q13464 | ROCK1 | SSKKILFyNDEQDKE | Y1153 | 0.048 | 93.29 | 1354 | 0.021057 | 0.146 | 0.0408 | 1 |
| O95631 | NTN1 | LKINMKKyCKKDYAV | Y490 | 0.003 | 91.16 | 604 | 0.001316 | 0.003 | 0.0013 | 1 |
| P36888 | FLT3 | HKHQPGEyIFHAEND | Y416 | 0.047 | 92.33 | 993 | 0.020618 | 0.047 | 0.0206 | 1 |
| Q8NEG7 | DENND6B | WHADDRHyNsRAPVA | Y115 | 0.081 | 91.02 | 585 | 0.035534 | 0.081 | 0.0355 | 1 |
| P12314 | FCGR1A | KLVyNVLyyRNGKAF | Y137 | 0.018 | 98.08 | 374 | 0.007896 | 0.018 | 0.0079 | 1 |
| Q12873 | CHD3 | TGGTLHMyQLEGLNW | Y739 | 0.169 | 90.91 | 2000 | 0.074138 | 0.169 | 0.0741 | 0.9 |
| Q12873 | CHD3 | QTIVFLYsLYKEGHT | S777 | 0 | 92.21 | 2000 | 0 | 0 | 0 | 1 |
| P35520 | CBS | LGAEIVrtPtNARFD | T191 | 0.029 | 90.05 | 551 | 0.019807 | 0.029 | 0.0198 | 1 |
| P35520 | CBS | LDQYRNAsNPLAHYD | S227 | 0.074 | 98.8 | 551 | 0.058824 | 0.074 | 0.0588 | 1 |
| Q7Z5S9 | TMEM144 | IIYIKDHsKRNDSIY | S214 | 0.144 | 93.44 | 345 | 0.114468 | 0.144 | 0.1145 | 0.9 |
| P33981 | TTK | LNkLQQHsDkIIRLY | S582 | 0.173 | 96.92 | 857 | 0.13752 | 0.173 | 0.1375 | 1 |
| Q86SQ9 | DHDDS | LNVCFAYtsRHEIsN | T157 | 0.049 | 98.77 | 333 | 0.033468 | 0.049 | 0.0335 | 1 |
| Q86SQ9 | DHDDS | NVCFAYtsRHEIsNA | S158 | 0.001 | 98.85 | 333 | 0.000795 | 0.001 | 0.0008 | 1 |
| Q9NP64 | ZCCHC17 | DRIKVsLsMKVVNQG | S86 | 0 | 95.61 | 241 | 0 | 0 | 0 | 0.8571 |
| Q8IY50 | SLC35F3 | GIALVVAsASMSALY | S241 | 0.001 | 95.94 | 421 | 0.000795 | 0.001 | 0.0008 | 0.8 |
| Q9H5P4 | PDZD7 | VTRHCSRyVHEGGIE | Y584 | 0.08 | 90.1 | 1033 | 0.035095 | 0.08 | 0.0351 | 0.8571 |
| O60242 | ADGRB3 | GDGMsQVtKTLLDLT | T599 | 0.065 | 94.63 | 1522 | 0.044396 | 0.065 | 0.0444 | 1 |
| O60242 | ADGRB3 | YAGDLLMsVEILRNV | S619 | 0 | 96.23 | 1522 | 0 | 0 | 0 | 1 |
| O60242 | ADGRB3 | FVLGAVLyKNLDLIL | Y767 | 0.043 | 95.96 | 1522 | 0.018863 | 0.043 | 0.0189 | 1 |
| P46736 | BRCC3 | IGkVCLEsAVELPkI | S231 | 0.17 | 95.41 | 316 | 0.135135 | 0.17 | 0.1351 | 0.875 |
| Q5M8T2 | SLC35D3 | TLVQCLTsSTAALSL | S49 | 0.132 | 96.18 | 416 | 0.104929 | 0.132 | 0.1049 | 0.9 |
| Q53F19 | NCBP3 | TRALINMssLPAQDk | S179 | 0.103 | 92.09 | 620 | 0.081876 | 0.103 | 0.0819 | 1 |
| Q53F19 | NCBP3 | LSQVEEEsLLRNDLR | S240 | 0.103 | 90.31 | 620 | 0.081876 | 0.103 | 0.0819 | 1 |
| Q5JPI9 | EEF1AKMT2 | EIWFGEEsMNRLIRW | S64 | 0.009 | 85.63 | 291 | 0.007154 | 0.168 | 0.1335 | 1 |
| P14174 | MIF | MPMFIVNtNVPrAsV | T8 | 0 | 98.94 | 115 | 0 | 0 | 0 | 1 |

| | | | | | | | | | | |
|---|---|---|---|---|---|---|---|---|---|---|
| P19827 | ITIH1 | EKRQAVDtAVDGVFI | T46 | 0.095 | 88.56 | 911 | 0.064886 | 0.19 | 0.1298 | 1 |
| P19827 | ITIH1 | ASILIMLtDGDPtEG | T402 | 0 | 98.4 | 911 | 0 | 0 | 0 | 1 |
| P19827 | ITIH1 | IRGRFPLyNLGFGHN | Y431 | 0.106 | 98.77 | 911 | 0.046501 | 0.106 | 0.0465 | 0.8462 |
| Q96RY7 | IFT140 | VISVAILsERAMSSH | S405 | 0.171 | 87.1 | 1462 | 0.13593 | 0.188 | 0.1494 | 1 |
| Q96RY7 | IFT140 | PDSKICFyDVEMDTV | Y602 | 0.056 | 93.33 | 1462 | 0.024566 | 0.056 | 0.0246 | 1 |
| Q7Z3B4 | NUP54 | QQLGVTLsMtRtELs | S282 | 0.148 | 92.54 | 507 | 0.117647 | 0.183 | 0.1455 | 1 |
| Q14839 | CHD4 | LIVRVELsPMQKKYY | S979 | 0.148 | 92.16 | 1912 | 0.117647 | 0.148 | 0.1176 | 1 |
| Q14315 | FLNC | VADFkVFtkGAGSGE | T501 | 0.123 | 87.49 | 2725 | 0.084011 | 0.164 | 0.1028 | 1 |
| Q14315 | FLNC | YPVVPGKyVVTITWG | Y545 | 0.005 | 94.59 | 2725 | 0.002193 | 0.005 | 0.0022 | 1 |
| Q9BZV1 | UBXN6 | GLRkYNytLLRVRLP | T337 | 0.041 | 89.36 | 441 | 0.028004 | 0.041 | 0.028 | 1 |
| Q9BZV1 | UBXN6 | RERLGAVyGFVREAL | Y363 | 0.052 | 89.97 | 441 | 0.022812 | 0.052 | 0.0228 | 1 |
| Q8N4C6 | NIN | QEEGIENsQEILKAL | S322 | 0.054 | 81.21 | 2090 | 0.042925 | 0.054 | 0.0429 | 1 |
| P78330 | PSPH | AEQPPHLtPGIRELV | T88 | 0.081 | 98.03 | 225 | 0.055324 | 0.081 | 0.0553 | 1 |
| Q15434 | RBMS2 | GNDQLSKtNLYIRGL | T56 | 0.045 | 92.89 | 407 | 0.030736 | 0.045 | 0.0307 | 1 |
| P28288 | ABCD3 | FDHVPLAtPNGDVLI | T449 | 0.196 | 84.73 | 659 | 0.133871 | 0.196 | 0.1339 | 1 |
| P28288 | ABCD3 | SDLVLkEyLDNVQLG | Y545 | 0.204 | 92.95 | 659 | 0.089492 | 0.204 | 0.0895 | 0.8 |
| P28288 | ABCD3 | LWkHHEyyLHMDGRG | Y636 | 0.174 | 94.82 | 659 | 0.076331 | 0.174 | 0.0763 | 1 |
| P10635 | CYP2D6 | WREQRRFsVSTLRNL | S135 | 0.003 | 96.45 | 497 | 0.002385 | 0.003 | 0.0024 | 0.8889 |
| Q8WTP8 | AEN | IDCEMVGtGPRGRVS | T120 | 0.006 | 97.84 | 325 | 0.004098 | 0.006 | 0.0041 | 1 |
| P35249 | RFC4 | MEKESKTtRFCLICN | T174 | 0.006 | 90.27 | 363 | 0.004098 | 0.139 | 0.0949 | 1 |
| Q7Z2T5 | TRMT1L | NMVEENPyRQLPCNC | Y508 | 0.24 | 81.62 | 733 | 0.105285 | 0.24 | 0.1053 | 1 |
| Q6S8J3 | POTEE | APrAVFPsIVGRPRQ | S733 | 0.064 | 96.13 | 1075 | 0.050875 | 0.064 | 0.0509 | 1 |
| Q6S8J3 | POTEE | MWISkQEyDEsGPsI | Y1062 | 0.113 | 97.33 | 1075 | 0.049571 | 0.113 | 0.0496 | 1 |
| P84095 | RHOG | DLRAQPDtLRRLkEQ | T125 | 0.131 | 98.19 | 191 | 0.089475 | 0.131 | 0.0895 | 1 |
| P19835 | CEL | AKLGAVytEGGFVEG | T28 | 0.029 | 98.68 | 753 | 0.019807 | 0.054 | 0.0364 | 1 |
| P19835 | CEL | GLIRRAIsQSGVALs | S238 | 0.004 | 98.96 | 753 | 0.00318 | 0.004 | 0.0032 | 1 |
| P19835 | CEL | sQSGVALsPWVIQKN | S245 | 0.046 | 98.64 | 753 | 0.036566 | 0.063 | 0.0533 | 0.8 |
| A9QM74 | KPNA7 | ITFLRNItWTLsNLC | T220 | 0.002 | 98.18 | 516 | 0.001366 | 0.002 | 0.0014 | 1 |
| A9QM74 | KPNA7 | RNItWTLsNLCRNKN | S224 | 0.039 | 97.59 | 516 | 0.031002 | 0.039 | 0.031 | 1 |
| A9QM74 | KPNA7 | ELNVLTPsLRTVGNI | S303 | 0 | 98.36 | 516 | 0 | 0 | 0 | 1 |
| Q15652 | JMJD1C | YFRRLsFsKNGVVRI | S1763 | 0.147 | 87.18 | 2540 | 0.116852 | 0.18 | 0.1431 | 1 |

| | | | | | | | | | | |
|---|---|---|---|---|---|---|---|---|---|---|
| Q9UPS6 | SETD1B | ADMREKRyEDEGIGS | Y1873 | 0.121 | 93.2 | 1966 | 0.053081 | 0.121 | 0.0531 | 1 |
| Q96DT5 | DNAH11 | IERVRAAtsELEHRV | T820 | 0.028 | 88.36 | 4516 | 0.019124 | 0.028 | 0.0191 | 1 |
| Q96AC1 | FERMT2 | TLkGYKQyWCTFKDT | Y395 | 0.25 | 95.85 | 680 | 0.109671 | 0.25 | 0.1097 | 1 |
| Q14956 | GPNMB | YRRHGRAyVPIAQVK | Y220 | 0.062 | 91.66 | 572 | 0.027199 | 0.062 | 0.0272 | 1 |
| P02679 | FGG | GRTSTADyAMFKVGP | Y288 | 0.006 | 98.29 | 453 | 0.002632 | 0.006 | 0.0026 | 1 |
| P02679 | FGG | VGPEADkyRLTYAYF | Y300 | 0.005 | 98.84 | 453 | 0.002193 | 0.005 | 0.0022 | 1 |
| P02679 | FGG | VyyQGGtySkASTPN | Y380 | 0.175 | 97.36 | 453 | 0.07677 | 0.175 | 0.0768 | 1 |
| Q9BXX2 | ANKRD30B | VLDGEGRtPLMKALQ | T108 | 0.024 | 94.54 | 1392 | 0.016392 | 0.024 | 0.0164 | 1 |
| P05062 | ALDOB | VPGICFLsGGMsEED | S272 | 0.079 | 96.93 | 364 | 0.062798 | 0.13 | 0.1033 | 1 |
| P05062 | ALDOB | PWKLSFSyGRALQAs | Y302 | 0 | 98.68 | 364 | 0 | 0 | 0 | 0.875 |
| Q9UNT1 | RABL2B | FDVQRKVtYRNLSTW | T108 | 0.008 | 95.22 | 228 | 0.005464 | 0.008 | 0.0055 | 0.8889 |
| A6NED2 | RCCD1 | DAAGQVFsWGGGRHG | S181 | 0 | 98.67 | 376 | 0 | 0 | 0 | 0.8333 |
| Q6TFL4 | KLHL24 | NEMMSPRtRPRRSTG | T303 | 0.041 | 86.18 | 600 | 0.028004 | 0.041 | 0.028 | 1 |
| Q6TFL4 | KLHL24 | PIAkRCItAVsLNNL | T496 | 0.124 | 96.64 | 600 | 0.084694 | 0.124 | 0.0847 | 1 |
| Q9UNX4 | WDR3 | LTKQYLRyVAsAVFG | Y10 | 0.087 | 94.87 | 943 | 0.038166 | 0.087 | 0.0382 | 1 |
| Q9UNX4 | WDR3 | kSFDLIHsPHGELKA | S373 | 0.165 | 79.1 | 943 | 0.131161 | 0.165 | 0.1312 | 1 |
| A4D0S4 | LAMB4 | CDCNRAGtQKPICDP | T1135 | 0.133 | 80.71 | 1761 | 0.090841 | 0.133 | 0.0908 | 0.9091 |
| Q5T160 | RARS2 | FDTMIYVtDKGQKKH | T346 | 0.166 | 95.53 | 578 | 0.11338 | 0.166 | 0.1134 | 1 |
| Q86VN1 | VPS36 | AGTLLLStHRLIWRD | T44 | 0.044 | 94.88 | 386 | 0.030053 | 0.044 | 0.0301 | 0.875 |
| P60660 | MYL6 | DsNGCINyEAFVRHI | Y141 | 0.092 | 97.17 | 151 | 0.040359 | 0.092 | 0.0404 | 1 |
| Q86SQ0 | PHLDB2 | LCNLEKkyssLsGGk | Y811 | 0.127 | 90.05 | 1253 | 0.055713 | 0.127 | 0.0557 | 1 |
| Q08174 | PCDH1 | GsDSKKKyFLQTTTP | Y456 | 0.279 | 88.58 | 1060 | 0.122393 | 0.279 | 0.1224 | 1 |
| Q14542 | SLC29A2 | LSVFPAItAMVTSsT | T312 | 0 | 94.75 | 456 | 0 | 0.144 | 0.0984 | 1 |
| Q96KR1 | ZFR | QVTITLTsPIIREEN | S869 | 0 | 92.98 | 1074 | 0 | 0.055 | 0.0437 | 1 |
| P04180 | LCAT | DTVATRstELCGLWQ | T377 | 0.007 | 98.55 | 440 | 0.004781 | 0.007 | 0.0048 | 1 |
| P62269 | RPS18 | IkGVGRRyAHVVLrk | Y40 | 0.102 | 94.84 | 152 | 0.044746 | 0.102 | 0.0447 | 1 |
| Q9H5J4 | ELOVL6 | GFYNGPVsKFWAYAF | S109 | 0 | 93.59 | 265 | 0 | 0 | 0 | 1 |
| Q9UPX0 | IGSF9B | AYPGNLTyTWYWQDE | Y261 | 0.304 | 93.26 | 1349 | 0.13336 | 0.304 | 0.1334 | 1 |
| Q9UPX0 | IGSF9B | WRKVGKPsRSKHSAL | S461 | 0.182 | 84.91 | 1349 | 0.144674 | 0.182 | 0.1447 | 1 |
| P42684 | ABL2 | QGWVPsNyItPVNSL | Y161 | 0.174 | 93.35 | 1182 | 0.076331 | 0.174 | 0.0763 | 1 |
| P42684 | ABL2 | LSSLINGsFLVRESE | S194 | 0.04 | 97.05 | 1182 | 0.031797 | 0.04 | 0.0318 | 1 |

| | | | | | | | | | | |
|---|---|---|---|---|---|---|---|---|---|---|
| P42684 | ABL2 | ESsPGQLsISLRyEG | S208 | 0.094 | 95.18 | 1182 | 0.074722 | 0.094 | 0.0747 | 1 |
| O43765 | SGTA | DNkKRLAyAIIQFLH | Y9 | 0.166 | 93.92 | 313 | 0.072822 | 0.166 | 0.0728 | 1 |
| P34913 | EPHX2 | LAMDMKGyGEsSAPP | Y296 | 0.05 | 98.87 | 555 | 0.021934 | 0.05 | 0.0219 | 1 |
| P34913 | EPHX2 | APPEIEEyCMEVLCK | Y308 | 0 | 98.85 | 555 | 0 | 0 | 0 | 0.8667 |
| P48169 | GABRA4 | YPKSEMIyTWTKGPE | Y202 | 0.107 | 96.9 | 554 | 0.046939 | 0.107 | 0.0469 | 1 |
| Q9BY44 | EIF2A | kINDFVLsPGPQPYk | S176 | 0.048 | 95.13 | 585 | 0.038156 | 0.048 | 0.0382 | 0.8571 |
| Q14994 | NR1I3 | CKGFFRRtVSKSIGP | T38 | 0.164 | 95.15 | 352 | 0.112014 | 0.164 | 0.112 | 1 |
| Q01484 | ANK2 | AATtNGytPLHISAR | T532 | 0.032 | 97.95 | 3957 | 0.021856 | 0.032 | 0.0219 | 1 |
| Q01484 | ANK2 | HVAAKyGsLDVAKLL | S575 | 0.128 | 96.52 | 3957 | 0.101749 | 0.128 | 0.1017 | 1 |
| Q01484 | ANK2 | VTPLHLAsQEGHTDM | S670 | 0 | 96.44 | 3957 | 0 | 0 | 0 | 1 |
| Q01484 | ANK2 | DAHTKLGytPLIVAC | Y729 | 0.198 | 93.32 | 3957 | 0.08686 | 0.198 | 0.0869 | 1 |
| Q01484 | ANK2 | AHTKLGytPLIVACH | T730 | 0.005 | 96.04 | 3957 | 0.003415 | 0.005 | 0.0034 | 1 |
| P13535 | MYH8 | AsADIETyLLEkSRV | Y269 | 0.255 | 76.16 | 1937 | 0.111865 | 0.255 | 0.1119 | 1 |
| Q5T2W1 | PDZK1 | DKETDNMyRLAHFSP | Y328 | 0.029 | 87.71 | 519 | 0.012722 | 0.029 | 0.0127 | 0.8667 |
| Q7Z3D6 | DGLUCY | KDELLKAsLsLSHAR | S324 | 0 | 98.76 | 616 | 0 | 0 | 0 | 0.9 |
| Q9GZT4 | SRR | LKPSVkVyAAEPsNA | Y207 | 0.19 | 98.93 | 340 | 0.08335 | 0.19 | 0.0833 | 1 |
| Q9GZT4 | SRR | NADDCyQskLKGkLM | S220 | 0 | 98.79 | 340 | 0 | 0 | 0 | 1 |
| Q86YG4 | NT5DC4 | ASGRPWRsyFDLIVV | S247 | 0.089 | 93.41 | 428 | 0.070747 | 0.089 | 0.0707 | 1 |
| Q86YG4 | NT5DC4 | VRGMDILyIGDHIFG | Y314 | 0.005 | 98.61 | 428 | 0.002193 | 0.005 | 0.0022 | 1 |
| Q8N6W0 | CELF5 | HKGCAFLtyCARDSA | T92 | 0.124 | 92.29 | 485 | 0.084694 | 0.124 | 0.0847 | 1 |
| Q8N6W0 | CELF5 | KGCAFLtyCARDSAI | Y93 | 0 | 93.38 | 485 | 0 | 0 | 0 | 0.9412 |
| Q9Y2G9 | SBNO2 | TGPyDGFyLSykVRG | Y1068 | 0.113 | 95.58 | 1366 | 0.049571 | 0.222 | 0.0964 | 1 |
| Q9Y2G9 | SBNO2 | LQIVRLKtKDRKKQV | T1198 | 0.064 | 85.92 | 1366 | 0.043713 | 0.064 | 0.0437 | 1 |
| P32418 | SLC8A1 | CTIGLKDsVTAVVFV | S834 | 0.135 | 90.54 | 973 | 0.107313 | 0.135 | 0.1073 | 0.8333 |
| Q8N7W2 | BEND7 | DDKtLANsLPNGKRK | S328 | 0.051 | 87 | 519 | 0.040541 | 0.051 | 0.0405 | 1 |
| P58173 | OR2B6 | YLLLAVMsFDRFVAI | S119 | 0.002 | 97.71 | 313 | 0.00159 | 0.002 | 0.0016 | 0.8182 |
| Q9NQT6 | FSCN3 | ECDGTVCyGRPRTSH | Y84 | 0.074 | 90.94 | 498 | 0.032463 | 0.074 | 0.0325 | 1 |
| P27449 | ATP6V0C | AAYGTAKsGtGIAAM | S37 | 0.001 | 92.74 | 155 | 0.000795 | 0.001 | 0.0008 | 1 |
| P09210 | GSTA2 | AEKPKLHysNIRGRM | Y9 | 0.191 | 98.74 | 222 | 0.083789 | 0.348 | 0.1406 | 1 |
| P09210 | GSTA2 | EKPKLHysNIRGRME | S10 | 0.034 | 97.11 | 222 | 0.027027 | 0.034 | 0.027 | 1 |
| O43542 | XRCC3 | HLQSLGAtLRELSSA | T241 | 0.213 | 96.83 | 346 | 0.145482 | 0.213 | 0.1455 | 1 |

| | | | | | | | | | | |
|---|---|---|---|---|---|---|---|---|---|---|
| O00748 | CES2 | KEMDREAsQAALQKM | S368 | 0.126 | 92.86 | 559 | 0.100159 | 0.126 | 0.1002 | 1 |
| Q99567 | NUP88 | sDEEDkDsLQELstE | S437 | 0.125 | 71.22 | 741 | 0.099364 | 0.125 | 0.0994 | 1 |
| Q6YBV0 | SLC36A4 | PRLDIVIsFVGAVSS | S417 | 0.006 | 96.38 | 504 | 0.004769 | 0.006 | 0.0048 | 1 |
| P06746 | POLB | AIHKyNAyRkAAsVI | Y39 | 0.199 | 96.24 | 335 | 0.087298 | 0.199 | 0.0873 | 1 |
| Q86T82 | USP37 | MLTLQDNsFLsIDKV | S80 | 0.019 | 88.47 | 979 | 0.015103 | 0.019 | 0.0151 | 0.8333 |
| P05164 | MPO | PNPRVPLsRVFFASW | S528 | 0.042 | 90.93 | 745 | 0.033386 | 0.042 | 0.0334 | 0.8571 |
| Q9NYV4 | CDK12 | FSEDHIKsFMKQLME | S838 | 0.153 | 97.75 | 1490 | 0.121622 | 0.153 | 0.1216 | 0.8889 |
| Q9NYV4 | CDK12 | NsEEsRPytNKVITL | Y892 | 0.24 | 94.21 | 1490 | 0.105285 | 0.24 | 0.1053 | 1 |
| Q9NYV4 | CDK12 | LISRLCGsPCPAVWP | S954 | 0.037 | 97.03 | 1490 | 0.029412 | 0.037 | 0.0294 | 1 |
| Q9BY50 | SEC11C | RGDLLFLtNFREDPI | T82 | 0.218 | 96.53 | 192 | 0.148897 | 0.218 | 0.1489 | 1 |
| Q9NYW3 | TAS2R7 | ILTsLAIsRICLLCV | S54 | 0.051 | 97.38 | 318 | 0.040541 | 0.051 | 0.0405 | 1 |
| O14792 | HS3ST1 | LLILRDPsERVLsDY | S150 | 0.03 | 98.46 | 307 | 0.023847 | 0.03 | 0.0238 | 0.8571 |
| P22607 | FGFR3 | ANCTHDLyMIMRECW | Y724 | 0.133 | 95.78 | 806 | 0.058345 | 0.296 | 0.1299 | 0.8333 |
| P07333 | CSF1R | QSQDYQCsALMGGRK | S178 | 0.006 | 94.97 | 972 | 0.004769 | 0.006 | 0.0048 | 1 |
| O60684 | KPNA6 | DSDLQLAttQKFRKL | T100 | 0.015 | 97.56 | 536 | 0.010245 | 0.015 | 0.0102 | 1 |
| O60684 | KPNA6 | SDLQLAttQKFRKLL | T101 | 0 | 97.96 | 536 | 0 | 0 | 0 | 1 |
| Q8NB12 | SMYD1 | IIFAERAysAVVFDs | Y39 | 0.071 | 96.75 | 490 | 0.031147 | 0.071 | 0.0311 | 1 |
| Q8NB12 | SMYD1 | IFAERAysAVVFDsL | S40 | 0.091 | 97.37 | 490 | 0.072337 | 0.091 | 0.0723 | 1 |
| Q8NB12 | SMYD1 | QEPVFADtNIyMLRM | T346 | 0.121 | 97.73 | 490 | 0.082645 | 0.151 | 0.093 | 1 |
| Q9H2Y9 | SLCO5A1 | DNVKKENssLYLAIM | S298 | 0.041 | 84.38 | 848 | 0.032592 | 0.041 | 0.0326 | 1 |
| P31943 | HNRNPH1 | rYVEVFksNNVEMdW | S88 | 0.072 | 82.74 | 449 | 0.057234 | 0.072 | 0.0572 | 1 |
| Q13099 | IFT88 | KMNMGNIyLkQRNys | Y243 | 0.341 | 95.77 | 824 | 0.149592 | 0.341 | 0.1496 | 0.8889 |
| B5MCN3 | SEC14L6 | VHSKRISyTVEVLLP | Y379 | 0.144 | 97.35 | 397 | 0.063171 | 0.144 | 0.0632 | 0.9 |
| Q96KR1 | ZFR | GRRHRLQyKKKVNPD | Y610 | 0.334 | 90.86 | 1074 | 0.146521 | 0.334 | 0.1465 | 1 |
| P26196 | DDX6 | GGAkVMAtTGGTNLR | T199 | 0.015 | 96.92 | 483 | 0.010245 | 0.015 | 0.0102 | 1 |
| P26196 | DDX6 | TLKGVTQyyAYVTER | Y312 | 0.273 | 98.34 | 483 | 0.119761 | 0.406 | 0.1439 | 1 |
| P26196 | DDX6 | GLAINLItyDDRFNL | T439 | 0.14 | 97.81 | 483 | 0.095622 | 0.14 | 0.0956 | 1 |
| Q96JJ6 | JPH4 | ESVSGLRyAGLWKDG | Y106 | 0.01 | 96.12 | 628 | 0.004387 | 0.01 | 0.0044 | 1 |
| Q96JJ6 | JPH4 | TYSDGGtyQGQWQAG | Y129 | 0 | 94.98 | 628 | 0 | 0 | 0 | 0.9286 |
| Q9P1Z9 | CCDC180 | LTKEVMEyPAIMLKE | Y613 | 0.054 | 88.45 | 1701 | 0.023689 | 0.383 | 0.1164 | 1 |
| Q16880 | UGT8 | PGHPVNRtIYWIDYI | T444 | 0.001 | 97.68 | 541 | 0.000683 | 0.001 | 0.0007 | 1 |

| | | | | | | | | | | |
|---|---|---|---|---|---|---|---|---|---|---|
| Q99758 | ABCA3 | NGAGKTTtLSMLTGL | T575 | 0.042 | 90.37 | 1704 | 0.028687 | 0.042 | 0.0287 | 1 |
| P62277 | RPS13 | DDVkEQIykLAkkGL | Y38 | 0.313 | 96.39 | 151 | 0.137309 | 0.313 | 0.1373 | 1 |
| P78562 | PHEX | KAKILYssCMNEKAI | S141 | 0.026 | 98.22 | 749 | 0.020668 | 0.026 | 0.0207 | 1 |
| P78562 | PHEX | RKFsLLQtLAtFRGQ | T190 | 0.027 | 97.87 | 749 | 0.018441 | 0.027 | 0.0184 | 1 |
| P78562 | PHEX | sLLQtLAtFRGQYSN | T193 | 0.212 | 97.33 | 749 | 0.144799 | 0.212 | 0.1448 | 1 |
| P78562 | PHEX | VVVRVPQyFKDLFRI | Y346 | 0.026 | 98.01 | 749 | 0.011406 | 0.026 | 0.0114 | 1 |
| P78562 | PHEX | LNVKGKRtLGENIAD | T639 | 0.05 | 97.69 | 749 | 0.034151 | 0.05 | 0.0342 | 1 |
| Q8IWV7 | UBR1 | ILFATTIyRIGLKVP | Y1302 | 0.084 | 93.51 | 1749 | 0.03685 | 0.084 | 0.0369 | 1 |
| O00534 | VWA5A | FIFLMDRsGsMQSPM | S289 | 0.008 | 94.6 | 786 | 0.006359 | 0.008 | 0.0064 | 1 |
| O00534 | VWA5A | GLSAKMLsPEQtVIF | S485 | 0.125 | 91.49 | 786 | 0.099364 | 0.125 | 0.0994 | 1 |
| Q7L5A8 | FA2H | HHFAHQKsGFGISTK | S346 | 0.127 | 95.83 | 372 | 0.100954 | 0.127 | 0.101 | 1 |
| Q7L5A8 | FA2H | DYCFHTLtPEKPHLK | T363 | 0.118 | 88.94 | 372 | 0.080596 | 0.118 | 0.0806 | 1 |
| Q9H040 | SPRTN | MHRINSLtGANITVY | T145 | 0.009 | 94.73 | 489 | 0.006147 | 0.009 | 0.0061 | 1 |
| P54886 | ALDH18A1 | IAASSQDsVGRVLRR | S450 | 0.016 | 94.3 | 795 | 0.012719 | 0.016 | 0.0127 | 0.8 |
| P54886 | ALDH18A1 | CVFWNAStRFSDGYR | T725 | 0 | 98.3 | 795 | 0 | 0 | 0 | 1 |
| Q9H0I9 | TKTL2 | SQVKNKPtAIVAKTF | T240 | 0.009 | 98.88 | 626 | 0.006147 | 0.009 | 0.0061 | 1 |
| Q9H0I9 | TKTL2 | KIATQKTyGLALAKL | Y325 | 0.008 | 96.54 | 626 | 0.003509 | 0.008 | 0.0035 | 1 |
| Q9H0I9 | TKTL2 | QLRMGAIsQANINLI | S410 | 0.124 | 97.92 | 626 | 0.098569 | 0.124 | 0.0986 | 1 |
| Q9H0I9 | TKTL2 | LLDMFGIstRHIIAA | S612 | 0.089 | 98.46 | 626 | 0.070747 | 0.089 | 0.0707 | 1 |
| Q9H0I9 | TKTL2 | LDMFGIstRHIIAAV | T613 | 0.137 | 98.51 | 626 | 0.093573 | 0.137 | 0.0936 | 1 |
| O15498 | YKT6 | SQLIVERsSKGTRAS | S51 | 0.098 | 95.37 | 198 | 0.077902 | 0.113 | 0.0898 | 0.875 |
| Q6ZVH7 | ESPNL | LTCLKLLtAAHGssV | T157 | 0.138 | 95.53 | 1005 | 0.094256 | 0.138 | 0.0943 | 1 |
| Q6ZVH7 | ESPNL | LtAAHGssVNRRtRS | S163 | 0.032 | 75.87 | 1005 | 0.025437 | 0.032 | 0.0254 | 1 |
| Q6ZVH7 | ESPNL | GssVNRRtRSGASPL | T168 | 0.213 | 95.29 | 1005 | 0.145482 | 0.213 | 0.1455 | 1 |
| Q6ZVH7 | ESPNL | FAARGGHtPILDRLL | T252 | 0.215 | 98.13 | 1005 | 0.146848 | 0.215 | 0.1468 | 1 |
| Q6ZVH7 | ESPNL | DRLLLMGtPILRDSW | T263 | 0.107 | 94.9 | 1005 | 0.073082 | 0.107 | 0.0731 | 1 |
| P08631 | HCK | IVVALYDyEAIHHED | Y89 | 0.055 | 94.27 | 526 | 0.024128 | 0.11 | 0.0483 | 1 |
| P08631 | HCK | ETTKGSysLSVRDYD | S181 | 0 | 97.49 | 526 | 0 | 0 | 0 | 1 |
| P08631 | HCK | AWEIPREsLKLEKKL | S261 | 0.142 | 96.83 | 526 | 0.112878 | 0.142 | 0.1129 | 1 |
| P55287 | CDH11 | AQTGIIRtALPNMDR | T219 | 0.054 | 96.95 | 796 | 0.036883 | 0.054 | 0.0369 | 1 |
| Q9UBL9 | P2RX2 | IADRTDGyLKRCTFH | Y222 | 0.298 | 88.96 | 471 | 0.130728 | 0.298 | 0.1307 | 1 |

| | | | | | | | | | | |
|---|---|---|---|---|---|---|---|---|---|---|
| Q92526 | CCT6B | QkGIDPFsLDsLAKH | S301 | 0.072 | 92.45 | 530 | 0.057234 | 0.072 | 0.0572 | 1 |
| P51530 | DNA2 | DIEEsIWsPRFGLKG | S267 | 0.001 | 93.7 | 1060 | 0.000795 | 0.001 | 0.0008 | 0.875 |
| P51530 | DNA2 | VVLYTLLsQERRADP | S321 | 0.019 | 95.06 | 1060 | 0.015103 | 0.019 | 0.0151 | 1 |
| O15205 | UBD | DGKMMADyGIRKGNL | Y148 | 0.171 | 93.3 | 165 | 0.075015 | 0.171 | 0.075 | 0.9091 |
| O75976 | CPD | RLySLGksVESRELY | S532 | 0.075 | 94.77 | 1380 | 0.059619 | 0.075 | 0.0596 | 1 |
| O75976 | CPD | PSMNPDGyEKSQEGD | Y611 | 0 | 98.14 | 1380 | 0 | 0 | 0 | 1 |
| Q9HAW9 | UGT1A8 | PQTVLWRyTGTRPSN | Y334 | 0.181 | 95.53 | 530 | 0.079402 | 0.181 | 0.0794 | 1 |
| Q9HBG6 | IFT122 | SkVkILFtLAkQskA | T1001 | 0.001 | 95.25 | 1241 | 0.000683 | 0.001 | 0.0007 | 1 |
| Q9HBG6 | IFT122 | LFtLAkQskALGAYR | S1006 | 0 | 95.31 | 1241 | 0 | 0 | 0 | 1 |
| P50213 | IDH3A | IPsEAKEsMDkNKMG | S87 | 0.043 | 97.91 | 366 | 0.034181 | 0.043 | 0.0342 | 1 |
| Q13976 | PRKG1 | FKDSKyLyMLMEACL | Y435 | 0.294 | 97.77 | 671 | 0.128974 | 0.294 | 0.129 | 1 |
| Q13976 | PRKG1 | LILDHRGyAKLVDFG | Y497 | 0.108 | 98.58 | 671 | 0.047378 | 0.108 | 0.0474 | 0.9286 |
| Q13976 | PRKG1 | KIGFGKKtWtFCGtP | T515 | 0.091 | 85.02 | 671 | 0.062154 | 0.091 | 0.0622 | 1 |
| Q13393 | PLD1 | ELALGINsEYTKRTL | S431 | 0.049 | 96.01 | 1074 | 0.038951 | 0.049 | 0.039 | 1 |
| Q13393 | PLD1 | TKIMKSKyRSLSYPF | Y711 | 0.035 | 92.3 | 1074 | 0.015354 | 0.035 | 0.0154 | 1 |
| O15537 | RS1 | GFGCAWLsKFQDsSQ | S114 | 0.002 | 89.1 | 224 | 0.00159 | 0.002 | 0.0016 | 1 |
| O15537 | RS1 | ERLNWIyyKDQtGNN | Y166 | 0.118 | 93.47 | 224 | 0.051765 | 0.118 | 0.0518 | 1 |
| Q9BWW4 | SSBP3 | EkLALyVyEYLLHVG | Y25 | 0.321 | 96.34 | 388 | 0.140818 | 0.321 | 0.1408 | 1 |
| P51689 | ARSD | TPDsEPLyHAVIARV | Y537 | 0.116 | 98.37 | 593 | 0.050888 | 0.116 | 0.0509 | 1 |
| Q13332 | PTPRS | RQESIIKyELLFREG | Y556 | 0.04 | 93.29 | 1948 | 0.017547 | 0.04 | 0.0175 | 1 |
| Q13332 | PTPRS | EVGPGPEsSPVVVRT | S708 | 0.107 | 92.07 | 1948 | 0.085056 | 0.182 | 0.1447 | 1 |
| Q13332 | PTPRS | ITNLQPEtAYSITVA | T803 | 0.103 | 81.01 | 1948 | 0.07035 | 0.203 | 0.1387 | 1 |
| Q13332 | PTPRS | SITVAAytMKGDGAR | T813 | 0.046 | 84.28 | 1948 | 0.031419 | 0.046 | 0.0314 | 0.9167 |
| Q15080 | NCF4 | VFVIEVKtKGGsKyL | T47 | 0.065 | 92.81 | 339 | 0.044396 | 0.065 | 0.0444 | 1 |
| P11166 | SLC2A1 | GFSKLGKsFEMLILG | S118 | 0.138 | 96.53 | 492 | 0.109698 | 0.138 | 0.1097 | 1 |
| Q96A58 | RERG | IkKPKNVtLILVGNK | T112 | 0.071 | 97.33 | 199 | 0.048494 | 0.142 | 0.097 | 1 |
| Q8NFY4 | SEMA6D | SDAVIYRsMGDGSAL | S201 | 0.067 | 95.45 | 1073 | 0.053259 | 0.067 | 0.0533 | 1 |
| Q96JQ0 | DCHS1 | DREERDAyNLRVTAT | Y438 | 0.08 | 88.24 | 3298 | 0.035095 | 0.08 | 0.0351 | 1 |
| Q9UHX1 | PUF60 | kGYGFIEyEKAQSsQ | Y274 | 0.035 | 92.8 | 559 | 0.015354 | 0.035 | 0.0154 | 1 |
| Q9UHX1 | PUF60 | EyEKAQSsQDAVSSM | S280 | 0.105 | 90.67 | 559 | 0.083466 | 0.105 | 0.0835 | 1 |
| Q96H96 | COQ2 | WGALLGWsAIKGSCD | S222 | 0 | 98.61 | 371 | 0 | 0 | 0 | 1 |

| | | | | | | | | | | |
|---|---|---|---|---|---|---|---|---|---|---|
| P09874 | PARP1 | DsQILDLsNRFytLI | S733 | 0 | 96.73 | 1014 | 0 | -0.003 | 0.014 | 1 |
| P09874 | PARP1 | LDLsNRFytLIPHDF | Y737 | 0.046 | 94.53 | 1014 | 0.02018 | 0.321 | 0.1328 | 1 |
| Q86X10 | RALGAPB | FDTVFIFyMKPGQKT | Y1167 | 0.01 | 96.31 | 1494 | 0.004387 | 0.01 | 0.0044 | 0.9 |
| Q96EH3 | MALSU1 | TDyFVIVsGTSTRHL | S128 | 0.061 | 97.1 | 234 | 0.04849 | 0.061 | 0.0485 | 1 |
| Q96L46 | CAPNS2 | YDRDHSGsLGSSQLR | S168 | 0.101 | 91.5 | 248 | 0.080286 | 0.101 | 0.0803 | 1 |
| Q86Y97 | KMT5C | LRAFLPEsGFtILPC | S104 | 0.029 | 97.55 | 462 | 0.023053 | 0.054 | 0.0444 | 0.8571 |
| Q8N960 | CEP120 | SVEILRVyLALQSKL | Y262 | 0.203 | 92.34 | 986 | 0.089053 | 0.203 | 0.0891 | 1 |
| B0I1T2 | MYO1G | GPEAIARyQGRELyE | Y66 | 0.233 | 91.48 | 1018 | 0.102214 | 0.233 | 0.1022 | 1 |
| B0I1T2 | MYO1G | LAskEPFyVRCIkPN | Y598 | 0.138 | 93.51 | 1018 | 0.060539 | 0.138 | 0.0605 | 1 |
| B0I1T2 | MYO1G | FLLRYkMtCEyTWPN | T654 | 0.116 | 92.73 | 1018 | 0.07923 | 0.116 | 0.0792 | 1 |
| B0I1T2 | MYO1G | RRAWARDyLSSATDN | Y839 | 0.108 | 92.8 | 1018 | 0.047378 | 0.108 | 0.0474 | 1 |
| B0I1T2 | MYO1G | LLTDQHLykLDPDRQ | Y897 | 0.01 | 94.9 | 1018 | 0.004387 | 0.01 | 0.0044 | 1 |
| B0I1T2 | MYO1G | kLDPDRQyRVMRAVP | Y905 | 0.047 | 91.48 | 1018 | 0.020618 | 0.084 | 0.0368 | 1 |
| B0I1T2 | MYO1G | RCARGsFtLLWPsR_ | T1012 | 0.163 | 92.6 | 1018 | 0.111331 | 0.207 | 0.1281 | 1 |
| Q92692 | NECTIN2 | TTVALCIsKEGRPPA | S185 | 0.011 | 98.38 | 538 | 0.008744 | 0.011 | 0.0087 | 1 |
| O14727 | APAF1 | REDIkDyyTDLSILQ | Y383 | 0.014 | 91.4 | 1248 | 0.006142 | 0.014 | 0.0061 | 1 |
| Q13569 | TDG | kCIYEIFsKEVFGVk | S239 | 0.001 | 95.67 | 410 | 0.000795 | 0.001 | 0.0008 | 0.8571 |
| Q9BUP3 | HTATIP2 | DFRMQNKsVFILGAS | S20 | 0.103 | 97.95 | 242 | 0.081876 | 0.103 | 0.0819 | 1 |
| Q7RTS9 | DYM | LILVVIRtIQyNMtR | T446 | 0.006 | 94.34 | 669 | 0.004098 | 0.006 | 0.0041 | 1 |
| Q96PF1 | TGM7 | RNNKEHHtQEMGVKR | T25 | 0.009 | 97.27 | 710 | 0.006147 | 0.009 | 0.0061 | 1 |
| Q96PF1 | TGM7 | EMGVKRLtVRRGQPF | T34 | 0.06 | 98.19 | 710 | 0.040981 | 0.06 | 0.041 | 0.875 |
| P00390 | GSR | tLSGLEVsMVTAVPG | S308 | 0.036 | 98.63 | 522 | 0.028617 | 0.036 | 0.0286 | 1 |
| P00390 | GSR | NTNVkGIyAVGDVCG | Y371 | 0.155 | 98.93 | 522 | 0.067996 | 0.155 | 0.068 | 1 |
| O95394 | PGM3 | STIGVMVtAsHNPEE | T62 | 0.02 | 98.48 | 542 | 0.01366 | 0.02 | 0.0137 | 0.8182 |
| O95394 | PGM3 | MkVPVyCtkTGVkHL | T349 | 0.116 | 96.66 | 542 | 0.07923 | 0.191 | 0.1259 | 0.875 |
| O95394 | PGM3 | SRAFVRPsGtEDVVR | S498 | 0.041 | 85.04 | 542 | 0.032592 | 0.041 | 0.0326 | 1 |
| P26006 | ITGA3 | KTVEDVGsPLKYEFQ | S793 | 0.067 | 88.74 | 1051 | 0.053259 | 0.067 | 0.0533 | 1 |
| P26006 | ITGA3 | VKARVWNstFIEDYR | S936 | 0.087 | 87.16 | 1051 | 0.069158 | 0.087 | 0.0692 | 1 |
| P13569 | CFTR | VDsADNLsEKLEREW | S50 | 0.066 | 92.55 | 1480 | 0.052464 | 0.171 | 0.1375 | 0.875 |
| P13569 | CFTR | FLYLGEVtKAVQPLL | T94 | 0.172 | 91.78 | 1480 | 0.117478 | 0.172 | 0.1175 | 1 |
| Q7Z392 | TRAPPC11 | GKKIEITsVDLALGN | S686 | 0.178 | 94.5 | 1133 | 0.141495 | 0.178 | 0.1415 | 1 |

| | | | | | | | | | | |
|---|---|---|---|---|---|---|---|---|---|---|
| Q7Z392 | TRAPPC11 | MFLVyVSyLINTTVE | Y859 | 0.028 | 92.97 | 1133 | 0.012283 | 0.028 | 0.0123 | 1 |
| Q93100 | PHKB | DEVLFsQtLDKVVRK | T316 | 0 | 97.14 | 1093 | 0 | 0 | 0 | 0.9 |
| Q86UE8 | TLK2 | RIHNEDNsQFKDHPT | S450 | 0.062 | 93.19 | 772 | 0.049285 | 0.062 | 0.0493 | 1 |
| P11047 | LAMC1 | FHTSRPEsFAIYkRt | S159 | 0 | 95.49 | 1609 | 0 | 0 | 0 | 1 |
| P11047 | LAMC1 | ALCTDEFsDISPLtG | S210 | 0.073 | 91.94 | 1609 | 0.058029 | 0.073 | 0.058 | 1 |
| P11047 | LAMC1 | DPkVLKsyyyAISDF | Y273 | 0.135 | 90.49 | 1609 | 0.059223 | 0.135 | 0.0592 | 1 |
| P11047 | LAMC1 | PkVLKsyyyAISDFA | Y274 | 0.156 | 92.62 | 1609 | 0.068435 | 0.156 | 0.0684 | 1 |
| P11047 | LAMC1 | kVLKsyyyAISDFAV | Y275 | 0.046 | 94.77 | 1609 | 0.02018 | 0.046 | 0.0202 | 1 |
| Q12874 | SF3A3 | EICGNyTyRGPkAFQ | Y416 | 0.151 | 91.72 | 501 | 0.066242 | 0.151 | 0.0662 | 1 |
| Q8TDW7 | FAT3 | IRFEKEVyDVSISEF | Y381 | 0.204 | 88.23 | 4557 | 0.089492 | 0.204 | 0.0895 | 1 |
| Q8TDW7 | FAT3 | DVSISEFsPPGVVVA | S389 | 0.051 | 87.44 | 4557 | 0.040541 | 0.144 | 0.1337 | 1 |
| Q9P2N5 | RBM27 | FVDKLFEsLYTKNyL | S71 | 0.116 | 85.27 | 1060 | 0.09221 | 0.116 | 0.0922 | 0.8571 |
| Q9P2N5 | RBM27 | EsLYTKNyLPLLEPV | Y77 | 0.119 | 78.56 | 1060 | 0.052204 | 0.182 | 0.0798 | 1 |
| P58418 | CLRN1 | yKEGTYVyKTQSEKY | Y178 | 0.001 | 97.47 | 232 | 0.000439 | 0.001 | 0.0004 | 0.9286 |
| O15492 | RGS16 | TLMEKDSyPRFLkSP | Y168 | 0.203 | 98.45 | 202 | 0.089053 | 0.203 | 0.0891 | 1 |
| O15492 | RGS16 | RFLkSPAyRDLAAQA | Y177 | 0.094 | 96.66 | 202 | 0.041236 | 0.094 | 0.0412 | 1 |
| P19174 | PLCG1 | VLTSSKIyYSEETSs | Y518 | 0.05 | 91.99 | 1290 | 0.021934 | 0.05 | 0.0219 | 1 |
| Q14195 | DPYSL3 | TNCPLyVtKVMsKsA | T253 | 0.043 | 98.54 | 570 | 0.02937 | 0.043 | 0.0294 | 0.875 |
| Q14195 | DPYSL3 | KsAADLIsQARKKGN | S265 | 0.069 | 98.67 | 570 | 0.054849 | 0.069 | 0.0548 | 0.8333 |
| Q14195 | DPYSL3 | FGEPITAsLGIDGTH | S282 | 0.022 | 98.62 | 570 | 0.017488 | 0.022 | 0.0175 | 0.9091 |
| Q9NZZ3 | CHMP5 | RDNLAQQsFNMEQAN | S86 | 0.061 | 96.31 | 219 | 0.04849 | 0.061 | 0.0485 | 1 |
| P06727 | APOA4 | EELKGRLtPYADEFK | T202 | 0.212 | 88.02 | 396 | 0.144799 | 0.212 | 0.1448 | 1 |
| P06727 | APOA4 | LKARIsAsAEELRQR | S259 | 0.081 | 95.13 | 396 | 0.064388 | 0.081 | 0.0644 | 1 |
| Q9UBF2 | COPG2 | HFGTTEAtEAFFAMT | T67 | 0.18 | 90.03 | 871 | 0.122942 | 0.18 | 0.1229 | 1 |
| Q9UBF2 | COPG2 | IKEMATIsEDVIIVT | S99 | 0.125 | 88.31 | 871 | 0.099364 | 0.125 | 0.0994 | 1 |
| P61106 | RAB14 | RAVTRSyyRGAAGAL | Y81 | 0.047 | 88.93 | 215 | 0.020618 | 0.047 | 0.0206 | 0.9167 |
| P06241 | FYN | ETtkGAysLsIRDWD | S186 | 0 | 97.25 | 537 | 0 | 0.12 | 0.0954 | 1 |
| P06241 | FYN | KLDNGGyyITtRAQF | Y214 | 0.303 | 91.32 | 537 | 0.132922 | 0.303 | 0.1329 | 1 |
| P06241 | FYN | FGEVWMGtWNGNtKV | T289 | 0.173 | 96.47 | 537 | 0.118161 | 0.173 | 0.1182 | 1 |
| P06241 | FYN | MGtWNGNtKVAIktL | T294 | 0.093 | 90.16 | 537 | 0.06352 | 0.093 | 0.0635 | 0.8333 |
| P06241 | FYN | NtKVAIktLkPGTMs | T300 | 0.174 | 93.99 | 537 | 0.118844 | 0.174 | 0.1188 | 1 |

| | | | | | | | | | | |
|---|---|---|---|---|---|---|---|---|---|---|
| P06241 | FYN | VVSEEPIyIVTEYMN | Y339 | 0.291 | 95.09 | 537 | 0.127658 | 0.291 | 0.1277 | 1 |
| Q9Y2R9 | MRPS7 | DPVISKFtNMMMIGG | T95 | 0.098 | 97.79 | 242 | 0.066935 | 0.098 | 0.0669 | 1 |
| Q05048 | CSTF1 | VFNHtEDyVLLPDER | Y367 | 0.124 | 95.08 | 431 | 0.054397 | 0.124 | 0.0544 | 1 |
| Q01415 | GALK2 | YQRAkHVySEAARVL | Y336 | 0.003 | 98.89 | 458 | 0.001316 | 0.003 | 0.0013 | 1 |
| Q9BR01 | SULT4A1 | PKDLVVSyYQFHRSL | Y142 | 0.05 | 93.73 | 284 | 0.021934 | 0.083 | 0.0364 | 0.8571 |
| Q06210 | GFPT1 | DFESETDtETIAkLV | T149 | 0.018 | 97.28 | 699 | 0.012294 | 0.018 | 0.0123 | 1 |
| Q06210 | GFPT1 | FPGQAVGtRRGsPLL | T201 | 0.002 | 98.33 | 699 | 0.001366 | 0.002 | 0.0014 | 1 |
| Q06210 | GFPT1 | AVGtRRGsPLLIGVR | S205 | 0.064 | 89.77 | 699 | 0.050875 | 0.064 | 0.0509 | 0.8889 |
| Q06210 | GFPT1 | VEEKAVEyyFASDAS | Y275 | 0.002 | 98.75 | 699 | 0.000877 | 0.002 | 0.0009 | 1 |
| Q06210 | GFPT1 | EEKAVEyyFASDASA | Y276 | 0.018 | 98.73 | 699 | 0.007896 | 0.018 | 0.0079 | 1 |
| Q06210 | GFPT1 | GITNtVGsSIsRETD | S470 | 0.022 | 88.33 | 699 | 0.017488 | 0.022 | 0.0175 | 1 |
| P06239 | LCK | GEWWkAQsLTTGQEG | S102 | 0.019 | 97.45 | 509 | 0.015103 | 0.019 | 0.0151 | 1 |
| P06239 | LCK | APGNTHGsFLIREsE | S150 | 0.027 | 97.98 | 509 | 0.021463 | 0.027 | 0.0215 | 1 |
| P06239 | LCK | GsFLIREsEstAGsF | S156 | 0.113 | 91.93 | 509 | 0.089825 | 0.113 | 0.0898 | 1 |
| P06239 | LCK | EsEstAGsFSLSVRD | S162 | 0.133 | 90.97 | 509 | 0.105724 | 0.133 | 0.1057 | 1 |
| P06239 | LCK | LHELVRHytNAsDGL | Y209 | 0.011 | 98.01 | 509 | 0.004826 | 0.011 | 0.0048 | 1 |
| P06239 | LCK | GEVWMGyyNGHtkVA | Y264 | 0.12 | 94.14 | 509 | 0.052642 | 0.12 | 0.0526 | 0.8462 |
| P11362 | FGFR1 | SKGNLREyLQARRPP | Y572 | 0.153 | 94.93 | 822 | 0.067119 | 0.209 | 0.0917 | 1 |
| B2RTY4 | MYO9A | PLMTFELyEEFLRAM | Y2151 | 0.034 | 88.42 | 2548 | 0.014915 | 0.034 | 0.0149 | 1 |
| P78356 | PIP4K2B | SSPKKEVyFMAIIDI | Y363 | 0 | 97.97 | 416 | 0 | 0 | 0 | 1 |
| Q6P3W7 | SCYL2 | NKQDIyKsFSRQLDQ | S277 | 0.163 | 82.69 | 929 | 0.129571 | 0.163 | 0.1296 | 1 |
| Q9Y5H3 | PCDHGA10 | RsGTVLVsVtVFDAN | S233 | 0.185 | 94.01 | 936 | 0.147059 | 0.185 | 0.1471 | 1 |
| Q01469 | FABP5 | NVTCTRIyEKVE___ | Y131 | 0.182 | 98.63 | 135 | 0.079841 | 0.27 | 0.1184 | 0.9167 |
| Q8NHH9 | ATL2 | LLDFMLRyMyNKDSQ | Y117 | 0.202 | 95.87 | 583 | 0.088614 | 0.202 | 0.0886 | 1 |
| Q8NHH9 | ATL2 | EAEIEETyANFIkHN | Y456 | 0.277 | 96.16 | 583 | 0.121516 | 0.277 | 0.1215 | 1 |
| Q9NSN8 | SNTG1 | TKAVLWRyKFSQLkG | Y434 | 0.15 | 97.12 | 517 | 0.065803 | 0.15 | 0.0658 | 0.8889 |
| O15270 | SPTLC2 | IDYLRTHsHSAVYAt | S402 | 0.005 | 90.6 | 562 | 0.003975 | 0.005 | 0.004 | 1 |
| O15270 | SPTLC2 | VVVGFPAtPIIESRA | T501 | 0.045 | 88.18 | 562 | 0.030736 | 0.045 | 0.0307 | 1 |
| Q2M3G0 | ABCB5 | RALNVRHyRDHIGVV | Y460 | 0.032 | 93.41 | 1257 | 0.014038 | 0.032 | 0.014 | 1 |
| Q2M3G0 | ABCB5 | VLFGTtIsNNIKYGR | S479 | 0.11 | 93.98 | 1257 | 0.087441 | 0.11 | 0.0874 | 1 |
| Q0Z7S8 | FABP9 | ADNRkVKsTITLENG | S83 | 0.001 | 98.84 | 132 | 0.000795 | 0.001 | 0.0008 | 0.8571 |

| | | | | | | | | | | |
|---|---|---|---|---|---|---|---|---|---|---|
| Q8NCC3 | PLA2G15 | PQAWKDKyIRAFVsL | Y219 | 0.279 | 98.83 | 412 | 0.122393 | 0.279 | 0.1224 | 1 |
| Q8NCC3 | PLA2G15 | KyIRAFVsLGAPWGG | S225 | 0.013 | 98.89 | 412 | 0.010334 | 0.013 | 0.0103 | 1 |
| Q96P16 | RPRD1A | QQSVQTLsLWLIHHR | S27 | 0.01 | 97.1 | 312 | 0.007949 | 0.01 | 0.0079 | 1 |
| Q96P16 | RPRD1A | LIHHRKHsRPIVTVW | S37 | 0.09 | 97.11 | 312 | 0.071542 | 0.09 | 0.0715 | 1 |
| P08100 | RHO | AIyNPVIyIMMNKQF | Y306 | 0.054 | 77.11 | 348 | 0.023689 | 0.054 | 0.0237 | 0.8333 |
| P43246 | MSH2 | LTsLNEEyTkNkTEy | Y563 | 0.196 | 92.27 | 934 | 0.085982 | 0.196 | 0.086 | 0.8333 |
| P43246 | MSH2 | LRsAtkDsLIIIDEL | S743 | 0 | 96.31 | 934 | 0 | 0 | 0 | 0.8 |
| Q04323 | UBXN1 | QDPVQLLsGFPRRAF | S262 | 0.182 | 92.19 | 297 | 0.144674 | 0.182 | 0.1447 | 0.875 |
| Q07864 | POLE | RLGSAVDyyFIQDDG | Y84 | 0 | 94.27 | 2286 | 0 | 0 | 0 | 1 |
| Q07864 | POLE | LGSAVDyyFIQDDGS | Y85 | 0.029 | 94.74 | 2286 | 0.012722 | 0.029 | 0.0127 | 1 |
| Q07864 | POLE | kRNYIRLsFHTVEDL | S156 | 0.072 | 95.17 | 2286 | 0.057234 | 0.144 | 0.1145 | 1 |
| P09972 | ALDOC | PEEIAMAtVtALRRT | T253 | 0.021 | 98.87 | 364 | 0.014343 | 0.042 | 0.0287 | 0.8 |
| P09972 | ALDOC | GQSEEEAsFNLNAIN | S281 | 0 | 97.62 | 364 | 0 | 0 | 0 | 1 |
| P48723 | HSPA13 | FDLKQRNstIEAANL | S185 | 0.105 | 97.73 | 471 | 0.083466 | 0.105 | 0.0835 | 1 |
| P48723 | HSPA13 | DLKQRNstIEAANLA | T186 | 0.007 | 97.29 | 471 | 0.004781 | 0.007 | 0.0048 | 1 |
| P48723 | HSPA13 | LRVINEPtAAAMAyG | T205 | 0.019 | 94.05 | 471 | 0.012977 | 0.019 | 0.013 | 1 |
| P48723 | HSPA13 | PtAAAMAyGLHKADV | Y211 | 0.218 | 91.43 | 471 | 0.095633 | 0.218 | 0.0956 | 1 |
| O00429 | DNM1L | GKSsVLEsLVGRDLL | S44 | 0.14 | 89.88 | 736 | 0.111288 | 0.14 | 0.1113 | 1 |
| O00429 | DNM1L | PIHLkIFsPNVVNLT | S136 | 0.021 | 87.65 | 736 | 0.016693 | 0.042 | 0.0334 | 1 |
| O00429 | DNM1L | RFISNPNsIILAVTA | S179 | 0 | 89.95 | 736 | 0 | 0 | 0 | 1 |
| O00429 | DNM1L | tSEALKIsREVDPDG | S200 | 0.023 | 92.68 | 736 | 0.018283 | 0.023 | 0.0183 | 1 |
| O00429 | DNM1L | ALFVPEVsFELLVKR | S412 | 0.025 | 89.09 | 736 | 0.019873 | 0.025 | 0.0199 | 1 |
| Q8TDG4 | HELQ | RGATLEMtLAKILYT | T480 | 0.009 | 96.76 | 1101 | 0.006147 | 0.009 | 0.0061 | 1 |
| Q8TDG4 | HELQ | EFLKRNQykQMIGRA | Y701 | 0.023 | 92.01 | 1101 | 0.01009 | 0.023 | 0.0101 | 1 |
| O60704 | TPST2 | DGRAsVHsMItRKVT | S191 | 0.038 | 98.15 | 377 | 0.030207 | 0.038 | 0.0302 | 0.875 |
| P00966 | ASS1 | QGyDVIAyLANIGQk | Y34 | 0.024 | 98.68 | 412 | 0.010528 | 0.024 | 0.0105 | 1 |
| P00966 | ASS1 | SALyEDRyLLGTsLA | Y87 | 0.049 | 98.55 | 412 | 0.021496 | 0.049 | 0.0215 | 0.8667 |
| P00966 | ASS1 | DQVRFELsCysLAPQ | S131 | 0.014 | 98.41 | 412 | 0.011129 | 0.014 | 0.0111 | 0.8333 |
| P00966 | ASS1 | GrNDLMEyAkQHGIP | Y163 | 0.071 | 95.59 | 412 | 0.031147 | 0.071 | 0.0311 | 1 |
| P00966 | ASS1 | NrFIGMKsRGIYETP | S278 | 0.166 | 95.06 | 412 | 0.131956 | 0.166 | 0.132 | 1 |
| Q9BYT8 | NLN | PEQIkTRtEELIVQT | T61 | 0.081 | 98.72 | 704 | 0.055324 | 0.118 | 0.0913 | 1 |

| | | | | | | | | | | |
|---|---|---|---|---|---|---|---|---|---|---|
| Q9BYT8 | NLN | QPVAGRPsLLRHDEV | S485 | 0.011 | 93.7 | 704 | 0.008744 | 0.011 | 0.0087 | 1 |
| Q9BYT8 | NLN | LVNtGLLtLRQIVLS | T575 | 0.078 | 96.92 | 704 | 0.053275 | 0.078 | 0.0533 | 0.8 |
| Q9BYT8 | NLN | NPEVGMkyRNLILKP | Y665 | 0.011 | 98.84 | 704 | 0.004826 | 0.011 | 0.0048 | 1 |
| Q9Y2A7 | NCKAP1 | LDLIITYttLMILLs | T141 | 0 | 98.36 | 1128 | 0 | 0 | 0 | 0.875 |
| Q9Y2A7 | NCKAP1 | HGAsDREyPRLGQMI | Y175 | 0.164 | 95.91 | 1128 | 0.071944 | 0.318 | 0.1371 | 1 |
| Q9Y2A7 | NCKAP1 | FHTKMVDsLVEMLVE | S528 | 0.052 | 97.79 | 1128 | 0.041336 | 0.052 | 0.0413 | 1 |
| Q8TDN6 | BRIX1 | AFDELPHyALLKELL | Y178 | 0.16 | 95.13 | 353 | 0.07019 | 0.16 | 0.0702 | 1 |
| Q9P260 | RELCH | LIPLILCtACLHPEP | T549 | 0.007 | 96 | 1216 | 0.004781 | 0.007 | 0.0048 | 1 |
| Q8NI27 | THOC2 | TkENVKPsGRQIGKL | S564 | 0.078 | 92.25 | 1593 | 0.062003 | 0.078 | 0.062 | 1 |
| Q8NI27 | THOC2 | LItPVVDsLKYLTSL | S600 | 0.016 | 92.53 | 1593 | 0.012719 | 0.016 | 0.0127 | 1 |
| Q96RK4 | BBS4 | NLNRHDLtyIMLGKI | T170 | 0 | 93.23 | 519 | 0 | 0 | 0 | 0.8182 |
| Q68CZ2 | TNS3 | CRPFLKLyQAMQPVY | Y204 | 0.008 | 96.2 | 1445 | 0.003509 | 0.008 | 0.0035 | 1 |
| Q9UHB6 | LIMA1 | AsLHGRIyCKPHFNQ | Y437 | 0.169 | 94.42 | 759 | 0.074138 | 0.169 | 0.0741 | 1 |
| Q02790 | FKBP4 | PMEGVDIsPkQDEGV | S26 | 0.018 | 96.66 | 459 | 0.014308 | 0.018 | 0.0143 | 1 |
| Q02790 | FKBP4 | RMEKGEHsIVyLkPS | S217 | 0.002 | 98.1 | 459 | 0.00159 | 0.002 | 0.0016 | 1 |
| Q02790 | FKBP4 | ykQALLQykKIVsWL | Y293 | 0.041 | 97.76 | 459 | 0.017986 | 0.041 | 0.018 | 1 |
| Q02790 | FKBP4 | AFsAAIEsCNkALEL | S341 | 0.111 | 98.62 | 459 | 0.088236 | 0.111 | 0.0882 | 1 |
| Q6ZMZ3 | SYNE3 | LLEQVAGsMRDRDLL | S525 | 0.133 | 88.57 | 975 | 0.105724 | 0.133 | 0.1057 | 1 |
| Q96MS0 | ROBO3 | VPGLLYRtLVAAATS | T843 | 0 | 88.41 | 1386 | 0 | 0 | 0 | 1 |
| Q9NQX3 | GPHN | GFAPRDVtPEAtKEV | T96 | 0.001 | 97.94 | 736 | 0.000683 | 0.001 | 0.0007 | 1 |
| Q9NQX3 | GPHN | VCGIRGKtLIINLPG | T139 | 0.027 | 96.91 | 736 | 0.018441 | 0.027 | 0.0184 | 1 |
| Q8NBZ7 | UXS1 | DRkRILItGGAGFVG | T94 | 0 | 98.91 | 420 | 0 | 0 | 0 | 1 |
| Q8NBZ7 | UXS1 | GGAGFVGsHLtDKLM | S102 | 0 | 98.72 | 420 | 0 | 0 | 0 | 1 |
| Q8NBZ7 | UXS1 | GFVGsHLtDKLMMDG | T105 | 0 | 98.87 | 420 | 0 | 0 | 0 | 1 |
| Q8NBZ7 | UXS1 | YMYNPIKtLKtNTIG | T175 | 0.021 | 98.35 | 420 | 0.014343 | 0.021 | 0.0143 | 1 |
| Q8NBZ7 | UXS1 | NPIKtLKtNTIGtLN | T178 | 0.122 | 98.67 | 420 | 0.083328 | 0.218 | 0.1489 | 1 |
| Q8NBZ7 | UXS1 | LKtNTIGtLNMLGLA | T183 | 0.026 | 98.85 | 420 | 0.017758 | 0.052 | 0.0355 | 1 |
| O14641 | DVL2 | ERREARkyASGLLkA | Y478 | 0.227 | 94.78 | 736 | 0.099582 | 0.343 | 0.1109 | 0.8667 |
| Q9HA77 | CARS2 | STAKGNVyFDLkSRG | Y194 | 0.225 | 98.56 | 564 | 0.098704 | 0.225 | 0.0987 | 1 |
| Q9GZN7 | ROGDI | DQVKGVLtLQGDALs | T76 | 0.031 | 95.39 | 287 | 0.021173 | 0.031 | 0.0212 | 1 |
| P09936 | UCHL1 | FPVNHGAssEDtLLk | S188 | 0.162 | 98.08 | 223 | 0.128776 | 0.162 | 0.1288 | 1 |

| | | | | | | | | | | |
|---|---|---|---|---|---|---|---|---|---|---|
| Q53EL9 | SEZ6 | RFHCATGyQLKGARH | Y387 | 0.252 | 91.05 | 994 | 0.110549 | 0.252 | 0.1105 | 1 |
| Q53EL9 | SEZ6 | PGDVLTFyDGDDLTA | Y649 | 0.293 | 96.76 | 994 | 0.128535 | 0.293 | 0.1285 | 1 |
| Q9UBM7 | DHCR7 | GIQEGAVtPAGVVNK | T135 | 0.101 | 89.6 | 475 | 0.068984 | 0.101 | 0.069 | 1 |
| Q9UBM7 | DHCR7 | HHSKLLVsGFWGVAR | S397 | 0.184 | 96.79 | 475 | 0.146264 | 0.184 | 0.1463 | 1 |
| Q9UHF3 | NAT8B | MAEHAPAtFRRLLKL | T32 | 0.061 | 92.84 | 227 | 0.041664 | 0.061 | 0.0417 | 0.8333 |
| P22459 | KCNA4 | FFDRNRPsFDAILyY | S227 | 0 | 96.88 | 653 | 0 | 0 | 0 | 0.875 |
| P02751 | FN1 | TTPDITGyRITTTPT | Y1206 | 0.014 | 90.91 | 2477 | 0.006142 | 0.014 | 0.0061 | 1 |
| Q504Q3 | PAN2 | TCEkYQPtIQtRNIR | T728 | 0.059 | 89.23 | 1202 | 0.040298 | 0.059 | 0.0403 | 1 |
| Q15063 | POSTN | IIKPAEKsLHEkLKQ | S498 | 0.17 | 94.23 | 836 | 0.135135 | 0.17 | 0.1351 | 1 |
| Q15063 | POSTN | SKESDIMtTNGVIHV | T616 | 0.004 | 96.02 | 836 | 0.002732 | 0.004 | 0.0027 | 1 |
| Q8TE68 | EPS8L1 | EPQLsPGsDASEVRA | S634 | 0.068 | 91.25 | 723 | 0.054054 | 0.068 | 0.0541 | 1 |
| Q8TE68 | EPS8L1 | KEELRAVsPEEGARV | S676 | 0.083 | 86.18 | 723 | 0.065978 | 0.083 | 0.066 | 1 |
| O95470 | SGPL1 | PEIVAPQsAHAAFNk | S240 | 0.071 | 98.68 | 568 | 0.056439 | 0.071 | 0.0564 | 1 |
| O95470 | SGPL1 | GSRPGGIsAACWAAL | S401 | 0.059 | 98.8 | 568 | 0.0469 | 0.059 | 0.0469 | 1 |
| O95470 | SGPL1 | VFGNPQLsVIALGsR | S449 | 0.028 | 98.68 | 568 | 0.022258 | 0.028 | 0.0223 | 1 |
| O95470 | SGPL1 | FLKDIREsVtQIMKN | S507 | 0 | 98.7 | 568 | 0 | 0 | 0 | 1 |
| Q2M389 | WASHC4 | RsMVVADsVSHITQH | S500 | 0.001 | 86.81 | 1173 | 0.000795 | 0.001 | 0.0008 | 1 |
| Q92997 | DVL3 | sYKFFFKsMDDDFGV | S48 | 0.071 | 91.62 | 716 | 0.056439 | 0.071 | 0.0564 | 0.8 |
| Q96DU9 | PABPC5 | IRDAsGKsKGFGFVR | S238 | 0.132 | 95.94 | 382 | 0.104929 | 0.132 | 0.1049 | 1 |
| P51798 | CLCN7 | CALIGILtGLVACFI | T137 | 0.061 | 96.43 | 805 | 0.041664 | 0.102 | 0.0674 | 0.9375 |
| O95260 | ATE1 | NQTCCPQyTIRCRPL | Y75 | 0.025 | 97.98 | 518 | 0.010967 | 0.05 | 0.0219 | 0.9333 |
| O95260 | ATE1 | FkATLLEsyQVYKRY | S291 | 0.028 | 98.41 | 518 | 0.022258 | 0.028 | 0.0223 | 1 |
| O95260 | ATE1 | HKRAIMPyGVYKKQQ | Y482 | 0.062 | 96.75 | 518 | 0.027199 | 0.062 | 0.0272 | 1 |
| O95260 | ATE1 | EEAAVLQyASLVGQk | Y501 | 0.001 | 97.62 | 518 | 0.000439 | 0.001 | 0.0004 | 0.8125 |
| O75325 | LRRN2 | ALPAGTQtLLLQSNS | T73 | 0.066 | 97.63 | 713 | 0.045079 | 0.066 | 0.0451 | 1 |
| O96019 | ACTL6A | LQQGIVksPLAGDFI | S195 | 0.025 | 95.28 | 429 | 0.019873 | 0.025 | 0.0199 | 1 |
| O94806 | PRKD3 | RLDSKCLtLFQNESG | T446 | 0.083 | 93.56 | 890 | 0.05669 | 0.083 | 0.0567 | 1 |
| Q00872 | MYBPC1 | AAEYSVMttGGQSSA | T417 | 0.213 | 89.4 | 1141 | 0.145482 | 0.213 | 0.1455 | 1 |
| Q00872 | MYBPC1 | AIGISKPsMPSRPFV | S706 | 0.068 | 97.04 | 1141 | 0.054054 | 0.068 | 0.0541 | 1 |
| O75976 | CPD | ATGIYSktsDDEVFk | T282 | 0.033 | 95.21 | 1380 | 0.022539 | 0.033 | 0.0225 | 0.8571 |
| O75976 | CPD | LELSCCKyPPASQLR | Y356 | 0.083 | 92.87 | 1380 | 0.036411 | 0.133 | 0.0503 | 0.8462 |

| | | | | | | | | | | |
|---|---|---|---|---|---|---|---|---|---|---|
| Q07890 | SOS2 | PVyHCWHyFELLKQL | Y351 | 0.014 | 94.43 | 1332 | 0.006142 | 0.014 | 0.0061 | 0.9286 |
| Q07890 | SOS2 | LKQLKACsEEQEDRE | S362 | 0.067 | 92.44 | 1332 | 0.053259 | 0.067 | 0.0533 | 1 |
| P68400 | CSNK2A1 | _MsGPVPsRARVytD | S7 | 0.069 | 95.59 | 391 | 0.054849 | 0.069 | 0.0548 | 0.8571 |
| P68400 | CSNK2A1 | GTEDLyDyIDkYNIE | Y257 | 0.048 | 98.35 | 391 | 0.021057 | 0.095 | 0.0417 | 0.9231 |
| P68400 | CSNK2A1 | REAMEHPyFYTVVkD | Y323 | 0.001 | 98.59 | 391 | 0.000439 | 0.001 | 0.0004 | 1 |
| Q13188 | STK3 | LKkLsEDsLTkQPEE | S18 | 0.167 | 90.89 | 491 | 0.132751 | 0.167 | 0.1328 | 1 |
| P62987 | UBA52 | DyNIQkEstLHLVLr | S65 | 0.17 | 96.63 | 128 | 0.135135 | 0.17 | 0.1351 | 1 |
| P35573 | AGL | NEKSGGGyIVVDPIL | Y106 | 0.211 | 96.1 | 1532 | 0.092563 | 0.211 | 0.0926 | 1 |
| P35573 | AGL | AEPGSEVyLRRELIC | Y479 | 0.029 | 96.15 | 1532 | 0.012722 | 0.029 | 0.0127 | 1 |
| P35573 | AGL | EAMsAyNsHEEGRLV | S586 | 0.041 | 92.77 | 1532 | 0.032592 | 0.041 | 0.0326 | 1 |
| P35573 | AGL | FPSLPILsPALMDVP | S1060 | 0.012 | 94.54 | 1532 | 0.009539 | 0.012 | 0.0095 | 1 |
| P35573 | AGL | IILAFAGtLRHGLIP | T1130 | 0 | 95.93 | 1532 | 0 | 0 | 0 | 1 |
| P35573 | AGL | TAGVDEEtGFVYGGN | T1245 | 0.111 | 96.77 | 1532 | 0.075815 | 0.111 | 0.0758 | 1 |
| P35573 | AGL | RNRGIPAtPRDGSAV | T1277 | 0.045 | 94.57 | 1532 | 0.030736 | 0.061 | 0.0487 | 1 |
| P35573 | AGL | AVEIVGLskSAVRWL | S1290 | 0 | 98.2 | 1532 | 0 | 0 | 0 | 1 |
| P35573 | AGL | LRAKLyFsRLMGPET | S1466 | 0.003 | 96.92 | 1532 | 0.002385 | 0.003 | 0.0024 | 1 |
| P25189 | MPZ | DAIsIFHyAKGQPYI | Y82 | 0.246 | 97.17 | 248 | 0.107917 | 0.246 | 0.1079 | 1 |
| Q9UH90 | FBXO40 | yGCPLsMsRHKLAKH | S77 | 0.103 | 97.21 | 709 | 0.081876 | 0.103 | 0.0819 | 1 |
| Q9UL12 | SARDH | FGFAIDKtIAyGyIH | T864 | 0.027 | 98.18 | 918 | 0.018441 | 0.027 | 0.0184 | 1 |
| Q9UL12 | SARDH | AIDKtIAyGyIHDPS | Y867 | 0.077 | 98.43 | 918 | 0.033779 | 0.077 | 0.0338 | 1 |
| Q9UL12 | SARDH | DKtIAyGyIHDPSGG | Y869 | 0.183 | 98.02 | 918 | 0.080279 | 0.183 | 0.0803 | 1 |
| P62258 | YWHAE | RkEAAENsLVAykAA | S148 | 0 | 98.77 | 255 | 0 | 0 | 0 | 1 |
| P62258 | YWHAE | AENsLVAykAASDIA | Y152 | 0 | 98.81 | 255 | 0 | 0 | 0 | 1 |
| Q13630 | GFUS | GsMRILVtGGsGLVG | T13 | 0 | 98.89 | 321 | 0 | 0 | 0 | 1 |
| Q9UMY4 | SNX12 | ESCVRRRysDFEWLk | Y72 | 0.273 | 97.03 | 162 | 0.119761 | 0.273 | 0.1198 | 1 |
| O76003 | GLRX3 | AYSsWPTyPQLYVSG | Y200 | 0.166 | 91.27 | 335 | 0.072822 | 0.166 | 0.0728 | 1 |
| Q96QE2 | SLC2A13 | PVYIAEVsPPNLRGR | S200 | 0.007 | 83.48 | 648 | 0.005564 | 0.007 | 0.0056 | 0.9 |
| Q96QE2 | SLC2A13 | DPDCGFCyKMNKSTV | Y455 | 0.034 | 92.87 | 648 | 0.014915 | 0.034 | 0.0149 | 1 |
| Q15147 | PLCB4 | SDKKIGTyVEVDMYG | Y721 | 0.219 | 97.58 | 1175 | 0.096072 | 0.219 | 0.0961 | 1 |
| P06213 | INSR | EIADGMAyLNAKKFV | Y1149 | 0.026 | 90.89 | 1382 | 0.011406 | 0.184 | 0.0807 | 0.8889 |
| O43181 | NDUFS4 | KSYGANFsWNkRTRV | S165 | 0.187 | 95.29 | 175 | 0.148649 | 0.187 | 0.1486 | 1 |

| | | | | | | | | | | |
|---|---|---|---|---|---|---|---|---|---|---|
| P46977 | STT3A | FDDFREAyyWLRHNT | Y509 | 0.024 | 95.72 | 705 | 0.010528 | 0.024 | 0.0105 | 1 |
| P46977 | STT3A | kHIkENDyytPtGEF | Y617 | 0.09 | 92.98 | 705 | 0.039482 | 0.09 | 0.0395 | 1 |
| P46977 | STT3A | IkENDyytPtGEFRV | T619 | 0.132 | 90.78 | 705 | 0.090158 | 0.132 | 0.0902 | 1 |
| P46977 | STT3A | YRFGQVytEAkRPPG | T652 | 0.12 | 82.76 | 705 | 0.081962 | 0.12 | 0.082 | 1 |
| Q13133 | NR1H3 | LLETSRRyNPGsEsI | Y306 | 0.181 | 96.91 | 447 | 0.079402 | 0.181 | 0.0794 | 1 |
| Q13133 | NR1H3 | RyNPGsEsItFLKDF | S312 | 0.102 | 96.73 | 447 | 0.081081 | 0.102 | 0.0811 | 1 |
| P11488 | GNAT1 | LLLGAGEsGkSTIVk | S40 | 0.031 | 97.47 | 350 | 0.024642 | 0.031 | 0.0246 | 1 |
| P08567 | PLEK | LQPAGDMsKSAVDGT | S197 | 0 | 90.75 | 350 | 0 | 0 | 0 | 0.9 |
| P10599 | TXN | kPFFHsLsEkYSNVI | S46 | 0.106 | 97.98 | 105 | 0.084261 | 0.106 | 0.0843 | 1 |
| Q7Z6Z7 | HUWE1 | IkAASLRtLtsIVHL | T330 | 0.013 | 93.195 | 4374 | 0.008879 | 0.013 | 0.0089 | 1 |
| Q7Z6Z7 | HUWE1 | ASLRtLtsIVHLERT | S333 | 0.028 | 89.66 | 4374 | 0.022258 | 0.028 | 0.0223 | 0.9 |
| P52333 | JAK3 | PAKkLQFyEDRQQLP | Y738 | 0.204 | 94.47 | 1124 | 0.089492 | 0.204 | 0.0895 | 0.8 |
| Q8IY47 | KBTBD2 | HkVGtVVtPDNDIyI | T313 | 0.159 | 94.85 | 623 | 0.108599 | 0.192 | 0.1311 | 1 |
| Q9Y697 | NFS1 | THAYGWEsEAAMERA | S99 | 0 | 98.57 | 457 | 0 | 0 | 0 | 1 |
| Q9Y697 | NFS1 | RFGIGRFttEEEVDy | T414 | 0.074 | 98.81 | 457 | 0.050543 | 0.147 | 0.1004 | 1 |
| Q9Y697 | NFS1 | tEEEVDytVEkCIQH | T422 | 0.001 | 98.62 | 457 | 0.000683 | 0.001 | 0.0007 | 0.9091 |
| Q9BZ29 | DOCK9 | VEtQVGysWLPLLKD | S758 | 0.013 | 95.05 | 2069 | 0.010334 | 0.013 | 0.0103 | 1 |
| O15144 | ARPC2 | LkRNCFAsVFEkYFQ | S123 | 0.042 | 96.78 | 300 | 0.033386 | 0.042 | 0.0334 | 1 |
| O15144 | ARPC2 | DEtMyVEskKDRVTV | S156 | 0.093 | 96.66 | 300 | 0.073927 | 0.093 | 0.0739 | 1 |
| Q9UHR6 | ZNHIT2 | ALGDLAQtLGrARkQ | T316 | 0.113 | 94.18 | 403 | 0.077181 | 0.113 | 0.0772 | 1 |
| P36776 | LONP1 | IIQCLKKtkTENPLI | T580 | 0.158 | 93.68 | 959 | 0.107916 | 0.158 | 0.1079 | 1 |
| O43252 | PAPSS1 | MVAGANFyIVGRDPA | Y518 | 0.048 | 98.84 | 624 | 0.021057 | 0.048 | 0.0211 | 1 |
| O43252 | PAPSS1 | PEtGKDLyEPSHGAK | Y537 | 0.282 | 98.29 | 624 | 0.123709 | 0.282 | 0.1237 | 1 |
| P09874 | PARP1 | RMAIMVQsPMFDGKV | S41 | 0.054 | 86.24 | 1014 | 0.042925 | 0.054 | 0.0429 | 1 |
| Q9NTM9 | CUTC | LGFERVLtSGCDSSA | T171 | 0 | 98.79 | 273 | 0 | 0 | 0 | 1 |
| Q96IP4 | TENT5A | RLNGSAAsHVLHQDS | S126 | 0.036 | 96.32 | 442 | 0.028617 | 0.036 | 0.0286 | 1 |
| Q96IP4 | TENT5A | LEDRKyEyLMTLHGV | Y354 | 0.006 | 98.16 | 442 | 0.002632 | 0.006 | 0.0026 | 1 |
| P07737 | PFN1 | FNVtVtktDktLVLL | T106 | 0.058 | 96.43 | 140 | 0.039615 | 0.058 | 0.0396 | 0.9 |
| P28074 | PSMB5 | DLEVEQAyDLARRAI | Y212 | 0.194 | 97.71 | 263 | 0.085105 | 0.194 | 0.0851 | 0.9091 |
| P28074 | PSMB5 | DLARRAIyQATYRDA | Y220 | 0.092 | 95.82 | 263 | 0.040359 | 0.092 | 0.0404 | 1 |
| P02549 | SPTA1 | YGKDLVAsEGLFHSH | S299 | 0 | 84.87 | 2419 | 0 | 0 | 0 | 1 |

| | | | | | | | | | | |
|---|---|---|---|---|---|---|---|---|---|---|
| Q9H0S4 | DDX47 | RLFALVLtPTRELAF | T99 | 0 | 94.98 | 455 | 0 | 0 | 0 | 0.8182 |
| Q9H0S4 | DDX47 | VIPRDRKtFLFSATM | T201 | 0.01 | 98.19 | 455 | 0.00683 | 0.01 | 0.0068 | 0.875 |
| Q96RV3 | PCNX1 | RKVLTTYyVKGIIYY | Y1700 | 0.02 | 95.29 | 2341 | 0.008774 | 0.02 | 0.0088 | 1 |
| Q07002 | CDK18 | GFGKLEtyVkLDkLG | Y144 | 0.231 | 91.77 | 474 | 0.101336 | 0.298 | 0.1307 | 1 |
| Q07002 | CDK18 | LGEGtyAtVFKGRSk | T157 | 0.116 | 84.47 | 474 | 0.07923 | 0.116 | 0.0792 | 1 |
| Q9UHN1 | POLG2 | NVVPCVLsVNGDLDR | S337 | 0.14 | 95.43 | 485 | 0.111288 | 0.14 | 0.1113 | 1 |
| O94830 | DDHD2 | KDNKYVPysESFSQV | Y128 | 0.146 | 94.27 | 711 | 0.064048 | 0.226 | 0.0991 | 1 |
| Q99941 | ATF6B | KTSRPKMsLVMPAMA | S618 | 0 | 94.05 | 703 | 0 | 0 | 0 | 1 |
| Q8NFU1 | BEST2 | LMCVVAGtVHGRDDR | T113 | 0.174 | 97.54 | 509 | 0.118844 | 0.174 | 0.1188 | 1 |
| Q96SB4 | SRPK1 | LIVAGkYskEFFTKK | S587 | 0.075 | 97.25 | 655 | 0.059619 | 0.075 | 0.0596 | 1 |
| Q92823 | NRCAM | PSEASEQyLTKASEP | Y739 | 0.264 | 90.01 | 1304 | 0.115813 | 0.264 | 0.1158 | 1 |
| Q92823 | NRCAM | IRGHLQGyRIYYWKT | Y881 | 0.01 | 93.22 | 1304 | 0.004387 | 0.071 | 0.0311 | 1 |
| Q96QS6 | PSKH2 | DPRVLARyDIKALIG | Y63 | 0.09 | 97.27 | 385 | 0.039482 | 0.09 | 0.0395 | 0.8571 |
| P08F94 | PKHD1 | ELLGGAMyYLEAEHH | Y452 | 0.167 | 95.81 | 4074 | 0.07326 | 0.167 | 0.0733 | 1 |
| P08F94 | PKHD1 | ADDFTSRyLNASDFT | Y828 | 0.312 | 90.68 | 4074 | 0.13687 | 0.312 | 0.1369 | 1 |
| P31997 | CEACAM8 | TRNDTGSyTLQVIkL | Y120 | 0.06 | 98.45 | 349 | 0.026321 | 0.06 | 0.0263 | 1 |
| Q8IWU9 | TPH2 | LLTKyCGyREDNVPQ | Y252 | 0.016 | 98.22 | 490 | 0.007019 | 0.016 | 0.007 | 1 |
| Q8IWU9 | TPH2 | AYGAGLLsSIGELKH | S382 | 0.044 | 94.35 | 490 | 0.034976 | 0.157 | 0.1248 | 1 |
| Q7Z7D3 | VTCN1 | QVIVGNAsLRLKNVQ | S114 | 0.102 | 98.36 | 282 | 0.081081 | 0.102 | 0.0811 | 1 |
| P30049 | ATP5F1D | QMsFTFAsPTQVFFN | S44 | 0.187 | 95.91 | 168 | 0.148649 | 0.187 | 0.1486 | 1 |
| Q8NHP8 | PLBD2 | AHNTWNNyQHMLRVI | Y267 | 0 | 97.93 | 589 | 0 | 0 | 0 | 1 |
| Q8NHP8 | PLBD2 | MSLARILsLLAASGP | S545 | 0.04 | 98.08 | 589 | 0.031797 | 0.04 | 0.0318 | 0.8571 |
| Q9NRM0 | SLC2A9 | LYFVLPEtKNRtYAE | T502 | 0.045 | 85.29 | 540 | 0.030736 | 0.045 | 0.0307 | 0.875 |
| Q9NRM0 | SLC2A9 | AEISQAFsKRNkAYP | S515 | 0.087 | 83.01 | 540 | 0.069158 | 0.087 | 0.0692 | 1 |
| P30532 | CHRNA5 | FEGTstktVIRYNGT | T150 | 0.001 | 95.66 | 468 | 0.000683 | 0.001 | 0.0007 | 1 |
| Q16825 | PTPN21 | EETIQFQtEDMETAK | T284 | 0.086 | 94.23 | 1174 | 0.058739 | 0.172 | 0.1175 | 1 |
| Q8N157 | AHI1 | EFkGIPIsyLEIHPN | S804 | 0.011 | 92.33 | 1196 | 0.008744 | 0.011 | 0.0087 | 1 |
| Q8N157 | AHI1 | LLIHTKDsTLRIMDL | S822 | 0.123 | 90.52 | 1196 | 0.097775 | 0.123 | 0.0978 | 1 |
| Q86WT6 | TRIM69 | DtLCPGLsPLTLDPK | S323 | 0.135 | 92.29 | 500 | 0.107313 | 0.135 | 0.1073 | 1 |
| Q659A1 | ICE2 | VKVIYINsPLPQKKM | S326 | 0.051 | 94.93 | 982 | 0.040541 | 0.11 | 0.0874 | 1 |
| Q9H2P9 | DPH5 | ______MLyLIGLGLG | Y3 | 0.077 | 98.37 | 285 | 0.033779 | 0.077 | 0.0338 | 1 |

| | | | | | | | | | | |
|---|---|---|---|---|---|---|---|---|---|---|
| Q6UWF7 | NXPE4 | DFLRARMsSPALMAG | S143 | 0.144 | 97.82 | 544 | 0.114468 | 0.144 | 0.1145 | 0.8333 |
| Q6UWF7 | NXPE4 | GYDRVIFtGQFVNGT | T205 | 0.196 | 98.21 | 544 | 0.133871 | 0.196 | 0.1339 | 1 |
| P20936 | RASA1 | KIMESKQsCELsPSK | S827 | 0.143 | 90.16 | 1047 | 0.113673 | 0.143 | 0.1137 | 1 |
| P20936 | RASA1 | SKQsCELsPSKLEKN | S831 | 0.091 | 88.93 | 1047 | 0.072337 | 0.091 | 0.0723 | 1 |
| Q10472 | GALNT1 | LFSIDRDyFQEIGTy | Y302 | 0.025 | 98.68 | 559 | 0.010967 | 0.025 | 0.011 | 1 |
| Q10472 | GALNT1 | yFQEIGTyDAGMDIW | Y309 | 0.019 | 96.5 | 559 | 0.008335 | 0.019 | 0.0083 | 1 |
| Q3SY69 | ALDH1L2 | HIGMSVQtFRYFAGW | T550 | 0 | 98.9 | 923 | 0 | 0 | 0 | 1 |
| Q6NUM6 | TYW1B | AVFGLGNsAyAsHFN | S136 | 0.167 | 87.53 | 668 | 0.132751 | 0.167 | 0.1328 | 1 |
| Q96JK2 | DCAF5 | RRLPPVLyDIHsRLP | Y260 | 0.08 | 95.36 | 942 | 0.035095 | 0.08 | 0.0351 | 1 |
| Q9BXI2 | SLC25A2 | LVKCRLQtMyEMEMS | T136 | 0 | 90.81 | 301 | 0 | 0 | 0 | 1 |
| P78417 | GSTO1 | PGPVPEGsIRIYSMr | S23 | 0.144 | 96.41 | 241 | 0.114468 | 0.144 | 0.1145 | 1 |
| P78417 | GSTO1 | LELyLQNsPEACDyG | S233 | 0.153 | 95.09 | 241 | 0.121622 | 0.153 | 0.1216 | 1 |
| P34896 | SHMT1 | MTDkkkIsATSIFFE | S161 | 0.124 | 97.34 | 483 | 0.098569 | 0.124 | 0.0986 | 1 |
| P34896 | SHMT1 | PDTGyINyDQLEENA | Y183 | 0.222 | 98.73 | 483 | 0.097388 | 0.222 | 0.0974 | 0.8182 |
| P34896 | SHMT1 | PSGLRLGtPALTSRG | T405 | 0.003 | 98.91 | 483 | 0.002049 | 0.003 | 0.002 | 1 |
| Q9P2E5 | CHPF2 | PGLVQkFsLRDCSPR | S729 | 0.187 | 85.94 | 772 | 0.148649 | 0.187 | 0.1486 | 1 |
| P07332 | FES | YRALARDsAQAKRKy | S149 | 0.066 | 93.55 | 822 | 0.052464 | 0.066 | 0.0525 | 1 |
| Q5VT52 | RPRD2 | NAIIFREsFADVLPE | S99 | 0.176 | 94.46 | 1461 | 0.139905 | 0.176 | 0.1399 | 0.8333 |
| Q9UBE8 | NLK | APEILMGsRHYSNAI | S315 | 0.016 | 88.61 | 527 | 0.012719 | 0.129 | 0.1025 | 1 |
| Q96J02 | ITCH | DQHGRVyyVDHVEkR | Y344 | 0.199 | 91.96 | 903 | 0.087298 | 0.199 | 0.0873 | 1 |
| Q96J02 | ITCH | DNMGRIyyVDHFTRT | Y376 | 0.122 | 95.91 | 903 | 0.05352 | 0.122 | 0.0535 | 0.9091 |
| Q9HAZ1 | CLK4 | EHHstLVstRHyRAP | S339 | 0.176 | 93.27 | 481 | 0.139905 | 0.176 | 0.1399 | 0.8889 |
| Q9Y5L0 | TNPO3 | MkMKIQTsFYELPTD | S74 | 0.132 | 90.08 | 923 | 0.104929 | 0.132 | 0.1049 | 1 |
| P30530 | AXL | QDDWIVVsQLRITSL | S100 | 0.022 | 92.92 | 894 | 0.017488 | 0.022 | 0.0175 | 1 |
| P30530 | AXL | LNKtssFsCEAHNAK | S204 | 0.129 | 86.3 | 894 | 0.102544 | 0.129 | 0.1025 | 1 |
| P10914 | IRF1 | WAIHTGRyKAGEKEP | Y65 | 0.18 | 94.02 | 325 | 0.078963 | 0.297 | 0.1303 | 1 |
| Q8TAF3 | WDR48 | NDkRHILtkDTNNNV | T377 | 0.039 | 98.77 | 677 | 0.026638 | 0.039 | 0.0266 | 1 |
| P12883 | MYH7 | GGkVtAEtEyGKTVt | T63 | 0.113 | 87.38 | 1935 | 0.077181 | 0.113 | 0.0772 | 1 |
| P12883 | MYH7 | GFPNRILyGDFRQRY | Y715 | 0.111 | 88.93 | 1935 | 0.048694 | 0.111 | 0.0487 | 0.9286 |
| P12883 | MYH7 | LDIDHNQykFGHTKV | Y756 | 0.056 | 87.18 | 1935 | 0.024566 | 0.056 | 0.0246 | 1 |
| Q6ZS86 | GK5 | sKRLFTAsLFTFTTQ | S159 | 0.1 | 89.51 | 529 | 0.079491 | 0.1 | 0.0795 | 1 |

| | | | | | | | | | | |
|---|---|---|---|---|---|---|---|---|---|---|
| Q6ZS86 | GK5 | FTFTTQQtSLRLVWI | T168 | 0 | 98.01 | 529 | 0 | 0 | 0 | 1 |
| Q14574 | DSC3 | LNDGSVYtARAVALS | T81 | 0.004 | 89.65 | 896 | 0.002732 | 0.004 | 0.0027 | 1 |
| P30520 | ADSS2 | GVGksREsMIQLF__ | S451 | 0.115 | 88.99 | 456 | 0.091415 | 0.115 | 0.0914 | 0.8889 |
| Q9Y2G3 | ATP11B | GIKVWVLtGDKHETA | T706 | 0.009 | 89.29 | 1177 | 0.006147 | 0.009 | 0.0061 | 1 |
| Q9P2E3 | ZNFX1 | SRAkKGMyCIGNMQM | Y1229 | 0.089 | 97.95 | 1918 | 0.039043 | 0.089 | 0.039 | 1 |
| Q69YN2 | CWF19L1 | VVEEVEKyKATLRRF | Y388 | 0 | 96.13 | 538 | 0 | 0 | 0 | 1 |
| Q08493 | PDE4C | VIDMVLAtDMsKHMN | T499 | 0.107 | 98.42 | 712 | 0.073082 | 0.107 | 0.0731 | 0.8889 |
| Q8WUA7 | TBC1D22A | LEELRRLsWsGIPKP | S219 | 0 | 98.42 | 517 | 0 | 0 | 0 | 1 |
| O95985 | TOP3B | TLDFLGKyNKWDkVD | Y70 | 0.26 | 90.92 | 862 | 0.114058 | 0.26 | 0.1141 | 1 |
| P0CK96 | SLC35E2B | VtVGVLLyNKARQHQ | Y367 | 0.052 | 86.84 | 405 | 0.022812 | 0.052 | 0.0228 | 1 |
| Q9Y6A2 | CYP46A1 | EDLGRLQyLsQVLKE | Y352 | 0.183 | 98.9 | 500 | 0.080279 | 0.183 | 0.0803 | 1 |
| Q9NYT0 | PLEK2 | GNLFKVItKDDTHyy | T326 | 0.103 | 94.78 | 353 | 0.07035 | 0.103 | 0.0703 | 0.8571 |
| P54274 | TERF1 | LTACQLRtIYICQFL | T122 | 0.01 | 95.34 | 439 | 0.00683 | 0.102 | 0.0697 | 0.8571 |
| Q6NUS8 | UGT3A1 | VTHGGQNsVMEAIRH | S374 | 0.118 | 94.78 | 523 | 0.0938 | 0.118 | 0.0938 | 1 |
| Q9HCR9 | PDE11A | LMQLLKQsILATDLT | S771 | 0.072 | 98.38 | 933 | 0.057234 | 0.072 | 0.0572 | 1 |
| Q9HCR9 | PDE11A | KLKPMLDsVAtNRSK | S897 | 0.176 | 97.1 | 933 | 0.139905 | 0.176 | 0.1399 | 1 |
| Q8NB46 | ANKRD52 | ARDkLWQtPLHVAAA | T109 | 0.075 | 95.2 | 1076 | 0.051226 | 0.075 | 0.0512 | 1 |
| Q04656 | ATP7A | VGDGINDsPALAMAN | S1306 | 0.089 | 78.14 | 1500 | 0.070747 | 0.089 | 0.0707 | 1 |
| O75439 | PMPCB | SRkDLVDyITTHYkG | Y229 | 0.053 | 93.46 | 489 | 0.02325 | 0.053 | 0.0232 | 1 |
| Q9HBK9 | AS3MT | VGEKGHVtGIDMtKG | T99 | 0.106 | 98.82 | 375 | 0.072399 | 0.106 | 0.0724 | 1 |
| Q8N6K7 | SAMD3 | IDKRMsQtLEIRRkM | T312 | 0.016 | 95.06 | 520 | 0.010928 | 0.016 | 0.0109 | 1 |
| Q96RD9 | FCRL5 | FYWCKAAtMPYSVIS | T264 | 0.02 | 82.54 | 977 | 0.01366 | 0.02 | 0.0137 | 1 |
| Q96RD9 | FCRL5 | PYSVISDsPRSWIQV | S273 | 0.058 | 91.11 | 977 | 0.046105 | 0.058 | 0.0461 | 1 |
| Q8WUA4 | GTF3C2 | GLLALACsDGKVLLF | S504 | 0 | 95.16 | 911 | 0 | 0 | 0 | 1 |
| Q9P265 | DIP2B | SRSLKLAytLLNKLG | Y383 | 0.259 | 93.41 | 1576 | 0.11362 | 0.259 | 0.1136 | 1 |
| O15554 | KCNN4 | FRQVRLKhRKLREQV | H358 | 0.245 | 95.42 | 427 | 0.127214 | 0.245 | 0.1272 | 1 |
| P24043 | LAMA2 | QSRISEIsMEVAEQG | S1358 | 0.168 | 94.37 | 3122 | 0.133546 | 0.168 | 0.1335 | 1 |
| Q9UKK3 | PARP4 | CLLDLIAtMLVLQFI | T1629 | 0 | 95.47 | 1724 | 0 | 0.151 | 0.1232 | 0.875 |
| O60942 | RNGTT | WkADGTRyMMLIDGT | Y300 | 0.007 | 96.78 | 597 | 0.003071 | 0.007 | 0.0031 | 1 |
| O60942 | RNGTT | NGQAVPRyLIyDIIK | Y359 | 0 | 97.2 | 597 | 0 | 0 | 0 | 1 |
| O15254 | ACOX3 | TEEFIIHsPDFEAAK | S187 | 0.052 | 98.59 | 700 | 0.041336 | 0.052 | 0.0413 | 1 |

| | | | | | | | | | | |
|---|---|---|---|---|---|---|---|---|---|---|
| Q709F0 | ACAD11 | MILVPMNtPGVKIIR | T576 | 0.186 | 98.75 | 780 | 0.127041 | 0.186 | 0.127 | 0.8333 |
| Q01581 | HMGCS1 | TDREDINsLCMTVVQ | S67 | 0.006 | 98.85 | 520 | 0.004769 | 0.006 | 0.0048 | 0.8889 |
| P52757 | CHN2 | YPDINIItGALKLYF | T349 | 0 | 98.61 | 468 | 0 | 0 | 0 | 1 |
| O15041 | SEMA3E | DGISCSRyYPTGTHA | Y547 | 0.183 | 91.39 | 775 | 0.080279 | 0.183 | 0.0803 | 1 |
| Q6UW01 | CBLN3 | GFDRASGsFVAPVRG | S115 | 0.069 | 95.6 | 205 | 0.054849 | 0.069 | 0.0548 | 0.8333 |
| Q2M385 | MPEG1 | DFSCPsGysPVHLLS | Y416 | 0.195 | 92.93 | 716 | 0.085544 | 0.195 | 0.0855 | 1 |
| Q99426 | TBCB | AyDQRQDtVRsFLKR | T120 | 0.163 | 94.61 | 244 | 0.111331 | 0.163 | 0.1113 | 1 |
| Q99426 | TBCB | QRQDtVRsFLKRsKL | S123 | 0.108 | 93.37 | 244 | 0.085851 | 0.108 | 0.0859 | 1 |
| Q9UBT6 | POLK | NkIIMEAtkGSRFYG | T44 | 0.093 | 90.19 | 870 | 0.06352 | 0.093 | 0.0635 | 1 |
| Q9UBT6 | POLK | STVSSVVsTAEEIFA | S479 | 0.183 | 95.76 | 870 | 0.145469 | 0.183 | 0.1455 | 1 |
| Q5JQF8 | PABPC1L2A | LGyAyVNyQQPVDAK | Y50 | 0.055 | 98.61 | 200 | 0.024128 | 0.055 | 0.0241 | 1 |
| O60551 | NMT2 | KKLVGFIsAIPANIR | S229 | 0.018 | 98.1 | 498 | 0.014308 | 0.018 | 0.0143 | 0.8889 |
| O60551 | NMT2 | EGIFQAVyTAGVVLP | Y283 | 0.236 | 97.5 | 498 | 0.10353 | 0.236 | 0.1035 | 0.8333 |
| P55060 | CSE1L | KALtLPGssENEyIM | S566 | 0 | 95.57 | 971 | 0 | 0 | 0 | 1 |
| P55060 | CSE1L | FLVFINLyCIKYGAL | Y797 | 0.064 | 97.8 | 971 | 0.028076 | 0.064 | 0.0281 | 0.8 |
| Q9BTZ2 | DHRS4 | GAHVVVSsRkQQNVD | S63 | 0.113 | 97.69 | 278 | 0.089825 | 0.113 | 0.0898 | 1 |
| P29374 | ARID4A | GAVWKQIyMDLGIPI | Y369 | 0.118 | 94.78 | 1257 | 0.051765 | 0.118 | 0.0518 | 0.9286 |
| Q9BZ72 | PITPNM2 | DLGYLIIyVTGRPDM | Y1149 | 0 | 92.02 | 1349 | 0 | 0 | 0 | 1 |
| Q9BZ72 | PITPNM2 | STKDVAVySAISLSP | Y1217 | 0.009 | 94.83 | 1349 | 0.003948 | 0.009 | 0.0039 | 1 |
| P35613 | BSG | LGSKILLtCSLNDSA | T156 | 0.145 | 97.41 | 385 | 0.099037 | 0.145 | 0.099 | 1 |
| P35613 | BSG | MEADPGQyRCNGTSS | Y299 | 0 | 97.03 | 385 | 0 | 0 | 0 | 1 |
| Q06124 | PTPN11 | LTRGVDGsFLARPsK | S28 | 0.07 | 96.81 | 593 | 0.055644 | 0.09 | 0.0771 | 1 |
| Q06124 | PTPN11 | DFVLSVRtGDDKGES | T153 | 0.05 | 88.82 | 593 | 0.034151 | 0.086 | 0.0634 | 1 |
| P04035 | HMGCR | RLPRACDsAEVkAWL | S602 | 0 | 97.69 | 888 | 0 | 0 | 0 | 1 |
| A0A0J9YWL9 | TEX13C | NGGsPAFytAFRSRP | Y35 | 0.071 | 87.51 | 993 | 0.031147 | 0.071 | 0.0311 | 1 |
| Q13542 | EIF4EBP2 | tPGGtLFsttPGGtR | S44 | 0.115 | 97.05 | 120 | 0.091415 | 0.115 | 0.0914 | 1 |
| Q13542 | EIF4EBP2 | PGGtRIIyDRKFLLD | Y54 | 0.234 | 96.74 | 120 | 0.102652 | 0.234 | 0.1027 | 1 |
| Q9Y365 | STARD10 | WLPMGADyIIMNySV | Y143 | 0.299 | 98.31 | 291 | 0.131167 | 0.299 | 0.1312 | 1 |
| Q9Y365 | STARD10 | KDLVRAVsIQTGYLI | S166 | 0.126 | 94.61 | 291 | 0.100159 | 0.126 | 0.1002 | 1 |
| O75419 | CDC45 | MkDMRVQtFSIHFGF | T360 | 0.066 | 97.55 | 566 | 0.045079 | 0.066 | 0.0451 | 0.9 |
| Q15048 | LRRC14 | LkELLRDsVAQAELR | S417 | 0.082 | 71.08 | 493 | 0.065183 | 0.186 | 0.1479 | 0.8571 |

| | | | | | | | | | | |
|---|---|---|---|---|---|---|---|---|---|---|
| P48730 | CSNK1D | HPQLHIEskIYkMMQ | S53 | 0.066 | 95.03 | 415 | 0.052464 | 0.066 | 0.0525 | 1 |
| P48730 | CSNK1D | tGtARYAsINTHLGI | S181 | 0.003 | 98.63 | 415 | 0.002385 | 0.003 | 0.0024 | 1 |
| P48730 | CSNK1D | THLGIEQsRRDDLES | S191 | 0.015 | 98.78 | 415 | 0.011924 | 0.015 | 0.0119 | 1 |
| P48730 | CSNK1D | RFDDkPDySYLRQLF | Y266 | 0.13 | 98.77 | 415 | 0.057029 | 0.13 | 0.057 | 1 |
| Q14507 | EDDM3A | kSFHMFIysLWFkIQ | Y70 | 0.332 | 98.39 | 147 | 0.145644 | 0.332 | 0.1456 | 1 |
| Q14507 | EDDM3A | SFHMFIysLWFkIQR | S71 | 0.105 | 97.45 | 147 | 0.083466 | 0.105 | 0.0835 | 0.8333 |
| O60637 | TSPAN3 | FKETKNQsVPLsCCR | S169 | 0.18 | 93.98 | 253 | 0.143085 | 0.18 | 0.1431 | 1 |
| O60637 | TSPAN3 | KNQsVPLsCCRETAS | S173 | 0.009 | 94.64 | 253 | 0.007154 | 0.009 | 0.0072 | 1 |
| Q9BYX4 | IFIH1 | EALRRTGsPLAARYM | S88 | 0.048 | 72.25 | 1025 | 0.038156 | 0.048 | 0.0382 | 0.875 |
| Q92502 | STARD8 | ALMLTSQyLMEPCGL | Y964 | 0.107 | 96.63 | 1023 | 0.046939 | 0.107 | 0.0469 | 1 |
| Q14315 | FLNC | LNPSGAktDtyVtDN | T1300 | 0.127 | 88.77 | 2725 | 0.086743 | 0.127 | 0.0867 | 0.9167 |
| Q14315 | FLNC | tDNGDGTyRVQYTAY | Y1312 | 0.035 | 93.72 | 2725 | 0.015354 | 0.035 | 0.0154 | 1 |
| P20020 | ATP2B1 | ICsDKtGtLTMNRMt | T479 | 0.016 | 76.26 | 1220 | 0.010928 | 0.016 | 0.0109 | 1 |
| Q92506 | HSD17B8 | GNVGQTNyAASkAGV | Y169 | 0.268 | 98.07 | 261 | 0.117568 | 0.268 | 0.1176 | 0.9333 |
| Q9H2G2 | SLK | IQVVCKQtLDALNYL | T139 | 0.007 | 97.36 | 1235 | 0.004781 | 0.126 | 0.0861 | 1 |
| P30043 | BLVRB | RGPsRVIskHDLGHF | S177 | 0 | 98.67 | 206 | 0 | 0 | 0 | 1 |
| P30043 | BLVRB | DGHSTYPsHQyQ___ | S202 | 0.01 | 98.7 | 206 | 0.007949 | 0.01 | 0.0079 | 1 |
| Q9H269 | VPS16 | NKYEAKkyAsRVGPE | Y775 | 0.025 | 91.26 | 839 | 0.010967 | 0.025 | 0.011 | 1 |
| O00499 | BIN1 | RkLVDyDsARHHyES | S152 | 0.157 | 90.95 | 593 | 0.124802 | 0.157 | 0.1248 | 1 |
| O43868 | SLC28A2 | APCALASsKLAYPEV | S377 | 0 | 98.33 | 658 | 0 | 0 | 0 | 1 |
| P51168 | SCNN1B | IyFQEFNyRTIEESA | Y504 | 0.3 | 90.93 | 640 | 0.131606 | 0.3 | 0.1316 | 1 |
| Q15418 | RPS6KA1 | RPDSGHLyAMkVLKK | Y91 | 0.059 | 94 | 735 | 0.025882 | 0.059 | 0.0259 | 1 |
| Q8TBY9 | CFAP251 | GFTEGTVyILDAMSL | Y674 | 0.284 | 96.85 | 1149 | 0.124587 | 0.284 | 0.1246 | 1 |
| Q8TBY9 | CFAP251 | RsFTVAVyMLVVRNG | Y720 | 0.046 | 96.61 | 1149 | 0.02018 | 0.046 | 0.0202 | 1 |
| Q4L235 | AASDH | WRGEGNKtQIFNVYG | T338 | 0.021 | 97.87 | 1098 | 0.014343 | 0.021 | 0.0143 | 1 |
| P30050 | RPL12 | LARELsGtIkEILGt | T128 | 0.02 | 80.29 | 165 | 0.01366 | 0.02 | 0.0137 | 1 |
| Q8TBZ6 | TRMT10A | HPVQFYLtSHGGQLK | T144 | 0.01 | 98.74 | 339 | 0.00683 | 0.01 | 0.0068 | 1 |
| Q9NR45 | NANS | GDTNNFPyLEkTAKK | Y142 | 0.118 | 98.33 | 359 | 0.051765 | 0.118 | 0.0518 | 1 |
| Q9H251 | CDH23 | GDGGLVNyRILSGAE | Y1246 | 0.152 | 92.22 | 3354 | 0.06668 | 0.152 | 0.0667 | 1 |
| Q15311 | RALBP1 | DLPENLLtkELMPRF | T280 | 0.154 | 87.31 | 655 | 0.105184 | 0.154 | 0.1052 | 0.875 |
| Q8IVF7 | FMNL3 | EVKLLRQyERERQPL | Y697 | 0.256 | 86.66 | 1028 | 0.112304 | 0.256 | 0.1123 | 1 |

| | | | | | | | | | | |
|---|---|---|---|---|---|---|---|---|---|---|
| Q86XW9 | NME9 | HHMCSGPsHLLILTR | S227 | 0 | 92.67 | 330 | 0 | 0 | 0 | 1 |
| P53618 | COPB1 | KLPGLLMtIIRFVLP | T61 | 0.068 | 90.47 | 953 | 0.046445 | 0.1 | 0.0771 | 1 |
| P53618 | COPB1 | VKYEAAGtLVtLSSA | T267 | 0.067 | 90.11 | 953 | 0.045762 | 0.067 | 0.0458 | 1 |
| O00418 | EEF2K | FDSGQNLsPDRCQDW | S627 | 0.019 | 85.15 | 725 | 0.015103 | 0.019 | 0.0151 | 1 |
| Q04637 | EIF4G1 | SKAIIEEyLHLNDMK | Y1253 | 0.07 | 95.49 | 1599 | 0.030708 | 0.07 | 0.0307 | 0.9167 |
| Q04637 | EIF4G1 | AEQkVEytLGEEsEA | T1425 | 0.059 | 87.21 | 1599 | 0.040298 | 0.059 | 0.0403 | 1 |
| Q06609 | RAD51 | HASTTRLyLRKGRGE | Y301 | 0.331 | 97.81 | 339 | 0.145205 | 0.331 | 0.1452 | 1 |
| Q06609 | RAD51 | LRKGRGEtRICKIyD | T309 | 0.203 | 97.31 | 339 | 0.138652 | 0.203 | 0.1387 | 1 |
| Q14103 | HNRNPD | kMFIGGLsWDTTkkD | S105 | 0.072 | 88.85 | 355 | 0.057234 | 0.072 | 0.0572 | 1 |
| Q14103 | HNRNPD | LDPItGRsrGFGFVL | S137 | 0.083 | 88.26 | 355 | 0.065978 | 0.083 | 0.066 | 1 |
| Q9UHC9 | NPC1L1 | LVSAVGMsVEFVSHI | S1205 | 0.011 | 92.15 | 1359 | 0.008744 | 0.011 | 0.0087 | 1 |
| Q9UHC9 | NPC1L1 | LERAKEAtISMGSAV | T1232 | 0.035 | 94.28 | 1359 | 0.023905 | 0.035 | 0.0239 | 1 |
| Q96ST3 | SIN3A | MDVDVEDyyPAFLDM | Y958 | 0.208 | 93.82 | 1273 | 0.091247 | 0.208 | 0.0912 | 0.9091 |
| Q86V81 | ALYREF | HYDrsGRsLGtADVH | S145 | 0.188 | 91.71 | 257 | 0.149444 | 0.188 | 0.1494 | 1 |
| Q7Z745 | MROH2B | sKIAKIVsKFIPNEE | S997 | 0.004 | 90.23 | 1585 | 0.00318 | 0.004 | 0.0032 | 0.8571 |
| Q9BWF3 | RBM4 | kLHVGNIsPtCtNkE | S86 | 0.101 | 89.57 | 364 | 0.080286 | 0.101 | 0.0803 | 1 |
| Q9H4Z3 | PCIF1 | SRSAsPDsRKVVKWN | S308 | 0.163 | 95.23 | 704 | 0.129571 | 0.163 | 0.1296 | 1 |
| Q9H4Z3 | PCIF1 | NNVVCIRyKGEMVKV | Y432 | 0.145 | 98.33 | 704 | 0.063609 | 0.145 | 0.0636 | 1 |
| Q9H4Z3 | PCIF1 | PEWREPPtPALTrME | T593 | 0.096 | 87.39 | 704 | 0.065569 | 0.096 | 0.0656 | 1 |
| Q7L8L6 | FASTKD5 | QAQVIVDyLCKLsSL | Y166 | 0.303 | 96.71 | 764 | 0.132922 | 0.303 | 0.1329 | 1 |
| Q7L8L6 | FASTKD5 | VDyLCKLsSLPAEQH | S171 | 0.103 | 94.18 | 764 | 0.081876 | 0.103 | 0.0819 | 0.8889 |
| Q7L8L6 | FASTKD5 | PPNAEEFySSLISEI | Y451 | 0.02 | 98.5 | 764 | 0.008774 | 0.02 | 0.0088 | 0.9286 |
| Q9H8Y8 | GORASP2 | VLRVQENsPGHRAGL | S26 | 0 | 94.76 | 452 | 0 | 0 | 0 | 1 |
| Q9H8Y8 | GORASP2 | kPVKMLIySSkTLEL | Y72 | 0.197 | 93.7 | 452 | 0.086421 | 0.197 | 0.0864 | 1 |
| P56377 | AP1S2 | MQFMLLFsRQGKLRL | S8 | 0.043 | 97.56 | 157 | 0.034181 | 0.043 | 0.0342 | 1 |
| Q13207 | TBX2 | RtyVFPEtDFIAVTA | T258 | 0 | 98.45 | 712 | 0 | 0 | 0 | 1 |
| P31689 | DNAJA1 | sVtLEDLyNGATRkL | Y119 | 0.067 | 94.99 | 397 | 0.029392 | 0.067 | 0.0294 | 0.9231 |
| P31689 | DNAJA1 | NRTIVITsHPGQIVk | S289 | 0.094 | 93.16 | 397 | 0.074722 | 0.184 | 0.1477 | 1 |
| Q9UF33 | EPHA6 | CSKCPPHsLTYMEAT | S300 | 0.102 | 95.99 | 1036 | 0.081081 | 0.102 | 0.0811 | 1 |
| O43173 | ST8SIA3 | RSLLPDVsPIMNKHY | S152 | 0.085 | 97.84 | 380 | 0.067568 | 0.085 | 0.0676 | 1 |
| Q16671 | AMHR2 | IKAFPPRsVAQFQAE | S236 | 0.064 | 89.6 | 573 | 0.050875 | 0.064 | 0.0509 | 1 |

| | | | | | | | | | | |
|---|---|---|---|---|---|---|---|---|---|---|
| Q9BV57 | ADI1 | YILDGsGyFDVRDkE | Y104 | 0.132 | 98.64 | 179 | 0.057907 | 0.132 | 0.0579 | 0.9286 |
| Q96CS3 | FAF2 | HPVFYQGtYSQALND | T158 | 0.19 | 91.53 | 445 | 0.129773 | 0.19 | 0.1298 | 1 |
| P51648 | ALDH3A2 | EAAAkHLtPVTLELG | T202 | 0.149 | 98.91 | 485 | 0.101769 | 0.149 | 0.1018 | 1 |
| P51648 | ALDH3A2 | YIAPTVLtDVDPkTk | T319 | 0.129 | 98.75 | 485 | 0.088109 | 0.129 | 0.0881 | 1 |
| Q8N7B6 | PACRGL | EQHGGsGsLsIIKSK | S232 | 0.001 | 92.05 | 248 | 0.000795 | 0.001 | 0.0008 | 1 |
| Q10570 | CPSF1 | sPHWLLVtGRGALRL | T956 | 0.023 | 83.65 | 1443 | 0.015709 | 0.023 | 0.0157 | 1 |
| Q10570 | CPSF1 | GQKIFLWsLRASELT | S1194 | 0.101 | 96.02 | 1443 | 0.080286 | 0.179 | 0.1449 | 1 |
| P27695 | APEX1 | DPPDQkTsPsGkPAT | S54 | 0.083 | 98.11 | 318 | 0.065978 | 0.083 | 0.066 | 1 |
| P27695 | APEX1 | NKkNAGFtPQERQGF | T233 | 0.025 | 98.5 | 318 | 0.017075 | 0.025 | 0.0171 | 1 |
| P49327 | FASN | FLRLGMLsPEGtCkA | S207 | 0.002 | 95.14 | 2511 | 0.00159 | 0.002 | 0.0016 | 1 |
| P49327 | FASN | FDTAGNGyCRsEGVV | Y222 | 0.132 | 96.51 | 2511 | 0.057907 | 0.132 | 0.0579 | 1 |
| P49327 | FASN | AGNGyCRsEGVVAVL | S225 | 0 | 97.08 | 2511 | 0 | 0 | 0 | 1 |
| P49327 | FASN | EQLIRSLyQsAGVAP | Y277 | 0.119 | 92.35 | 2511 | 0.052204 | 0.201 | 0.075 | 1 |
| P49327 | FASN | VAPEsFEyIEAHGTG | Y289 | 0.011 | 97.42 | 2511 | 0.004826 | 0.011 | 0.0048 | 1 |
| P49327 | FASN | PRGtPLIsPLIKWDH | S831 | 0.019 | 96.84 | 2511 | 0.015103 | 0.019 | 0.0151 | 1 |
| Q93050 | ATP6V0A1 | QTNQtPPtyNkTNkF | T363 | 0.147 | 91.7 | 837 | 0.100403 | 0.147 | 0.1004 | 0.8 |
| Q93050 | ATP6V0A1 | VEFQNKFysGtGFkF | Y815 | 0.258 | 87.51 | 837 | 0.113181 | 0.258 | 0.1132 | 0.9375 |
| P53708 | ITGA8 | IPDARTAsVLVGAPK | S72 | 0.004 | 92.68 | 1063 | 0.00318 | 0.004 | 0.0032 | 1 |
| P12111 | COL6A3 | IFVKSAGsRIEDGVL | S934 | 0.063 | 93.72 | 3177 | 0.05008 | 0.063 | 0.0501 | 1 |
| Q7Z4R8 | C6orf120 | GHPSHLEsEFEMKVY | S127 | 0.003 | 97.34 | 191 | 0.002385 | 0.003 | 0.0024 | 1 |
| O43424 | GRID2 | DIGISALtITPDREN | T525 | 0.051 | 93.17 | 1007 | 0.034834 | 0.18 | 0.1229 | 0.8 |
| Q9H336 | CRISPLD1 | TRVYSDLssICRAAV | S442 | 0 | 93.69 | 500 | 0 | 0 | 0 | 1 |
| Q9H336 | CRISPLD1 | RVYSDLssICRAAVH | S443 | 0.005 | 94.69 | 500 | 0.003975 | 0.005 | 0.004 | 1 |
| Q3SXM5 | HSDL1 | RADLIkQyGRWAVVS | Y66 | 0.175 | 97.37 | 330 | 0.07677 | 0.175 | 0.0768 | 1 |
| Q6MZM0 | HEPHL1 | ANKADKVySILPHGV | Y481 | 0 | 97.99 | 1159 | 0 | 0 | 0 | 0.8 |
| Q96I51 | RCC1L | GMGLNKDsQLGFHRS | S140 | 0.025 | 97.25 | 464 | 0.019873 | 0.025 | 0.0199 | 0.9091 |
| Q96I51 | RCC1L | VSCGRAHsLVLtDRE | S186 | 0 | 98.87 | 464 | 0 | 0 | 0 | 0.8182 |
| Q96I51 | RCC1L | RAHsLVLtDREGVFs | T190 | 0.03 | 98.71 | 464 | 0.02049 | 0.03 | 0.0205 | 1 |
| Q96I51 | RCC1L | tDREGVFsMGNNSYG | S197 | 0.023 | 98.79 | 464 | 0.018283 | 0.023 | 0.0183 | 1 |
| Q96I51 | RCC1L | kVVENEIysESHRVH | Y216 | 0.238 | 96.18 | 464 | 0.104407 | 0.238 | 0.1044 | 1 |
| O15374 | SLC16A4 | tFLLAPFtkFLIDLY | T165 | 0.002 | 93.49 | 487 | 0.001366 | 0.002 | 0.0014 | 1 |

| | | | | | | | | | | |
|---|---|---|---|---|---|---|---|---|---|---|
| O15374 | SLC16A4 | ETVSQIIsGWVADQN | S354 | 0.065 | 98.23 | 487 | 0.051669 | 0.065 | 0.0517 | 1 |
| P26998 | CRYBB3 | GSYKVILyELENFQG | Y29 | 0.191 | 98.01 | 211 | 0.083789 | 0.191 | 0.0838 | 0.9091 |
| Q9NTI5 | PDS5B | LIKQVNksIDGTADD | S628 | 0.053 | 93.32 | 1447 | 0.04213 | 0.132 | 0.1199 | 1 |
| Q9NTI5 | PDS5B | LLsLLPEyVVPYTIH | Y990 | 0.09 | 96.84 | 1447 | 0.039482 | 0.09 | 0.0395 | 1 |
| Q9NTI5 | PDS5B | LLAHDPDyVKVQDIE | Y1005 | 0.05 | 96.96 | 1447 | 0.021934 | 0.05 | 0.0219 | 1 |
| Q9NTI5 | PDS5B | MVENIKQtKDAQGPD | T1050 | 0.005 | 94.76 | 1447 | 0.003415 | 0.005 | 0.0034 | 1 |
| Q9NRD8 | DUOX2 | TTRLREIyssPKGNG | Y1347 | 0.04 | 83.18 | 1548 | 0.017547 | 0.134 | 0.0588 | 1 |
| P11487 | FGF3 | TRRTQKSsLFLPRVL | S176 | 0 | 98.28 | 239 | 0 | 0 | 0 | 1 |
| Q5U5R9 | HECTD2 | GLDMGGLtKEWFLLL | T461 | 0.075 | 90.93 | 776 | 0.051226 | 0.075 | 0.0512 | 0.8889 |
| Q9BZ19 | ANKRD60 | HRLNHTLsERQMVLL | S301 | 0.028 | 92.42 | 345 | 0.022258 | 0.028 | 0.0223 | 1 |
| P07315 | CRYGC | YLLRRGEyPDyQQWM | Y63 | 0.33 | 98.37 | 174 | 0.144766 | 0.33 | 0.1448 | 1 |
| Q16549 | PCSK7 | HyQINDIySCSWGPD | Y284 | 0 | 98.74 | 785 | 0 | 0 | 0 | 0.8 |
| P29372 | MPG | ARKPLRFyVRGSPWV | Y278 | 0.011 | 98.47 | 298 | 0.004826 | 0.175 | 0.0768 | 1 |
| O00507 | USP9Y | RELDMGPytVAGVAN | Y1829 | 0.205 | 92.07 | 2555 | 0.089931 | 0.205 | 0.0899 | 1 |
| Q6UWM7 | LCTL | HVNHYRFsLsWPRLL | S111 | 0.021 | 98.89 | 567 | 0.016693 | 0.021 | 0.0167 | 1 |
| Q6UWM7 | LCTL | NHYRFsLsWPRLLPt | S113 | 0.063 | 98.85 | 567 | 0.05008 | 0.063 | 0.0501 | 1 |
| P41180 | CASR | QFKSFLRtIPNDEHQ | T186 | 0.035 | 94.27 | 1078 | 0.023905 | 0.035 | 0.0239 | 1 |
| P41180 | CASR | PNDEHQAtAMADIIE | T195 | 0.185 | 93.57 | 1078 | 0.126358 | 0.185 | 0.1264 | 1 |
| Q92890 | UFD1 | STQYRCFsVsMLAGP | S27 | 0.085 | 89.98 | 307 | 0.067568 | 0.085 | 0.0676 | 1 |
| Q92890 | UFD1 | NkNSDRMtHCGVLEF | T78 | 0.004 | 94.2 | 307 | 0.002732 | 0.004 | 0.0027 | 0.8571 |
| Q9UKV8 | AGO2 | HPTEFDFyLCsHAGI | Y749 | 0.053 | 98.83 | 859 | 0.02325 | 0.053 | 0.0232 | 0.8667 |
| Q9UKV8 | AGO2 | HAGIQGtsRPSHYHV | S760 | 0.086 | 96.74 | 859 | 0.068363 | 0.086 | 0.0684 | 1 |
| P55259 | GP2 | GEGGVRMsEtCVQVH | S104 | 0.036 | 94.07 | 537 | 0.028617 | 0.036 | 0.0286 | 1 |
| Q96PE1 | ADGRA2 | GDFRWPRtLAGITAy | T370 | 0.116 | 87.6 | 1338 | 0.07923 | 0.116 | 0.0792 | 1 |
| Q9P0K1 | ADAM22 | HRLSVVHtNtYAKSV | T262 | 0.005 | 90.95 | 906 | 0.003415 | 0.005 | 0.0034 | 1 |
| Q9P0K1 | ADAM22 | IsENPLItLREFMKY | T308 | 0.004 | 96.24 | 906 | 0.002732 | 0.004 | 0.0027 | 1 |
| Q9P2N4 | ADAMTS9 | FSINLyGtGLSLTES | T1881 | 0.182 | 86.54 | 1935 | 0.124308 | 0.182 | 0.1243 | 0.8571 |
| P68104 | EEF1A1 | GHVDSGksttGHLI | S21 | 0.131 | 89.83 | 462 | 0.104134 | 0.131 | 0.1041 | 1 |
| P68104 | EEF1A1 | DSGksttGHLIykC | T24 | 0.028 | 91.11 | 462 | 0.019124 | 0.028 | 0.0191 | 1 |
| P68104 | EEF1A1 | RGITIDIsLWkFEtS | S76 | 0.134 | 75.64 | 462 | 0.106519 | 0.134 | 0.1065 | 1 |
| P68104 | EEF1A1 | WkFEtSkyyVtIIDA | Y85 | 0.154 | 88.29 | 462 | 0.067558 | 0.25 | 0.0955 | 0.9091 |

| | | | | | | | | | | |
|---|---|---|---|---|---|---|---|---|---|---|
| P68104 | EEF1A1 | EHALLAytLGVkQLI | T142 | 0.154 | 82.36 | 462 | 0.105184 | 0.194 | 0.1325 | 0.875 |
| P68104 | EEF1A1 | GVNkMDstEPPysQk | T158 | 0.042 | 95.18 | 462 | 0.028687 | 0.042 | 0.0287 | 1 |
| P68104 | EEF1A1 | ykIGGIGtVPVGrVE | T261 | 0.114 | 94.09 | 462 | 0.077864 | 0.09 | 0.0952 | 1 |
| P68104 | EEF1A1 | TFAPVNVtTEVksVE | T286 | 0.134 | 95.29 | 462 | 0.091524 | 0.218 | 0.1489 | 1 |
| P68104 | EEF1A1 | GkPMCVEsFsDyPPL | S414 | 0.089 | 94.12 | 462 | 0.070747 | 0.089 | 0.0707 | 1 |
| P68104 | EEF1A1 | AAGAGkVtksAQkAQ | T452 | 0.05 | 85.75 | 462 | 0.034151 | 0.05 | 0.0342 | 1 |
| P68104 | EEF1A1 | GAGkVtksAQkAQkA | S454 | 0.093 | 82.25 | 462 | 0.073927 | 0.183 | 0.1458 | 1 |
| Q9H9P8 | L2HGDH | SDRISELsGCTPDPR | S270 | 0.088 | 90.33 | 463 | 0.069952 | 0.088 | 0.07 | 1 |
| Q9BW91 | NUDT9 | NKARTsPyPGskVER | Y70 | 0.006 | 97.75 | 350 | 0.002632 | 0.006 | 0.0026 | 1 |
| Q9BW91 | NUDT9 | RKSKNGLyEIENGRP | Y140 | 0.122 | 98.71 | 350 | 0.05352 | 0.122 | 0.0535 | 1 |
| P58107 | EPPK1 | PFSGkQVsVFQAMKk | S1021 | 0.041 | 93.38 | 5088 | 0.032592 | 0.041 | 0.0326 | 1 |
| P51649 | ALDH5A1 | AALLRtDsFVGGRWL | S63 | 0.037 | 98.82 | 535 | 0.029412 | 0.037 | 0.0294 | 1 |
| P51649 | ALDH5A1 | SSLLRkWyNLMIQNk | Y128 | 0.307 | 98.89 | 535 | 0.134676 | 0.307 | 0.1347 | 1 |
| P51649 | ALDH5A1 | SKYGIDEyLELKYVC | Y524 | 0.052 | 98.93 | 535 | 0.022812 | 0.052 | 0.0228 | 1 |
| O75144 | ICOSLG | DVYVYWQtSESKTVV | T56 | 0.014 | 98.06 | 302 | 0.009562 | 0.014 | 0.0096 | 1 |
| Q9H346 | OR52D1 | GHRVMtHtyCEHMGI | T180 | 0.013 | 90.52 | 318 | 0.008879 | 0.213 | 0.1455 | 1 |
| Q14103 | HNRNPD | VGGLsPDtPEEkIRE | T193 | 0.156 | 77.57 | 355 | 0.10655 | 0.156 | 0.1066 | 0.8889 |
| P53597 | SUCLG1 | QQALEYGtkLVGGTt | T80 | 0.079 | 98.03 | 346 | 0.053958 | 0.079 | 0.054 | 1 |
| P05108 | CYP11A1 | EVMAPEAtKNFLPLL | T164 | 0.135 | 98.04 | 521 | 0.092207 | 0.135 | 0.0922 | 0.8889 |
| P31930 | UQCRC1 | EHLAFkGtkNRPGsA | T101 | 0.019 | 93.9 | 480 | 0.012977 | 0.019 | 0.013 | 1 |
| P31930 | UQCRC1 | StREHTAyyIkALSk | Y131 | 0.052 | 97.68 | 480 | 0.022812 | 0.052 | 0.0228 | 1 |
| P31930 | UQCRC1 | sRADLTEyLstHykA | Y219 | 0.039 | 93.94 | 480 | 0.017109 | 0.039 | 0.0171 | 1 |
| P31930 | UQCRC1 | VCEDIGRsLLTYGRR | S416 | 0.039 | 96.76 | 480 | 0.031002 | 0.039 | 0.031 | 0.8889 |
| P31930 | UQCRC1 | QCPAVAGyGPIEQLP | Y459 | 0.26 | 98.2 | 480 | 0.114058 | 0.26 | 0.1141 | 1 |
| O95999 | BCL10 | GLDTLVEsIRREkTQ | S85 | 0.067 | 92.8 | 233 | 0.053259 | 0.067 | 0.0533 | 1 |
| Q9NUV7 | SPTLC3 | GDLLQLKySRHKKSA | Y531 | 0.289 | 93.28 | 552 | 0.12678 | 0.289 | 0.1268 | 1 |
| Q58F21 | BRDT | EKMDNQEyKDAykFA | Y331 | 0.155 | 96.38 | 947 | 0.067996 | 0.155 | 0.068 | 1 |
| O43602 | DCX | PQGVRyIyTIDGSRK | Y105 | 0.189 | 93.2 | 365 | 0.082912 | 0.189 | 0.0829 | 1 |
| P16662 | UGT2B7 | MKKWDQFySEVLGRP | Y236 | 0 | 98.26 | 529 | 0 | 0 | 0 | 1 |
| P16662 | UGT2B7 | RVINDPSyKENVMKL | Y438 | 0.184 | 98.7 | 529 | 0.080718 | 0.184 | 0.0807 | 1 |
| Q9ULD0 | OGDHL | RVTNRNItLsLVANP | T354 | 0.165 | 96.23 | 1010 | 0.112697 | 0.165 | 0.1127 | 1 |

| | | | | | | | | | | |
|---|---|---|---|---|---|---|---|---|---|---|
| Q9ULD0 | OGDHL | TNRNItLsLVANPsH | S356 | 0.049 | 96.2 | 1010 | 0.038951 | 0.049 | 0.039 | 1 |
| Q9ULD0 | OGDHL | LRGRADMtKNRtVDW | T633 | 0.159 | 97.67 | 1010 | 0.108599 | 0.159 | 0.1086 | 1 |
| Q9UL01 | DSE | KFVMRNGsGNWLADQ | S338 | 0.076 | 98.76 | 958 | 0.060414 | 0.076 | 0.0604 | 0.8889 |
| Q9UL01 | DSE | WMDDtGysEKATFAS | S625 | 0.073 | 96.41 | 958 | 0.058029 | 0.073 | 0.058 | 1 |
| Q9UL01 | DSE | yPyNGTNyVNVtMHL | Y646 | 0.322 | 98.78 | 958 | 0.141257 | 0.322 | 0.1413 | 1 |
| O60522 | TDRD6 | VAKDPDGtLIIELyG | T1136 | 0.076 | 86.19 | 2096 | 0.051909 | 0.076 | 0.0519 | 1 |
| O60347 | TBC1D12 | GLGILRLyEDILLQM | Y707 | 0.268 | 97.63 | 775 | 0.117568 | 0.268 | 0.1176 | 1 |
| P11717 | IGF2R | SDKSkVVssCQEKRE | S1280 | 0 | 89.76 | 2491 | 0 | 0 | 0 | 1 |
| P11717 | IGF2R | DKSkVVssCQEKREP | S1281 | 0.008 | 92 | 2491 | 0.006359 | 0.008 | 0.0064 | 1 |
| O15439 | ABCC4 | SVERVIEyTDLEkEA | Y1016 | 0.19 | 90.7 | 1325 | 0.08335 | 0.341 | 0.1401 | 1 |
| Q7L1W4 | LRRC8D | YKLYVVQtVIKTAKF | T312 | 0.094 | 92.35 | 858 | 0.064203 | 0.094 | 0.0642 | 1 |
| P53396 | ACLY | sLRSAYDstMETMNy | S581 | 0.066 | 98.1 | 1101 | 0.052464 | 0.092 | 0.0731 | 1 |
| P53396 | ACLY | LRSAYDstMETMNyA | T582 | 0 | 98.75 | 1101 | 0 | 0 | 0 | 1 |
| P53396 | ACLY | MNyAQIRtIAIIAEG | T593 | 0.041 | 98.77 | 1101 | 0.028004 | 0.041 | 0.028 | 1 |
| P53396 | ACLY | GCFkIGNtGGMLDNI | T639 | 0.034 | 98.54 | 1101 | 0.023222 | 0.034 | 0.0232 | 1 |
| P53396 | ACLY | MLDNILAskLyRPGs | S649 | 0.038 | 97.45 | 1101 | 0.030207 | 0.077 | 0.0612 | 1 |
| P53396 | ACLY | yRPGsVAyVSRsGGM | Y659 | 0.013 | 98.83 | 1101 | 0.005703 | 0.013 | 0.0057 | 1 |
| P53396 | ACLY | sVAyVSRsGGMsNEL | S663 | 0.092 | 92.67 | 1101 | 0.073132 | -0.012 | 0.1069 | 1 |
| P53396 | ACLY | VSRsGGMsNELNNII | S667 | 0.03 | 97.79 | 1101 | 0.023847 | 0.03 | 0.0238 | 1 |
| P53396 | ACLY | FMDHVLRyQDtPGVK | Y704 | 0.011 | 98.56 | 1101 | 0.004826 | 0.011 | 0.0048 | 1 |
| P53396 | ACLY | PAtPLLDyALEVEkI | Y1006 | 0.21 | 98.14 | 1101 | 0.092124 | 0.21 | 0.0921 | 1 |
| P53396 | ACLY | SMGFIGHyLDQkRLk | Y1073 | 0.31 | 98.18 | 1101 | 0.135993 | 0.385 | 0.15 | 1 |
| P09769 | FGR | ETtkGAysLsIRDWD | S181 | 0 | 97.33 | 529 | 0 | 0.119 | 0.0946 | 1 |
| P09769 | FGR | KLDMGGyyIttRVQF | Y209 | 0.291 | 94.14 | 529 | 0.127658 | 0.291 | 0.1277 | 1 |
| P22732 | SLC2A5 | WPILLGLtGVPAALQ | T199 | 0.018 | 93.23 | 501 | 0.012294 | 0.018 | 0.0123 | 1 |
| P17844 | DDX5 | VQQVAAEyCRACRLk | Y190 | 0.027 | 93.7 | 614 | 0.011845 | 0.027 | 0.0118 | 1 |
| P17844 | DDX5 | RLkstCIyGGAPkGP | Y202 | 0.337 | 95.02 | 614 | 0.147837 | 0.337 | 0.1478 | 0.8333 |
| P17844 | DDX5 | GVEICIAtPGRLIDF | T224 | 0.134 | 94.54 | 614 | 0.091524 | 0.134 | 0.0915 | 1 |
| P17844 | DDX5 | kTNLRRTtyLVLDEA | T243 | 0.052 | 96.26 | 614 | 0.035517 | 0.052 | 0.0355 | 1 |
| P17844 | DDX5 | TNLRRTtyLVLDEAD | Y244 | 0.008 | 97.56 | 614 | 0.003509 | 0.008 | 0.0035 | 1 |
| P17844 | DDX5 | DrQTLMWsATWPkEV | S279 | 0.017 | 90.09 | 614 | 0.013514 | 0.017 | 0.0135 | 0.8889 |

| | | | | | | | | | | |
|---|---|---|---|---|---|---|---|---|---|---|
| P17844 | DDX5 | MsEkENktIVFVETk | T344 | 0 | 94.78 | 614 | 0 | 0 | 0 | 1 |
| P17844 | DDX5 | DVkFVINyDYPNsSE | Y416 | 0.122 | 96.95 | 614 | 0.05352 | 0.122 | 0.0535 | 1 |
| P17844 | DDX5 | YPNsSEDyIHRIGRT | Y425 | 0.015 | 90.2 | 614 | 0.00658 | 0.015 | 0.0066 | 1 |
| P17844 | DDX5 | ARSTktGtAyTFFtP | T440 | 0.18 | 95.66 | 614 | 0.122942 | 0.18 | 0.1229 | 1 |
| P17844 | DDX5 | GtAyTFFtPNNIkQV | T446 | 0.034 | 92.74 | 614 | 0.023222 | 0.034 | 0.0232 | 1 |
| Q0VDG4 | SCRN3 | KGNITFEtMMEILRD | T258 | 0.148 | 98.04 | 424 | 0.101086 | 0.148 | 0.1011 | 1 |
| Q6TCH7 | PAQR3 | NPyITDGyRAYLPSR | Y50 | 0.171 | 95.77 | 311 | 0.075015 | 0.171 | 0.075 | 0.9286 |
| Q8IXJ9 | ASXL1 | TDGLLRLsSSALNNE | S304 | 0.14 | 82.94 | 1541 | 0.111288 | 0.14 | 0.1113 | 1 |
| P03951 | F11 | RIVGGTAsVRGEWPW | S394 | 0.067 | 84.71 | 625 | 0.053259 | 0.067 | 0.0533 | 1 |
| Q9NSA2 | KCND1 | RFETWKNtLDRYPDT | T57 | 0.102 | 93.14 | 647 | 0.069667 | 0.199 | 0.1354 | 0.8333 |
| O15371 | EIF3D | DEGNSFNsPRNLAME | S308 | 0.072 | 92.61 | 548 | 0.057234 | 0.072 | 0.0572 | 1 |
| O15371 | EIF3D | RNLAMEAtyINHNFS | T317 | 0.161 | 97.75 | 548 | 0.109965 | 0.161 | 0.11 | 0.8571 |
| O15371 | EIF3D | VYSLPDGtFssDEDE | T526 | 0.049 | 81.86 | 548 | 0.033468 | 0.049 | 0.0335 | 1 |
| P13489 | RNH1 | sCkIQkLsLQNCCLt | S91 | 0.032 | 98.51 | 461 | 0.025437 | 0.032 | 0.0254 | 1 |
| P13489 | RNH1 | DFkELTVsNNDINEA | S178 | 0.02 | 98.8 | 461 | 0.015898 | 0.02 | 0.0159 | 1 |
| Q14896 | MYBPC3 | IGRVHKLtIDDVtPA | T602 | 0.111 | 95.64 | 1274 | 0.075815 | 0.111 | 0.0758 | 1 |
| Q14896 | MYBPC3 | FSKQGVLtLEIRKPC | T1237 | 0.187 | 91.56 | 1274 | 0.127724 | 0.187 | 0.1277 | 1 |
| Q9UPI3 | FLVCR2 | GMLGAVIsGIWLDRS | S364 | 0.109 | 86.89 | 526 | 0.086646 | 0.109 | 0.0866 | 1 |
| Q9UQP3 | TNN | GLHPGTEyKITVVPM | Y419 | 0.15 | 89.42 | 1299 | 0.065803 | 0.23 | 0.1009 | 1 |
| Q9UQP3 | TNN | KYVVRYTsADGDTKE | S479 | 0.094 | 86.13 | 1299 | 0.074722 | 0.094 | 0.0747 | 1 |
| Q9UQP3 | TNN | KADtNALtEIDSPAN | T529 | 0.009 | 86.43 | 1299 | 0.006147 | 0.009 | 0.0061 | 0.9286 |
| Q9UQP3 | TNN | IDKYVVRyTSADDQE | Y565 | 0.098 | 88.57 | 1299 | 0.042991 | 0.187 | 0.0798 | 1 |
| Q9H497 | TOR3A | TQQLCHQtLFIFDEA | T230 | 0 | 95.81 | 397 | 0 | 0 | 0 | 1 |
| Q9HCH5 | SYTL2 | KktLNPVyNEILRYK | Y696 | 0.237 | 96.99 | 934 | 0.103968 | 0.372 | 0.1269 | 0.8667 |
| B2RTY4 | MYO9A | GEKLILPyKLAEAVT | Y486 | 0.022 | 83.1 | 2548 | 0.009651 | 0.217 | 0.0952 | 1 |
| P26358 | DNMT1 | LNRFTEDsLLRHAQF | S549 | 0.05 | 87.83 | 1616 | 0.039746 | 0.05 | 0.0397 | 1 |
| O95932 | TGM6 | ITISySKyKEDLTED | Y572 | 0.003 | 94.83 | 706 | 0.001316 | 0.003 | 0.0013 | 0.9333 |
| P49753 | ACOT2 | GVGLLGIsKGGELCL | S294 | 0.144 | 98.45 | 483 | 0.114468 | 0.144 | 0.1145 | 1 |
| Q96LJ7 | DHRS1 | GHYFCSVyGARLMVP | Y133 | 0.236 | 98.92 | 313 | 0.10353 | 0.236 | 0.1035 | 1 |
| Q96LJ7 | DHRS1 | DPNILsLsGKVLPSC | S249 | 0.065 | 98.63 | 313 | 0.051669 | 0.065 | 0.0517 | 1 |
| Q8N1F7 | NUP93 | VRLLMLYtRKFESTD | T536 | 0.001 | 91.45 | 819 | 0.000683 | 0.001 | 0.0007 | 0.8889 |

| | | | | | | | | | | |
|---|---|---|---|---|---|---|---|---|---|---|
| Q8N1F7 | NUP93 | NMFLRCVsELVIESR | S571 | 0.057 | 88.34 | 819 | 0.04531 | 0.057 | 0.0453 | 1 |
| Q8N1F7 | NUP93 | ELMNkLLsPVVPQIs | S647 | 0.119 | 88.51 | 819 | 0.094595 | 0.119 | 0.0946 | 1 |
| Q15005 | SPCS2 | DkYTLKLtFIsGRTK | T175 | 0.085 | 78.86 | 226 | 0.058056 | 0.085 | 0.0581 | 0.8571 |
| Q15005 | SPCS2 | TLKLtFIsGRTKQQR | S178 | 0.128 | 68.06 | 226 | 0.101749 | 0.165 | 0.1312 | 1 |
| P55060 | CSE1L | EQIQKQLsDAISIIG | S112 | 0.072 | 86.92 | 971 | 0.057234 | 0.072 | 0.0572 | 0.9 |
| P55060 | CSE1L | DAFALPLtNLFKAtI | T185 | 0.014 | 97.93 | 971 | 0.009562 | 0.014 | 0.0096 | 1 |
| P55060 | CSE1L | LtNLFKAtIELCSTH | T191 | 0.004 | 96.11 | 971 | 0.002732 | 0.004 | 0.0027 | 1 |
| P55060 | CSE1L | GLLELLksQICDNAA | S269 | 0.058 | 95.7 | 971 | 0.046105 | 0.058 | 0.0461 | 1 |
| P0DPD8 | F1AKMT4-E( | VEyWDQRyQGAADSA | Y30 | 0.289 | 91.65 | 883 | 0.12678 | 0.289 | 0.1268 | 0.8462 |
| Q96JS3 | PGBD1 | FTSVKLLsALKKKGV | S637 | 0.128 | 93.35 | 809 | 0.101749 | 0.165 | 0.1312 | 0.8571 |
| Q99062 | CSF3R | QRQLDPRtVQLFWKP | T351 | 0.16 | 87.31 | 836 | 0.109282 | 0.16 | 0.1093 | 1 |
| Q99062 | CSF3R | PVPLEEDsGRIQGYV | S365 | 0.067 | 74.27 | 836 | 0.053259 | 0.067 | 0.0533 | 1 |
| O95948 | ONECUT2 | GQLEEINtKEVAQRI | T335 | 0.057 | 95.59 | 504 | 0.038932 | 0.057 | 0.0389 | 1 |
| P21673 | SAT1 | SIVGFAMyyFTYDPW | Y77 | 0.212 | 98.88 | 171 | 0.093001 | 0.212 | 0.093 | 1 |
| Q14527 | HLTF | NIKSLVVsQFTTFLS | S858 | 0.001 | 92.25 | 1009 | 0.000795 | 0.144 | 0.1145 | 0.8571 |
| Q9NXZ2 | DDX43 | IIQEQPEsLVKIFGS | S110 | 0.03 | 86.1 | 648 | 0.023847 | 0.03 | 0.0238 | 1 |
| Q96QE4 | LRRC37B | KyLDMGTtHItLTTL | T688 | 0.035 | 95.86 | 947 | 0.023905 | 0.035 | 0.0239 | 1 |
| Q96G23 | CERS2 | NATLEHFyLTSGkQP | Y88 | 0.107 | 94.74 | 380 | 0.046939 | 0.107 | 0.0469 | 1 |
| P43490 | NAMPT | kyEEtVFyGLQyILN | Y60 | 0.019 | 98.79 | 491 | 0.008335 | 0.019 | 0.0083 | 1 |
| P43490 | NAMPT | QKKILAkyLLETSGN | Y175 | 0.169 | 97.53 | 491 | 0.074138 | 0.169 | 0.0741 | 1 |
| P43490 | NAMPT | GMkQkMWsIENIAFG | S374 | 0 | 98.47 | 491 | 0 | 0 | 0 | 1 |
| P43490 | NAMPT | IENIAFGsGGGLLQk | S382 | 0.016 | 96.85 | 491 | 0.012719 | 0.016 | 0.0127 | 0.8182 |
| Q96BD0 | SLCO4A1 | PHFTAGRyEVELDAG | Y197 | 0.284 | 83.94 | 722 | 0.124587 | 0.284 | 0.1246 | 1 |
| Q5S007 | LRRK2 | RVEKLHLsHNKLKEI | S1253 | 0.066 | 92.07 | 2527 | 0.052464 | 0.066 | 0.0525 | 1 |
| P16284 | PECAM1 | HASsVPRsKILTVRV | S587 | 0.065 | 94.85 | 738 | 0.051669 | 0.065 | 0.0517 | 1 |
| O60231 | DHX16 | IRVRKAItAGYFYHT | T944 | 0 | 94.92 | 1041 | 0 | 0 | 0 | 1 |
| P18621 | RPL17 | DPENPTksCKSRGSN | S14 | 0.018 | 94.72 | 184 | 0.014308 | 0.018 | 0.0143 | 1 |
| Q9Y5G2 | PCDHGB2 | DFEIAssyTLsIEAK | Y315 | 0.13 | 94.31 | 931 | 0.057029 | 0.13 | 0.057 | 1 |
| O75096 | LRP4 | HFPMDIHtLHPQRQP | T689 | 0.021 | 95.09 | 1905 | 0.014343 | 0.021 | 0.0143 | 1 |
| O75096 | LRP4 | DYGSQRLyWADAGMK | Y918 | 0.007 | 95.94 | 1905 | 0.003071 | 0.007 | 0.0031 | 1 |
| O75096 | LRP4 | ITTGLQTtDGLAVDA | T1126 | 0.014 | 91.49 | 1905 | 0.009562 | 0.014 | 0.0096 | 1 |

| | | | | | | | | | | |
|---|---|---|---|---|---|---|---|---|---|---|
| Q8N3P4 | VPS8 | HsLLKGIsAQIVSAA | S142 | 0.099 | 93.69 | 1428 | 0.078697 | 0.187 | 0.1486 | 1 |
| P18505 | GABRB1 | VNMDYTLtMYFQQSW | T85 | 0.09 | 96.72 | 474 | 0.061471 | 0.159 | 0.1137 | 1 |
| P18505 | GABRB1 | VFLALLEyAFVNYIF | Y324 | 0.21 | 87.44 | 474 | 0.092124 | 0.33 | 0.1228 | 1 |
| Q96KX2 | CAPZA3 | LLCALKLyVNDHYPK | Y128 | 0.147 | 97.92 | 299 | 0.064487 | 0.244 | 0.107 | 0.9167 |
| Q8IYK2 | TEKTL1 | MERDMEKsEVLLKTL | S220 | 0.003 | 95.96 | 499 | 0.002385 | 0.114 | 0.0906 | 1 |
| P48643 | CCT5 | VANtMrTsLGPNGLD | S51 | 0.028 | 92.81 | 541 | 0.022258 | 0.033 | 0.0344 | 1 |
| P48643 | CCT5 | PIrIADGyEQAARVA | Y137 | 0.023 | 96.63 | 541 | 0.01009 | 0.023 | 0.0101 | 1 |
| P48643 | CCT5 | DTEPLIQtAkTTLGS | T168 | 0.016 | 90.83 | 541 | 0.010928 | 0.016 | 0.0109 | 1 |
| P48643 | CCT5 | EVIPMALsENsGMNP | S463 | 0.006 | 94.9 | 541 | 0.004769 | 0.006 | 0.0048 | 1 |
| P48643 | CCT5 | PMALsENsGMNPIQt | S466 | 0.176 | 92.82 | 541 | 0.139905 | 0.176 | 0.1399 | 1 |
| P48643 | CCT5 | kQQHVIEtLIGkkQQ | T509 | 0.122 | 93.28 | 541 | 0.083328 | 0.216 | 0.1444 | 1 |
| P48643 | CCT5 | kQQIsLAtQMVRMIL | T521 | 0.103 | 96.62 | 541 | 0.07035 | 0.103 | 0.0703 | 1 |
| P09110 | ACAA1 | TVNRQCssGLQAVAs | S125 | 0 | 98.79 | 424 | 0 | 0 | 0 | 1 |
| P30048 | PRDX3 | TKQISRDyGVLLEGs | Y172 | 0.024 | 98.45 | 256 | 0.010528 | 0.024 | 0.0105 | 1 |
| P30048 | PRDX3 | VGrSVEEtLrLVKAF | T212 | 0.073 | 98.78 | 256 | 0.04986 | 0.073 | 0.0499 | 0.8 |
| P56282 | POLE2 | AIVLGMItQLkEGkF | T188 | 0.113 | 96.98 | 527 | 0.077181 | 0.113 | 0.0772 | 1 |
| P56282 | POLE2 | PVYWAyDyALRVYPV | Y467 | 0.151 | 97.29 | 527 | 0.066242 | 0.151 | 0.0662 | 0.8333 |
| P63173 | RPL38 | EIkDFLLtArRKDAK | T14 | 0.155 | 95.92 | 70 | 0.105867 | 0.155 | 0.1059 | 1 |
| Q8IVS2 | MCAT | LLAQQLVsPVKWEQT | S322 | 0.18 | 98.76 | 390 | 0.143085 | 0.18 | 0.1431 | 0.8 |
| O75900 | MMP23B | WDHFNLTyRILSFPR | Y95 | 0.13 | 98.7 | 390 | 0.057029 | 0.13 | 0.057 | 0.9167 |
| Q9Y394 | DHRS7 | NGGMSQRsLCMDTSL | S145 | 0.091 | 94.1 | 339 | 0.072337 | 0.091 | 0.0723 | 1 |
| Q9Y394 | DHRS7 | RKQGKIVtVNsILGI | T187 | 0.001 | 98.76 | 339 | 0.000683 | 0.001 | 0.0007 | 0.8182 |
| Q9Y394 | DHRS7 | GKIVtVNsILGIISV | S190 | 0.1 | 98.02 | 339 | 0.079491 | 0.1 | 0.0795 | 1 |
| Q9Y394 | DHRS7 | LSIGYCAsKHALRGF | S206 | 0 | 98.48 | 339 | 0 | 0 | 0 | 1 |
| Q7Z5L0 | VMO1 | GAYLVAFsLRVEAPT | S124 | 0 | 98.32 | 202 | 0 | 0 | 0 | 0.875 |
| Q9NYY3 | PLK2 | DLTNNkVyAAKIIPH | Y108 | 0.209 | 97.95 | 685 | 0.091685 | 0.209 | 0.0917 | 1 |
| Q9NYY3 | PLK2 | WALGCVMyTMLLGRP | Y271 | 0.011 | 98.45 | 685 | 0.004826 | 0.011 | 0.0048 | 1 |
| O75417 | POLQ | INQFMTEtVKNCKRD | T2430 | 0.101 | 90.3 | 2590 | 0.068984 | 0.101 | 0.069 | 1 |
| Q8NHP1 | AKR7L | rRMDAPtsAAVtRAF | S28 | 0 | 98.36 | 331 | 0 | 0 | 0 | 1 |
| Q8NHP1 | AKR7L | RHFGLRFyAyNPLAG | Y195 | 0.063 | 98.94 | 331 | 0.027637 | 0.126 | 0.0553 | 0.9286 |
| P78329 | CYP4F2 | CQEKPSEyIAAILEL | Y221 | 0.091 | 93.14 | 520 | 0.03992 | 0.091 | 0.0399 | 1 |

| | | | | | | | | | | |
|---|---|---|---|---|---|---|---|---|---|---|
| Q96MP8 | KCTD7 | FLGLMPyyKDHLERI | Y163 | 0.149 | 94.23 | 289 | 0.065364 | 0.149 | 0.0654 | 1 |
| Q9NW81 | DMAC2 | VHEKNRSyTWLEKQH | Y74 | 0.244 | 86.23 | 257 | 0.107039 | 0.244 | 0.107 | 0.9167 |
| Q9NVE5 | USP40 | DYLQGAPyyLKDGDT | Y1156 | 0.297 | 84.17 | 1235 | 0.13029 | 0.297 | 0.1303 | 1 |
| Q13516 | OLIG2 | RKLSKIAtLLLARNY | T151 | 0 | 96.3 | 323 | 0 | 0 | 0 | 1 |
| Q9UHJ3 | SFMBT1 | QDFDWADyLkQCGAE | Y361 | 0.077 | 95.77 | 866 | 0.033779 | 0.077 | 0.0338 | 1 |
| P25054 | APC | ASVLRNLsWRADVNS | S552 | 0.05 | 98.17 | 2843 | 0.039746 | 0.05 | 0.0397 | 1 |
| P35568 | IRS1 | GPARLEyyENEKkWR | Y47 | 0.159 | 98.24 | 1242 | 0.069751 | 0.159 | 0.0698 | 0.9091 |
| Q9UI17 | DMGDH | NVVNGPItySPDILP | T373 | 0.088 | 98.7 | 866 | 0.060105 | 0.088 | 0.0601 | 1 |
| Q9UI17 | DMGDH | RKVLQKLtSEDLSDD | T645 | 0.187 | 98.73 | 866 | 0.127724 | 0.187 | 0.1277 | 1 |
| Q9UI17 | DMGDH | REDSVALyDAIMNAG | Y695 | 0.111 | 98.87 | 866 | 0.048694 | 0.111 | 0.0487 | 1 |
| Q14690 | PDCD11 | VPEGTSPtTKLKVGK | T1079 | 0.042 | 86.47 | 1871 | 0.028687 | 0.042 | 0.0287 | 0.9 |
| Q8N0U4 | FAM185A | GDNCKIEtEHGTSIL | T176 | 0.054 | 93.02 | 392 | 0.036883 | 0.054 | 0.0369 | 0.9286 |
| Q9HDC9 | APMAP | EVKLLLssETPIEGk | S188 | 0.056 | 97.6 | 416 | 0.044515 | 0.056 | 0.0445 | 1 |
| Q9HDC9 | APMAP | TQDGRkIyFtDsSSk | Y213 | 0.123 | 98.8 | 416 | 0.053958 | 0.123 | 0.054 | 1 |
| Q9HDC9 | APMAP | DGRkIyFtDsSSkWQ | T215 | 0 | 98.84 | 416 | 0 | 0 | 0 | 1 |
| Q9HDC9 | APMAP | RkIyFtDsSSkWQRR | S217 | 0.002 | 98.59 | 416 | 0.00159 | 0.002 | 0.0016 | 1 |
| Q9HDC9 | APMAP | DDGRLLEyDtVTREV | Y242 | 0.042 | 98.7 | 416 | 0.018425 | 0.042 | 0.0184 | 1 |
| Q9HDC9 | APMAP | FPNGVQLsPAEDFVL | S265 | 0.062 | 97.44 | 416 | 0.049285 | 0.062 | 0.0493 | 1 |
| O15232 | MATN3 | LDIGPADtRVAVVNY | T120 | 0.017 | 90.9 | 486 | 0.011611 | 0.063 | 0.043 | 1 |
| O15519 | CFLAR | SFVCVLVsRGGSQSV | S314 | 0.004 | 94.63 | 480 | 0.00318 | 0.004 | 0.0032 | 1 |
| Q7Z3E1 | TIPARP | FAkkASysHNFSkKS | S572 | 0.043 | 96.96 | 657 | 0.034181 | 0.043 | 0.0342 | 0.9091 |
| Q9UNH7 | SNX6 | LLYRRSRsLVDyENA | S316 | 0.073 | 94.48 | 406 | 0.058029 | 0.073 | 0.058 | 1 |
| Q15464 | SHB | CSYLVRNsQTSKHDY | S437 | 0.172 | 93.33 | 509 | 0.136725 | 0.172 | 0.1367 | 0.8571 |
| P43405 | SYK | LPMDtEVyEsPyADP | Y348 | 0.039 | 96.48 | 635 | 0.017109 | 0.3 | 0.1316 | 1 |
| P43405 | SYK | ARNVLLVtQHyAkIS | T504 | 0.018 | 96.84 | 635 | 0.012294 | 0.036 | 0.0246 | 1 |
| P43405 | SYK | VLLVtQHyAkISDFG | Y507 | 0 | 98.42 | 635 | 0 | 0 | 0 | 0.9444 |
| Q92830 | KAT2A | ERLRSRYyVTRkLFV | Y787 | 0.112 | 93.35 | 837 | 0.049133 | 0.112 | 0.0491 | 0.9286 |
| A6NHN0 | OTOL1 | NCSIPGTyVFSyHIT | Y388 | 0 | 97.73 | 477 | 0 | 0 | 0 | 0.8125 |
| P33121 | ACSL1 | FQRGIQVsNNGPCLG | S103 | 0.161 | 97.79 | 698 | 0.127981 | 0.161 | 0.128 | 1 |
| P33121 | ACSL1 | RWLLDFAskRKEAEL | S394 | 0.154 | 92.89 | 698 | 0.122417 | 0.154 | 0.1224 | 1 |
| P33121 | ACSL1 | SGIIRNNsLWDRLIF | S410 | 0.036 | 91.06 | 698 | 0.028617 | 0.036 | 0.0286 | 1 |

| | | | | | | | | | | |
|---|---|---|---|---|---|---|---|---|---|---|
| Q8TF46 | DIS3L | DMLPSVLsADLCSLL | S544 | 0.016 | 92.48 | 1054 | 0.012719 | 0.016 | 0.0127 | 1 |
| Q8TF46 | DIS3L | FTSPIRRysDIVVHR | Y802 | 0 | 98.4 | 1054 | 0 | 0.248 | 0.1088 | 1 |
| Q8TF46 | DIS3L | TSPIRRysDIVVHRL | S803 | 0.016 | 98.24 | 1054 | 0.012719 | 0.157 | 0.1248 | 1 |
| P05813 | CRYBA1 | YHIERLMsFRPICSA | S112 | 0 | 98.43 | 215 | 0 | 0 | 0 | 1 |
| P05813 | CRYBA1 | WFNNEVGsMKIQSGA | S160 | 0 | 97.91 | 215 | 0 | 0 | 0 | 1 |
| Q8WUM4 | PDCD6IP | GGEEQAQyCRAAEEL | Y39 | 0.077 | 95.41 | 868 | 0.033779 | 0.077 | 0.0338 | 0.9 |
| Q8WUM4 | PDCD6IP | ISPDTVGtLSLIMLA | T186 | 0.127 | 97.16 | 868 | 0.086743 | 0.127 | 0.0867 | 1 |
| Q8WUM4 | PDCD6IP | IMQANAEyHQSILAK | Y258 | 0.221 | 98.61 | 868 | 0.09695 | 0.369 | 0.1359 | 1 |
| Q8WUM4 | PDCD6IP | VPVSVQQsLAAyNQR | S366 | 0.073 | 93.25 | 868 | 0.058029 | 0.125 | 0.1068 | 1 |
| Q8WUM4 | PDCD6IP | VQQsLAAyNQRKADL | Y370 | 0.021 | 94.94 | 868 | 0.009212 | 0.021 | 0.0092 | 1 |
| Q8WUM4 | PDCD6IP | ERWQRtPsNELykPL | S481 | 0.002 | 92.63 | 868 | 0.00159 | 0.002 | 0.0016 | 1 |
| Q8WUM4 | PDCD6IP | RtPsNELykPLRAEG | Y485 | 0 | 94.75 | 868 | 0 | 0 | 0 | 1 |
| Q8WUM4 | PDCD6IP | VTELDrVyGGLttkV | Y608 | 0.024 | 95.65 | 868 | 0.010528 | 0.024 | 0.0105 | 1 |
| Q96RW7 | HMCN1 | AGTTERKyNLKVHVP | Y1351 | 0.023 | 89.25 | 5635 | 0.01009 | 0.023 | 0.0101 | 1 |
| Q96FL9 | GALNT14 | KARRLDPtEPIRTPI | T272 | 0.145 | 94.75 | 552 | 0.099037 | 0.145 | 0.099 | 1 |
| Q96FL9 | GALNT14 | AEVWMDEyKQYYYAA | Y366 | 0.27 | 96.63 | 552 | 0.118445 | 0.27 | 0.1184 | 1 |
| Q66K74 | MAP1S | VLDALLAskQHWDRD | S999 | 0.004 | 95.22 | 1059 | 0.00318 | 0.004 | 0.0032 | 1 |
| Q8WWR8 | NEU4 | PHsFAFYsDDHGRTW | S202 | 0.013 | 98.65 | 484 | 0.010334 | 0.013 | 0.0103 | 1 |
| Q96AG4 | LRRC59 | kATILDLsCNkLTTL | S47 | 0.102 | 98.2 | 307 | 0.081081 | 0.102 | 0.0811 | 1 |
| Q9NRH2 | SNRK | kQGLVKLtDFGFsNk | T157 | 0.189 | 89.05 | 765 | 0.12909 | 0.189 | 0.1291 | 0.8333 |
| Q6PI47 | KCTD18 | SLCRFKDsMLASMFs | S36 | 0.005 | 89.74 | 426 | 0.003975 | 0.005 | 0.004 | 1 |
| Q6PI47 | KCTD18 | PTVWVLHyLNtsGAS | Y147 | 0.188 | 85.77 | 426 | 0.082473 | 0.188 | 0.0825 | 1 |
| Q99543 | DNAJC2 | DWKNQDHyAVLGLGH | Y90 | 0.001 | 95.91 | 621 | 0.000439 | 0.001 | 0.0004 | 1 |
| Q99543 | DNAJC2 | IkEGDNDyFTCItkA | Y137 | 0.097 | 90.62 | 621 | 0.042553 | 0.097 | 0.0426 | 1 |
| Q86VW2 | ARHGEF25 | LKDFLKYyNRAGMDT | Y309 | 0.06 | 94.38 | 580 | 0.026321 | 0.06 | 0.0263 | 1 |
| Q7Z2W4 | ZC3HAV1 | IHHLHPEyVRVSEHF | Y740 | 0.263 | 96.61 | 902 | 0.115374 | 0.263 | 0.1154 | 0.8 |
| P11177 | PDHB | VINSAAKtYYMSGGL | T130 | 0.208 | 98.89 | 359 | 0.142067 | 0.208 | 0.1421 | 1 |
| P11177 | PDHB | DMEtIEAsVMKTNHL | S282 | 0.033 | 98.89 | 359 | 0.026232 | 0.033 | 0.0262 | 1 |
| Q8N6C8 | LILRA3 | VSPSRRWsYRCYGYD | S194 | 0.078 | 91.04 | 439 | 0.062003 | 0.078 | 0.062 | 1 |
| P05023 | ATP1A1 | TRNIAFFsTNCVEGt | S246 | 0.177 | 80.37 | 1023 | 0.1407 | 0.177 | 0.1407 | 0.8333 |
| P05023 | ATP1A1 | VVytGDrtVMGRIAT | T265 | 0.001 | 87.98 | 1023 | 0.000683 | 0.001 | 0.0007 | 0.9091 |

| | | | | | | | | | | |
|---|---|---|---|---|---|---|---|---|---|---|
| P05023 | ATP1A1 | EAVETLGsTSTICSD | S369 | 0.028 | 95.75 | 1023 | 0.022258 | 0.028 | 0.0223 | 1 |
| P05023 | ATP1A1 | STICSDktGTLTQNR | T378 | 0.087 | 87.67 | 1023 | 0.059422 | 0.087 | 0.0594 | 0.8889 |
| P05023 | ATP1A1 | RAVAGDAsESALLkC | S452 | 0.137 | 89.08 | 1023 | 0.108903 | 0.137 | 0.1089 | 1 |
| P05023 | ATP1A1 | PFNsTNkyQLSIHkN | Y488 | 0.176 | 92.33 | 1023 | 0.077209 | 0.176 | 0.0772 | 1 |
| P05023 | ATP1A1 | RILDrCSsILLHGkE | S520 | 0.174 | 95.69 | 1023 | 0.138315 | 0.174 | 0.1383 | 1 |
| P05023 | ATP1A1 | GIkVIMVtGDHPItA | T617 | 0.021 | 92.67 | 1023 | 0.014343 | 0.021 | 0.0143 | 1 |
| P05023 | ATP1A1 | IISEGNEtVEDIAAR | T640 | 0.016 | 92.96 | 1023 | 0.010928 | 0.016 | 0.0109 | 1 |
| P05023 | ATP1A1 | TGDGVNDsPALkkAD | S722 | 0 | 86.12 | 1023 | 0 | 0 | 0 | 1 |
| P05023 | ATP1A1 | ICKTRRNsVFQQGMk | S943 | 0.075 | 96.6 | 1023 | 0.059619 | 0.075 | 0.0596 | 1 |
| Q8N5I3 | KCNRG | IFTTRFStIKQFPAs | T22 | 0.177 | 94.33 | 272 | 0.120893 | 0.177 | 0.1209 | 0.8333 |
| P31937 | HIBADH | KNLMKHGyPLIIYDV | Y63 | 0.065 | 98.13 | 336 | 0.028515 | 0.122 | 0.0507 | 1 |
| P31937 | HIBADH | LkKVKKGsLLIDsST | S127 | 0.008 | 98.51 | 336 | 0.006359 | 0.008 | 0.0064 | 1 |
| P31937 | HIBADH | KGsLLIDsSTIDPAV | S132 | 0 | 98.83 | 336 | 0 | 0 | 0 | 1 |
| P31937 | HIBADH | STIDPAVsKELAkEV | S140 | 0 | 98.55 | 336 | 0 | 0 | 0 | 1 |
| P31937 | HIBADH | GSLAHQIyRMMCAKG | Y310 | 0 | 98.78 | 336 | 0 | 0 | 0 | 1 |
| Q96JB8 | MPP4 | FQSAVPHtTRTKKSY | T462 | 0 | 95.12 | 637 | 0 | 0 | 0 | 1 |
| Q9P2D3 | HEATR5B | AAQNsRLsLQRTQAG | S548 | 0.141 | 93.09 | 2071 | 0.112083 | 0.141 | 0.1121 | 1 |
| Q13621 | SLC12A1 | KDCKLRIyVGGKINR | Y944 | 0.018 | 94.07 | 1099 | 0.007896 | 0.018 | 0.0079 | 1 |
| Q13621 | SLC12A1 | AANLIVLsLPVARKG | S1056 | 0.04 | 93.84 | 1099 | 0.031797 | 0.062 | 0.0557 | 1 |
| O94823 | ATP10B | GIQLWVLtGDKQETA | T915 | 0.022 | 86.19 | 1461 | 0.015026 | 0.132 | 0.0902 | 1 |
| P19224 | UGT1A6 | FyCLFsKyEELASAV | Y229 | 0.025 | 95.2 | 532 | 0.010967 | 0.025 | 0.011 | 1 |
| P53396 | ACLY | YkFICTTsAIQNRFK | S23 | 0.11 | 88.31 | 1101 | 0.087441 | 0.11 | 0.0874 | 1 |
| P53396 | ACLY | VVTKDGVyVLDLAAK | Y213 | 0.272 | 98.01 | 1101 | 0.119322 | 0.334 | 0.1465 | 1 |
| P53396 | ACLY | LDAksGAsLkLTLLN | S263 | 0.04 | 93.48 | 1101 | 0.031797 | -0.019 | 0.0953 | 1 |
| P53396 | ACLY | QTyDYAKtILsLMTR | T322 | 0.097 | 98.58 | 1101 | 0.066252 | 0.097 | 0.0663 | 1 |
| Q14CX7 | NAA25 | LKtELQFsDYYCLLA | S464 | 0 | 96.53 | 972 | 0 | 0 | 0 | 1 |
| Q14CX7 | NAA25 | IQHDTIGyLLTRYAE | Y544 | 0.106 | 95.02 | 972 | 0.046501 | 0.296 | 0.1299 | 0.8333 |
| Q14CX7 | NAA25 | TLWLRIRsLTLRLIS | S691 | 0.048 | 97.92 | 972 | 0.038156 | 0.048 | 0.0382 | 1 |
| Q6IEV9 | OR4C11 | LKLACMDtyMINLLL | T190 | 0.194 | 91.32 | 310 | 0.132505 | 0.194 | 0.1325 | 1 |
| Q9BZG1 | RAB34 | FSLQLWDtAGQERFK | T108 | 0.026 | 83.19 | 259 | 0.017758 | 0.026 | 0.0178 | 1 |
| Q99683 | MAP3K5 | ySGVLKIsDFGtSkR | S821 | 0.144 | 92.86 | 1374 | 0.114468 | 0.062 | 0.1298 | 1 |

| | | | | | | | | | | |
|---|---|---|---|---|---|---|---|---|---|---|
| Q8HWS3 | RFX6 | GIGIKEssAYYHsVy | S201 | 0.064 | 85.3 | 928 | 0.050875 | 0.105 | 0.0835 | 1 |
| Q9Y4E8 | USP15 | HLIDELDyILLPTEG | Y89 | 0.145 | 92.5 | 981 | 0.063609 | 0.288 | 0.1263 | 1 |
| Q9BY49 | PECR | KELLELGsNVVIAsR | S43 | 0 | 97.76 | 303 | 0 | 0 | 0 | 1 |
| Q9BY49 | PECR | RAGVyNLtKSLALEW | T182 | 0 | 98.86 | 303 | 0 | 0 | 0 | 1 |
| P08243 | ASNS | WRPkEAFsDGITsVk | S471 | 0.185 | 90.11 | 561 | 0.147059 | 0.185 | 0.1471 | 0.8 |
| Q9H7C9 | AAMDC | ______MtsPEIASLS | S3 | 0.1 | 98 | 122 | 0.079491 | 0.1 | 0.0795 | 0.9 |
| Q9H7C9 | AAMDC | WDWRETGtEHsPGVQ | T43 | 0.084 | 97.02 | 122 | 0.057373 | 0.084 | 0.0574 | 0.875 |
| Q9BXA7 | TSSK1B | DSGRMALsKTFCGSP | S172 | 0.056 | 92.58 | 367 | 0.044515 | 0.056 | 0.0445 | 1 |
| Q9BXA7 | TSSK1B | IPYQPKVyDIWSLGV | Y197 | 0.032 | 97.47 | 367 | 0.014038 | 0.032 | 0.014 | 1 |
| Q9BXA7 | TSSK1B | WSLGVILyIMVCGSM | Y207 | 0.076 | 98.76 | 367 | 0.03334 | 0.076 | 0.0333 | 1 |
| Q9BXA7 | TSSK1B | MVCGSMPyDDSNIKK | Y216 | 0.107 | 96.72 | 367 | 0.046939 | 0.107 | 0.0469 | 1 |
| O94923 | GLCE | ISTLVRAyLLTKDHI | Y458 | 0.34 | 98.9 | 617 | 0.149153 | 0.34 | 0.1492 | 1 |
| O94923 | GLCE | DTGSGTIyDLRHFML | Y560 | 0.21 | 98.8 | 617 | 0.092124 | 0.21 | 0.0921 | 1 |
| P49736 | MCM2 | YSDLrkEsMAtGSIP | S754 | 0.074 | 82.45 | 904 | 0.058824 | 0.074 | 0.0588 | 0.875 |
| Q5T4D3 | TMTC4 | AIKHRRKyPDCyyNL | Y583 | 0.183 | 97.46 | 741 | 0.080279 | 0.183 | 0.0803 | 1 |
| Q5T4D3 | TMTC4 | RKyPDCyyNLGRLyA | Y588 | 0.327 | 98.54 | 741 | 0.14345 | 0.327 | 0.1434 | 1 |
| Q5T4D3 | TMTC4 | yyNLGRLyADLNRHV | Y594 | 0.095 | 98.25 | 741 | 0.041675 | 0.095 | 0.0417 | 1 |
| Q5T4D3 | TMTC4 | KSQKYKEsEALFLKA | S671 | 0 | 97.63 | 741 | 0 | 0 | 0 | 1 |
| Q14004 | CDK13 | QDCHELWsKKRRRQK | S1022 | 0.055 | 89.4 | 1512 | 0.04372 | 0.055 | 0.0437 | 1 |
| Q9Y490 | TLN1 | HSRTVsDsIKKLITS | S1643 | 0.08 | 90.18 | 2541 | 0.063593 | 0.08 | 0.0636 | 1 |
| Q9Y5B8 | NME7 | rLLGPANsGVARTDA | S181 | 0 | 96.43 | 376 | 0 | 0 | 0 | 1 |
| Q96AX1 | VPS33A | LRLVCLQsVCNsGLk | S414 | 0.01 | 98.16 | 596 | 0.007949 | 0.01 | 0.0079 | 1 |
| P53794 | SLC5A3 | NIAHAkGstLMAGFL | S298 | 0 | 96.89 | 718 | 0 | 0 | 0 | 1 |
| P53794 | SLC5A3 | IAHAkGstLMAGFLK | T299 | 0 | 97.07 | 718 | 0 | 0 | 0 | 1 |
| P53794 | SLC5A3 | IVVPGMIsRILFTDD | S320 | 0 | 96.84 | 718 | 0 | 0 | 0 | 0.9 |
| P41091 | EIF2S3 | kAISGVHtVRFkNEL | T66 | 0.089 | 78.26 | 472 | 0.060788 | 0.218 | 0.1499 | 0.8182 |
| P41091 | EIF2S3 | VNIGsLstGGRVsAV | T413 | 0.045 | 90.84 | 472 | 0.030736 | 0.045 | 0.0307 | 1 |
| O43463 | SUV39H1 | LDyVEDVyTVDAAYY | Y308 | 0.299 | 94.62 | 412 | 0.131167 | 0.299 | 0.1312 | 0.9091 |
| Q9H0R6 | QRSL1 | GRSLREVsAALkQGQ | S10 | 0.017 | 96.42 | 528 | 0.013514 | 0.017 | 0.0135 | 0.9 |
| Q9H0R6 | QRSL1 | GsDtGGStRNPAAHC | T196 | 0 | 98.88 | 528 | 0 | 0 | 0 | 0.875 |
| Q13753 | LAMC2 | AQRLDPVyFVAPAKF | Y250 | 0.123 | 95.98 | 1193 | 0.053958 | 0.123 | 0.054 | 1 |

| | | | | | | | | | | |
|---|---|---|---|---|---|---|---|---|---|---|
| Q13753 | LAMC2 | GQSLSFDyRVDRGGR | Y273 | 0.038 | 94.87 | 1193 | 0.01667 | 0.038 | 0.0167 | 1 |
| C9JR72 | KBTBD13 | VVRGDTVytVNRMFT | Y404 | 0.117 | 96 | 458 | 0.051326 | 0.233 | 0.1022 | 1 |
| C9JR72 | KBTBD13 | VRGDTVytVNRMFTL | T405 | 0.059 | 94.37 | 458 | 0.040298 | 0.059 | 0.0403 | 1 |
| P02042 | HBD | kGtFsQLsELHCDkL | S90 | 0.011 | 97.18 | 147 | 0.008744 | 0.011 | 0.0087 | 1 |
| P02042 | HBD | TPQMQAAyQKVVAGV | Y131 | 0.003 | 98.35 | 147 | 0.001316 | 0.003 | 0.0013 | 0.9412 |
| Q9ULC4 | MCTS1 | REGPFYPtLRLLHkY | T81 | 0.054 | 98.79 | 181 | 0.036883 | 0.054 | 0.0369 | 0.9 |
| P17174 | GOT1 | FLFPFFDsAyQGFAS | S224 | 0.043 | 98.43 | 413 | 0.034181 | 0.043 | 0.0342 | 1 |
| P17174 | GOT1 | PsGRINVsGLTTkNL | S391 | 0.007 | 94.86 | 413 | 0.005564 | 0.007 | 0.0056 | 0.8889 |
| Q13126 | MTAP | RFSSRAEsFMFRtWG | S183 | 0 | 97.47 | 283 | 0 | 0 | 0 | 1 |
| Q2TAL8 | QRICH1 | LGAHSPStLLTTLMF | T621 | 0.063 | 95.45 | 776 | 0.04303 | 0.063 | 0.043 | 0.8182 |
| P47755 | CAPZA2 | QYNLDQFtPVKIEGY | T63 | 0.129 | 98.27 | 286 | 0.088109 | 0.213 | 0.1455 | 1 |
| P47755 | CAPZA2 | HYPNGVCtVyGkKID | T142 | 0.045 | 98.7 | 286 | 0.030736 | 0.045 | 0.0307 | 1 |
| P47755 | CAPZA2 | PNGVCtVyGkKIDGQ | Y144 | 0.049 | 97.98 | 286 | 0.021496 | 0.049 | 0.0215 | 1 |
| P13807 | GYS1 | PEFLSstsPLLPVDY | S486 | 0.164 | 92.43 | 737 | 0.130366 | 0.164 | 0.1304 | 1 |
| Q6UXS9 | CASP12 | VVIKENLtAQEMETA | T142 | 0.175 | 98.38 | 341 | 0.119527 | 0.175 | 0.1195 | 1 |
| Q9NUV9 | GIMAP4 | TEEEHKAtEKILKMF | T134 | 0.091 | 91.16 | 329 | 0.062154 | 0.091 | 0.0622 | 1 |
| Q9NUV9 | GIMAP4 | GCYTNRMyQRAEEEI | Y226 | 0.026 | 91.09 | 329 | 0.011406 | 0.026 | 0.0114 | 1 |
| Q9NUV9 | GIMAP4 | EEEIQkQtQAMQELH | T237 | 0.175 | 94.94 | 329 | 0.119527 | 0.175 | 0.1195 | 1 |
| Q9NUV9 | GIMAP4 | kARIREEyEEkIRKL | Y259 | 0.272 | 97.24 | 329 | 0.119322 | 0.272 | 0.1193 | 1 |
| Q9NUV9 | GIMAP4 | LAEQEAHyAVRQQRA | Y290 | 0.222 | 96.3 | 329 | 0.097388 | 0.222 | 0.0974 | 1 |
| Q6IS24 | GALNT17 | WWDAGDPsLPIRtPA | S317 | 0.158 | 96.35 | 598 | 0.125597 | 0.158 | 0.1256 | 1 |
| Q92747 | ARPC1A | DRTQIALsPNNHEVH | S26 | 0 | 96.45 | 370 | 0 | 0 | 0 | 1 |
| Q5SRE5 | NUP188 | RQQFEELykTEAPTW | Y181 | 0.108 | 91.94 | 1749 | 0.047378 | 0.108 | 0.0474 | 1 |
| Q9NX05 | FAM120C | LkLARTVsRQQQQQH | S31 | 0.047 | 80.52 | 1096 | 0.037361 | 0.174 | 0.1383 | 1 |
| Q9NX05 | FAM120C | LHRHELDtFLAQAVS | T789 | 0 | 98.11 | 1096 | 0 | 0 | 0 | 1 |
| Q9NRA0 | SPHK2 | GLAtLHtyRGRLSyL | Y372 | 0.213 | 97.59 | 654 | 0.09344 | 0.213 | 0.0934 | 1 |
| Q9NRA0 | SPHK2 | tyRGRLSyLPATVEP | Y378 | 0.141 | 98.04 | 654 | 0.061855 | 0.248 | 0.1088 | 1 |
| P50749 | RASSF2 | SAEEFALyVVHTSGE | Y224 | 0.037 | 98.18 | 326 | 0.016231 | 0.037 | 0.0162 | 1 |
| Q9UH77 | KLHL3 | IKDVDGQtLSKLIDY | T100 | 0.09 | 95.38 | 587 | 0.061471 | 0.09 | 0.0615 | 1 |
| Q9UH77 | KLHL3 | LLIKNPRtKPRTPVS | T291 | 0.148 | 85.68 | 587 | 0.101086 | 0.148 | 0.1011 | 1 |
| Q02388 | COL7A1 | QASSSSVtITWTRVP | T703 | 0.219 | 89.8 | 2944 | 0.14958 | 0.219 | 0.1496 | 1 |

| | | | | | | | | | | |
|---|---|---|---|---|---|---|---|---|---|---|
| Q9BSE4 | HERPUD2 | KTHLSNVyPskPLtk | Y45 | 0.054 | 87.91 | 406 | 0.023689 | 0.054 | 0.0237 | 0.9286 |
| Q8TCU6 | PREX1 | IFRGRINtEVMEVEN | T333 | 0.033 | 90.9 | 1659 | 0.022539 | 0.033 | 0.0225 | 1 |
| P42696 | RBM34 | TSSRDKRsVFVGNLP | S288 | 0.134 | 93.3 | 430 | 0.106519 | 0.134 | 0.1065 | 1 |
| P39059 | COL15A1 | FFRDFAIsVVVKPSS | S94 | 0.091 | 95.62 | 1388 | 0.072337 | 0.091 | 0.0723 | 0.9 |
| Q9BRQ8 | AIFM2 | GALTFLLsMGRNDGV | S331 | 0.01 | 97.22 | 373 | 0.007949 | 0.01 | 0.0079 | 1 |
| Q15699 | ALX1 | REQLALRtELtEARV | T169 | 0.06 | 97.54 | 326 | 0.040981 | 0.06 | 0.041 | 1 |
| Q8N8U9 | BMPER | EISWDGDsFVEVMAA | S479 | 0.003 | 81.27 | 685 | 0.002385 | 0.152 | 0.1208 | 1 |
| A6NKT7 | RGPD3 | MMACDRLsQsGHMLL | S353 | 0.001 | 95.69 | 1758 | 0.000795 | 0.001 | 0.0008 | 1 |
| P62937 | PPIA | GEPLGRVsFELFADk | S21 | 0.051 | 98.75 | 165 | 0.040541 | 0.051 | 0.0405 | 1 |
| P04040 | CAT | FEHIGkKtPIAVRFs | T107 | 0.018 | 98.91 | 527 | 0.012294 | 0.018 | 0.0123 | 1 |
| P04040 | CAT | PIAVRFstVAGESGS | T115 | 0.045 | 98.73 | 527 | 0.030736 | 0.045 | 0.0307 | 1 |
| P04040 | CAT | WDLVGNNtPIFFIRD | T150 | 0.004 | 98.74 | 527 | 0.002732 | 0.004 | 0.0027 | 1 |
| P04040 | CAT | LFPSFIHsQKRNPQT | S167 | 0.073 | 98.72 | 527 | 0.058029 | 0.073 | 0.058 | 1 |
| P04040 | CAT | NANGEAVyCkFHYkT | Y231 | 0.105 | 98.93 | 527 | 0.046062 | 0.105 | 0.0461 | 1 |
| P04040 | CAT | KVWPHkDyPLIPVGk | Y308 | 0.187 | 98.77 | 527 | 0.082034 | 0.187 | 0.082 | 1 |
| P04040 | CAT | RHRLGPNyLHIPVNC | Y370 | 0.118 | 98.89 | 527 | 0.051765 | 0.118 | 0.0518 | 1 |
| P04040 | CAT | VTQVRAFyVNVLNEE | Y447 | 0.135 | 98.88 | 527 | 0.059223 | 0.135 | 0.0592 | 1 |
| Q9UBF8 | PI4KB | LPTkEQKtQRLISEL | T349 | 0.151 | 94.48 | 816 | 0.103135 | 0.151 | 0.1031 | 0.8889 |
| Q8NBF2 | NHLRC2 | APGGrGRsLsGLLPA | S10 | 0.054 | 91.89 | 726 | 0.042925 | 0.125 | 0.0994 | 1 |
| Q8NBF2 | NHLRC2 | HKDKLFLyTSIALKY | Y190 | 0 | 98.81 | 726 | 0 | 0 | 0 | 1 |
| Q8NBF2 | NHLRC2 | TSIALKYyKDRGQIR | Y198 | 0.002 | 98.79 | 726 | 0.000877 | 0.002 | 0.0009 | 1 |
| O43861 | ATP9B | EVNsQLYsKLtVRGK | S217 | 0.149 | 92.96 | 1147 | 0.118442 | 0.149 | 0.1184 | 1 |
| Q9H1I8 | ASCC2 | MQkRLHRsVFLTFLR | S122 | 0.032 | 97.82 | 757 | 0.025437 | 0.032 | 0.0254 | 1 |
| Q86TB9 | PATL1 | YERRYLLsLEEERPA | S546 | 0.117 | 87.64 | 770 | 0.093005 | 0.117 | 0.093 | 1 |
| O14939 | PLD2 | LGINSGysKRALMLL | S416 | 0.035 | 98.2 | 933 | 0.027822 | 0.035 | 0.0278 | 1 |
| O14939 | PLD2 | FFWLGKDySNLITKD | Y511 | 0.013 | 98.62 | 933 | 0.005703 | 0.013 | 0.0057 | 0.875 |
| O14939 | PLD2 | FIQRWNFtKTTKAKy | T566 | 0 | 96.02 | 933 | 0 | 0 | 0 | 1 |
| O14939 | PLD2 | tKTTKAKyKTPTyPy | Y573 | 0.077 | 95.51 | 933 | 0.033779 | 0.077 | 0.0338 | 1 |
| O14939 | PLD2 | AKyKTPTyPyLLPKS | Y578 | 0.057 | 97.52 | 933 | 0.025005 | 0.057 | 0.025 | 1 |
| Q5JS13 | RALGPS1 | EDDNYkLsLRIEPGS | S283 | 0 | 97.54 | 557 | 0 | 0 | 0 | 1 |
| Q9NT68 | TENM2 | SLLKItMtQstVPLN | T1088 | 0.08 | 95.205 | 2774 | 0.054641 | 0.08 | 0.0546 | 1 |

| | | | | | | | | | | |
|---|---|---|---|---|---|---|---|---|---|---|
| Q9NT68 | TENM2 | VVSVGFEyETCPSLI | Y1150 | 0.044 | 92.23 | 2774 | 0.019302 | 0.044 | 0.0193 | 1 |
| Q9NT68 | TENM2 | AIDATLMsPRGIAVD | S1357 | 0.155 | 93.8 | 2774 | 0.123212 | 0.155 | 0.1232 | 1 |
| P51797 | CLCN6 | KVLGVLFsVAGGLFV | S190 | 0 | 94.33 | 869 | 0 | 0 | 0 | 1 |
| Q96PD5 | PGLYRP2 | QsHPDLGtEGCWDQL | T245 | 0.168 | 93.08 | 576 | 0.114746 | 0.168 | 0.1147 | 1 |
| Q8TAV3 | CYP2W1 | LPHVPRCtAADtQLG | T368 | 0.03 | 93.28 | 490 | 0.02049 | 0.03 | 0.0205 | 1 |
| Q8TAV3 | CYP2W1 | PRCtAADtQLGGFLL | T372 | 0.098 | 91.34 | 490 | 0.066935 | 0.148 | 0.1011 | 1 |
| Q9H6S0 | YTHDC2 | RIGQtIGyQIRLEsR | Y274 | 0.063 | 95.88 | 1430 | 0.027637 | 0.063 | 0.0276 | 1 |
| Q15631 | TSN | VVQSLEQtAREILTL | T34 | 0.063 | 98.47 | 228 | 0.04303 | 0.063 | 0.043 | 1 |
| Q9Y3P9 | RABGAP1 | DNDEPLLsGSGDVSK | S524 | 0.083 | 89.87 | 1069 | 0.065978 | 0.083 | 0.066 | 1 |
| Q9UL54 | TAOK2 | VAIKKMSySGkQSNE | Y61 | 0.075 | 91.46 | 1235 | 0.032901 | 0.075 | 0.0329 | 1 |
| Q15542 | TAF5 | ELKGHtDtVCsLRFS | T713 | 0.215 | 95.8 | 800 | 0.146848 | 0.215 | 0.1468 | 1 |
| Q15542 | TAF5 | GHtDtVCsLRFSRDG | S716 | 0.065 | 95.06 | 800 | 0.051669 | 0.065 | 0.0517 | 1 |
| Q9UFW8 | CGGBP1 | NGGsIPKsDQLRRAY | S143 | 0.078 | 87.67 | 167 | 0.062003 | 0.078 | 0.062 | 1 |
| Q9H6Q4 | CIAO3 | HLTPDkIyHVtVMPC | Y239 | 0.164 | 98.52 | 476 | 0.071944 | 0.328 | 0.1439 | 1 |
| Q9H6Q4 | CIAO3 | PDkIyHVtVMPCyDk | T242 | 0 | 98.87 | 476 | 0 | 0 | 0 | 1 |
| P55786 | NPEPPS | PkDRVALsNMNVIDR | S214 | 0.016 | 98.82 | 919 | 0.012719 | 0.016 | 0.0127 | 1 |
| P55786 | NPEPPS | ADLRsPVyLTVLKHG | Y748 | 0.002 | 97.75 | 919 | 0.000877 | 0.002 | 0.0009 | 1 |
| P55786 | NPEPPS | IERVLGAtLLPDLIQ | T786 | 0.041 | 94.69 | 919 | 0.028004 | 0.041 | 0.028 | 1 |
| P34903 | GABRA3 | PGLGDAVtEVKTDIY | T91 | 0.062 | 96.66 | 492 | 0.042347 | 0.062 | 0.0423 | 1 |
| Q9NQC7 | CYLD | LFKkIFPsLELNITD | S772 | 0.092 | 96.24 | 956 | 0.073132 | 0.144 | 0.1145 | 1 |
| P06239 | LCK | DNCPEELyQLMRLCW | Y470 | 0.326 | 97.75 | 509 | 0.143012 | 0.326 | 0.143 | 0.8182 |
| Q8TAQ2 | SMARCC2 | NNAsHVVyPVPGNLE | Y179 | 0.225 | 93.33 | 1214 | 0.098704 | 0.225 | 0.0987 | 1 |
| O15303 | GRM6 | VRLAGGLtLGGLFPV | T40 | 0.118 | 92.95 | 877 | 0.080596 | 0.118 | 0.0806 | 1 |
| O15303 | GRM6 | LGLGTTLsYSALLtK | S664 | 0.047 | 95.64 | 877 | 0.037361 | 0.067 | 0.0533 | 1 |
| O15303 | GRM6 | LsYSALLtKtNRIyR | T670 | 0.085 | 94.6 | 877 | 0.058056 | 0.085 | 0.0581 | 1 |
| O15303 | GRM6 | YSALLtKtNRIyRIF | T672 | 0.029 | 91.44 | 877 | 0.019807 | 0.208 | 0.1421 | 1 |
| Q8N119 | MMP21 | ALAFRMWsEVtPLDF | S206 | 0.024 | 96.66 | 569 | 0.019078 | 0.024 | 0.0191 | 1 |
| Q8N119 | MMP21 | FRMWsEVtPLDFRED | T209 | 0.009 | 96.73 | 569 | 0.006147 | 0.009 | 0.0061 | 1 |
| Q8N119 | MMP21 | VCDVHIstLNM____ | T566 | 0.061 | 86.79 | 569 | 0.041664 | 0.061 | 0.0417 | 1 |
| P02689 | PMP2 | GVVCtRIyEKV____ | Y129 | 0.185 | 98.69 | 132 | 0.081157 | 0.185 | 0.0812 | 0.8333 |
| Q92616 | GCN1 | VPEVRTVsAkALGAM | S1677 | 0.001 | 90.48 | 2671 | 0.000795 | 0.001 | 0.0008 | 0.9091 |

| | | | | | | | | | | |
|---|---|---|---|---|---|---|---|---|---|---|
| Q8IZ83 | ALDH16A1 | kHQRLLWtLEsLVTG | T121 | 0 | 96.48 | 802 | 0 | 0 | 0 | 0.8182 |
| Q8IZ83 | ALDH16A1 | RLLWtLEsLVTGRAV | S124 | 0.025 | 94.65 | 802 | 0.019873 | 0.025 | 0.0199 | 1 |
| Q9Y399 | MRPS2 | KGIILFIsRNRQFSY | S146 | 0 | 96.72 | 296 | 0 | 0 | 0 | 1 |
| Q9Y399 | MRPS2 | RQQVEALyRLQGQkE | Y271 | 0.299 | 85.89 | 296 | 0.131167 | 0.299 | 0.1312 | 1 |
| Q8N158 | GPC2 | QENSAKVsAQVFQEC | S339 | 0.076 | 94.13 | 579 | 0.060414 | 0.076 | 0.0604 | 1 |
| Q9UKU6 | TRHDE | ItQQHFIyDISAKTK | Y666 | 0.222 | 87.72 | 1024 | 0.097388 | 0.222 | 0.0974 | 1 |
| P06396 | GSN | GQFYGGDsYIILyNy | S473 | 0 | 98.44 | 782 | 0 | 0 | 0 | 1 |
| P06396 | GSN | sYIILyNyRHGGRQG | Y480 | 0.161 | 97.96 | 782 | 0.070628 | 0.161 | 0.0706 | 0.8462 |
| P06396 | GSN | KEPAHLMsLFGGKPM | S537 | 0.079 | 97.44 | 782 | 0.062798 | 0.079 | 0.0628 | 1 |
| P06396 | GSN | PkAGALNsNDAFVLK | S590 | 0.038 | 95.75 | 782 | 0.030207 | 0.038 | 0.0302 | 1 |
| P06396 | GSN | LKTPsAAyLWVGTGA | Y603 | 0.314 | 98.14 | 782 | 0.137747 | 0.314 | 0.1377 | 1 |
| P06396 | GSN | ALTSAkRyIETDPAN | Y730 | 0.041 | 94.28 | 782 | 0.017986 | 0.147 | 0.0645 | 1 |
| P06396 | GSN | PANRDRRtPITVVKQ | T742 | 0.067 | 98.12 | 782 | 0.045762 | 0.067 | 0.0458 | 0.8889 |
| Q8NEK8 | TENT5D | PDIMKDAyVQKLVKV | Y130 | 0.001 | 97.46 | 389 | 0.000439 | 0.001 | 0.0004 | 1 |
| Q8NEK8 | TENT5D | EGMTKyDyLMTLHGV | Y298 | 0.001 | 97.82 | 389 | 0.000439 | 0.001 | 0.0004 | 1 |
| O43847 | NRDC | NSEVTVYyQsGtRSL | Y924 | 0.006 | 96.85 | 1151 | 0.002632 | 0.006 | 0.0026 | 1 |
| O43847 | NRDC | EVTVYyQsGtRSLRE | S926 | 0.151 | 93.98 | 1151 | 0.120032 | 0.151 | 0.12 | 1 |
| Q5W0Q7 | USPL1 | KMVLEKVsPIFMLHF | S415 | 0.13 | 96.31 | 1092 | 0.103339 | 0.13 | 0.1033 | 0.8571 |
| P55036 | PSMD4 | RIIAFVGsPVEDNEk | S115 | 0.013 | 84.49 | 377 | 0.010334 | 0.013 | 0.0103 | 1 |
| Q9NX20 | MRPL16 | GKGAIDHyVTPVkAG | Y150 | 0.24 | 96.99 | 251 | 0.105285 | 0.24 | 0.1053 | 1 |
| P17858 | PFKL | PTPFDrNyGTkLGVK | Y674 | 0.158 | 95.61 | 780 | 0.069312 | 0.158 | 0.0693 | 1 |
| O75832 | PSMD10 | DKDDAGWsPLHIAAS | S75 | 0.022 | 97.98 | 226 | 0.017488 | 0.022 | 0.0175 | 1 |
| Q9HAB8 | PPCS | VLVTSGGtkVPLEAR | T45 | 0 | 96.78 | 311 | 0 | 0 | 0 | 1 |
| Q9Y6C9 | MTCH2 | QLPGLFsyAQHIAsI | Y57 | 0.013 | 93.64 | 303 | 0.005703 | 0.013 | 0.0057 | 0.9167 |
| P16403 | H1-2 | kGtGAsGsFkLNkkA | S104 | 0.134 | 96.24 | 213 | 0.106519 | 0.134 | 0.1065 | 0.875 |
| Q9HCU4 | CELSR2 | DYKLERQyVLAVTAS | Y681 | 0.068 | 89.235 | 2923 | 0.029831 | 0.068 | 0.0298 | 1 |
| O95067 | CCNB2 | ITALLLAsKYEEMFS | S220 | 0 | 98.5 | 398 | 0 | 0 | 0 | 1 |
| O95067 | CCNB2 | LHFLRRAskAGEVDV | S274 | 0.143 | 96.75 | 398 | 0.113673 | 0.143 | 0.1137 | 1 |
| O75899 | GABBR2 | PPSSPPLsIMGLMPL | S57 | 0.071 | 95.36 | 941 | 0.056439 | 0.121 | 0.0985 | 1 |
| O75899 | GABBR2 | YQWKRVGtLtQDVQR | T200 | 0.021 | 98.04 | 941 | 0.014343 | 0.021 | 0.0143 | 1 |
| O75899 | GABBR2 | WKRVGtLtQDVQRFS | T202 | 0.012 | 97.99 | 941 | 0.008196 | 0.012 | 0.0082 | 1 |

| | | | | | | | | | | |
|---|---|---|---|---|---|---|---|---|---|---|
| Q9H410 | DSN1 | IsVDLAEskRLGCLL | S131 | 0.126 | 89.3 | 356 | 0.100159 | 0.126 | 0.1002 | 1 |
| Q86U86 | PBRM1 | KsYYkPDsPEyKAAC | S131 | 0.175 | 92.6 | 1689 | 0.13911 | 0.175 | 0.1391 | 1 |
| Q58FG0 | HSP90AA5P | KANQKHIyyITGETK | Y177 | 0.066 | 86.99 | 334 | 0.028953 | 0.066 | 0.029 | 1 |
| Q58FG0 | HSP90AA5P | ANQKHIyyITGETKD | Y178 | 0.17 | 88.8 | 334 | 0.074577 | 0.17 | 0.0746 | 1 |
| Q96FA3 | PELI1 | PSkAPVKyGELIVLG | Y17 | 0.107 | 98.47 | 418 | 0.046939 | 0.107 | 0.0469 | 1 |
| Q96FA3 | PELI1 | GDRGRRKsRFALFKR | S39 | 0.019 | 95.15 | 418 | 0.015103 | 0.019 | 0.0151 | 0.8571 |
| Q96FA3 | PELI1 | IsNKDQHsIsYtLsR | S76 | 0.028 | 95.11 | 418 | 0.022258 | 0.028 | 0.0223 | 1 |
| Q96FA3 | PELI1 | NKDQHsIsYtLsRAQ | S78 | 0.124 | 93.79 | 418 | 0.098569 | 0.124 | 0.0986 | 0.875 |
| Q96FA3 | PELI1 | YtLsRAQtVVVEYTH | T86 | 0.016 | 89.96 | 418 | 0.010928 | 0.016 | 0.0109 | 1 |
| Q8ND23 | CARMIL3 | TKLCLAKtAISPRGL | T312 | 0 | 91.03 | 1372 | 0 | 0 | 0 | 1 |
| Q8ND23 | CARMIL3 | DIGAKMLsKALQINS | S590 | 0.057 | 94.22 | 1372 | 0.04531 | 0.057 | 0.0453 | 1 |
| Q8ND23 | CARMIL3 | KALQINSsLRTILWD | S598 | 0.177 | 85.96 | 1372 | 0.1407 | 0.177 | 0.1407 | 1 |
| P98066 | TNFAIP6 | DPKQIFKsPGFPNEy | S148 | 0.052 | 95.14 | 277 | 0.041336 | 0.059 | 0.0629 | 1 |
| P98066 | TNFAIP6 | sPGFPNEyEDNQICY | Y155 | 0.045 | 94.5 | 277 | 0.019741 | 0.045 | 0.0197 | 1 |
| Q8IYJ3 | SYTL1 | AAARRRRsDPyVKSY | S301 | 0.001 | 92.69 | 562 | 0.000795 | 0.001 | 0.0008 | 1 |
| Q96MR6 | CFAP57 | TLELFKEyQEEAYSI | Y429 | 0.248 | 91.26 | 1250 | 0.108794 | 0.323 | 0.1417 | 1 |
| Q9NZ52 | GGA3 | kVTkRLHtLEEVNNN | T217 | 0.184 | 94.23 | 723 | 0.125675 | 0.184 | 0.1257 | 1 |
| Q9NZ52 | GGA3 | ETEDNDNsLGDILQA | S275 | 0.041 | 89.64 | 723 | 0.032592 | 0.041 | 0.0326 | 1 |
| Q8WUM0 | NUP133 | SKLAALAsDFSEDML | S981 | 0.153 | 91.37 | 1156 | 0.121622 | 0.153 | 0.1216 | 1 |
| O43315 | AQP9 | MNPARDLsPRLFTAL | S222 | 0.048 | 98.73 | 295 | 0.038156 | 0.048 | 0.0382 | 0.8333 |
| O95271 | TNKS | DTQGRNStPLHLAAG | T839 | 0.009 | 97.88 | 1327 | 0.006147 | 0.009 | 0.0061 | 1 |
| Q6ZMV9 | KIF6 | SHIPYRNsMMTSVLR | S300 | 0.035 | 90.65 | 814 | 0.027822 | 0.035 | 0.0278 | 1 |
| Q6ZMV9 | KIF6 | MTSVLRDsLGGNCMT | S309 | 0.027 | 92.56 | 814 | 0.021463 | 0.027 | 0.0215 | 1 |
| Q9Y2E4 | DIP2C | YGKLWTRsMKVAYSI | S359 | 0 | 95.58 | 1556 | 0 | 0 | 0 | 1 |
| Q9Y2E4 | DIP2C | LWHGILTsVMNMMHV | S555 | 0.001 | 96.91 | 1556 | 0.000795 | 0.001 | 0.0008 | 1 |
| Q9Y2E4 | DIP2C | LMKVNPLsWIQKVCQ | S576 | 0.01 | 92.11 | 1556 | 0.007949 | 0.01 | 0.0079 | 1 |
| P22492 | H1-6 | RGtGAsGsFkLSkKV | S108 | 0.136 | 95.47 | 207 | 0.108108 | 0.136 | 0.1081 | 0.875 |
| Q33E94 | RFX4 | HCQRILDtVIRANFD | T215 | 0.012 | 89.96 | 735 | 0.008196 | 0.012 | 0.0082 | 1 |
| Q33E94 | RFX4 | LNHLCQAsRTVIHSA | S333 | 0 | 96.85 | 735 | 0 | 0 | 0 | 1 |
| Q96S94 | CCNL2 | LPQVAMAtGQVLFQR | T101 | 0.005 | 97.26 | 520 | 0.003415 | 0.005 | 0.0034 | 0.9 |
| Q9BQW3 | EBF4 | HYRLRLVyNNGLRTE | Y117 | 0.1 | 95.92 | 602 | 0.043869 | 0.1 | 0.0439 | 1 |

| | | | | | | | | | | |
|---|---|---|---|---|---|---|---|---|---|---|
| Q8IZF2 | ADGRF5 | TPDPISVsEGQNFSI | S483 | 0 | 87.23 | 1346 | 0 | 0 | 0 | 1 |
| Q8IZF2 | ADGRF5 | DYKVTFHtGSssLPA | T604 | 0.139 | 80.85 | 1346 | 0.094939 | 0.188 | 0.1284 | 1 |
| Q9Y678 | COPG1 | LrYAAVRtLNKVAMK | T310 | 0.1 | 96.04 | 874 | 0.068301 | 0.1 | 0.0683 | 1 |
| O00186 | STXBP3 | ASkSENKykAAYIYF | Y103 | 0.036 | 95.84 | 592 | 0.015793 | 0.036 | 0.0158 | 1 |
| P27348 | YWHAQ | RkQtIDNsQGAyQEA | S145 | 0 | 98.59 | 245 | 0 | 0 | 0 | 1 |
| P27348 | YWHAQ | IDNsQGAyQEAFDIs | Y149 | 0 | 98.85 | 245 | 0 | 0 | 0 | 0.9333 |
| P27348 | YWHAQ | yQEAFDIskkEMQPT | S156 | 0.001 | 98.33 | 245 | 0.000795 | 0.001 | 0.0008 | 1 |
| Q9P283 | SEMA5B | YPGARDFsQLALDPS | S127 | 0.122 | 95.42 | 1151 | 0.09698 | 0.122 | 0.097 | 1 |
| Q9P283 | SEMA5B | RKVFMCGtNAFSPMC | T201 | 0.004 | 97.09 | 1151 | 0.002732 | 0.004 | 0.0027 | 1 |
| Q9P283 | SEMA5B | ISSQGELyAATVIDF | Y248 | 0.023 | 98.09 | 1151 | 0.01009 | 0.023 | 0.0101 | 1 |
| Q8N7B9 | EFCAB3 | DSRNESFyDTFSTYT | Y372 | 0.259 | 83.98 | 438 | 0.11362 | 0.259 | 0.1136 | 1 |
| O75369 | FLNB | GVRIREttDFkVDTk | T473 | 0 | 81.73 | 2602 | 0 | 0 | 0 | 1 |
| O75369 | FLNB | DGVYAFEyYPstPGR | Y515 | 0.054 | 88.38 | 2602 | 0.023689 | 0.054 | 0.0237 | 1 |
| O75369 | FLNB | DDEDIKDsPyMAFIH | S630 | 0 | 88.2 | 2602 | 0 | 0 | 0 | 0.8571 |
| O75369 | FLNB | DGTyACsytPVKAIK | Y708 | 0.035 | 91.35 | 2602 | 0.015354 | 0.035 | 0.0154 | 1 |
| O75369 | FLNB | GGVNIPHsPyRVNIG | S730 | 0.076 | 89.16 | 2602 | 0.060414 | 0.076 | 0.0604 | 0.8333 |
| Q8TAT6 | NPLOC4 | HPNMCRLsPDGHFGS | S358 | 0 | 93.39 | 608 | 0 | 0.091 | 0.0723 | 1 |
| O15438 | ABCC3 | KALNPKRyQQTLEAC | Y720 | 0.128 | 93.95 | 1527 | 0.056152 | 0.128 | 0.0562 | 1 |
| Q13085 | ACACA | kLAKMVGyVSAGTVE | Y417 | 0.061 | 91.38 | 2346 | 0.02676 | 0.061 | 0.0268 | 1 |
| P02768 | ALB | VLIAFAQyLQQCPFE | Y54 | 0.161 | 97.62 | 609 | 0.070628 | 0.243 | 0.1066 | 1 |
| P02768 | ALB | VkLVNEVtEFAkTCV | T71 | 0.005 | 96.92 | 609 | 0.003415 | 0.005 | 0.0034 | 1 |
| P02768 | ALB | TFLkkyLyEIArRHP | Y164 | 0.079 | 98.45 | 609 | 0.034656 | 0.079 | 0.0347 | 0.8462 |
| P02768 | ALB | AWAVARLsQRFPkAE | S244 | 0 | 98.48 | 609 | 0 | 0 | 0 | 1 |
| P02768 | ALB | skLVtDLtkVHTECC | T263 | 0.005 | 98.25 | 609 | 0.003415 | 0.005 | 0.0034 | 1 |
| P02768 | ALB | CENQDsIsskLkECC | S296 | 0.046 | 97.21 | 609 | 0.036566 | 0.046 | 0.0366 | 1 |
| P02768 | ALB | EkPLLEKsHCIAEVE | S311 | 0.11 | 97.64 | 609 | 0.087441 | 0.11 | 0.0874 | 1 |
| P02768 | ALB | LLRLAKtyEttLEKC | Y377 | 0 | 97.96 | 609 | 0 | 0 | 0 | 1 |
| P02768 | ALB | LAKtyEttLEKCCAA | T380 | 0.078 | 97.5 | 609 | 0.053275 | 0.146 | 0.0997 | 1 |
| P02768 | ALB | AADPHECyAkVFDEF | Y394 | 0.02 | 96.34 | 609 | 0.008774 | 0.02 | 0.0088 | 1 |
| P02768 | ALB | NALLVRYtKkVPQVs | T436 | 0.01 | 97.56 | 609 | 0.00683 | 0.01 | 0.0068 | 1 |
| P02768 | ALB | tPtLVEVsRNLGkVG | S451 | 0.004 | 96.95 | 609 | 0.00318 | 0.004 | 0.0032 | 1 |

| | | | | | | | | | | |
|---|---|---|---|---|---|---|---|---|---|---|
| O00219 | HAS3 | SLGYRTKyTARSKCL | Y333 | 0.076 | 98.67 | 553 | 0.03334 | 0.251 | 0.1101 | 1 |
| Q9UNX3 | RPL26L1 | VrRkIMssPLskELR | S32 | 0.004 | 97.38 | 145 | 0.00318 | 0.004 | 0.0032 | 1 |
| Q9H7Z6 | KAT8 | QPPGkEIyRkSNISV | Y241 | 0.05 | 98.74 | 458 | 0.021934 | 0.05 | 0.0219 | 1 |
| P32754 | HPD | LQFHRFWsVDDTQVH | S211 | 0.175 | 89.78 | 393 | 0.13911 | 0.175 | 0.1391 | 1 |
| P32754 | HPD | TQVHTEyssLRSIVV | S222 | 0.067 | 77.63 | 393 | 0.053259 | 0.067 | 0.0533 | 1 |
| P32754 | HPD | QVHTEyssLRSIVVA | S223 | 0.045 | 80.75 | 393 | 0.035771 | 0.045 | 0.0358 | 1 |
| P32754 | HPD | LsVPSTyykQLREKL | Y296 | 0 | 95.21 | 393 | 0 | 0 | 0 | 0.9286 |
