## Supplementary Table 2 for "Mapping Cryptic Phosphorylation Sites in the Human Proteome"

### SUPP. TABLE 2

| PROT_ID | TAXID | MNEMONIC | BUSCO_SCORE | BUSCO_TOTAL | CPD_STATUS | SCI_NAME | COMM_NAME |
|---|---|---|---|---|---|---|---|
| UP000248482 | 391180 | ENHLU | 99 | 14502 | Standard | Enhydra lutris kenyoni | northern sea otter |
| UP000002356 | 9940 | SHEEP | 29 | 13335 | Close to standard (low value) | Ovis aries | Sheep |
| UP000694554 | 42100 | PHOSS | 5 | 13335 | Standard | Phocoena sinus | Vaquita |
| UP000079721 | 9365 | ERIEU | 66 | 12234 | Standard | Erinaceus europaeus | Western European hedgehog |
| UP000694387 | 9793 | EQUAS | 98 | 12234 | Close to standard (high value) | Equus asinus | Donkey |
| UP000000589 | 10090 | MOUSE | 99 | 13798 | Close to standard (high value) | Mus musculus | Mouse |
| UP000005640 | 9606 | HUMAN | 99 | 13780 | Outlier (high value) | Homo sapiens | Human |
| UP000515140 | 38626 | PHACI | 98 | 9226 | Close to standard (high value) | Phascolarctos cinereus | Koala |
| UP000189705 | 38654 | ALLSI | 98 | 7480 | Outlier (high value) | Alligator sinensis | Chinese alligator |
| UP000504639 | 219594 | AYTFU | 99 | 8338 | Outlier (high value) | Aythya fuligula | Tufted duck |
| UP000504619 | 103695 | 9SAUR | 98 | 7480 | Close to standard (high value) | Pogona vitticeps | central bearded dragon |
| UP000186698 | 8355 | XENLA | 98 | 5310 | Outlier (high value) | Xenopus laevis | African clawed frog |
| UP000192220 | 52670 | AUSLI | 96 | 15213 | Standard | Austrofundulus limnaeus | Annual killifish |
| UP000695023 | 303518 | 9CICH | 98 | 3640 | Close to standard (high value) | Pundamilia nyererei | |
| UP000000437 | 7955 | DANRE | 98 | 3640 | Outlier (high value) | Danio rerio | Zebrafish |
| UP000000803 | 7227 | DROME | 100 | 3285 | Close to standard (high value) | Drosophila melanogaster | Fruit fly |
| UP000594260 | 109461 | VARDE | 99 | 2934 | Close to standard (high value) | Varroa destructor | Honeybee mite |
| UP000694843 | 294128 | HYAAZ | 97 | 1013 | Standard | Hyalella azteca | Amphipod |
| UP000515163 | 6105 | ACTTE | 98 | 954 | Unknown | Actinia tenebrosa | Australian red waratah sea anemone |
| UP000001940 | 6239 | CAEEL | 100 | 3131 | Close to standard (high value) | Caenorhabditis elegans | |
| UP000014254 | 1220926 | MUCC1 | 100 | 2449 | Unknown | Mucor circinelloides f. circinelloides (strain 1006PhL) | Mucormycosis agent |
| UP000000819 | 284813 | ENCCU | 94 | 600 | Unknown | Encephalitozoon cuniculi (strain GB-M1) | Microsporidian parasite |
| UP000053201 | 645134 | SPIPD | 96 | 758 | Unknown | Spizellomyces punctatus (strain DAOM BR117) | |
| UP000006548 | 3702 | ARATH | 98 | 4596 | Standard | Arabidopsis thaliana | Mouse-ear cress |
| UP000006906 | 3055 | CHLRE | 99 | 1519 | Close to standard (high value) | Chlamydomonas reinhardtii | |
| UP000002226 | 432359 | TOXGV | 100 | 502 | Standard | Toxoplasma gondii (strain ATCC 50861 / VEG) | |
| UP000000600 | 5888 | PARTE | 99 | 171 | Outlier (high value) | Paramecium tetraurelia | |
| UP000000542 | 5664 | LEIMA | 100 | 130 | Standard | Leishmania major | |
| UP000006643 | 403677 | PHYIT | 100 | 100 | Standard | Phytophthora infestans (strain T30-4) | Potato late blight agent |
| UP000241769 | 1890364 | 9EUKA | 97 | 255 | Unknown | Planoprotostelium fungivorum | |
| UP000000559 | 237561 | CANAL | 99 | 2137 | Standard | Candida albicans (strain SC5314 / ATCC MYA-2876) | Yeast |
| UP000261680 | 29073 | URSMA | 99 | 14502 | Standard | Ursus maritimus | Polar bear |
| UP000694540 | 51154 | 9CETA | 96 | 13335 | Standard | Catagonus wagneri | Chacoan peccary |
| UP000694857 | 9771 | BALMU | 99 | 13335 | Close to standard (high value) | Balaenoptera musculus | Blue whale |
| UP000002281 | 9796 | HORSE | 97 | 12234 | Outlier (high value) | Equus caballus | Horse |
| UP000504640 | 9515 | SAPAP | 99 | 13780 | Close to standard (high value) | Sapajus apella | Brown-capped capuchin |
| UP000472275 | 223781 | AQUCH | 97 | 8338 | Outlier (high value) | Aquila chrysaetos chrysaetos | |
| UP000504624 | 321398 | 9PASS | 99 | 10844 | Outlier (high value) | Lepidothrix coronata | blue-crowned manakin |
| UP000008143 | 8364 | XENTR | 97 | 5310 | Close to standard (high value) | Xenopus tropicalis | Western clawed frog |
| UP000028760 | 48698 | POEFO | 95 | 15213 | Close to standard (low value) | Poecilia formosa | Amazon molly |

| | | | | | | | |
|---|---|---|---|---|---|---|---|
| UP000005207 | 8128 | ORENI | 98 | 3640 | Outlier (high value) | Oreochromis niloticus | Nile tilapia |
| UP000515145 | 210632 | 9TELE | 99 | 3640 | Close to standard (high value) | Parambassis ranga | Indian glassy fish |
| UP000887116 | 2740835 | TRICU | 97 | 2934 | Outlier (high value) | Trichonephila clavata | Joro spider |
| UP000000305 | 6669 | DAPPU | 96 | 1013 | Close to standard (high value) | Daphnia pulex | Water flea |
| UP000085678 | 7574 | LINUN | 99 | 954 | Close to standard (high value) | Lingula unguis | |
| UP000887567 | 2652724 | EXADI | 96 | 954 | Unknown | Exaiptasia diaphana | Tropical sea anemone |
| UP000006672 | 6279 | BRUMA | 99 | 3131 | Standard | Brugia malayi | Filarial nematode worm |
| UP000000715 | 9669 | MUSPF | 99 | 14502 | Close to standard (high value) | Mustela putorius furo | European domestic ferret |
| UP001055712 | 3077 | CHLVU | 99 | 1519 | Standard | Chlorella vulgaris | Green alga |
| UP000248481 | 29088 | NEOSC | 98 | 14502 | Standard | Neomonachus schauinslandi | Hawaiian monk seal |
| UP000291000 | 9925 | CAPHI | 97 | 13335 | Standard | Capra hircus | Goat |
| UP000245320 | 9739 | TURTR | 99 | 13335 | Close to standard (high value) | Tursiops truncatus | Atlantic bottle-nosed dolphin |
| UP000694399 | 9689 | PANLE | 96 | 14502 | Standard | Panthera leo | Lion |
| UP000000539 | 9031 | CHICK | 4 | 8338 | Outlier (high value) | Gallus gallus | Chicken |
| UP000694421 | 96440 | SALMN | 95 | 7480 | Standard | Salvator merianae | Argentine black and white tegu |
| UP000515156 | 1415580 | 9AMPH | 97 | 5310 | Close to standard (high value) | Microcaecilia unicolor | |
| UP000265160 | 106582 | 9CICH | 98 | 3640 | Close to standard (high value) | Maylandia zebra | zebra mbuna |
| UP000515150 | 158456 | BETSP | 99 | 3640 | Close to standard (high value) | Betta splendens | Siamese fighting fish |
| UP000494040 | 79782 | CIMLE | 99 | 2510 | Close to standard (high value) | Cimex lectularius | Bed bug |
| UP000790347 | 6954 | DERFA | 95 | 2934 | Standard | Dermatophagoides farinae | American house dust mite |
| UP000008827 | 3847 | SOYBN | 95 | 5366 | Outlier (high value) | Glycine max | Soybean |
| UP000694844 | 6565 | CRAVI | 98 | 5295 | Outlier (high value) | Crassostrea virginica | Eastern oyster |
| UP000694398 | 34839 | CHILA | 95 | 13798 | Standard | Chinchilla lanigera | Long-tailed chinchilla |
| UP000006591 | 4536 | ORYNI | 96 | 4896 | Standard | Oryza nivara | Indian wild rice |
| UP000504605 | 30538 | VICPA | 98 | 13335 | Standard | Vicugna pacos | Alpaca |
| UP000252040 | 1706337 | NEOAA | 98 | 13335 | Standard | Neophocaena asiaeorientalis asiaeorientalis | Yangtze finless porpoise |
| UP000002494 | 10116 | RAT | 98 | 13798 | Close to standard (high value) | Rattus norvegicus | Rat |
| UP000002277 | 9598 | PANTR | 98 | 13780 | Close to standard (high value) | Pan troglodytes | Chimpanzee |
| UP000694562 | 100819 | FALTI | 96 | 8338 | Outlier (high value) | Falco tinnunculus | Common kestrel |
| UP000504627 | 649802 | 9PASS | 98 | 10844 | Outlier (high value) | Pipra filicauda | Wire-tailed manakin |
| UP000515157 | 94885 | PANGU | 86 | 7480 | Close to standard (high value) | Pantherophis guttatus | Corn snake |
| UP000515159 | 260995 | GEOSA | 97 | 5310 | Close to standard (high value) | Geotrypetes seraphini | Gaboon caecilian |
| UP000515129 | 7957 | CARAU | 99 | 3640 | Outlier (high value) | Carassius auratus | Goldfish |
| UP000008820 | 7159 | AEDAE | 99 | 3285 | Close to standard (high value) | Aedes aegypti | Yellowfever mosquito |
| UP000515146 | 6956 | DERPT | 95 | 2934 | Close to standard (low value) | Dermatophagoides pteronyssinus | European house dust mite |
| UP000024404 | 6282 | ONCVO | 98 | 3131 | Close to standard (low value) | Onchocerca volvulus | |
| UP000286641 | 34884 | CALUR | 98 | 14502 | Close to standard (high value) | Callorhinus ursinus | Northern fur seal |
| UP000694544 | 68415 | MOSMO | 97 | 13335 | Standard | Moschus moschiferus | Siberian musk deer |
| UP000515203 | 10160 | OCTDE | 98 | 13798 | Standard | Octodon degus | Degu |
| UP000006718 | 9544 | MACMU | 97 | 13780 | Close to standard (high value) | Macaca mulatta | Rhesus macaque |
| UP000694380 | 8478 | CHRPI | 97 | 7480 | Outlier (high value) | Chrysemys picta bellii | Western painted turtle |
| UP000695026 | 176946 | PYTBI | 95 | 7480 | Standard | Python bivittatus | Burmese python |
| UP000002311 | 559292 | YEAST | 99 | 2137 | Close to standard (high value) | Saccharomyces cerevisiae (strain ATCC 204508 / S288c) | Baker's yeast |

| | | | | | | | |
|---|---|---|---|---|---|---|---|
| UP000472276 | 47969 | OREAU | 97 | 3640 | Outlier (high value) | Oreochromis aureus | Israeli tilapia |
| UP000504606 | 133901 | FRAOC | 99 | 1367 | Close to standard (high value) | Frankliniella occidentalis | Western flower thrips |
| UP000233120 | 9545 | MACNE | 96 | 13780 | Close to standard (high value) | Macaca nemestrina | Pig-tailed macaque |
| UP000749559 | 6347 | OWEFU | 97 | 954 | Standard | Owenia fusiformis | Polychaete worm |
| UP000286640 | 9627 | VULVU | 98 | 14502 | Standard | Vulpes vulpes | Red fox |
| UP000248483 | 9749 | DELLE | 99 | 13335 | Close to standard (high value) | Delphinapterus leucas | Beluga whale |
| UP000001595 | 9601 | PONAB | 97 | 13780 | Standard | Pongo abelii | Sumatran orangutan |
| UP000694412 | 93934 | COTJA | 97 | 8338 | Outlier (high value) | Coturnix japonica | Japanese quail |
| UP000694841 | 202946 | 9AVES | 98 | 8338 | Outlier (high value) | Apteryx mantelli mantelli | |
| UP000007305 | 4577 | MAIZE | 97 | 4896 | Close to standard (high value) | Zea mays | Maize |
| UP000011712 | 9685 | FELCA | 0 | 14502 | Standard | Felis catus | Cat |
| UP000694564 | 55149 | SCIVU | 96 | 13798 | Standard | Sciurus vulgaris | Eurasian red squirrel |
| UP000694411 | 9565 | THEGE | 97 | 13780 | Standard | Theropithecus gelada | Gelada baboon |
| UP000009136 | 9913 | BOVIN | 96 | 13335 | Close to standard (high value) | Bos taurus | Bovine |
| UP000694417 | 9999 | UROPR | 95 | 13798 | Standard | Urocitellus parryii | Arctic ground squirrel |
| UP000081671 | 10020 | DIPOR | 97 | 13798 | Standard | Dipodomys ordii | Ord's kangaroo rat |
| UP000233060 | 9531 | CERAT | 96 | 13780 | Close to standard (high value) | Cercocebus atys | Sooty mangabey |
| UP000504613 | 9880 | ODOVR | 98 | 13335 | Close to standard (high value) | Odocoileus virginianus texanus | |
| UP000008227 | 9823 | PIG | 97 | 13335 | Standard | Sus scrofa | Pig |
| UP000504631 | 207650 | 9HYME | 99 | 5991 | Close to standard (high value) | Bombus vosnesenskii | |
| UP000193067 | 1353009 | TRAC3 | 99 | 4464 | Unknown | Trametes coccinea (strain BRFM310) | |
| UP000092730 | 1296100 | 9TREE | 99 | 4284 | Outlier (high value) | Kwoniella bestiolae CBS 10118 | |
| UP000224006 | 94643 | 9APIC | 100 | 502 | Standard | Besnoitia besnoiti | |
| UP000001519 | 9595 | GORGO | 97 | 13780 | Standard | Gorilla gorilla gorilla | Western lowland gorilla |
| UP000001300 | 284591 | YARLI | 93 | 2137 | Unknown | Yarrowia lipolytica (strain CLIB 122 / E 150) | Yeast |
| UP000472241 | 61383 | LYNCA | 94 | 14502 | Standard | Lynx canadensis | Canada lynx |
| UP000314987 | 29139 | VOMUR | 93 | 9226 | Standard | Vombatus ursinus | Common wombat |
| UP000007648 | 9305 | SARHA | 92 | 9226 | Close to standard (high value) | Sarcophilus harrisii | Tasmanian devil |
| UP000001593 | 45351 | NEMVE | 94 | 954 | Unknown | Nematostella vectensis | Starlet sea anemone |
| UP000275408 | 46731 | POCDA | 93 | 954 | Unknown | Pocillopora damicornis | Cauliflower coral |
| UP001108240 | 630221 | CYPCA | 97 | 3640 | Outlier (high value) | Cyprinus carpio carpio | |
| UP000472260 | 1608454 | 9TELE | 97 | 3640 | Outlier (high value) | Sinocyclocheilus anshuiensis | |
| UP000694393 | 367368 | 9SAUR | 96 | 7480 | Outlier (high value) | Pelusios castaneus | West African mud turtle |
| UP000694390 | 1825980 | 9SAUR | 95 | 7480 | Outlier (high value) | Gopherus evgoodei | Goodes thornscrub tortoise |
| UP000472274 | 2587831 | TERCA | 95 | 7480 | Outlier (high value) | Terrapene carolina triunguis | Three-toed box turtle |
| UP000594220 | 8502 | CROPO | 94 | 7480 | Outlier (high value) | Crocodylus porosus | Saltwater crocodile |
| UP000694557 | 8019 | ONCKI | 98 | 3640 | Outlier (high value) | Oncorhynchus kisutch | Coho salmon |
| UP000472277 | 8032 | SALTR | 98 | 3640 | Outlier (high value) | Salmo trutta | Brown trout |
| UP000265140 | 8010 | ESOLU | 19 | 3640 | Outlier (high value) | Esox lucius | Northern pike |
| UP000318571 | 6832 | TIGCA | 94 | 1013 | Standard | Tigriopus californicus | Marine copepod |
| UP000007819 | 7029 | ACYPI | 99 | 2510 | Close to standard (high value) | Acyrthosiphon pisum | Pea aphid |
| UP000515158 | 161013 | THRPL | 97 | 1367 | Close to standard (high value) | Thrips palmi | Melon thrips |
| UP000301870 | 69820 | SPOLT | 99 | 5286 | Close to standard (high value) | Spodoptera litura | Asian cotton leafworm |

| | | | | | | | |
|---|---|---|---|---|---|---|---|
| UP000746747 | 2874296 | 9BILA | 96 | 3131 | Close to standard (low value) | Cercopithifilaria johnstoni | |
| UP000789390 | 27404 | 9CRUS | 93 | 1013 | Standard | Daphnia galeata | |
| UP000007110 | 7668 | STRPU | 99 | 954 | Unknown | Strongylocentrotus purpuratus | Purple sea urchin |
| UP000694845 | 133434 | ACAPL | 99 | 954 | Unknown | Acanthaster planci | Crown-of-thorns starfish |
| UP000887568 | 46514 | PATMI | 98 | 954 | Unknown | Patiria miniata | Bat star |
| UP000014500 | 126957 | STRMM | 93 | 1013 | Standard | Strigamia maritima | European centipede |
| UP000805418 | 9615 | CANLF | 96 | 14502 | Standard | Canis lupus familiaris | Dog |
| UP000009183 | 29760 | VITVI | 96 | 2326 | Close to standard (low value) | Vitis vinifera | Grape |
| UP000084051 | 4097 | TOBAC | 99 | 5950 | Outlier (high value) | Nicotiana tabacum | Common tobacco |
| UP000005204 | 7091 | BOMMO | 99 | 5286 | Close to standard (high value) | Bombyx mori | Silk moth |
| UP000005203 | 7460 | APIME | 99 | 5991 | Standard | Apis mellifera | Honeybee |
