## Supplementary Table 3 for "Mapping Cryptic Phosphorylation Sites in the Human Proteome"

| ROW_ID | GENE_SYMBOL | GENOMIC_MUTATION_ID | MUTATION_CDS | MUTATION_AA | MUTATION_DESCRIPTION | COUNT | SUB_ACCID | ONC_TSG |
|---|---|---|---|---|---|---|---|---|
| 1 | SLC12A8 | COSV58863520 | c.320A>C | p.Y107S | missense_variant | 1 | A0AV02 | |
| 2 | CD300LB | COSV58725218 | c.258G>A | p.T86= | synonymous_variant | 2 | A8K4G0 | |
| 3 | SGK1 | COSV105103895 | c.372T>G | p.Y124* | stop_gained | 6 | O00141 | oncogene |
| 4 | SGK1 | COSV105103614 | c.371A>C | p.Y124S | missense_variant | 5 | O00141 | oncogene |
| 5 | WWP2 | COSV63123024 | c.1892A>T | p.Y631F | missense_variant | 3 | O00308 | |
| 6 | DDX3X | COSV67864564 | c.1475C>G | p.S492* | stop_gained | 14 | O00571 | TSG |
| 7 | PPP6C | COSV65207327 | c.590C>T | p.S197L | missense_variant | 3 | O00743 | TSG |
| 8 | PTPRT | COSV62026099 | c.329C>G | p.S110C | missense_variant | 7 | O14522 | TSG |
| 9 | TERT | COSV57211055 | c.2036C>T | p.S679F | missense_variant | 3 | O14746 | oncogene, TSG |
| 10 | TERT | COSV99717329 | c.2035T>G | p.S679A | missense_variant | 3 | O14746 | oncogene, TSG |
| 11 | CHEK1 | COSV54026317 | c.1145C>G | p.T382S | missense_variant | 8 | O14757 | |
| 12 | INPP4B | COSV53752823 | c.1057A>G | p.S353G | missense_variant | 4 | O15327 | |
| 13 | XRCC3 | COSV57974380 | c.722C>T | p.T241M | missense_variant | 5 | O43542 | |
| 14 | SPOP | COSV61653556 | c.355A>G | p.S119G | ssense_variant,splice_region_vari | 4 | O43791 | TSG |
| 15 | SPOP | COSV61653897 | c.356G>A | p.S119N | missense_variant | 4 | O43791 | TSG |
| 16 | RAD21 | COSV52062390 | c.120_123del | p.S40Rfs*9 | frameshift_variant | 1 | O60216 | oncogene, TSG |
| 17 | JAK2 | COSV67677042 | c.2818dup | p.Y940Lfs*6 | frameshift_variant | 1 | O60674 | oncogene, fusion |
| 18 | JAK2 | COSV104701274 | c.1911T>C | p.Y637= | synonymous_variant | 1 | O60674 | oncogene, fusion |
| 19 | ALOX12B | COSV59871334 | c.28A>G | p.T10A | missense_variant | 1 | O75342 | |
| 20 | ALOX12B | COSV59872899 | c.53C>T | p.T18I | missense_variant | 1 | O75342 | |
| 21 | SF3B1 | COSV99067054 | c.3408del | p.Y1136* | frameshift_variant | 1 | O75533 | oncogene |
| 22 | IDH1 | COSV105912838 | c.691T>C | p.Y231H | missense_variant | 3 | O75874 | oncogene |
| 23 | IDH1 | COSV61616499 | c.623A>G | p.Y208C | missense_variant | 3 | O75874 | oncogene |
| 24 | IDH1 | COSV61617838 | c.55A>G | p.T19A | missense_variant | 3 | O75874 | oncogene |
| 25 | EGFR | COSV104368883 | c.339C>T | p.Y113= | synonymous_variant | 6 | P00533 | oncogene |
| 26 | SERPINA1 | COSV105916564 | c.782G>C | p.S261T | missense_variant | 10 | P01009 | |
| 27 | KRAS | COSV55570738 | c.266C>A | p.S89* | stop_gained | 3 | P01116 | oncogene |
| 28 | HLA-A | COSV65144279 | c.439T>G | p.Y147D | missense_variant | 5 | P04439 | fusion |
| 29 | HLA-A | COSV65139971 | c.440A>G | p.Y147C | missense_variant | 5 | P04439 | fusion |
| 30 | HLA-A | COSV65142314 | c.248A>G | p.Y83C | missense_variant | 5 | P04439 | fusion |
| 31 | HLA-A | COSV65145457 | c.247T>A | p.Y83N | missense_variant | 5 | P04439 | fusion |
| 32 | TP53 | COSV52661985 | c.463A>G | p.T155A | missense_variant | 15 | P04637 | oncogene, TSG, fusion |
| 33 | TP53 | COSV52775381 | c.660T>G | p.Y220* | stop_gained | 16 | P04637 | oncogene, TSG, fusion |
| 34 | TP53 | COSV52858547 | c.463_464del | p.T155Pfs*25 | frameshift_variant | 18 | P04637 | oncogene, TSG, fusion |
| 35 | TP53 | COSV52838787 | c.463_470del | p.T155Pfs*23 | frameshift_variant | 14 | P04637 | oncogene, TSG, fusion |
| 36 | TP53 | COSV52782333 | c.464C>G | p.T155S | missense_variant | 16 | P04637 | oncogene, TSG, fusion |
| 37 | TP53 | COSV53706115 | c.658_664del | p.Y220Rfs*25 | frameshift_variant | 15 | P04637 | oncogene, TSG, fusion |
| 38 | TP53 | COSV52969178 | c.465C>T | p.T155= | synonymous_variant | 14 | P04637 | oncogene, TSG, fusion |
| 39 | TP53 | COSV53050162 | c.465C>A | p.T155= | synonymous_variant | 16 | P04637 | oncogene, TSG, fusion |
| 40 | TP53 | COSV52871989 | c.463_476del | p.T155Hfs*21 | frameshift_variant | 16 | P04637 | oncogene, TSG, fusion |
| 41 | PRKCG | COSV54734426 | c.1561T>A | p.Y521N | missense_variant | 1 | P05129 | |
| 42 | INSR | COSV57162809 | c.2787C>T | p.S929= | synonymous_variant | 2 | P06213 | |
| 43 | FYN | COSV99991463 | c.557C>T | p.S186L | missense_variant | 6 | P06241 | |
| 44 | POLB | COSV107205381 | c.117C>A | p.Y39* | splice_region_variant,stop_gainec | 1 | P06746 | |
| 45 | HSP90AA1 | COSV53488088 | c.632C>T | p.S211F | missense_variant | 2 | P07900 | fusion |
| 46 | FH | COSV63818767 | c.203A>G | p.Y68C | missense_variant | 1 | P07954 | TSG |
| 47 | FH | COSV63818522 | c.328T>C | p.Y110H | missense_variant | 1 | P07954 | TSG |
| 48 | IGF1R | COSV105089218 | c.3834C>T | p.S1278= | synonymous_variant | 2 | P08069 | |

| | | | | | | | | |
|---|---|---|---|---|---|---|---|---|
| 49 | IGF1R | COSV106351687 | c.3833C>T | p.S1278F | missense_variant | 2 | P08069 | |
| 50 | PTPRC | COSV106061710 | c.2579C>G | p.T860R | missense_variant | 2 | P08575 | TSG |
| 51 | PARP1 | COSV64687449 | c.122C>T | p.S41L | ssense_variant,splice_region_vari | 4 | P09874 | |
| 52 | PARP1 | COSV64692750 | c.1436C>T | p.S479F | missense_variant | 1 | P09874 | |
| 53 | PTPRF | COSV63448717 | c.4214A>G | p.Y1405C | missense_variant | 2 | P10586 | |
| 54 | PRKAR1A | COSV62235529 | c.737_738del | p.Y246* | frameshift_variant | 5 | P10644 | oncogene, TSG, fusion |
| 55 | IRF1 | COSV55377752 | c.194dup | p.Y65* | frameshift_variant,stop_gained | 2 | P10914 | |
| 56 | CDH1 | COSV55738933 | c.208dup | p.S70Ffs*24 | frameshift_variant | 5 | P12830 | TSG |
| 57 | CDH1 | COSV55745796 | c.207_208insA | p.S70Ifs*24 | frameshift_variant | 5 | P12830 | TSG |
| 58 | CDH1 | COSV55734761 | c.209C>T | p.S70F | missense_variant | 5 | P12830 | TSG |
| 59 | CDH1 | COSV99932516 | c.208_220del | p.S70Dfs*9 | frameshift_variant | 5 | P12830 | TSG |
| 60 | XRCC6 | COSV63752864 | c.270C>T | p.T90= | synonymous_variant | 6 | P12956 | |
| 61 | CPM | COSV107435844 | c.372C>T | p.S124= | synonymous_variant | 3 | P14384 | |
| 62 | INSRR | COSV63875065 | c.3371C>T | p.S1124F | missense_variant | 1 | P14616 | |
| 63 | PLCG2 | COSV63869801 | c.2047A>G | p.T683A | missense_variant | 2 | P16885 | |
| 64 | PLCG2 | COSV63871984 | c.2036C>T | p.S679F | missense_variant | 2 | P16885 | |
| 65 | ITGA2 | COSV99670474 | c.3014A>T | p.Y1005F | missense_variant | 1 | P17301 | |
| 66 | FLT1 | COSV107287933 | c.1853C>T | p.T618I | missense_variant | 4 | P17948 | |
| 67 | LIG1 | COSV54389293 | c.2142C>A | p.S714= | synonymous_variant | 4 | P18858 | |
| 68 | PLCG1 | COSV54823575 | c.1137C>T | p.Y379= | synonymous_variant | 2 | P19174 | oncogene, fusion |
| 69 | TNFAIP3 | COSV52797875 | c.382del | p.S128Afs*88 | frameshift_variant | 7 | P21580 | TSG |
| 70 | JAK1 | COSV104650478 | c.651C>T | p.Y217= | synonymous_variant | 1 | P23458 | oncogene, TSG |
| 71 | DNMT1 | COSV61584945 | c.1646C>G | p.S549C | missense_variant | 2 | P26358 | |
| 72 | AKT1 | COSV104666470 | c.528C>T | p.Y176= | synonymous_variant | 6 | P31749 | oncogene |
| 73 | CTNNB1 | COSV105912830 | c.932G>A | p.S311N | missense_variant | 22 | P35222 | oncogene, fusion |
| 74 | CTNNB1 | COSV62714200 | c.1415C>G | p.T472S | missense_variant | 23 | P35222 | oncogene, fusion |
| 75 | IRS1 | COSV59337520 | c.140A>G | p.Y47C | missense_variant | 1 | P35568 | |
| 76 | SOX11 | COSV58983416 | c.179C>T | p.S60F | missense_variant | 1 | P35716 | |
| 77 | VHL | COSV56544742 | c.214T>C | p.S72P | missense_variant | 2 | P40337 | TSG |
| 78 | VHL | COSV56542656 | c.333C>G | p.S111R | missense_variant | 2 | P40337 | TSG |
| 79 | VHL | COSV56547849 | c.331A>C | p.S111R | missense_variant | 2 | P40337 | TSG |
| 80 | VHL | COSV56544837 | c.333C>A | p.S111R | missense_variant | 2 | P40337 | TSG |
| 81 | VHL | COSV56570422 | c.238_244del | p.S80Afs*77 | frameshift_variant | 2 | P40337 | TSG |
| 82 | VHL | COSV56551476 | c.331A>G | p.S111G | missense_variant | 2 | P40337 | TSG |
| 83 | VHL | COSV56547599 | c.238A>C | p.S80R | missense_variant | 2 | P40337 | TSG |
| 84 | VHL | COSV56542846 | c.240T>A | p.S80R | missense_variant | 2 | P40337 | TSG |
| 85 | VHL | COSV56543658 | c.240T>G | p.S80R | missense_variant | 2 | P40337 | TSG |
| 86 | VHL | COSV56556318 | c.220_239dup | p.S80Rfs*45 | frameshift_variant | 2 | P40337 | TSG |
| 87 | VHL | COSV56565170 | c.328_331dup | p.S111Tfs*23 | frameshift_variant | 2 | P40337 | TSG |
| 88 | VHL | COSV56545944 | c.524del | p.Y175Sfs*27 | frameshift_variant | 2 | P40337 | TSG |
| 89 | VHL | COSV56561202 | c.330_331dup | p.S111Tfs*8 | frameshift_variant | 2 | P40337 | TSG |
| 90 | VHL | COSV56543942 | c.326_329dup | p.S111Pfs*22 | frameshift_variant | 2 | P40337 | TSG |
| 91 | VHL | COSV56562607 | c.238A>G | p.S80G | missense_variant | 2 | P40337 | TSG |
| 92 | VHL | COSV56547241 | c.239G>A | p.S80N | missense_variant | 2 | P40337 | TSG |
| 93 | VHL | COSV56545566 | c.215_216delinsA | p.S72Yfs*46 | frameshift_variant | 2 | P40337 | TSG |
| 94 | VHL | COSV56555146 | c.237del | p.S80Vfs*38 | frameshift_variant | 2 | P40337 | TSG |
| 95 | VHL | COSV56573078 | c.238_256del | p.S80Pfs*73 | frameshift_variant | 2 | P40337 | TSG |
| 96 | VHL | COSV56559539 | c.330_331delinsT | p.S111Afs*7 | frameshift_variant | 2 | P40337 | TSG |
| 97 | VHL | COSV56553653 | c.331_333delinsCCT | p.S111P | missense_variant | 2 | P40337 | TSG |

| | | | | | | | | |
|---|---|---|---|---|---|---|---|---|
| 98 | VHL | COSV56553885 | c.330_333del | p.S111Tfs*47 | frameshift_variant | 2 | P40337 | TSG |
| 99 | VHL | COSV56572126 | c.523T>G | p.Y175D | missense_variant | 2 | P40337 | TSG |
| 100 | VHL | COSV56563065 | c.214dup | p.S72Ffs*60 | frameshift_variant | 2 | P40337 | TSG |
| 101 | VHL | COSV56543581 | c.213del | p.S72Pfs*46 | frameshift_variant | 2 | P40337 | TSG |
| 102 | VHL | COSV56570092 | c.331_340del | p.S111Cfs*4 | ıeshift_variant,splice_region_vari | 2 | P40337 | TSG |
| 103 | VHL | COSV56542714 | c.332G>A | p.S111N | missense_variant | 2 | P40337 | TSG |
| 104 | VHL | COSV56572109 | c.212_237dup | p.S80Pfs*47 | frameshift_variant | 2 | P40337 | TSG |
| 105 | VHL | COSV56544133 | c.525C>A | p.Y175* | stop_gained | 2 | P40337 | TSG |
| 106 | VHL | COSV56543727 | c.525C>G | p.Y175* | stop_gained | 2 | P40337 | TSG |
| 107 | VHL | COSV56552913 | c.238_239delinsTT | p.S80F | missense_variant | 2 | P40337 | TSG |
| 108 | VHL | COSV56553040 | c.333C>T | p.S111= | synonymous_variant | 2 | P40337 | TSG |
| 109 | VHL | COSV56543921 | c.332G>T | p.S111I | missense_variant | 2 | P40337 | TSG |
| 110 | VHL | COSV56558946 | c.331del | p.S111Afs*48 | frameshift_variant | 2 | P40337 | TSG |
| 111 | MTOR | COSV63875903 | c.2638A>G | p.T880A | missense_variant | 1 | P42345 | oncogene |
| 112 | MSH2 | COSV51878184 | c.2228C>T | p.S743L | missense_variant | 4 | P43246 | TSG |
| 113 | NOTCH1 | COSV99072385 | c.5788A>G | p.T1930A | missense_variant | 1 | P46531 | oncogene, TSG, fusion |
| 114 | NOTCH1 | COSV105113181 | c.5790C>T | p.T1930= | synonymous_variant | 1 | P46531 | oncogene, TSG, fusion |
| 115 | PIP4K2A | COSV105307937 | c.279C>T | p.Y93= | synonymous_variant | 1 | P48426 | |
| 116 | MRE11 | COSV60580154 | c.1146C>T | p.S382= | synonymous_variant | 4 | P49959 | |
| 117 | JAK3 | COSV71686142 | c.713A>G | p.Y238C | missense_variant | 3 | P52333 | oncogene |
| 118 | JAK3 | COSV105359963 | c.714C>A | p.Y238* | stop_gained | 3 | P52333 | oncogene |
| 119 | JAK3 | COSV105940680 | c.1192T>A | p.S398T | missense_variant | 3 | P52333 | oncogene |
| 120 | JAK3 | COSV71688160 | c.1193C>G | p.S398C | missense_variant | 3 | P52333 | oncogene |
| 121 | BLM | COSV61925297 | c.2489C>T | p.T830M | missense_variant | 2 | P54132 | TSG |
| 122 | EPHB1 | COSV105333076 | c.275C>T | p.T92I | missense_variant | 1 | P54762 | |
| 123 | CDH8 | COSV54875451 | c.1599C>A | p.Y533* | stop_gained | 3 | P55286 | |
| 124 | CDH8 | COSV54863386 | c.1598A>G | p.Y533C | missense_variant | 3 | P55286 | |
| 125 | PTEN | COSV64297907 | c.532del | p.Y178Ifs*5 | frameshift_variant | 1 | P60484 | TSG |
| 126 | PTEN | COSV64290082 | c.463T>C | p.Y155H | missense_variant | 1 | P60484 | TSG |
| 127 | PTEN | COSV64302106 | c.534T>A | p.Y178* | stop_gained | 1 | P60484 | TSG |
| 128 | PTEN | COSV64309475 | c.534T>G | p.Y178* | stop_gained | 1 | P60484 | TSG |
| 129 | PTEN | COSV64297713 | c.204C>G | p.Y68* | stop_gained | 1 | P60484 | TSG |
| 130 | PTEN | COSV64292424 | c.532dup | p.Y178Lfs*2 | frameshift_variant | 1 | P60484 | TSG |
| 131 | PTEN | COSV64294248 | c.680C>T | p.S227F | missense_variant | 1 | P60484 | TSG |
| 132 | PTEN | COSV64291974 | c.411del | p.Y138Ifs*9 | frameshift_variant | 1 | P60484 | TSG |
| 133 | PTEN | COSV64292590 | c.202T>A | p.Y68N | missense_variant | 1 | P60484 | TSG |
| 134 | PTEN | COSV64297519 | c.412T>G | p.Y138D | missense_variant | 1 | P60484 | TSG |
| 135 | PTEN | COSV100910380 | c.532T>A | p.Y178N | missense_variant | 1 | P60484 | TSG |
| 136 | PTEN | COSV64291751 | c.464A>G | p.Y155C | missense_variant | 1 | P60484 | TSG |
| 137 | PTEN | COSV64290045 | c.720C>A | p.Y240* | stop_gained | 1 | P60484 | TSG |
| 138 | PTEN | COSV64293993 | c.203A>G | p.Y68C | missense_variant | 1 | P60484 | TSG |
| 139 | PTEN | COSV64306562 | c.413A>G | p.Y138C | missense_variant | 1 | P60484 | TSG |
| 140 | PTEN | COSV99057849 | c.413_420del | p.Y138Sfs*39 | frameshift_variant | 1 | P60484 | TSG |
| 141 | PTEN | COSV64292375 | c.464A>C | p.Y155S | missense_variant | 1 | P60484 | TSG |
| 142 | PTEN | COSV64291335 | c.718_719insCCTT | p.Y240Sfs*4 | frameshift_variant | 1 | P60484 | TSG |
| 143 | PTEN | COSV64304856 | c.532T>G | p.Y178D | missense_variant | 1 | P60484 | TSG |
| 144 | PTEN | COSV64307317 | c.463T>A | p.Y155N | missense_variant | 1 | P60484 | TSG |
| 145 | PTEN | COSV64304988 | c.532T>C | p.Y178H | missense_variant | 1 | P60484 | TSG |
| 146 | PTEN | COSV64293233 | c.531_532dup | p.Y178Ffs*6 | frameshift_variant | 1 | P60484 | TSG |

| | | | | | | | | |
|---|---|---|---|---|---|---|---|---|
| 147 | PTEN | COSV107457816 | c.414T>G | p.Y138* | stop_gained | 1 | P60484 | TSG |
| 148 | PTEN | COSV64306097 | c.414T>A | p.Y138* | stop_gained | 1 | P60484 | TSG |
| 149 | PTEN | COSV64289561 | c.202T>C | p.Y68H | missense_variant | 1 | P60484 | TSG |
| 150 | PTEN | COSV64297752 | c.678del | p.S227Pfs*29 | frameshift_variant | 1 | P60484 | TSG |
| 151 | PTEN | COSV64301130 | c.679dup | p.S227Ffs*16 | frameshift_variant | 1 | P60484 | TSG |
| 152 | PTEN | COSV64303044 | c.202_203del | p.Y68Qfs*5 | frameshift_variant | 1 | P60484 | TSG |
| 153 | ACTB | COSV104405261 | c.564C>A | p.Y188* | stop_gained | 3 | P60709 | |
| 154 | RHOA | COSV69041959 | c.479C>T | p.S160L | missense_variant | 3 | P61586 | oncogene, TSG |
| 155 | MAP2K1 | COSV61072444 | c.636C>T | p.S212= | synonymous_variant | 2 | Q02750 | oncogene |
| 156 | MAP2K1 | COSV61069356 | c.635G>A | p.S212N | missense_variant | 2 | Q02750 | oncogene |
| 157 | MAP2K1 | COSV61070930 | c.692C>T | p.S231L | ssense_variant,splice_region_vari | 2 | Q02750 | oncogene |
| 158 | AHNAK | COSV57213871 | c.122C>T | p.S41F | missense_variant | 3 | Q09666 | |
| 159 | ATM | COSV104377472 | c.115A>G | p.T39A | missense_variant | 7 | Q13315 | TSG |
| 160 | PTPRS | COSV53627207 | c.2408C>T | p.T803I | missense_variant | 5 | Q13332 | |
| 161 | PTPRS | COSV53641670 | c.4451C>T | p.S1484L | missense_variant | 4 | Q13332 | |
| 162 | ATR | COSV106502669 | c.870_871del | p.Y291* | frameshift_variant | 1 | Q13535 | TSG |
| 163 | CUL3 | COSV52372604 | c.1160A>T | p.Y387F | missense_variant | 4 | Q13618 | TSG |
| 164 | CBFB | COSV51998442 | c.255_267del | p.Y85* | frameshift_variant | 3 | Q13951 | TSG, fusion |
| 165 | KCNJ12 | COSV59163708 | c.1061C>T | p.T354M | missense_variant | 2 | Q14500 | |
| 166 | FAT1 | COSV71671686 | c.3528C>G | p.Y1176* | stop_gained | 2 | Q14517 | TSG |
| 167 | FAT1 | COSV71671751 | c.2379C>T | p.T793= | synonymous_variant | 2 | Q14517 | TSG |
| 168 | IRF3 | COSV59192429 | c.1215C>T | p.S405= | synonymous_variant | 9 | Q14653 | |
| 169 | TMEM132B | COSV54750164 | c.1907C>T | p.S636L | missense_variant | 2 | Q14DG7 | |
| 170 | ERBB4 | COSV53535326 | c.470A>G | p.Y157C | missense_variant | 3 | Q15303 | oncogene, TSG |
| 171 | STK11 | COSV58827761 | c.719C>A | p.S240* | stop_gained | 2 | Q15831 | TSG |
| 172 | STK11 | COSV58830737 | c.719C>T | p.S240L | missense_variant | 2 | Q15831 | TSG |
| 173 | MAP3K21 | COSV64046257 | c.425C>T | p.T142I | missense_variant | 2 | Q5TCX8 | |
| 174 | RIF1 | COSV105843215 | c.428C>T | p.S143L | missense_variant | 4 | Q5UIP0 | |
| 175 | TET2 | COSV106092004 | c.4035T>A | p.Y1345* | stop_gained | 3 | Q6N021 | TSG |
| 176 | TET2 | COSV54435975 | c.4035T>G | p.Y1345* | stop_gained | 3 | Q6N021 | TSG |
| 177 | TET2 | COSV54398617 | c.4034A>C | p.Y1345S | missense_variant | 3 | Q6N021 | TSG |
| 178 | PAXIP1 | COSV68186146 | c.2756C>A | p.S919Y | missense_variant | 2 | Q6ZW49 | |
| 179 | CATSPERD | COSV58465122 | c.950A>G | p.Y317C | missense_variant | 1 | Q86XM0 | |
| 180 | KLB | COSV57301509 | c.2639A>C | p.Y880S | missense_variant | 1 | Q86Z14 | |
| 181 | NUP93 | COSV57429064 | c.1940C>T | p.S647L | missense_variant | 4 | Q8N1F7 | |
| 182 | ANKK1 | COSV100392625 | c.2081C>A | p.T694K | missense_variant | 1 | Q8NFD2 | |
| 183 | CERK | COSV99348566 | c.1071C>G | p.S357R | missense_variant | 1 | Q8TCT0 | |
| 184 | FAT3 | COSV104590297 | c.1167C>A | p.S389= | synonymous_variant | 2 | Q8TDW7 | |
| 185 | USP13 | COSV55862159 | c.1162A>G | p.T388A | ssense_variant,splice_region_vari | 2 | Q92995 | |
| 186 | FBXW7 | COSV55967533 | c.2063G>T | p.S688I | missense_variant | 6 | Q969H0 | TSG |
| 187 | MOGAT1 | COSV71479842 | c.487T>C | p.S163P | missense_variant | 1 | Q96PD6 | |
| 188 | HMCN1 | COSV54883450 | c.1938C>T | p.T646= | synonymous_variant | 1 | Q96RW7 | |
| 189 | PARK7 | COSV100096631 | c.461C>T | p.T154I | missense_variant | 5 | Q99497 | |
| 190 | SYT3 | COSV58899619 | c.1421C>T | p.S474F | missense_variant | 3 | Q9BQG1 | |
| 191 | MKNK1 | COSV57862830 | c.203G>T | p.S68I | missense_variant | 5 | Q9BUB5 | |
| 192 | TSSK1B | COSV101180936 | c.591C>T | p.Y197= | synonymous_variant | 1 | Q9BXA7 | |
| 193 | CARD11 | COSV99052450 | c.2251T>C | p.Y751H | missense_variant | 1 | Q9BXL7 | oncogene |
| 194 | SHARPIN | COSV59643602 | c.845G>C | p.S282T | missense_variant | 1 | Q9H0F6 | |
| 195 | SLITRK6 | COSV68389758 | c.1217C>T | p.S406L | missense_variant | 3 | Q9H5Y7 | |

| | | | | | | | | |
|---|---|---|---|---|---|---|---|---|
| 196 | FANCE | COSV100003273 | c.1466A>T | p.Y489F | missense_variant | 1 | Q9HB96 | TSG |
| 197 | ARHGAP35 | COSV105936964 | c.923A>T | p.Y308F | missense_variant | 2 | Q9NRY4 | TSG |
| 198 | CDK12 | COSV71001656 | c.2860A>T | p.S954C | missense_variant | 2 | Q9NYV4 | TSG |
| 199 | NTM | COSV66178941 | c.597C>T | p.Y199= | synonymous_variant | 6 | Q9P121 | |
| 200 | RPS6KB2 | COSV57048174 | c.407A>G | p.Y136C | missense_variant | 1 | Q9UBS0 | |
| 201 | AASS | COSV62789805 | c.789G>T | p.T263= | synonymous_variant | 2 | Q9UDR5 | |
| 202 | CDC23 | COSV67508291 | c.362C>T | p.S121F | missense_variant | 3 | Q9UJX2 | |
| 203 | MTO1 | COSV64773367 | c.1464C>A | p.Y488* | stop_gained | 4 | Q9Y2Z2 | |
| 204 | DNMT3A | COSV53068654 | c.1913C>G | p.S638C | missense_variant | 4 | Q9Y6K1 | TSG |
