## Supplementary Table 4 for "Mapping Cryptic Phosphorylation Sites in the Human Proteome"

| ROW_ID | GENE | UPID | FTID | WT_AA | VAR_AA | VAR_TYPE | DISEASE(s) | MUT_SOURCE | MOD_RSD | MOD_TYPE | VAR_CLASS | ONC_TSG |
|---|---|---|---|---|---|---|---|---|---|---|---|---|
| 1 | A2ML1 | A8K2U0 | VAR_071856 | T | M | LB/B | - | Uniprot Humsavar | 1312 | Phosphorylation | I | |
| 2 | ABHD12 | Q8N2K0 | VAR_081591 | T | R | LP/P | Polyneuropathy, hearing loss, ataxia, retinitis pigmentosa, and cataract (PHARC) [MIM:612674] | Uniprot Humsavar | 253 | Phosphorylation | I | |
| 3 | ACTG1 | P63261 | VAR_067816 | S | F | LP/P | Baraitser-Winter syndrome 2 (BRWS2) [MIM:614583] | Uniprot Humsavar | 155 | Phosphorylation | I | |
| 4 | ACTG1 | P63261 | VAR_032434 | T | I | LP/P | Deafness, autosomal dominant, 20 (DFNA20) [MIM:604717] | Uniprot Humsavar | 89 | Phosphorylation | I | |
| 5 | ADGRF5 | Q8IZF2 | VAR_025326 | T | M | LB/B | - | Uniprot Humsavar | 604 | Phosphorylation | I | |
| 6 | AGO2 | Q9UKV8 | VAR_085427 | S | R | LP/P | Lessel-Kreienkamp syndrome (LESKRES) [MIM:619149] | Uniprot Humsavar | 760 | Phosphorylation | I | |
| 7 | ANOS1 | P23352 | VAR_069207 | Y | D | US | - | Uniprot Humsavar | 217 | Phosphorylation | I | |
| 8 | AR | P10275 | VAR_009824 | S | P | US | Prostate cancer | Uniprot Humsavar | 792 | Phosphorylation | I | oncogene |
| 9 | ATP2A2 | P16615 | VAR_008622 | S | F | LP/P | Darier disease (DD) [MIM:124200] | Uniprot Humsavar | 495 | Phosphorylation | I | |
| 10 | ATP7B | P35670 | VAR_000778 | T | M | LP/P | Wilson disease (WD) [MIM:277900] | Uniprot Humsavar | 1220 | Phosphorylation | I | |
| 11 | BBS12 | Q6ZW61 | VAR_062965 | T | M | LP/P | Bardet-Biedl syndrome 12 (BBS12) [MIM:615989] | Uniprot Humsavar | 501 | Phosphorylation | I | |
| 12 | BCS1L | Q9Y276 | VAR_018162 | S | N | LP/P | Mitochondrial complex III deficiency, nuclear type 1 (MC3DN1) [MIM:124000] | Uniprot Humsavar | 277 | Phosphorylation | I | |
| 13 | CAD | P27708 | VAR_035898; Co74[COSMIC] | Y | C | US | A colorectal cancer sample; colorectal_cancer[COSMIC] | Uniprot Humsavar; COSMIC | 735 | Phosphorylation | I | |
| 14 | CAMK2A | Q9UQM7 | VAR_080587 | T | P | LP/P | Intellectual developmental disorder, autosomal dominant 53 (MRD53) [MIM:617798] | Uniprot Humsavar | 286 | Phosphorylation | I | |
| 15 | CAPN15 | O75808 | VAR_085922 | S | F | US | Oculogastrointestinal neurodevelopmental syndrome (OGIN) [MIM:619318] | Uniprot Humsavar | 720 | Phosphorylation | I | |
| 16 | CBS | P35520 | VAR_008068 | T | M | LP/P | Cystathionine beta-synthase deficiency (CBSD) [MIM:236200] | Uniprot Humsavar | 191 | Phosphorylation | I | |
| 17 | CPS1 | P31327 | VAR_030676 | S | P | LP/P | Carbamoyl phosphate synthetase 1 deficiency (CPS1D) [MIM:237300] | Uniprot Humsavar | 918 | Phosphorylation | I | |
| 18 | CPS1 | P31327 | VAR_066168 | S | P | LP/P | Carbamoyl phosphate synthetase 1 deficiency (CPS1D) [MIM:237300] | Uniprot Humsavar | 1331 | Phosphorylation | I | |
| 19 | CPS1 | P31327 | VAR_064069 | Y | H | LP/P | Carbamoyl phosphate synthetase 1 deficiency (CPS1D) [MIM:237300] | Uniprot Humsavar | 1491 | Phosphorylation | I | |
| 20 | CPS1 | P31327 | VAR_006835 | T | M | LP/P | Carbamoyl phosphate synthetase 1 deficiency (CPS1D) [MIM:237300] | Uniprot Humsavar | 544 | Phosphorylation | I | |
| 21 | CYBB | P04839 | VAR_025613 | Y | D | LP/P | Granulomatous disease, chronic, X-linked (CGDX) [MIM:306400] | Uniprot Humsavar | 41 | Phosphorylation | I | |
| 22 | DHCR7 | Q9UBM7 | VAR_023176 | S | L | LP/P | Smith-Lemli-Opitz syndrome (SLOS) [MIM:270400] | Uniprot Humsavar | 397 | Phosphorylation | I | |
| 23 | DHCR7 | Q9UBM7 | TCGA-10-0930-01A | T | I | Disease | ovarian_cancer[TCGA] | TCGA | 135 | Phosphorylation | I | |
| 24 | DSP | P15924 | VAR_015402 | S | R | US | Arrhythmogenic right ventricular dysplasia, familial, 8 (ARVD8) [MIM:607450] | Uniprot Humsavar | 299 | Phosphorylation | I | |
| 25 | EIF2B5 | Q13144 | VAR_068469 | S | L | LP/P | Leukoencephalopathy with vanishing white matter 5 (VWM5) [MIM:620315] | Uniprot Humsavar | 447 | Phosphorylation | I | |
| 26 | F5 | P12259 | TCGA-13-0760-01A | Y | C | Disease | ovarian_cancer[TCGA] | TCGA | 2149 | Phosphorylation | I | |
| 27 | F8 | P00451 | VAR_001099 | Y | N | LP/P | Hemophilia A (HEMA) [MIM:306700] | Uniprot Humsavar | 450 | Phosphorylation | I | |
| 28 | F8 | P00451 | VAR_028533 | Y | D | LP/P | Hemophilia A (HEMA) [MIM:306700] | Uniprot Humsavar | 450 | Phosphorylation | I | |
| 29 | FBXO8 | Q9NRD0 | VAR_071119 | Y | H | LB/B | - | Uniprot Humsavar | 156 | Phosphorylation | I | |
| 30 | FCSK | Q8N0W3 | VAR_081646 | S | P | US | Congenital disorder of glycosylation with defective fucosylation 2 (CDGF2) [MIM:618324] | Uniprot Humsavar | 223 | Phosphorylation | I | |
| 31 | FECH | P22830 | VAR_030556 | S | P | LP/P | Protoporphyria, erythropoietic, 1 (EPP1) [MIM:177000] | Uniprot Humsavar | 151 | Phosphorylation | I | |
| 32 | FGF23 | Q9GZV9 | VAR_023831 | S | G | LP/P | Tumoral calcinosis, hyperphosphatemic, familial, 2 (HFTC2) [MIM:617993] | Uniprot Humsavar | 71 | Phosphorylation | I | |
| 33 | FMNL2 | Q96PY5 | VAR_032570 | Y | C | LB/B | - | Uniprot Humsavar | 352 | Phosphorylation | I | |
| 34 | G6PD | P11413 | VAR_020535 | Y | H | LB/B | - | Uniprot Humsavar | 322 | Phosphorylation | I | |
| 35 | GALC | P54803 | VAR_013957 | S | F | LP/P | Krabbe disease (KRB) [MIM:245200] | Uniprot Humsavar | 68 | Phosphorylation | I | |
| 36 | GAN | Q9H2C0 | VAR_010760 | S | G | LP/P | Giant axonal neuropathy 1, autosomal recessive (GAN1) [MIM:256850] | Uniprot Humsavar | 52 | Phosphorylation | I | |
| 37 | GCK | P35557 | VAR_079468 | S | L | LP/P | Maturity-onset diabetes of the young 2 (MODY2) [MIM:125851] | Uniprot Humsavar | 383 | Phosphorylation | I | |
| 38 | GHR | P10912 | VAR_018434 | Y | C | LP/P | Laron syndrome (LARS) [MIM:262500] | Uniprot Humsavar | 226 | Phosphorylation | I | |
| 39 | GLDC | P23378 | VAR_079313 | Y | C | LP/P | Non-ketotic hyperglycinemia (NKH) [MIM:605899] | Uniprot Humsavar | 839 | Phosphorylation | I | |
| 40 | GNA12 | Q03113 | VAR_071044 | S | G | LB/B | - | Uniprot Humsavar | 68 | Phosphorylation | I | |
| 41 | GPI | P06744 | VAR_002522 | T | I | LP/P | Hemolytic anemia, non-spherocytic, due to glucose phosphate isomerase deficiency (HA-GPID) [MIM:613470] | Uniprot Humsavar | 195 | Phosphorylation | I | |
| 42 | GRM7 | Q14831 | VAR_084625 | T | K | US | Neurodevelopmental disorder with seizures, hypotonia, and brain imaging abnormalities (NEDSHBA) [MIM:618922] | Uniprot Humsavar | 675 | Phosphorylation | I | |
| 43 | HAAO | P46952 | TCGA-13-1498-01A | S | F | Disease | ovarian_cancer[TCGA] | TCGA | 270 | Phosphorylation | I | |
| 44 | HBB | P68871 | VAR_002995 | S | N | LB/B | - | Uniprot Humsavar | 90 | Phosphorylation | I | |
| 45 | HBB | P68871 | VAR_002996 | S | R | LB/B | - | Uniprot Humsavar | 90 | Phosphorylation | I | |
| 46 | HBB | P68871 | VAR_003063 | Y | D | LB/B | - | Uniprot Humsavar | 131 | Phosphorylation | I | |
| 47 | HLA-DPB1 | P04440 | VAR_060642 | Y | D | LB/B | - | Uniprot Humsavar | 57 | Phosphorylation | I | |
| 48 | HMBS | P08397 | VAR_003670 | T | I | LP/P | Acute intermittent porphyria (AIP) [MIM:176000] | Uniprot Humsavar | 269 | Phosphorylation | I | |
| 49 | HOXD13 | P35453 | VAR_075400 | T | R | LP/P | Synpolydactyly 1 (SPD1) [MIM:186000] | Uniprot Humsavar | 313 | Phosphorylation | I | oncogene, fusion |
| 50 | IDS | P22304 | VAR_007351 | Y | D | LP/P | Mucopolysaccharidosis 2 (MPS2) [MIM:309900] | Uniprot Humsavar | 225 | Phosphorylation | I | |
| 51 | INSR | P06213 | VAR_079542 | Y | C | LP/P | Leprechaunism (LEPRCH) [MIM:246200] | Uniprot Humsavar | 818 | Phosphorylation | I | |
| 52 | IRS1 | P35568 | Br02X | Y | C | Disease | glioblastoma_jhu[COSMIC] | COSMIC | 47 | Phosphorylation | I | |
| 53 | KCNB1 | Q14721 | VAR_071992 | T | I | LP/P | Developmental and epileptic encephalopathy 26 (DEE26) [MIM:616056] | Uniprot Humsavar | 374 | Phosphorylation | I | |
| 54 | KLB | Q86Z14 | VAR_049296 | Y | H | LB/B | - | Uniprot Humsavar | 906 | Phosphorylation | I | |
| 55 | KPNB1 | Q14974 | Pa28C | S | L | Disease | pancreatic_cancer[COSMIC] | COSMIC | 853 | Phosphorylation | I | |
| 56 | LZTR1 | Q8N653 | VAR_075657 | Y | C | LP/P | Noonan syndrome 10 (NS10) [MIM:616564] | Uniprot Humsavar | 119 | Phosphorylation | I | TSG |
| 57 | MARK2 | Q7KZI7 | TCGA-24-1427-01A | S | N | Disease | ovarian_cancer[TCGA] | TCGA | 197 | Phosphorylation | I | |
| 58 | MARS1 | P56192 | VAR_075361 | Y | C | LP/P | Interstitial lung and liver disease (ILLD) [MIM:615486] | Uniprot Humsavar | 344 | Phosphorylation | I | |
| 59 | MATN3 | O15232 | VAR_019882 | T | M | LP/P | Multiple epiphyseal dysplasia 5 (EDM5) [MIM:607078] | Uniprot Humsavar | 120 | Phosphorylation | I | |

| | | | | | | | | | | | | |
|---|---|---|---|---|---|---|---|---|---|---|---|---|
| 60 | MITF | O75030 | VAR_010300 | S | P | LP/P | Waardenburg syndrome 2A (WS2A) [MIM:193510] | Uniprot Humsavar | 357 | Phosphorylation | I | oncogene |
| 61 | MME | P08473 | VAR_077690 | Y | H | LB/B | - | Uniprot Humsavar | 497 | Phosphorylation | I | |
| 62 | MOGAT1 | Q96PD6 | VAR_027389 | S | P | LB/B | - | Uniprot Humsavar | 163 | Phosphorylation | I | |
| 63 | MPI | P34949 | VAR_012338 | S | L | LP/P | Congenital disorder of glycosylation 1B (CDG1B) [MIM:602579] | Uniprot Humsavar | 102 | Phosphorylation | I | |
| 64 | MPZ | P25189 | VAR_004514 | Y | C | LP/P | Charcot-Marie-Tooth disease, demyelinating, 1B (CMT1B) [MIM:118200]; Dejerine-Sottas syndrome (DSS) [MIM:145900] | Uniprot Humsavar | 82 | Phosphorylation | I | |
| 65 | MTHFD1 | P11586 | VAR_074077 | T | I | LP/P | Combined immunodeficiency and megaloblastic anemia with or without hyperhomocysteinemia (CIMAH) [MIM:617780] | Uniprot Humsavar | 269 | Phosphorylation | I | |
| 66 | MYH9 | P35579 | HX17T | Y | C | Disease | liver_cancer[COSMIC] | COSMIC | 151 | Phosphorylation | I | TSG, fusion |
| 67 | MYO9A | B2RTY4 | VAR_081671 | Y | C | US | Myasthenic syndrome, congenital, 24, presynaptic (CMS24) [MIM:618198] | Uniprot Humsavar | 203 | Phosphorylation | I | |
| 68 | NME1 | P15531 | VAR_004625 | S | G | LB/B | - | Uniprot Humsavar | 120 | Phosphorylation | I | |
| 69 | NRCAM | Q92823 | TCGA-36-1574-01A | Y | H | Disease | ovarian_cancer[TCGA] | TCGA | 739 | Phosphorylation | I | |
| 70 | PAH | P00439 | VAR_000888 | S | R | LP/P | Non-phenylketonuria hyperphenylalaninemia (Non-PKU HPA) [MIM:261600] | Uniprot Humsavar | 87 | Phosphorylation | I | |
| 71 | PC | P11498 | VAR_058959 | Y | C | LP/P | Pyruvate carboxylase deficiency (PC deficiency) [MIM:266150] | Uniprot Humsavar | 304 | Phosphorylation | I | |
| 72 | PDHA1 | P08559 | VAR_021053 | Y | N | LP/P | Pyruvate dehydrogenase E1-alpha deficiency (PDHAD) [MIM:312170] | Uniprot Humsavar | 243 | Phosphorylation | I | |
| 73 | PDPR | Q8NCN5 | VAR_042504 | Y | H | LB/B | - | Uniprot Humsavar | 109 | Phosphorylation | I | |
| 74 | PHEX | P78562 | VAR_010619 | S | P | LP/P | Hypophosphatemic rickets, X-linked dominant (XLHR) [MIM:307800] | Uniprot Humsavar | 141 | Phosphorylation | I | |
| 75 | PIGW | Q7Z7B1 | VAR_071933 | T | P | LP/P | Glycosylphosphatidylinositol biosynthesis defect 11 (GPIBD11) [MIM:616025] | Uniprot Humsavar | 71 | Phosphorylation | I | |
| 76 | PINK1 | Q9BXM7 | VAR_046589 | T | M | LP/P | Parkinson disease 6 (PARK6) [MIM:605909] | Uniprot Humsavar | 313 | Phosphorylation | I | |
| 77 | PLCG2 | P16885 | TCGA-13-1498-01A | Y | D | Disease | ovarian_cancer[TCGA] | TCGA | 1036 | Phosphorylation | I | |
| 78 | PLOD2 | O00469 | VAR_022166 | T | I | LP/P | Bruck syndrome 2 (BRKS2) [MIM:609220] | Uniprot Humsavar | 608 | Phosphorylation | I | |
| 79 | PMS2 | P54278 | VAR_078537 | S | L | LP/P | Mismatch repair cancer syndrome 4 (MMRCS4) [MIM:619101] | Uniprot Humsavar | 815 | Phosphorylation | I | TSG |
| 80 | POLB | P06746 | VAR_088254 | Y | C | US | - | Uniprot Humsavar | 39 | Phosphorylation | I | |
| 81 | POLR2A | P24928 | VAR_082994 | T | M | LP/P | Neurodevelopmental disorder with hypotonia and variable intellectual and behavioral abnormalities (NEDHIB) [MIM:618603] | Uniprot Humsavar | 736 | Phosphorylation | I | |
| 82 | POR | P16435 | VAR_021159 | Y | C | LP/P | Antley-Bixler syndrome, with genital anomalies and disordered steroidogenesis (ABS1) [MIM:201750] | Uniprot Humsavar | 575 | Phosphorylation | I | |
| 83 | PRKCG | P05129 | 16905 | Y | N | Disease | lung_cancer[cBio] | cBio | 521 | Phosphorylation | I | |
| 84 | PROS1 | P07225 | VAR_014666 | T | M | LP/P | Thrombophilia due to protein S deficiency, autosomal dominant (THPH5) [MIM:612336] | Uniprot Humsavar | 78 | Phosphorylation | I | |
| 85 | PTEN | P60484 | VAR_026272 | S | F | LB/B | - | Uniprot Humsavar | 227 | Phosphorylation | I | TSG |
| 86 | PTEN | P60484 | VAR_007462 | Y | H | LP/P | Cowden syndrome 1 (CWS1) [MIM:158350] | Uniprot Humsavar | 68 | Phosphorylation | I | TSG |
| 87 | PTEN | P60484 | TCGA-06-0208-01A | Y | C | Disease | glioblastoma_tcga[TCGA] | TCGA | 68 | Phosphorylation | I | TSG |
| 88 | PTEN | P60484 | VAR_026263 | Y | C | LP/P | Cowden syndrome 1 (CWS1) [MIM:158350] | Uniprot Humsavar | 155 | Phosphorylation | I | TSG |
| 89 | RAC1 | P63000 | VAR_014546 | T | I | LB/B | - | Uniprot Humsavar | 108 | Phosphorylation | I | oncogene |
| 90 | RANBP2 | P49792 | VAR_050575 | S | G | LB/B | - | Uniprot Humsavar | 725 | Phosphorylation | I | TSG, fusion |
| 91 | RPS19 | P39019 | VAR_055442 | T | P | LP/P | Diamond-Blackfan anemia 1 (DBA1) [MIM:105650] | Uniprot Humsavar | 76 | Phosphorylation | I | |
| 92 | RUNX2 | Q13950 | VAR_012133 | S | R | LP/P | Cleidocranial dysplasia 1 (CLCD1) [MIM:119600] | Uniprot Humsavar | 118 | Phosphorylation | I | |
| 93 | RUNX2 | Q13950 | VAR_064082 | S | N | LP/P | Cleidocranial dysplasia 1 (CLCD1) [MIM:119600] | Uniprot Humsavar | 118 | Phosphorylation | I | |
| 94 | SH2D1A | O60880 | VAR_048009 | S | R | LP/P | Lymphoproliferative syndrome, X-linked, 1 (XLP1) [MIM:308240] | Uniprot Humsavar | 28 | Phosphorylation | I | |
| 95 | SLC13A5 | Q86YT5 | VAR_078916 | S | L | LP/P | Developmental and epileptic encephalopathy 25, with amelogenesis imperfecta (DEE25) [MIM:615905] | Uniprot Humsavar | 427 | Phosphorylation | I | |
| 96 | SLC25A42 | Q86VD7 | VAR_032970 | S | P | LB/B | - | Uniprot Humsavar | 39 | Phosphorylation | I | |
| 97 | SLC26A3 | P40879 | VAR_066072 | Y | C | LP/P | Diarrhea 1, secretory chloride, congenital (DIAR1) [MIM:214700] | Uniprot Humsavar | 520 | Phosphorylation | I | |
| 98 | SLC26A4 | O43511 | VAR_021650 | S | P | LP/P | Pendred syndrome (PDS) [MIM:274600] | Uniprot Humsavar | 137 | Phosphorylation | I | |
| 99 | SLC3A1 | Q07837 | VAR_011426 | Y | H | LP/P | Cystinuria (CSNU) [MIM:220100] | Uniprot Humsavar | 461 | Phosphorylation | I | |
| 100 | SLC52A2 | Q9HAB3 | VAR_077438 | Y | C | LP/P | Brown-Vialetto-Van Laere syndrome 2 (BVVLS2) [MIM:614707] | Uniprot Humsavar | 305 | Phosphorylation | I | |
| 101 | SMC1A | Q14683 | VAR_052438 | T | P | LB/B | - | Uniprot Humsavar | 28 | Phosphorylation | I | TSG |
| 102 | SMPD1 | P17405 | VAR_077323 | S | F | LB/B | - | Uniprot Humsavar | 510 | Phosphorylation | I | |
| 103 | SOD1 | P00441 | VAR_013535 | S | L | LP/P | Amyotrophic lateral sclerosis 1 (ALS1) [MIM:105400] | Uniprot Humsavar | 106 | Phosphorylation | I | |
| 104 | SOX11 | P35716 | VAR_071461 | S | P | LP/P | evelopmental disorder with microcephaly and with or without ocular malformations or hypogonadotropic hypogonadism (IDDMOH) [M | Uniprot Humsavar | 60 | Phosphorylation | I | |
| 105 | SPAST | Q9UBP0 | VAR_027215 | S | F | LP/P | Spastic paraplegia 4, autosomal dominant (SPG4) [MIM:182601] | Uniprot Humsavar | 436 | Phosphorylation | I | |
| 106 | SPAST | Q9UBP0 | VAR_067644 | S | P | LP/P | Spastic paraplegia 4, autosomal dominant (SPG4) [MIM:182601] | Uniprot Humsavar | 436 | Phosphorylation | I | |
| 107 | STAT3 | P40763 | VAR_037377 | T | I | LP/P | Hyper-IgE syndrome 1, autosomal dominant, with recurrent infections (HIES1) [MIM:147060] | Uniprot Humsavar | 622 | Phosphorylation | I | oncogene |
| 108 | SYT3 | Q9BQG1 | VAR_036389; HCC1395[COSMIC] | S | F | US | A breast cancer sample; breast_cancer[COSMIC] | Uniprot Humsavar; COSMIC | 474 | Phosphorylation | I | |
| 109 | TBK1 | Q9UHD2 | VAR_069754 | S | F | LB/B | - | Uniprot Humsavar | 151 | Phosphorylation | I | |
| 110 | TGFBR2 | P37173 | VAR_022352 | Y | N | LP/P | Loeys-Dietz syndrome 2 (LDS2) [MIM:610168] | Uniprot Humsavar | 336 | Phosphorylation | I | TSG |
| 111 | TGM1 | P22735 | VAR_058673 | Y | D | LP/P | Ichthyosis, congenital, autosomal recessive 1 (ARCI1) [MIM:242300] | Uniprot Humsavar | 365 | Phosphorylation | I | |
| 112 | TIFA | Q96CG3 | VAR_051422 | T | M | LB/B | - | Uniprot Humsavar | 19 | Phosphorylation | I | |
| 113 | TLN2 | Q9Y4G6 | VAR_076545 | S | L | US | - | Uniprot Humsavar | 339 | Phosphorylation | I | |
| 114 | TMPRSS6 | Q8IU80 | VAR_072902 | T | N | LP/P | Iron-refractory iron deficiency anemia (IRIDA) [MIM:206200] | Uniprot Humsavar | 287 | Phosphorylation | I | |
| 115 | TP53 | P04637 | 6-0644-01A; TCGA-08-0351-01A; TC | Y | C | LP/P | Li-Fraumeni syndrome (LFS) [MIM:151623]; lung_cancer[cBio]; glioblastoma_tcga[TCGA]; ovarian_cancer[TCGA] | Uniprot Humsavar; cBio; TCGA | 220 | Phosphorylation | I | oncogene, TSG, fusion |
| 116 | TP53 | P04637 | VAR_005958 | Y | H | US | Sporadic cancers | Uniprot Humsavar | 220 | Phosphorylation | I | oncogene, TSG, fusion |
| 117 | TP53 | P04637 | VAR_045116 | Y | D | US | Sporadic cancers | Uniprot Humsavar | 220 | Phosphorylation | I | oncogene, TSG, fusion |
| 118 | TP53 | P04637 | VAR_045117 | Y | F | US | A sporadic cancer | Uniprot Humsavar | 220 | Phosphorylation | I | oncogene, TSG, fusion |
| 119 | TP53 | P04637 | VAR_045118 | Y | N | US | Sporadic cancers | Uniprot Humsavar | 220 | Phosphorylation | I | oncogene, TSG, fusion |

| | | | | | | | | | | | | |
|---|---|---|---|---|---|---|---|---|---|---|---|---|
| 120 | TP53 | P04637 | VAR_005901 | T | A | US | Sporadic cancers | Uniprot Humsavar | 155 | Phosphorylation | I | oncogene, TSG, fusion |
| 121 | TP53 | P04637 | VAR_044834 | T | I | US | Sporadic cancers | Uniprot Humsavar | 155 | Phosphorylation | I | oncogene, TSG, fusion |
| 122 | TP53 | P04637 | VAR_044835 | T | M | US | A sporadic cancer | Uniprot Humsavar | 155 | Phosphorylation | I | oncogene, TSG, fusion |
| 123 | TP53 | P04637 | VAR_044836 | T | N | LP/P | Li-Fraumeni syndrome (LFS) [MIM:151623] | Uniprot Humsavar | 155 | Phosphorylation | I | oncogene, TSG, fusion |
| 124 | TP53 | P04637 | 4837; 16827; 17290; Pa01C[cBio; CC | T | P | US | Sporadic cancers; lung_cancer[cBio]; pancreatic_cancer[COSMIC] | Uniprot Humsavar; cBio; COSMIC | 155 | Phosphorylation | I | oncogene, TSG, fusion |
| 125 | TSC2 | P49815 | VAR_005664 | Y | C | LP/P | Tuberous sclerosis 2 (TSC2) [MIM:613254] | Uniprot Humsavar | 1650 | Phosphorylation | I | TSG |
| 126 | TSC2 | P49815 | VAR_018603 | S | F | LP/P | Tuberous sclerosis 2 (TSC2) [MIM:613254] | Uniprot Humsavar | 1653 | Phosphorylation | I | TSG |
| 127 | UHMK1 | Q8TAS1 | VAR_041273 | Y | D | LB/B | - | Uniprot Humsavar | 197 | Phosphorylation | I | |
| 128 | UROS | P10746 | VAR_003675 | Y | C | LP/P | Congenital erythropoietic porphyria (CEP) [MIM:263700] | Uniprot Humsavar | 19 | Phosphorylation | I | |
| 129 | VHL | P40337 | VAR_005766 | Y | D | LP/P | Von Hippel-Lindau disease (VHLD) [MIM:193300] | Uniprot Humsavar | 175 | Phosphorylation | I | TSG |
| 130 | VHL | P40337 | VAR_005714 | S | C | LP/P | Von Hippel-Lindau disease (VHLD) [MIM:193300] | Uniprot Humsavar | 111 | Phosphorylation | I | TSG |
| 131 | VHL | P40337 | VAR_005715 | S | N | LP/P | Von Hippel-Lindau disease (VHLD) [MIM:193300] | Uniprot Humsavar | 111 | Phosphorylation | I | TSG |
| 132 | VHL | P40337 | VAR_005716 | S | R | LP/P | Von Hippel-Lindau disease (VHLD) [MIM:193300] | Uniprot Humsavar | 111 | Phosphorylation | I | TSG |
| 133 | VHL | P40337 | VAR_005686 | S | I | LP/P | Von Hippel-Lindau disease (VHLD) [MIM:193300] | Uniprot Humsavar | 80 | Phosphorylation | I | TSG |
| 134 | VHL | P40337 | VAR_005687 | S | R | LP/P | Von Hippel-Lindau disease (VHLD) [MIM:193300] | Uniprot Humsavar | 80 | Phosphorylation | I | TSG |
| 135 | VHL | P40337 | VAR_005688 | S | N | LP/P | Pheochromocytoma (PCC) [MIM:171300]; Von Hippel-Lindau disease (VHLD) [MIM:193300] | Uniprot Humsavar | 80 | Phosphorylation | I | TSG |
| 136 | VKORC1 | Q9BQB6 | VAR_065788 | S | W | LP/P | Coumarin resistance (CMRES) [MIM:122700] | Uniprot Humsavar | 52 | Phosphorylation | I | |
| 137 | XRCC3 | O43542 | VAR_013006 | T | M | LP/P | Melanoma, cutaneous malignant 6 (CMM6) [MIM:613972] | Uniprot Humsavar | 241 | Phosphorylation | I | |
| 138 | YWHAZ | P63104 | VAR_082642 | S | L | US | - | Uniprot Humsavar | 145 | Phosphorylation | I | |
