## Supplementary Table 5 for "Mapping Cryptic Phosphorylation Sites in the Human Proteome"

| ROW_ID | GENE | UPID | FTID | WT_AA | VAR_AA | VAR_TYPE | DISEASE(s) | MUT_SOURCE | MOD_RSD | MOD_TYPE | VAR_CLASS | ONC_TSG |
|---|---|---|---|---|---|---|---|---|---|---|---|---|
| 1 | ANOS1 | P23352 | VAR_069207 | Y | D | US | - | Uniprot Humsavar | 217 | Phosphorylation | I | |
| 2 | CYBB | P04839 | VAR_025613 | Y | D | LP/P | Granulomatous disease, chronic, X-linked (CGDX) [MIM:306400] | Uniprot Humsavar | 41 | Phosphorylation | I | |
| 3 | F8 | P00451 | VAR_028533 | Y | D | LP/P | Hemophilia A (HEMA) [MIM:306700] | Uniprot Humsavar | 450 | Phosphorylation | I | |
| 4 | HBB | P68871 | VAR_003063 | Y | D | LB/B | - | Uniprot Humsavar | 131 | Phosphorylation | I | |
| 5 | HLA-DPB1 | P04440 | VAR_060642 | Y | D | LB/B | - | Uniprot Humsavar | 57 | Phosphorylation | I | |
| 6 | IDS | P22304 | VAR_007351 | Y | D | LP/P | Mucopolysaccharidosis 2 (MPS2) [MIM:309900] | Uniprot Humsavar | 225 | Phosphorylation | I | |
| 7 | PLCG2 | P16885 | TCGA-13-1498-01A | Y | D | Disease | ovarian_cancer[TCGA] | TCGA | 1036 | Phosphorylation | I | |
| 8 | TGM1 | P22735 | VAR_058673 | Y | D | LP/P | Ichthyosis, congenital, autosomal recessive 1 (ARCI1) [MIM:242300] | Uniprot Humsavar | 365 | Phosphorylation | I | |
| 9 | TP53 | P04637 | VAR_045116 | Y | D | US | Sporadic cancers | Uniprot Humsavar | 220 | Phosphorylation | I | oncogene, TSG, fusion |
| 10 | UHMK1 | Q8TAS1 | VAR_041273 | Y | D | LB/B | - | Uniprot Humsavar | 197 | Phosphorylation | I | |
| 11 | VHL | P40337 | VAR_005766 | Y | D | LP/P | Von Hippel-Lindau disease (VHLD) [MIM:193300] | Uniprot Humsavar | 175 | Phosphorylation | I | TSG |
